# Polygenic risk modeling with latent trait-related genetic components

**DOI:** 10.1101/808675

**Authors:** Matthew Aguirre, Yosuke Tanigawa, Guhan Ram Venkataraman, Rob Tibshirani, Trevor Hastie, Manuel A. Rivas

**Author notes:** Correspondence to Manuel A. Rivas.

## Abstract

Polygenic risk models have led to significant advances in understanding complex diseases and their clinical presentation. While models like polygenic risk scores (PRS) can effectively predict outcomes, they do not generally account for disease subtypes or pathways which underlie within-trait diversity. Here, we introduce a latent factor model of genetic risk based on components from Decomposition of Genetic Associations (DeGAs), which we call the DeGAs polygenic risk score (dPRS). We compute DeGAs using genetic associations for 977 traits in the UK Biobank and find that dPRS performs comparably to standard PRS while offering greater interpretability. We show how to decompose an individual’s genetic risk for a trait across DeGAs components, highlighting specific results for body mass index (BMI), myocardial infarction (heart attack), and gout in 337,151 white British individuals, with replication in a further set of 25,486 non-British white individuals from the Biobank. We find that BMI polygenic risk factorizes into components relating to fat-free mass, fat mass, and overall health indicators like physical activity measures. Most individuals with high dPRS for BMI have strong contributions from both a fat mass component and a fat-free mass component, whereas a few ‘outlier’ individuals have strong contributions from only one of the two components. Overall, our method enables fine-scale interpretation of the drivers of genetic risk for complex traits.

## Introduction

Common diseases like diabetes and heart disease are leading causes of death and financial burden in the developed world^1^. Polygenic risk scores (PRS), which sum effects from many risk loci for a trait, have been used to identify individuals at high risk for conditions like cancer, diabetes, heart disease, and obesity^2–5^. Although many versions of PRS can be used to estimate risk^6–8^, previous work suggests that a “palette” model which decomposes genetic risk into pathways could better describe the clinical manifestations of complex disease^9^.

Here, we present a polygenic model based on latent trait-related genetic components identified using Decomposition of Genetic Associations (DeGAs) ^10^. Where standard PRS for a trait models genetic risk as a sum of effects from genetic variants, the DeGAs polygenic risk score (dPRS) models risk as a sum of contributions from DeGAs components^10^. Each DeGAs component consists of a set of variants which affect a subset of the traits (**Figure 1**). The component’s genetic loading is a component PRS (cPRS) that approximates risk for a weighted combination of relevant traits. Instantiated cPRS values for an individual can then be used to create a profile that describes disease risk and informs genetic subtyping.

**Figure 1:**
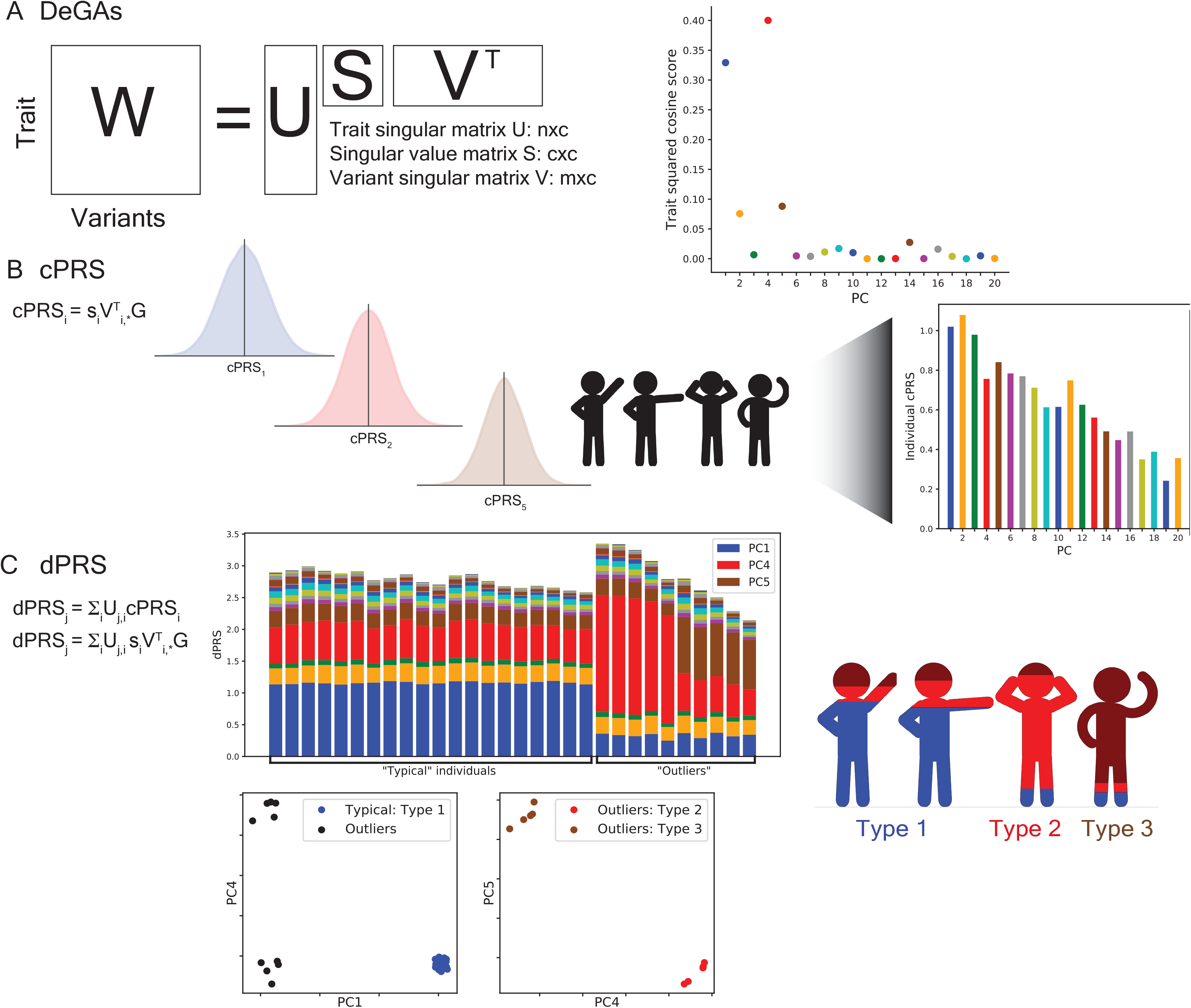
Study overview. (A) Matrix Decomposition of Genetic Associations (DeGAs) is performed by taking the truncated singular value decomposition (TSVD) of a matrix W (n xm) containing summary statistics from GWAS of n=977 traits over m=469,341 variants from the UK Biobank. The squared columns of the resulting singular matrices U (n x c) and V (m x c) measure the importance of traits (variants) to each component; the rows map traits (variants) back to components. The squared cosine score (a unit-normalized row of US) for some hypothetical trait indicates high contribution from PC1, PC4, and PC5. (B) Component polygenic risk scores (cPRS) for the ith component is defined as siVTi,*G (i-th singular value in S and i-th row in VT), for an individual with genotypes G. (C) DeGAs polygenic risk scores (dPRS) for trait j are recovered by taking a weighted sum of cPRSi, with weights from U (j,i-th entry). We also compute DeGAs risk profiles for each individual (Methods), which measure the relative contribution of each component to genetic risk. We “paint” the dPRS high risk individuals with these profiles and label them “typical” or “outliers” based on similarity to the mean risk profile (driven by PC1, in blue). Outliers are clustered on their profiles to find additional genetic subtypes: this identifies “Type 2” and “Type 3”, with risk driven by PC4 (red) and PC5 (tan). Clusters visually separate each subtype along relevant cPRS (below). Image credit: VectorStock.com/1143365.

As proof of concept, we compute DeGAs using summary statistics generated from genome-wide associations between 977 traits and 469,341 independent common variants in a subset of unrelated white British individuals (*n*=236,005) in the UK Biobank^11^ (**Methods**). We then develop a series of dPRS models and evaluate their performance in additional independent samples of unrelated white British individuals (*n*=33,716 validation set; *n*=67,430 test set), and in UK Biobank non-British whites (*n*=25,486 extra test set). We highlight results for body mass index (BMI), myocardial infarction (MI/heart attack), and gout, motivated by their high prevalence among older individuals in this cohort^12^.

## Material and Methods

### Study Population

The UK Biobank is a large longitudinal cohort study consisting of 502,560 individuals aged 37-73 at recruitment during 2006-2010^11^. The data acquisition and study development protocols are online (http://www.ukbiobank.ac.uk/wp-content/uploads/2011/11/UK-Biobank-Protocol.pdf). In short, participants visited a nearby center for an in-person baseline assessment where various anthropometric data, blood samples, and survey responses were collected. Additional data were linked from registries and collected during follow-up visits.

We used a subsample consisting of 337,151 unrelated individuals of white British ancestry for genetic analysis. We split this cohort at random into three groups: a 70% training population (*n*=236,005), a 10% validation population (*n*=33,716), and a 20% test population (*n*=67,430). We used the training population to conduct genome-wide association studies for DeGAs and the validation population to evaluate dPRS model performance for various DeGAs hyperparameters. We report final associations and performance measures in the test population. An additional cohort of unrelated non-British White individuals (*n*=25,486) was used as an extra test set. The “white British” and “non-British white” populations were defined using a combination of genotype PCs from UK Biobank’s PCA calculation and self-reported ancestry (UK Biobank Field 21000)^11,13^.

In brief, individuals were subject to the following filtration criteria from the UK Biobank sample quality control file (ukb_sqc_v2.txt): “putative sex chromosome aneuploidy=0”, “het_missing_outliers=0”, “excess_relatives=0”, and “used_in_pca_calculation=1”. Individuals were labeled “white” based on two thresholds in Biobank’s PCA calculation: −20 <= PC1 <= 40 and −25 <= PC2 <= 10. We further designate individuals in the “non-British white” set as those that self-reported white but not British ancestry in Field 21000. The “white British” set was taken to be individuals with “in_white_British_ancestry_subset=1”.

### Genome-wide association studies in the UK Biobank

PLINK v2.00a^14^ [2 April 2019] was used for genome-wide associations of 805,426 directly genotyped variants, 362 HLA allelotypes, and 1,815 non-rare (AF > 0.01%) copy number variants^15^ (CNV) in the UKB training population. We used the --glm Firth-fallback option to apply an additive-effect model across all sites. Quantitative trait values were rank-normalized using the --pheno-quantile-normalize flag. The following covariates were used: age, sex, the first four genetic principal components, and, for variants present on both of the UK Biobank’s genotyping arrays, the array which was used for each sample.

Prior to public release, genotyped sites and samples were subject to rigorous quality control by the UK Biobank^11^. In brief, markers were subject to outlier-based filtration on effects due to batch, plate, sex, genotyping array, as well as discordance across control replicates. Samples with excess heterozygosity (thresholds varied by ancestry) or missingness (> 5%) were excluded from the data release. Prior to use in downstream methods, we performed additional variant quality control on array-genotyped variants, including more stringent filters on missingness (> 1%), gross departures (p < 10^−7^) from Hardy-Weinberg Equilibrium, and other indicators of unreliable genotyping^16^. As with previous versions of DeGAs, we further filtered variants by minor allele frequency (MAF > 0.01%), array-specific missingness (< 5%), and LD-independence^10^. The LD independent set was computed with “--indep-pairwise 50 5 0.5” in PLINK v1.90b4.4 [21 May 2017]. MAF and LD filters were applied within and across each array-genotyped group. This process resulted in a set of 469,341 variants (467,427 genotyped variants, 118 HLA allelotypes, and 1,796 CNVs) for our analysis.

Binary disease outcomes were defined from UK Biobank resources using a previously described method which combines self-reported questionnaire data and diagnostic codes from hospital inpatient data^20,21^. Additional traits like biomarkers, environmental variables, and self-reported questionnaire data like health outcomes and lifestyle measures were collected from fields curated by the UK Biobank and processed using previously described methods^16,17^. Multiple observations were processed by taking the median of quantitative values, or by defining an individual as a binary case if any recorded instance met the trait’s defining criteria. In all, we collected 977 traits with at least 1000 observations (quantitative traits) or cases (binary traits). These comprise most common traits in the Global Biobank Engine^18^, excluding imaging features and traits which were subject to manual curation. A full list of traits and their Global Biobank Engine IDs is in **Data S1**. Summary statistics from all GWAS described here are publicly available on the Global Biobank Engine (**Web Resources**). In this work, we highlight results for body mass index (GBE ID: INI21001), myocardial infarction (HC326), and gout (HC328).

### Risk modeling using Decomposition of Genetic Associations (DeGAs)

Given GWAS summary statistics, we computed DeGAs as previously described^10^. First, a sparse matrix of genetic associations *W* (*n* × *m*) was populated with effect size estimates (or *z*-statistics) between the *n* =977 traits and *m* =469,341 variants. Only variants with at least two associations were used (*p* < 10^−6^; **Figure S1** has additional cutoffs). After filtration, rows of *W* were standardized to zero mean and unit variance, to give traits equal relative weight.

Next, we performed a truncated singular value decomposition (TSVD) on *W* using the TruncatedSVD function in the scikit-learn python module^19,20^ to identify the top *c*=500 trait-related genetic components. TSVD outputs three matrices whose product approximates *W*: a trait singular matrix *U* (*n* × *c*), a variant singular matrix *V* (*m* × *c*), and a diagonal matrix *S* (*c* × *c*) of singular values *s_i_* (**Figure 1a**). *W* is approximated by *U*, *S*, and *V* as below:

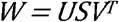

The matrices *U, S*, and *V* are then used to compute component polygenic risk scores (cPRS). The component PRS for the *i-*th DeGAs component can be written:

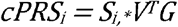

for an individual with genotypes *G* (*m* × 1) over the variants used in DeGAs. Here, *S_i,*_* is the *i*-th row of *S*. With cPRS, we define the DeGAs polygenic risk score (dPRS) for the *j*-th trait:

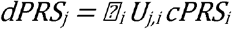

where *U_j,i_* is the (j,i)’th entry of *U*. In terms of the matrices *U*, *S*, and *V*, this can be rewritten:

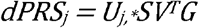

For interpretability, the population distribution of dPRS for each trait *j* is scaled to zero mean and unit variance, independently of the distributions of dPRS for other traits.

We further relate individuals to traits via components using a measure we call the DeGAs risk profile (dRP). An individual’s DeGAs risk profile for a phenotype *j* is a vector over the *c* DeGAs components, where the value for the *i*-th component is proportional to

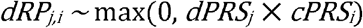

with a denominator introduced for normalization so that these values sum to one. Note we only consider component scores which have the same sign as the overall risk score when estimating their contribution to an individual’s genetic risk, hence the max operator. The DeGAs risk profile is therefore a normalized measure which, for high risk individuals with positive dPRS, is the fraction of risk owing to driving components. Analogously, for low risk individuals with negative dPRS, it measures the contribution from protective components.

### Computing polygenic risk scores

As a baseline model for dPRS, we computed single-trait polygenic risk scores (PRS) with a pruning and thresholding approach using the same summary statistics which were input to DeGAs. As DeGAs requires variants filtered on LD independence and for *p* < *p\** based on a critical value *p\** (see above), we simply used the variants present in the DeGAs input matrix *W*. Specifically, the PRS weights for trait *j* were taken from the *j*-th row of *W*. The PRS was then computed with PLINK v1.90b4.4 [21 May 2017] using the --score flag, with the `sum`, `center`, and `double-dosage` modifiers. These correspond to the assumptions that variants make additive contributions across sites; that the mean distribution of risk is zero; and that the effect alleles have additive effects. These are the same assumptions used in our GWAS.

In a similar fashion, polygenic scores (cPRS) for all DeGAs components were computed with PLINK2 v2.00a2 (2 Apr 2019) using the --score flag, with ‘center’ and ‘cols=scoresums’ modifiers. These modifiers correspond to the same assumptions as in the PRS: that genetic effects are additive across sites (this is the default genotype model for -- score); that each component is zero-centered; and that alleles make additive contributions. Given population-wide estimates of cPRS for every component, we computed dPRS and DeGAs risk profiles for each trait using the formulas listed above.

In some analysis, PRS and dPRS were further adjusted by age, sex, and four genetic principal components from UK Biobank’s PCA calculation. Covariate adjustment was performed by fitting a multiple regression model with dPRS (or PRS) and covariates in the validation population. We also fit a covariate-only model using the same procedure (without either polygenic score) and used its performance as baseline for the joint models (**Figure 2**).

**Figure 2:**
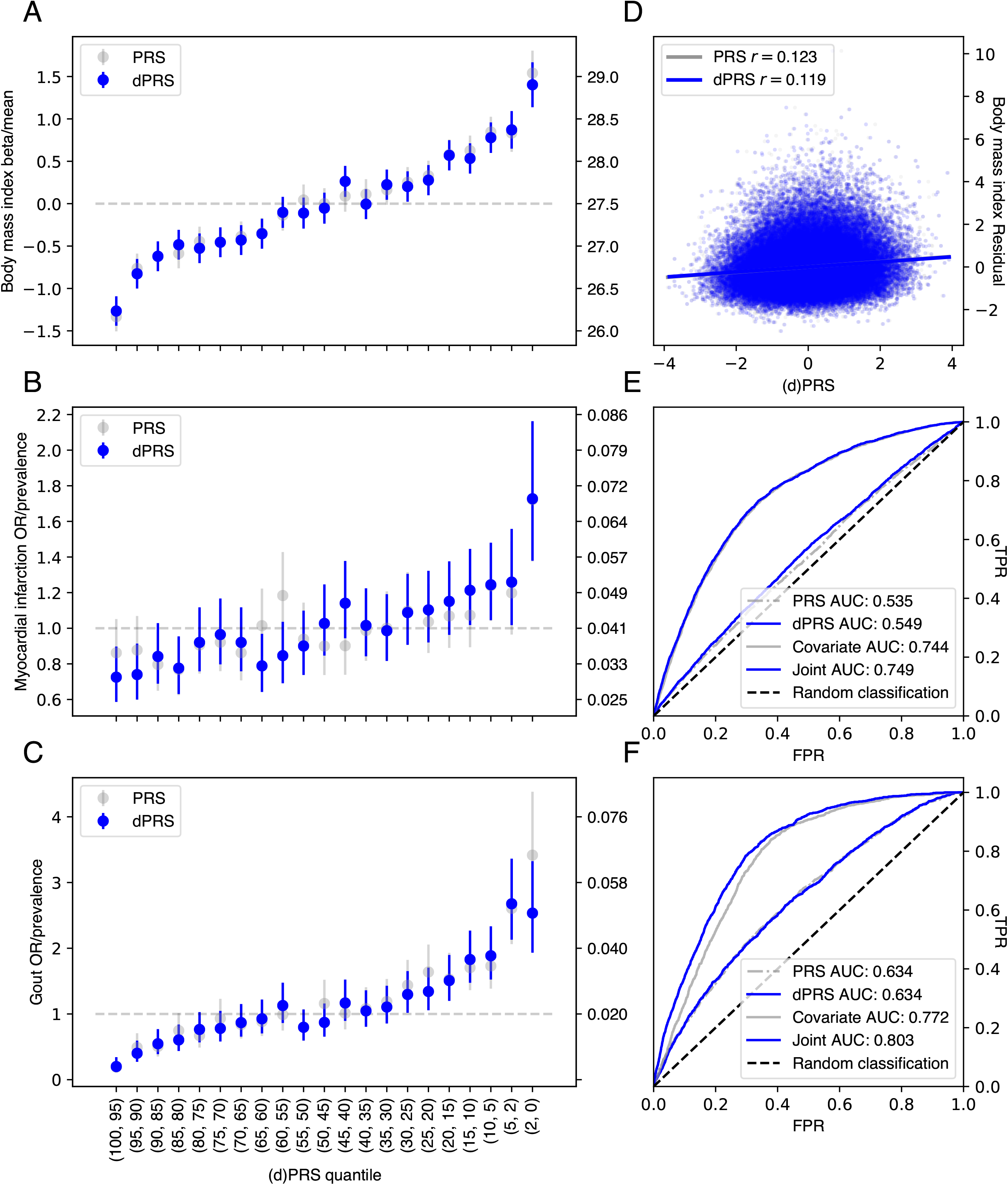
Performance of dPRS. (A-C) Effect of increased risk (dPRS or PRS) on BMI, MI, and gout. Beta/OR (left axis) were estimated by comparing the quantile of interest (x-axis) with a middle quantile (40-60%), adjusted for these covariates: age, sex, 4PCs (Methods). Trait mean or prevalence (right axis) was computed within each quantile; error bars denote the 95% confidence interval of each estimate. (D) Correlation between dPRS or PRS and covariate adjusted BMI. Receiver operating curves with area under curve (AUC) values for MI (E) or gout (F) for dPRS, PRS, covariates, and a joint model with covariates and dPRS. Models with covariates were fit in the validation set; all evaluation was in the test set. (Methods).

### Model validation

To select DeGAs hyperparameters (the input *p*-value filter, and whether to use GWAS betas or *z*-statistics as weights) we performed a grid search over a range of filtering *p*-values for both betas and *z*-statistics. Performance of a DeGAs instance was assessed using the average correlation between its dPRS models and their respective traits. For all traits used in the decomposition, we computed Spearman’s rho (rank correlation) between dPRS and covariate-adjusted trait residuals in the training and validation sets. Residual traits are the result of regressing out age, sex, and four genetic principal components. To avoid overfitting, we further required modest correlation (Pearson’s *r*^2^ > 0.5) between training and validation set performance. With this constraint, we find optimal performance using betas, a *p*-value cutoff of 10^−6^, and 500 components (**Figure S1**), and note that this model explains nearly all (>99%) of the variance of its corresponding input matrix *W* (**Figure S2**).

For this best-performing DeGAs instance, we present several assessment metrics for each polygenic score (dPRS and PRS) in each study population (**Figure 2**). For each score and population, we estimated disease prevalence and mean quantitative trait values at various population risk strata. We also estimated the effect of dPRS (or PRS) quantiles on traits using a two-step approach. First, in the training set we compute the below regression:

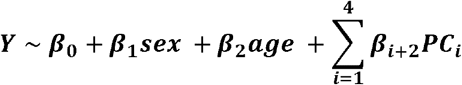

with PC_i_ the i’th population-genetic PC, not the i’th DeGAs component. Second, in the test/ validation set we estimate the effect β due to PRS quantile using the above parameters:

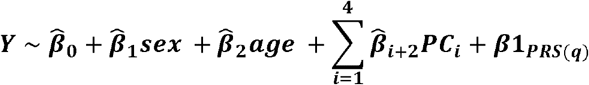

where ***1**_PRS(q)_* is an indicator function that equals 1 if an individual is in the quantile of interest *q* (e.g. 0-2%), and 0 if the individual is in the baseline group (40-60% risk quantile). Individuals in neither the quantile of interest nor the baseline group were excluded; if individuals were in both *q* and the baseline group (e.g. if *q* were 45-40%) they were counted in *q* and removed from the baseline group.

We then assessed how well dPRS and PRS predicted quantitative trait values, and how well they performed binary classification on disease status. For quantitative traits, we report Pearson’s *r* between each score and residual trait values. For binary traits, we report the area under the receiver operating curve (AUROC/AUC) for each score as a classifying metric, both alone and in a joint model with covariates. As baseline, we also report AUC for a covariate-only model (see above for a description of this model and covariate adjustment).

### Classifying genetic risk profiles from DeGAs components

In order to assess whether our method could identify subtypes of genetic risk, we analyzed the DeGAs risk profiles of high-risk individuals whose dPRS is driven by an “atypical” combination of DeGAs components. We used the Mahalanobis distance (D_M_) to identify outlier individuals whose *z*-scored distance from the mean DeGAs risk profile exceeded 1:

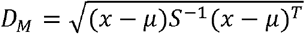

where *x* is the DeGAs risk profile; *μ* is the mean profile; and *S* is the identity matrix. Traditionally, *S* is taken to be the covariance matrix for each of the features across all *x*. However, for this measure we treat each component as having equal variance. This results in the above formula reducing to the Euclidean distance between a profile *x* and the mean profile *μ*, which can be used to identify “atypical” individuals rather than statistical outliers.

We then intersected this set of outlier individuals with the top 5% of dPRS to create the “high risk outlier” group. Here, we define the mean risk profile for a trait as the component-wise mean across all individuals’ DeGAs risk profiles in a high risk set (top 5% of dPRS). To identify subtypes among high risk outliers, we performed a *k*-means clustering of their DeGAs risk profiles using the KMeans function from the python scikit-learn module^19^. The number of clusters *k* was determined by optimizing a statistic over putative values of *k* ranging from 1 to 20. Specifically, we used the gap-star statistic in the python “gap-statistic” module^21,22^ as the statistic for selecting a value *k*. We then evaluated which components drove risk in each cluster by computing a mean risk profile for the group (defined as above). The mean risk profile was then renormalized to one for visualization (**Figure 4g-i**).

## Results

Genome-wide associations between 977 traits and 469,341 independent human leukocyte antigen (HLA) allelotypes, copy-number variants^14^, and array-genotyped variants were computed in a training set of 236,005 unrelated white British individuals from the UK Biobank study^11^ (**Methods**). We applied DeGAs^10^ to scaled beta-or *z*-statistics from these GWAS with varying *p*-value thresholds for input (**Figure 1a**). We then defined polygenic risk scores for each DeGAs component (cPRS; **Figure 1b**) and used them to build the DeGAs polygenic risk score (dPRS; **Figure 1c**). The model with optimal out-of-sample prediction (**Figure S1**) corresponded to DeGAs on beta values with significant (*p* < 10^−6^) associations.

To validate this model, we estimated disease prevalence (or, for BMI, mean BMI) at several quantiles of risk in a held-out test set of white British individuals in the UK Biobank (*n*=67,430). For all example traits we observed increasing severity (quantitative) or prevalence (binary) at increasing quantiles of dPRS (**Figure 2a-c**) adjusted for age, sex, and the first 4 genotype principal components from UK Biobank’s PCA calculation^11^. This trend was most pronounced at the highest risk quantile (2%) for each trait. At this stratum we observed 1.40 kg/m^2^ higher BMI (95% CI: 1.14-1.67); 1.73-fold increased odds of MI (CI: 1.38-2.16); and 2.53-fold increased odds of gout (CI: 1.93-3.33) over the population average in the test set (overall *n*=67,235 individuals for BMI; 2,812 MI cases; and 1,484 gout cases).

Further, we found dPRS to be comparable to prune- and threshold-based PRS using the same input data (**Figure S2**). Although there was some discrepancy between the individuals considered high risk by each model (**Figure S3, Table S2**), we observe similar effects at the extreme tail of PRS as with dPRS. The top 2% of PRS for each trait had 1.54 kg/m^2^ higher BMI (CI: 1.28-1.81); 1.72-fold increased odds of MI (CI: 1.38-2.16); and 3.42-fold odds of gout (CI: 2.67-4.38) (**Figure 2a-c**) using the same covariate adjustment as dPRS. Population-wide predictive measures were also similar, with BMI residual *r*=0.21, and PRS AUC (not adjusted for covariates) 0.54 for MI and 0.63 for gout (**Figure 2d-f**). We also note similar performance for BMI dPRS predicting obesity (defined as BMI > 30; **Figure S5**), with OR=1.7 at the 2% tail, and AUC=0.56. On balance, despite the reduced rank of the DeGAs risk models — the input matrix *W* is reduced from ~1,000 traits to a 500-dimensional representation — we achieve performance equivalent to traditional PRS for these example traits and observe a similar trend for the other traits (**Figure S2, Data S1**).

However, we note that dPRS and PRS add little population-wide predictive value over factors such as age, sex, and demographic effects that are captured by genomic PCs (**Figure 2d-f**). At the population level, we found *r*=0.12 between covariate-adjusted dPRS and residualized BMI. For binary traits, the area-under the receiver operating curve (AUC) was 0.55 for MI and 0.63 for gout, with unadjusted dPRS as the classifying score. Adjusting for covariates, the marginal increase in AUC is modest: 0.005 for MI and 0.03 for gout.

### Characterizing DeGAs Components

We describe the latent structure identified through DeGAs by annotating each component with its contributing traits and variants, aggregated by gene. The relative importance of traits to components is measured using the trait contribution score^10^, which corresponds to a squared column of the trait singular matrix *U*. The relative importance of components to each trait is measured using the trait squared cosine score^10^, which is a normalized squared row of *US*. The contribution and squared cosine score are defined analogously for variants and genes using the variant singular matrix *V*. For each example trait, we highlight 5 components of interest (ranked by the trait squared cosine score) and describe them by their respective trait contribution scores (**Figure 3**) and gene contribution scores. The trait and gene contribution scores for all components can be found in **Data S2** and **Data S3**, respectively.

**Figure 3:**
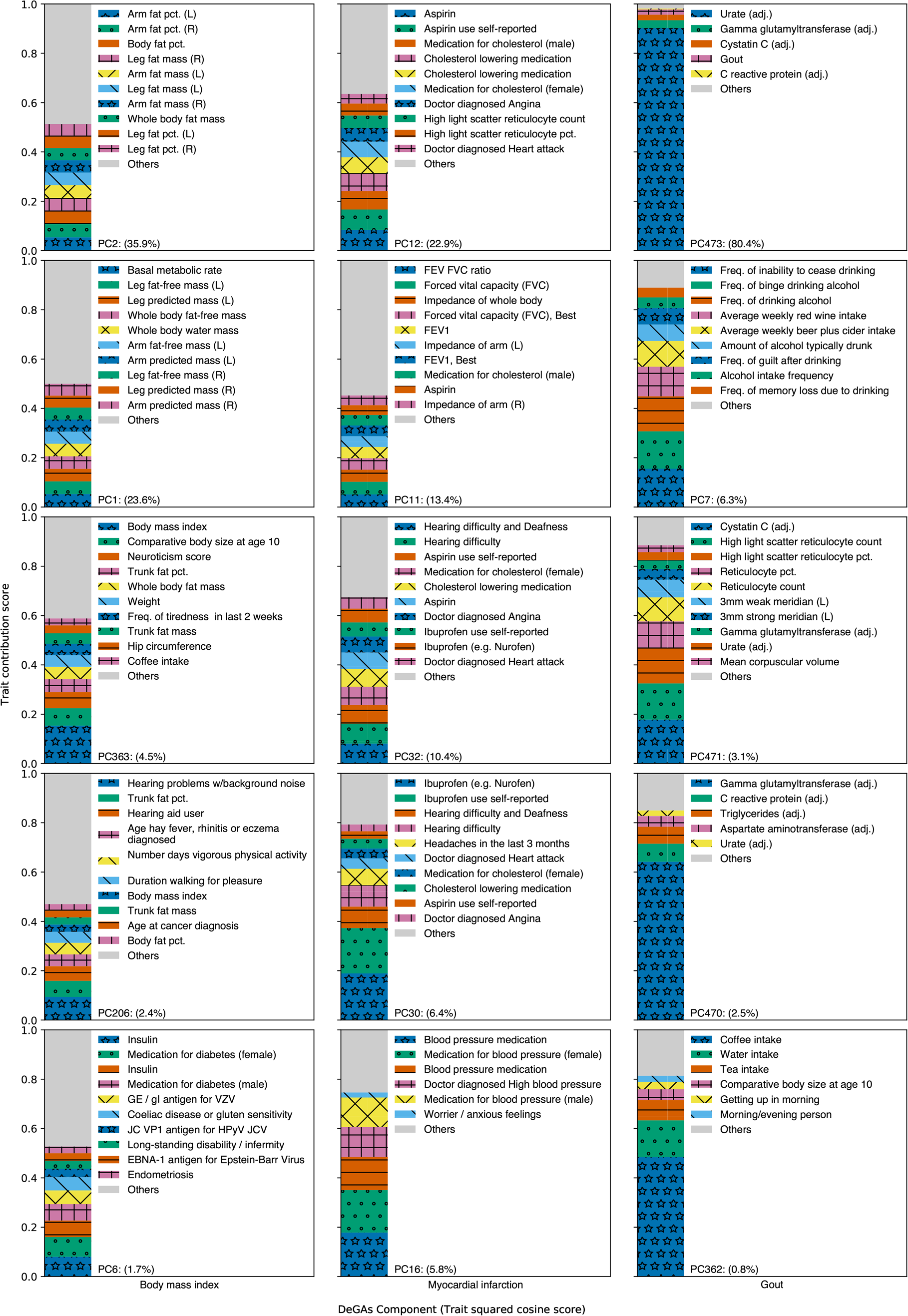
Top five DeGAs components for each example trait. Top five DeGAs components for BMI (left), MI (center), and gout (right), as ranked by the trait squared cosine score. Each component is labeled with its top ten traits, as determined by the trait contribution score (squared column of U), and with its relative importance (squared cosine score). Traits are displayed for a component if their contribution score for the component exceeds 0.02.

Body mass index is a polygenic trait with associated genetic variation relevant to adipogenesis, insulin secretion, energy metabolism, and synaptic function^10,23^. Here, the DeGAs trait squared cosine score (**Figure 3**) indicates strong contribution from components related to body size and fat-free mass (PC1; 23.6%), fat mass (PC2; 35.9%), as well as risk factors for obesity like body size at age 10 and trunk fat percentage (PC363; 4,5%). Components related to exercise (PC206; 2.4%) and diabetes (PC6; 1.7%) also contribute.

Genetic variation proximal to *STC2* and *MC4R* contribute strongly to both PC1 and PC2 (**Data S3**). *STC2* is a stanniocalcin-related protein most highly expressed in cardiomyocytes and skeletal muscle. It has previously been associated with lean mass traits in humans^24,25^ and has been shown to restrict post-natal growth in mouse^26^. *MC4R* is a melanocortin receptor in the G-protein coupled receptor family. It is primarily expressed in the brain, is known to play a role in energy homeostasis and somatic growth^27,28^, and has been associated with fat-mass and obesity-related traits in humans^29^. Both components also have contribution from variation proximal to *FTO* and *DLEU1*, both of which associate with traits affecting body size in adults^30,31^. *FTO* is an alpha-ketoglutarate dependent dioxygenase whose causal role in BMI has been questioned^32^; *DLEU1* is a tumor-suppressing lncRNA named for its frequent deletion in patients with chronic lymphocytic leukemia^33^. These components reflect of the roles of adipogenesis and growth regulation pathways in high BMI.

Myocardial infarction is a polygenic outcome with well-established risk factors attributable to common and rare genetic variation^5^, age, sex, and lifestyle attributes like diet and smoking. DeGAs components important to this trait are related to measures of lung function, as well as usage of medications for an array of conditions which are comorbid with MI (**Figure 3**). These medications include aspirin, ibuprofen, and cholesterol-lowering medications (e.g. statins), which are represented across PC12 (22.9%; also includes reticulocyte measurements), PC32 (10.4%; also includes hearing problems and angina), and PC30 (6.4%; also includes headaches). A component related to blood pressure medications also contributes (PC16; 5.8%). Another relevant component (PC11; 13.4%) has contribution from measures of lung function like forced expiratory volume in 1 second (FEV1), forced vital capacity (FVC), and the ratio of the two (FEV FVC ratio).

Two of these components, PC11 and PC12, have contribution from variation proximal to the lipoprotein gene *LPA*, at the *9p21.3* susceptibility locus (*CDKN2B*), and in the brain-expressed solute carrier *SLC22A3*^34^ (**Data S3**). Variation in these three genes also contributes to PC32, as does variation proximal to the transcription factor *STAT6* (which has been associated with adult-onset asthma and inflammatory response to mosquito bites^35,36^. PC30 also has contribution from *STAT6*, as well as the phosphatase and actin regulator *PHACTR1*, which has been identified in prior coronary artery disease GWAS^37^. These components reflect the diversity of conditions and risk factors which are comorbid with MI.

Gout is a heritable (*h^2^*=17.0-35.1%) common complex form of arthritis characterized by severe sudden onset joint pain and tenderness, which is believed to arise due to excessive blood uric acid which crystallizes and forms deposits in the joints^38^. The top component for gout (**Figure 3**) has strong contribution from covariate-adjusted blood urate^13^ (PC473; 80.4%). This component also explains most of the variance in the input genetic associations for gout. Meanwhile, two other important components measure dietary intake. One of them (PC7; 6.3%) is driven by measures of alcohol use and abuse; the other (PC362; 0.8%) is related to coffee, water, and tea intake. Two other components are related to covariate and statin-adjusted biomarkers^13^, namely, cystatin C (PC471; 3.1%) and gamma glutamyltransferase (PC470; 2.5%). The key gene for PC473 is *SLC2A9*, which is involved in uric acid transport and is associated with gout^39^. The alcohol dehydrogenase *ADH1B* is key to PC7 and is associated with alcoholism and blood urea nitrogen (BUN)^40,41^. Indeed, alcohol use has been identified as a lifestyle risk factor for gout^42^. These components reflect the clinical pathogenesis of gout by urate buildup, as well as some of its lifestyle risk factors.

### Painting DeGAs Risk Profiles

To further characterize the genetic architecture of these traits, we “painted” the genetic risk profiles of each high-risk individual (top 5% of dPRS). For this, we decomposed each individual’s dPRS across DeGAs components into a vector we call the DeGAs risk profile (**Methods**). The DeGAs risk profile is an individual-level measure over the (in this case) *c*=500 DeGAs components and is normalized such that the entries in the vector sum to one. For individuals with higher than average risk (dPRS > 0), it describes the contributions from components which contribute positively to risk; for individuals with lower than average risk (dPRS < 0) it describes the contributions from components which have protective effects. As an individual rather than population-level measure, the DeGAs risk profile can be used to further examine the underlying genetic diversity among high risk individuals (**Figure 4a-c**) in a way which complements the trait and gene squared scores from DeGAs.

We therefore investigated the diversity of components which drive risk among high risk individuals, using their DeGAs risk profiles. We used the Mahalanobis criterion (**Methods**) to find individuals in the test population whose risk profiles significantly differed from average. We then found high-risk individuals (top 5% of dPRS) among these outliers (*z*-scored Mahalanobis distance > 2) to identify a group of “high-risk outliers”. For these individuals (**Figure 4d-f**), genetic risk is often driven by the same components as for other high-risk individuals (**Figure 4a-c**), but the degree to which certain components contribute can differ. For example, while the trait squared cosine score for gout identifies PC473 (urate) as the top component (**Figure 3**), the DeGAs risk profile suggests PC7 (alcohol use traits) has a key role in driving genetic risk for some individuals with high gout dPRS (**Figure 4c**). This suggests that the DeGAs risk profile can identify individuals with high genetic risk whose pathology may differ from “typical”.

**Figure 4:**
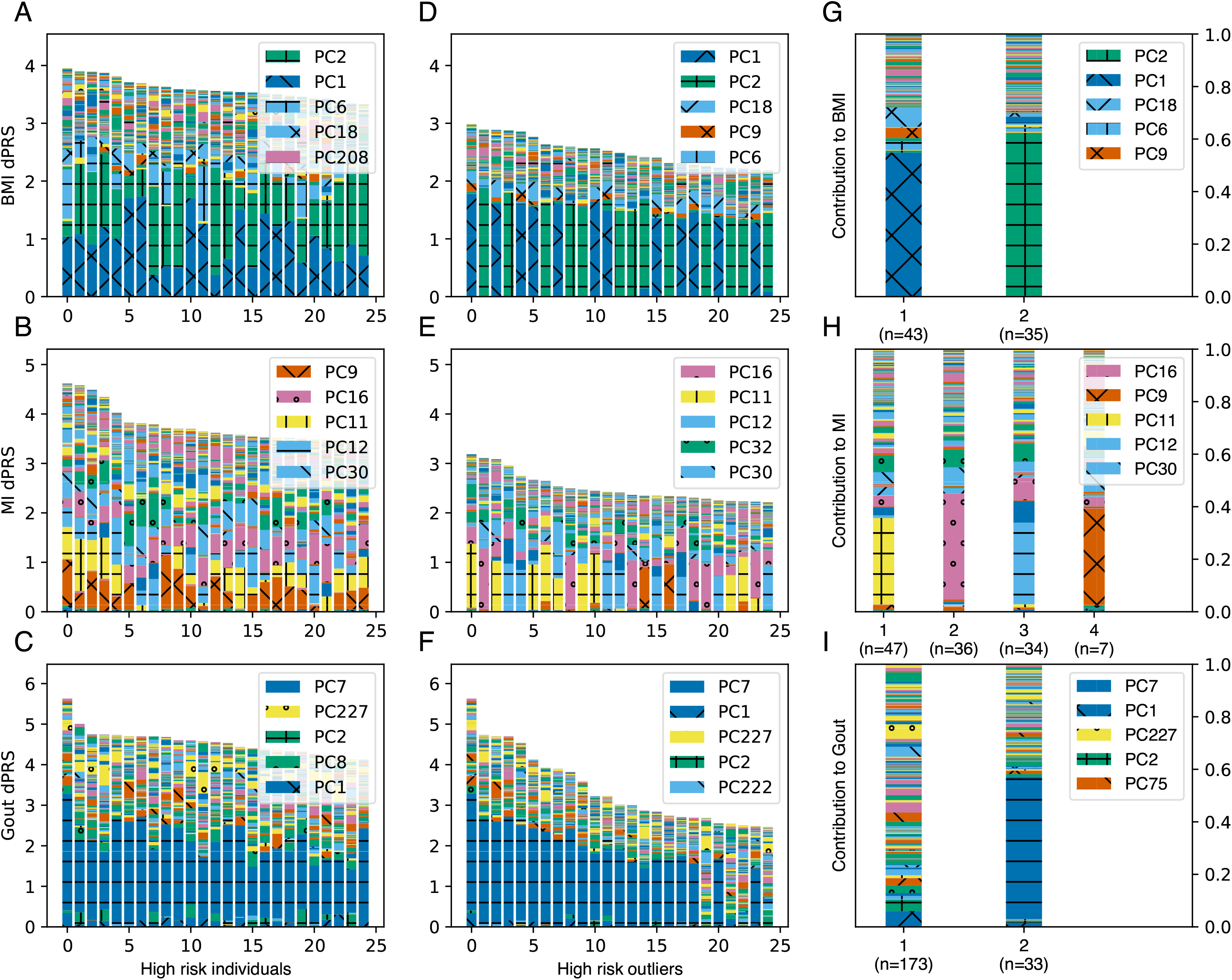
Painting components of genetic risk. (A-C) Component-painted risk for the 25 individuals or (D-F) outliers with highest dPRS for each trait in the test set. Each bar represents one individual; the height of the bar is the covariate-adjusted dPRS, and the colored components of the plot are the individual’s DeGAs risk profile, scaled to fit bar height. Colors for the five most represented components in each box are shown in its legend in rank order. (G-I) Mean DeGAs risk profiles from k-means clustering of high-risk outlier risk profiles, annotated with cluster size (n). Phenotype groups for selected components in this figure include: PC1 (Fat free mass); PC2 (Fat mass); PC7 (Alcohol use); PC9 (leukocytes and viral antigens); PC11 (Lung function); PC12 (Aspirin and cholesterol medication); PC16 (Blood pressure medication); PC32 (Hearing, ibuprofen, and cholesterol medication);

To better describe genetic diversity among outlying individuals, we attempted to identify genetic subtypes of each example trait in the high-risk outlier population. We performed a *k*-means clustering of this group using DeGAs risk profiles as the input; *k* was chosen by optimizing the gap-star statistic across an array of potential values (**Methods**). We described each cluster using its mean risk profile (**Figure 4a-c**) and noticed that cluster membership divides individuals based on cPRS for relevant components (**Figure 4d-f**).

For body mass index, we identify two risk clusters (**Figure 4g**): one driven by the fat mass component (PC2; 59.0%, *n*=43) and the other by the fat-free mass component (PC1; 70.4%, *n*=35). Some outlying individuals at risk for high BMI have genetic contribution from the near exclusively fat-related component (PC2), hence their deviation from “typical”. However, other individuals are outlying due to contribution from the lean mass component (PC1). Genetic risk from this cluster comes mainly from variant loadings related to fat-free mass-related traits like whole-body water and fat-free mass. That this cluster is distinct from other outliers at risk for high BMI implies relevant differences between individuals, which may suggest alternative preventative and therapeutic approaches across groups.

We also find five clusters of risk for myocardial infarction, four of which are driven primarily by components which were identified as important via the phenotype cosine score (**Figure 3**). These were PC11 (lung function; 34.8%; *n*=47), PC12 (high cholesterol; 32.9%; *n*=33), PC16 (blood pressure; 40.2%; *n*=31), and PC32 (hearing and cholesterol; 27.0%; *n=*6), all of which have additional contribution from medications commonly used for conditions comorbid with MI (**Figure 4h**). The fifth cluster is driven primarily by PC9 (37.0%; *n*=7), which has high phenotype contribution from leukocyte measures, vitamin B9, and an array of viral antigens. Its genetic contribution is primarily from variants proximal to the HLA genes, and other genes in *6p21.3* like the butrophylin-like protein *BTNL2* and the testis sperm-binding protein *TSBP1*. Though these clusters could offer therapeutic insights for MI, the components are less clear to interpret than those underlying risk for BMI.

We find two clusters of outliers for gout (**Figure 4i**): one is driven by the alcohol trait component (PC7; 54.3%; *n*=33), and the other has a profile which does not have a single dominant component, and instead is driven by several (PC1, PC2, PC227; 6.0, 5.4, 5.5%; *n*=173). The cluster of outliers with risk driven by PC7 is not surprising, as the component is identified as important for gout by its trait cosine score. Furthermore, genetic variation in *ADH1B* (one of the key genes for PC7) has been associated with gout in prior study^43^, suggesting there may be shared genetic risk between both traits. The other cluster is harder to interpret, due to the number of relevant components. Several components related to BMI (PC1, PC2, PC227) are highly represented in the mean DeGAs risk profile, and high BMI is a risk factor for gout^44,45^, but the data are not conclusive enough for definitive interpretation. Instead, we note challenges in interpreting components as a limitation of our approach.

## Discussion

In this study, we describe a novel technique to model polygenic traits using components of genetic associations. We build an example model using data from unrelated white British individuals in the UK Biobank to show that our method adds an interpretable dimension to traditional polygenic risk models by expressing disease, lifestyle, and biomarker-level elements in trait-related genetic components. Predicting genetic risk with these components led us to infer disease pathology beyond variant-trait associations without loss of predictive power from reducing model rank (**Figure S2**).

For three phenotypes of interest (BMI, MI, and gout), we showed that the DeGAs risk profile offers meaningful insight into the genetic drivers of trait risk for an individual. We then used this measure to identify clusters of high-risk individuals who share similar genetic risk profiles for each of the traits. We find, as in previous work^10^, that genetic risk for BMI can be decomposed into fat-mass and fat-free mass related components. We also show that while many individuals have risk for BMI driven by a combination of the two components, there exist “outlier” individuals who have strong contributions from only one of them. Our results further indicate that this diversity of contributory genetic risk is not limited to BMI. However, extracting biological insights for other traits will likely require deeper phenotyping, or other rich resources like single cell data.

We further demonstrated the generalizability of dPRS by assessing its performance in independent test sets of white British and non-British white individuals (**Figure S4**; all traits in **Data S1**) from the UK Biobank. Among non-British whites, the top 2% of dPRS carries OR=1.9 for MI and 5.1 for gout (**Figure S4**), compared to 1.7 and 2.5 in the test set individuals (**Figure 2**). Likewise, the top 2% of dPRS risk has 1.63 kg/m^2^ higher BMI in non-British whites (1.40 kg/m^2^) in the test set. Though we find similar performance for these traits across these two groups, concerns about the generalizability of traditional clump-and-threshold PRS across groups also apply to dPRS. Though methods exist to identify suspected causal variants via fine-mapping, we decided to LD-prune variants prior to analysis with DeGAs. One benefit of this approach is that it is agnostic to fine-scale patterns of association within LD blocks, which avoids the problem of having distinct (but highly correlated) causal variants across traits. However, LD pruning may leave dPRS slightly more vulnerable to overfitting patterns of LD in the GWAS population compared to approaches which use fine mapped variants. This may be worth revisiting in future work.

We also note that our analysis of subtypes may not be robust to different choices of input traits or study population. Taking gout as an example, our study finds two clusters of outliers (**Figure 4c**), one of which is due to a component related to a clinical risk factor for the trait (namely, alcohol use). Our ability to identify such clusters is clearly limited by the inclusion or exclusion of related traits and their degree of correlation in our analysis cohort. The components of genetic associations which can be identified by DeGAs also depend on trait selection. Here, we excluded traits which may have noisy or confounded patterns of genetic associations: specifically, rare conditions (*n* < 1000 in the UK Biobank) or traits which correlate with social measures like socioeconomic status. In future work, careful selection and curation of phenotypes may provide further insights than those we offer in this study. We encourage replication efforts using similar methods and have made all DeGAs risk models from this work available on the Global Biobank Engine^18^ (**Web Resources**).

Looking forward, we anticipate many potential applications of component-aware polygenic risk models like dPRS. Heritable conditions with known or putative biomarkers would be good candidates for follow-up studies that jointly investigate an outcome with its related features. For example, brain and liver images, metabolomics, and serum and urine biomarkers have been collected in resources like the UK Biobank, and may be of interest for future work. Since DeGAs requires only summary-level data, it is possible to build a component model of genetic risk in one cohort (or across several) and use it to estimate genetic risk and identify trait subtypes in another. Such analyses will help elucidate the diversity of polygenic risk for complex traits across individuals and populations.

## Supporting information

Data S1: Performance Summary

Data S2: Component annotation (phenotypes)

Data S3: Component annotation (genes)

## Acknowledgements

This research has been conducted using the UK Biobank Resource under Application Number 24983, “Generating effective therapeutic hypotheses from genomic and hospital linkage data” (http://www.ukbiobank.ac.uk/wp-content/uploads/2017/06/24983-Dr-Manuel-Rivas.pdf). Based on the information provided in Protocol 44532 the Stanford IRB has determined that the research does not involve human subjects as defined in 45 CFR 46.102(f) or 21 CFR 50.3(g). All participants in the UK Biobank provided written informed consent (more information is available at https://www.ukbiobank.ac.uk/2018/02/gdpr/). We thank all the participants in the UK Biobank study.

Some of the computing for this project was performed on the Sherlock cluster. We would like to thank Stanford University and the Stanford Research Computing Center for providing computational resources and support that contributed to these research results.

Figure 1 image credit: VectorStock.com/1143365.

## Author Contributions

M.A.R. conceived and designed the study. M.A. and M.A.R. carried out statistical and computational analyses. M.A., Y.T., G.R.V., and M.A.R. carried out quality control of the data. R.T. and T.H. aided in statistical design and conception. We thank Johanne Justesen, Michael Wainberg, and members of the Rivas Lab for comments on the manuscript. The manuscript was written by M.A. and M.A.R. and revised by all the co-authors. All co-authors have approved of the final version of the manuscript.

## Web Resources

Supplemental data, including weights for the final DeGAs model, are available on the Global Biobank Engine^18^: https://biobankengine.stanford.edu/downloads

## References

1. GBD 2017 Disease and Injury Incidence and Prevalence Collaborators. Global, regional, and national incidence, prevalence, and years lived with disability for 354 diseases and injuries for 195 countries and territories, 1990-2017: a systematic analysis for the Global Burden of Disease Study 2017. Lancet 2018; 392: 1789–1858.

2. Fritsche LG, Gruber SB, Wu Z et al. Association of Polygenic Risk Scores for Multiple Cancers in a Phenome-wide Study: Results from The Michigan Genomics Initiative. Am J Hum Genet 2018; 102: 1048–1061.

3. Läll K, Mägi R, Morris A, Metspalu A, Fischer K. Personalized risk prediction for type 2 diabetes: the potential of genetic risk scores. Genet Med 2016; 19: 322.

4. Khera AV, Chaffin M, Zekavat SM et al. Whole-Genome Sequencing to Characterize Monogenic and Polygenic Contributions in Patients Hospitalized With Early-Onset Myocardial Infarction. Circulation 2019; 139: 1593–1602.

5. Belsky DW, Moffitt TE, Sugden K et al. Development and evaluation of a genetic risk score for obesity. Biodemography Soc Biol 2013; 59: 85–100.

6. Euesden J, Lewis CM, O’Reilly PF. PRSice: Polygenic Risk Score software. Bioinformatics 2015; 31: 1466–1468.

7. Vilhjálmsson BJ, Yang J, Finucane HK et al. Modeling Linkage Disequilibrium Increases Accuracy of Polygenic Risk Scores. Am J Hum Genet 2015; 97: 576–592.

8. Qian J, Du W, Tanigawa Y et al. A Fast and Flexible Algorithm for Solving the Lasso in Large-scale and Ultrahigh-dimensional Problems. bioRxiv. 2019; 630079.

9. McCarthy MI. Painting a new picture of personalised medicine for diabetes. Diabetologia 2017; 60: 793–799.

10. Tanigawa Y, Li J, Justesen JM et al. Components of genetic associations across 2,138 phenotypes in the UK Biobank highlight novel adipocyte biology. Nat Commun 2019; 10: 2064.

11. Bycroft C, Freeman C, Petkova D et al. The UK Biobank resource with deep phenotyping and genomic data. Nature 2018; 562: 203–209.

12. Fry A, Littlejohns TJ, Sudlow C et al. Comparison of Sociodemographic and Health-Related Characteristics of UK Biobank Participants With Those of the General Population. Am J Epidemiol 2017; 186: 1026–1034.

13. Sinnott-Armstrong N, Tanigawa Y, Amar D et al. Genetics of 38 blood and urine biomarkers in the UK Biobank. doi:10.1101/660506.

14. Chang CC, Chow CC, Tellier LC, Vattikuti S, Purcell SM, Lee JJ. Second-generation PLINK: rising to the challenge of larger and richer datasets. Gigascience 2015; 4: 7.

15. Aguirre M, Rivas MA, Priest J. Phenome-wide Burden of Copy-Number Variation in the UK Biobank. Am J Hum Genet 2019; 105: 373–383.

16. DeBoever C, Tanigawa Y, Lindholm ME et al. Medical relevance of protein-truncating variants across 337,205 individuals in the UK Biobank study. Nat Commun 2018; 9: 1612.

17. DeBoever C, Tanigawa Y, Aguirre M, McInnes G, Lavertu A, Rivas MA. Assessing digital phenotyping to enhance genetic studies of human diseases. Am J Hum Genet 2020; 106: 611–622.

18. McInnes G, Tanigawa Y, DeBoever C et al. Global Biobank Engine: enabling genotype-phenotype browsing for biobank summary statistics. Bioinformatics 2018. doi:10.1093/bioinformatics/bty999.

19. Pedregosa F, Varoquaux G, Gramfort A et al. Scikit-learn: Machine Learning in Python. J Mach Learn Res 2011; 12: 2825–2830.

20. Halko N, Martinsson PG, Tropp JA. Finding Structure with Randomness: Probabilistic Algorithms for Constructing Approximate Matrix Decompositions. SIAM Review. 2011; 53: 217–288.

21. Tibshirani R, Walther G, Hastie T. Estimating the number of clusters in a data set via the gap statistic. Journal of the Royal Statistical Society: Series B (Statistical Methodology). 2001; 63: 411–423.

22. Mohajer M, Englmeier K-H, Schmid VJ. A comparison of Gap statistic definitions with and without logarithm function. 2011. http://arxiv.org/abs/1103.4767 (accessed 25May2020).

23. Locke AE, Kahali B, Berndt SI et al. Genetic studies of body mass index yield new insights for obesity biology. Nature 2015; 518: 197–206.

24. Kichaev G, Bhatia G, Loh P-R et al. Leveraging Polygenic Functional Enrichment to Improve GWAS Power. Am J Hum Genet 2019; 104: 65–75.

25. Hernandez Cordero AI, Gonzales NM, Parker CC et al. Genome-wide Associations Reveal Human-Mouse Genetic Convergence and Modifiers of Myogenesis, CPNE1 and STC2. Am J Hum Genet 2019; 105: 1222–1236.

26. Chang AC-M, Hook J, Lemckert FA et al. The murine stanniocalcin 2 gene is a negative regulator of postnatal growth. Endocrinology 2008; 149: 2403–2410.

27. Xu B, Goulding EH, Zang K et al. Brain-derived neurotrophic factor regulates energy balance downstream of melanocortin-4 receptor. Nat Neurosci 2003; 6: 736–742.

28. Tao Y-X. Molecular mechanisms of the neural melanocortin receptor dysfunction in severe early onset obesity. Mol Cell Endocrinol 2005; 239: 1–14.

29. Shungin D, Winkler TW, Croteau-Chonka DC et al. New genetic loci link adipose and insulin biology to body fat distribution. Nature 2015; 518: 187–196.

30. Frayling TM, Timpson NJ, Weedon MN et al. A common variant in the FTO gene is associated with body mass index and predisposes to childhood and adult obesity. Science 2007; 316: 889–894.

31. Weedon MN, Lango H, Lindgren CM et al. Genome-wide association analysis identifies 20 loci that influence adult height. Nat Genet 2008; 40: 575–583.

32. Claussnitzer M, Dankel SN, Kim K-H et al. FTO Obesity Variant Circuitry and Adipocyte Browning in Humans. N Engl J Med 2015; 373: 895–907.

33. Liu Y, Corcoran M, Rasool O et al. Cloning of two candidate tumor suppressor genes within a 10 kb region on chromosome 13q14, frequently deleted in chronic lymphocytic leukemia. Oncogene 1997; 15: 2463–2473.

34. Paquette M, Bernard S, Baass A. SLC22A3 is associated with lipoprotein (a) concentration and cardiovascular disease in familial hypercholesterolemia. Clin Biochem 2019; 66: 44–48.

35. Gao PS, Mao XQ, Roberts MH et al. Variants of STAT6 (signal transducer and activator of transcription 6) in atopic asthma. J Med Genet 2000; 37: 380–382.

36. Jones AV, Tilley M, Gutteridge A et al. GWAS of self-reported mosquito bite size, itch intensity and attractiveness to mosquitoes implicates immune-related predisposition loci. Hum Mol Genet 2017; 26: 1391–1406.

37. Myocardial Infarction Genetics Consortium, Kathiresan S, Voight BF et al. Genome-wide association of early-onset myocardial infarction with single nucleotide polymorphisms and copy number variants. Nat Genet 2009; 41: 334–341.

38. Kuo C-F, Grainge MJ, See L-C et al. Familial aggregation of gout and relative genetic and environmental contributions: a nationwide population study in Taiwan. Ann Rheum Dis 2015; 74: 369–374.

39. Vitart V, Rudan I, Hayward C et al. SLC2A9 is a newly identified urate transporter influencing serum urate concentration, urate excretion and gout. Nat Genet 2008; 40: 437–442.

40. Tolstrup JS, Nordestgaard BG, Rasmussen S, Tybjaerg-Hansen A, Grønbaek M. Alcoholism and alcohol drinking habits predicted from alcohol dehydrogenase genes. Pharmacogenomics J 2008; 8: 220–227.

41. Wuttke M, Li Y, Li M et al. A catalog of genetic loci associated with kidney function from analyses of a million individuals. Nat Genet 2019; 51: 957–972.

42. Choi HK, Atkinson K, Karlson EW, Willett W, Curhan G. Alcohol intake and risk of incident gout in men: a prospective study. Lancet 2004; 363: 1277–1281.

43. Sakiyama M, Matsuo H, Akashi A et al. Independent effects of ADH1B and ALDH2 common dysfunctional variants on gout risk. Sci Rep 2017; 7: 2500.

44. Bhole V, de Vera M, Rahman MM, Krishnan E, Choi H. Epidemiology of gout in women: Fifty-two-year followup of a prospective cohort. Arthritis Rheum 2010; 62: 1069–1076.

45. Choi HK, Atkinson K, Karlson EW, Curhan G. Obesity, weight change, hypertension, diuretic use, and risk of gout in men: the health professionals follow-up study. Arch Intern Med 2005; 165: 742–748.

