## Supplementary material for "Polygenic risk modeling with latent trait-related genetic components": Data S2: Component annotation (phenotypes)

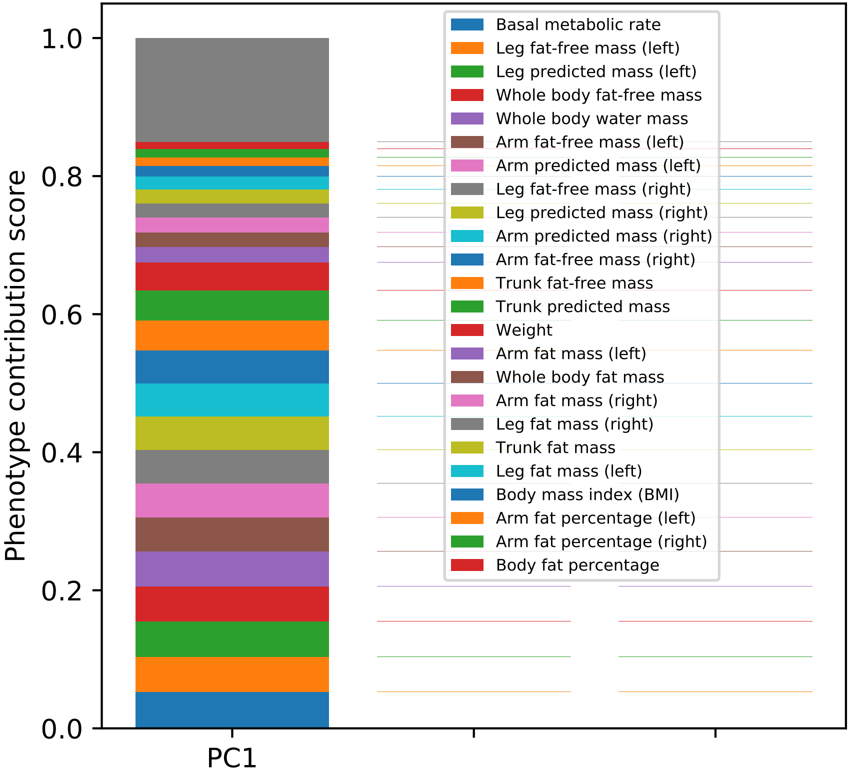

Phenotype contribution score
1.0
0.8
0.6
0.4
0.2
0.0
PC1
Basal metabolic rate
Leg fat-free mass (left)
Leg predicted mass (left)
Whole body fat-free mass
Whole body water mass
Arm fat-free mass (left)
Arm predicted mass (left)
Leg fat-free mass (right)
Leg predicted mass (right)
Arm predicted mass (right)
Arm fat-free mass (right)
Trunk fat-free mass
Trunk predicted mass
Weight
Arm fat mass (left)
Whole body fat mass
Arm fat mass (right)
Leg fat mass (right)
Trunk fat mass
Leg fat mass (left)
Body mass index (BMI)
Arm fat percentage (left)
Arm fat percentage (right)
Body fat percentage

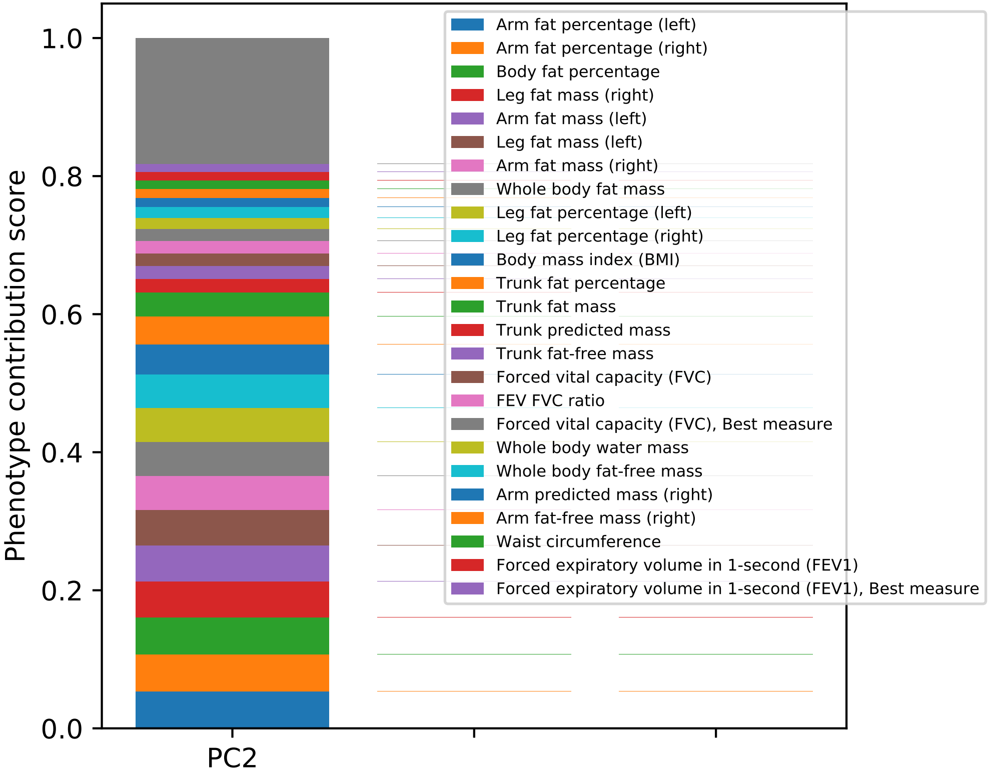

Phenotype contribution score
1.0
0.8
0.6
0.4
0.2
0.0
PC2
Arm fat percentage (left)
Arm fat percentage (right)
Body fat percentage
Leg fat mass (right)
Arm fat mass (left)
Leg fat mass (left)
Arm fat mass (right)
Whole body fat mass
Leg fat percentage (left)
Leg fat percentage (right)
Body mass index (BMI)
Trunk fat percentage
Trunk fat mass
Trunk predicted mass
Trunk fat-free mass
Forced vital capacity (FVC)
FEV FVC ratio
Forced vital capacity (FVC), Best measure
Whole body water mass
Whole body fat-free mass
Arm predicted mass (right)
Arm fat-free mass (right)
Waist circumference
Forced expiratory volume in 1-second (FEV1)
Forced expiratory volume in 1-second (FEV1), Best measure

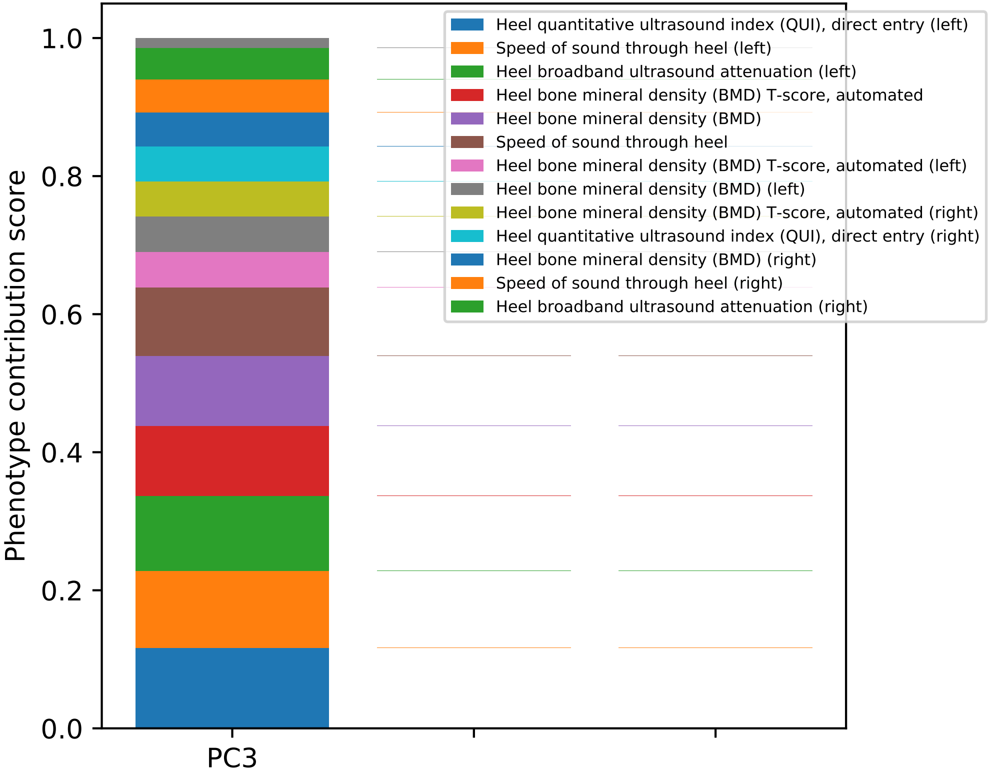

1.0
0.8
0.6
0.4
0.2
0.0
Phenotype contribution score
PC3
Heel quantitative ultrasound index (QUI), direct entry (left)
Speed of sound through heel (left)
Heel broadband ultrasound attenuation (left)
Heel bone mineral density (BMD) T-score, automated
Heel bone mineral density (BMD)
Speed of sound through heel
Heel bone mineral density (BMD) T-score, automated (left)
Heel bone mineral density (BMD) (left)
Heel bone mineral density (BMD) T-score, automated (right)
Heel quantitative ultrasound index (QUI), direct entry (right)
Heel bone mineral density (BMD) (right)
Speed of sound through heel (right)
Heel broadband ultrasound attenuation (right)

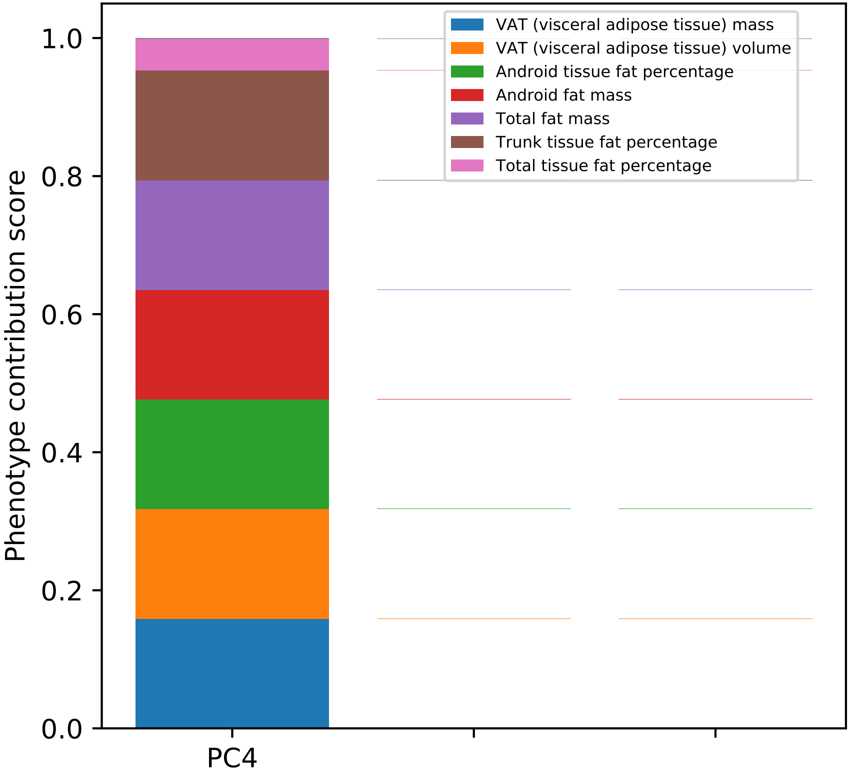

VAT (visceral adipose tissue) mass
VAT (visceral adipose tissue) volume
Android tissue fat percentage
Android fat mass
Total fat mass
Trunk tissue fat percentage
Total tissue fat percentage
Phenotype contribution score
1.0
0.8
0.6
0.4
0.2
0.0
PC4

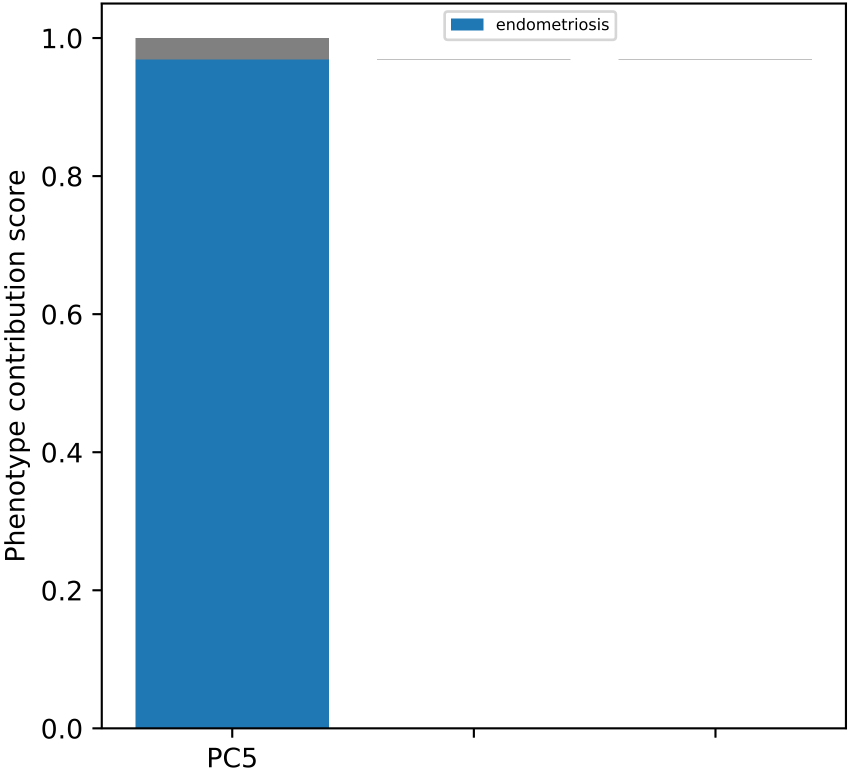
endometriosis
Phenotype contribution score
1.0
0.8
0.6
0.4
0.2
0.0
PC5

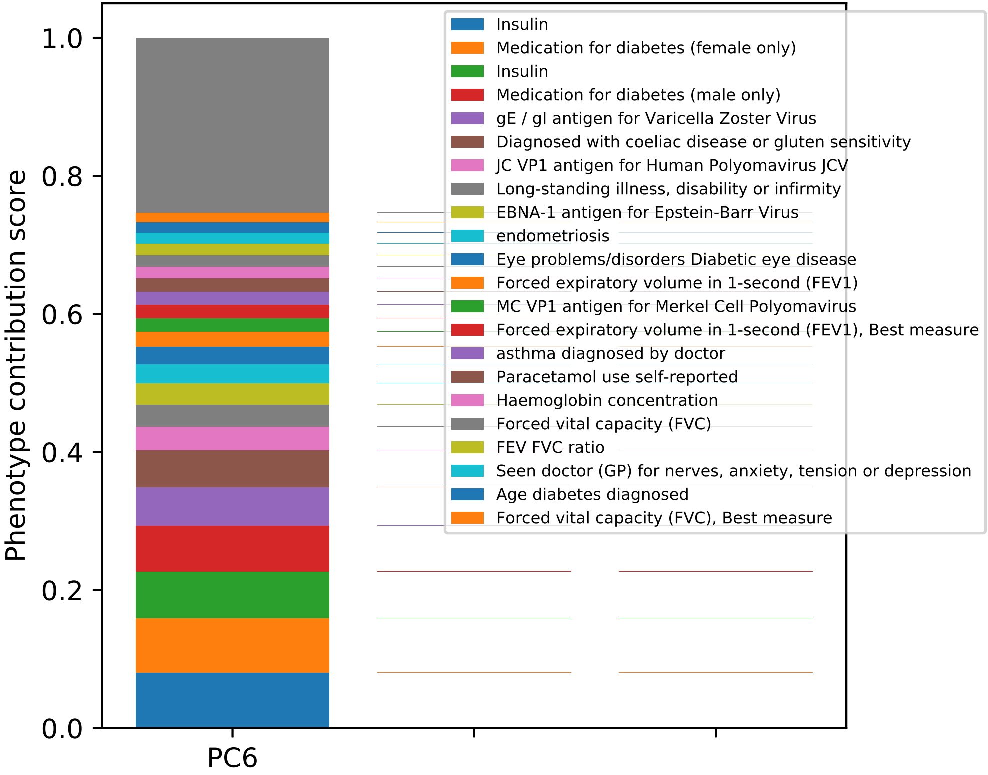
Phenotype contribution score
1.0
0.8
0.6
0.4
0.2
0.0
PC6
Insulin
Medication for diabetes (female only)
Insulin
Medication for diabetes (male only)
gE / gI antigen for Varicella Zoster Virus
Diagnosed with coeliac disease or gluten sensitivity
JC VP1 antigen for Human Polyomavirus JCV
Long-standing illness, disability or infirmity
EBNA-1 antigen for Epstein-Barr Virus
endometriosis
Eye problems/disorders Diabetic eye disease
Forced expiratory volume in 1-second (FEV1)
MC VP1 antigen for Merkel Cell Polyomavirus
Forced expiratory volume in 1-second (FEV1), Best measure
asthma diagnosed by doctor
Paracetamol use self-reported
Haemoglobin concentration
Forced vital capacity (FVC)
FEV FVC ratio
Seen doctor (GP) for nerves, anxiety, tension or depression
Age diabetes diagnosed
Forced vital capacity (FVC), Best measure

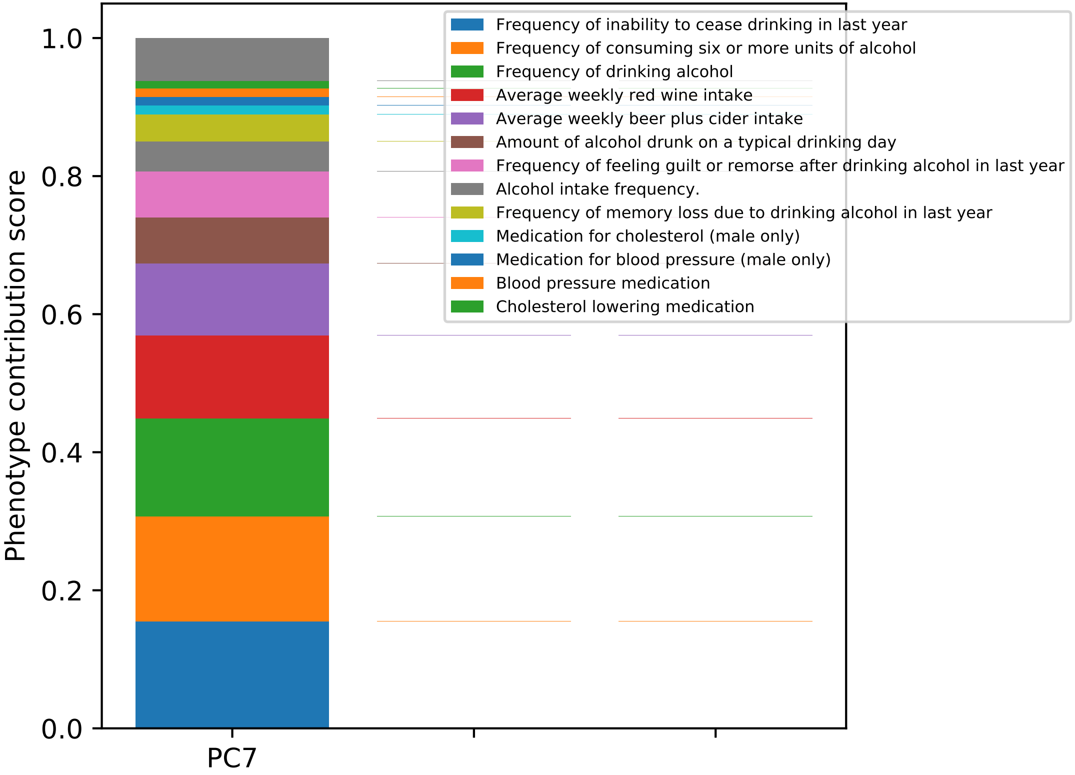
Phenotype contribution score
1.0
0.8
0.6
0.4
0.2
0.0
PC7
Frequency of inability to cease drinking in last year
Frequency of consuming six or more units of alcohol
Frequency of drinking alcohol
Average weekly red wine intake
Average weekly beer plus cider intake
Amount of alcohol drunk on a typical drinking day
Frequency of feeling guilt or remorse after drinking alcohol in last year
Alcohol intake frequency.
Frequency of memory loss due to drinking alcohol in last year
Medication for cholesterol (male only)
Medication for blood pressure (male only)
Blood pressure medication
Cholesterol lowering medication

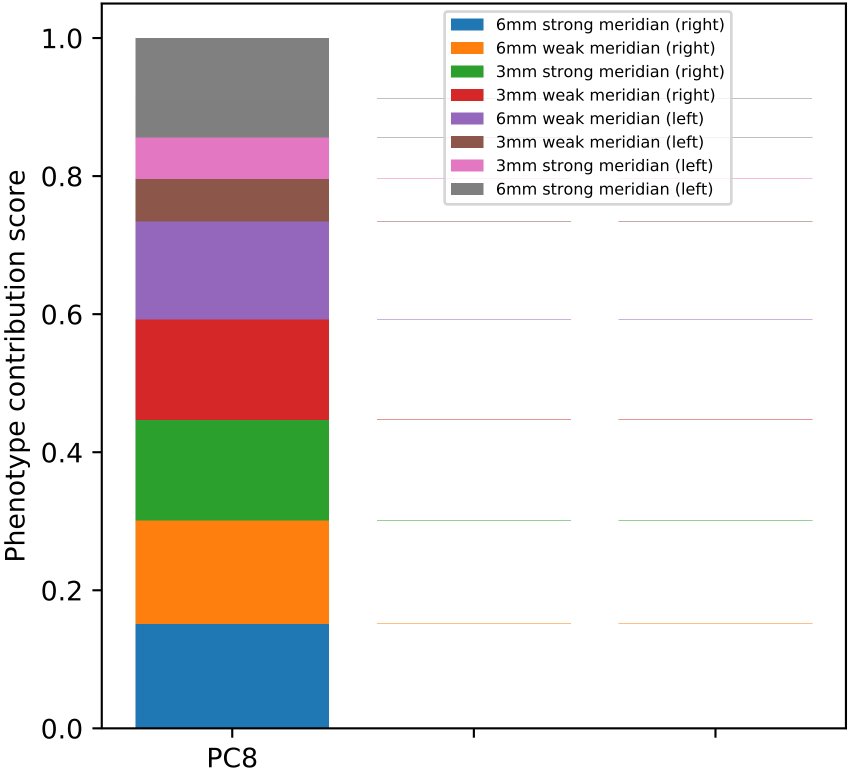

6mm strong meridian (right)
6mm weak meridian (right)
3mm strong meridian (right)
3mm weak meridian (right)
6mm weak meridian (left)
3mm weak meridian (left)
3mm strong meridian (left)
6mm strong meridian (left)
Phenotype contribution score
1.0
0.8
0.6
0.4
0.2
0.0
PC8

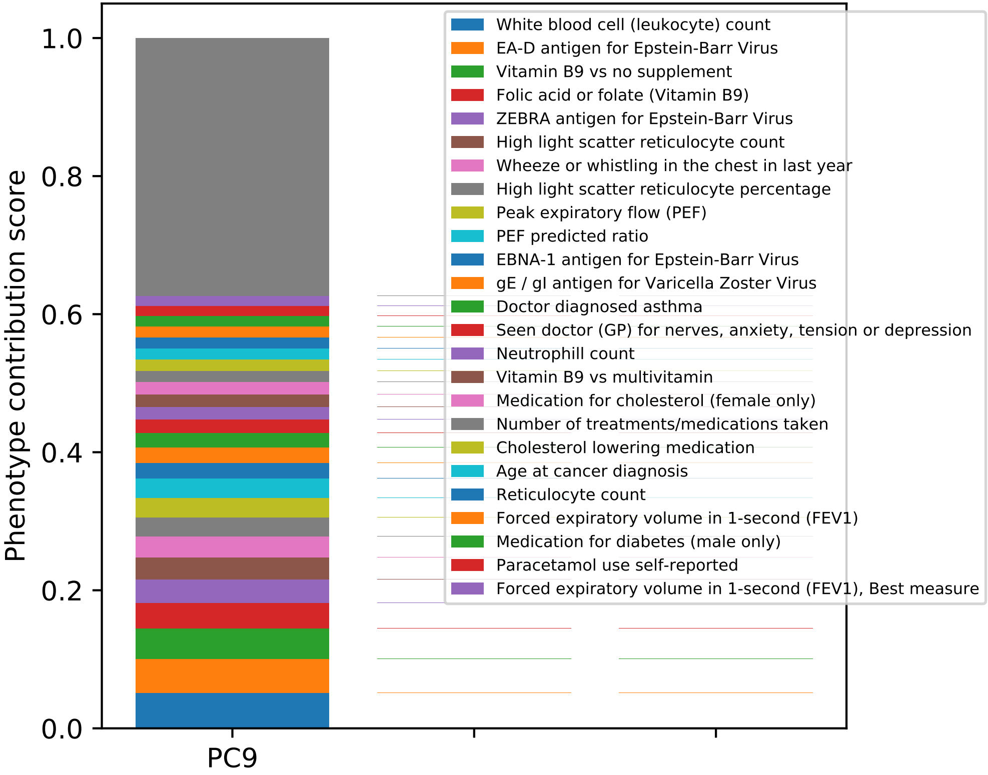

Phenotype contribution score
0.0
0.2
0.4
0.6
0.8
1.0
PC9
White blood cell (leukocyte) count
EA-D antigen for Epstein-Barr Virus
Vitamin B9 vs no supplement
Folic acid or folate (Vitamin B9)
ZEBRA antigen for Epstein-Barr Virus
High light scatter reticulocyte count
Wheeze or whistling in the chest in last year
High light scatter reticulocyte percentage
Peak expiratory flow (PEF)
PEF predicted ratio
EBNA-1 antigen for Epstein-Barr Virus
gE / gI antigen for Varicella Zoster Virus
Doctor diagnosed asthma
Seen doctor (GP) for nerves, anxiety, tension or depression
Neutrophill count
Vitamin B9 vs multivitamin
Medication for cholesterol (female only)
Number of treatments/medications taken
Cholesterol lowering medication
Age at cancer diagnosis
Reticulocyte count
Forced expiratory volume in 1-second (FEV1)
Medication for diabetes (male only)
Paracetamol use self-reported
Forced expiratory volume in 1-second (FEV1), Best measure

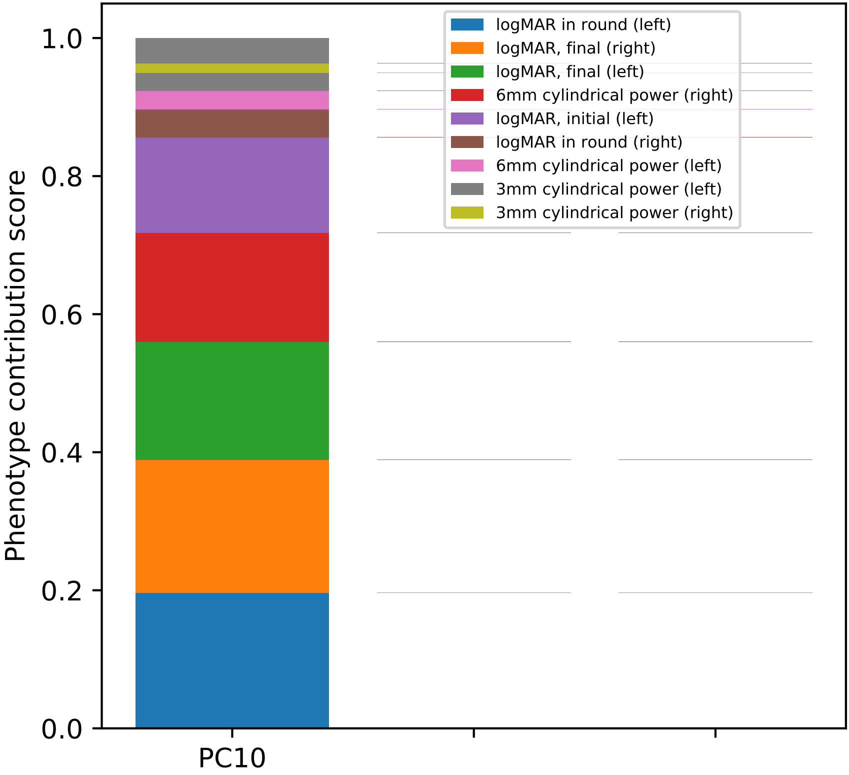

Phenotype contribution score
1.0
0.8
0.6
0.4
0.2
0.0
PC10
logMAR in round (left)
logMAR, final (right)
logMAR, final (left)
6mm cylindrical power (right)
logMAR, initial (left)
logMAR in round (right)
6mm cylindrical power (left)
3mm cylindrical power (left)
3mm cylindrical power (right)

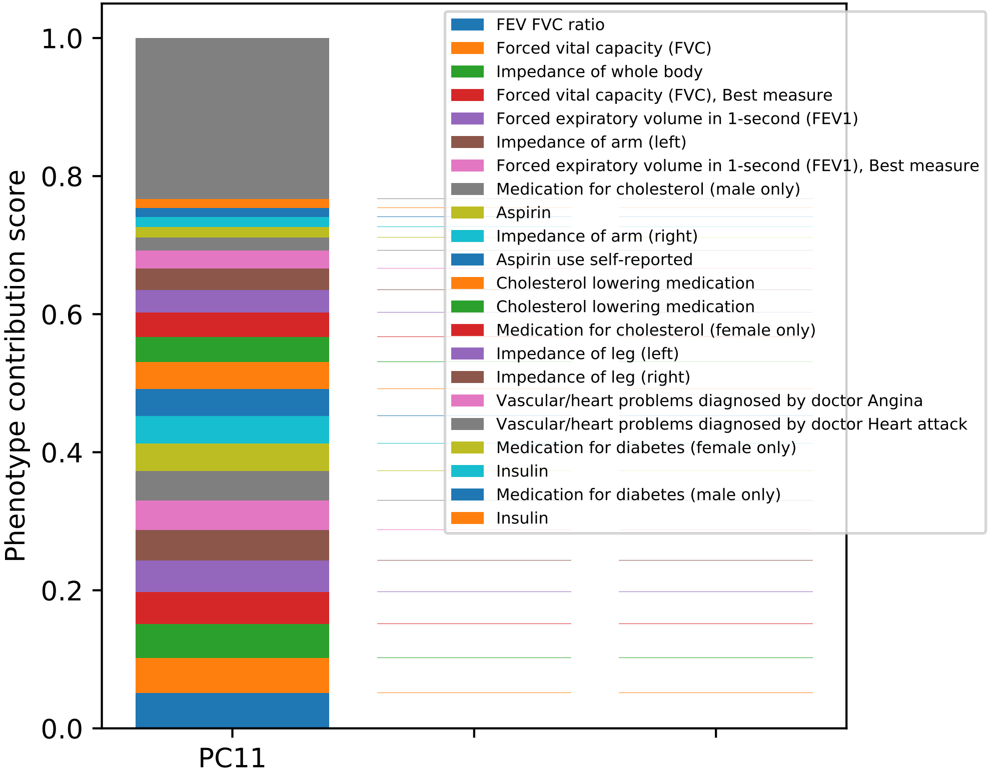
FEV FVC ratio
Forced vital capacity (FVC)
Impedance of whole body
Forced vital capacity (FVC), Best measure
Forced expiratory volume in 1-second (FEV1)
Impedance of arm (left)
Forced expiratory volume in 1-second (FEV1), Best measure
Medication for cholesterol (male only)
Aspirin
Impedance of arm (right)
Aspirin use self-reported
Cholesterol lowering medication
Cholesterol lowering medication
Medication for cholesterol (female only)
Impedance of leg (left)
Impedance of leg (right)
Vascular/heart problems diagnosed by doctor Angina
Vascular/heart problems diagnosed by doctor Heart attack
Medication for diabetes (female only)
Insulin
Medication for diabetes (male only)
Insulin
Phenotype contribution score
1.0
0.8
0.6
0.4
0.2
0.0
PC11

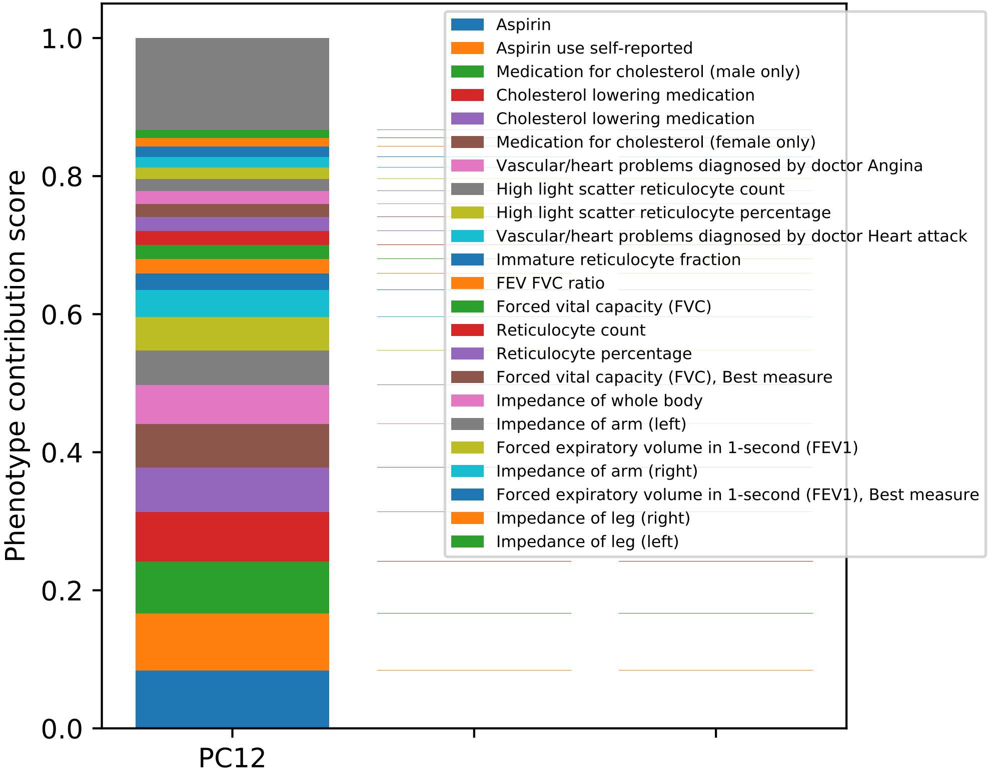

Phenotype contribution score
1.0
0.8
0.6
0.4
0.2
0.0
PC12
Aspirin
Aspirin use self-reported
Medication for cholesterol (male only)
Cholesterol lowering medication
Cholesterol lowering medication
Medication for cholesterol (female only)
Vascular/heart problems diagnosed by doctor Angina
High light scatter reticulocyte count
High light scatter reticulocyte percentage
Vascular/heart problems diagnosed by doctor Heart attack
Immature reticulocyte fraction
FEV FVC ratio
Forced vital capacity (FVC)
Reticulocyte count
Reticulocyte percentage
Forced vital capacity (FVC), Best measure
Impedance of whole body
Impedance of arm (left)
Forced expiratory volume in 1-second (FEV1)
Impedance of arm (right)
Forced expiratory volume in 1-second (FEV1), Best measure
Impedance of leg (right)
Impedance of leg (left)

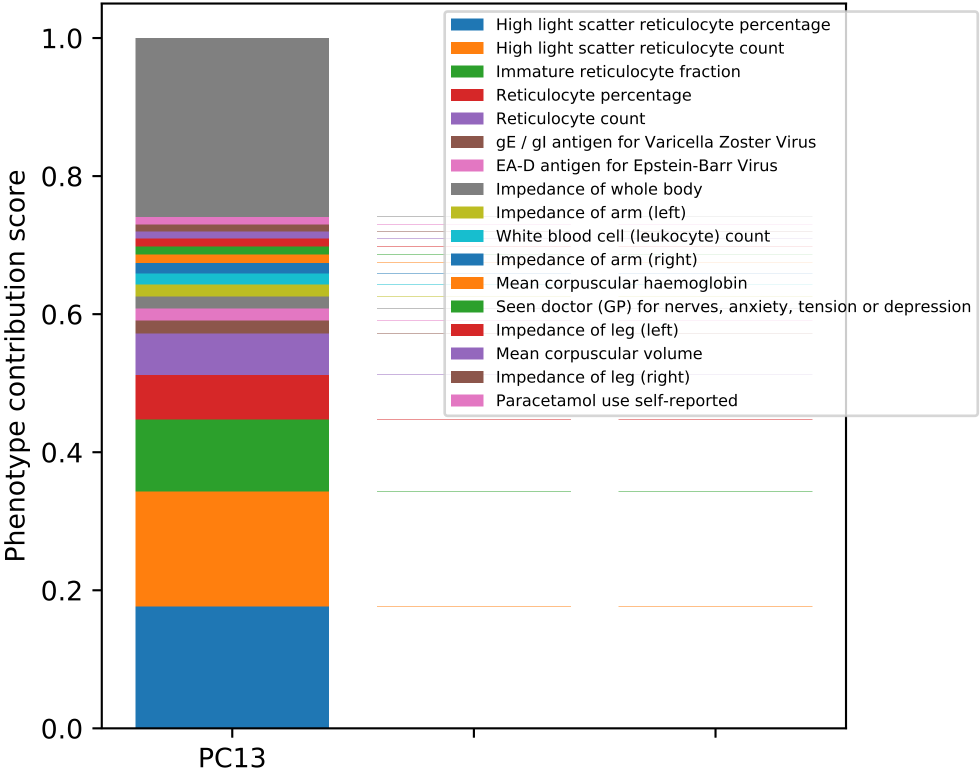
High light scatter reticulocyte percentage
High light scatter reticulocyte count
Immature reticulocyte fraction
Reticulocyte percentage
Reticulocyte count
gE / gI antigen for Varicella Zoster Virus
EA-D antigen for Epstein-Barr Virus
Impedance of whole body
Impedance of arm (left)
White blood cell (leukocyte) count
Impedance of arm (right)
Mean corpuscular haemoglobin
Seen doctor (GP) for nerves, anxiety, tension or depression
Impedance of leg (left)
Mean corpuscular volume
Impedance of leg (right)
Paracetamol use self-reported
Phenotype contribution score
1.0
0.8
0.6
0.4
0.2
0.0
PC13

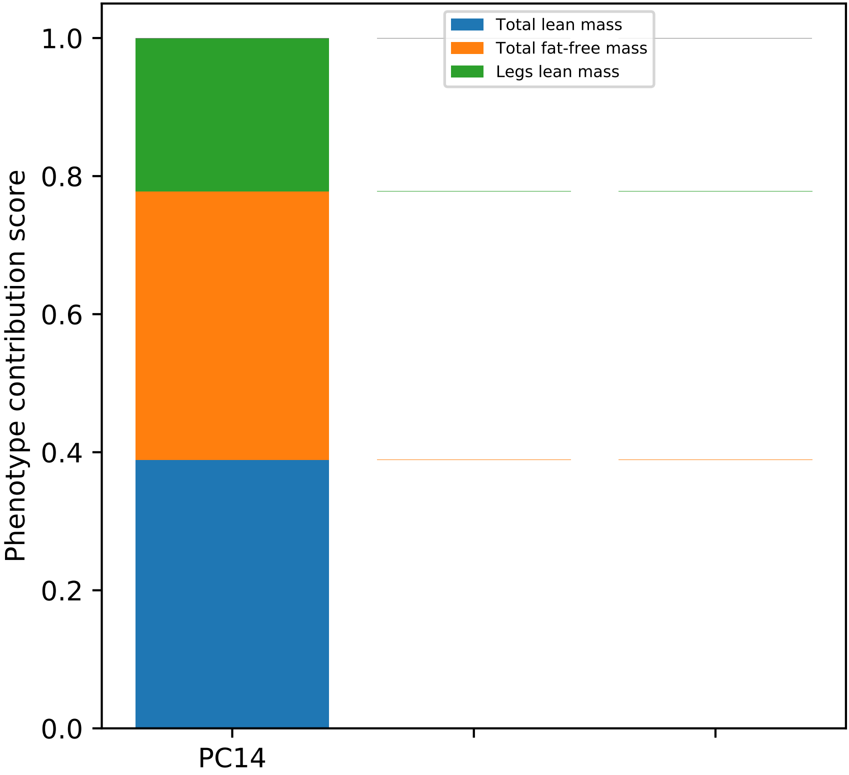
Total lean mass
Total fat-free mass
Legs lean mass
Phenotype contribution score
1.0
0.8
0.6
0.4
0.2
0.0
PC14

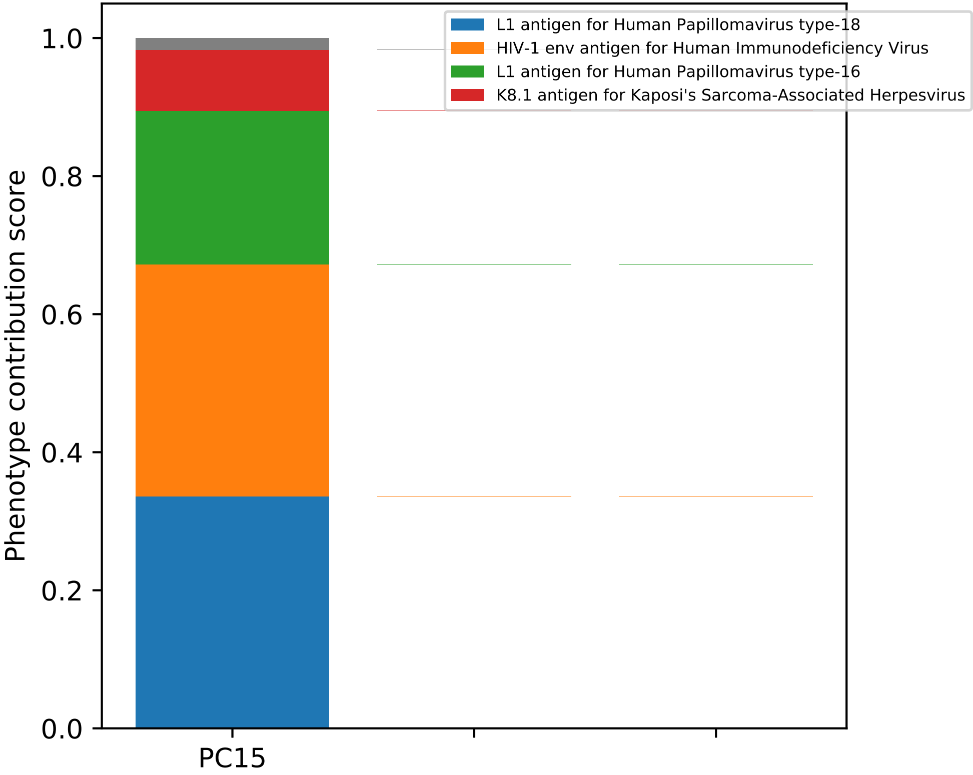

L1 antigen for Human Papillomavirus type-18
HIV-1 env antigen for Human Immunodeficiency Virus
L1 antigen for Human Papillomavirus type-16
K8.1 antigen for Kaposi's Sarcoma-Associated Herpesvirus
Phenotype contribution score
1.0
0.8
0.6
0.4
0.2
0.0
PC15

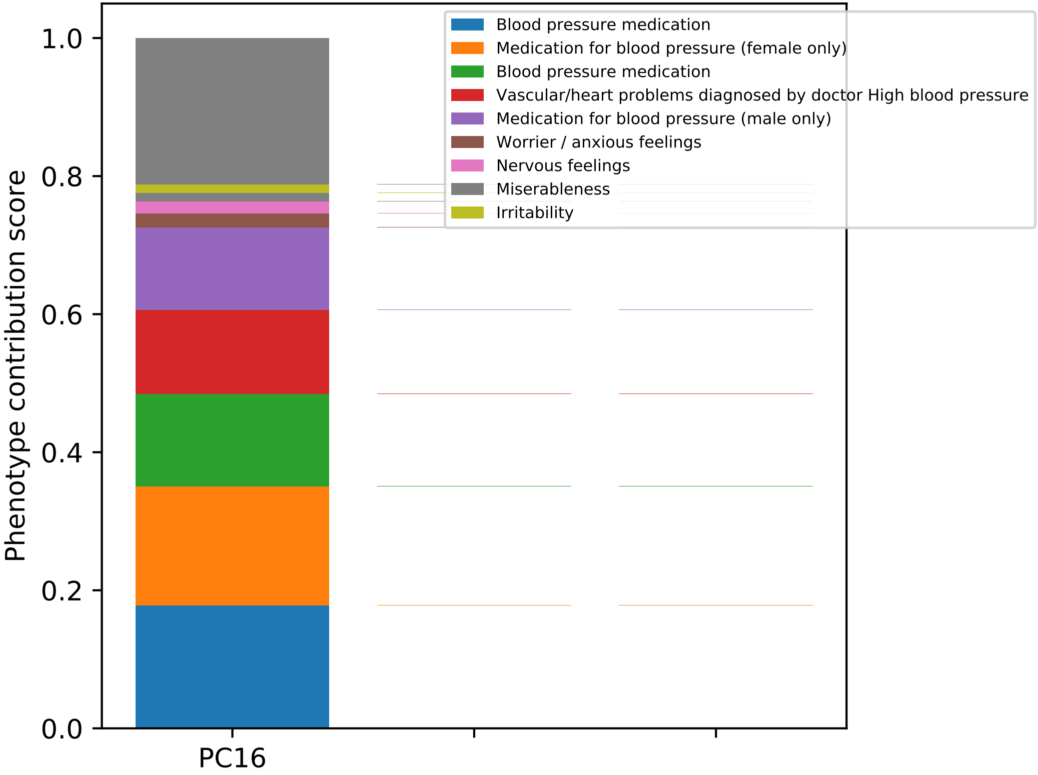

Phenotype contribution score
1.0
0.8
0.6
0.4
0.2
0.0
PC16
Blood pressure medication
Medication for blood pressure (female only)
Blood pressure medication
Vascular/heart problems diagnosed by doctor High blood pressure
Medication for blood pressure (male only)
Worrier / anxious feelings
Nervous feelings
Miserableness
Irritability

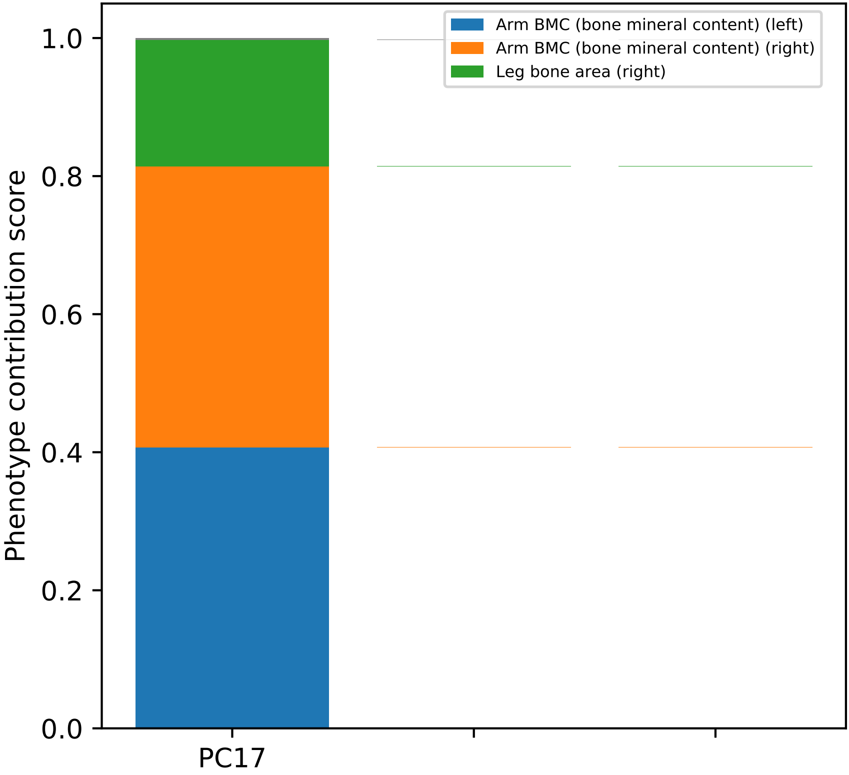
Arm BMC (bone mineral content) (left)
Arm BMC (bone mineral content) (right)
Leg bone area (right)
Phenotype contribution score
1.0
0.8
0.6
0.4
0.2
0.0
PC17

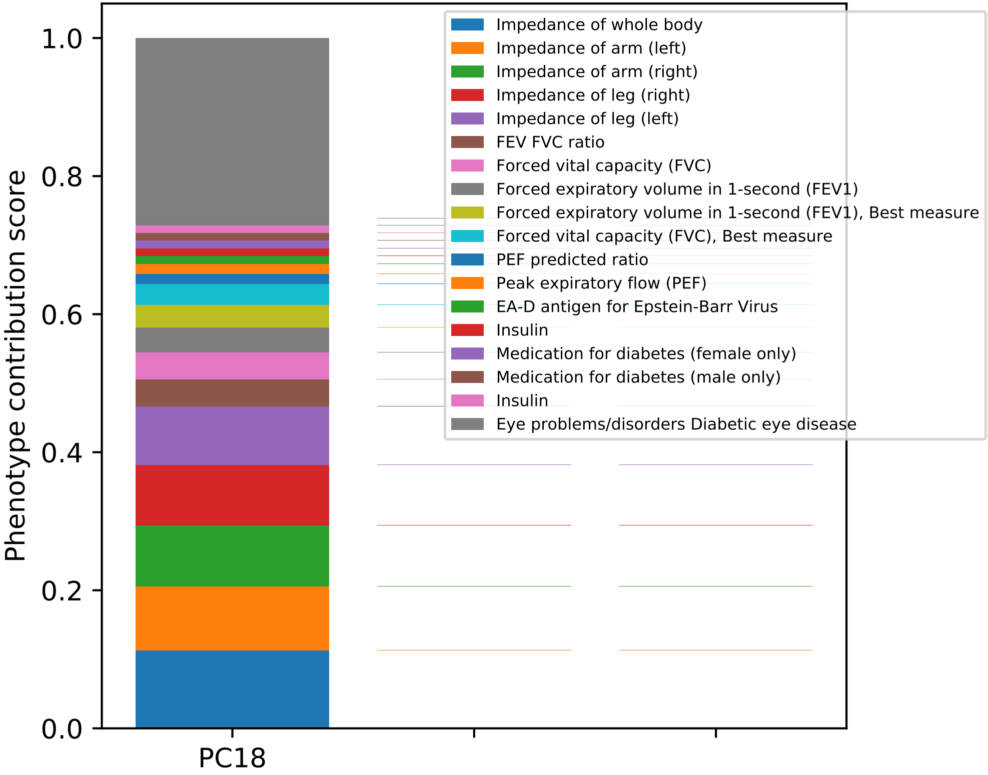

Phenotype contribution score
0.0
0.2
0.4
0.6
0.8
1.0
PC18
Impedance of whole body
Impedance of arm (left)
Impedance of arm (right)
Impedance of leg (right)
Impedance of leg (left)
FEV FVC ratio
Forced vital capacity (FVC)
Forced expiratory volume in 1-second (FEV1)
Forced expiratory volume in 1-second (FEV1), Best measure
Forced vital capacity (FVC), Best measure
PEF predicted ratio
Peak expiratory flow (PEF)
EA-D antigen for Epstein-Barr Virus
Insulin
Medication for diabetes (female only)
Medication for diabetes (male only)
Insulin
Eye problems/disorders Diabetic eye disease

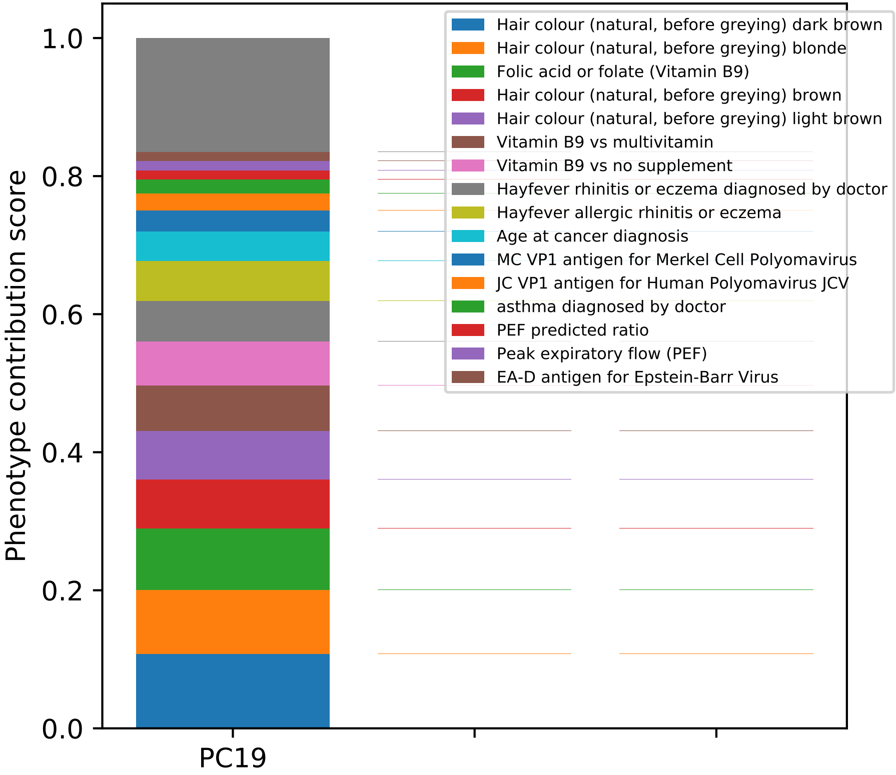

Phenotype contribution score
0.0
0.2
0.4
0.6
0.8
1.0
PC19
Hair colour (natural, before greying) dark brown
Hair colour (natural, before greying) blonde
Folic acid or folate (Vitamin B9)
Hair colour (natural, before greying) brown
Hair colour (natural, before greying) light brown
Vitamin B9 vs multivitamin
Vitamin B9 vs no supplement
Hayfever rhinitis or eczema diagnosed by doctor
Hayfever allergic rhinitis or eczema
Age at cancer diagnosis
MC VP1 antigen for Merkel Cell Polyomavirus
JC VP1 antigen for Human Polyomavirus JCV
asthma diagnosed by doctor
PEF predicted ratio
Peak expiratory flow (PEF)
EA-D antigen for Epstein-Barr Virus

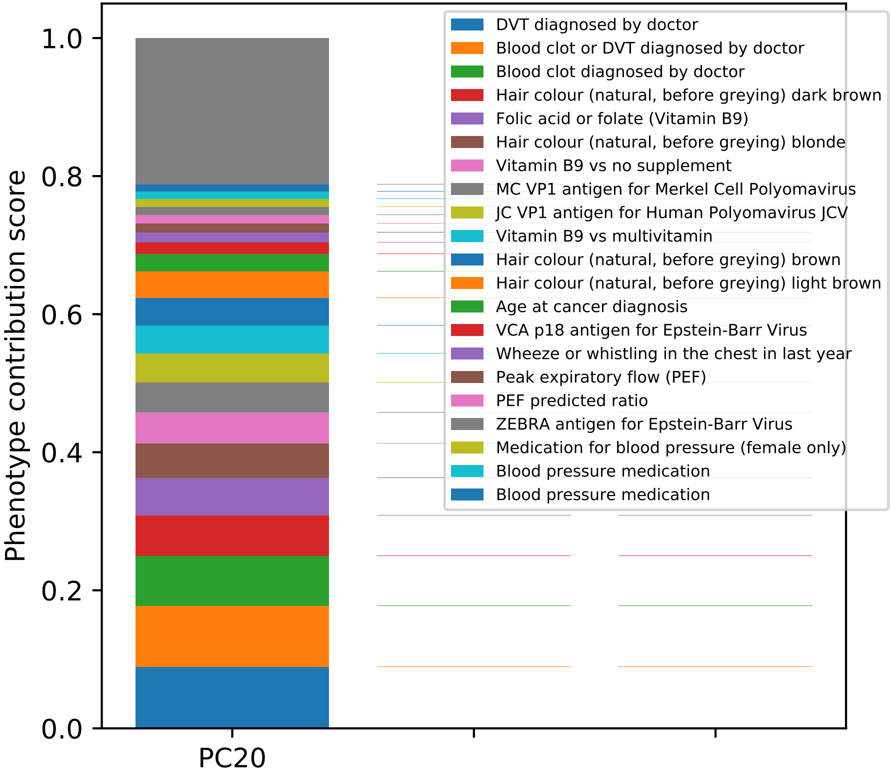

Phenotype contribution score
1.0
0.8
0.6
0.4
0.2
0.0
PC20
DVT diagnosed by doctor
Blood clot or DVT diagnosed by doctor
Blood clot diagnosed by doctor
Hair colour (natural, before greying) dark brown
Folic acid or folate (Vitamin B9)
Hair colour (natural, before greying) blonde
Vitamin B9 vs no supplement
MC VP1 antigen for Merkel Cell Polyomavirus
JC VP1 antigen for Human Polyomavirus JCV
Vitamin B9 vs multivitamin
Hair colour (natural, before greying) brown
Hair colour (natural, before greying) light brown
Age at cancer diagnosis
VCA p18 antigen for Epstein-Barr Virus
Wheeze or whistling in the chest in last year
Peak expiratory flow (PEF)
PEF predicted ratio
ZEBRA antigen for Epstein-Barr Virus
Medication for blood pressure (female only)
Blood pressure medication
Blood pressure medication

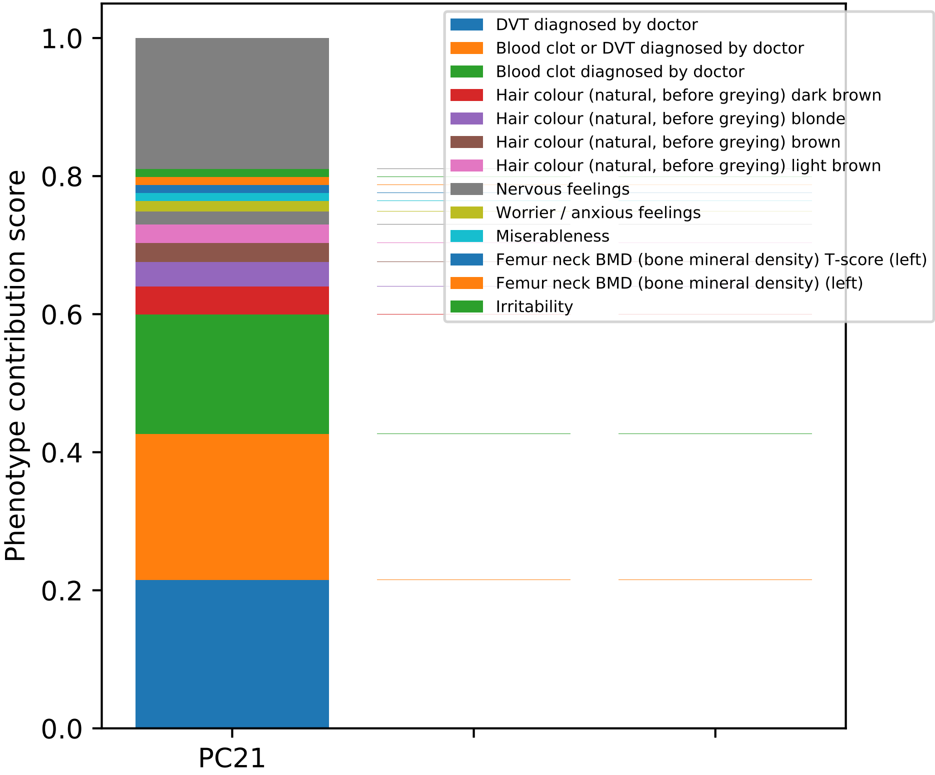
Phenotype contribution score
1.0
0.8
0.6
0.4
0.2
0.0
PC21
DVT diagnosed by doctor
Blood clot or DVT diagnosed by doctor
Blood clot diagnosed by doctor
Hair colour (natural, before greying) dark brown
Hair colour (natural, before greying) blonde
Hair colour (natural, before greying) brown
Hair colour (natural, before greying) light brown
Nervous feelings
Worrier / anxious feelings
Miserableness
Femur neck BMD (bone mineral density) T-score (left)
Femur neck BMD (bone mineral density) (left)
Irritability

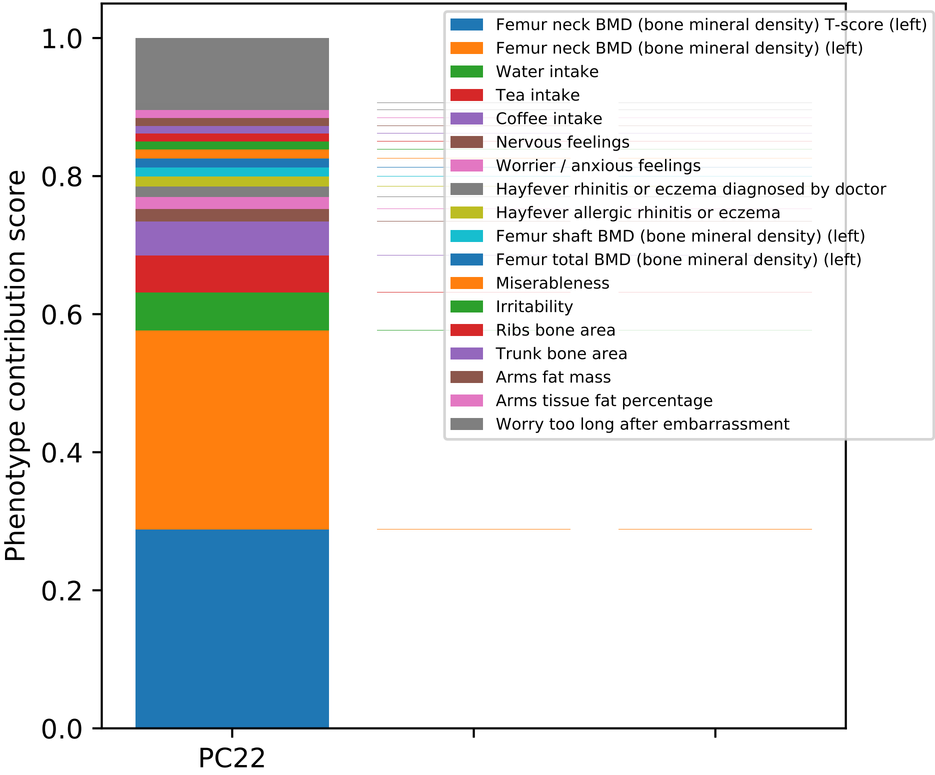

Phenotype contribution score
0.0
0.2
0.4
0.6
0.8
1.0
PC22
Femur neck BMD (bone mineral density) T-score (left)
Femur neck BMD (bone mineral density) (left)
Water intake
Tea intake
Coffee intake
Nervous feelings
Worrier / anxious feelings
Hayfever rhinitis or eczema diagnosed by doctor
Hayfever allergic rhinitis or eczema
Femur shaft BMD (bone mineral density) (left)
Femur total BMD (bone mineral density) (left)
Miserableness
Irritability
Ribs bone area
Trunk bone area
Arms fat mass
Arms tissue fat percentage
Worry too long after embarrassment

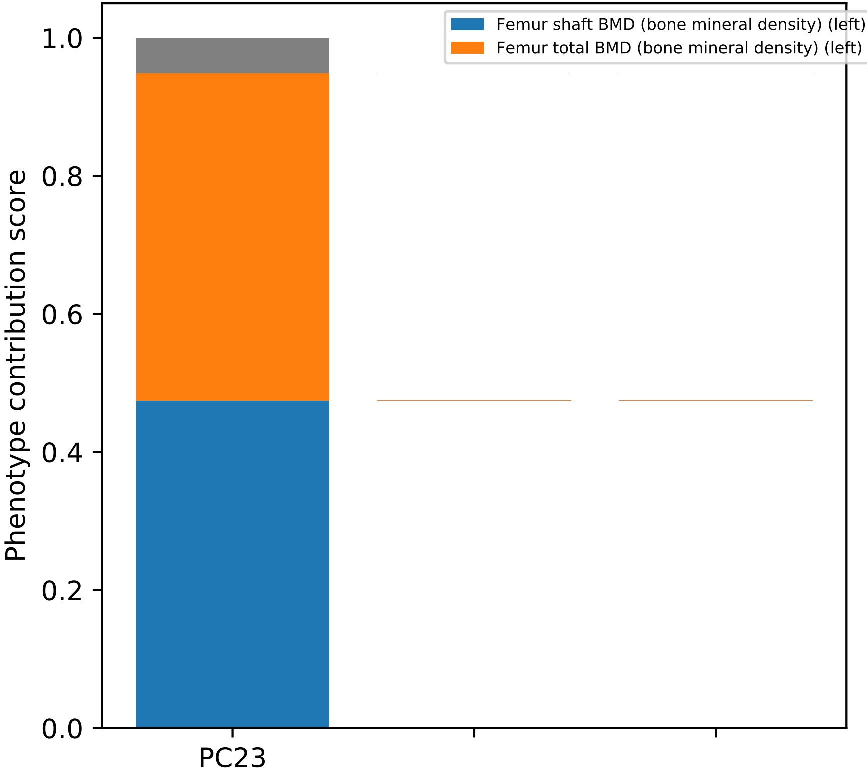

Femur shaft BMD (bone mineral density) (left)
Femur total BMD (bone mineral density) (left)
Phenotype contribution score
1.0
0.8
0.6
0.4
0.2
0.0
PC23

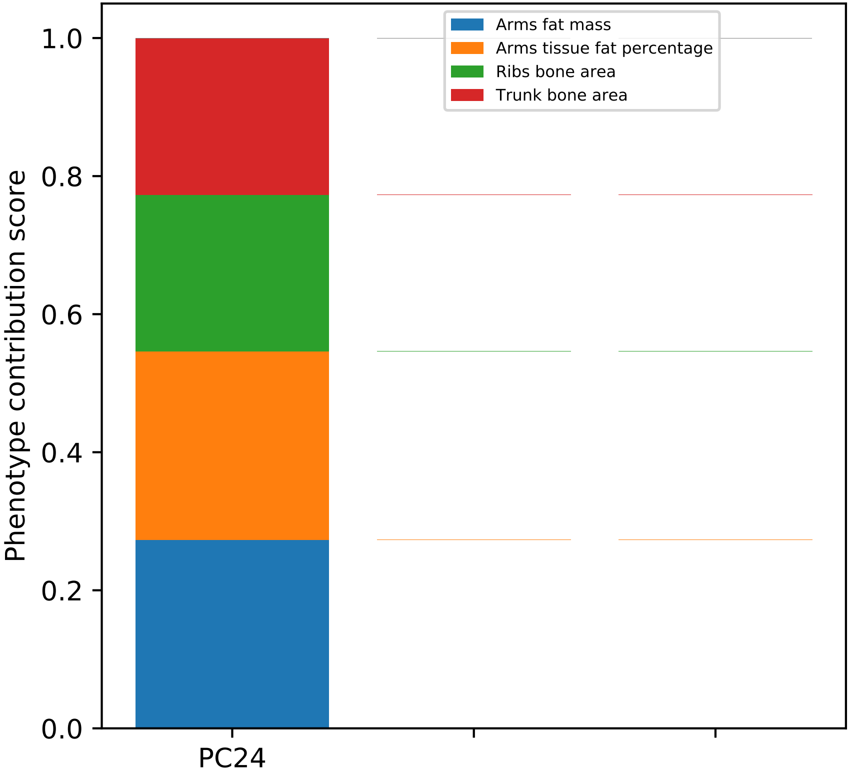
Arms fat mass
Arms tissue fat percentage
Ribs bone area
Trunk bone area
1.0
0.8
0.6
0.4
0.2
0.0
Phenotype contribution score
PC24

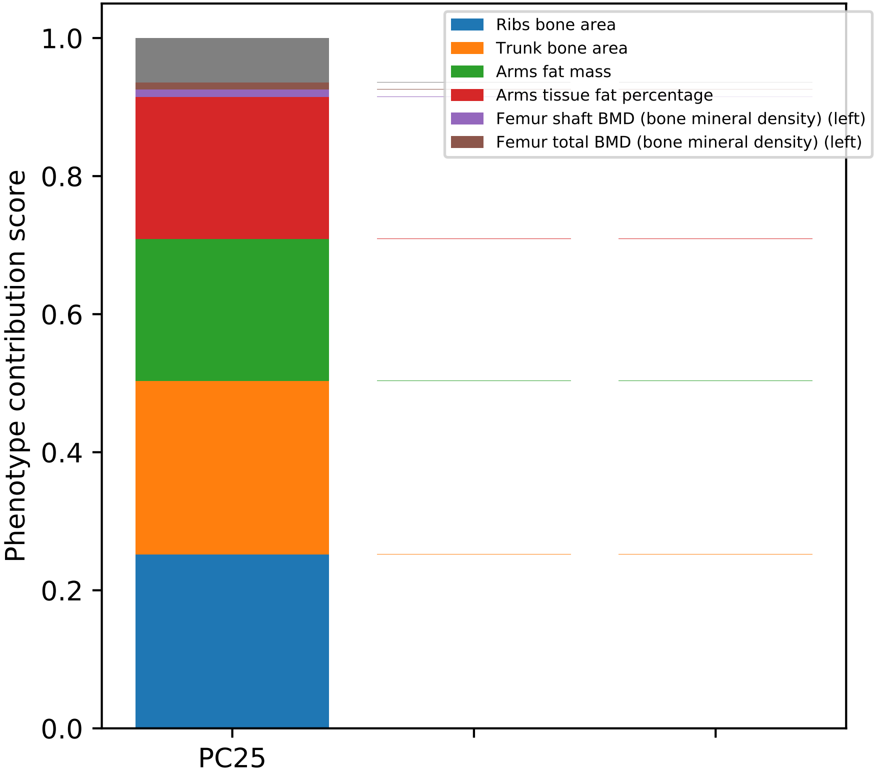

Ribs bone area
Trunk bone area
Arms fat mass
Arms tissue fat percentage
Femur shaft BMD (bone mineral density) (left)
Femur total BMD (bone mineral density) (left)
Phenotype contribution score
0.0
0.2
0.4
0.6
0.8
1.0
PC25

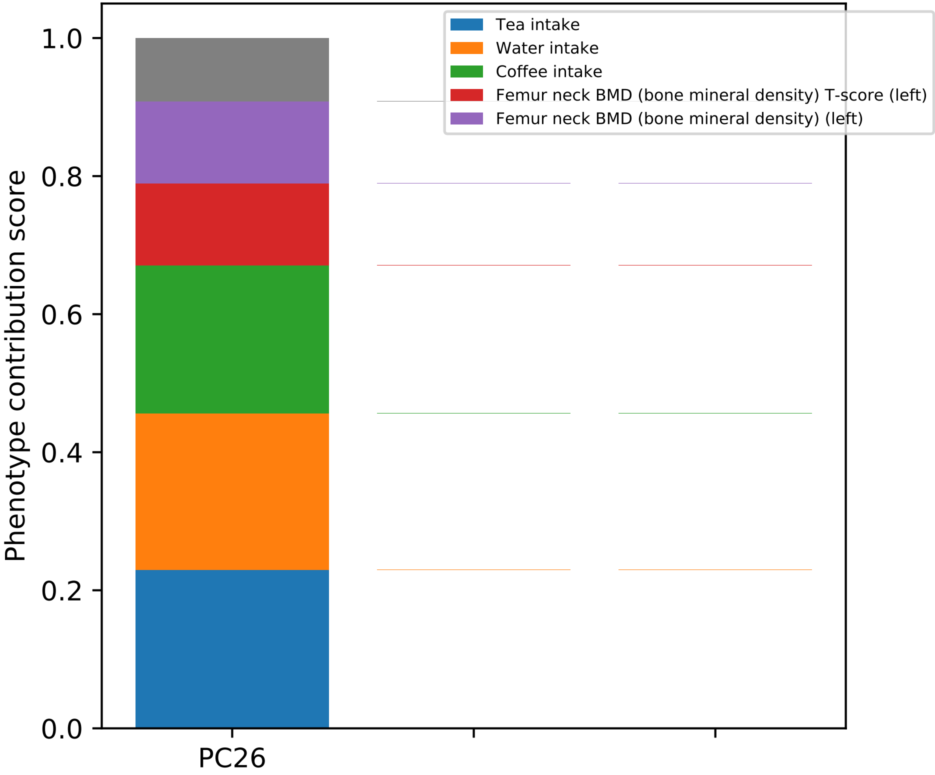

Phenotype contribution score
1.0
0.8
0.6
0.4
0.2
0.0
PC26
Tea intake
Water intake
Coffee intake
Femur neck BMD (bone mineral density) T-score (left)
Femur neck BMD (bone mineral density) (left)

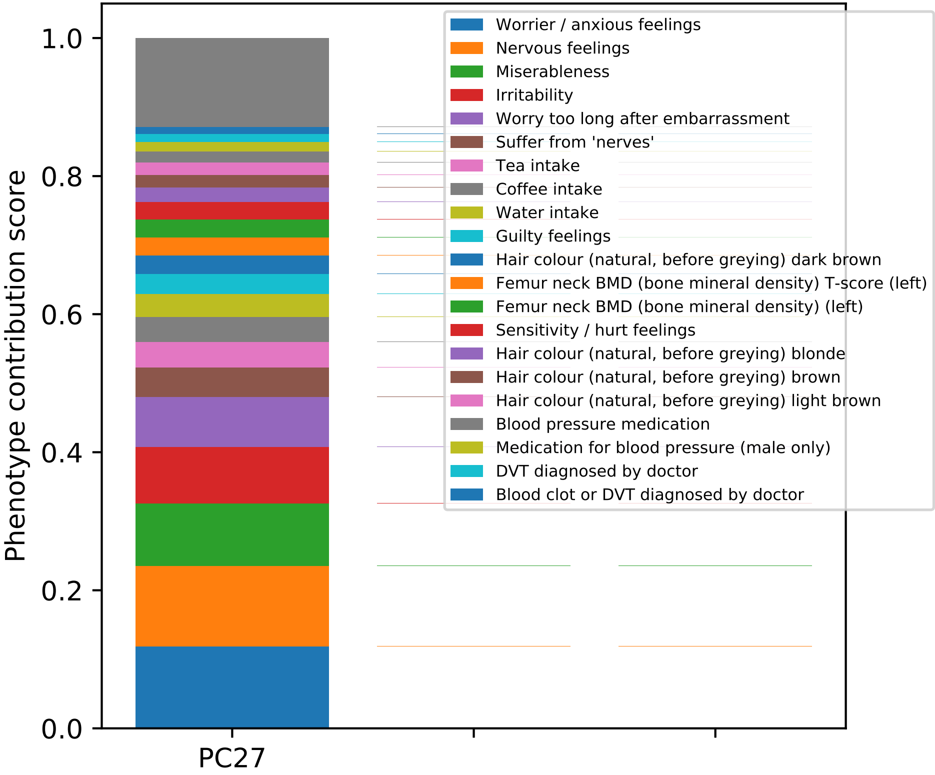
Phenotype contribution score
1.0
0.8
0.6
0.4
0.2
0.0
PC27
Worrier / anxious feelings
Nervous feelings
Miserableness
Irritability
Worry too long after embarrassment
Suffer from 'nerves'
Tea intake
Coffee intake
Water intake
Guilty feelings
Hair colour (natural, before greying) dark brown
Femur neck BMD (bone mineral density) T-score (left)
Femur neck BMD (bone mineral density) (left)
Sensitivity / hurt feelings
Hair colour (natural, before greying) blonde
Hair colour (natural, before greying) brown
Hair colour (natural, before greying) light brown
Blood pressure medication
Medication for blood pressure (male only)
DVT diagnosed by doctor
Blood clot or DVT diagnosed by doctor

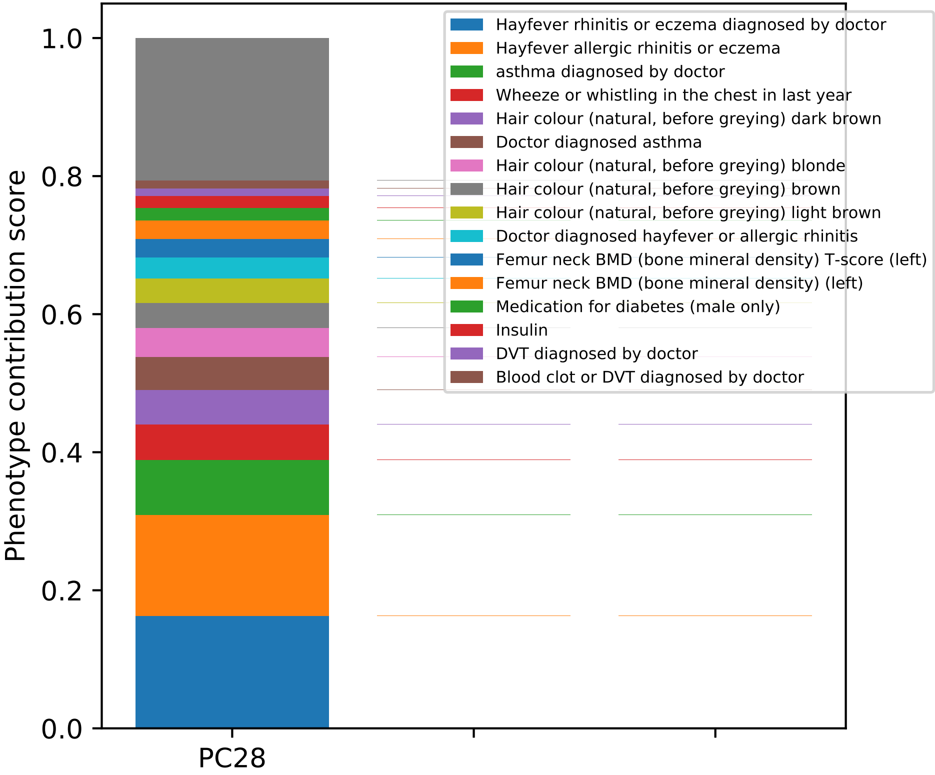

Phenotype contribution score
0.0
0.2
0.4
0.6
0.8
1.0
PC28
Hayfever rhinitis or eczema diagnosed by doctor
Hayfever allergic rhinitis or eczema
asthma diagnosed by doctor
Wheeze or whistling in the chest in last year
Hair colour (natural, before greying) dark brown
Doctor diagnosed asthma
Hair colour (natural, before greying) blonde
Hair colour (natural, before greying) brown
Hair colour (natural, before greying) light brown
Doctor diagnosed hayfever or allergic rhinitis
Femur neck BMD (bone mineral density) T-score (left)
Femur neck BMD (bone mineral density) (left)
Medication for diabetes (male only)
Insulin
DVT diagnosed by doctor
Blood clot or DVT diagnosed by doctor

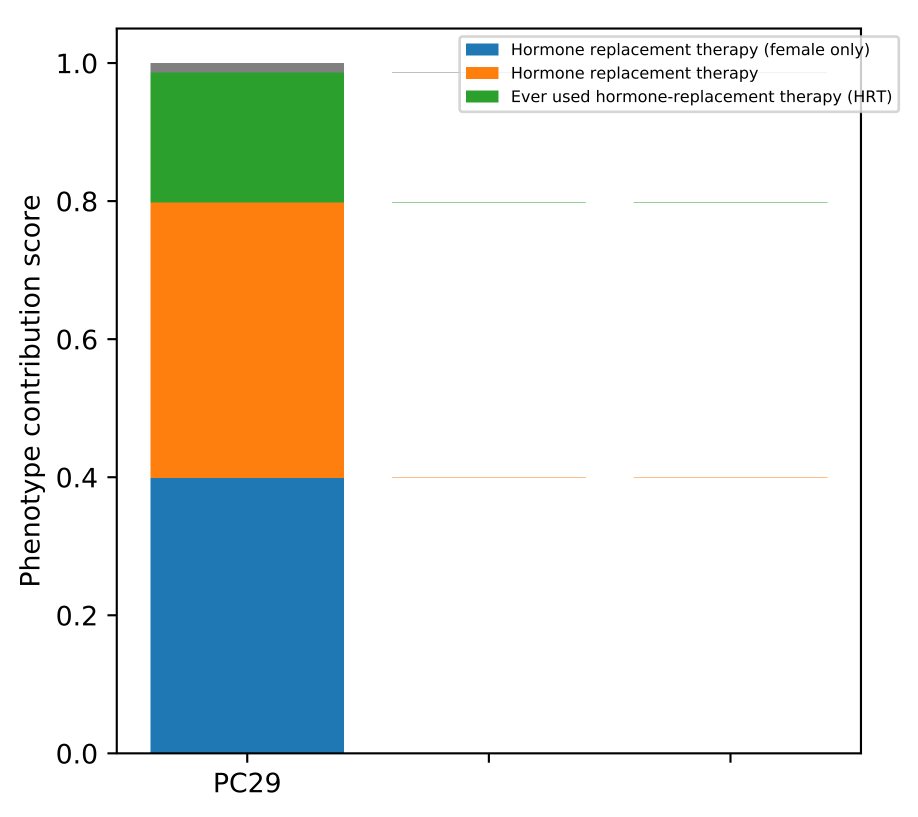

Hormone replacement therapy (female only)
Hormone replacement therapy
Ever used hormone-replacement therapy (HRT)
Phenotype contribution score
0.0
0.2
0.4
0.6
0.8
1.0
PC29

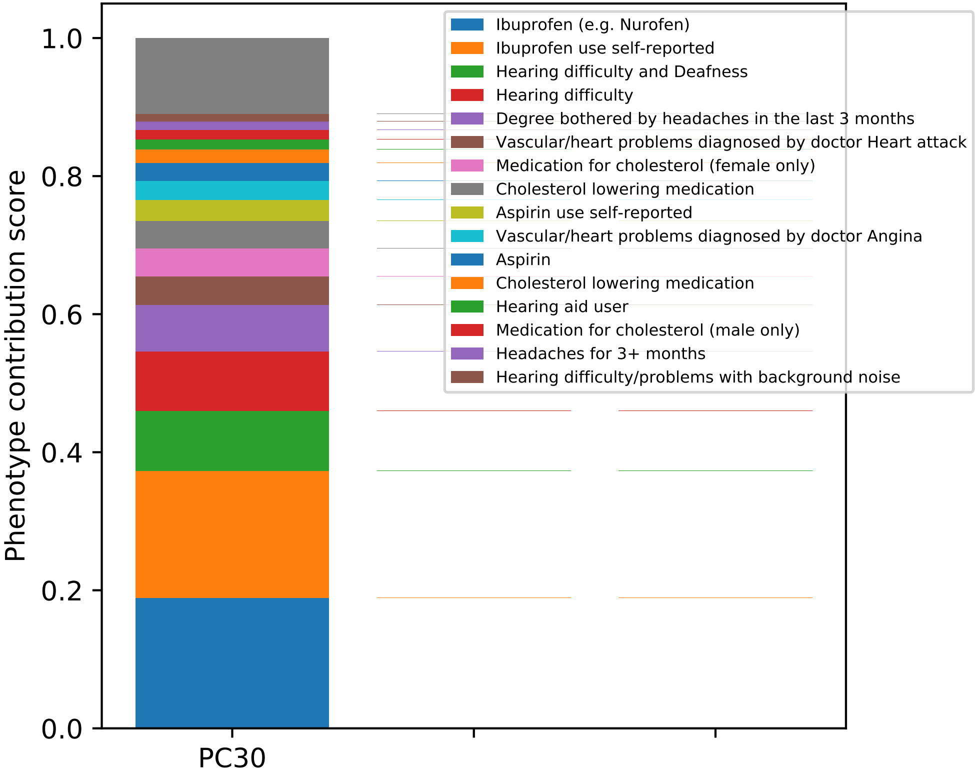

Phenotype contribution score
1.0
0.8
0.6
0.4
0.2
0.0
PC30
Ibuprofen (e.g. Nurofen)
Ibuprofen use self-reported
Hearing difficulty and Deafness
Hearing difficulty
Degree bothered by headaches in the last 3 months
Vascular/heart problems diagnosed by doctor Heart attack
Medication for cholesterol (female only)
Cholesterol lowering medication
Aspirin use self-reported
Vascular/heart problems diagnosed by doctor Angina
Aspirin
Cholesterol lowering medication
Hearing aid user
Medication for cholesterol (male only)
Headaches for 3+ months
Hearing difficulty/problems with background noise

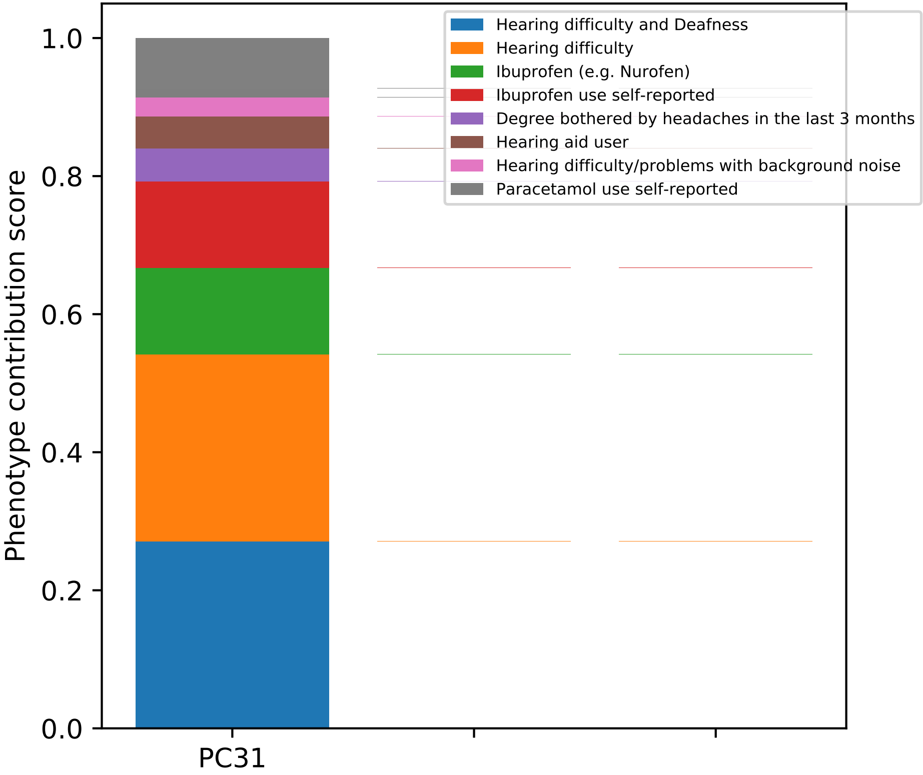
Hearing difficulty and Deafness
Hearing difficulty
Ibuprofen (e.g. Nurofen)
Ibuprofen use self-reported
Degree bothered by headaches in the last 3 months
Hearing aid user
Hearing difficulty/problems with background noise
Paracetamol use self-reported
Phenotype contribution score
1.0
0.8
0.6
0.4
0.2
0.0
PC31

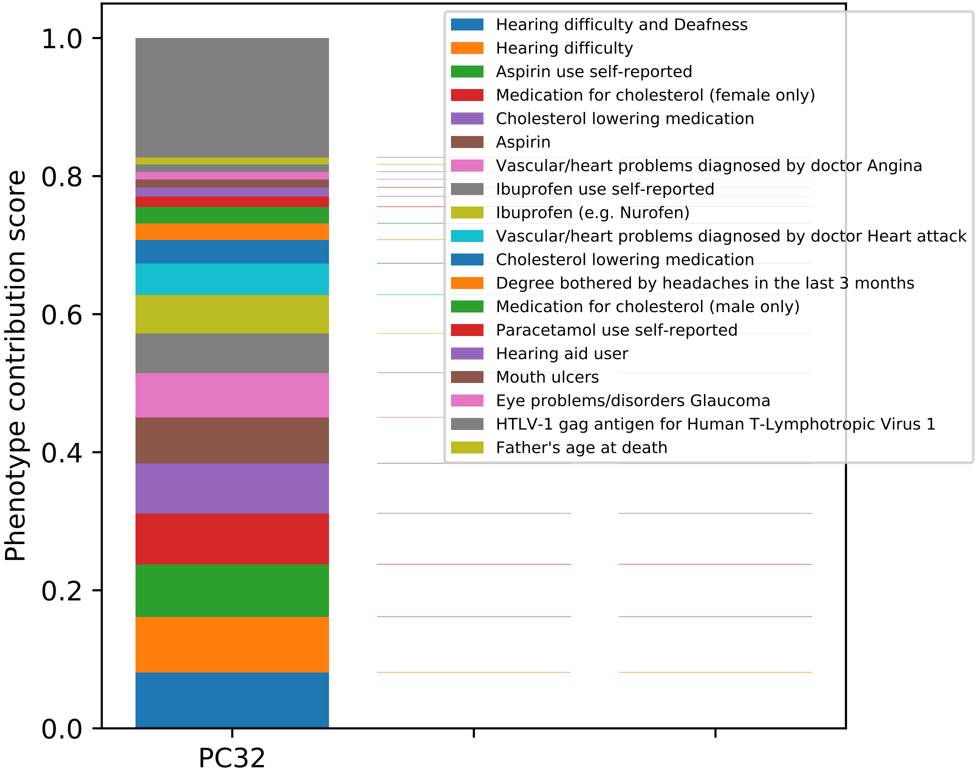

Phenotype contribution score
1.0
0.8
0.6
0.4
0.2
0.0
PC32
Hearing difficulty and Deafness
Hearing difficulty
Aspirin use self-reported
Medication for cholesterol (female only)
Cholesterol lowering medication
Aspirin
Vascular/heart problems diagnosed by doctor Angina
Ibuprofen use self-reported
Ibuprofen (e.g. Nurofen)
Vascular/heart problems diagnosed by doctor Heart attack
Cholesterol lowering medication
Degree bothered by headaches in the last 3 months
Medication for cholesterol (male only)
Paracetamol use self-reported
Hearing aid user
Mouth ulcers
Eye problems/disorders Glaucoma
HTLV-1 gag antigen for Human T-Lymphotropic Virus 1
Father's age at death

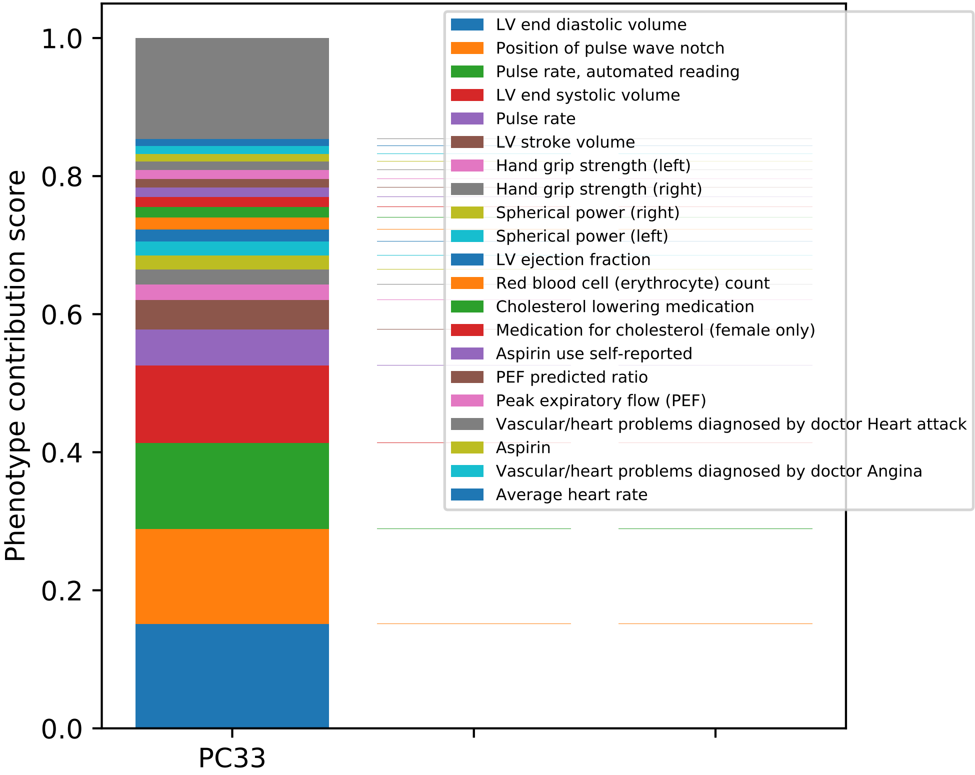

Phenotype contribution score
1.0
0.8
0.6
0.4
0.2
0.0
PC33
LV end diastolic volume
Position of pulse wave notch
Pulse rate, automated reading
LV end systolic volume
Pulse rate
LV stroke volume
Hand grip strength (left)
Hand grip strength (right)
Spherical power (right)
Spherical power (left)
LV ejection fraction
Red blood cell (erythrocyte) count
Cholesterol lowering medication
Medication for cholesterol (female only)
Aspirin use self-reported
PEF predicted ratio
Peak expiratory flow (PEF)
Vascular/heart problems diagnosed by doctor Heart attack
Aspirin
Vascular/heart problems diagnosed by doctor Angina
Average heart rate

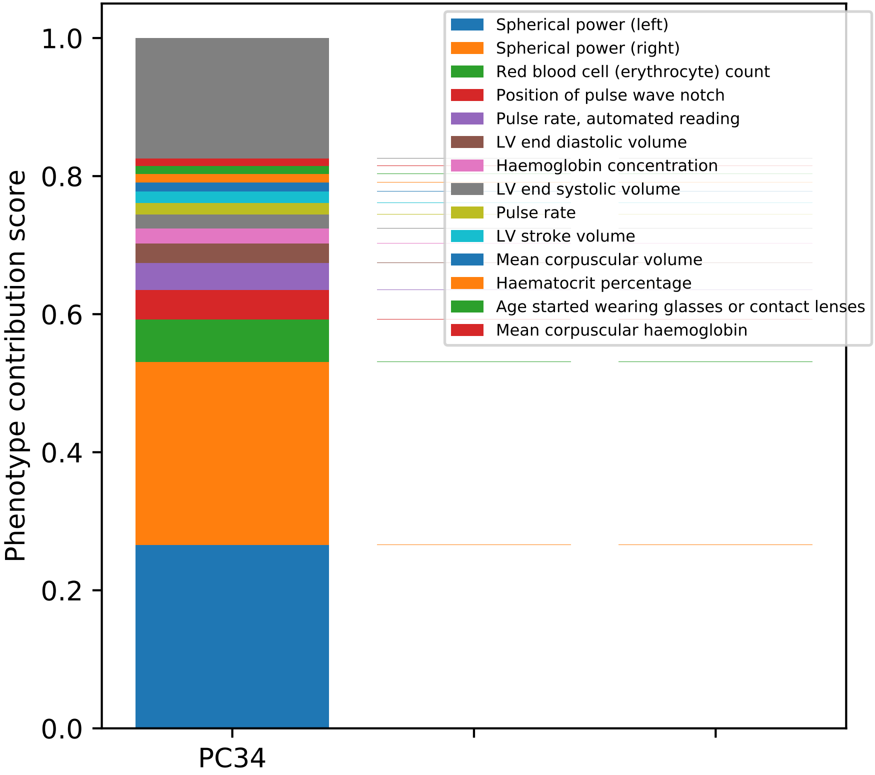
Spherical power (left)
Spherical power (right)
Red blood cell (erythrocyte) count
Position of pulse wave notch
Pulse rate, automated reading
LV end diastolic volume
Haemoglobin concentration
LV end systolic volume
Pulse rate
LV stroke volume
Mean corpuscular volume
Haematocrit percentage
Age started wearing glasses or contact lenses
Mean corpuscular haemoglobin
Phenotype contribution score
1.0
0.8
0.6
0.4
0.2
0.0
PC34

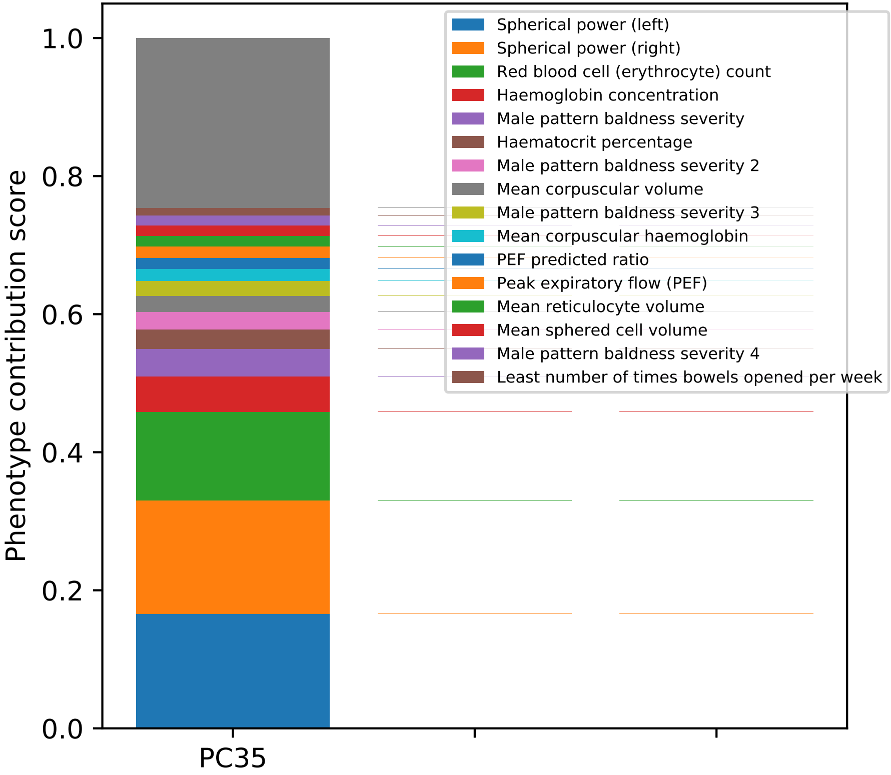
Phenotype contribution score
0.0
0.2
0.4
0.6
0.8
1.0
PC35
Spherical power (left)
Spherical power (right)
Red blood cell (erythrocyte) count
Haemoglobin concentration
Male pattern baldness severity
Haematocrit percentage
Male pattern baldness severity 2
Mean corpuscular volume
Male pattern baldness severity 3
Mean corpuscular haemoglobin
PEF predicted ratio
Peak expiratory flow (PEF)
Mean reticulocyte volume
Mean sphered cell volume
Male pattern baldness severity 4
Least number of times bowels opened per week

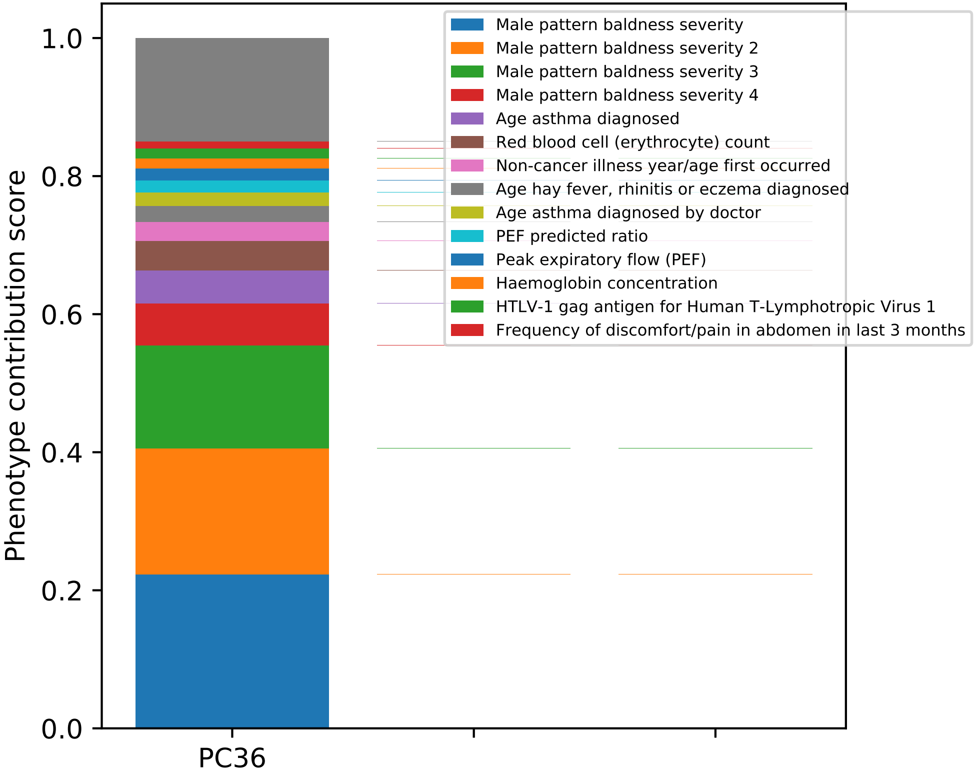

Phenotype contribution score
1.0
0.8
0.6
0.4
0.2
0.0
PC36
Male pattern baldness severity
Male pattern baldness severity 2
Male pattern baldness severity 3
Male pattern baldness severity 4
Age asthma diagnosed
Red blood cell (erythrocyte) count
Non-cancer illness year/age first occurred
Age hay fever, rhinitis or eczema diagnosed
Age asthma diagnosed by doctor
PEF predicted ratio
Peak expiratory flow (PEF)
Haemoglobin concentration
HTLV-1 gag antigen for Human T-Lymphotropic Virus 1
Frequency of discomfort/pain in abdomen in last 3 months

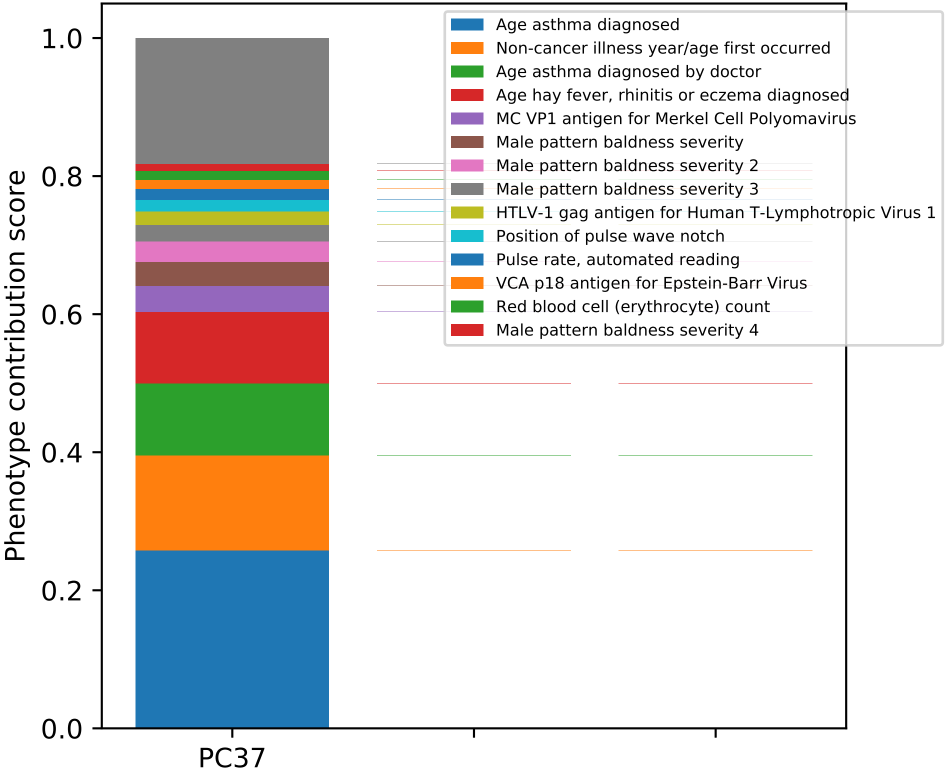

Phenotype contribution score
1.0
0.8
0.6
0.4
0.2
0.0
PC37
Age asthma diagnosed
Non-cancer illness year/age first occurred
Age asthma diagnosed by doctor
Age hay fever, rhinitis or eczema diagnosed
MC VP1 antigen for Merkel Cell Polyomavirus
Male pattern baldness severity
Male pattern baldness severity 2
Male pattern baldness severity 3
HTLV-1 gag antigen for Human T-Lymphotropic Virus 1
Position of pulse wave notch
Pulse rate, automated reading
VCA p18 antigen for Epstein-Barr Virus
Red blood cell (erythrocyte) count
Male pattern baldness severity 4

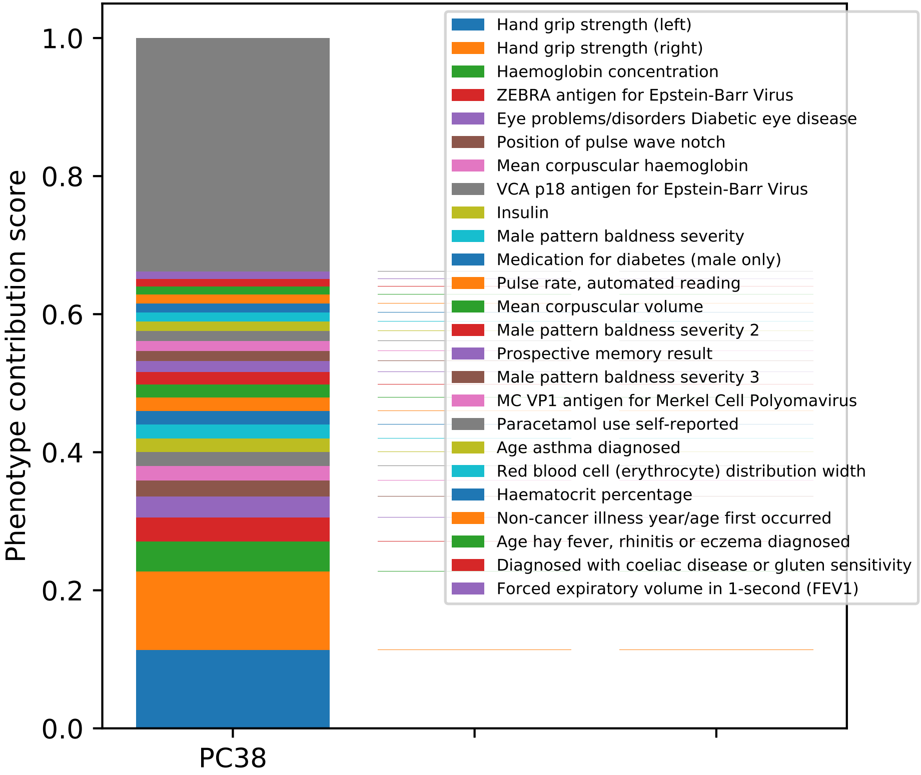

Phenotype contribution score
0.0
0.2
0.4
0.6
0.8
1.0
PC38
Hand grip strength (left)
Hand grip strength (right)
Haemoglobin concentration
ZEBRA antigen for Epstein-Barr Virus
Eye problems/disorders Diabetic eye disease
Position of pulse wave notch
Mean corpuscular haemoglobin
VCA p18 antigen for Epstein-Barr Virus
Insulin
Male pattern baldness severity
Medication for diabetes (male only)
Pulse rate, automated reading
Mean corpuscular volume
Male pattern baldness severity 2
Prospective memory result
Male pattern baldness severity 3
MC VP1 antigen for Merkel Cell Polyomavirus
Paracetamol use self-reported
Age asthma diagnosed
Red blood cell (erythrocyte) distribution width
Haematocrit percentage
Non-cancer illness year/age first occurred
Age hay fever, rhinitis or eczema diagnosed
Diagnosed with coeliac disease or gluten sensitivity
Forced expiratory volume in 1-second (FEV1)

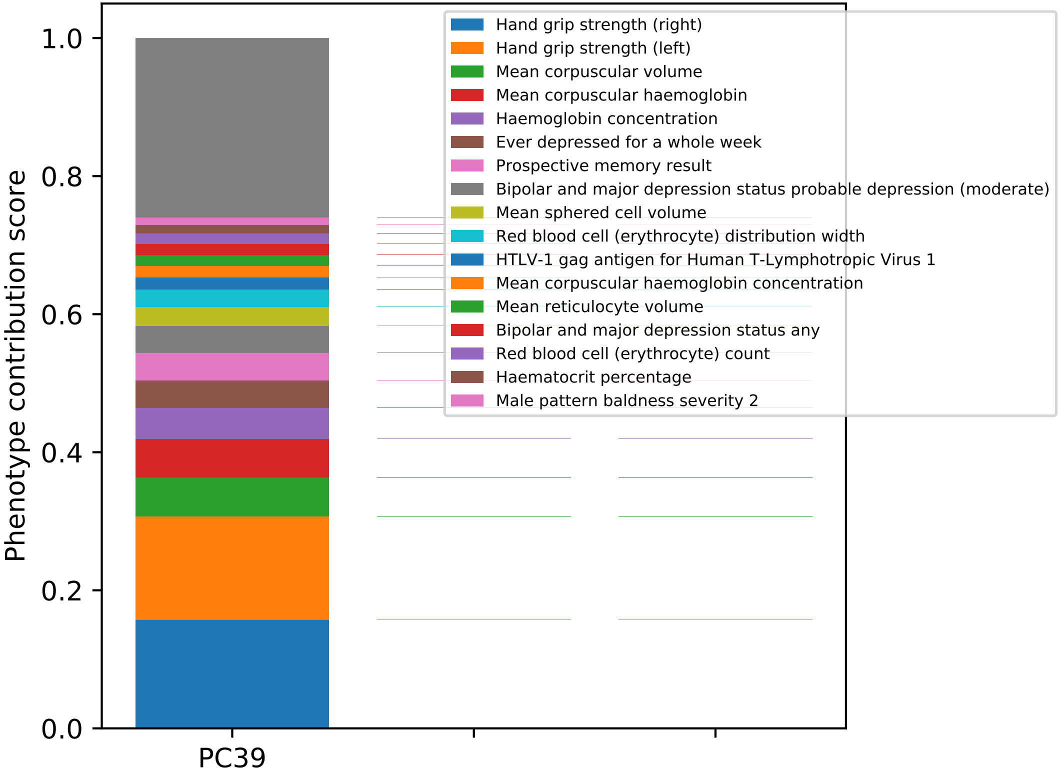

Phenotype contribution score
1.0
0.8
0.6
0.4
0.2
0.0
PC39
Hand grip strength (right)
Hand grip strength (left)
Mean corpuscular volume
Mean corpuscular haemoglobin
Haemoglobin concentration
Ever depressed for a whole week
Prospective memory result
Bipolar and major depression status probable depression (moderate)
Mean sphered cell volume
Red blood cell (erythrocyte) distribution width
HTLV-1 gag antigen for Human T-Lymphotropic Virus 1
Mean corpuscular haemoglobin concentration
Mean reticulocyte volume
Bipolar and major depression status any
Red blood cell (erythrocyte) count
Haematocrit percentage
Male pattern baldness severity 2

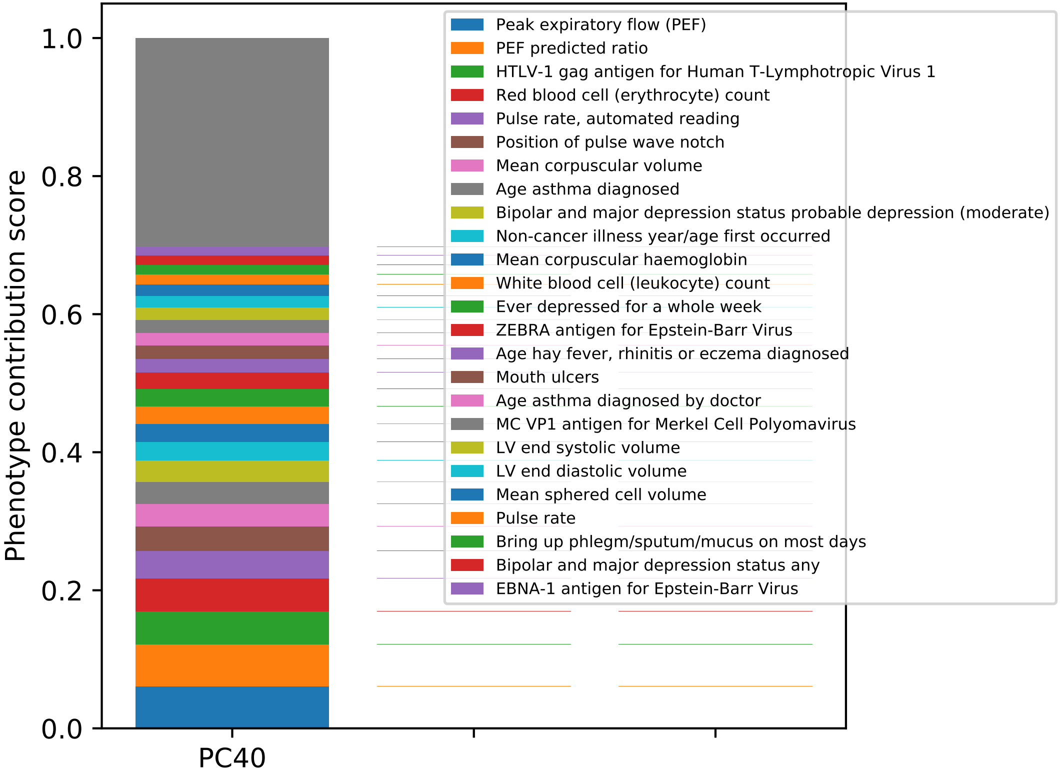

Phenotype contribution score
0.0
0.2
0.4
0.6
0.8
1.0
PC40
Peak expiratory flow (PEF)
PEF predicted ratio
HTLV-1 gag antigen for Human T-Lymphotropic Virus 1
Red blood cell (erythrocyte) count
Pulse rate, automated reading
Position of pulse wave notch
Mean corpuscular volume
Age asthma diagnosed
Bipolar and major depression status probable depression (moderate)
Non-cancer illness year/age first occurred
Mean corpuscular haemoglobin
White blood cell (leukocyte) count
Ever depressed for a whole week
ZEBRA antigen for Epstein-Barr Virus
Age hay fever, rhinitis or eczema diagnosed
Mouth ulcers
Age asthma diagnosed by doctor
MC VP1 antigen for Merkel Cell Polyomavirus
LV end systolic volume
LV end diastolic volume
Mean sphered cell volume
Pulse rate
Bring up phlegm/sputum/mucus on most days
Bipolar and major depression status any
EBNA-1 antigen for Epstein-Barr Virus

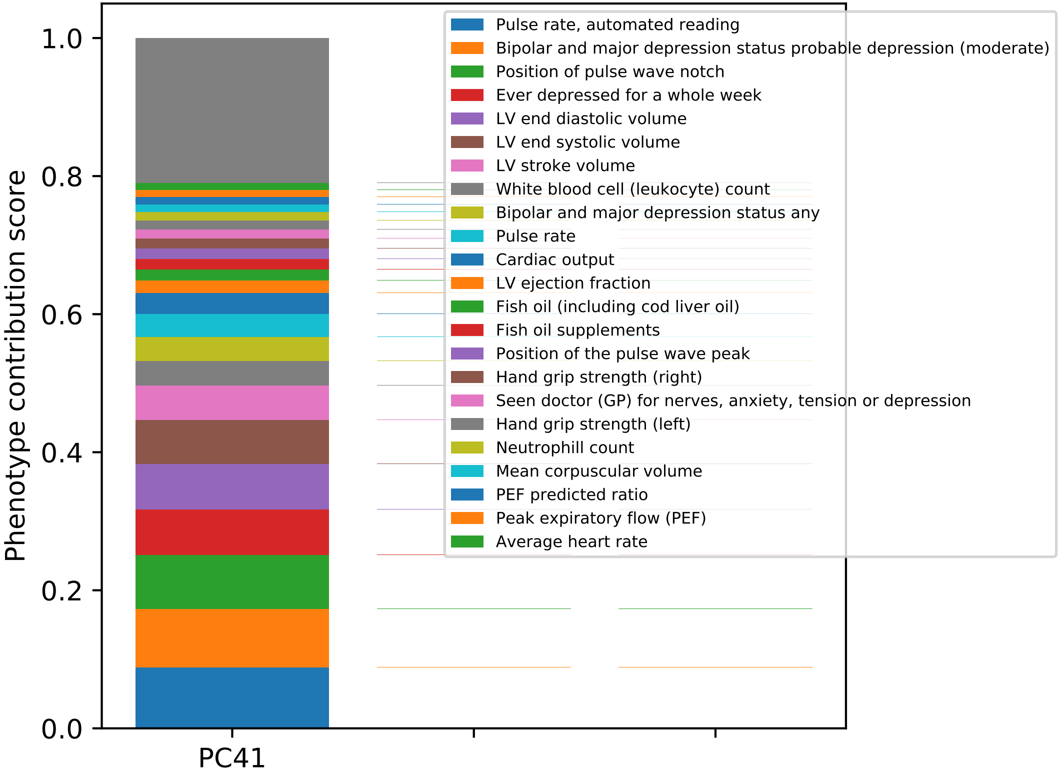
Phenotype contribution score
1.0
0.8
0.6
0.4
0.2
0.0
PC41
Pulse rate, automated reading
Bipolar and major depression status probable depression (moderate)
Position of pulse wave notch
Ever depressed for a whole week
LV end diastolic volume
LV end systolic volume
LV stroke volume
White blood cell (leukocyte) count
Bipolar and major depression status any
Pulse rate
Cardiac output
LV ejection fraction
Fish oil (including cod liver oil)
Fish oil supplements
Position of the pulse wave peak
Hand grip strength (right)
Seen doctor (GP) for nerves, anxiety, tension or depression
Hand grip strength (left)
Neutrophill count
Mean corpuscular volume
PEF predicted ratio
Peak expiratory flow (PEF)
Average heart rate

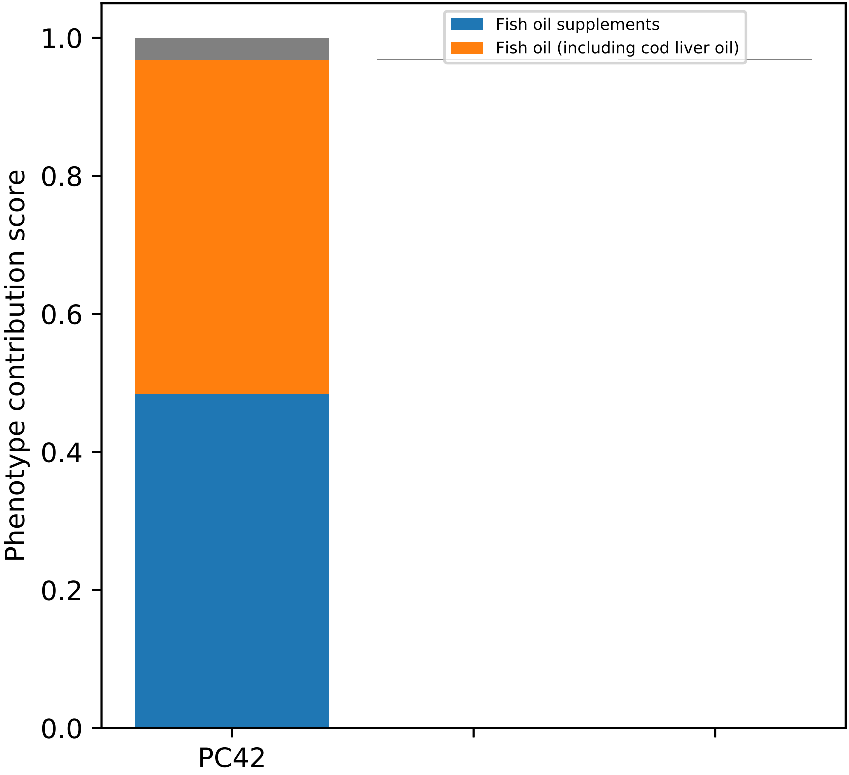
Fish oil supplements
Fish oil (including cod liver oil)
Phenotype contribution score
1.0
0.8
0.6
0.4
0.2
0.0
PC42

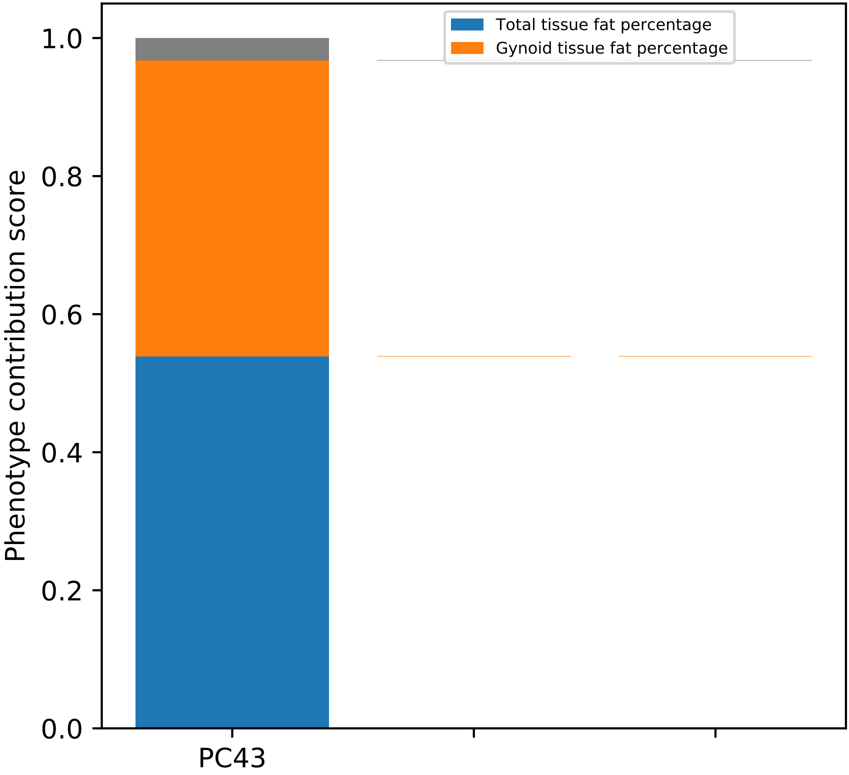
Total tissue fat percentage
Gynoid tissue fat percentage
1.0
0.8
0.6
0.4
0.2
0.0
Phenotype contribution score
PC43

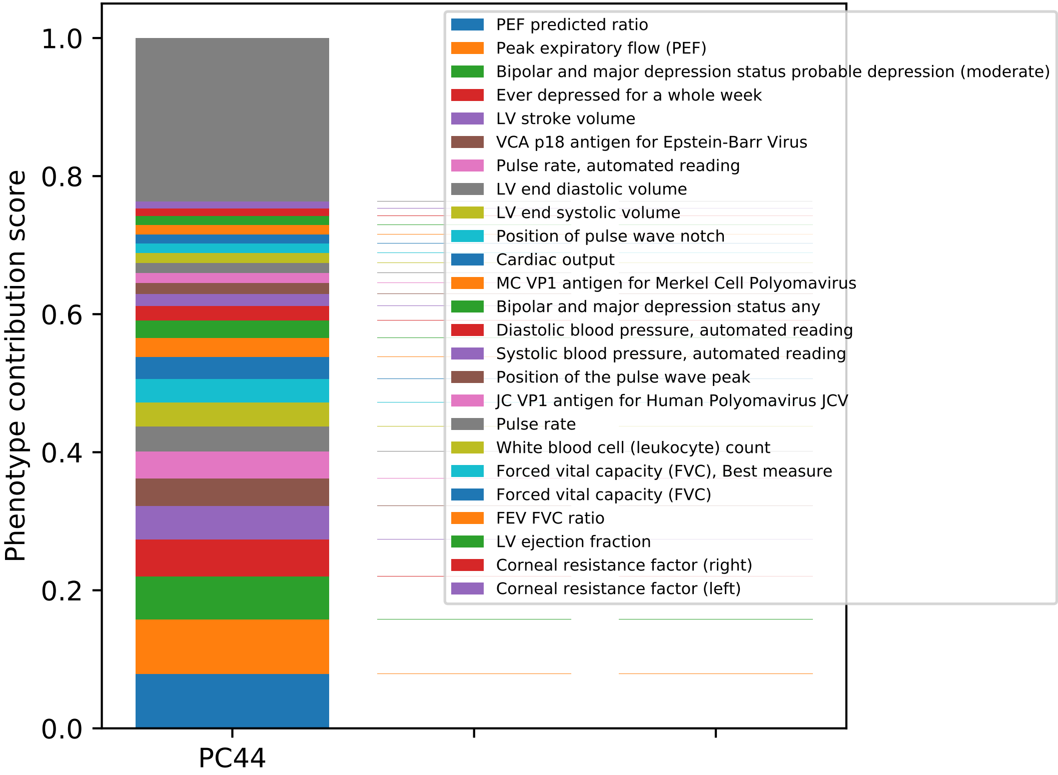
Phenotype contribution score
1.0
0.8
0.6
0.4
0.2
0.0
PC44
PEF predicted ratio
Peak expiratory flow (PEF)
Bipolar and major depression status probable depression (moderate)
Ever depressed for a whole week
LV stroke volume
VCA p18 antigen for Epstein-Barr Virus
Pulse rate, automated reading
LV end diastolic volume
LV end systolic volume
Position of pulse wave notch
Cardiac output
MC VP1 antigen for Merkel Cell Polyomavirus
Bipolar and major depression status any
Diastolic blood pressure, automated reading
Systolic blood pressure, automated reading
Position of the pulse wave peak
JC VP1 antigen for Human Polyomavirus JCV
Pulse rate
White blood cell (leukocyte) count
Forced vital capacity (FVC), Best measure
Forced vital capacity (FVC)
FEV FVC ratio
LV ejection fraction
Corneal resistance factor (right)
Corneal resistance factor (left)

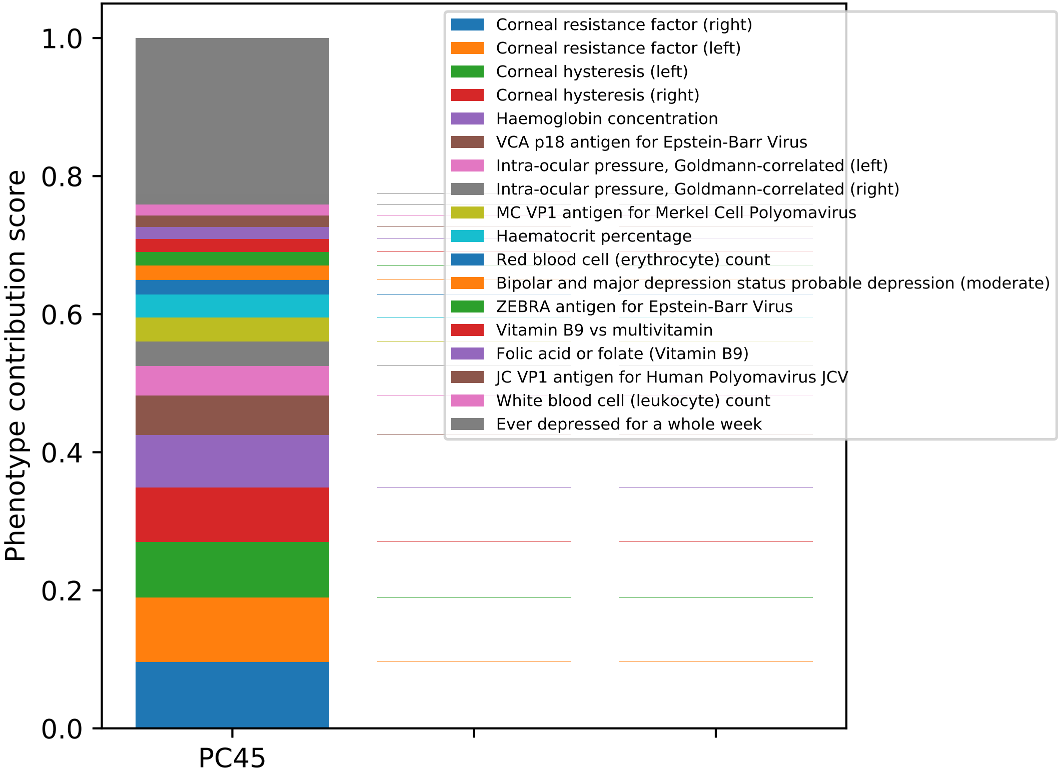
Phenotype contribution score
0.0
0.2
0.4
0.6
0.8
1.0
PC45
Corneal resistance factor (right)
Corneal resistance factor (left)
Corneal hysteresis (left)
Corneal hysteresis (right)
Haemoglobin concentration
VCA p18 antigen for Epstein-Barr Virus
Intra-ocular pressure, Goldmann-correlated (left)
Intra-ocular pressure, Goldmann-correlated (right)
MC VP1 antigen for Merkel Cell Polyomavirus
Haematocrit percentage
Red blood cell (erythrocyte) count
Bipolar and major depression status probable depression (moderate)
ZEBRA antigen for Epstein-Barr Virus
Vitamin B9 vs multivitamin
Folic acid or folate (Vitamin B9)
JC VP1 antigen for Human Polyomavirus JCV
White blood cell (leukocyte) count
Ever depressed for a whole week

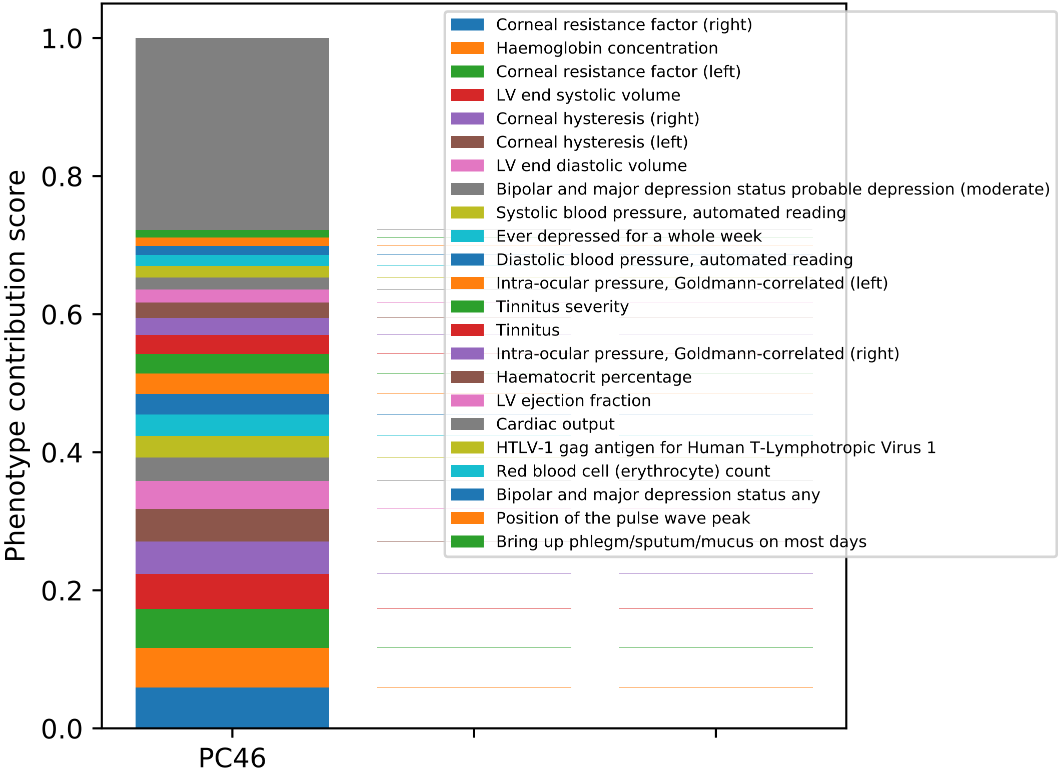

Phenotype contribution score
1.0
0.8
0.6
0.4
0.2
0.0
PC46
Corneal resistance factor (right)
Haemoglobin concentration
Corneal resistance factor (left)
LV end systolic volume
Corneal hysteresis (right)
Corneal hysteresis (left)
LV end diastolic volume
Bipolar and major depression status probable depression (moderate)
Systolic blood pressure, automated reading
Ever depressed for a whole week
Diastolic blood pressure, automated reading
Intra-ocular pressure, Goldmann-correlated (left)
Tinnitus severity
Tinnitus
Intra-ocular pressure, Goldmann-correlated (right)
Haematocrit percentage
LV ejection fraction
Cardiac output
HTLV-1 gag antigen for Human T-Lymphotropic Virus 1
Red blood cell (erythrocyte) count
Bipolar and major depression status any
Position of the pulse wave peak
Bring up phlegm/sputum/mucus on most days

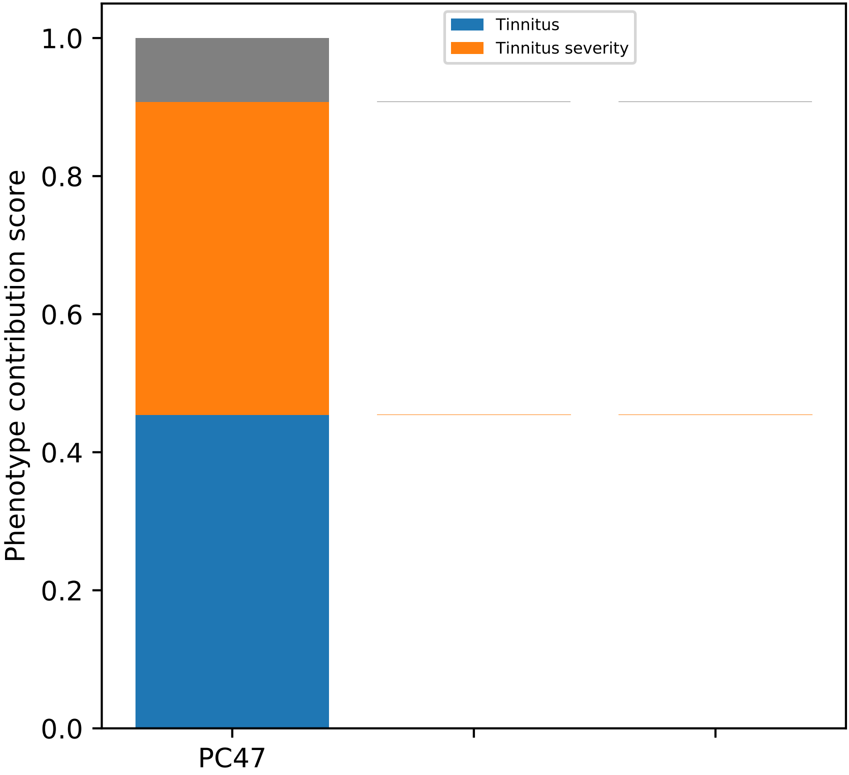
Tinnitus
Tinnitus severity
1.0
0.8
0.6
0.4
0.2
0.0
Phenotype contribution score
PC47

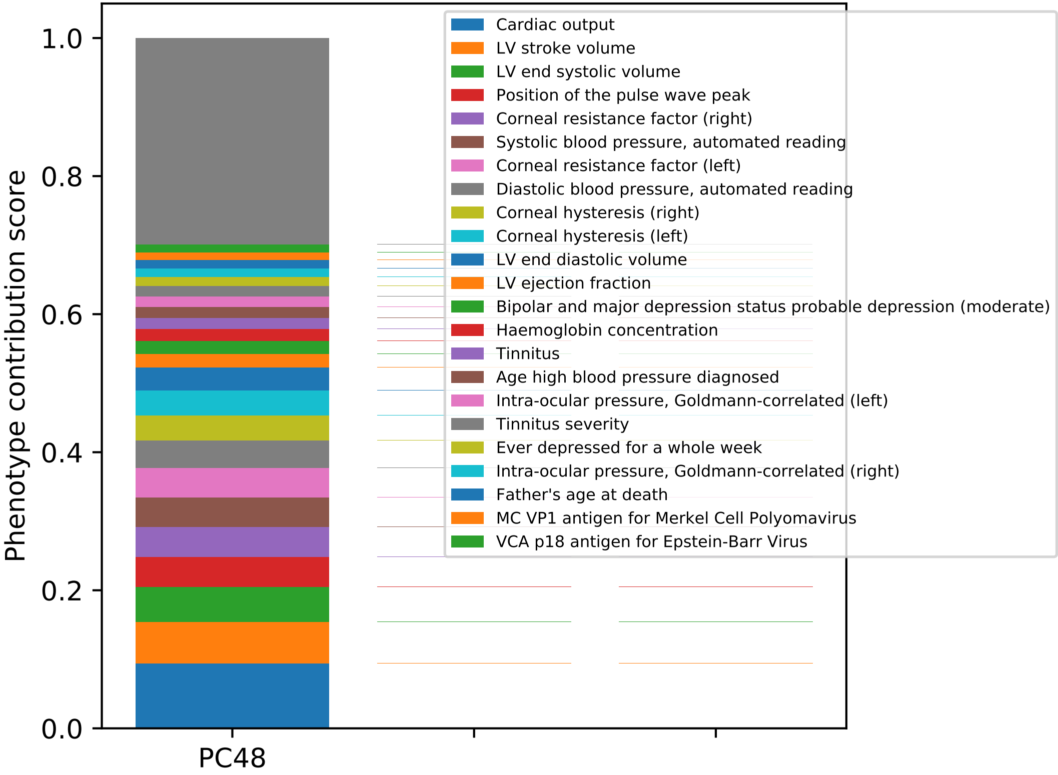

Phenotype contribution score
0.0
0.2
0.4
0.6
0.8
1.0
PC48
Cardiac output
LV stroke volume
LV end systolic volume
Position of the pulse wave peak
Corneal resistance factor (right)
Systolic blood pressure, automated reading
Corneal resistance factor (left)
Diastolic blood pressure, automated reading
Corneal hysteresis (right)
Corneal hysteresis (left)
LV end diastolic volume
LV ejection fraction
Bipolar and major depression status probable depression (moderate)
Haemoglobin concentration
Tinnitus
Age high blood pressure diagnosed
Intra-ocular pressure, Goldmann-correlated (left)
Tinnitus severity
Ever depressed for a whole week
Intra-ocular pressure, Goldmann-correlated (right)
Father's age at death
MC VP1 antigen for Merkel Cell Polyomavirus
VCA p18 antigen for Epstein-Barr Virus

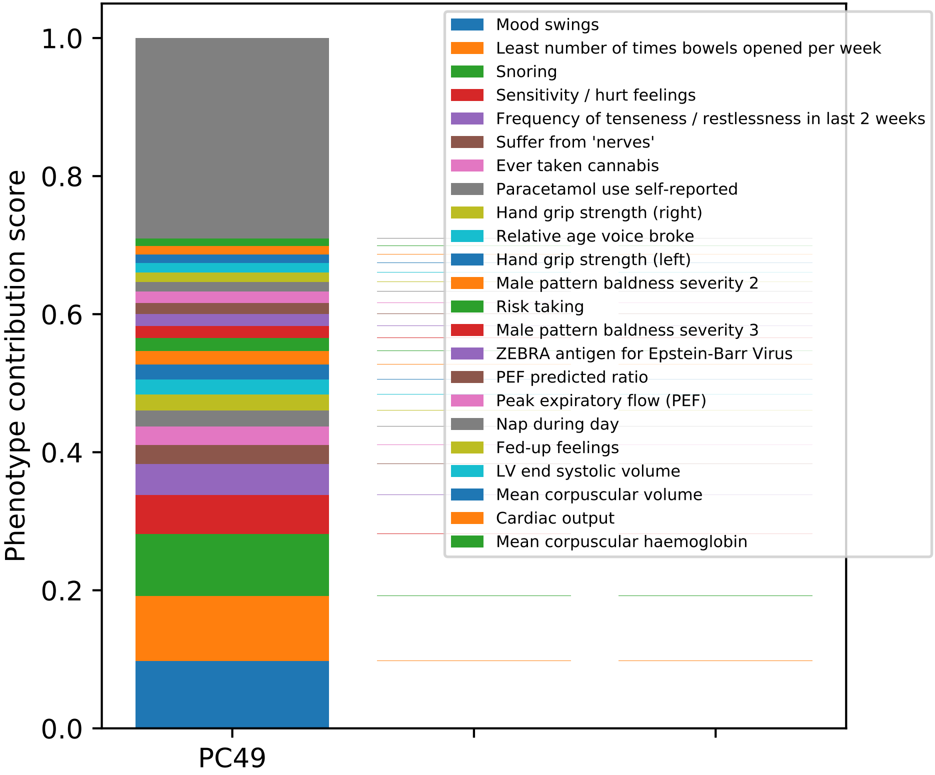

1.0
0.8
0.6
0.4
0.2
0.0
Phenotype contribution score
PC49
Mood swings
Least number of times bowels opened per week
Snoring
Sensitivity / hurt feelings
Frequency of tenseness / restlessness in last 2 weeks
Suffer from 'nerves'
Ever taken cannabis
Paracetamol use self-reported
Hand grip strength (right)
Relative age voice broke
Hand grip strength (left)
Male pattern baldness severity 2
Risk taking
Male pattern baldness severity 3
ZEBRA antigen for Epstein-Barr Virus
PEF predicted ratio
Peak expiratory flow (PEF)
Nap during day
Fed-up feelings
LV end systolic volume
Mean corpuscular volume
Cardiac output
Mean corpuscular haemoglobin

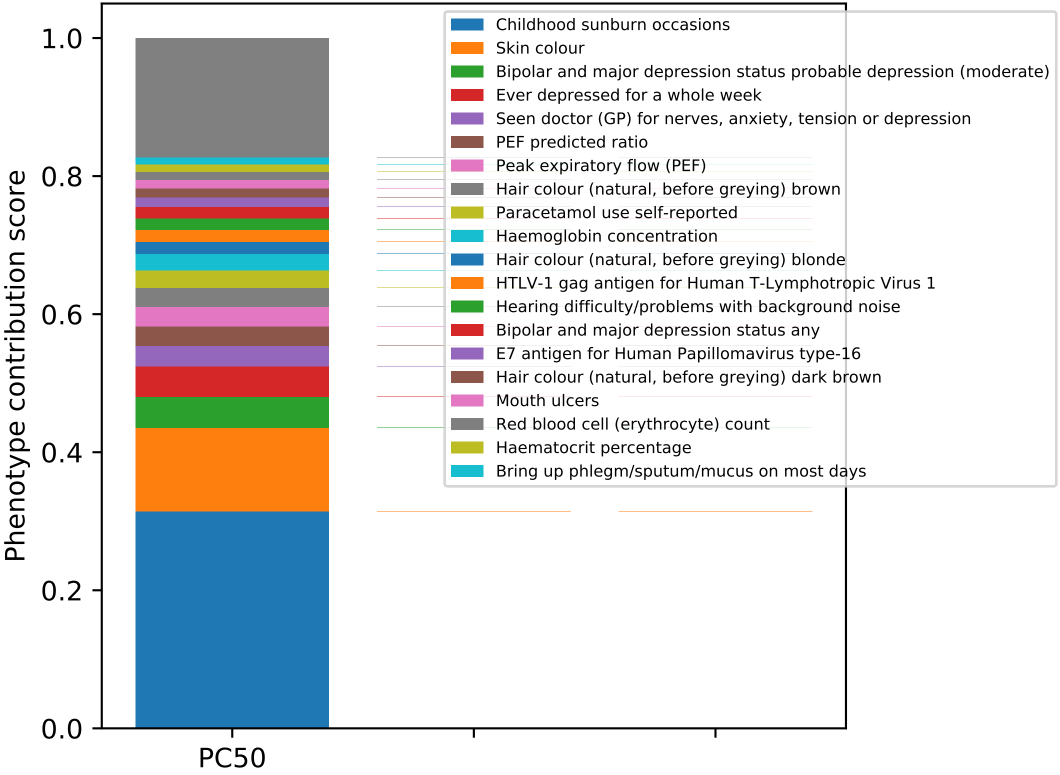
Phenotype contribution score
1.0
0.8
0.6
0.4
0.2
0.0
PC50
Childhood sunburn occasions
Skin colour
Bipolar and major depression status probable depression (moderate)
Ever depressed for a whole week
Seen doctor (GP) for nerves, anxiety, tension or depression
PEF predicted ratio
Peak expiratory flow (PEF)
Hair colour (natural, before greying) brown
Paracetamol use self-reported
Haemoglobin concentration
Hair colour (natural, before greying) blonde
HTLV-1 gag antigen for Human T-Lymphotropic Virus 1
Hearing difficulty/problems with background noise
Bipolar and major depression status any
E7 antigen for Human Papillomavirus type-16
Hair colour (natural, before greying) dark brown
Mouth ulcers
Red blood cell (erythrocyte) count
Haematocrit percentage
Bring up phlegm/sputum/mucus on most days

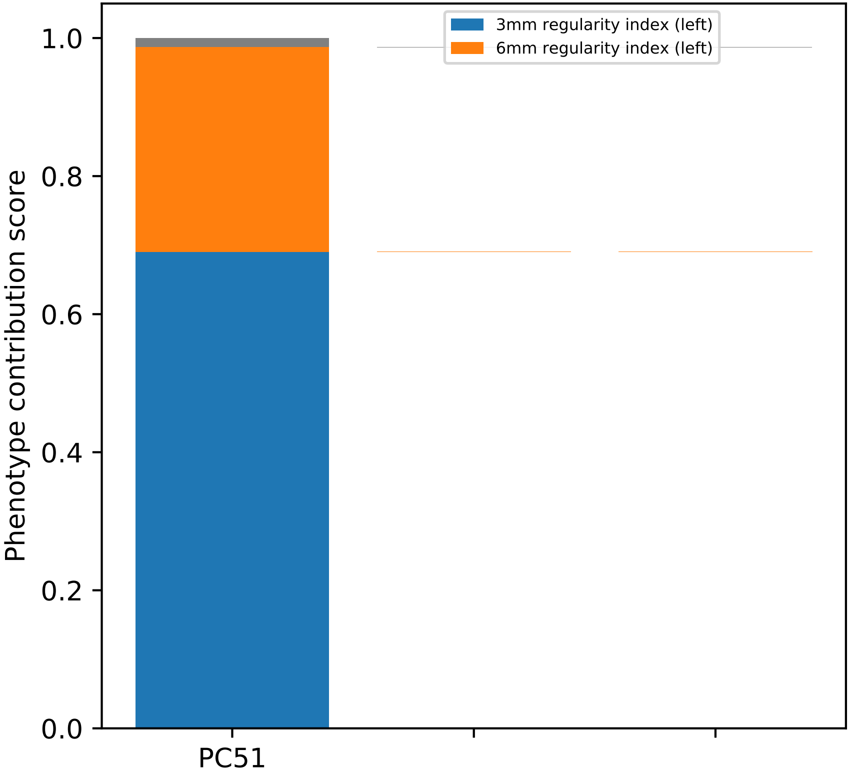

3mm regularity index (left)
6mm regularity index (left)
Phenotype contribution score
1.0
0.8
0.6
0.4
0.2
0.0
PC51

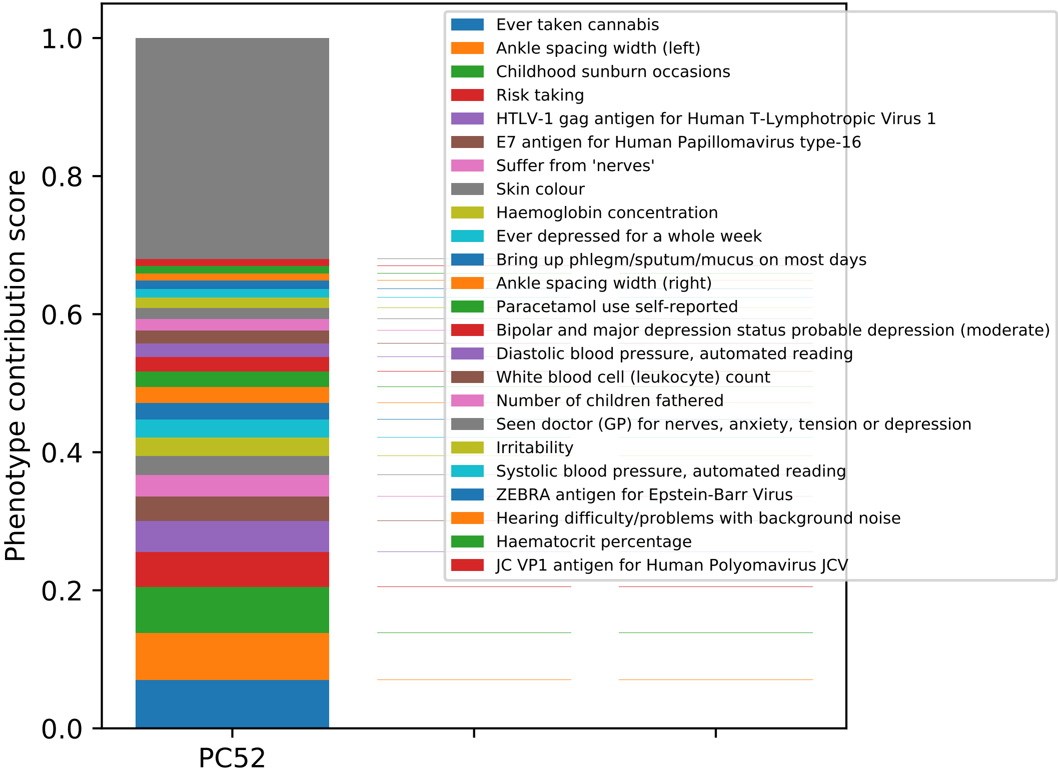
Phenotype contribution score
1.0
0.8
0.6
0.4
0.2
0.0
PC52
Ever taken cannabis
Ankle spacing width (left)
Childhood sunburn occasions
Risk taking
HTLV-1 gag antigen for Human T-Lymphotropic Virus 1
E7 antigen for Human Papillomavirus type-16
Suffer from 'nerves'
Skin colour
Haemoglobin concentration
Ever depressed for a whole week
Bring up phlegm/sputum/mucus on most days
Ankle spacing width (right)
Paracetamol use self-reported
Bipolar and major depression status probable depression (moderate)
Diastolic blood pressure, automated reading
White blood cell (leukocyte) count
Number of children fathered
Seen doctor (GP) for nerves, anxiety, tension or depression
Irritability
Systolic blood pressure, automated reading
ZEBRA antigen for Epstein-Barr Virus
Hearing difficulty/problems with background noise
Haematocrit percentage
JC VP1 antigen for Human Polyomavirus JCV

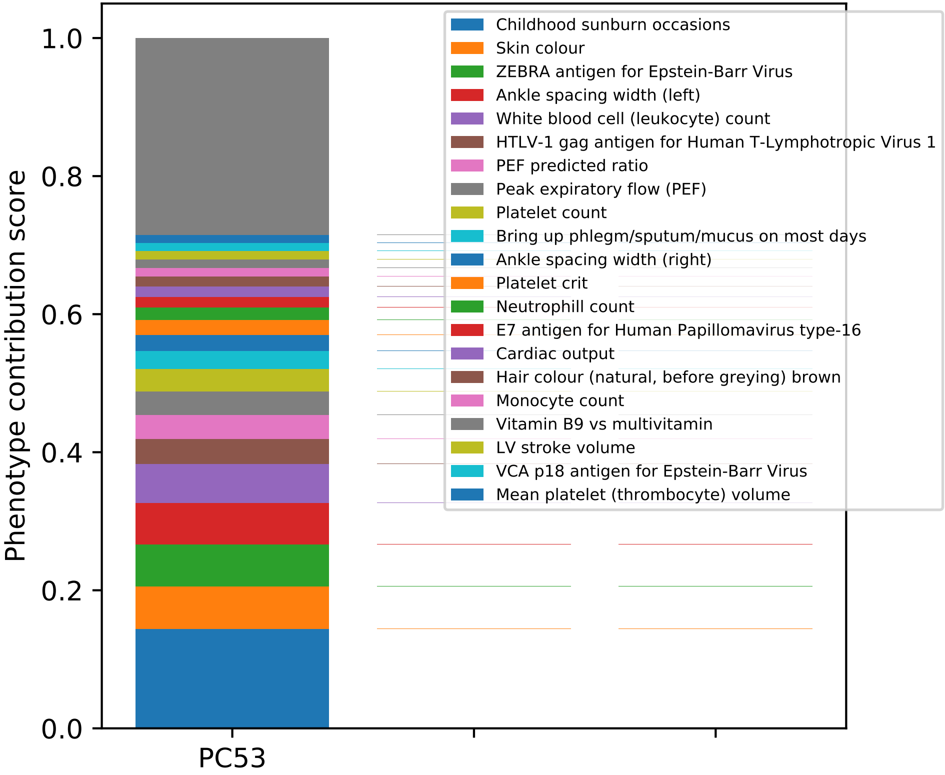

Phenotype contribution score
1.0
0.8
0.6
0.4
0.2
0.0
PC53
Childhood sunburn occasions
Skin colour
ZEBRA antigen for Epstein-Barr Virus
Ankle spacing width (left)
White blood cell (leukocyte) count
HTLV-1 gag antigen for Human T-Lymphotropic Virus 1
PEF predicted ratio
Peak expiratory flow (PEF)
Platelet count
Bring up phlegm/sputum/mucus on most days
Ankle spacing width (right)
Platelet crit
Neutrophill count
E7 antigen for Human Papillomavirus type-16
Cardiac output
Hair colour (natural, before greying) brown
Monocyte count
Vitamin B9 vs multivitamin
LV stroke volume
VCA p18 antigen for Epstein-Barr Virus
Mean platelet (thrombocyte) volume

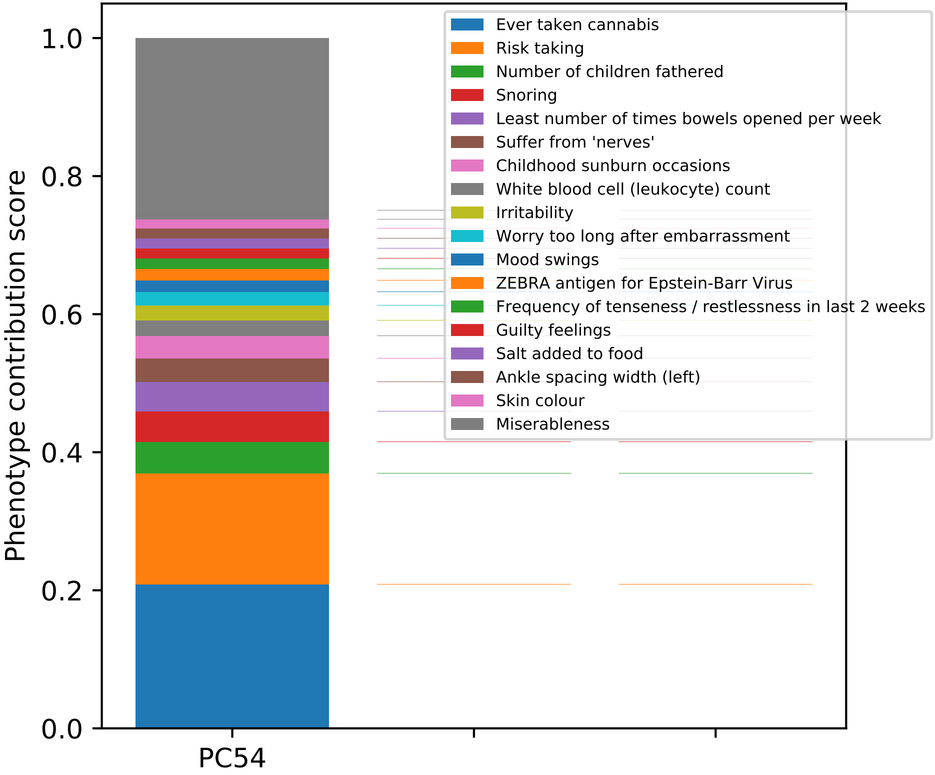
Phenotype contribution score
1.0
0.8
0.6
0.4
0.2
0.0
PC54
Ever taken cannabis
Risk taking
Number of children fathered
Snoring
Least number of times bowels opened per week
Suffer from 'nerves'
Childhood sunburn occasions
White blood cell (leukocyte) count
Irritability
Worry too long after embarrassment
Mood swings
ZEBRA antigen for Epstein-Barr Virus
Frequency of tenseness / restlessness in last 2 weeks
Guilty feelings
Salt added to food
Ankle spacing width (left)
Skin colour
Miserableness

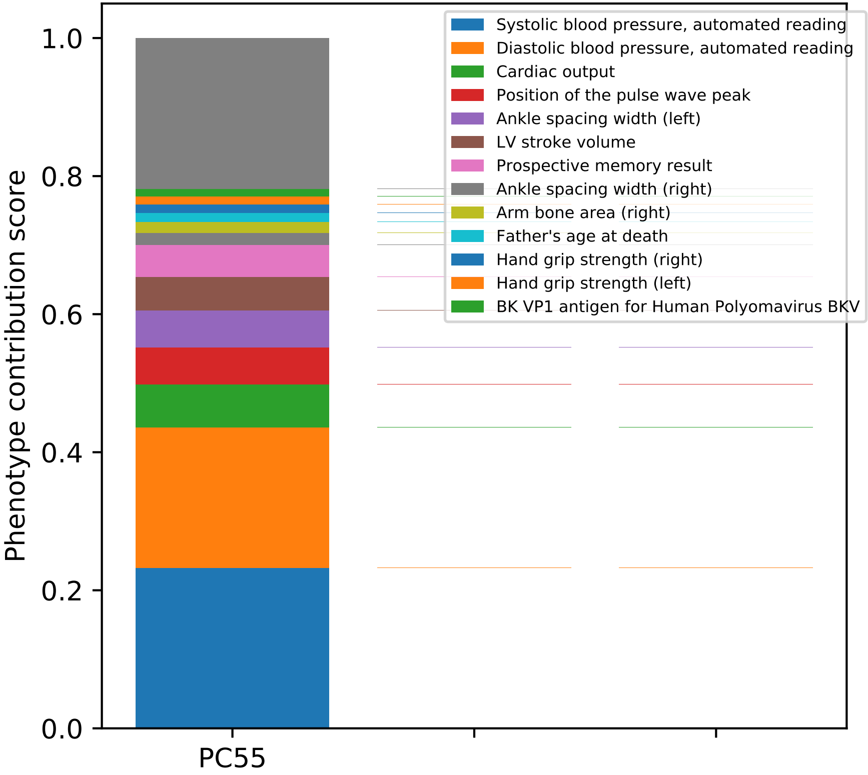

Phenotype contribution score
1.0
0.8
0.6
0.4
0.2
0.0
PC55
Systolic blood pressure, automated reading
Diastolic blood pressure, automated reading
Cardiac output
Position of the pulse wave peak
Ankle spacing width (left)
LV stroke volume
Prospective memory result
Ankle spacing width (right)
Arm bone area (right)
Father's age at death
Hand grip strength (right)
Hand grip strength (left)
BK VP1 antigen for Human Polyomavirus BKV

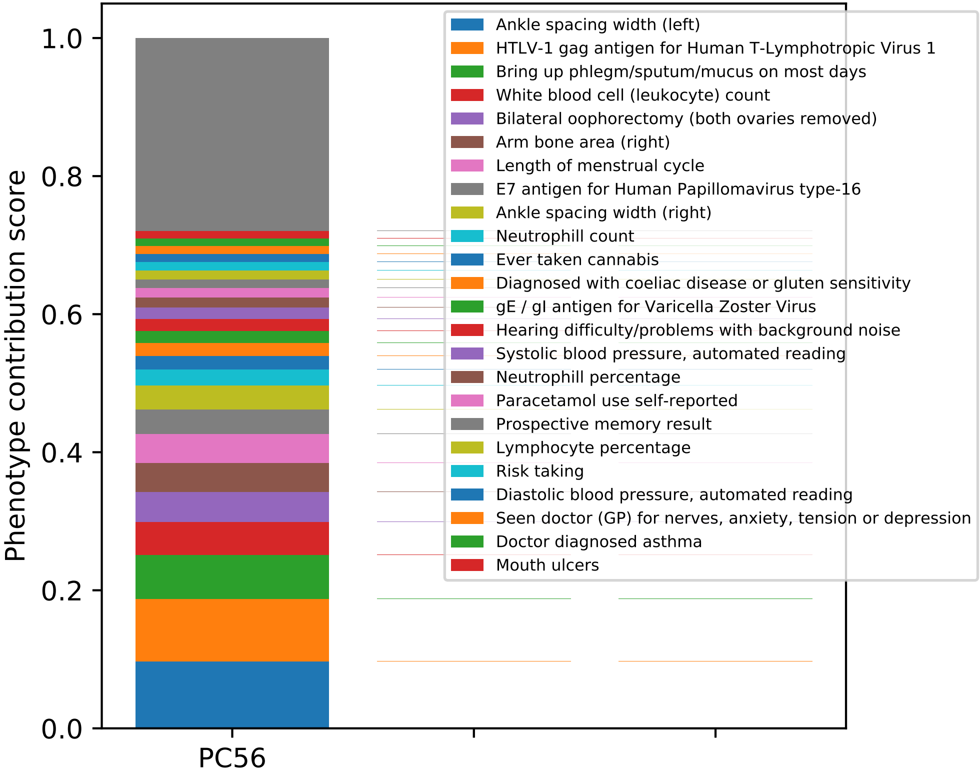
Phenotype contribution score
0.0
0.2
0.4
0.6
0.8
1.0
PC56
Ankle spacing width (left)
HTLV-1 gag antigen for Human T-Lymphotropic Virus 1
Bring up phlegm/sputum/mucus on most days
White blood cell (leukocyte) count
Bilateral oophorectomy (both ovaries removed)
Arm bone area (right)
Length of menstrual cycle
E7 antigen for Human Papillomavirus type-16
Ankle spacing width (right)
Neutrophill count
Ever taken cannabis
Diagnosed with coeliac disease or gluten sensitivity
gE / gI antigen for Varicella Zoster Virus
Hearing difficulty/problems with background noise
Systolic blood pressure, automated reading
Neutrophill percentage
Paracetamol use self-reported
Prospective memory result
Lymphocyte percentage
Risk taking
Diastolic blood pressure, automated reading
Seen doctor (GP) for nerves, anxiety, tension or depression
Doctor diagnosed asthma
Mouth ulcers

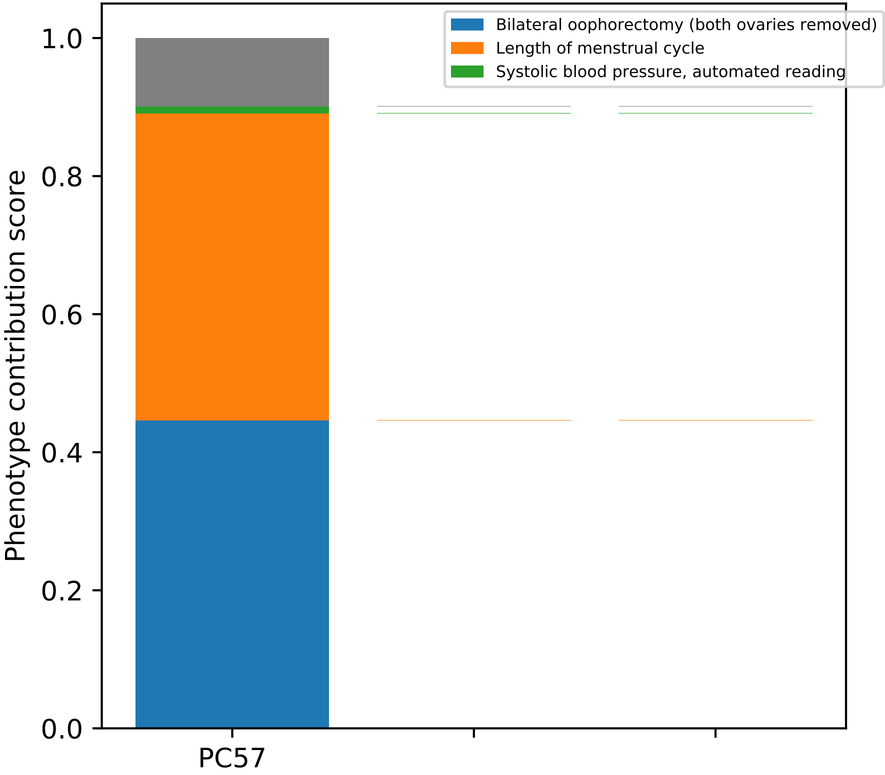

Bilateral oophorectomy (both ovaries removed)
Length of menstrual cycle
Systolic blood pressure, automated reading
Phenotype contribution score
0.0
0.2
0.4
0.6
0.8
1.0
PC57

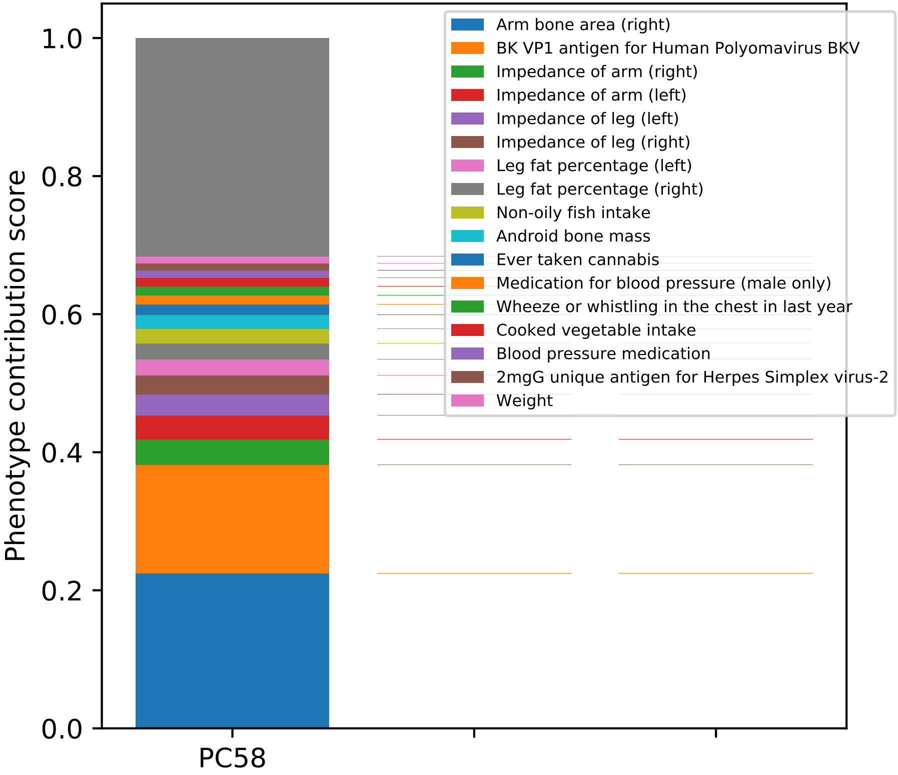

Arm bone area (right)
BK VP1 antigen for Human Polyomavirus BKV
Impedance of arm (right)
Impedance of arm (left)
Impedance of leg (left)
Impedance of leg (right)
Leg fat percentage (left)
Leg fat percentage (right)
Non-oily fish intake
Android bone mass
Ever taken cannabis
Medication for blood pressure (male only)
Wheeze or whistling in the chest in last year
Cooked vegetable intake
Blood pressure medication
2mgG unique antigen for Herpes Simplex virus-2
Weight
Phenotype contribution score
1.0
0.8
0.6
0.4
0.2
0.0
PC58

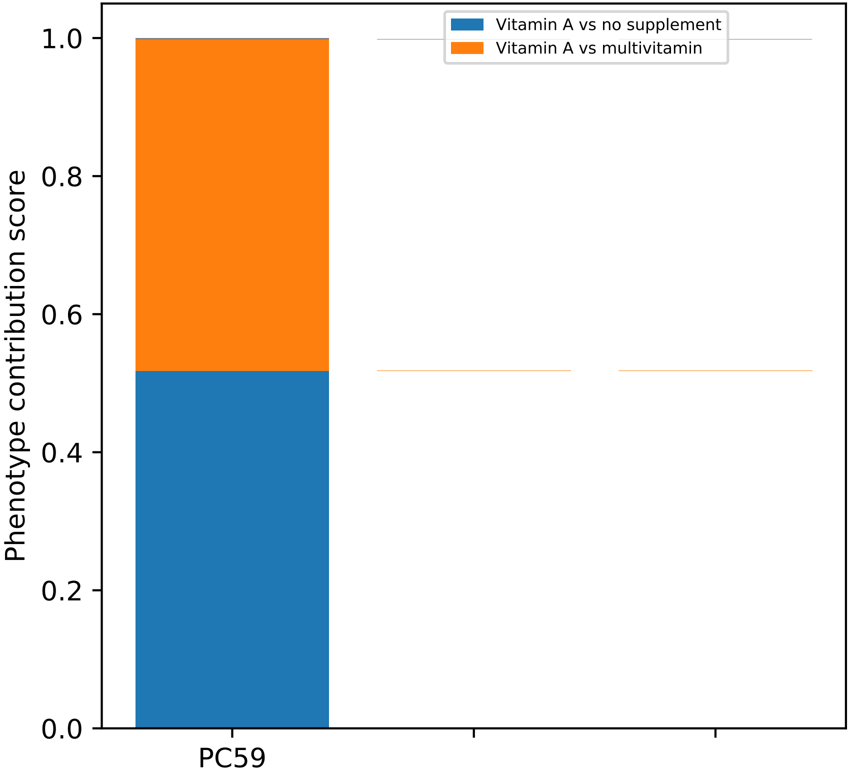

Vitamin A vs no supplement
Vitamin A vs multivitamin
1.0
0.8
0.6
0.4
0.2
0.0
Phenotype contribution score
PC59

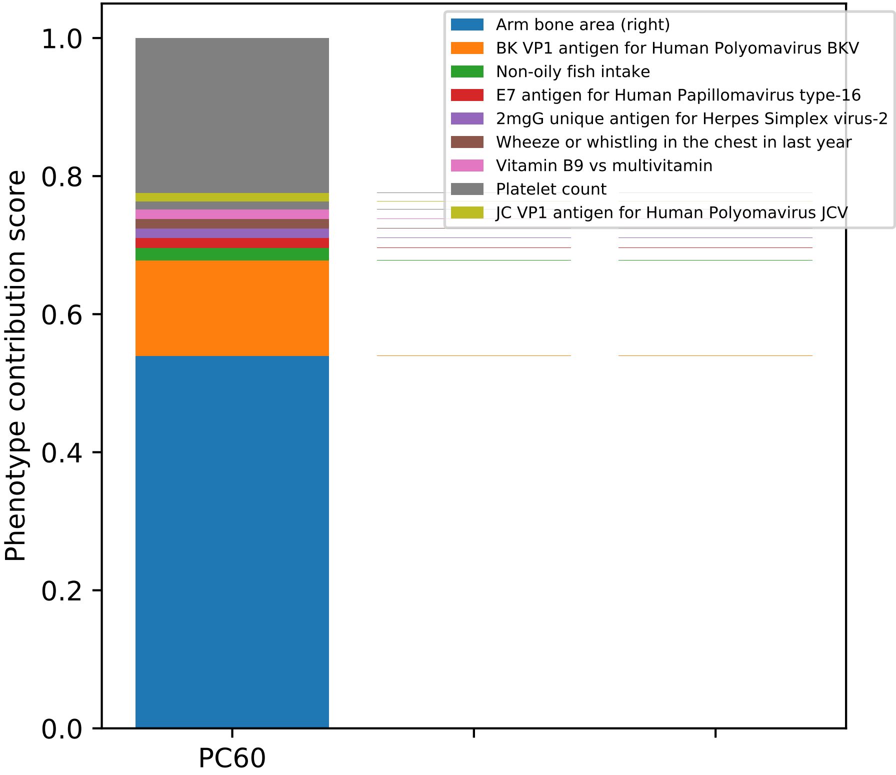

Arm bone area (right)
BK VP1 antigen for Human Polyomavirus BKV
Non-oily fish intake
E7 antigen for Human Papillomavirus type-16
2mgG unique antigen for Herpes Simplex virus-2
Wheeze or whistling in the chest in last year
Vitamin B9 vs multivitamin
Platelet count
JC VP1 antigen for Human Polyomavirus JCV
Phenotype contribution score
0.0
0.2
0.4
0.6
0.8
1.0
PC60

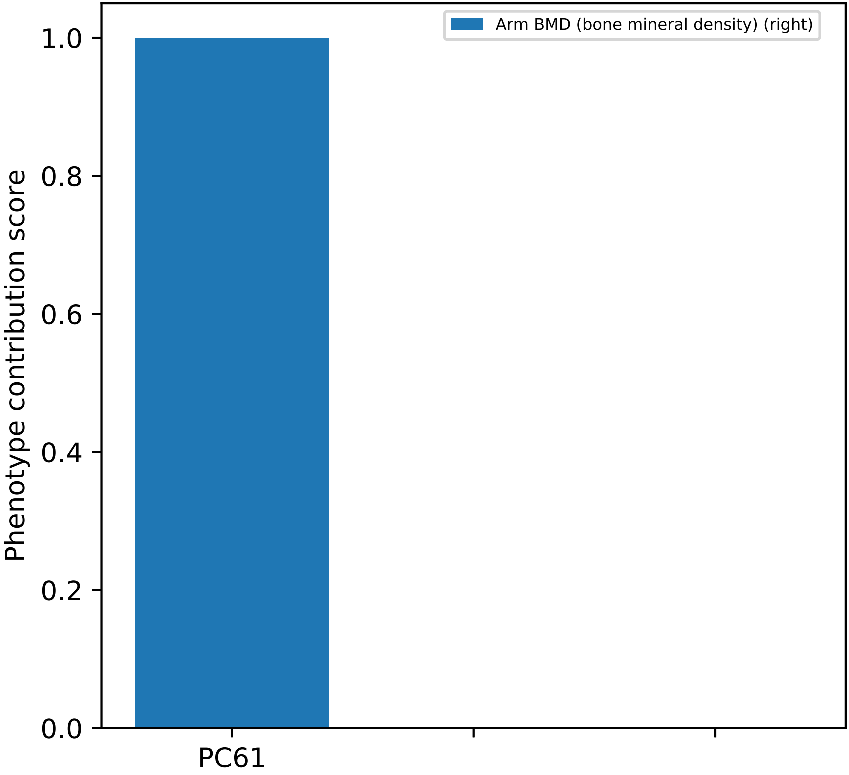
Arm BMD (bone mineral density) (right)
1.0
0.8
0.6
0.4
0.2
0.0
Phenotype contribution score
PC61

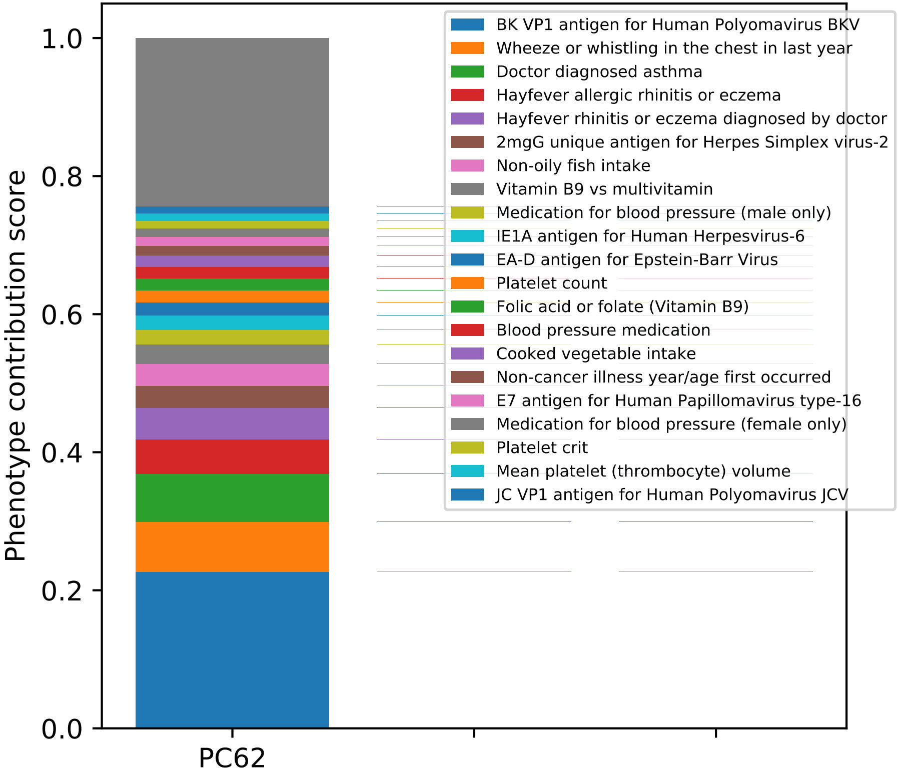

BK VP1 antigen for Human Polyomavirus BKV
Wheeze or whistling in the chest in last year
Doctor diagnosed asthma
Hayfever allergic rhinitis or eczema
Hayfever rhinitis or eczema diagnosed by doctor
2mgG unique antigen for Herpes Simplex virus-2
Non-oily fish intake
Vitamin B9 vs multivitamin
Medication for blood pressure (male only)
IE1A antigen for Human Herpesvirus-6
EA-D antigen for Epstein-Barr Virus
Platelet count
Folic acid or folate (Vitamin B9)
Blood pressure medication
Cooked vegetable intake
Non-cancer illness year/age first occurred
E7 antigen for Human Papillomavirus type-16
Medication for blood pressure (female only)
Platelet crit
Mean platelet (thrombocyte) volume
JC VP1 antigen for Human Polyomavirus JCV
Phenotype contribution score
1.0
0.8
0.6
0.4
0.2
0.0
PC62

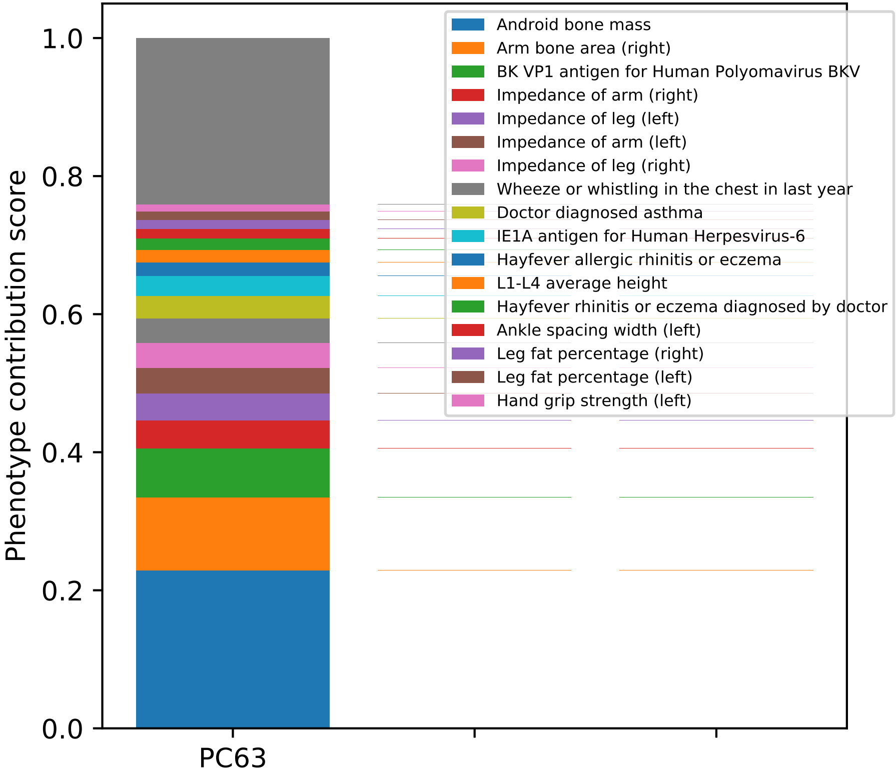

Phenotype contribution score
1.0
0.8
0.6
0.4
0.2
0.0
PC63
Android bone mass
Arm bone area (right)
BK VP1 antigen for Human Polyomavirus BKV
Impedance of arm (right)
Impedance of leg (left)
Impedance of arm (left)
Impedance of leg (right)
Wheeze or whistling in the chest in last year
Doctor diagnosed asthma
IE1A antigen for Human Herpesvirus-6
Hayfever allergic rhinitis or eczema
L1-L4 average height
Hayfever rhinitis or eczema diagnosed by doctor
Ankle spacing width (left)
Leg fat percentage (right)
Leg fat percentage (left)
Hand grip strength (left)

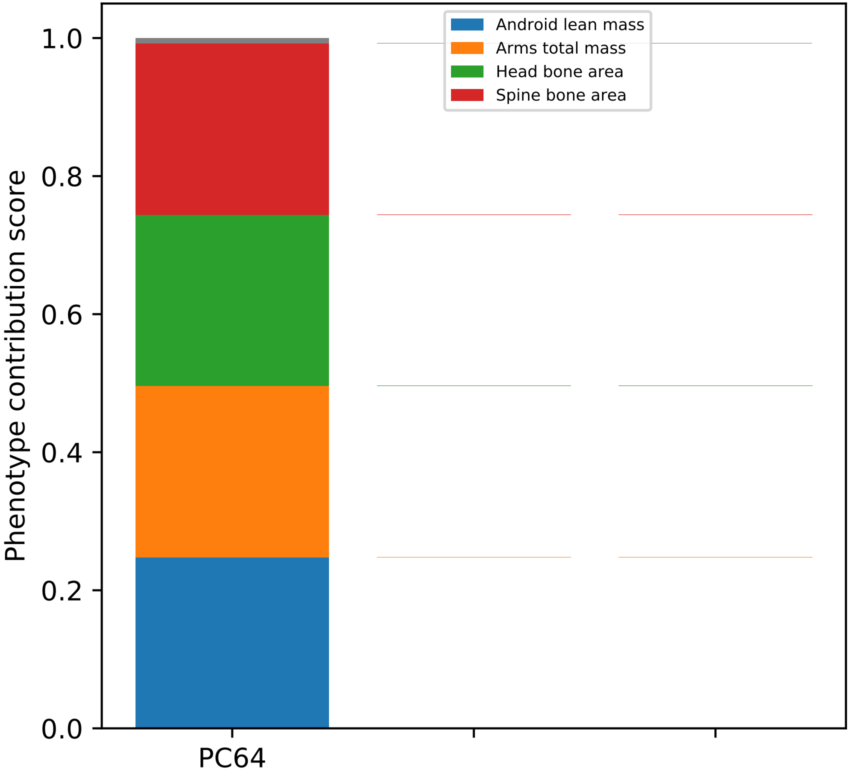

Android lean mass
Arms total mass
Head bone area
Spine bone area
Phenotype contribution score
1.0
0.8
0.6
0.4
0.2
0.0
PC64

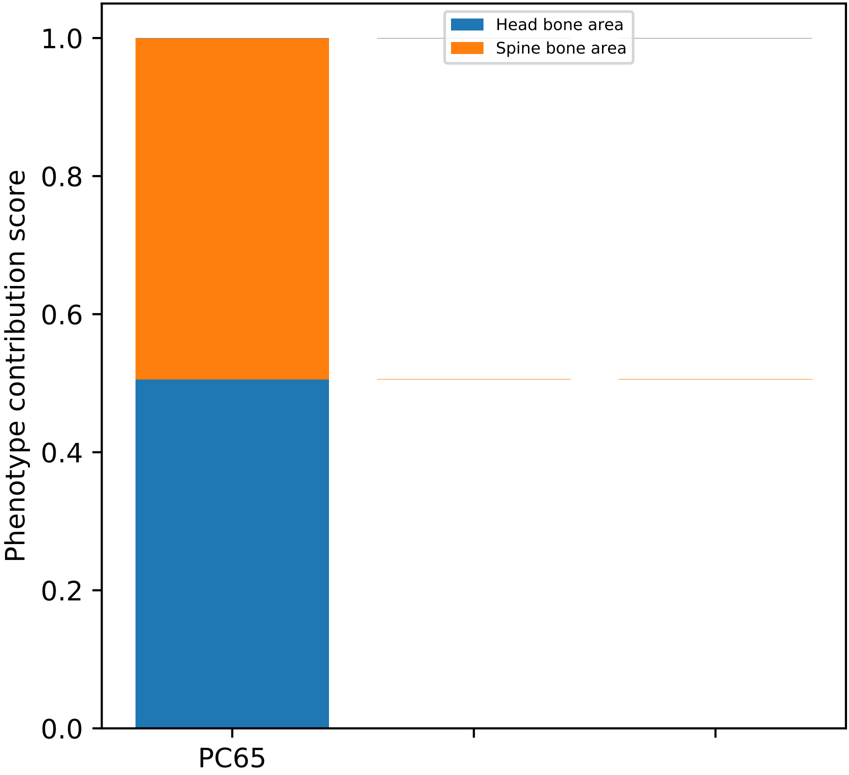

Head bone area
Spine bone area
Phenotype contribution score
1.0
0.8
0.6
0.4
0.2
0.0
PC65

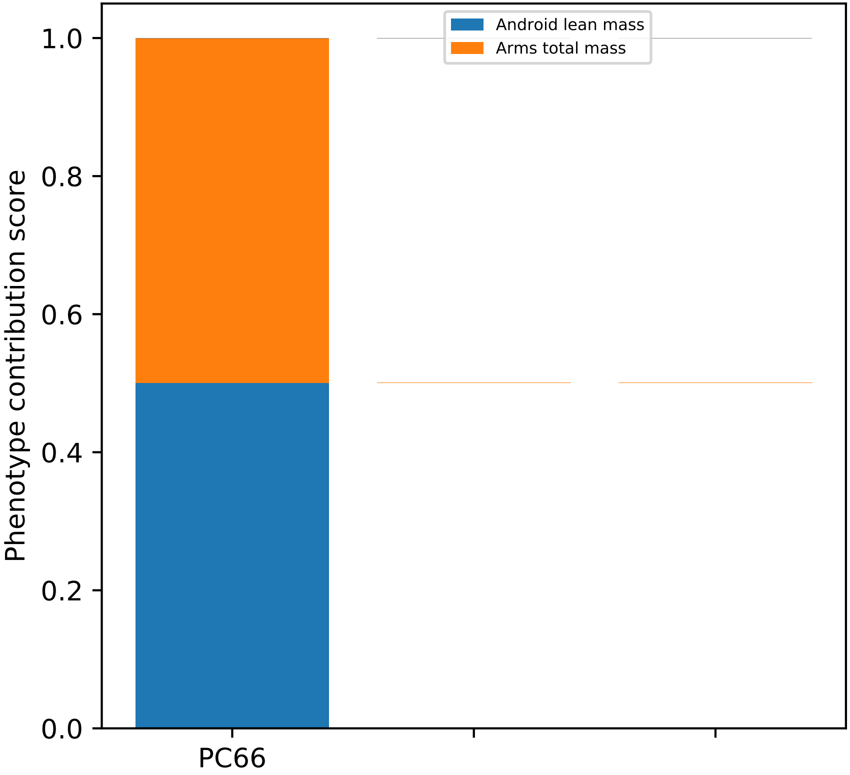
Android lean mass
Arms total mass
Phenotype contribution score
1.0
0.8
0.6
0.4
0.2
0.0
PC66

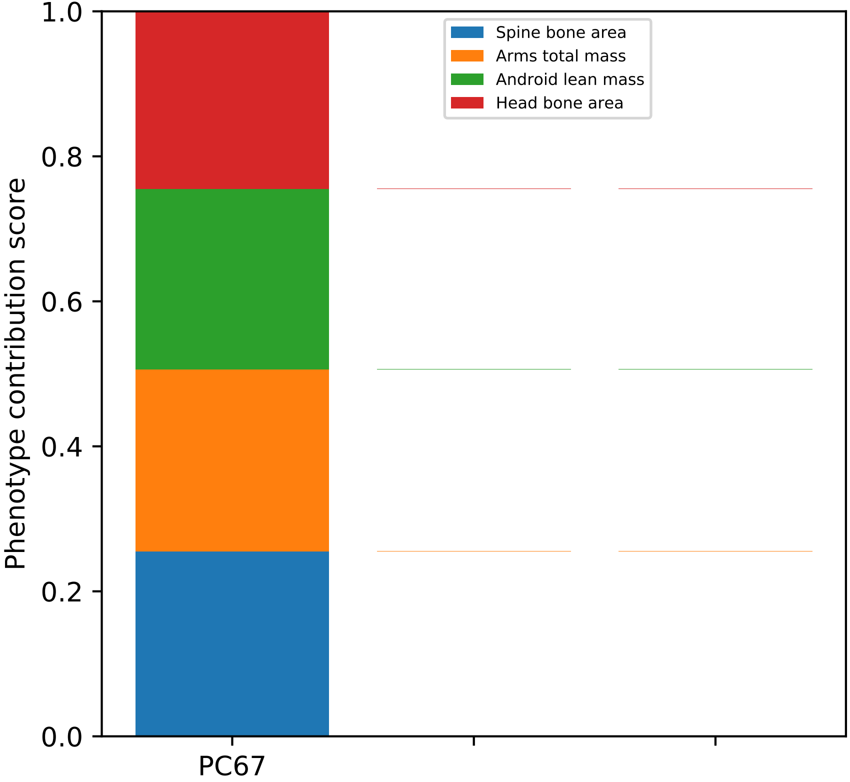

Spine bone area
Arms total mass
Android lean mass
Head bone area
Phenotype contribution score
1.0
0.8
0.6
0.4
0.2
0.0
PC67

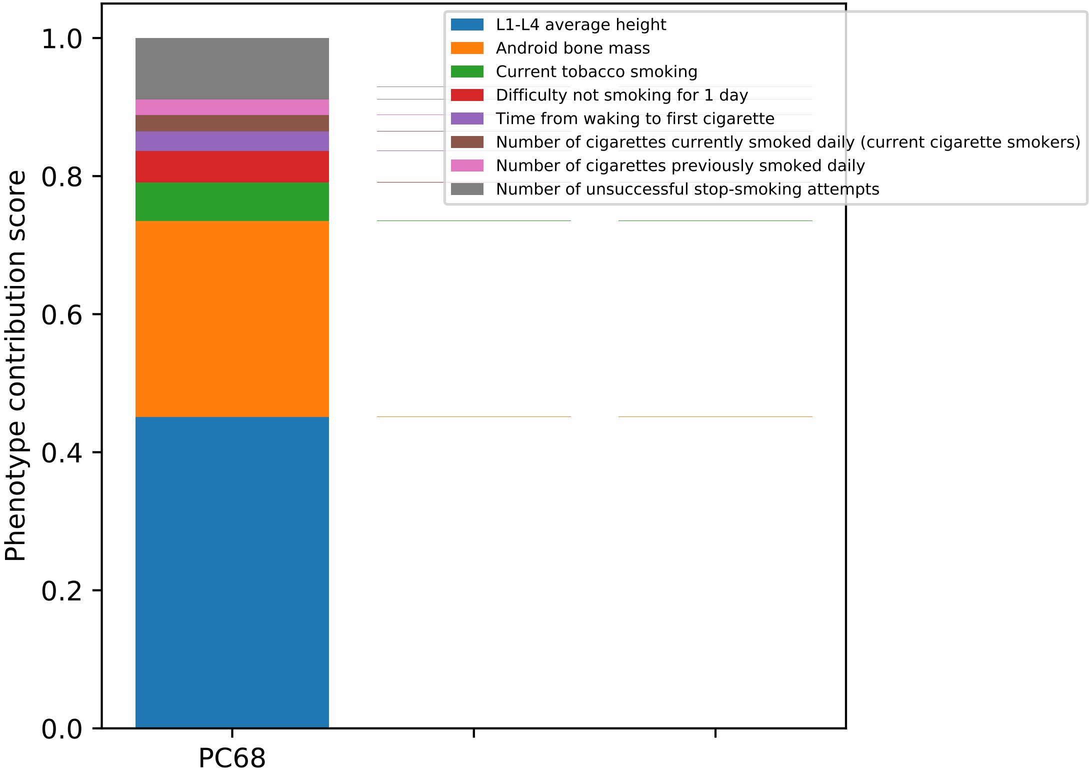
L1-L4 average height
Android bone mass
Current tobacco smoking
Difficulty not smoking for 1 day
Time from waking to first cigarette
Number of cigarettes currently smoked daily (current cigarette smokers)
Number of cigarettes previously smoked daily
Number of unsuccessful stop-smoking attempts
Phenotype contribution score
0.0
0.2
0.4
0.6
0.8
1.0
PC68

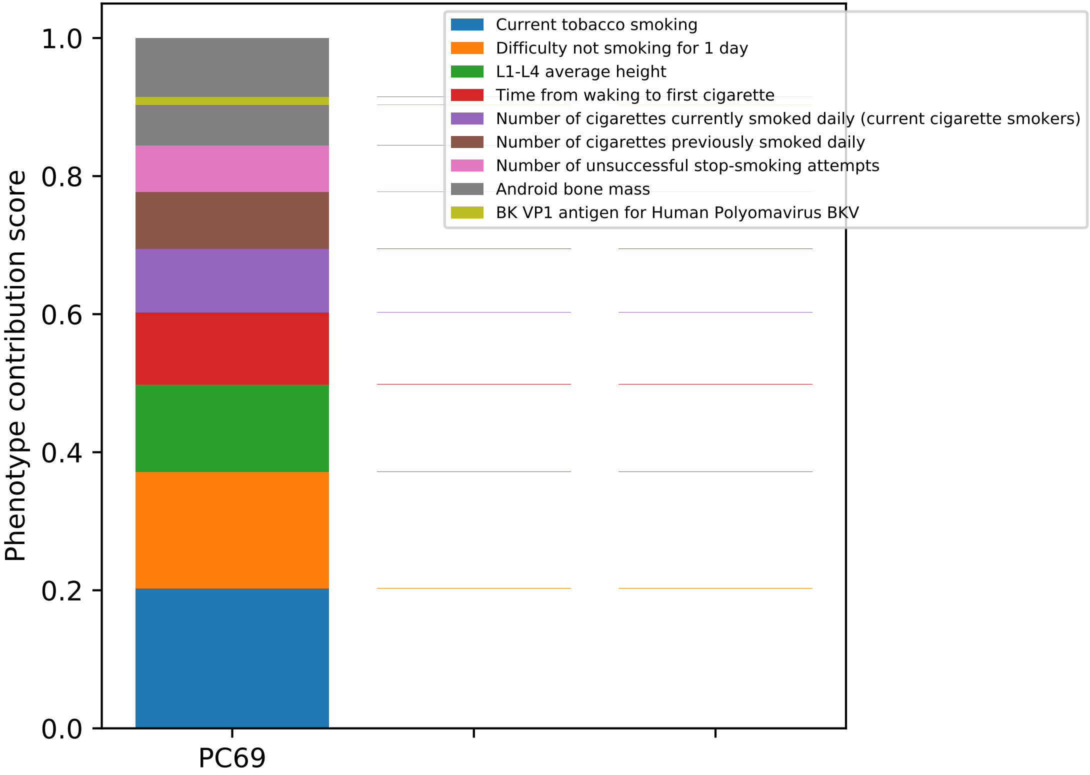
Current tobacco smoking
Difficulty not smoking for 1 day
L1-L4 average height
Time from waking to first cigarette
Number of cigarettes currently smoked daily (current cigarette smokers)
Number of cigarettes previously smoked daily
Number of unsuccessful stop-smoking attempts
Android bone mass
BK VP1 antigen for Human Polyomavirus BKV
Phenotype contribution score
1.0
0.8
0.6
0.4
0.2
0.0
PC69

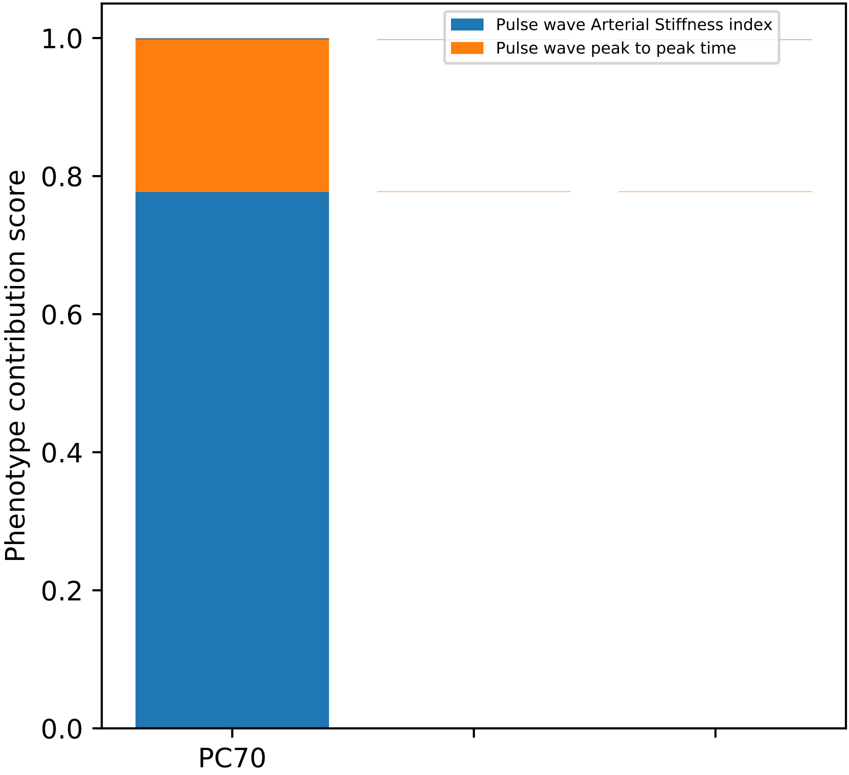

Pulse wave Arterial Stiffness index
Pulse wave peak to peak time
1.0
0.8
0.6
0.4
0.2
0.0
Phenotype contribution score
PC70

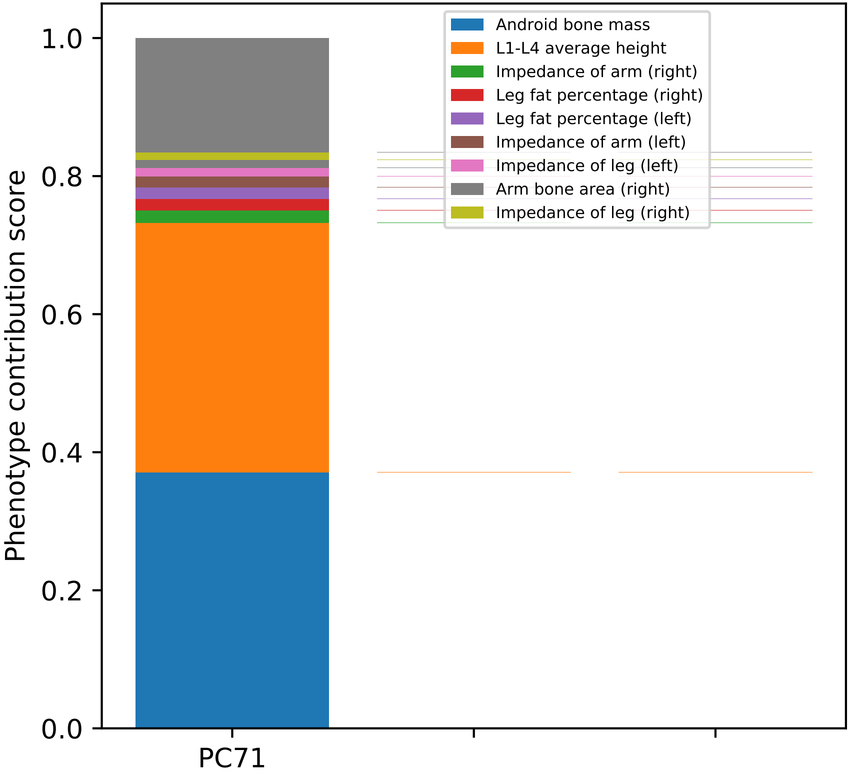

Android bone mass
L1-L4 average height
Impedance of arm (right)
Leg fat percentage (right)
Leg fat percentage (left)
Impedance of arm (left)
Impedance of leg (left)
Arm bone area (right)
Impedance of leg (right)
Phenotype contribution score
0.0
0.2
0.4
0.6
0.8
1.0
PC71

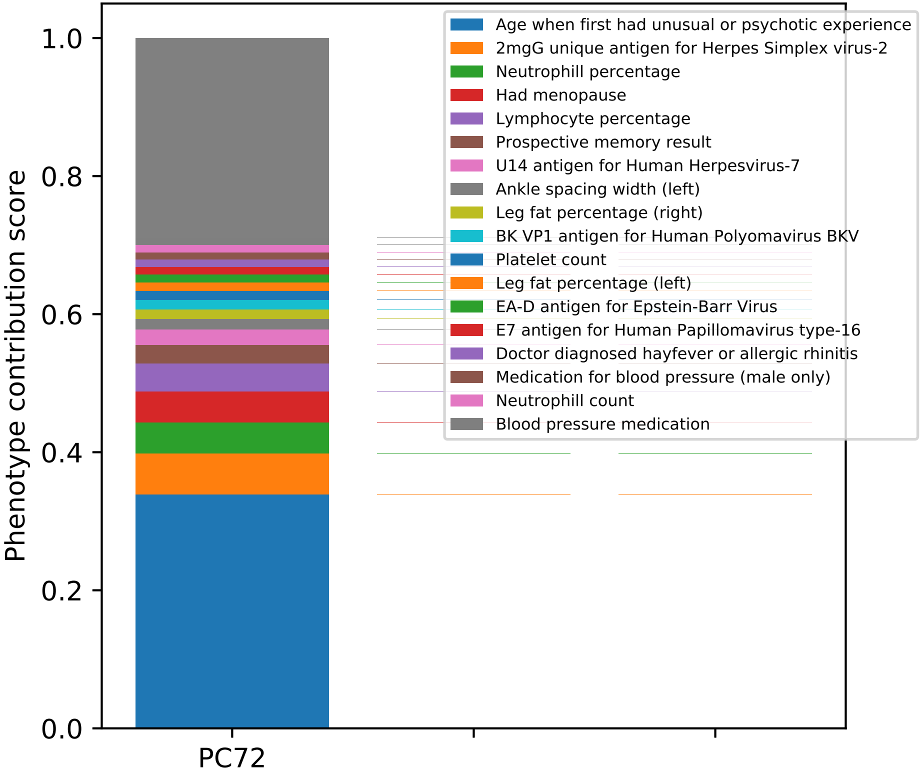

Age when first had unusual or psychotic experience
2mgG unique antigen for Herpes Simplex virus-2
Neutrophill percentage
Had menopause
Lymphocyte percentage
Prospective memory result
U14 antigen for Human Herpesvirus-7
Ankle spacing width (left)
Leg fat percentage (right)
BK VP1 antigen for Human Polyomavirus BKV
Platelet count
Leg fat percentage (left)
EA-D antigen for Epstein-Barr Virus
E7 antigen for Human Papillomavirus type-16
Doctor diagnosed hayfever or allergic rhinitis
Medication for blood pressure (male only)
Neutrophill count
Blood pressure medication
Phenotype contribution score
1.0
0.8
0.6
0.4
0.2
0.0
PC72

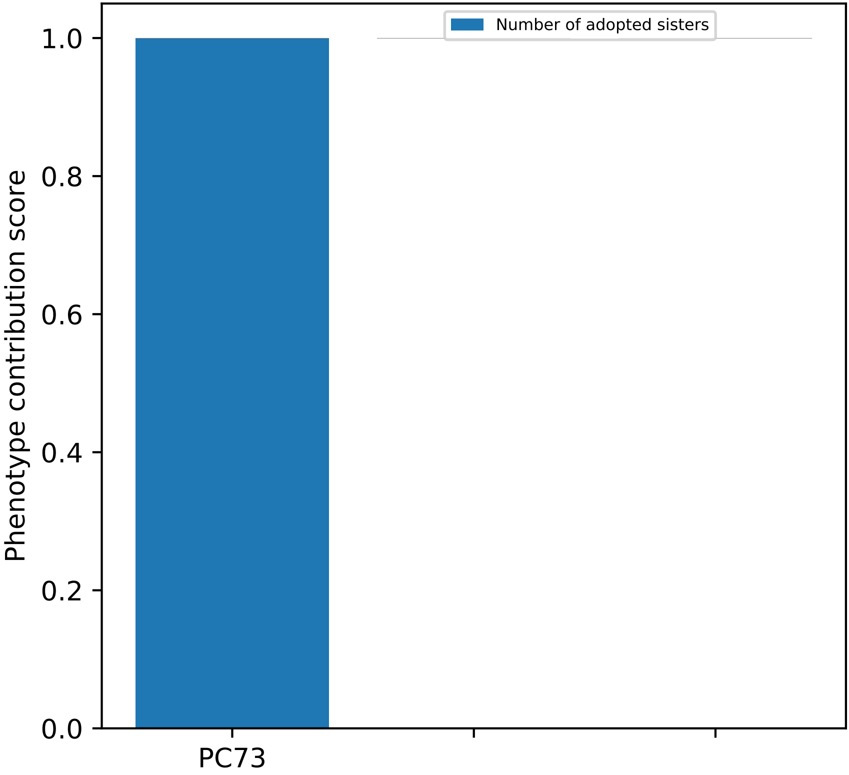
Number of adopted sisters
1.0
0.8
0.6
0.4
0.2
0.0
Phenotype contribution score
PC73

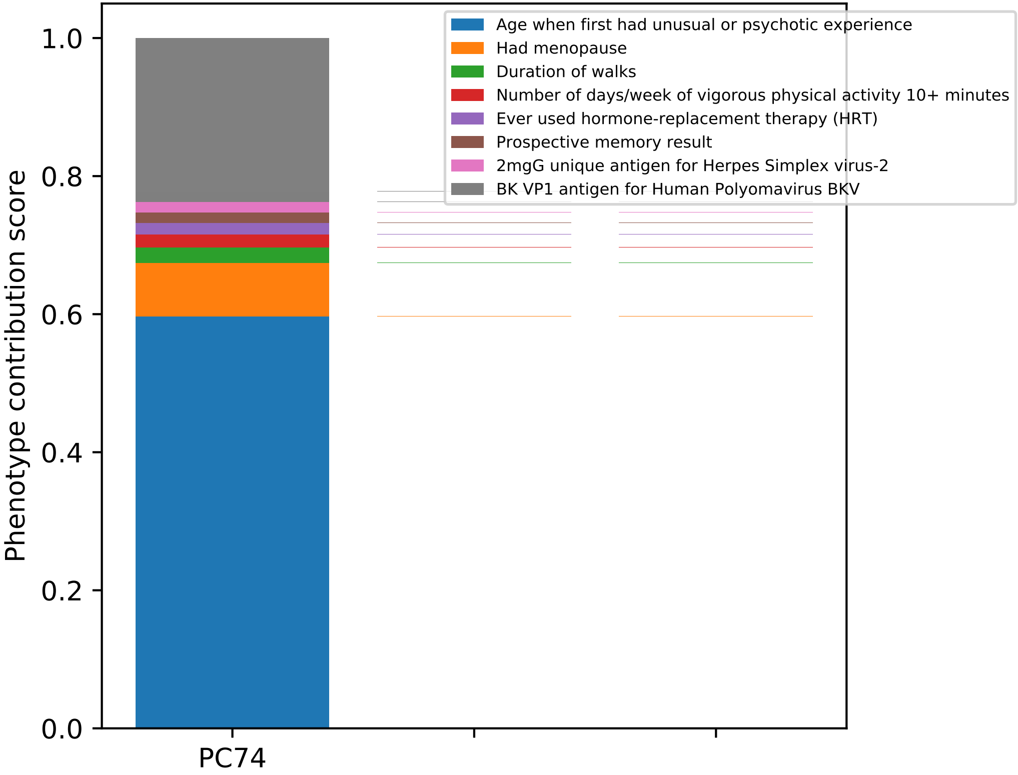

Phenotype contribution score
1.0
0.8
0.6
0.4
0.2
0.0
PC74
Age when first had unusual or psychotic experience
Had menopause
Duration of walks
Number of days/week of vigorous physical activity 10+ minutes
Ever used hormone-replacement therapy (HRT)
Prospective memory result
2mgG unique antigen for Herpes Simplex virus-2
BK VP1 antigen for Human Polyomavirus BKV

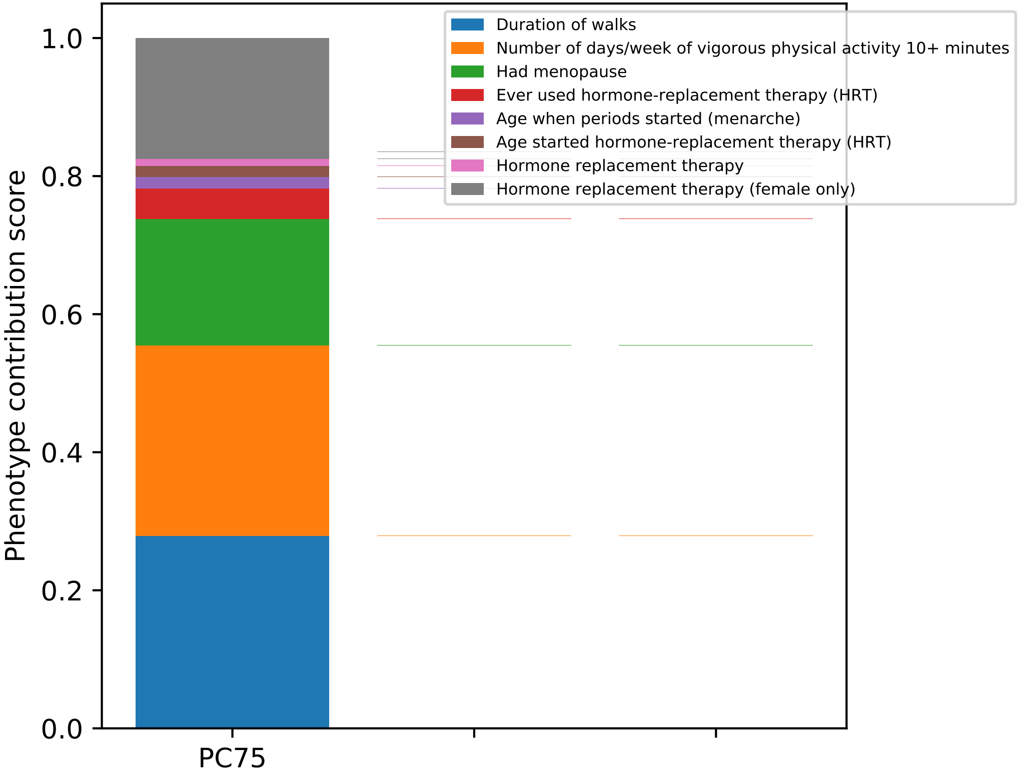

Duration of walks
Number of days/week of vigorous physical activity 10+ minutes
Had menopause
Ever used hormone-replacement therapy (HRT)
Age when periods started (menarche)
Age started hormone-replacement therapy (HRT)
Hormone replacement therapy
Hormone replacement therapy (female only)
Phenotype contribution score
1.0
0.8
0.6
0.4
0.2
0.0
PC75

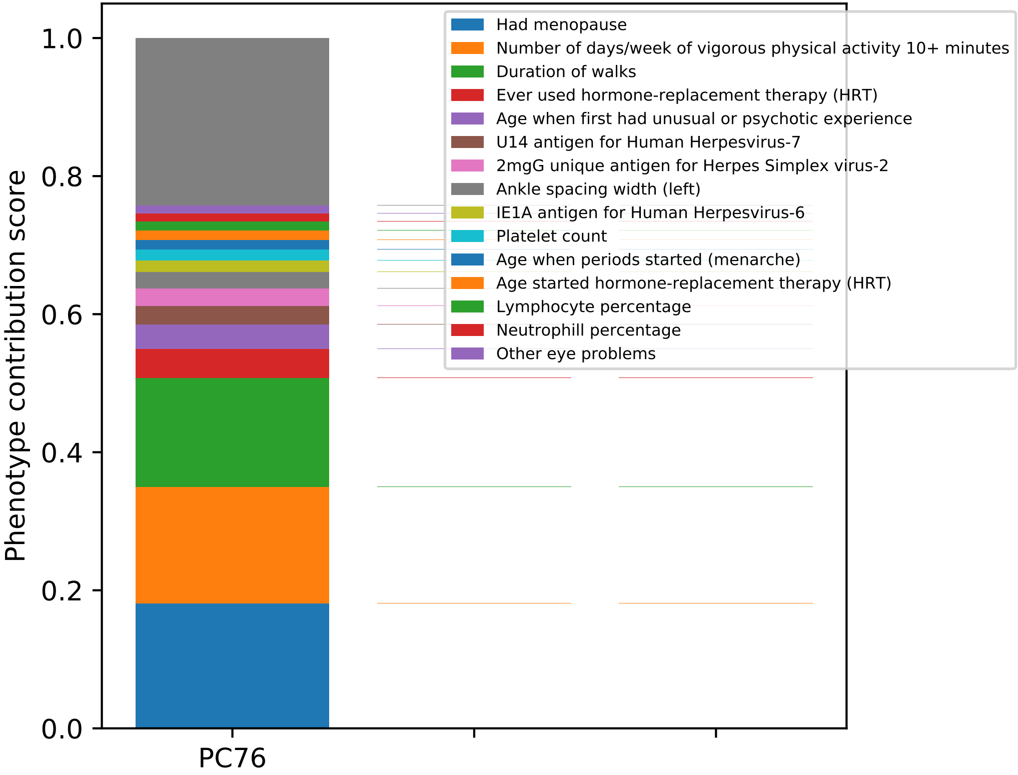

Phenotype contribution score
1.0
0.8
0.6
0.4
0.2
0.0
PC76
Had menopause
Number of days/week of vigorous physical activity 10+ minutes
Duration of walks
Ever used hormone-replacement therapy (HRT)
Age when first had unusual or psychotic experience
U14 antigen for Human Herpesvirus-7
2mgG unique antigen for Herpes Simplex virus-2
Ankle spacing width (left)
IE1A antigen for Human Herpesvirus-6
Platelet count
Age when periods started (menarche)
Age started hormone-replacement therapy (HRT)
Lymphocyte percentage
Neutrophill percentage
Other eye problems

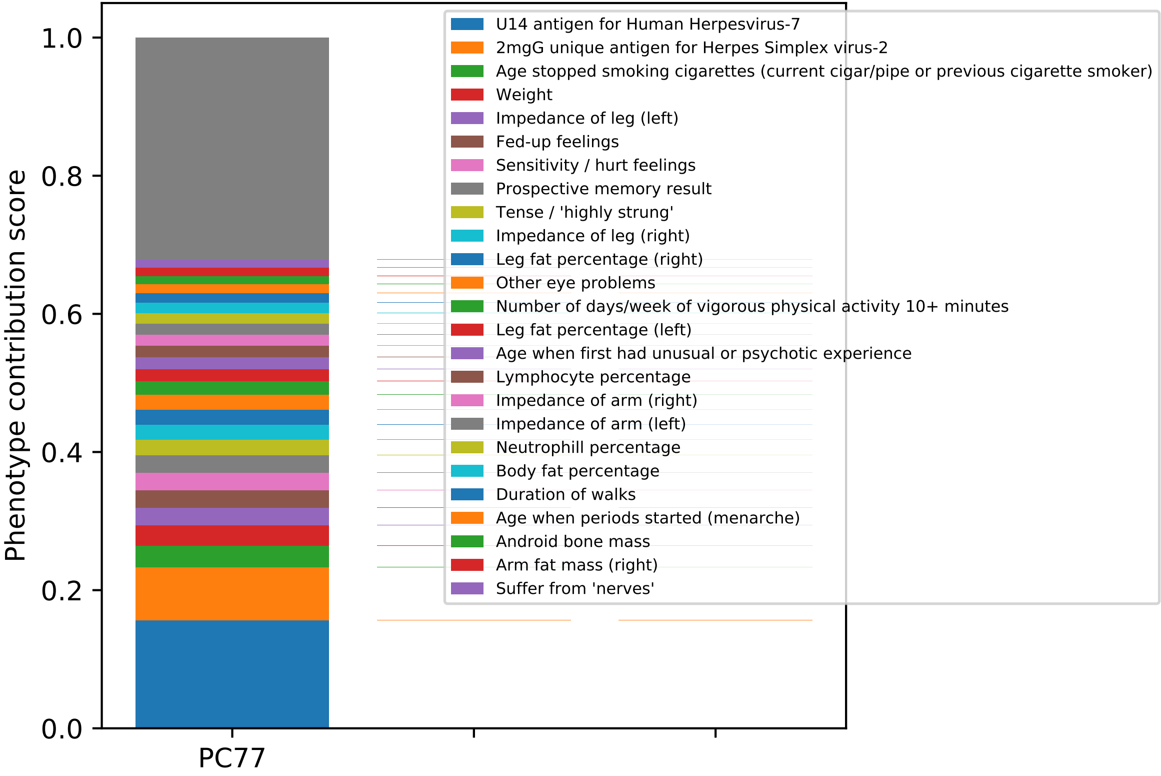

1.0
0.8
0.6
0.4
0.2
0.0
Phenotype contribution score
PC77
U14 antigen for Human Herpesvirus-7
2mgG unique antigen for Herpes Simplex virus-2
Age stopped smoking cigarettes (current cigar/pipe or previous cigarette smoker)
Weight
Impedance of leg (left)
Fed-up feelings
Sensitivity / hurt feelings
Prospective memory result
Tense / 'highly strung'
Impedance of leg (right)
Leg fat percentage (right)
Other eye problems
Number of days/week of vigorous physical activity 10+ minutes
Leg fat percentage (left)
Age when first had unusual or psychotic experience
Lymphocyte percentage
Impedance of arm (right)
Impedance of arm (left)
Neutrophill percentage
Body fat percentage
Duration of walks
Age when periods started (menarche)
Android bone mass
Arm fat mass (right)
Suffer from 'nerves'

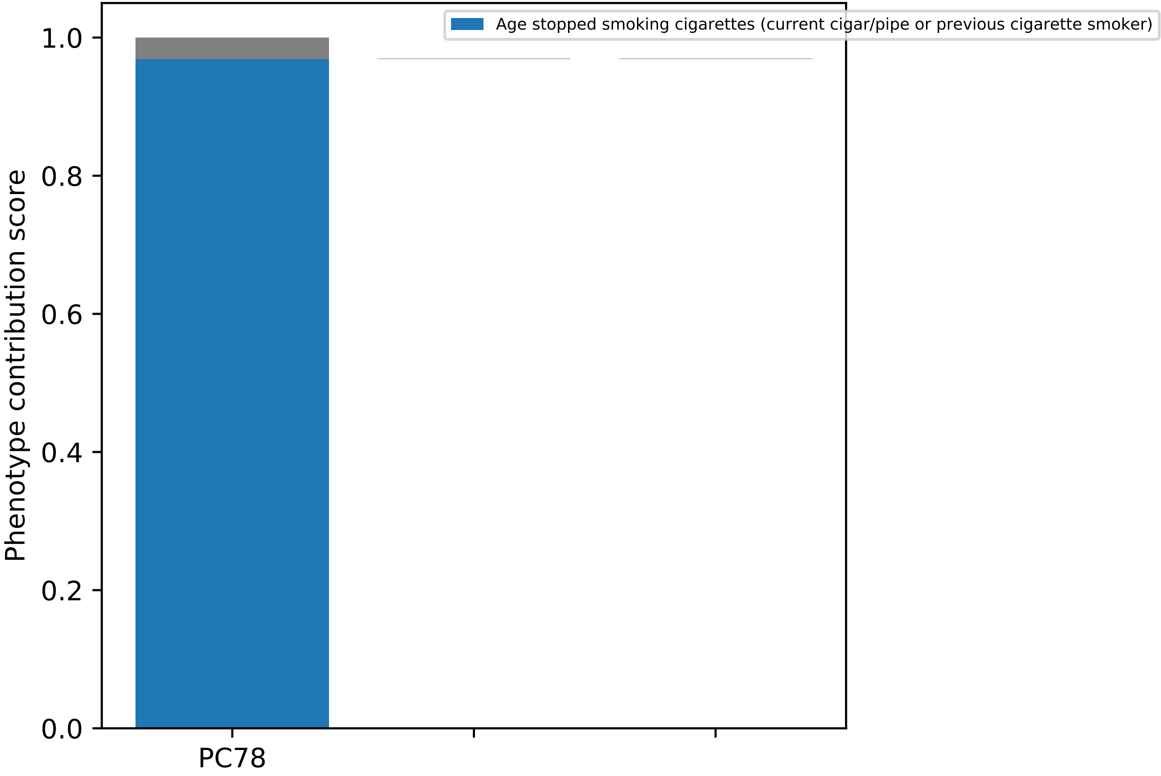

Age stopped smoking cigarettes (current cigar/pipe or previous cigarette smoker)
Phenotype contribution score
0.0
0.2
0.4
0.6
0.8
1.0
PC78

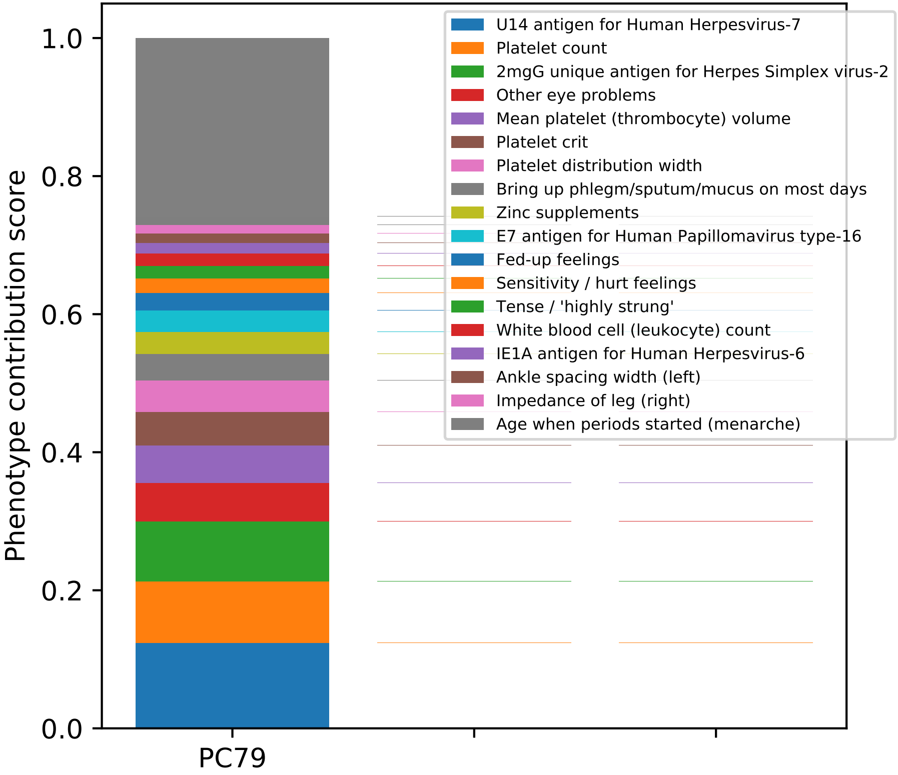

U14 antigen for Human Herpesvirus-7
Platelet count
2mgG unique antigen for Herpes Simplex virus-2
Other eye problems
Mean platelet (thrombocyte) volume
Platelet crit
Platelet distribution width
Bring up phlegm/sputum/mucus on most days
Zinc supplements
E7 antigen for Human Papillomavirus type-16
Fed-up feelings
Sensitivity / hurt feelings
Tense / 'highly strung'
White blood cell (leukocyte) count
IE1A antigen for Human Herpesvirus-6
Ankle spacing width (left)
Impedance of leg (right)
Age when periods started (menarche)
Phenotype contribution score
0.0
0.2
0.4
0.6
0.8
1.0
PC79

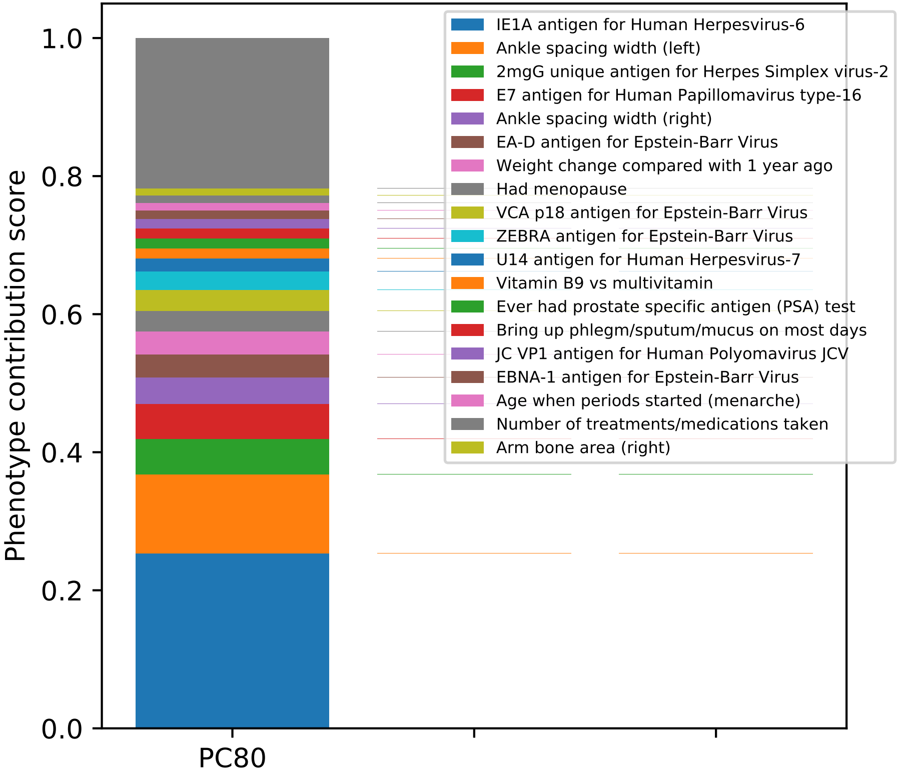

IE1A antigen for Human Herpesvirus-6
Ankle spacing width (left)
2mgG unique antigen for Herpes Simplex virus-2
E7 antigen for Human Papillomavirus type-16
Ankle spacing width (right)
EA-D antigen for Epstein-Barr Virus
Weight change compared with 1 year ago
Had menopause
VCA p18 antigen for Epstein-Barr Virus
ZEBRA antigen for Epstein-Barr Virus
U14 antigen for Human Herpesvirus-7
Vitamin B9 vs multivitamin
Ever had prostate specific antigen (PSA) test
Bring up phlegm/sputum/mucus on most days
JC VP1 antigen for Human Polyomavirus JCV
EBNA-1 antigen for Epstein-Barr Virus
Age when periods started (menarche)
Number of treatments/medications taken
Arm bone area (right)
Phenotype contribution score
1.0
0.8
0.6
0.4
0.2
0.0
PC80

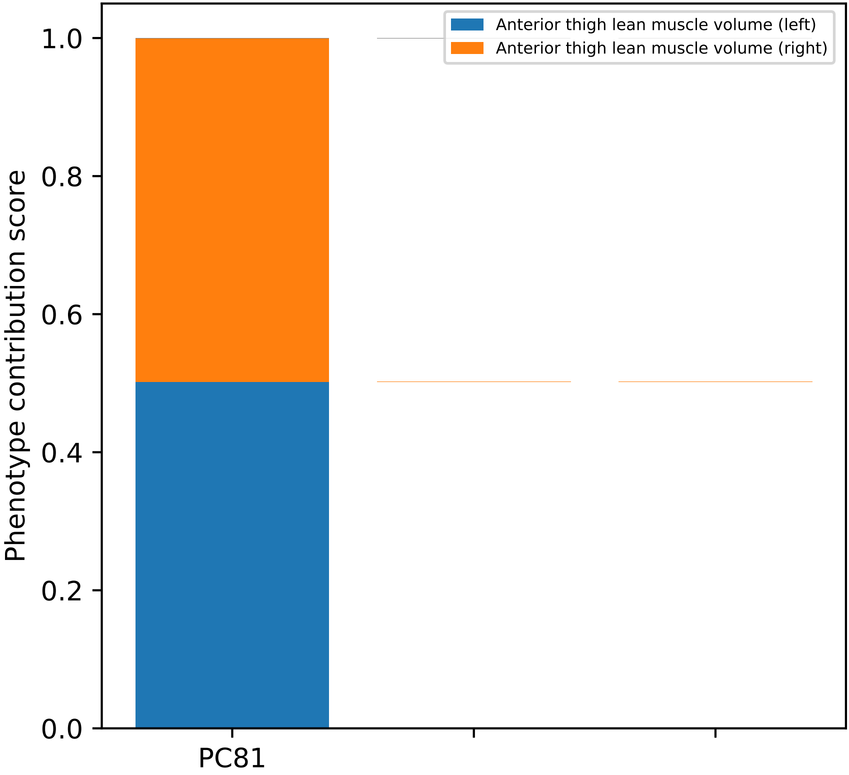

Anterior thigh lean muscle volume (left)
Anterior thigh lean muscle volume (right)
Phenotype contribution score
1.0
0.8
0.6
0.4
0.2
0.0
PC81

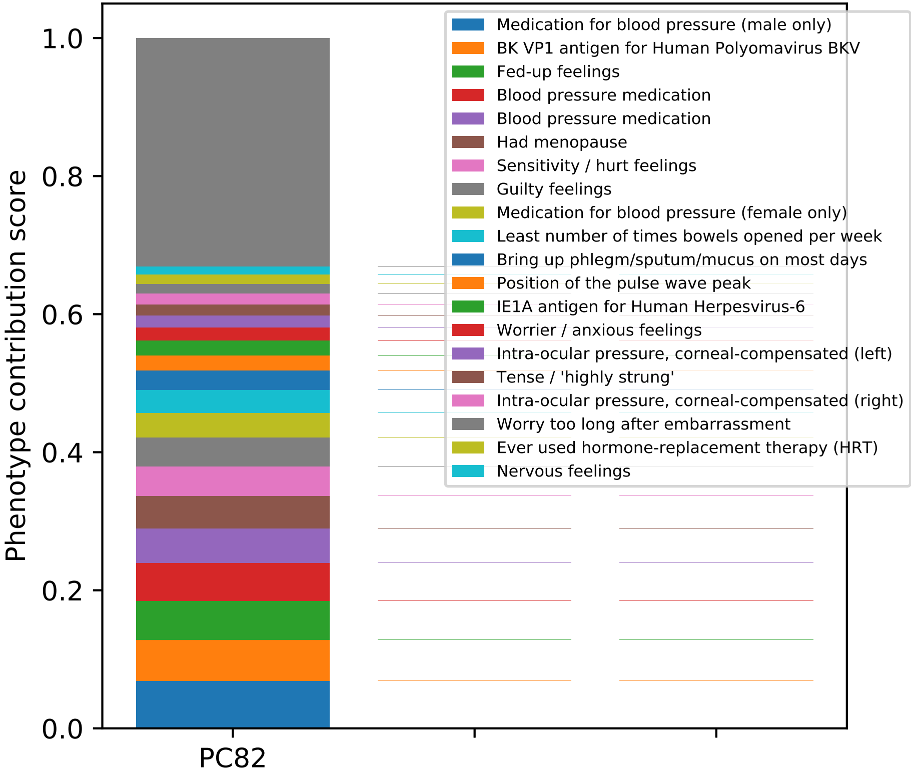

Phenotype contribution score
1.0
0.8
0.6
0.4
0.2
0.0
PC82
Medication for blood pressure (male only)
BK VP1 antigen for Human Polyomavirus BKV
Fed-up feelings
Blood pressure medication
Blood pressure medication
Had menopause
Sensitivity / hurt feelings
Guilty feelings
Medication for blood pressure (female only)
Least number of times bowels opened per week
Bring up phlegm/sputum/mucus on most days
Position of the pulse wave peak
IE1A antigen for Human Herpesvirus-6
Worrier / anxious feelings
Intra-ocular pressure, corneal-compensated (left)
Tense / 'highly strung'
Intra-ocular pressure, corneal-compensated (right)
Worry too long after embarrassment
Ever used hormone-replacement therapy (HRT)
Nervous feelings

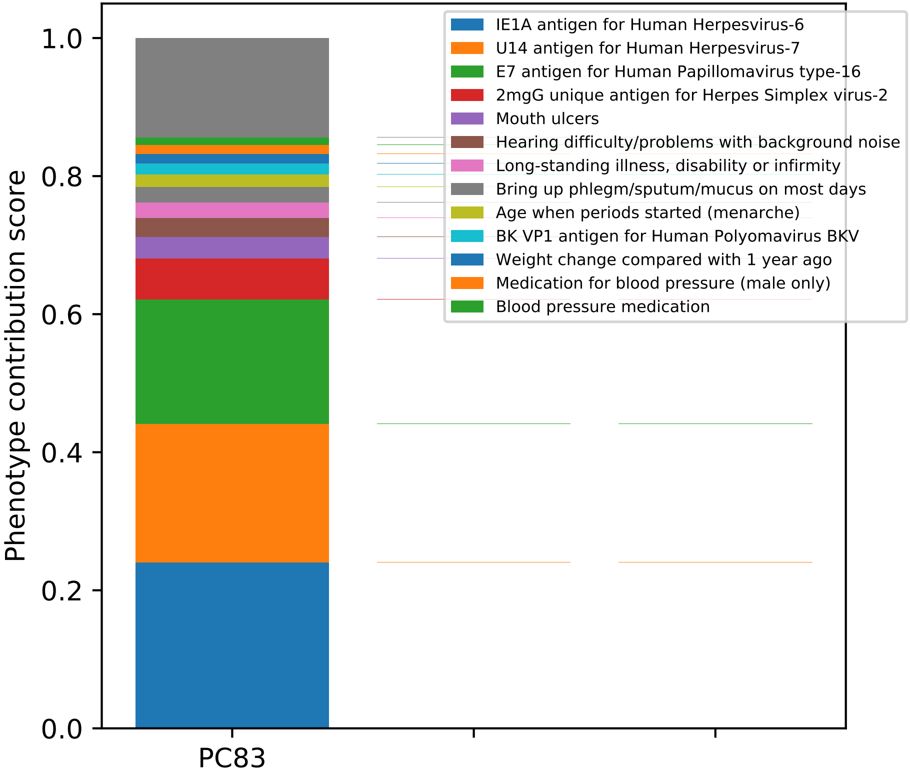

IE1A antigen for Human Herpesvirus-6
U14 antigen for Human Herpesvirus-7
E7 antigen for Human Papillomavirus type-16
2mgG unique antigen for Herpes Simplex virus-2
Mouth ulcers
Hearing difficulty/problems with background noise
Long-standing illness, disability or infirmity
Bring up phlegm/sputum/mucus on most days
Age when periods started (menarche)
BK VP1 antigen for Human Polyomavirus BKV
Weight change compared with 1 year ago
Medication for blood pressure (male only)
Blood pressure medication
Phenotype contribution score
1.0
0.8
0.6
0.4
0.2
0.0
PC83

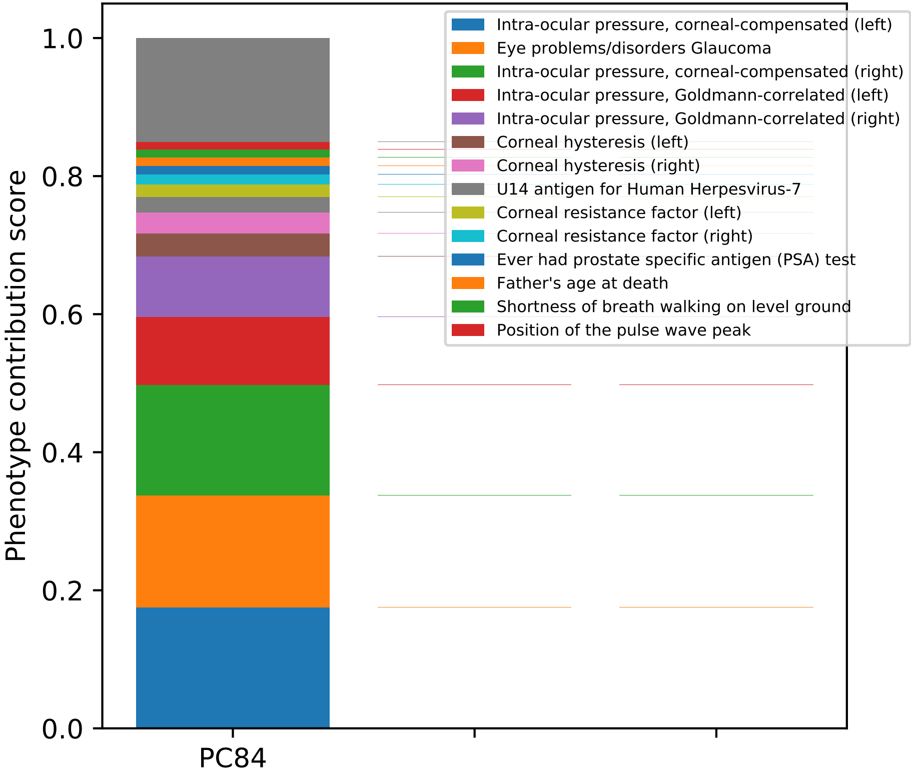

Phenotype contribution score
0.0
0.2
0.4
0.6
0.8
1.0
PC84
Intra-ocular pressure, corneal-compensated (left)
Eye problems/disorders Glaucoma
Intra-ocular pressure, corneal-compensated (right)
Intra-ocular pressure, Goldmann-correlated (left)
Intra-ocular pressure, Goldmann-correlated (right)
Corneal hysteresis (left)
Corneal hysteresis (right)
U14 antigen for Human Herpesvirus-7
Corneal resistance factor (left)
Corneal resistance factor (right)
Ever had prostate specific antigen (PSA) test
Father's age at death
Shortness of breath walking on level ground
Position of the pulse wave peak

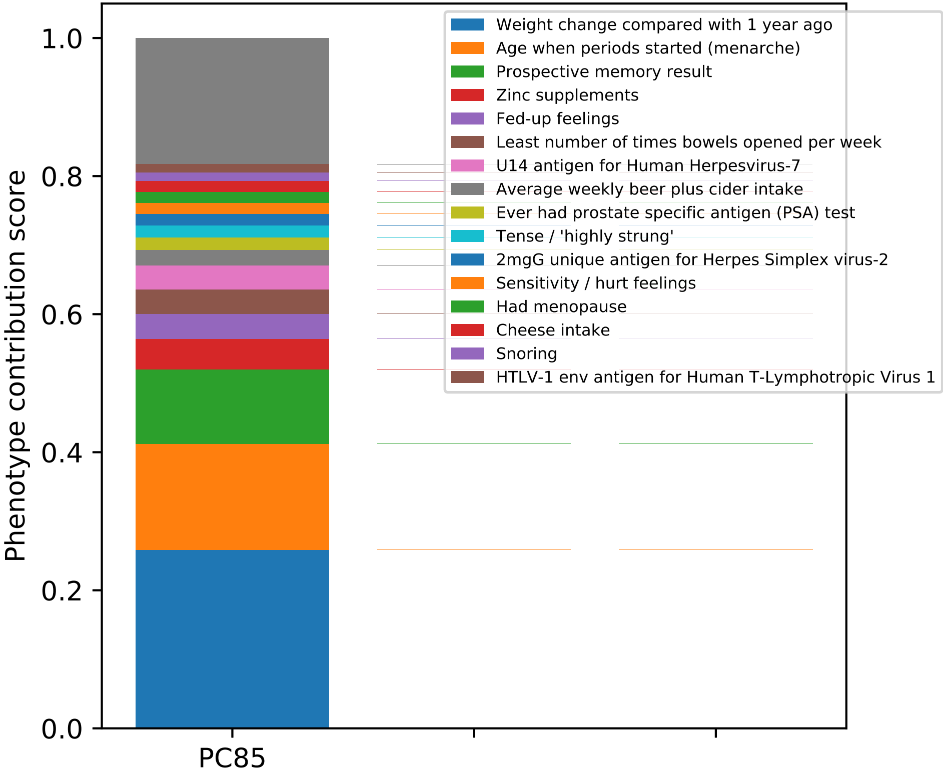

Phenotype contribution score
1.0
0.8
0.6
0.4
0.2
0.0
PC85
Weight change compared with 1 year ago
Age when periods started (menarche)
Prospective memory result
Zinc supplements
Fed-up feelings
Least number of times bowels opened per week
U14 antigen for Human Herpesvirus-7
Average weekly beer plus cider intake
Ever had prostate specific antigen (PSA) test
Tense / 'highly strung'
2mgG unique antigen for Herpes Simplex virus-2
Sensitivity / hurt feelings
Had menopause
Cheese intake
Snoring
HTLV-1 env antigen for Human T-Lymphotropic Virus 1

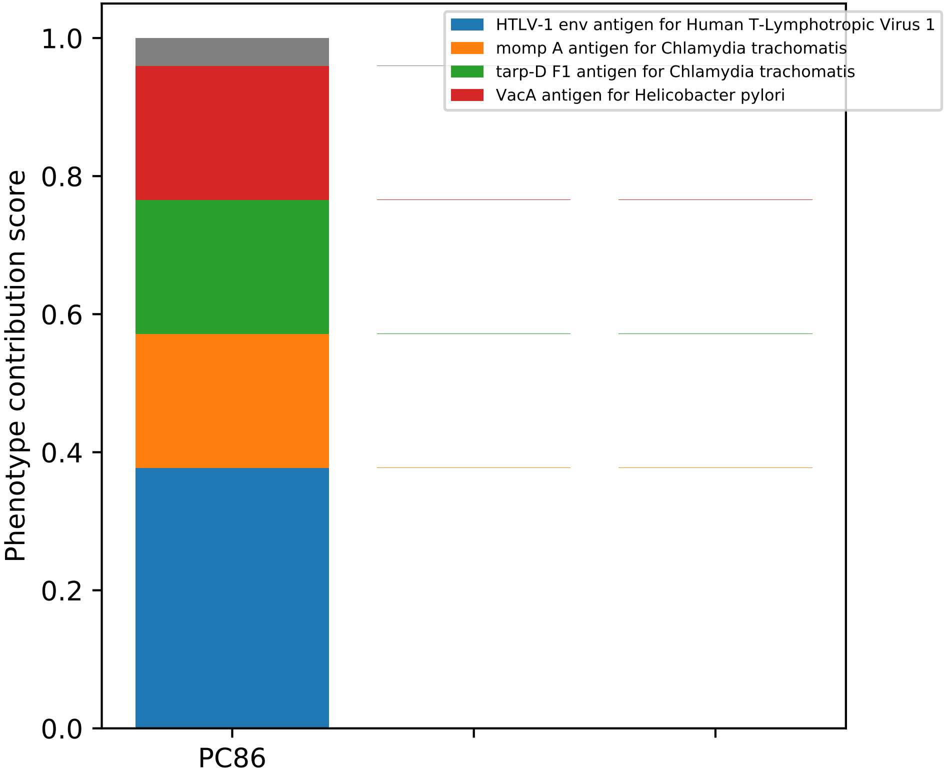

HTLV-1 env antigen for Human T-Lymphotropic Virus 1
momp A antigen for Chlamydia trachomatis
tarp-D F1 antigen for Chlamydia trachomatis
VacA antigen for Helicobacter pylori
Phenotype contribution score
1.0
0.8
0.6
0.4
0.2
0.0
PC86

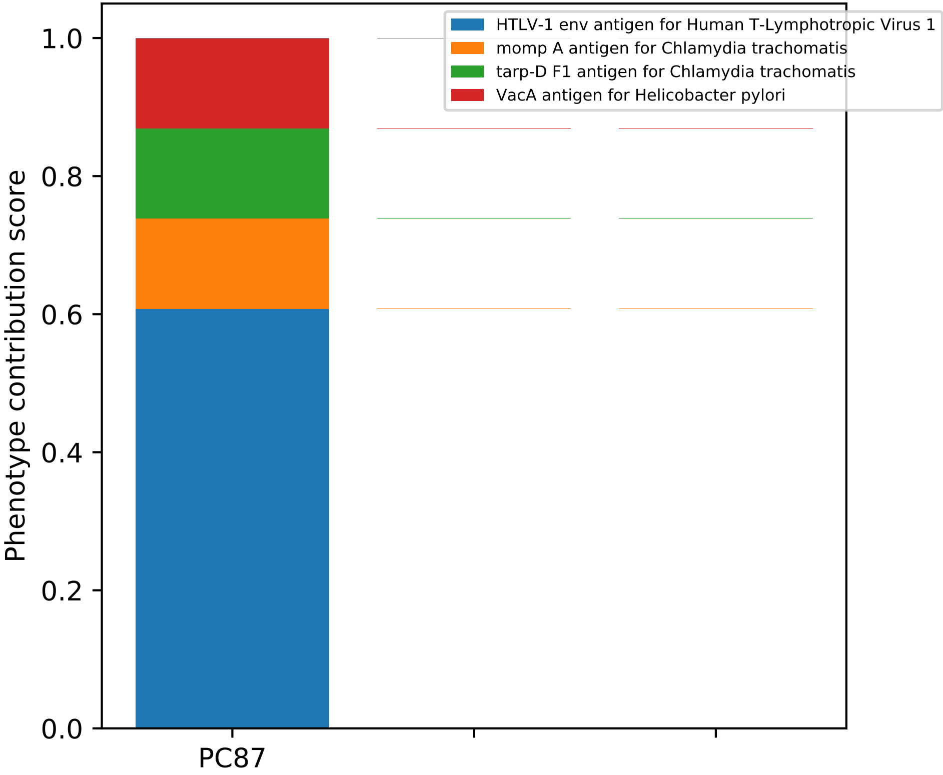

HTLV-1 env antigen for Human T-Lymphotropic Virus 1
momp A antigen for Chlamydia trachomatis
tarp-D F1 antigen for Chlamydia trachomatis
VacA antigen for Helicobacter pylori
Phenotype contribution score
1.0
0.8
0.6
0.4
0.2
0.0
PC87

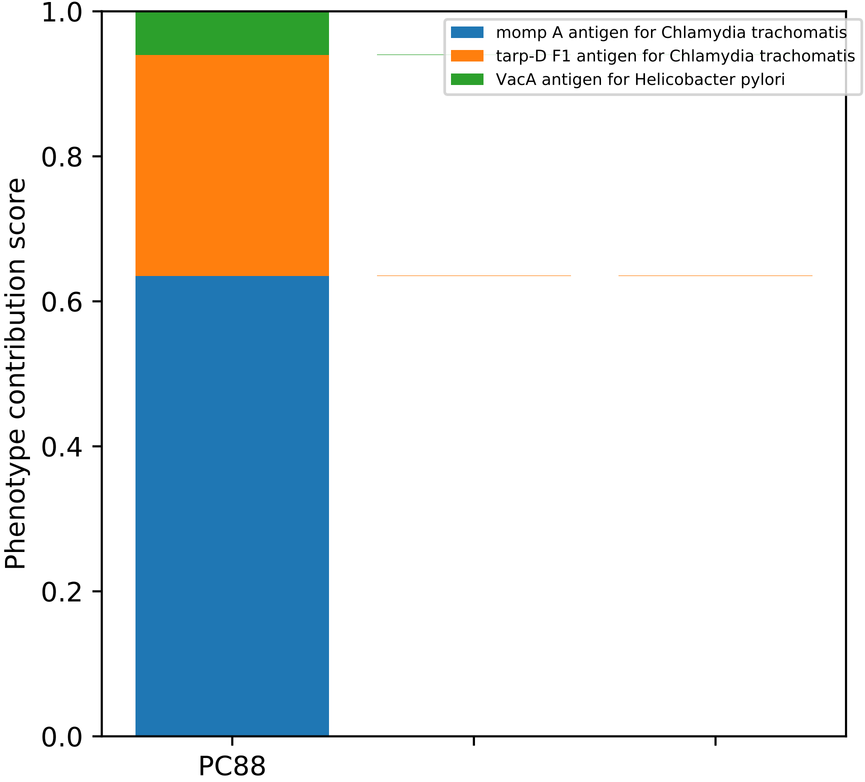

1.0
0.8
0.6
0.4
0.2
0.0
Phenotype contribution score
PC88
momp A antigen for Chlamydia trachomatis
tarp-D F1 antigen for Chlamydia trachomatis
VacA antigen for Helicobacter pylori

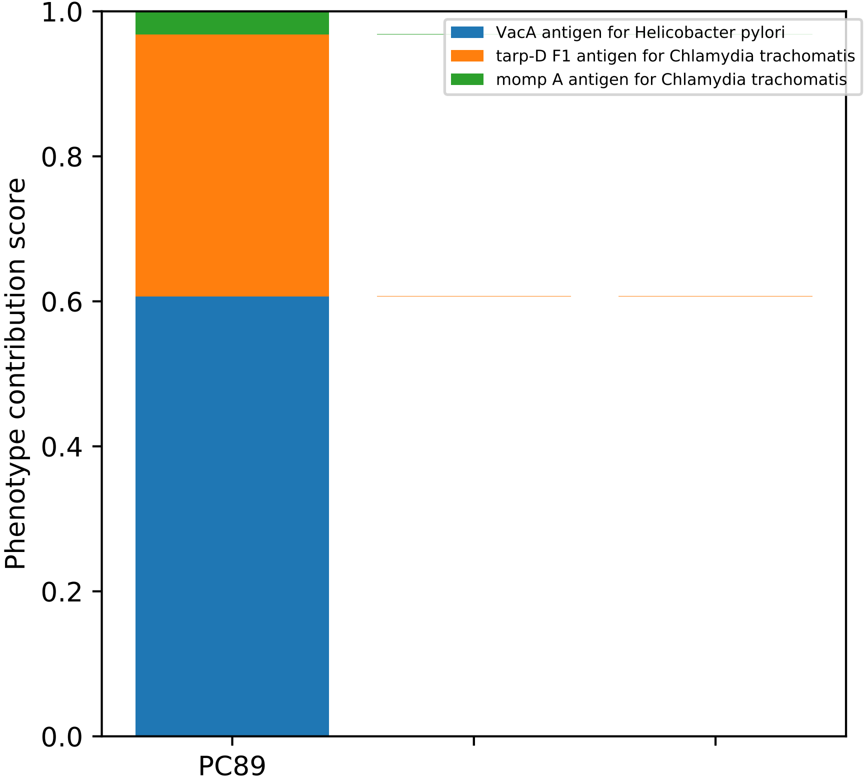
Phenotype contribution score
1.0
0.8
0.6
0.4
0.2
0.0
PC89
VacA antigen for Helicobacter pylori
tarp-D F1 antigen for Chlamydia trachomatis
momp A antigen for Chlamydia trachomatis

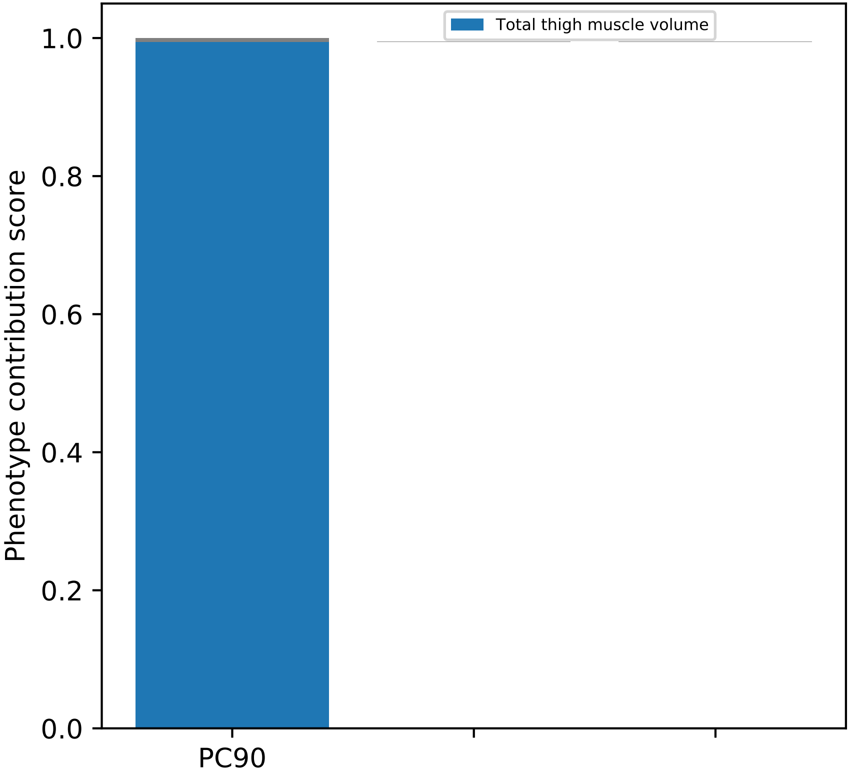
Total thigh muscle volume
Phenotype contribution score
0.0
0.2
0.4
0.6
0.8
1.0
PC90

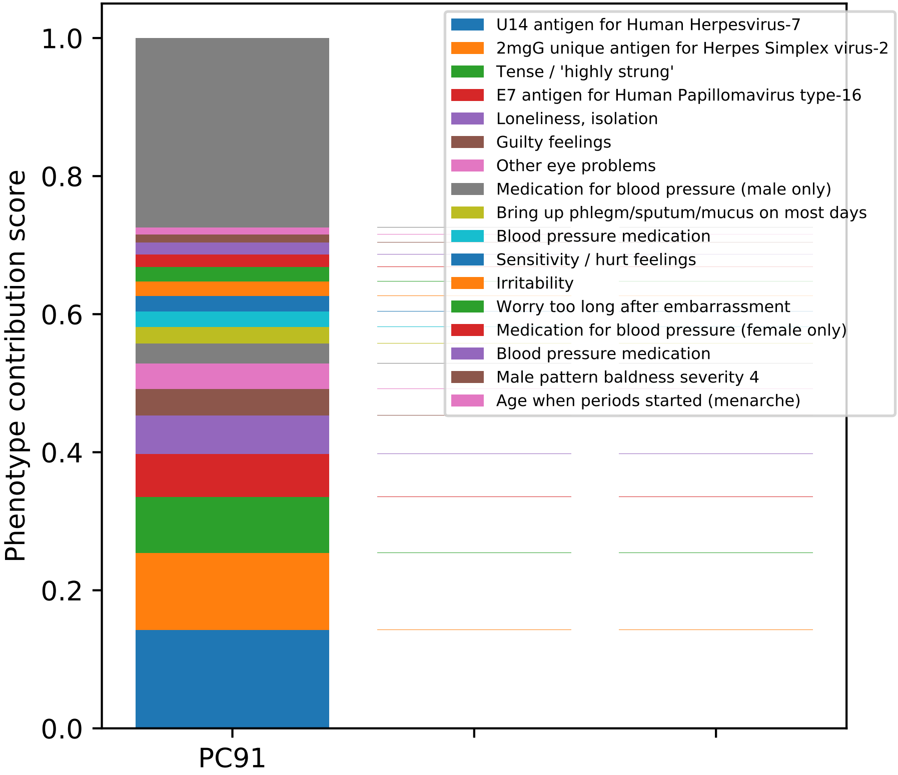

Phenotype contribution score
1.0
0.8
0.6
0.4
0.2
0.0
PC91
U14 antigen for Human Herpesvirus-7
2mgG unique antigen for Herpes Simplex virus-2
Tense / 'highly strung'
E7 antigen for Human Papillomavirus type-16
Loneliness, isolation
Guilty feelings
Other eye problems
Medication for blood pressure (male only)
Bring up phlegm/sputum/mucus on most days
Blood pressure medication
Sensitivity / hurt feelings
Irritability
Worry too long after embarrassment
Medication for blood pressure (female only)
Blood pressure medication
Male pattern baldness severity 4
Age when periods started (menarche)

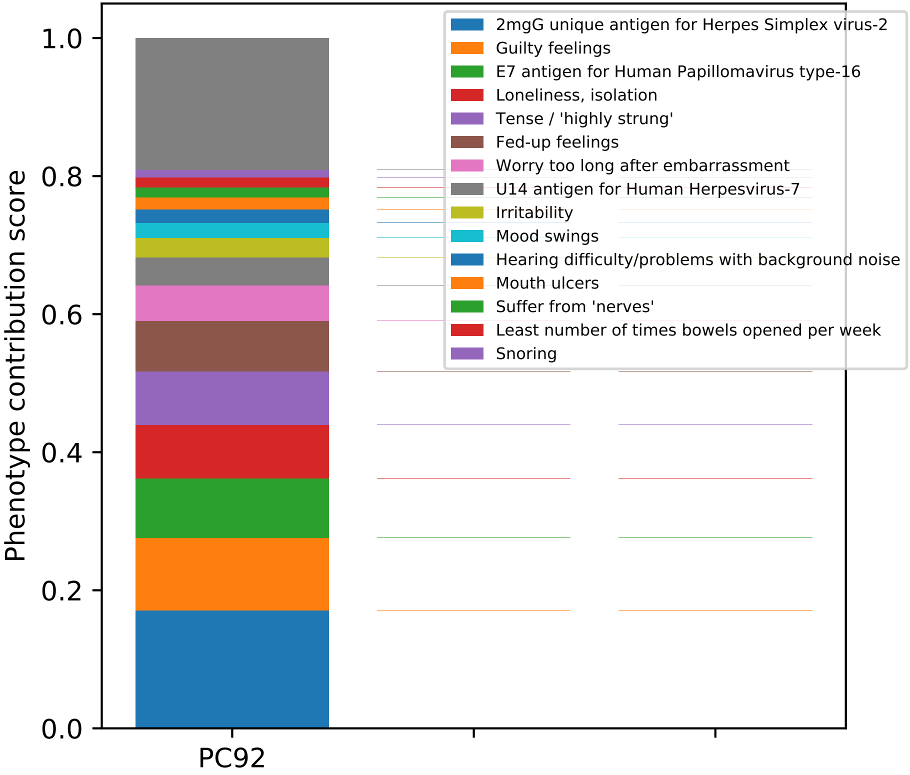

Phenotype contribution score
0.0
0.2
0.4
0.6
0.8
1.0
PC92
2mgG unique antigen for Herpes Simplex virus-2
Guilty feelings
E7 antigen for Human Papillomavirus type-16
Loneliness, isolation
Tense / 'highly strung'
Fed-up feelings
Worry too long after embarrassment
U14 antigen for Human Herpesvirus-7
Irritability
Mood swings
Hearing difficulty/problems with background noise
Mouth ulcers
Suffer from 'nerves'
Least number of times bowels opened per week
Snoring

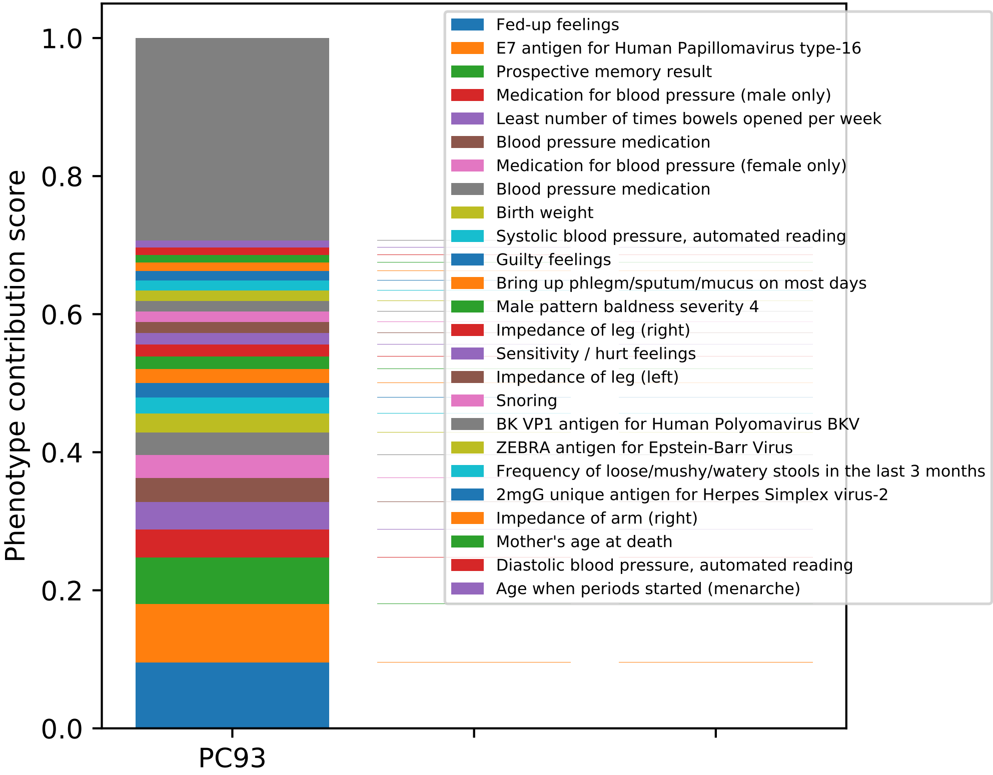

Phenotype contribution score
1.0
0.8
0.6
0.4
0.2
0.0
PC93
Fed-up feelings
E7 antigen for Human Papillomavirus type-16
Prospective memory result
Medication for blood pressure (male only)
Least number of times bowels opened per week
Blood pressure medication
Medication for blood pressure (female only)
Blood pressure medication
Birth weight
Systolic blood pressure, automated reading
Guilty feelings
Bring up phlegm/sputum/mucus on most days
Male pattern baldness severity 4
Impedance of leg (right)
Sensitivity / hurt feelings
Impedance of leg (left)
Snoring
BK VP1 antigen for Human Polyomavirus BKV
ZEBRA antigen for Epstein-Barr Virus
Frequency of loose/mushy/watery stools in the last 3 months
2mgG unique antigen for Herpes Simplex virus-2
Impedance of arm (right)
Mother's age at death
Diastolic blood pressure, automated reading
Age when periods started (menarche)

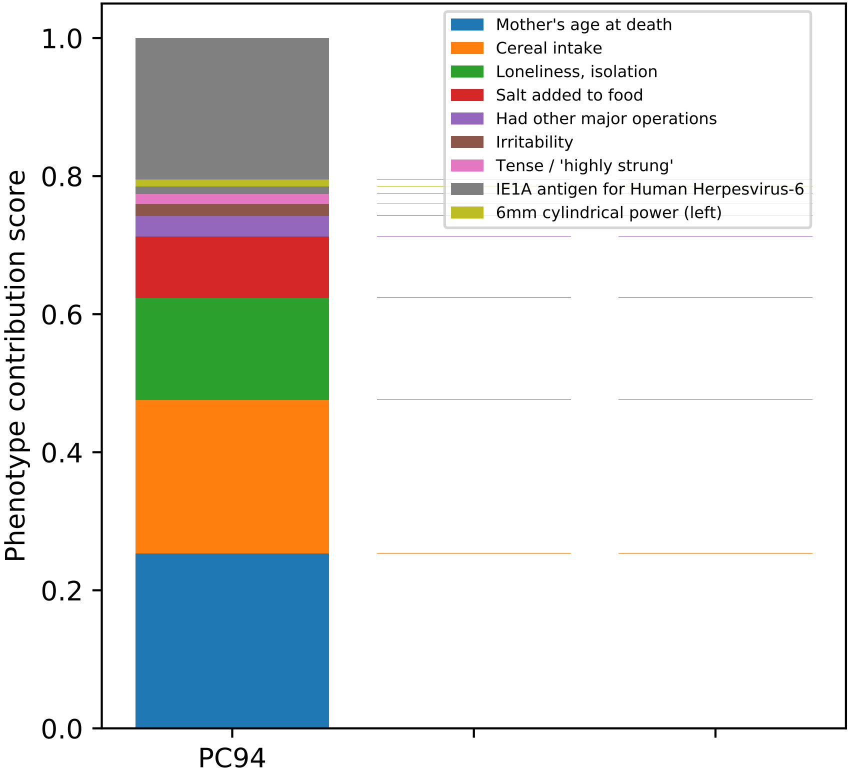
Mother's age at death
Cereal intake
Loneliness, isolation
Salt added to food
Had other major operations
Irritability
Tense / 'highly strung'
IE1A antigen for Human Herpesvirus-6
6mm cylindrical power (left)
Phenotype contribution score
1.0
0.8
0.6
0.4
0.2
0.0
PC94

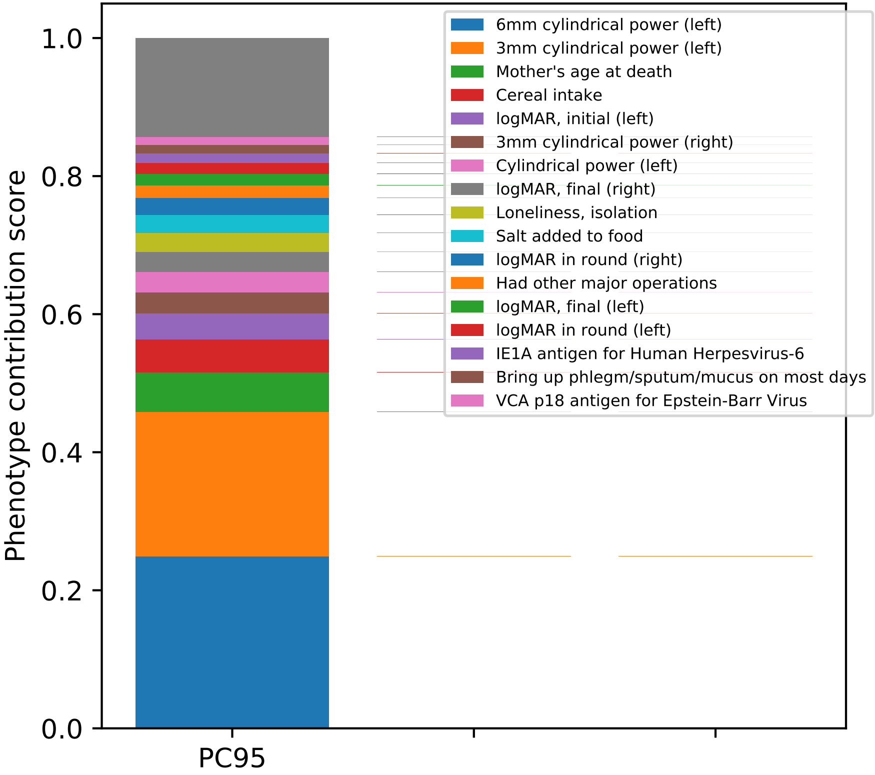
Phenotype contribution score
1.0
0.8
0.6
0.4
0.2
0.0
PC95
6mm cylindrical power (left)
3mm cylindrical power (left)
Mother's age at death
Cereal intake
logMAR, initial (left)
3mm cylindrical power (right)
Cylindrical power (left)
logMAR, final (right)
Loneliness, isolation
Salt added to food
logMAR in round (right)
Had other major operations
logMAR, final (left)
logMAR in round (left)
IE1A antigen for Human Herpesvirus-6
Bring up phlegm/sputum/mucus on most days
VCA p18 antigen for Epstein-Barr Virus

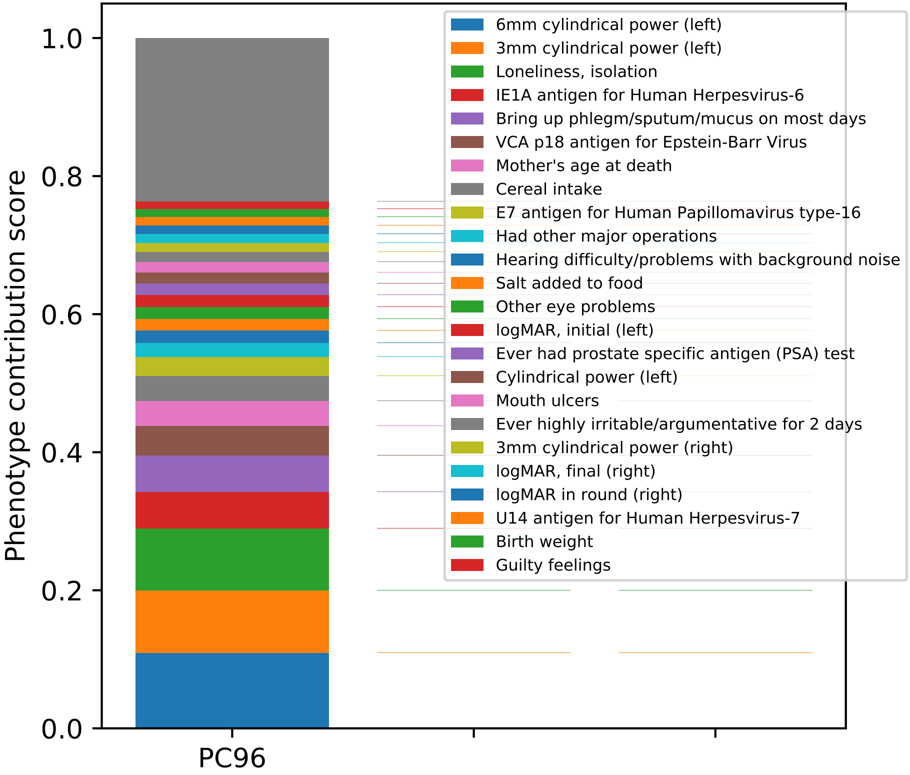

Phenotype contribution score
0.0
0.2
0.4
0.6
0.8
1.0
PC96
6mm cylindrical power (left)
3mm cylindrical power (left)
Loneliness, isolation
IE1A antigen for Human Herpesvirus-6
Bring up phlegm/sputum/mucus on most days
VCA p18 antigen for Epstein-Barr Virus
Mother's age at death
Cereal intake
E7 antigen for Human Papillomavirus type-16
Had other major operations
Hearing difficulty/problems with background noise
Salt added to food
Other eye problems
logMAR, initial (left)
Ever had prostate specific antigen (PSA) test
Cylindrical power (left)
Mouth ulcers
Ever highly irritable/argumentative for 2 days
3mm cylindrical power (right)
logMAR, final (right)
logMAR in round (right)
U14 antigen for Human Herpesvirus-7
Birth weight
Guilty feelings

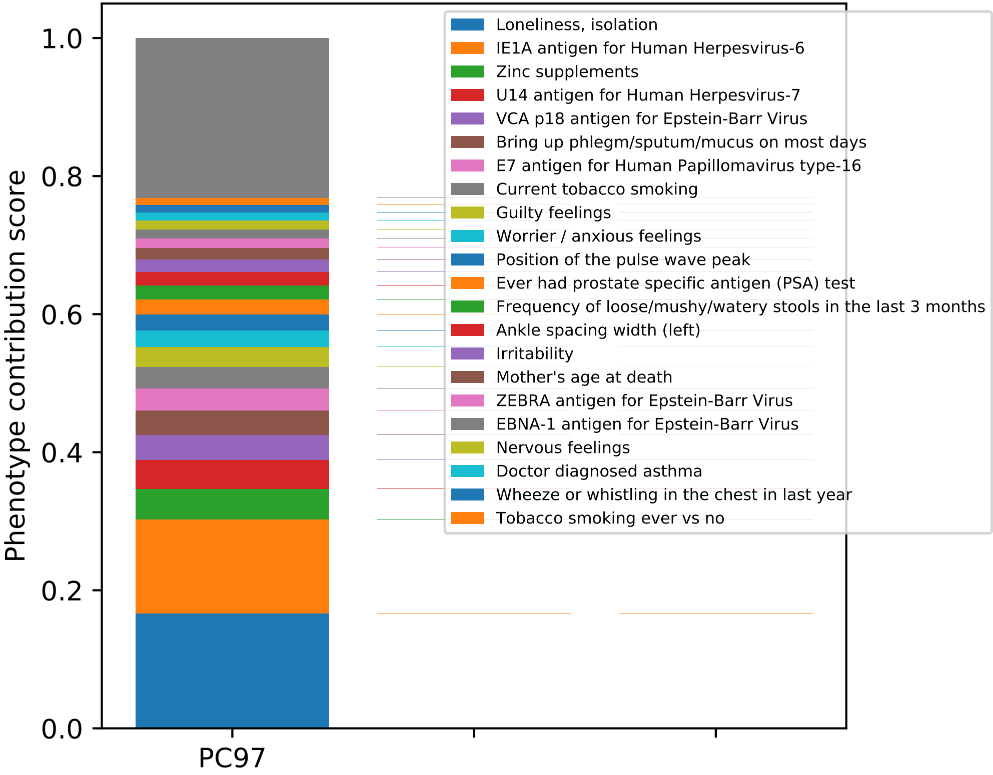

Phenotype contribution score
0.0
0.2
0.4
0.6
0.8
1.0
PC97
Loneliness, isolation
IE1A antigen for Human Herpesvirus-6
Zinc supplements
U14 antigen for Human Herpesvirus-7
VCA p18 antigen for Epstein-Barr Virus
Bring up phlegm/sputum/mucus on most days
E7 antigen for Human Papillomavirus type-16
Current tobacco smoking
Guilty feelings
Worrier / anxious feelings
Position of the pulse wave peak
Ever had prostate specific antigen (PSA) test
Frequency of loose/mushy/watery stools in the last 3 months
Ankle spacing width (left)
Irritability
Mother's age at death
ZEBRA antigen for Epstein-Barr Virus
EBNA-1 antigen for Epstein-Barr Virus
Nervous feelings
Doctor diagnosed asthma
Wheeze or whistling in the chest in last year
Tobacco smoking ever vs no

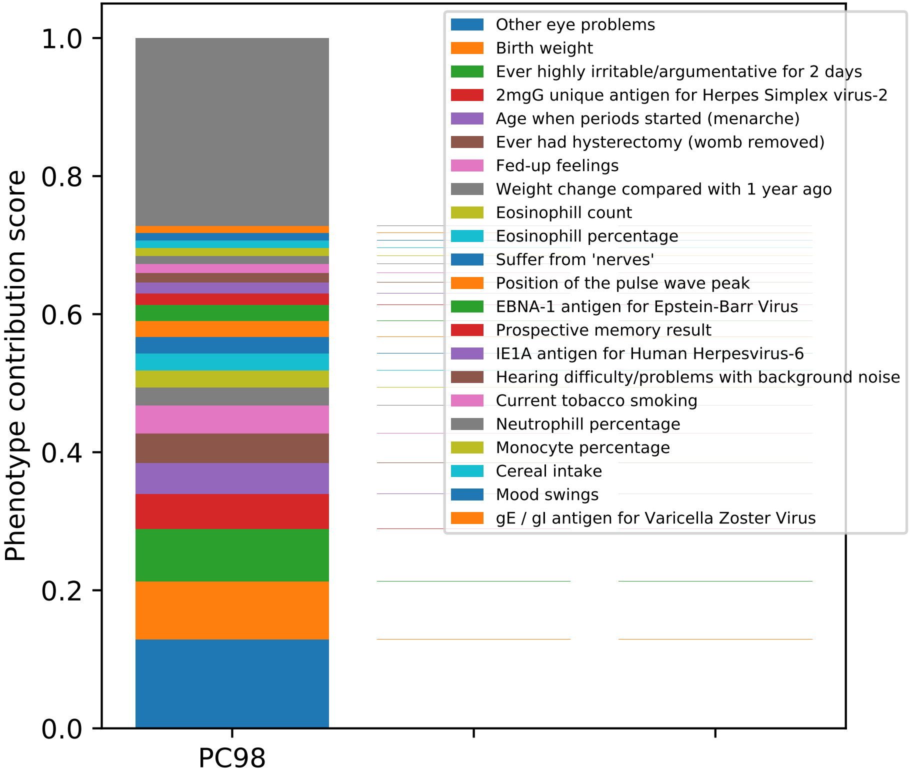
Phenotype contribution score
1.0
0.8
0.6
0.4
0.2
0.0
PC98
Other eye problems
Birth weight
Ever highly irritable/argumentative for 2 days
2mgG unique antigen for Herpes Simplex virus-2
Age when periods started (menarche)
Ever had hysterectomy (womb removed)
Fed-up feelings
Weight change compared with 1 year ago
Eosinophill count
Eosinophill percentage
Suffer from 'nerves'
Position of the pulse wave peak
EBNA-1 antigen for Epstein-Barr Virus
Prospective memory result
IE1A antigen for Human Herpesvirus-6
Hearing difficulty/problems with background noise
Current tobacco smoking
Neutrophill percentage
Monocyte percentage
Cereal intake
Mood swings
gE / gI antigen for Varicella Zoster Virus

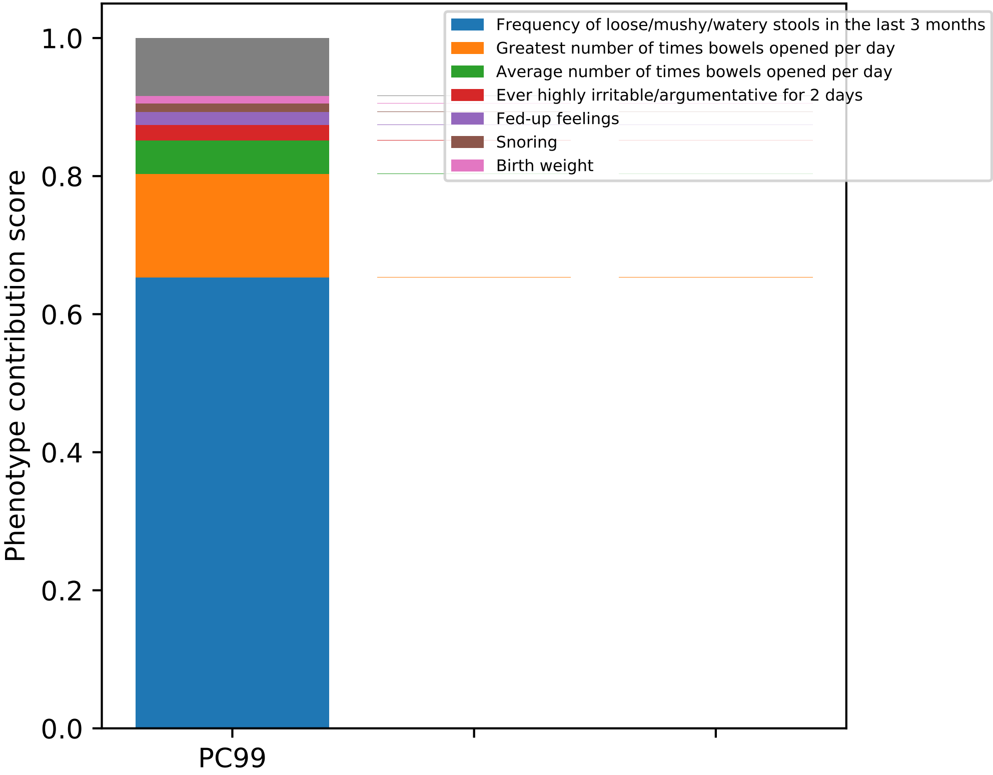

Frequency of loose/mushy/watery stools in the last 3 months
Greatest number of times bowels opened per day
Average number of times bowels opened per day
Ever highly irritable/argumentative for 2 days
Fed-up feelings
Snoring
Birth weight
Phenotype contribution score
0.0
0.2
0.4
0.6
0.8
1.0
PC99

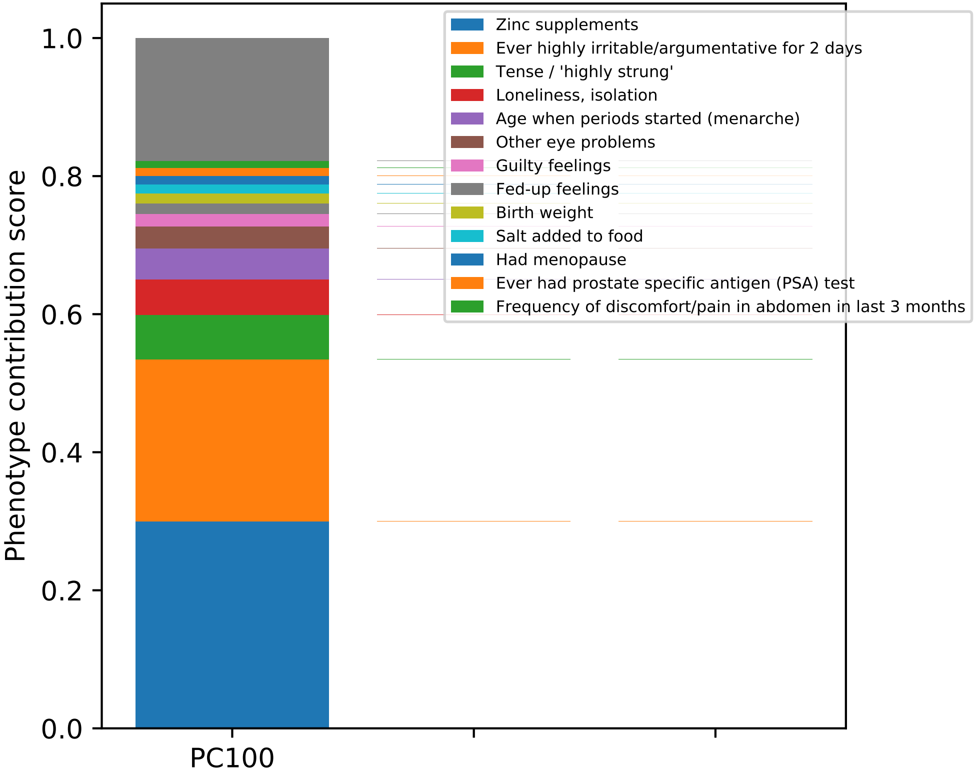

Phenotype contribution score
1.0
0.8
0.6
0.4
0.2
0.0
PC100
Zinc supplements
Ever highly irritable/argumentative for 2 days
Tense / 'highly strung'
Loneliness, isolation
Age when periods started (menarche)
Other eye problems
Guilty feelings
Fed-up feelings
Birth weight
Salt added to food
Had menopause
Ever had prostate specific antigen (PSA) test
Frequency of discomfort/pain in abdomen in last 3 months

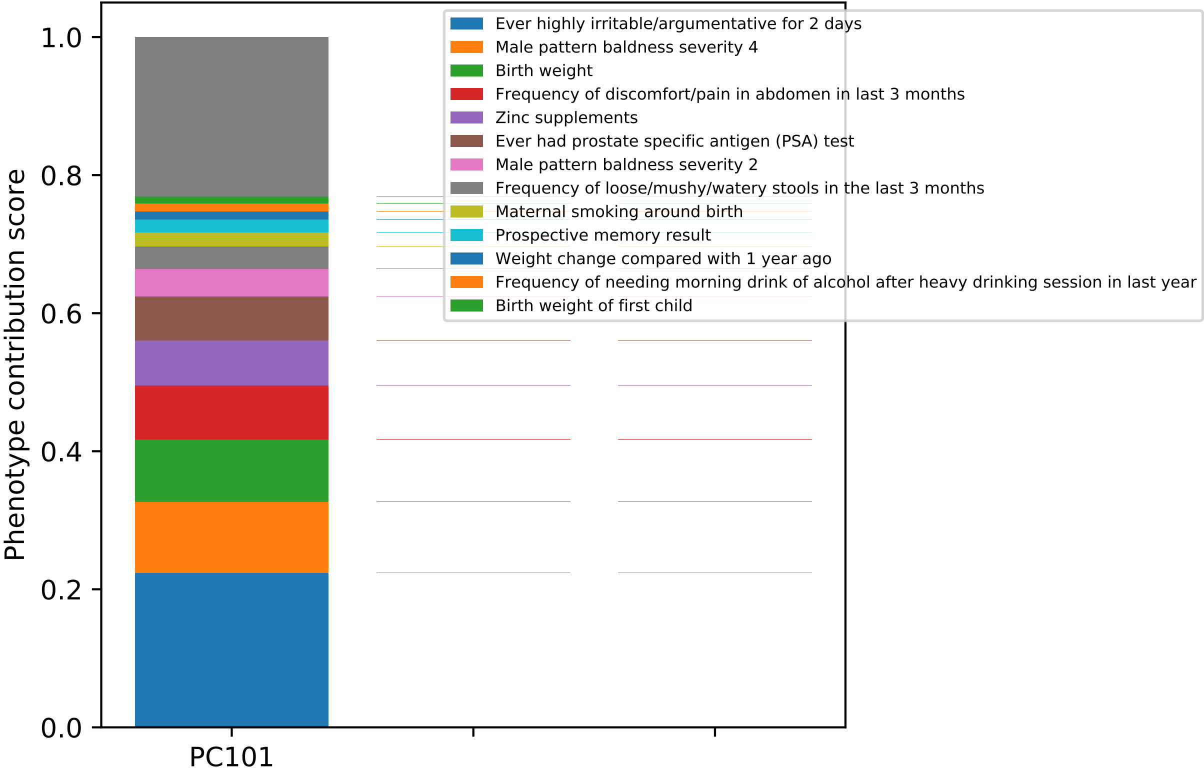

Phenotype contribution score
1.0
0.8
0.6
0.4
0.2
0.0
PC101
Ever highly irritable/argumentative for 2 days
Male pattern baldness severity 4
Birth weight
Frequency of discomfort/pain in abdomen in last 3 months
Zinc supplements
Ever had prostate specific antigen (PSA) test
Male pattern baldness severity 2
Frequency of loose/mushy/watery stools in the last 3 months
Maternal smoking around birth
Prospective memory result
Weight change compared with 1 year ago
Frequency of needing morning drink of alcohol after heavy drinking session in last year
Birth weight of first child

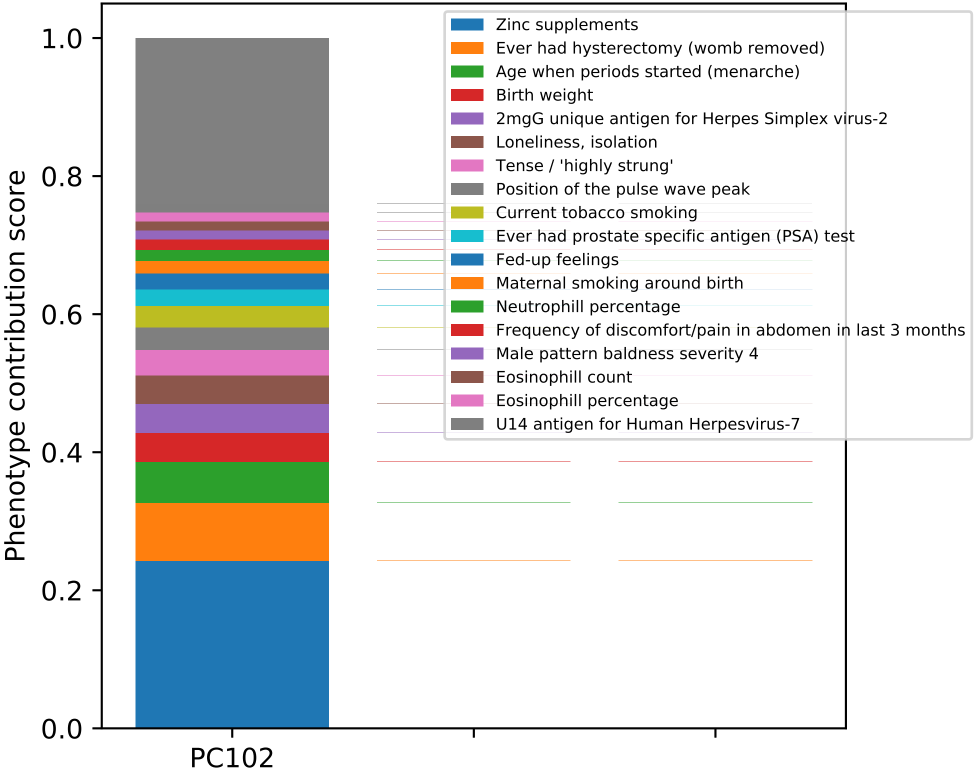

Phenotype contribution score
1.0
0.8
0.6
0.4
0.2
0.0
PC102
Zinc supplements
Ever had hysterectomy (womb removed)
Age when periods started (menarche)
Birth weight
2mgG unique antigen for Herpes Simplex virus-2
Loneliness, isolation
Tense / 'highly strung'
Position of the pulse wave peak
Current tobacco smoking
Ever had prostate specific antigen (PSA) test
Fed-up feelings
Maternal smoking around birth
Neutrophill percentage
Frequency of discomfort/pain in abdomen in last 3 months
Male pattern baldness severity 4
Eosinophill count
Eosinophill percentage
U14 antigen for Human Herpesvirus-7

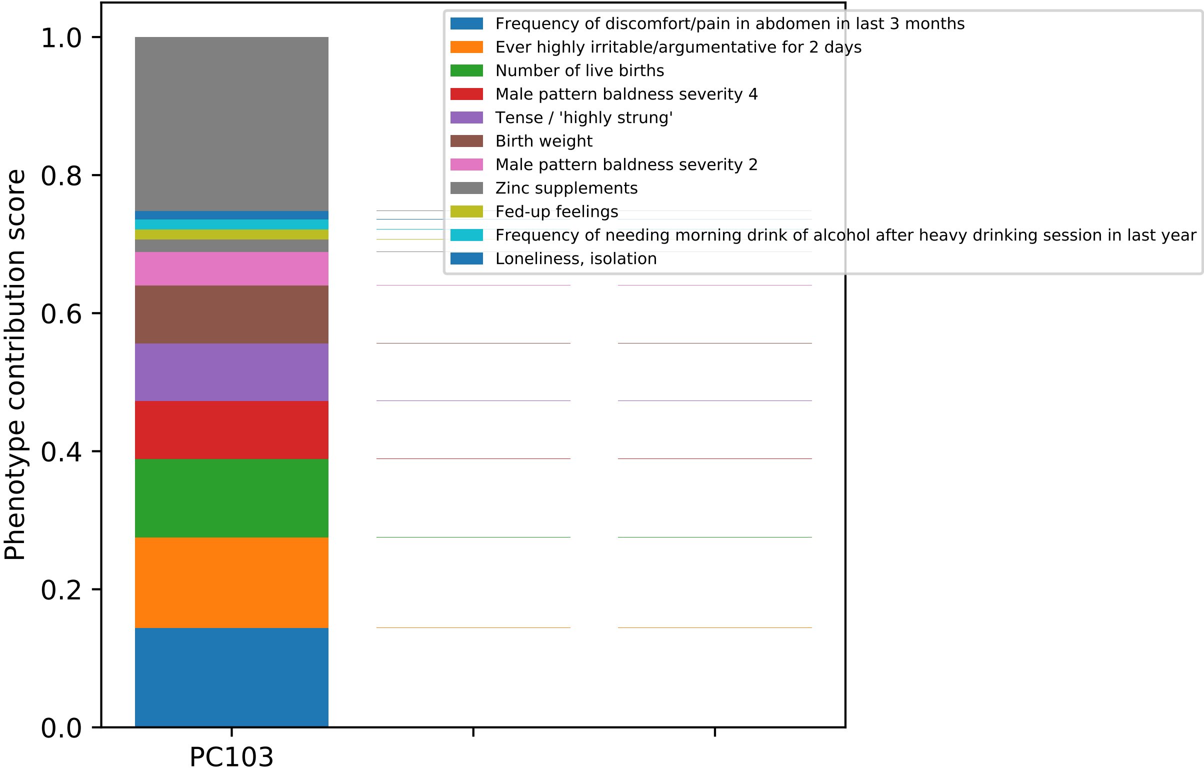

Frequency of discomfort/pain in abdomen in last 3 months
Ever highly irritable/argumentative for 2 days
Number of live births
Male pattern baldness severity 4
Tense / 'highly strung'
Birth weight
Male pattern baldness severity 2
Zinc supplements
Fed-up feelings
Frequency of needing morning drink of alcohol after heavy drinking session in last year
Loneliness, isolation
Phenotype contribution score
0.0
0.2
0.4
0.6
0.8
1.0
PC103

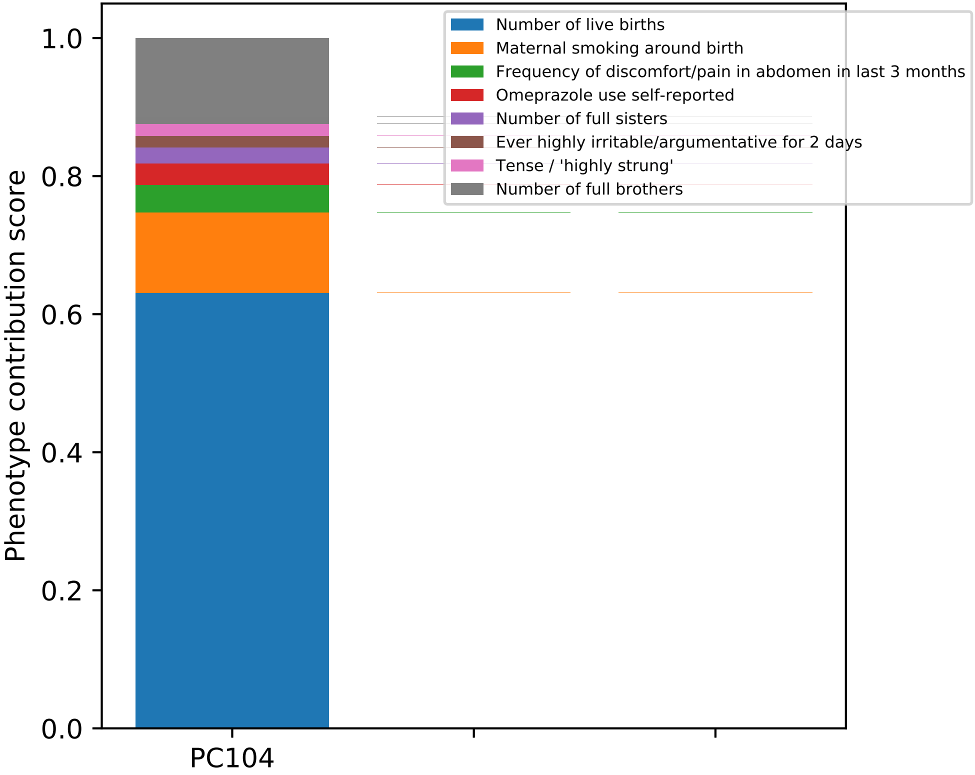

Phenotype contribution score
1.0
0.8
0.6
0.4
0.2
0.0
PC104
Number of live births
Maternal smoking around birth
Frequency of discomfort/pain in abdomen in last 3 months
Omeprazole use self-reported
Number of full sisters
Ever highly irritable/argumentative for 2 days
Tense / 'highly strung'
Number of full brothers

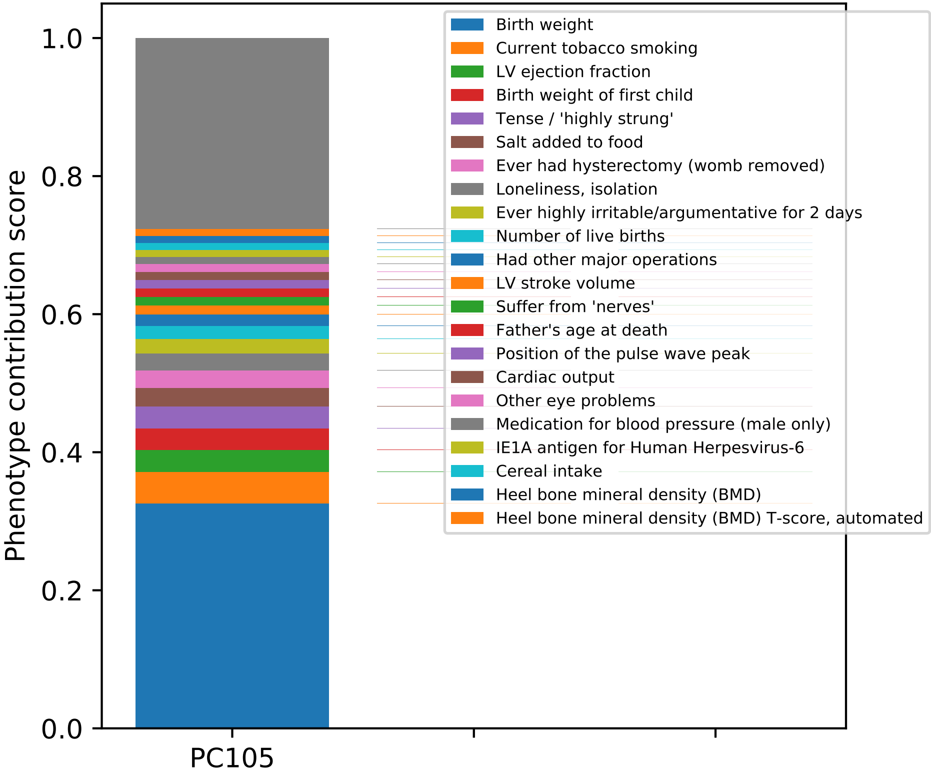
Phenotype contribution score
1.0
0.8
0.6
0.4
0.2
0.0
PC105
Birth weight
Current tobacco smoking
LV ejection fraction
Birth weight of first child
Tense / 'highly strung'
Salt added to food
Ever had hysterectomy (womb removed)
Loneliness, isolation
Ever highly irritable/argumentative for 2 days
Number of live births
Had other major operations
LV stroke volume
Suffer from 'nerves'
Father's age at death
Position of the pulse wave peak
Cardiac output
Other eye problems
Medication for blood pressure (male only)
IE1A antigen for Human Herpesvirus-6
Cereal intake
Heel bone mineral density (BMD)
Heel bone mineral density (BMD) T-score, automated

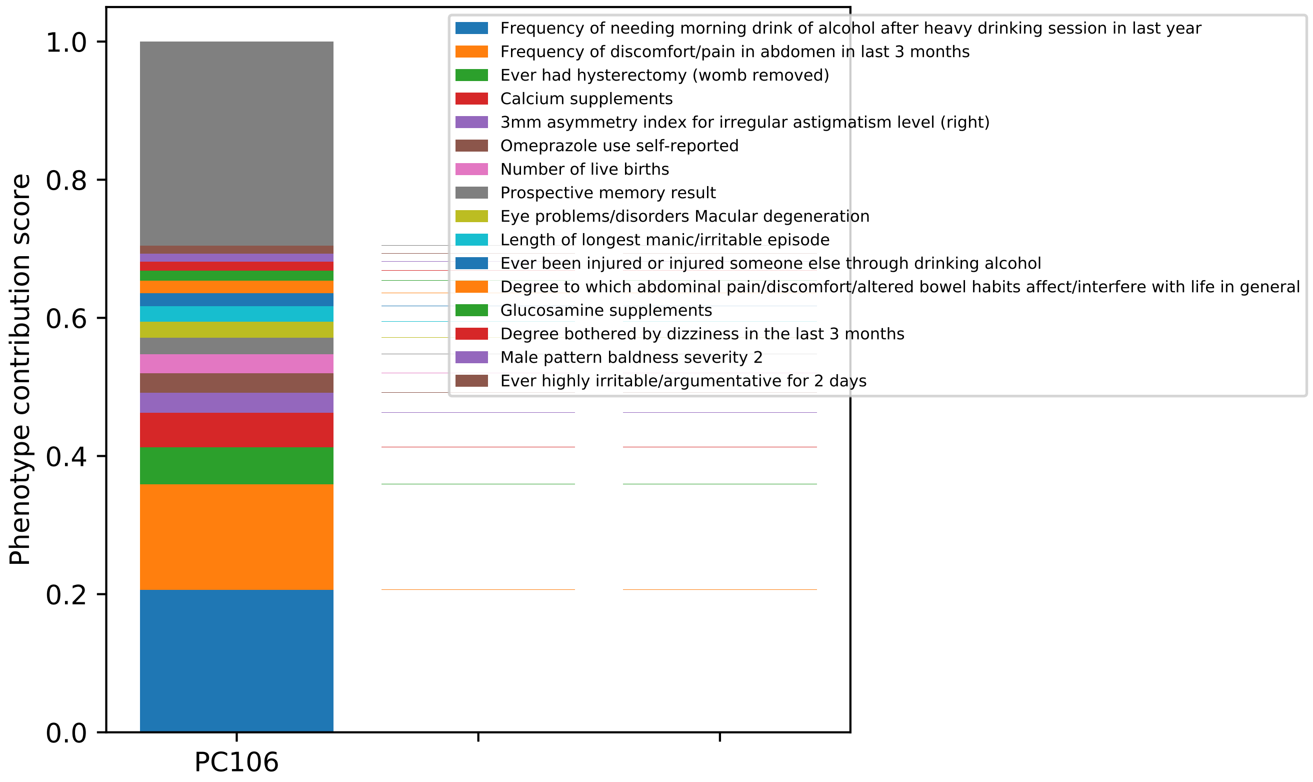

Phenotype contribution score
1.0
0.8
0.6
0.4
0.2
0.0
PC106
Frequency of needing morning drink of alcohol after heavy drinking session in last year
Frequency of discomfort/pain in abdomen in last 3 months
Ever had hysterectomy (womb removed)
Calcium supplements
3mm asymmetry index for irregular astigmatism level (right)
Omeprazole use self-reported
Number of live births
Prospective memory result
Eye problems/disorders Macular degeneration
Length of longest manic/irritable episode
Ever been injured or injured someone else through drinking alcohol
Degree to which abdominal pain/discomfort/altered bowel habits affect/interfere with life in general
Glucosamine supplements
Degree bothered by dizziness in the last 3 months
Male pattern baldness severity 2
Ever highly irritable/argumentative for 2 days

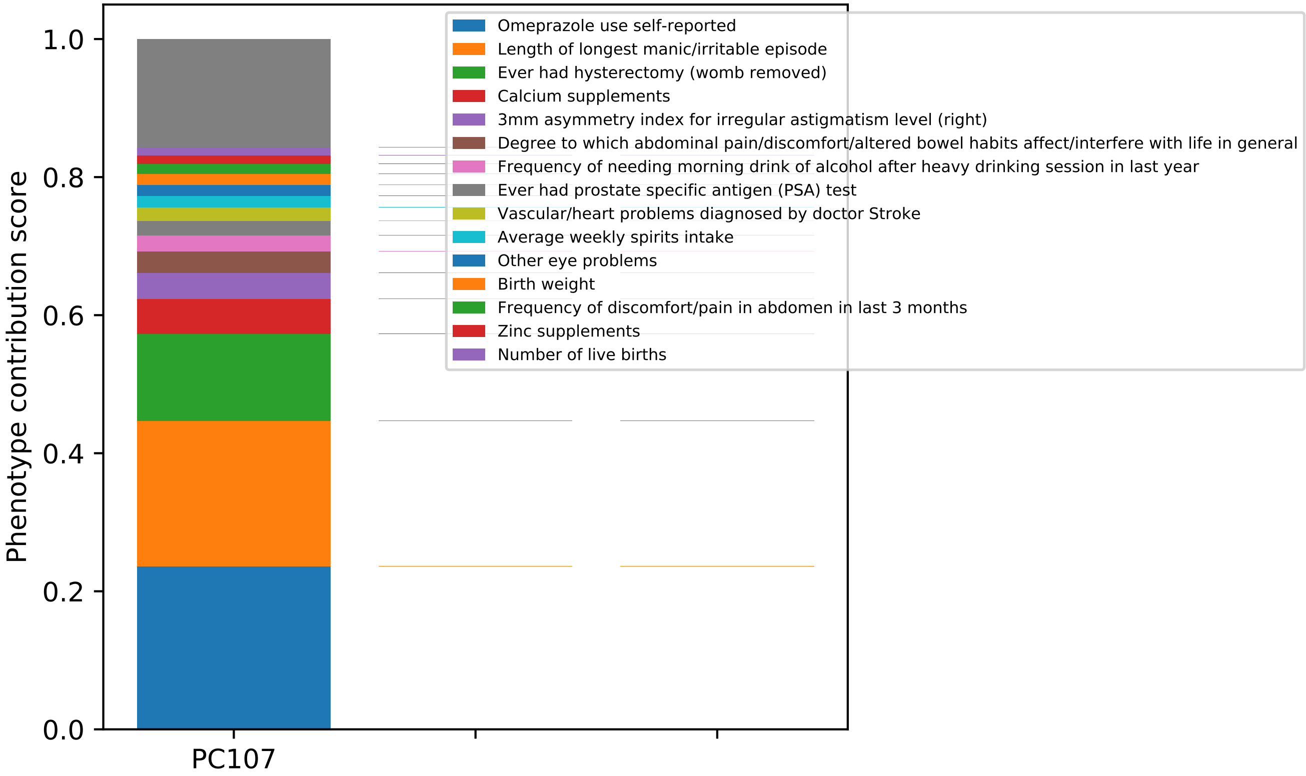

Phenotype contribution score
1.0
0.8
0.6
0.4
0.2
0.0
PC107
Omeprazole use self-reported
Length of longest manic/irritable episode
Ever had hysterectomy (womb removed)
Calcium supplements
3mm asymmetry index for irregular astigmatism level (right)
Degree to which abdominal pain/discomfort/altered bowel habits affect/interfere with life in general
Frequency of needing morning drink of alcohol after heavy drinking session in last year
Ever had prostate specific antigen (PSA) test
Vascular/heart problems diagnosed by doctor Stroke
Average weekly spirits intake
Other eye problems
Birth weight
Frequency of discomfort/pain in abdomen in last 3 months
Zinc supplements
Number of live births

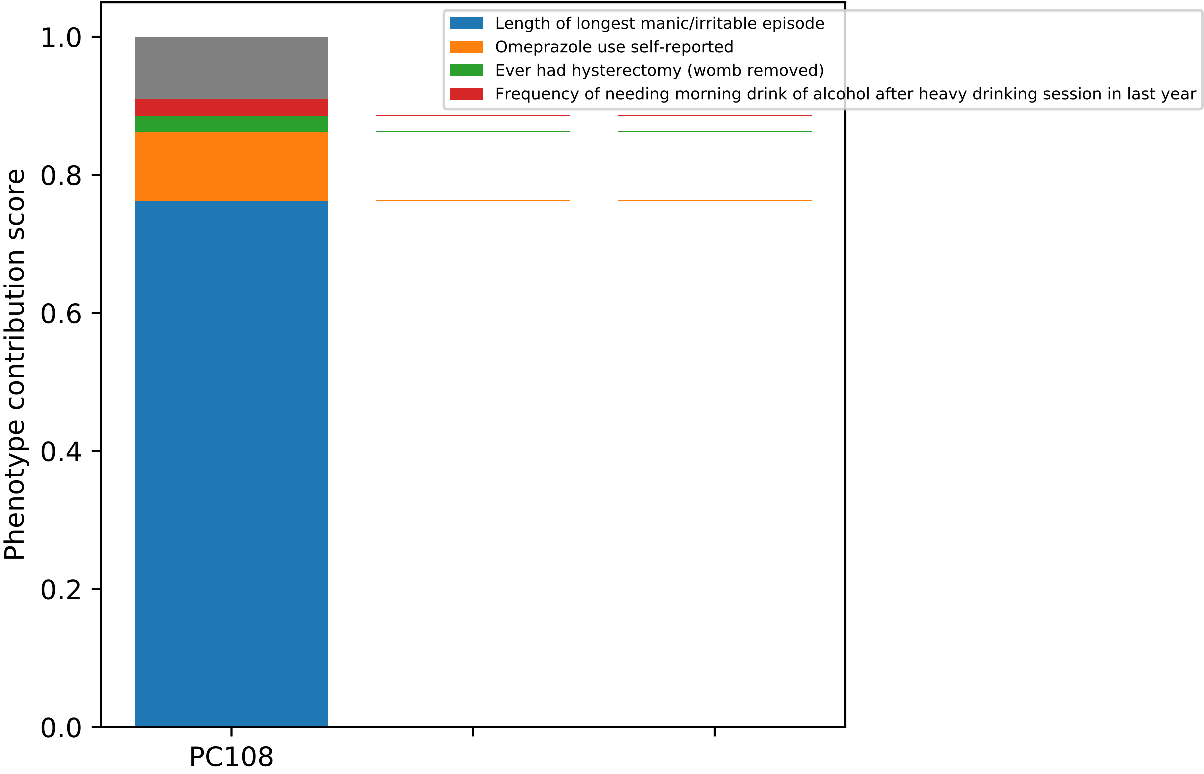

Length of longest manic/irritable episode
Omeprazole use self-reported
Ever had hysterectomy (womb removed)
Frequency of needing morning drink of alcohol after heavy drinking session in last year
Phenotype contribution score
1.0
0.8
0.6
0.4
0.2
0.0
PC108

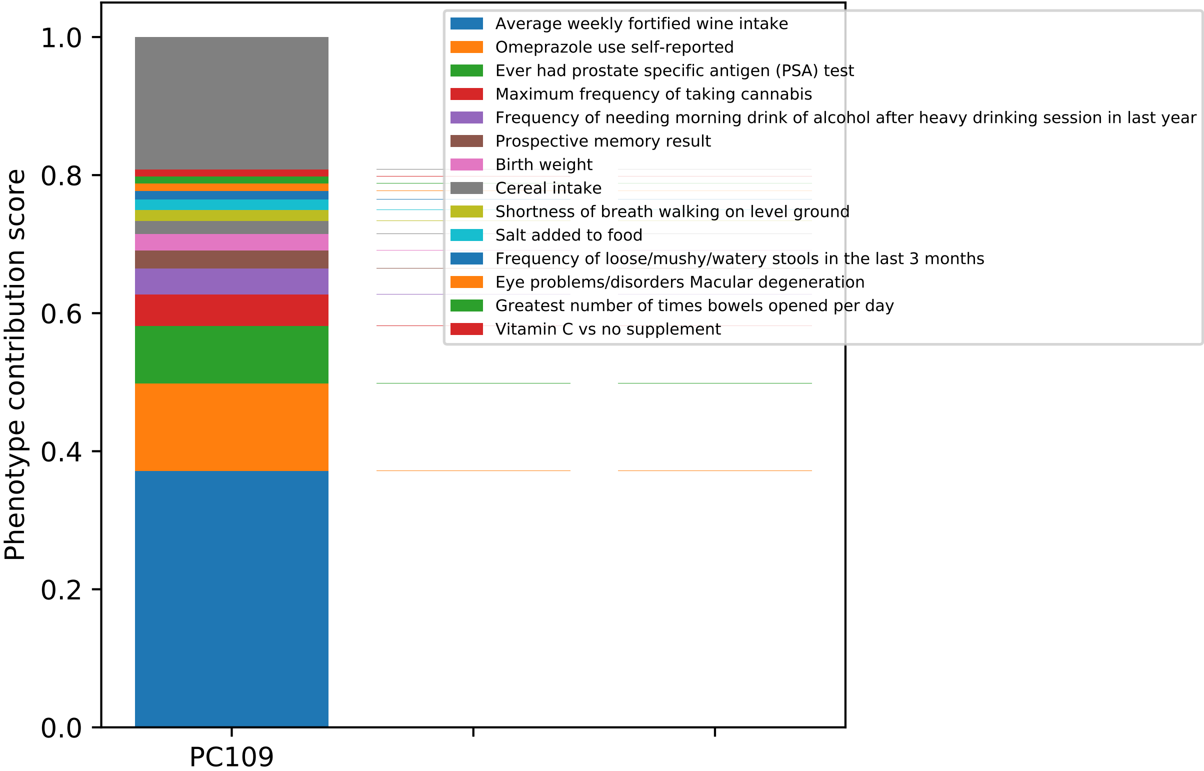

Phenotype contribution score
1.0
0.8
0.6
0.4
0.2
0.0
PC109
Average weekly fortified wine intake
Omeprazole use self-reported
Ever had prostate specific antigen (PSA) test
Maximum frequency of taking cannabis
Frequency of needing morning drink of alcohol after heavy drinking session in last year
Prospective memory result
Birth weight
Cereal intake
Shortness of breath walking on level ground
Salt added to food
Frequency of loose/mushy/watery stools in the last 3 months
Eye problems/disorders Macular degeneration
Greatest number of times bowels opened per day
Vitamin C vs no supplement

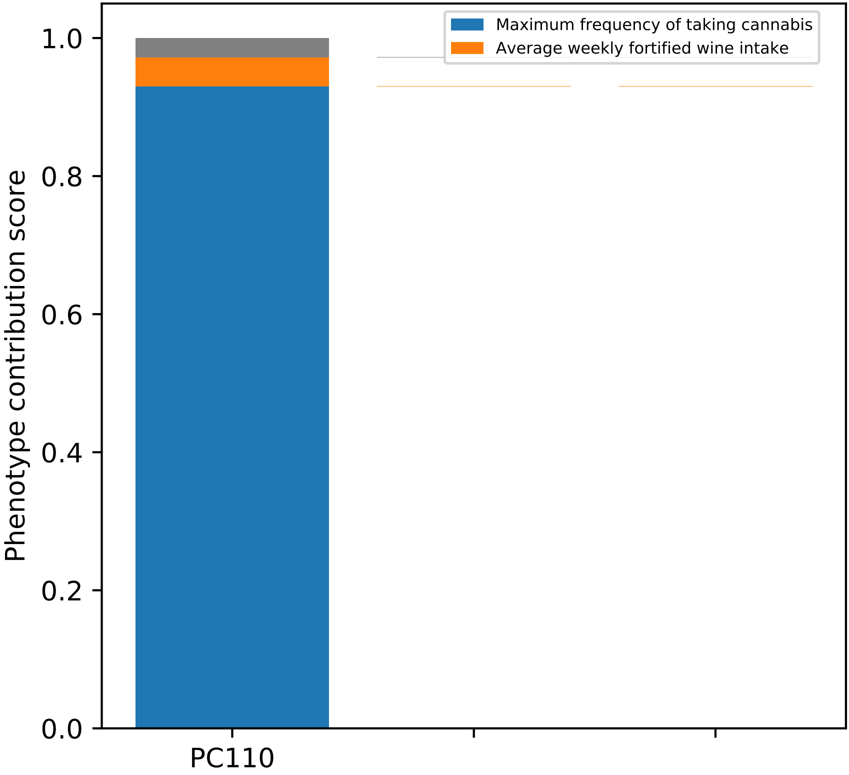

Maximum frequency of taking cannabis
Average weekly fortified wine intake
Phenotype contribution score
1.0
0.8
0.6
0.4
0.2
0.0
PC110

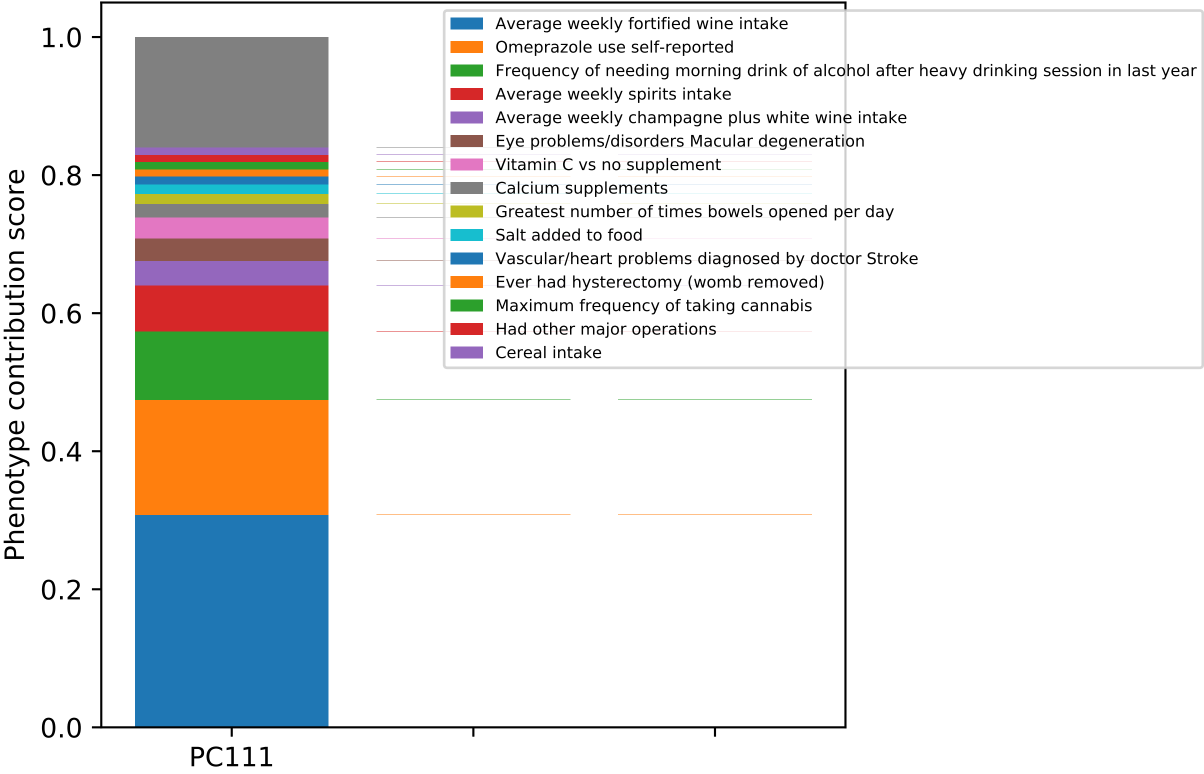
Phenotype contribution score
0.0
0.2
0.4
0.6
0.8
1.0
PC111
Average weekly fortified wine intake
Omeprazole use self-reported
Frequency of needing morning drink of alcohol after heavy drinking session in last year
Average weekly spirits intake
Average weekly champagne plus white wine intake
Eye problems/disorders Macular degeneration
Vitamin C vs no supplement
Calcium supplements
Greatest number of times bowels opened per day
Salt added to food
Vascular/heart problems diagnosed by doctor Stroke
Ever had hysterectomy (womb removed)
Maximum frequency of taking cannabis
Had other major operations
Cereal intake

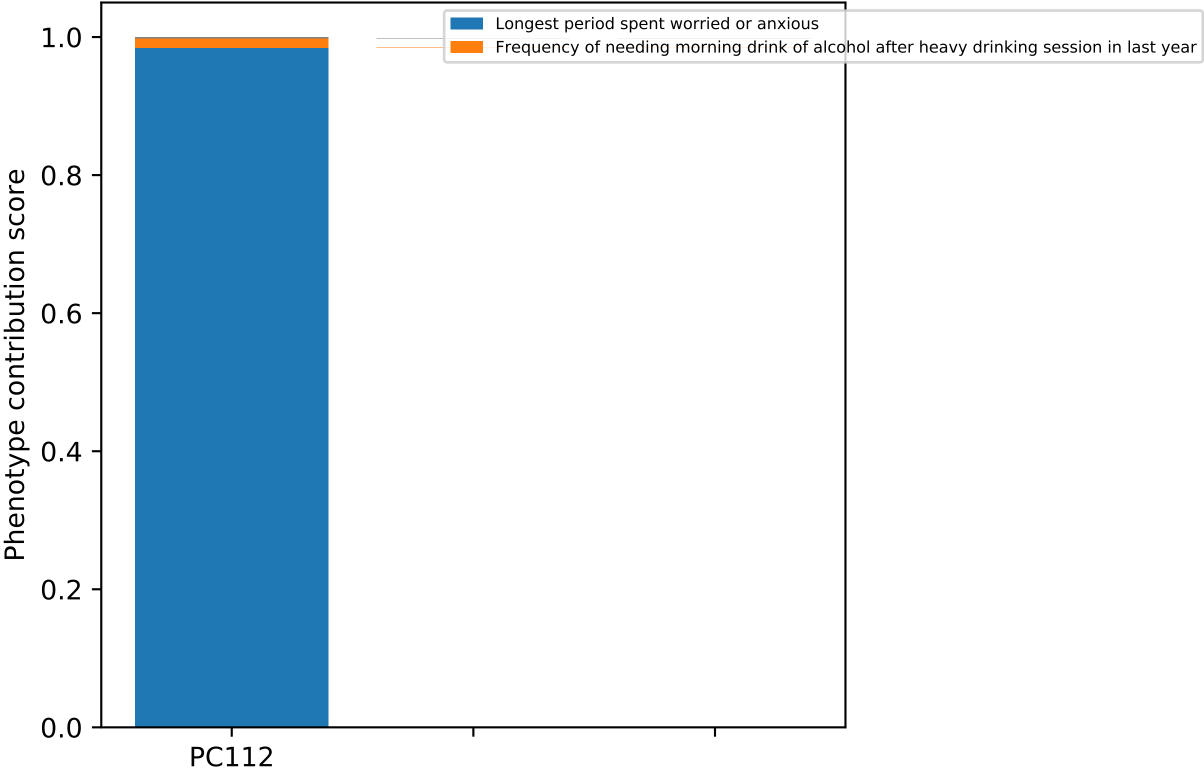

Longest period spent worried or anxious
Frequency of needing morning drink of alcohol after heavy drinking session in last year
Phenotype contribution score
1.0
0.8
0.6
0.4
0.2
0.0
PC112

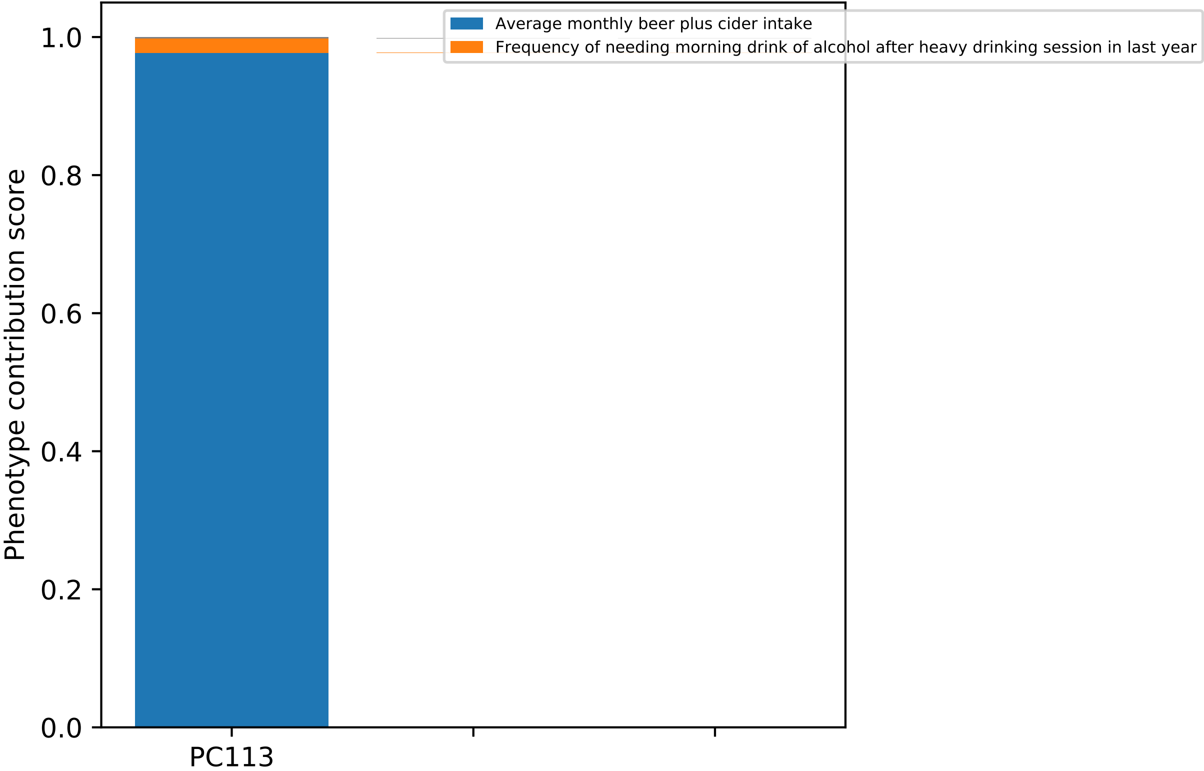

Average monthly beer plus cider intake
Frequency of needing morning drink of alcohol after heavy drinking session in last year
Phenotype contribution score
1.0
0.8
0.6
0.4
0.2
0.0
PC113

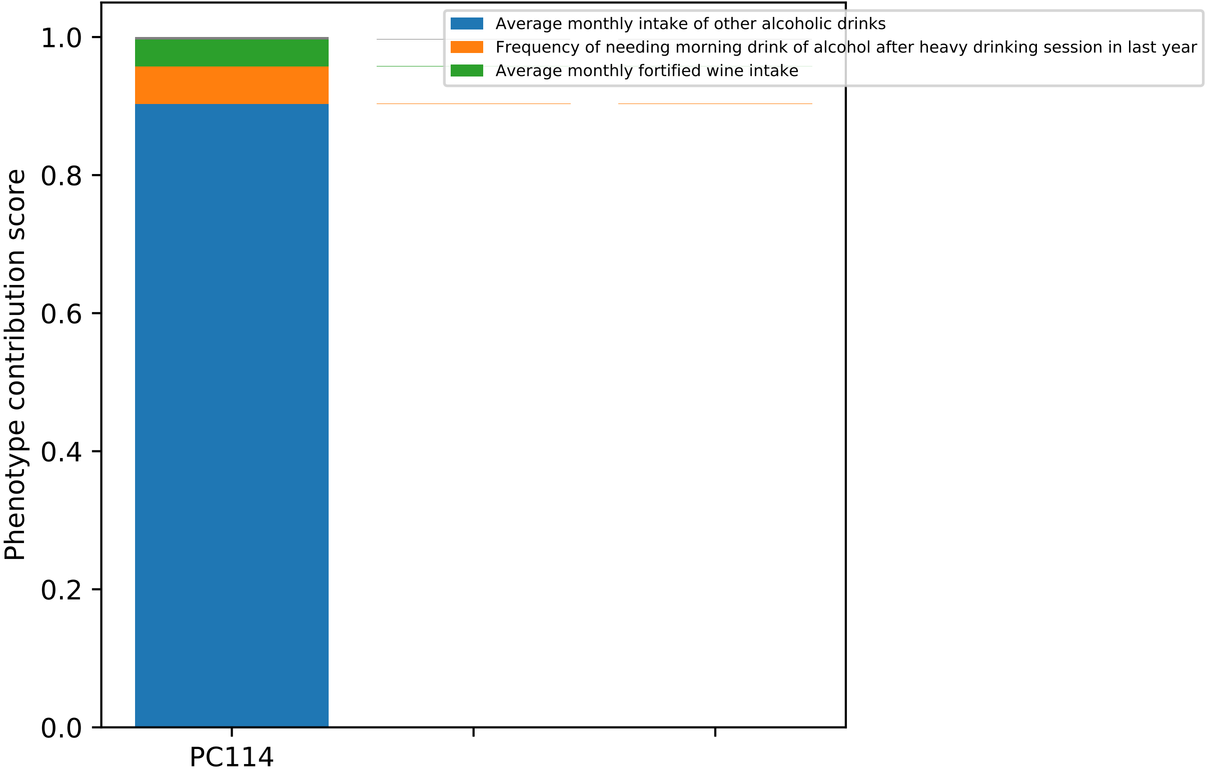

Average monthly intake of other alcoholic drinks
Frequency of needing morning drink of alcohol after heavy drinking session in last year
Average monthly fortified wine intake
Phenotype contribution score
1.0
0.8
0.6
0.4
0.2
0.0
PC114

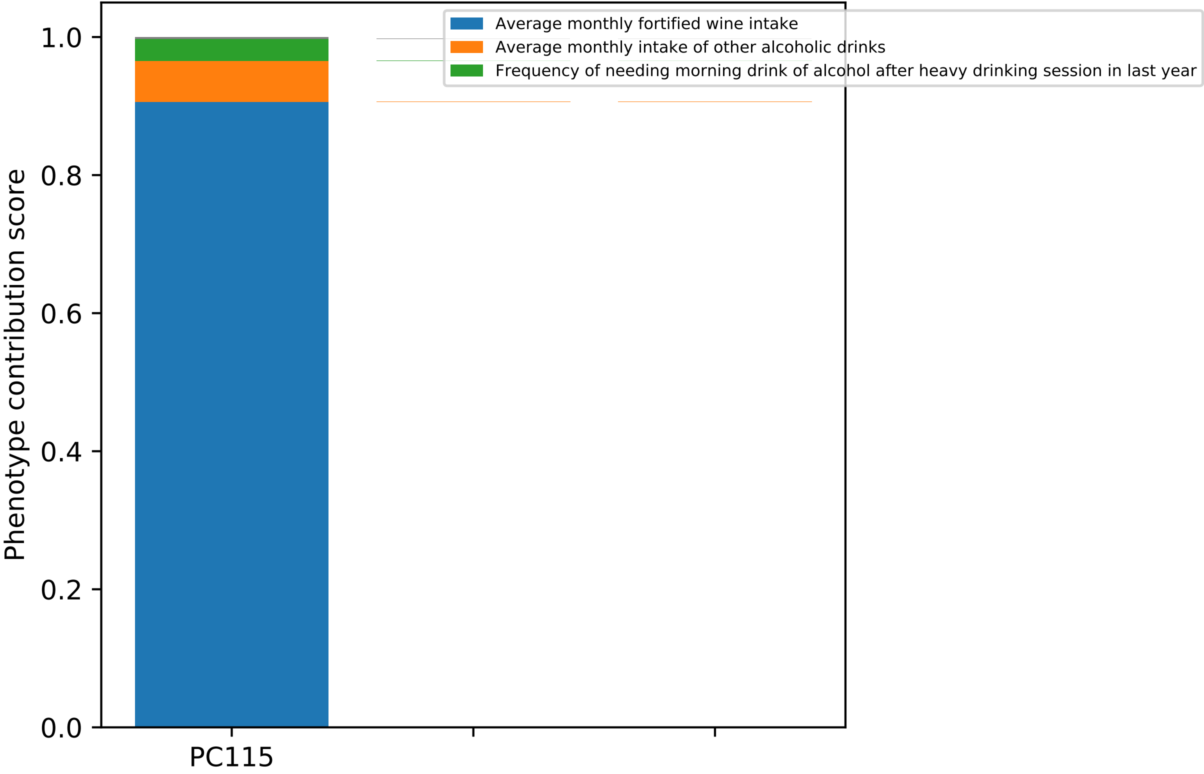

Average monthly fortified wine intake
Average monthly intake of other alcoholic drinks
Frequency of needing morning drink of alcohol after heavy drinking session in last year
Phenotype contribution score
1.0
0.8
0.6
0.4
0.2
0.0
PC115

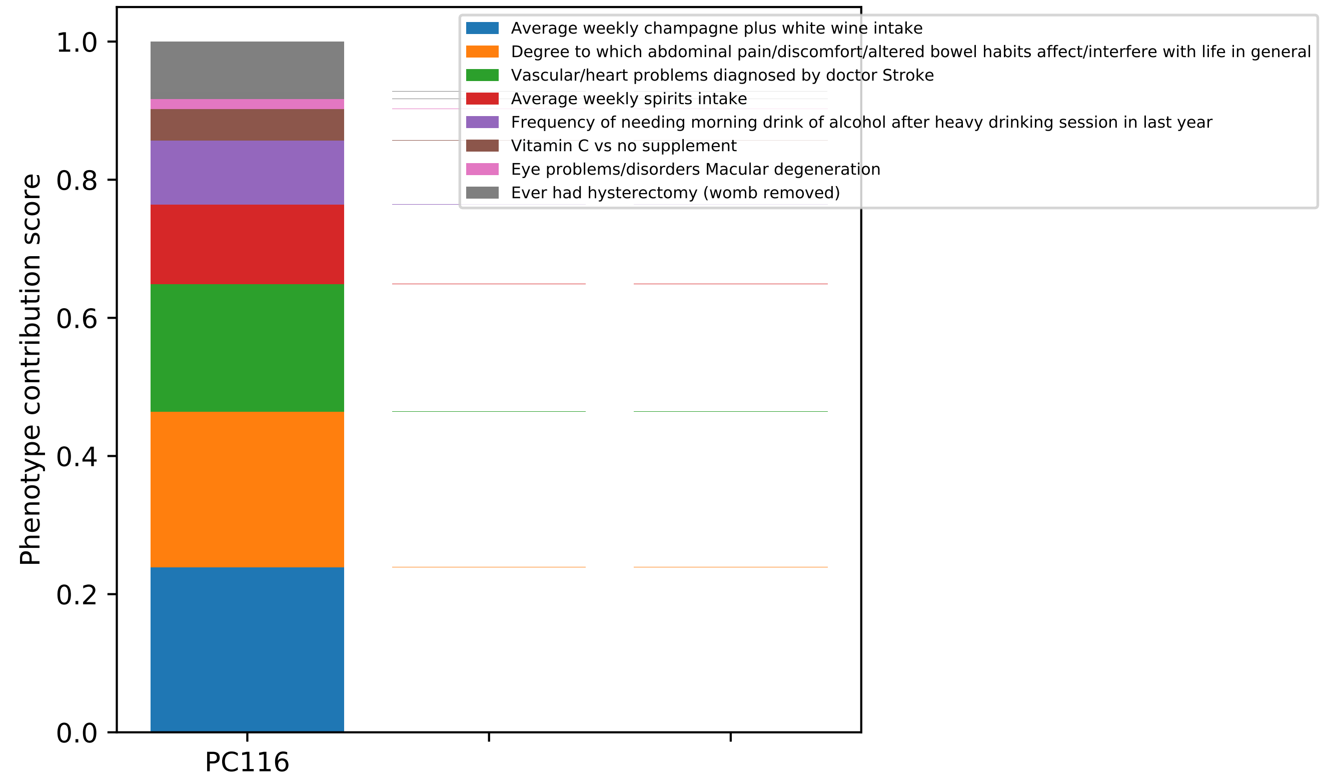

Phenotype contribution score
0.0
0.2
0.4
0.6
0.8
1.0
PC116
Average weekly champagne plus white wine intake
Degree to which abdominal pain/discomfort/altered bowel habits affect/interfere with life in general
Vascular/heart problems diagnosed by doctor Stroke
Average weekly spirits intake
Frequency of needing morning drink of alcohol after heavy drinking session in last year
Vitamin C vs no supplement
Eye problems/disorders Macular degeneration
Ever had hysterectomy (womb removed)

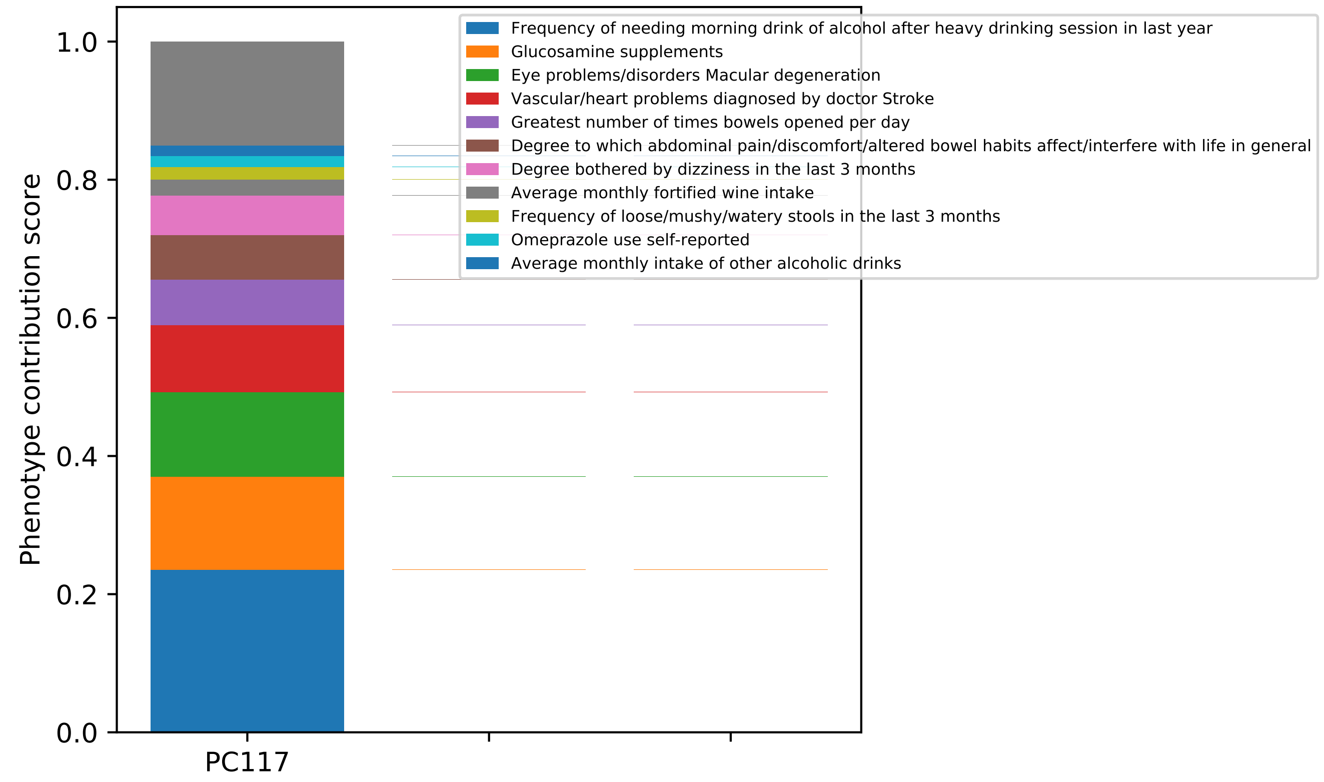

Phenotype contribution score
1.0
0.8
0.6
0.4
0.2
0.0
PC117
Frequency of needing morning drink of alcohol after heavy drinking session in last year
Glucosamine supplements
Eye problems/disorders Macular degeneration
Vascular/heart problems diagnosed by doctor Stroke
Greatest number of times bowels opened per day
Degree to which abdominal pain/discomfort/altered bowel habits affect/interfere with life in general
Degree bothered by dizziness in the last 3 months
Average monthly fortified wine intake
Frequency of loose/mushy/watery stools in the last 3 months
Omeprazole use self-reported
Average monthly intake of other alcoholic drinks

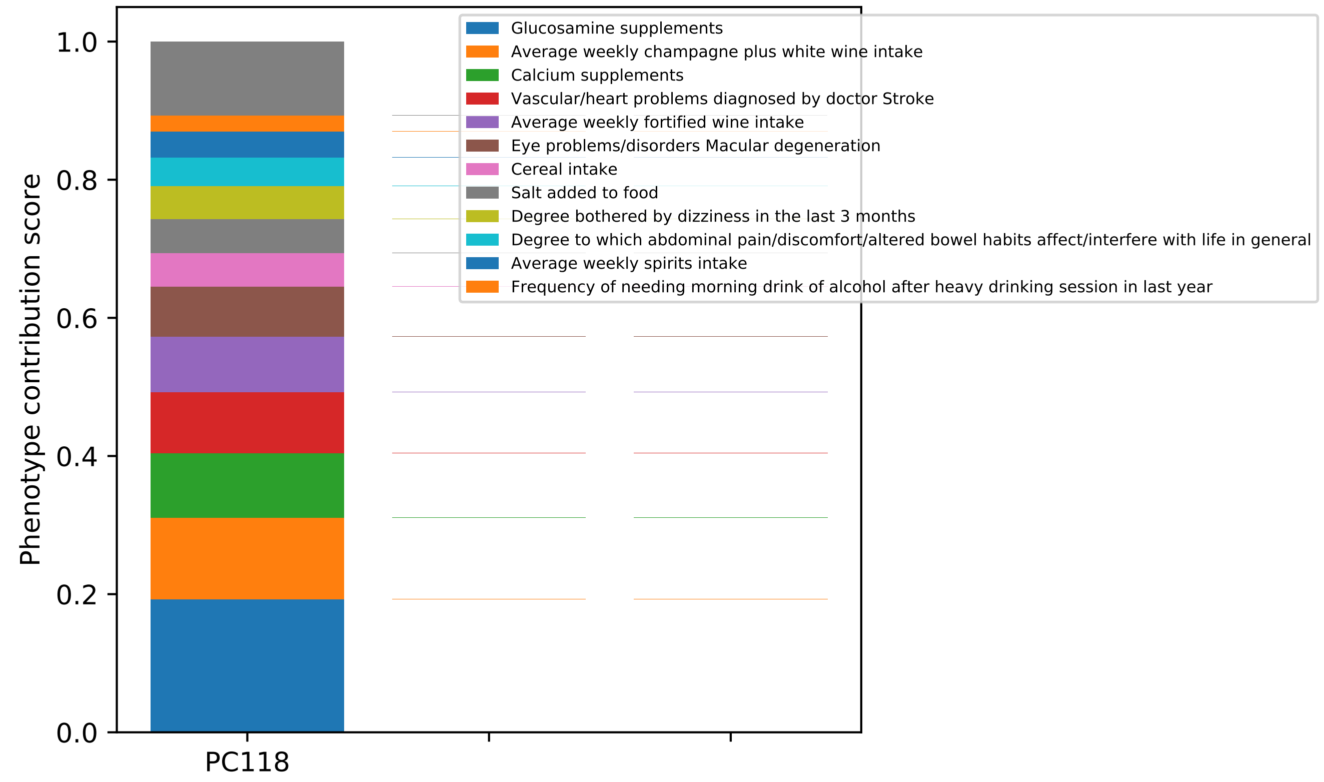

Phenotype contribution score
1.0
0.8
0.6
0.4
0.2
0.0
PC118
Glucosamine supplements
Average weekly champagne plus white wine intake
Calcium supplements
Vascular/heart problems diagnosed by doctor Stroke
Average weekly fortified wine intake
Eye problems/disorders Macular degeneration
Cereal intake
Salt added to food
Degree bothered by dizziness in the last 3 months
Degree to which abdominal pain/discomfort/altered bowel habits affect/interfere with life in general
Average weekly spirits intake
Frequency of needing morning drink of alcohol after heavy drinking session in last year

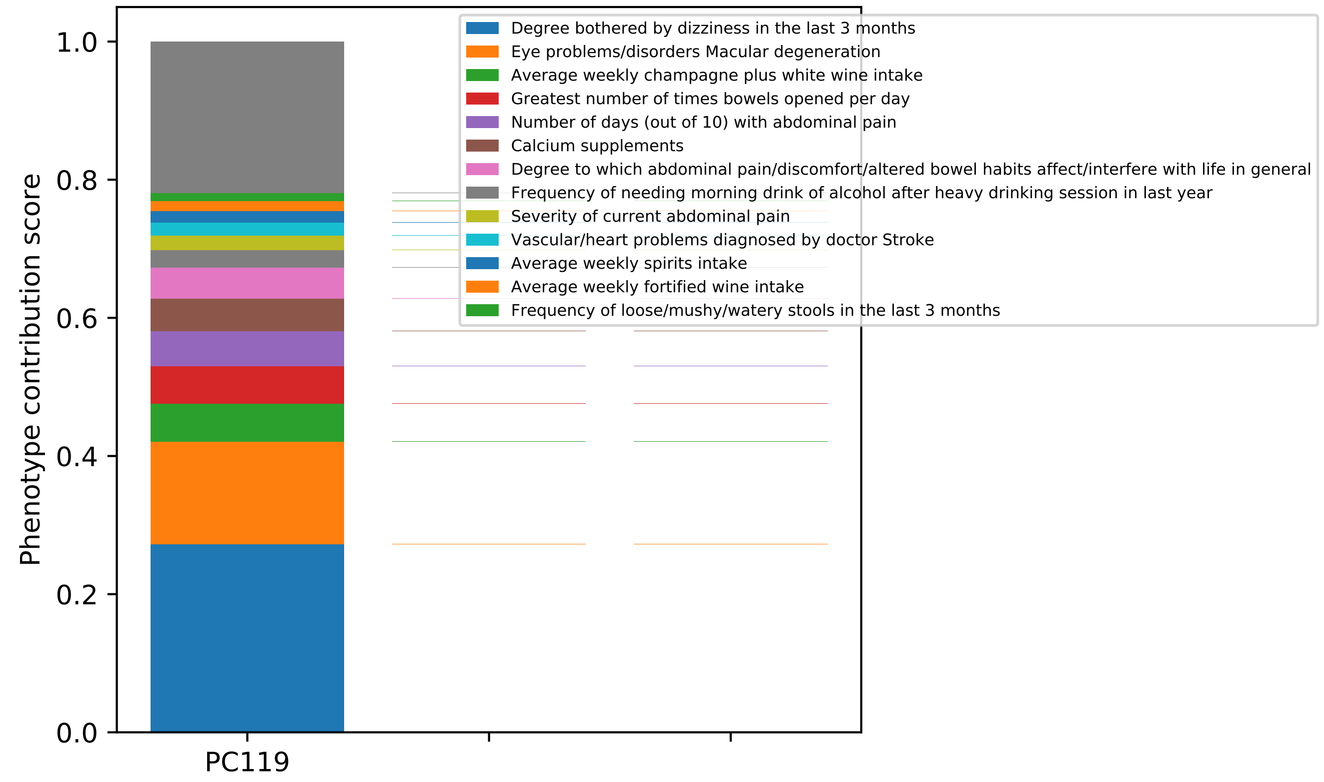
Phenotype contribution score
0.0
0.2
0.4
0.6
0.8
1.0
PC119
Degree bothered by dizziness in the last 3 months
Eye problems/disorders Macular degeneration
Average weekly champagne plus white wine intake
Greatest number of times bowels opened per day
Number of days (out of 10) with abdominal pain
Calcium supplements
Degree to which abdominal pain/discomfort/altered bowel habits affect/interfere with life in general
Frequency of needing morning drink of alcohol after heavy drinking session in last year
Severity of current abdominal pain
Vascular/heart problems diagnosed by doctor Stroke
Average weekly spirits intake
Average weekly fortified wine intake
Frequency of loose/mushy/watery stools in the last 3 months

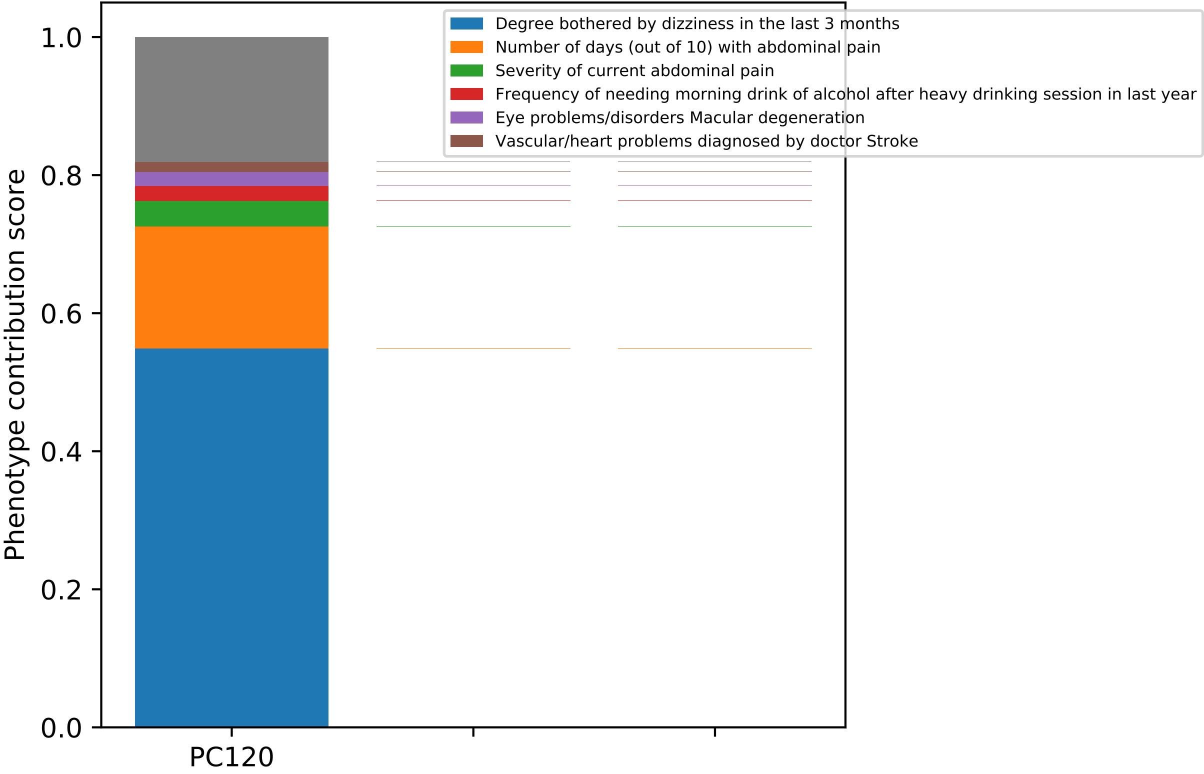

Degree bothered by dizziness in the last 3 months
Number of days (out of 10) with abdominal pain
Severity of current abdominal pain
Frequency of needing morning drink of alcohol after heavy drinking session in last year
Eye problems/disorders Macular degeneration
Vascular/heart problems diagnosed by doctor Stroke
Phenotype contribution score
1.0
0.8
0.6
0.4
0.2
0.0
PC120

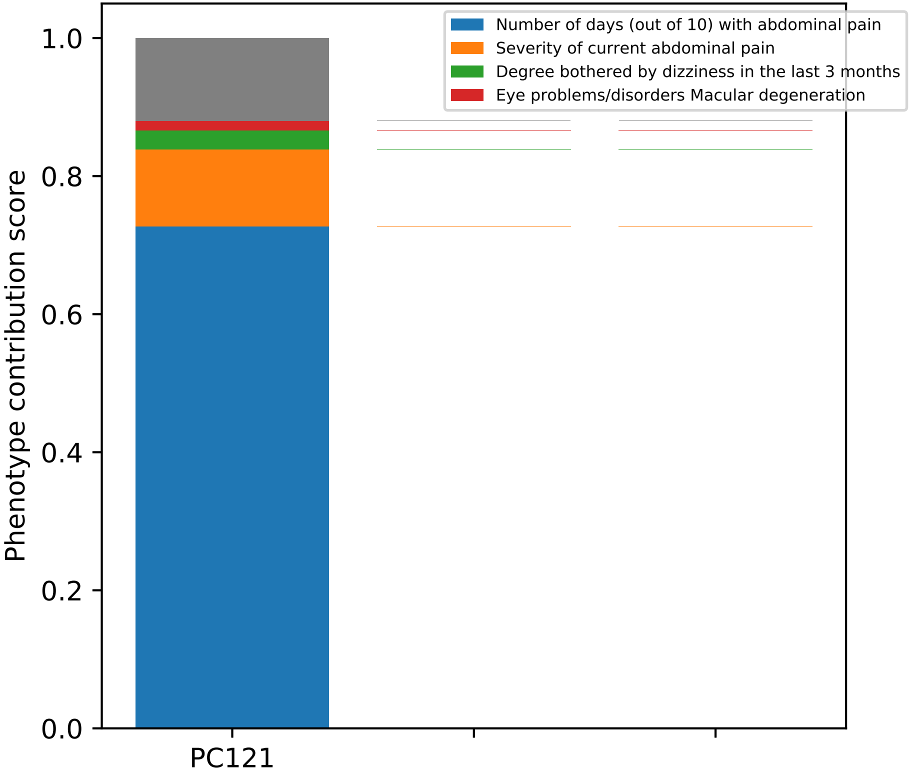
Number of days (out of 10) with abdominal pain
Severity of current abdominal pain
Degree bothered by dizziness in the last 3 months
Eye problems/disorders Macular degeneration
Phenotype contribution score
1.0
0.8
0.6
0.4
0.2
0.0
PC121

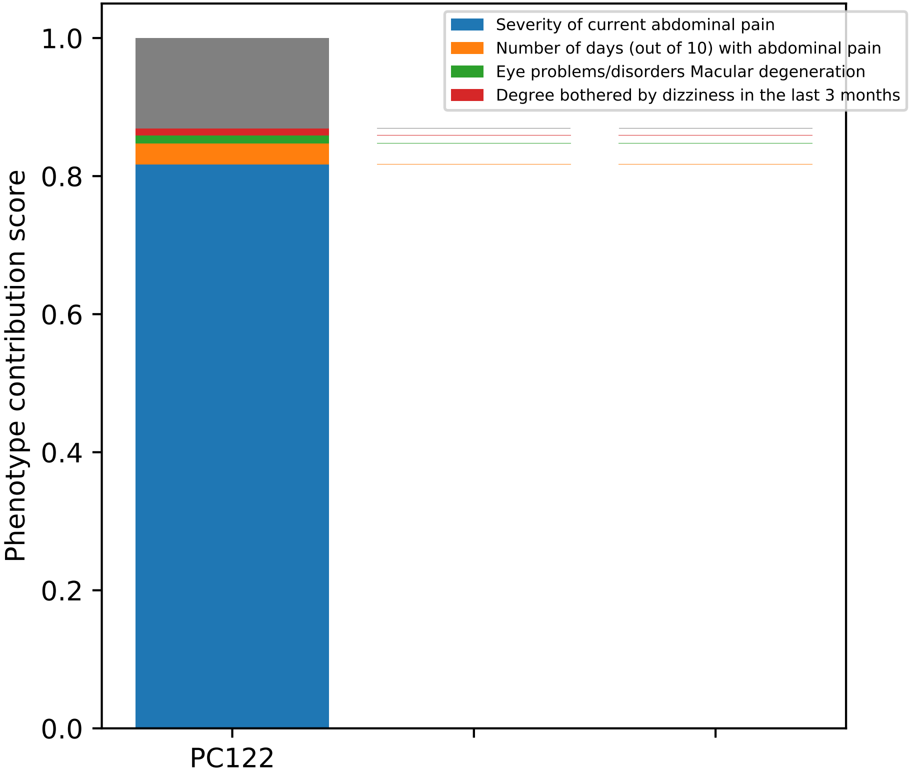
Severity of current abdominal pain
Number of days (out of 10) with abdominal pain
Eye problems/disorders Macular degeneration
Degree bothered by dizziness in the last 3 months
Phenotype contribution score
1.0
0.8
0.6
0.4
0.2
0.0
PC122

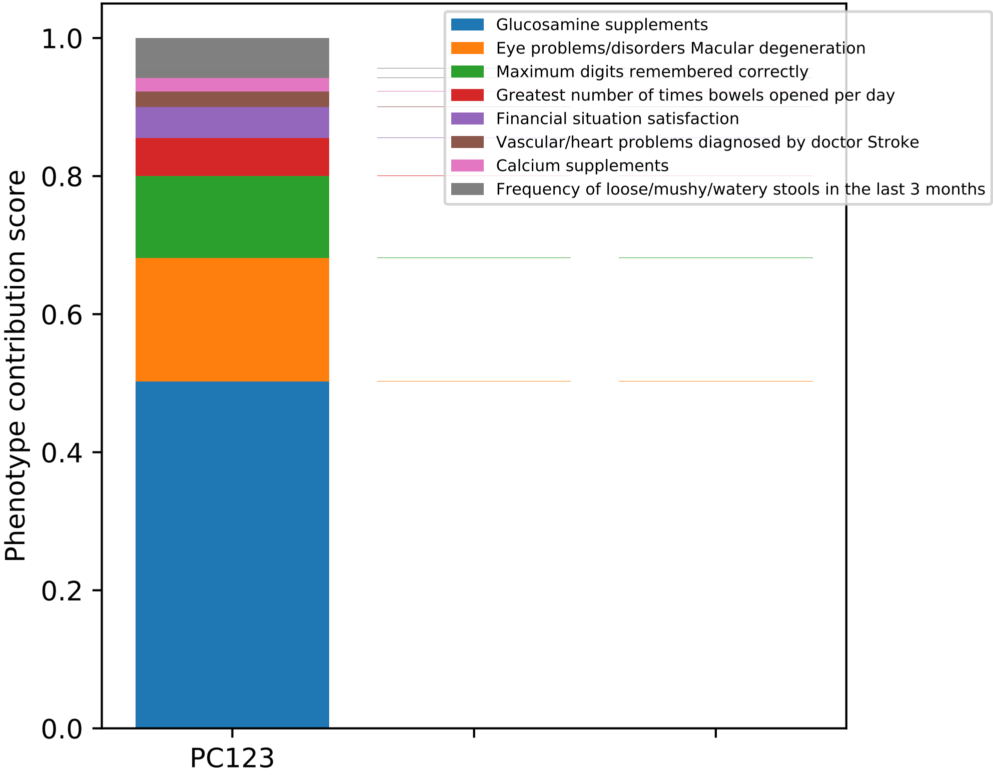
Glucosamine supplements
Eye problems/disorders Macular degeneration
Maximum digits remembered correctly
Greatest number of times bowels opened per day
Financial situation satisfaction
Vascular/heart problems diagnosed by doctor Stroke
Calcium supplements
Frequency of loose/mushy/watery stools in the last 3 months
Phenotype contribution score
1.0
0.8
0.6
0.4
0.2
0.0
PC123

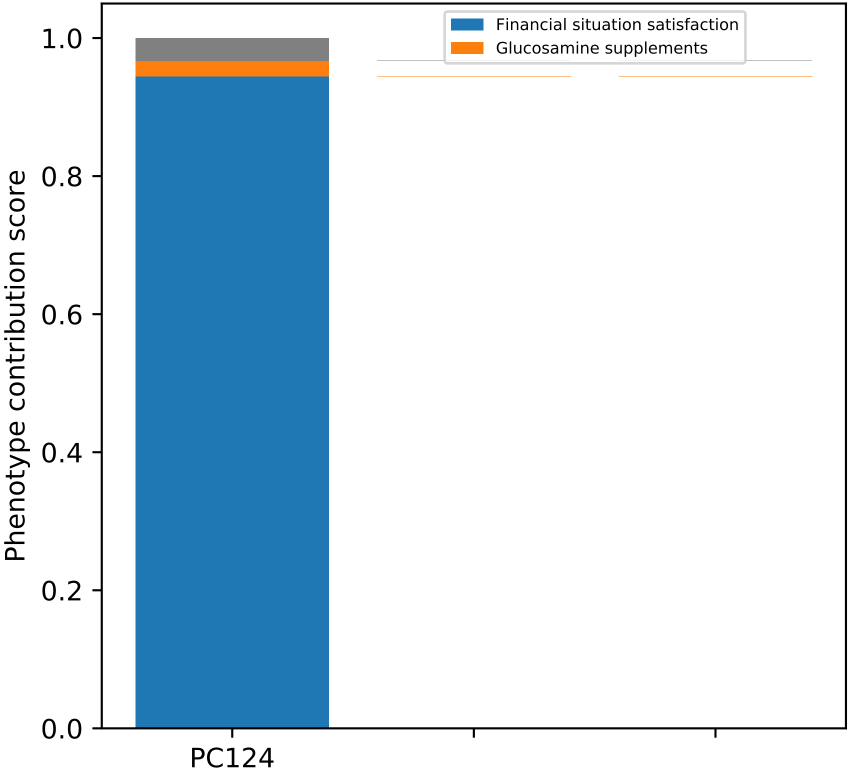
Financial situation satisfaction
Glucosamine supplements
Phenotype contribution score
1.0
0.8
0.6
0.4
0.2
0.0
PC124

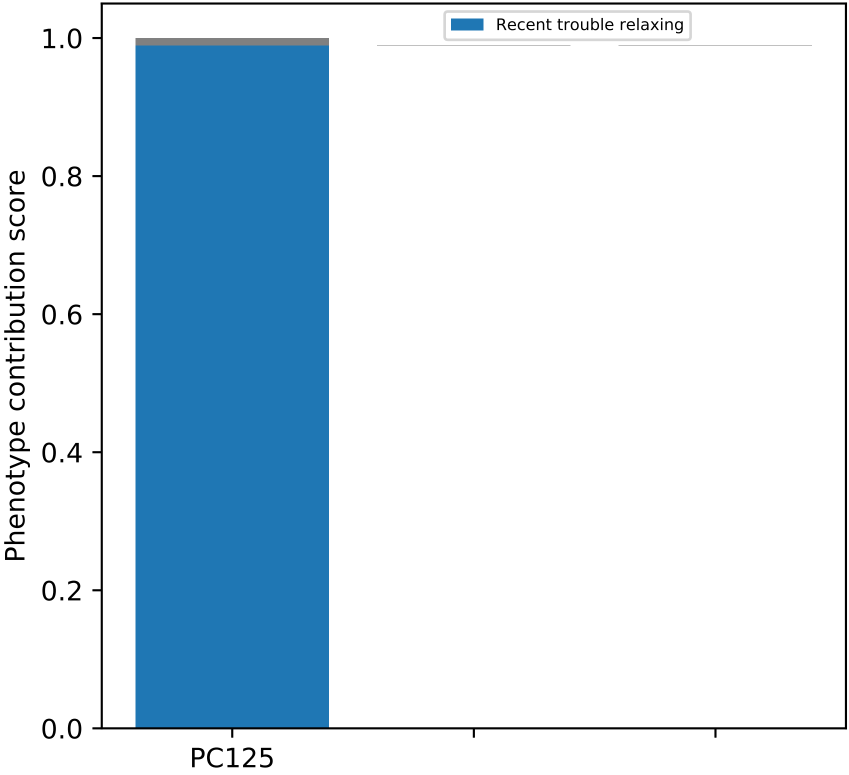
Recent trouble relaxing
Phenotype contribution score
1.0
0.8
0.6
0.4
0.2
0.0
PC125

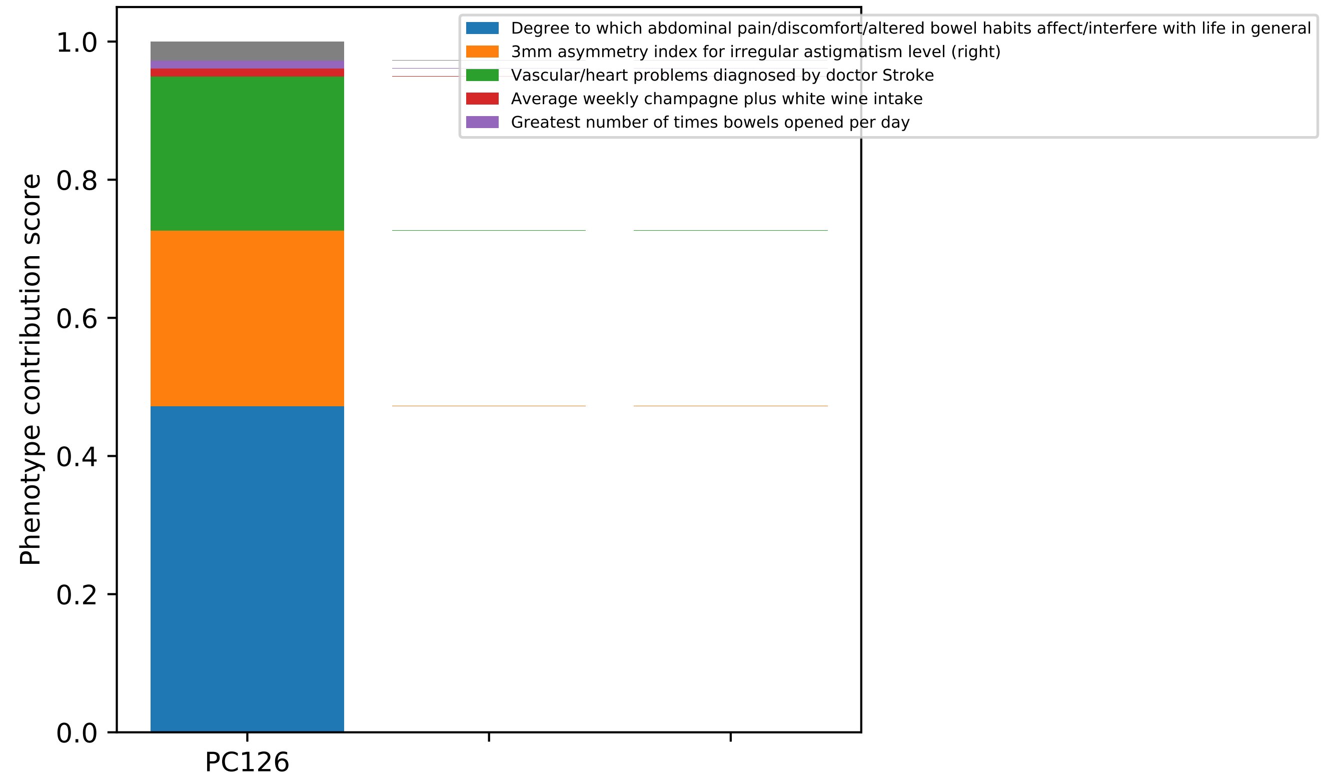

Phenotype contribution score
1.0
0.8
0.6
0.4
0.2
0.0
PC126
Degree to which abdominal pain/discomfort/altered bowel habits affect/interfere with life in general
3mm asymmetry index for irregular astigmatism level (right)
Vascular/heart problems diagnosed by doctor Stroke
Average weekly champagne plus white wine intake
Greatest number of times bowels opened per day

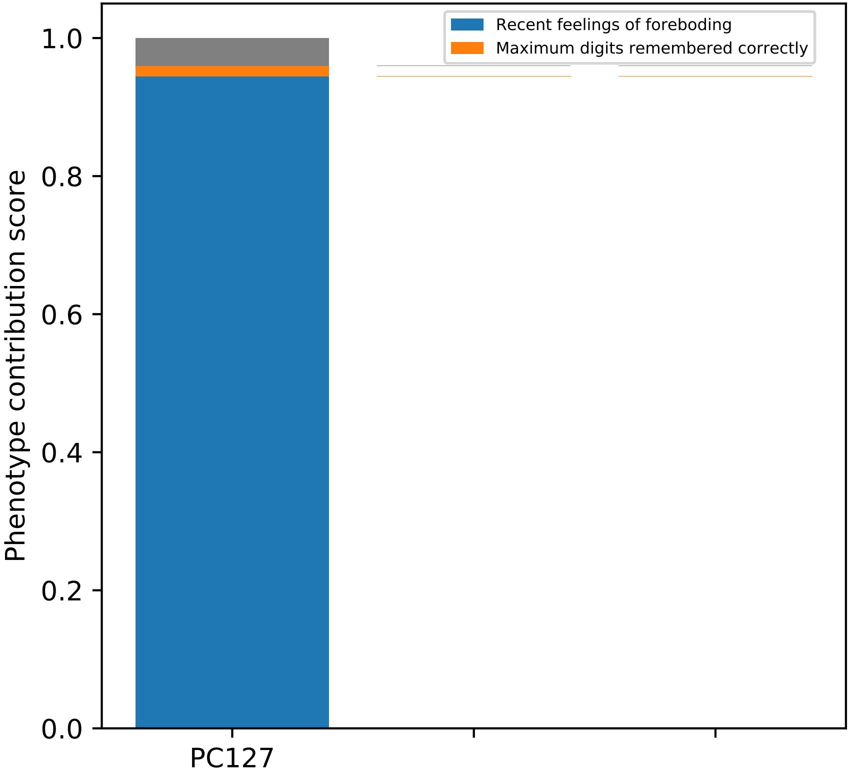
Recent feelings of foreboding
Maximum digits remembered correctly
Phenotype contribution score
1.0
0.8
0.6
0.4
0.2
0.0
PC127

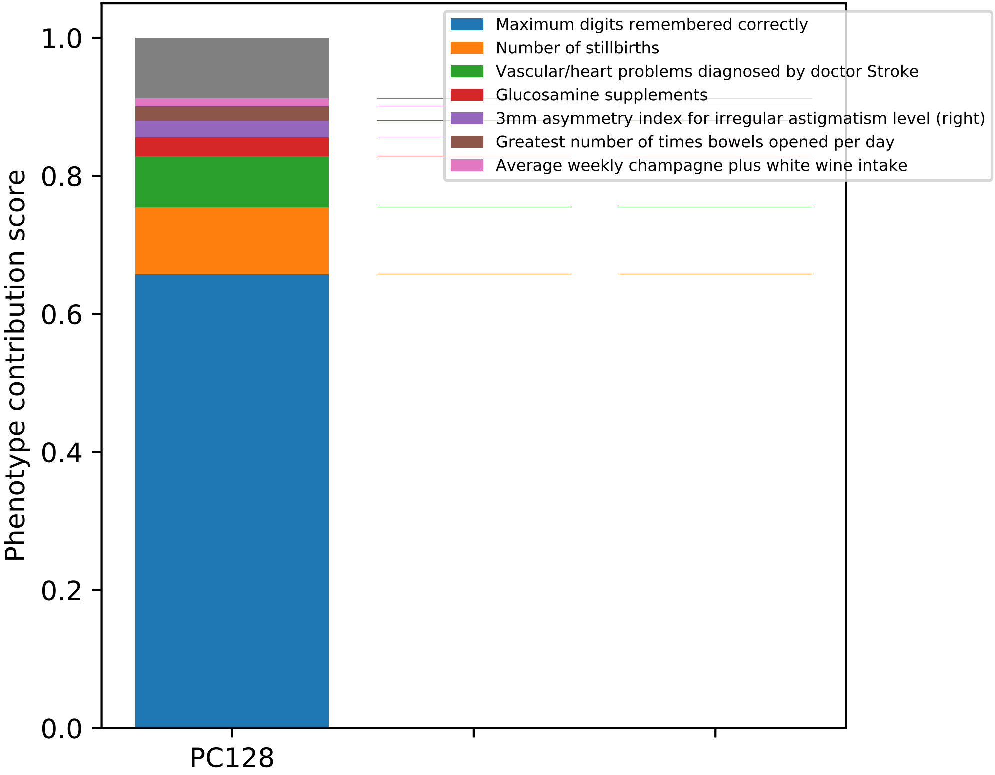
Maximum digits remembered correctly
Number of stillbirths
Vascular/heart problems diagnosed by doctor Stroke
Glucosamine supplements
3mm asymmetry index for irregular astigmatism level (right)
Greatest number of times bowels opened per day
Average weekly champagne plus white wine intake
Phenotype contribution score
1.0
0.8
0.6
0.4
0.2
0.0
PC128

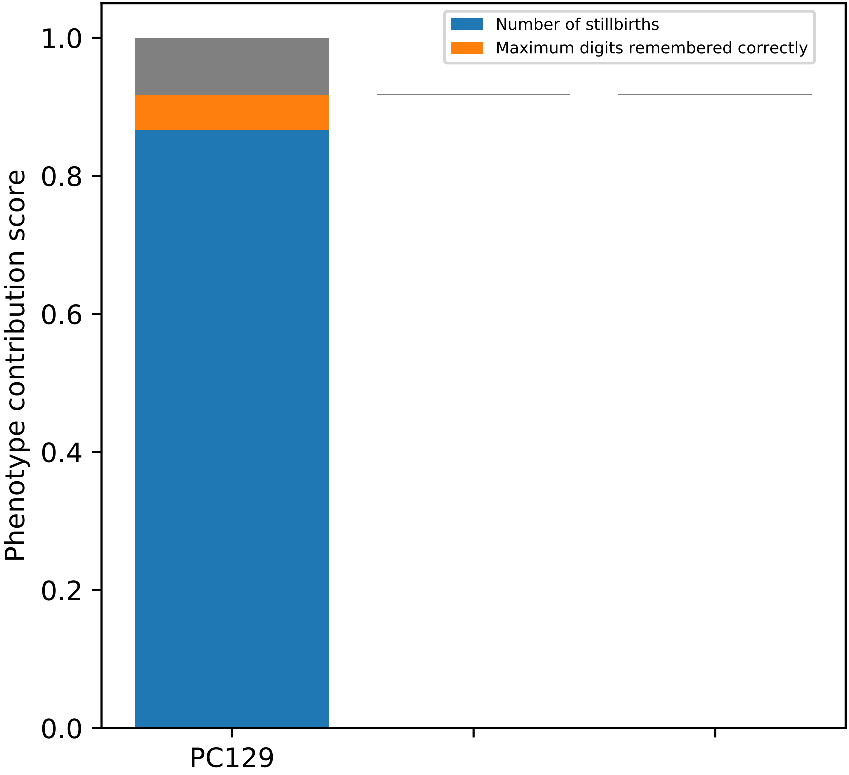
Number of stillbirths
Maximum digits remembered correctly
Phenotype contribution score
0.0
0.2
0.4
0.6
0.8
1.0
PC129

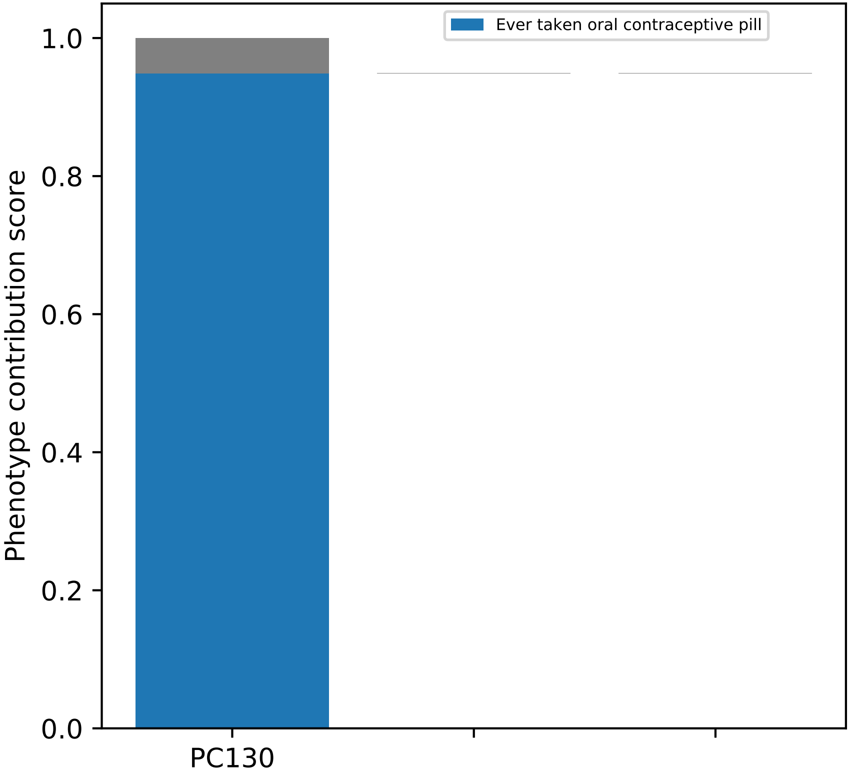

Ever taken oral contraceptive pill
Phenotype contribution score
1.0
0.8
0.6
0.4
0.2
0.0
PC130

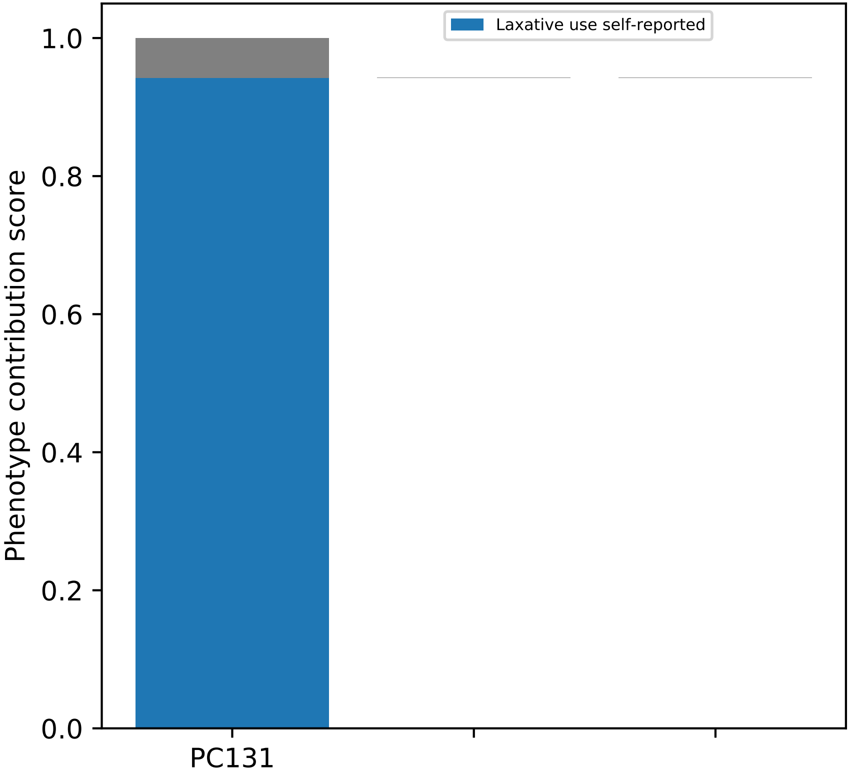
Laxative use self-reported
Phenotype contribution score
1.0
0.8
0.6
0.4
0.2
0.0
PC131

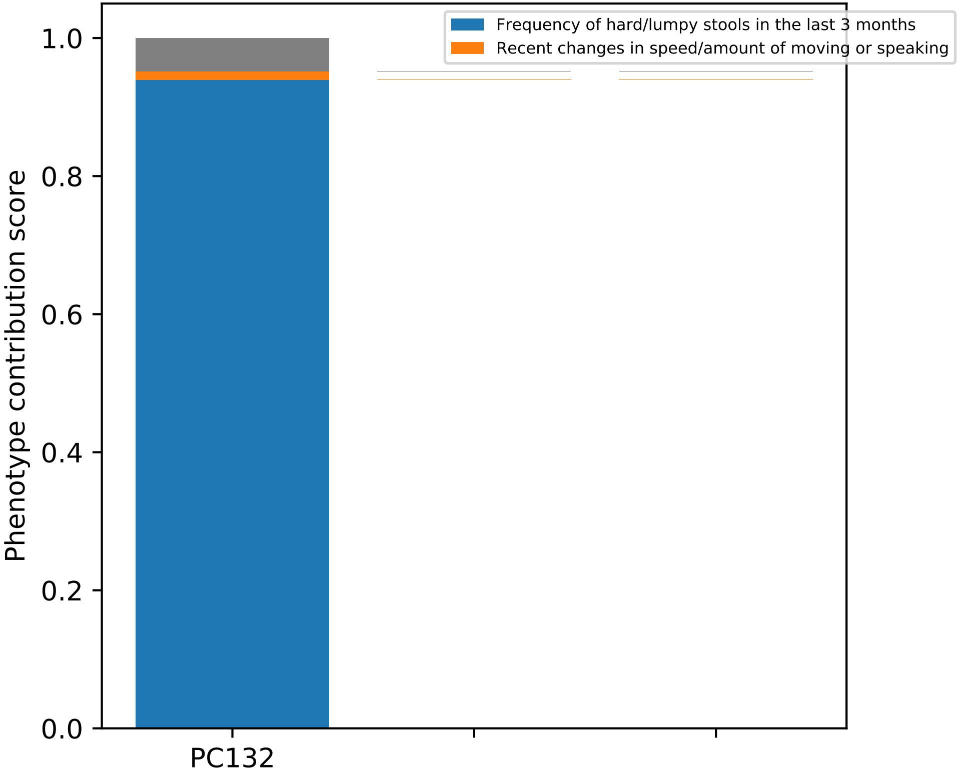
Frequency of hard/lumpy stools in the last 3 months
Recent changes in speed/amount of moving or speaking
Phenotype contribution score
0.0
0.2
0.4
0.6
0.8
1.0
PC132

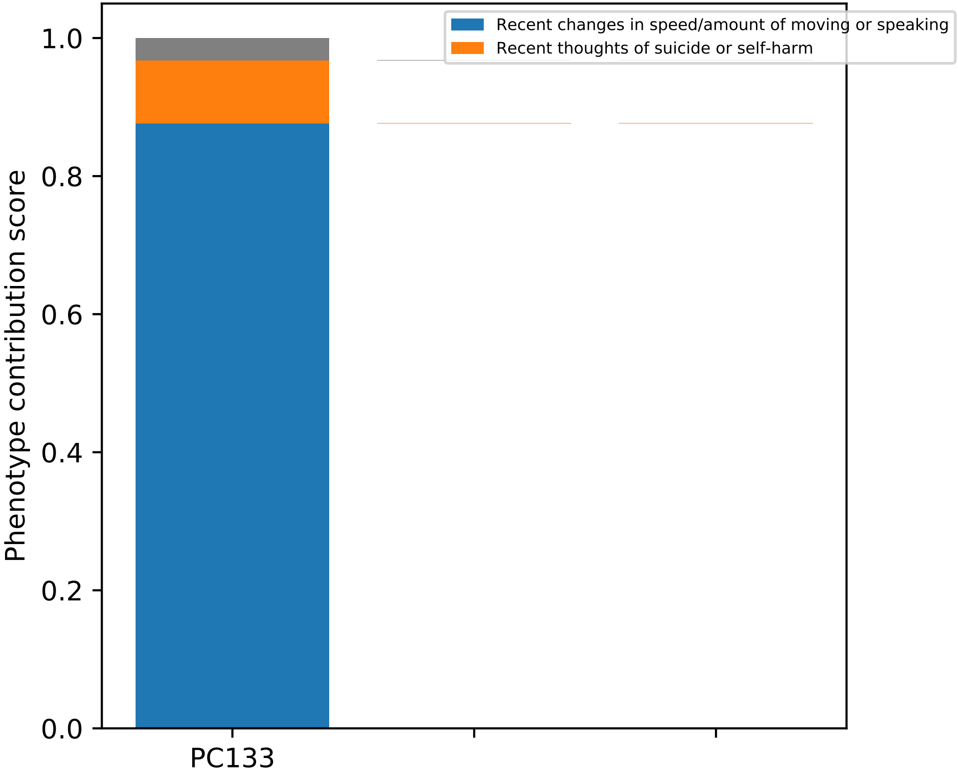
Recent changes in speed/amount of moving or speaking
Recent thoughts of suicide or self-harm
Phenotype contribution score
1.0
0.8
0.6
0.4
0.2
0.0
PC133

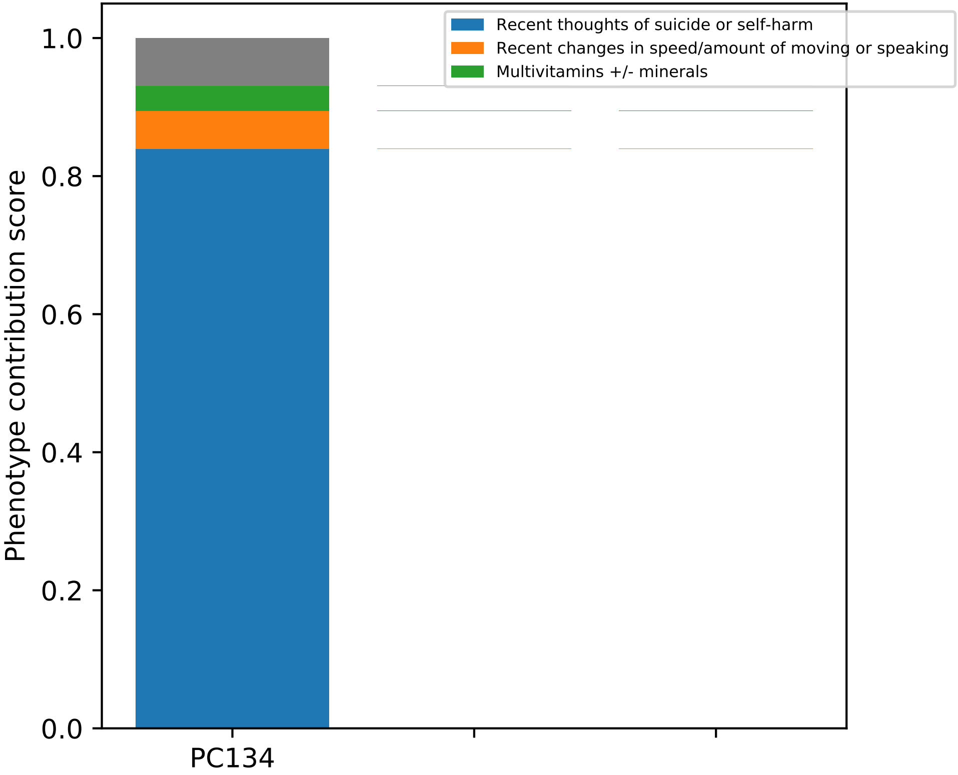

Recent thoughts of suicide or self-harm
Recent changes in speed/amount of moving or speaking
Multivitamins +/- minerals
Phenotype contribution score
1.0
0.8
0.6
0.4
0.2
0.0
PC134

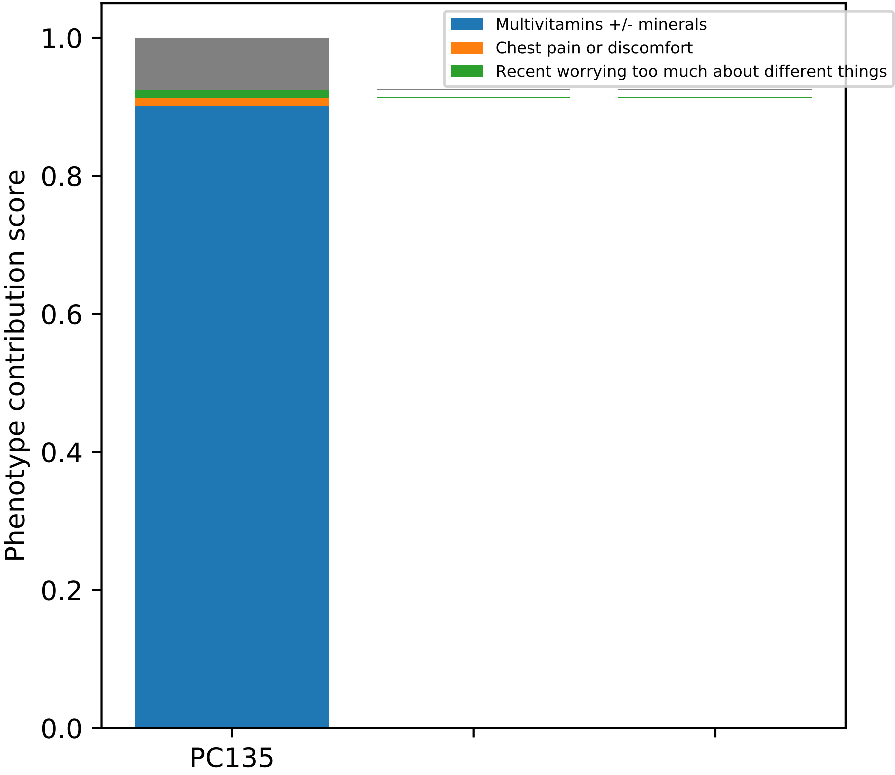

Multivitamins +/- minerals
Chest pain or discomfort
Recent worrying too much about different things
Phenotype contribution score
1.0
0.8
0.6
0.4
0.2
0.0
PC135

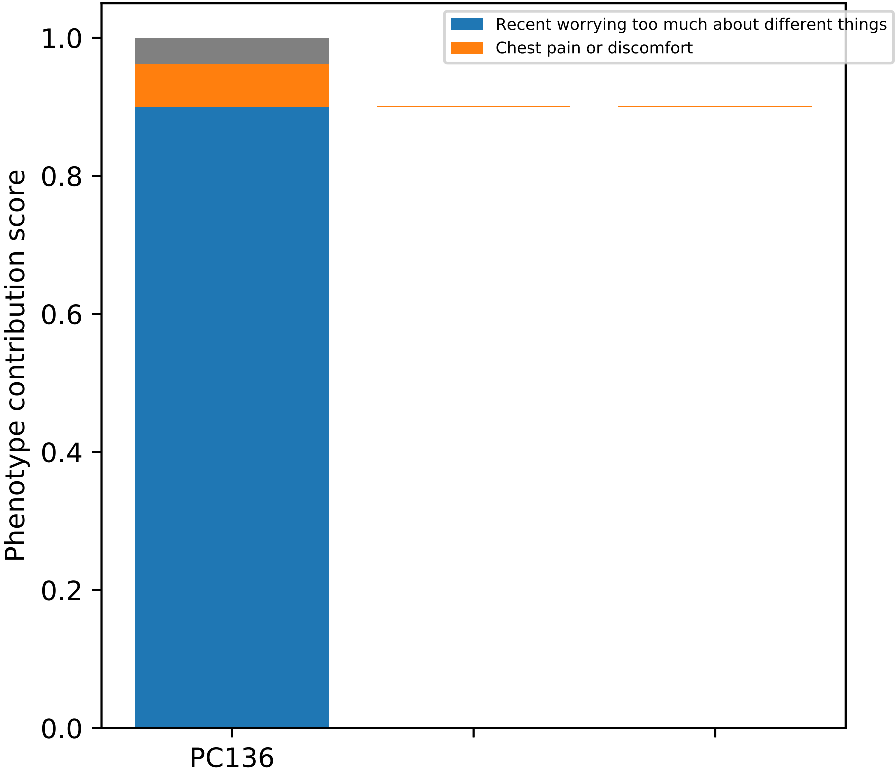

Recent worrying too much about different things
Chest pain or discomfort
Phenotype contribution score
1.0
0.8
0.6
0.4
0.2
0.0
PC136

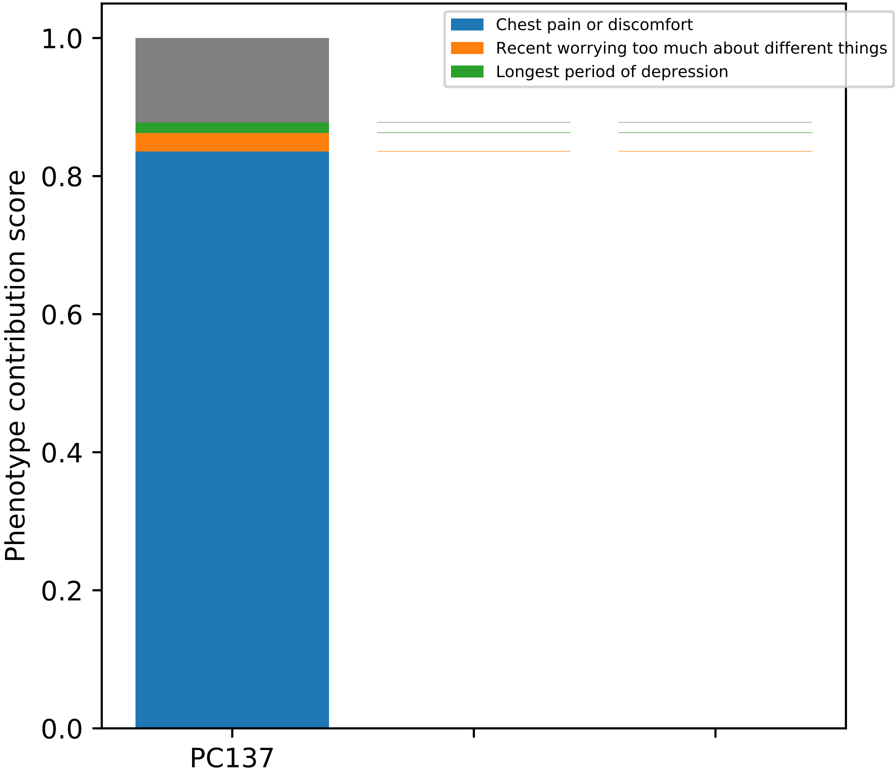
Chest pain or discomfort
Recent worrying too much about different things
Longest period of depression
1.0
0.8
0.6
0.4
0.2
0.0
Phenotype contribution score
PC137

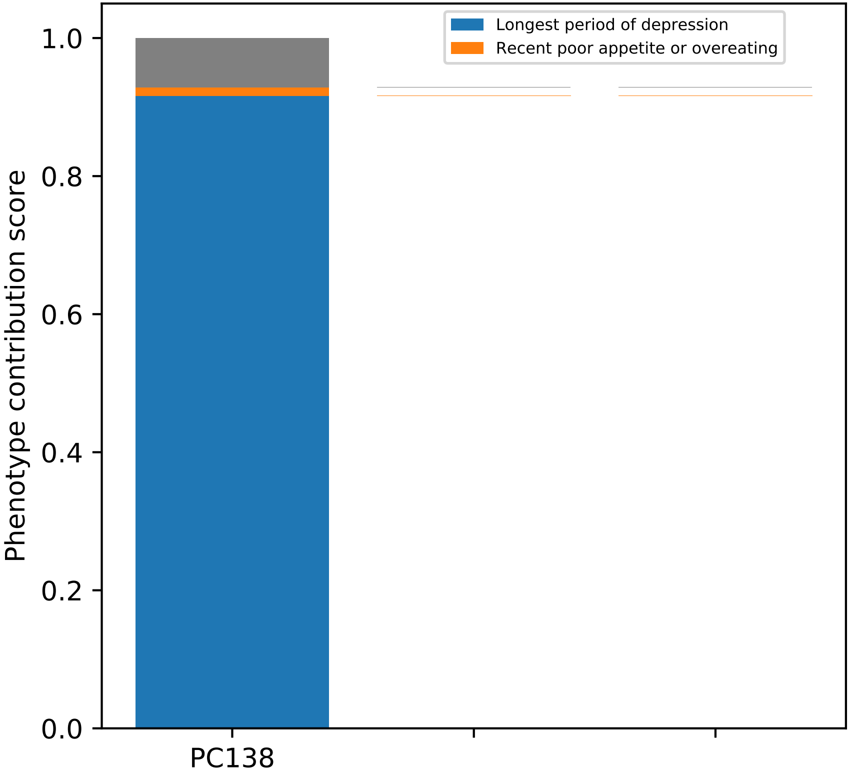
Longest period of depression
Recent poor appetite or overeating
Phenotype contribution score
1.0
0.8
0.6
0.4
0.2
0.0
PC138

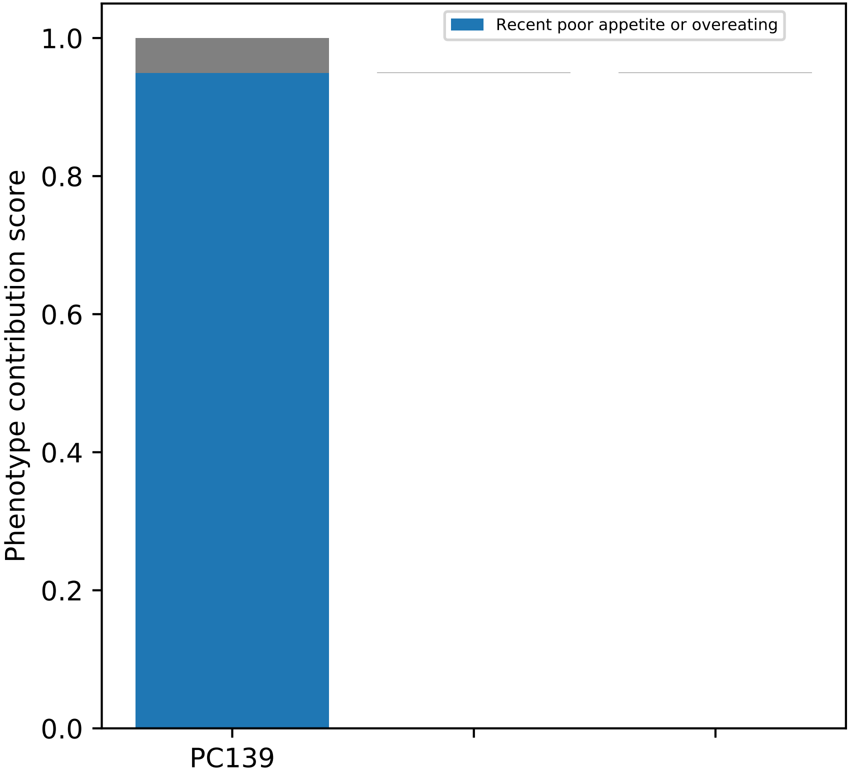
Recent poor appetite or overeating
Phenotype contribution score
1.0
0.8
0.6
0.4
0.2
0.0
PC139

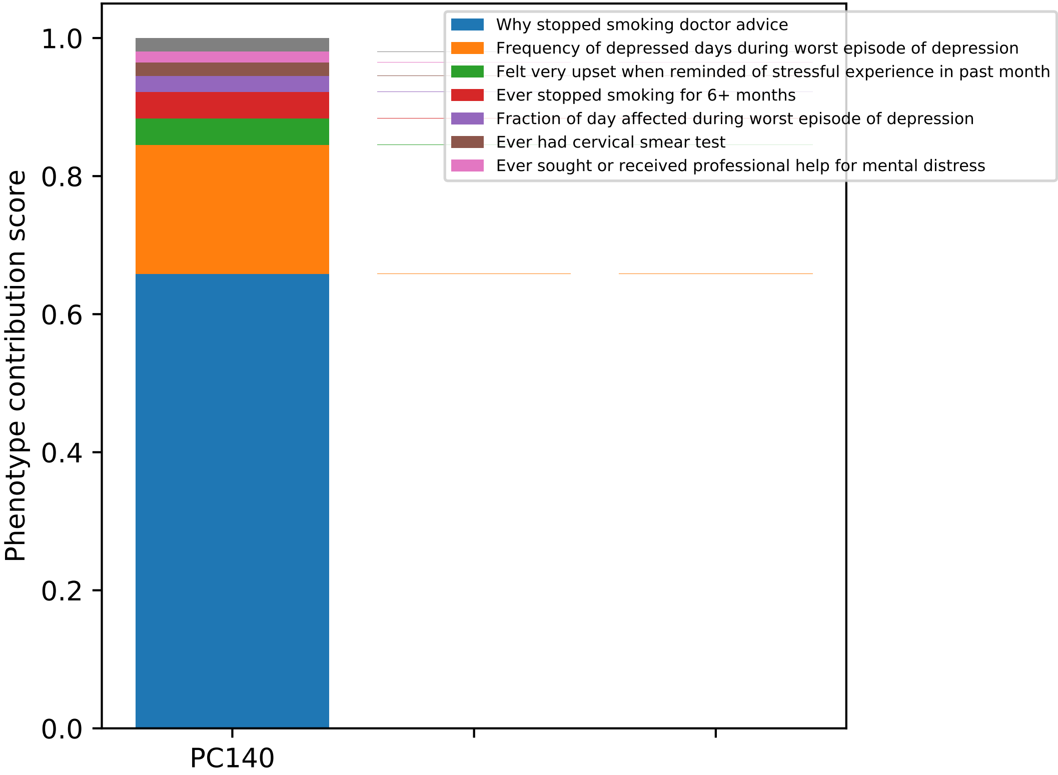

Phenotype contribution score
1.0
0.8
0.6
0.4
0.2
0.0
PC140
Why stopped smoking doctor advice
Frequency of depressed days during worst episode of depression
Felt very upset when reminded of stressful experience in past month
Ever stopped smoking for 6+ months
Fraction of day affected during worst episode of depression
Ever had cervical smear test
Ever sought or received professional help for mental distress

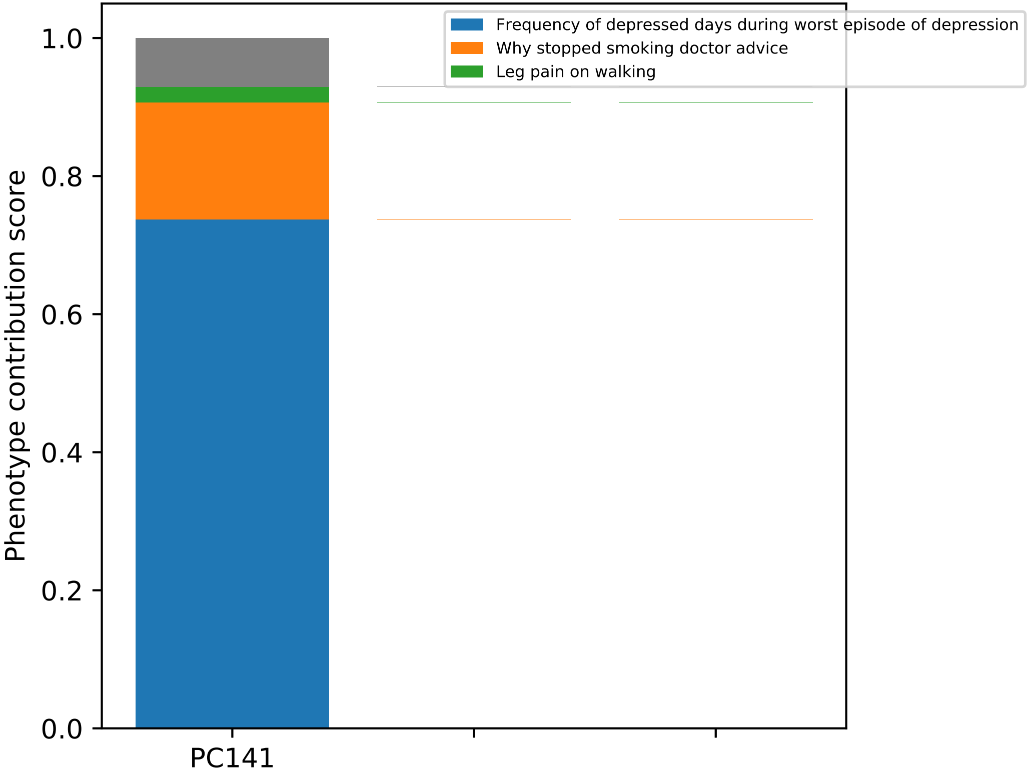

Frequency of depressed days during worst episode of depression
Why stopped smoking doctor advice
Leg pain on walking
Phenotype contribution score
1.0
0.8
0.6
0.4
0.2
0.0
PC141

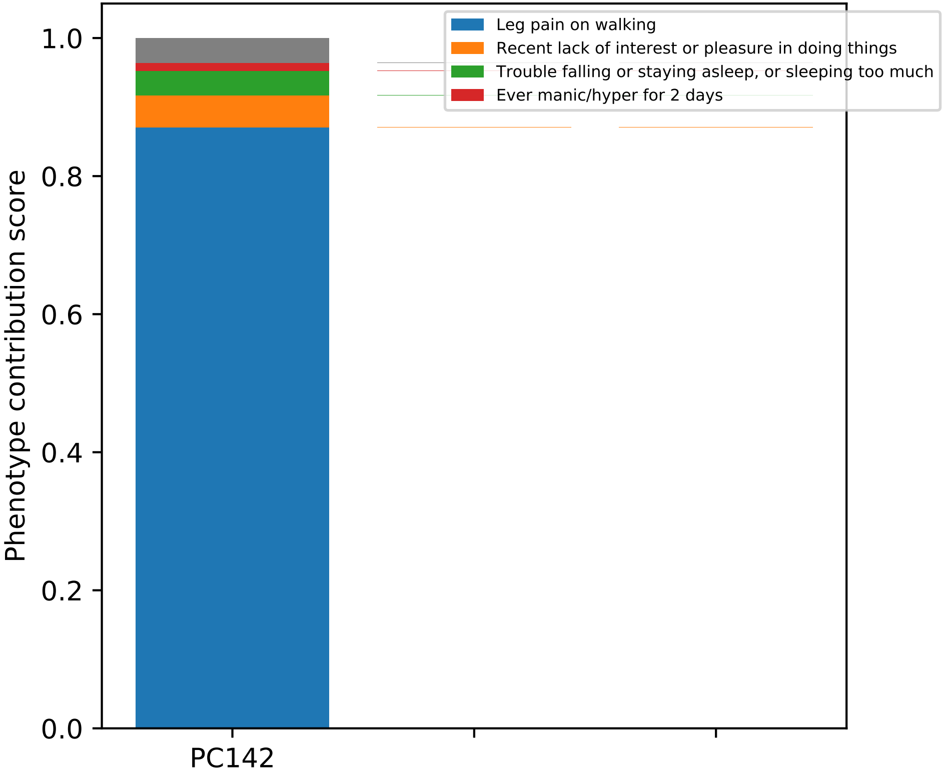

Leg pain on walking
Recent lack of interest or pleasure in doing things
Trouble falling or staying asleep, or sleeping too much
Ever manic/hyper for 2 days
Phenotype contribution score
1.0
0.8
0.6
0.4
0.2
0.0
PC142

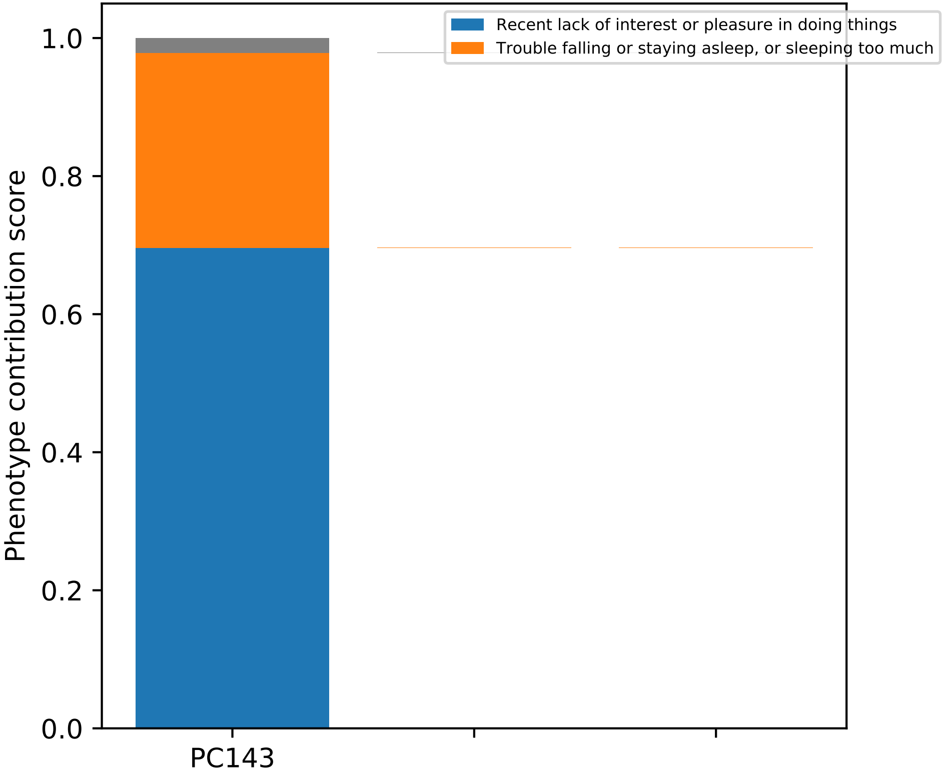
Recent lack of interest or pleasure in doing things
Trouble falling or staying asleep, or sleeping too much
Phenotype contribution score
0.0
0.2
0.4
0.6
0.8
1.0
PC143

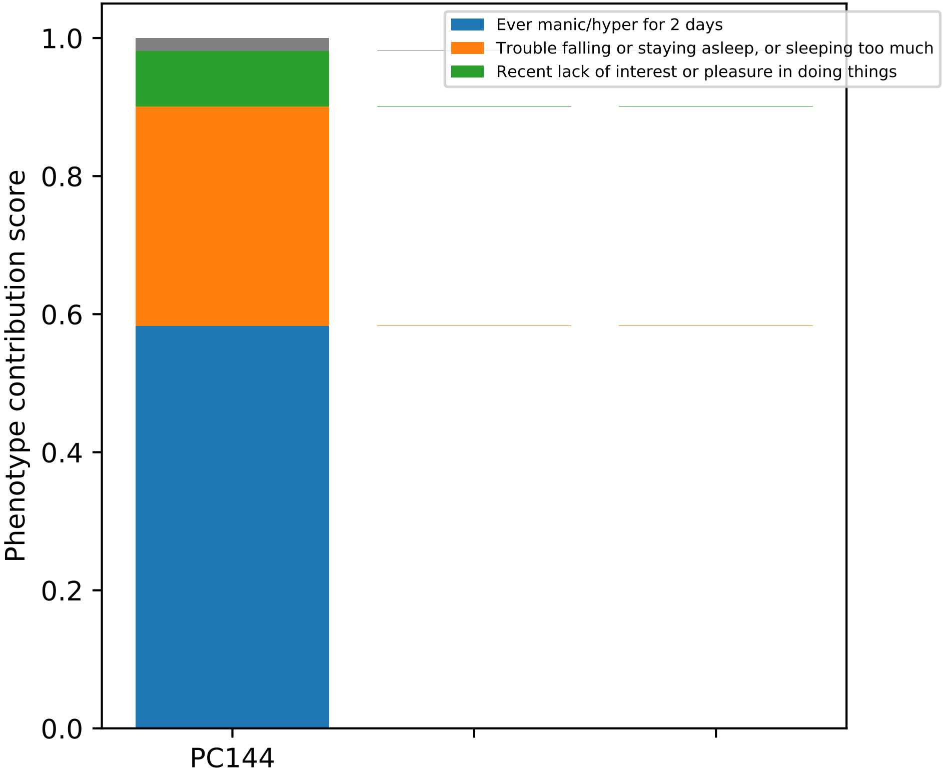

Ever manic/hyper for 2 days
Trouble falling or staying asleep, or sleeping too much
Recent lack of interest or pleasure in doing things
Phenotype contribution score
1.0
0.8
0.6
0.4
0.2
0.0
PC144

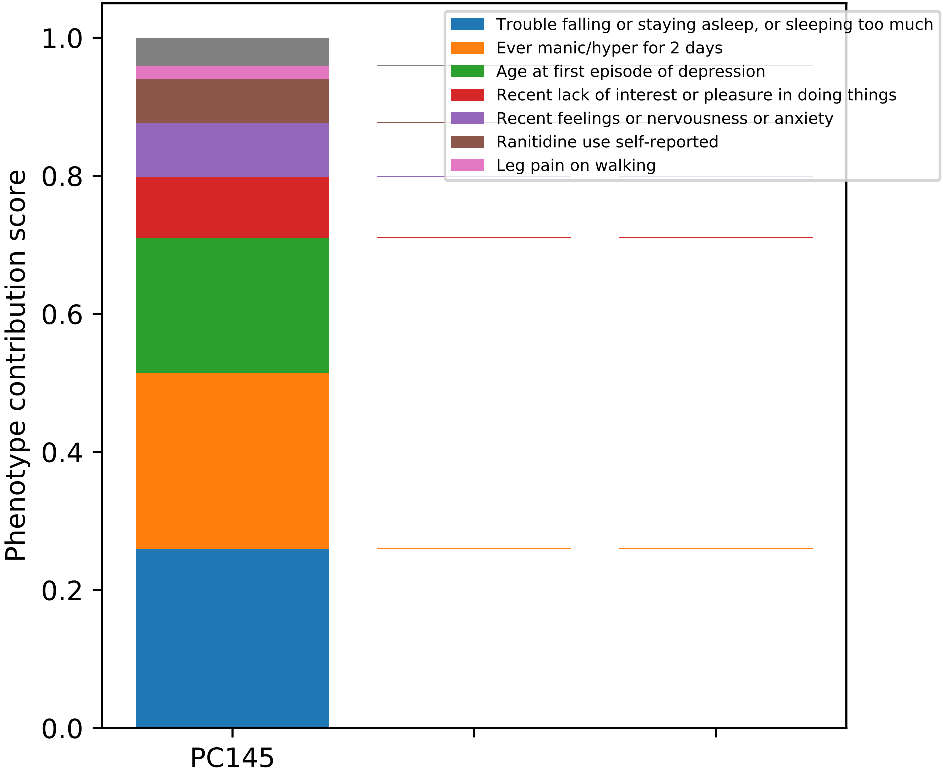

Phenotype contribution score
1.0
0.8
0.6
0.4
0.2
0.0
PC145
Trouble falling or staying asleep, or sleeping too much
Ever manic/hyper for 2 days
Age at first episode of depression
Recent lack of interest or pleasure in doing things
Recent feelings or nervousness or anxiety
Ranitidine use self-reported
Leg pain on walking

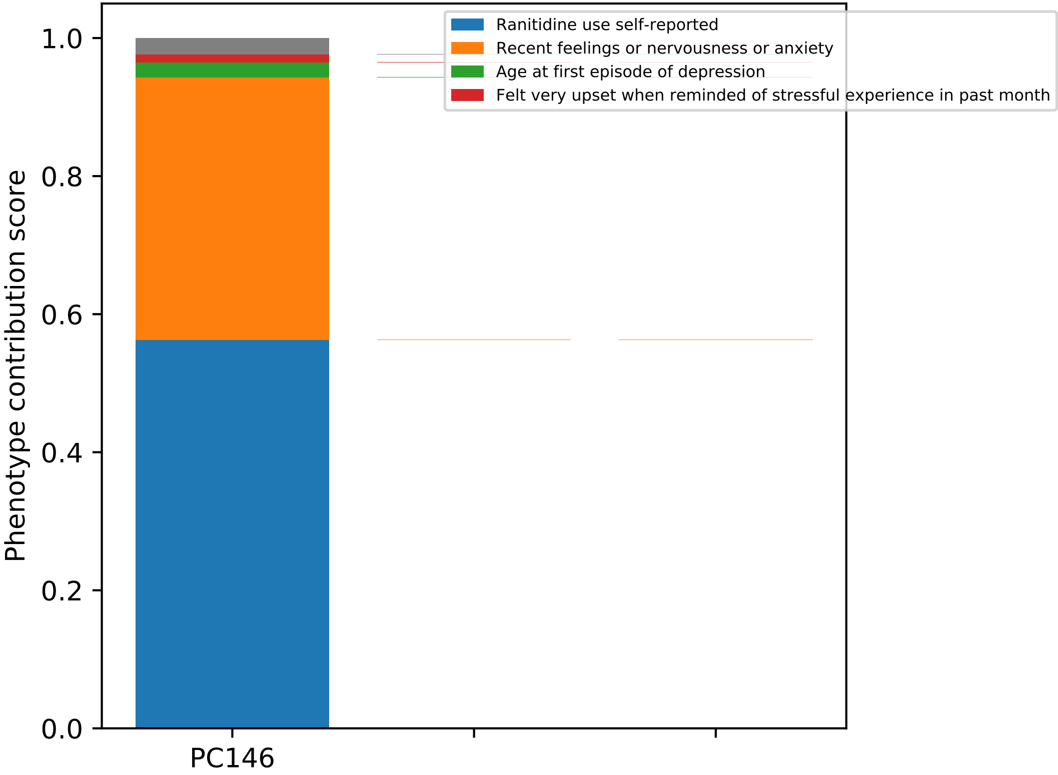

Ranitidine use self-reported
Recent feelings or nervousness or anxiety
Age at first episode of depression
Felt very upset when reminded of stressful experience in past month
Phenotype contribution score
1.0
0.8
0.6
0.4
0.2
0.0
PC146

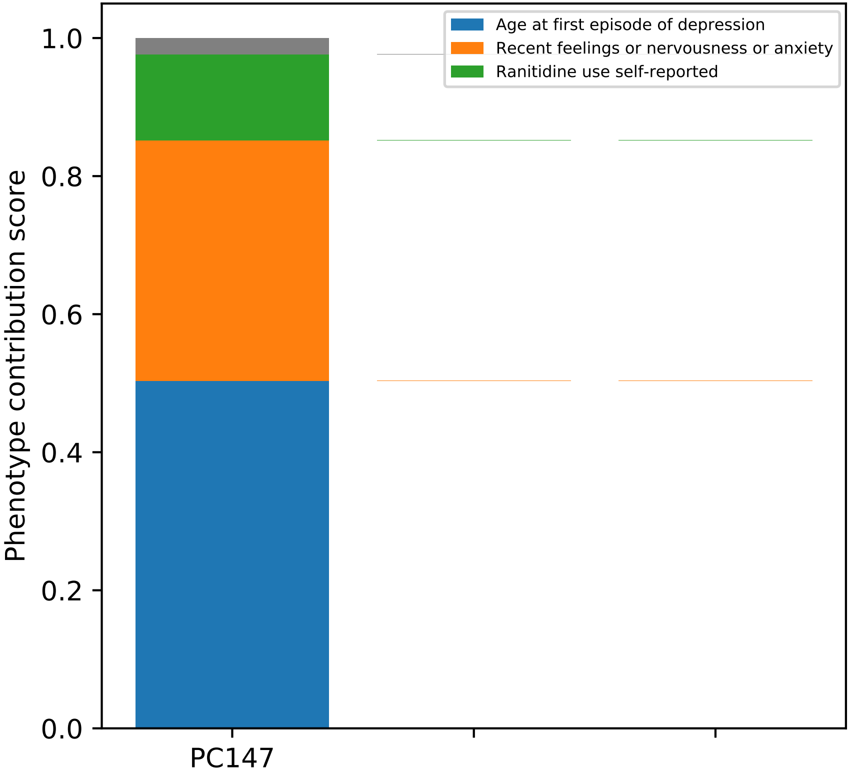

Age at first episode of depression
Recent feelings or nervousness or anxiety
Ranitidine use self-reported
Phenotype contribution score
1.0
0.8
0.6
0.4
0.2
0.0
PC147

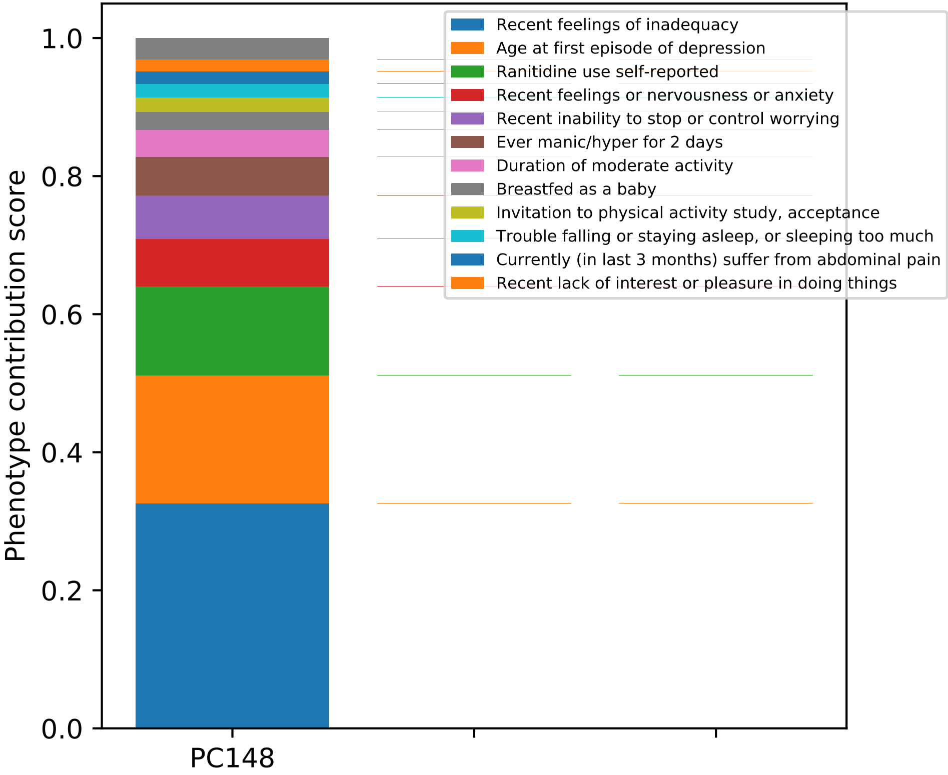

Recent feelings of inadequacy
Age at first episode of depression
Ranitidine use self-reported
Recent feelings or nervousness or anxiety
Recent inability to stop or control worrying
Ever manic/hyper for 2 days
Duration of moderate activity
Breastfed as a baby
Invitation to physical activity study, acceptance
Trouble falling or staying asleep, or sleeping too much
Currently (in last 3 months) suffer from abdominal pain
Recent lack of interest or pleasure in doing things
Phenotype contribution score
0.0
0.2
0.4
0.6
0.8
1.0
PC148

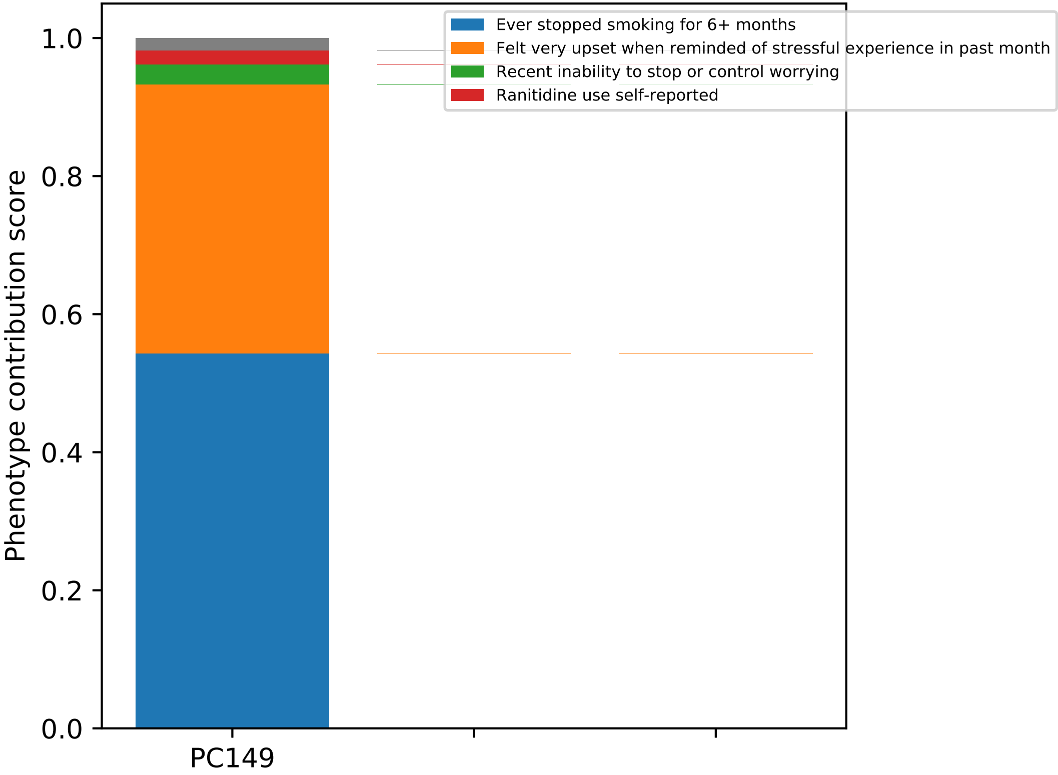

Ever stopped smoking for 6+ months
Felt very upset when reminded of stressful experience in past month
Recent inability to stop or control worrying
Ranitidine use self-reported
Phenotype contribution score
1.0
0.8
0.6
0.4
0.2
0.0
PC149

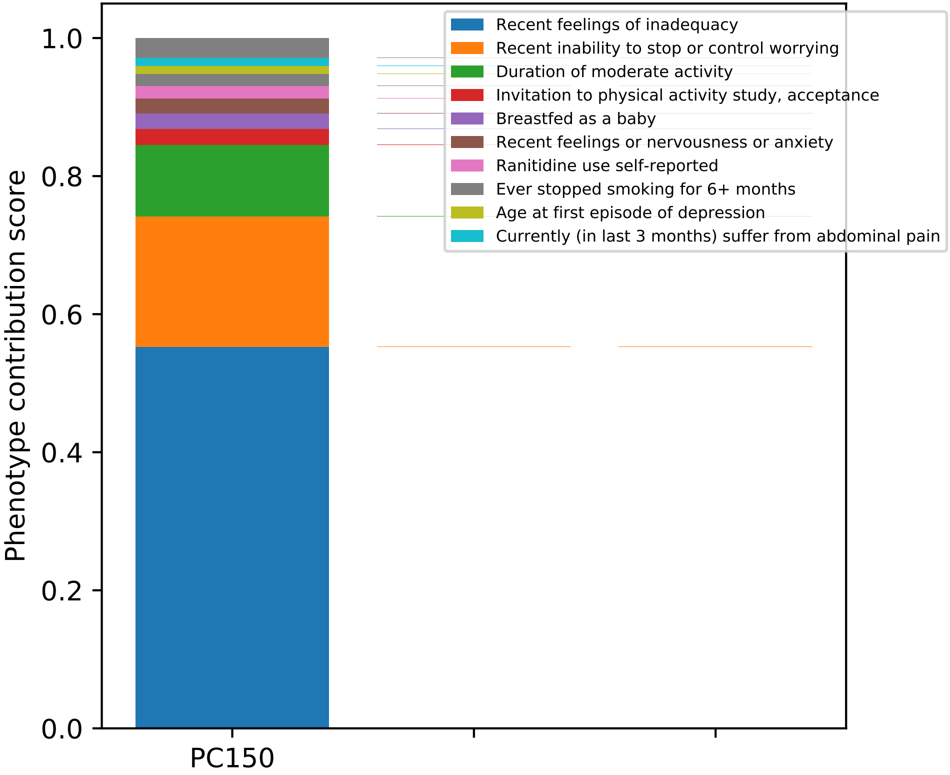

Phenotype contribution score
0.0
0.2
0.4
0.6
0.8
1.0
PC150
Recent feelings of inadequacy
Recent inability to stop or control worrying
Duration of moderate activity
Invitation to physical activity study, acceptance
Breastfed as a baby
Recent feelings or nervousness or anxiety
Ranitidine use self-reported
Ever stopped smoking for 6+ months
Age at first episode of depression
Currently (in last 3 months) suffer from abdominal pain

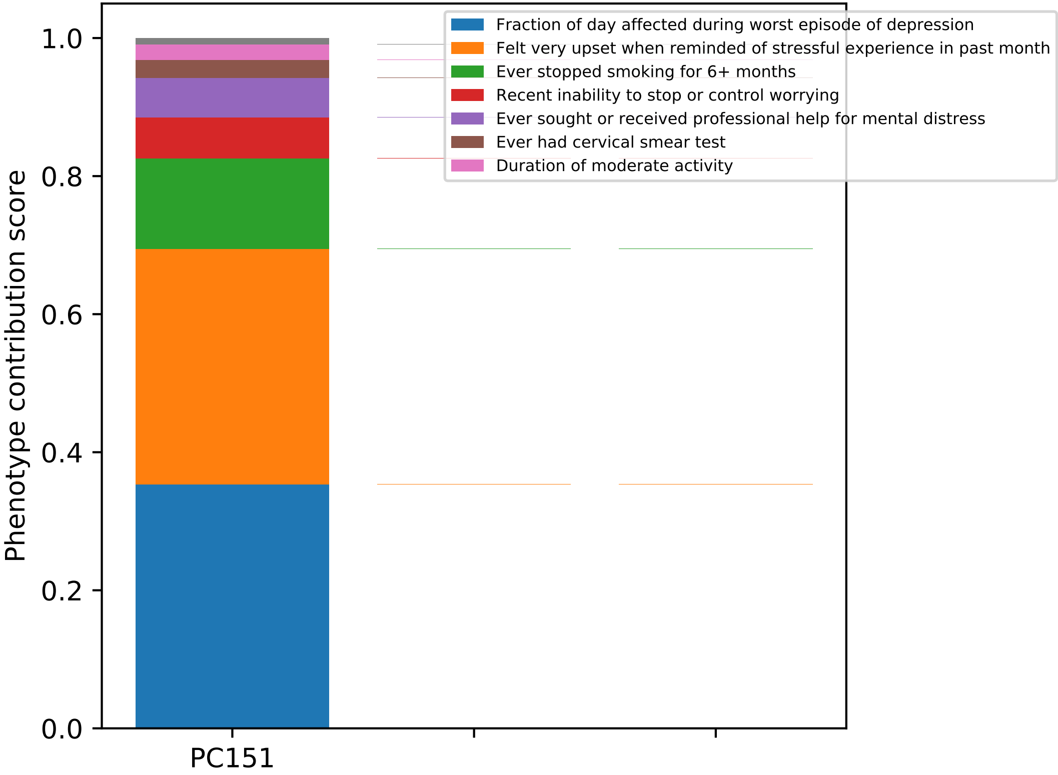

Fraction of day affected during worst episode of depression
Felt very upset when reminded of stressful experience in past month
Ever stopped smoking for 6+ months
Recent inability to stop or control worrying
Ever sought or received professional help for mental distress
Ever had cervical smear test
Duration of moderate activity
Phenotype contribution score
0.0
0.2
0.4
0.6
0.8
1.0
PC151

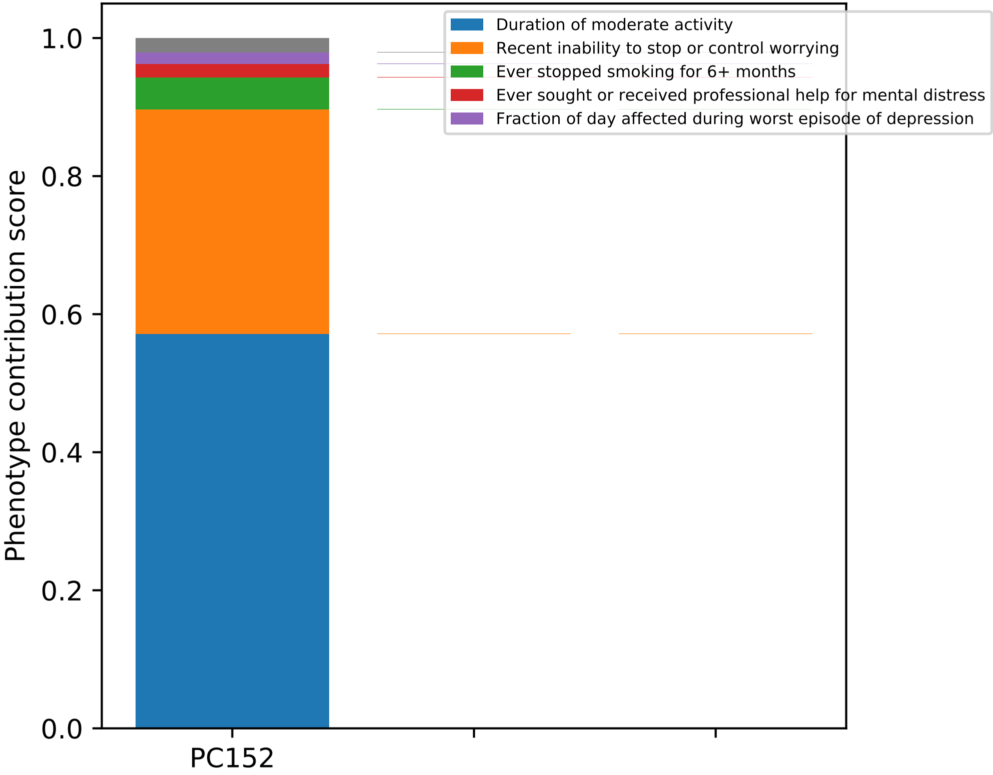
Duration of moderate activity
Recent inability to stop or control worrying
Ever stopped smoking for 6+ months
Ever sought or received professional help for mental distress
Fraction of day affected during worst episode of depression
Phenotype contribution score
1.0
0.8
0.6
0.4
0.2
0.0
PC152

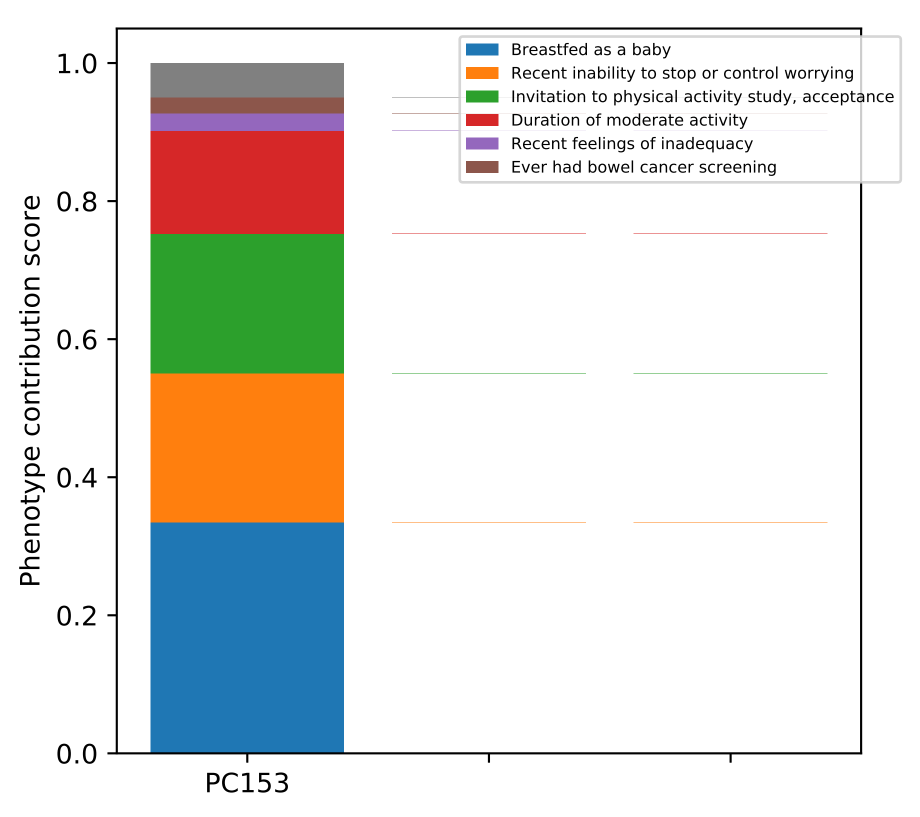
Breastfed as a baby
Recent inability to stop or control worrying
Invitation to physical activity study, acceptance
Duration of moderate activity
Recent feelings of inadequacy
Ever had bowel cancer screening
Phenotype contribution score
0.0
0.2
0.4
0.6
0.8
1.0
PC153

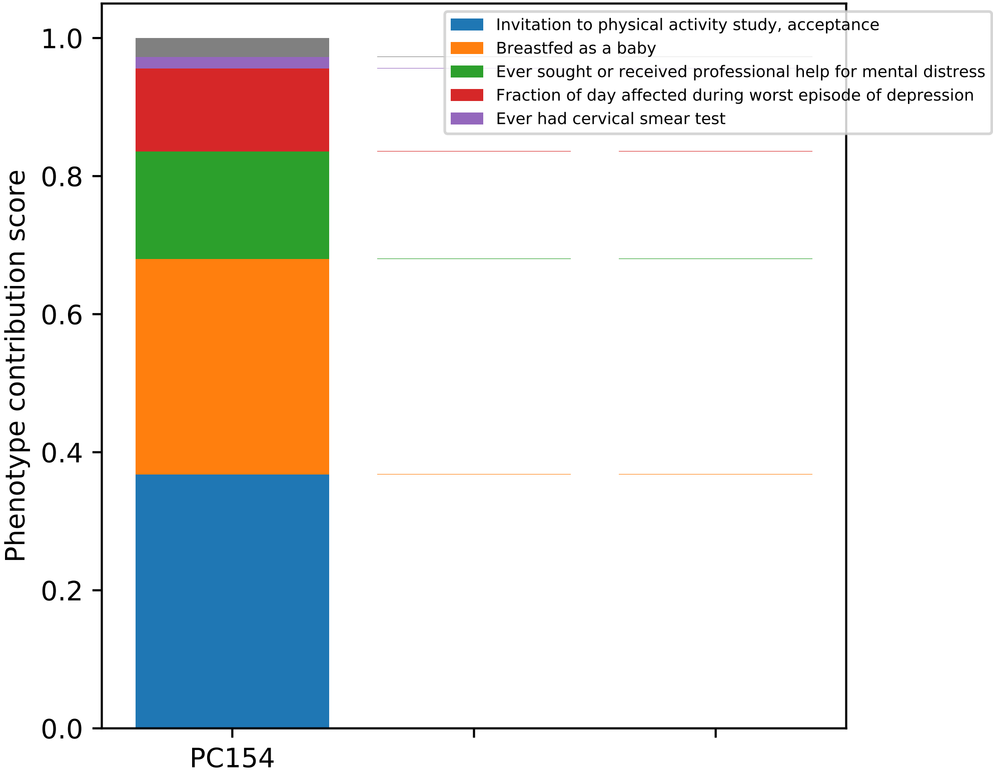
Invitation to physical activity study, acceptance
Breastfed as a baby
Ever sought or received professional help for mental distress
Fraction of day affected during worst episode of depression
Ever had cervical smear test
Phenotype contribution score
1.0
0.8
0.6
0.4
0.2
0.0
PC154

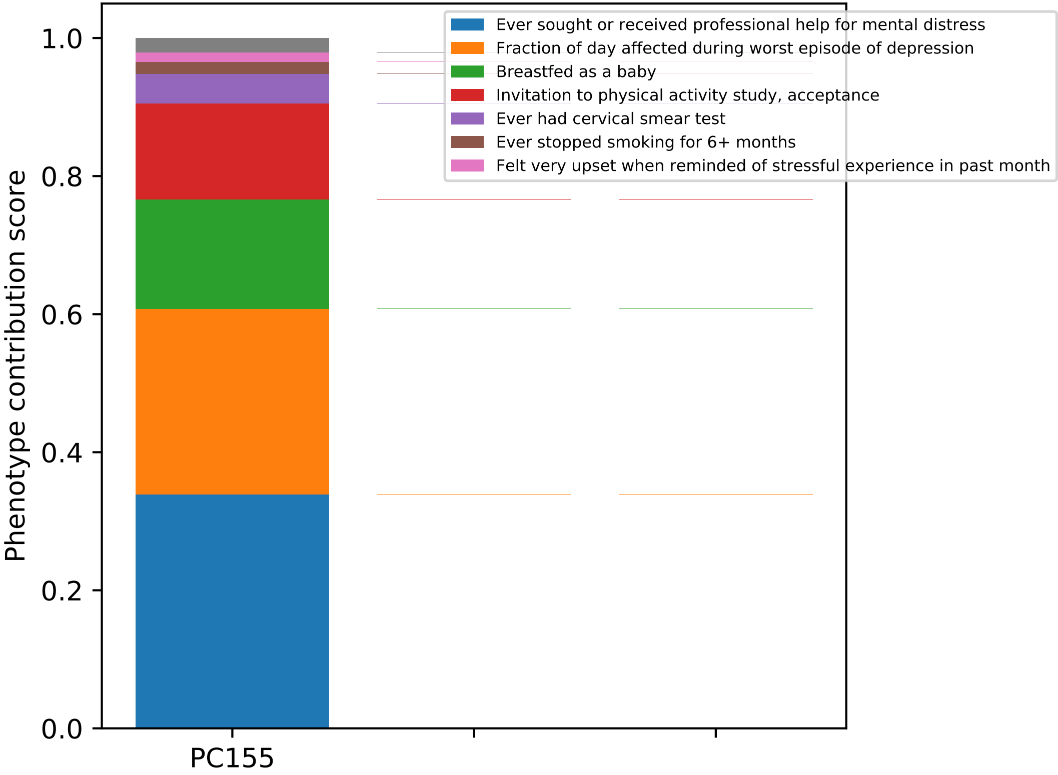

Phenotype contribution score
1.0
0.8
0.6
0.4
0.2
0.0
PC155
Ever sought or received professional help for mental distress
Fraction of day affected during worst episode of depression
Breastfed as a baby
Invitation to physical activity study, acceptance
Ever had cervical smear test
Ever stopped smoking for 6+ months
Felt very upset when reminded of stressful experience in past month

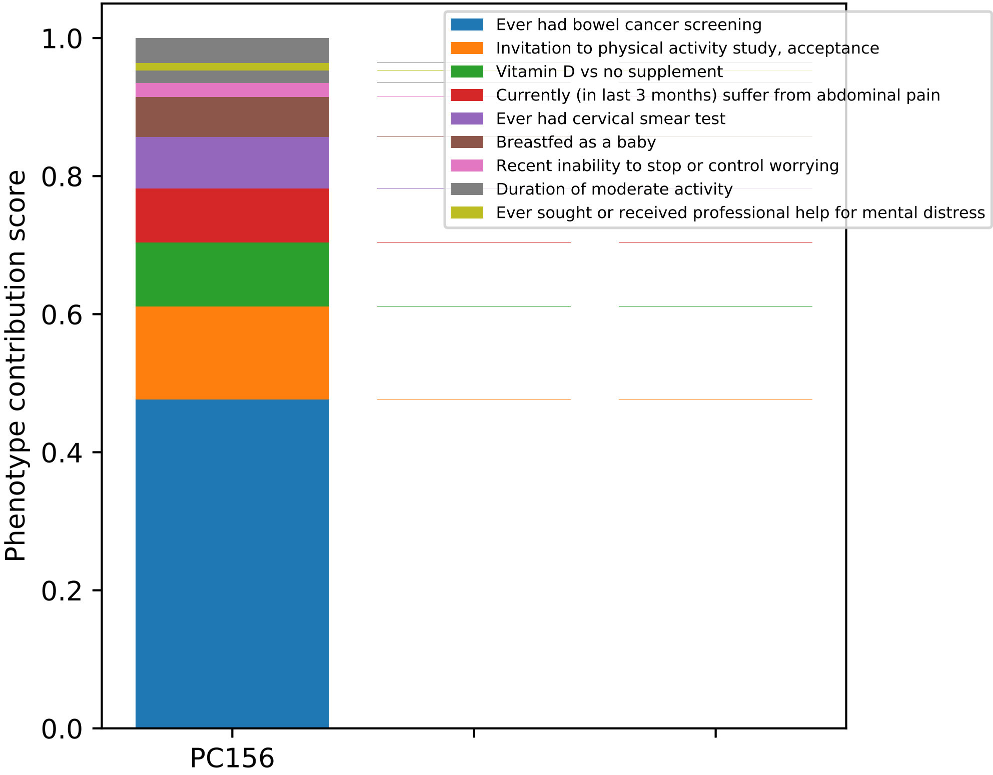

Ever had bowel cancer screening
Invitation to physical activity study, acceptance
Vitamin D vs no supplement
Currently (in last 3 months) suffer from abdominal pain
Ever had cervical smear test
Breastfed as a baby
Recent inability to stop or control worrying
Duration of moderate activity
Ever sought or received professional help for mental distress
Phenotype contribution score
1.0
0.8
0.6
0.4
0.2
0.0
PC156

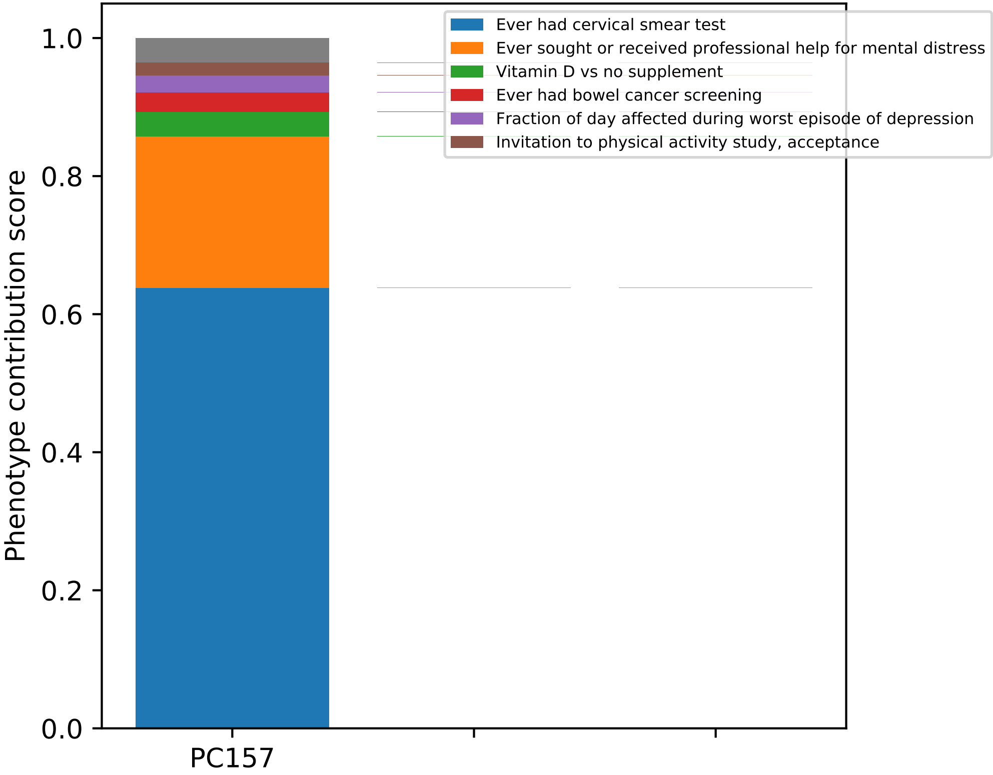
Ever had cervical smear test
Ever sought or received professional help for mental distress
Vitamin D vs no supplement
Ever had bowel cancer screening
Fraction of day affected during worst episode of depression
Invitation to physical activity study, acceptance
Phenotype contribution score
0.0
0.2
0.4
0.6
0.8
1.0
PC157

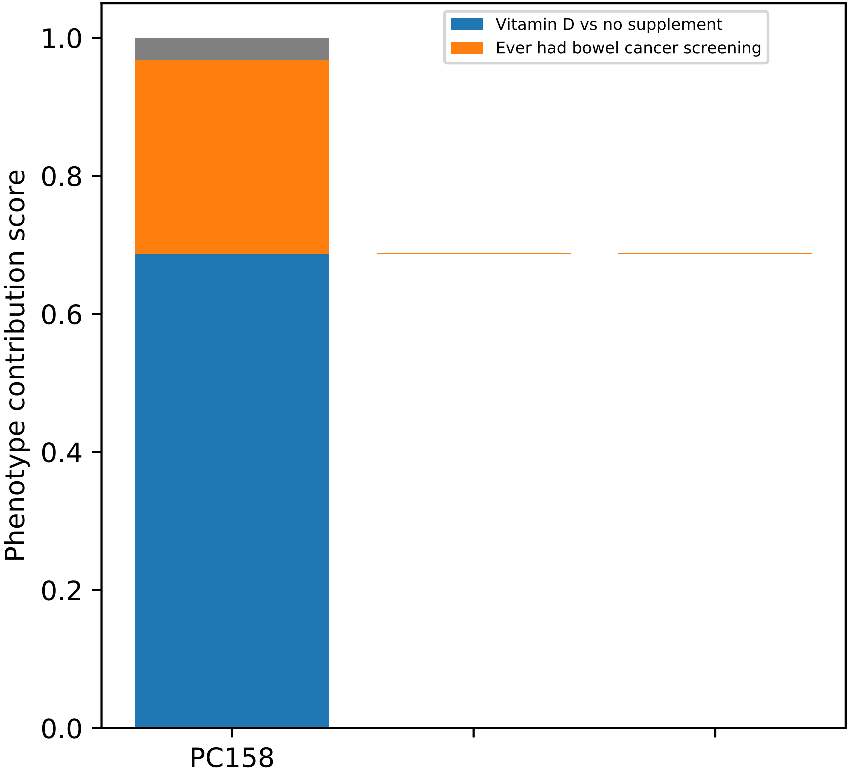

Vitamin D vs no supplement
Ever had bowel cancer screening
Phenotype contribution score
1.0
0.8
0.6
0.4
0.2
0.0
PC158

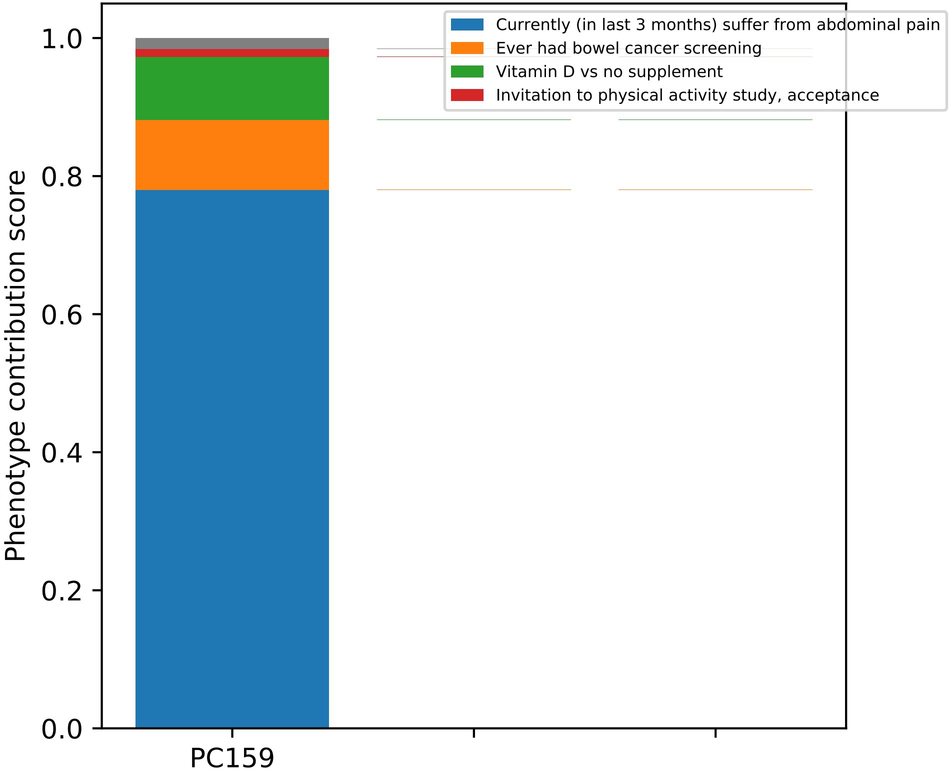
Phenotype contribution score
0.0
0.2
0.4
0.6
0.8
1.0
PC159
Currently (in last 3 months) suffer from abdominal pain
Ever had bowel cancer screening
Vitamin D vs no supplement
Invitation to physical activity study, acceptance

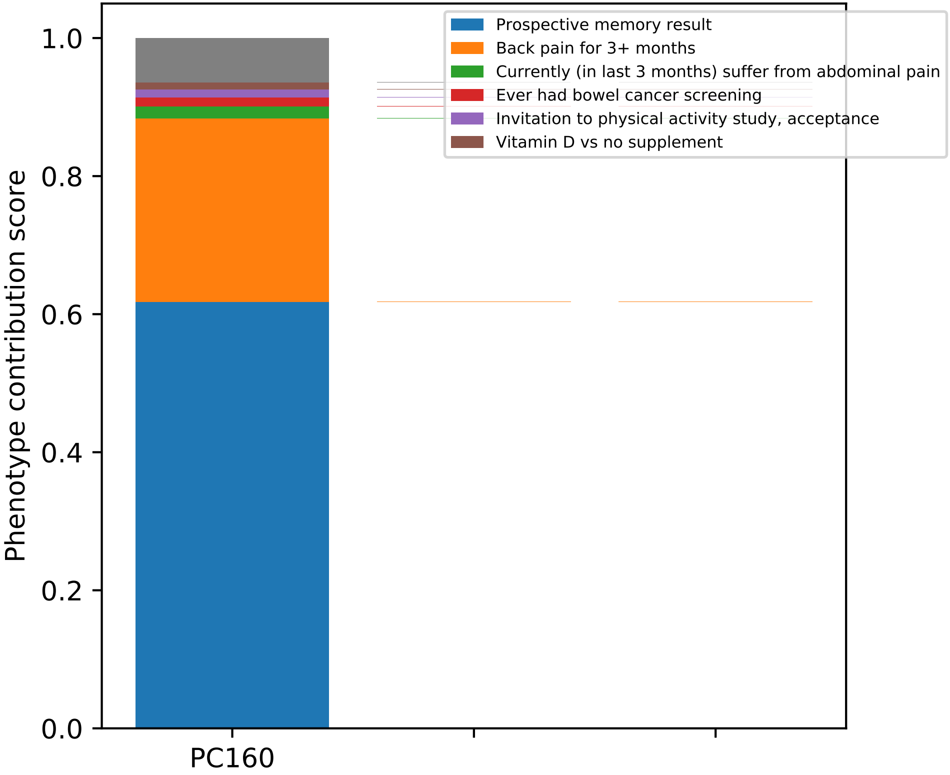
Prospective memory result
Back pain for 3+ months
Currently (in last 3 months) suffer from abdominal pain
Ever had bowel cancer screening
Invitation to physical activity study, acceptance
Vitamin D vs no supplement
Phenotype contribution score
0.0
0.2
0.4
0.6
0.8
1.0
PC160

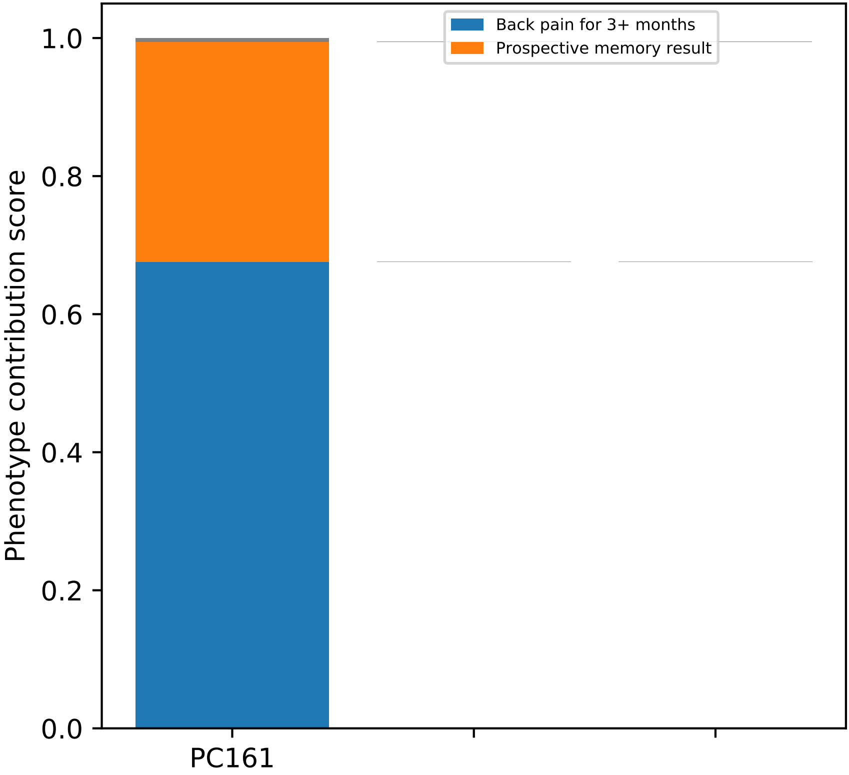

Back pain for 3+ months
Prospective memory result
Phenotype contribution score
1.0
0.8
0.6
0.4
0.2
0.0
PC161

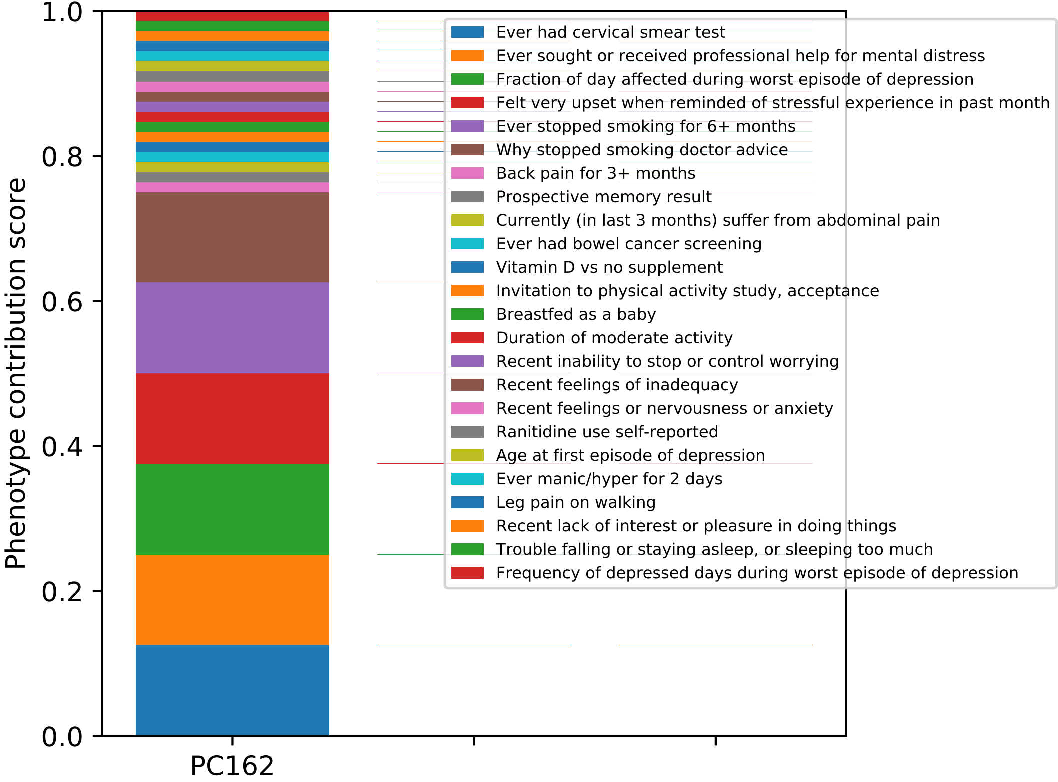
Phenotype contribution score
1.0
0.8
0.6
0.4
0.2
0.0
PC162
Ever had cervical smear test
Ever sought or received professional help for mental distress
Fraction of day affected during worst episode of depression
Felt very upset when reminded of stressful experience in past month
Ever stopped smoking for 6+ months
Why stopped smoking doctor advice
Back pain for 3+ months
Prospective memory result
Currently (in last 3 months) suffer from abdominal pain
Ever had bowel cancer screening
Vitamin D vs no supplement
Invitation to physical activity study, acceptance
Breastfed as a baby
Duration of moderate activity
Recent inability to stop or control worrying
Recent feelings of inadequacy
Recent feelings or nervousness or anxiety
Ranitidine use self-reported
Age at first episode of depression
Ever manic/hyper for 2 days
Leg pain on walking
Recent lack of interest or pleasure in doing things
Trouble falling or staying asleep, or sleeping too much
Frequency of depressed days during worst episode of depression

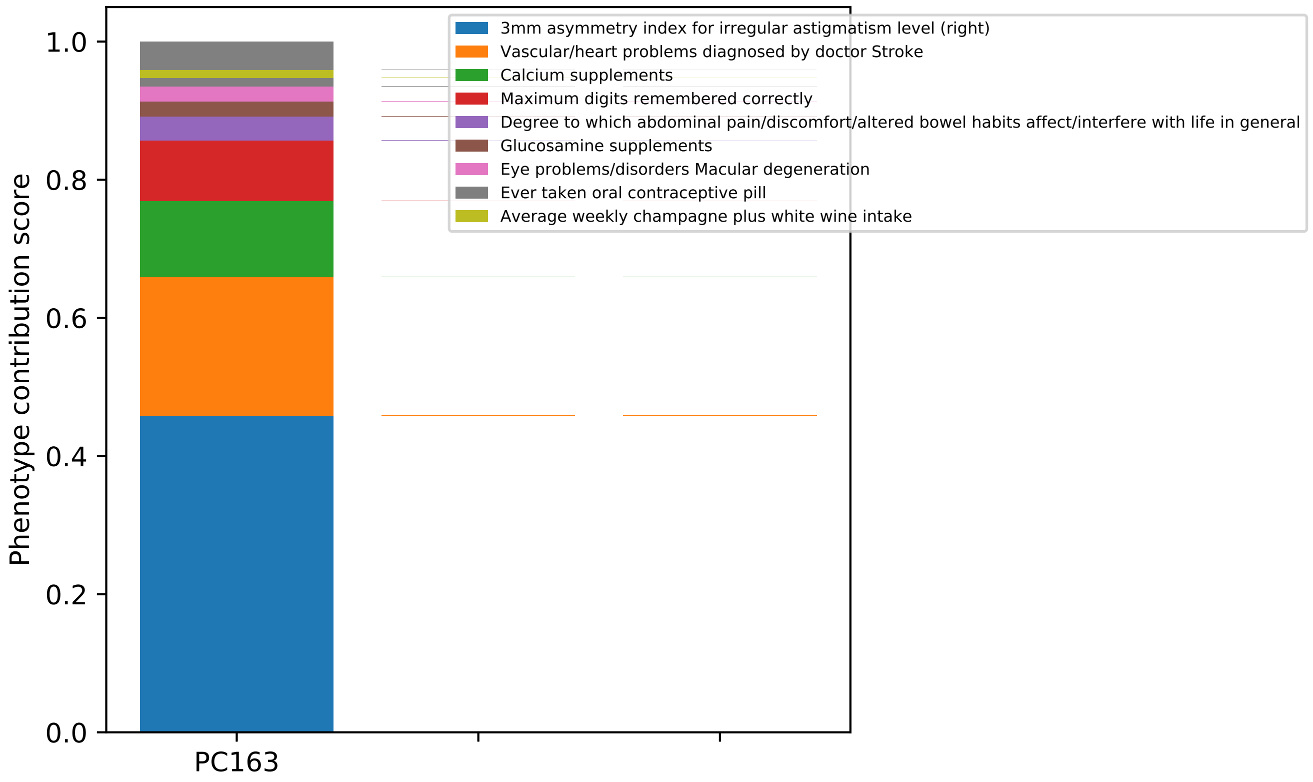

Phenotype contribution score
1.0
0.8
0.6
0.4
0.2
0.0
PC163
3mm asymmetry index for irregular astigmatism level (right)
Vascular/heart problems diagnosed by doctor Stroke
Calcium supplements
Maximum digits remembered correctly
Degree to which abdominal pain/discomfort/altered bowel habits affect/interfere with life in general
Glucosamine supplements
Eye problems/disorders Macular degeneration
Ever taken oral contraceptive pill
Average weekly champagne plus white wine intake

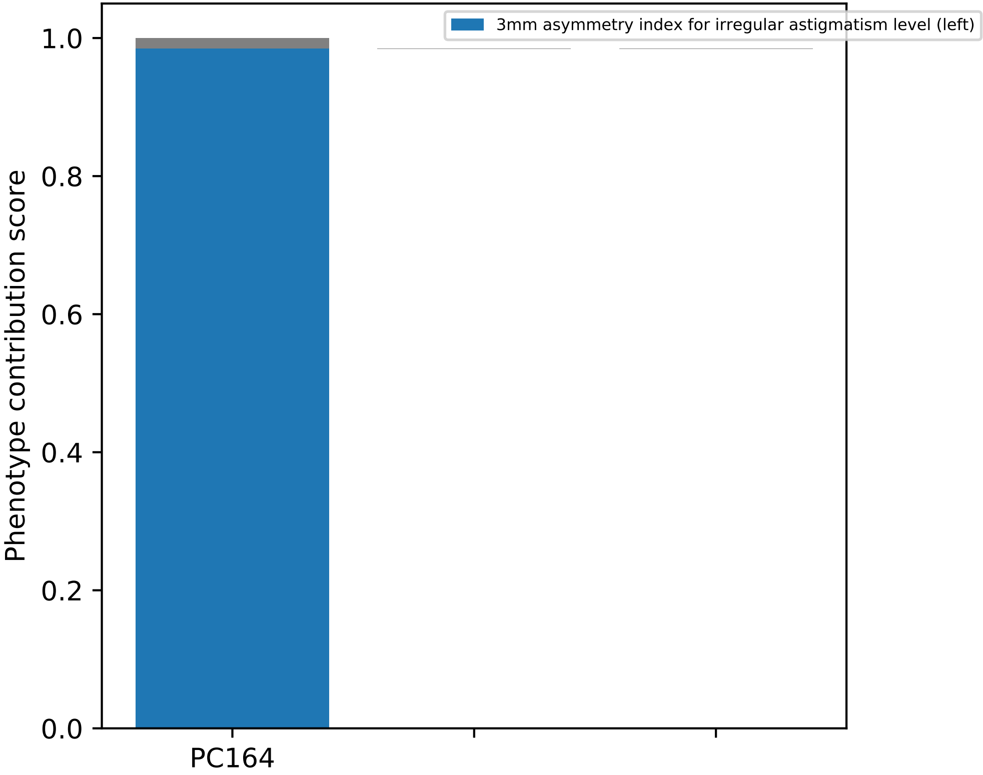
3mm asymmetry index for irregular astigmatism level (left)
1.0
0.8
0.6
0.4
0.2
0.0
Phenotype contribution score
PC164

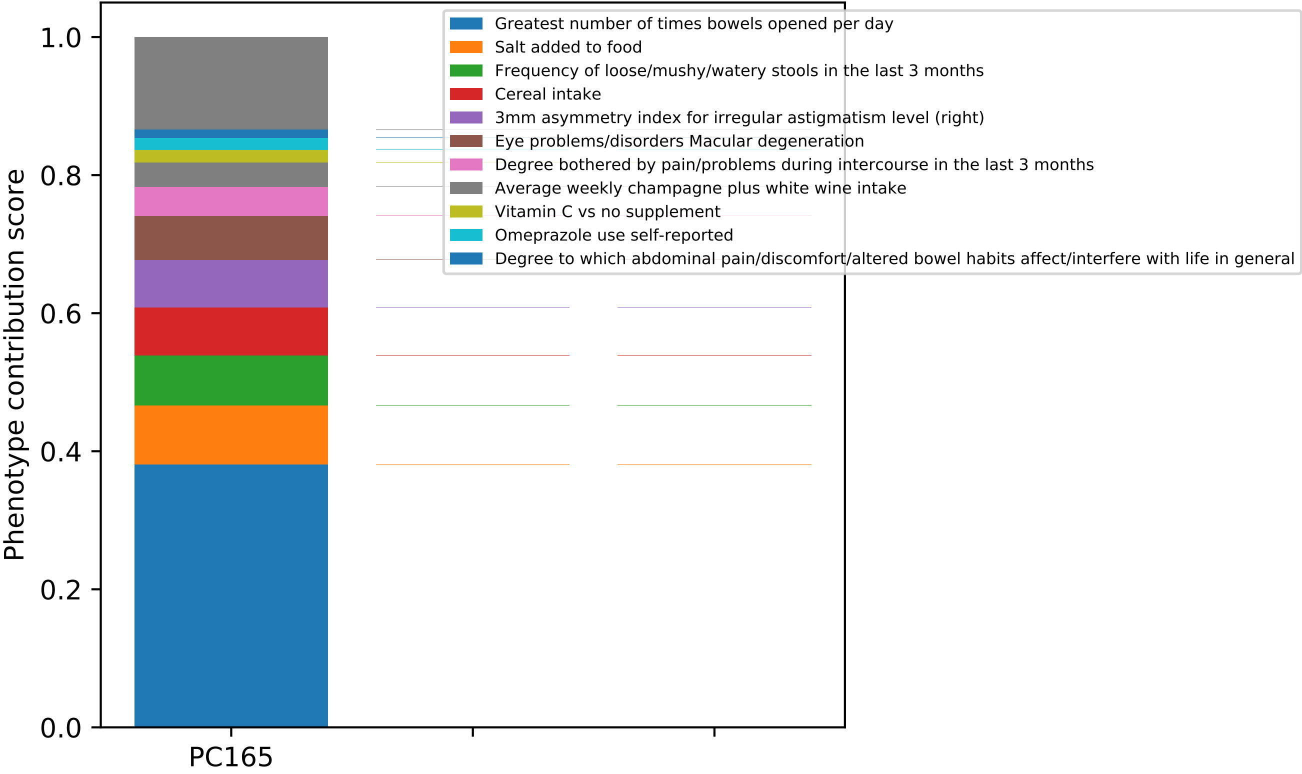

Greatest number of times bowels opened per day
Salt added to food
Frequency of loose/mushy/watery stools in the last 3 months
Cereal intake
3mm asymmetry index for irregular astigmatism level (right)
Eye problems/disorders Macular degeneration
Degree bothered by pain/problems during intercourse in the last 3 months
Average weekly champagne plus white wine intake
Vitamin C vs no supplement
Omeprazole use self-reported
Degree to which abdominal pain/discomfort/altered bowel habits affect/interfere with life in general
Phenotype contribution score
1.0
0.8
0.6
0.4
0.2
0.0
PC165

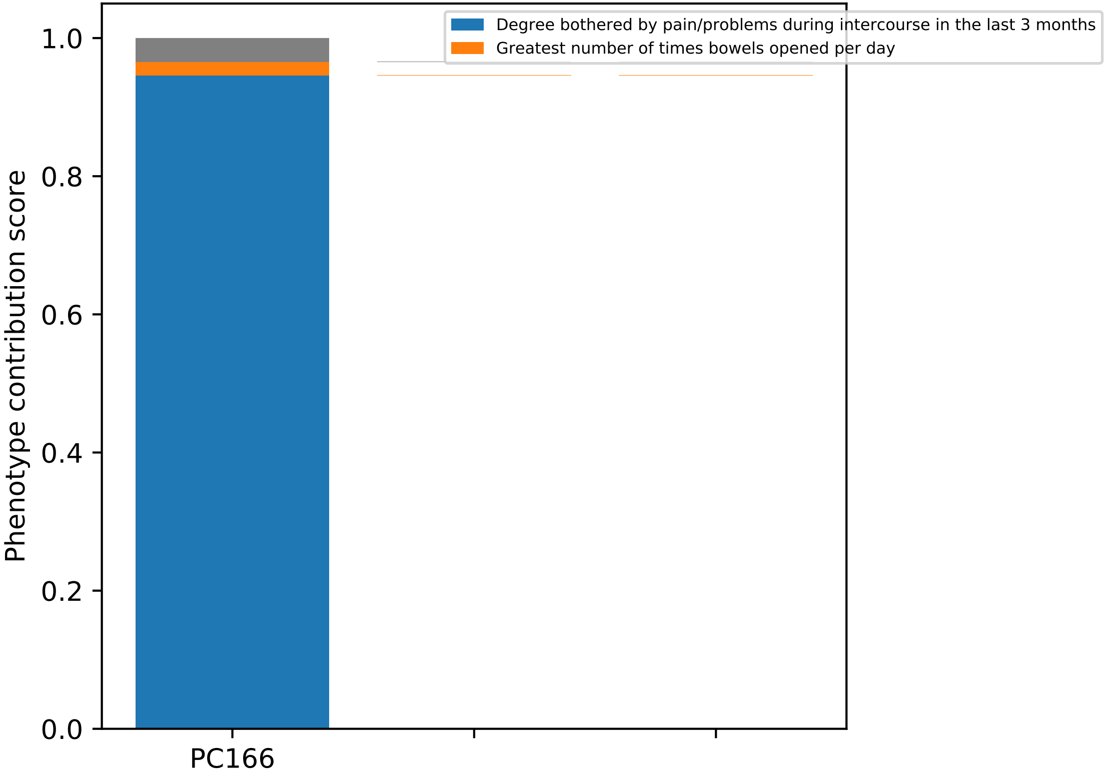

Degree bothered by pain/problems during intercourse in the last 3 months
Greatest number of times bowels opened per day
Phenotype contribution score
1.0
0.8
0.6
0.4
0.2
0.0
PC166

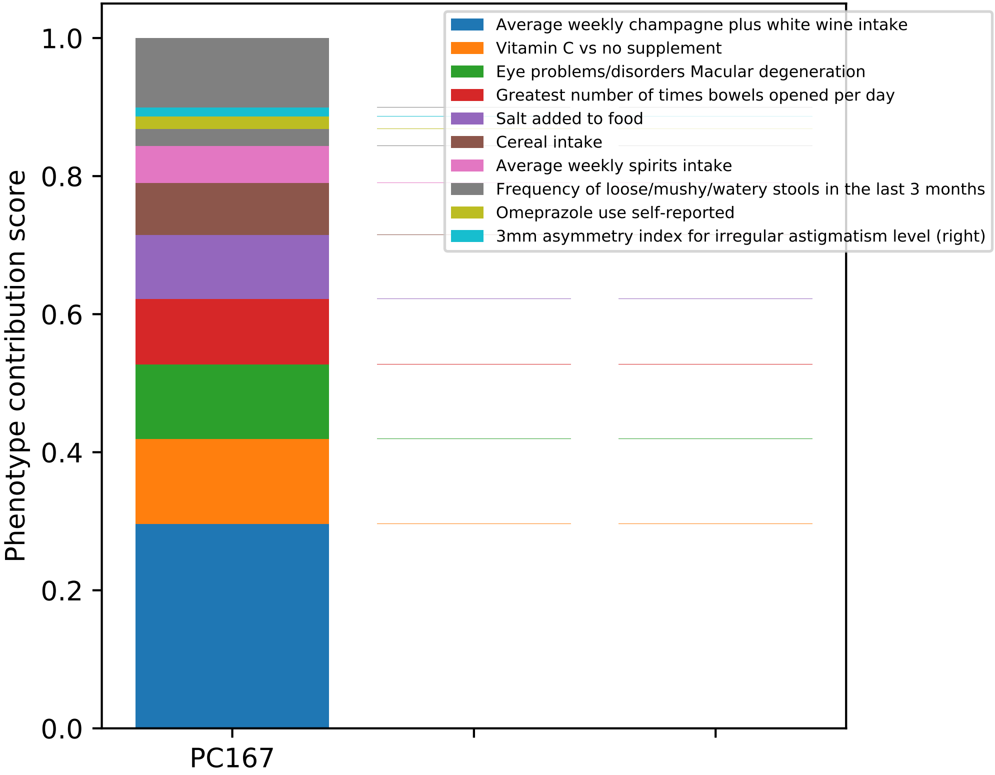

Average weekly champagne plus white wine intake
Vitamin C vs no supplement
Eye problems/disorders Macular degeneration
Greatest number of times bowels opened per day
Salt added to food
Cereal intake
Average weekly spirits intake
Frequency of loose/mushy/watery stools in the last 3 months
Omeprazole use self-reported
3mm asymmetry index for irregular astigmatism level (right)
Phenotype contribution score
1.0
0.8
0.6
0.4
0.2
0.0
PC167

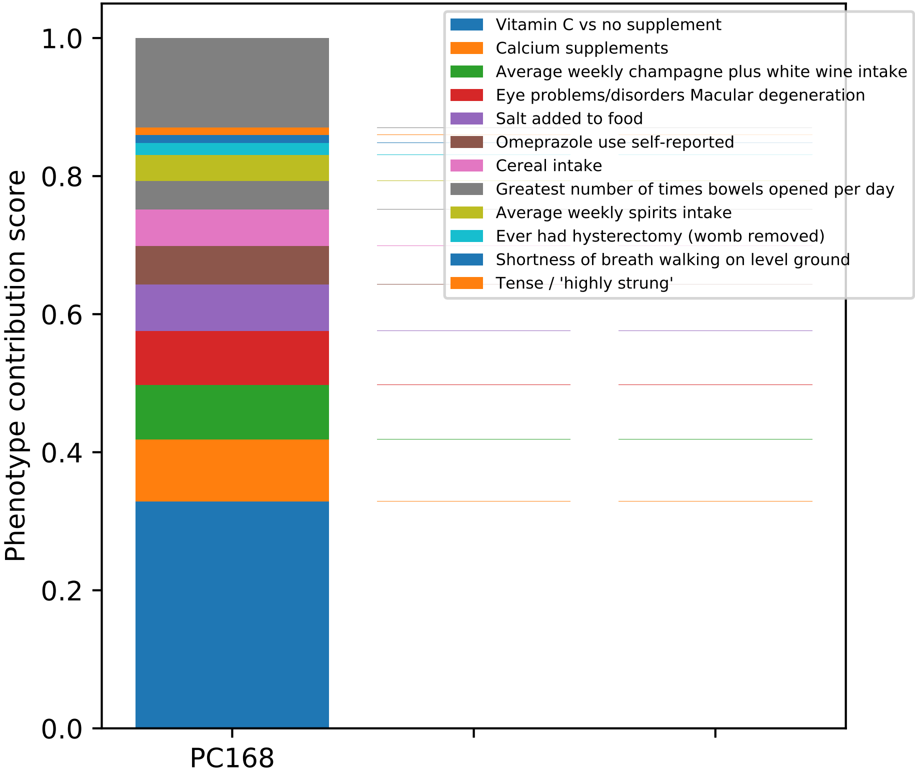

Vitamin C vs no supplement
Calcium supplements
Average weekly champagne plus white wine intake
Eye problems/disorders Macular degeneration
Salt added to food
Omeprazole use self-reported
Cereal intake
Greatest number of times bowels opened per day
Average weekly spirits intake
Ever had hysterectomy (womb removed)
Shortness of breath walking on level ground
Tense / 'highly strung'
Phenotype contribution score
1.0
0.8
0.6
0.4
0.2
0.0
PC168

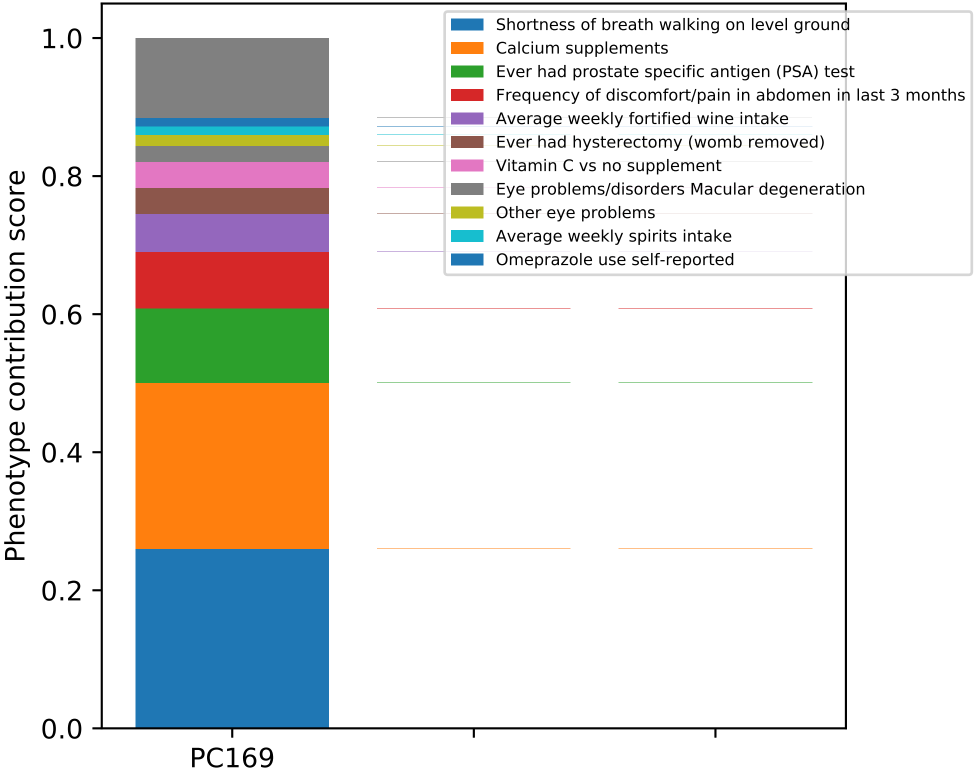

Phenotype contribution score
0.0
0.2
0.4
0.6
0.8
1.0
PC169
Shortness of breath walking on level ground
Calcium supplements
Ever had prostate specific antigen (PSA) test
Frequency of discomfort/pain in abdomen in last 3 months
Average weekly fortified wine intake
Ever had hysterectomy (womb removed)
Vitamin C vs no supplement
Eye problems/disorders Macular degeneration
Other eye problems
Average weekly spirits intake
Omeprazole use self-reported

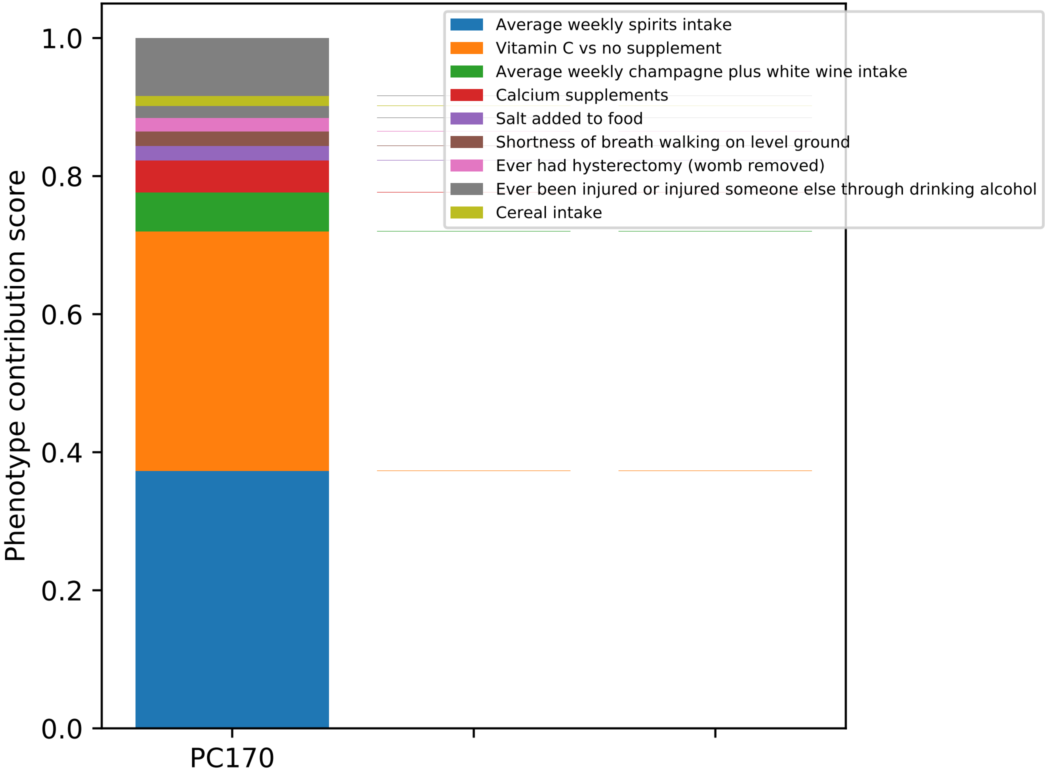

Average weekly spirits intake
Vitamin C vs no supplement
Average weekly champagne plus white wine intake
Calcium supplements
Salt added to food
Shortness of breath walking on level ground
Ever had hysterectomy (womb removed)
Ever been injured or injured someone else through drinking alcohol
Cereal intake
Phenotype contribution score
0.0
0.2
0.4
0.6
0.8
1.0
PC170

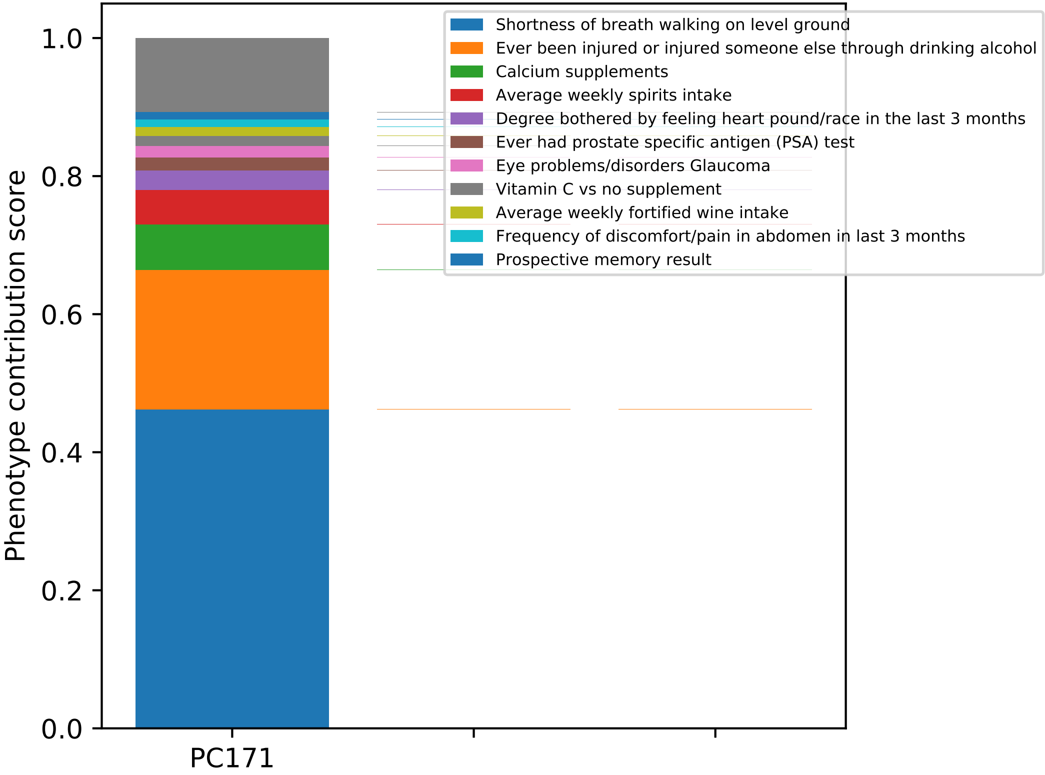

Phenotype contribution score
1.0
0.8
0.6
0.4
0.2
0.0
PC171
Shortness of breath walking on level ground
Ever been injured or injured someone else through drinking alcohol
Calcium supplements
Average weekly spirits intake
Degree bothered by feeling heart pound/race in the last 3 months
Ever had prostate specific antigen (PSA) test
Eye problems/disorders Glaucoma
Vitamin C vs no supplement
Average weekly fortified wine intake
Frequency of discomfort/pain in abdomen in last 3 months
Prospective memory result

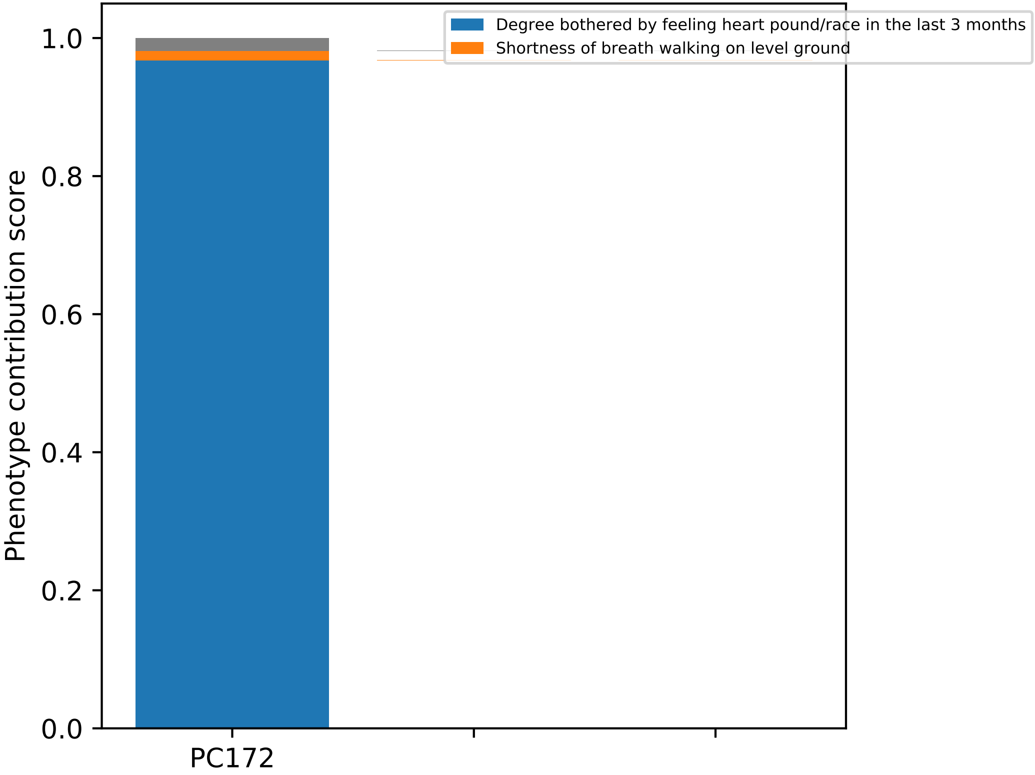

Degree bothered by feeling heart pound/race in the last 3 months
Shortness of breath walking on level ground
Phenotype contribution score
1.0
0.8
0.6
0.4
0.2
0.0
PC172

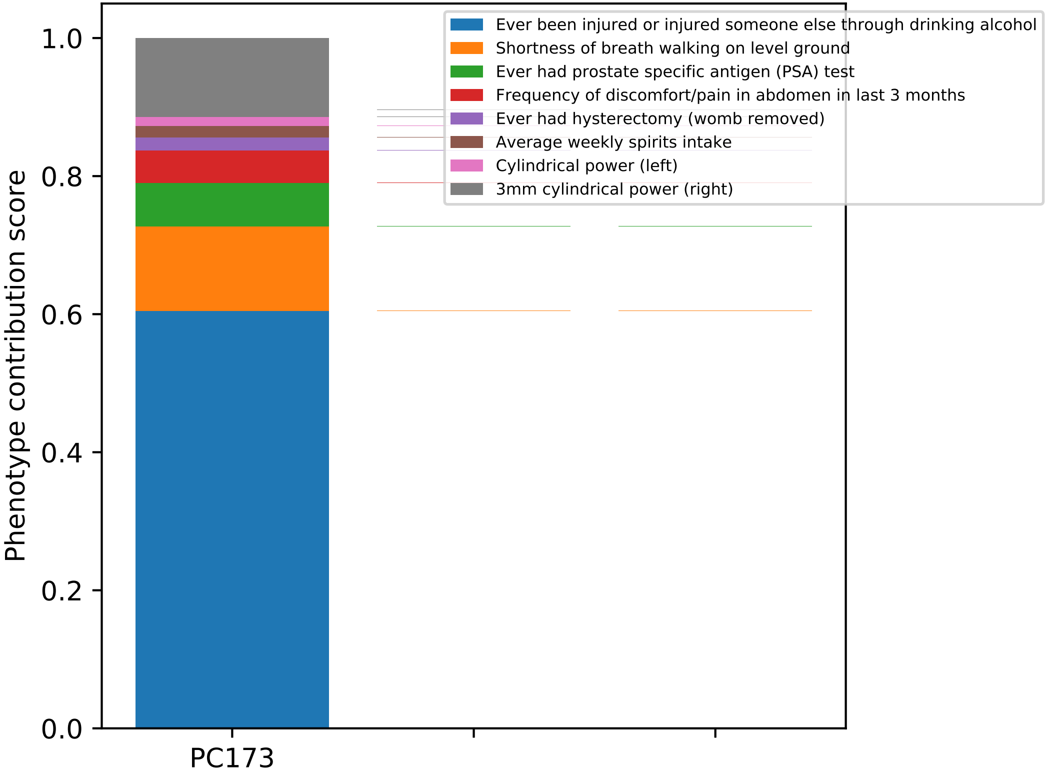

Ever been injured or injured someone else through drinking alcohol
Shortness of breath walking on level ground
Ever had prostate specific antigen (PSA) test
Frequency of discomfort/pain in abdomen in last 3 months
Ever had hysterectomy (womb removed)
Average weekly spirits intake
Cylindrical power (left)
3mm cylindrical power (right)
Phenotype contribution score
1.0
0.8
0.6
0.4
0.2
0.0
PC173

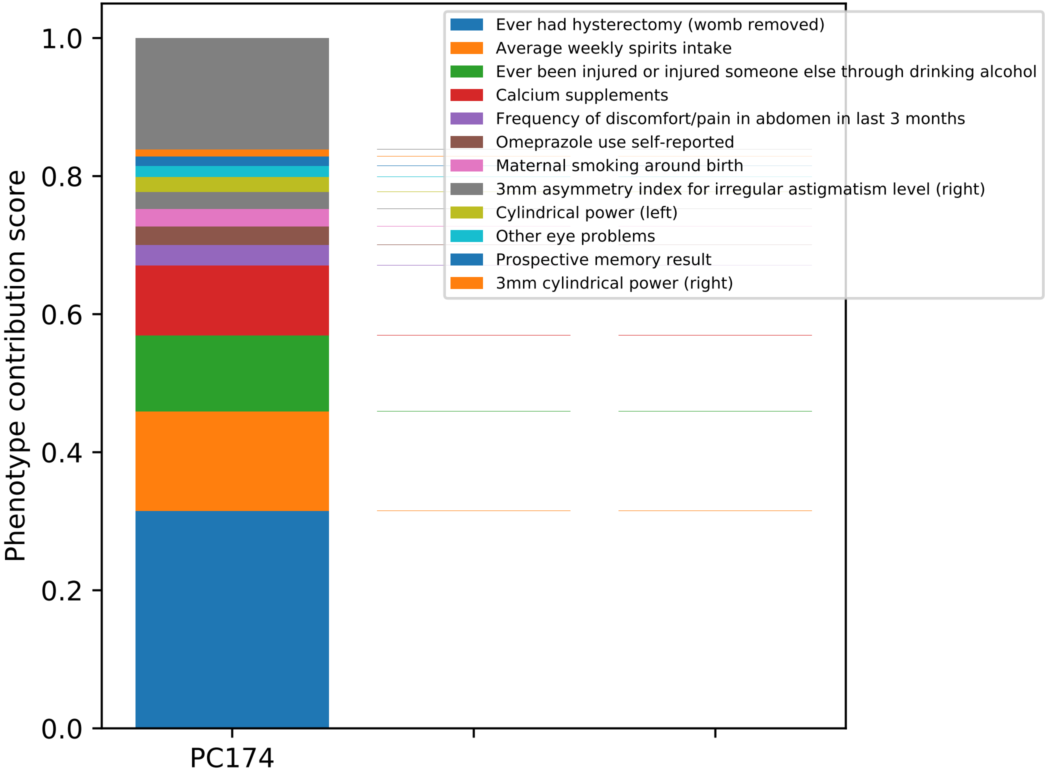
Phenotype contribution score
1.0
0.8
0.6
0.4
0.2
0.0
PC174
Ever had hysterectomy (womb removed)
Average weekly spirits intake
Ever been injured or injured someone else through drinking alcohol
Calcium supplements
Frequency of discomfort/pain in abdomen in last 3 months
Omeprazole use self-reported
Maternal smoking around birth
3mm asymmetry index for irregular astigmatism level (right)
Cylindrical power (left)
Other eye problems
Prospective memory result
3mm cylindrical power (right)

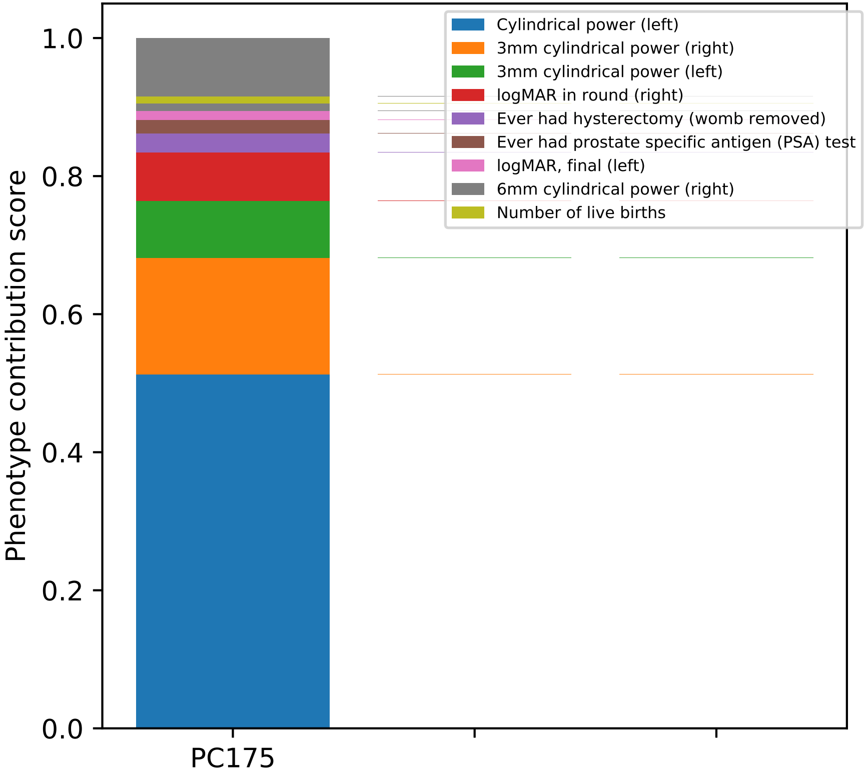

Cylindrical power (left)
3mm cylindrical power (right)
3mm cylindrical power (left)
logMAR in round (right)
Ever had hysterectomy (womb removed)
Ever had prostate specific antigen (PSA) test
logMAR, final (left)
6mm cylindrical power (right)
Number of live births
Phenotype contribution score
0.0
0.2
0.4
0.6
0.8
1.0
PC175

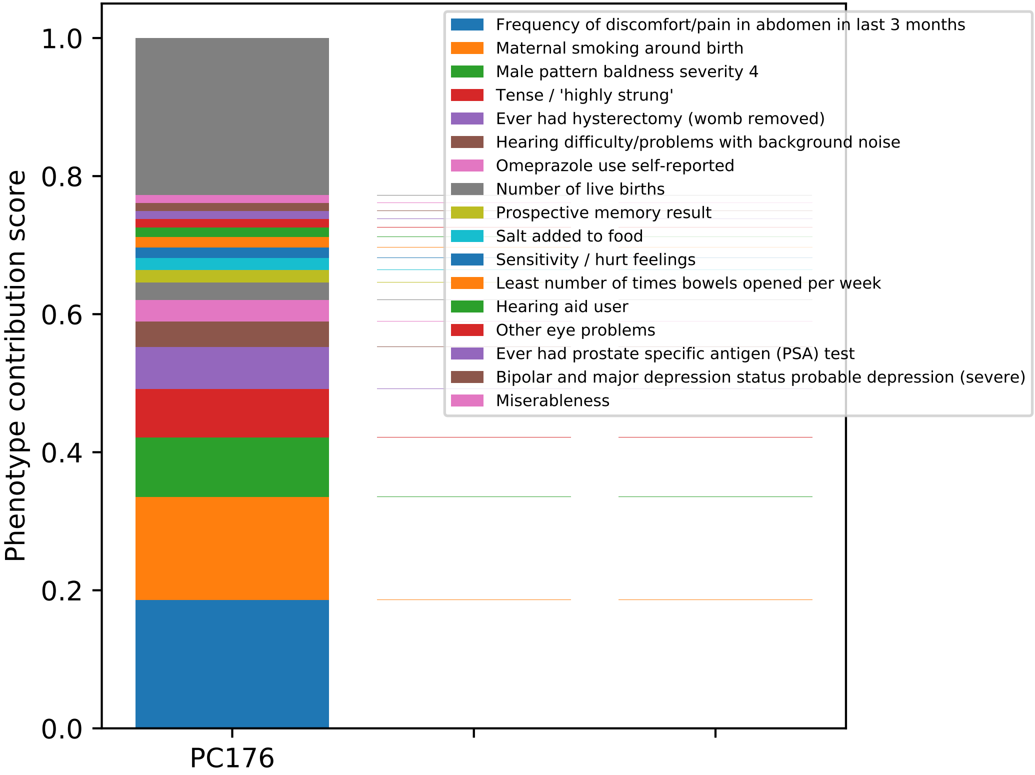

Phenotype contribution score
1.0
0.8
0.6
0.4
0.2
0.0
PC176
Frequency of discomfort/pain in abdomen in last 3 months
Maternal smoking around birth
Male pattern baldness severity 4
Tense / 'highly strung'
Ever had hysterectomy (womb removed)
Hearing difficulty/problems with background noise
Omeprazole use self-reported
Number of live births
Prospective memory result
Salt added to food
Sensitivity / hurt feelings
Least number of times bowels opened per week
Hearing aid user
Other eye problems
Ever had prostate specific antigen (PSA) test
Bipolar and major depression status probable depression (severe)
Miserableness

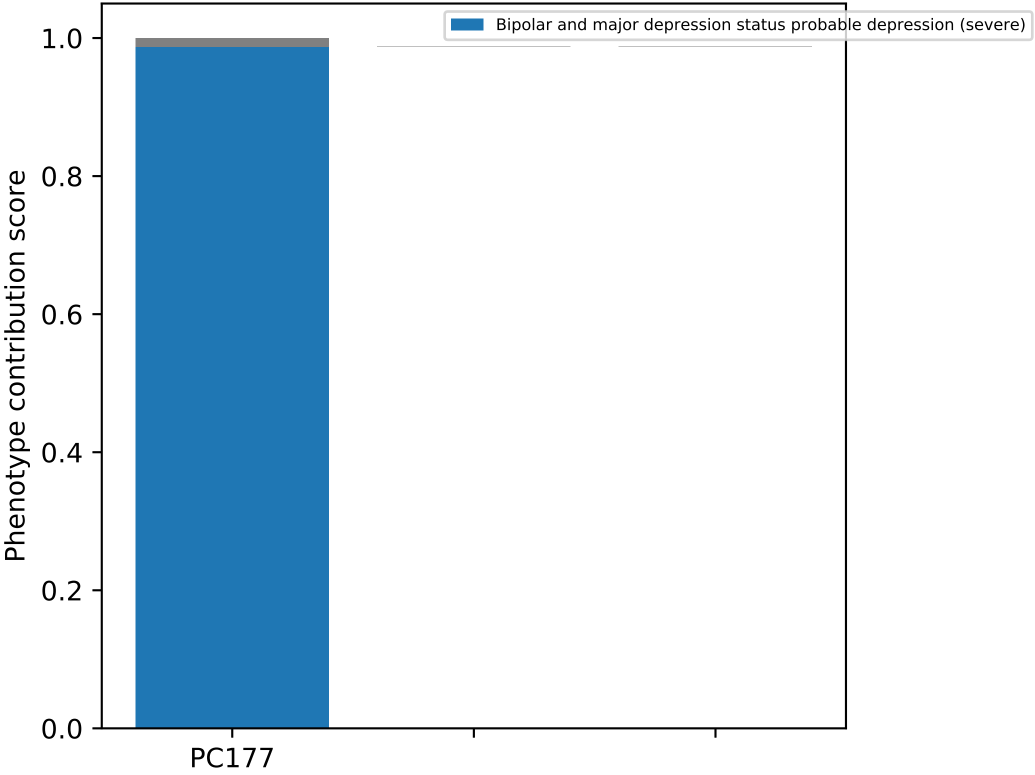
Bipolar and major depression status probable depression (severe)
Phenotype contribution score
1.0
0.8
0.6
0.4
0.2
0.0
PC177

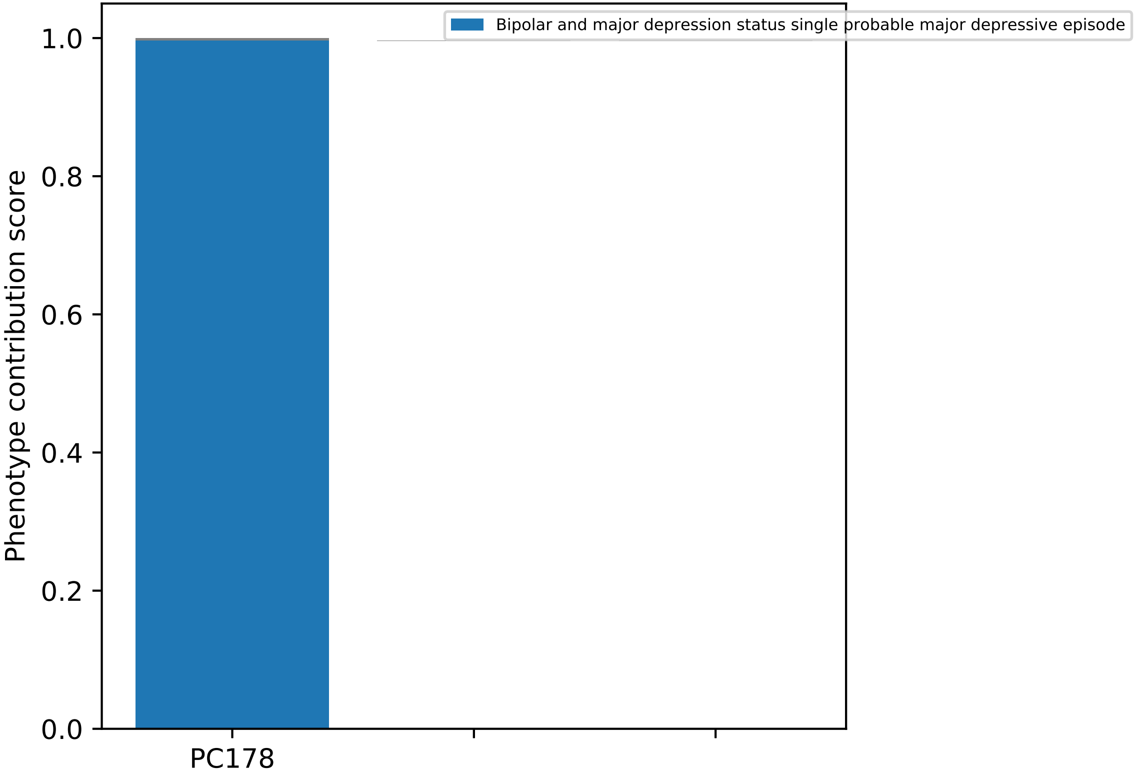

Bipolar and major depression status single probable major depressive episode
Phenotype contribution score
1.0
0.8
0.6
0.4
0.2
0.0
PC178

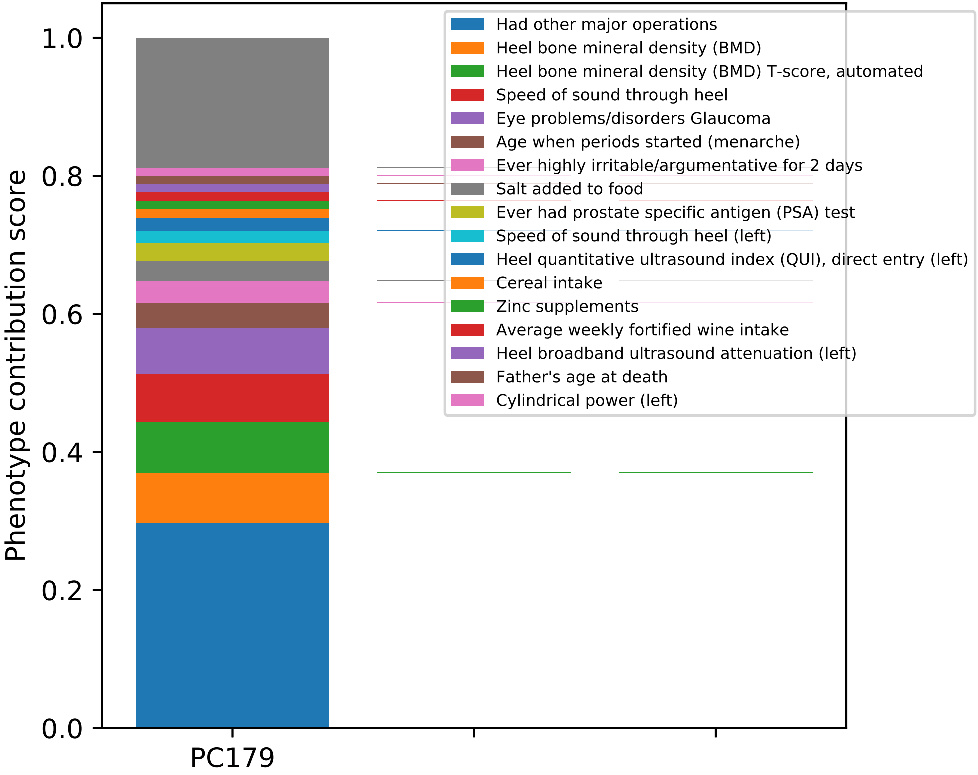

Phenotype contribution score
1.0
0.8
0.6
0.4
0.2
0.0
PC179
Had other major operations
Heel bone mineral density (BMD)
Heel bone mineral density (BMD) T-score, automated
Speed of sound through heel
Eye problems/disorders Glaucoma
Age when periods started (menarche)
Ever highly irritable/argumentative for 2 days
Salt added to food
Ever had prostate specific antigen (PSA) test
Speed of sound through heel (left)
Heel quantitative ultrasound index (QUI), direct entry (left)
Cereal intake
Zinc supplements
Average weekly fortified wine intake
Heel broadband ultrasound attenuation (left)
Father's age at death
Cylindrical power (left)

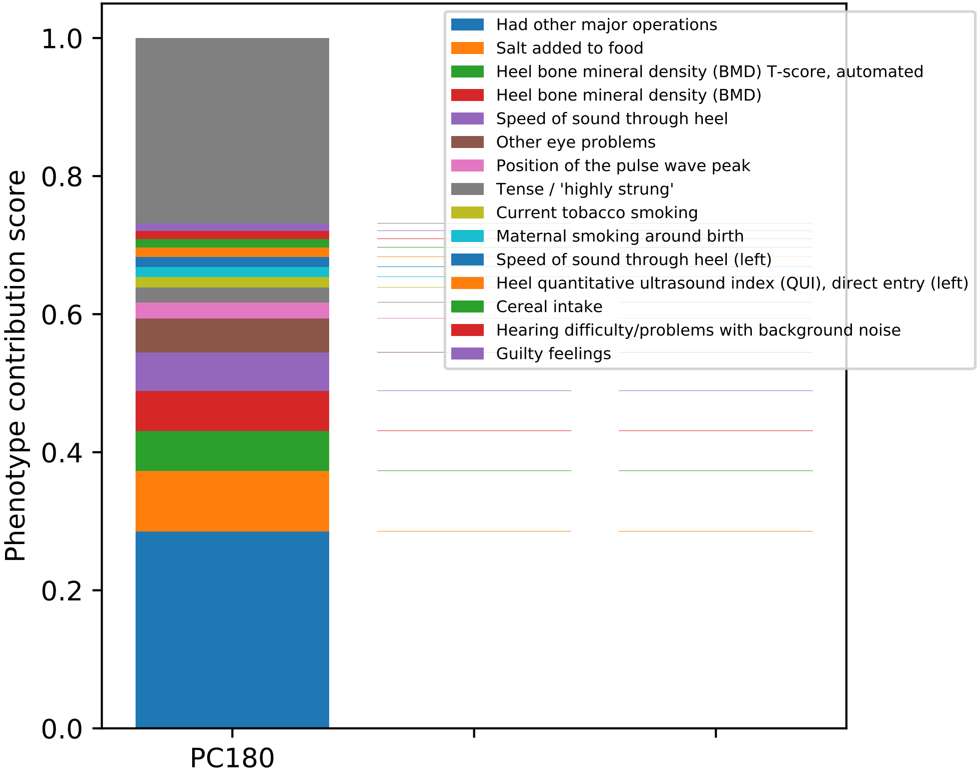
Had other major operations
Salt added to food
Heel bone mineral density (BMD) T-score, automated
Heel bone mineral density (BMD)
Speed of sound through heel
Other eye problems
Position of the pulse wave peak
Tense / 'highly strung'
Current tobacco smoking
Maternal smoking around birth
Speed of sound through heel (left)
Heel quantitative ultrasound index (QUI), direct entry (left)
Cereal intake
Hearing difficulty/problems with background noise
Guilty feelings
Phenotype contribution score
1.0
0.8
0.6
0.4
0.2
0.0
PC180

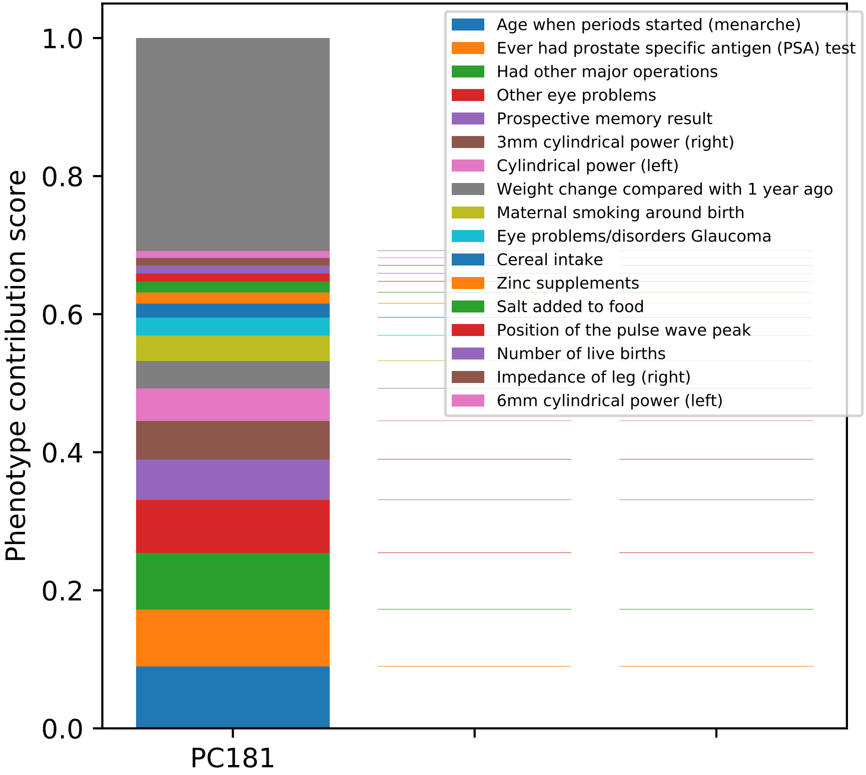
Age when periods started (menarche)
Ever had prostate specific antigen (PSA) test
Had other major operations
Other eye problems
Prospective memory result
3mm cylindrical power (right)
Cylindrical power (left)
Weight change compared with 1 year ago
Maternal smoking around birth
Eye problems/disorders Glaucoma
Cereal intake
Zinc supplements
Salt added to food
Position of the pulse wave peak
Number of live births
Impedance of leg (right)
6mm cylindrical power (left)
Phenotype contribution score
1.0
0.8
0.6
0.4
0.2
0.0
PC181

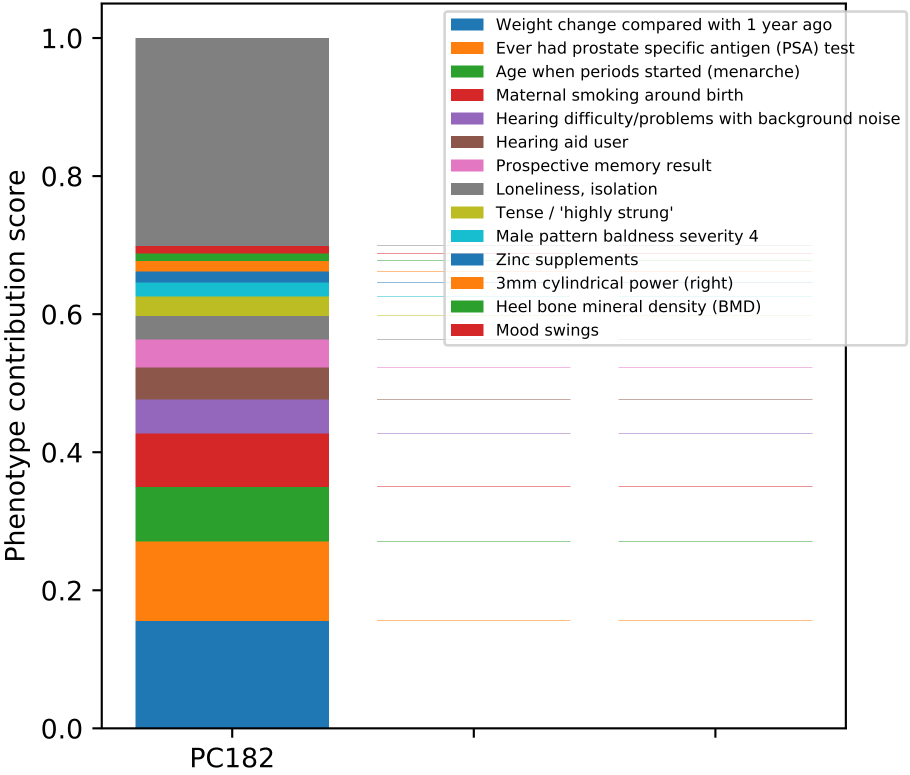

Phenotype contribution score
0.0
0.2
0.4
0.6
0.8
1.0
PC182
Weight change compared with 1 year ago
Ever had prostate specific antigen (PSA) test
Age when periods started (menarche)
Maternal smoking around birth
Hearing difficulty/problems with background noise
Hearing aid user
Prospective memory result
Loneliness, isolation
Tense / 'highly strung'
Male pattern baldness severity 4
Zinc supplements
3mm cylindrical power (right)
Heel bone mineral density (BMD)
Mood swings

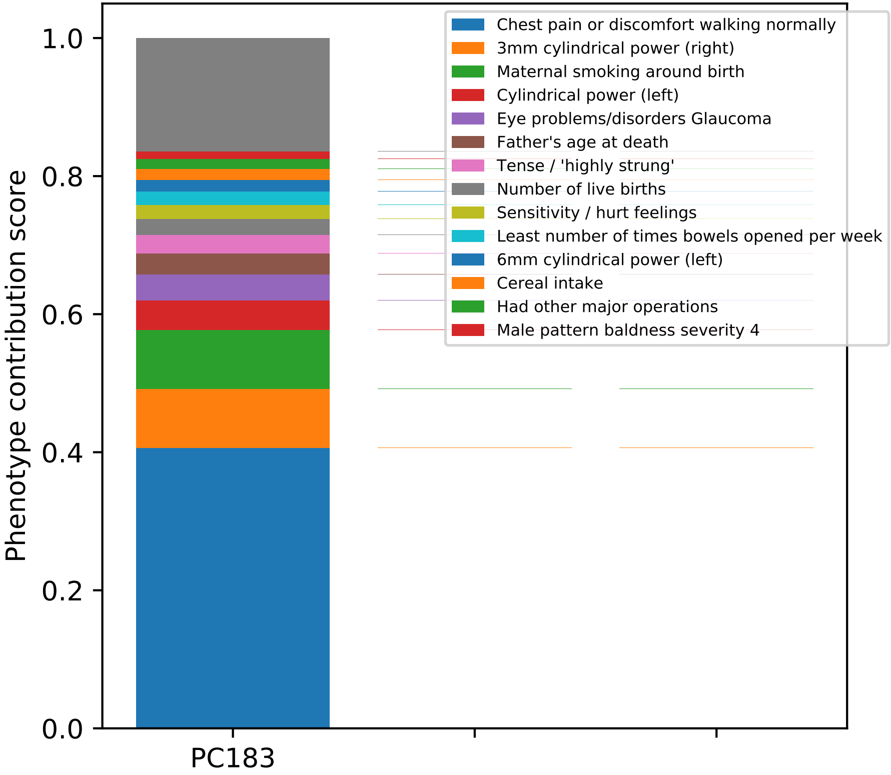

Phenotype contribution score
1.0
0.8
0.6
0.4
0.2
0.0
PC183
Chest pain or discomfort walking normally
3mm cylindrical power (right)
Maternal smoking around birth
Cylindrical power (left)
Eye problems/disorders Glaucoma
Father's age at death
Tense / 'highly strung'
Number of live births
Sensitivity / hurt feelings
Least number of times bowels opened per week
6mm cylindrical power (left)
Cereal intake
Had other major operations
Male pattern baldness severity 4

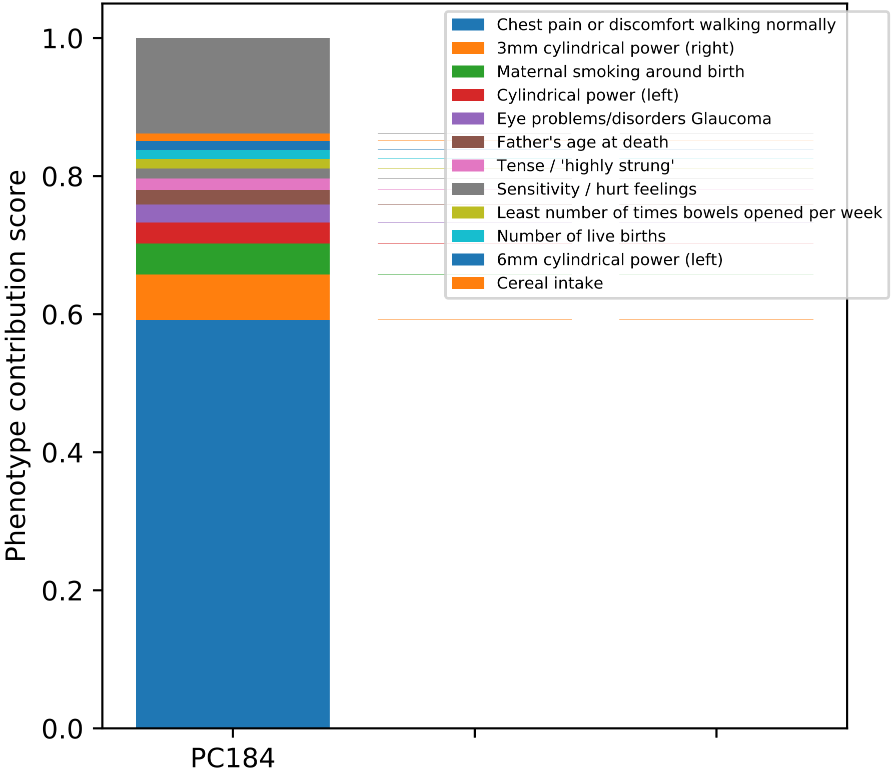

Chest pain or discomfort walking normally
3mm cylindrical power (right)
Maternal smoking around birth
Cylindrical power (left)
Eye problems/disorders Glaucoma
Father's age at death
Tense / 'highly strung'
Sensitivity / hurt feelings
Least number of times bowels opened per week
Number of live births
6mm cylindrical power (left)
Cereal intake
Phenotype contribution score
1.0
0.8
0.6
0.4
0.2
0.0
PC184

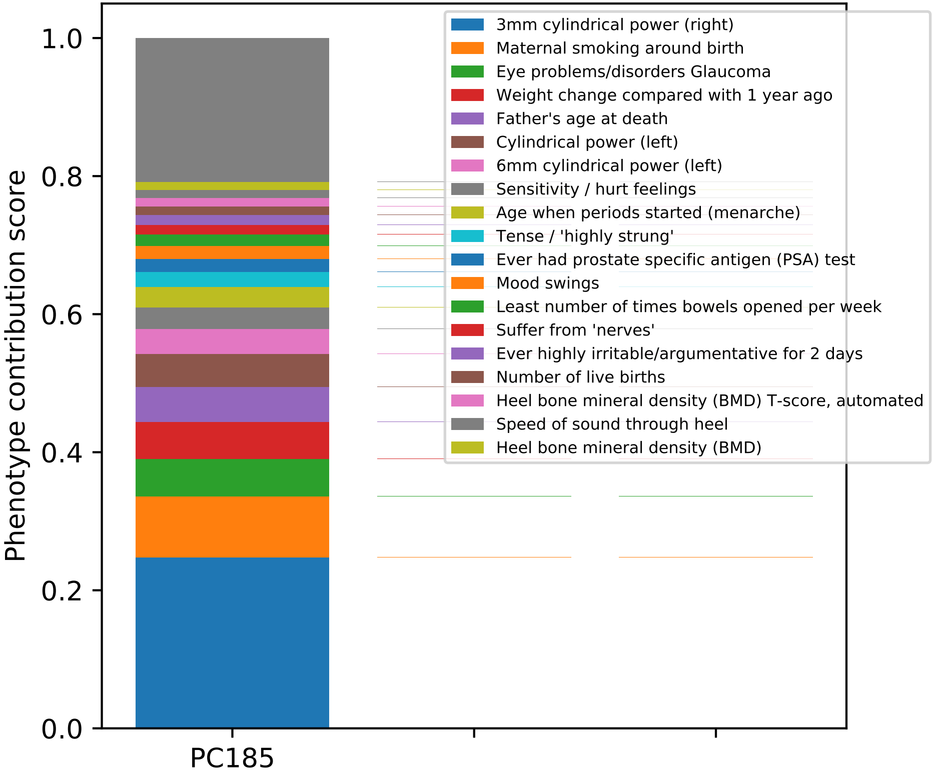

Phenotype contribution score
0.0
0.2
0.4
0.6
0.8
1.0
PC185
3mm cylindrical power (right)
Maternal smoking around birth
Eye problems/disorders Glaucoma
Weight change compared with 1 year ago
Father's age at death
Cylindrical power (left)
6mm cylindrical power (left)
Sensitivity / hurt feelings
Age when periods started (menarche)
Tense / 'highly strung'
Ever had prostate specific antigen (PSA) test
Mood swings
Least number of times bowels opened per week
Suffer from 'nerves'
Ever highly irritable/argumentative for 2 days
Number of live births
Heel bone mineral density (BMD) T-score, automated
Speed of sound through heel
Heel bone mineral density (BMD)

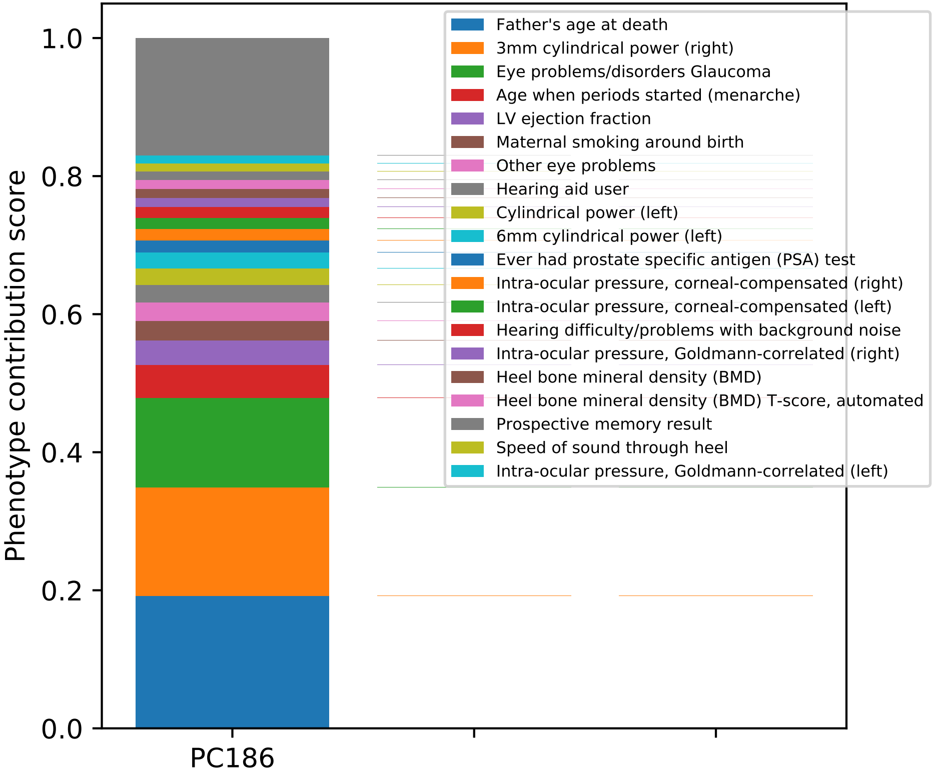

Phenotype contribution score
0.0
0.2
0.4
0.6
0.8
1.0
PC186
Father's age at death
3mm cylindrical power (right)
Eye problems/disorders Glaucoma
Age when periods started (menarche)
LV ejection fraction
Maternal smoking around birth
Other eye problems
Hearing aid user
Cylindrical power (left)
6mm cylindrical power (left)
Ever had prostate specific antigen (PSA) test
Intra-ocular pressure, corneal-compensated (right)
Intra-ocular pressure, corneal-compensated (left)
Hearing difficulty/problems with background noise
Intra-ocular pressure, Goldmann-correlated (right)
Heel bone mineral density (BMD)
Heel bone mineral density (BMD) T-score, automated
Prospective memory result
Speed of sound through heel
Intra-ocular pressure, Goldmann-correlated (left)

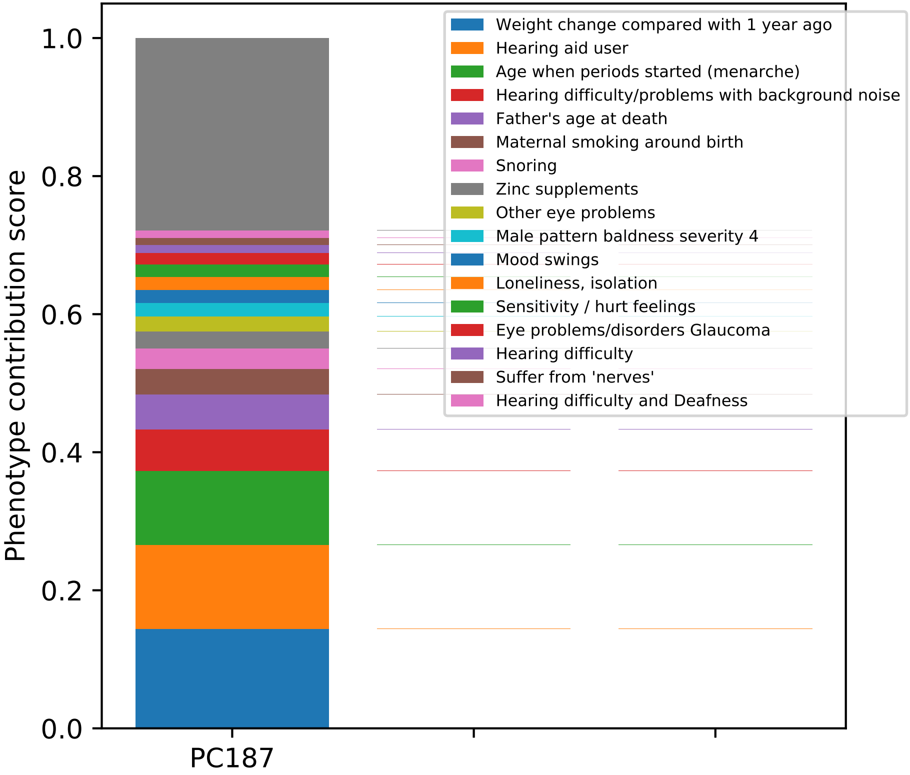

Phenotype contribution score
0.0
0.2
0.4
0.6
0.8
1.0
PC187
Weight change compared with 1 year ago
Hearing aid user
Age when periods started (menarche)
Hearing difficulty/problems with background noise
Father's age at death
Maternal smoking around birth
Snoring
Zinc supplements
Other eye problems
Male pattern baldness severity 4
Mood swings
Loneliness, isolation
Sensitivity / hurt feelings
Eye problems/disorders Glaucoma
Hearing difficulty
Suffer from 'nerves'
Hearing difficulty and Deafness

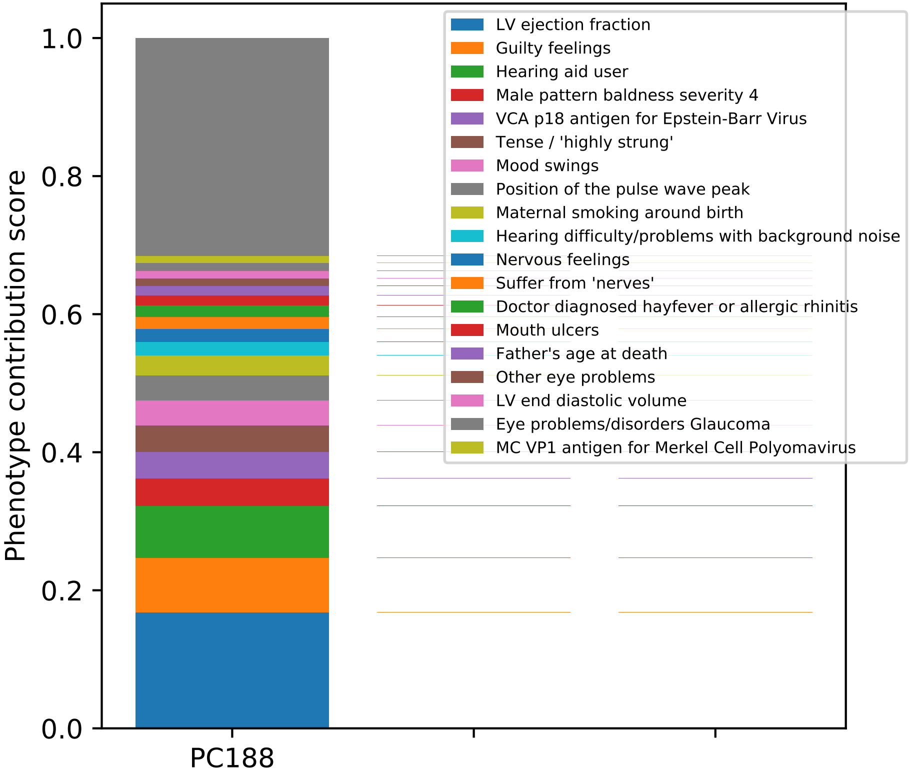

Phenotype contribution score
0.0
0.2
0.4
0.6
0.8
1.0
PC188
LV ejection fraction
Guilty feelings
Hearing aid user
Male pattern baldness severity 4
VCA p18 antigen for Epstein-Barr Virus
Tense / 'highly strung'
Mood swings
Position of the pulse wave peak
Maternal smoking around birth
Hearing difficulty/problems with background noise
Nervous feelings
Suffer from 'nerves'
Doctor diagnosed hayfever or allergic rhinitis
Mouth ulcers
Father's age at death
Other eye problems
LV end diastolic volume
Eye problems/disorders Glaucoma
MC VP1 antigen for Merkel Cell Polyomavirus

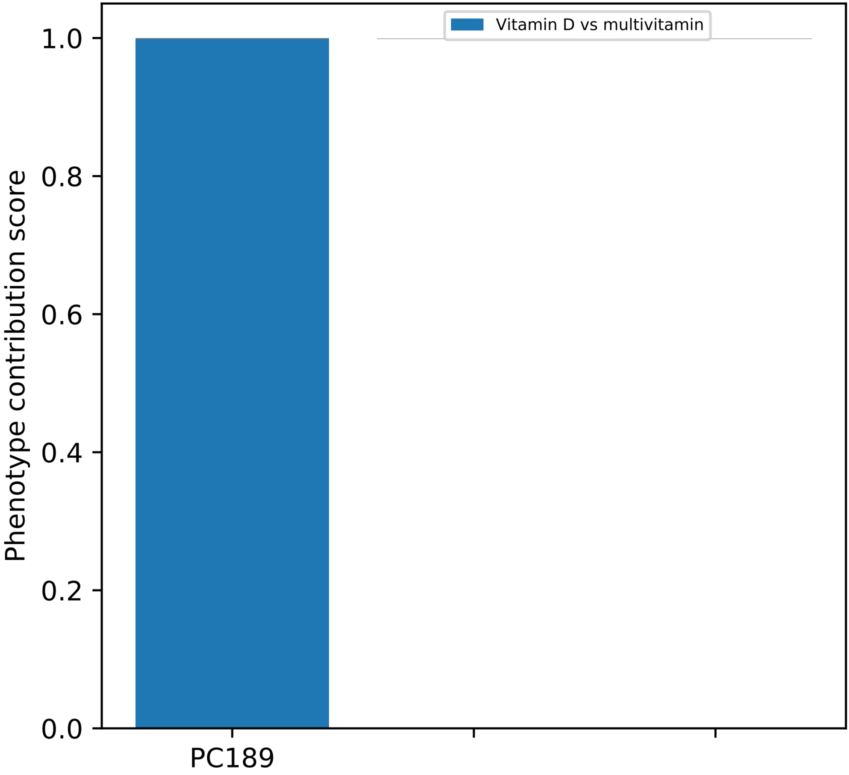
Vitamin D vs multivitamin
1.0
0.8
0.6
0.4
0.2
0.0
Phenotype contribution score
PC189

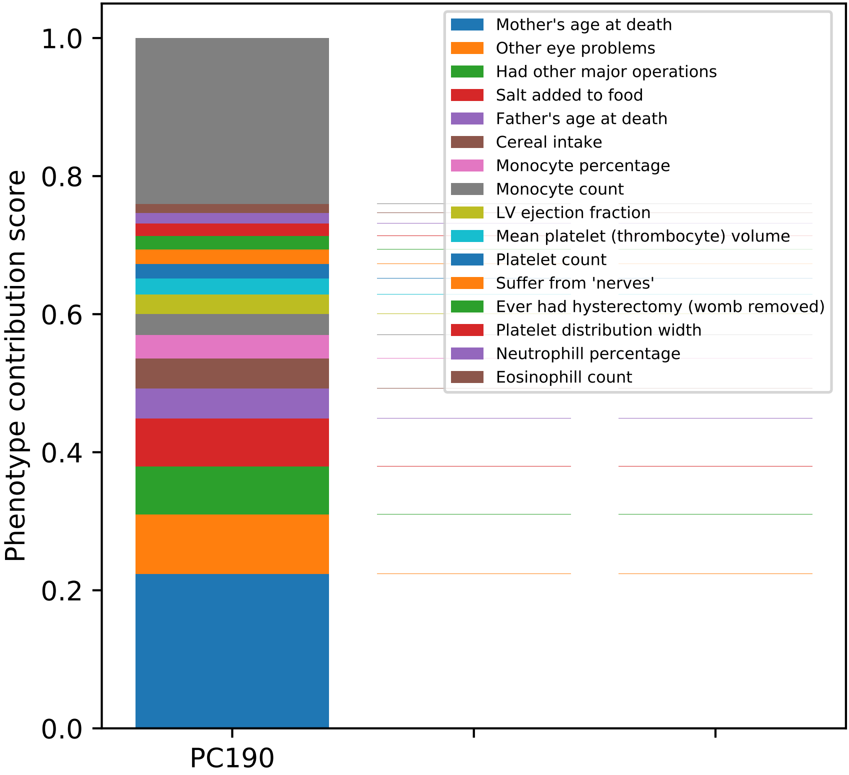

Phenotype contribution score
1.0
0.8
0.6
0.4
0.2
0.0
PC190
Mother's age at death
Other eye problems
Had other major operations
Salt added to food
Father's age at death
Cereal intake
Monocyte percentage
Monocyte count
LV ejection fraction
Mean platelet (thrombocyte) volume
Platelet count
Suffer from 'nerves'
Ever had hysterectomy (womb removed)
Platelet distribution width
Neutrophill percentage
Eosinophill count

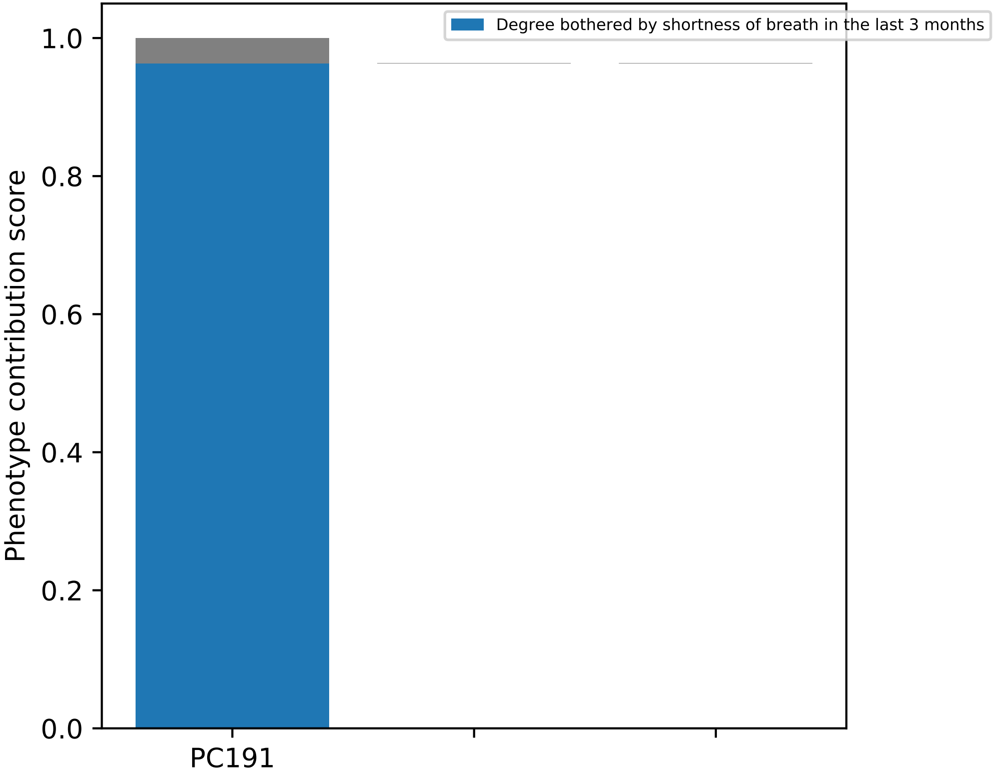
Degree bothered by shortness of breath in the last 3 months
Phenotype contribution score
1.0
0.8
0.6
0.4
0.2
0.0
PC191

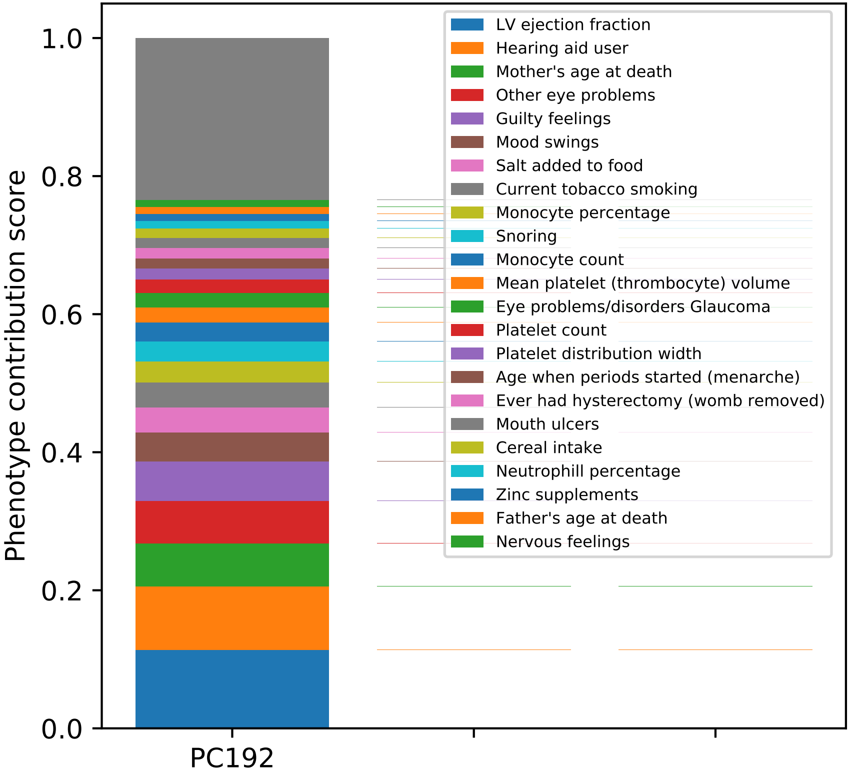
Phenotype contribution score
0.0
0.2
0.4
0.6
0.8
1.0
PC192
LV ejection fraction
Hearing aid user
Mother's age at death
Other eye problems
Guilty feelings
Mood swings
Salt added to food
Current tobacco smoking
Monocyte percentage
Snoring
Monocyte count
Mean platelet (thrombocyte) volume
Eye problems/disorders Glaucoma
Platelet count
Platelet distribution width
Age when periods started (menarche)
Ever had hysterectomy (womb removed)
Mouth ulcers
Cereal intake
Neutrophill percentage
Zinc supplements
Father's age at death
Nervous feelings

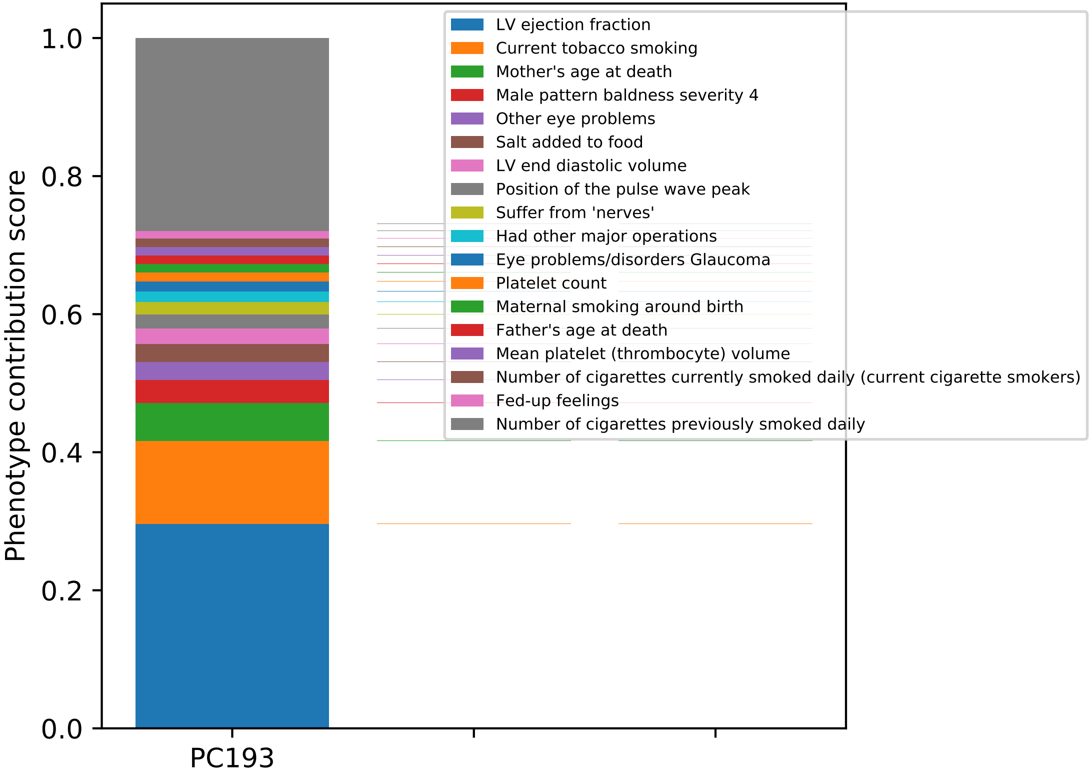

Phenotype contribution score
0.0
0.2
0.4
0.6
0.8
1.0
PC193
LV ejection fraction
Current tobacco smoking
Mother's age at death
Male pattern baldness severity 4
Other eye problems
Salt added to food
LV end diastolic volume
Position of the pulse wave peak
Suffer from 'nerves'
Had other major operations
Eye problems/disorders Glaucoma
Platelet count
Maternal smoking around birth
Father's age at death
Mean platelet (thrombocyte) volume
Number of cigarettes currently smoked daily (current cigarette smokers)
Fed-up feelings
Number of cigarettes previously smoked daily

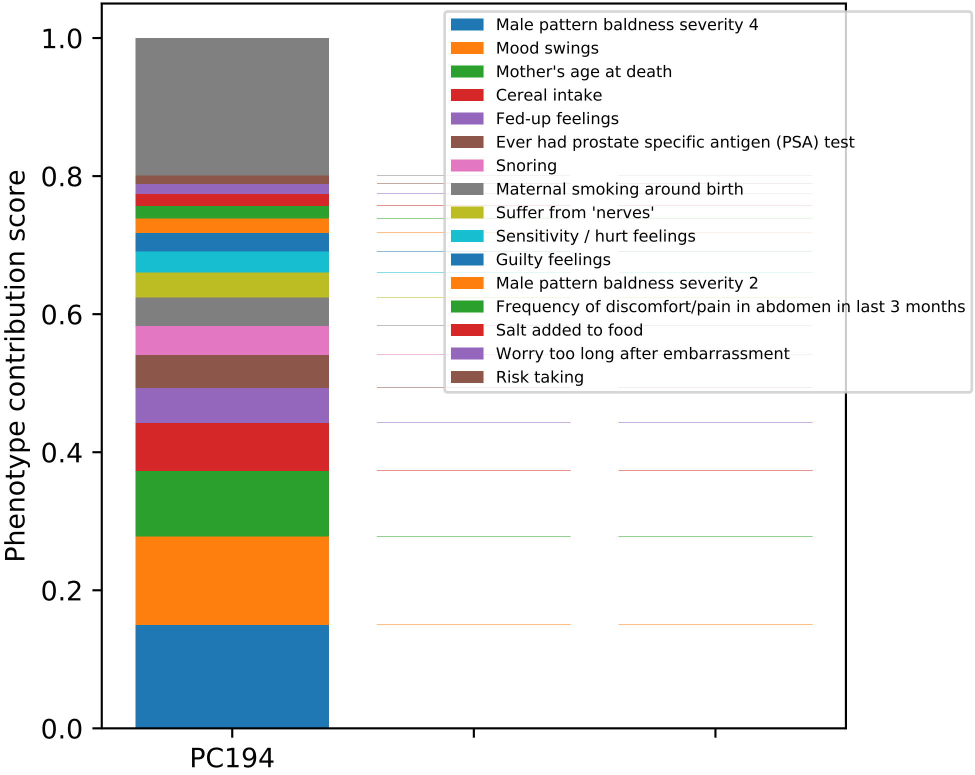
Phenotype contribution score
1.0
0.8
0.6
0.4
0.2
0.0
PC194
Male pattern baldness severity 4
Mood swings
Mother's age at death
Cereal intake
Fed-up feelings
Ever had prostate specific antigen (PSA) test
Snoring
Maternal smoking around birth
Suffer from 'nerves'
Sensitivity / hurt feelings
Guilty feelings
Male pattern baldness severity 2
Frequency of discomfort/pain in abdomen in last 3 months
Salt added to food
Worry too long after embarrassment
Risk taking

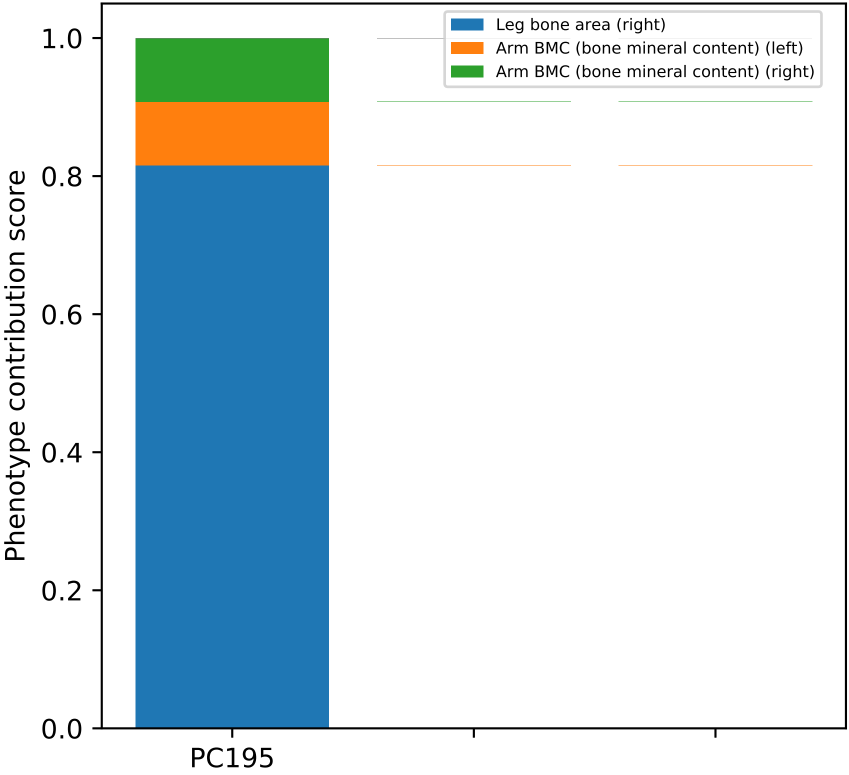
Leg bone area (right)
Arm BMC (bone mineral content) (left)
Arm BMC (bone mineral content) (right)
Phenotype contribution score
1.0
0.8
0.6
0.4
0.2
0.0
PC195

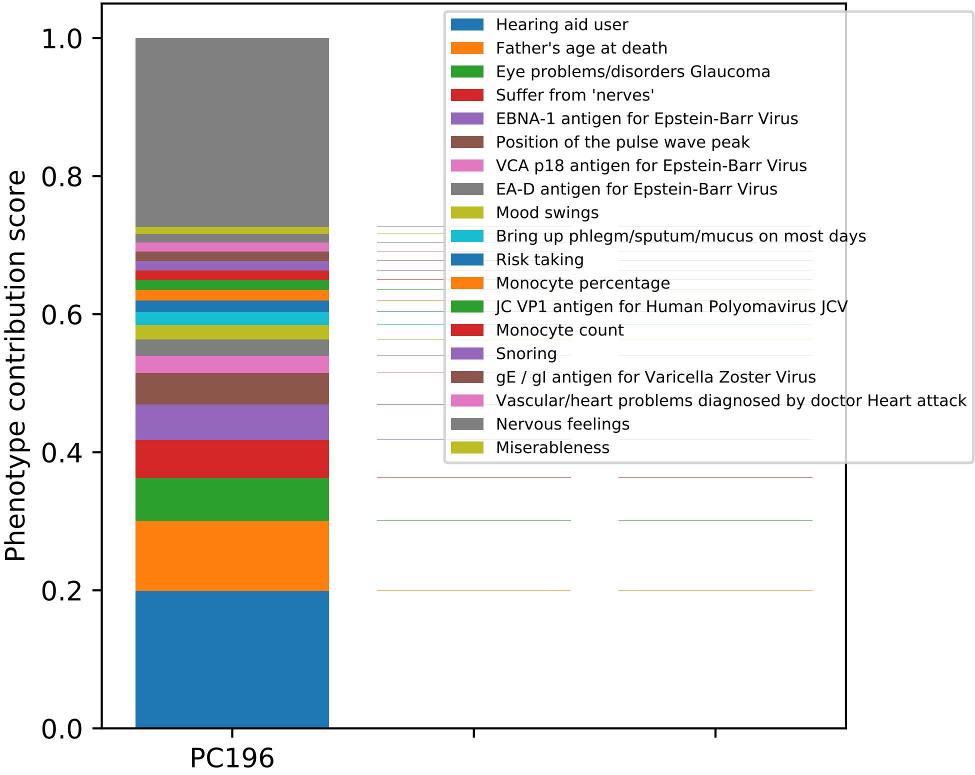
Phenotype contribution score
1.0
0.8
0.6
0.4
0.2
0.0
PC196
Hearing aid user
Father's age at death
Eye problems/disorders Glaucoma
Suffer from 'nerves'
EBNA-1 antigen for Epstein-Barr Virus
Position of the pulse wave peak
VCA p18 antigen for Epstein-Barr Virus
EA-D antigen for Epstein-Barr Virus
Mood swings
Bring up phlegm/sputum/mucus on most days
Risk taking
Monocyte percentage
JC VP1 antigen for Human Polyomavirus JCV
Monocyte count
Snoring
gE / gI antigen for Varicella Zoster Virus
Vascular/heart problems diagnosed by doctor Heart attack
Nervous feelings
Miserableness

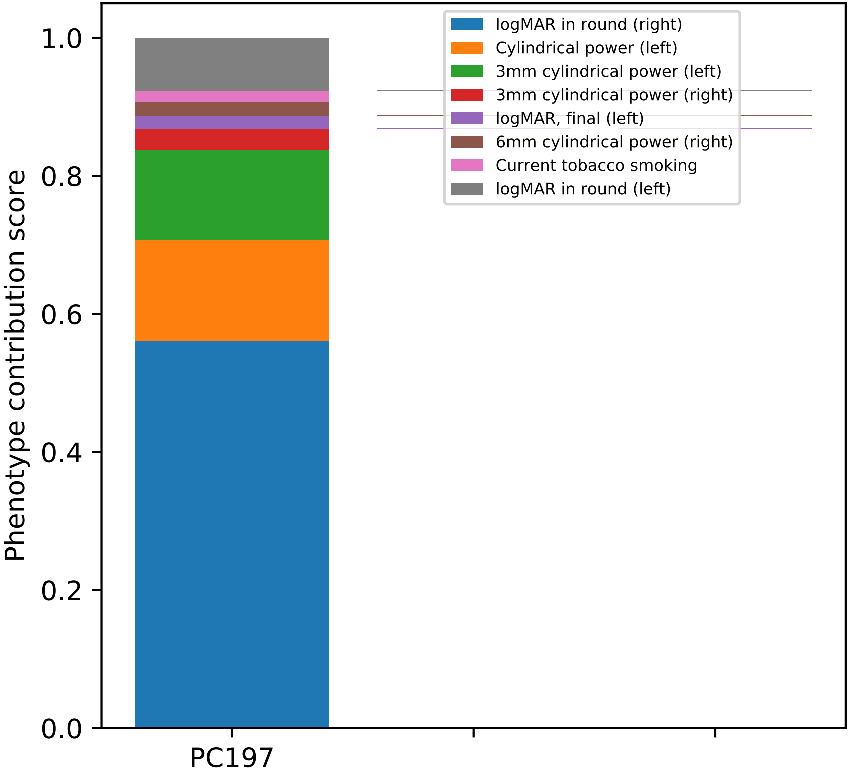

Phenotype contribution score
1.0
0.8
0.6
0.4
0.2
0.0
PC197
logMAR in round (right)
Cylindrical power (left)
3mm cylindrical power (left)
3mm cylindrical power (right)
logMAR, final (left)
6mm cylindrical power (right)
Current tobacco smoking
logMAR in round (left)

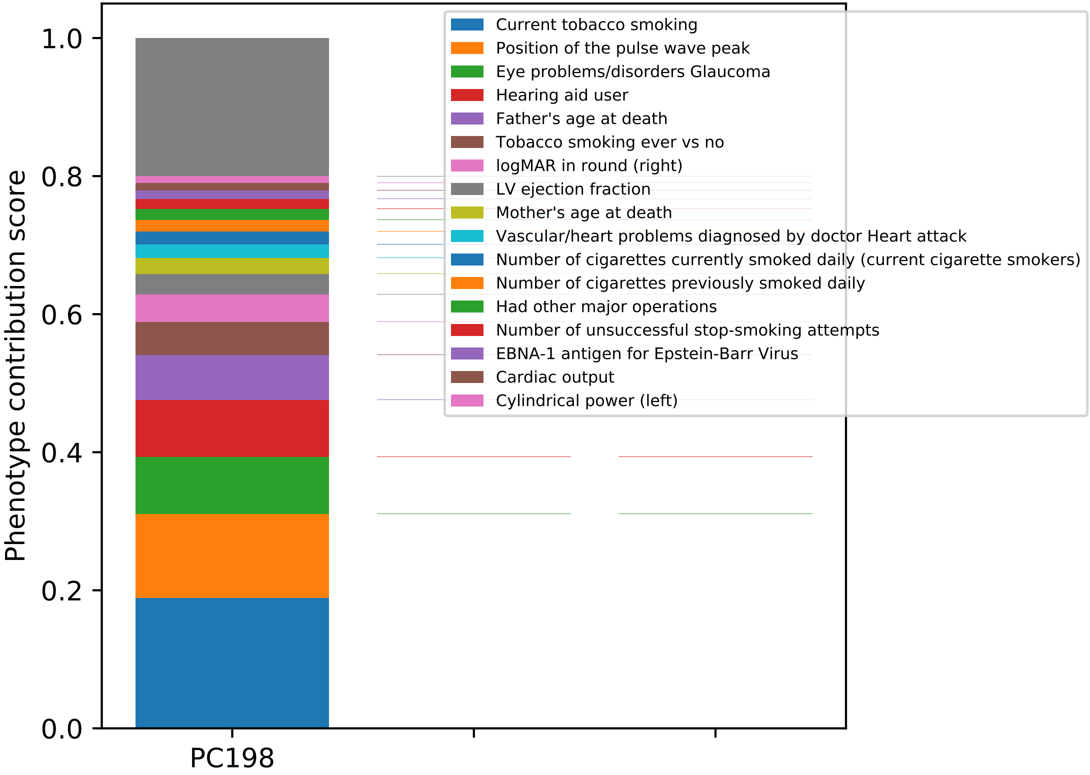
Phenotype contribution score
0.0
0.2
0.4
0.6
0.8
1.0
PC198
Current tobacco smoking
Position of the pulse wave peak
Eye problems/disorders Glaucoma
Hearing aid user
Father's age at death
Tobacco smoking ever vs no
logMAR in round (right)
LV ejection fraction
Mother's age at death
Vascular/heart problems diagnosed by doctor Heart attack
Number of cigarettes currently smoked daily (current cigarette smokers)
Number of cigarettes previously smoked daily
Had other major operations
Number of unsuccessful stop-smoking attempts
EBNA-1 antigen for Epstein-Barr Virus
Cardiac output
Cylindrical power (left)

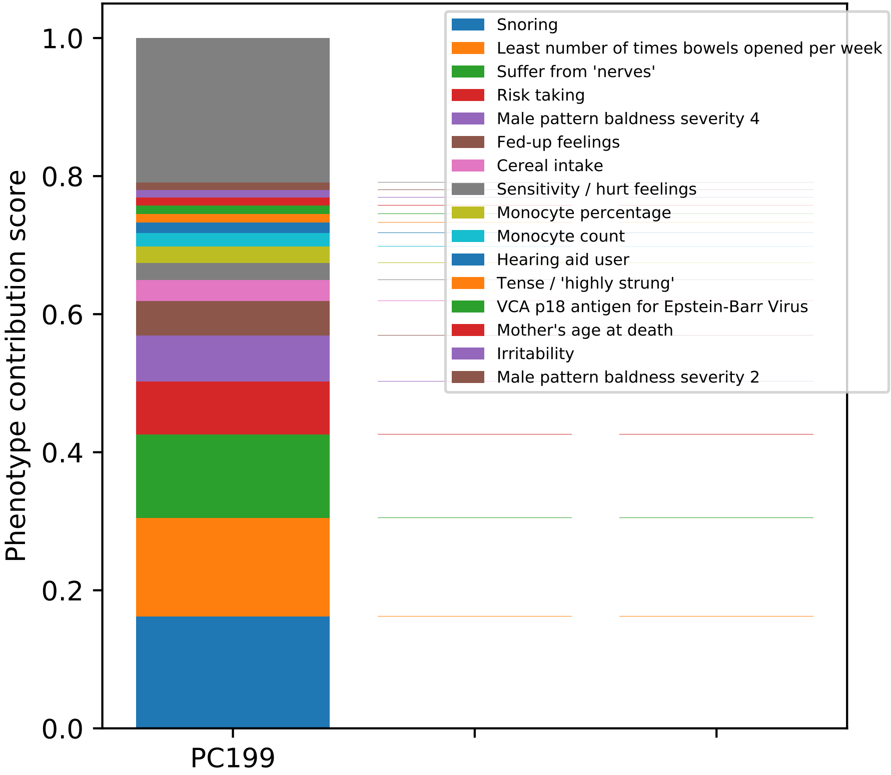
Phenotype contribution score
0.0
0.2
0.4
0.6
0.8
1.0
PC199
Snoring
Least number of times bowels opened per week
Suffer from 'nerves'
Risk taking
Male pattern baldness severity 4
Fed-up feelings
Cereal intake
Sensitivity / hurt feelings
Monocyte percentage
Monocyte count
Hearing aid user
Tense / 'highly strung'
VCA p18 antigen for Epstein-Barr Virus
Mother's age at death
Irritability
Male pattern baldness severity 2

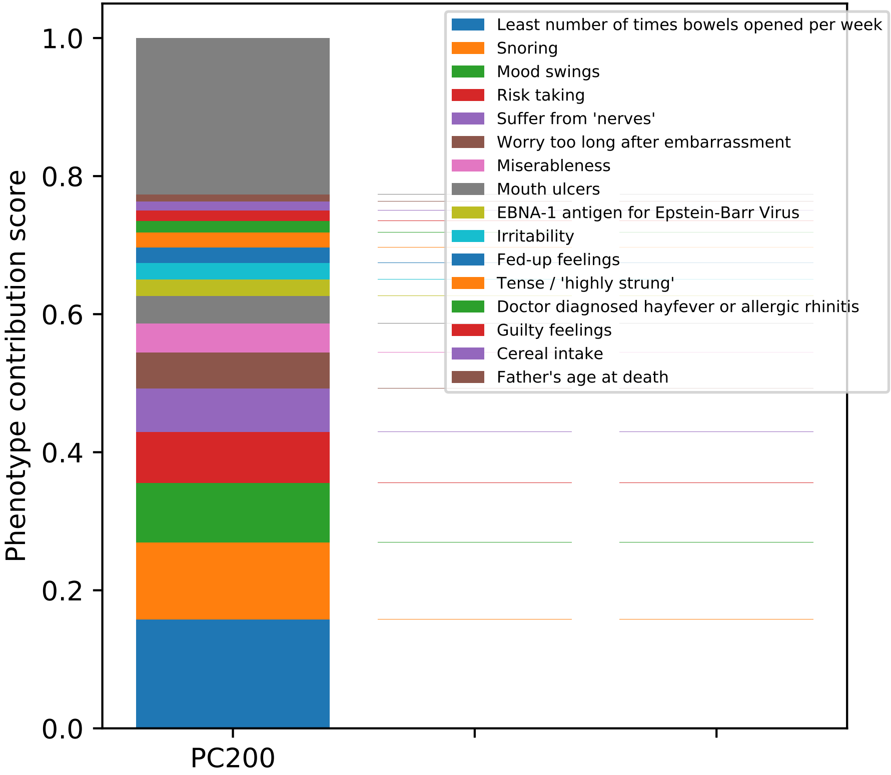
Phenotype contribution score
0.0
0.2
0.4
0.6
0.8
1.0
PC200
Least number of times bowels opened per week
Snoring
Mood swings
Risk taking
Suffer from 'nerves'
Worry too long after embarrassment
Miserableness
Mouth ulcers
EBNA-1 antigen for Epstein-Barr Virus
Irritability
Fed-up feelings
Tense / 'highly strung'
Doctor diagnosed hayfever or allergic rhinitis
Guilty feelings
Cereal intake
Father's age at death

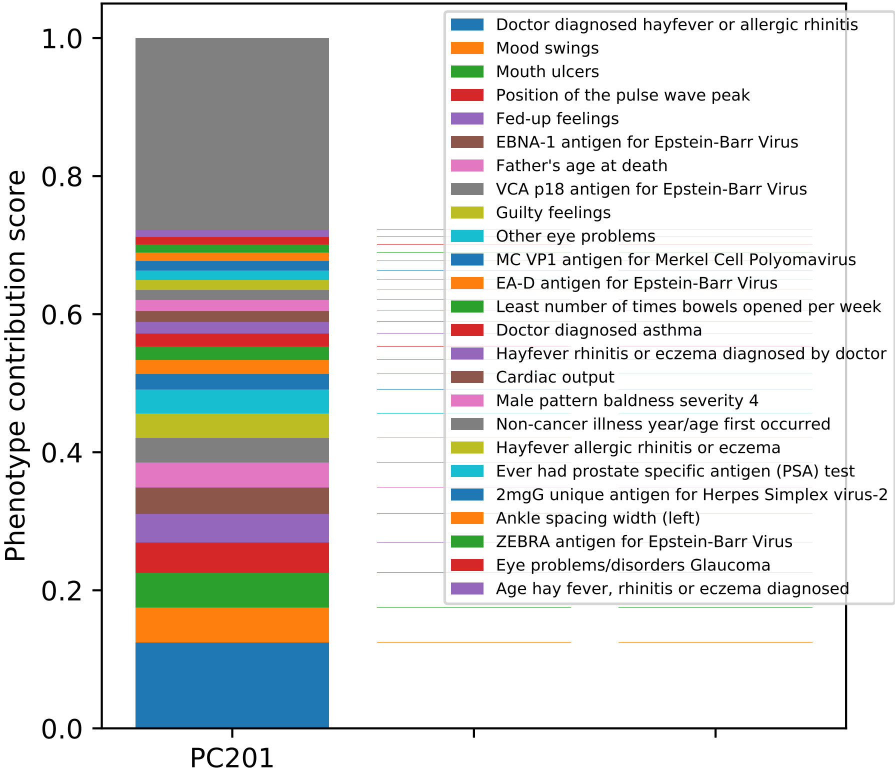
Phenotype contribution score
1.0
0.8
0.6
0.4
0.2
0.0
PC201
Doctor diagnosed hayfever or allergic rhinitis
Mood swings
Mouth ulcers
Position of the pulse wave peak
Fed-up feelings
EBNA-1 antigen for Epstein-Barr Virus
Father's age at death
VCA p18 antigen for Epstein-Barr Virus
Guilty feelings
Other eye problems
MC VP1 antigen for Merkel Cell Polyomavirus
EA-D antigen for Epstein-Barr Virus
Least number of times bowels opened per week
Doctor diagnosed asthma
Hayfever rhinitis or eczema diagnosed by doctor
Cardiac output
Male pattern baldness severity 4
Non-cancer illness year/age first occurred
Hayfever allergic rhinitis or eczema
Ever had prostate specific antigen (PSA) test
2mgG unique antigen for Herpes Simplex virus-2
Ankle spacing width (left)
ZEBRA antigen for Epstein-Barr Virus
Eye problems/disorders Glaucoma
Age hay fever, rhinitis or eczema diagnosed

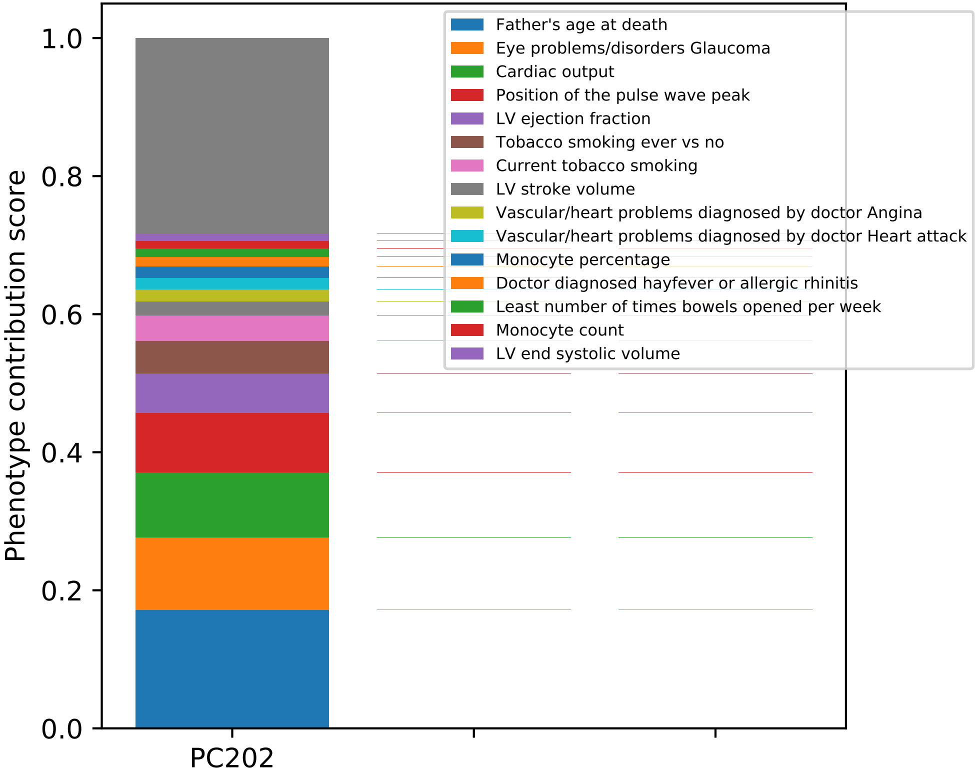

Phenotype contribution score
1.0
0.8
0.6
0.4
0.2
0.0
PC202
Father's age at death
Eye problems/disorders Glaucoma
Cardiac output
Position of the pulse wave peak
LV ejection fraction
Tobacco smoking ever vs no
Current tobacco smoking
LV stroke volume
Vascular/heart problems diagnosed by doctor Angina
Vascular/heart problems diagnosed by doctor Heart attack
Monocyte percentage
Doctor diagnosed hayfever or allergic rhinitis
Least number of times bowels opened per week
Monocyte count
LV end systolic volume

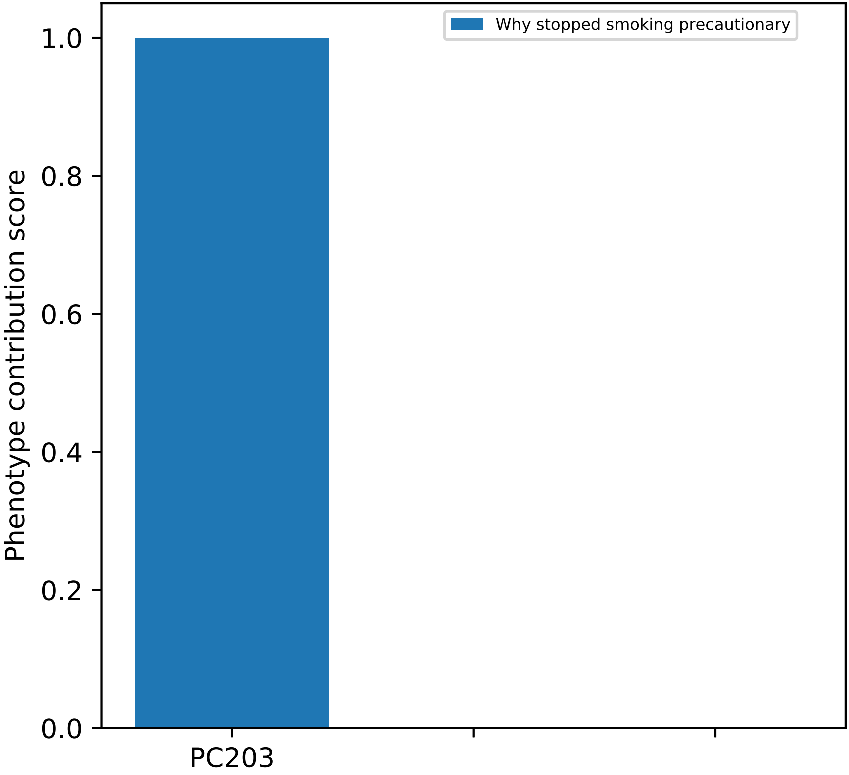
Why stopped smoking precautionary
Phenotype contribution score
1.0
0.8
0.6
0.4
0.2
0.0
PC203

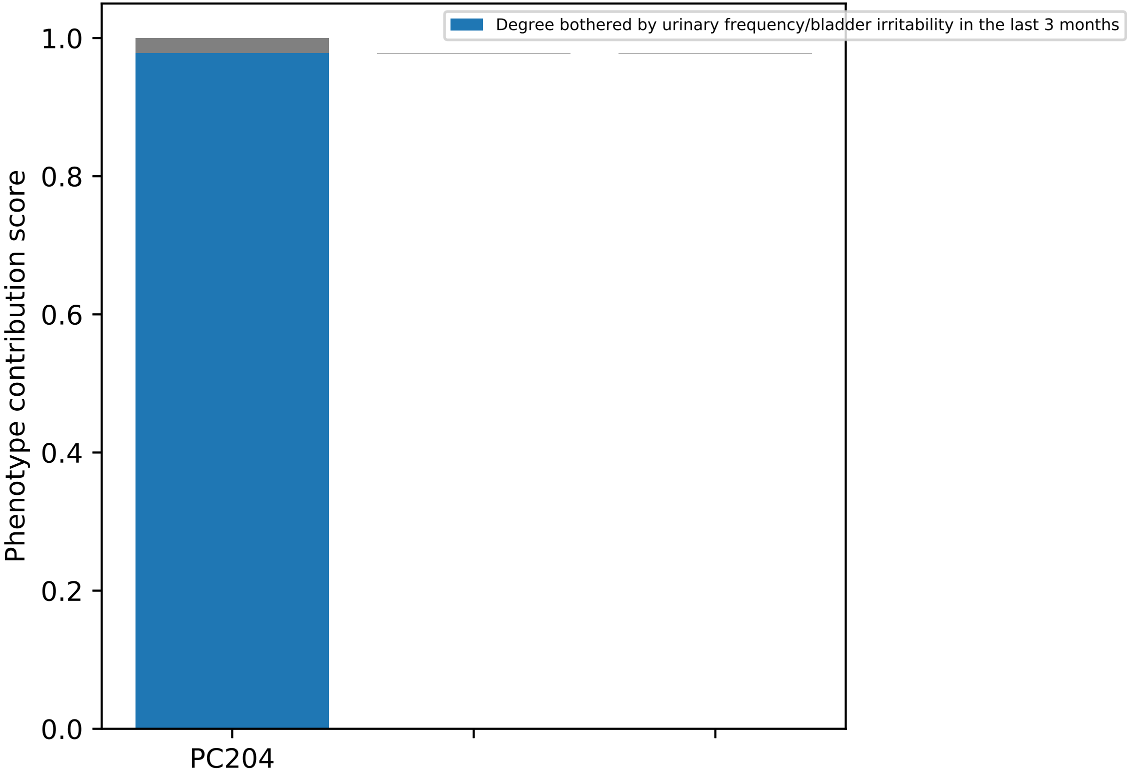

Degree bothered by urinary frequency/bladder irritability in the last 3 months
Phenotype contribution score
1.0
0.8
0.6
0.4
0.2
0.0
PC204

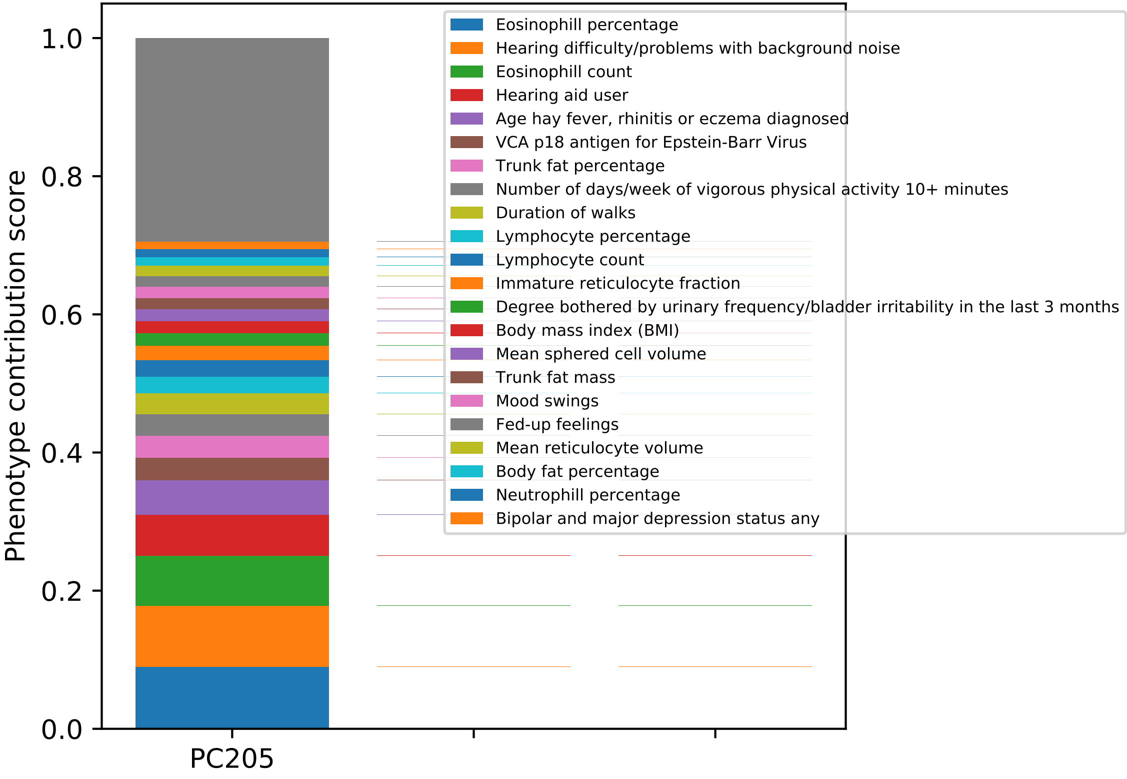
Phenotype contribution score
0.0
0.2
0.4
0.6
0.8
1.0
PC205
Eosinophill percentage
Hearing difficulty/problems with background noise
Eosinophill count
Hearing aid user
Age hay fever, rhinitis or eczema diagnosed
VCA p18 antigen for Epstein-Barr Virus
Trunk fat percentage
Number of days/week of vigorous physical activity 10+ minutes
Duration of walks
Lymphocyte percentage
Lymphocyte count
Immature reticulocyte fraction
Degree bothered by urinary frequency/bladder irritability in the last 3 months
Body mass index (BMI)
Mean sphered cell volume
Trunk fat mass
Mood swings
Fed-up feelings
Mean reticulocyte volume
Body fat percentage
Neutrophill percentage
Bipolar and major depression status any

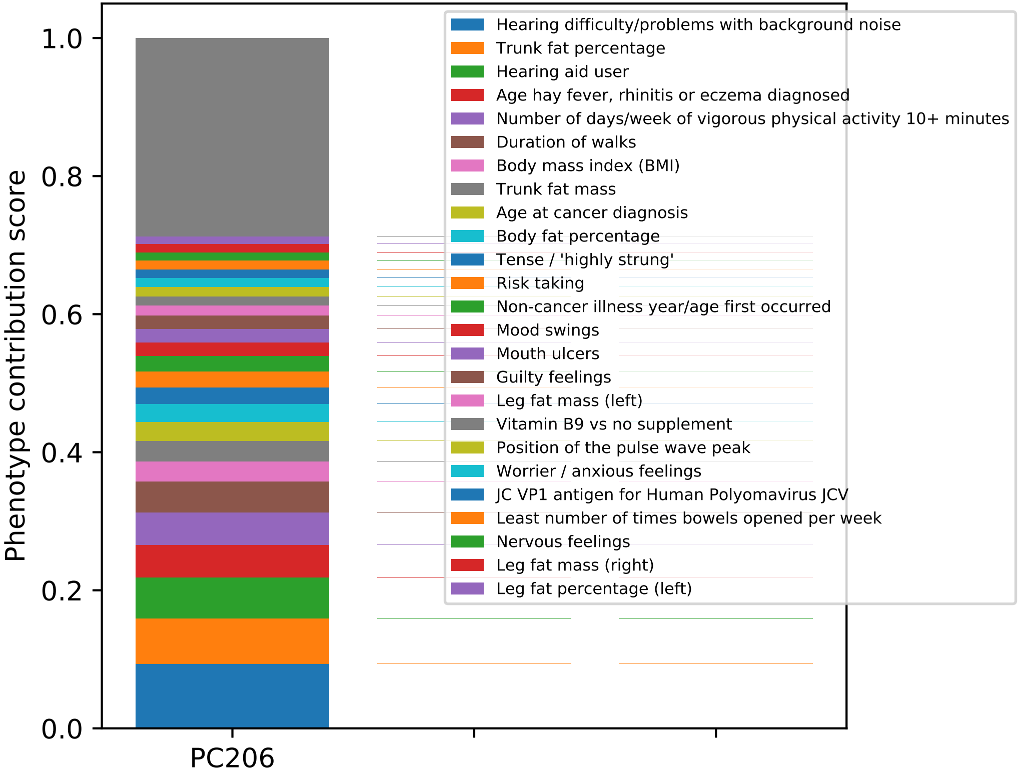
Phenotype contribution score
0.0
0.2
0.4
0.6
0.8
1.0
PC206
Hearing difficulty/problems with background noise
Trunk fat percentage
Hearing aid user
Age hay fever, rhinitis or eczema diagnosed
Number of days/week of vigorous physical activity 10+ minutes
Duration of walks
Body mass index (BMI)
Trunk fat mass
Age at cancer diagnosis
Body fat percentage
Tense / 'highly strung'
Risk taking
Non-cancer illness year/age first occurred
Mood swings
Mouth ulcers
Guilty feelings
Leg fat mass (left)
Vitamin B9 vs no supplement
Position of the pulse wave peak
Worrier / anxious feelings
JC VP1 antigen for Human Polyomavirus JCV
Least number of times bowels opened per week
Nervous feelings
Leg fat mass (right)
Leg fat percentage (left)

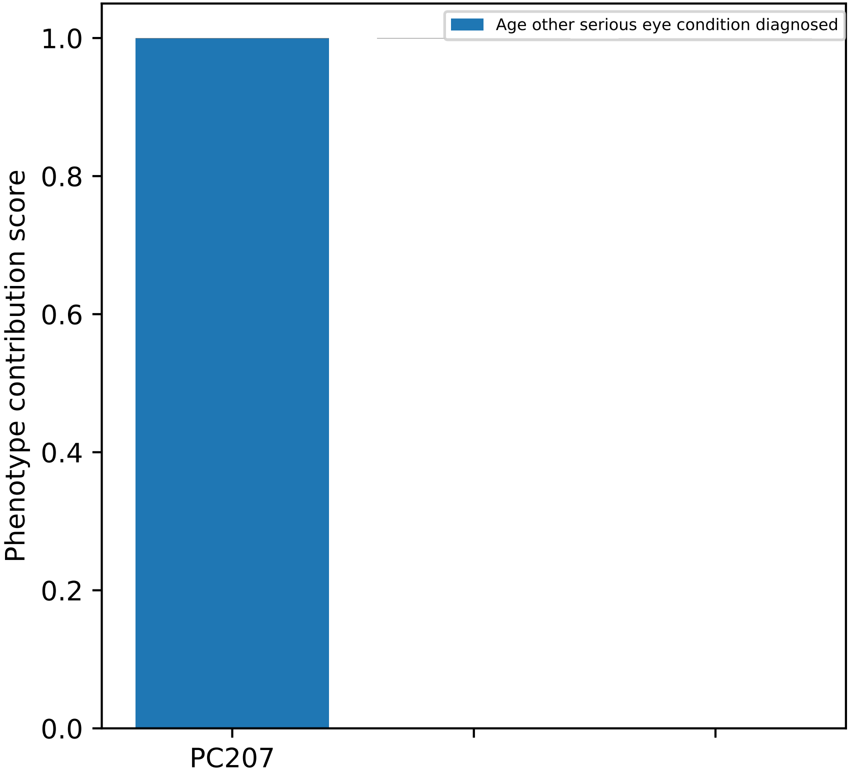

Age other serious eye condition diagnosed
Phenotype contribution score
1.0
0.8
0.6
0.4
0.2
0.0
PC207

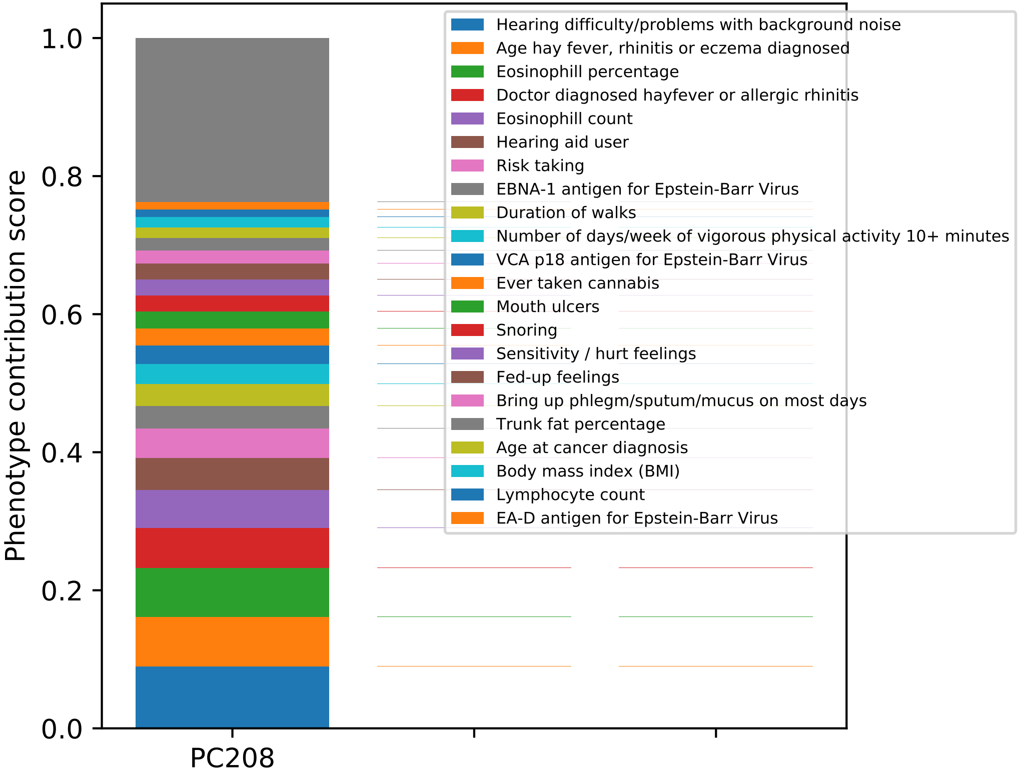

Phenotype contribution score
1.0
0.8
0.6
0.4
0.2
0.0
PC208
Hearing difficulty/problems with background noise
Age hay fever, rhinitis or eczema diagnosed
Eosinophill percentage
Doctor diagnosed hayfever or allergic rhinitis
Eosinophill count
Hearing aid user
Risk taking
EBNA-1 antigen for Epstein-Barr Virus
Duration of walks
Number of days/week of vigorous physical activity 10+ minutes
VCA p18 antigen for Epstein-Barr Virus
Ever taken cannabis
Mouth ulcers
Snoring
Sensitivity / hurt feelings
Fed-up feelings
Bring up phlegm/sputum/mucus on most days
Trunk fat percentage
Age at cancer diagnosis
Body mass index (BMI)
Lymphocyte count
EA-D antigen for Epstein-Barr Virus

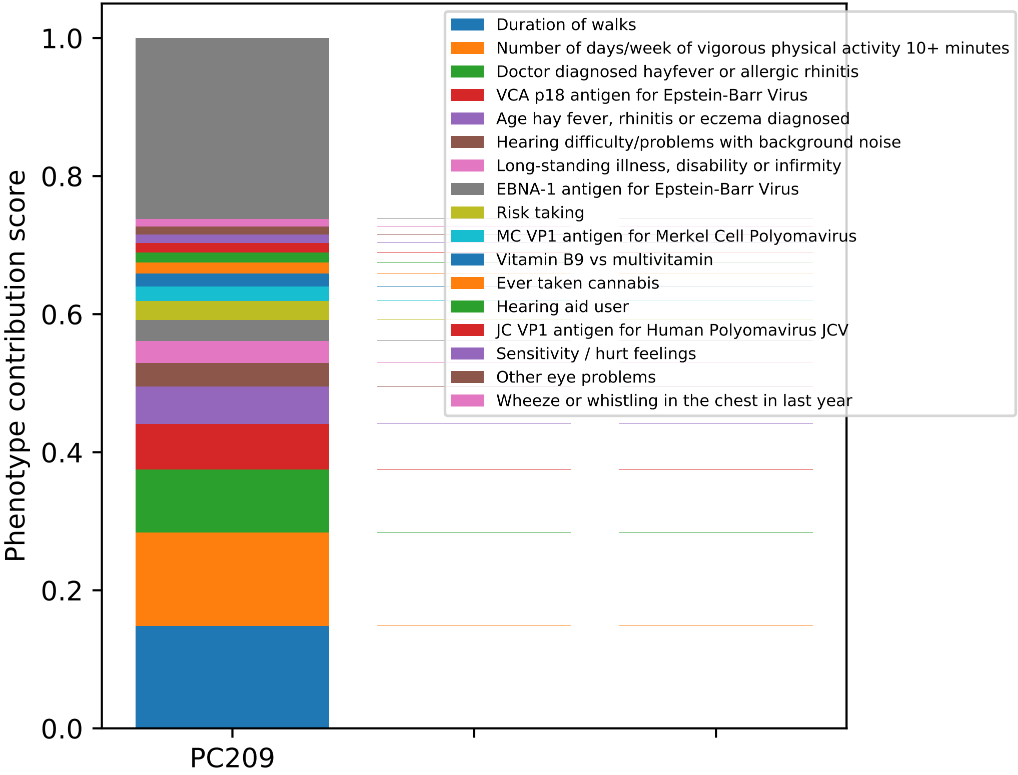

Phenotype contribution score
0.0
0.2
0.4
0.6
0.8
1.0
PC209
Duration of walks
Number of days/week of vigorous physical activity 10+ minutes
Doctor diagnosed hayfever or allergic rhinitis
VCA p18 antigen for Epstein-Barr Virus
Age hay fever, rhinitis or eczema diagnosed
Hearing difficulty/problems with background noise
Long-standing illness, disability or infirmity
EBNA-1 antigen for Epstein-Barr Virus
Risk taking
MC VP1 antigen for Merkel Cell Polyomavirus
Vitamin B9 vs multivitamin
Ever taken cannabis
Hearing aid user
JC VP1 antigen for Human Polyomavirus JCV
Sensitivity / hurt feelings
Other eye problems
Wheeze or whistling in the chest in last year

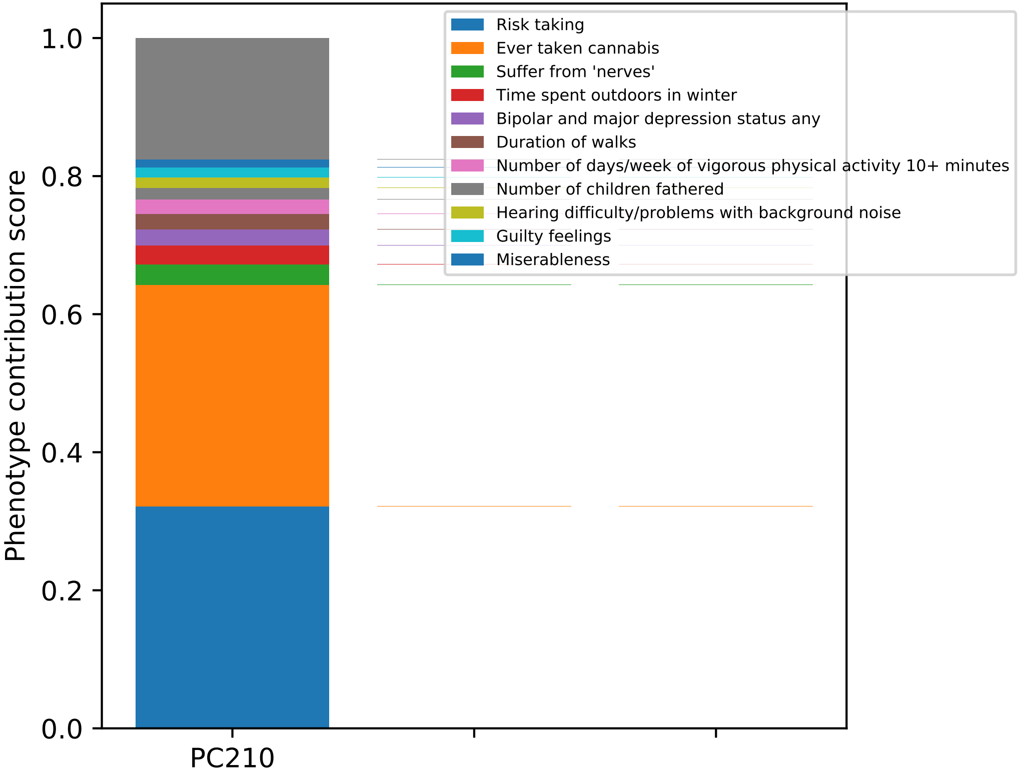

Risk taking
Ever taken cannabis
Suffer from 'nerves'
Time spent outdoors in winter
Bipolar and major depression status any
Duration of walks
Number of days/week of vigorous physical activity 10+ minutes
Number of children fathered
Hearing difficulty/problems with background noise
Guilty feelings
Miserableness
Phenotype contribution score
1.0
0.8
0.6
0.4
0.2
0.0
PC210

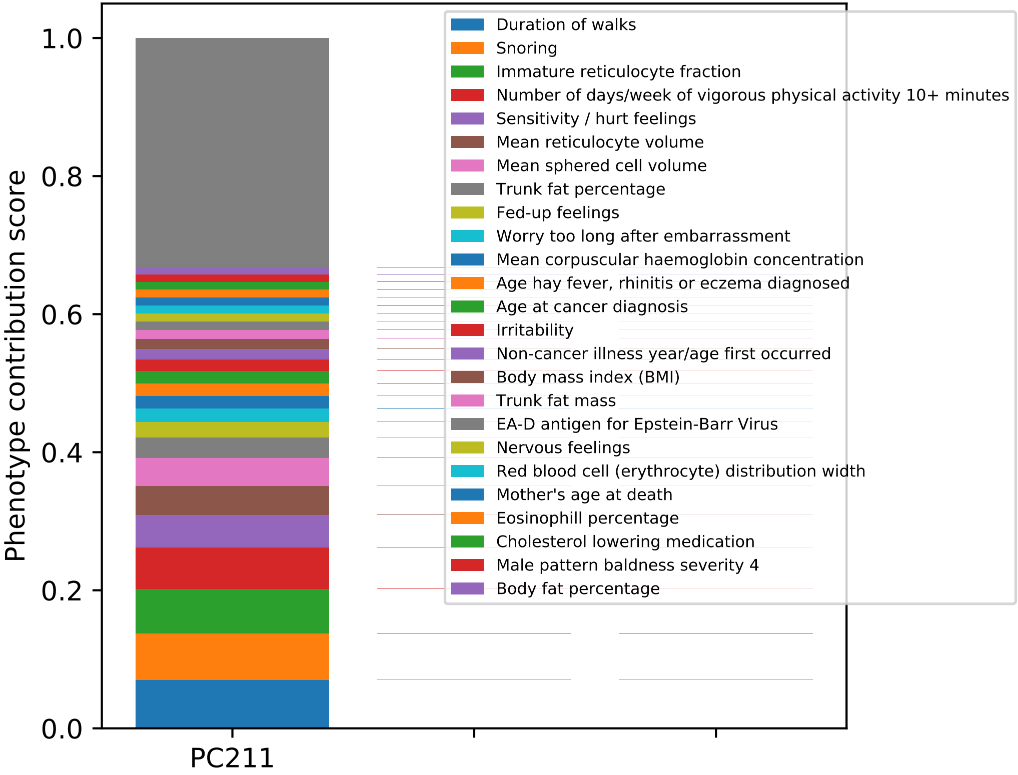

Phenotype contribution score
1.0
0.8
0.6
0.4
0.2
0.0
PC211
Duration of walks
Snoring
Immature reticulocyte fraction
Number of days/week of vigorous physical activity 10+ minutes
Sensitivity / hurt feelings
Mean reticulocyte volume
Mean sphered cell volume
Trunk fat percentage
Fed-up feelings
Worry too long after embarrassment
Mean corpuscular haemoglobin concentration
Age hay fever, rhinitis or eczema diagnosed
Age at cancer diagnosis
Irritability
Non-cancer illness year/age first occurred
Body mass index (BMI)
Trunk fat mass
EA-D antigen for Epstein-Barr Virus
Nervous feelings
Red blood cell (erythrocyte) distribution width
Mother's age at death
Eosinophill percentage
Cholesterol lowering medication
Male pattern baldness severity 4
Body fat percentage

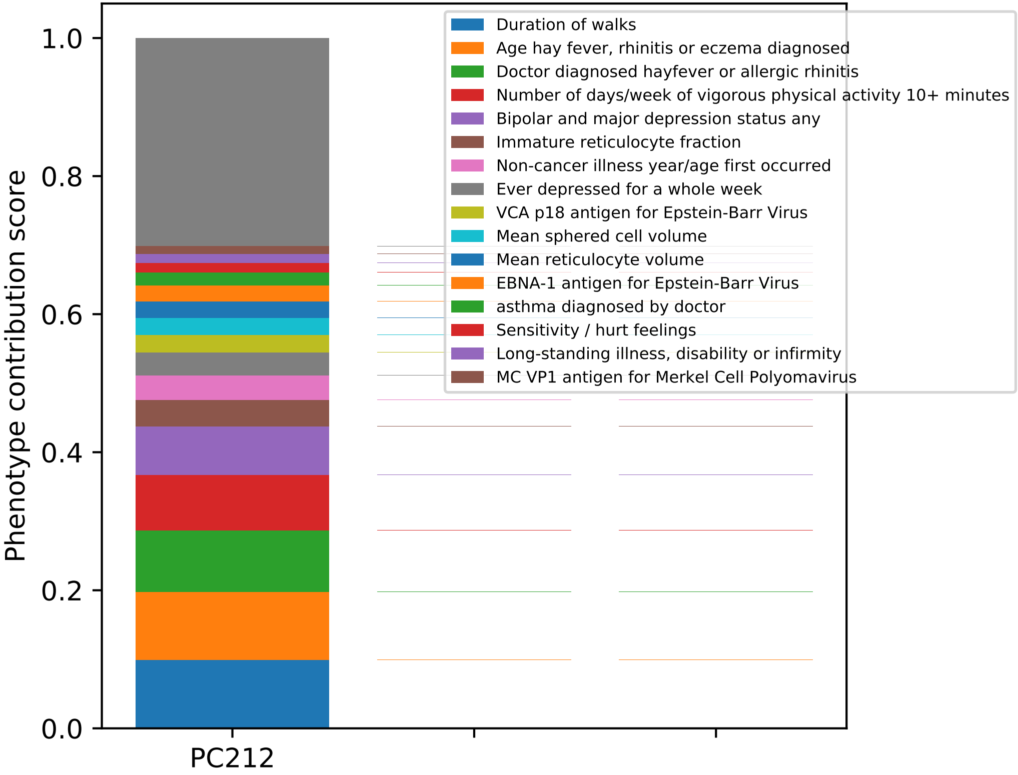

Phenotype contribution score
1.0
0.8
0.6
0.4
0.2
0.0
PC212
Duration of walks
Age hay fever, rhinitis or eczema diagnosed
Doctor diagnosed hayfever or allergic rhinitis
Number of days/week of vigorous physical activity 10+ minutes
Bipolar and major depression status any
Immature reticulocyte fraction
Non-cancer illness year/age first occurred
Ever depressed for a whole week
VCA p18 antigen for Epstein-Barr Virus
Mean sphered cell volume
Mean reticulocyte volume
EBNA-1 antigen for Epstein-Barr Virus
asthma diagnosed by doctor
Sensitivity / hurt feelings
Long-standing illness, disability or infirmity
MC VP1 antigen for Merkel Cell Polyomavirus

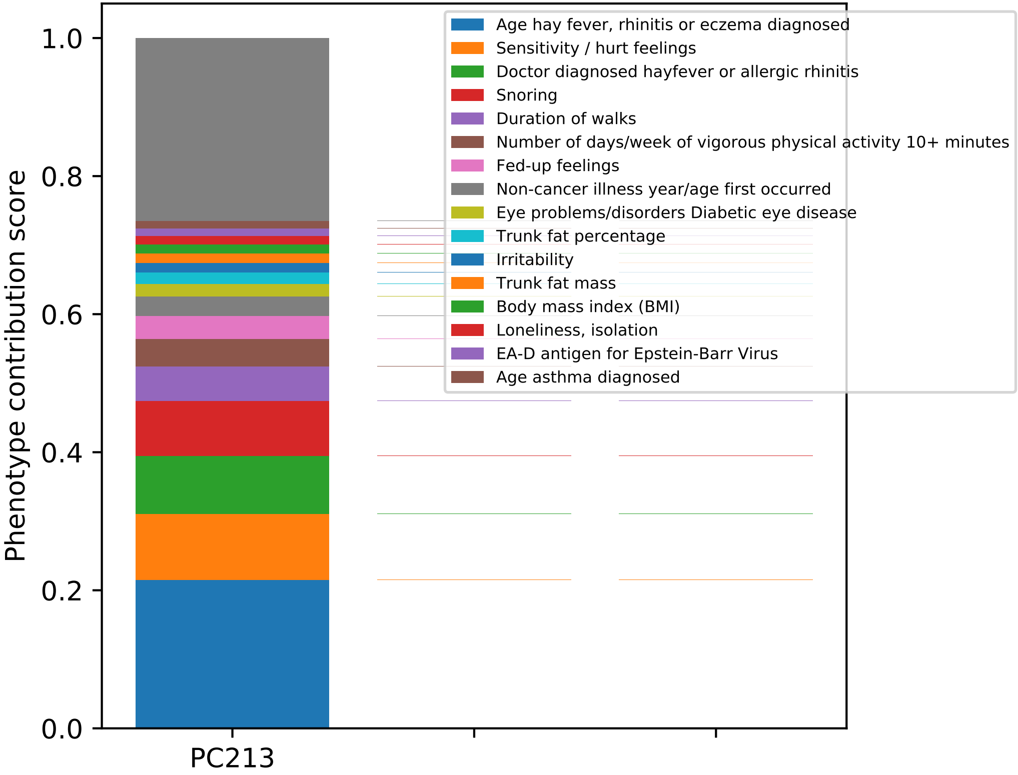
Phenotype contribution score
0.0
0.2
0.4
0.6
0.8
1.0
PC213
Age hay fever, rhinitis or eczema diagnosed
Sensitivity / hurt feelings
Doctor diagnosed hayfever or allergic rhinitis
Snoring
Duration of walks
Number of days/week of vigorous physical activity 10+ minutes
Fed-up feelings
Non-cancer illness year/age first occurred
Eye problems/disorders Diabetic eye disease
Trunk fat percentage
Irritability
Trunk fat mass
Body mass index (BMI)
Loneliness, isolation
EA-D antigen for Epstein-Barr Virus
Age asthma diagnosed

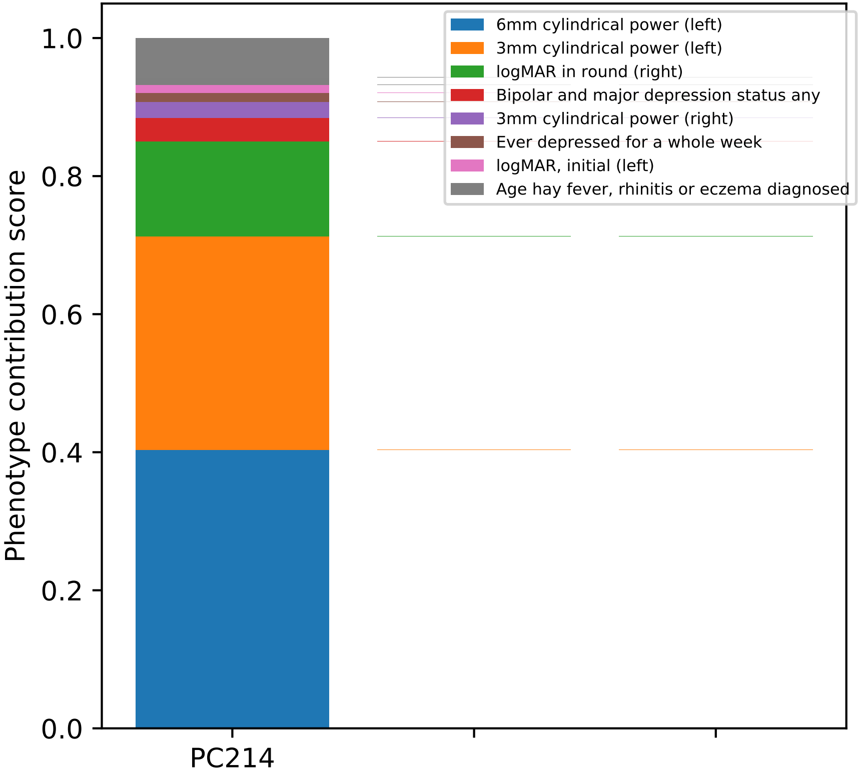

Phenotype contribution score
1.0
0.8
0.6
0.4
0.2
0.0
PC214
6mm cylindrical power (left)
3mm cylindrical power (left)
logMAR in round (right)
Bipolar and major depression status any
3mm cylindrical power (right)
Ever depressed for a whole week
logMAR, initial (left)
Age hay fever, rhinitis or eczema diagnosed

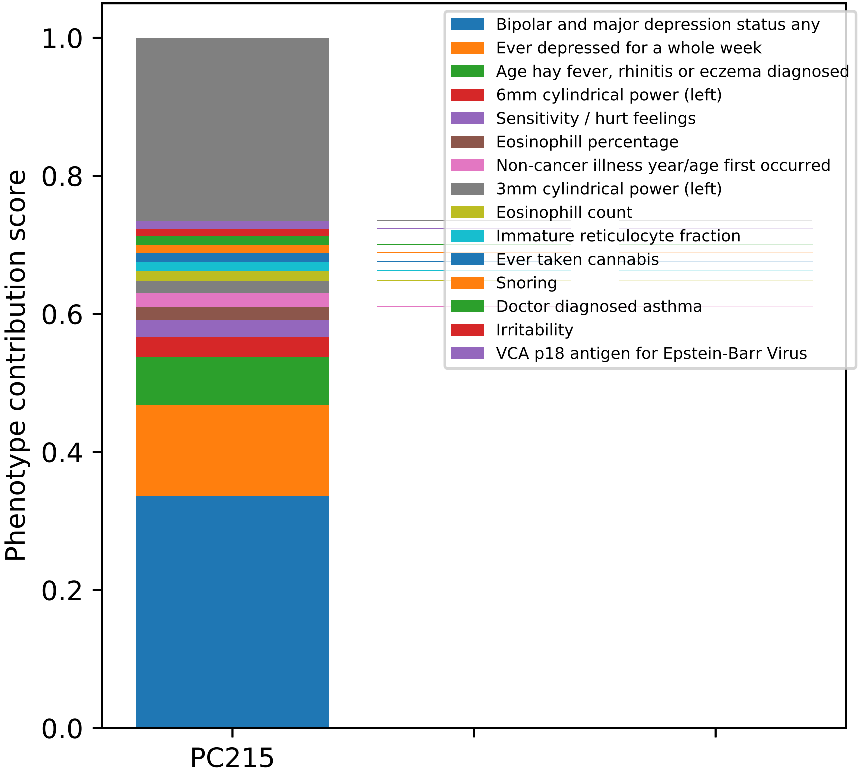

Bipolar and major depression status any
Ever depressed for a whole week
Age hay fever, rhinitis or eczema diagnosed
6mm cylindrical power (left)
Sensitivity / hurt feelings
Eosinophill percentage
Non-cancer illness year/age first occurred
3mm cylindrical power (left)
Eosinophill count
Immature reticulocyte fraction
Ever taken cannabis
Snoring
Doctor diagnosed asthma
Irritability
VCA p18 antigen for Epstein-Barr Virus
Phenotype contribution score
1.0
0.8
0.6
0.4
0.2
0.0
PC215

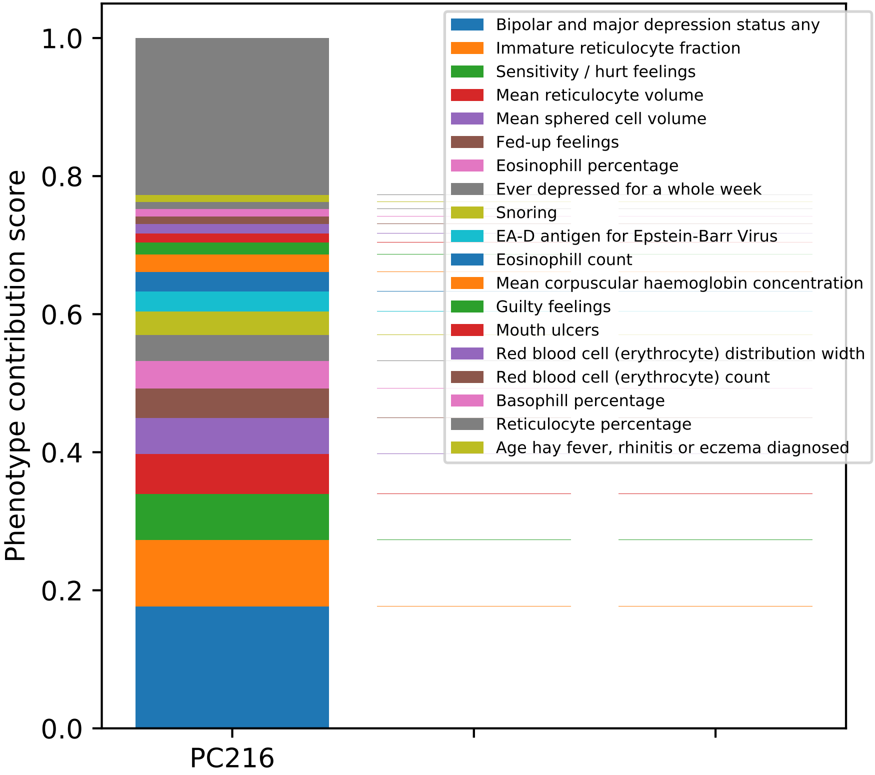

Phenotype contribution score
0.0
0.2
0.4
0.6
0.8
1.0
PC216
Bipolar and major depression status any
Immature reticulocyte fraction
Sensitivity / hurt feelings
Mean reticulocyte volume
Mean sphered cell volume
Fed-up feelings
Eosinophill percentage
Ever depressed for a whole week
Snoring
EA-D antigen for Epstein-Barr Virus
Eosinophill count
Mean corpuscular haemoglobin concentration
Guilty feelings
Mouth ulcers
Red blood cell (erythrocyte) distribution width
Red blood cell (erythrocyte) count
Basophill percentage
Reticulocyte percentage
Age hay fever, rhinitis or eczema diagnosed

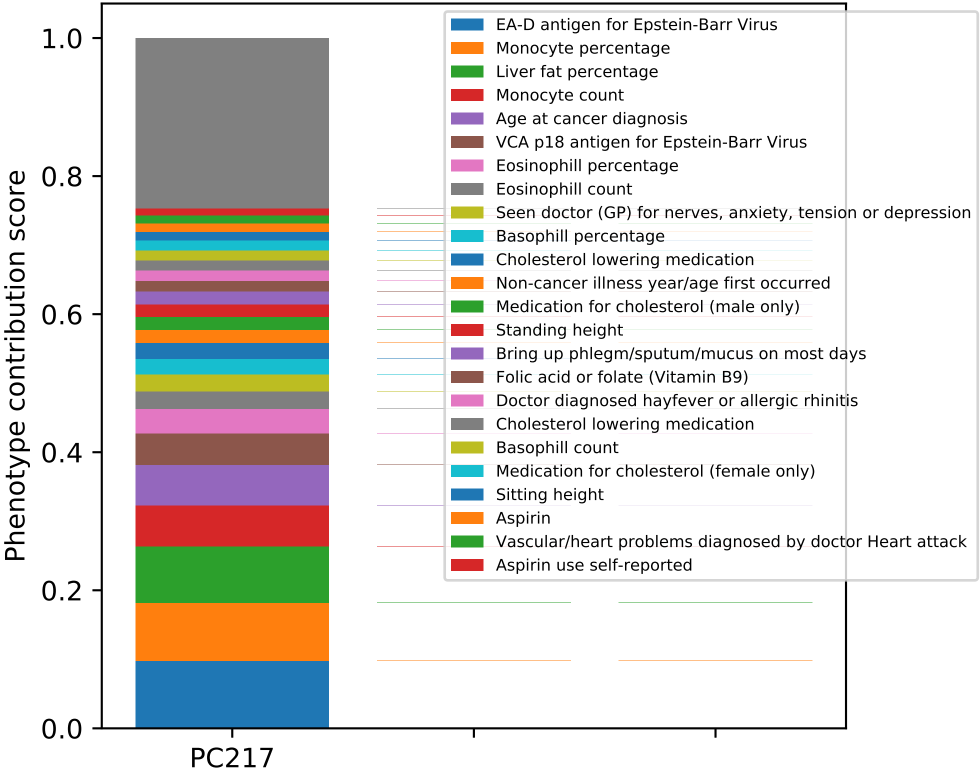

Phenotype contribution score
1.0
0.8
0.6
0.4
0.2
0.0
PC217
EA-D antigen for Epstein-Barr Virus
Monocyte percentage
Liver fat percentage
Monocyte count
Age at cancer diagnosis
VCA p18 antigen for Epstein-Barr Virus
Eosinophill percentage
Eosinophill count
Seen doctor (GP) for nerves, anxiety, tension or depression
Basophill percentage
Cholesterol lowering medication
Non-cancer illness year/age first occurred
Medication for cholesterol (male only)
Standing height
Bring up phlegm/sputum/mucus on most days
Folic acid or folate (Vitamin B9)
Doctor diagnosed hayfever or allergic rhinitis
Cholesterol lowering medication
Basophill count
Medication for cholesterol (female only)
Sitting height
Aspirin
Vascular/heart problems diagnosed by doctor Heart attack
Aspirin use self-reported

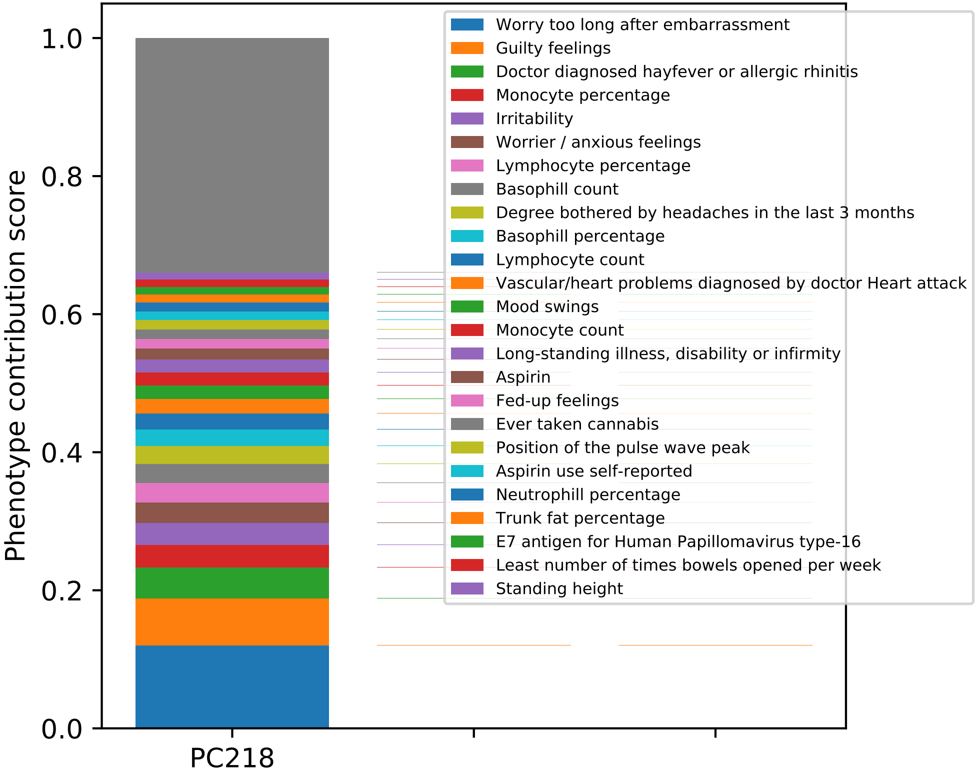
Phenotype contribution score
1.0
0.8
0.6
0.4
0.2
0.0
PC218
Worry too long after embarrassment
Guilty feelings
Doctor diagnosed hayfever or allergic rhinitis
Monocyte percentage
Irritability
Worrier / anxious feelings
Lymphocyte percentage
Basophill count
Degree bothered by headaches in the last 3 months
Basophill percentage
Lymphocyte count
Vascular/heart problems diagnosed by doctor Heart attack
Mood swings
Monocyte count
Long-standing illness, disability or infirmity
Aspirin
Fed-up feelings
Ever taken cannabis
Position of the pulse wave peak
Aspirin use self-reported
Neutrophill percentage
Trunk fat percentage
E7 antigen for Human Papillomavirus type-16
Least number of times bowels opened per week
Standing height

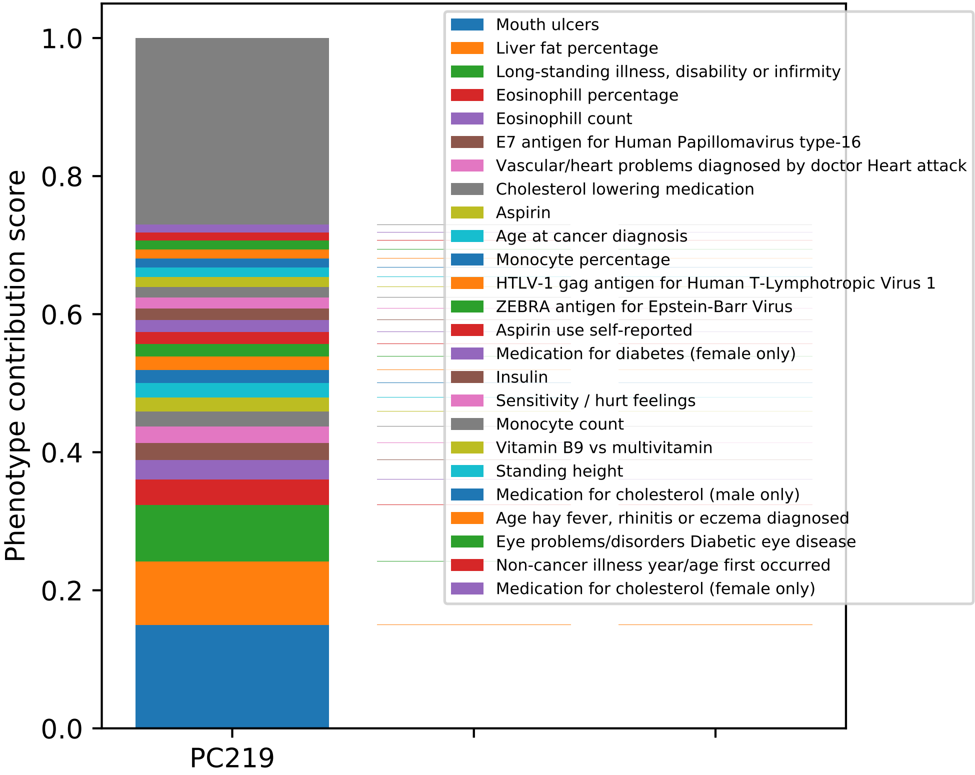

Phenotype contribution score
0.0
0.2
0.4
0.6
0.8
1.0
PC219
Mouth ulcers
Liver fat percentage
Long-standing illness, disability or infirmity
Eosinophill percentage
Eosinophill count
E7 antigen for Human Papillomavirus type-16
Vascular/heart problems diagnosed by doctor Heart attack
Cholesterol lowering medication
Aspirin
Age at cancer diagnosis
Monocyte percentage
HTLV-1 gag antigen for Human T-Lymphotropic Virus 1
ZEBRA antigen for Epstein-Barr Virus
Aspirin use self-reported
Medication for diabetes (female only)
Insulin
Sensitivity / hurt feelings
Monocyte count
Vitamin B9 vs multivitamin
Standing height
Medication for cholesterol (male only)
Age hay fever, rhinitis or eczema diagnosed
Eye problems/disorders Diabetic eye disease
Non-cancer illness year/age first occurred
Medication for cholesterol (female only)

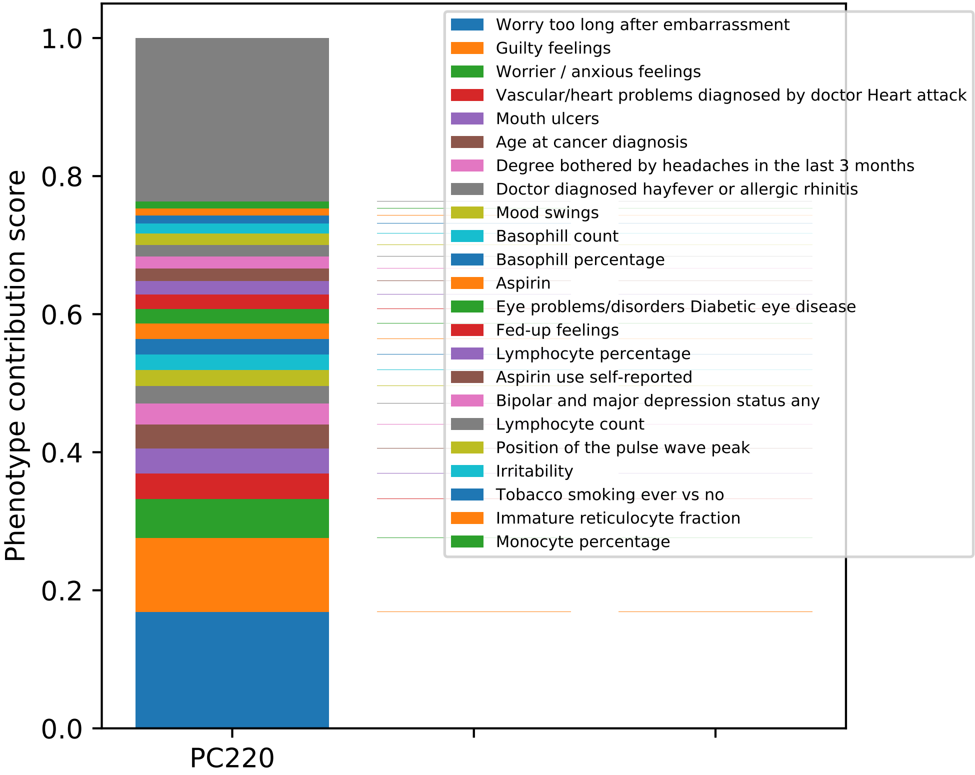
Phenotype contribution score
1.0
0.8
0.6
0.4
0.2
0.0
PC220
Worry too long after embarrassment
Guilty feelings
Worrier / anxious feelings
Vascular/heart problems diagnosed by doctor Heart attack
Mouth ulcers
Age at cancer diagnosis
Degree bothered by headaches in the last 3 months
Doctor diagnosed hayfever or allergic rhinitis
Mood swings
Basophill count
Basophill percentage
Aspirin
Eye problems/disorders Diabetic eye disease
Fed-up feelings
Lymphocyte percentage
Aspirin use self-reported
Bipolar and major depression status any
Lymphocyte count
Position of the pulse wave peak
Irritability
Tobacco smoking ever vs no
Immature reticulocyte fraction
Monocyte percentage

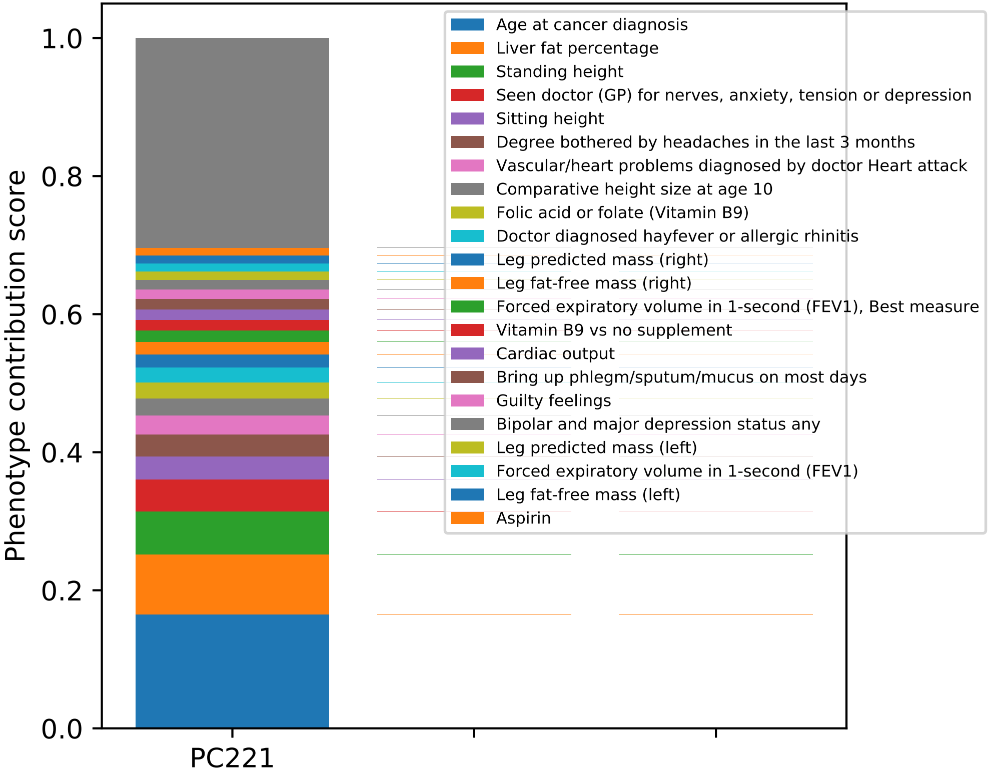

Phenotype contribution score
1.0
0.8
0.6
0.4
0.2
0.0
PC221
Age at cancer diagnosis
Liver fat percentage
Standing height
Seen doctor (GP) for nerves, anxiety, tension or depression
Sitting height
Degree bothered by headaches in the last 3 months
Vascular/heart problems diagnosed by doctor Heart attack
Comparative height size at age 10
Folic acid or folate (Vitamin B9)
Doctor diagnosed hayfever or allergic rhinitis
Leg predicted mass (right)
Leg fat-free mass (right)
Forced expiratory volume in 1-second (FEV1), Best measure
Vitamin B9 vs no supplement
Cardiac output
Bring up phlegm/sputum/mucus on most days
Guilty feelings
Bipolar and major depression status any
Leg predicted mass (left)
Forced expiratory volume in 1-second (FEV1)
Leg fat-free mass (left)
Aspirin

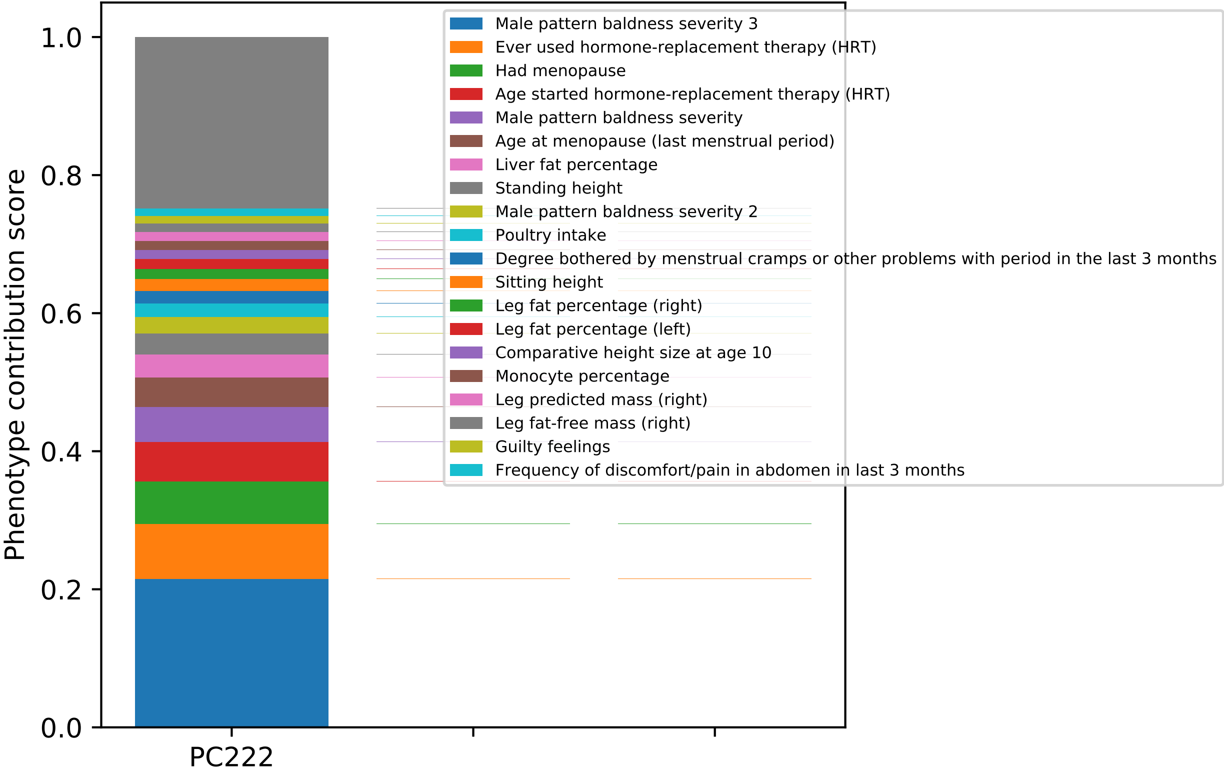

Phenotype contribution score
0.0
0.2
0.4
0.6
0.8
1.0
PC222
Male pattern baldness severity 3
Ever used hormone-replacement therapy (HRT)
Had menopause
Age started hormone-replacement therapy (HRT)
Male pattern baldness severity
Age at menopause (last menstrual period)
Liver fat percentage
Standing height
Male pattern baldness severity 2
Poultry intake
Degree bothered by menstrual cramps or other problems with period in the last 3 months
Sitting height
Leg fat percentage (right)
Leg fat percentage (left)
Comparative height size at age 10
Monocyte percentage
Leg predicted mass (right)
Leg fat-free mass (right)
Guilty feelings
Frequency of discomfort/pain in abdomen in last 3 months

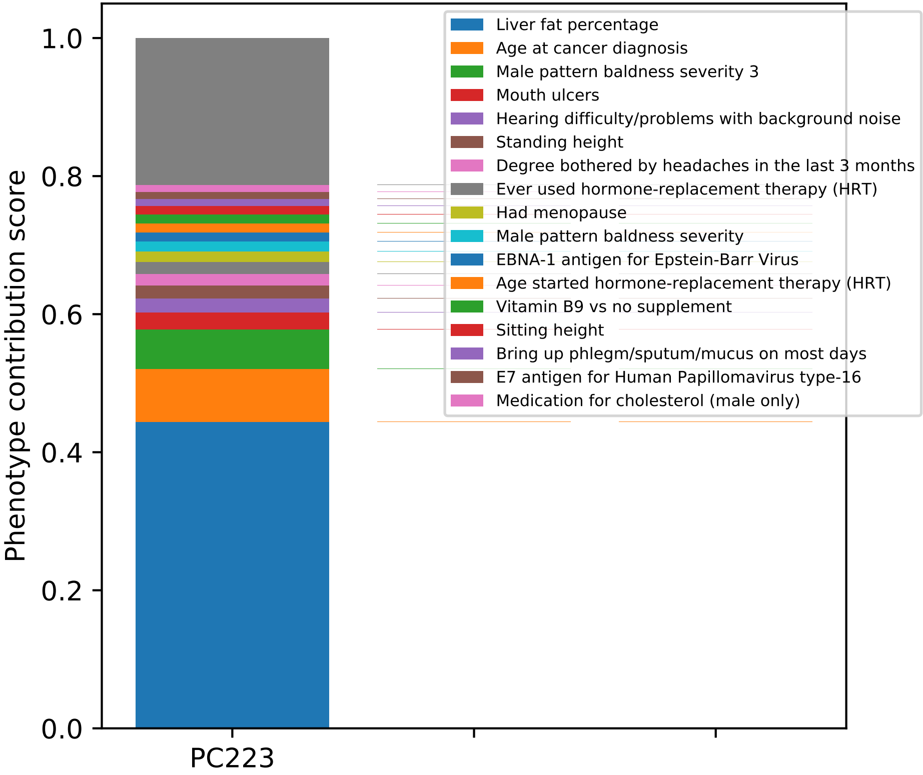

Phenotype contribution score
1.0
0.8
0.6
0.4
0.2
0.0
PC223
Liver fat percentage
Age at cancer diagnosis
Male pattern baldness severity 3
Mouth ulcers
Hearing difficulty/problems with background noise
Standing height
Degree bothered by headaches in the last 3 months
Ever used hormone-replacement therapy (HRT)
Had menopause
Male pattern baldness severity
EBNA-1 antigen for Epstein-Barr Virus
Age started hormone-replacement therapy (HRT)
Vitamin B9 vs no supplement
Sitting height
Bring up phlegm/sputum/mucus on most days
E7 antigen for Human Papillomavirus type-16
Medication for cholesterol (male only)

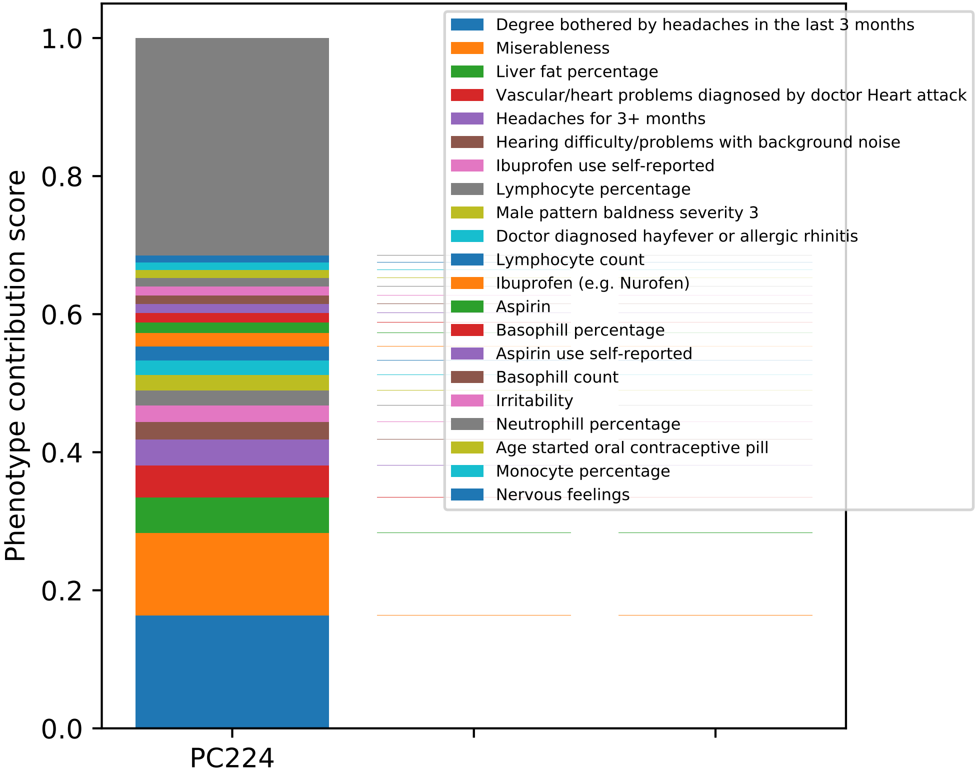

Phenotype contribution score
0.0
0.2
0.4
0.6
0.8
1.0
PC224
Degree bothered by headaches in the last 3 months
Miserableness
Liver fat percentage
Vascular/heart problems diagnosed by doctor Heart attack
Headaches for 3+ months
Hearing difficulty/problems with background noise
Ibuprofen use self-reported
Lymphocyte percentage
Male pattern baldness severity 3
Doctor diagnosed hayfever or allergic rhinitis
Lymphocyte count
Ibuprofen (e.g. Nurofen)
Aspirin
Basophill percentage
Aspirin use self-reported
Basophill count
Irritability
Neutrophill percentage
Age started oral contraceptive pill
Monocyte percentage
Nervous feelings

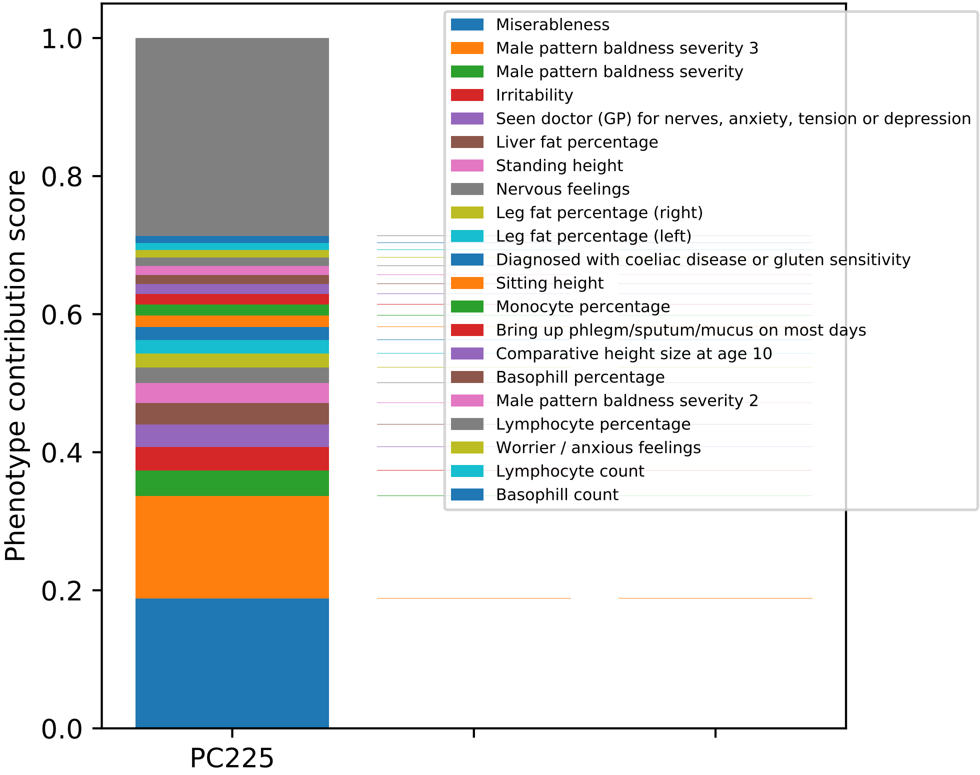

Phenotype contribution score
1.0
0.8
0.6
0.4
0.2
0.0
PC225
Miserableness
Male pattern baldness severity 3
Male pattern baldness severity
Irritability
Seen doctor (GP) for nerves, anxiety, tension or depression
Liver fat percentage
Standing height
Nervous feelings
Leg fat percentage (right)
Leg fat percentage (left)
Diagnosed with coeliac disease or gluten sensitivity
Sitting height
Monocyte percentage
Bring up phlegm/sputum/mucus on most days
Comparative height size at age 10
Basophill percentage
Male pattern baldness severity 2
Lymphocyte percentage
Worrier / anxious feelings
Lymphocyte count
Basophill count

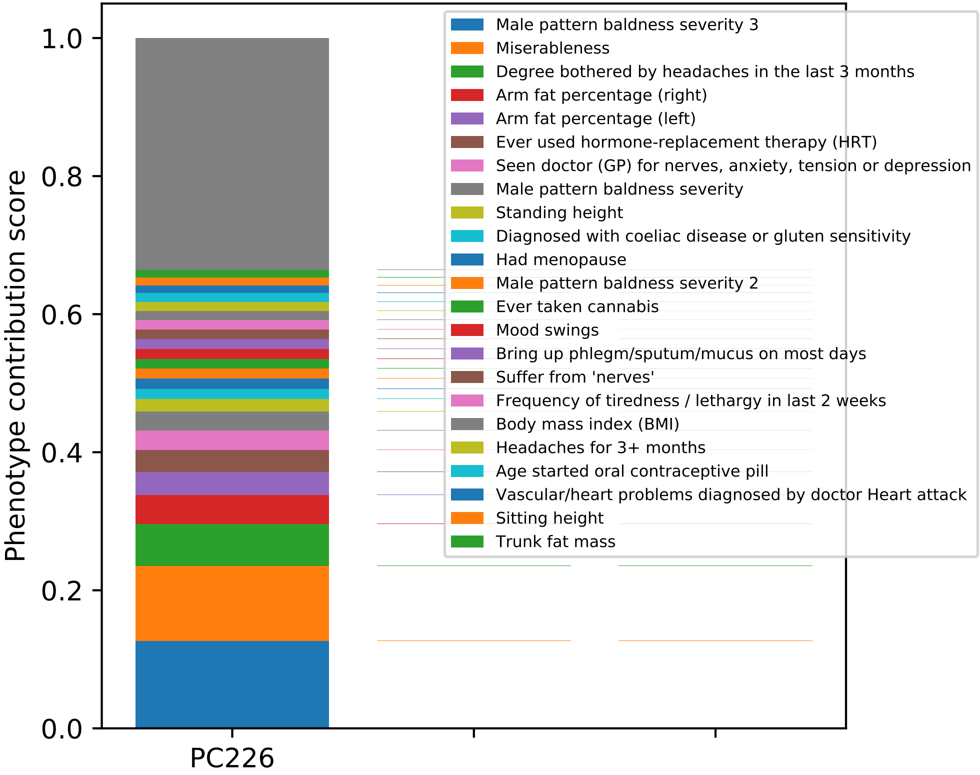

Phenotype contribution score
1.0
0.8
0.6
0.4
0.2
0.0
PC226
Male pattern baldness severity 3
Miserableness
Degree bothered by headaches in the last 3 months
Arm fat percentage (right)
Arm fat percentage (left)
Ever used hormone-replacement therapy (HRT)
Seen doctor (GP) for nerves, anxiety, tension or depression
Male pattern baldness severity
Standing height
Diagnosed with coeliac disease or gluten sensitivity
Had menopause
Male pattern baldness severity 2
Ever taken cannabis
Mood swings
Bring up phlegm/sputum/mucus on most days
Suffer from 'nerves'
Frequency of tiredness / lethargy in last 2 weeks
Body mass index (BMI)
Headaches for 3+ months
Age started oral contraceptive pill
Vascular/heart problems diagnosed by doctor Heart attack
Sitting height
Trunk fat mass

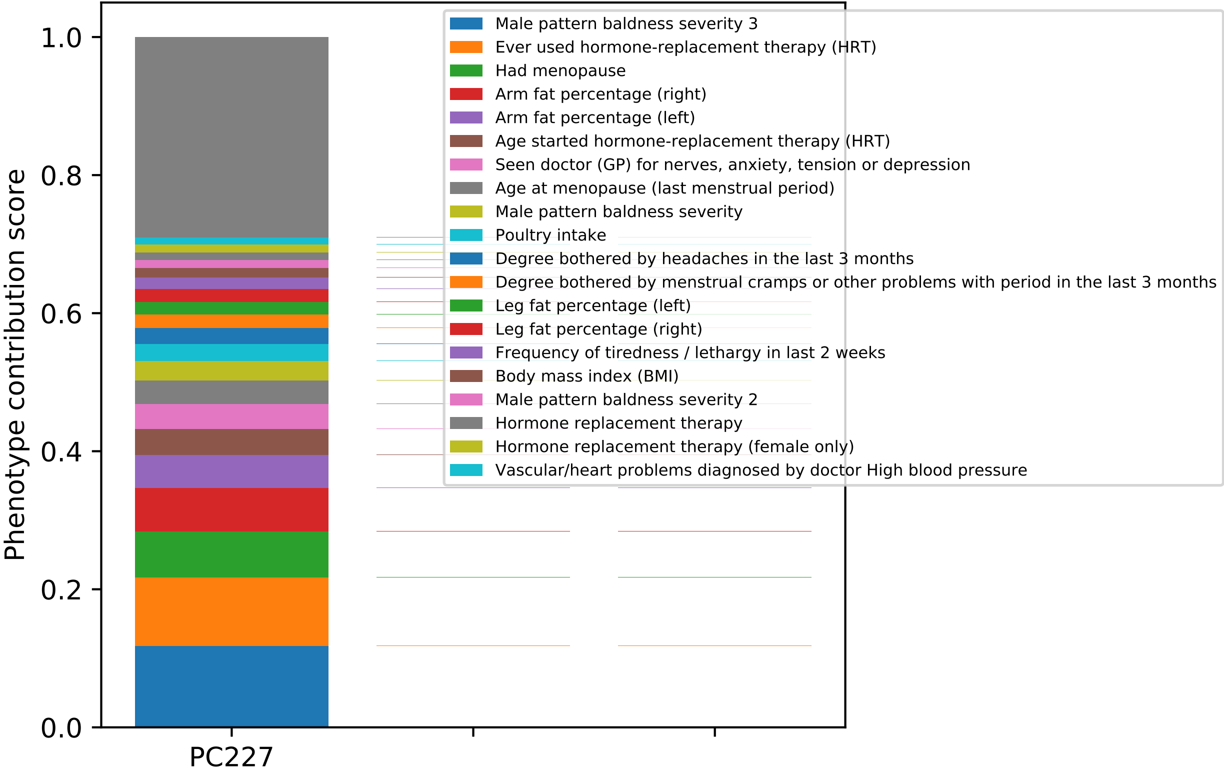

1.0
0.8
0.6
0.4
0.2
0.0
Phenotype contribution score
PC227
Male pattern baldness severity 3
Ever used hormone-replacement therapy (HRT)
Had menopause
Arm fat percentage (right)
Arm fat percentage (left)
Age started hormone-replacement therapy (HRT)
Seen doctor (GP) for nerves, anxiety, tension or depression
Age at menopause (last menstrual period)
Male pattern baldness severity
Poultry intake
Degree bothered by headaches in the last 3 months
Degree bothered by menstrual cramps or other problems with period in the last 3 months
Leg fat percentage (left)
Leg fat percentage (right)
Frequency of tiredness / lethargy in last 2 weeks
Body mass index (BMI)
Male pattern baldness severity 2
Hormone replacement therapy
Hormone replacement therapy (female only)
Vascular/heart problems diagnosed by doctor High blood pressure

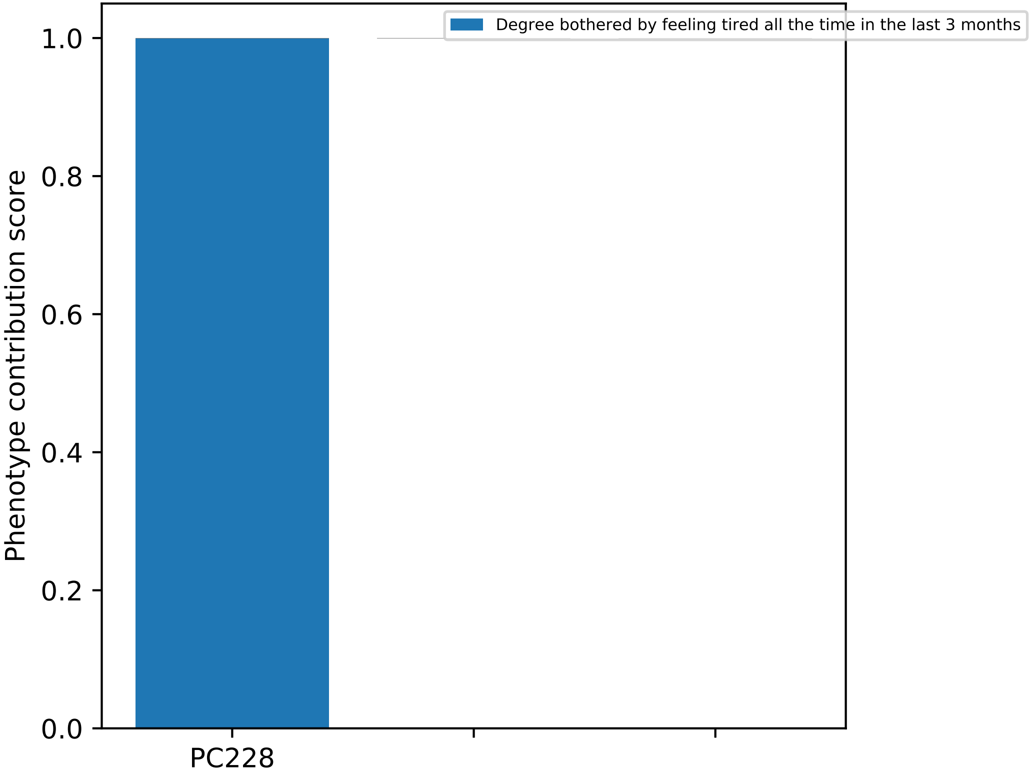

Degree bothered by feeling tired all the time in the last 3 months
Phenotype contribution score
1.0
0.8
0.6
0.4
0.2
0.0
PC228

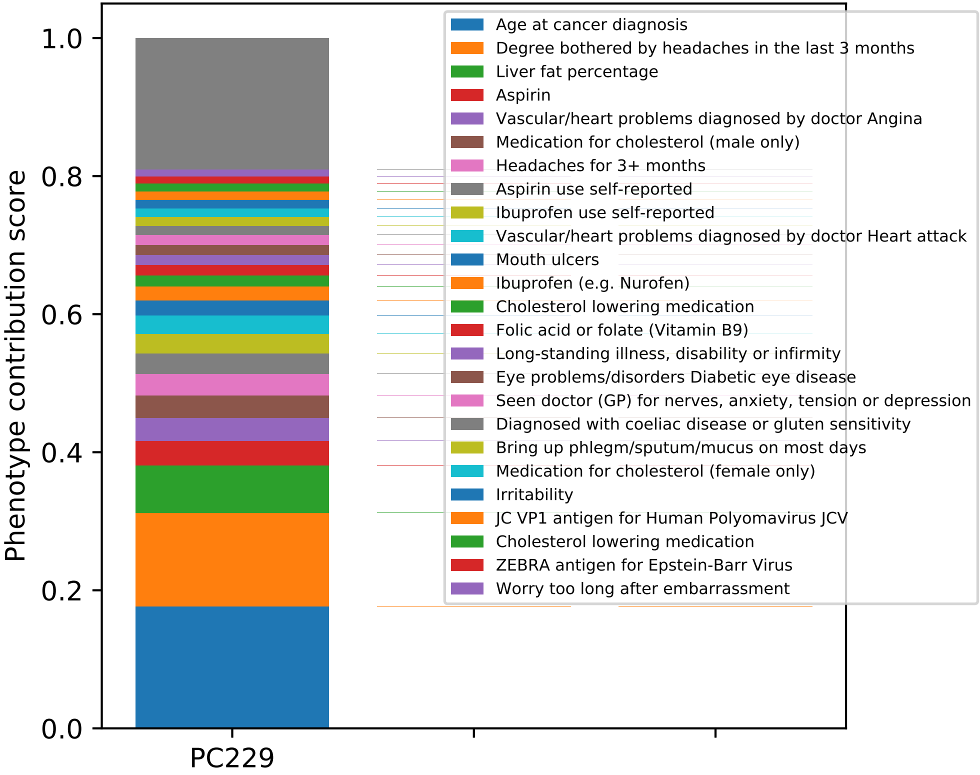

Phenotype contribution score
1.0
0.8
0.6
0.4
0.2
0.0
PC229
Age at cancer diagnosis
Degree bothered by headaches in the last 3 months
Liver fat percentage
Aspirin
Vascular/heart problems diagnosed by doctor Angina
Medication for cholesterol (male only)
Headaches for 3+ months
Aspirin use self-reported
Ibuprofen use self-reported
Vascular/heart problems diagnosed by doctor Heart attack
Mouth ulcers
Ibuprofen (e.g. Nurofen)
Cholesterol lowering medication
Folic acid or folate (Vitamin B9)
Long-standing illness, disability or infirmity
Eye problems/disorders Diabetic eye disease
Seen doctor (GP) for nerves, anxiety, tension or depression
Diagnosed with coeliac disease or gluten sensitivity
Bring up phlegm/sputum/mucus on most days
Medication for cholesterol (female only)
Irritability
JC VP1 antigen for Human Polyomavirus JCV
Cholesterol lowering medication
ZEBRA antigen for Epstein-Barr Virus
Worry too long after embarrassment

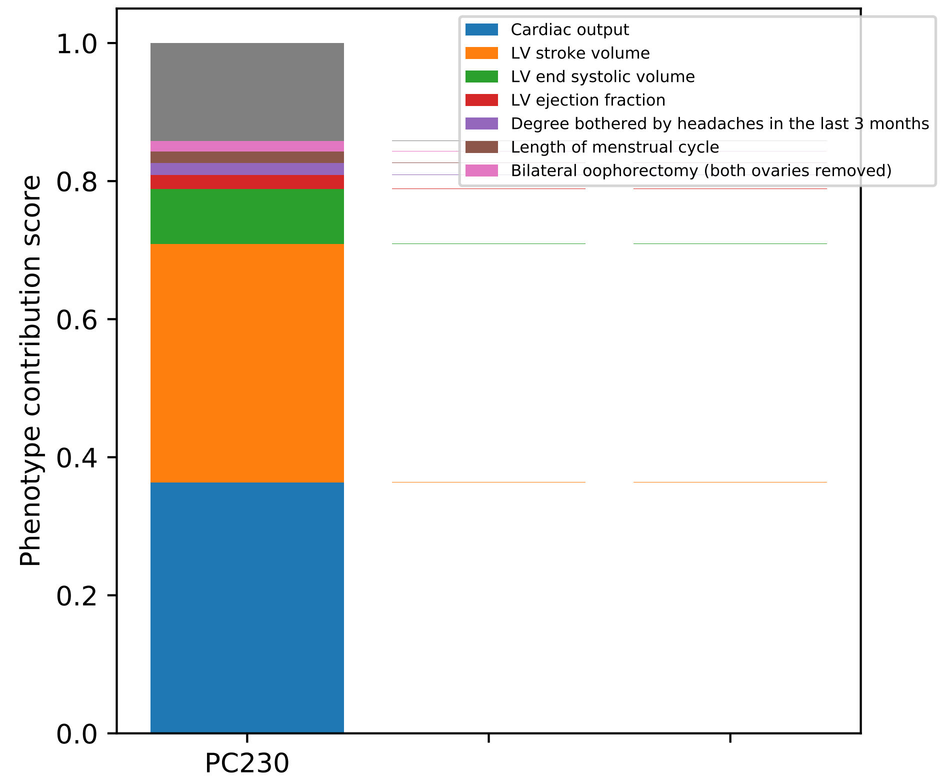

Phenotype contribution score
1.0
0.8
0.6
0.4
0.2
0.0
PC230
Cardiac output
LV stroke volume
LV end systolic volume
LV ejection fraction
Degree bothered by headaches in the last 3 months
Length of menstrual cycle
Bilateral oophorectomy (both ovaries removed)

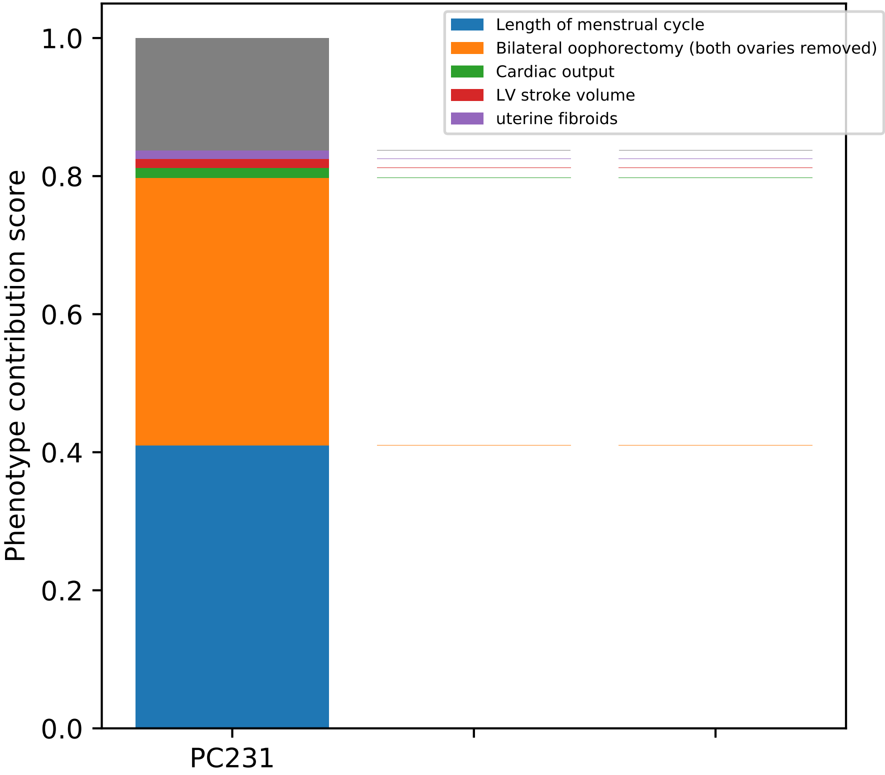

Length of menstrual cycle
Bilateral oophorectomy (both ovaries removed)
Cardiac output
LV stroke volume
uterine fibroids
Phenotype contribution score
1.0
0.8
0.6
0.4
0.2
0.0
PC231

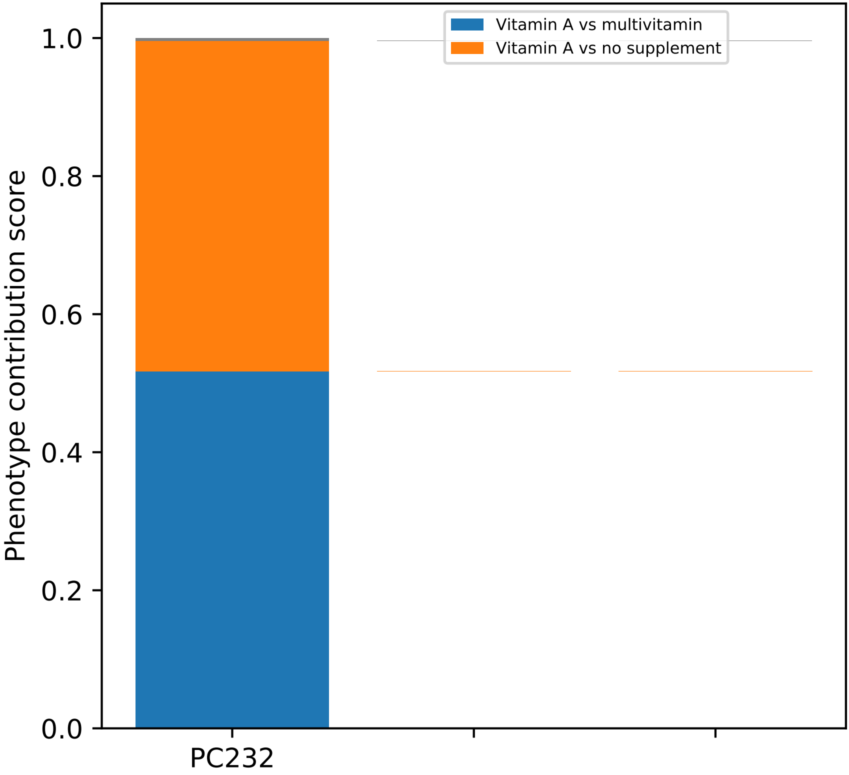

Vitamin A vs multivitamin
Vitamin A vs no supplement
Phenotype contribution score
0.0
0.2
0.4
0.6
0.8
1.0
PC232

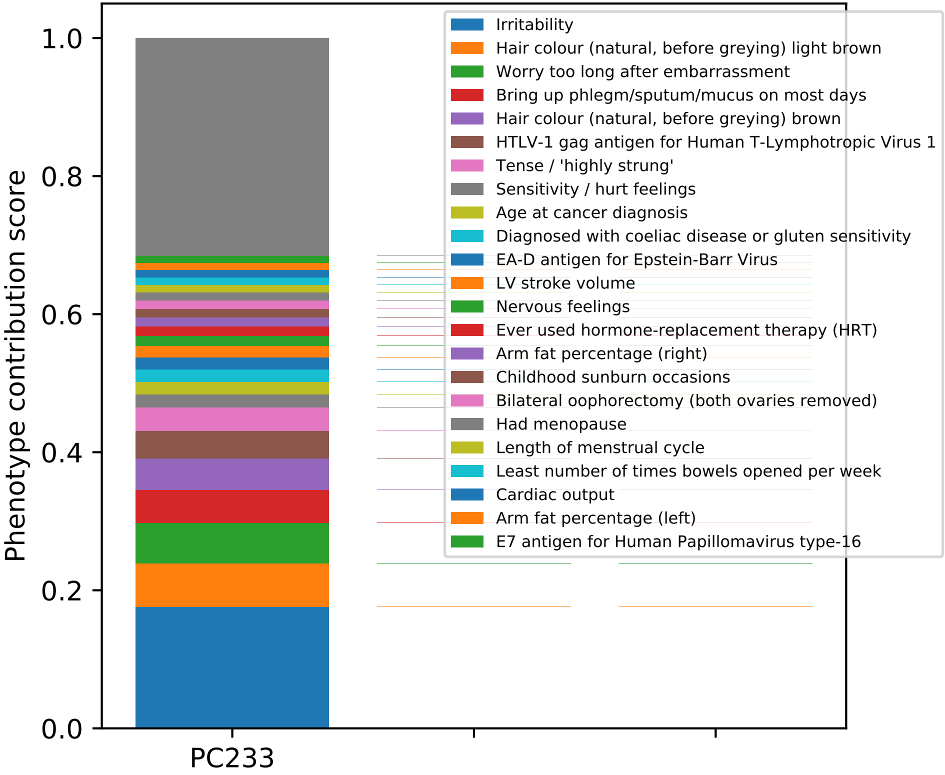

Phenotype contribution score
1.0
0.8
0.6
0.4
0.2
0.0
PC233
Irritability
Hair colour (natural, before greying) light brown
Worry too long after embarrassment
Bring up phlegm/sputum/mucus on most days
Hair colour (natural, before greying) brown
HTLV-1 gag antigen for Human T-Lymphotropic Virus 1
Tense / 'highly strung'
Sensitivity / hurt feelings
Age at cancer diagnosis
Diagnosed with coeliac disease or gluten sensitivity
EA-D antigen for Epstein-Barr Virus
LV stroke volume
Nervous feelings
Ever used hormone-replacement therapy (HRT)
Arm fat percentage (right)
Childhood sunburn occasions
Bilateral oophorectomy (both ovaries removed)
Had menopause
Length of menstrual cycle
Least number of times bowels opened per week
Cardiac output
Arm fat percentage (left)
E7 antigen for Human Papillomavirus type-16

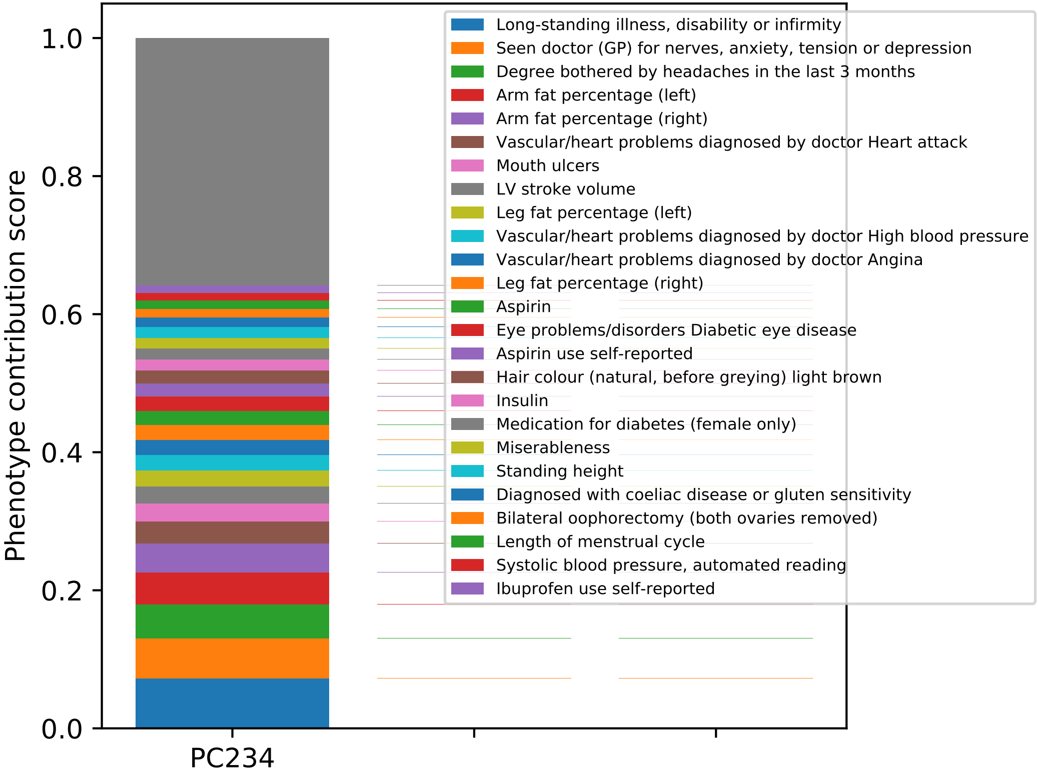
Phenotype contribution score
1.0
0.8
0.6
0.4
0.2
0.0
PC234
Long-standing illness, disability or infirmity
Seen doctor (GP) for nerves, anxiety, tension or depression
Degree bothered by headaches in the last 3 months
Arm fat percentage (left)
Arm fat percentage (right)
Vascular/heart problems diagnosed by doctor Heart attack
Mouth ulcers
LV stroke volume
Leg fat percentage (left)
Vascular/heart problems diagnosed by doctor High blood pressure
Vascular/heart problems diagnosed by doctor Angina
Leg fat percentage (right)
Aspirin
Eye problems/disorders Diabetic eye disease
Aspirin use self-reported
Hair colour (natural, before greying) light brown
Insulin
Medication for diabetes (female only)
Miserableness
Standing height
Diagnosed with coeliac disease or gluten sensitivity
Bilateral oophorectomy (both ovaries removed)
Length of menstrual cycle
Systolic blood pressure, automated reading
Ibuprofen use self-reported

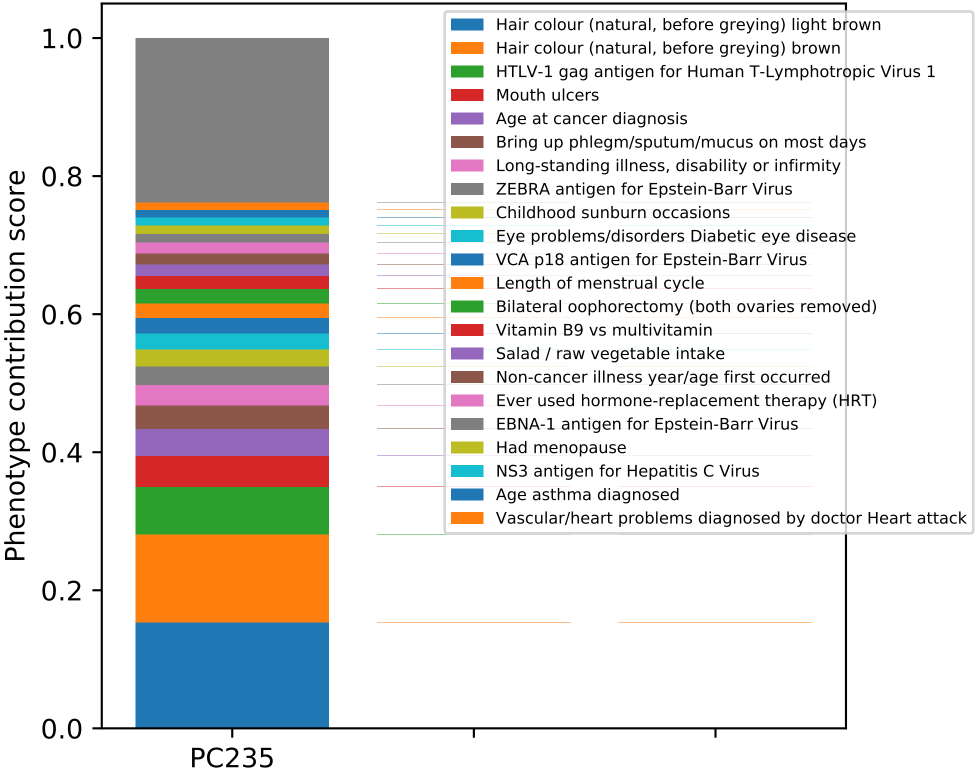

Phenotype contribution score
0.0
0.2
0.4
0.6
0.8
1.0
PC235
Hair colour (natural, before greying) light brown
Hair colour (natural, before greying) brown
HTLV-1 gag antigen for Human T-Lymphotropic Virus 1
Mouth ulcers
Age at cancer diagnosis
Bring up phlegm/sputum/mucus on most days
Long-standing illness, disability or infirmity
ZEBRA antigen for Epstein-Barr Virus
Childhood sunburn occasions
Eye problems/disorders Diabetic eye disease
VCA p18 antigen for Epstein-Barr Virus
Length of menstrual cycle
Bilateral oophorectomy (both ovaries removed)
Vitamin B9 vs multivitamin
Salad / raw vegetable intake
Non-cancer illness year/age first occurred
Ever used hormone-replacement therapy (HRT)
EBNA-1 antigen for Epstein-Barr Virus
Had menopause
NS3 antigen for Hepatitis C Virus
Age asthma diagnosed
Vascular/heart problems diagnosed by doctor Heart attack

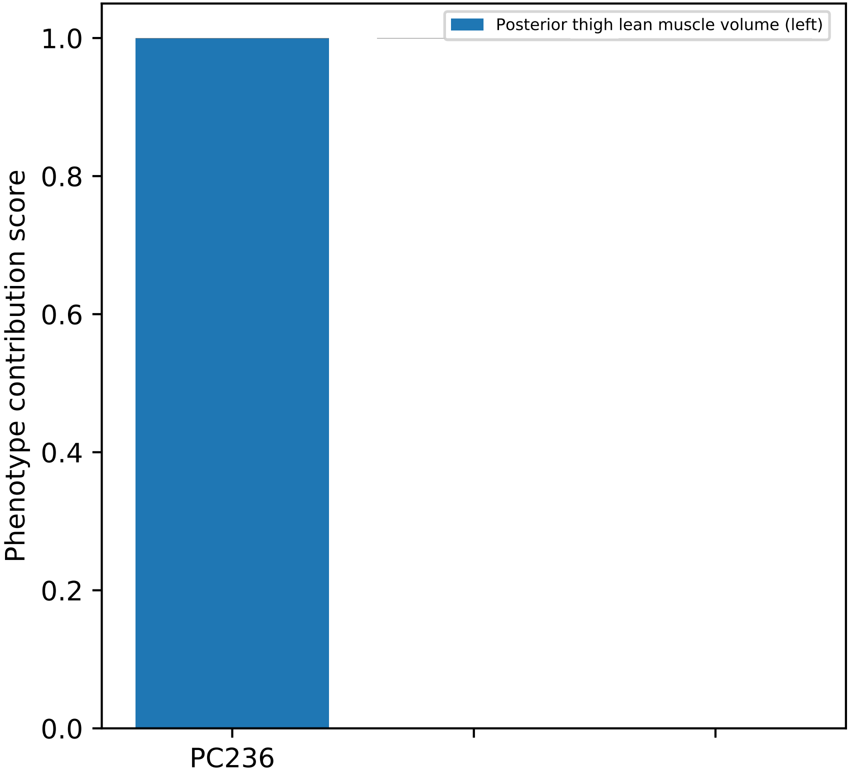
Posterior thigh lean muscle volume (left)
Phenotype contribution score
1.0
0.8
0.6
0.4
0.2
0.0
PC236

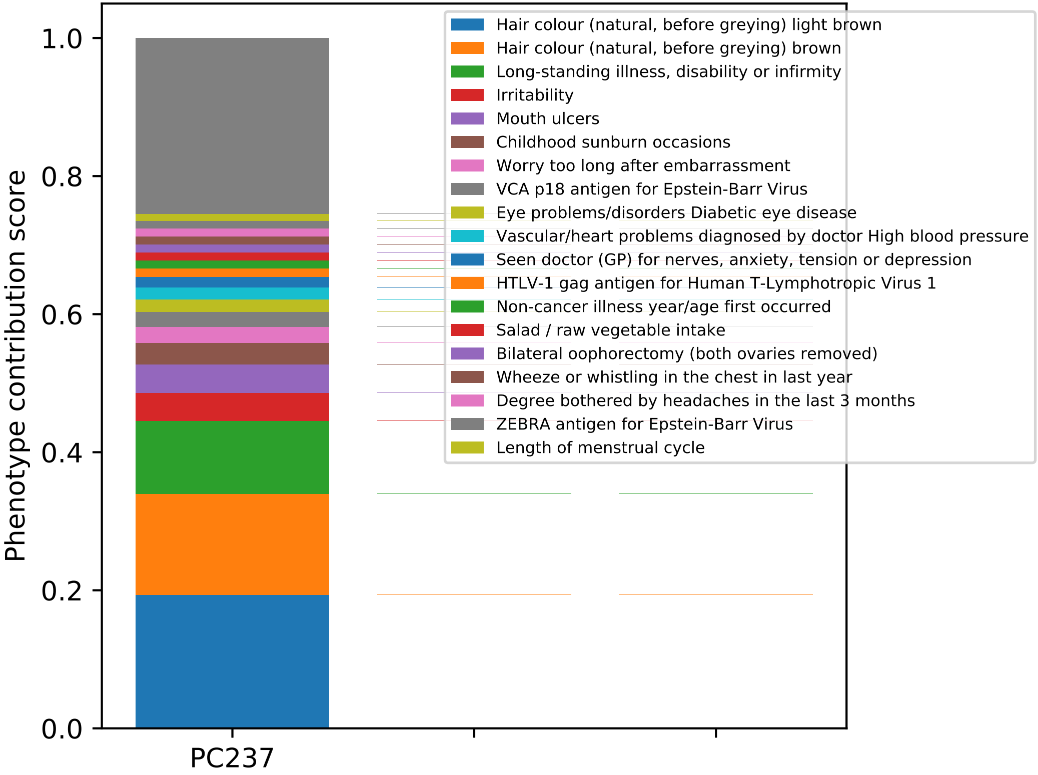

Phenotype contribution score
0.0
0.2
0.4
0.6
0.8
1.0
PC237
Hair colour (natural, before greying) light brown
Hair colour (natural, before greying) brown
Long-standing illness, disability or infirmity
Irritability
Mouth ulcers
Childhood sunburn occasions
Worry too long after embarrassment
VCA p18 antigen for Epstein-Barr Virus
Eye problems/disorders Diabetic eye disease
Vascular/heart problems diagnosed by doctor High blood pressure
Seen doctor (GP) for nerves, anxiety, tension or depression
HTLV-1 gag antigen for Human T-Lymphotropic Virus 1
Non-cancer illness year/age first occurred
Salad / raw vegetable intake
Bilateral oophorectomy (both ovaries removed)
Wheeze or whistling in the chest in last year
Degree bothered by headaches in the last 3 months
ZEBRA antigen for Epstein-Barr Virus
Length of menstrual cycle

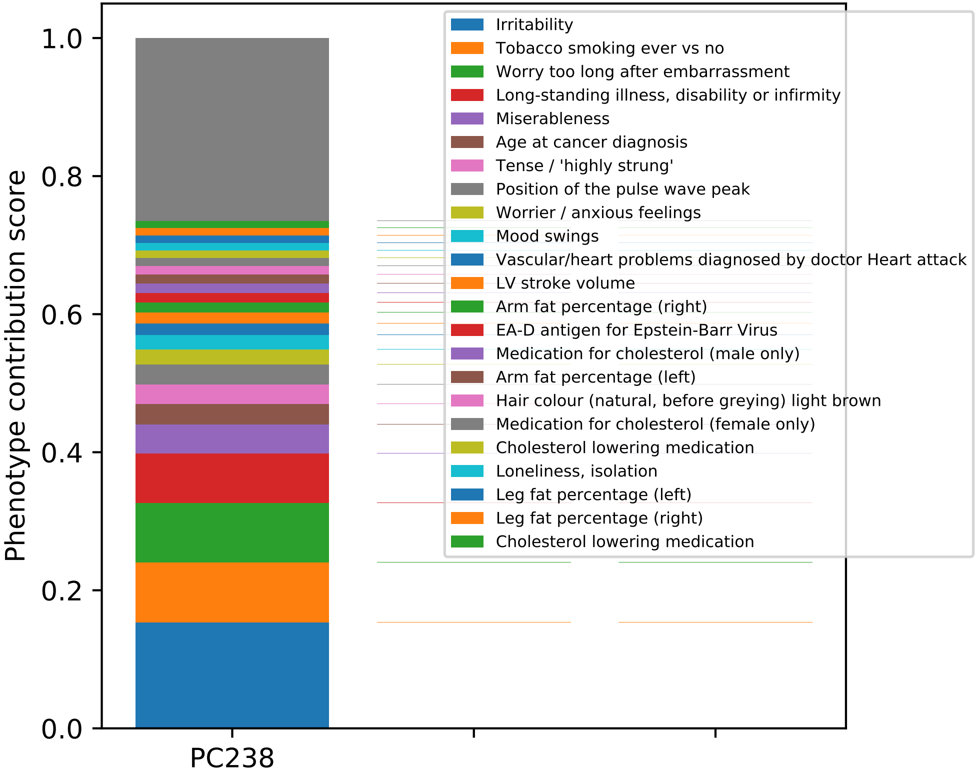

Irritability
Tobacco smoking ever vs no
Worry too long after embarrassment
Long-standing illness, disability or infirmity
Miserableness
Age at cancer diagnosis
Tense / 'highly strung'
Position of the pulse wave peak
Worrier / anxious feelings
Mood swings
Vascular/heart problems diagnosed by doctor Heart attack
LV stroke volume
Arm fat percentage (right)
EA-D antigen for Epstein-Barr Virus
Medication for cholesterol (male only)
Arm fat percentage (left)
Hair colour (natural, before greying) light brown
Medication for cholesterol (female only)
Cholesterol lowering medication
Loneliness, isolation
Leg fat percentage (left)
Leg fat percentage (right)
Cholesterol lowering medication
Phenotype contribution score
1.0
0.8
0.6
0.4
0.2
0.0
PC238

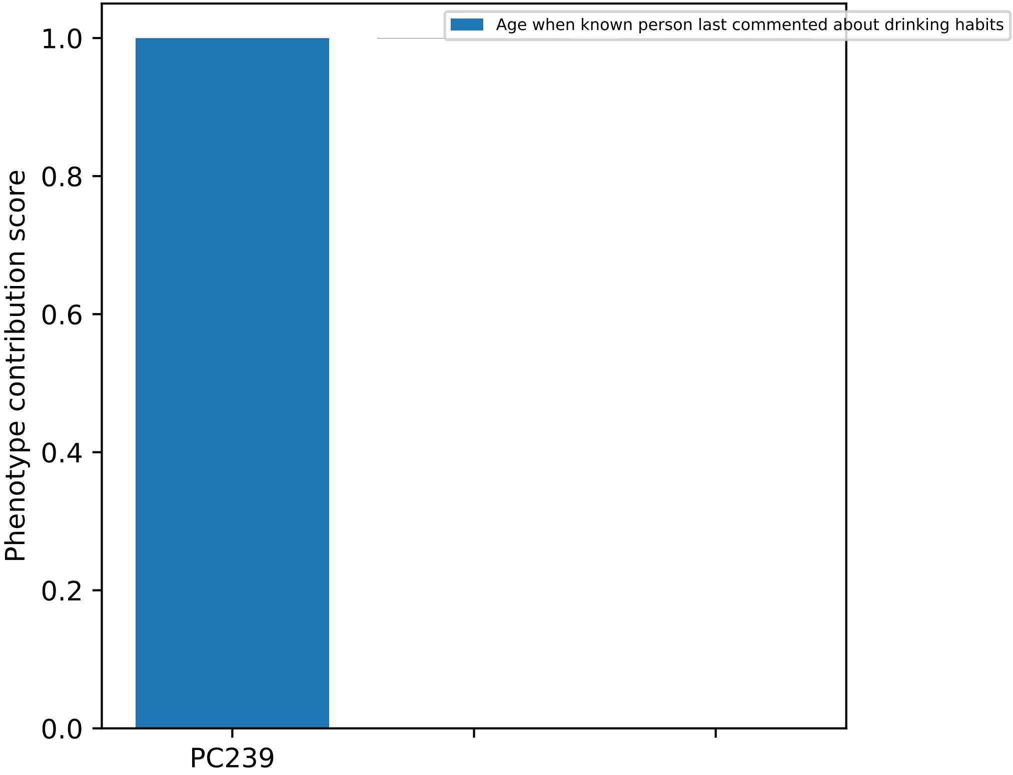
Age when known person last commented about drinking habits
1.0
0.8
0.6
0.4
0.2
0.0
Phenotype contribution score
PC239

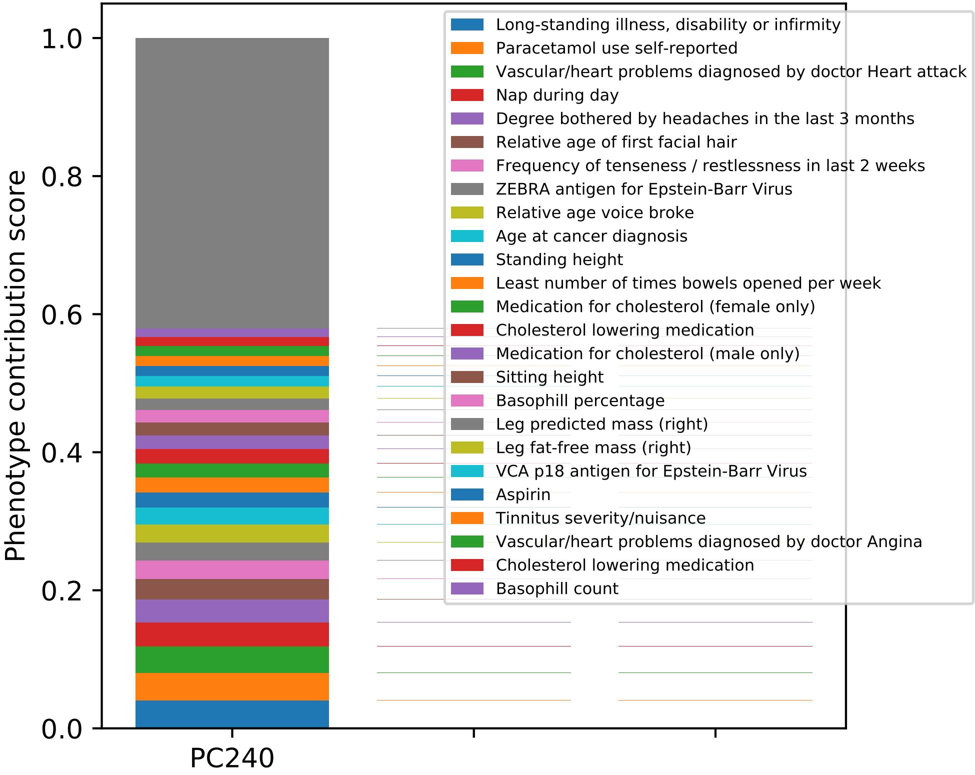

Phenotype contribution score
0.0
0.2
0.4
0.6
0.8
1.0
PC240
Long-standing illness, disability or infirmity
Paracetamol use self-reported
Vascular/heart problems diagnosed by doctor Heart attack
Nap during day
Degree bothered by headaches in the last 3 months
Relative age of first facial hair
Frequency of tenseness / restlessness in last 2 weeks
ZEBRA antigen for Epstein-Barr Virus
Relative age voice broke
Age at cancer diagnosis
Standing height
Least number of times bowels opened per week
Medication for cholesterol (female only)
Cholesterol lowering medication
Medication for cholesterol (male only)
Sitting height
Basophill percentage
Leg predicted mass (right)
Leg fat-free mass (right)
VCA p18 antigen for Epstein-Barr Virus
Aspirin
Tinnitus severity/nuisance
Vascular/heart problems diagnosed by doctor Angina
Cholesterol lowering medication
Basophill count

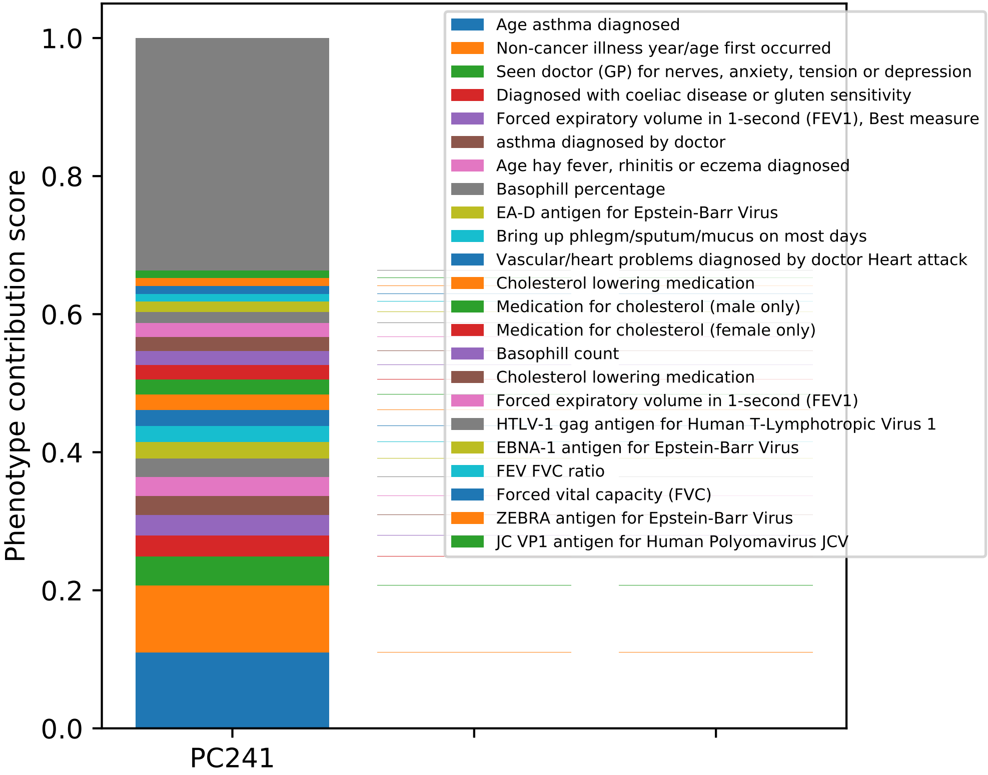

Phenotype contribution score
1.0
0.8
0.6
0.4
0.2
0.0
PC241
Age asthma diagnosed
Non-cancer illness year/age first occurred
Seen doctor (GP) for nerves, anxiety, tension or depression
Diagnosed with coeliac disease or gluten sensitivity
Forced expiratory volume in 1-second (FEV1), Best measure
asthma diagnosed by doctor
Age hay fever, rhinitis or eczema diagnosed
Basophill percentage
EA-D antigen for Epstein-Barr Virus
Bring up phlegm/sputum/mucus on most days
Vascular/heart problems diagnosed by doctor Heart attack
Cholesterol lowering medication
Medication for cholesterol (male only)
Medication for cholesterol (female only)
Basophill count
Cholesterol lowering medication
Forced expiratory volume in 1-second (FEV1)
HTLV-1 gag antigen for Human T-Lymphotropic Virus 1
EBNA-1 antigen for Epstein-Barr Virus
FEV FVC ratio
Forced vital capacity (FVC)
ZEBRA antigen for Epstein-Barr Virus
JC VP1 antigen for Human Polyomavirus JCV

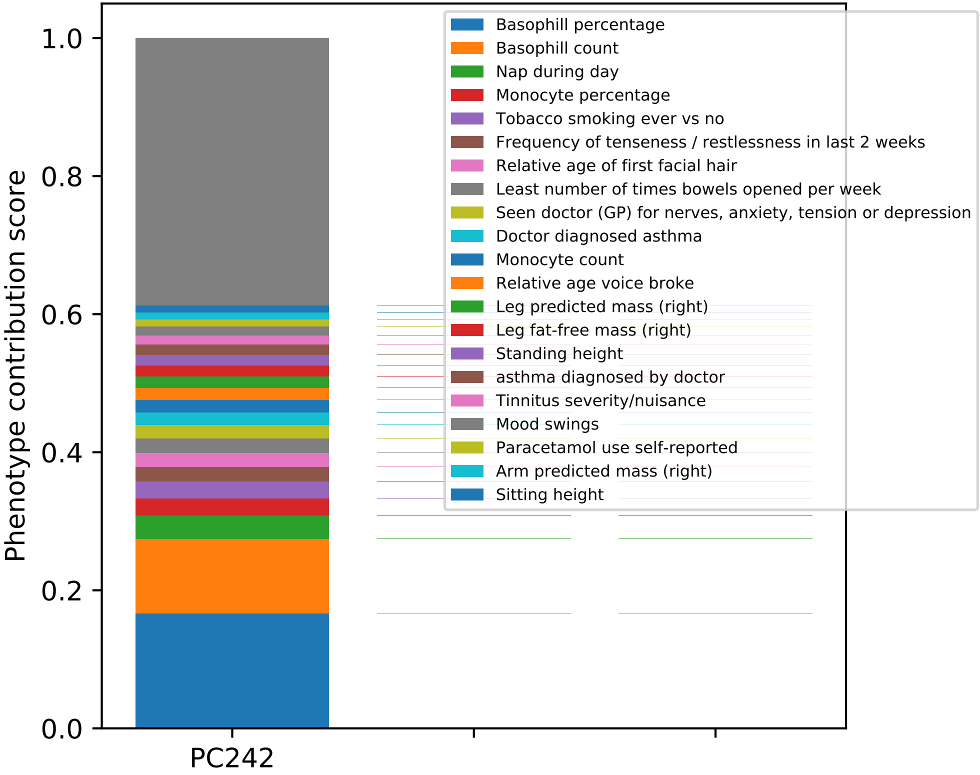
Phenotype contribution score
1.0
0.8
0.6
0.4
0.2
0.0
PC242
Basophill percentage
Basophill count
Nap during day
Monocyte percentage
Tobacco smoking ever vs no
Frequency of tenseness / restlessness in last 2 weeks
Relative age of first facial hair
Least number of times bowels opened per week
Seen doctor (GP) for nerves, anxiety, tension or depression
Doctor diagnosed asthma
Monocyte count
Relative age voice broke
Leg predicted mass (right)
Leg fat-free mass (right)
Standing height
asthma diagnosed by doctor
Tinnitus severity/nuisance
Mood swings
Paracetamol use self-reported
Arm predicted mass (right)
Sitting height

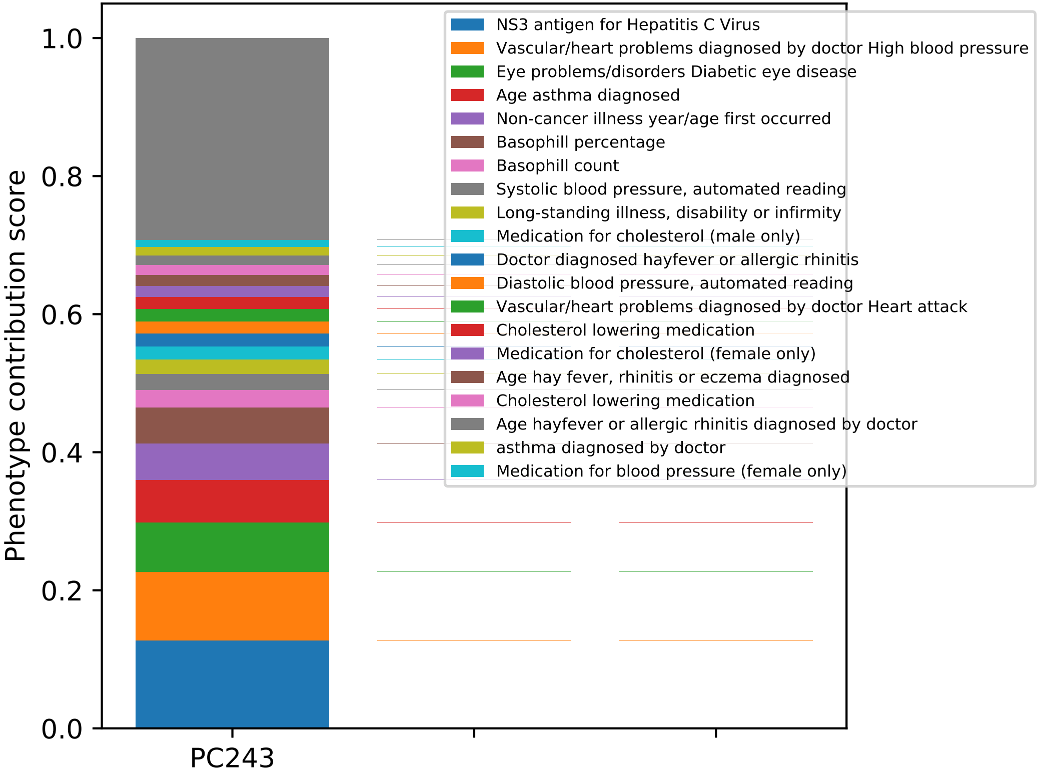
Phenotype contribution score
1.0
0.8
0.6
0.4
0.2
0.0
PC243
NS3 antigen for Hepatitis C Virus
Vascular/heart problems diagnosed by doctor High blood pressure
Eye problems/disorders Diabetic eye disease
Age asthma diagnosed
Non-cancer illness year/age first occurred
Basophill percentage
Basophill count
Systolic blood pressure, automated reading
Long-standing illness, disability or infirmity
Medication for cholesterol (male only)
Doctor diagnosed hayfever or allergic rhinitis
Diastolic blood pressure, automated reading
Vascular/heart problems diagnosed by doctor Heart attack
Cholesterol lowering medication
Medication for cholesterol (female only)
Age hay fever, rhinitis or eczema diagnosed
Cholesterol lowering medication
Age hayfever or allergic rhinitis diagnosed by doctor
asthma diagnosed by doctor
Medication for blood pressure (female only)

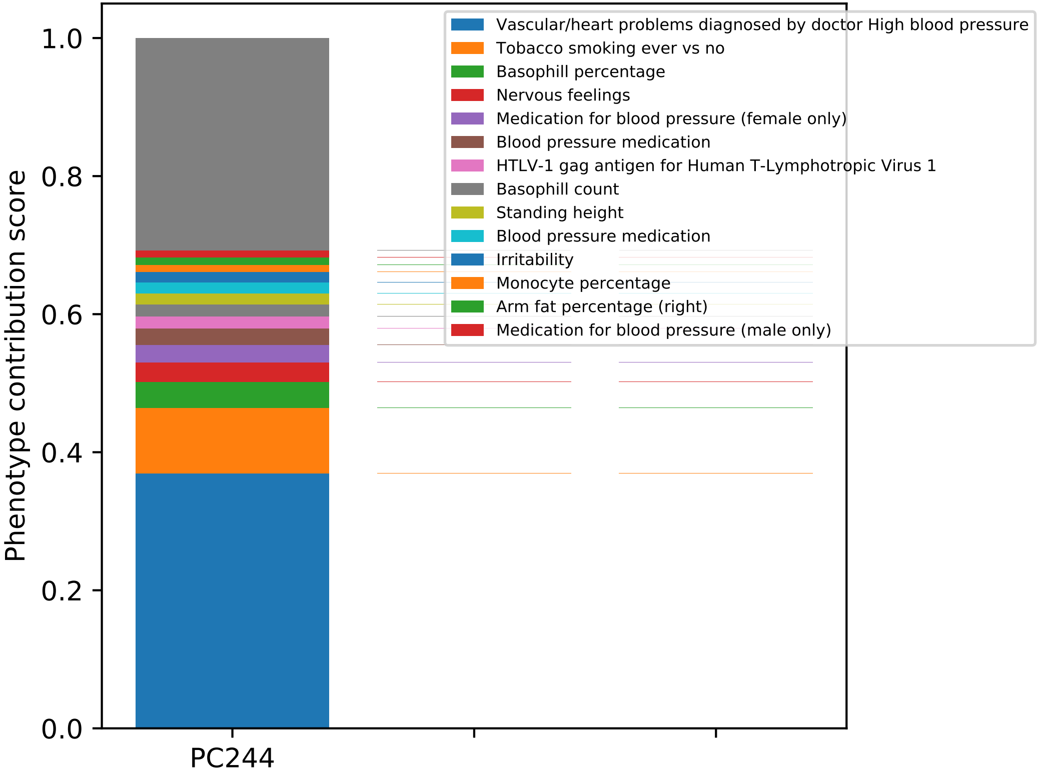

Phenotype contribution score
0.0
0.2
0.4
0.6
0.8
1.0
PC244
Vascular/heart problems diagnosed by doctor High blood pressure
Tobacco smoking ever vs no
Basophill percentage
Nervous feelings
Medication for blood pressure (female only)
Blood pressure medication
HTLV-1 gag antigen for Human T-Lymphotropic Virus 1
Basophill count
Standing height
Blood pressure medication
Irritability
Monocyte percentage
Arm fat percentage (right)
Medication for blood pressure (male only)

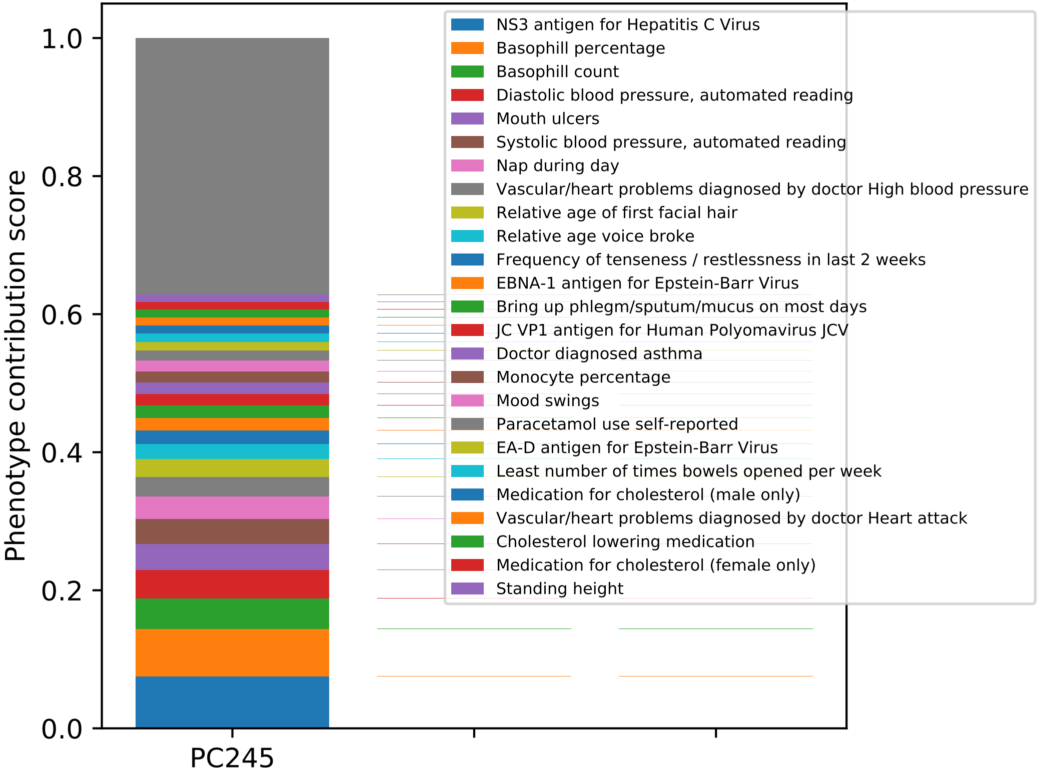

Phenotype contribution score
1.0
0.8
0.6
0.4
0.2
0.0
PC245
NS3 antigen for Hepatitis C Virus
Basophill percentage
Basophill count
Diastolic blood pressure, automated reading
Mouth ulcers
Systolic blood pressure, automated reading
Nap during day
Vascular/heart problems diagnosed by doctor High blood pressure
Relative age of first facial hair
Relative age voice broke
Frequency of tenseness / restlessness in last 2 weeks
EBNA-1 antigen for Epstein-Barr Virus
Bring up phlegm/sputum/mucus on most days
JC VP1 antigen for Human Polyomavirus JCV
Doctor diagnosed asthma
Monocyte percentage
Mood swings
Paracetamol use self-reported
EA-D antigen for Epstein-Barr Virus
Least number of times bowels opened per week
Medication for cholesterol (male only)
Vascular/heart problems diagnosed by doctor Heart attack
Cholesterol lowering medication
Medication for cholesterol (female only)
Standing height

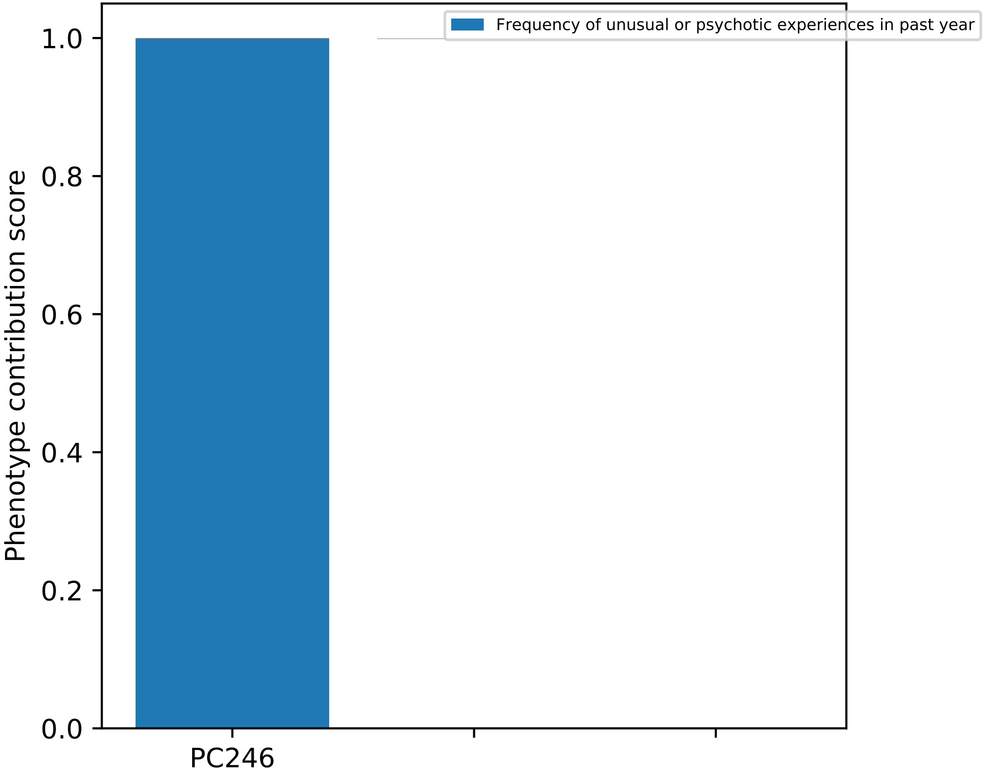
Frequency of unusual or psychotic experiences in past year
1.0
0.8
0.6
0.4
0.2
0.0
Phenotype contribution score
PC246

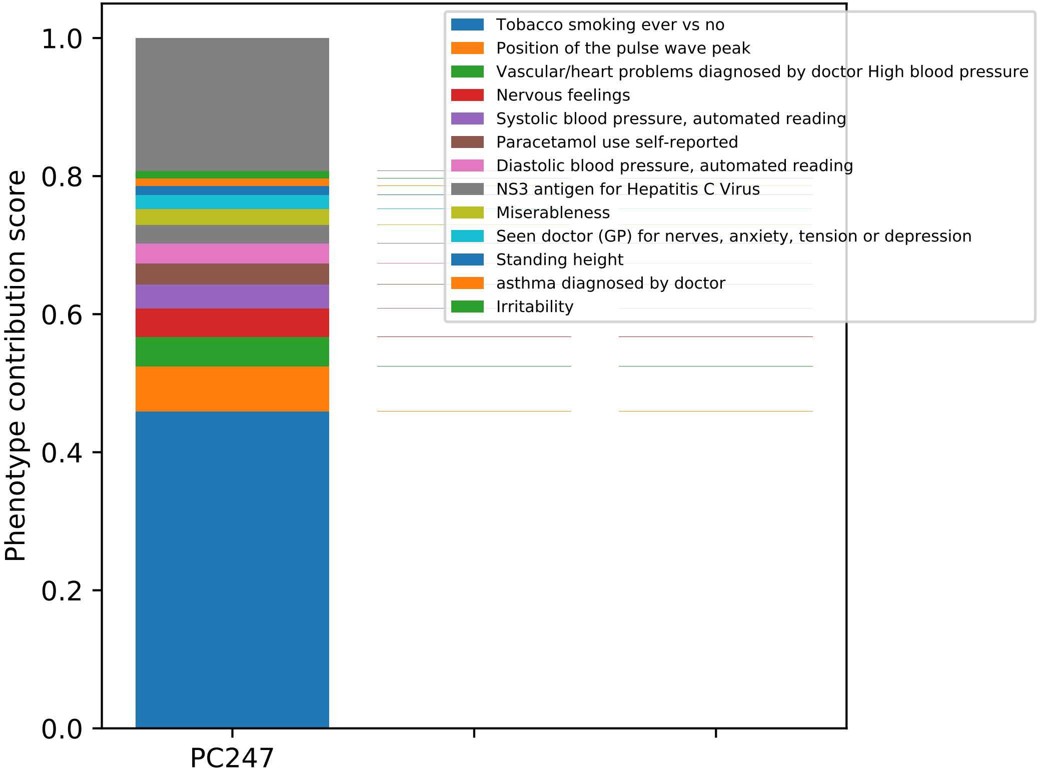

Phenotype contribution score
1.0
0.8
0.6
0.4
0.2
0.0
PC247
Tobacco smoking ever vs no
Position of the pulse wave peak
Vascular/heart problems diagnosed by doctor High blood pressure
Nervous feelings
Systolic blood pressure, automated reading
Paracetamol use self-reported
Diastolic blood pressure, automated reading
NS3 antigen for Hepatitis C Virus
Miserableness
Seen doctor (GP) for nerves, anxiety, tension or depression
Standing height
asthma diagnosed by doctor
Irritability

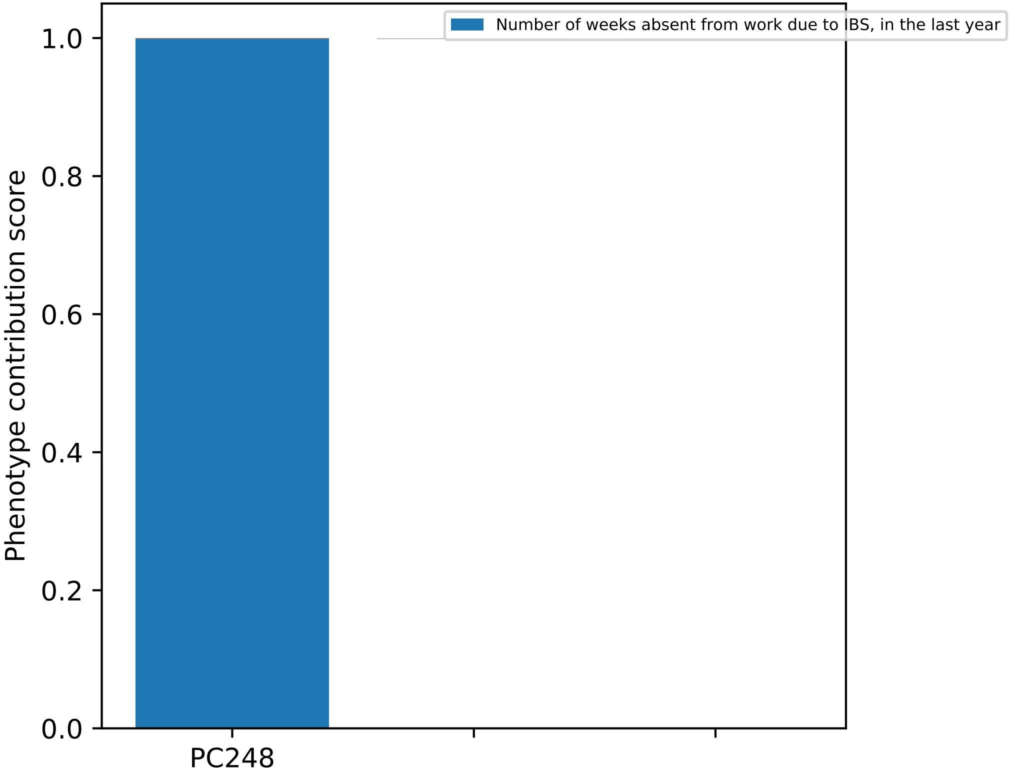

Number of weeks absent from work due to IBS, in the last year
Phenotype contribution score
1.0
0.8
0.6
0.4
0.2
0.0
PC248

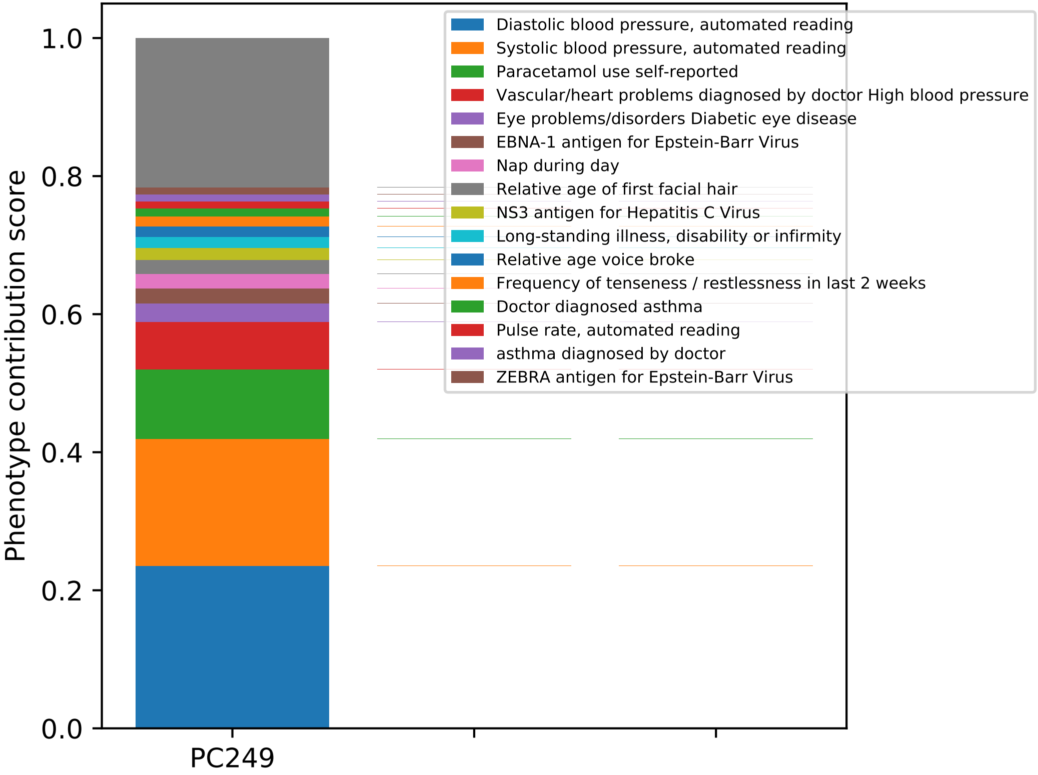
Phenotype contribution score
1.0
0.8
0.6
0.4
0.2
0.0
PC249
Diastolic blood pressure, automated reading
Systolic blood pressure, automated reading
Paracetamol use self-reported
Vascular/heart problems diagnosed by doctor High blood pressure
Eye problems/disorders Diabetic eye disease
EBNA-1 antigen for Epstein-Barr Virus
Nap during day
Relative age of first facial hair
NS3 antigen for Hepatitis C Virus
Long-standing illness, disability or infirmity
Relative age voice broke
Frequency of tenseness / restlessness in last 2 weeks
Doctor diagnosed asthma
Pulse rate, automated reading
asthma diagnosed by doctor
ZEBRA antigen for Epstein-Barr Virus

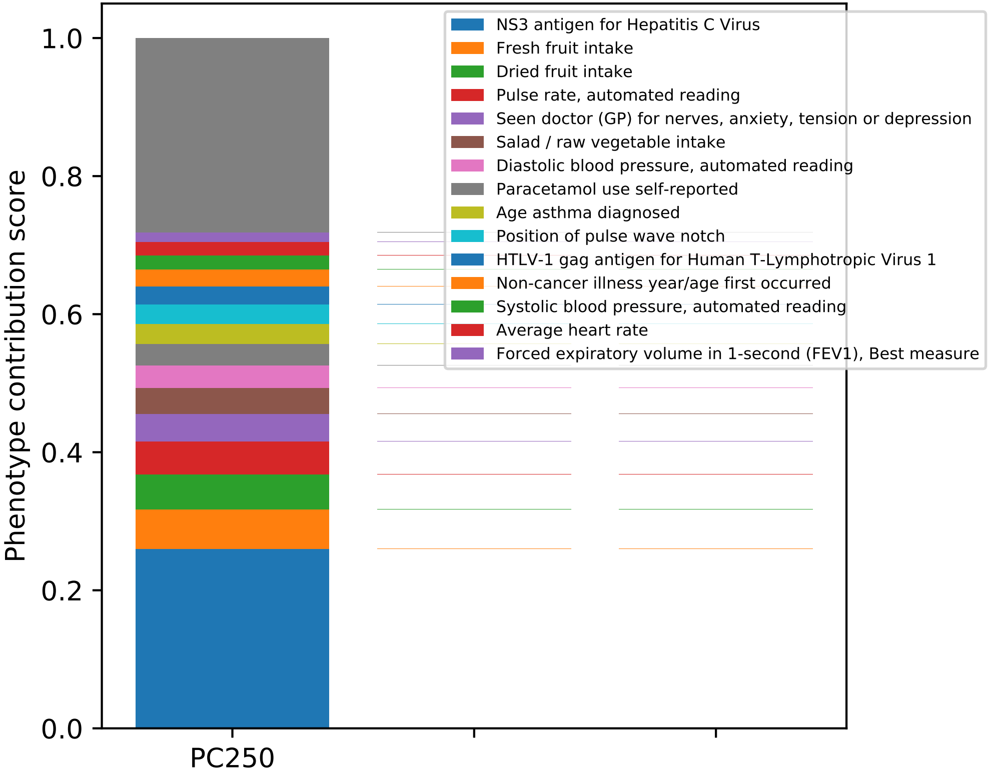
NS3 antigen for Hepatitis C Virus
Fresh fruit intake
Dried fruit intake
Pulse rate, automated reading
Seen doctor (GP) for nerves, anxiety, tension or depression
Salad / raw vegetable intake
Diastolic blood pressure, automated reading
Paracetamol use self-reported
Age asthma diagnosed
Position of pulse wave notch
HTLV-1 gag antigen for Human T-Lymphotropic Virus 1
Non-cancer illness year/age first occurred
Systolic blood pressure, automated reading
Average heart rate
Forced expiratory volume in 1-second (FEV1), Best measure
Phenotype contribution score
1.0
0.8
0.6
0.4
0.2
0.0
PC250

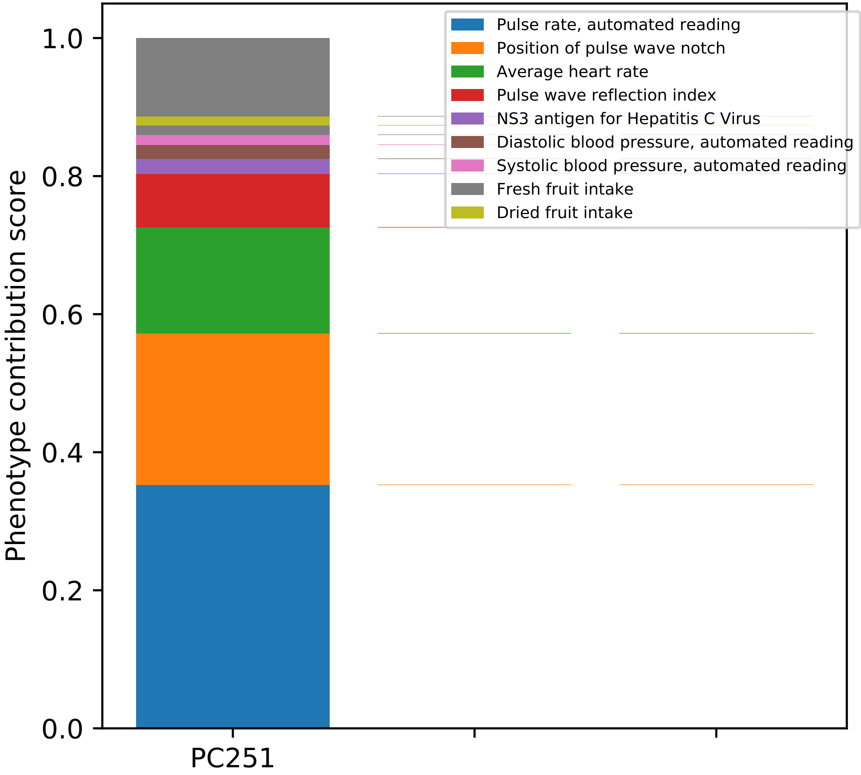
Pulse rate, automated reading
Position of pulse wave notch
Average heart rate
Pulse wave reflection index
NS3 antigen for Hepatitis C Virus
Diastolic blood pressure, automated reading
Systolic blood pressure, automated reading
Fresh fruit intake
Dried fruit intake
Phenotype contribution score
1.0
0.8
0.6
0.4
0.2
0.0
PC251

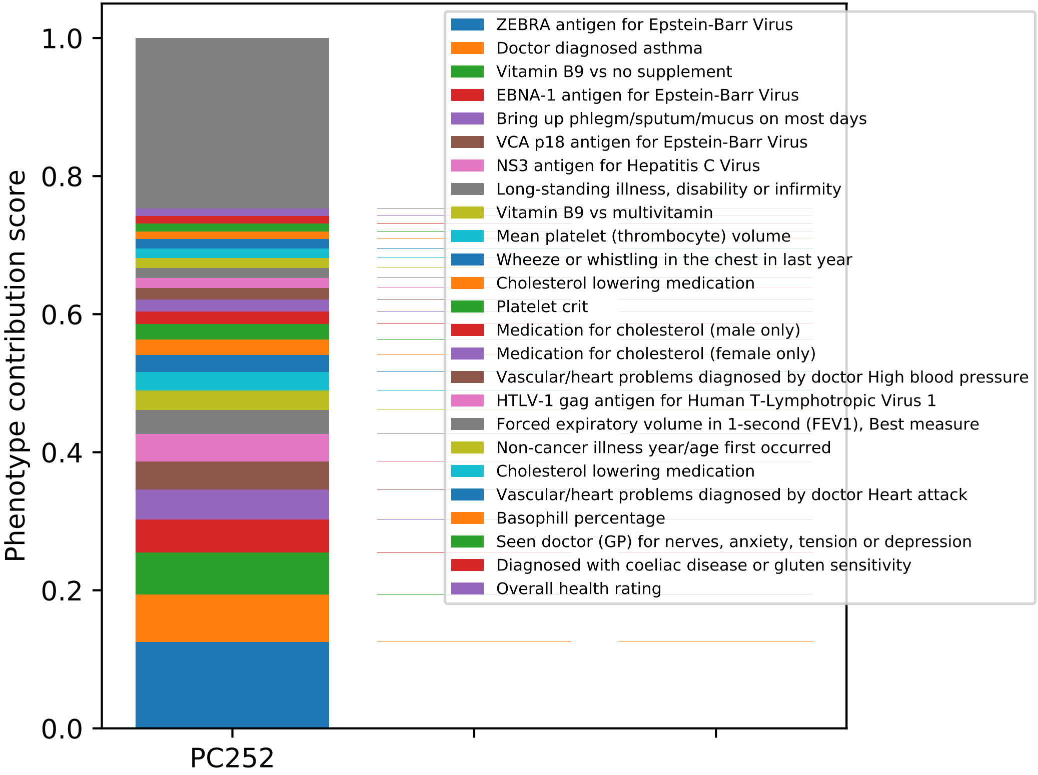

Phenotype contribution score
1.0
0.8
0.6
0.4
0.2
0.0
PC252
ZEBRA antigen for Epstein-Barr Virus
Doctor diagnosed asthma
Vitamin B9 vs no supplement
EBNA-1 antigen for Epstein-Barr Virus
Bring up phlegm/sputum/mucus on most days
VCA p18 antigen for Epstein-Barr Virus
NS3 antigen for Hepatitis C Virus
Long-standing illness, disability or infirmity
Vitamin B9 vs multivitamin
Mean platelet (thrombocyte) volume
Wheeze or whistling in the chest in last year
Cholesterol lowering medication
Platelet crit
Medication for cholesterol (male only)
Medication for cholesterol (female only)
Vascular/heart problems diagnosed by doctor High blood pressure
HTLV-1 gag antigen for Human T-Lymphotropic Virus 1
Forced expiratory volume in 1-second (FEV1), Best measure
Non-cancer illness year/age first occurred
Cholesterol lowering medication
Vascular/heart problems diagnosed by doctor Heart attack
Basophill percentage
Seen doctor (GP) for nerves, anxiety, tension or depression
Diagnosed with coeliac disease or gluten sensitivity
Overall health rating

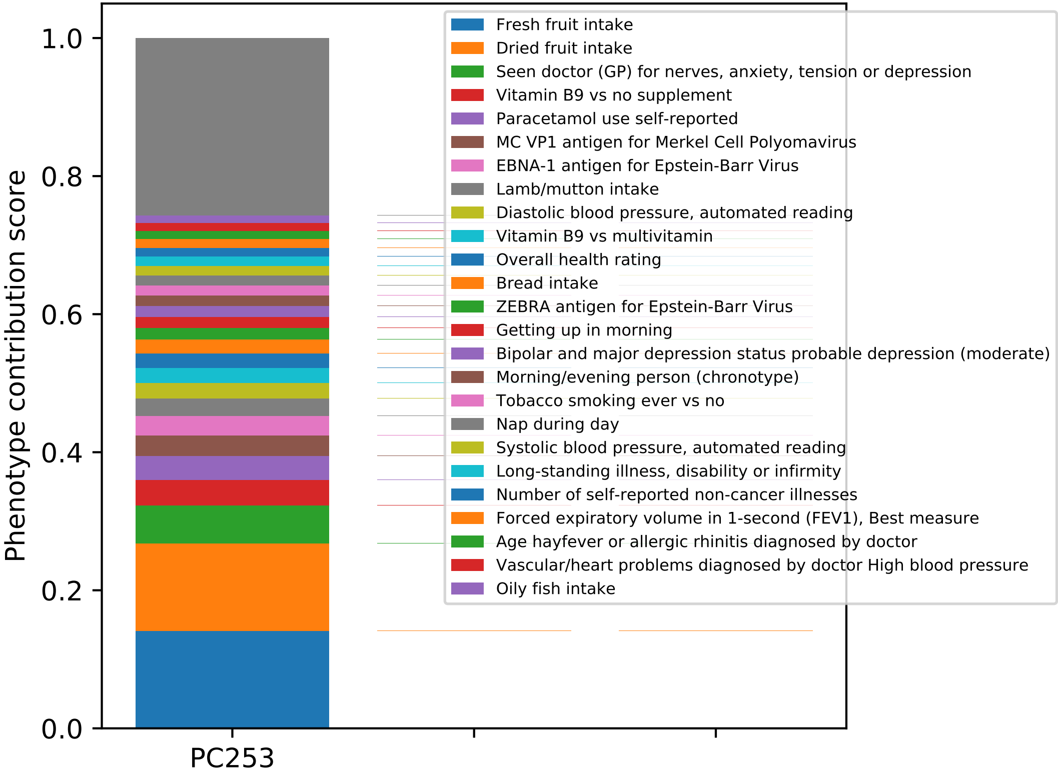

Phenotype contribution score
1.0
0.8
0.6
0.4
0.2
0.0
PC253
Fresh fruit intake
Dried fruit intake
Seen doctor (GP) for nerves, anxiety, tension or depression
Vitamin B9 vs no supplement
Paracetamol use self-reported
MC VP1 antigen for Merkel Cell Polyomavirus
EBNA-1 antigen for Epstein-Barr Virus
Lamb/mutton intake
Diastolic blood pressure, automated reading
Vitamin B9 vs multivitamin
Overall health rating
Bread intake
ZEBRA antigen for Epstein-Barr Virus
Getting up in morning
Bipolar and major depression status probable depression (moderate)
Morning/evening person (chronotype)
Tobacco smoking ever vs no
Nap during day
Systolic blood pressure, automated reading
Long-standing illness, disability or infirmity
Number of self-reported non-cancer illnesses
Forced expiratory volume in 1-second (FEV1), Best measure
Age hayfever or allergic rhinitis diagnosed by doctor
Vascular/heart problems diagnosed by doctor High blood pressure
Oily fish intake

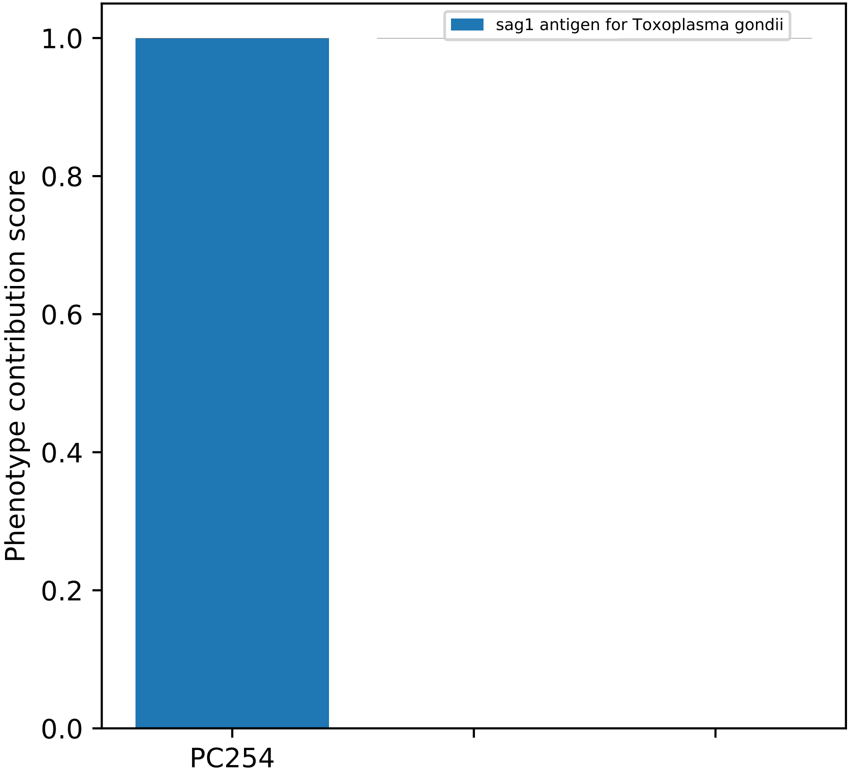
sag1 antigen for Toxoplasma gondii
Phenotype contribution score
1.0
0.8
0.6
0.4
0.2
0.0
PC254

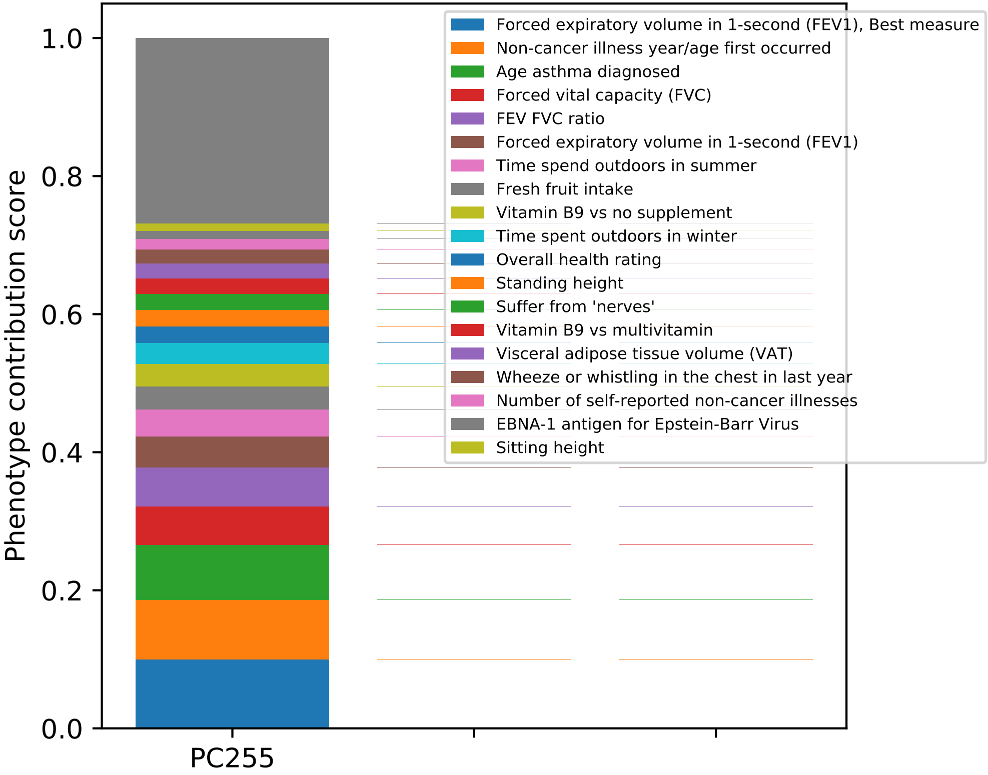
Phenotype contribution score
1.0
0.8
0.6
0.4
0.2
0.0
PC255
Forced expiratory volume in 1-second (FEV1), Best measure
Non-cancer illness year/age first occurred
Age asthma diagnosed
Forced vital capacity (FVC)
FEV FVC ratio
Forced expiratory volume in 1-second (FEV1)
Time spend outdoors in summer
Fresh fruit intake
Vitamin B9 vs no supplement
Time spent outdoors in winter
Overall health rating
Standing height
Suffer from 'nerves'
Vitamin B9 vs multivitamin
Visceral adipose tissue volume (VAT)
Wheeze or whistling in the chest in last year
Number of self-reported non-cancer illnesses
EBNA-1 antigen for Epstein-Barr Virus
Sitting height

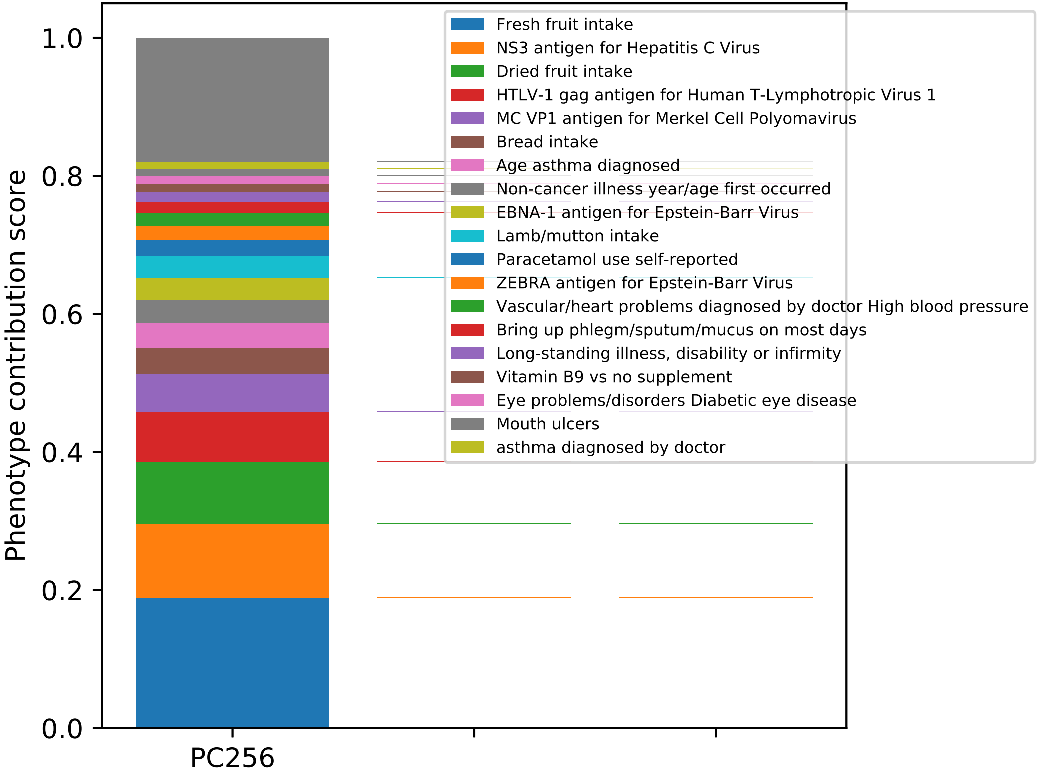

Fresh fruit intake
NS3 antigen for Hepatitis C Virus
Dried fruit intake
HTLV-1 gag antigen for Human T-Lymphotropic Virus 1
MC VP1 antigen for Merkel Cell Polyomavirus
Bread intake
Age asthma diagnosed
Non-cancer illness year/age first occurred
EBNA-1 antigen for Epstein-Barr Virus
Lamb/mutton intake
Paracetamol use self-reported
ZEBRA antigen for Epstein-Barr Virus
Vascular/heart problems diagnosed by doctor High blood pressure
Bring up phlegm/sputum/mucus on most days
Long-standing illness, disability or infirmity
Vitamin B9 vs no supplement
Eye problems/disorders Diabetic eye disease
Mouth ulcers
asthma diagnosed by doctor
1.0
0.8
0.6
0.4
0.2
0.0
Phenotype contribution score
PC256

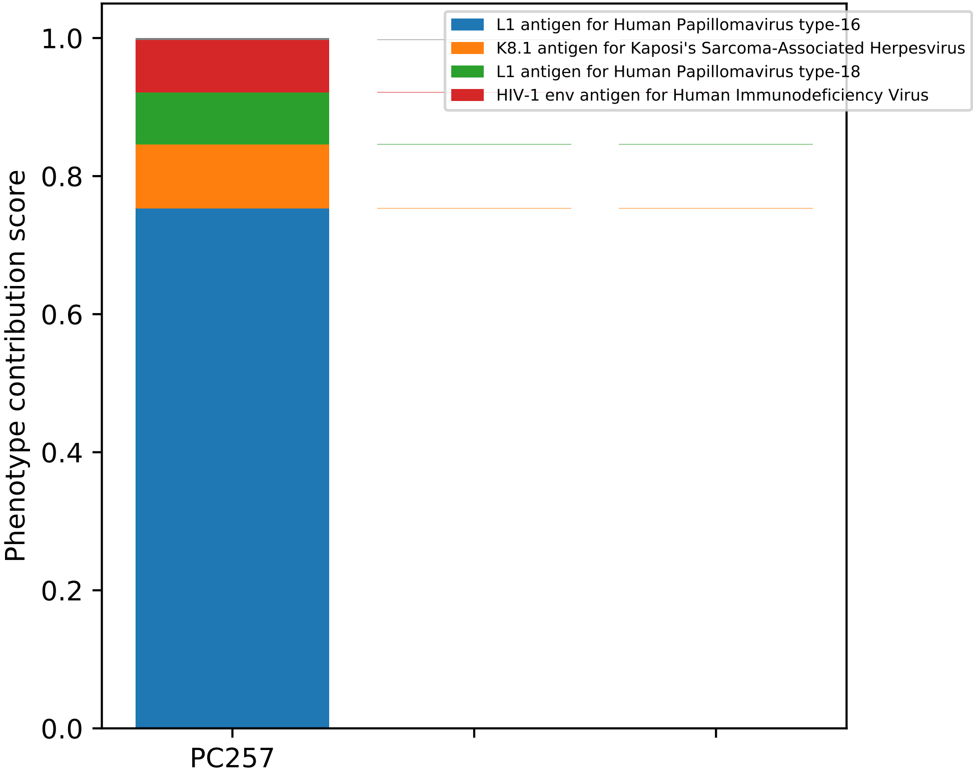

L1 antigen for Human Papillomavirus type-16
K8.1 antigen for Kaposi's Sarcoma-Associated Herpesvirus
L1 antigen for Human Papillomavirus type-18
HIV-1 env antigen for Human Immunodeficiency Virus
Phenotype contribution score
0.0
0.2
0.4
0.6
0.8
1.0
PC257

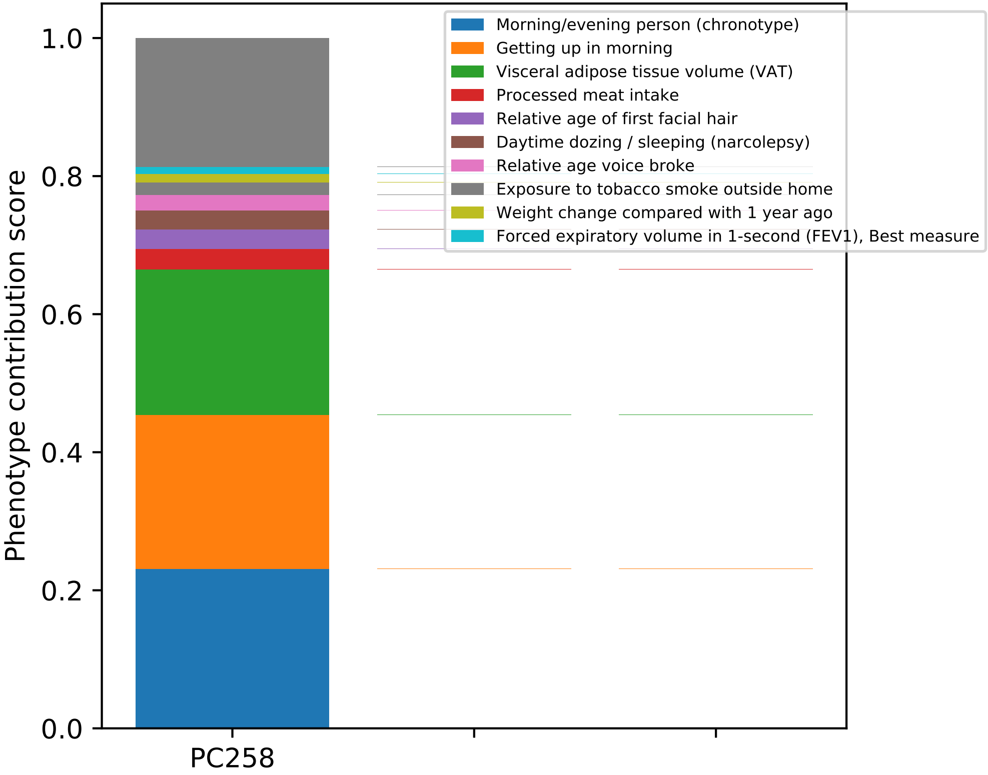
Phenotype contribution score
0.0
0.2
0.4
0.6
0.8
1.0
PC258
Morning/evening person (chronotype)
Getting up in morning
Visceral adipose tissue volume (VAT)
Processed meat intake
Relative age of first facial hair
Daytime dozing / sleeping (narcolepsy)
Relative age voice broke
Exposure to tobacco smoke outside home
Weight change compared with 1 year ago
Forced expiratory volume in 1-second (FEV1), Best measure

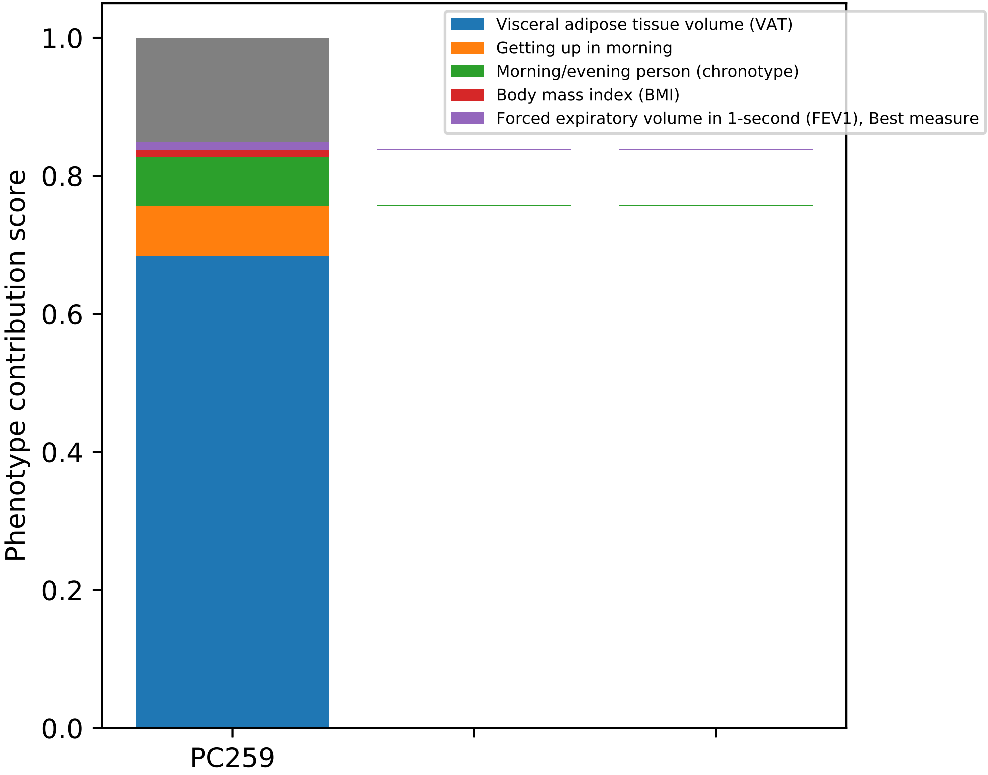

Visceral adipose tissue volume (VAT)
Getting up in morning
Morning/evening person (chronotype)
Body mass index (BMI)
Forced expiratory volume in 1-second (FEV1), Best measure
Phenotype contribution score
1.0
0.8
0.6
0.4
0.2
0.0
PC259

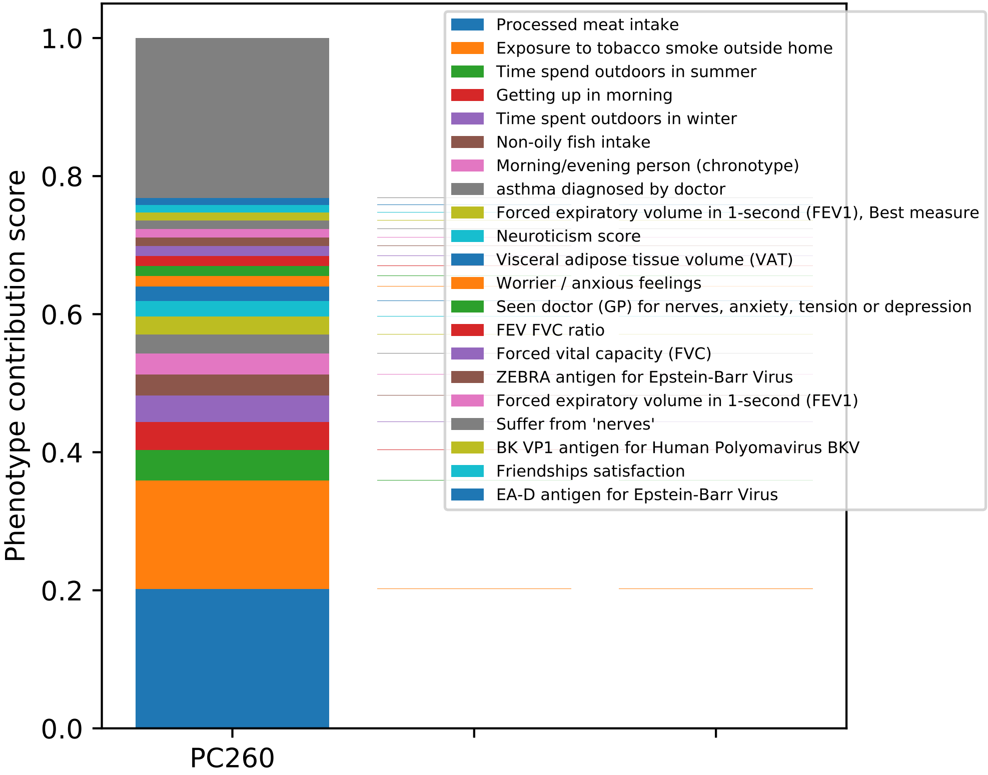

Phenotype contribution score
1.0
0.8
0.6
0.4
0.2
0.0
PC260
Processed meat intake
Exposure to tobacco smoke outside home
Time spend outdoors in summer
Getting up in morning
Time spent outdoors in winter
Non-oily fish intake
Morning/evening person (chronotype)
asthma diagnosed by doctor
Forced expiratory volume in 1-second (FEV1), Best measure
Neuroticism score
Visceral adipose tissue volume (VAT)
Worrier / anxious feelings
Seen doctor (GP) for nerves, anxiety, tension or depression
FEV FVC ratio
Forced vital capacity (FVC)
ZEBRA antigen for Epstein-Barr Virus
Forced expiratory volume in 1-second (FEV1)
Suffer from 'nerves'
BK VP1 antigen for Human Polyomavirus BKV
Friendships satisfaction
EA-D antigen for Epstein-Barr Virus

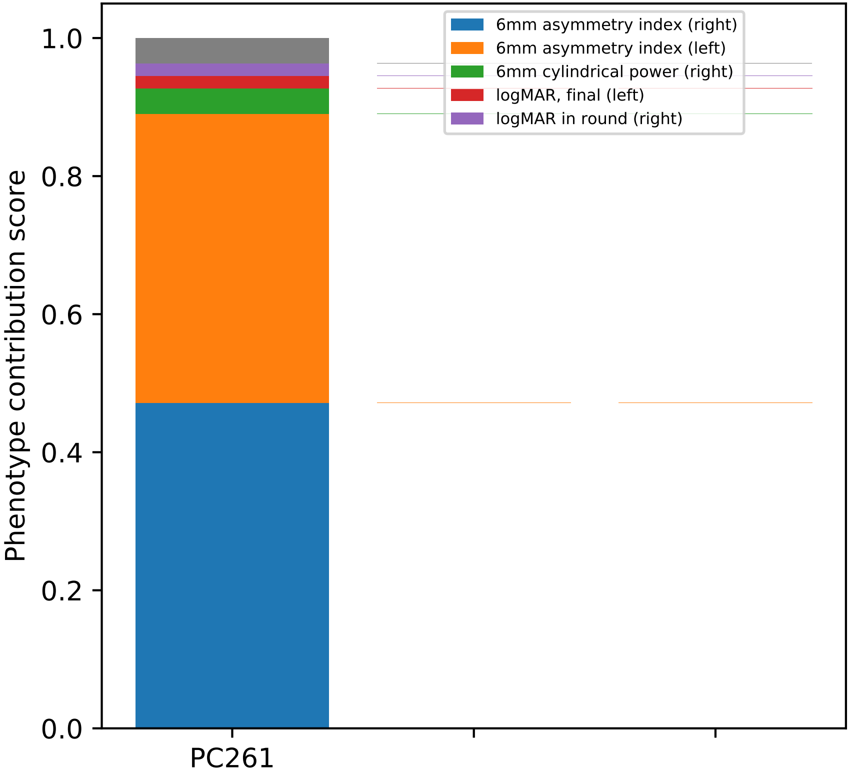

6mm asymmetry index (right)
6mm asymmetry index (left)
6mm cylindrical power (right)
logMAR, final (left)
logMAR in round (right)
Phenotype contribution score
0.0
0.2
0.4
0.6
0.8
1.0
PC261

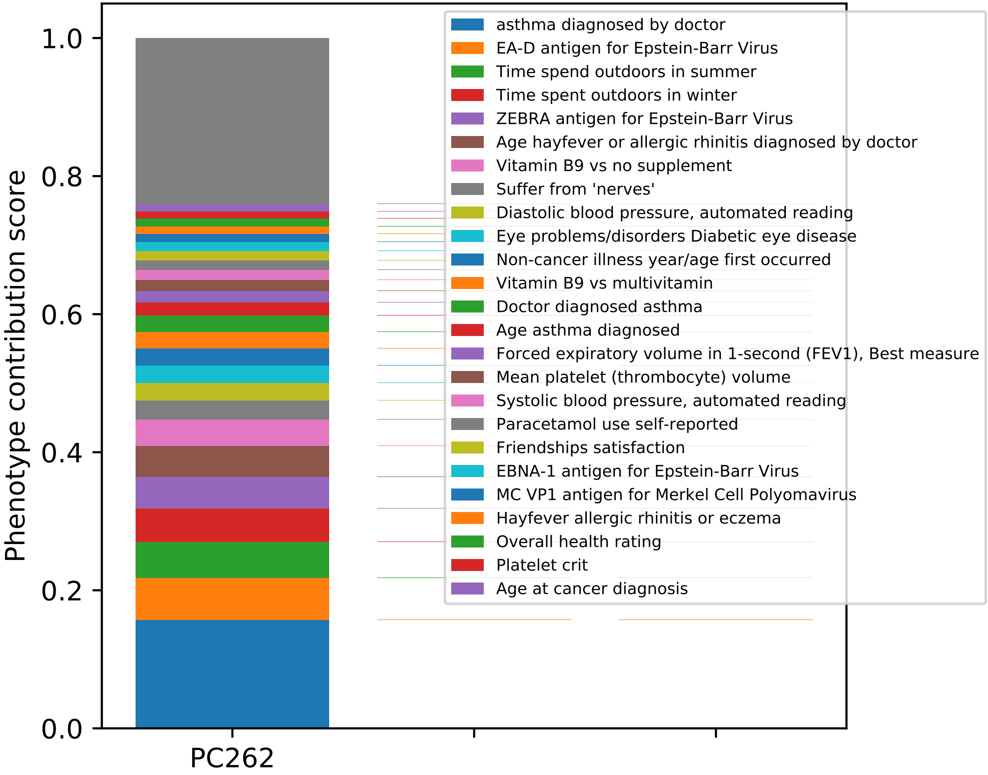

Phenotype contribution score
0.0
0.2
0.4
0.6
0.8
1.0
PC262
asthma diagnosed by doctor
EA-D antigen for Epstein-Barr Virus
Time spend outdoors in summer
Time spent outdoors in winter
ZEBRA antigen for Epstein-Barr Virus
Age hayfever or allergic rhinitis diagnosed by doctor
Vitamin B9 vs no supplement
Suffer from 'nerves'
Diastolic blood pressure, automated reading
Eye problems/disorders Diabetic eye disease
Non-cancer illness year/age first occurred
Vitamin B9 vs multivitamin
Doctor diagnosed asthma
Age asthma diagnosed
Forced expiratory volume in 1-second (FEV1), Best measure
Mean platelet (thrombocyte) volume
Systolic blood pressure, automated reading
Paracetamol use self-reported
Friendships satisfaction
EBNA-1 antigen for Epstein-Barr Virus
MC VP1 antigen for Merkel Cell Polyomavirus
Hayfever allergic rhinitis or eczema
Overall health rating
Platelet crit
Age at cancer diagnosis

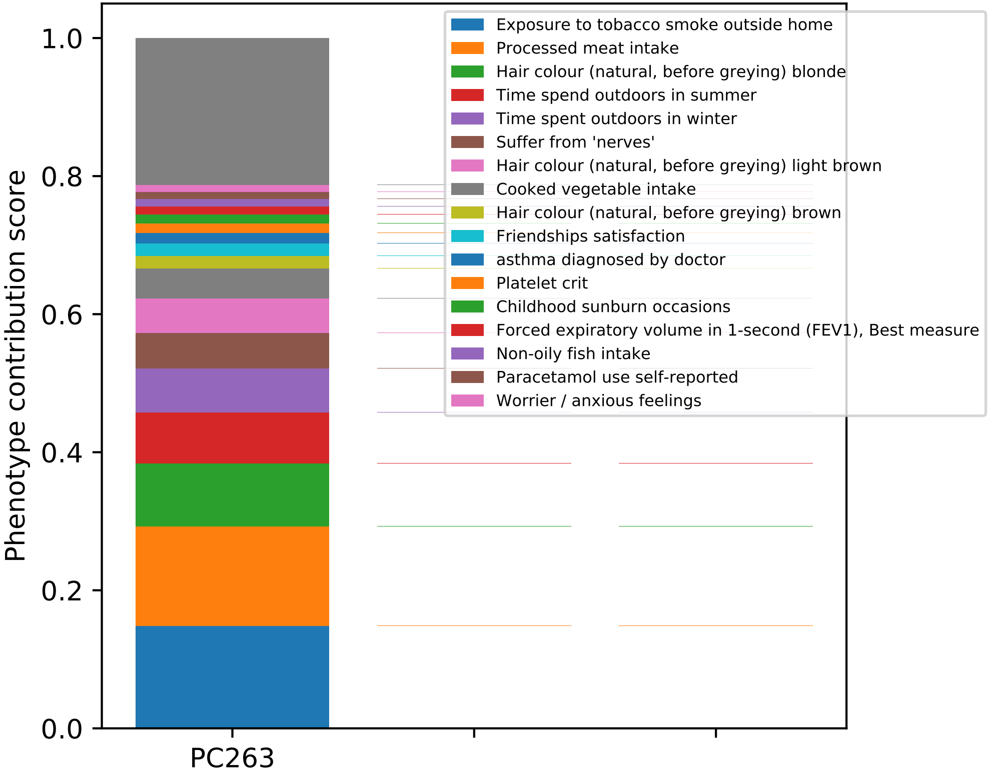

Phenotype contribution score
1.0
0.8
0.6
0.4
0.2
0.0
PC263
Exposure to tobacco smoke outside home
Processed meat intake
Hair colour (natural, before greying) blonde
Time spend outdoors in summer
Time spent outdoors in winter
Suffer from 'nerves'
Hair colour (natural, before greying) light brown
Cooked vegetable intake
Hair colour (natural, before greying) brown
Friendships satisfaction
asthma diagnosed by doctor
Platelet crit
Childhood sunburn occasions
Forced expiratory volume in 1-second (FEV1), Best measure
Non-oily fish intake
Paracetamol use self-reported
Worrier / anxious feelings

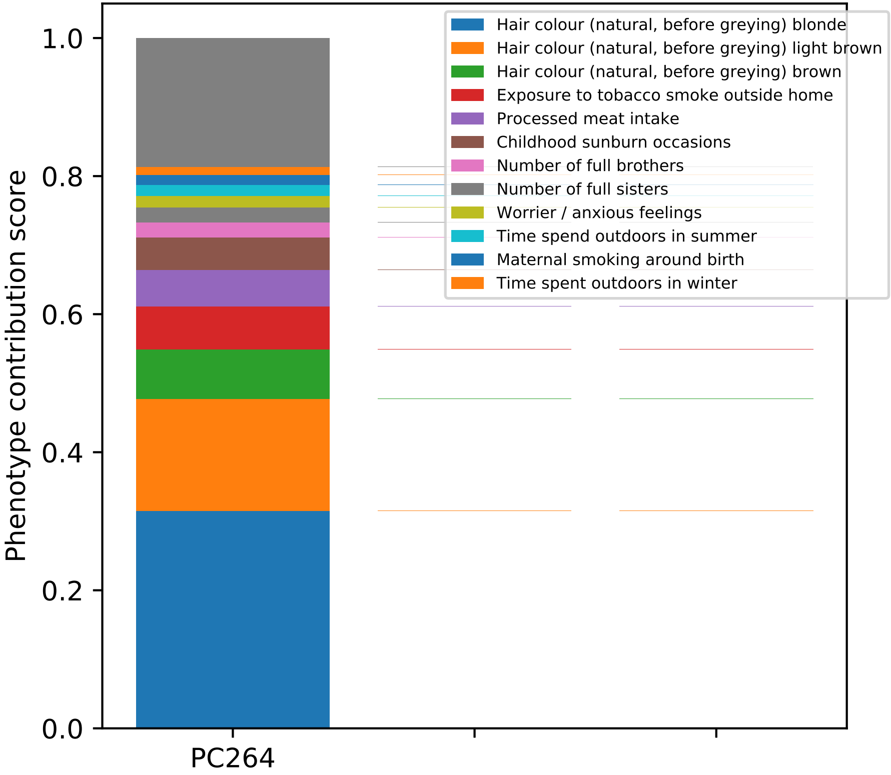

Phenotype contribution score
1.0
0.8
0.6
0.4
0.2
0.0
PC264
Hair colour (natural, before greying) blonde
Hair colour (natural, before greying) light brown
Hair colour (natural, before greying) brown
Exposure to tobacco smoke outside home
Processed meat intake
Childhood sunburn occasions
Number of full brothers
Number of full sisters
Worrier / anxious feelings
Time spend outdoors in summer
Maternal smoking around birth
Time spent outdoors in winter

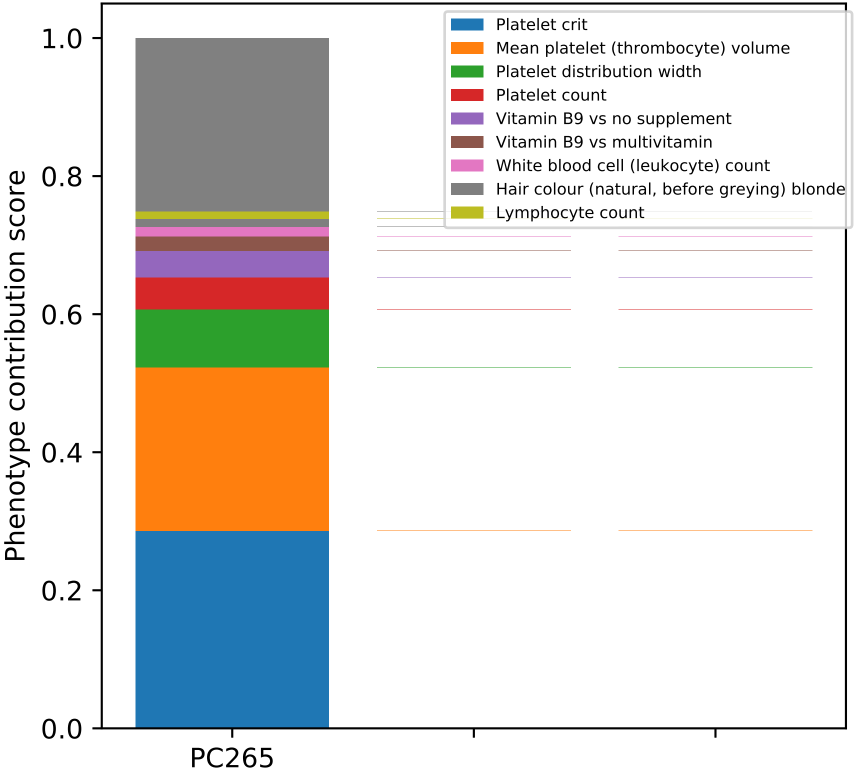

Platelet crit
Mean platelet (thrombocyte) volume
Platelet distribution width
Platelet count
Vitamin B9 vs no supplement
Vitamin B9 vs multivitamin
White blood cell (leukocyte) count
Hair colour (natural, before greying) blonde
Lymphocyte count
Phenotype contribution score
1.0
0.8
0.6
0.4
0.2
0.0
PC265

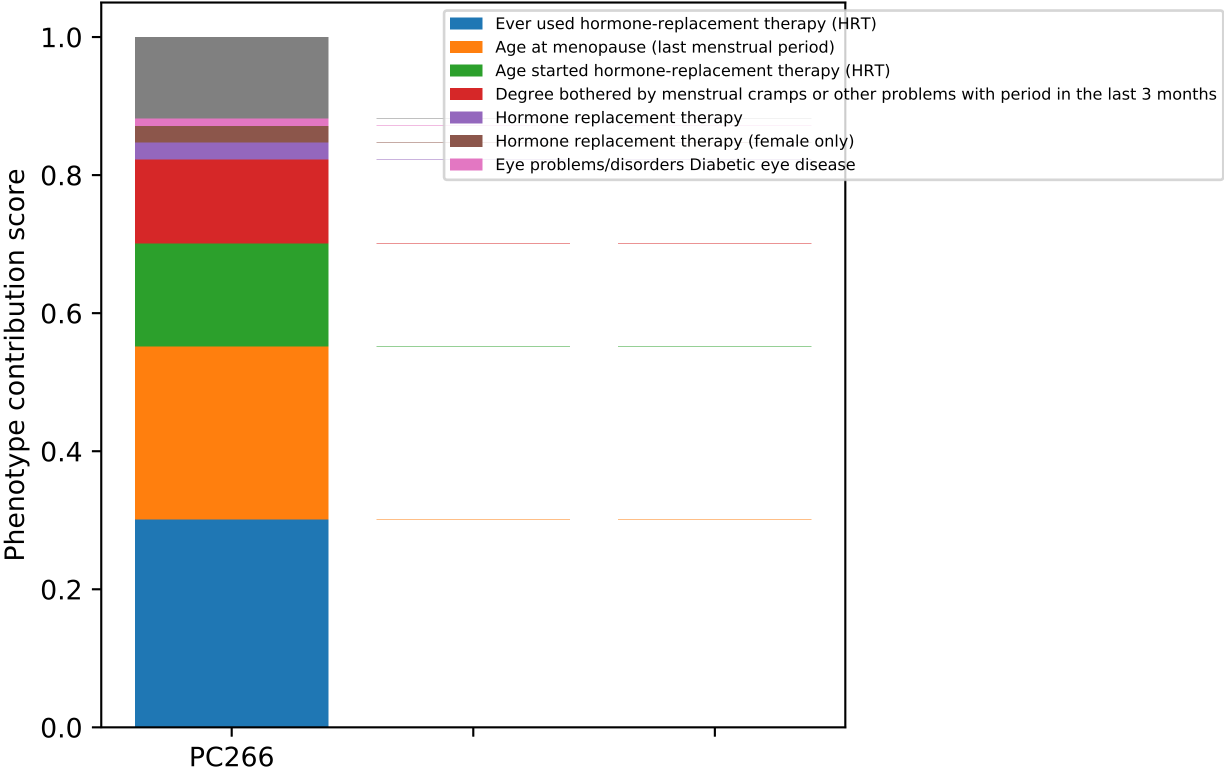

Ever used hormone-replacement therapy (HRT)
Age at menopause (last menstrual period)
Age started hormone-replacement therapy (HRT)
Degree bothered by menstrual cramps or other problems with period in the last 3 months
Hormone replacement therapy
Hormone replacement therapy (female only)
Eye problems/disorders Diabetic eye disease
Phenotype contribution score
1.0
0.8
0.6
0.4
0.2
0.0
PC266

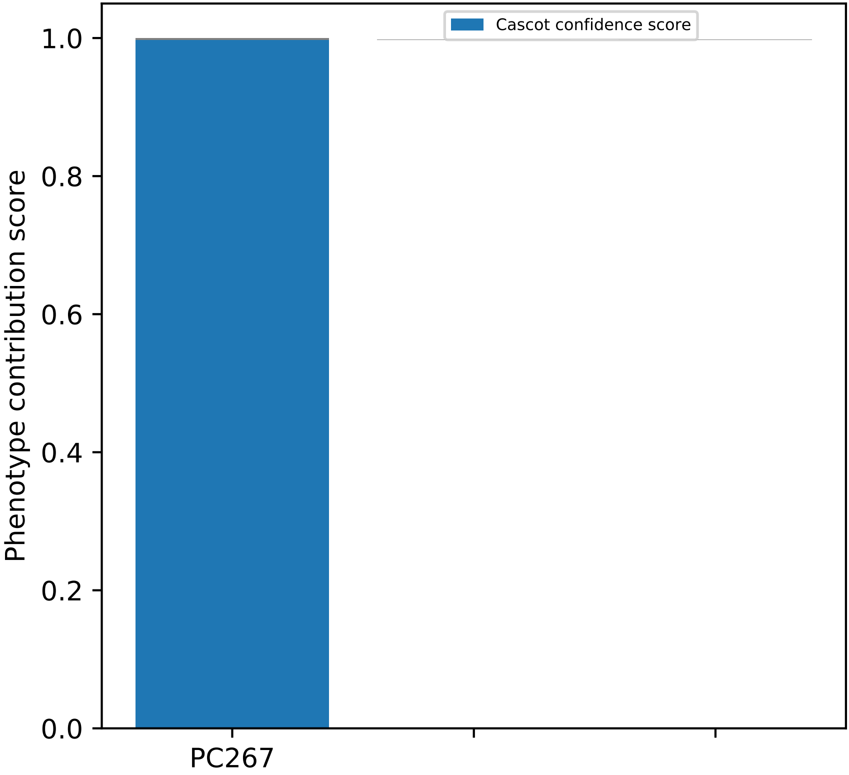
Cascot confidence score
Phenotype contribution score
1.0
0.8
0.6
0.4
0.2
0.0
PC267

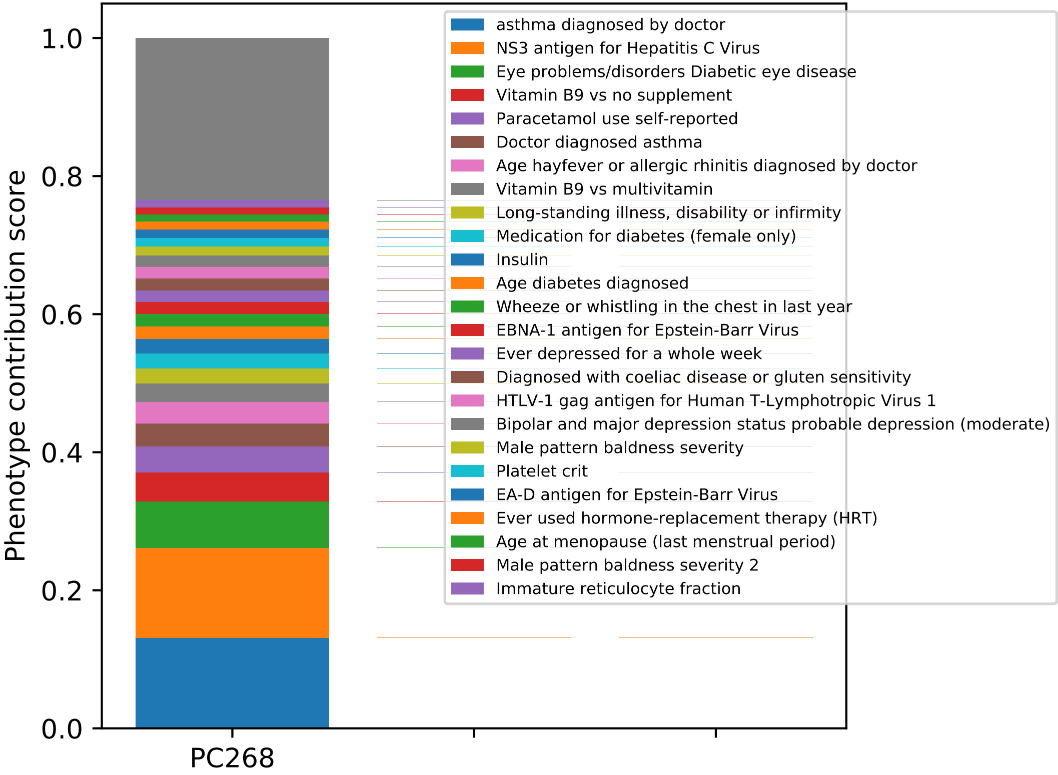

Phenotype contribution score
1.0
0.8
0.6
0.4
0.2
0.0
PC268
asthma diagnosed by doctor
NS3 antigen for Hepatitis C Virus
Eye problems/disorders Diabetic eye disease
Vitamin B9 vs no supplement
Paracetamol use self-reported
Doctor diagnosed asthma
Age hayfever or allergic rhinitis diagnosed by doctor
Vitamin B9 vs multivitamin
Long-standing illness, disability or infirmity
Medication for diabetes (female only)
Insulin
Age diabetes diagnosed
Wheeze or whistling in the chest in last year
EBNA-1 antigen for Epstein-Barr Virus
Ever depressed for a whole week
Diagnosed with coeliac disease or gluten sensitivity
HTLV-1 gag antigen for Human T-Lymphotropic Virus 1
Bipolar and major depression status probable depression (moderate)
Male pattern baldness severity
Platelet crit
EA-D antigen for Epstein-Barr Virus
Ever used hormone-replacement therapy (HRT)
Age at menopause (last menstrual period)
Male pattern baldness severity 2
Immature reticulocyte fraction

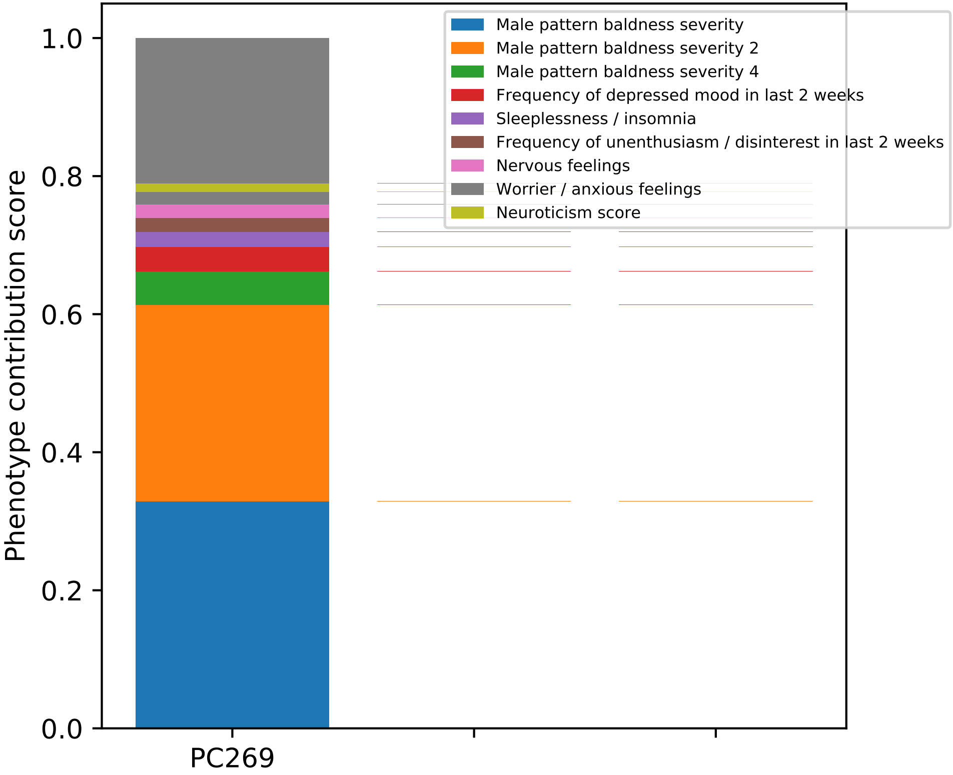

Phenotype contribution score
1.0
0.8
0.6
0.4
0.2
0.0
PC269
Male pattern baldness severity
Male pattern baldness severity 2
Male pattern baldness severity 4
Frequency of depressed mood in last 2 weeks
Sleeplessness / insomnia
Frequency of unenthusiasm / disinterest in last 2 weeks
Nervous feelings
Worrier / anxious feelings
Neuroticism score

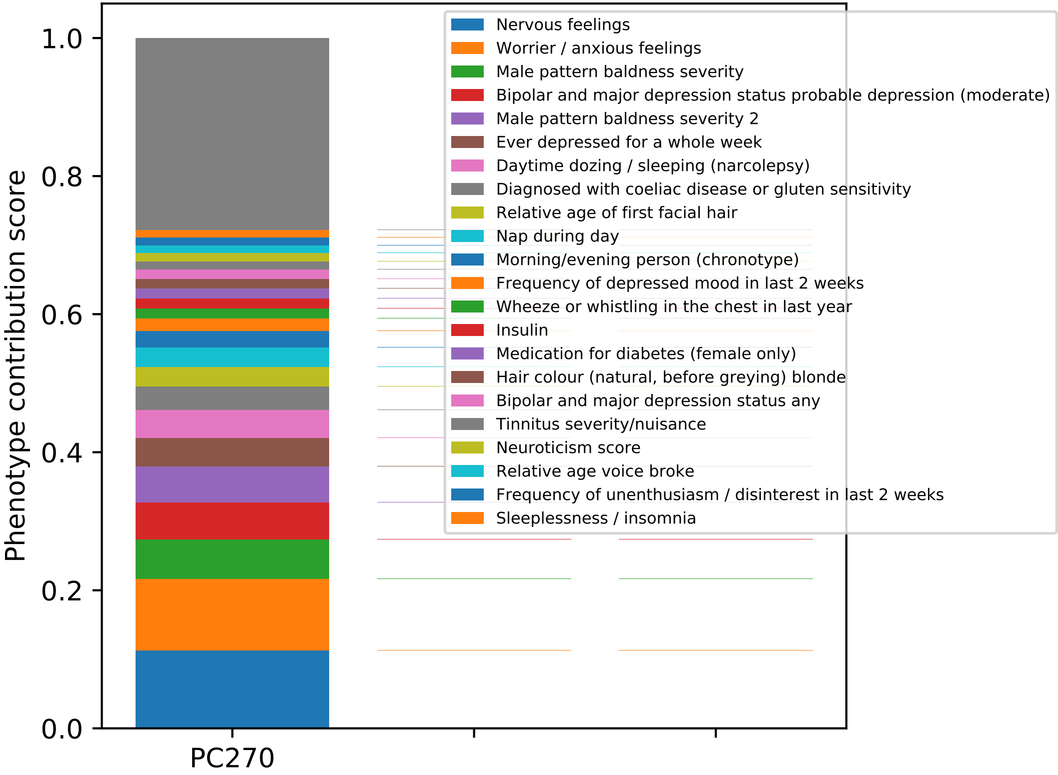

Phenotype contribution score
1.0
0.8
0.6
0.4
0.2
0.0
PC270
Nervous feelings
Worrier / anxious feelings
Male pattern baldness severity
Bipolar and major depression status probable depression (moderate)
Male pattern baldness severity 2
Ever depressed for a whole week
Daytime dozing / sleeping (narcolepsy)
Diagnosed with coeliac disease or gluten sensitivity
Relative age of first facial hair
Nap during day
Morning/evening person (chronotype)
Frequency of depressed mood in last 2 weeks
Wheeze or whistling in the chest in last year
Insulin
Medication for diabetes (female only)
Hair colour (natural, before greying) blonde
Bipolar and major depression status any
Tinnitus severity/nuisance
Neuroticism score
Relative age voice broke
Frequency of unenthusiasm / disinterest in last 2 weeks
Sleeplessness / insomnia

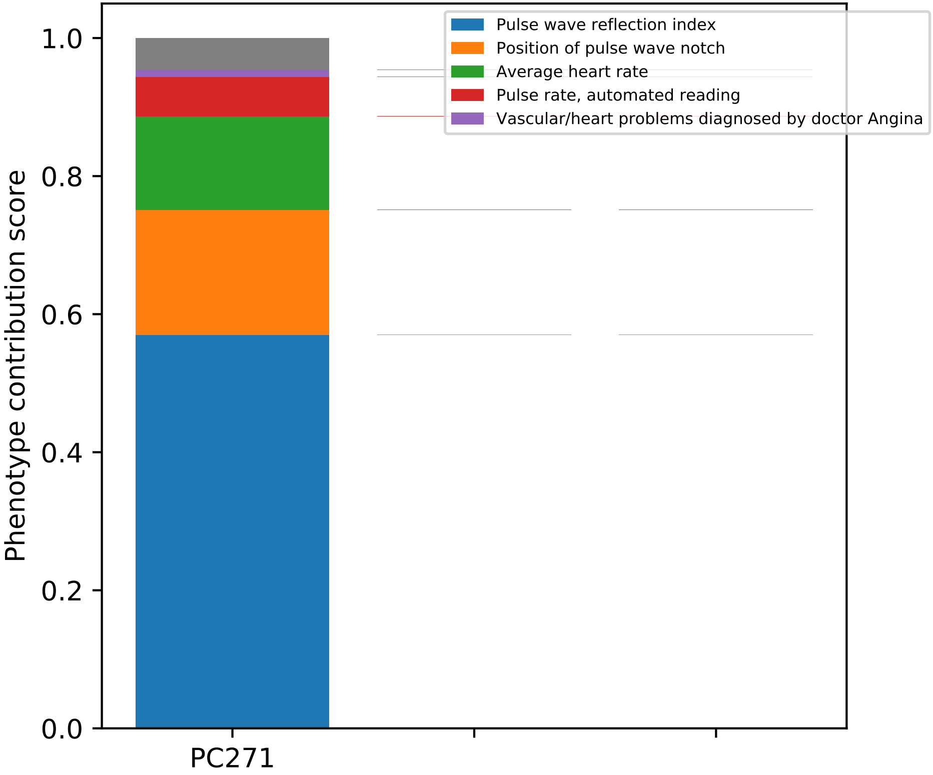
Pulse wave reflection index
Position of pulse wave notch
Average heart rate
Pulse rate, automated reading
Vascular/heart problems diagnosed by doctor Angina
Phenotype contribution score
1.0
0.8
0.6
0.4
0.2
0.0
PC271

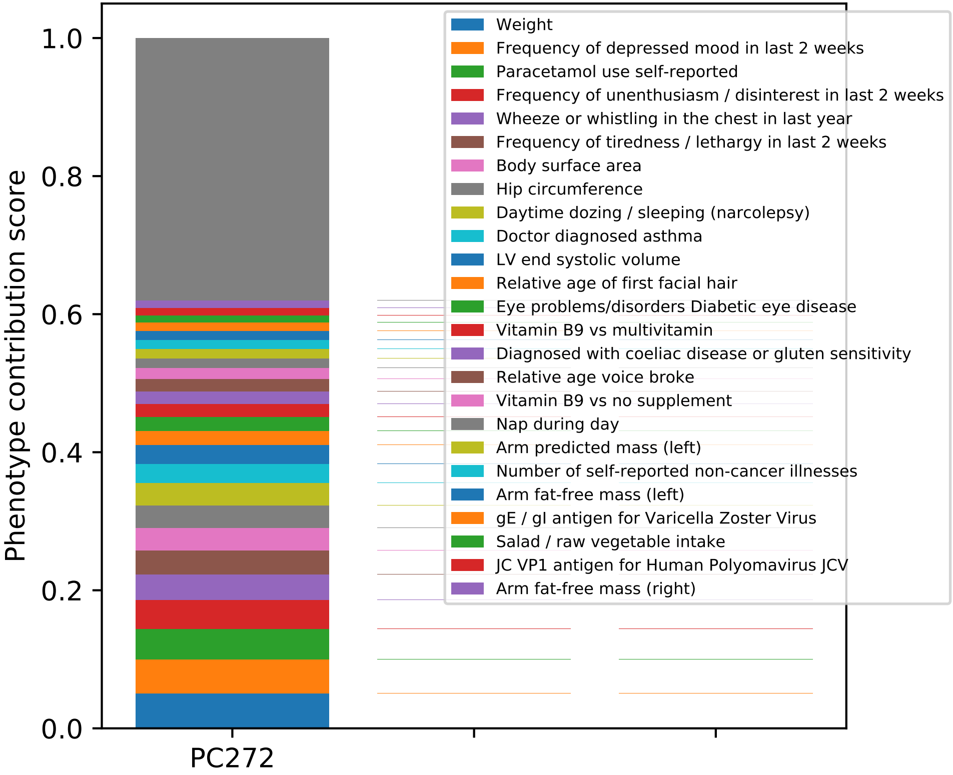

Phenotype contribution score
1.0
0.8
0.6
0.4
0.2
0.0
PC272
Weight
Frequency of depressed mood in last 2 weeks
Paracetamol use self-reported
Frequency of unenthusiasm / disinterest in last 2 weeks
Wheeze or whistling in the chest in last year
Frequency of tiredness / lethargy in last 2 weeks
Body surface area
Hip circumference
Daytime dozing / sleeping (narcolepsy)
Doctor diagnosed asthma
LV end systolic volume
Relative age of first facial hair
Eye problems/disorders Diabetic eye disease
Vitamin B9 vs multivitamin
Diagnosed with coeliac disease or gluten sensitivity
Relative age voice broke
Vitamin B9 vs no supplement
Nap during day
Arm predicted mass (left)
Number of self-reported non-cancer illnesses
Arm fat-free mass (left)
gE / gI antigen for Varicella Zoster Virus
Salad / raw vegetable intake
JC VP1 antigen for Human Polyomavirus JCV
Arm fat-free mass (right)

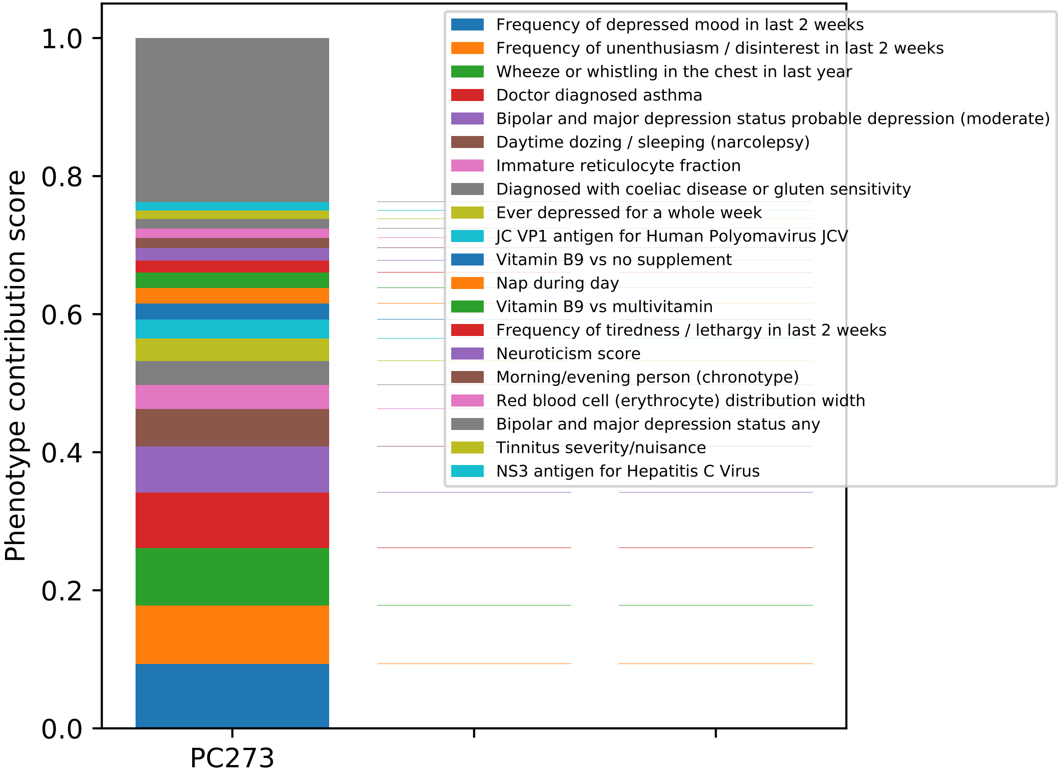

Phenotype contribution score
1.0
0.8
0.6
0.4
0.2
0.0
PC273
Frequency of depressed mood in last 2 weeks
Frequency of unenthusiasm / disinterest in last 2 weeks
Wheeze or whistling in the chest in last year
Doctor diagnosed asthma
Bipolar and major depression status probable depression (moderate)
Daytime dozing / sleeping (narcolepsy)
Immature reticulocyte fraction
Diagnosed with coeliac disease or gluten sensitivity
Ever depressed for a whole week
JC VP1 antigen for Human Polyomavirus JCV
Vitamin B9 vs no supplement
Nap during day
Vitamin B9 vs multivitamin
Frequency of tiredness / lethargy in last 2 weeks
Neuroticism score
Morning/evening person (chronotype)
Red blood cell (erythrocyte) distribution width
Bipolar and major depression status any
Tinnitus severity/nuisance
NS3 antigen for Hepatitis C Virus

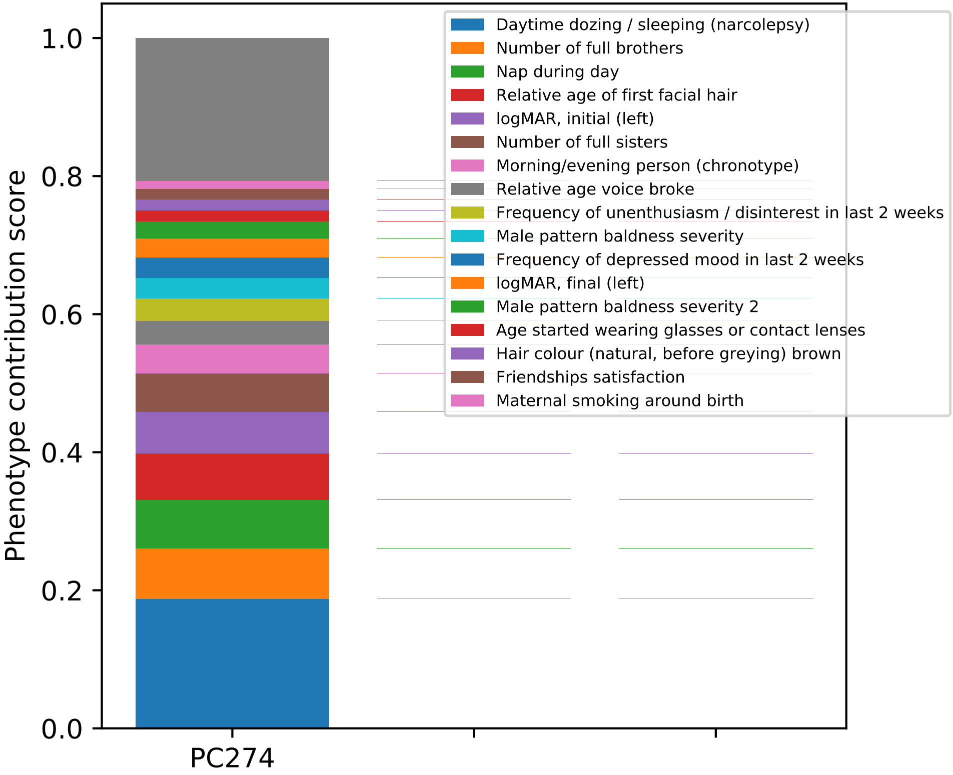
1.0
0.8
0.6
0.4
0.2
0.0
Phenotype contribution score
PC274
Daytime dozing / sleeping (narcolepsy)
Number of full brothers
Nap during day
Relative age of first facial hair
logMAR, initial (left)
Number of full sisters
Morning/evening person (chronotype)
Relative age voice broke
Frequency of unenthusiasm / disinterest in last 2 weeks
Male pattern baldness severity
Frequency of depressed mood in last 2 weeks
logMAR, final (left)
Male pattern baldness severity 2
Age started wearing glasses or contact lenses
Hair colour (natural, before greying) brown
Friendships satisfaction
Maternal smoking around birth

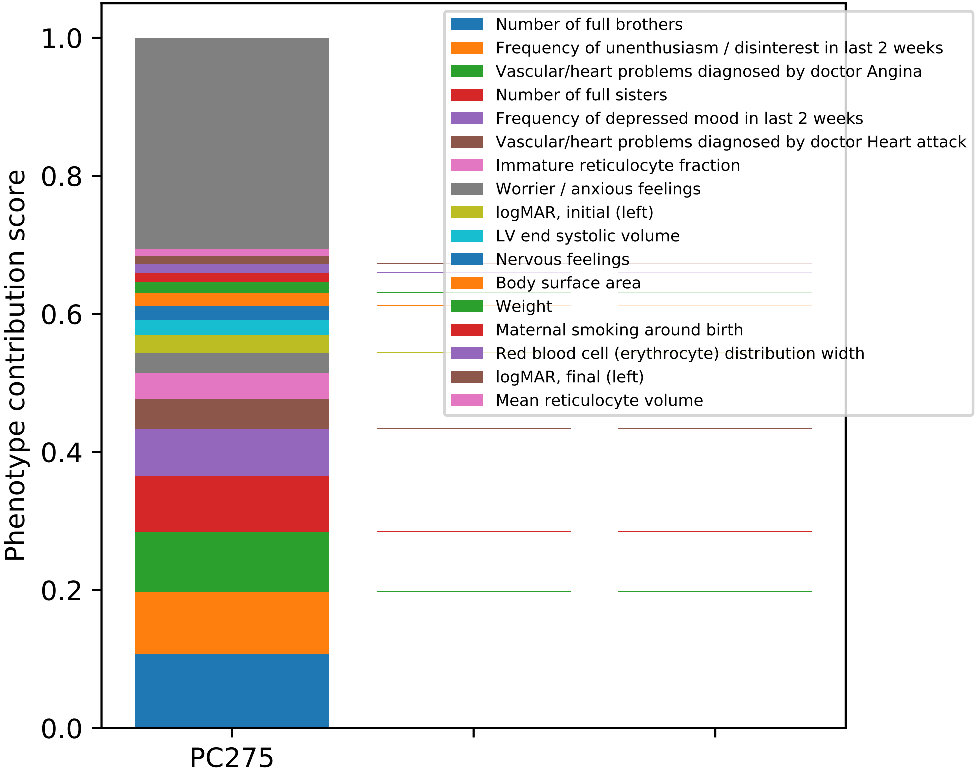
Phenotype contribution score
1.0
0.8
0.6
0.4
0.2
0.0
PC275
Number of full brothers
Frequency of unenthusiasm / disinterest in last 2 weeks
Vascular/heart problems diagnosed by doctor Angina
Number of full sisters
Frequency of depressed mood in last 2 weeks
Vascular/heart problems diagnosed by doctor Heart attack
Immature reticulocyte fraction
Worrier / anxious feelings
logMAR, initial (left)
LV end systolic volume
Nervous feelings
Body surface area
Weight
Maternal smoking around birth
Red blood cell (erythrocyte) distribution width
logMAR, final (left)
Mean reticulocyte volume

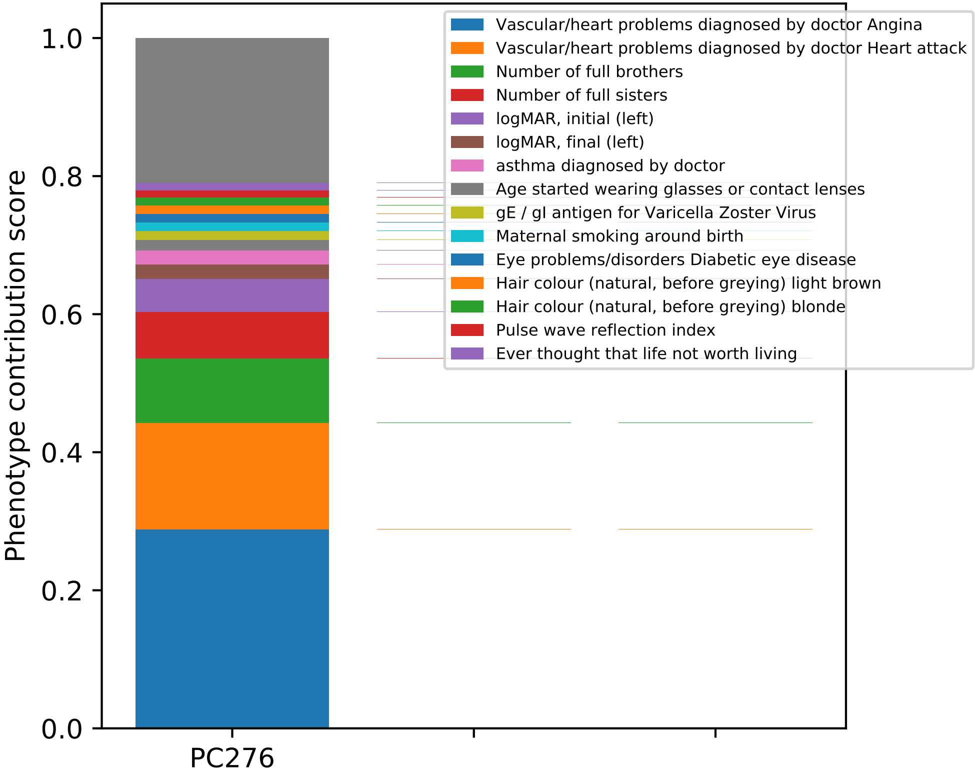

Vascular/heart problems diagnosed by doctor Angina
Vascular/heart problems diagnosed by doctor Heart attack
Number of full brothers
Number of full sisters
logMAR, initial (left)
logMAR, final (left)
asthma diagnosed by doctor
Age started wearing glasses or contact lenses
gE / gI antigen for Varicella Zoster Virus
Maternal smoking around birth
Eye problems/disorders Diabetic eye disease
Hair colour (natural, before greying) light brown
Hair colour (natural, before greying) blonde
Pulse wave reflection index
Ever thought that life not worth living
Phenotype contribution score
0.0
0.2
0.4
0.6
0.8
1.0
PC276

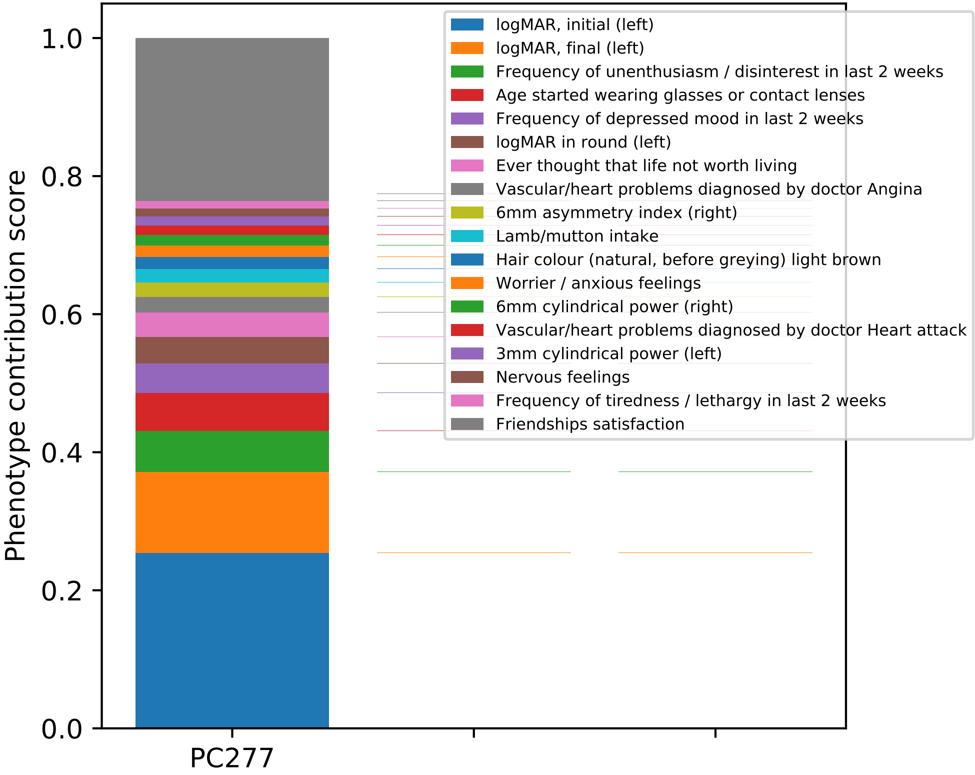

Phenotype contribution score
0.0
0.2
0.4
0.6
0.8
1.0
PC277
logMAR, initial (left)
logMAR, final (left)
Frequency of unenthusiasm / disinterest in last 2 weeks
Age started wearing glasses or contact lenses
Frequency of depressed mood in last 2 weeks
logMAR in round (left)
Ever thought that life not worth living
Vascular/heart problems diagnosed by doctor Angina
6mm asymmetry index (right)
Lamb/mutton intake
Hair colour (natural, before greying) light brown
Worrier / anxious feelings
6mm cylindrical power (right)
Vascular/heart problems diagnosed by doctor Heart attack
3mm cylindrical power (left)
Nervous feelings
Frequency of tiredness / lethargy in last 2 weeks
Friendships satisfaction

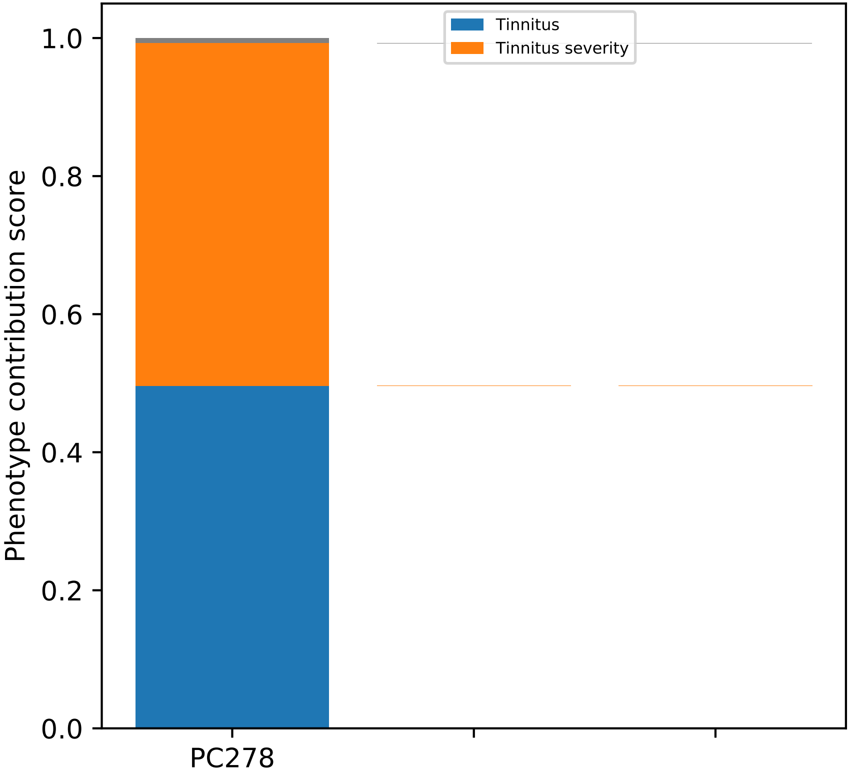
Tinnitus
Tinnitus severity
Phenotype contribution score
1.0
0.8
0.6
0.4
0.2
0.0
PC278

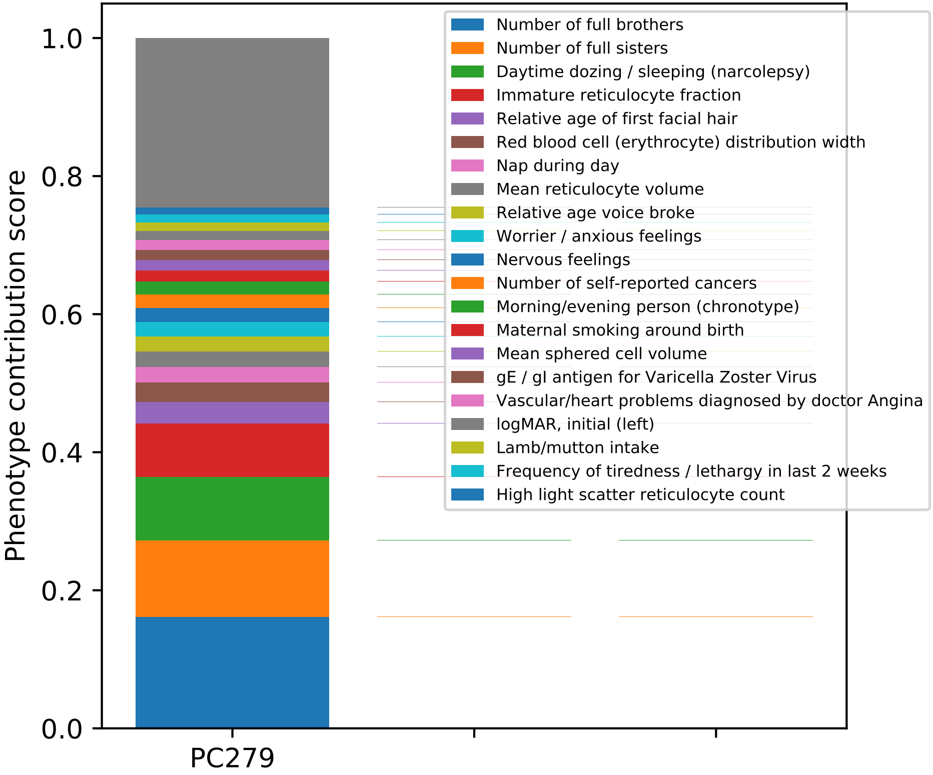

Phenotype contribution score
0.0
0.2
0.4
0.6
0.8
1.0
PC279
Number of full brothers
Number of full sisters
Daytime dozing / sleeping (narcolepsy)
Immature reticulocyte fraction
Relative age of first facial hair
Red blood cell (erythrocyte) distribution width
Nap during day
Mean reticulocyte volume
Relative age voice broke
Worrier / anxious feelings
Nervous feelings
Number of self-reported cancers
Morning/evening person (chronotype)
Maternal smoking around birth
Mean sphered cell volume
gE / gI antigen for Varicella Zoster Virus
Vascular/heart problems diagnosed by doctor Angina
logMAR, initial (left)
Lamb/mutton intake
Frequency of tiredness / lethargy in last 2 weeks
High light scatter reticulocyte count

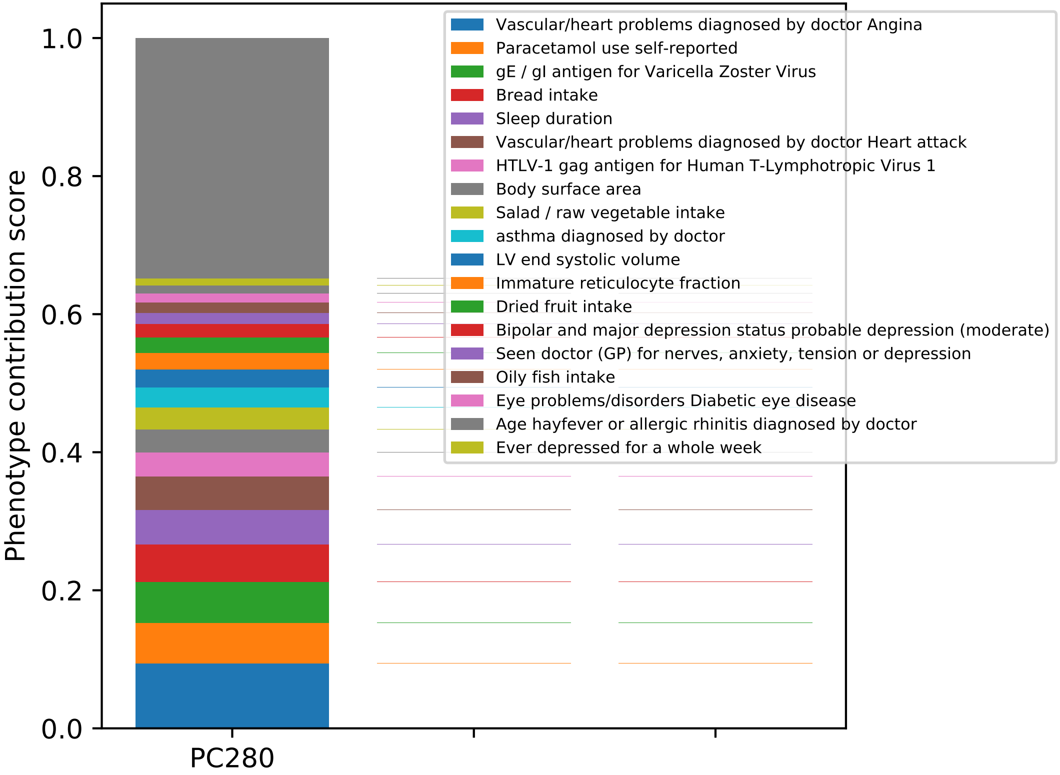
Phenotype contribution score
1.0
0.8
0.6
0.4
0.2
0.0
PC280
Vascular/heart problems diagnosed by doctor Angina
Paracetamol use self-reported
gE / gI antigen for Varicella Zoster Virus
Bread intake
Sleep duration
Vascular/heart problems diagnosed by doctor Heart attack
HTLV-1 gag antigen for Human T-Lymphotropic Virus 1
Body surface area
Salad / raw vegetable intake
asthma diagnosed by doctor
LV end systolic volume
Immature reticulocyte fraction
Dried fruit intake
Bipolar and major depression status probable depression (moderate)
Seen doctor (GP) for nerves, anxiety, tension or depression
Oily fish intake
Eye problems/disorders Diabetic eye disease
Age hayfever or allergic rhinitis diagnosed by doctor
Ever depressed for a whole week

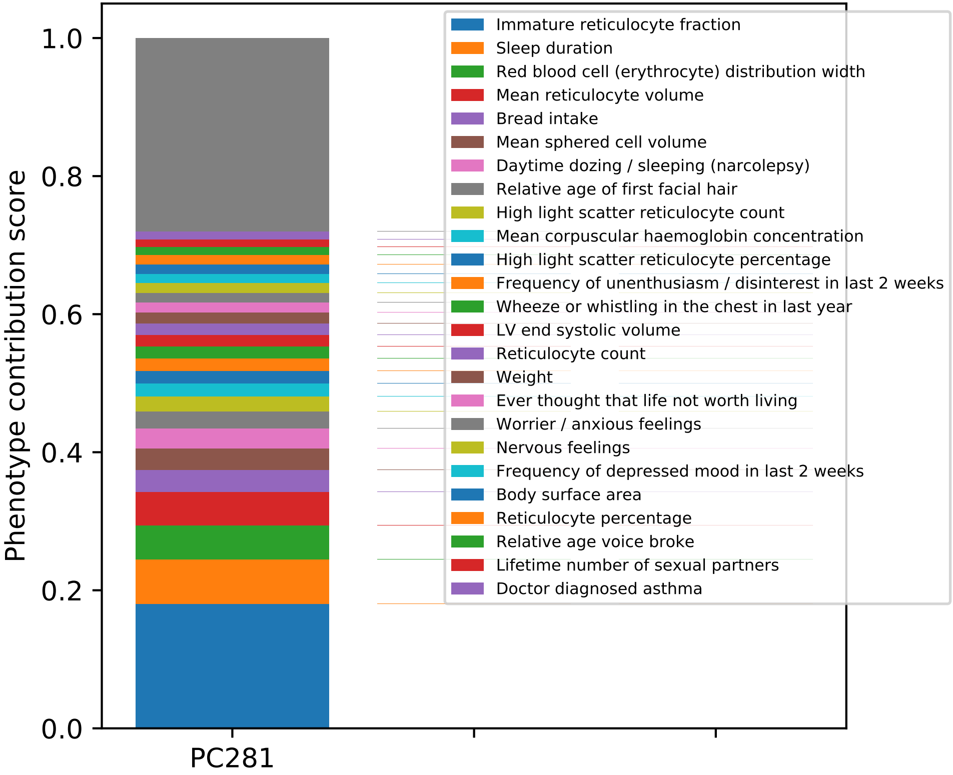

Phenotype contribution score
1.0
0.8
0.6
0.4
0.2
0.0
PC281
Immature reticulocyte fraction
Sleep duration
Red blood cell (erythrocyte) distribution width
Mean reticulocyte volume
Bread intake
Mean sphered cell volume
Daytime dozing / sleeping (narcolepsy)
Relative age of first facial hair
High light scatter reticulocyte count
Mean corpuscular haemoglobin concentration
High light scatter reticulocyte percentage
Frequency of unenthusiasm / disinterest in last 2 weeks
Wheeze or whistling in the chest in last year
LV end systolic volume
Reticulocyte count
Weight
Ever thought that life not worth living
Worrier / anxious feelings
Nervous feelings
Frequency of depressed mood in last 2 weeks
Body surface area
Reticulocyte percentage
Relative age voice broke
Lifetime number of sexual partners
Doctor diagnosed asthma

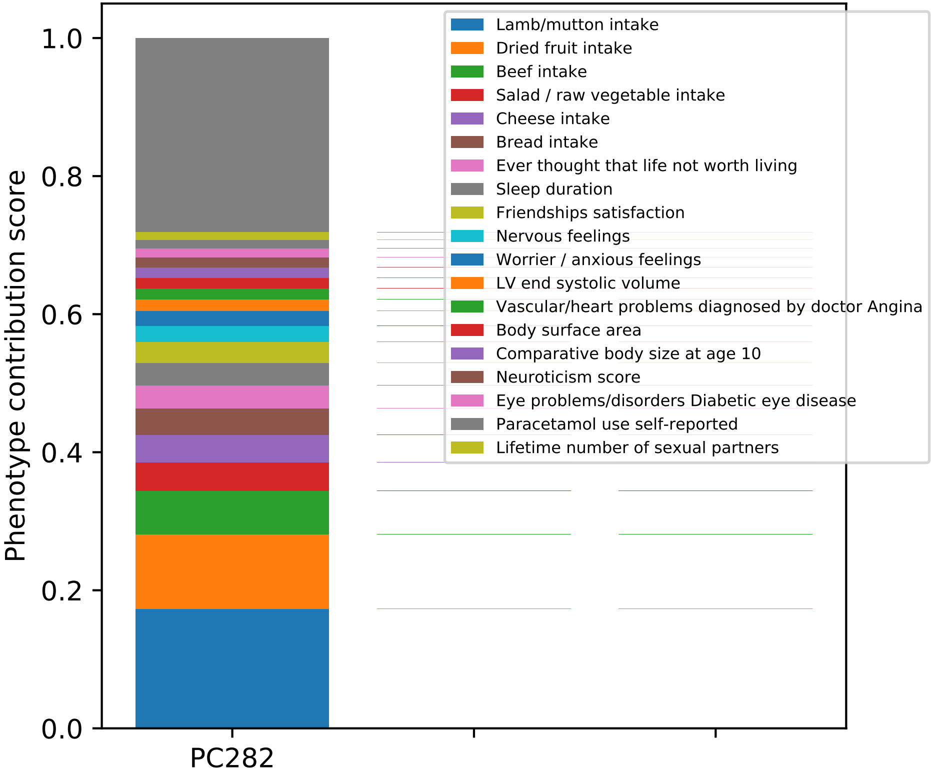

Lamb/mutton intake
Dried fruit intake
Beef intake
Salad / raw vegetable intake
Cheese intake
Bread intake
Ever thought that life not worth living
Sleep duration
Friendships satisfaction
Nervous feelings
Worrier / anxious feelings
LV end systolic volume
Vascular/heart problems diagnosed by doctor Angina
Body surface area
Comparative body size at age 10
Neuroticism score
Eye problems/disorders Diabetic eye disease
Paracetamol use self-reported
Lifetime number of sexual partners
Phenotype contribution score
1.0
0.8
0.6
0.4
0.2
0.0
PC282

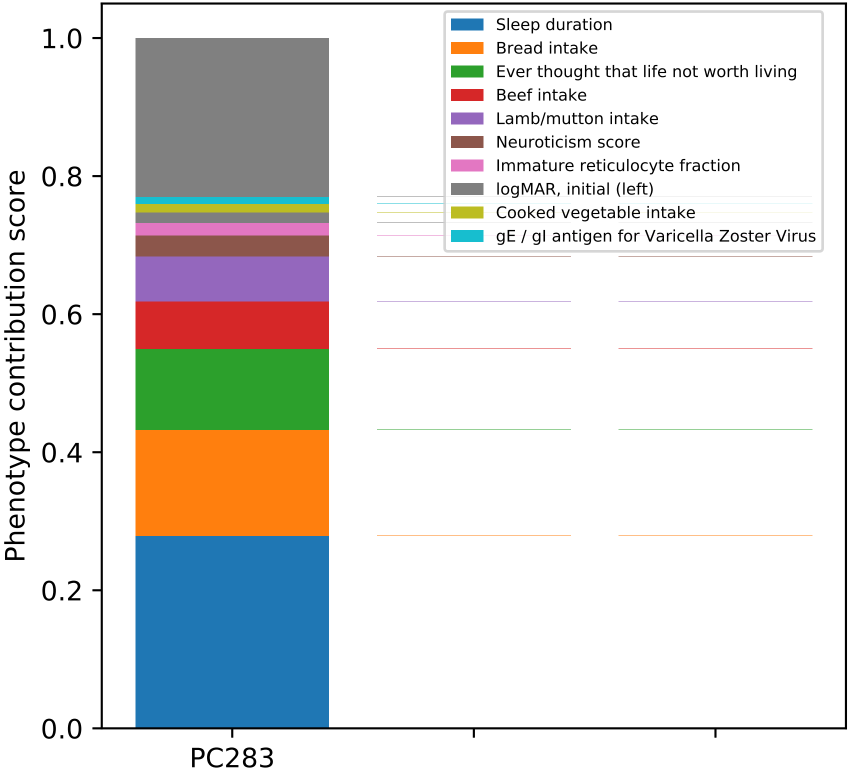

Sleep duration
Bread intake
Ever thought that life not worth living
Beef intake
Lamb/mutton intake
Neuroticism score
Immature reticulocyte fraction
logMAR, initial (left)
Cooked vegetable intake
gE / gI antigen for Varicella Zoster Virus
Phenotype contribution score
0.0
0.2
0.4
0.6
0.8
1.0
PC283

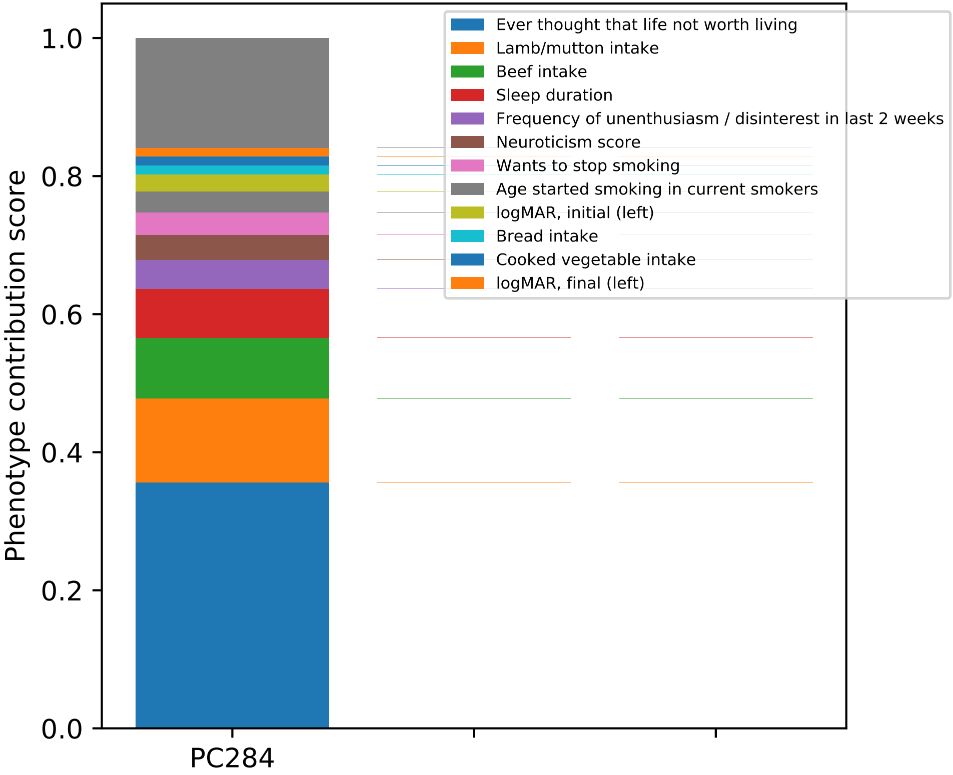

Ever thought that life not worth living
Lamb/mutton intake
Beef intake
Sleep duration
Frequency of unenthusiasm / disinterest in last 2 weeks
Neuroticism score
Wants to stop smoking
Age started smoking in current smokers
logMAR, initial (left)
Bread intake
Cooked vegetable intake
logMAR, final (left)
Phenotype contribution score
1.0
0.8
0.6
0.4
0.2
0.0
PC284

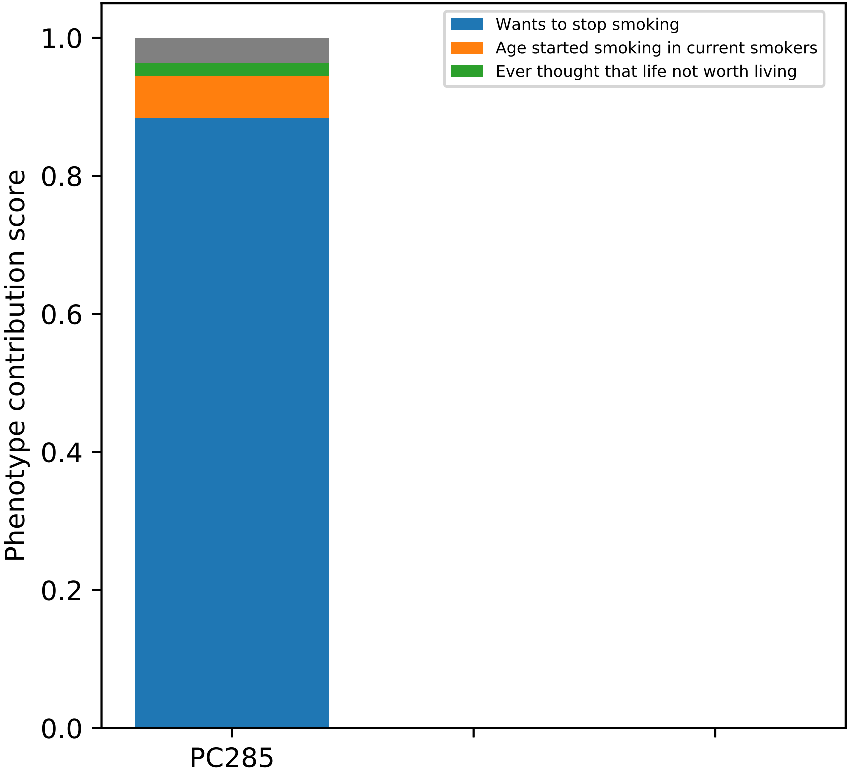

Wants to stop smoking
Age started smoking in current smokers
Ever thought that life not worth living
Phenotype contribution score
1.0
0.8
0.6
0.4
0.2
0.0
PC285

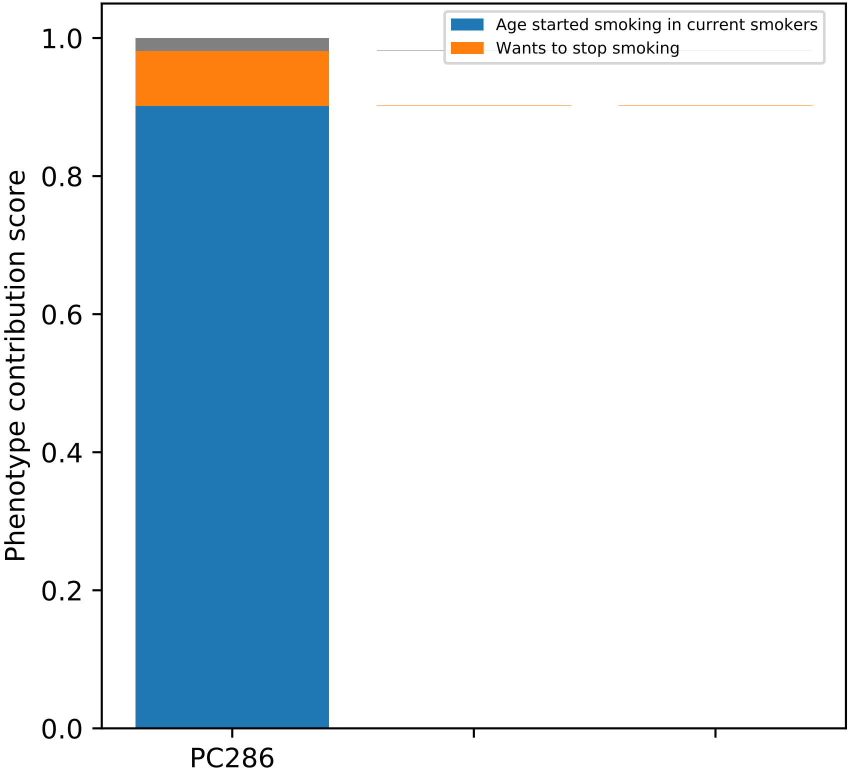

Age started smoking in current smokers
Wants to stop smoking
Phenotype contribution score
0.0
0.2
0.4
0.6
0.8
1.0
PC286

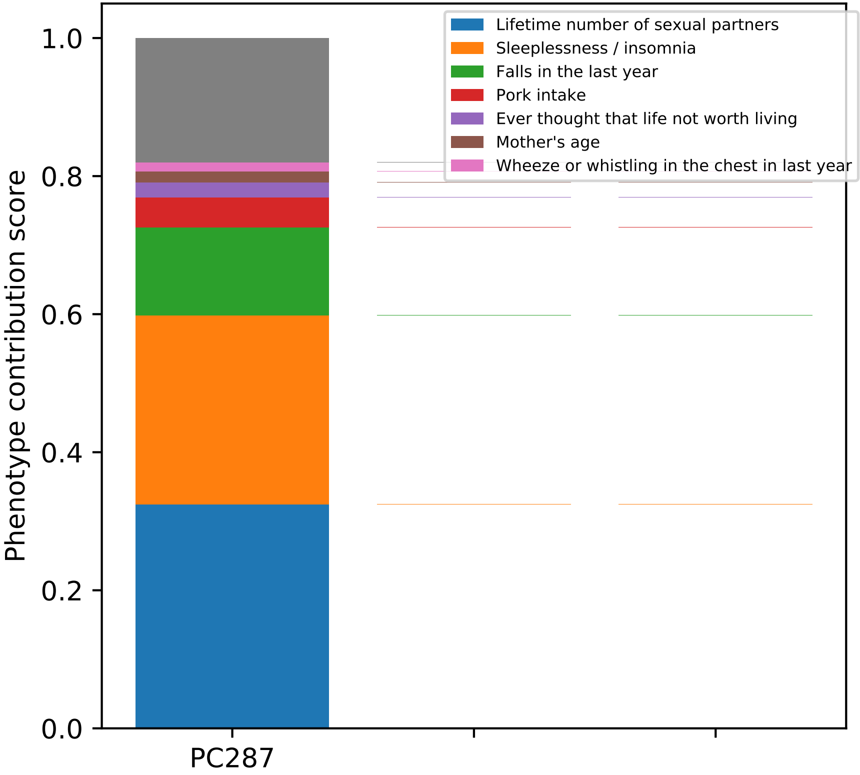

Phenotype contribution score
1.0
0.8
0.6
0.4
0.2
0.0
PC287
Lifetime number of sexual partners
Sleeplessness / insomnia
Falls in the last year
Pork intake
Ever thought that life not worth living
Mother's age
Wheeze or whistling in the chest in last year

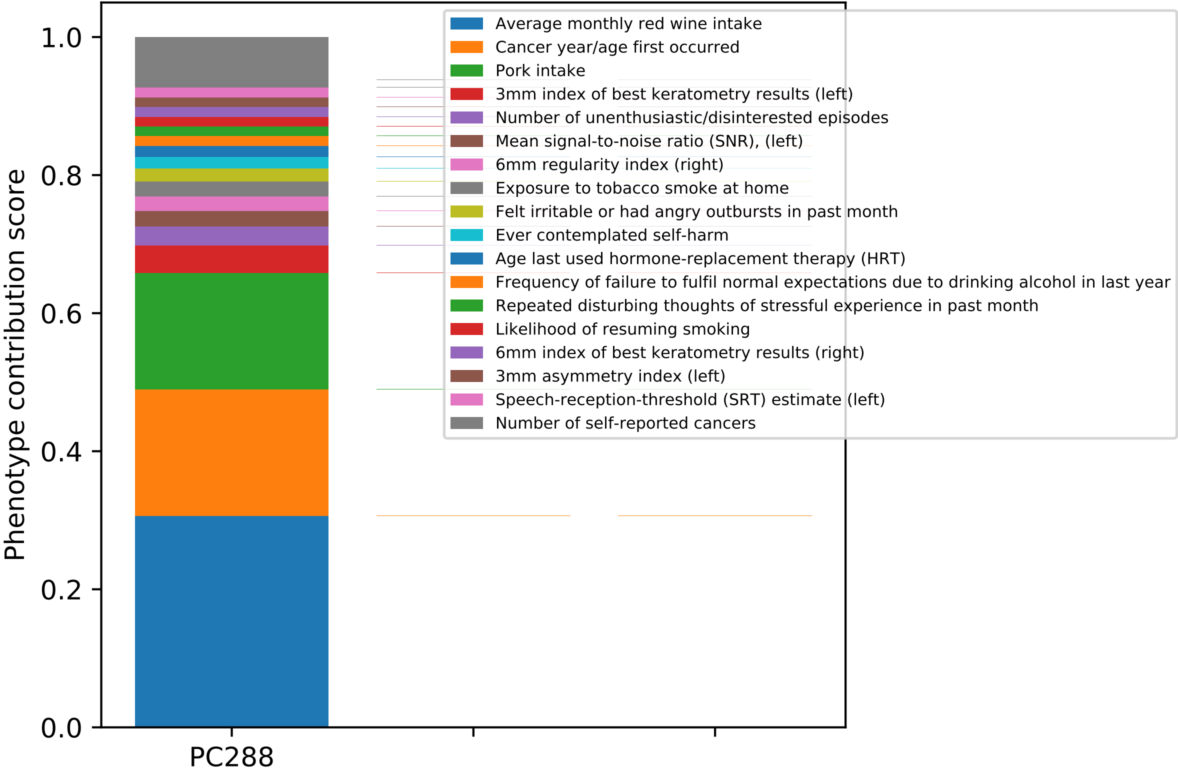

Phenotype contribution score
1.0
0.8
0.6
0.4
0.2
0.0
PC288
Average monthly red wine intake
Cancer year/age first occurred
Pork intake
3mm index of best keratometry results (left)
Number of unenthusiastic/disinterested episodes
Mean signal-to-noise ratio (SNR), (left)
6mm regularity index (right)
Exposure to tobacco smoke at home
Felt irritable or had angry outbursts in past month
Ever contemplated self-harm
Age last used hormone-replacement therapy (HRT)
Frequency of failure to fulfil normal expectations due to drinking alcohol in last year
Repeated disturbing thoughts of stressful experience in past month
Likelihood of resuming smoking
6mm index of best keratometry results (right)
3mm asymmetry index (left)
Speech-reception-threshold (SRT) estimate (left)
Number of self-reported cancers

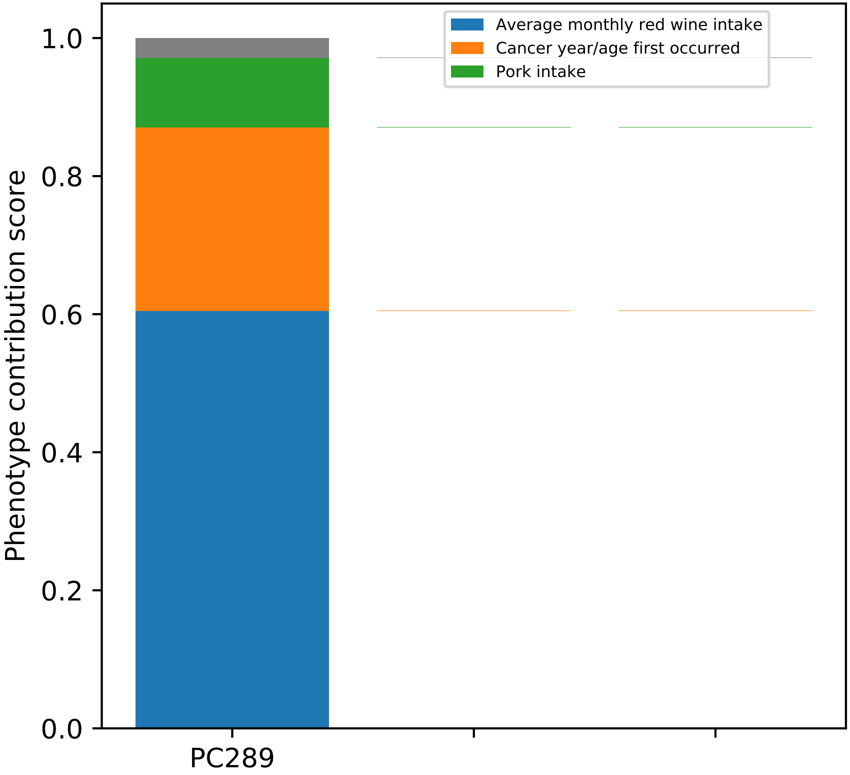
Average monthly red wine intake
Cancer year/age first occurred
Pork intake
1.0
0.8
0.6
0.4
0.2
0.0
Phenotype contribution score
PC289

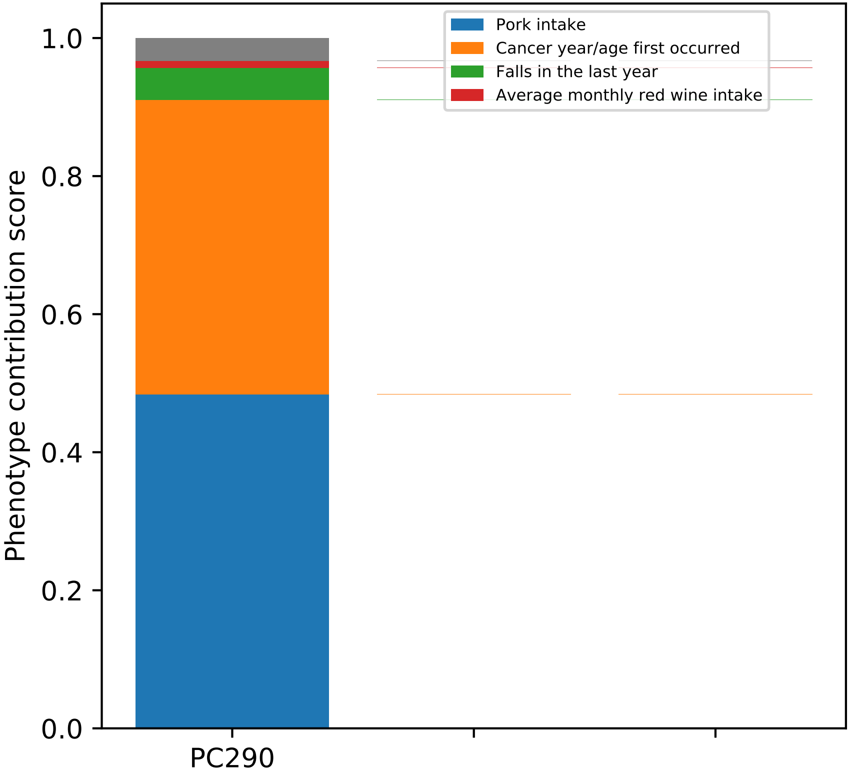

Pork intake
Cancer year/age first occurred
Falls in the last year
Average monthly red wine intake
Phenotype contribution score
1.0
0.8
0.6
0.4
0.2
0.0
PC290

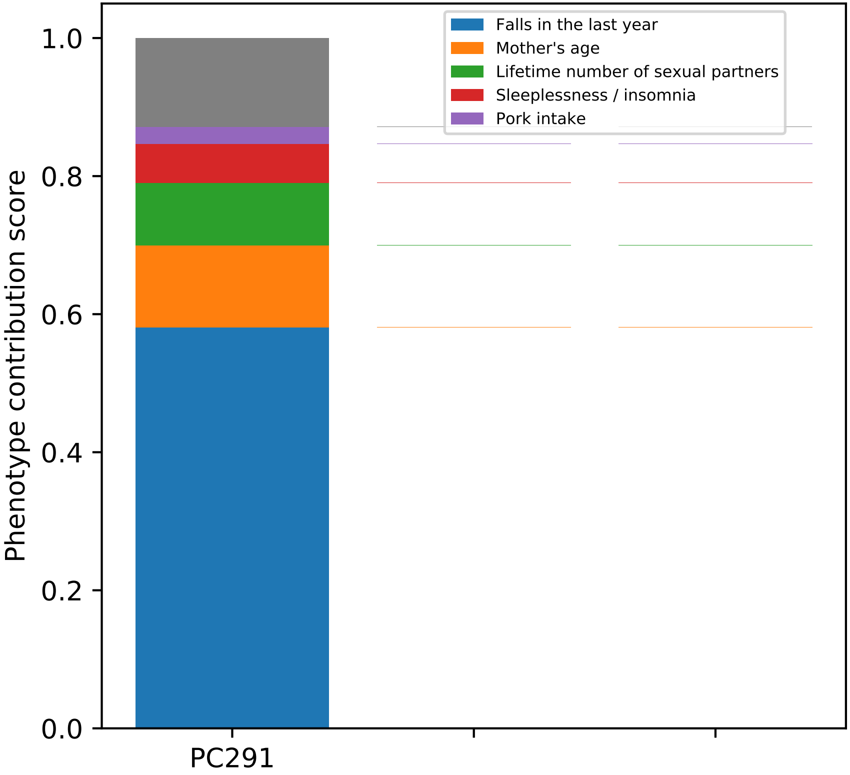
Falls in the last year
Mother's age
Lifetime number of sexual partners
Sleeplessness / insomnia
Pork intake
Phenotype contribution score
1.0
0.8
0.6
0.4
0.2
0.0
PC291

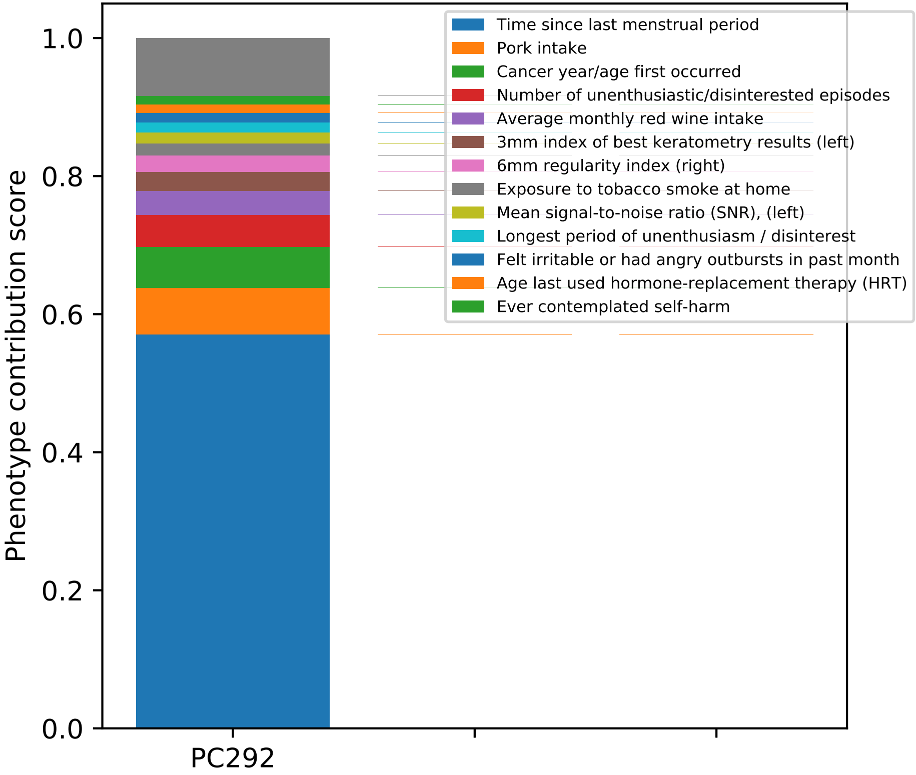

Phenotype contribution score
1.0
0.8
0.6
0.4
0.2
0.0
PC292
Time since last menstrual period
Pork intake
Cancer year/age first occurred
Number of unenthusiastic/disinterested episodes
Average monthly red wine intake
3mm index of best keratometry results (left)
6mm regularity index (right)
Exposure to tobacco smoke at home
Mean signal-to-noise ratio (SNR), (left)
Longest period of unenthusiasm / disinterest
Felt irritable or had angry outbursts in past month
Age last used hormone-replacement therapy (HRT)
Ever contemplated self-harm

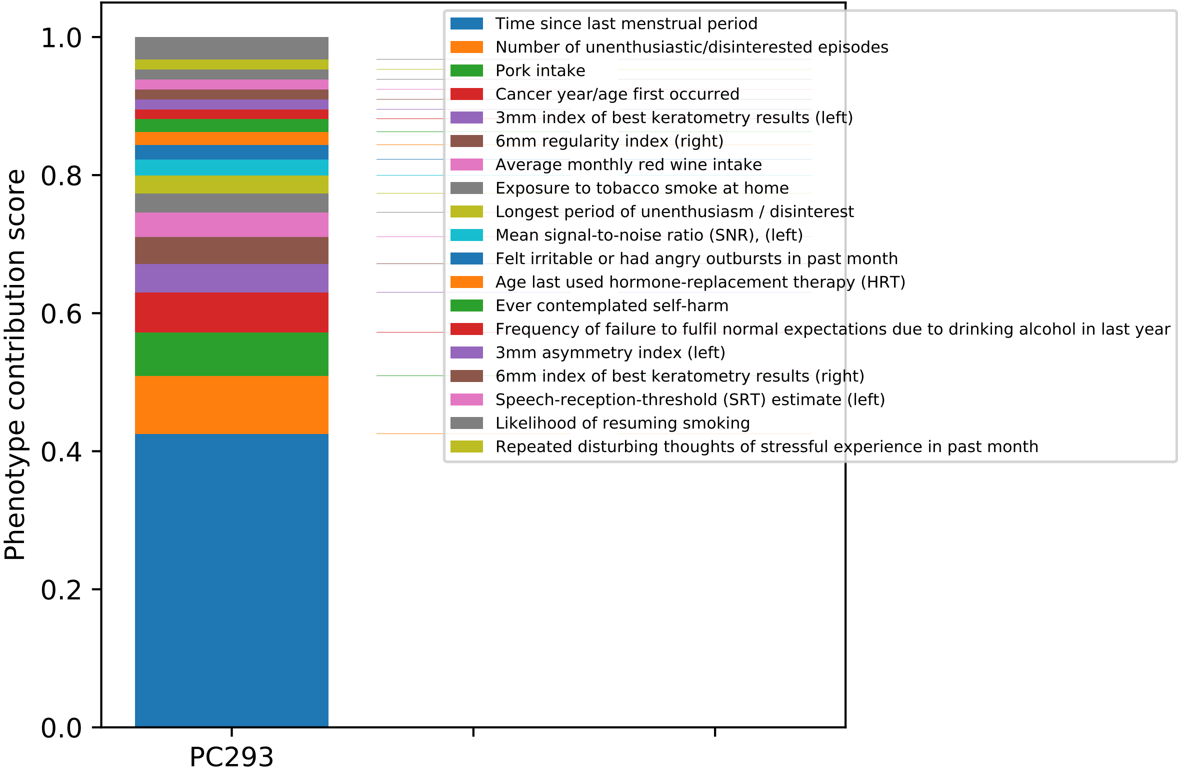
Phenotype contribution score
0.0
0.2
0.4
0.6
0.8
1.0
PC293
Time since last menstrual period
Number of unenthusiastic/disinterested episodes
Pork intake
Cancer year/age first occurred
3mm index of best keratometry results (left)
6mm regularity index (right)
Average monthly red wine intake
Exposure to tobacco smoke at home
Longest period of unenthusiasm / disinterest
Mean signal-to-noise ratio (SNR), (left)
Felt irritable or had angry outbursts in past month
Age last used hormone-replacement therapy (HRT)
Ever contemplated self-harm
Frequency of failure to fulfil normal expectations due to drinking alcohol in last year
3mm asymmetry index (left)
6mm index of best keratometry results (right)
Speech-reception-threshold (SRT) estimate (left)
Likelihood of resuming smoking
Repeated disturbing thoughts of stressful experience in past month

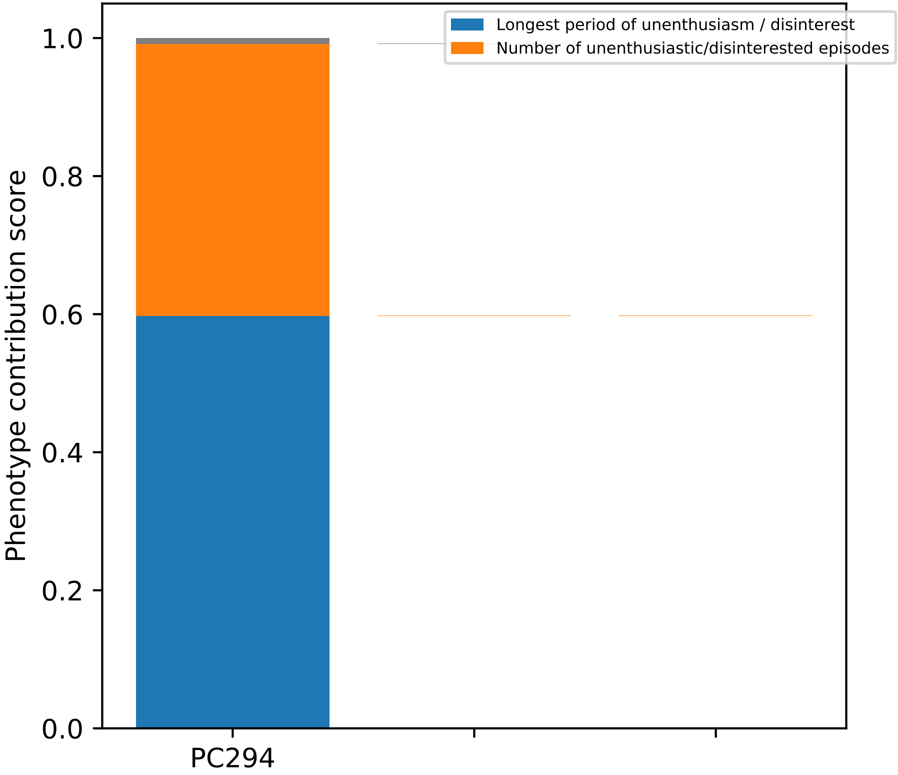
Longest period of unenthusiasm / disinterest
Number of unenthusiastic/disinterested episodes
Phenotype contribution score
1.0
0.8
0.6
0.4
0.2
0.0
PC294

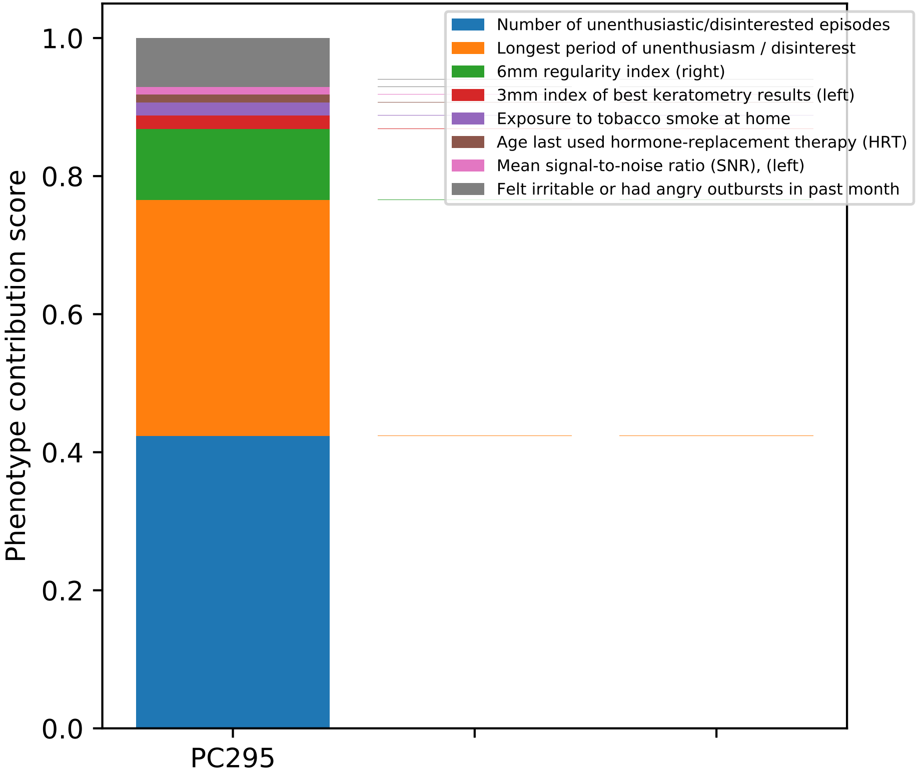
Phenotype contribution score
1.0
0.8
0.6
0.4
0.2
0.0
PC295
Number of unenthusiastic/disinterested episodes
Longest period of unenthusiasm / disinterest
6mm regularity index (right)
3mm index of best keratometry results (left)
Exposure to tobacco smoke at home
Age last used hormone-replacement therapy (HRT)
Mean signal-to-noise ratio (SNR), (left)
Felt irritable or had angry outbursts in past month

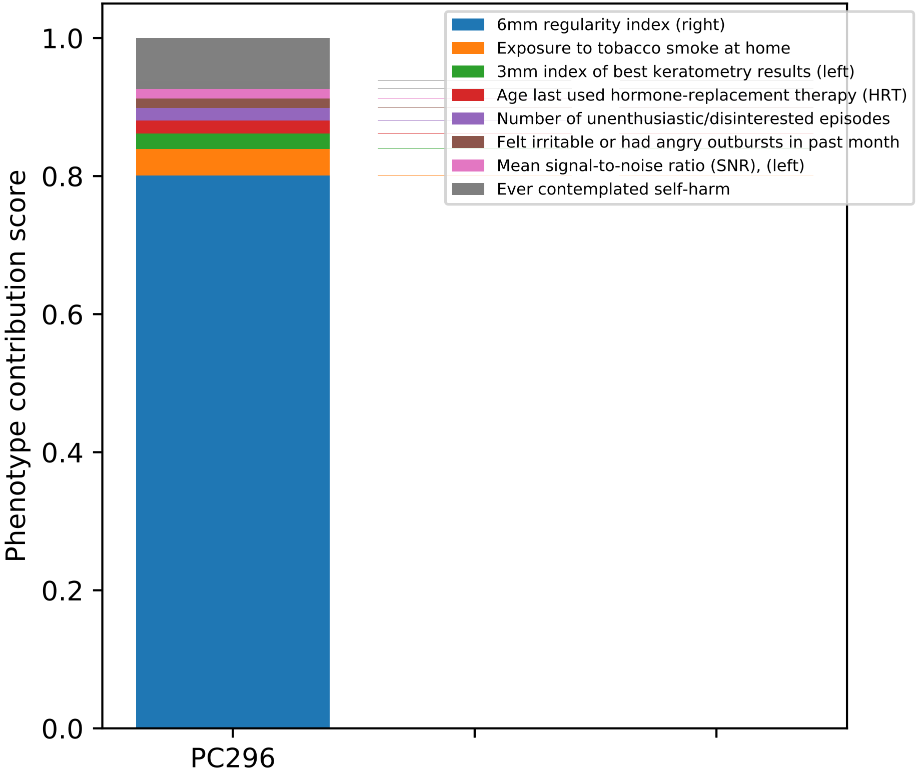

6mm regularity index (right)
Exposure to tobacco smoke at home
3mm index of best keratometry results (left)
Age last used hormone-replacement therapy (HRT)
Number of unenthusiastic/disinterested episodes
Felt irritable or had angry outbursts in past month
Mean signal-to-noise ratio (SNR), (left)
Ever contemplated self-harm
Phenotype contribution score
1.0
0.8
0.6
0.4
0.2
0.0
PC296

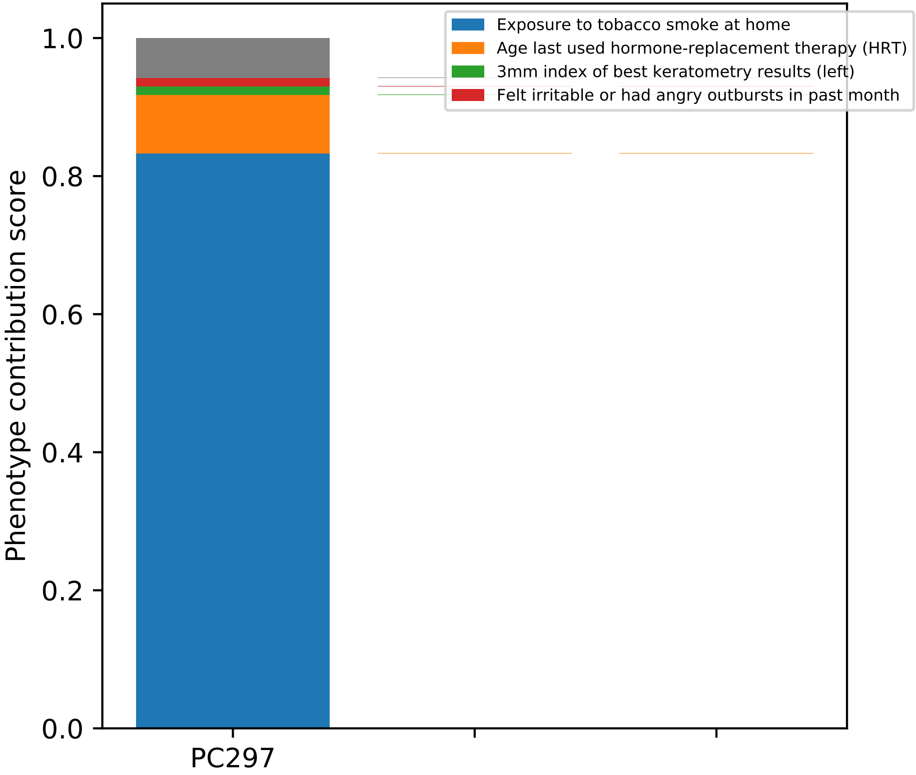

Exposure to tobacco smoke at home
Age last used hormone-replacement therapy (HRT)
3mm index of best keratometry results (left)
Felt irritable or had angry outbursts in past month
Phenotype contribution score
0.0
0.2
0.4
0.6
0.8
1.0
PC297

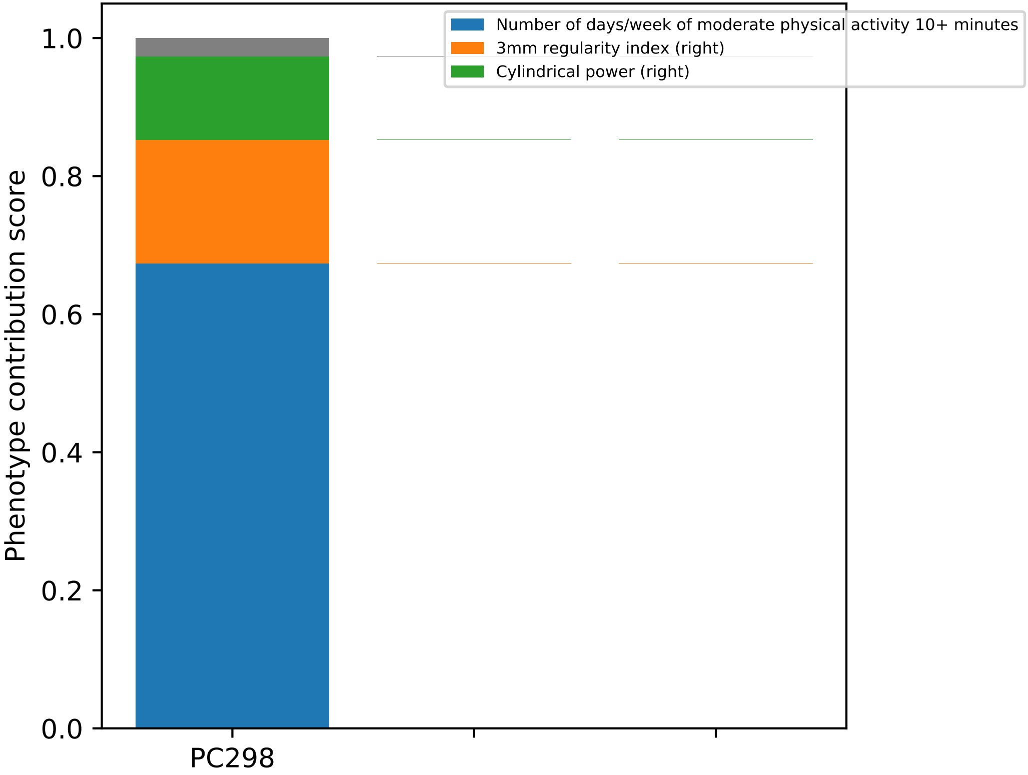

Number of days/week of moderate physical activity 10+ minutes
3mm regularity index (right)
Cylindrical power (right)
Phenotype contribution score
0.0
0.2
0.4
0.6
0.8
1.0
PC298

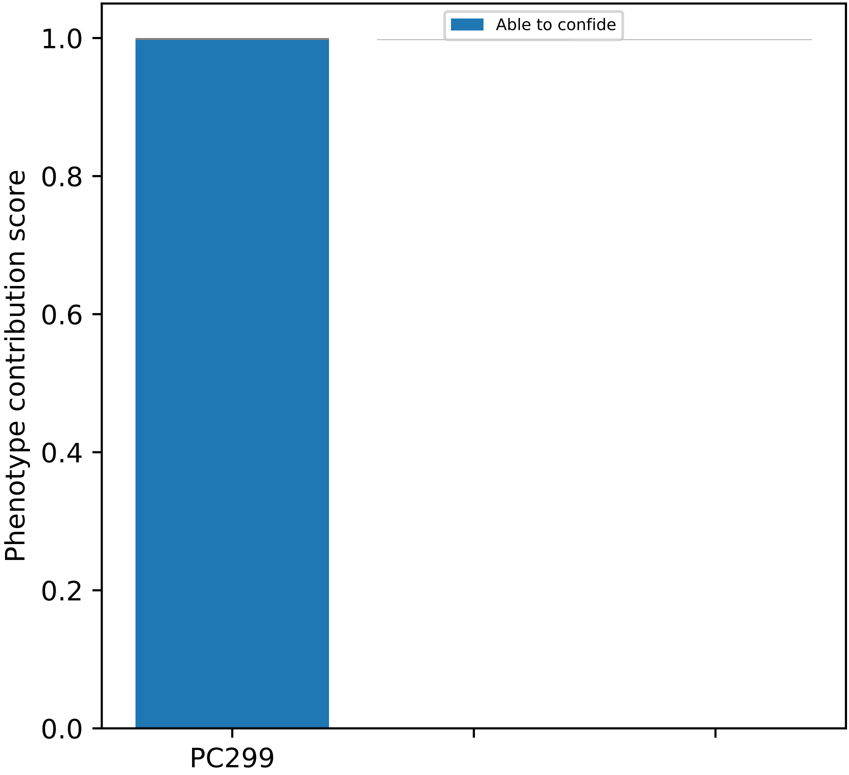
Able to confide
Phenotype contribution score
1.0
0.8
0.6
0.4
0.2
0.0
PC299

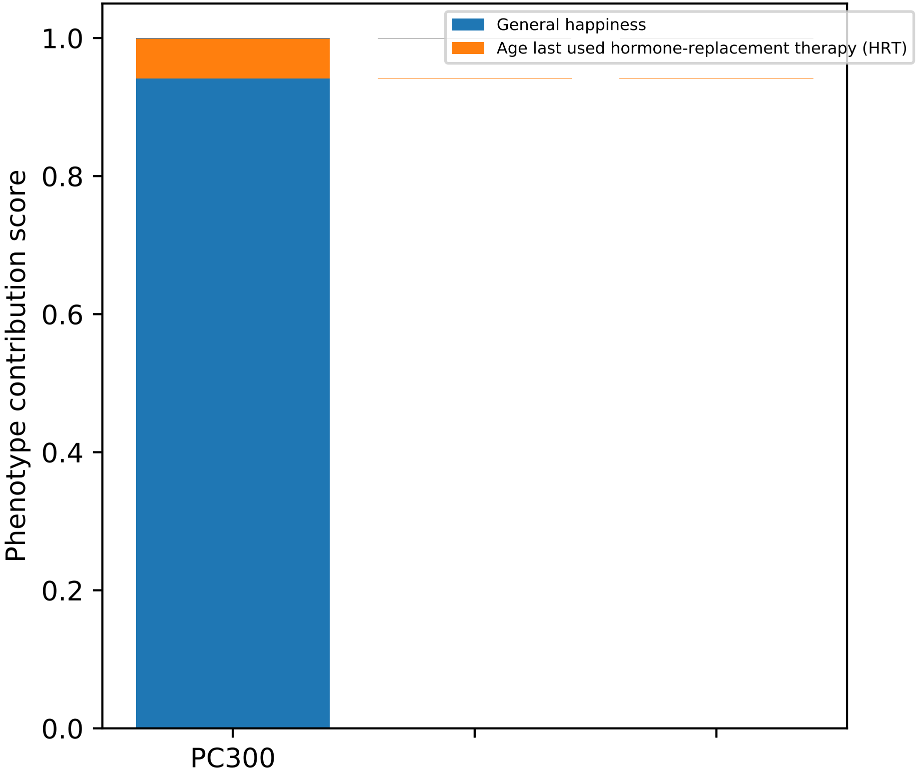

General happiness
Age last used hormone-replacement therapy (HRT)
Phenotype contribution score
1.0
0.8
0.6
0.4
0.2
0.0
PC300

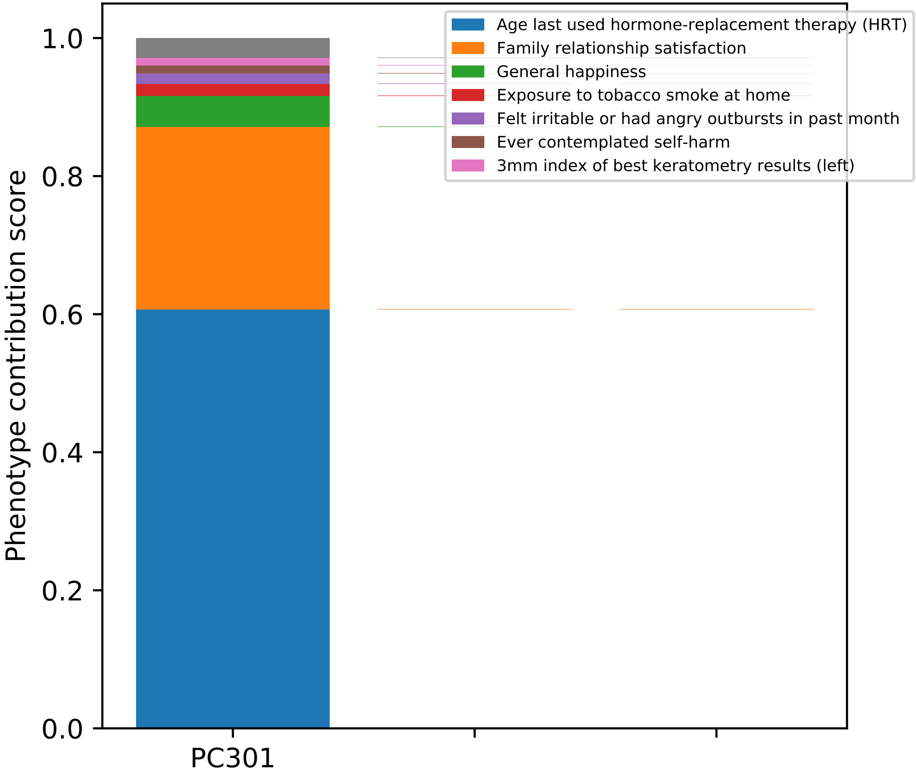

Age last used hormone-replacement therapy (HRT)
Family relationship satisfaction
General happiness
Exposure to tobacco smoke at home
Felt irritable or had angry outbursts in past month
Ever contemplated self-harm
3mm index of best keratometry results (left)
Phenotype contribution score
1.0
0.8
0.6
0.4
0.2
0.0
PC301

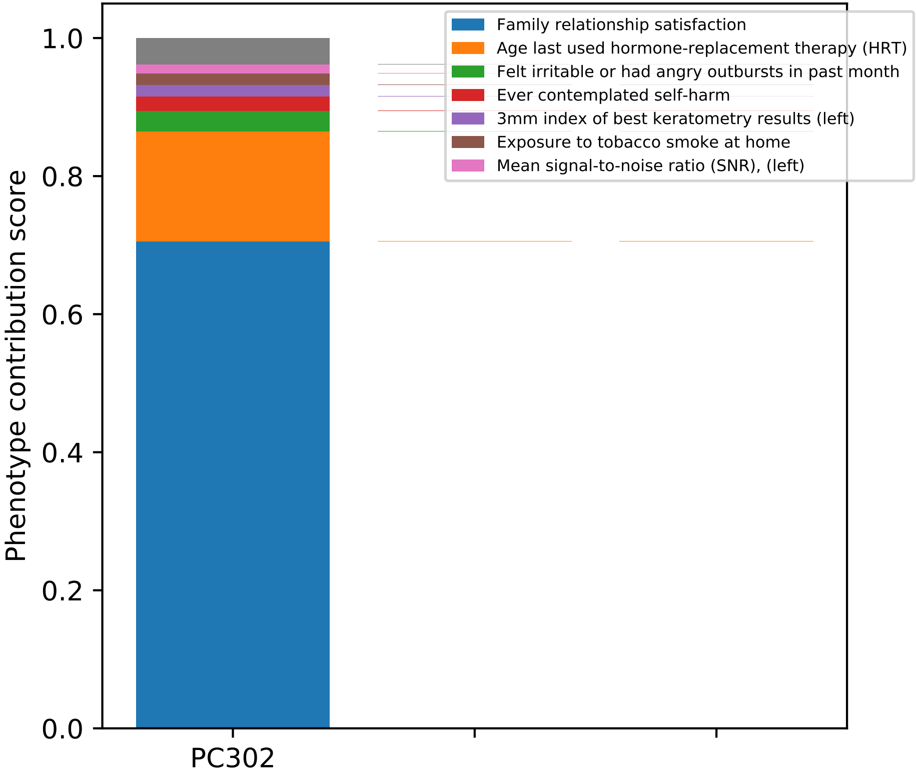

Phenotype contribution score
1.0
0.8
0.6
0.4
0.2
0.0
PC302
Family relationship satisfaction
Age last used hormone-replacement therapy (HRT)
Felt irritable or had angry outbursts in past month
Ever contemplated self-harm
3mm index of best keratometry results (left)
Exposure to tobacco smoke at home
Mean signal-to-noise ratio (SNR), (left)

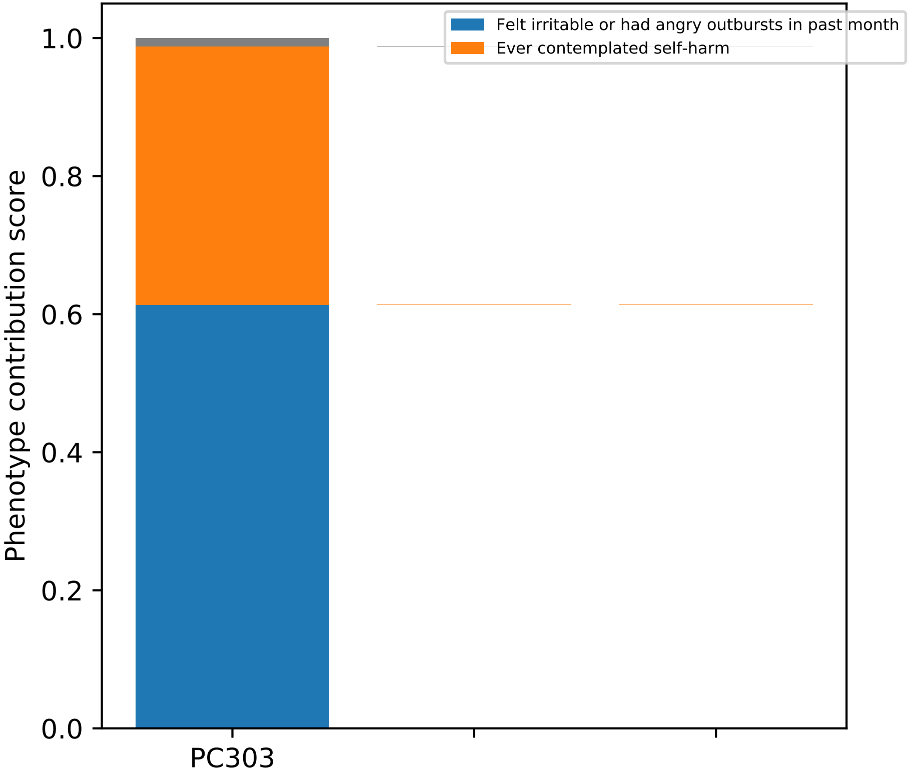
Felt irritable or had angry outbursts in past month
Ever contemplated self-harm
Phenotype contribution score
1.0
0.8
0.6
0.4
0.2
0.0
PC303

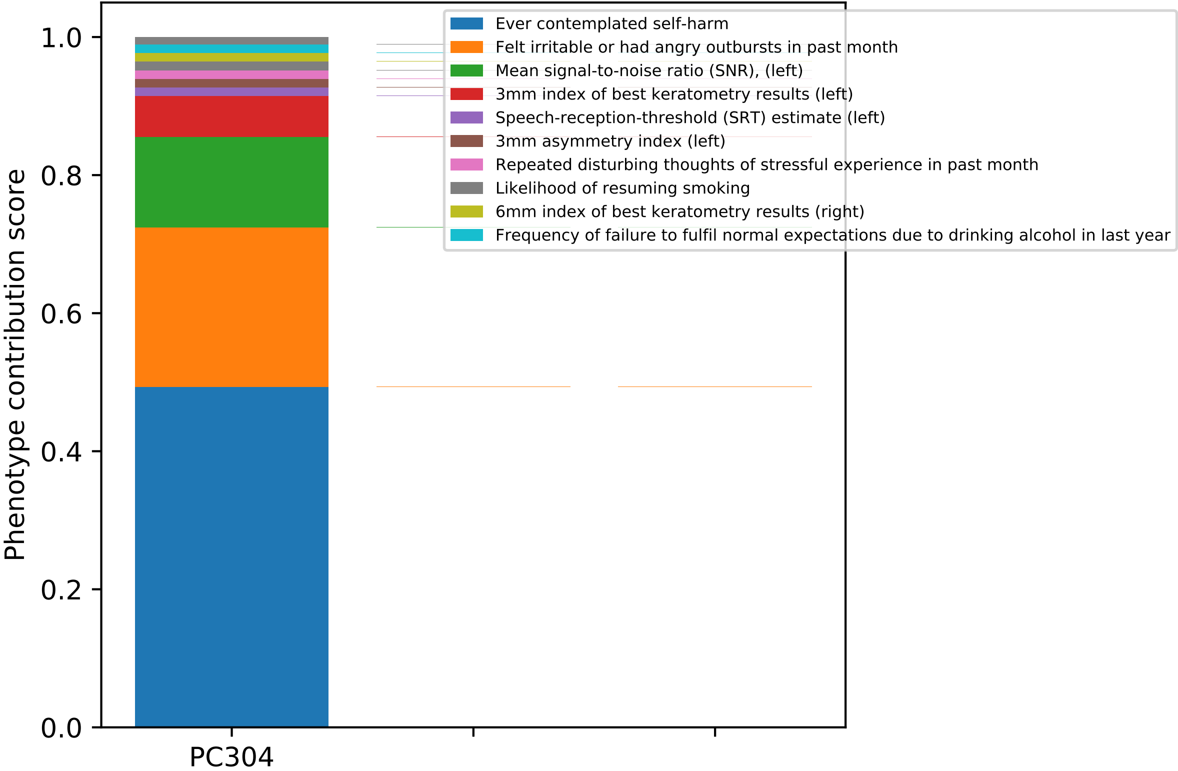

Phenotype contribution score
1.0
0.8
0.6
0.4
0.2
0.0
PC304
Ever contemplated self-harm
Felt irritable or had angry outbursts in past month
Mean signal-to-noise ratio (SNR), (left)
3mm index of best keratometry results (left)
Speech-reception-threshold (SRT) estimate (left)
3mm asymmetry index (left)
Repeated disturbing thoughts of stressful experience in past month
Likelihood of resuming smoking
6mm index of best keratometry results (right)
Frequency of failure to fulfil normal expectations due to drinking alcohol in last year

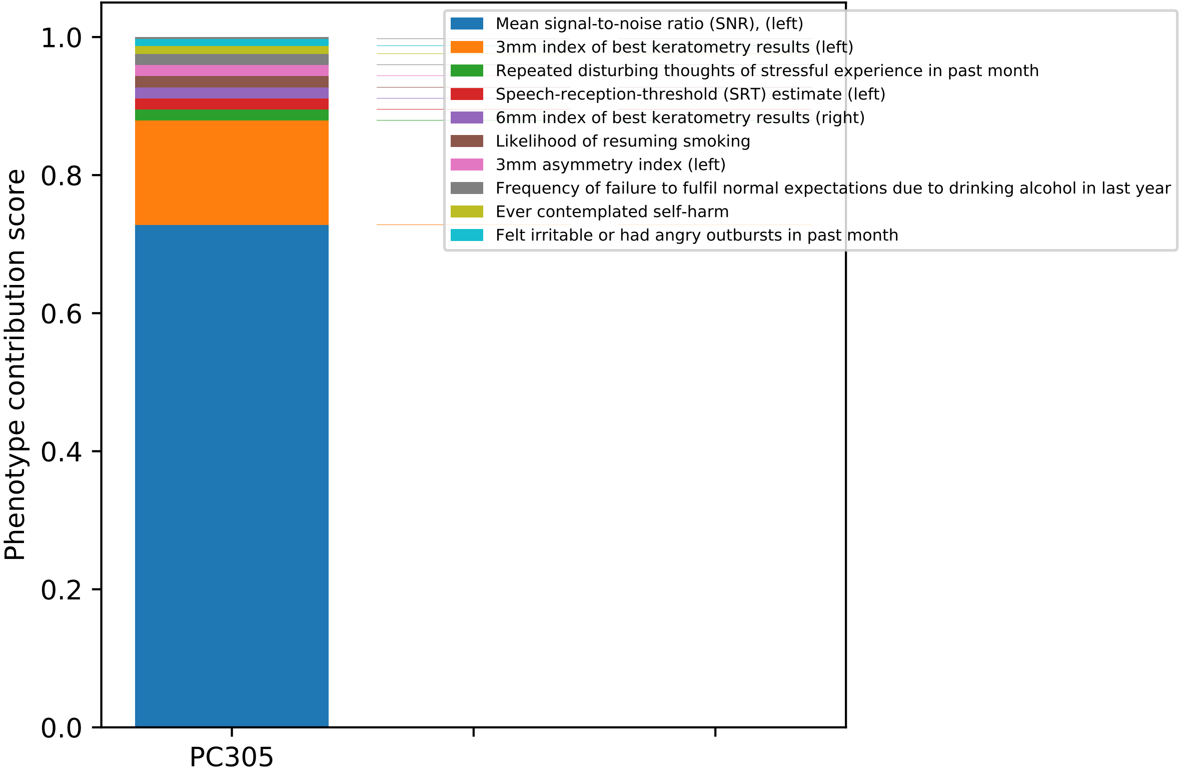

Phenotype contribution score
1.0
0.8
0.6
0.4
0.2
0.0
PC305
Mean signal-to-noise ratio (SNR), (left)
3mm index of best keratometry results (left)
Repeated disturbing thoughts of stressful experience in past month
Speech-reception-threshold (SRT) estimate (left)
6mm index of best keratometry results (right)
Likelihood of resuming smoking
3mm asymmetry index (left)
Frequency of failure to fulfil normal expectations due to drinking alcohol in last year
Ever contemplated self-harm
Felt irritable or had angry outbursts in past month

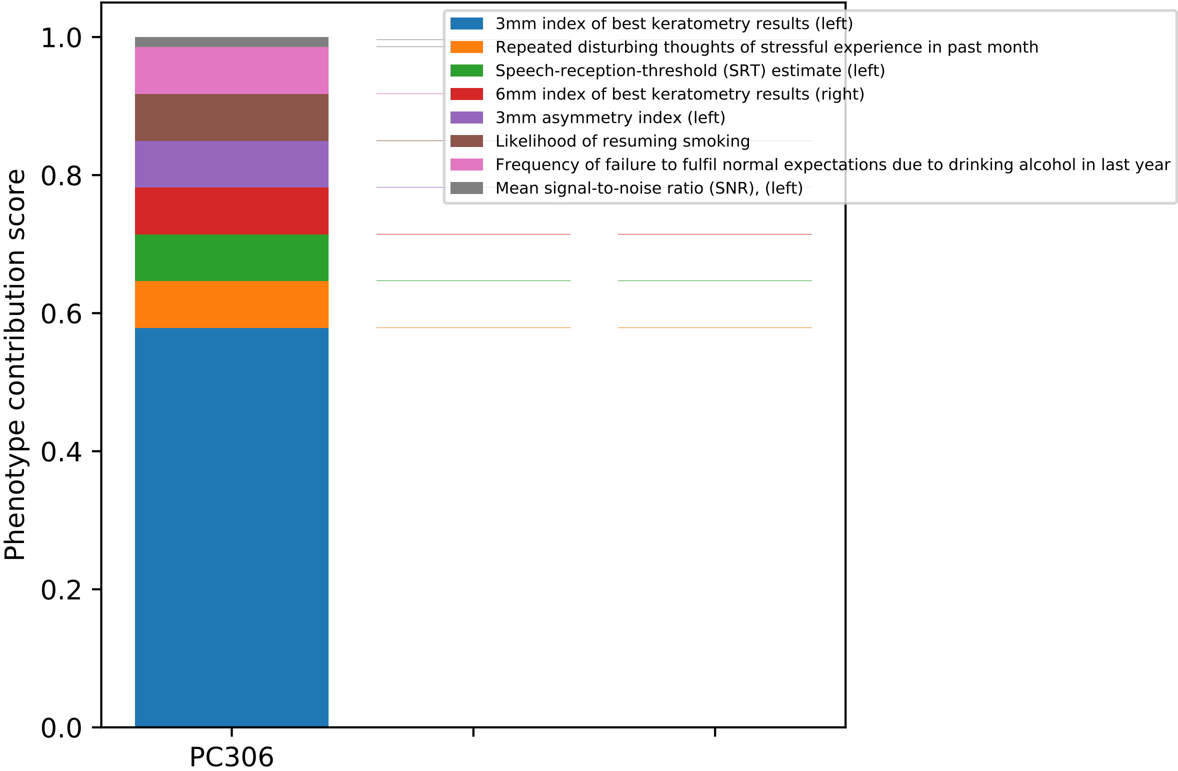

3mm index of best keratometry results (left)
Repeated disturbing thoughts of stressful experience in past month
Speech-reception-threshold (SRT) estimate (left)
6mm index of best keratometry results (right)
3mm asymmetry index (left)
Likelihood of resuming smoking
Frequency of failure to fulfil normal expectations due to drinking alcohol in last year
Mean signal-to-noise ratio (SNR), (left)
Phenotype contribution score
0.0
0.2
0.4
0.6
0.8
1.0
PC306

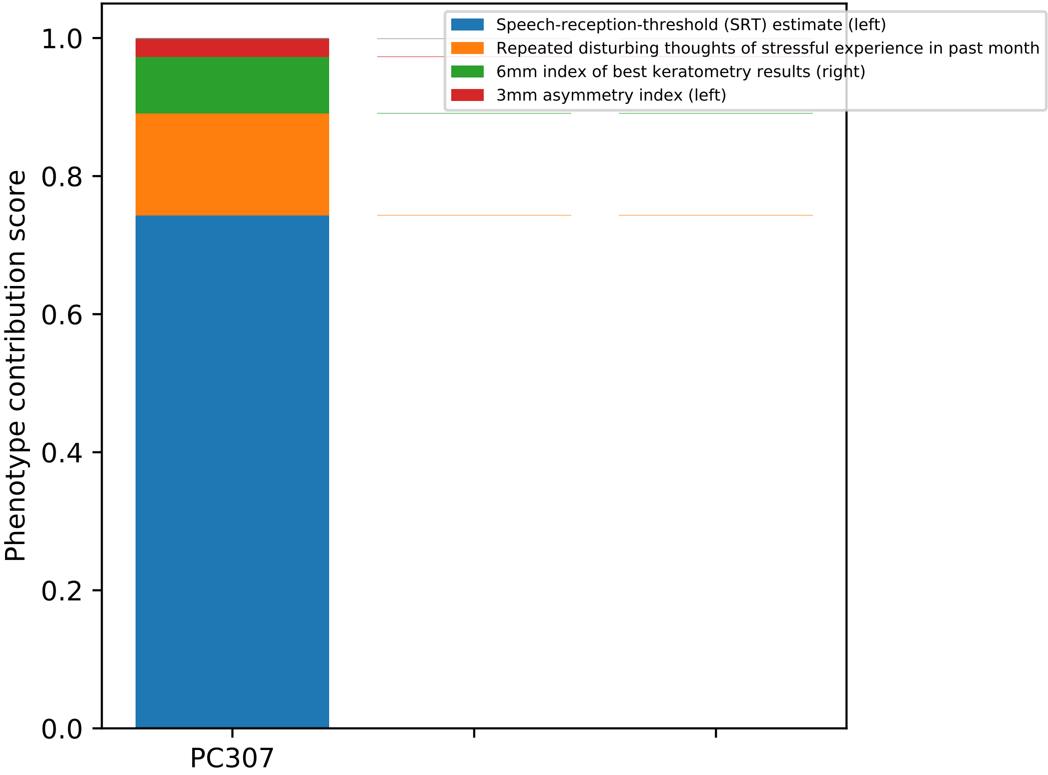
Speech-reception-threshold (SRT) estimate (left)
Repeated disturbing thoughts of stressful experience in past month
6mm index of best keratometry results (right)
3mm asymmetry index (left)
Phenotype contribution score
1.0
0.8
0.6
0.4
0.2
0.0
PC307

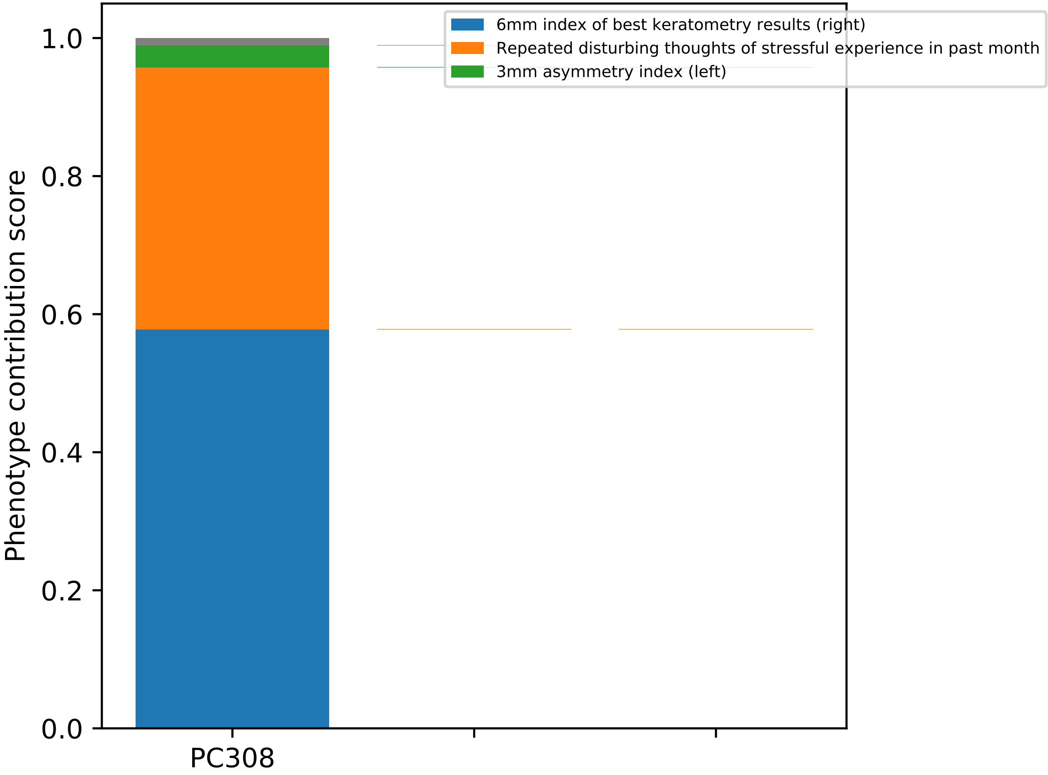

Phenotype contribution score
0.0
0.2
0.4
0.6
0.8
1.0
PC308
6mm index of best keratometry results (right)
Repeated disturbing thoughts of stressful experience in past month
3mm asymmetry index (left)

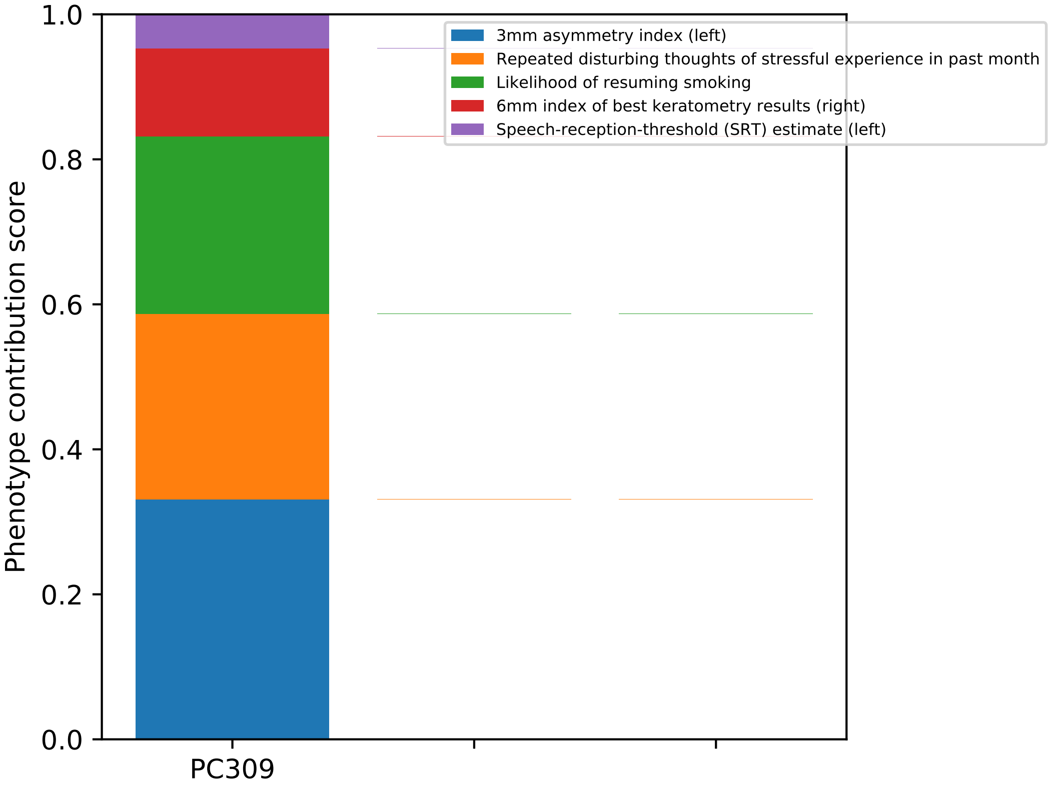

1.0
0.8
0.6
0.4
0.2
0.0
Phenotype contribution score
PC309
3mm asymmetry index (left)
Repeated disturbing thoughts of stressful experience in past month
Likelihood of resuming smoking
6mm index of best keratometry results (right)
Speech-reception-threshold (SRT) estimate (left)

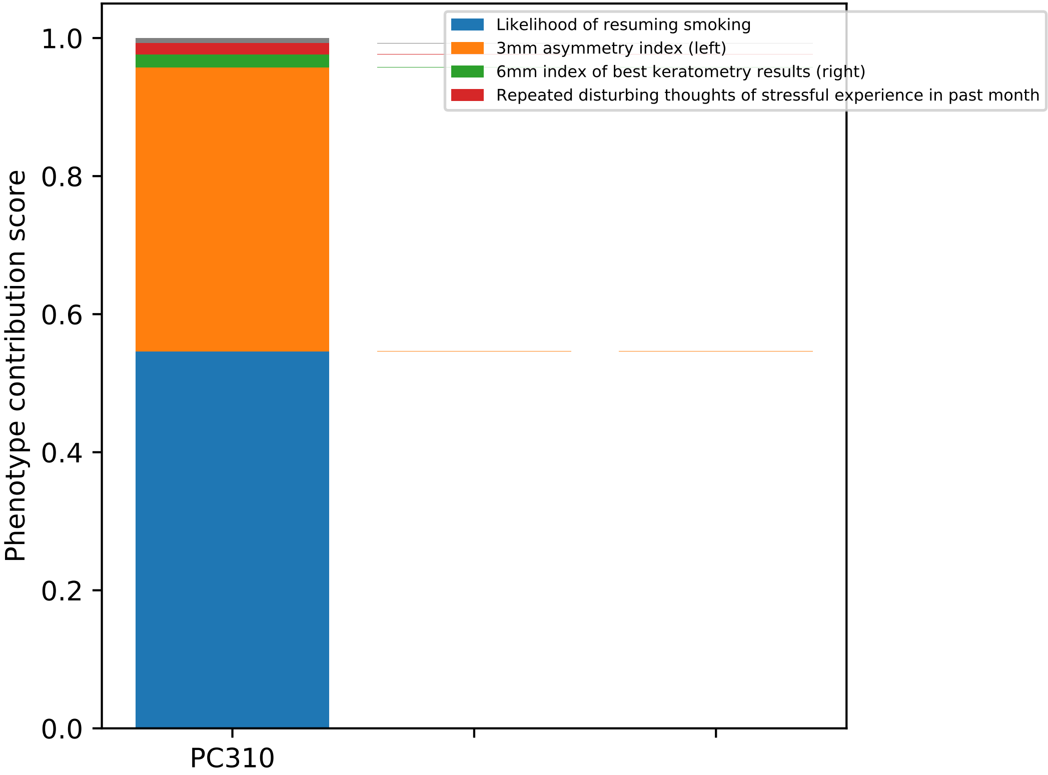

Likelihood of resuming smoking
3mm asymmetry index (left)
6mm index of best keratometry results (right)
Repeated disturbing thoughts of stressful experience in past month
Phenotype contribution score
1.0
0.8
0.6
0.4
0.2
0.0
PC310

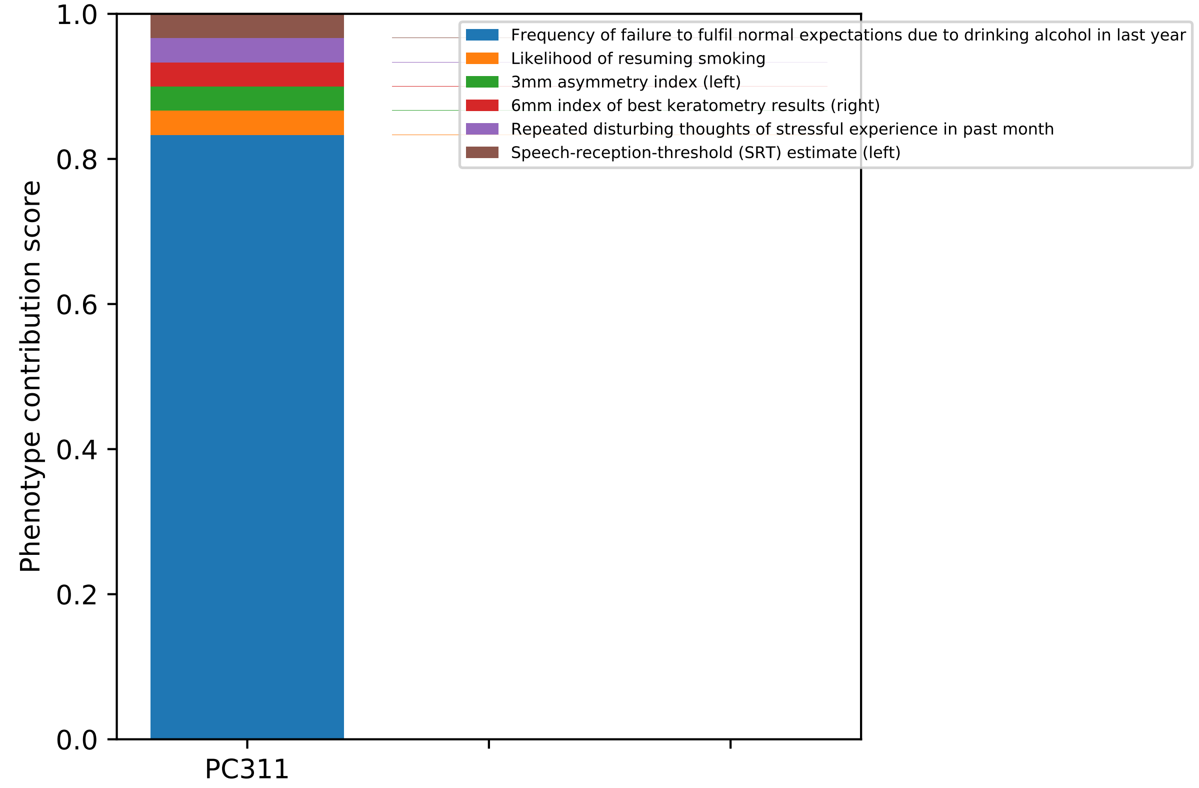

1.0
0.8
0.6
0.4
0.2
0.0
Phenotype contribution score
PC311
Frequency of failure to fulfil normal expectations due to drinking alcohol in last year
Likelihood of resuming smoking
3mm asymmetry index (left)
6mm index of best keratometry results (right)
Repeated disturbing thoughts of stressful experience in past month
Speech-reception-threshold (SRT) estimate (left)

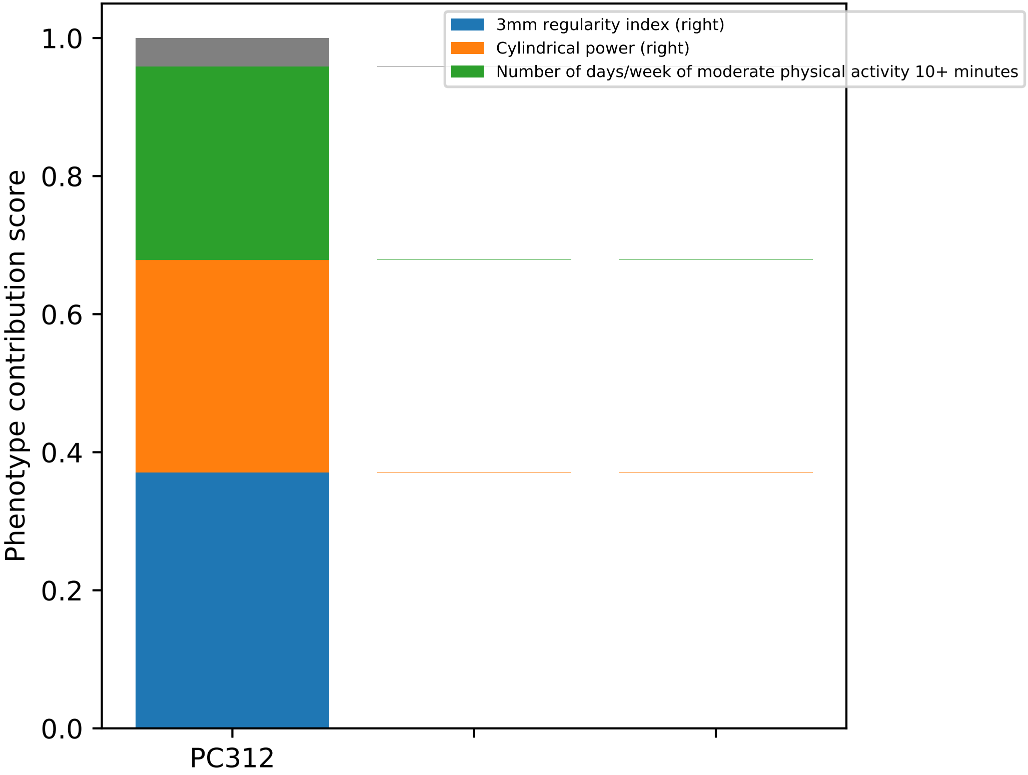
3mm regularity index (right)
Cylindrical power (right)
Number of days/week of moderate physical activity 10+ minutes
Phenotype contribution score
0.0
0.2
0.4
0.6
0.8
1.0
PC312

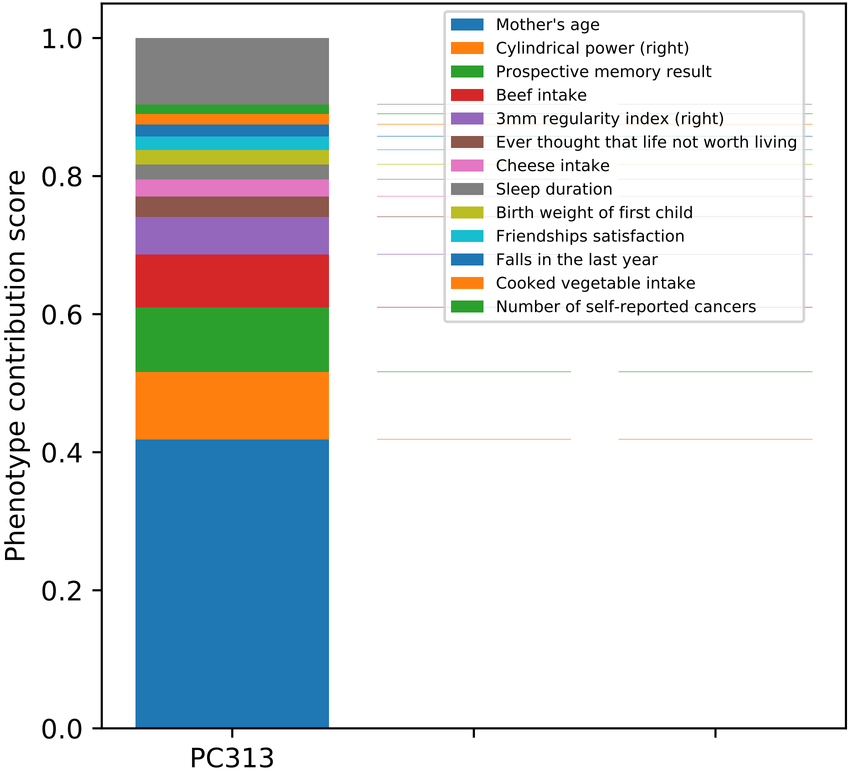
Phenotype contribution score
0.0
0.2
0.4
0.6
0.8
1.0
PC313
Mother's age
Cylindrical power (right)
Prospective memory result
Beef intake
3mm regularity index (right)
Ever thought that life not worth living
Cheese intake
Sleep duration
Birth weight of first child
Friendships satisfaction
Falls in the last year
Cooked vegetable intake
Number of self-reported cancers

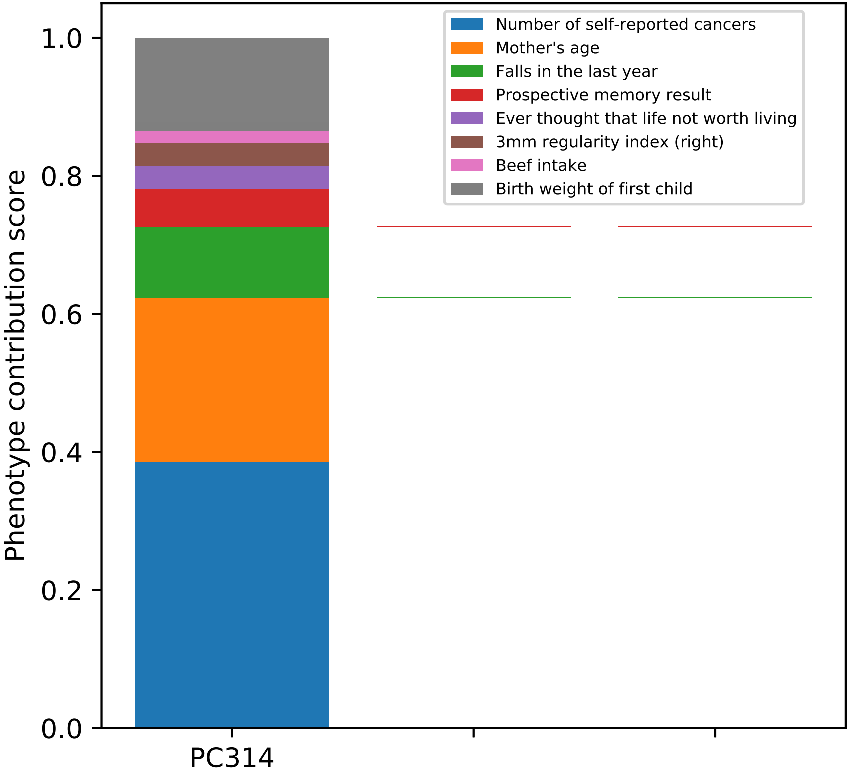

Number of self-reported cancers
Mother's age
Falls in the last year
Prospective memory result
Ever thought that life not worth living
3mm regularity index (right)
Beef intake
Birth weight of first child
Phenotype contribution score
1.0
0.8
0.6
0.4
0.2
0.0
PC314

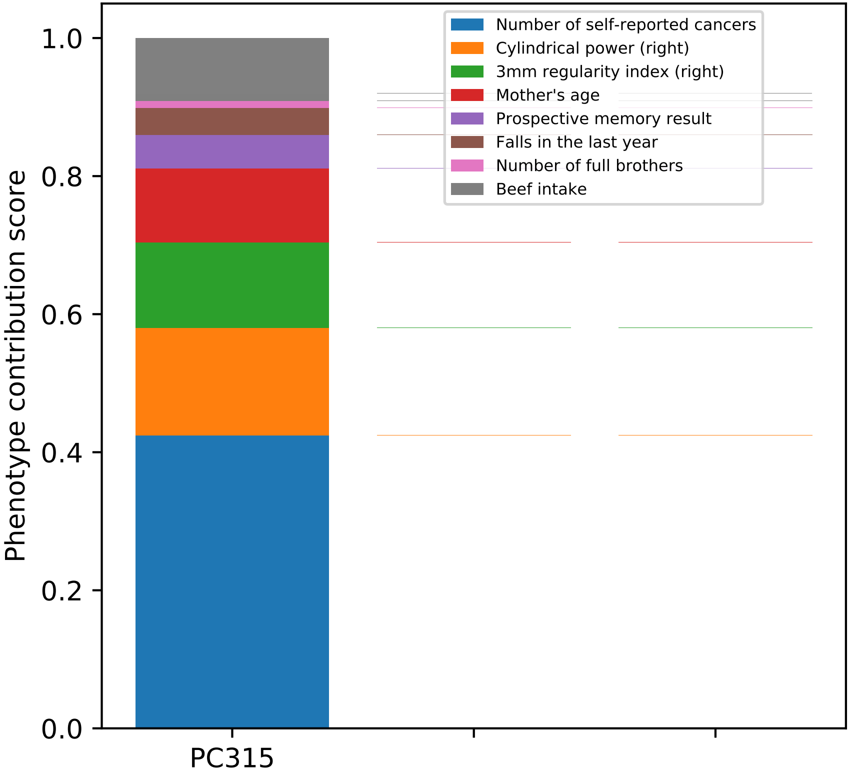
Number of self-reported cancers
Cylindrical power (right)
3mm regularity index (right)
Mother's age
Prospective memory result
Falls in the last year
Number of full brothers
Beef intake
Phenotype contribution score
0.0
0.2
0.4
0.6
0.8
1.0
PC315

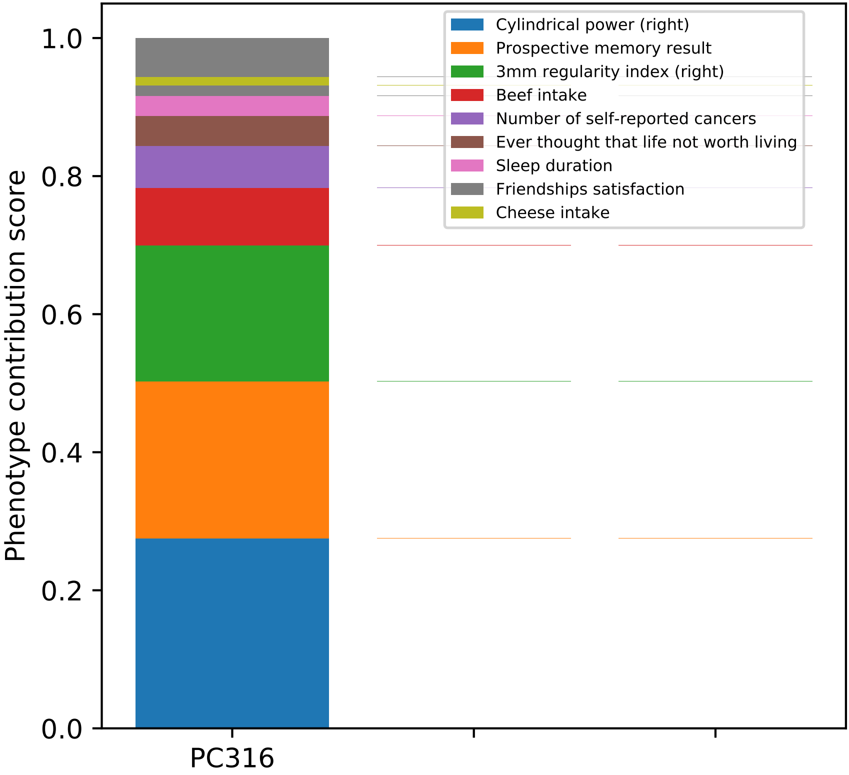

Phenotype contribution score
1.0
0.8
0.6
0.4
0.2
0.0
PC316
Cylindrical power (right)
Prospective memory result
3mm regularity index (right)
Beef intake
Number of self-reported cancers
Ever thought that life not worth living
Sleep duration
Friendships satisfaction
Cheese intake

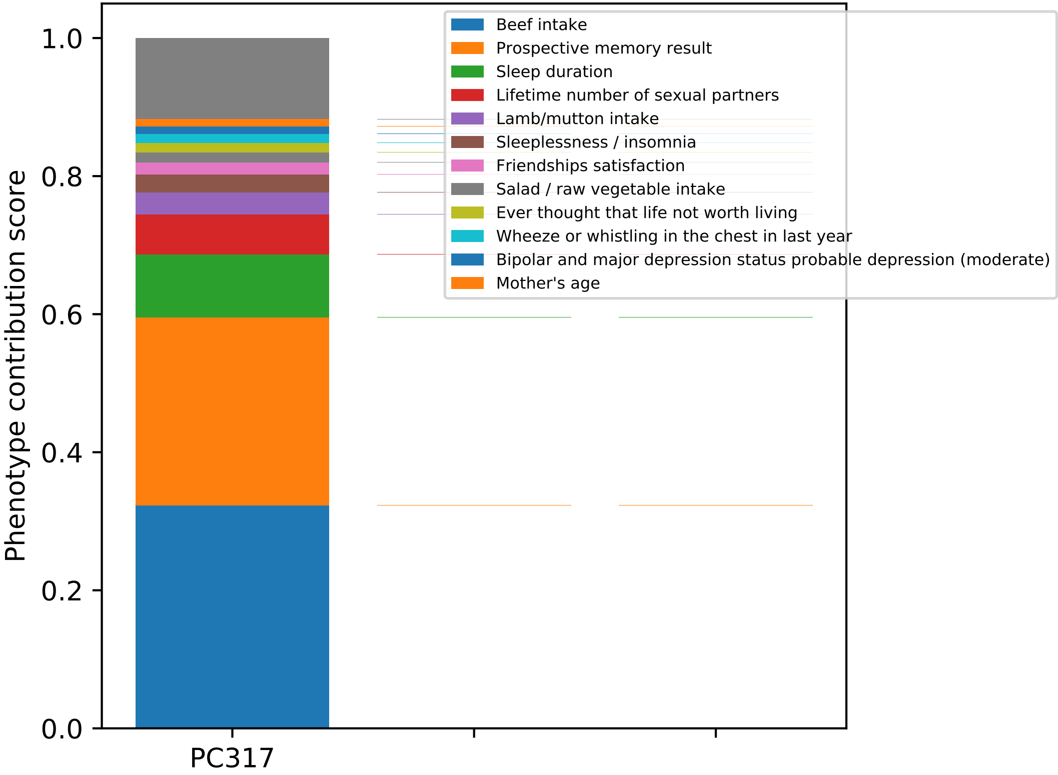
Phenotype contribution score
0.0
0.2
0.4
0.6
0.8
1.0
PC317
Beef intake
Prospective memory result
Sleep duration
Lifetime number of sexual partners
Lamb/mutton intake
Sleeplessness / insomnia
Friendships satisfaction
Salad / raw vegetable intake
Ever thought that life not worth living
Wheeze or whistling in the chest in last year
Bipolar and major depression status probable depression (moderate)
Mother's age

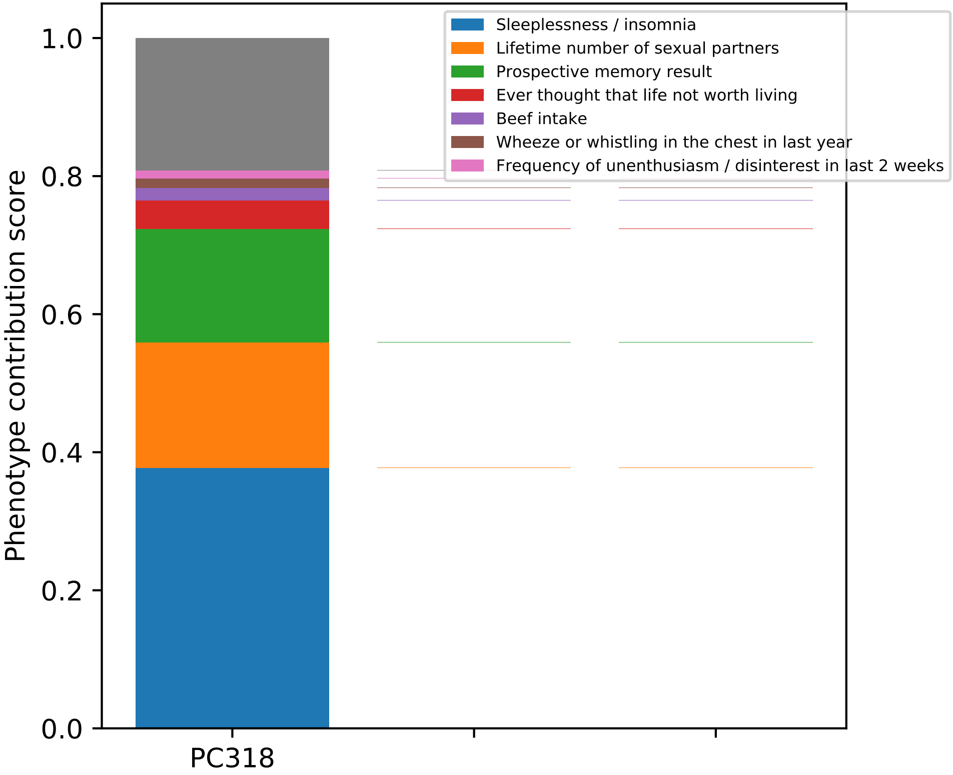

Sleeplessness / insomnia
Lifetime number of sexual partners
Prospective memory result
Ever thought that life not worth living
Beef intake
Wheeze or whistling in the chest in last year
Frequency of unenthusiasm / disinterest in last 2 weeks
Phenotype contribution score
0.0
0.2
0.4
0.6
0.8
1.0
PC318

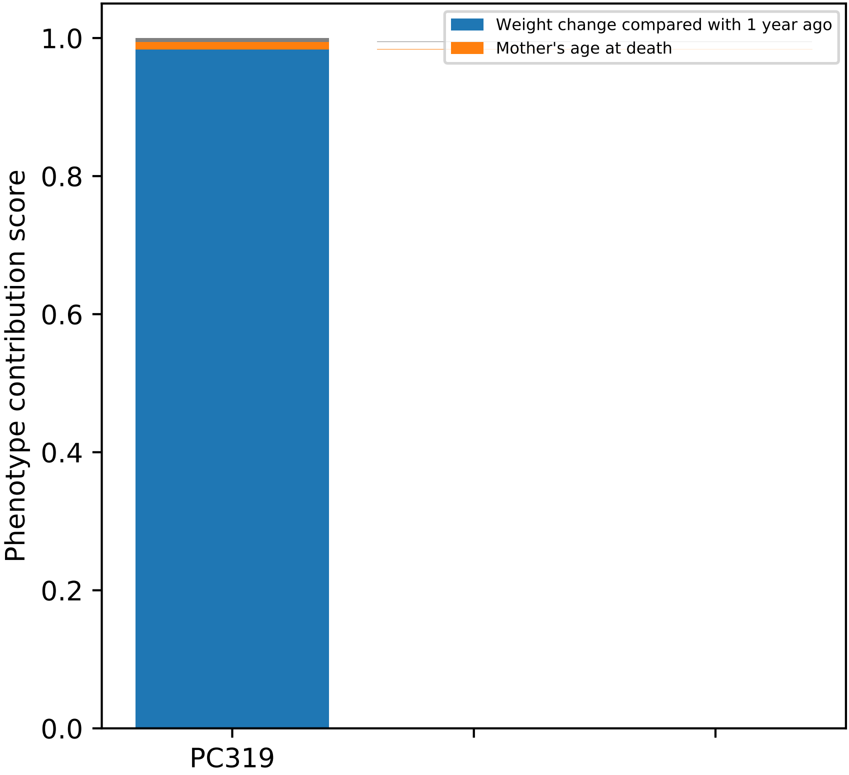

Weight change compared with 1 year ago
Mother's age at death
Phenotype contribution score
1.0
0.8
0.6
0.4
0.2
0.0
PC319

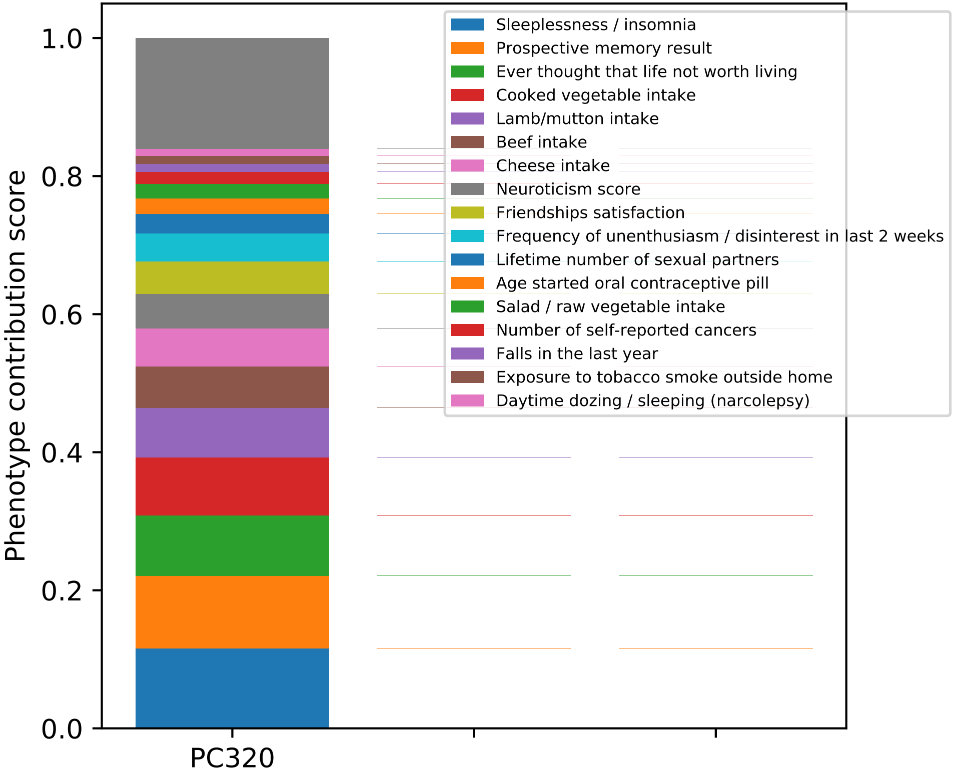

Phenotype contribution score
0.0
0.2
0.4
0.6
0.8
1.0
PC320
Sleeplessness / insomnia
Prospective memory result
Ever thought that life not worth living
Cooked vegetable intake
Lamb/mutton intake
Beef intake
Cheese intake
Neuroticism score
Friendships satisfaction
Frequency of unenthusiasm / disinterest in last 2 weeks
Lifetime number of sexual partners
Age started oral contraceptive pill
Salad / raw vegetable intake
Number of self-reported cancers
Falls in the last year
Exposure to tobacco smoke outside home
Daytime dozing / sleeping (narcolepsy)

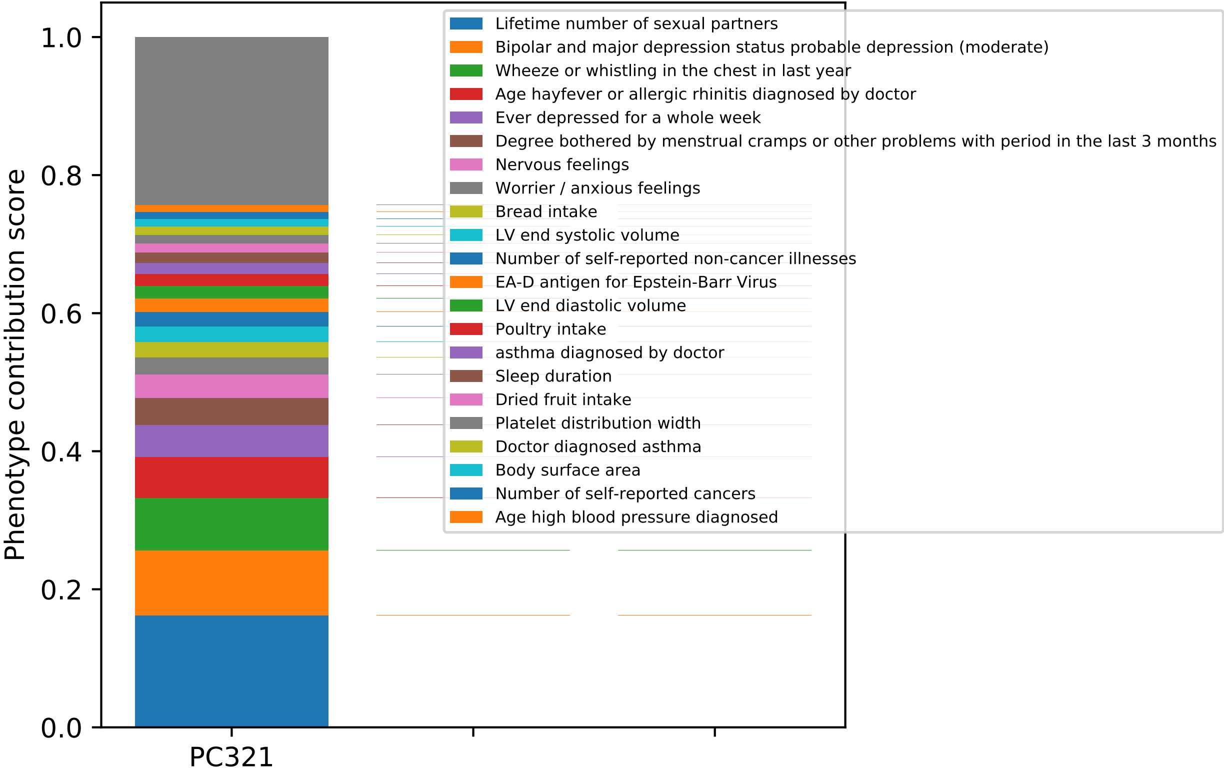

Phenotype contribution score
1.0
0.8
0.6
0.4
0.2
0.0
PC321
Lifetime number of sexual partners
Bipolar and major depression status probable depression (moderate)
Wheeze or whistling in the chest in last year
Age hayfever or allergic rhinitis diagnosed by doctor
Ever depressed for a whole week
Degree bothered by menstrual cramps or other problems with period in the last 3 months
Nervous feelings
Worrier / anxious feelings
Bread intake
LV end systolic volume
Number of self-reported non-cancer illnesses
EA-D antigen for Epstein-Barr Virus
LV end diastolic volume
Poultry intake
asthma diagnosed by doctor
Sleep duration
Dried fruit intake
Platelet distribution width
Doctor diagnosed asthma
Body surface area
Number of self-reported cancers
Age high blood pressure diagnosed

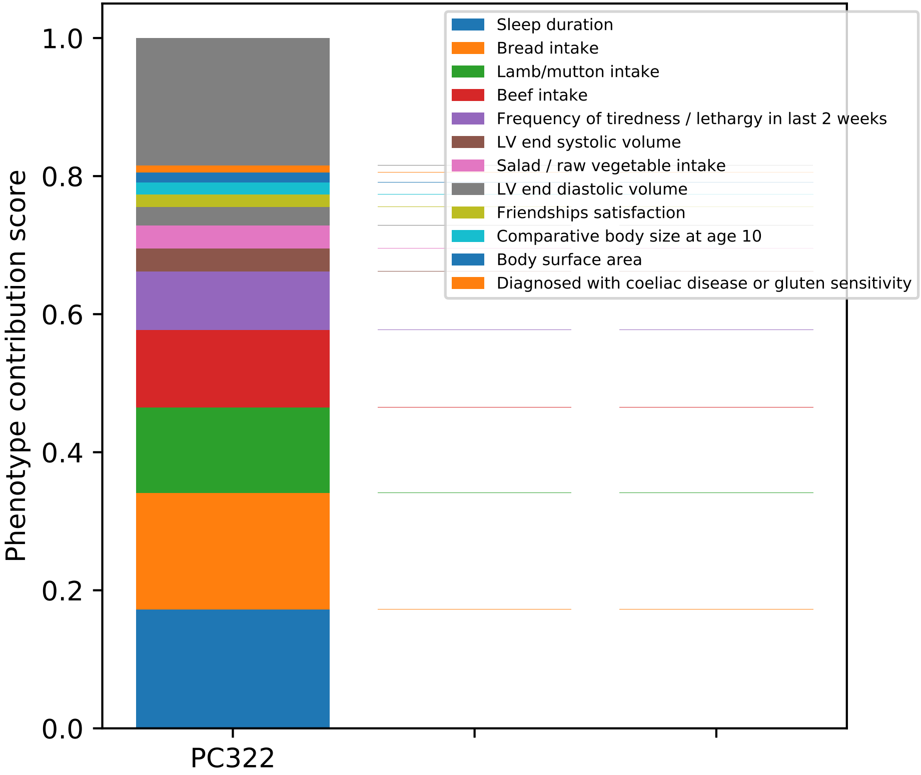
Phenotype contribution score
1.0
0.8
0.6
0.4
0.2
0.0
PC322
Sleep duration
Bread intake
Lamb/mutton intake
Beef intake
Frequency of tiredness / lethargy in last 2 weeks
LV end systolic volume
Salad / raw vegetable intake
LV end diastolic volume
Friendships satisfaction
Comparative body size at age 10
Body surface area
Diagnosed with coeliac disease or gluten sensitivity

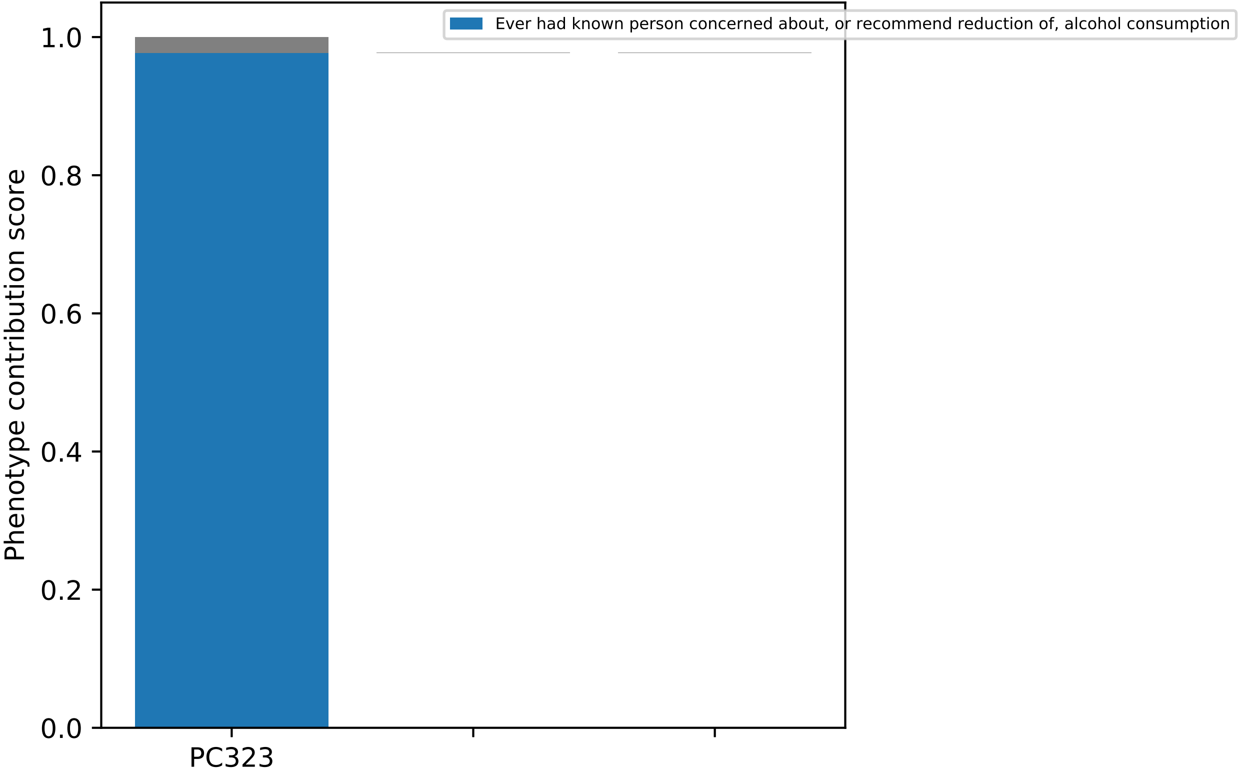

Ever had known person concerned about, or recommend reduction of, alcohol consumption
Phenotype contribution score
1.0
0.8
0.6
0.4
0.2
0.0
PC323

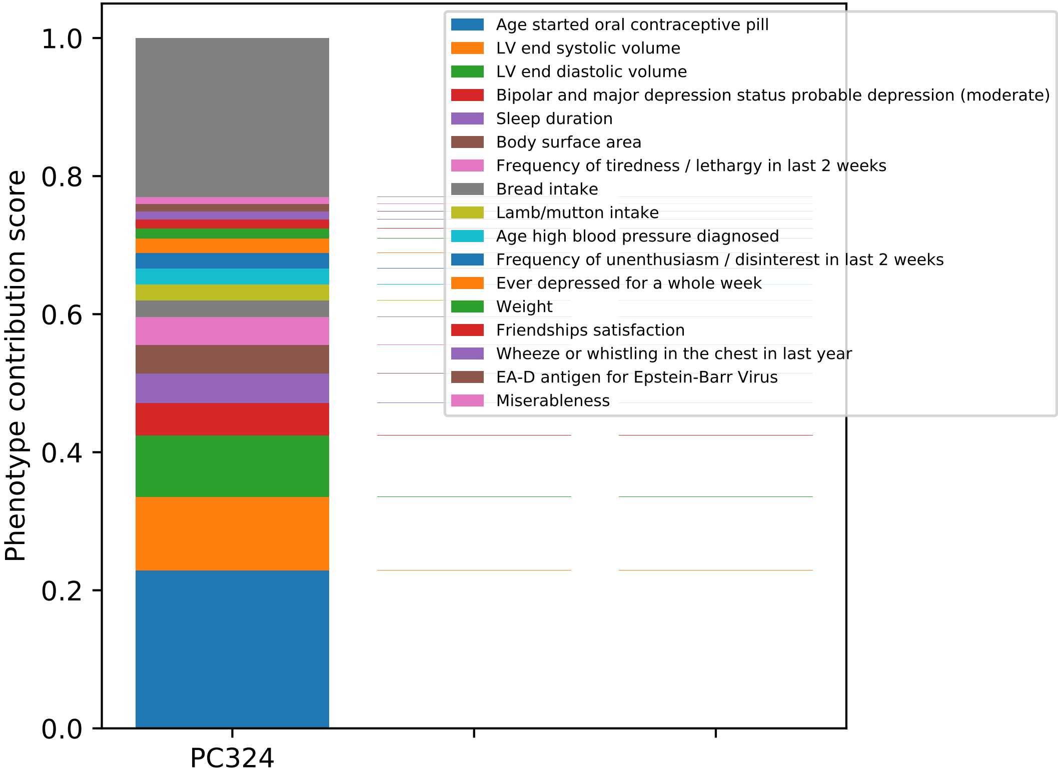

Phenotype contribution score
1.0
0.8
0.6
0.4
0.2
0.0
PC324
Age started oral contraceptive pill
LV end systolic volume
LV end diastolic volume
Bipolar and major depression status probable depression (moderate)
Sleep duration
Body surface area
Frequency of tiredness / lethargy in last 2 weeks
Bread intake
Lamb/mutton intake
Age high blood pressure diagnosed
Frequency of unenthusiasm / disinterest in last 2 weeks
Ever depressed for a whole week
Weight
Friendships satisfaction
Wheeze or whistling in the chest in last year
EA-D antigen for Epstein-Barr Virus
Miserableness

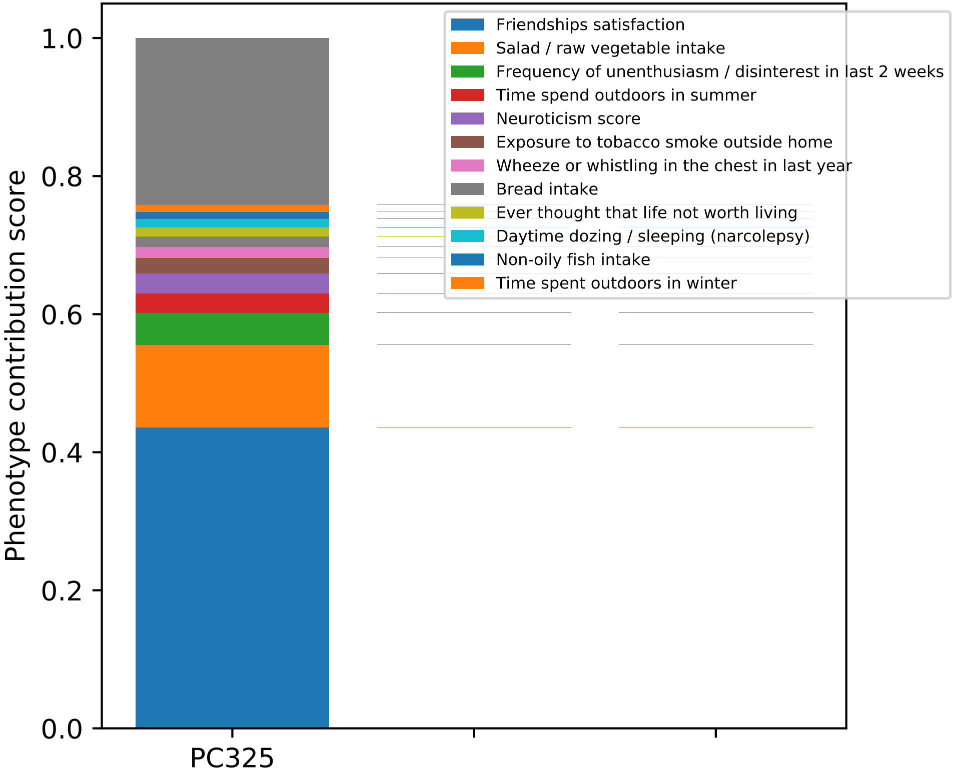

Phenotype contribution score
1.0
0.8
0.6
0.4
0.2
0.0
PC325
Friendships satisfaction
Salad / raw vegetable intake
Frequency of unenthusiasm / disinterest in last 2 weeks
Time spend outdoors in summer
Neuroticism score
Exposure to tobacco smoke outside home
Wheeze or whistling in the chest in last year
Bread intake
Ever thought that life not worth living
Daytime dozing / sleeping (narcolepsy)
Non-oily fish intake
Time spent outdoors in winter

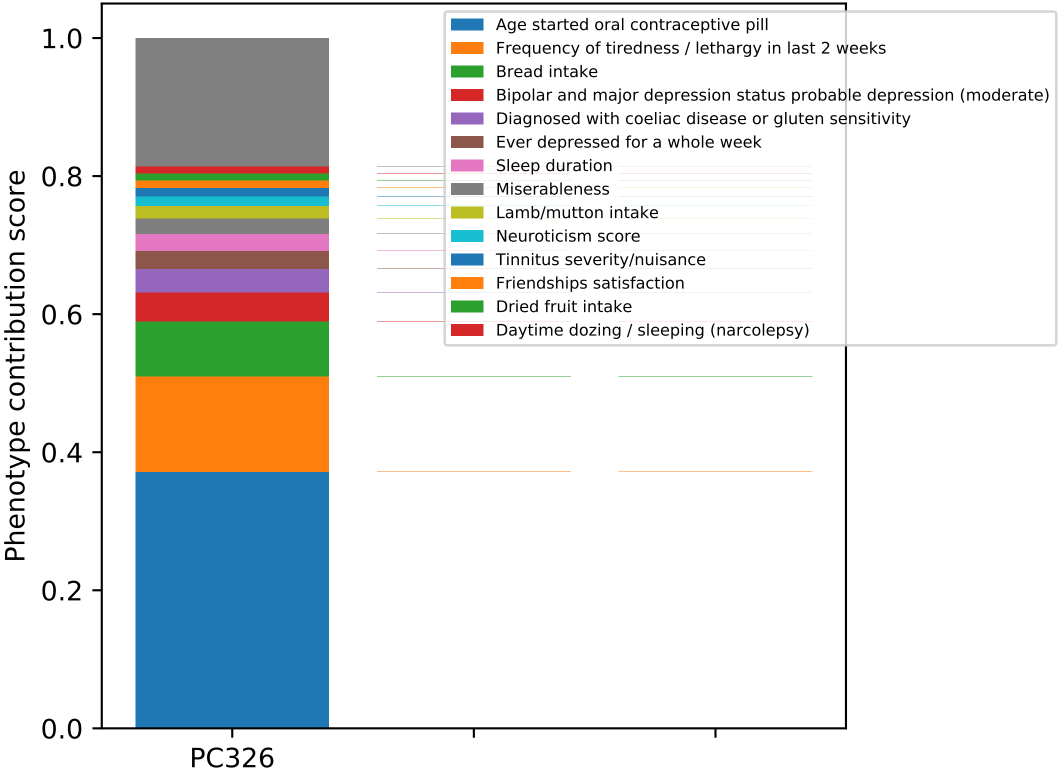

Phenotype contribution score
1.0
0.8
0.6
0.4
0.2
0.0
PC326
Age started oral contraceptive pill
Frequency of tiredness / lethargy in last 2 weeks
Bread intake
Bipolar and major depression status probable depression (moderate)
Diagnosed with coeliac disease or gluten sensitivity
Ever depressed for a whole week
Sleep duration
Miserableness
Lamb/mutton intake
Neuroticism score
Tinnitus severity/nuisance
Friendships satisfaction
Dried fruit intake
Daytime dozing / sleeping (narcolepsy)

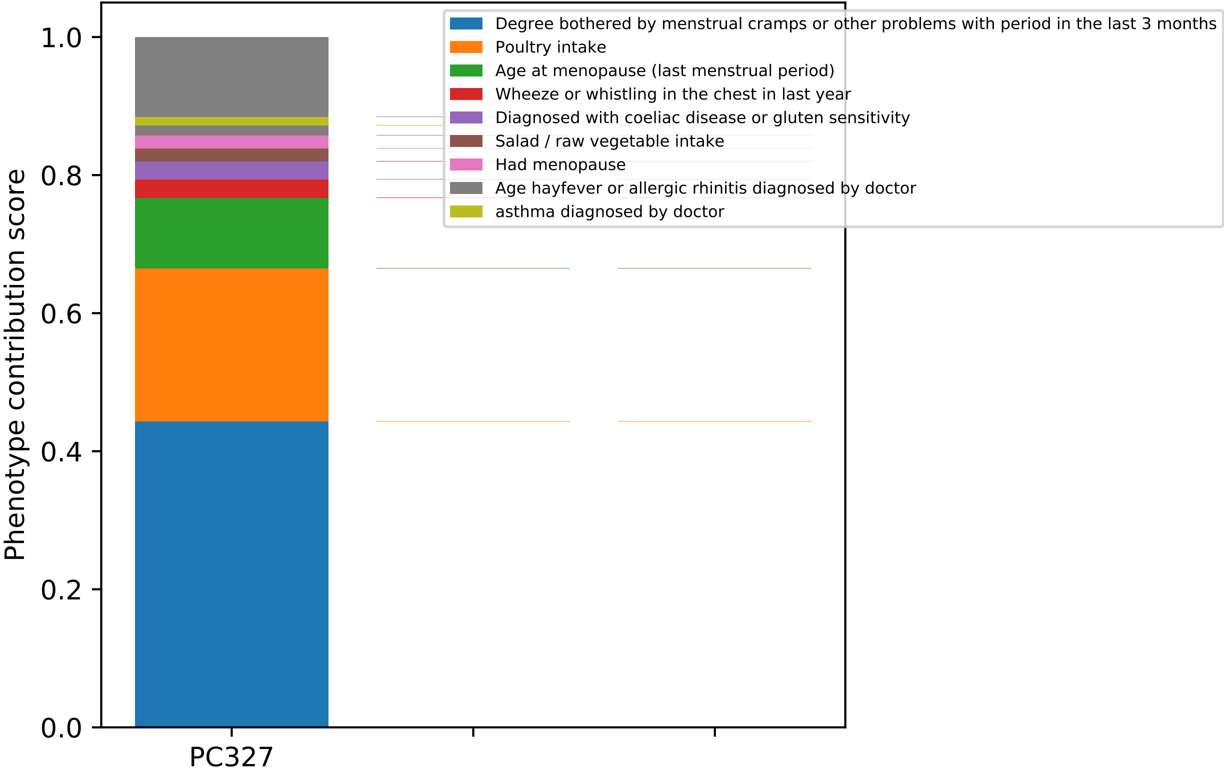
Degree bothered by menstrual cramps or other problems with period in the last 3 months
Poultry intake
Age at menopause (last menstrual period)
Wheeze or whistling in the chest in last year
Diagnosed with coeliac disease or gluten sensitivity
Salad / raw vegetable intake
Had menopause
Age hayfever or allergic rhinitis diagnosed by doctor
asthma diagnosed by doctor
Phenotype contribution score
1.0
0.8
0.6
0.4
0.2
0.0
PC327

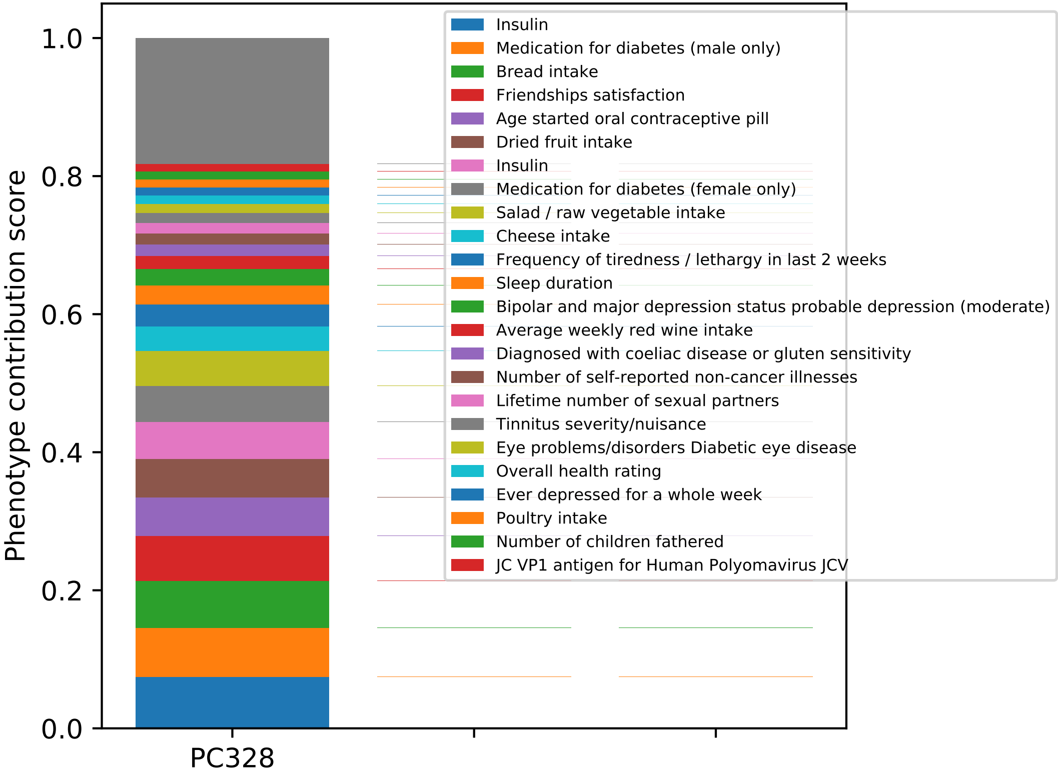
Phenotype contribution score
1.0
0.8
0.6
0.4
0.2
0.0
PC328
Insulin
Medication for diabetes (male only)
Bread intake
Friendships satisfaction
Age started oral contraceptive pill
Dried fruit intake
Insulin
Medication for diabetes (female only)
Salad / raw vegetable intake
Cheese intake
Frequency of tiredness / lethargy in last 2 weeks
Sleep duration
Bipolar and major depression status probable depression (moderate)
Average weekly red wine intake
Diagnosed with coeliac disease or gluten sensitivity
Number of self-reported non-cancer illnesses
Lifetime number of sexual partners
Tinnitus severity/nuisance
Eye problems/disorders Diabetic eye disease
Overall health rating
Ever depressed for a whole week
Poultry intake
Number of children fathered
JC VP1 antigen for Human Polyomavirus JCV

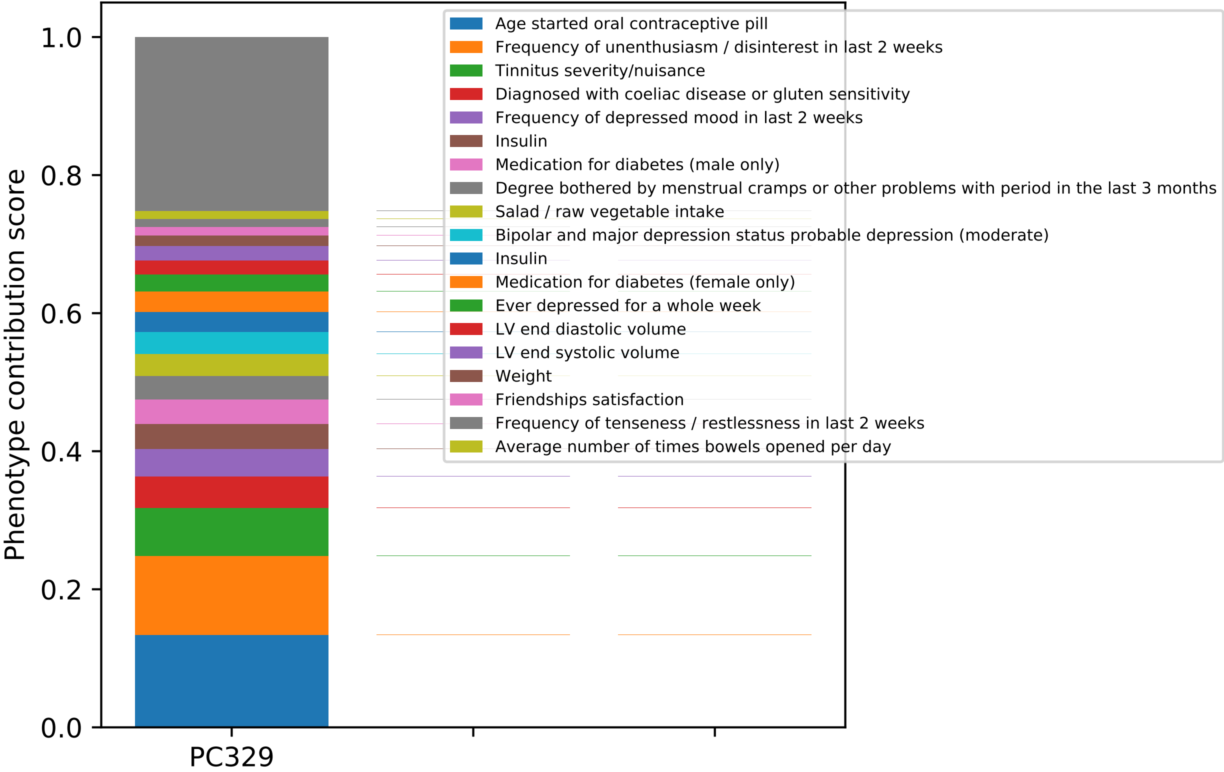

Phenotype contribution score
1.0
0.8
0.6
0.4
0.2
0.0
PC329
Age started oral contraceptive pill
Frequency of unenthusiasm / disinterest in last 2 weeks
Tinnitus severity/nuisance
Diagnosed with coeliac disease or gluten sensitivity
Frequency of depressed mood in last 2 weeks
Insulin
Medication for diabetes (male only)
Degree bothered by menstrual cramps or other problems with period in the last 3 months
Salad / raw vegetable intake
Bipolar and major depression status probable depression (moderate)
Insulin
Medication for diabetes (female only)
Ever depressed for a whole week
LV end diastolic volume
LV end systolic volume
Weight
Friendships satisfaction
Frequency of tenseness / restlessness in last 2 weeks
Average number of times bowels opened per day

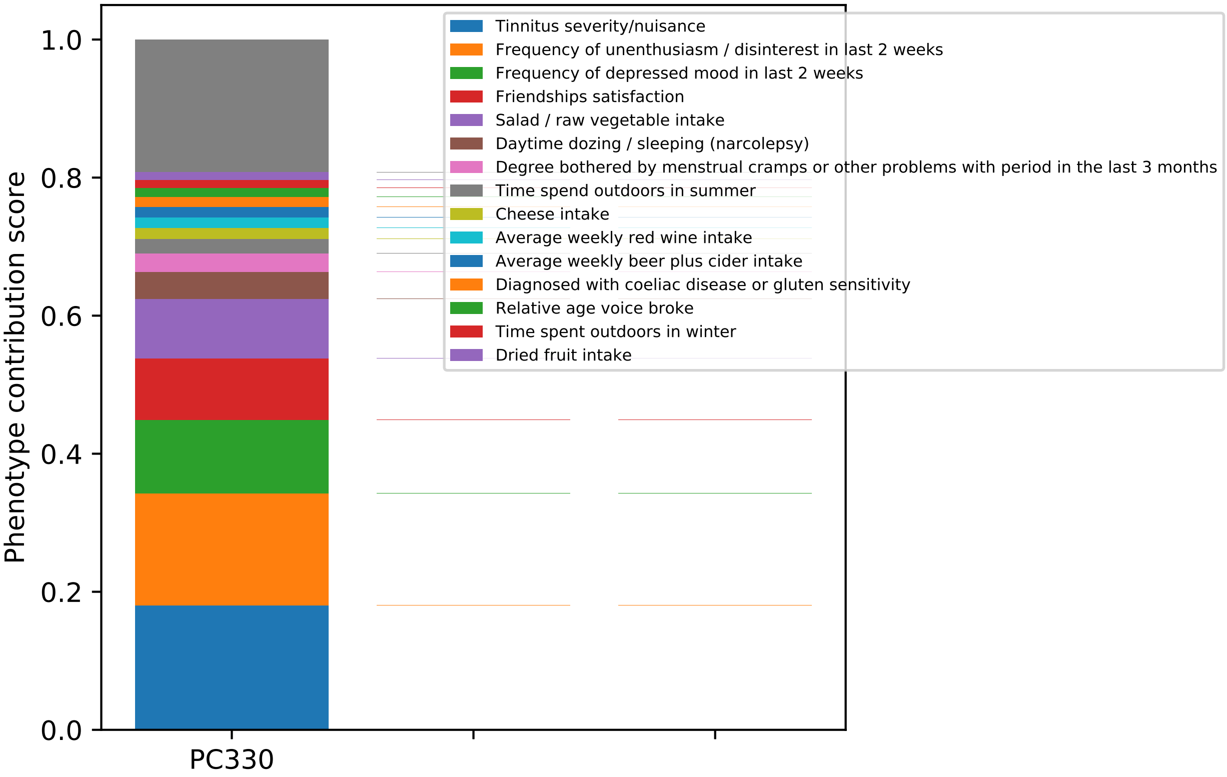

Phenotype contribution score
0.0
0.2
0.4
0.6
0.8
1.0
PC330
Tinnitus severity/nuisance
Frequency of unenthusiasm / disinterest in last 2 weeks
Frequency of depressed mood in last 2 weeks
Friendships satisfaction
Salad / raw vegetable intake
Daytime dozing / sleeping (narcolepsy)
Degree bothered by menstrual cramps or other problems with period in the last 3 months
Time spend outdoors in summer
Cheese intake
Average weekly red wine intake
Average weekly beer plus cider intake
Diagnosed with coeliac disease or gluten sensitivity
Relative age voice broke
Time spent outdoors in winter
Dried fruit intake

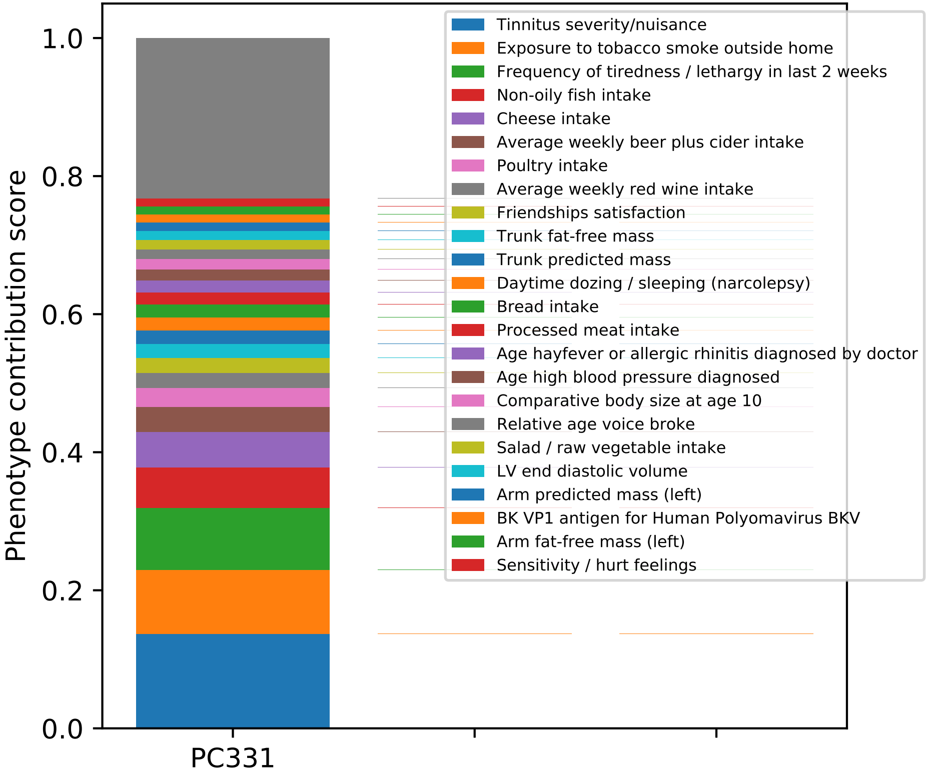

Phenotype contribution score
0.0
0.2
0.4
0.6
0.8
1.0
PC331
Tinnitus severity/nuisance
Exposure to tobacco smoke outside home
Frequency of tiredness / lethargy in last 2 weeks
Non-oily fish intake
Cheese intake
Average weekly beer plus cider intake
Poultry intake
Average weekly red wine intake
Friendships satisfaction
Trunk fat-free mass
Trunk predicted mass
Daytime dozing / sleeping (narcolepsy)
Bread intake
Processed meat intake
Age hayfever or allergic rhinitis diagnosed by doctor
Age high blood pressure diagnosed
Comparative body size at age 10
Relative age voice broke
Salad / raw vegetable intake
LV end diastolic volume
Arm predicted mass (left)
BK VP1 antigen for Human Polyomavirus BKV
Arm fat-free mass (left)
Sensitivity / hurt feelings

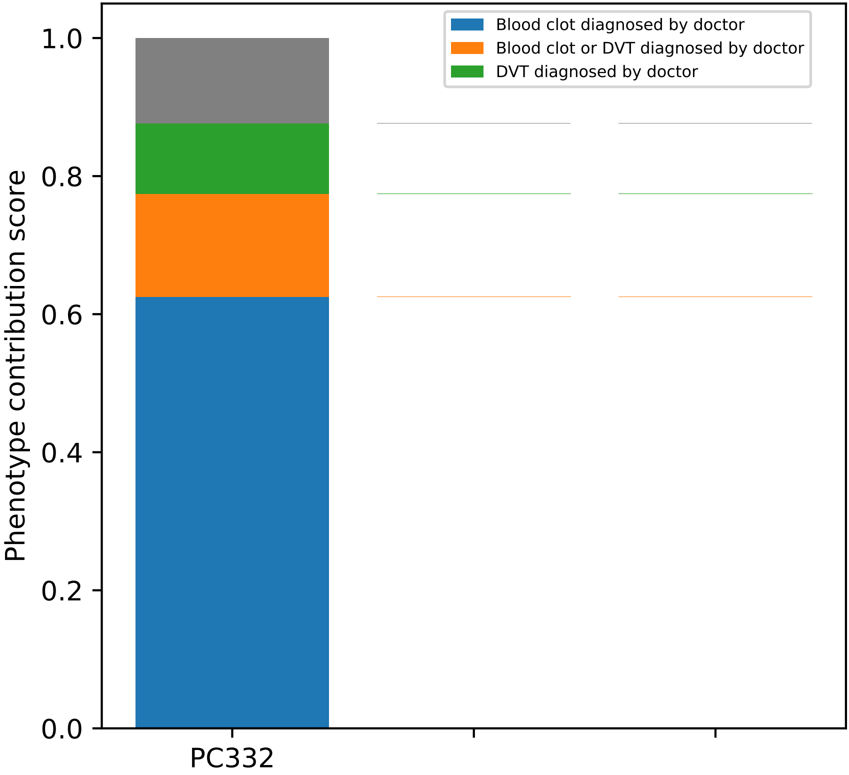

Blood clot diagnosed by doctor
Blood clot or DVT diagnosed by doctor
DVT diagnosed by doctor
Phenotype contribution score
1.0
0.8
0.6
0.4
0.2
0.0
PC332

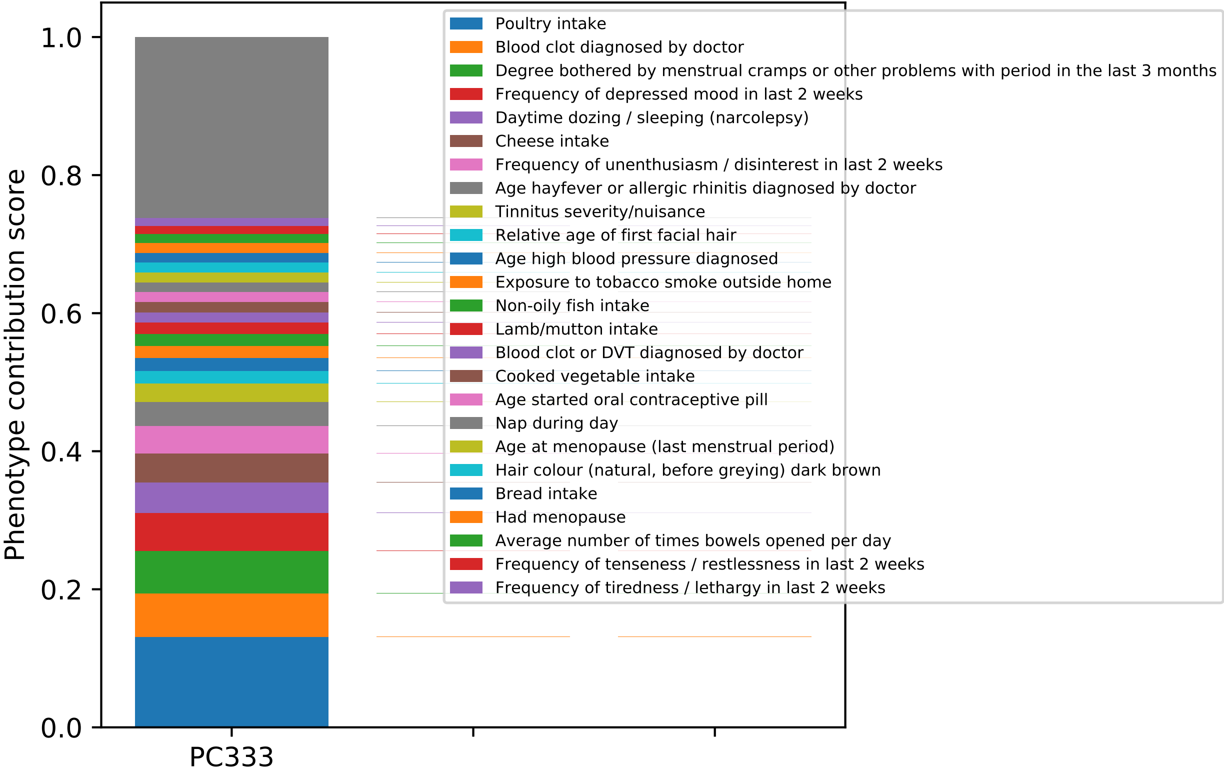

Phenotype contribution score
0.0
0.2
0.4
0.6
0.8
1.0
PC333
Poultry intake
Blood clot diagnosed by doctor
Degree bothered by menstrual cramps or other problems with period in the last 3 months
Frequency of depressed mood in last 2 weeks
Daytime dozing / sleeping (narcolepsy)
Cheese intake
Frequency of unenthusiasm / disinterest in last 2 weeks
Age hayfever or allergic rhinitis diagnosed by doctor
Tinnitus severity/nuisance
Relative age of first facial hair
Age high blood pressure diagnosed
Exposure to tobacco smoke outside home
Non-oily fish intake
Lamb/mutton intake
Blood clot or DVT diagnosed by doctor
Cooked vegetable intake
Age started oral contraceptive pill
Nap during day
Age at menopause (last menstrual period)
Hair colour (natural, before greying) dark brown
Bread intake
Had menopause
Average number of times bowels opened per day
Frequency of tenseness / restlessness in last 2 weeks
Frequency of tiredness / lethargy in last 2 weeks

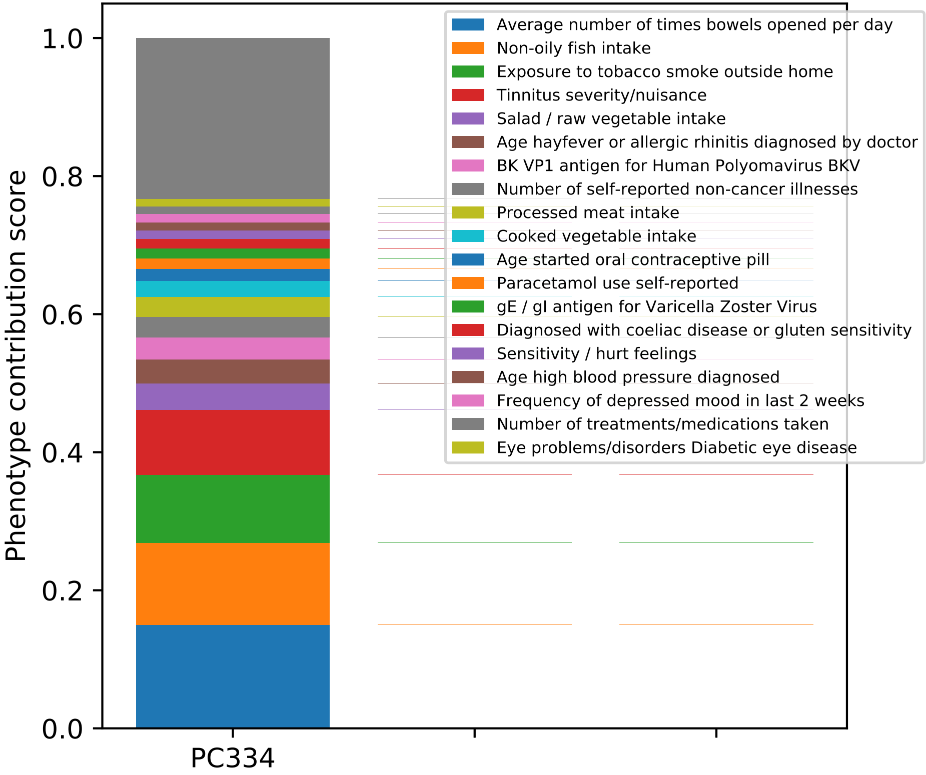

Phenotype contribution score
1.0
0.8
0.6
0.4
0.2
0.0
PC334
Average number of times bowels opened per day
Non-oily fish intake
Exposure to tobacco smoke outside home
Tinnitus severity/nuisance
Salad / raw vegetable intake
Age hayfever or allergic rhinitis diagnosed by doctor
BK VP1 antigen for Human Polyomavirus BKV
Number of self-reported non-cancer illnesses
Processed meat intake
Cooked vegetable intake
Age started oral contraceptive pill
Paracetamol use self-reported
gE / gI antigen for Varicella Zoster Virus
Diagnosed with coeliac disease or gluten sensitivity
Sensitivity / hurt feelings
Age high blood pressure diagnosed
Frequency of depressed mood in last 2 weeks
Number of treatments/medications taken
Eye problems/disorders Diabetic eye disease

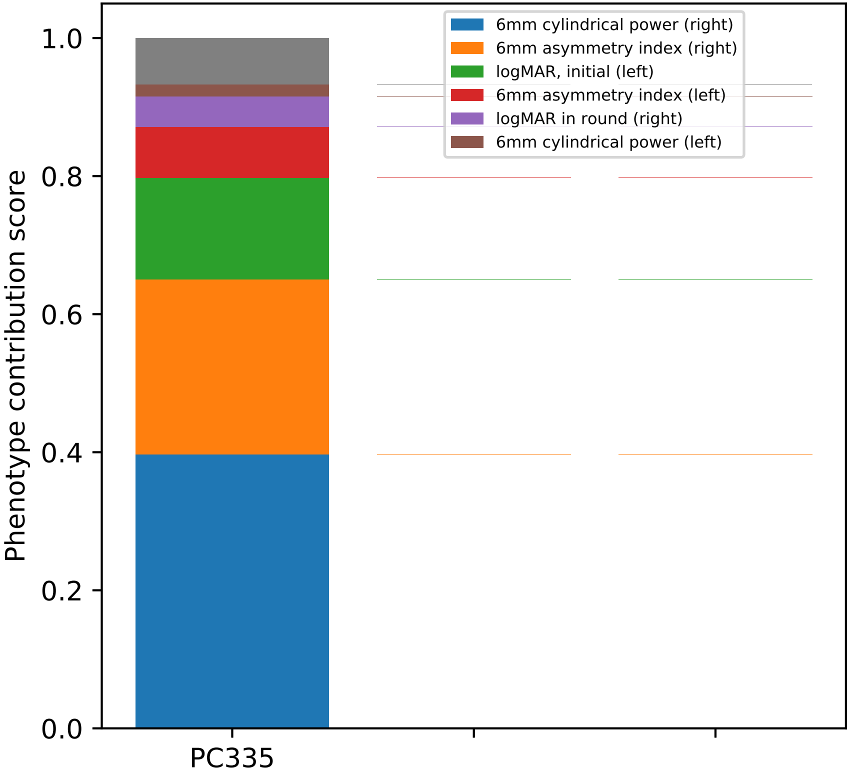
6mm cylindrical power (right)
6mm asymmetry index (right)
logMAR, initial (left)
6mm asymmetry index (left)
logMAR in round (right)
6mm cylindrical power (left)
1.0
0.8
0.6
0.4
0.2
0.0
Phenotype contribution score
PC335

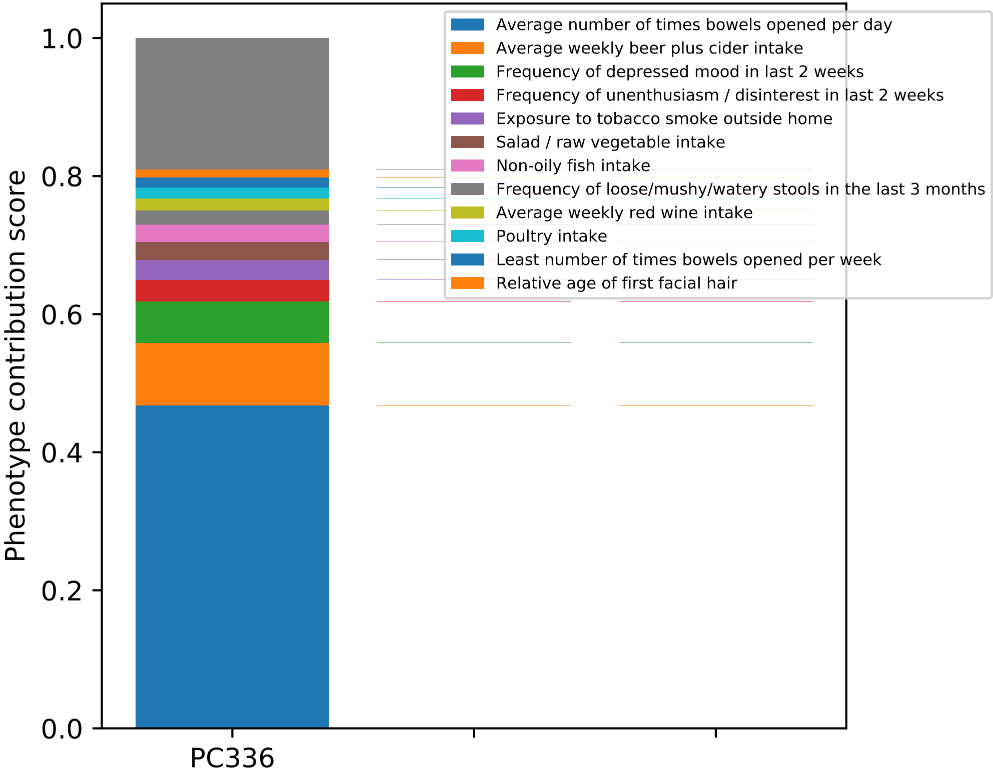

Phenotype contribution score
0.0
0.2
0.4
0.6
0.8
1.0
PC336
Average number of times bowels opened per day
Average weekly beer plus cider intake
Frequency of depressed mood in last 2 weeks
Frequency of unenthusiasm / disinterest in last 2 weeks
Exposure to tobacco smoke outside home
Salad / raw vegetable intake
Non-oily fish intake
Frequency of loose/mushy/watery stools in the last 3 months
Average weekly red wine intake
Poultry intake
Least number of times bowels opened per week
Relative age of first facial hair

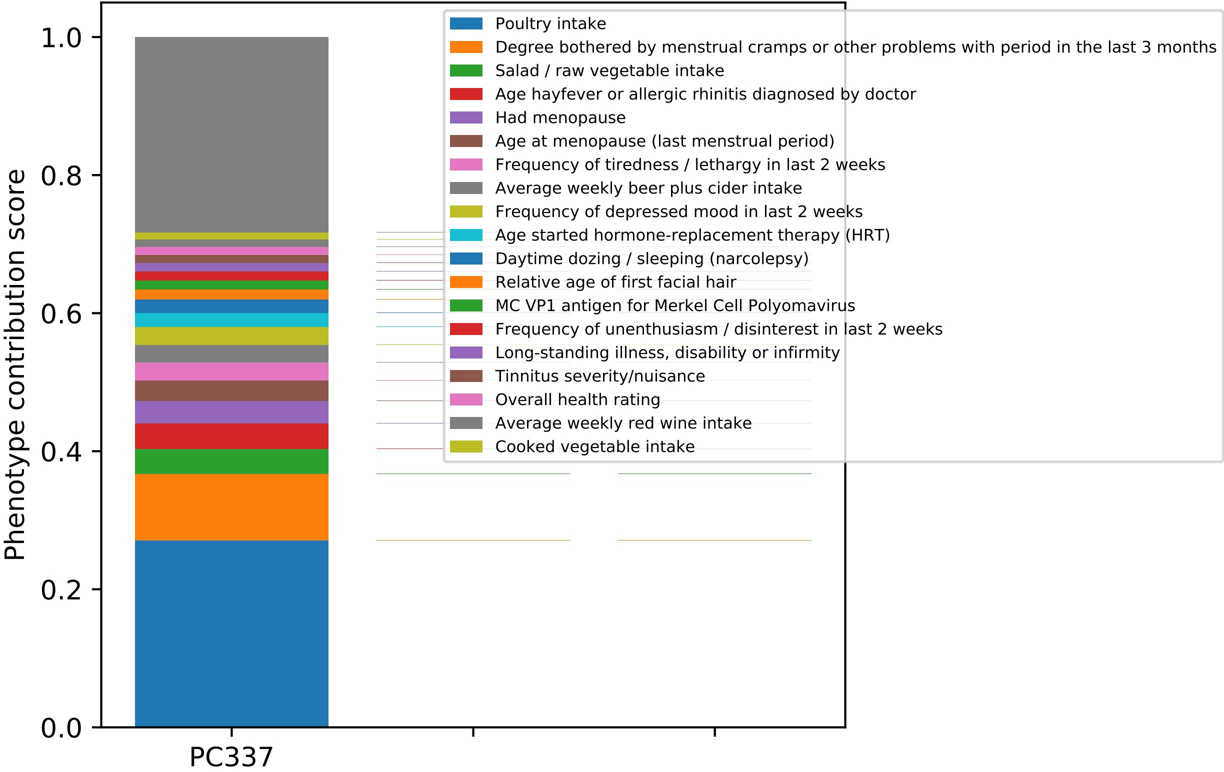

Phenotype contribution score
0.0
0.2
0.4
0.6
0.8
1.0
PC337
Poultry intake
Degree bothered by menstrual cramps or other problems with period in the last 3 months
Salad / raw vegetable intake
Age hayfever or allergic rhinitis diagnosed by doctor
Had menopause
Age at menopause (last menstrual period)
Frequency of tiredness / lethargy in last 2 weeks
Average weekly beer plus cider intake
Frequency of depressed mood in last 2 weeks
Age started hormone-replacement therapy (HRT)
Daytime dozing / sleeping (narcolepsy)
Relative age of first facial hair
MC VP1 antigen for Merkel Cell Polyomavirus
Frequency of unenthusiasm / disinterest in last 2 weeks
Long-standing illness, disability or infirmity
Tinnitus severity/nuisance
Overall health rating
Average weekly red wine intake
Cooked vegetable intake

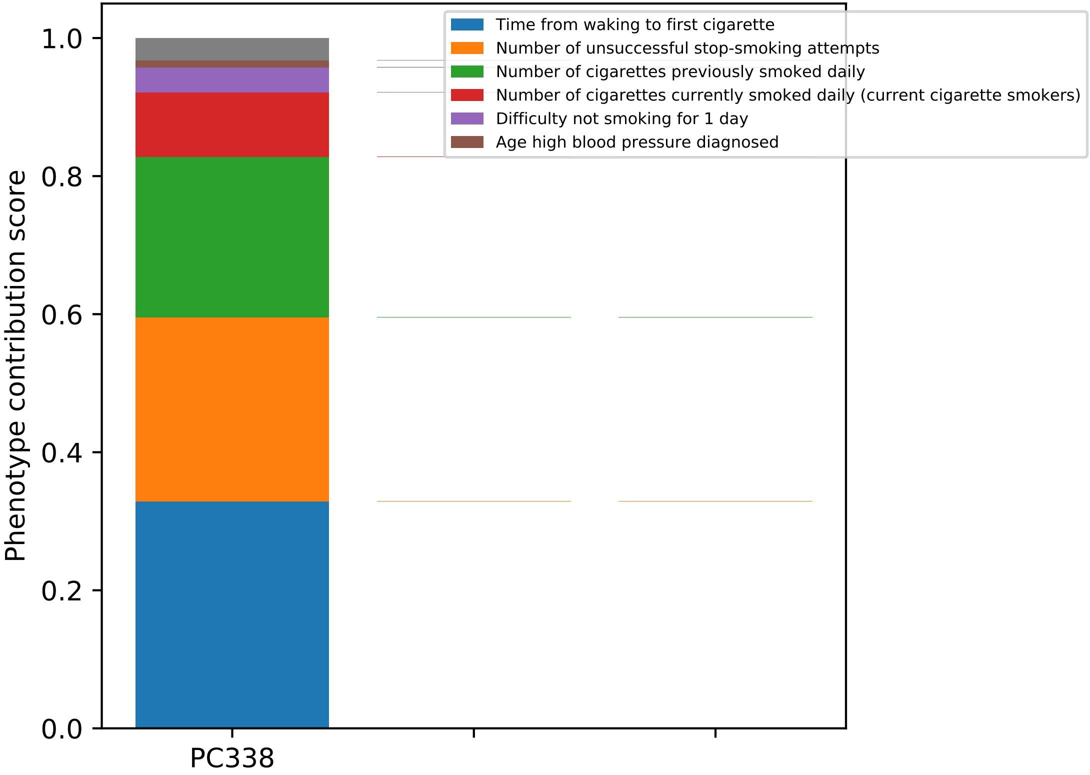
Time from waking to first cigarette
Number of unsuccessful stop-smoking attempts
Number of cigarettes previously smoked daily
Number of cigarettes currently smoked daily (current cigarette smokers)
Difficulty not smoking for 1 day
Age high blood pressure diagnosed
Phenotype contribution score
1.0
0.8
0.6
0.4
0.2
0.0
PC338

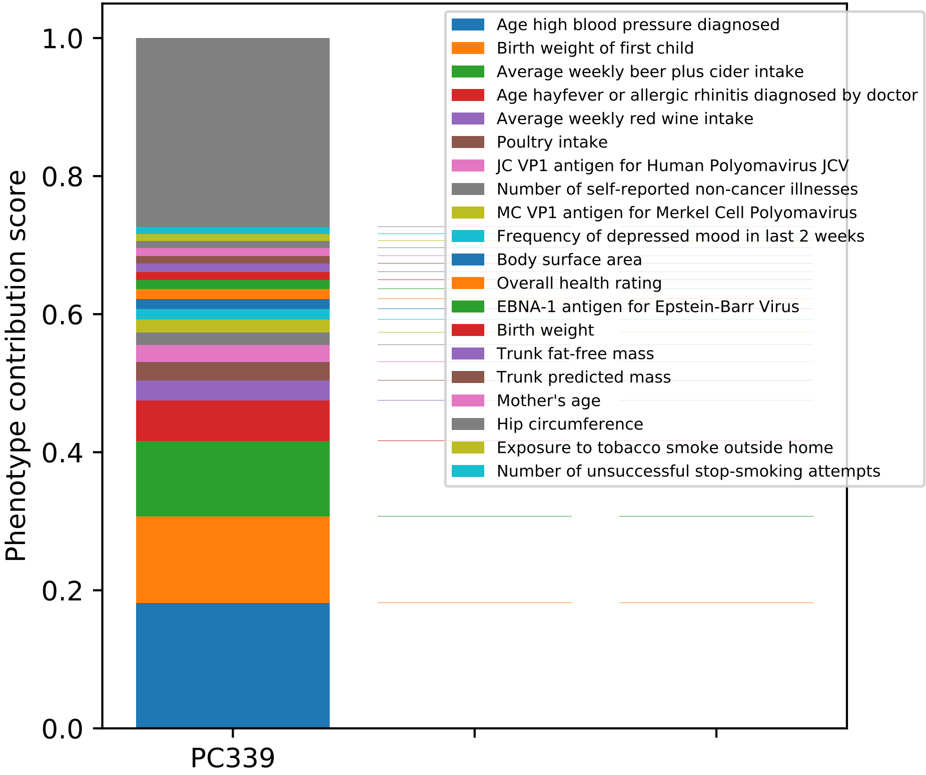
Phenotype contribution score
1.0
0.8
0.6
0.4
0.2
0.0
PC339
Age high blood pressure diagnosed
Birth weight of first child
Average weekly beer plus cider intake
Age hayfever or allergic rhinitis diagnosed by doctor
Average weekly red wine intake
Poultry intake
JC VP1 antigen for Human Polyomavirus JCV
Number of self-reported non-cancer illnesses
MC VP1 antigen for Merkel Cell Polyomavirus
Frequency of depressed mood in last 2 weeks
Body surface area
Overall health rating
EBNA-1 antigen for Epstein-Barr Virus
Birth weight
Trunk fat-free mass
Trunk predicted mass
Mother's age
Hip circumference
Exposure to tobacco smoke outside home
Number of unsuccessful stop-smoking attempts

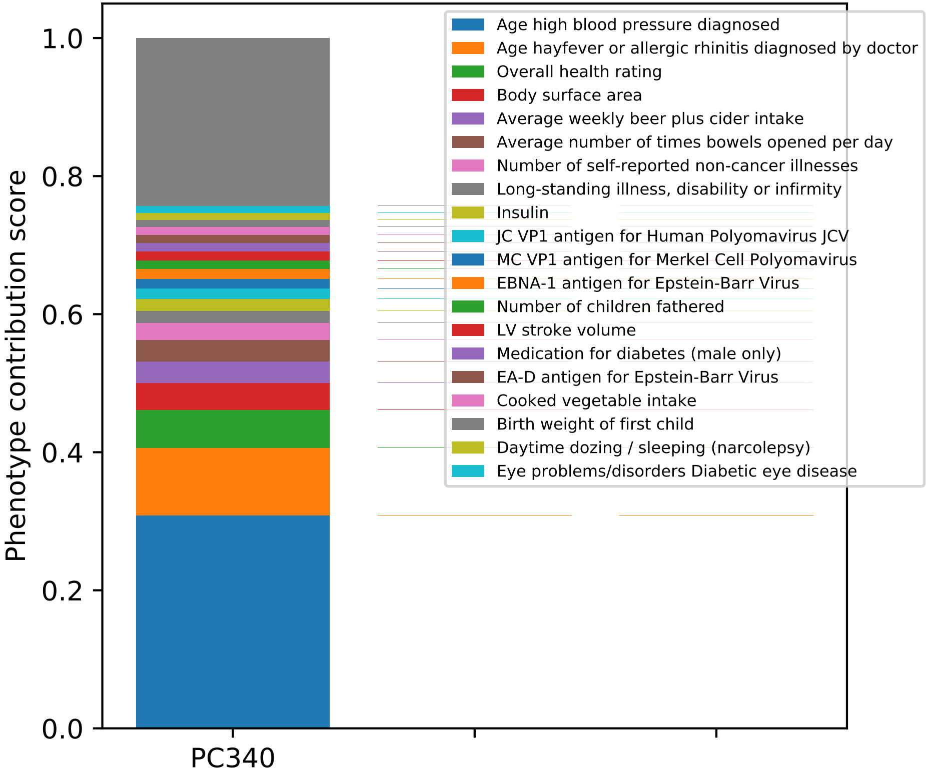
Phenotype contribution score
1.0
0.8
0.6
0.4
0.2
0.0
PC340
Age high blood pressure diagnosed
Age hayfever or allergic rhinitis diagnosed by doctor
Overall health rating
Body surface area
Average weekly beer plus cider intake
Average number of times bowels opened per day
Number of self-reported non-cancer illnesses
Long-standing illness, disability or infirmity
Insulin
JC VP1 antigen for Human Polyomavirus JCV
MC VP1 antigen for Merkel Cell Polyomavirus
EBNA-1 antigen for Epstein-Barr Virus
Number of children fathered
LV stroke volume
Medication for diabetes (male only)
EA-D antigen for Epstein-Barr Virus
Cooked vegetable intake
Birth weight of first child
Daytime dozing / sleeping (narcolepsy)
Eye problems/disorders Diabetic eye disease

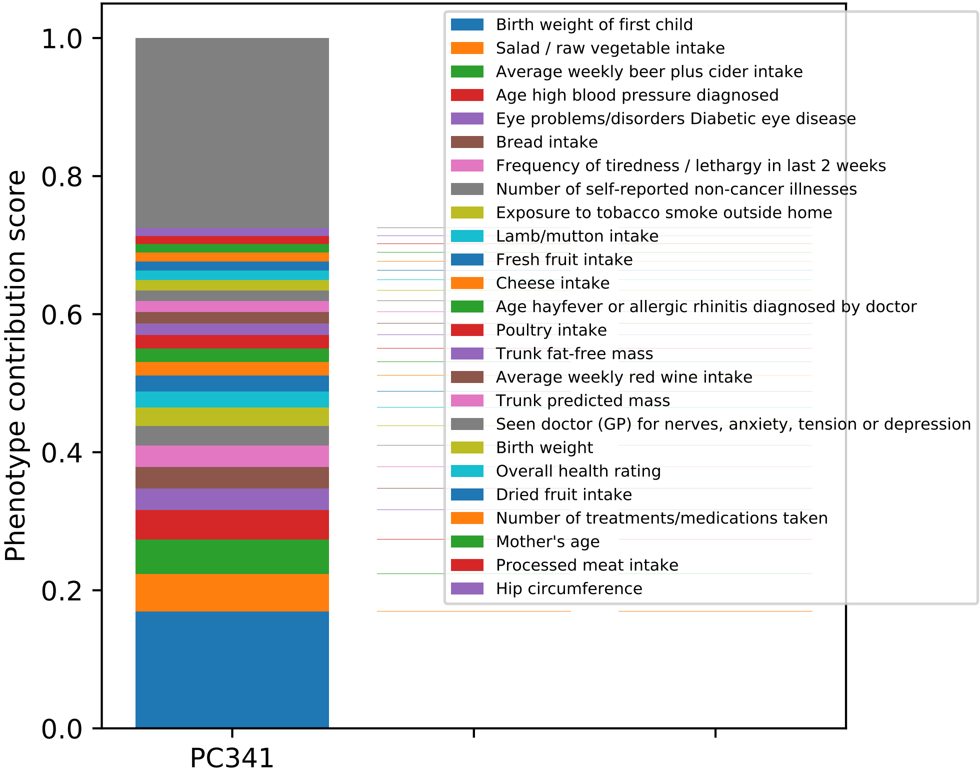

Phenotype contribution score
1.0
0.8
0.6
0.4
0.2
0.0
PC341
Birth weight of first child
Salad / raw vegetable intake
Average weekly beer plus cider intake
Age high blood pressure diagnosed
Eye problems/disorders Diabetic eye disease
Bread intake
Frequency of tiredness / lethargy in last 2 weeks
Number of self-reported non-cancer illnesses
Exposure to tobacco smoke outside home
Lamb/mutton intake
Fresh fruit intake
Cheese intake
Age hayfever or allergic rhinitis diagnosed by doctor
Poultry intake
Trunk fat-free mass
Average weekly red wine intake
Trunk predicted mass
Seen doctor (GP) for nerves, anxiety, tension or depression
Birth weight
Overall health rating
Dried fruit intake
Number of treatments/medications taken
Mother's age
Processed meat intake
Hip circumference

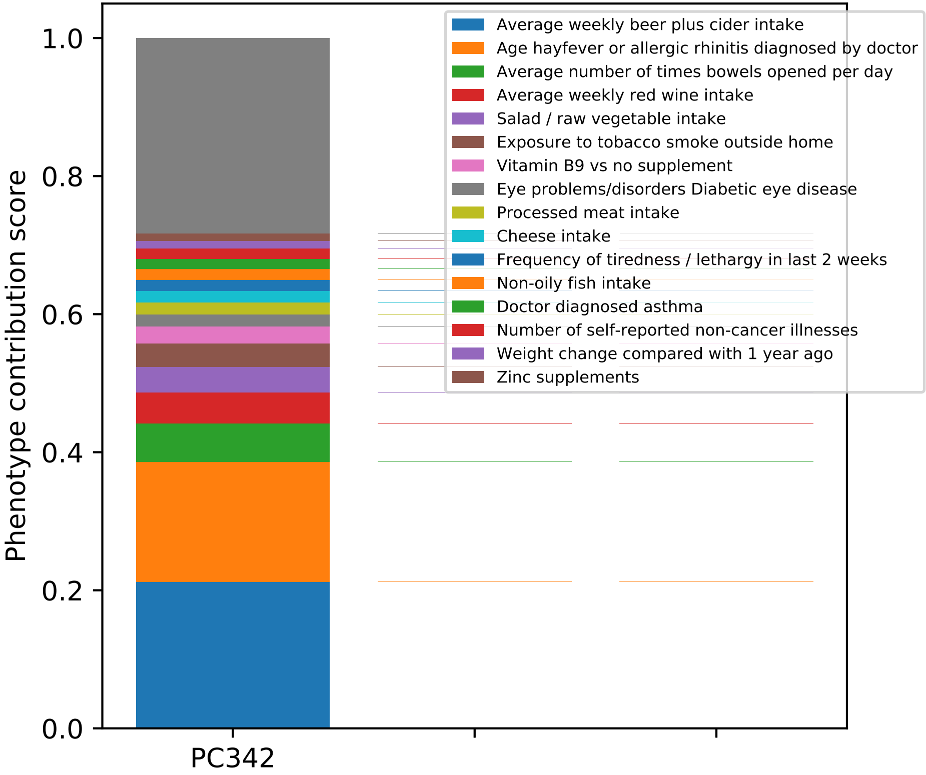
Average weekly beer plus cider intake
Age hayfever or allergic rhinitis diagnosed by doctor
Average number of times bowels opened per day
Average weekly red wine intake
Salad / raw vegetable intake
Exposure to tobacco smoke outside home
Vitamin B9 vs no supplement
Eye problems/disorders Diabetic eye disease
Processed meat intake
Cheese intake
Frequency of tiredness / lethargy in last 2 weeks
Non-oily fish intake
Doctor diagnosed asthma
Number of self-reported non-cancer illnesses
Weight change compared with 1 year ago
Zinc supplements
Phenotype contribution score
1.0
0.8
0.6
0.4
0.2
0.0
PC342

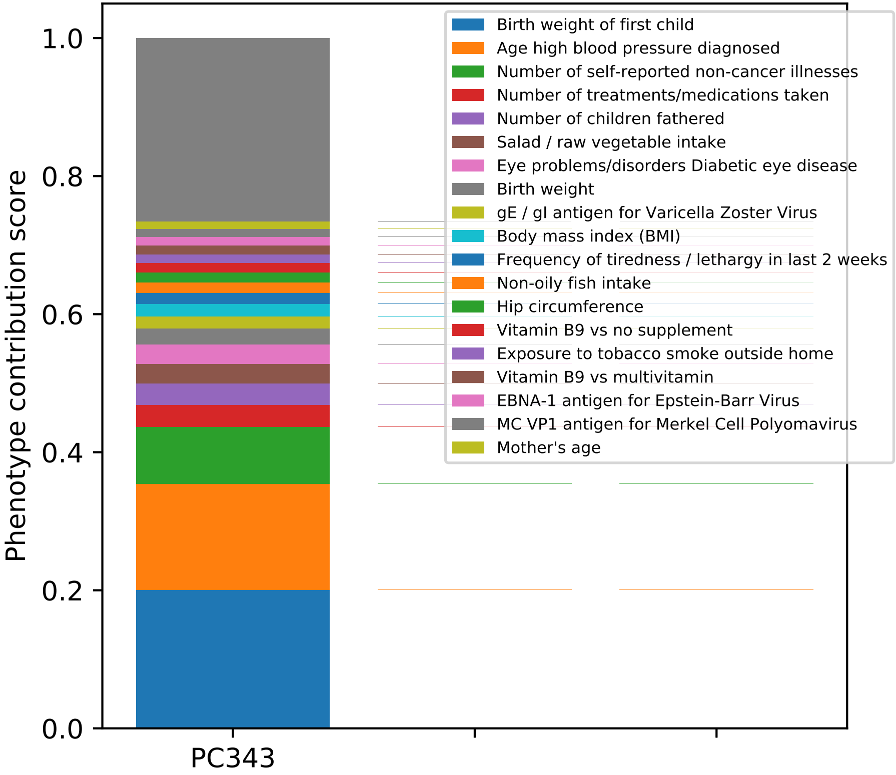

Phenotype contribution score
0.0
0.2
0.4
0.6
0.8
1.0
PC343
Birth weight of first child
Age high blood pressure diagnosed
Number of self-reported non-cancer illnesses
Number of treatments/medications taken
Number of children fathered
Salad / raw vegetable intake
Eye problems/disorders Diabetic eye disease
Birth weight
gE / gI antigen for Varicella Zoster Virus
Body mass index (BMI)
Frequency of tiredness / lethargy in last 2 weeks
Non-oily fish intake
Hip circumference
Vitamin B9 vs no supplement
Exposure to tobacco smoke outside home
Vitamin B9 vs multivitamin
EBNA-1 antigen for Epstein-Barr Virus
MC VP1 antigen for Merkel Cell Polyomavirus
Mother's age

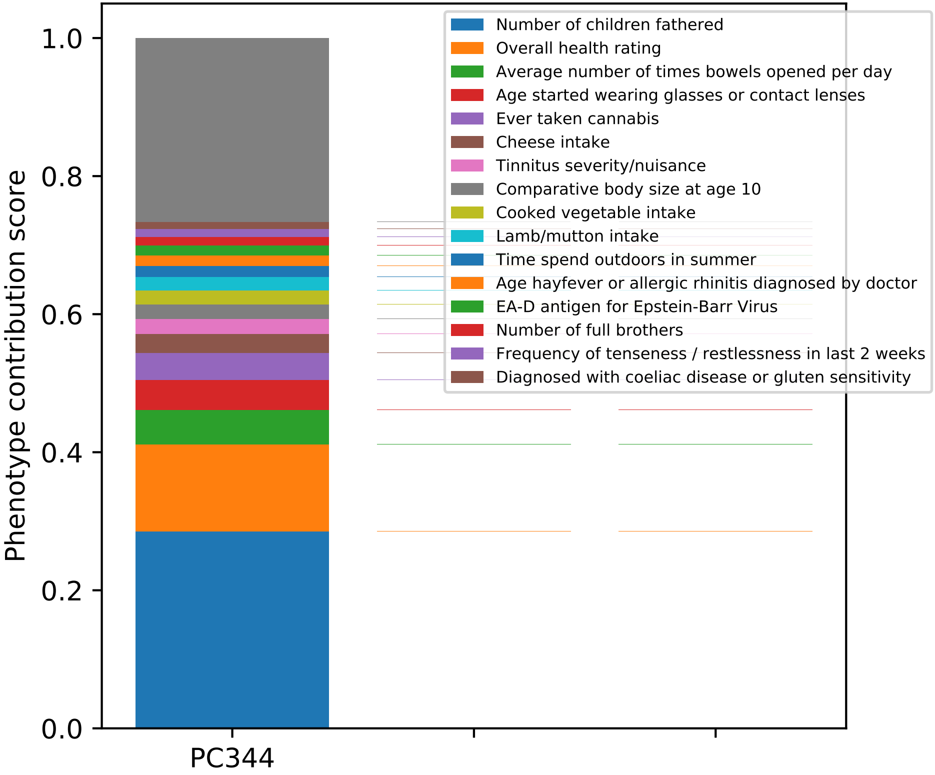

Number of children fathered
Overall health rating
Average number of times bowels opened per day
Age started wearing glasses or contact lenses
Ever taken cannabis
Cheese intake
Tinnitus severity/nuisance
Comparative body size at age 10
Cooked vegetable intake
Lamb/mutton intake
Time spend outdoors in summer
Age hayfever or allergic rhinitis diagnosed by doctor
EA-D antigen for Epstein-Barr Virus
Number of full brothers
Frequency of tenseness / restlessness in last 2 weeks
Diagnosed with coeliac disease or gluten sensitivity
Phenotype contribution score
0.0
0.2
0.4
0.6
0.8
1.0
PC344

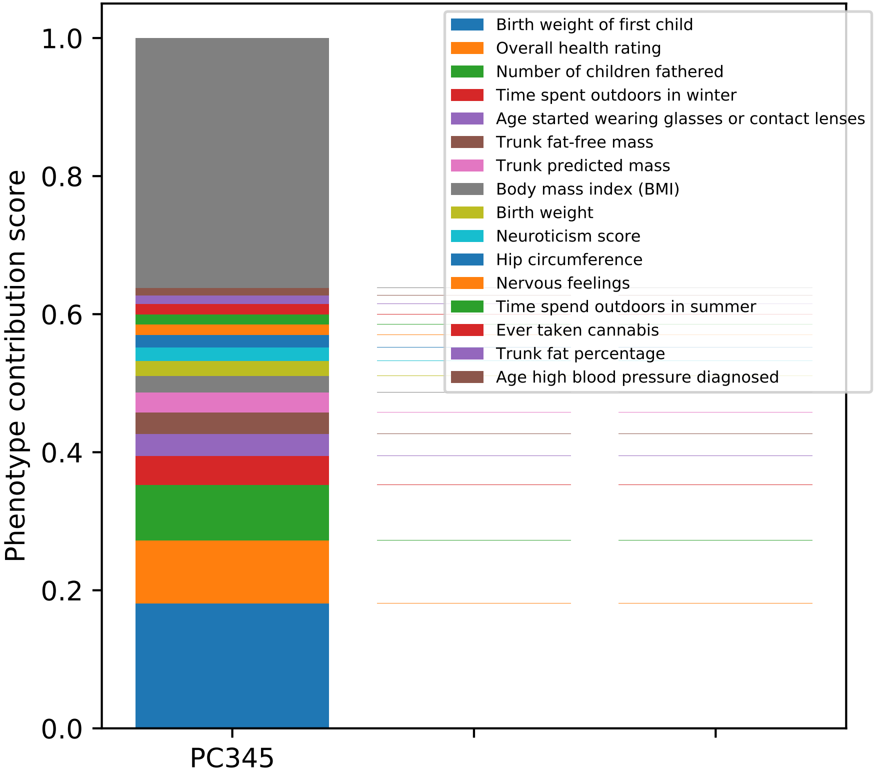

Phenotype contribution score
1.0
0.8
0.6
0.4
0.2
0.0
PC345
Birth weight of first child
Overall health rating
Number of children fathered
Time spent outdoors in winter
Age started wearing glasses or contact lenses
Trunk fat-free mass
Trunk predicted mass
Body mass index (BMI)
Birth weight
Neuroticism score
Hip circumference
Nervous feelings
Time spend outdoors in summer
Ever taken cannabis
Trunk fat percentage
Age high blood pressure diagnosed

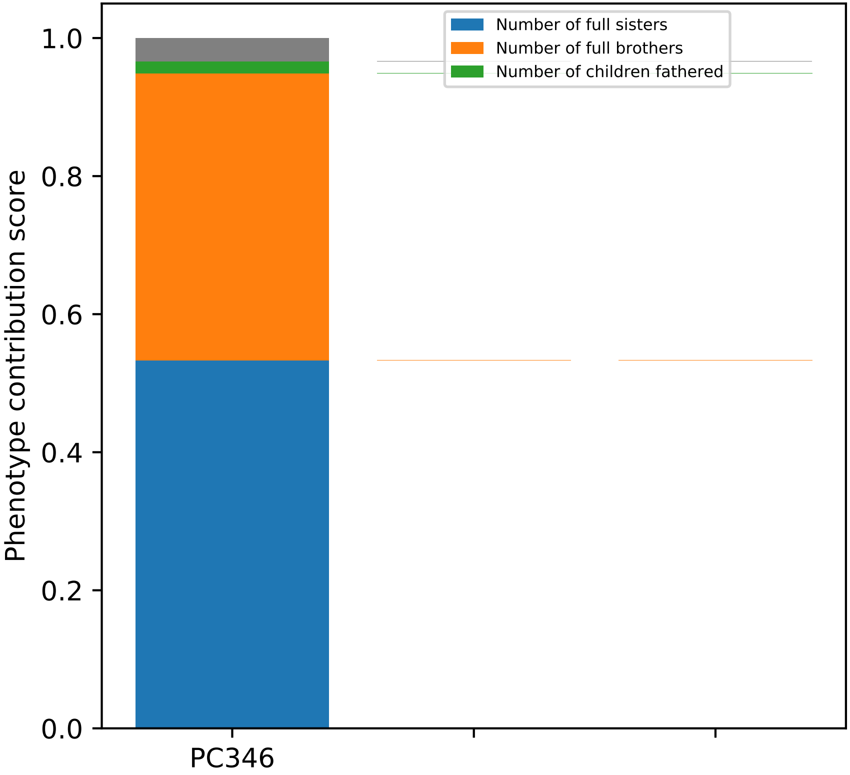

Number of full sisters
Number of full brothers
Number of children fathered
Phenotype contribution score
1.0
0.8
0.6
0.4
0.2
0.0
PC346

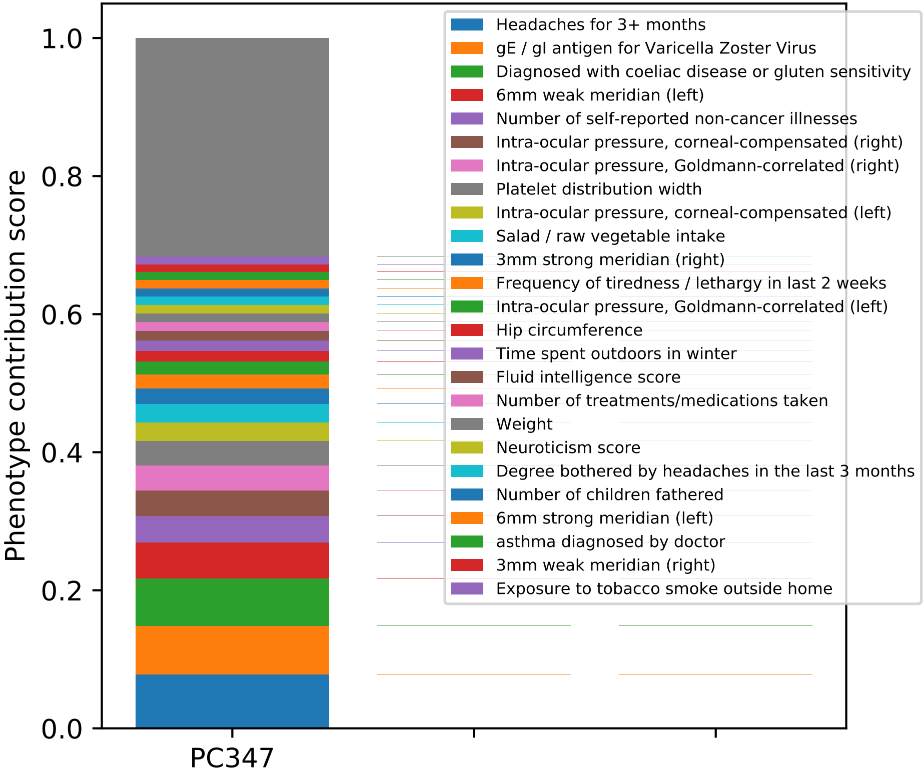

Phenotype contribution score
0.0
0.2
0.4
0.6
0.8
1.0
PC347
Headaches for 3+ months
gE / gI antigen for Varicella Zoster Virus
Diagnosed with coeliac disease or gluten sensitivity
6mm weak meridian (left)
Number of self-reported non-cancer illnesses
Intra-ocular pressure, corneal-compensated (right)
Intra-ocular pressure, Goldmann-correlated (right)
Platelet distribution width
Intra-ocular pressure, corneal-compensated (left)
Salad / raw vegetable intake
3mm strong meridian (right)
Frequency of tiredness / lethargy in last 2 weeks
Intra-ocular pressure, Goldmann-correlated (left)
Hip circumference
Time spent outdoors in winter
Fluid intelligence score
Number of treatments/medications taken
Weight
Neuroticism score
Degree bothered by headaches in the last 3 months
Number of children fathered
6mm strong meridian (left)
asthma diagnosed by doctor
3mm weak meridian (right)
Exposure to tobacco smoke outside home

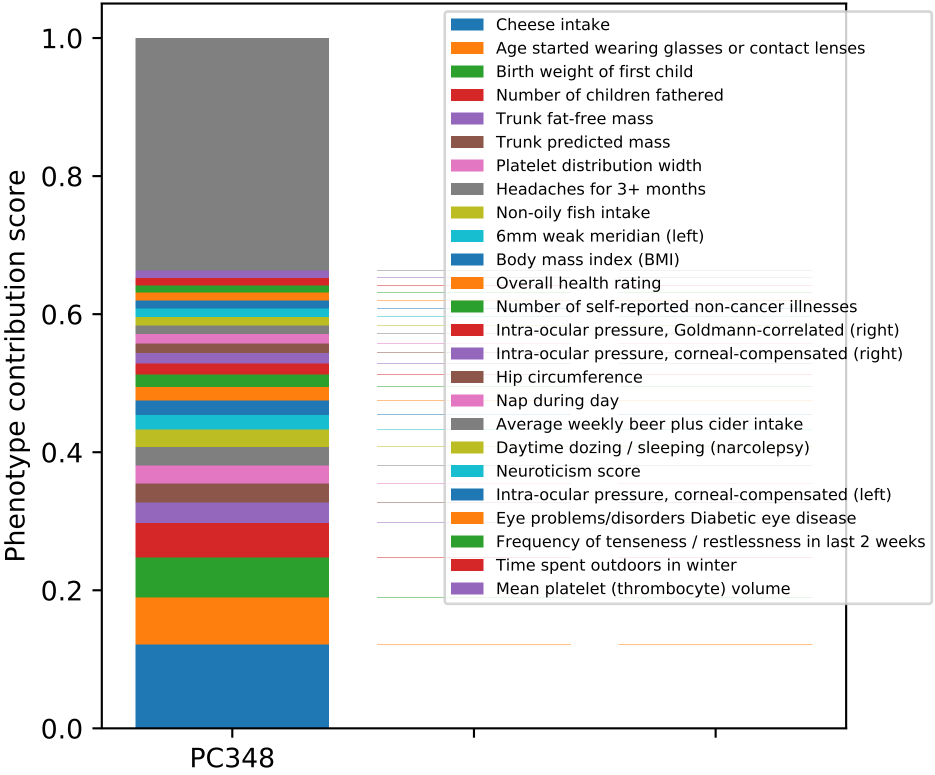

Phenotype contribution score
1.0
0.8
0.6
0.4
0.2
0.0
PC348
Cheese intake
Age started wearing glasses or contact lenses
Birth weight of first child
Number of children fathered
Trunk fat-free mass
Trunk predicted mass
Platelet distribution width
Headaches for 3+ months
Non-oily fish intake
6mm weak meridian (left)
Body mass index (BMI)
Overall health rating
Number of self-reported non-cancer illnesses
Intra-ocular pressure, Goldmann-correlated (right)
Intra-ocular pressure, corneal-compensated (right)
Hip circumference
Nap during day
Average weekly beer plus cider intake
Daytime dozing / sleeping (narcolepsy)
Neuroticism score
Intra-ocular pressure, corneal-compensated (left)
Eye problems/disorders Diabetic eye disease
Frequency of tenseness / restlessness in last 2 weeks
Time spent outdoors in winter
Mean platelet (thrombocyte) volume

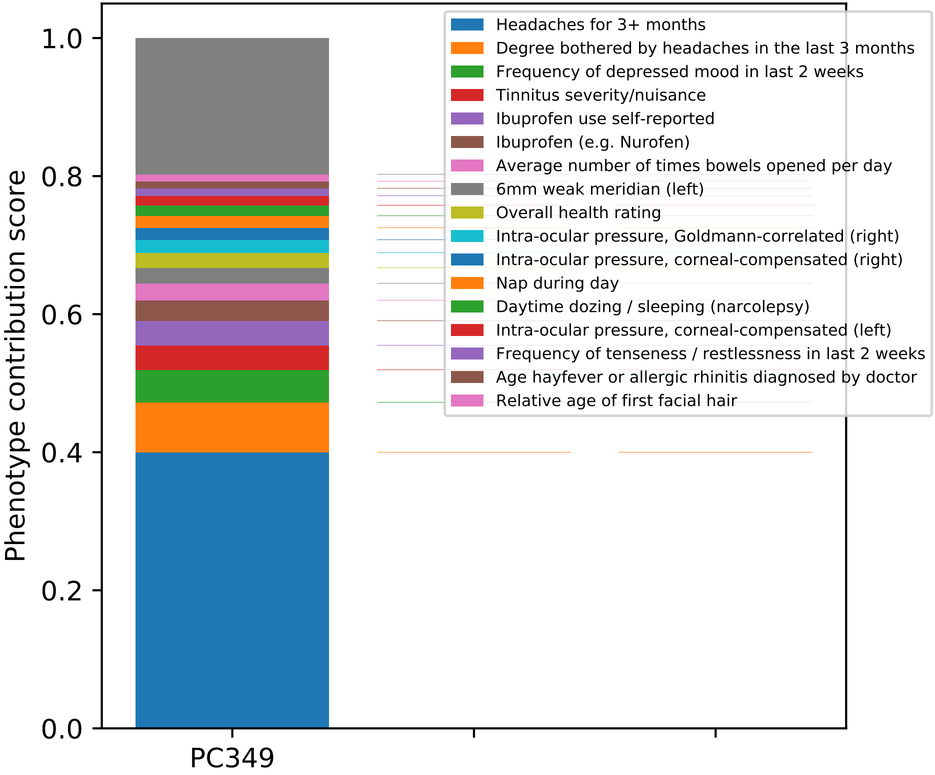

Headaches for 3+ months
Degree bothered by headaches in the last 3 months
Frequency of depressed mood in last 2 weeks
Tinnitus severity/nuisance
Ibuprofen use self-reported
Ibuprofen (e.g. Nurofen)
Average number of times bowels opened per day
6mm weak meridian (left)
Overall health rating
Intra-ocular pressure, Goldmann-correlated (right)
Intra-ocular pressure, corneal-compensated (right)
Nap during day
Daytime dozing / sleeping (narcolepsy)
Intra-ocular pressure, corneal-compensated (left)
Frequency of tenseness / restlessness in last 2 weeks
Age hayfever or allergic rhinitis diagnosed by doctor
Relative age of first facial hair
Phenotype contribution score
0.0
0.2
0.4
0.6
0.8
1.0
PC349

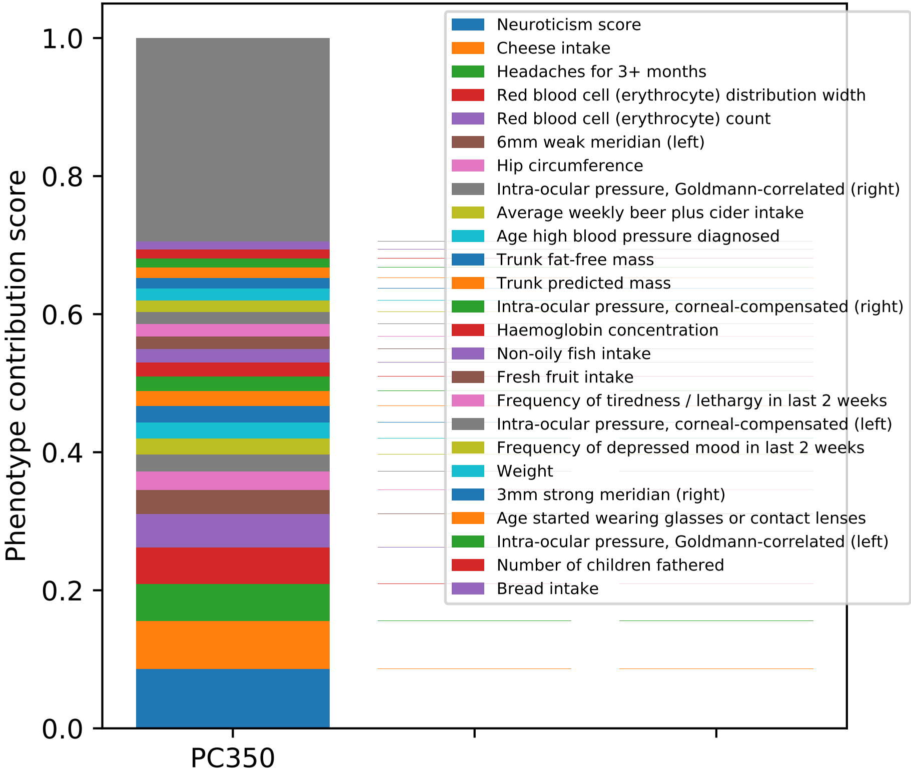

Phenotype contribution score
0.0
0.2
0.4
0.6
0.8
1.0
PC350
Neuroticism score
Cheese intake
Headaches for 3+ months
Red blood cell (erythrocyte) distribution width
Red blood cell (erythrocyte) count
6mm weak meridian (left)
Hip circumference
Intra-ocular pressure, Goldmann-correlated (right)
Average weekly beer plus cider intake
Age high blood pressure diagnosed
Trunk fat-free mass
Trunk predicted mass
Intra-ocular pressure, corneal-compensated (right)
Haemoglobin concentration
Non-oily fish intake
Fresh fruit intake
Frequency of tiredness / lethargy in last 2 weeks
Intra-ocular pressure, corneal-compensated (left)
Frequency of depressed mood in last 2 weeks
Weight
3mm strong meridian (right)
Age started wearing glasses or contact lenses
Intra-ocular pressure, Goldmann-correlated (left)
Number of children fathered
Bread intake

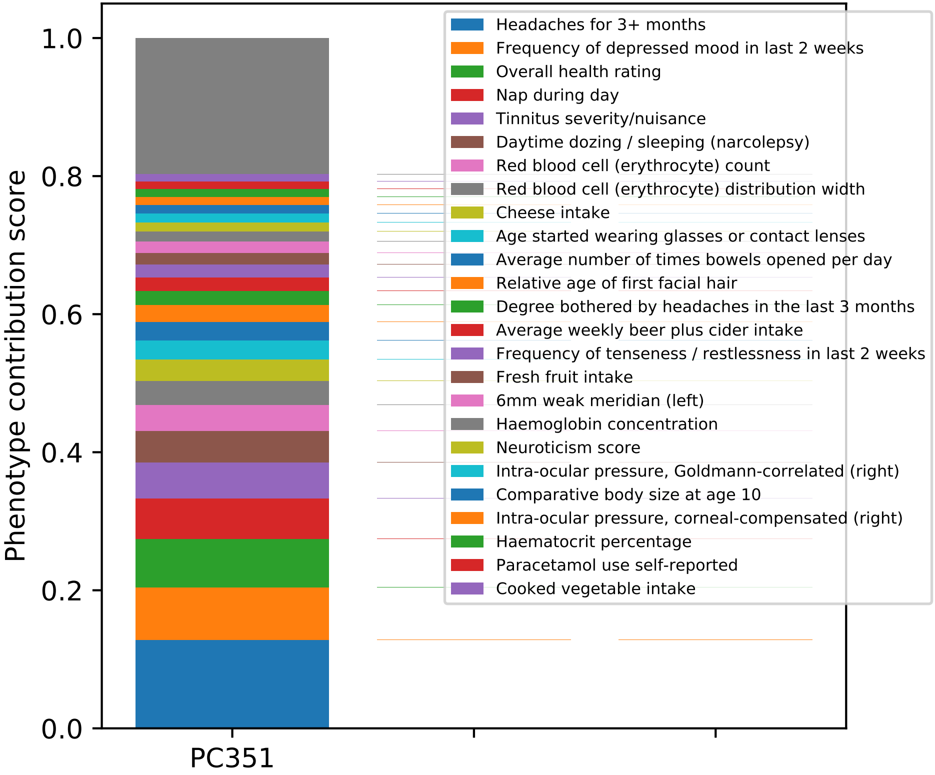

Phenotype contribution score
1.0
0.8
0.6
0.4
0.2
0.0
PC351
Headaches for 3+ months
Frequency of depressed mood in last 2 weeks
Overall health rating
Nap during day
Tinnitus severity/nuisance
Daytime dozing / sleeping (narcolepsy)
Red blood cell (erythrocyte) count
Red blood cell (erythrocyte) distribution width
Cheese intake
Age started wearing glasses or contact lenses
Average number of times bowels opened per day
Relative age of first facial hair
Degree bothered by headaches in the last 3 months
Average weekly beer plus cider intake
Frequency of tenseness / restlessness in last 2 weeks
Fresh fruit intake
6mm weak meridian (left)
Haemoglobin concentration
Neuroticism score
Intra-ocular pressure, Goldmann-correlated (right)
Comparative body size at age 10
Intra-ocular pressure, corneal-compensated (right)
Haematocrit percentage
Paracetamol use self-reported
Cooked vegetable intake

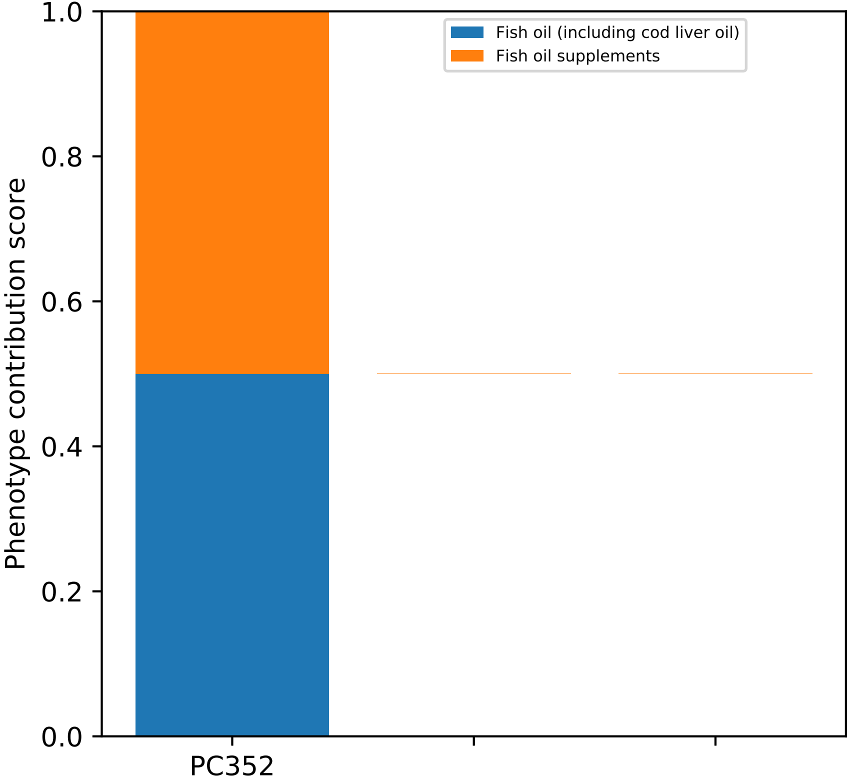
Fish oil (including cod liver oil)
Fish oil supplements
Phenotype contribution score
1.0
0.8
0.6
0.4
0.2
0.0
PC352

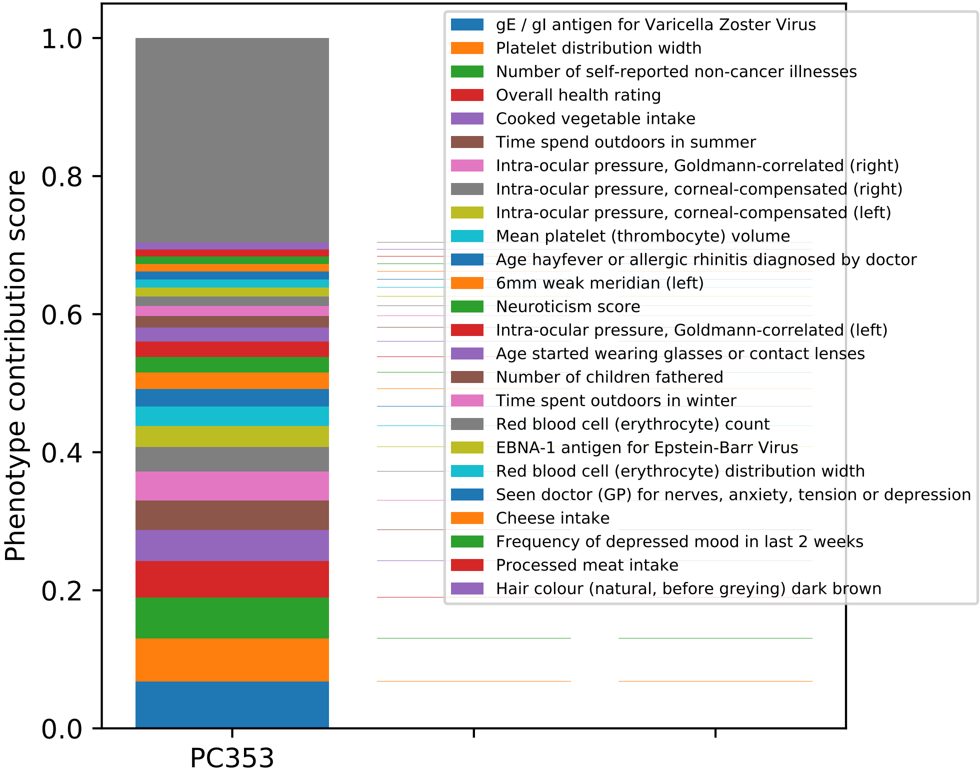
1.0
0.8
0.6
0.4
0.2
0.0
Phenotype contribution score
PC353
gE / gI antigen for Varicella Zoster Virus
Platelet distribution width
Number of self-reported non-cancer illnesses
Overall health rating
Cooked vegetable intake
Time spend outdoors in summer
Intra-ocular pressure, Goldmann-correlated (right)
Intra-ocular pressure, corneal-compensated (right)
Intra-ocular pressure, corneal-compensated (left)
Mean platelet (thrombocyte) volume
Age hayfever or allergic rhinitis diagnosed by doctor
6mm weak meridian (left)
Neuroticism score
Intra-ocular pressure, Goldmann-correlated (left)
Age started wearing glasses or contact lenses
Number of children fathered
Time spent outdoors in winter
Red blood cell (erythrocyte) count
EBNA-1 antigen for Epstein-Barr Virus
Red blood cell (erythrocyte) distribution width
Seen doctor (GP) for nerves, anxiety, tension or depression
Cheese intake
Frequency of depressed mood in last 2 weeks
Processed meat intake
Hair colour (natural, before greying) dark brown

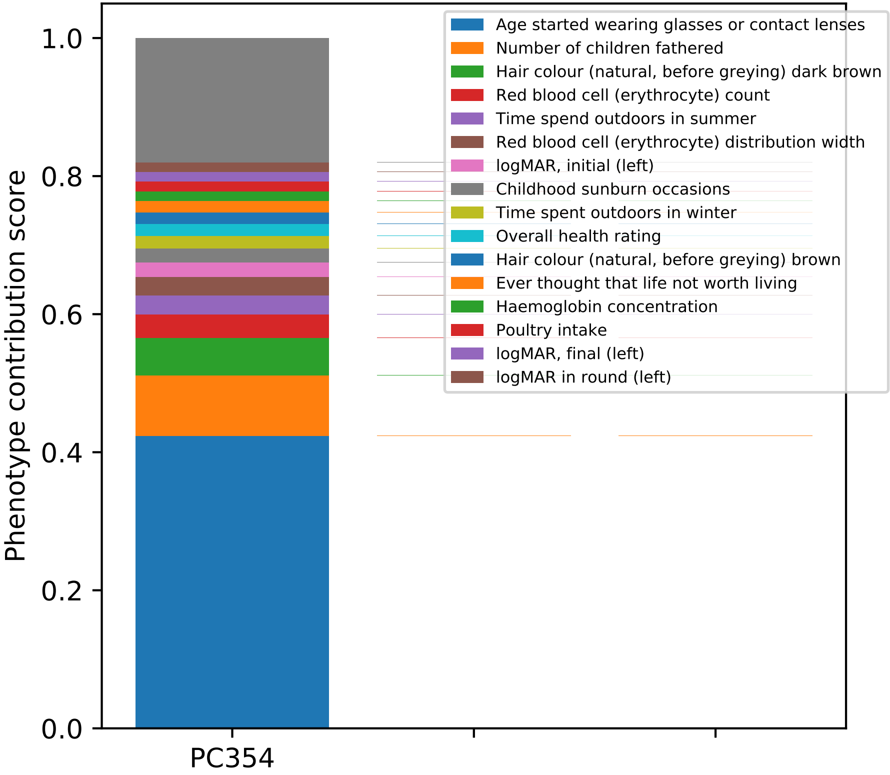

Phenotype contribution score
0.0
0.2
0.4
0.6
0.8
1.0
PC354
Age started wearing glasses or contact lenses
Number of children fathered
Hair colour (natural, before greying) dark brown
Red blood cell (erythrocyte) count
Time spend outdoors in summer
Red blood cell (erythrocyte) distribution width
logMAR, initial (left)
Childhood sunburn occasions
Time spent outdoors in winter
Overall health rating
Hair colour (natural, before greying) brown
Ever thought that life not worth living
Haemoglobin concentration
Poultry intake
logMAR, final (left)
logMAR in round (left)

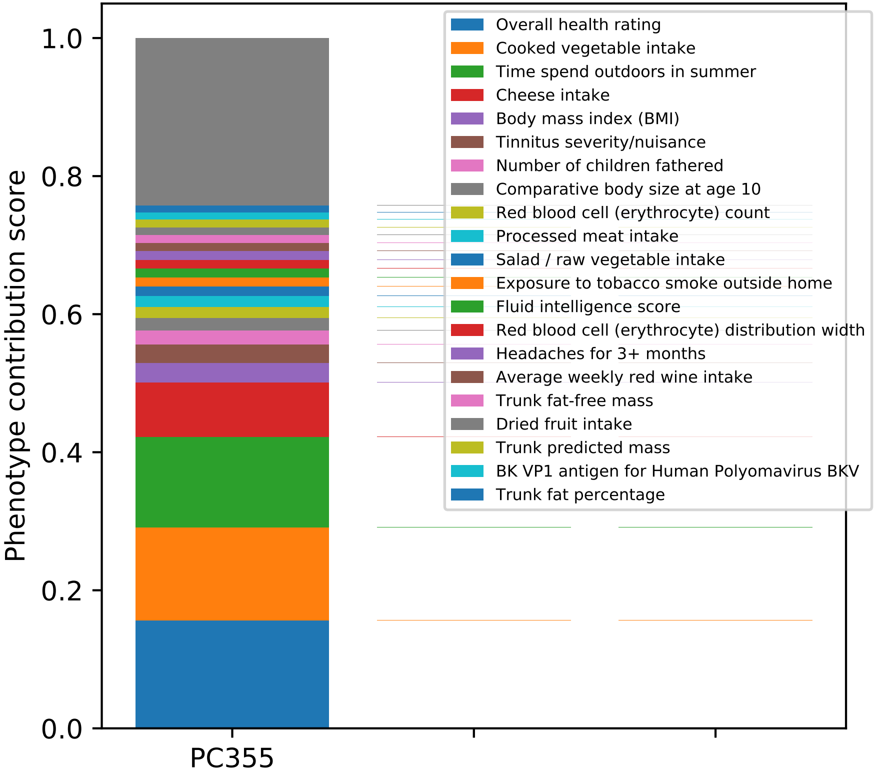

Phenotype contribution score
1.0
0.8
0.6
0.4
0.2
0.0
PC355
Overall health rating
Cooked vegetable intake
Time spend outdoors in summer
Cheese intake
Body mass index (BMI)
Tinnitus severity/nuisance
Number of children fathered
Comparative body size at age 10
Red blood cell (erythrocyte) count
Processed meat intake
Salad / raw vegetable intake
Exposure to tobacco smoke outside home
Fluid intelligence score
Red blood cell (erythrocyte) distribution width
Headaches for 3+ months
Average weekly red wine intake
Trunk fat-free mass
Dried fruit intake
Trunk predicted mass
BK VP1 antigen for Human Polyomavirus BKV
Trunk fat percentage

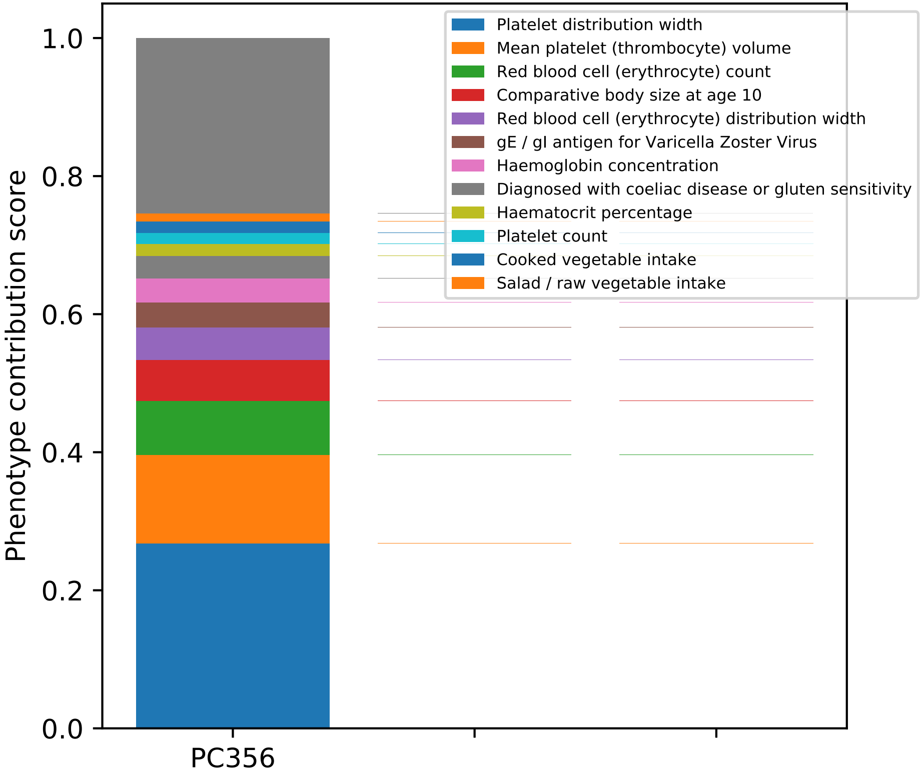
Phenotype contribution score
0.0
0.2
0.4
0.6
0.8
1.0
PC356
Platelet distribution width
Mean platelet (thrombocyte) volume
Red blood cell (erythrocyte) count
Comparative body size at age 10
Red blood cell (erythrocyte) distribution width
gE / gI antigen for Varicella Zoster Virus
Haemoglobin concentration
Diagnosed with coeliac disease or gluten sensitivity
Haematocrit percentage
Platelet count
Cooked vegetable intake
Salad / raw vegetable intake

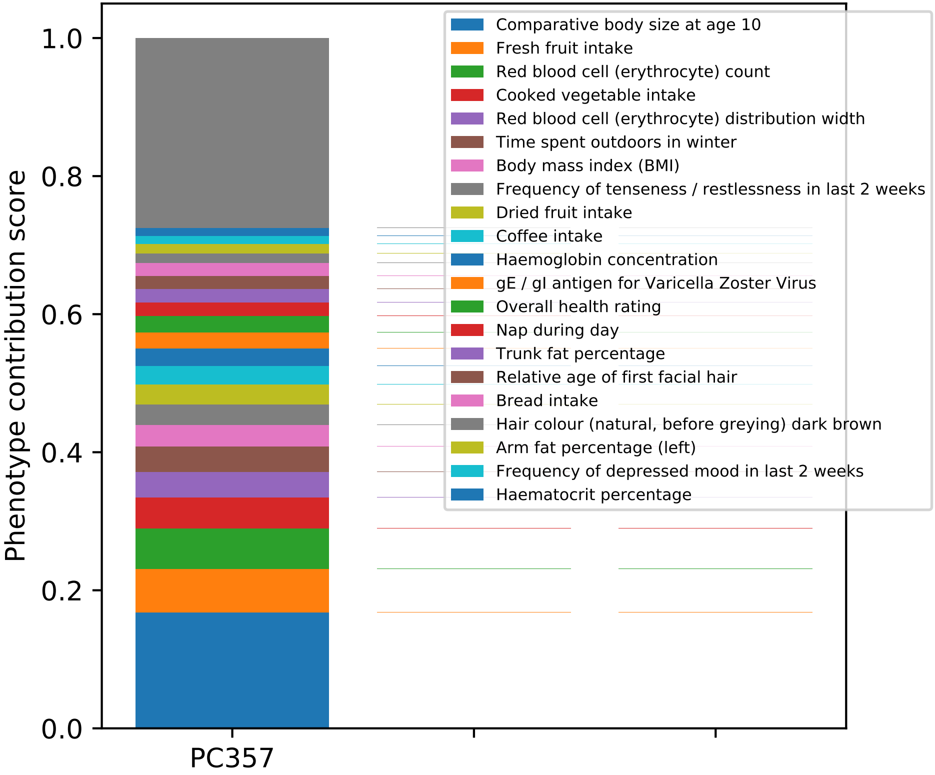

Phenotype contribution score
0.0
0.2
0.4
0.6
0.8
1.0
PC357
Comparative body size at age 10
Fresh fruit intake
Red blood cell (erythrocyte) count
Cooked vegetable intake
Red blood cell (erythrocyte) distribution width
Time spent outdoors in winter
Body mass index (BMI)
Frequency of tenseness / restlessness in last 2 weeks
Dried fruit intake
Coffee intake
Haemoglobin concentration
gE / gI antigen for Varicella Zoster Virus
Overall health rating
Nap during day
Trunk fat percentage
Relative age of first facial hair
Bread intake
Hair colour (natural, before greying) dark brown
Arm fat percentage (left)
Frequency of depressed mood in last 2 weeks
Haematocrit percentage

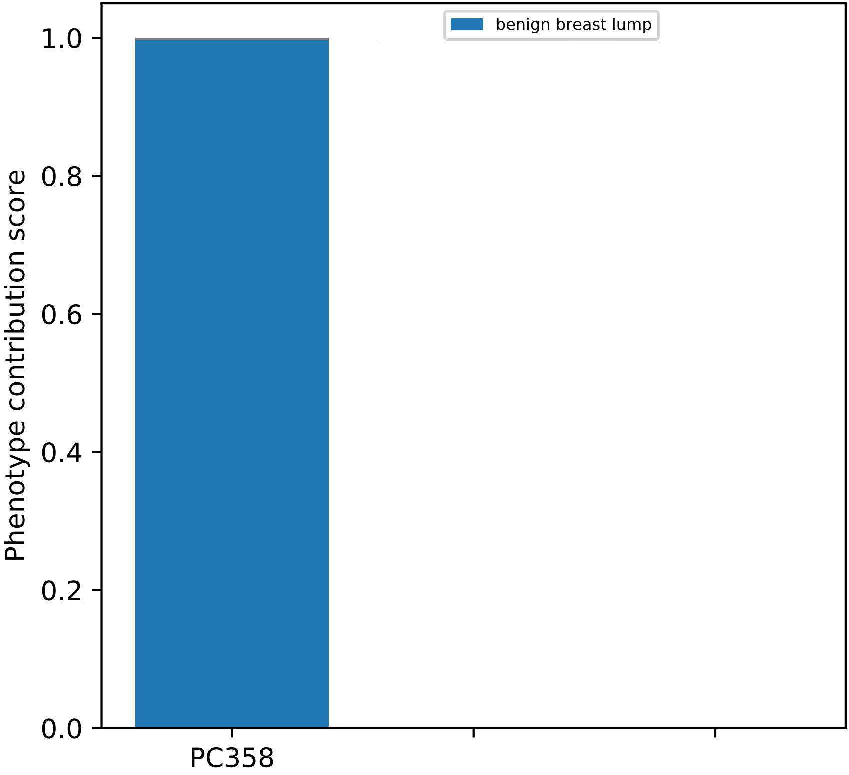
benign breast lump
Phenotype contribution score
1.0
0.8
0.6
0.4
0.2
0.0
PC358

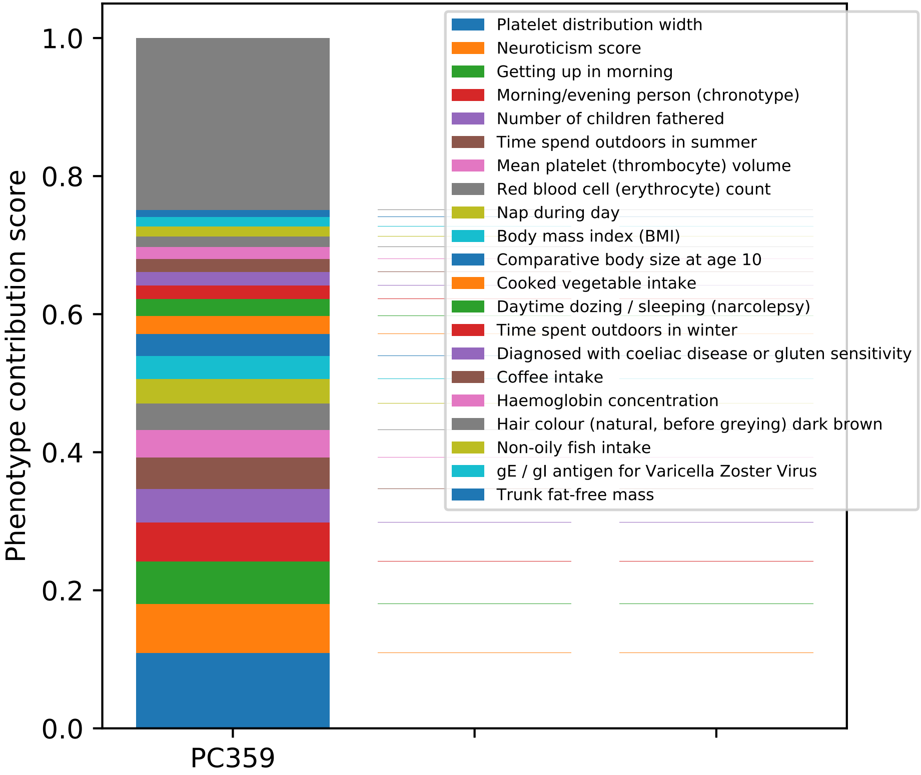

Phenotype contribution score
1.0
0.8
0.6
0.4
0.2
0.0
PC359
Platelet distribution width
Neuroticism score
Getting up in morning
Morning/evening person (chronotype)
Number of children fathered
Time spend outdoors in summer
Mean platelet (thrombocyte) volume
Red blood cell (erythrocyte) count
Nap during day
Body mass index (BMI)
Comparative body size at age 10
Cooked vegetable intake
Daytime dozing / sleeping (narcolepsy)
Time spent outdoors in winter
Diagnosed with coeliac disease or gluten sensitivity
Coffee intake
Haemoglobin concentration
Hair colour (natural, before greying) dark brown
Non-oily fish intake
gE / gI antigen for Varicella Zoster Virus
Trunk fat-free mass

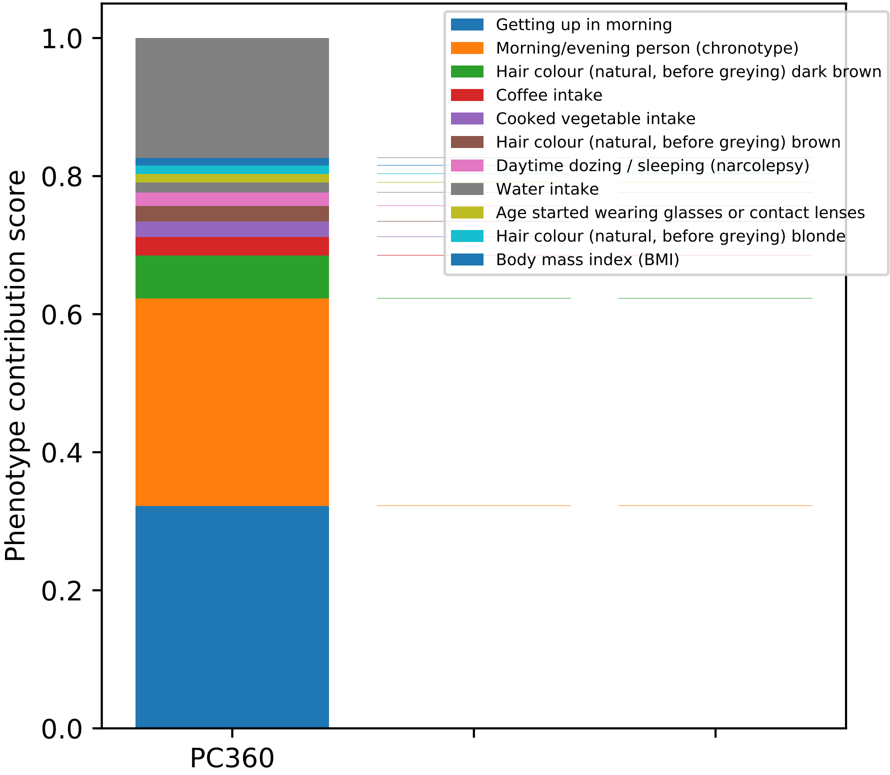

Phenotype contribution score
0.0
0.2
0.4
0.6
0.8
1.0
PC360
Getting up in morning
Morning/evening person (chronotype)
Hair colour (natural, before greying) dark brown
Coffee intake
Cooked vegetable intake
Hair colour (natural, before greying) brown
Daytime dozing / sleeping (narcolepsy)
Water intake
Age started wearing glasses or contact lenses
Hair colour (natural, before greying) blonde
Body mass index (BMI)

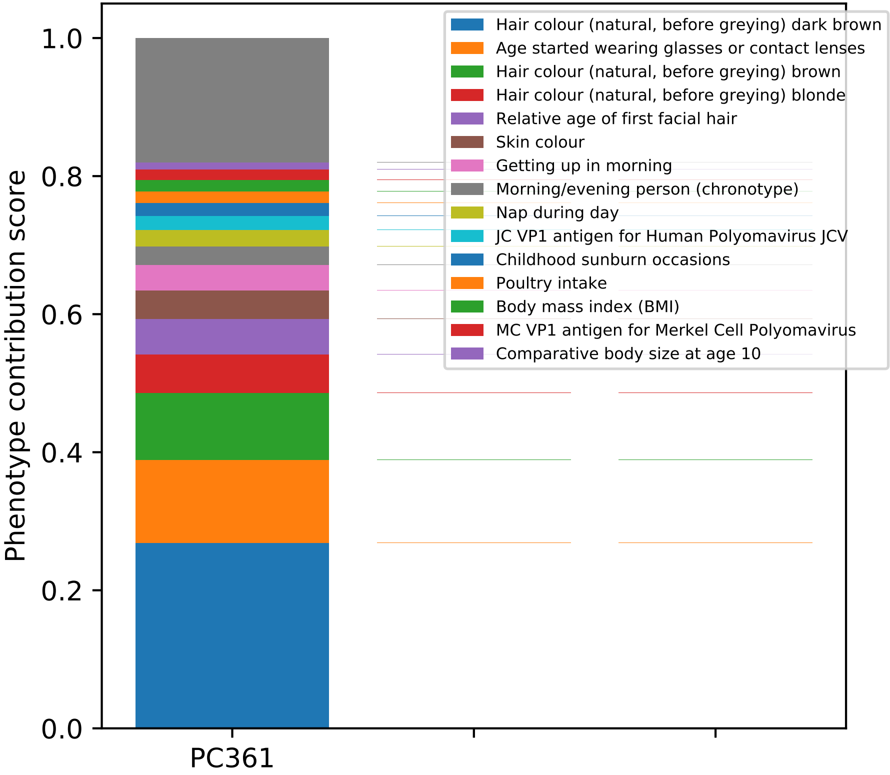

Phenotype contribution score
1.0
0.8
0.6
0.4
0.2
0.0
PC361
Hair colour (natural, before greying) dark brown
Age started wearing glasses or contact lenses
Hair colour (natural, before greying) brown
Hair colour (natural, before greying) blonde
Relative age of first facial hair
Skin colour
Getting up in morning
Morning/evening person (chronotype)
Nap during day
JC VP1 antigen for Human Polyomavirus JCV
Childhood sunburn occasions
Poultry intake
Body mass index (BMI)
MC VP1 antigen for Merkel Cell Polyomavirus
Comparative body size at age 10

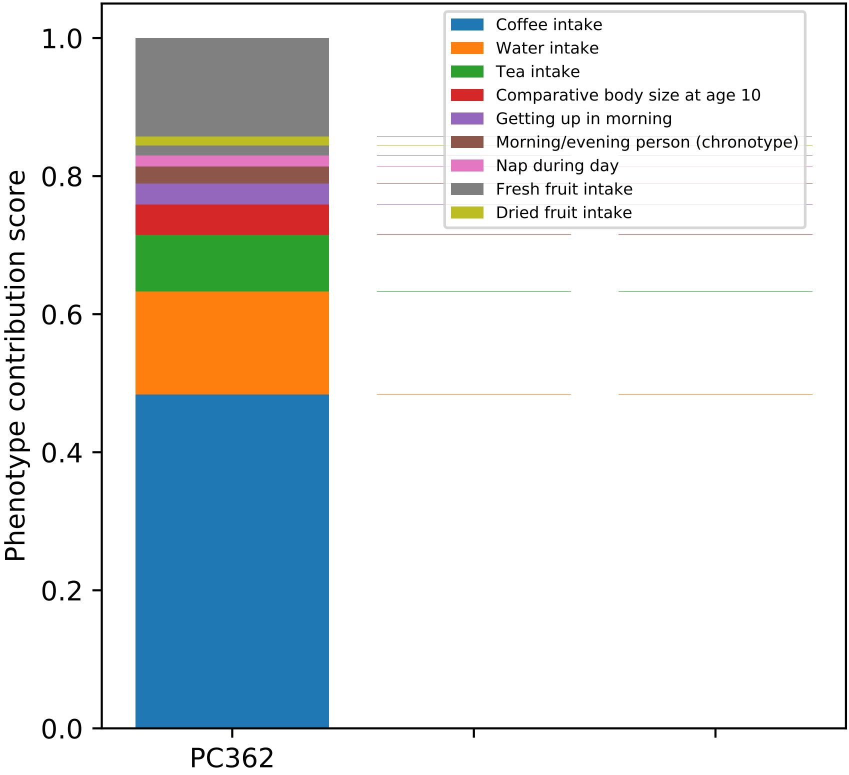

1.0
0.8
0.6
0.4
0.2
0.0
Phenotype contribution score
PC362
Coffee intake
Water intake
Tea intake
Comparative body size at age 10
Getting up in morning
Morning/evening person (chronotype)
Nap during day
Fresh fruit intake
Dried fruit intake

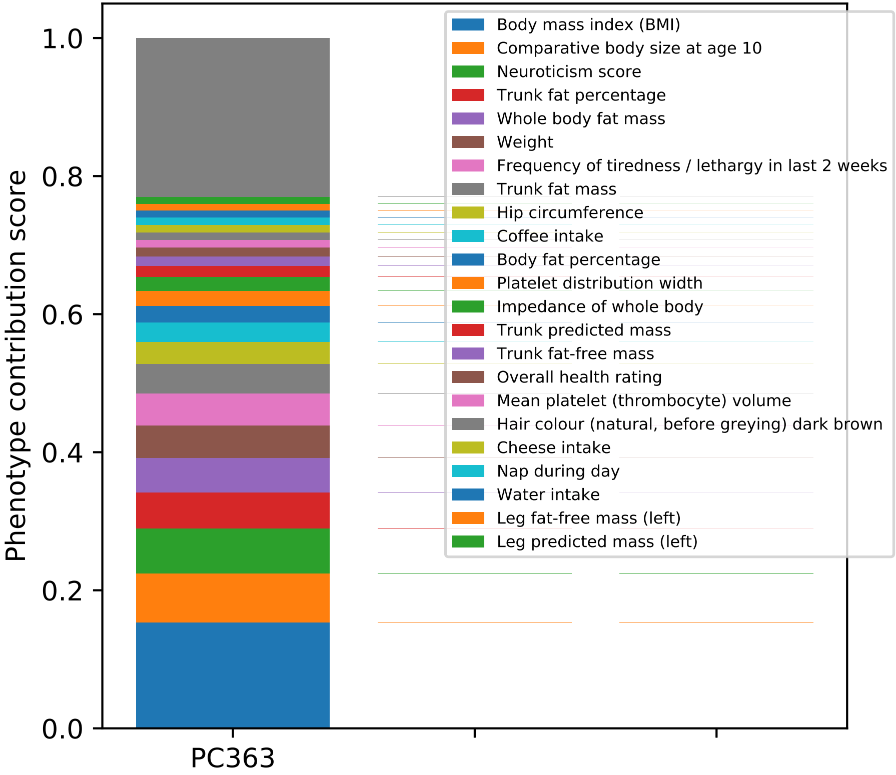

Phenotype contribution score
0.0
0.2
0.4
0.6
0.8
1.0
PC363
Body mass index (BMI)
Comparative body size at age 10
Neuroticism score
Trunk fat percentage
Whole body fat mass
Weight
Frequency of tiredness / lethargy in last 2 weeks
Trunk fat mass
Hip circumference
Coffee intake
Body fat percentage
Platelet distribution width
Impedance of whole body
Trunk predicted mass
Trunk fat-free mass
Overall health rating
Mean platelet (thrombocyte) volume
Hair colour (natural, before greying) dark brown
Cheese intake
Nap during day
Water intake
Leg fat-free mass (left)
Leg predicted mass (left)

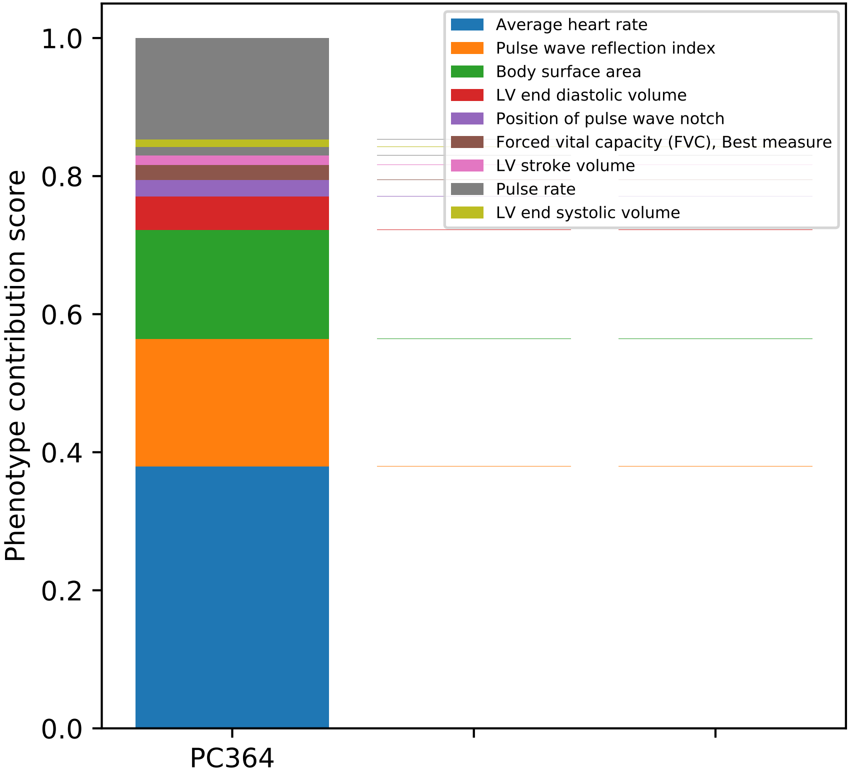

Average heart rate
Pulse wave reflection index
Body surface area
LV end diastolic volume
Position of pulse wave notch
Forced vital capacity (FVC), Best measure
LV stroke volume
Pulse rate
LV end systolic volume
Phenotype contribution score
0.0
0.2
0.4
0.6
0.8
1.0
PC364

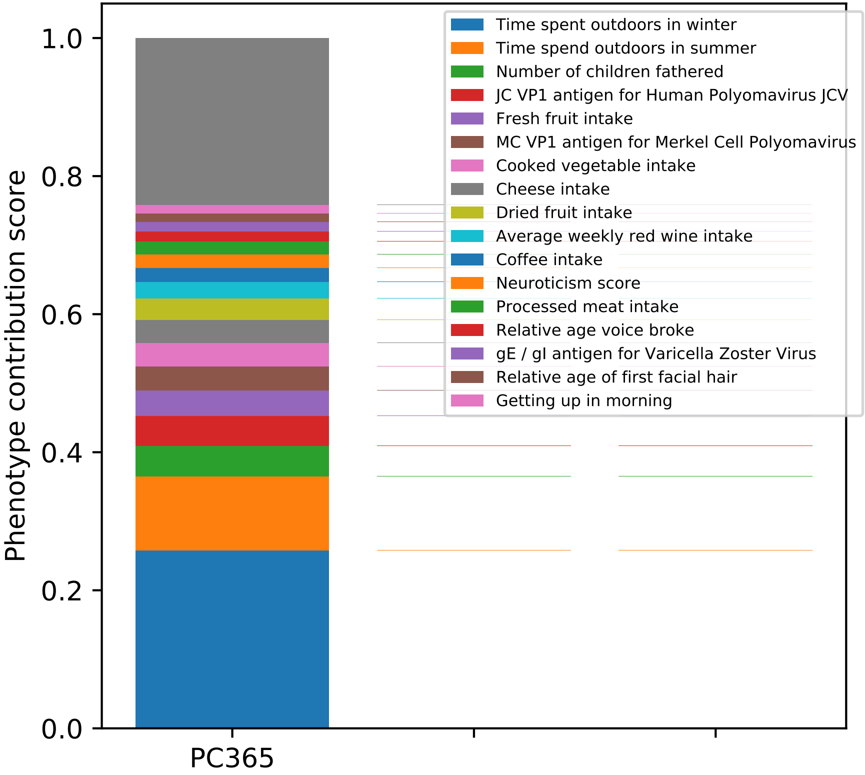

Phenotype contribution score
1.0
0.8
0.6
0.4
0.2
0.0
PC365
Time spent outdoors in winter
Time spend outdoors in summer
Number of children fathered
JC VP1 antigen for Human Polyomavirus JCV
Fresh fruit intake
MC VP1 antigen for Merkel Cell Polyomavirus
Cooked vegetable intake
Cheese intake
Dried fruit intake
Average weekly red wine intake
Coffee intake
Neuroticism score
Processed meat intake
Relative age voice broke
gE / gI antigen for Varicella Zoster Virus
Relative age of first facial hair
Getting up in morning

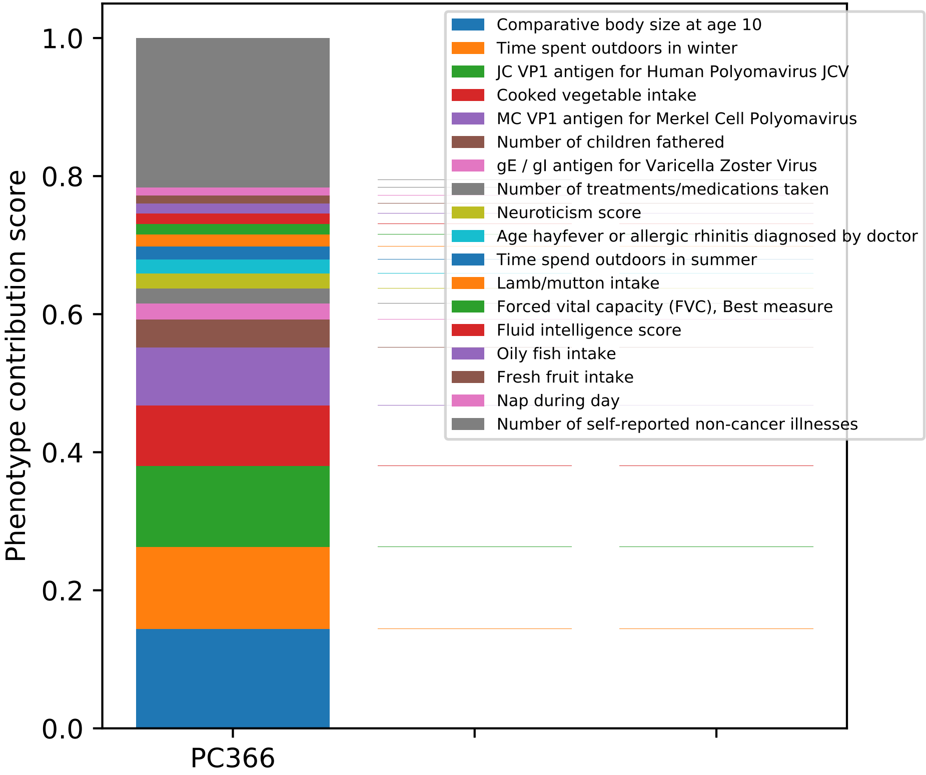

Phenotype contribution score
1.0
0.8
0.6
0.4
0.2
0.0
PC366
Comparative body size at age 10
Time spent outdoors in winter
JC VP1 antigen for Human Polyomavirus JCV
Cooked vegetable intake
MC VP1 antigen for Merkel Cell Polyomavirus
Number of children fathered
gE / gI antigen for Varicella Zoster Virus
Number of treatments/medications taken
Neuroticism score
Age hayfever or allergic rhinitis diagnosed by doctor
Time spend outdoors in summer
Lamb/mutton intake
Forced vital capacity (FVC), Best measure
Fluid intelligence score
Oily fish intake
Fresh fruit intake
Nap during day
Number of self-reported non-cancer illnesses

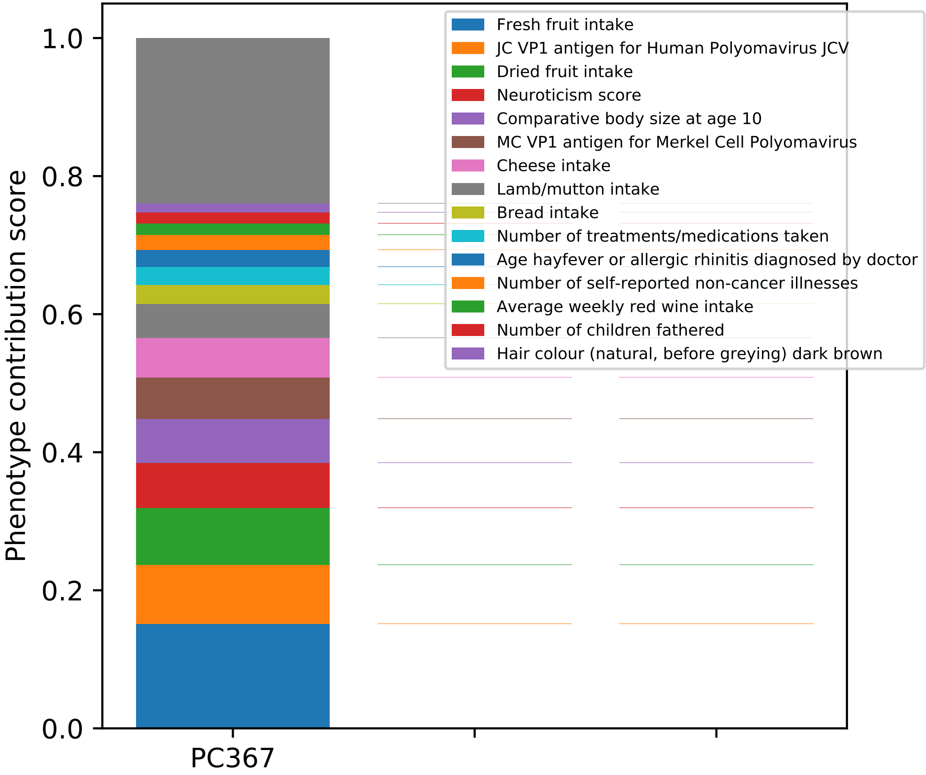

Phenotype contribution score
0.0
0.2
0.4
0.6
0.8
1.0
PC367
Fresh fruit intake
JC VP1 antigen for Human Polyomavirus JCV
Dried fruit intake
Neuroticism score
Comparative body size at age 10
MC VP1 antigen for Merkel Cell Polyomavirus
Cheese intake
Lamb/mutton intake
Bread intake
Number of treatments/medications taken
Age hayfever or allergic rhinitis diagnosed by doctor
Number of self-reported non-cancer illnesses
Average weekly red wine intake
Number of children fathered
Hair colour (natural, before greying) dark brown

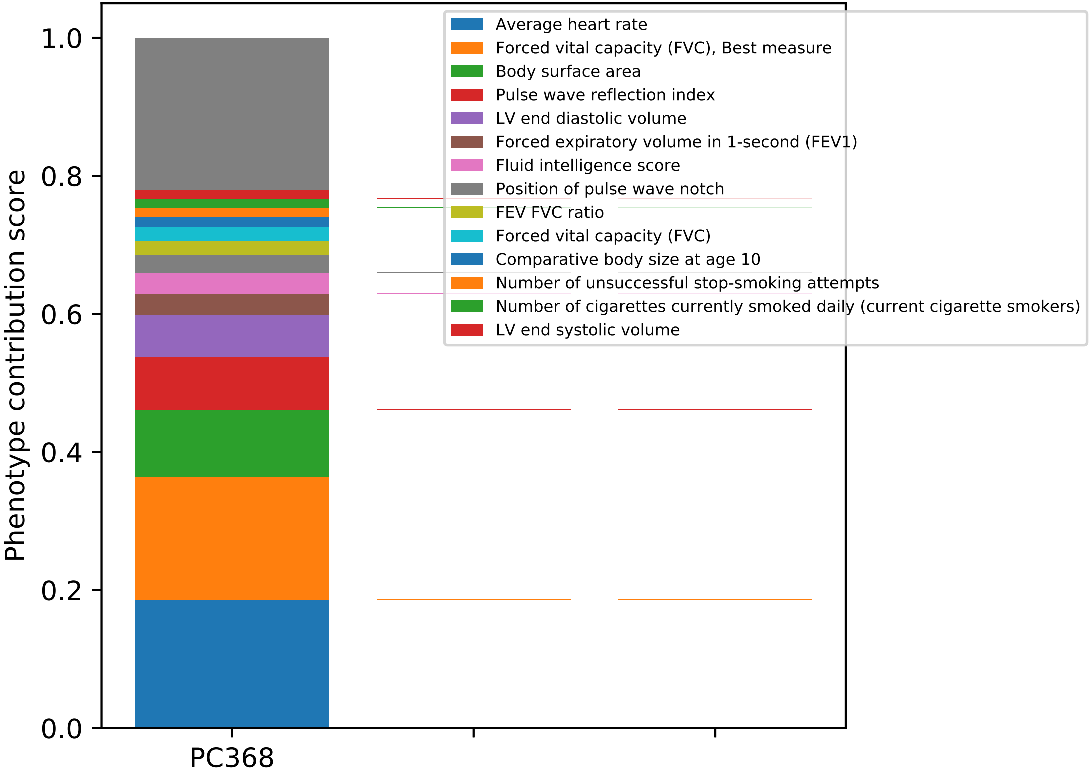
Phenotype contribution score
1.0
0.8
0.6
0.4
0.2
0.0
PC368
Average heart rate
Forced vital capacity (FVC), Best measure
Body surface area
Pulse wave reflection index
LV end diastolic volume
Forced expiratory volume in 1-second (FEV1)
Fluid intelligence score
Position of pulse wave notch
FEV FVC ratio
Forced vital capacity (FVC)
Comparative body size at age 10
Number of unsuccessful stop-smoking attempts
Number of cigarettes currently smoked daily (current cigarette smokers)
LV end systolic volume

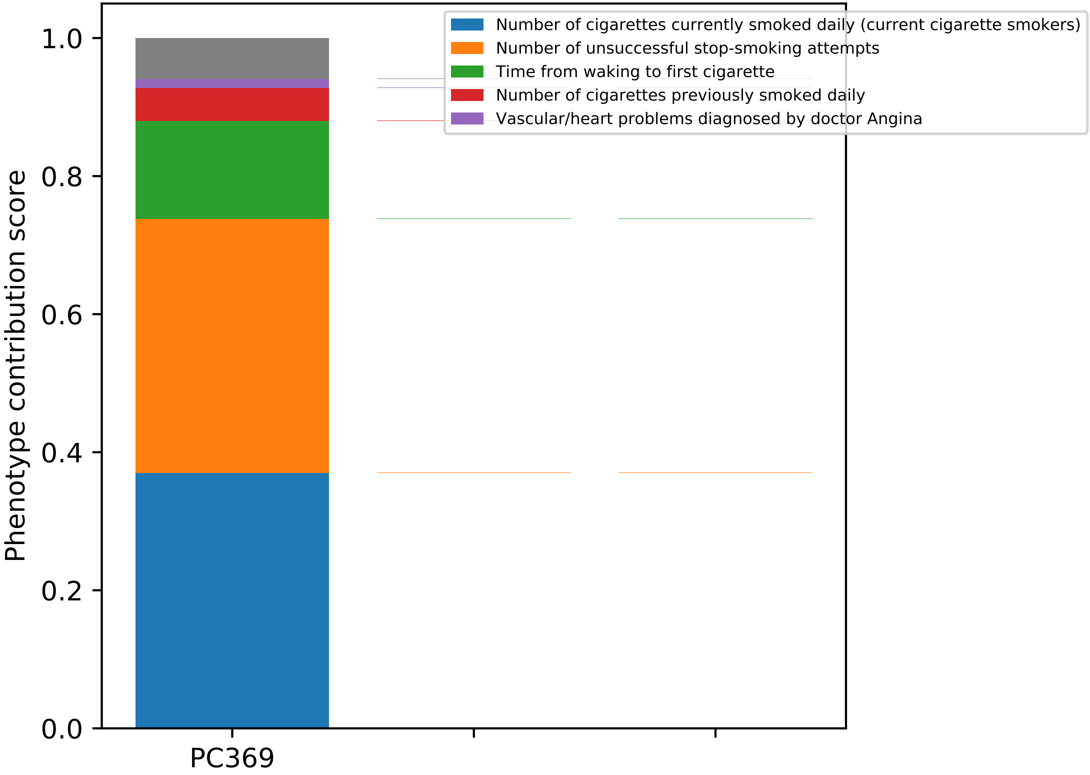
Number of cigarettes currently smoked daily (current cigarette smokers)
Number of unsuccessful stop-smoking attempts
Time from waking to first cigarette
Number of cigarettes previously smoked daily
Vascular/heart problems diagnosed by doctor Angina
Phenotype contribution score
1.0
0.8
0.6
0.4
0.2
0.0
PC369

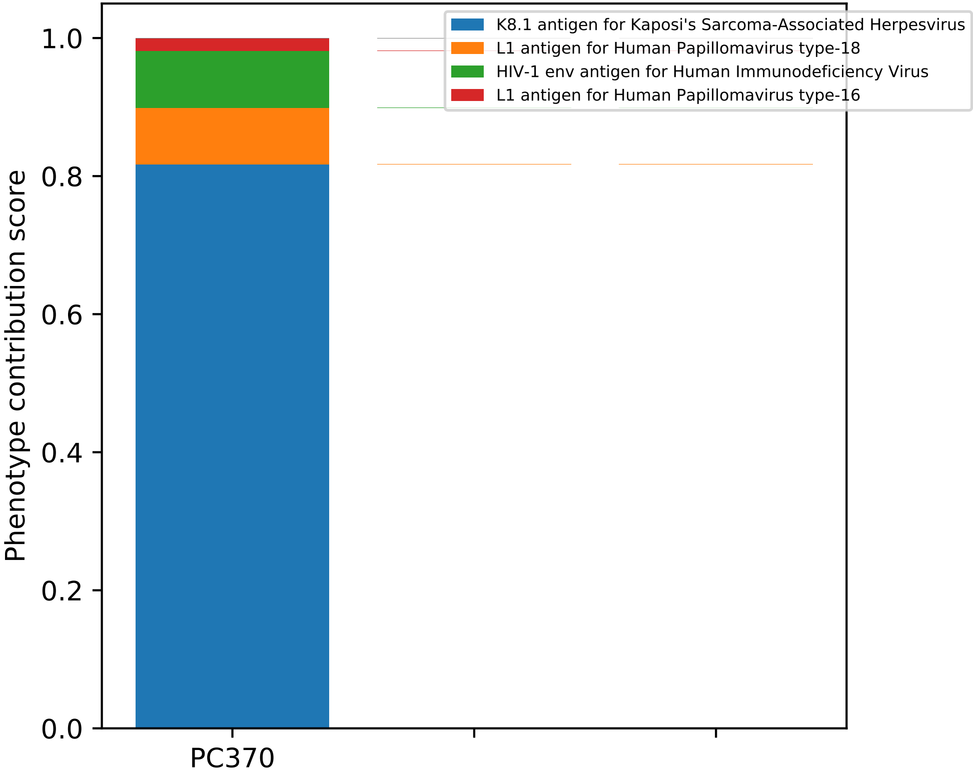
K8.1 antigen for Kaposi's Sarcoma-Associated Herpesvirus
L1 antigen for Human Papillomavirus type-18
HIV-1 env antigen for Human Immunodeficiency Virus
L1 antigen for Human Papillomavirus type-16
Phenotype contribution score
1.0
0.8
0.6
0.4
0.2
0.0
PC370

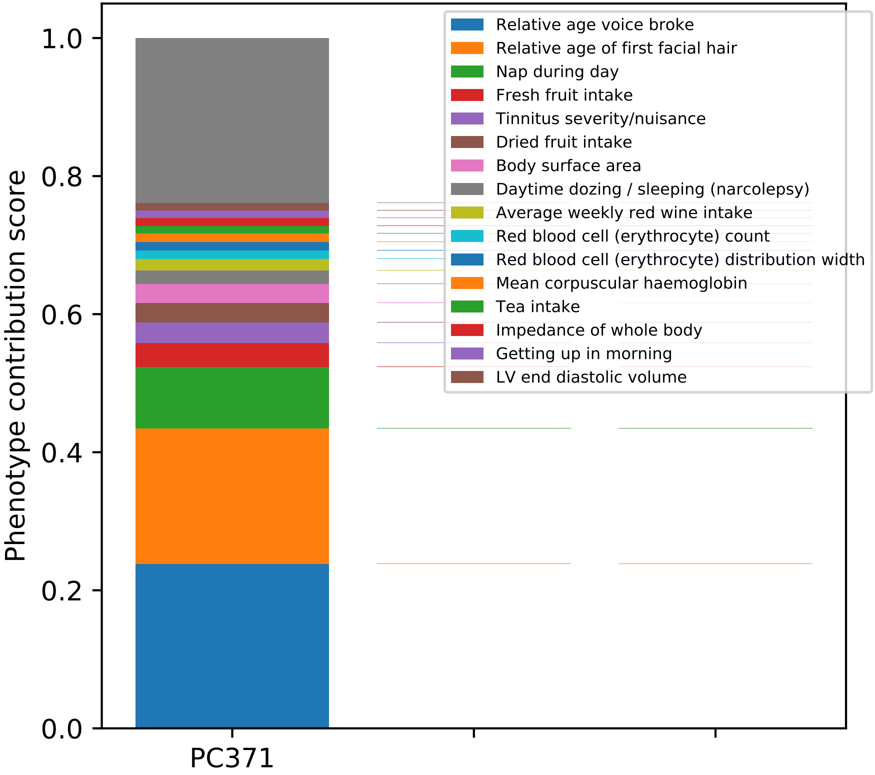

Phenotype contribution score
0.0
0.2
0.4
0.6
0.8
1.0
PC371
Relative age voice broke
Relative age of first facial hair
Nap during day
Fresh fruit intake
Tinnitus severity/nuisance
Dried fruit intake
Body surface area
Daytime dozing / sleeping (narcolepsy)
Average weekly red wine intake
Red blood cell (erythrocyte) count
Red blood cell (erythrocyte) distribution width
Mean corpuscular haemoglobin
Tea intake
Impedance of whole body
Getting up in morning
LV end diastolic volume

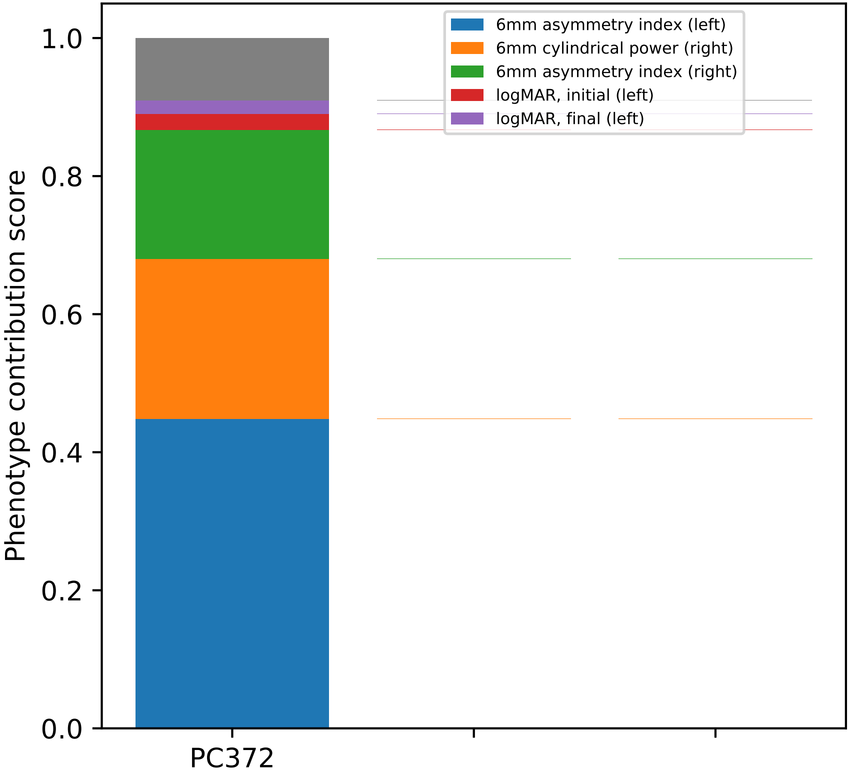

6mm asymmetry index (left)
6mm cylindrical power (right)
6mm asymmetry index (right)
logMAR, initial (left)
logMAR, final (left)
Phenotype contribution score
1.0
0.8
0.6
0.4
0.2
0.0
PC372

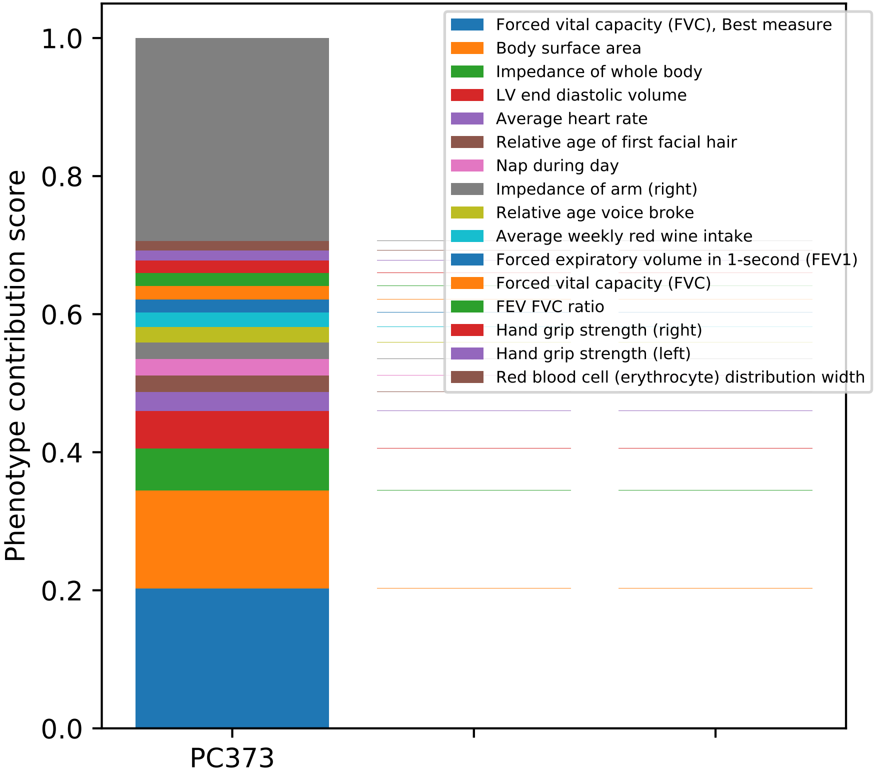
Phenotype contribution score
1.0
0.8
0.6
0.4
0.2
0.0
PC373
Forced vital capacity (FVC), Best measure
Body surface area
Impedance of whole body
LV end diastolic volume
Average heart rate
Relative age of first facial hair
Nap during day
Impedance of arm (right)
Relative age voice broke
Average weekly red wine intake
Forced expiratory volume in 1-second (FEV1)
Forced vital capacity (FVC)
FEV FVC ratio
Hand grip strength (right)
Hand grip strength (left)
Red blood cell (erythrocyte) distribution width

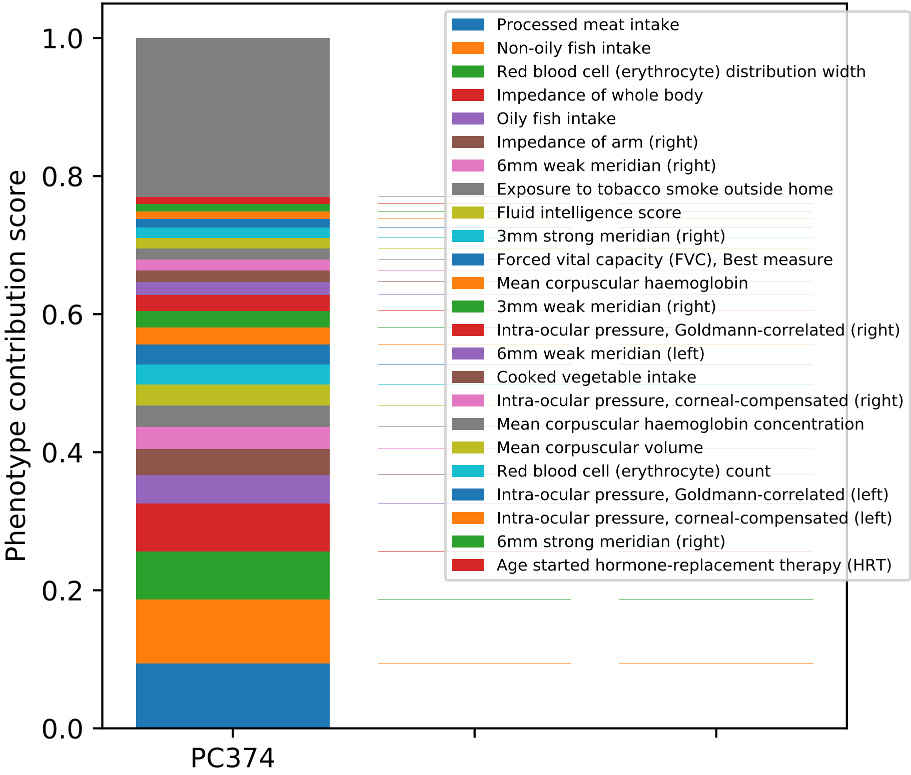

Phenotype contribution score
0.0
0.2
0.4
0.6
0.8
1.0
PC374
Processed meat intake
Non-oily fish intake
Red blood cell (erythrocyte) distribution width
Impedance of whole body
Oily fish intake
Impedance of arm (right)
6mm weak meridian (right)
Exposure to tobacco smoke outside home
Fluid intelligence score
3mm strong meridian (right)
Forced vital capacity (FVC), Best measure
Mean corpuscular haemoglobin
3mm weak meridian (right)
Intra-ocular pressure, Goldmann-correlated (right)
6mm weak meridian (left)
Cooked vegetable intake
Intra-ocular pressure, corneal-compensated (right)
Mean corpuscular haemoglobin concentration
Mean corpuscular volume
Red blood cell (erythrocyte) count
Intra-ocular pressure, Goldmann-correlated (left)
Intra-ocular pressure, corneal-compensated (left)
6mm strong meridian (right)
Age started hormone-replacement therapy (HRT)

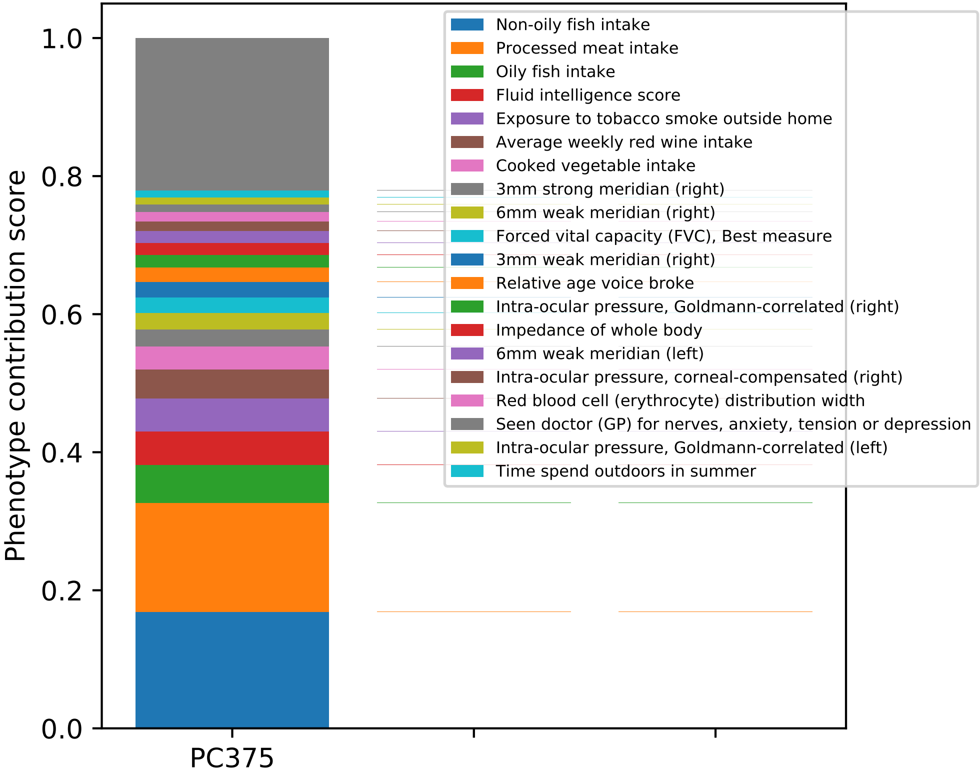

Phenotype contribution score
0.0
0.2
0.4
0.6
0.8
1.0
PC375
Non-oily fish intake
Processed meat intake
Oily fish intake
Fluid intelligence score
Exposure to tobacco smoke outside home
Average weekly red wine intake
Cooked vegetable intake
3mm strong meridian (right)
6mm weak meridian (right)
Forced vital capacity (FVC), Best measure
3mm weak meridian (right)
Relative age voice broke
Intra-ocular pressure, Goldmann-correlated (right)
Impedance of whole body
6mm weak meridian (left)
Intra-ocular pressure, corneal-compensated (right)
Red blood cell (erythrocyte) distribution width
Seen doctor (GP) for nerves, anxiety, tension or depression
Intra-ocular pressure, Goldmann-correlated (left)
Time spend outdoors in summer

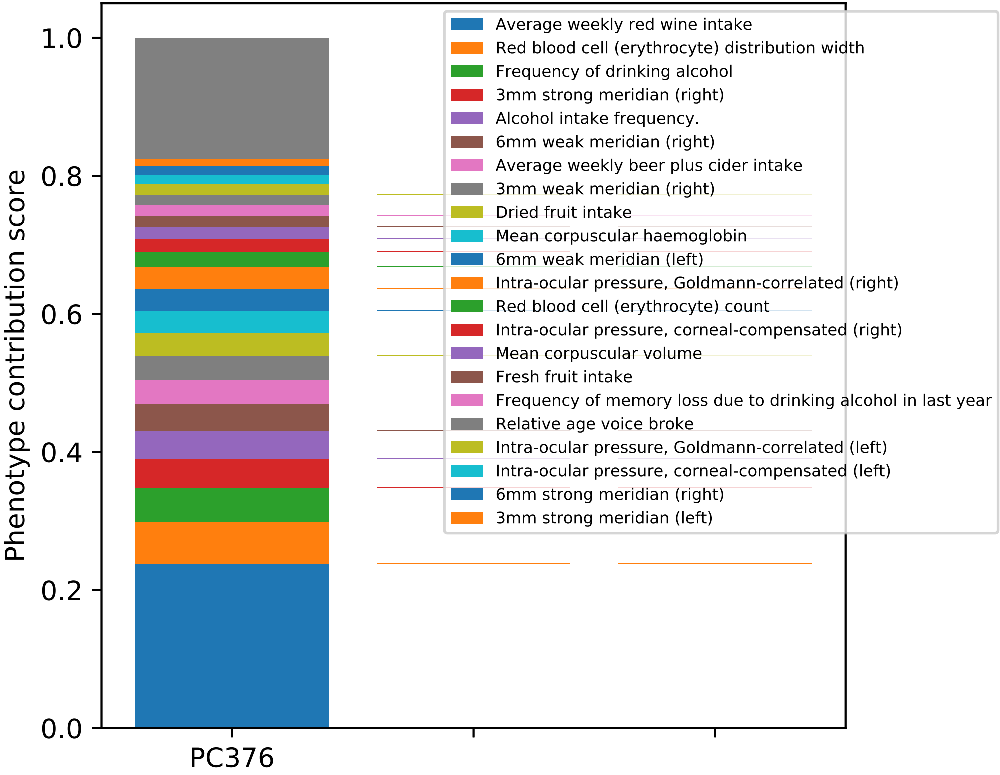

Phenotype contribution score
1.0
0.8
0.6
0.4
0.2
0.0
PC376
Average weekly red wine intake
Red blood cell (erythrocyte) distribution width
Frequency of drinking alcohol
3mm strong meridian (right)
Alcohol intake frequency.
6mm weak meridian (right)
Average weekly beer plus cider intake
3mm weak meridian (right)
Dried fruit intake
Mean corpuscular haemoglobin
6mm weak meridian (left)
Intra-ocular pressure, Goldmann-correlated (right)
Red blood cell (erythrocyte) count
Intra-ocular pressure, corneal-compensated (right)
Mean corpuscular volume
Fresh fruit intake
Frequency of memory loss due to drinking alcohol in last year
Relative age voice broke
Intra-ocular pressure, Goldmann-correlated (left)
Intra-ocular pressure, corneal-compensated (left)
6mm strong meridian (right)
3mm strong meridian (left)

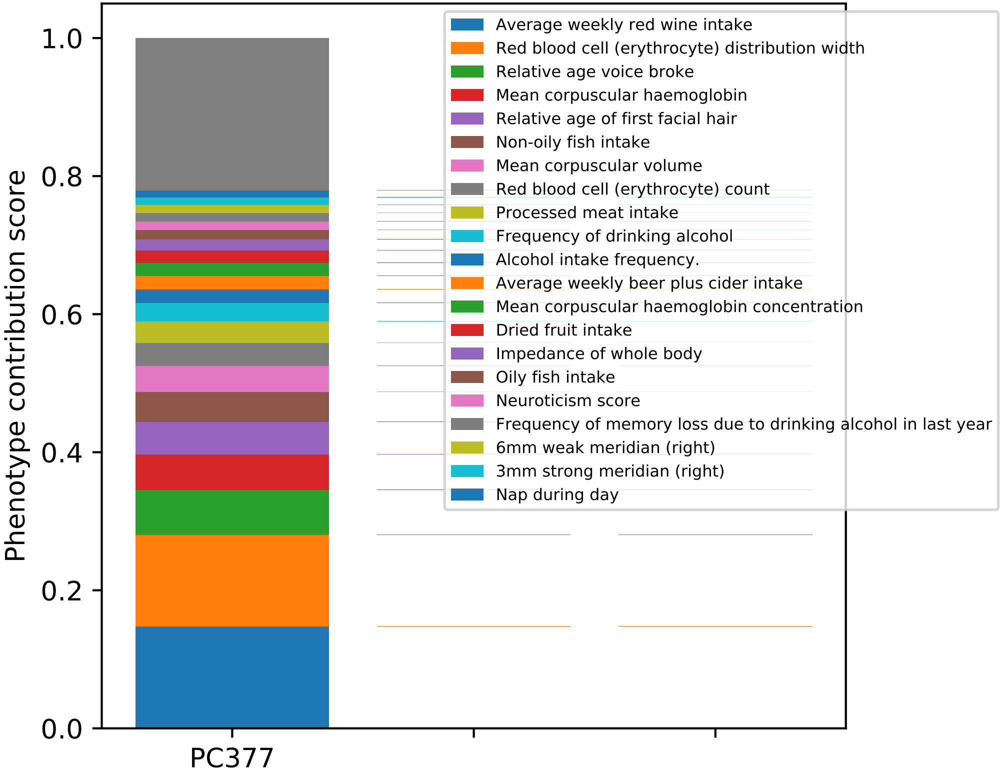

1.0
0.8
0.6
0.4
0.2
0.0
Phenotype contribution score
PC377
Average weekly red wine intake
Red blood cell (erythrocyte) distribution width
Relative age voice broke
Mean corpuscular haemoglobin
Relative age of first facial hair
Non-oily fish intake
Mean corpuscular volume
Red blood cell (erythrocyte) count
Processed meat intake
Frequency of drinking alcohol
Alcohol intake frequency.
Average weekly beer plus cider intake
Mean corpuscular haemoglobin concentration
Dried fruit intake
Impedance of whole body
Oily fish intake
Neuroticism score
Frequency of memory loss due to drinking alcohol in last year
6mm weak meridian (right)
3mm strong meridian (right)
Nap during day

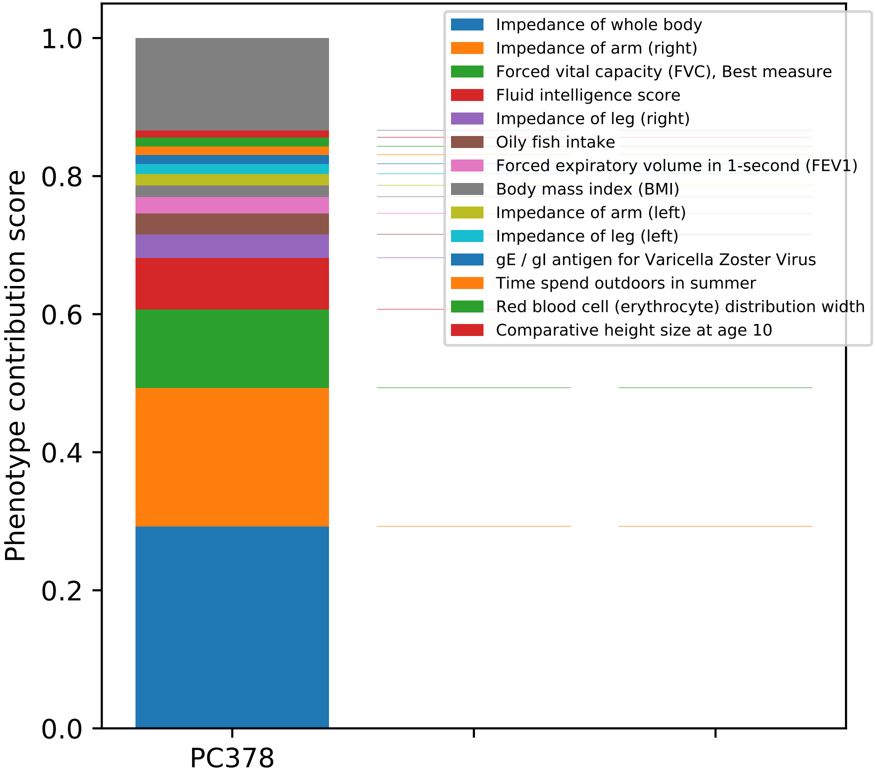

Phenotype contribution score
0.0
0.2
0.4
0.6
0.8
1.0
PC378
Impedance of whole body
Impedance of arm (right)
Forced vital capacity (FVC), Best measure
Fluid intelligence score
Impedance of leg (right)
Oily fish intake
Forced expiratory volume in 1-second (FEV1)
Body mass index (BMI)
Impedance of arm (left)
Impedance of leg (left)
gE / gI antigen for Varicella Zoster Virus
Time spend outdoors in summer
Red blood cell (erythrocyte) distribution width
Comparative height size at age 10

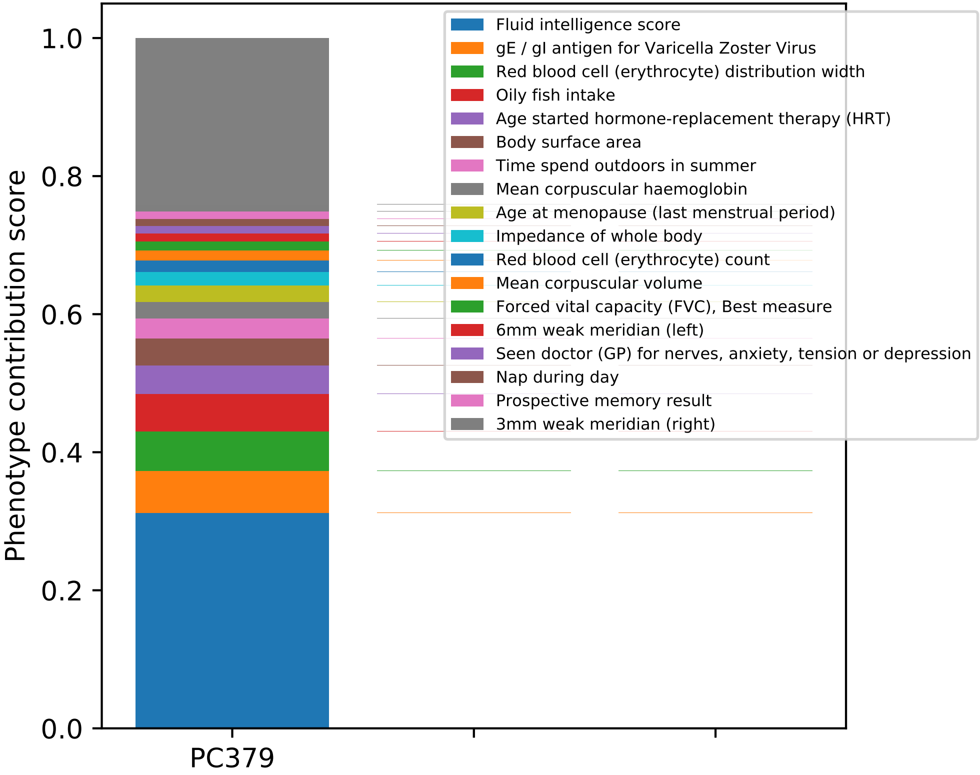

Phenotype contribution score
0.0
0.2
0.4
0.6
0.8
1.0
PC379
Fluid intelligence score
gE / gI antigen for Varicella Zoster Virus
Red blood cell (erythrocyte) distribution width
Oily fish intake
Age started hormone-replacement therapy (HRT)
Body surface area
Time spend outdoors in summer
Mean corpuscular haemoglobin
Age at menopause (last menstrual period)
Impedance of whole body
Red blood cell (erythrocyte) count
Mean corpuscular volume
Forced vital capacity (FVC), Best measure
6mm weak meridian (left)
Seen doctor (GP) for nerves, anxiety, tension or depression
Nap during day
Prospective memory result
3mm weak meridian (right)

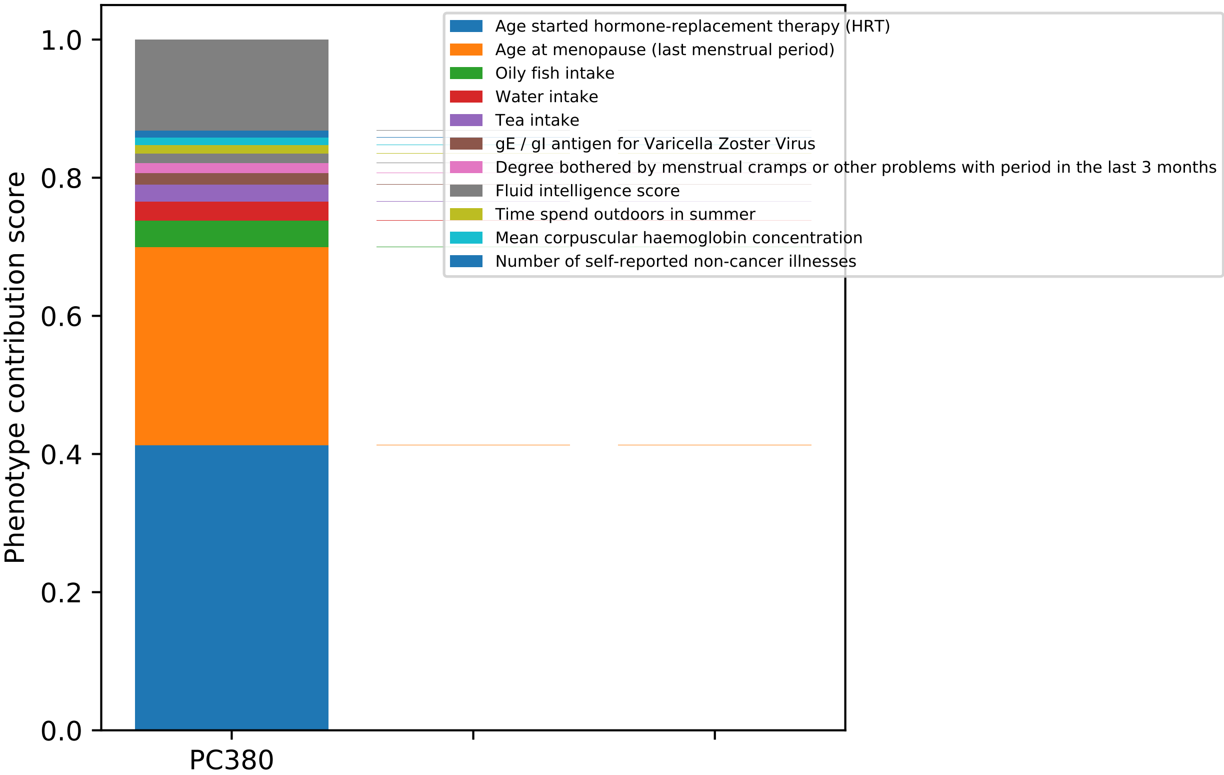
Age started hormone-replacement therapy (HRT)
Age at menopause (last menstrual period)
Oily fish intake
Water intake
Tea intake
gE / gI antigen for Varicella Zoster Virus
Degree bothered by menstrual cramps or other problems with period in the last 3 months
Fluid intelligence score
Time spend outdoors in summer
Mean corpuscular haemoglobin concentration
Number of self-reported non-cancer illnesses
Phenotype contribution score
1.0
0.8
0.6
0.4
0.2
0.0
PC380

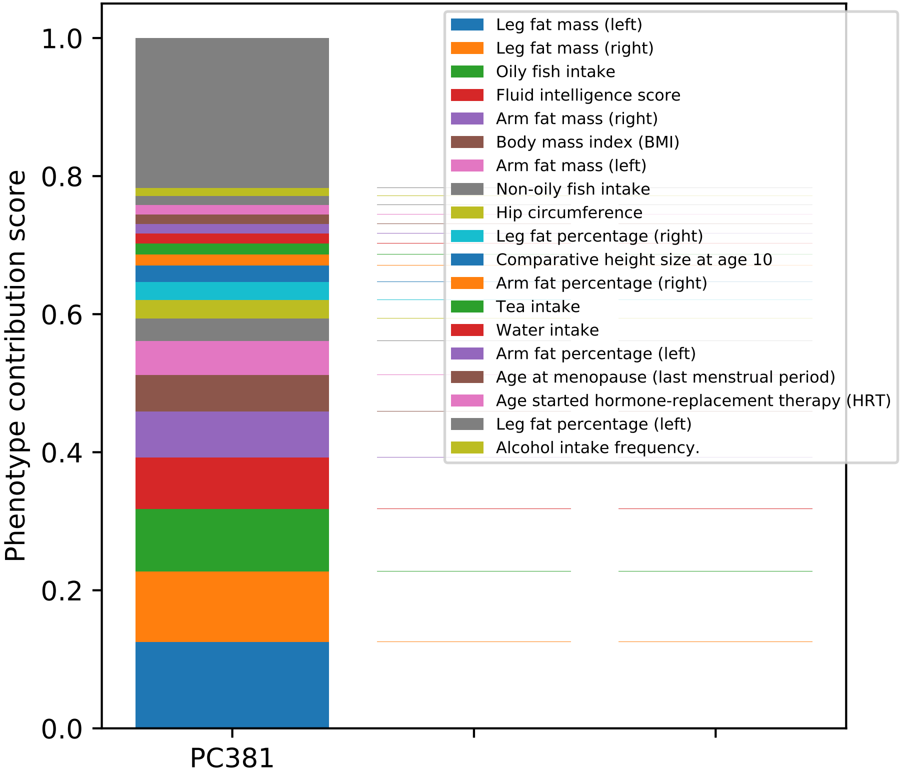
Phenotype contribution score
0.0
0.2
0.4
0.6
0.8
1.0
PC381
Leg fat mass (left)
Leg fat mass (right)
Oily fish intake
Fluid intelligence score
Arm fat mass (right)
Body mass index (BMI)
Arm fat mass (left)
Non-oily fish intake
Hip circumference
Leg fat percentage (right)
Comparative height size at age 10
Arm fat percentage (right)
Tea intake
Water intake
Arm fat percentage (left)
Age at menopause (last menstrual period)
Age started hormone-replacement therapy (HRT)
Leg fat percentage (left)
Alcohol intake frequency.

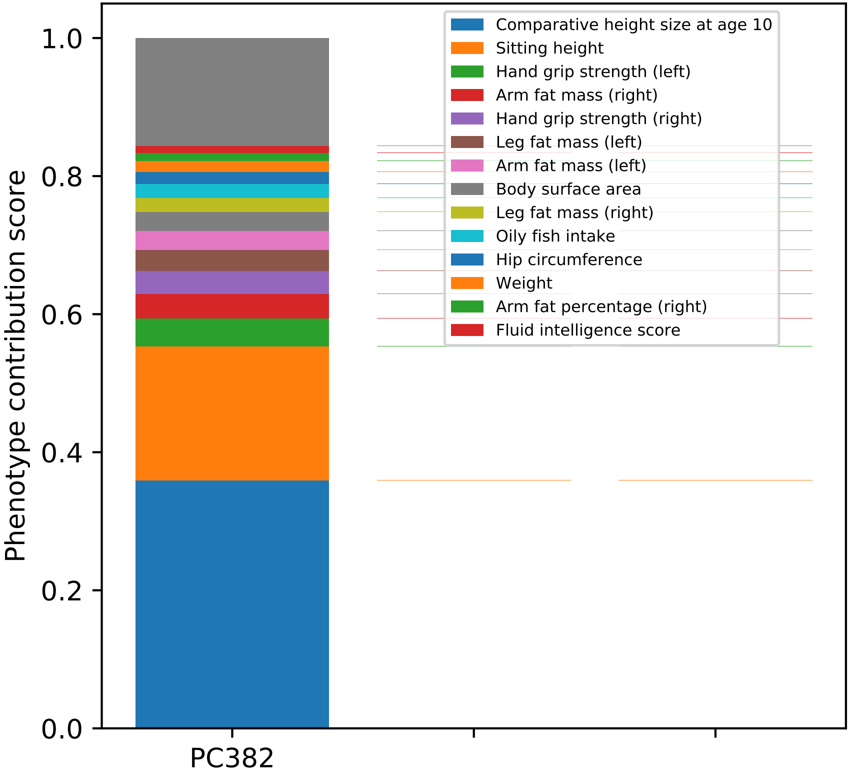

Phenotype contribution score
1.0
0.8
0.6
0.4
0.2
0.0
PC382
Comparative height size at age 10
Sitting height
Hand grip strength (left)
Arm fat mass (right)
Hand grip strength (right)
Leg fat mass (left)
Arm fat mass (left)
Body surface area
Leg fat mass (right)
Oily fish intake
Hip circumference
Weight
Arm fat percentage (right)
Fluid intelligence score

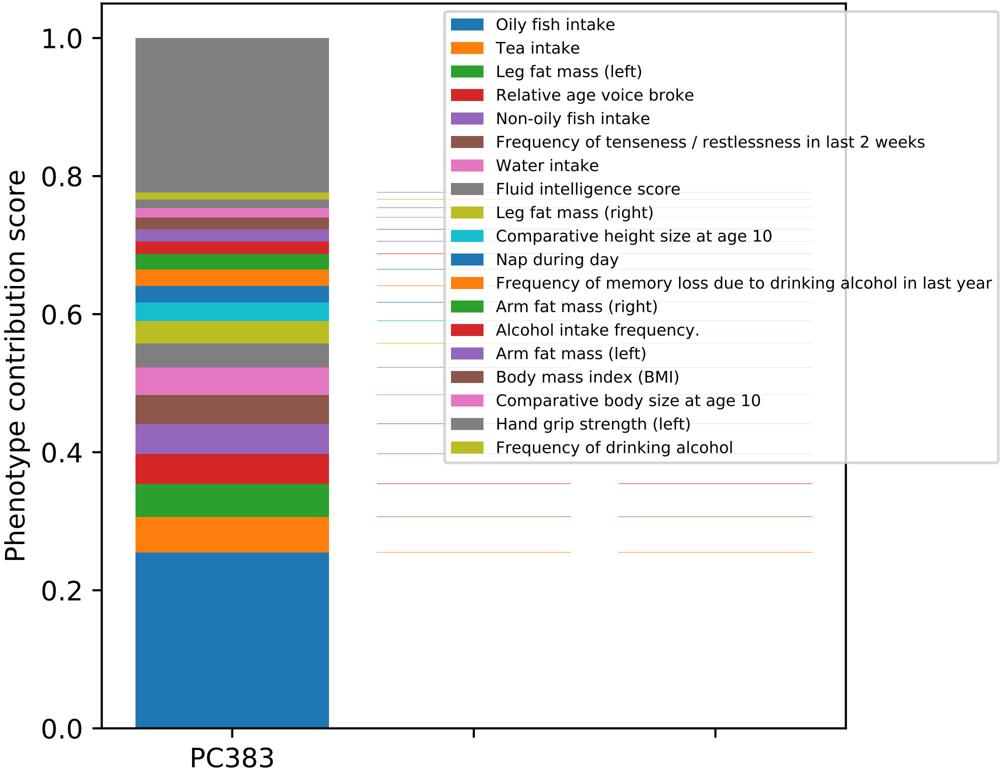

Phenotype contribution score
1.0
0.8
0.6
0.4
0.2
0.0
PC383
Oily fish intake
Tea intake
Leg fat mass (left)
Relative age voice broke
Non-oily fish intake
Frequency of tenseness / restlessness in last 2 weeks
Water intake
Fluid intelligence score
Leg fat mass (right)
Comparative height size at age 10
Nap during day
Frequency of memory loss due to drinking alcohol in last year
Arm fat mass (right)
Alcohol intake frequency.
Arm fat mass (left)
Body mass index (BMI)
Comparative body size at age 10
Hand grip strength (left)
Frequency of drinking alcohol

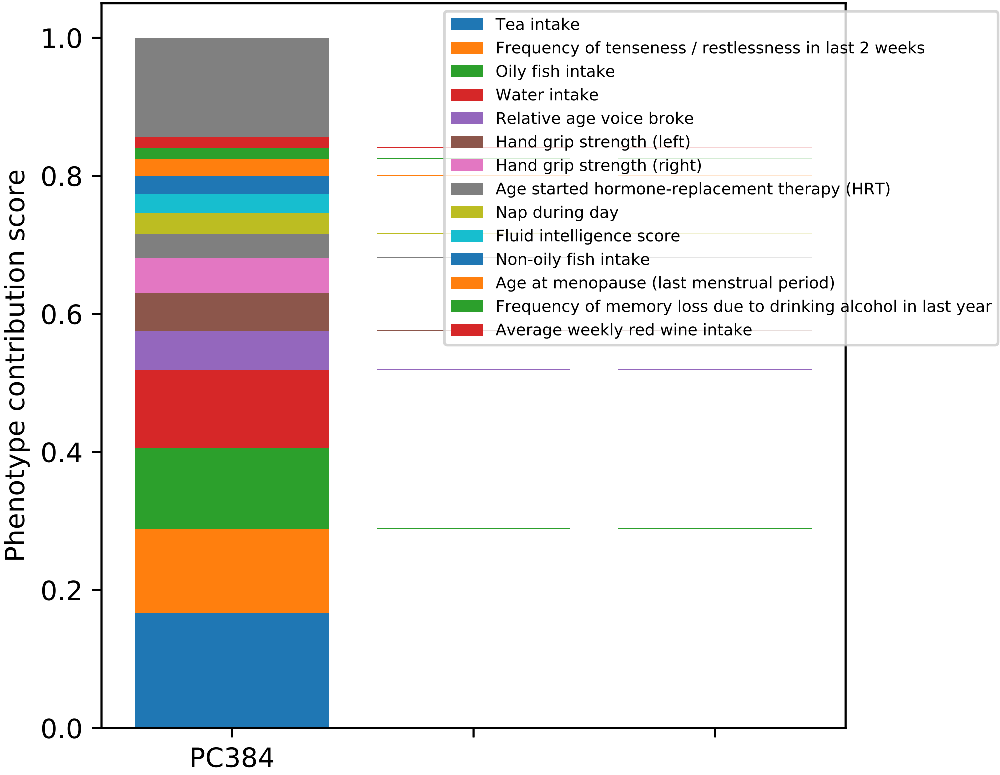

Phenotype contribution score
1.0
0.8
0.6
0.4
0.2
0.0
PC384
Tea intake
Frequency of tenseness / restlessness in last 2 weeks
Oily fish intake
Water intake
Relative age voice broke
Hand grip strength (left)
Hand grip strength (right)
Age started hormone-replacement therapy (HRT)
Nap during day
Fluid intelligence score
Non-oily fish intake
Age at menopause (last menstrual period)
Frequency of memory loss due to drinking alcohol in last year
Average weekly red wine intake

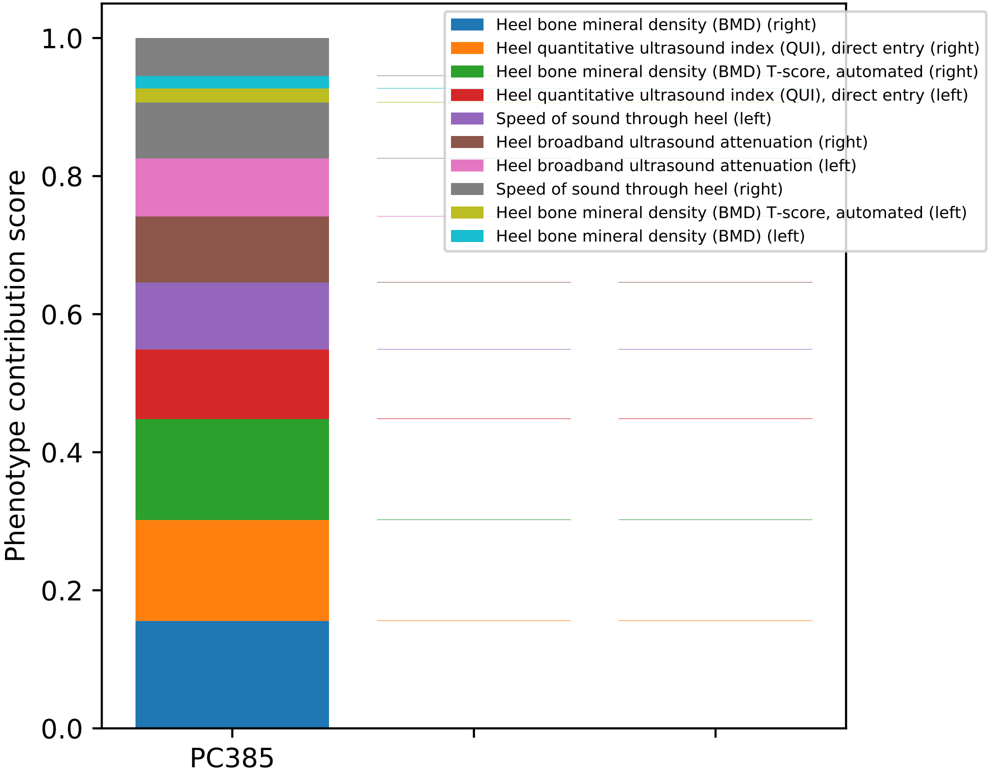
Phenotype contribution score
0.0
0.2
0.4
0.6
0.8
1.0
PC385
Heel bone mineral density (BMD) (right)
Heel quantitative ultrasound index (QUI), direct entry (right)
Heel bone mineral density (BMD) T-score, automated (right)
Heel quantitative ultrasound index (QUI), direct entry (left)
Speed of sound through heel (left)
Heel broadband ultrasound attenuation (right)
Heel broadband ultrasound attenuation (left)
Speed of sound through heel (right)
Heel bone mineral density (BMD) T-score, automated (left)
Heel bone mineral density (BMD) (left)

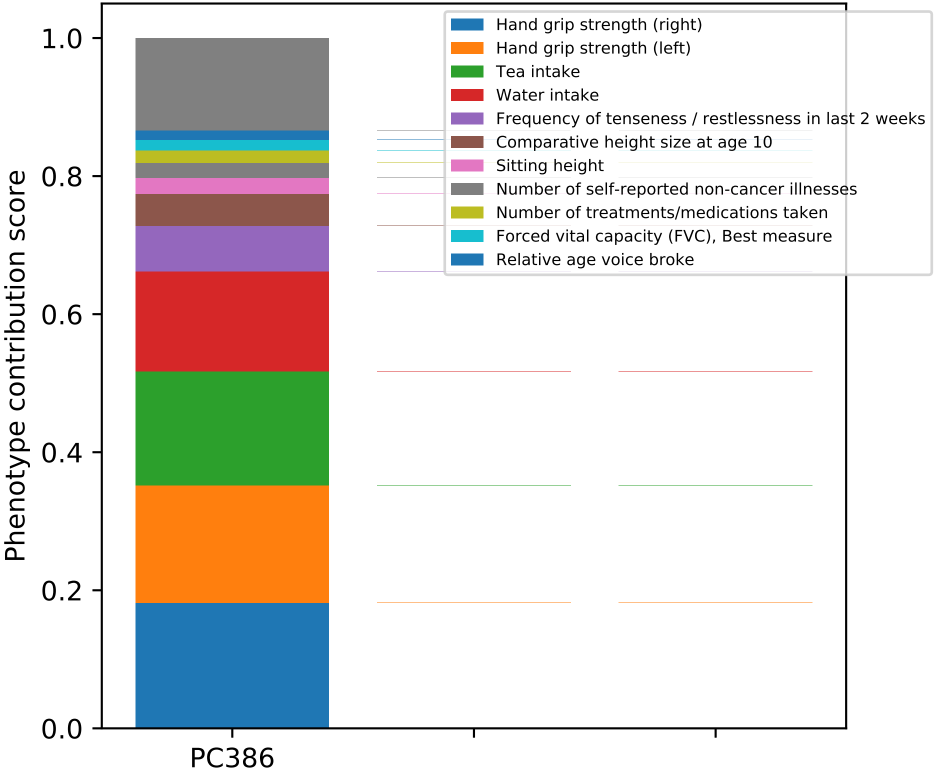
Phenotype contribution score
1.0
0.8
0.6
0.4
0.2
0.0
PC386
Hand grip strength (right)
Hand grip strength (left)
Tea intake
Water intake
Frequency of tenseness / restlessness in last 2 weeks
Comparative height size at age 10
Sitting height
Number of self-reported non-cancer illnesses
Number of treatments/medications taken
Forced vital capacity (FVC), Best measure
Relative age voice broke

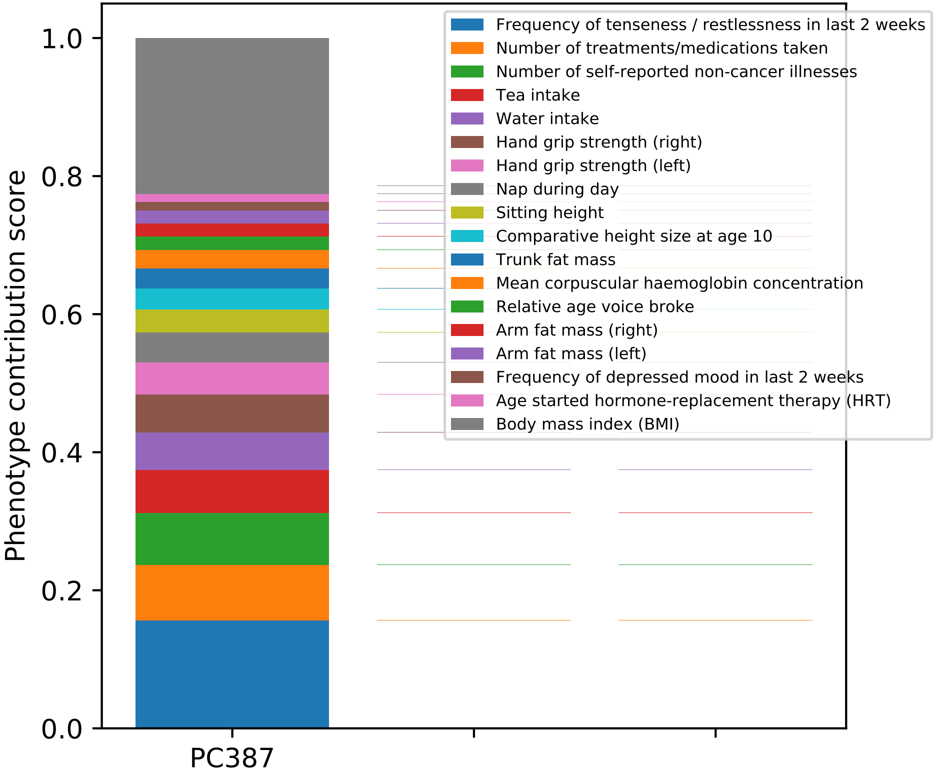
Phenotype contribution score
1.0
0.8
0.6
0.4
0.2
0.0
PC387
Frequency of tenseness / restlessness in last 2 weeks
Number of treatments/medications taken
Number of self-reported non-cancer illnesses
Tea intake
Water intake
Hand grip strength (right)
Hand grip strength (left)
Nap during day
Sitting height
Comparative height size at age 10
Trunk fat mass
Mean corpuscular haemoglobin concentration
Relative age voice broke
Arm fat mass (right)
Arm fat mass (left)
Frequency of depressed mood in last 2 weeks
Age started hormone-replacement therapy (HRT)
Body mass index (BMI)

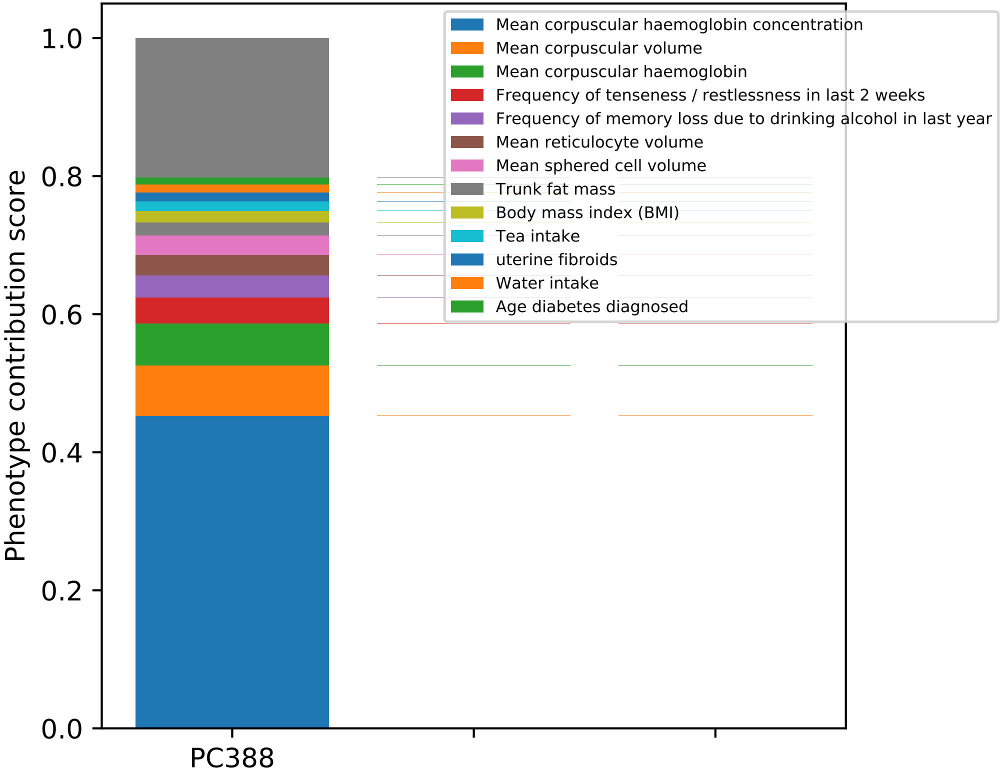

Mean corpuscular haemoglobin concentration
Mean corpuscular volume
Mean corpuscular haemoglobin
Frequency of tenseness / restlessness in last 2 weeks
Frequency of memory loss due to drinking alcohol in last year
Mean reticulocyte volume
Mean sphered cell volume
Trunk fat mass
Body mass index (BMI)
Tea intake
uterine fibroids
Water intake
Age diabetes diagnosed
Phenotype contribution score
1.0
0.8
0.6
0.4
0.2
0.0
PC388

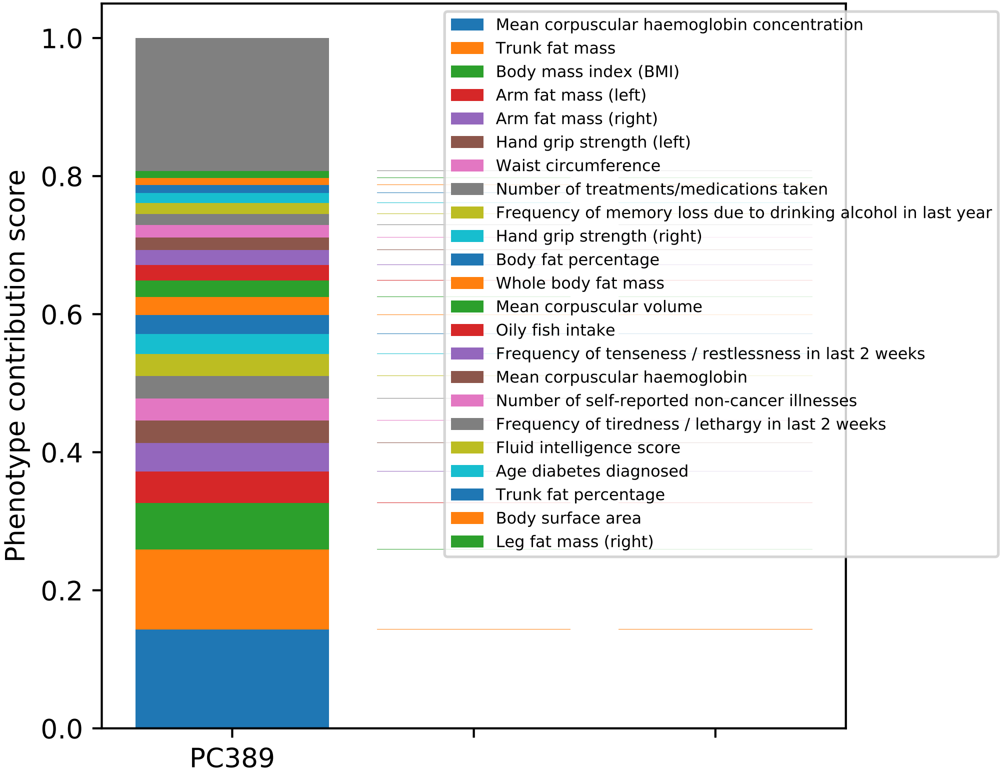

Phenotype contribution score
0.0
0.2
0.4
0.6
0.8
1.0
PC389
Mean corpuscular haemoglobin concentration
Trunk fat mass
Body mass index (BMI)
Arm fat mass (left)
Arm fat mass (right)
Hand grip strength (left)
Waist circumference
Number of treatments/medications taken
Frequency of memory loss due to drinking alcohol in last year
Hand grip strength (right)
Body fat percentage
Whole body fat mass
Mean corpuscular volume
Oily fish intake
Frequency of tenseness / restlessness in last 2 weeks
Mean corpuscular haemoglobin
Number of self-reported non-cancer illnesses
Frequency of tiredness / lethargy in last 2 weeks
Fluid intelligence score
Age diabetes diagnosed
Trunk fat percentage
Body surface area
Leg fat mass (right)

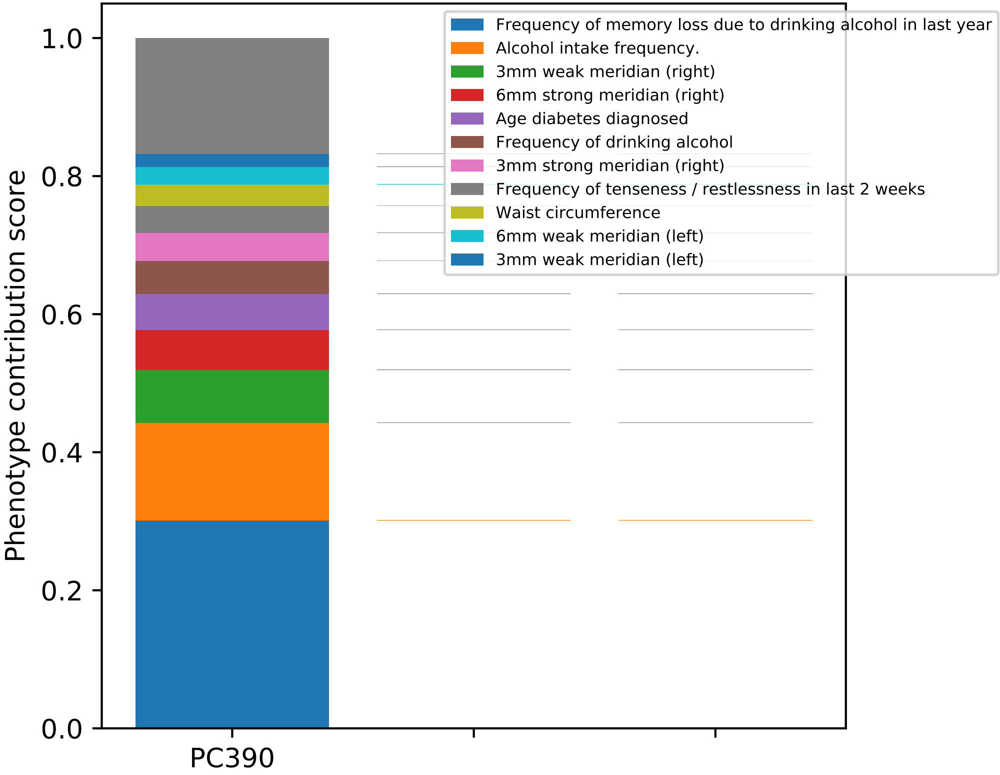

Phenotype contribution score
1.0
0.8
0.6
0.4
0.2
0.0
PC390
Frequency of memory loss due to drinking alcohol in last year
Alcohol intake frequency.
3mm weak meridian (right)
6mm strong meridian (right)
Age diabetes diagnosed
Frequency of drinking alcohol
3mm strong meridian (right)
Frequency of tenseness / restlessness in last 2 weeks
Waist circumference
6mm weak meridian (left)
3mm weak meridian (left)

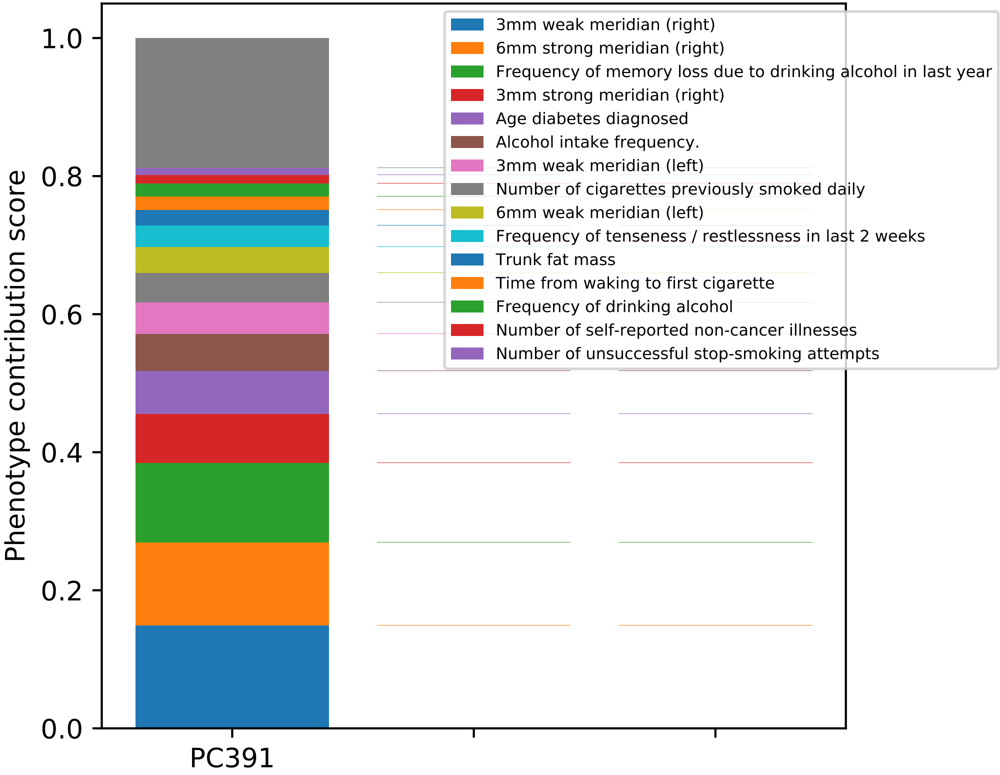

Phenotype contribution score
1.0
0.8
0.6
0.4
0.2
0.0
PC391
3mm weak meridian (right)
6mm strong meridian (right)
Frequency of memory loss due to drinking alcohol in last year
3mm strong meridian (right)
Age diabetes diagnosed
Alcohol intake frequency.
3mm weak meridian (left)
Number of cigarettes previously smoked daily
6mm weak meridian (left)
Frequency of tenseness / restlessness in last 2 weeks
Trunk fat mass
Time from waking to first cigarette
Frequency of drinking alcohol
Number of self-reported non-cancer illnesses
Number of unsuccessful stop-smoking attempts

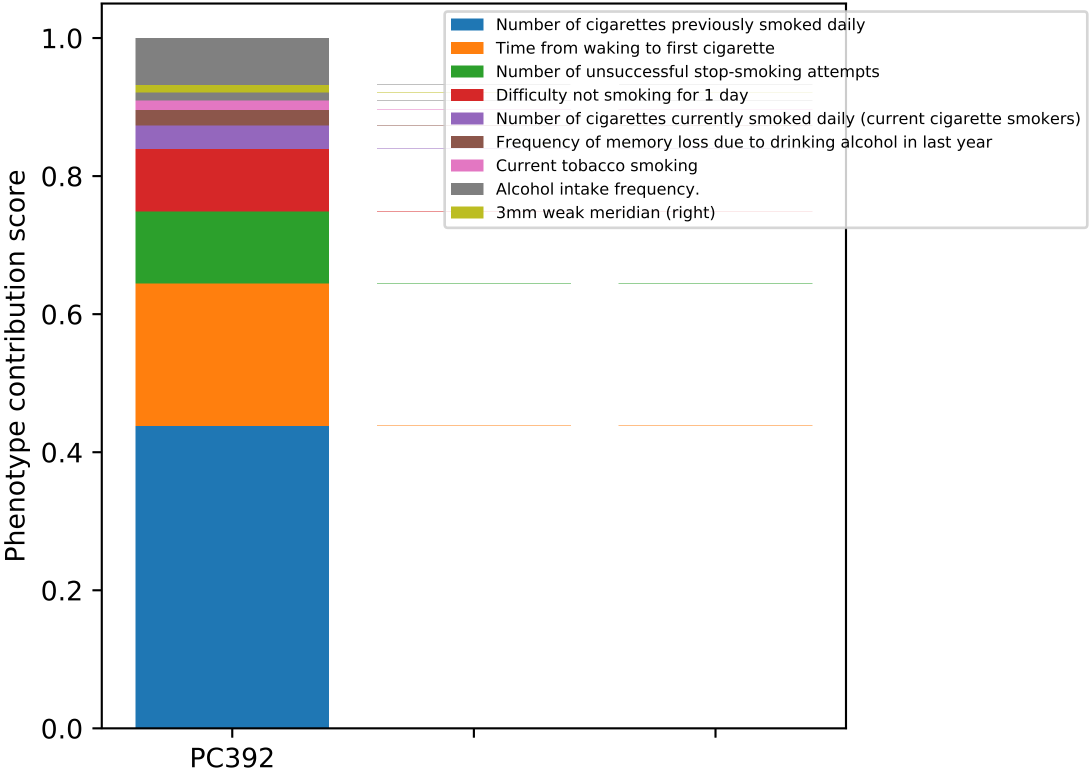

Phenotype contribution score
0.0
0.2
0.4
0.6
0.8
1.0
PC392
Number of cigarettes previously smoked daily
Time from waking to first cigarette
Number of unsuccessful stop-smoking attempts
Difficulty not smoking for 1 day
Number of cigarettes currently smoked daily (current cigarette smokers)
Frequency of memory loss due to drinking alcohol in last year
Current tobacco smoking
Alcohol intake frequency.
3mm weak meridian (right)

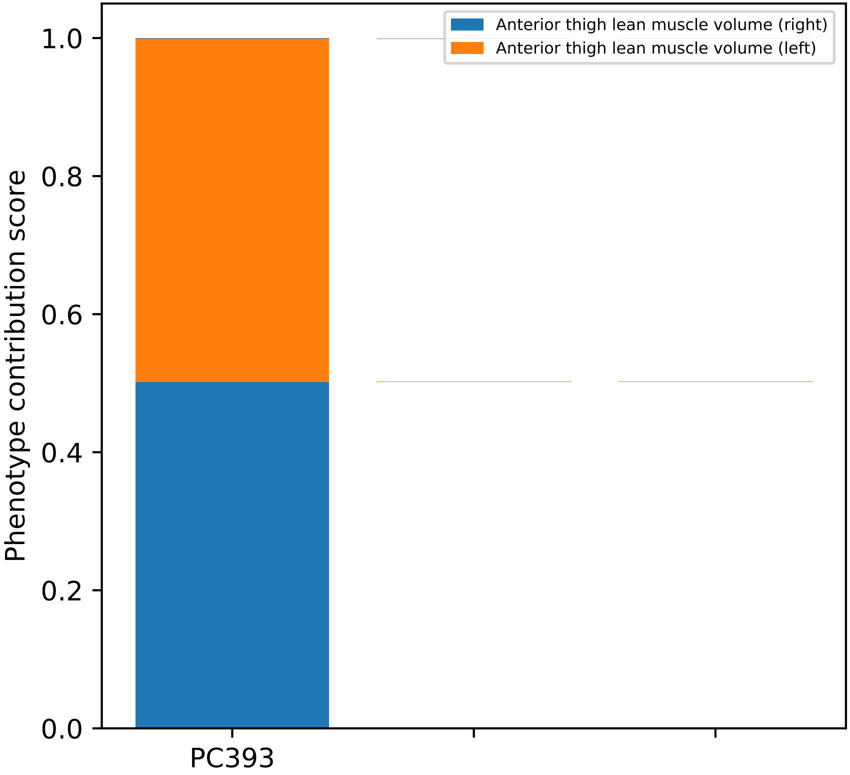
Anterior thigh lean muscle volume (right)
Anterior thigh lean muscle volume (left)
Phenotype contribution score
0.0
0.2
0.4
0.6
0.8
1.0
PC393

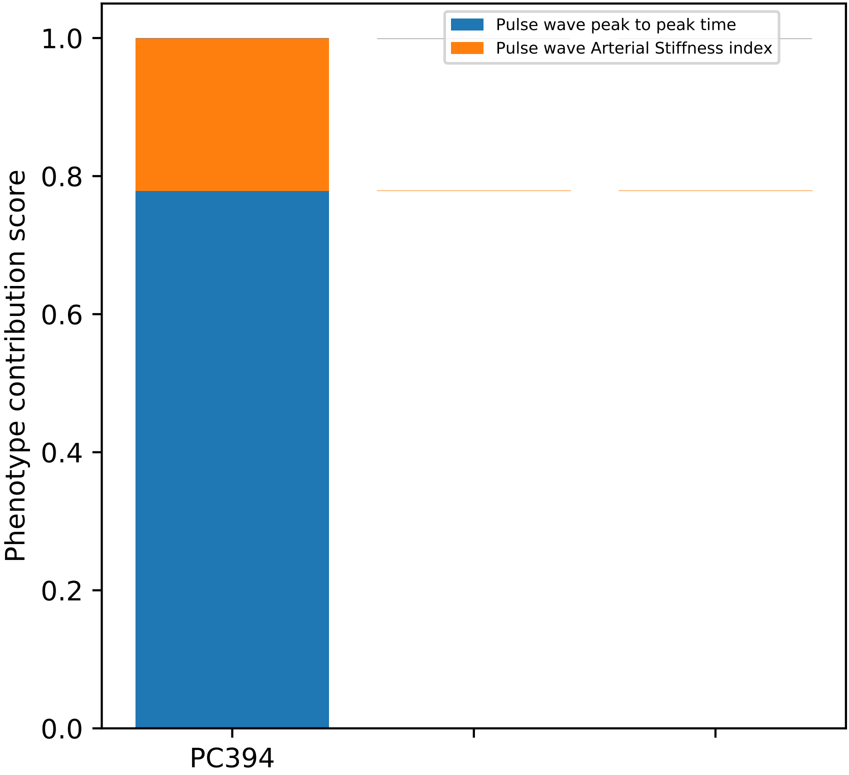
Pulse wave peak to peak time
Pulse wave Arterial Stiffness index
Phenotype contribution score
1.0
0.8
0.6
0.4
0.2
0.0
PC394

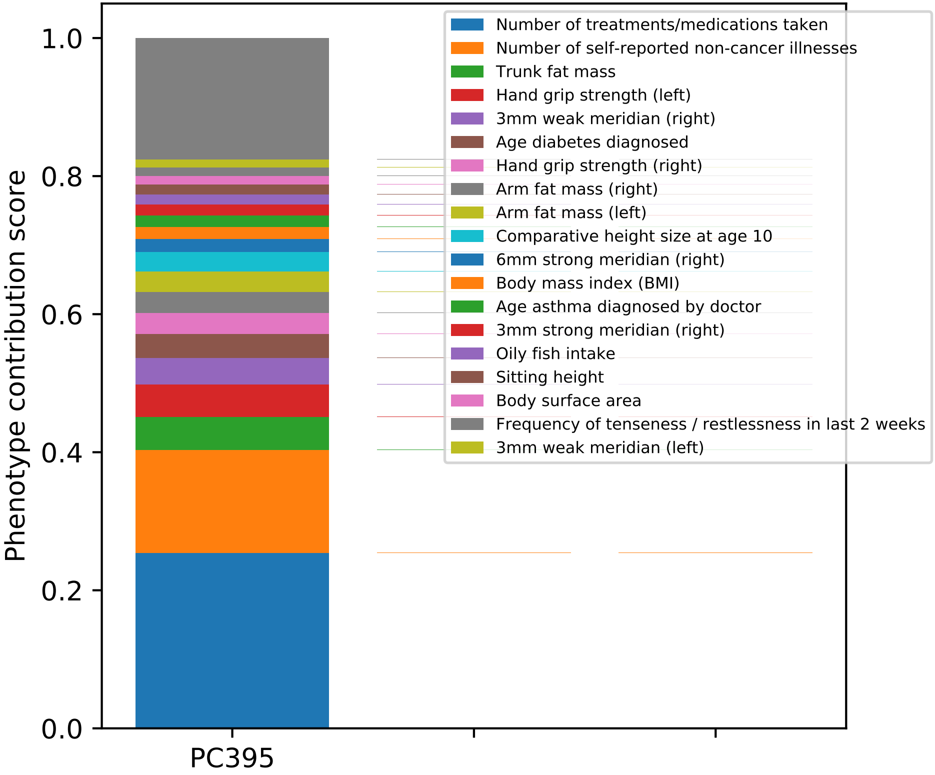
Phenotype contribution score
1.0
0.8
0.6
0.4
0.2
0.0
PC395
Number of treatments/medications taken
Number of self-reported non-cancer illnesses
Trunk fat mass
Hand grip strength (left)
3mm weak meridian (right)
Age diabetes diagnosed
Hand grip strength (right)
Arm fat mass (right)
Arm fat mass (left)
Comparative height size at age 10
6mm strong meridian (right)
Body mass index (BMI)
Age asthma diagnosed by doctor
3mm strong meridian (right)
Oily fish intake
Sitting height
Body surface area
Frequency of tenseness / restlessness in last 2 weeks
3mm weak meridian (left)

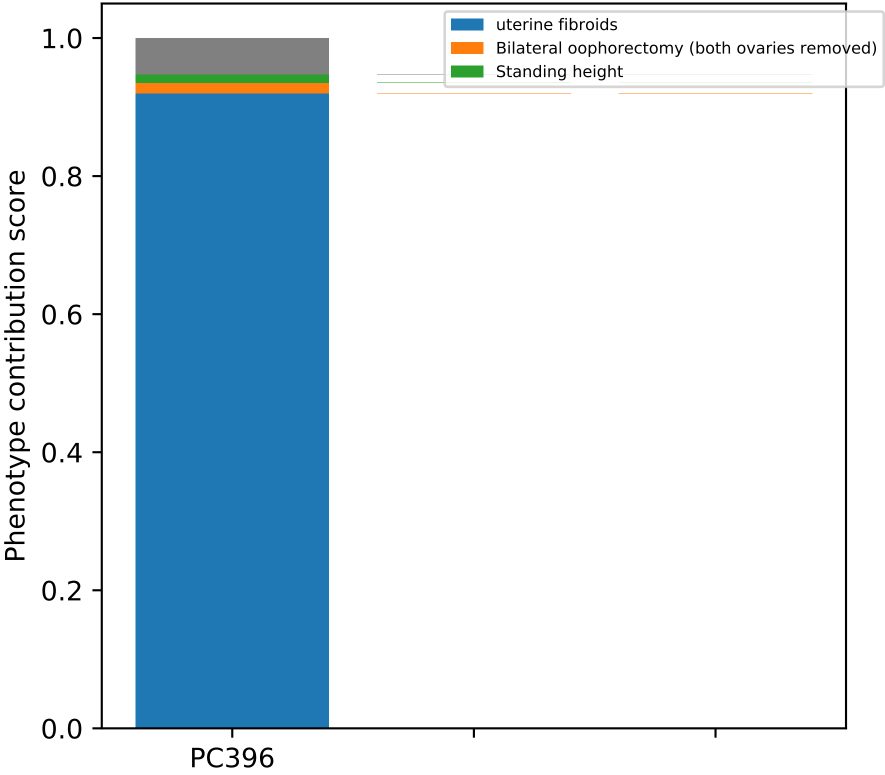
uterine fibroids
Bilateral oophorectomy (both ovaries removed)
Standing height
Phenotype contribution score
1.0
0.8
0.6
0.4
0.2
0.0
PC396

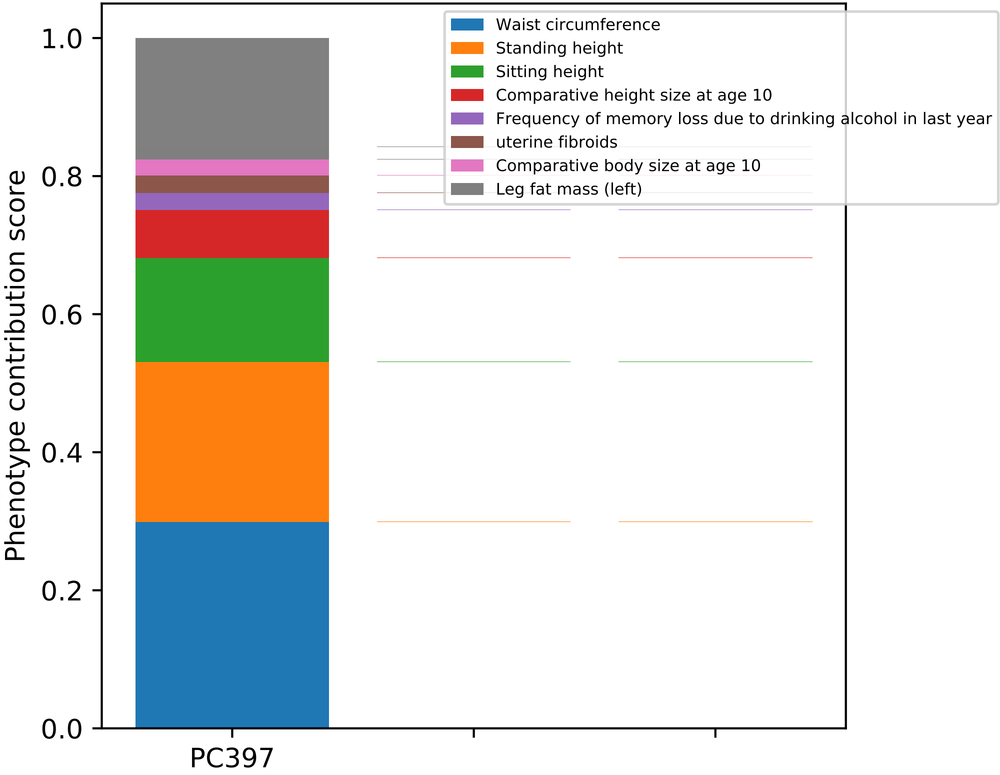

Waist circumference
Standing height
Sitting height
Comparative height size at age 10
Frequency of memory loss due to drinking alcohol in last year
uterine fibroids
Comparative body size at age 10
Leg fat mass (left)
Phenotype contribution score
0.0
0.2
0.4
0.6
0.8
1.0
PC397

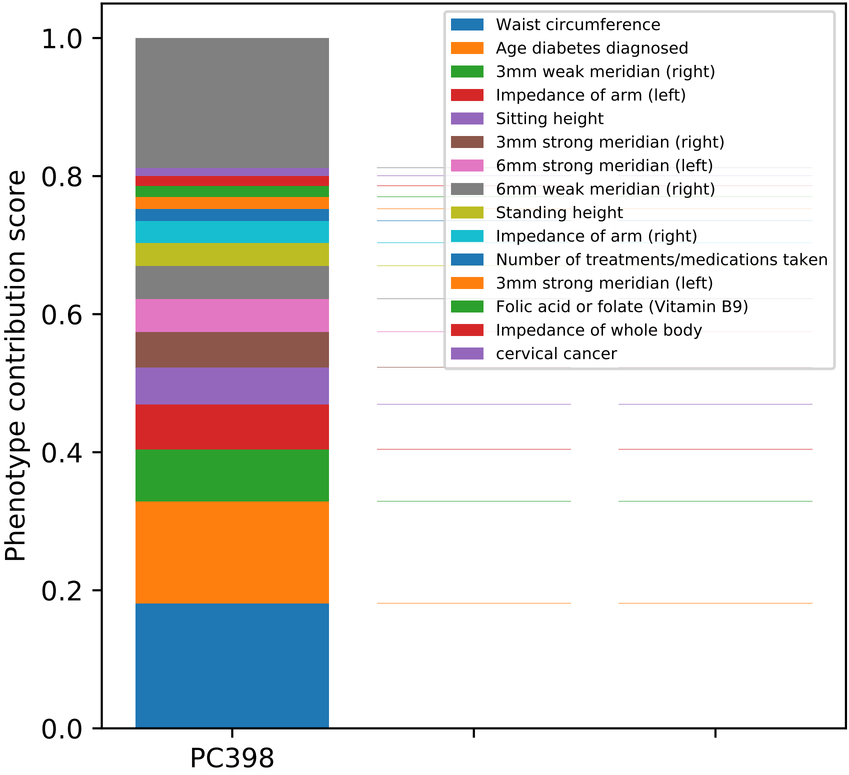

Phenotype contribution score
1.0
0.8
0.6
0.4
0.2
0.0
PC398
Waist circumference
Age diabetes diagnosed
3mm weak meridian (right)
Impedance of arm (left)
Sitting height
3mm strong meridian (right)
6mm strong meridian (left)
6mm weak meridian (right)
Standing height
Impedance of arm (right)
Number of treatments/medications taken
3mm strong meridian (left)
Folic acid or folate (Vitamin B9)
Impedance of whole body
cervical cancer

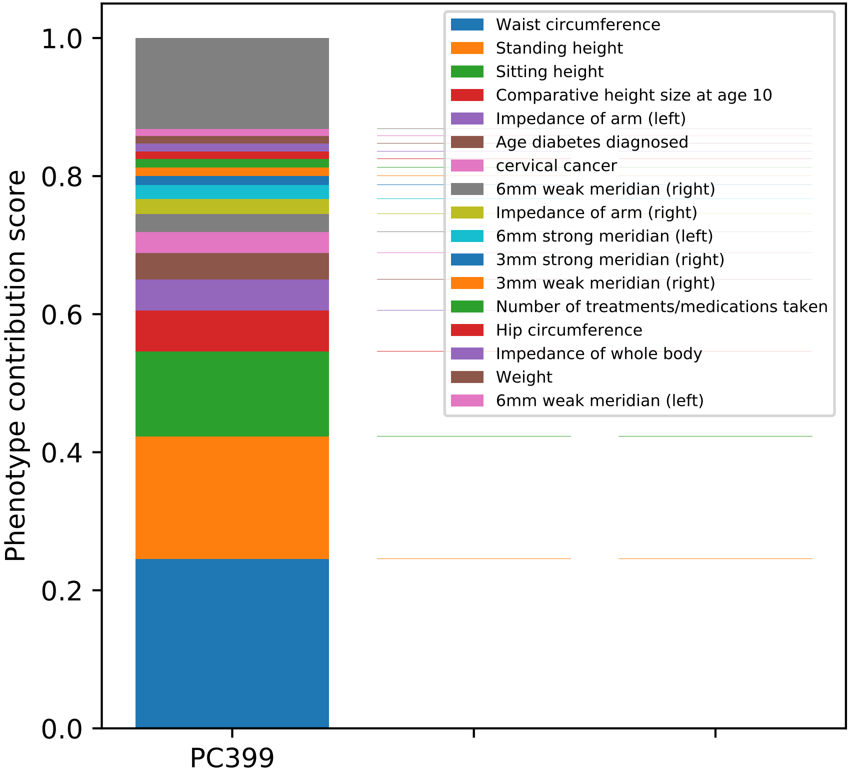
Waist circumference
Standing height
Sitting height
Comparative height size at age 10
Impedance of arm (left)
Age diabetes diagnosed
cervical cancer
6mm weak meridian (right)
Impedance of arm (right)
6mm strong meridian (left)
3mm strong meridian (right)
3mm weak meridian (right)
Number of treatments/medications taken
Hip circumference
Impedance of whole body
Weight
6mm weak meridian (left)
Phenotype contribution score
1.0
0.8
0.6
0.4
0.2
0.0
PC399

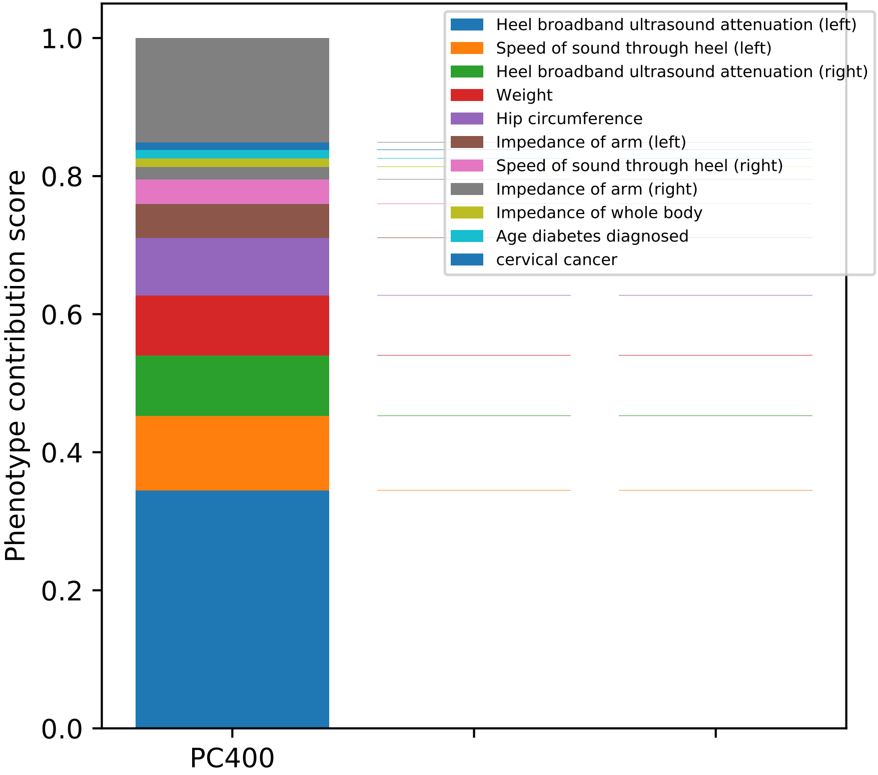
Phenotype contribution score
1.0
0.8
0.6
0.4
0.2
0.0
PC400
Heel broadband ultrasound attenuation (left)
Speed of sound through heel (left)
Heel broadband ultrasound attenuation (right)
Weight
Hip circumference
Impedance of arm (left)
Speed of sound through heel (right)
Impedance of arm (right)
Impedance of whole body
Age diabetes diagnosed
cervical cancer

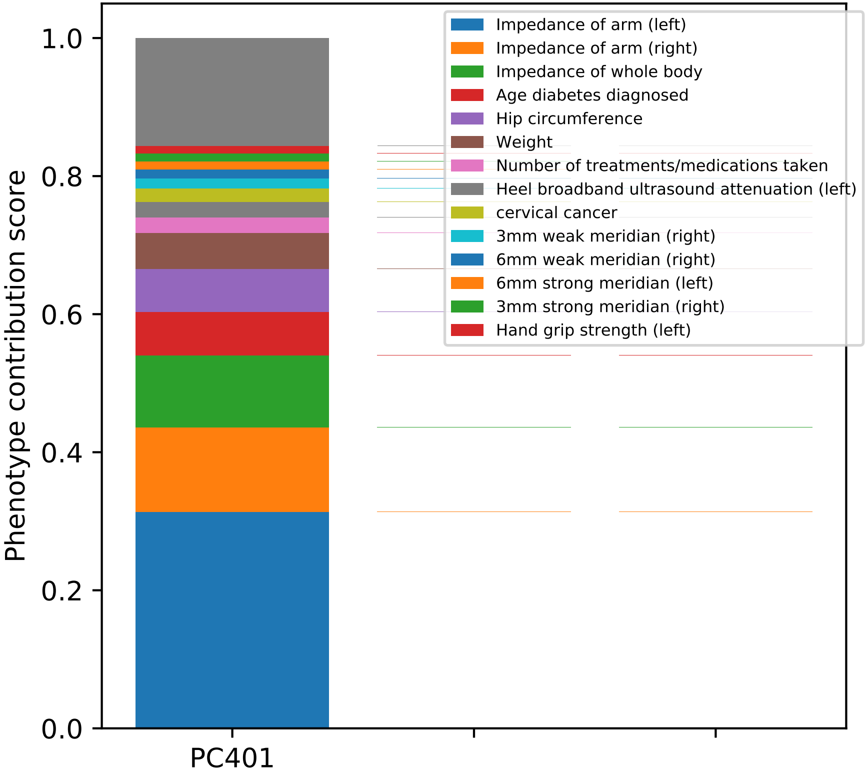
Phenotype contribution score
0.0
0.2
0.4
0.6
0.8
1.0
PC401
Impedance of arm (left)
Impedance of arm (right)
Impedance of whole body
Age diabetes diagnosed
Hip circumference
Weight
Number of treatments/medications taken
Heel broadband ultrasound attenuation (left)
cervical cancer
3mm weak meridian (right)
6mm weak meridian (right)
6mm strong meridian (left)
3mm strong meridian (right)
Hand grip strength (left)

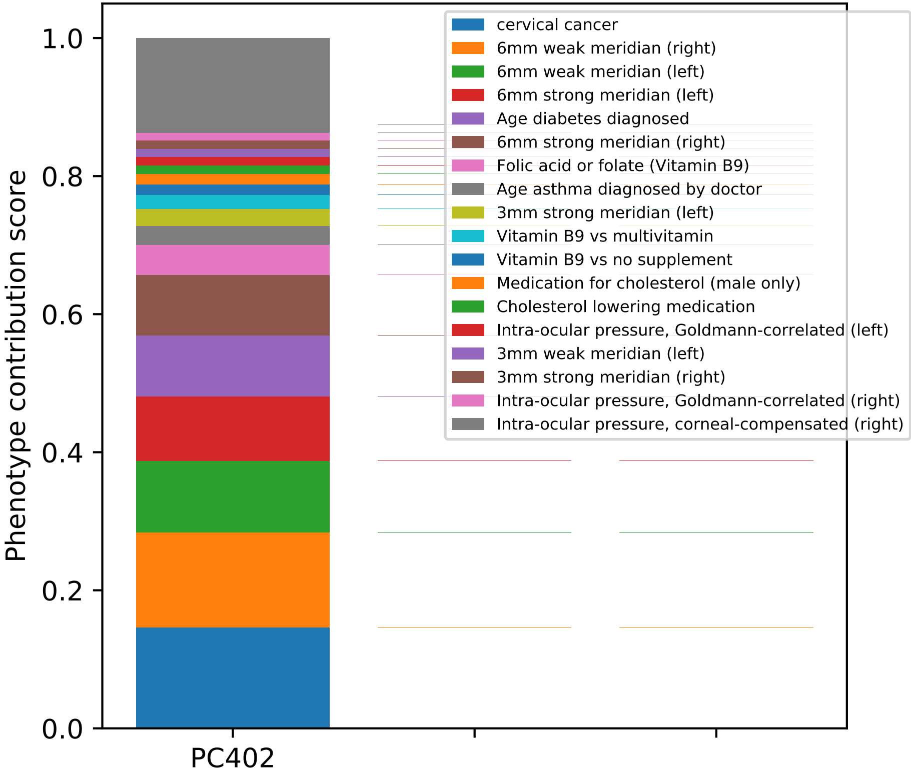

Phenotype contribution score
0.0
0.2
0.4
0.6
0.8
1.0
PC402
cervical cancer
6mm weak meridian (right)
6mm weak meridian (left)
6mm strong meridian (left)
Age diabetes diagnosed
6mm strong meridian (right)
Folic acid or folate (Vitamin B9)
Age asthma diagnosed by doctor
3mm strong meridian (left)
Vitamin B9 vs multivitamin
Vitamin B9 vs no supplement
Medication for cholesterol (male only)
Cholesterol lowering medication
Intra-ocular pressure, Goldmann-correlated (left)
3mm weak meridian (left)
3mm strong meridian (right)
Intra-ocular pressure, Goldmann-correlated (right)
Intra-ocular pressure, corneal-compensated (right)

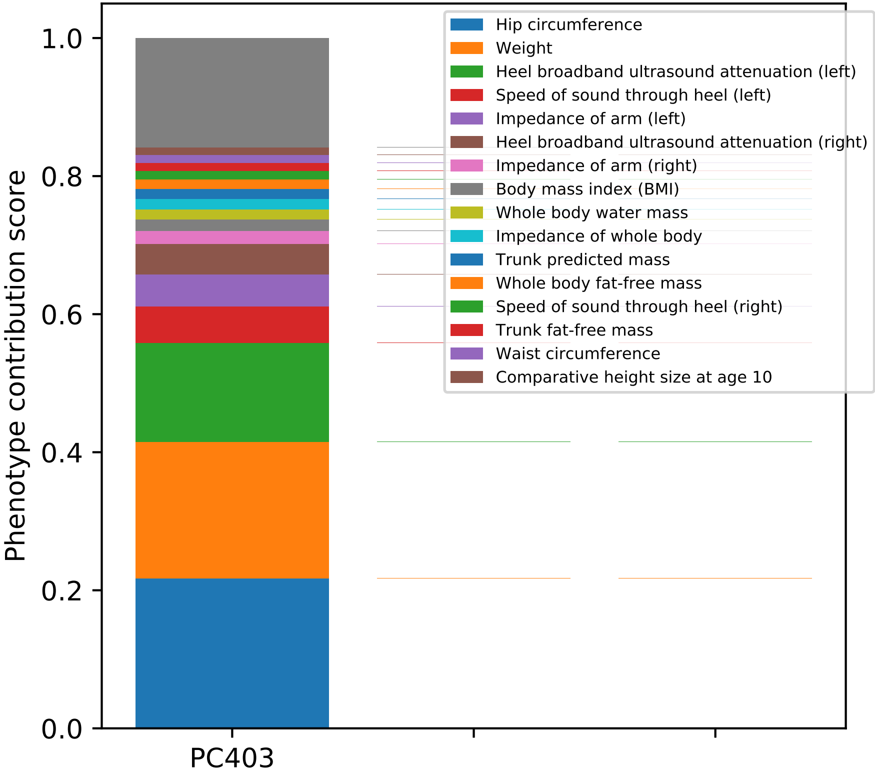

Phenotype contribution score
0.0
0.2
0.4
0.6
0.8
1.0
PC403
Hip circumference
Weight
Heel broadband ultrasound attenuation (left)
Speed of sound through heel (left)
Impedance of arm (left)
Heel broadband ultrasound attenuation (right)
Impedance of arm (right)
Body mass index (BMI)
Whole body water mass
Impedance of whole body
Trunk predicted mass
Whole body fat-free mass
Speed of sound through heel (right)
Trunk fat-free mass
Waist circumference
Comparative height size at age 10

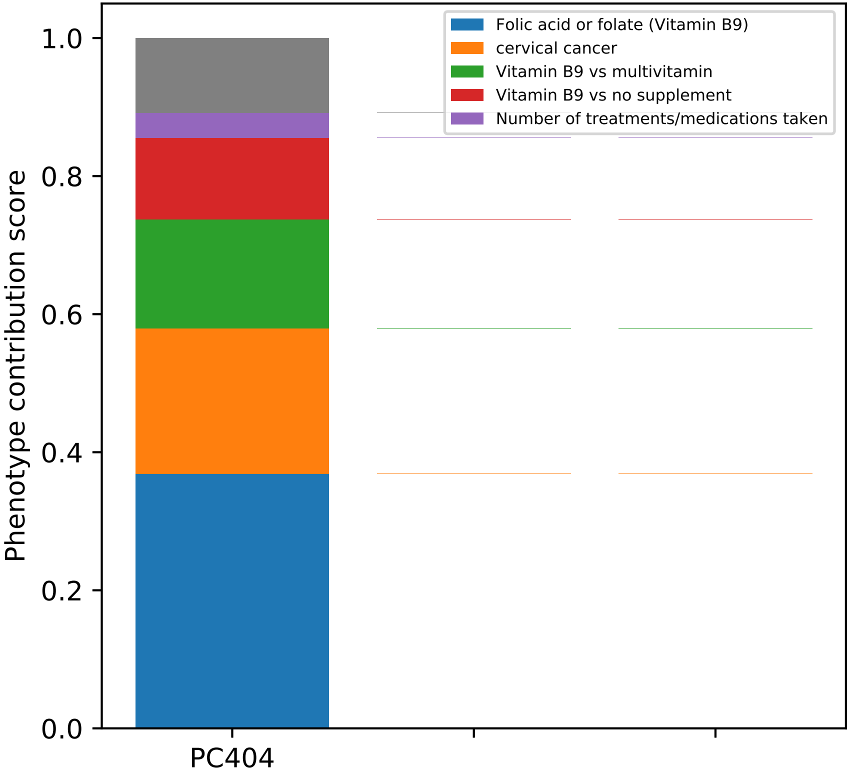
Folic acid or folate (Vitamin B9)
cervical cancer
Vitamin B9 vs multivitamin
Vitamin B9 vs no supplement
Number of treatments/medications taken
Phenotype contribution score
1.0
0.8
0.6
0.4
0.2
0.0
PC404

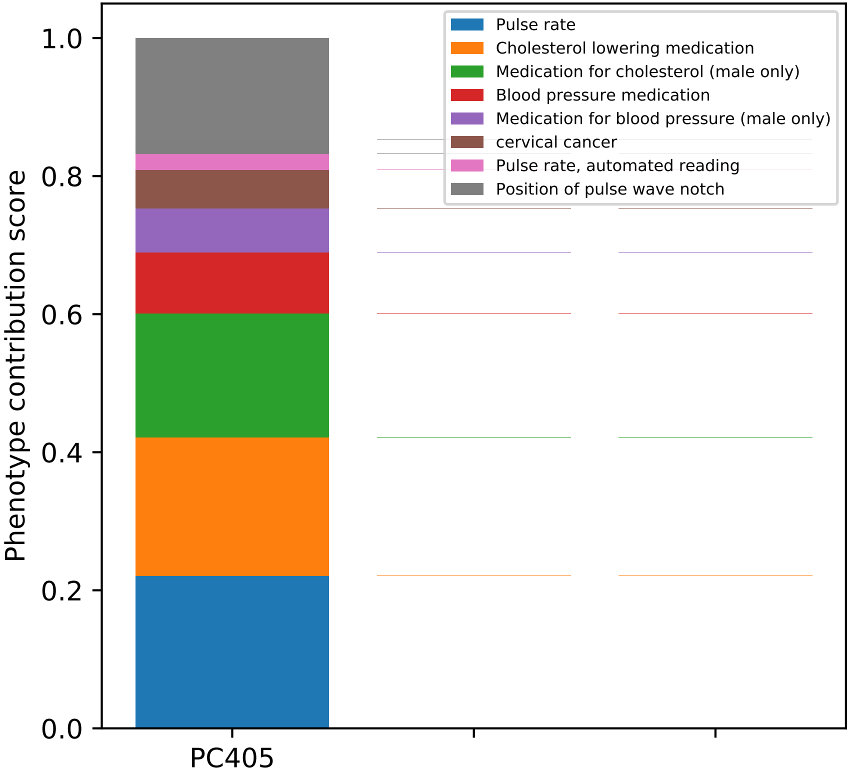

1.0
0.8
0.6
0.4
0.2
0.0
Phenotype contribution score
PC405
Pulse rate
Cholesterol lowering medication
Medication for cholesterol (male only)
Blood pressure medication
Medication for blood pressure (male only)
cervical cancer
Pulse rate, automated reading
Position of pulse wave notch

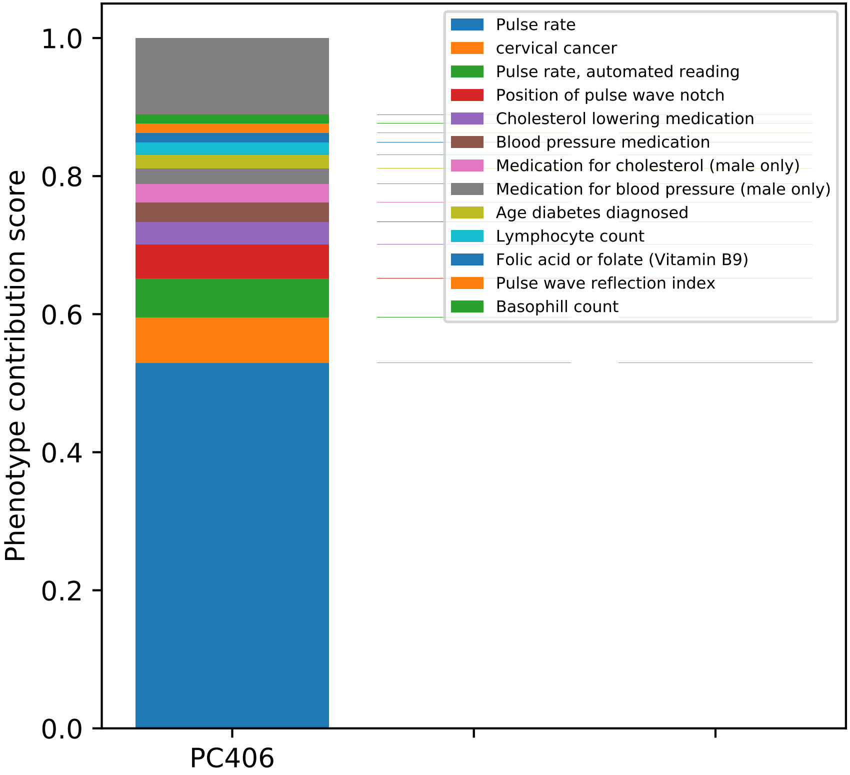
Phenotype contribution score
0.0
0.2
0.4
0.6
0.8
1.0
PC406
Pulse rate
cervical cancer
Pulse rate, automated reading
Position of pulse wave notch
Cholesterol lowering medication
Blood pressure medication
Medication for cholesterol (male only)
Medication for blood pressure (male only)
Age diabetes diagnosed
Lymphocyte count
Folic acid or folate (Vitamin B9)
Pulse wave reflection index
Basophill count

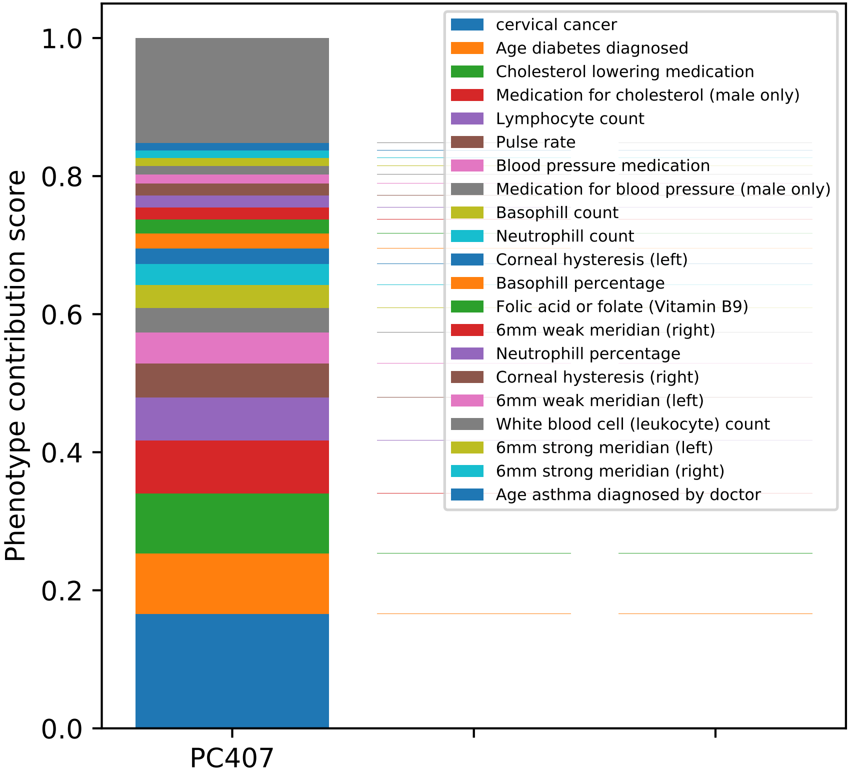

Phenotype contribution score
0.0
0.2
0.4
0.6
0.8
1.0
PC407
cervical cancer
Age diabetes diagnosed
Cholesterol lowering medication
Medication for cholesterol (male only)
Lymphocyte count
Pulse rate
Blood pressure medication
Medication for blood pressure (male only)
Basophill count
Neutrophill count
Corneal hysteresis (left)
Basophill percentage
Folic acid or folate (Vitamin B9)
6mm weak meridian (right)
Neutrophill percentage
Corneal hysteresis (right)
6mm weak meridian (left)
White blood cell (leukocyte) count
6mm strong meridian (left)
6mm strong meridian (right)
Age asthma diagnosed by doctor

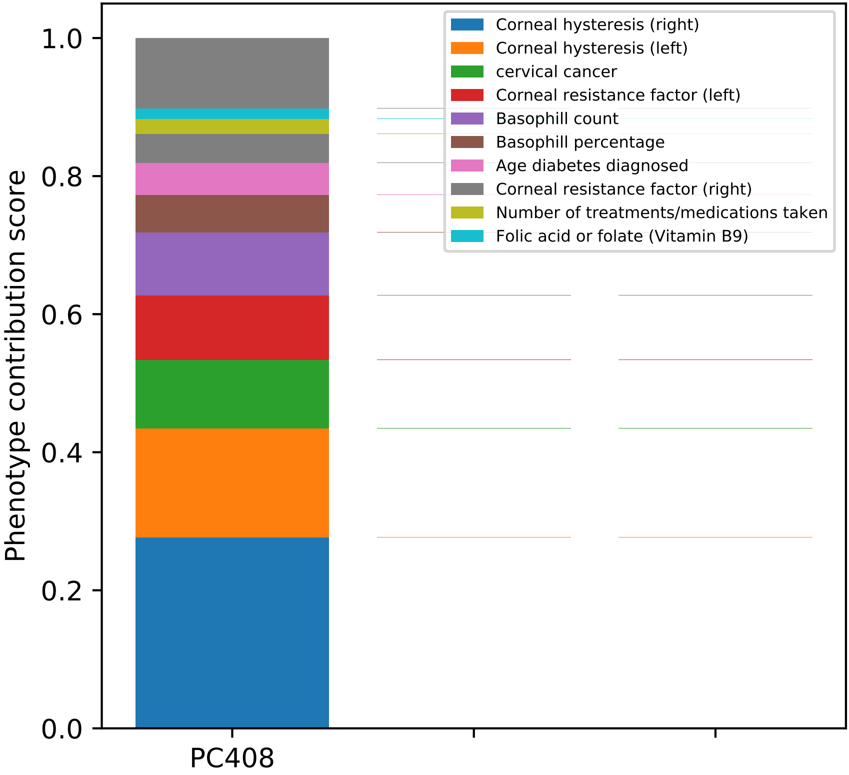
Phenotype contribution score
0.0
0.2
0.4
0.6
0.8
1.0
PC408
Corneal hysteresis (right)
Corneal hysteresis (left)
cervical cancer
Corneal resistance factor (left)
Basophill count
Basophill percentage
Age diabetes diagnosed
Corneal resistance factor (right)
Number of treatments/medications taken
Folic acid or folate (Vitamin B9)

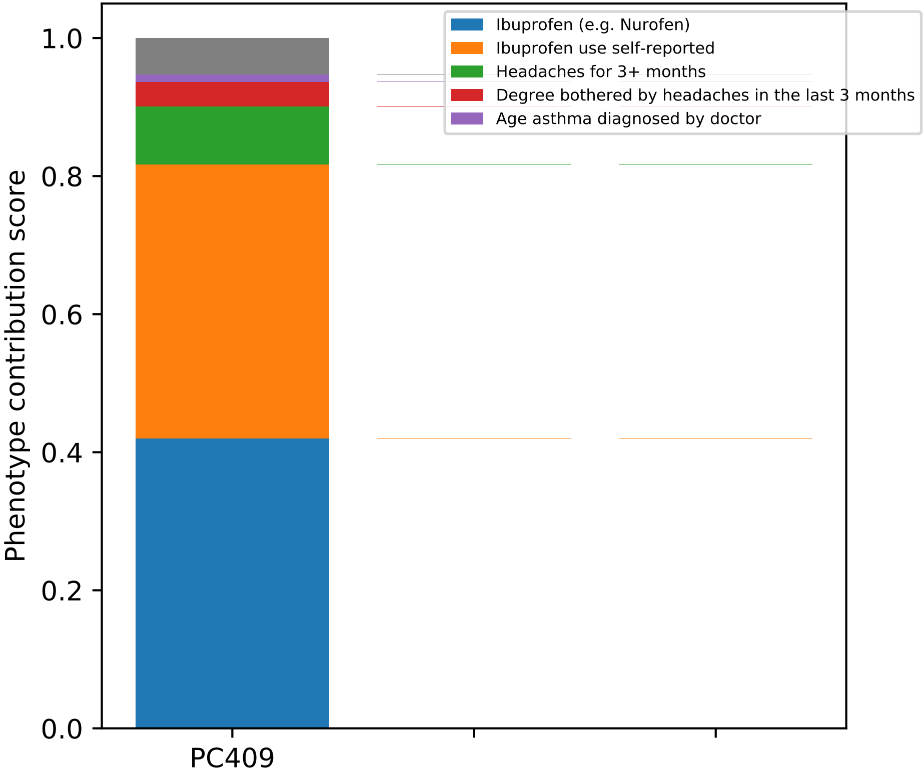

Ibuprofen (e.g. Nurofen)
Ibuprofen use self-reported
Headaches for 3+ months
Degree bothered by headaches in the last 3 months
Age asthma diagnosed by doctor
Phenotype contribution score
1.0
0.8
0.6
0.4
0.2
0.0
PC409

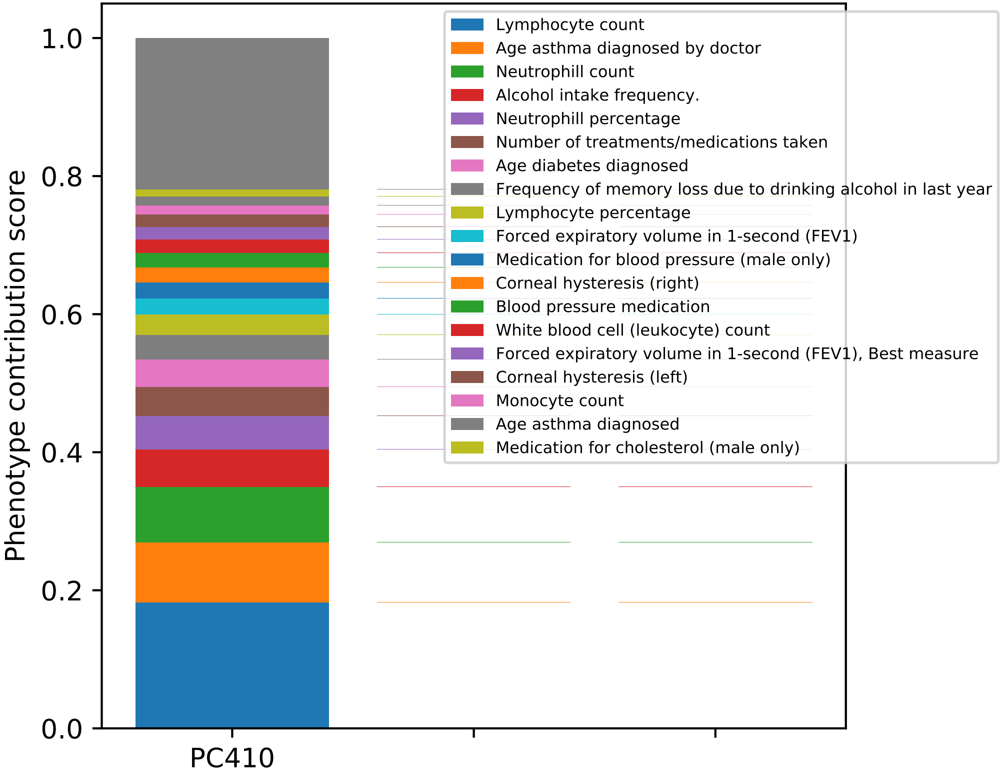
Phenotype contribution score
0.0
0.2
0.4
0.6
0.8
1.0
PC410
Lymphocyte count
Age asthma diagnosed by doctor
Neutrophill count
Alcohol intake frequency.
Neutrophill percentage
Number of treatments/medications taken
Age diabetes diagnosed
Frequency of memory loss due to drinking alcohol in last year
Lymphocyte percentage
Forced expiratory volume in 1-second (FEV1)
Medication for blood pressure (male only)
Corneal hysteresis (right)
Blood pressure medication
White blood cell (leukocyte) count
Forced expiratory volume in 1-second (FEV1), Best measure
Corneal hysteresis (left)
Monocyte count
Age asthma diagnosed
Medication for cholesterol (male only)

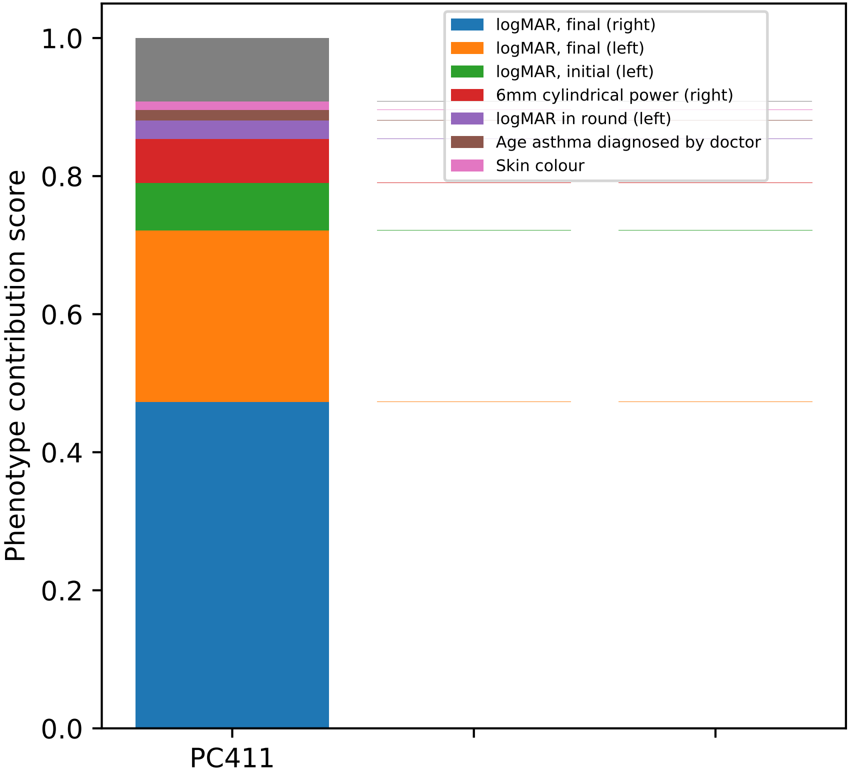

logMAR, final (right)
logMAR, final (left)
logMAR, initial (left)
6mm cylindrical power (right)
logMAR in round (left)
Age asthma diagnosed by doctor
Skin colour
Phenotype contribution score
1.0
0.8
0.6
0.4
0.2
0.0
PC411

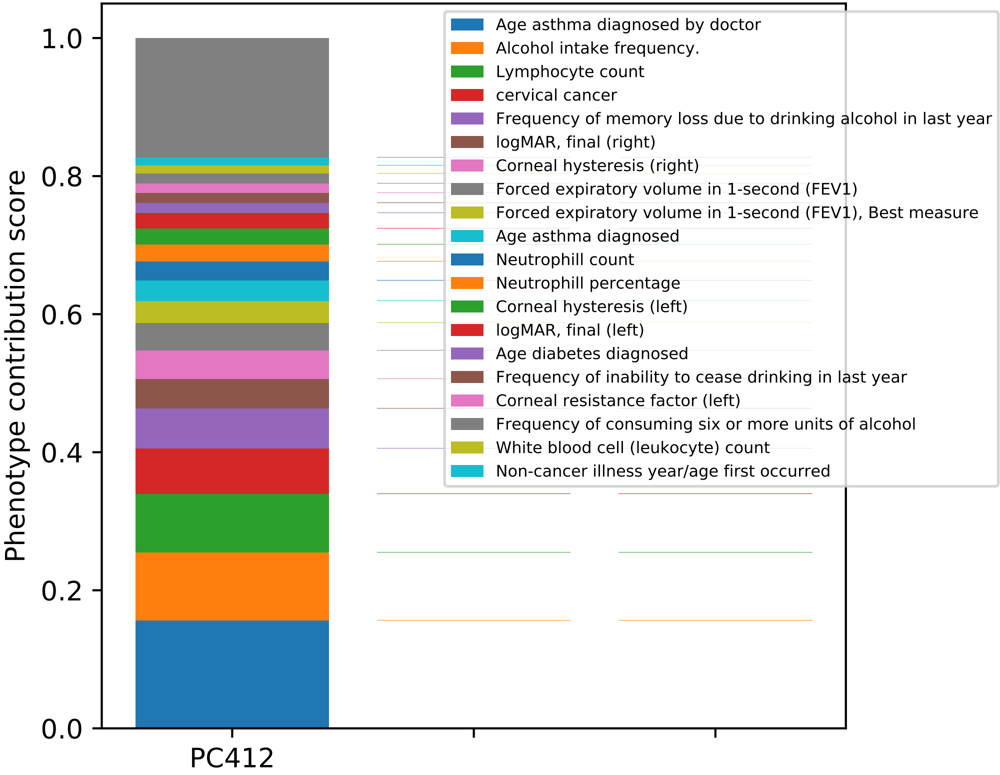

Phenotype contribution score
1.0
0.8
0.6
0.4
0.2
0.0
PC412
Age asthma diagnosed by doctor
Alcohol intake frequency.
Lymphocyte count
cervical cancer
Frequency of memory loss due to drinking alcohol in last year
logMAR, final (right)
Corneal hysteresis (right)
Forced expiratory volume in 1-second (FEV1)
Forced expiratory volume in 1-second (FEV1), Best measure
Age asthma diagnosed
Neutrophill count
Neutrophill percentage
Corneal hysteresis (left)
logMAR, final (left)
Age diabetes diagnosed
Frequency of inability to cease drinking in last year
Corneal resistance factor (left)
Frequency of consuming six or more units of alcohol
White blood cell (leukocyte) count
Non-cancer illness year/age first occurred

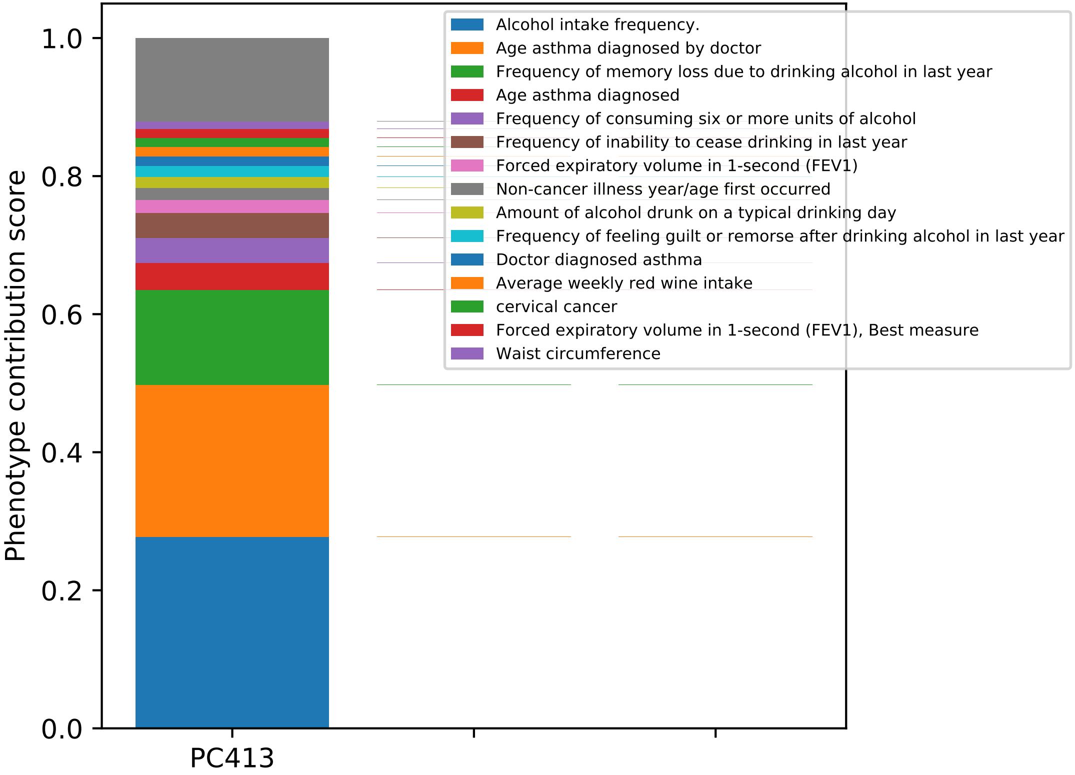
Phenotype contribution score
1.0
0.8
0.6
0.4
0.2
0.0
PC413
Alcohol intake frequency.
Age asthma diagnosed by doctor
Frequency of memory loss due to drinking alcohol in last year
Age asthma diagnosed
Frequency of consuming six or more units of alcohol
Frequency of inability to cease drinking in last year
Forced expiratory volume in 1-second (FEV1)
Non-cancer illness year/age first occurred
Amount of alcohol drunk on a typical drinking day
Frequency of feeling guilt or remorse after drinking alcohol in last year
Doctor diagnosed asthma
Average weekly red wine intake
cervical cancer
Forced expiratory volume in 1-second (FEV1), Best measure
Waist circumference

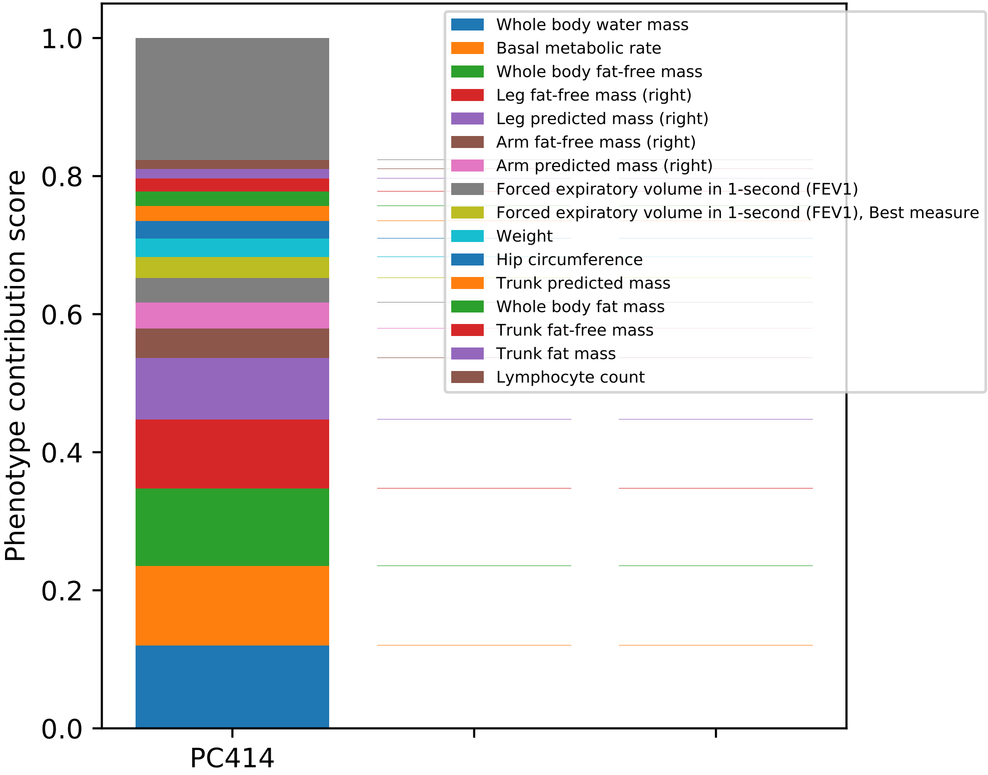

Phenotype contribution score
1.0
0.8
0.6
0.4
0.2
0.0
PC414
Whole body water mass
Basal metabolic rate
Whole body fat-free mass
Leg fat-free mass (right)
Leg predicted mass (right)
Arm fat-free mass (right)
Arm predicted mass (right)
Forced expiratory volume in 1-second (FEV1)
Forced expiratory volume in 1-second (FEV1), Best measure
Weight
Hip circumference
Trunk predicted mass
Whole body fat mass
Trunk fat-free mass
Trunk fat mass
Lymphocyte count

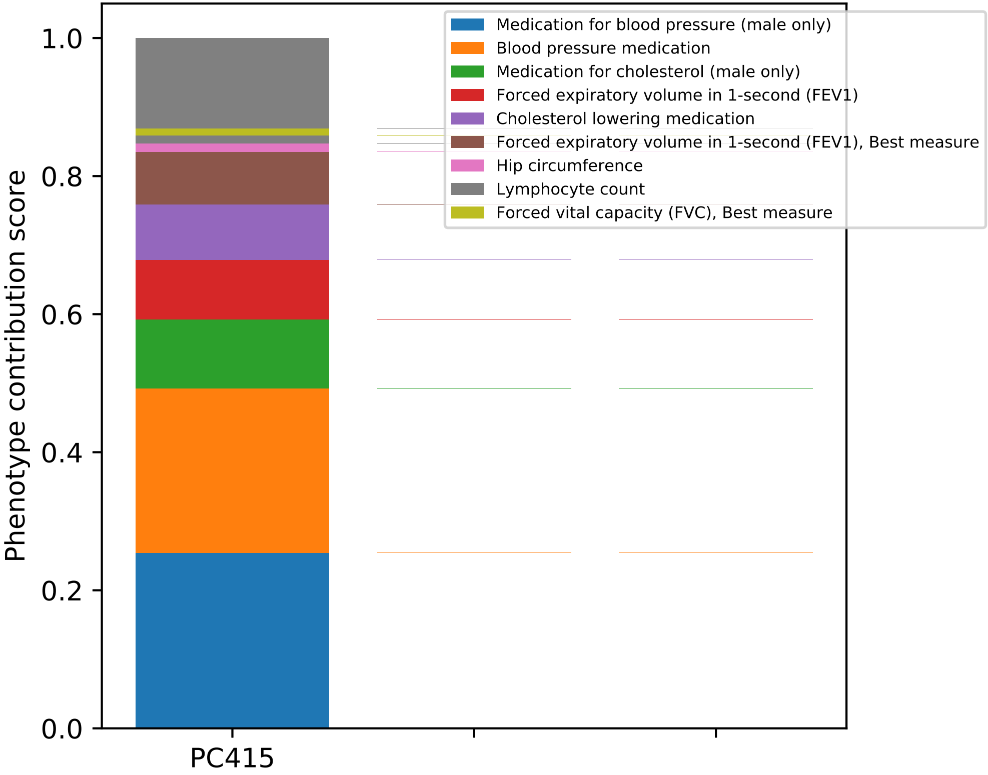
Medication for blood pressure (male only)
Blood pressure medication
Medication for cholesterol (male only)
Forced expiratory volume in 1-second (FEV1)
Cholesterol lowering medication
Forced expiratory volume in 1-second (FEV1), Best measure
Hip circumference
Lymphocyte count
Forced vital capacity (FVC), Best measure
Phenotype contribution score
1.0
0.8
0.6
0.4
0.2
0.0
PC415

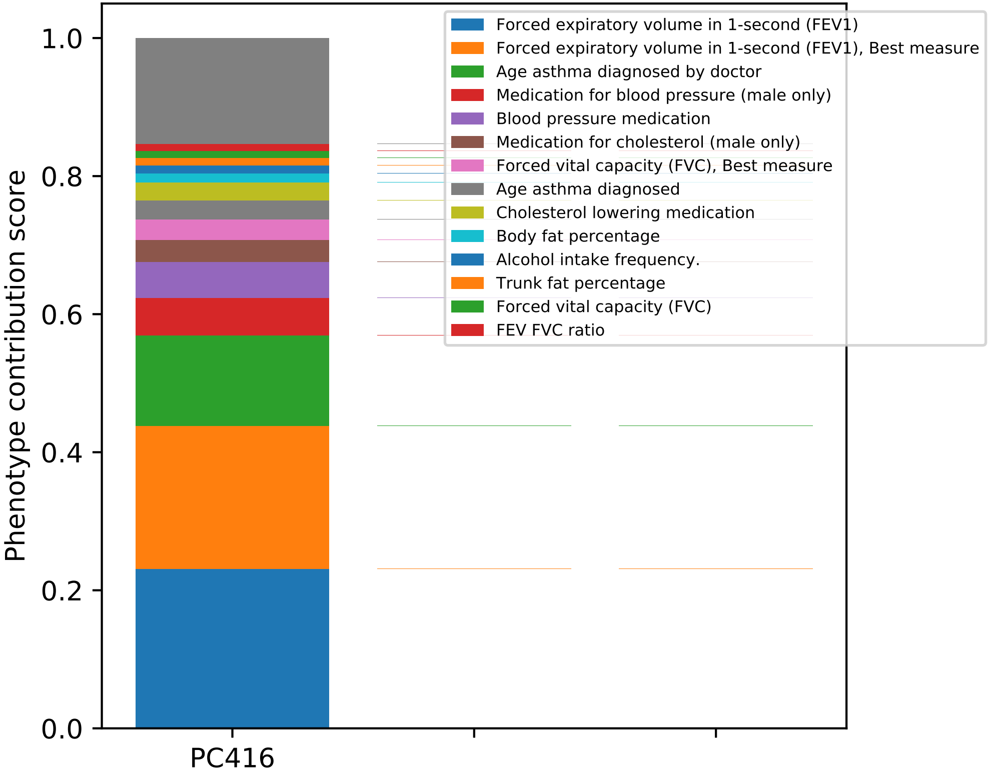

Phenotype contribution score
1.0
0.8
0.6
0.4
0.2
0.0
PC416
Forced expiratory volume in 1-second (FEV1)
Forced expiratory volume in 1-second (FEV1), Best measure
Age asthma diagnosed by doctor
Medication for blood pressure (male only)
Blood pressure medication
Medication for cholesterol (male only)
Forced vital capacity (FVC), Best measure
Age asthma diagnosed
Cholesterol lowering medication
Body fat percentage
Alcohol intake frequency.
Trunk fat percentage
Forced vital capacity (FVC)
FEV FVC ratio

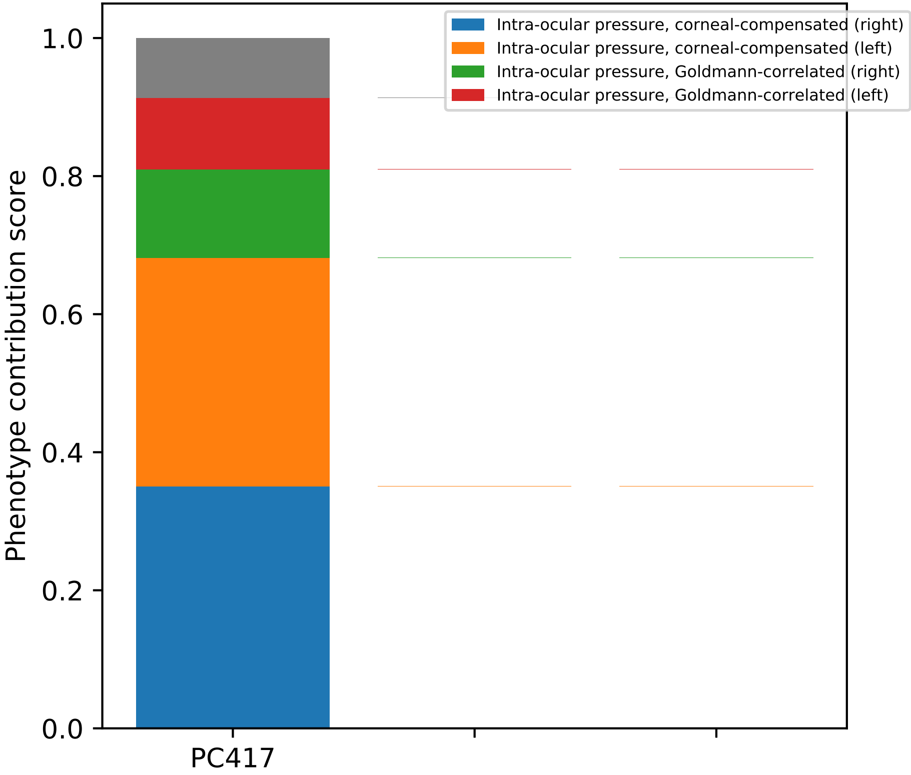

Intra-ocular pressure, corneal-compensated (right)
Intra-ocular pressure, corneal-compensated (left)
Intra-ocular pressure, Goldmann-correlated (right)
Intra-ocular pressure, Goldmann-correlated (left)
Phenotype contribution score
0.0
0.2
0.4
0.6
0.8
1.0
PC417

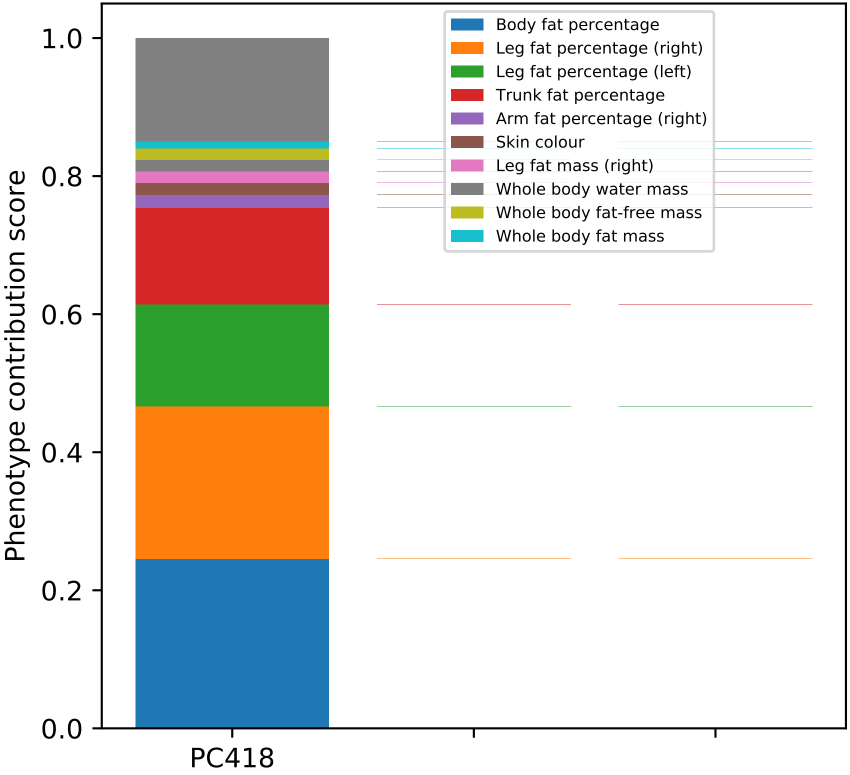

Phenotype contribution score
1.0
0.8
0.6
0.4
0.2
0.0
PC418
Body fat percentage
Leg fat percentage (right)
Leg fat percentage (left)
Trunk fat percentage
Arm fat percentage (right)
Skin colour
Leg fat mass (right)
Whole body water mass
Whole body fat-free mass
Whole body fat mass

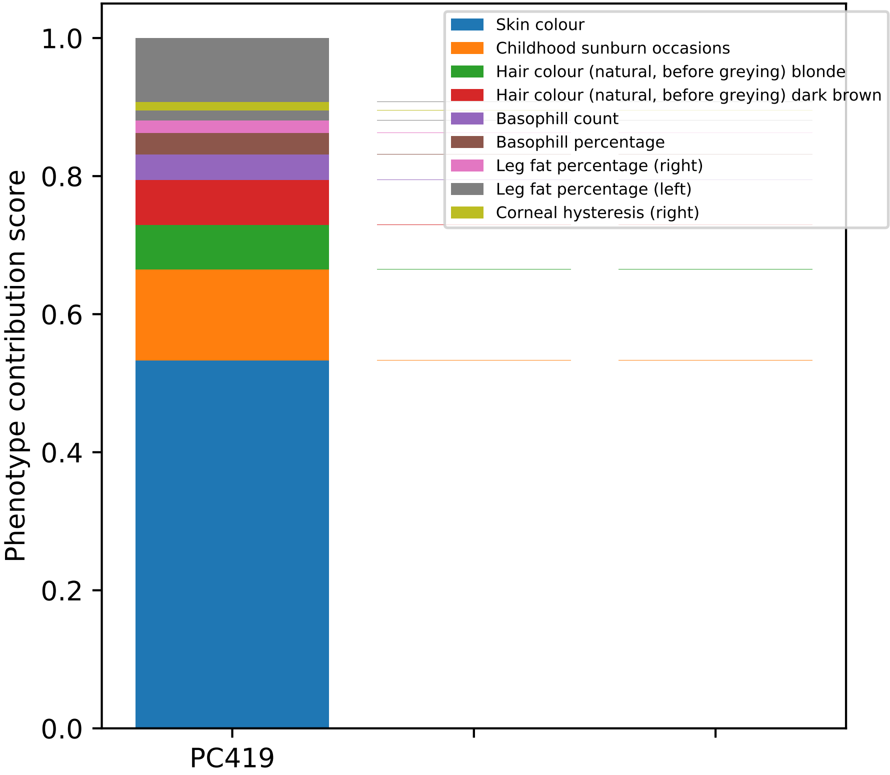

Skin colour
Childhood sunburn occasions
Hair colour (natural, before greying) blonde
Hair colour (natural, before greying) dark brown
Basophill count
Basophill percentage
Leg fat percentage (right)
Leg fat percentage (left)
Corneal hysteresis (right)
Phenotype contribution score
0.0
0.2
0.4
0.6
0.8
1.0
PC419

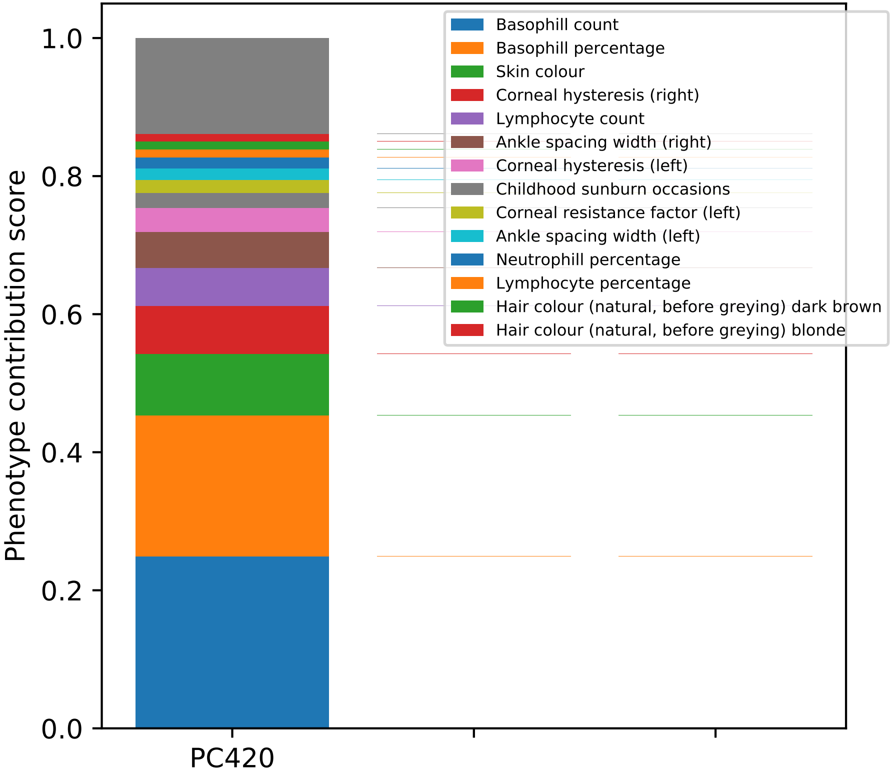

Phenotype contribution score
1.0
0.8
0.6
0.4
0.2
0.0
PC420
Basophill count
Basophill percentage
Skin colour
Corneal hysteresis (right)
Lymphocyte count
Ankle spacing width (right)
Corneal hysteresis (left)
Childhood sunburn occasions
Corneal resistance factor (left)
Ankle spacing width (left)
Neutrophill percentage
Lymphocyte percentage
Hair colour (natural, before greying) dark brown
Hair colour (natural, before greying) blonde

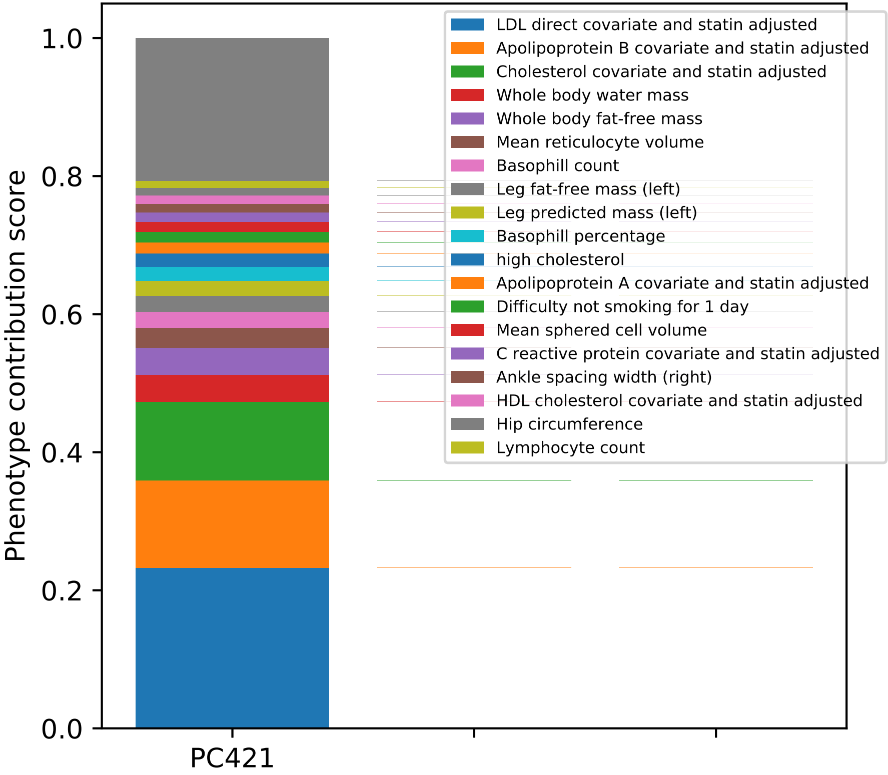

Phenotype contribution score
1.0
0.8
0.6
0.4
0.2
0.0
PC421
LDL direct covariate and statin adjusted
Apolipoprotein B covariate and statin adjusted
Cholesterol covariate and statin adjusted
Whole body water mass
Whole body fat-free mass
Mean reticulocyte volume
Basophill count
Leg fat-free mass (left)
Leg predicted mass (left)
Basophill percentage
high cholesterol
Apolipoprotein A covariate and statin adjusted
Difficulty not smoking for 1 day
Mean sphered cell volume
C reactive protein covariate and statin adjusted
Ankle spacing width (right)
HDL cholesterol covariate and statin adjusted
Hip circumference
Lymphocyte count

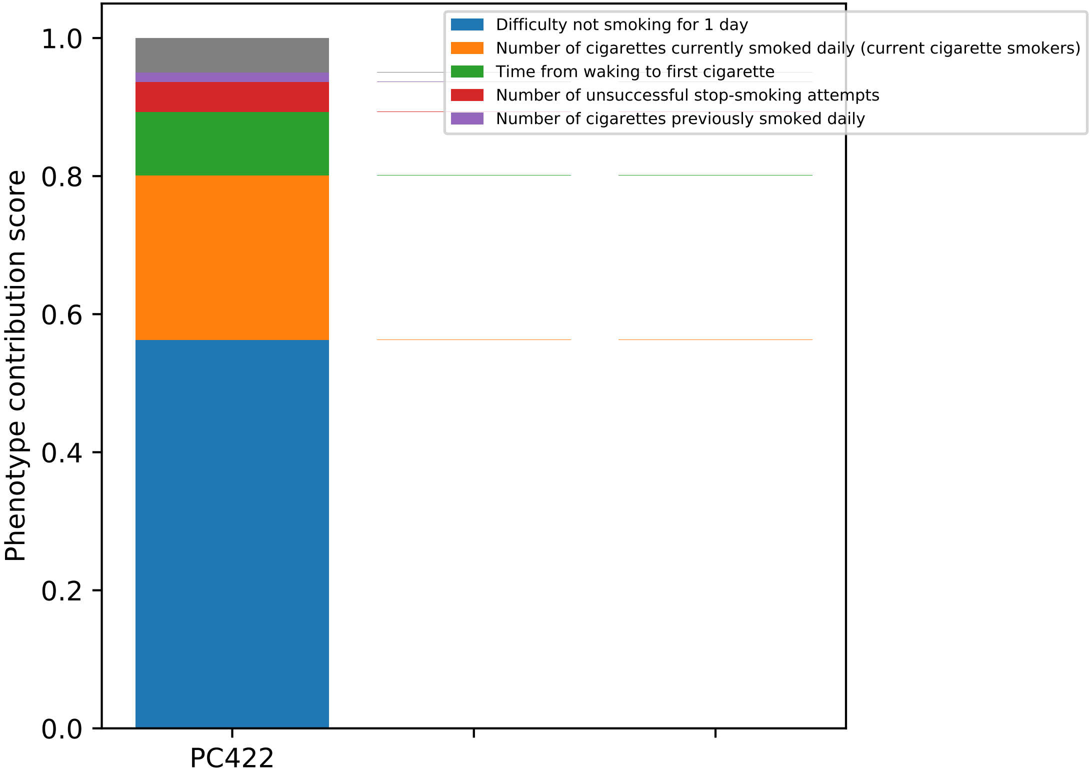
Difficulty not smoking for 1 day
Number of cigarettes currently smoked daily (current cigarette smokers)
Time from waking to first cigarette
Number of unsuccessful stop-smoking attempts
Number of cigarettes previously smoked daily
Phenotype contribution score
1.0
0.8
0.6
0.4
0.2
0.0
PC422

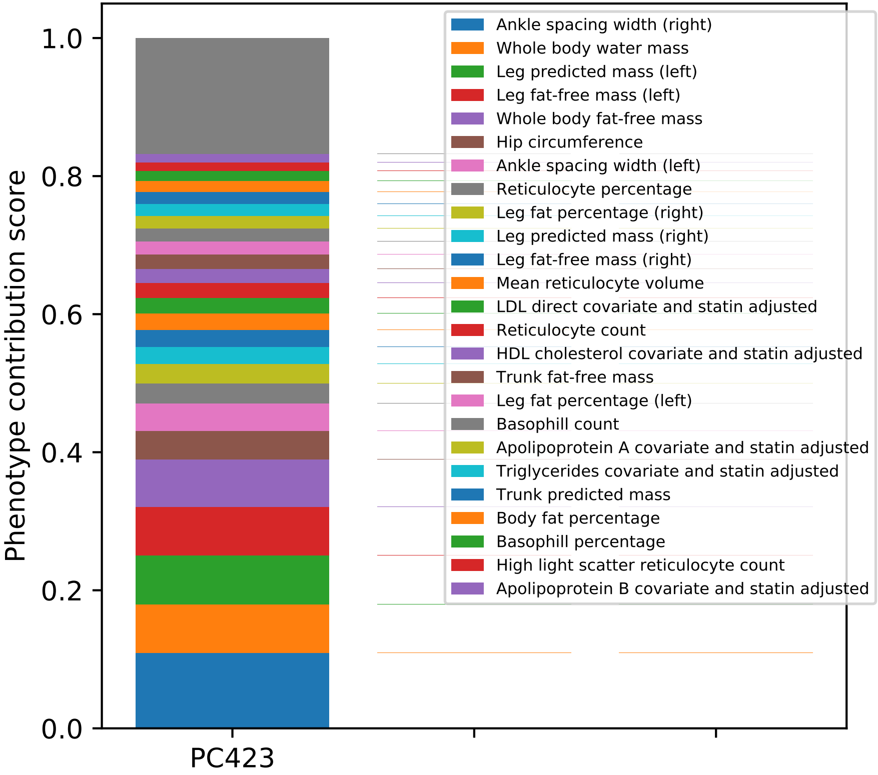

Phenotype contribution score
0.0
0.2
0.4
0.6
0.8
1.0
PC423
Ankle spacing width (right)
Whole body water mass
Leg predicted mass (left)
Leg fat-free mass (left)
Whole body fat-free mass
Hip circumference
Ankle spacing width (left)
Reticulocyte percentage
Leg fat percentage (right)
Leg predicted mass (right)
Leg fat-free mass (right)
Mean reticulocyte volume
LDL direct covariate and statin adjusted
Reticulocyte count
HDL cholesterol covariate and statin adjusted
Trunk fat-free mass
Leg fat percentage (left)
Basophill count
Apolipoprotein A covariate and statin adjusted
Triglycerides covariate and statin adjusted
Trunk predicted mass
Body fat percentage
Basophill percentage
High light scatter reticulocyte count
Apolipoprotein B covariate and statin adjusted

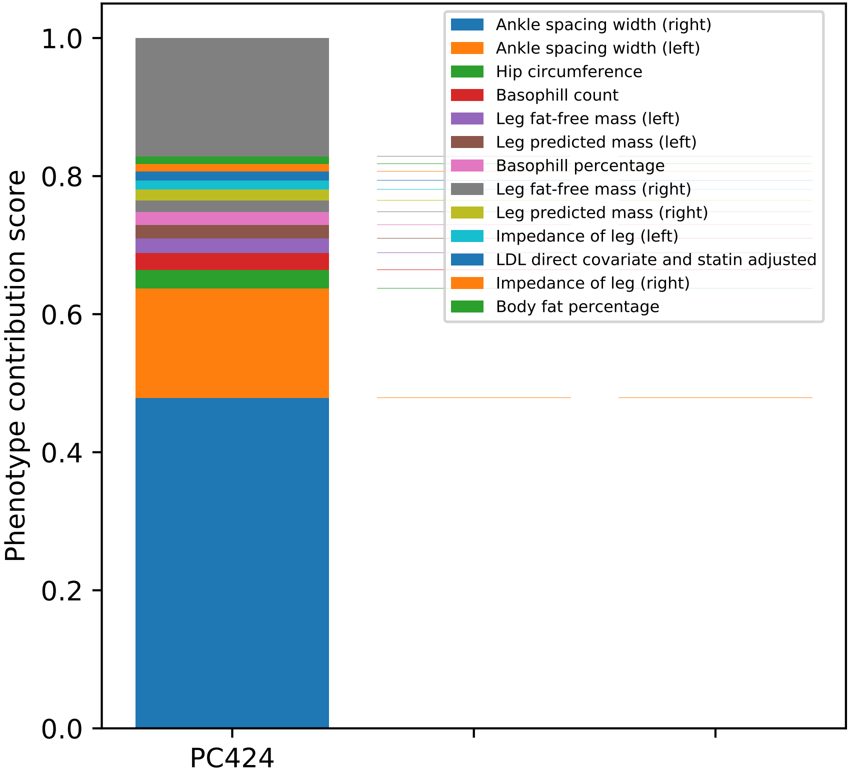

Phenotype contribution score
1.0
0.8
0.6
0.4
0.2
0.0
PC424
Ankle spacing width (right)
Ankle spacing width (left)
Hip circumference
Basophill count
Leg fat-free mass (left)
Leg predicted mass (left)
Basophill percentage
Leg fat-free mass (right)
Leg predicted mass (right)
Impedance of leg (left)
LDL direct covariate and statin adjusted
Impedance of leg (right)
Body fat percentage

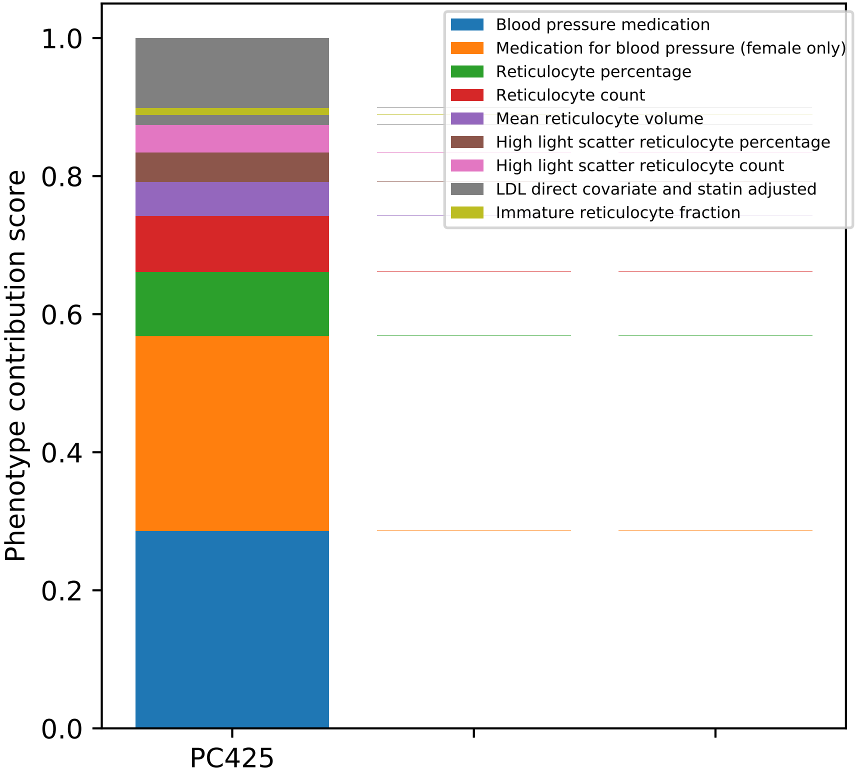
Phenotype contribution score
1.0
0.8
0.6
0.4
0.2
0.0
PC425
Blood pressure medication
Medication for blood pressure (female only)
Reticulocyte percentage
Reticulocyte count
Mean reticulocyte volume
High light scatter reticulocyte percentage
High light scatter reticulocyte count
LDL direct covariate and statin adjusted
Immature reticulocyte fraction

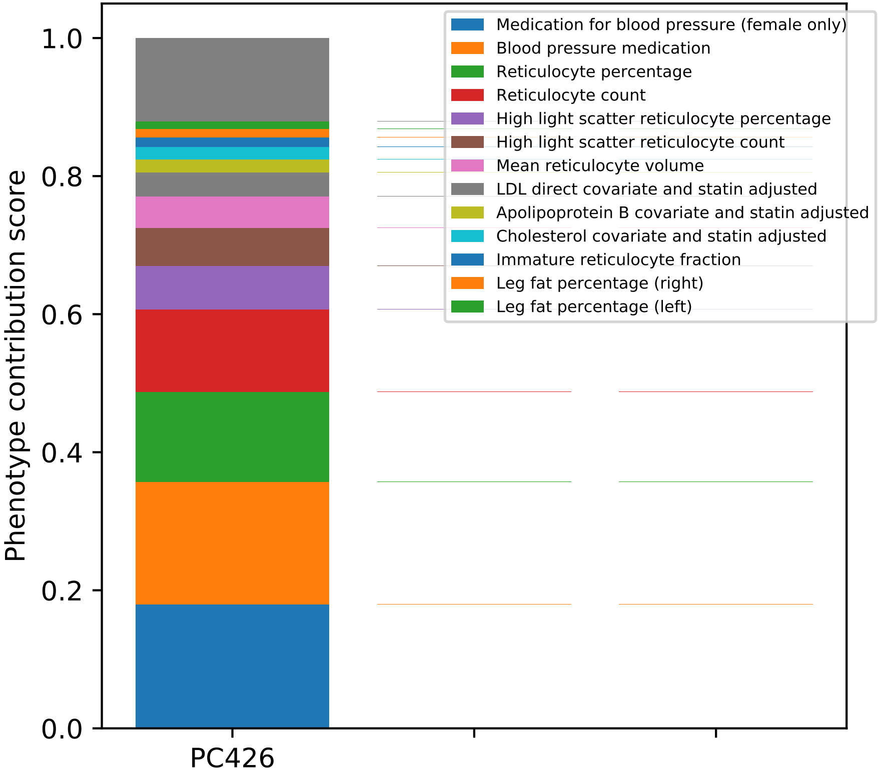

Medication for blood pressure (female only)
Blood pressure medication
Reticulocyte percentage
Reticulocyte count
High light scatter reticulocyte percentage
High light scatter reticulocyte count
Mean reticulocyte volume
LDL direct covariate and statin adjusted
Apolipoprotein B covariate and statin adjusted
Cholesterol covariate and statin adjusted
Immature reticulocyte fraction
Leg fat percentage (right)
Leg fat percentage (left)
Phenotype contribution score
1.0
0.8
0.6
0.4
0.2
0.0
PC426

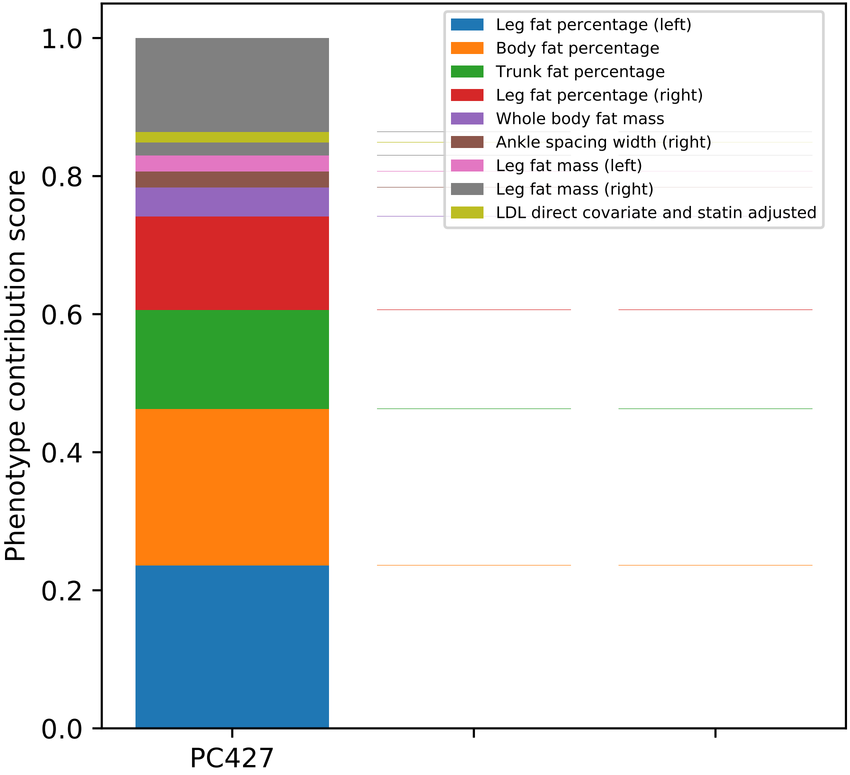
Leg fat percentage (left)
Body fat percentage
Trunk fat percentage
Leg fat percentage (right)
Whole body fat mass
Ankle spacing width (right)
Leg fat mass (left)
Leg fat mass (right)
LDL direct covariate and statin adjusted
1.0
0.8
0.6
0.4
0.2
0.0
Phenotype contribution score
PC427

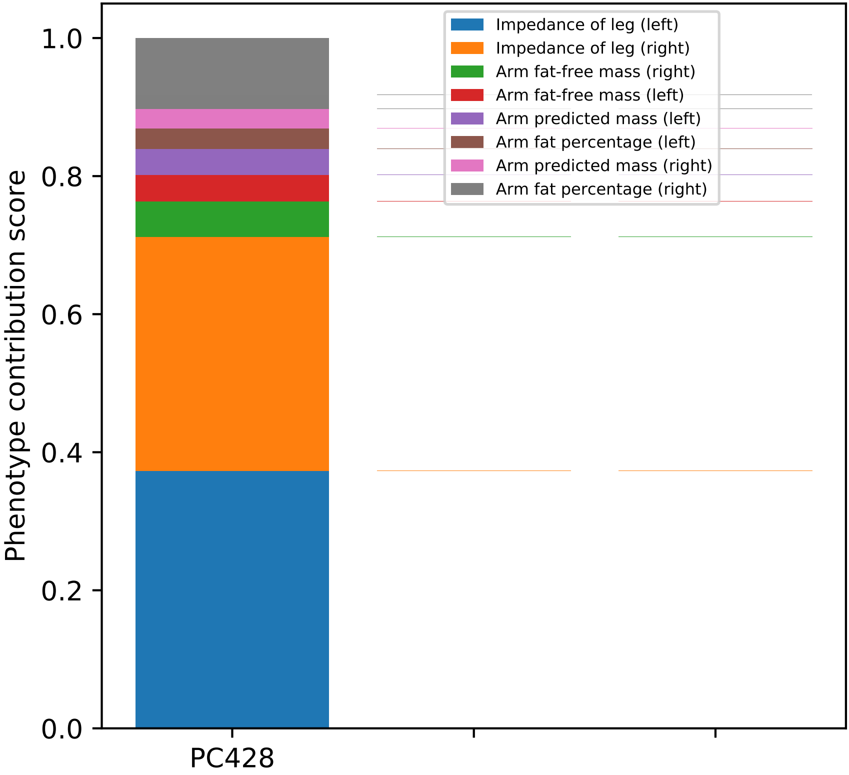

Impedance of leg (left)
Impedance of leg (right)
Arm fat-free mass (right)
Arm fat-free mass (left)
Arm predicted mass (left)
Arm fat percentage (left)
Arm predicted mass (right)
Arm fat percentage (right)
1.0
0.8
0.6
0.4
0.2
0.0
Phenotype contribution score
PC428

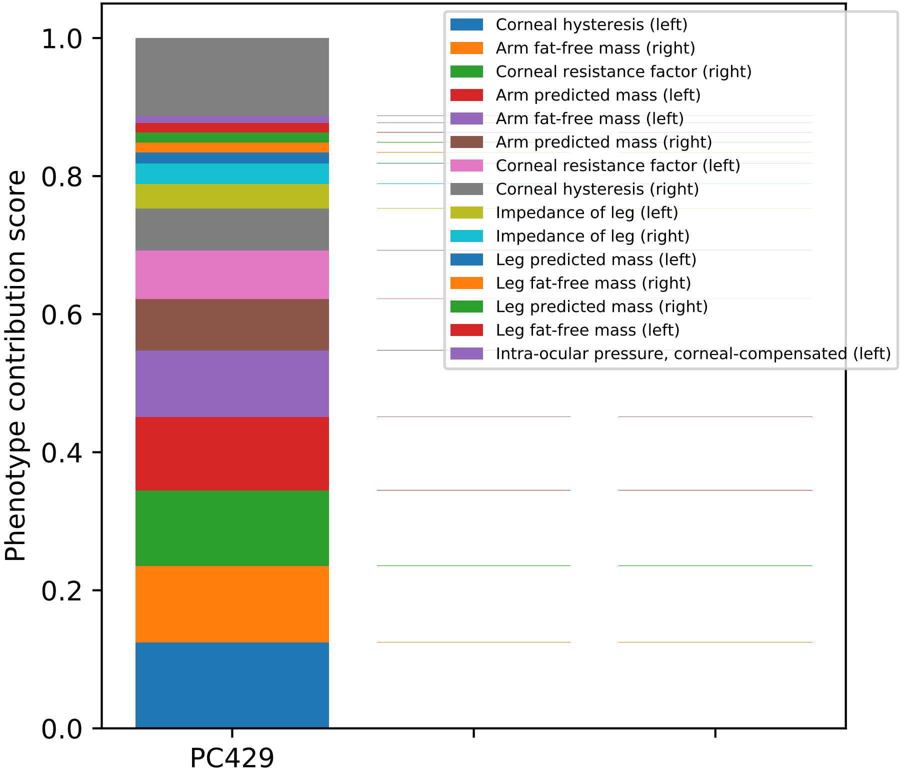

Phenotype contribution score
1.0
0.8
0.6
0.4
0.2
0.0
PC429
Corneal hysteresis (left)
Arm fat-free mass (right)
Corneal resistance factor (right)
Arm predicted mass (left)
Arm fat-free mass (left)
Arm predicted mass (right)
Corneal resistance factor (left)
Corneal hysteresis (right)
Impedance of leg (left)
Impedance of leg (right)
Leg predicted mass (left)
Leg fat-free mass (right)
Leg predicted mass (right)
Leg fat-free mass (left)
Intra-ocular pressure, corneal-compensated (left)

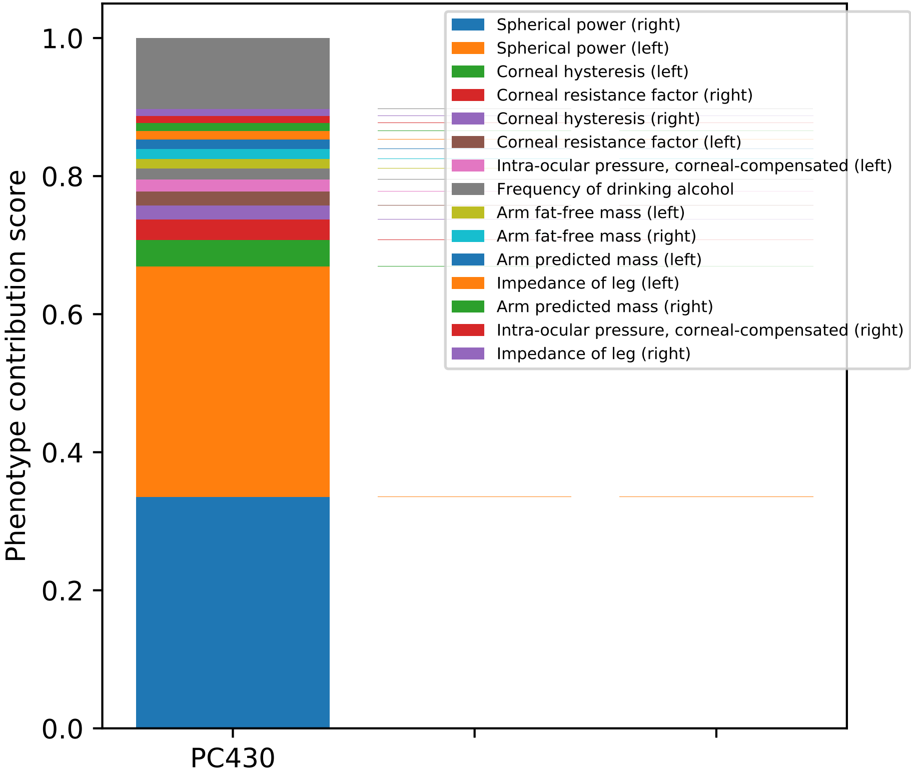

Phenotype contribution score
1.0
0.8
0.6
0.4
0.2
0.0
PC430
Spherical power (right)
Spherical power (left)
Corneal hysteresis (left)
Corneal resistance factor (right)
Corneal hysteresis (right)
Corneal resistance factor (left)
Intra-ocular pressure, corneal-compensated (left)
Frequency of drinking alcohol
Arm fat-free mass (left)
Arm fat-free mass (right)
Arm predicted mass (left)
Impedance of leg (left)
Arm predicted mass (right)
Intra-ocular pressure, corneal-compensated (right)
Impedance of leg (right)

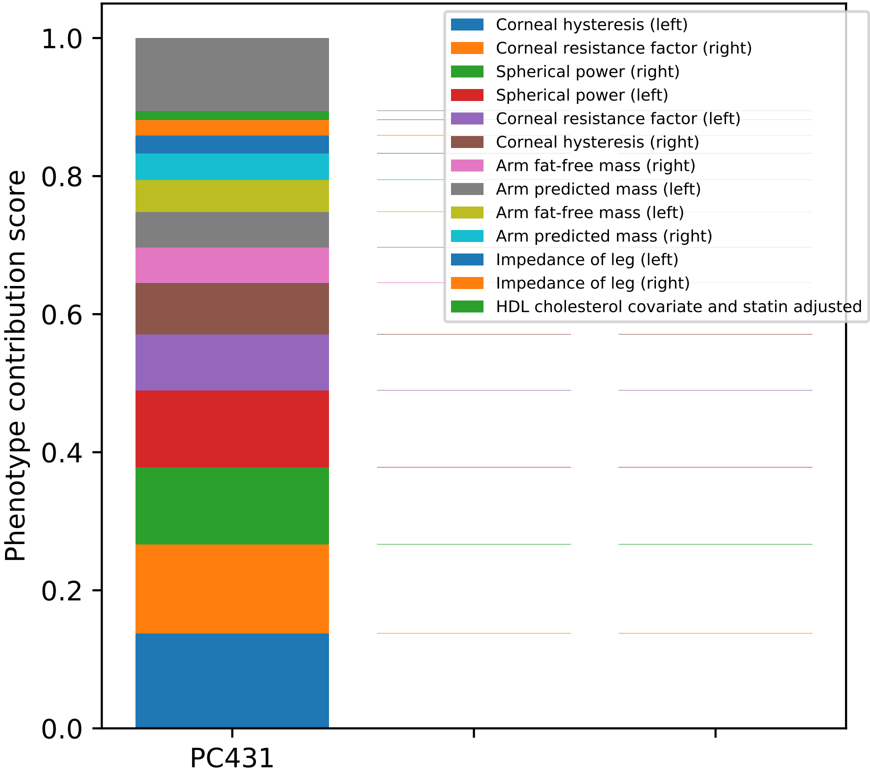
Phenotype contribution score
0.0
0.2
0.4
0.6
0.8
1.0
PC431
Corneal hysteresis (left)
Corneal resistance factor (right)
Spherical power (right)
Spherical power (left)
Corneal resistance factor (left)
Corneal hysteresis (right)
Arm fat-free mass (right)
Arm predicted mass (left)
Arm fat-free mass (left)
Arm predicted mass (right)
Impedance of leg (left)
Impedance of leg (right)
HDL cholesterol covariate and statin adjusted

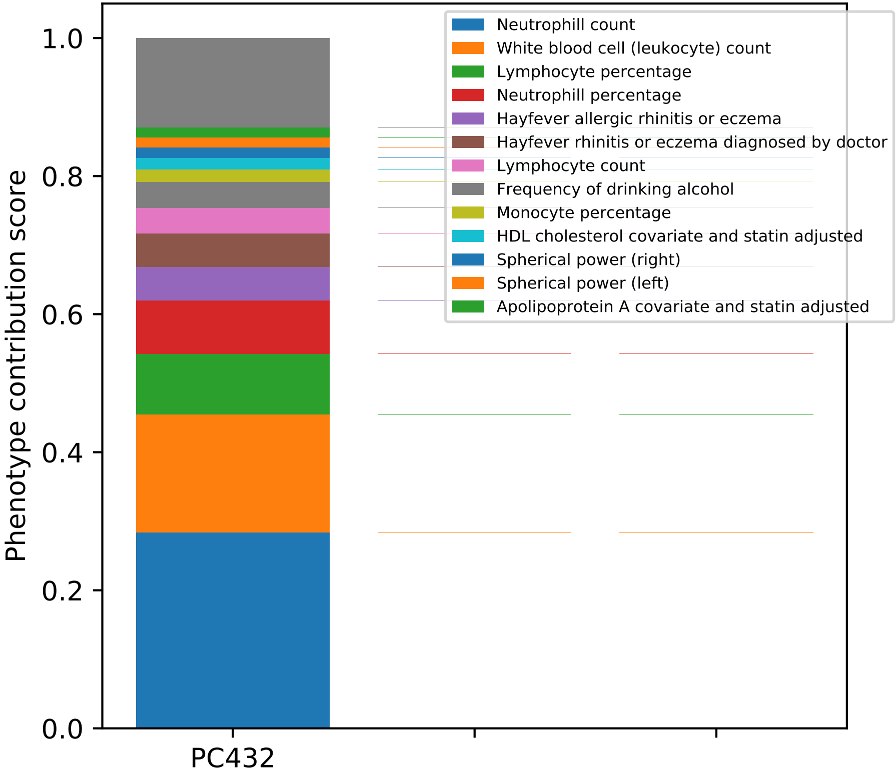

Neutrophill count
White blood cell (leukocyte) count
Lymphocyte percentage
Neutrophill percentage
Hayfever allergic rhinitis or eczema
Hayfever rhinitis or eczema diagnosed by doctor
Lymphocyte count
Frequency of drinking alcohol
Monocyte percentage
HDL cholesterol covariate and statin adjusted
Spherical power (right)
Spherical power (left)
Apolipoprotein A covariate and statin adjusted
Phenotype contribution score
0.0
0.2
0.4
0.6
0.8
1.0
PC432

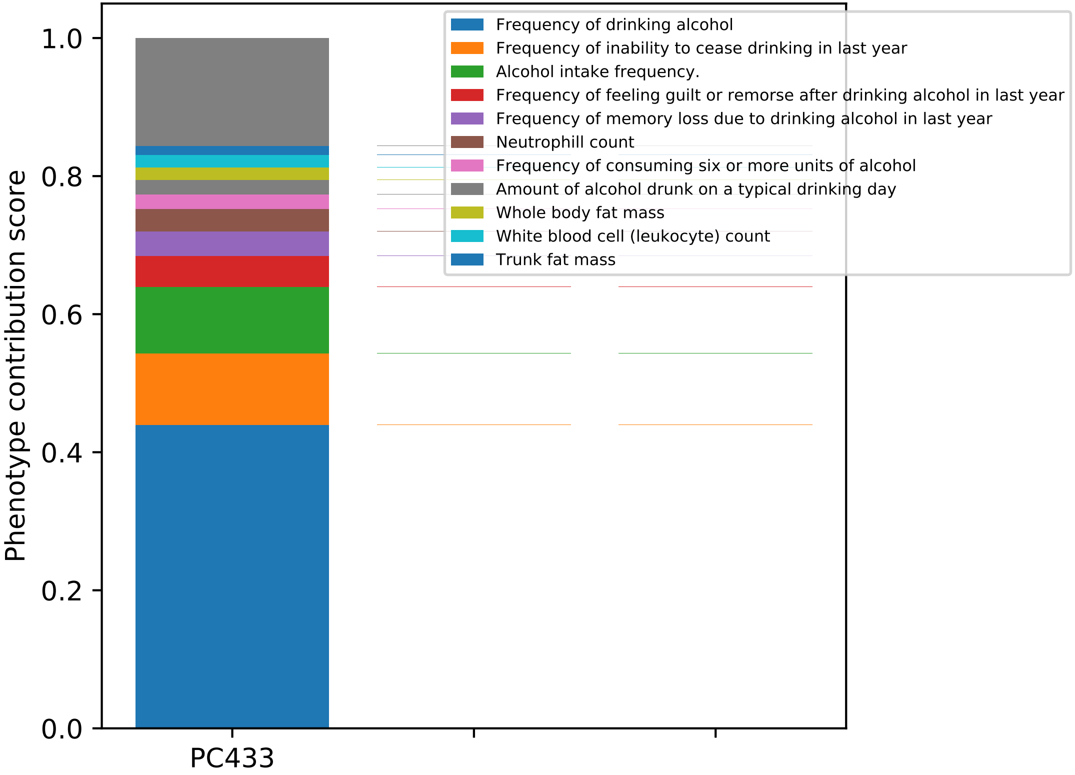

Phenotype contribution score
0.0
0.2
0.4
0.6
0.8
1.0
PC433
Frequency of drinking alcohol
Frequency of inability to cease drinking in last year
Alcohol intake frequency.
Frequency of feeling guilt or remorse after drinking alcohol in last year
Frequency of memory loss due to drinking alcohol in last year
Neutrophill count
Frequency of consuming six or more units of alcohol
Amount of alcohol drunk on a typical drinking day
Whole body fat mass
White blood cell (leukocyte) count
Trunk fat mass

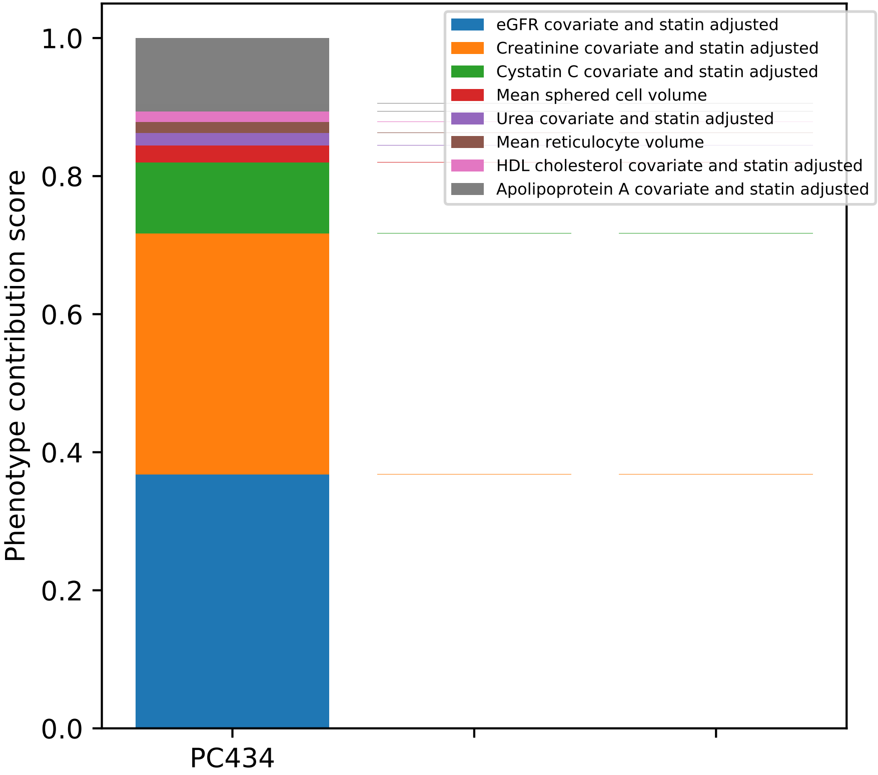

eGFR covariate and statin adjusted
Creatinine covariate and statin adjusted
Cystatin C covariate and statin adjusted
Mean sphered cell volume
Urea covariate and statin adjusted
Mean reticulocyte volume
HDL cholesterol covariate and statin adjusted
Apolipoprotein A covariate and statin adjusted
Phenotype contribution score
0.0
0.2
0.4
0.6
0.8
1.0
PC434

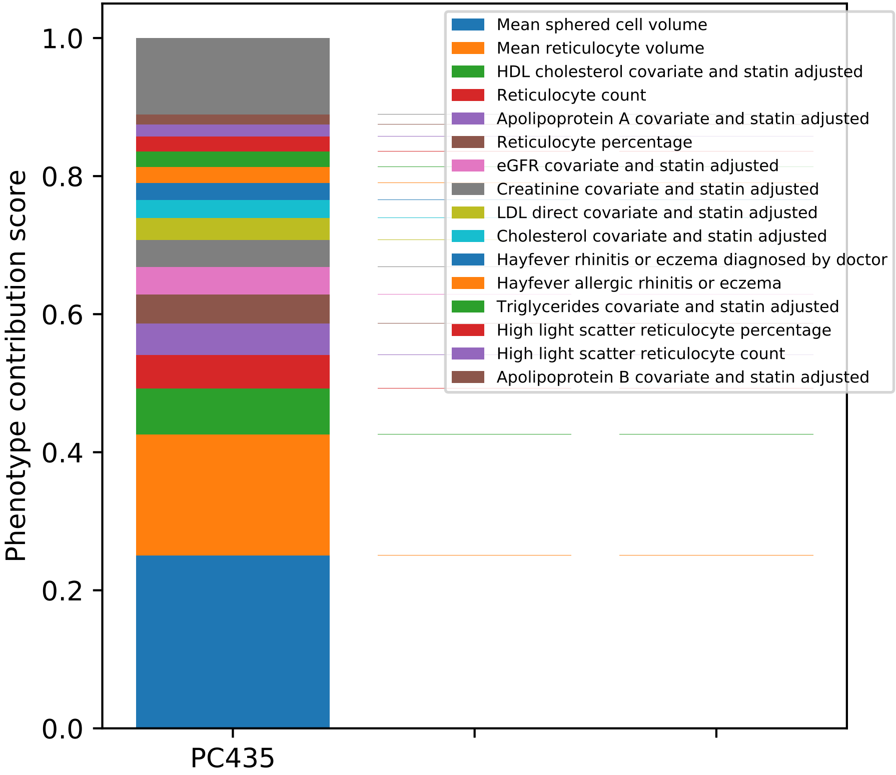
Mean sphered cell volume
Mean reticulocyte volume
HDL cholesterol covariate and statin adjusted
Reticulocyte count
Apolipoprotein A covariate and statin adjusted
Reticulocyte percentage
eGFR covariate and statin adjusted
Creatinine covariate and statin adjusted
LDL direct covariate and statin adjusted
Cholesterol covariate and statin adjusted
Hayfever rhinitis or eczema diagnosed by doctor
Hayfever allergic rhinitis or eczema
Triglycerides covariate and statin adjusted
High light scatter reticulocyte percentage
High light scatter reticulocyte count
Apolipoprotein B covariate and statin adjusted
Phenotype contribution score
0.0
0.2
0.4
0.6
0.8
1.0
PC435

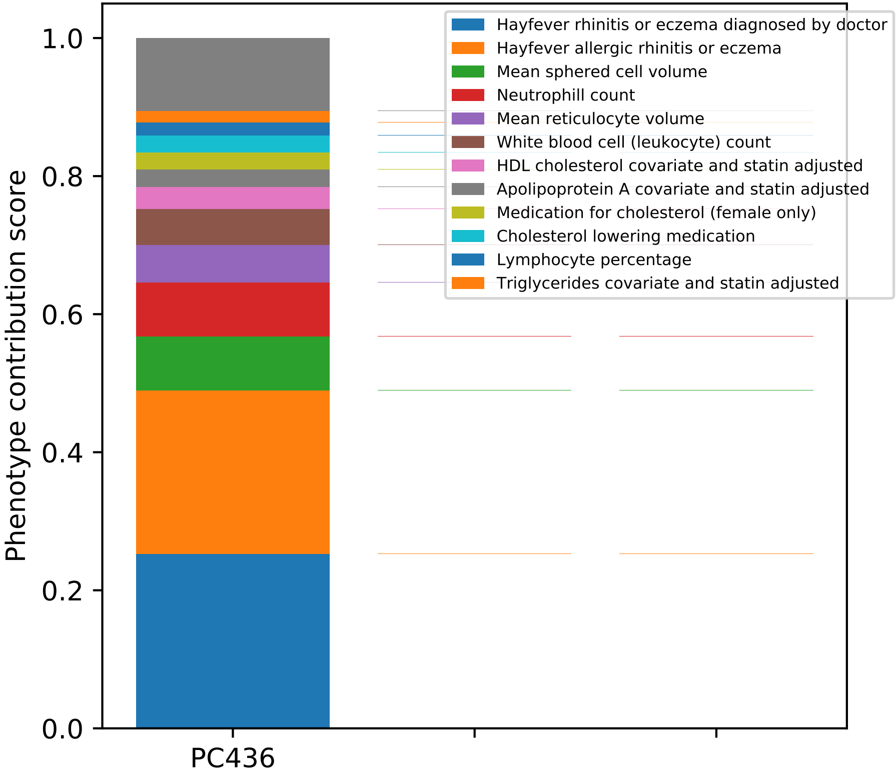

Phenotype contribution score
0.0
0.2
0.4
0.6
0.8
1.0
PC436
Hayfever rhinitis or eczema diagnosed by doctor
Hayfever allergic rhinitis or eczema
Mean sphered cell volume
Neutrophill count
Mean reticulocyte volume
White blood cell (leukocyte) count
HDL cholesterol covariate and statin adjusted
Apolipoprotein A covariate and statin adjusted
Medication for cholesterol (female only)
Cholesterol lowering medication
Lymphocyte percentage
Triglycerides covariate and statin adjusted

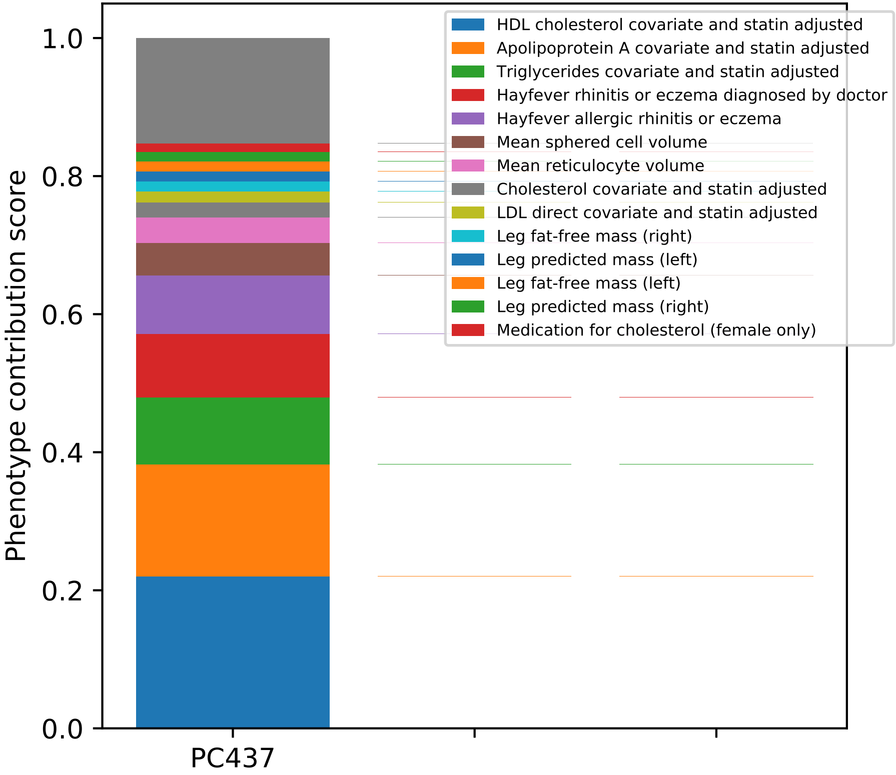

Phenotype contribution score
0.0
0.2
0.4
0.6
0.8
1.0
PC437
HDL cholesterol covariate and statin adjusted
Apolipoprotein A covariate and statin adjusted
Triglycerides covariate and statin adjusted
Hayfever rhinitis or eczema diagnosed by doctor
Hayfever allergic rhinitis or eczema
Mean sphered cell volume
Mean reticulocyte volume
Cholesterol covariate and statin adjusted
LDL direct covariate and statin adjusted
Leg fat-free mass (right)
Leg predicted mass (left)
Leg fat-free mass (left)
Leg predicted mass (right)
Medication for cholesterol (female only)

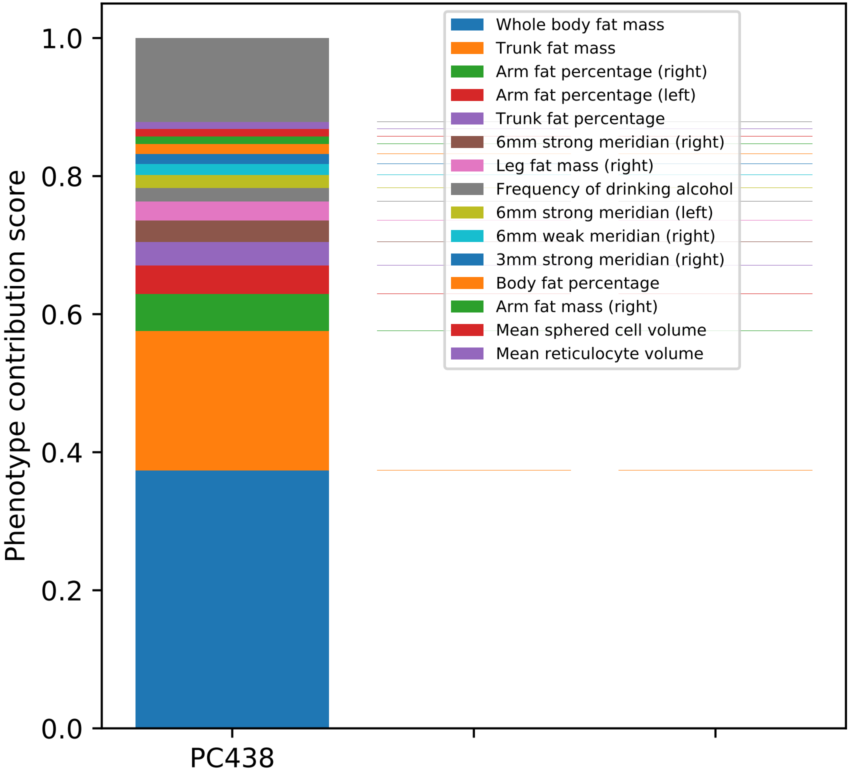
Phenotype contribution score
0.0
0.2
0.4
0.6
0.8
1.0
PC438
Whole body fat mass
Trunk fat mass
Arm fat percentage (right)
Arm fat percentage (left)
Trunk fat percentage
6mm strong meridian (right)
Leg fat mass (right)
Frequency of drinking alcohol
6mm strong meridian (left)
6mm weak meridian (right)
3mm strong meridian (right)
Body fat percentage
Arm fat mass (right)
Mean sphered cell volume
Mean reticulocyte volume

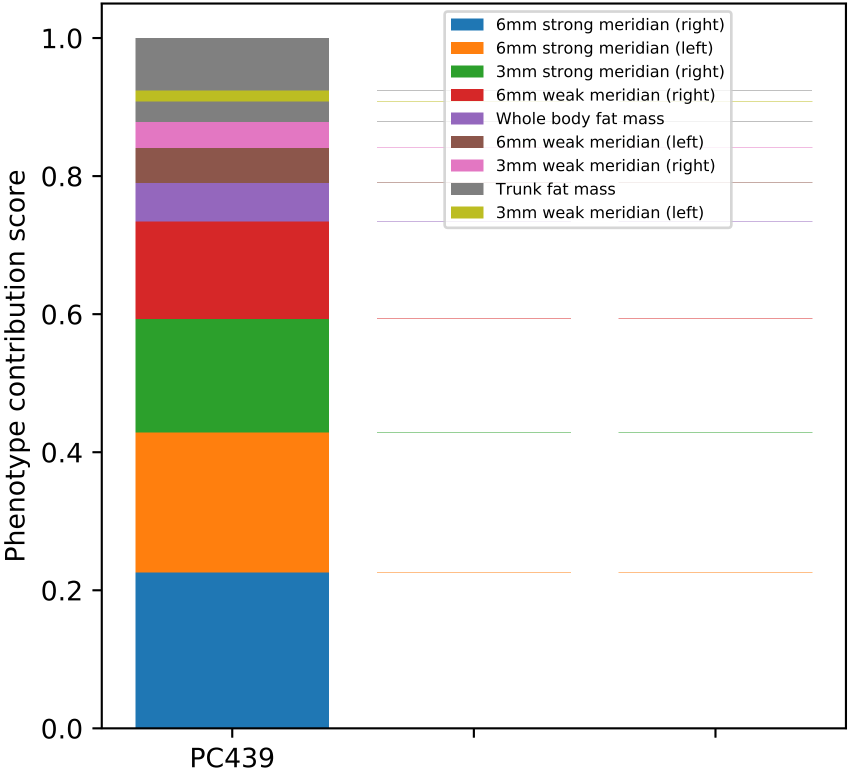
Phenotype contribution score
0.0
0.2
0.4
0.6
0.8
1.0
PC439
6mm strong meridian (right)
6mm strong meridian (left)
3mm strong meridian (right)
6mm weak meridian (right)
Whole body fat mass
6mm weak meridian (left)
3mm weak meridian (right)
Trunk fat mass
3mm weak meridian (left)

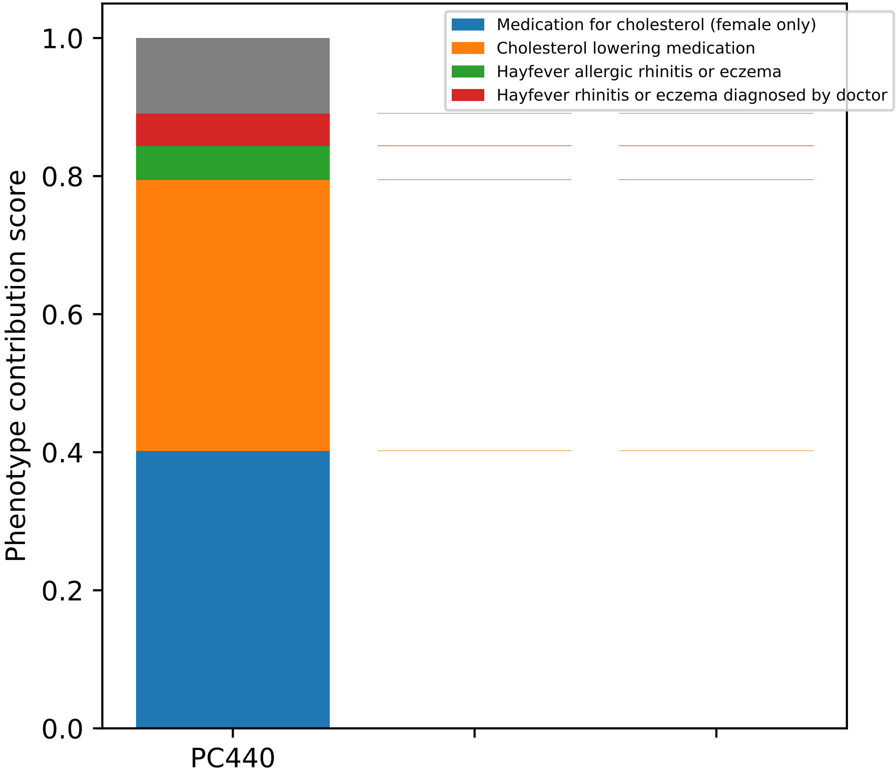

Medication for cholesterol (female only)
Cholesterol lowering medication
Hayfever allergic rhinitis or eczema
Hayfever rhinitis or eczema diagnosed by doctor
1.0
0.8
0.6
0.4
0.2
0.0
Phenotype contribution score
PC440

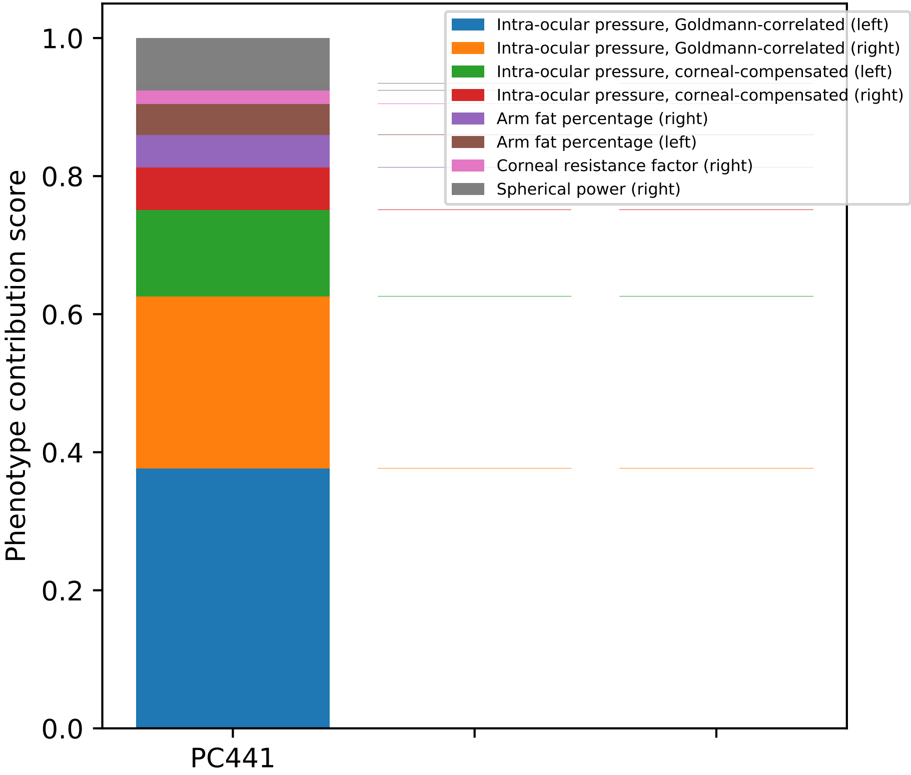
Phenotype contribution score
0.0
0.2
0.4
0.6
0.8
1.0
PC441
Intra-ocular pressure, Goldmann-correlated (left)
Intra-ocular pressure, Goldmann-correlated (right)
Intra-ocular pressure, corneal-compensated (left)
Intra-ocular pressure, corneal-compensated (right)
Arm fat percentage (right)
Arm fat percentage (left)
Corneal resistance factor (right)
Spherical power (right)

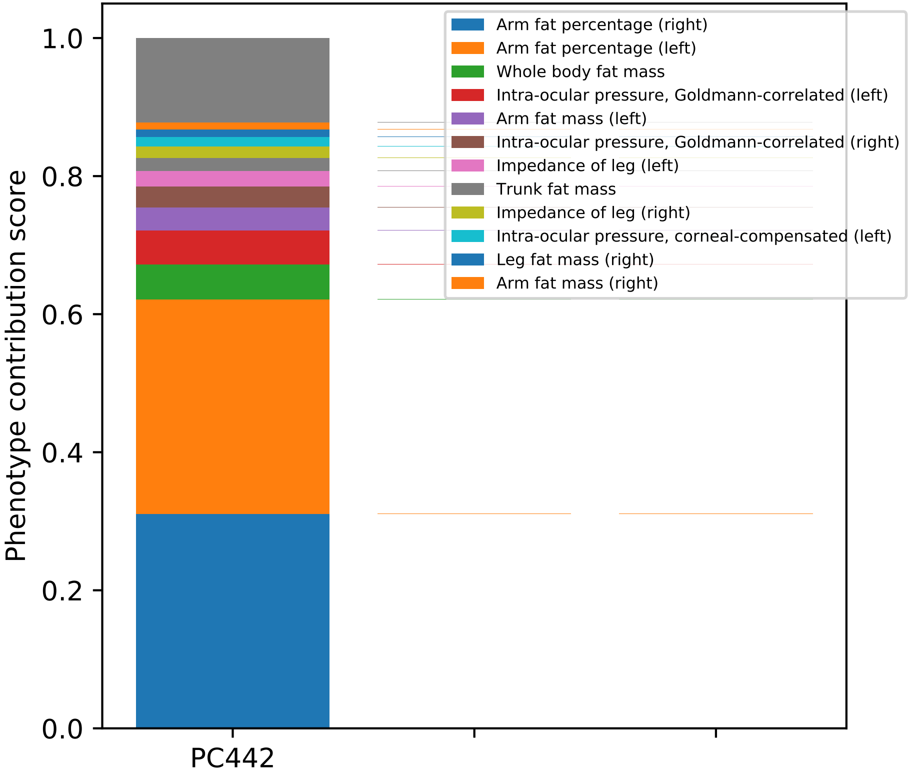

Phenotype contribution score
1.0
0.8
0.6
0.4
0.2
0.0
PC442
Arm fat percentage (right)
Arm fat percentage (left)
Whole body fat mass
Intra-ocular pressure, Goldmann-correlated (left)
Arm fat mass (left)
Intra-ocular pressure, Goldmann-correlated (right)
Impedance of leg (left)
Trunk fat mass
Impedance of leg (right)
Intra-ocular pressure, corneal-compensated (left)
Leg fat mass (right)
Arm fat mass (right)

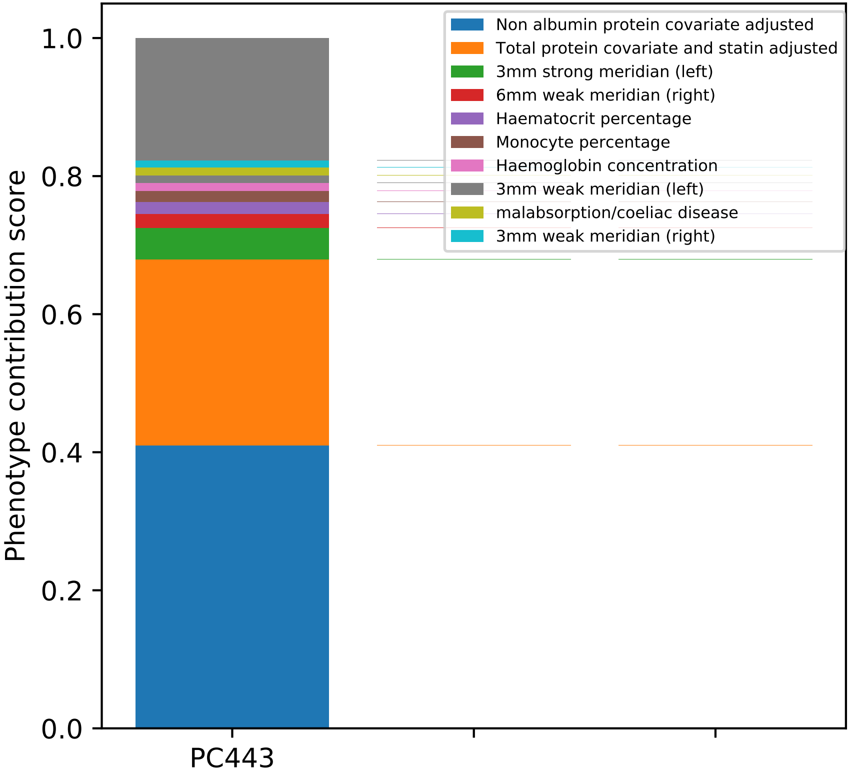

Phenotype contribution score
1.0
0.8
0.6
0.4
0.2
0.0
PC443
Non albumin protein covariate adjusted
Total protein covariate and statin adjusted
3mm strong meridian (left)
6mm weak meridian (right)
Haematocrit percentage
Monocyte percentage
Haemoglobin concentration
3mm weak meridian (left)
malabsorption/coeliac disease
3mm weak meridian (right)

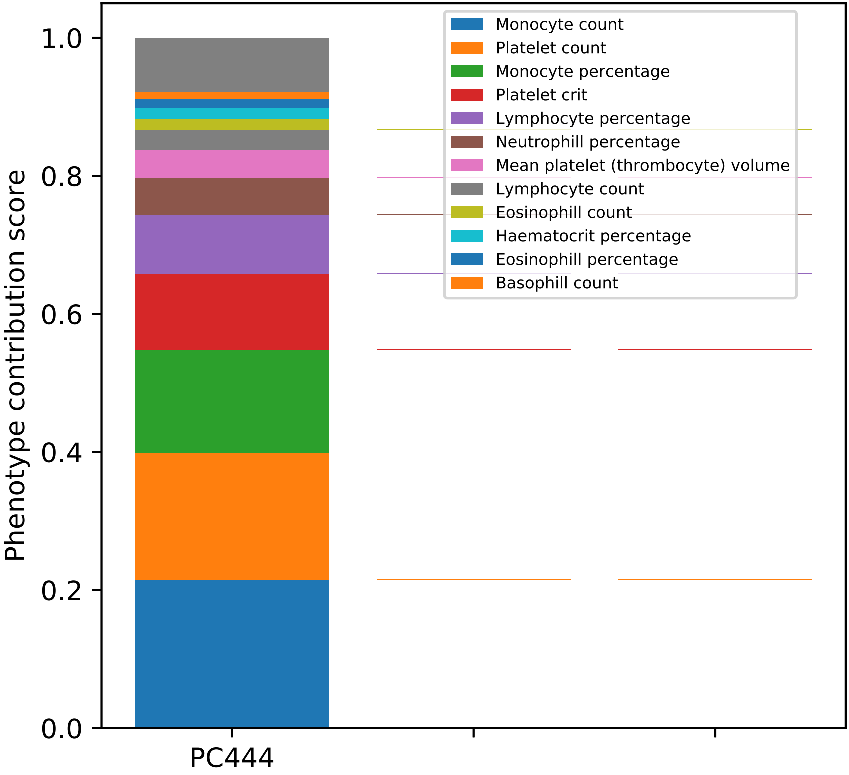
Phenotype contribution score
1.0
0.8
0.6
0.4
0.2
0.0
PC444
Monocyte count
Platelet count
Monocyte percentage
Platelet crit
Lymphocyte percentage
Neutrophill percentage
Mean platelet (thrombocyte) volume
Lymphocyte count
Eosinophill count
Haematocrit percentage
Eosinophill percentage
Basophill count

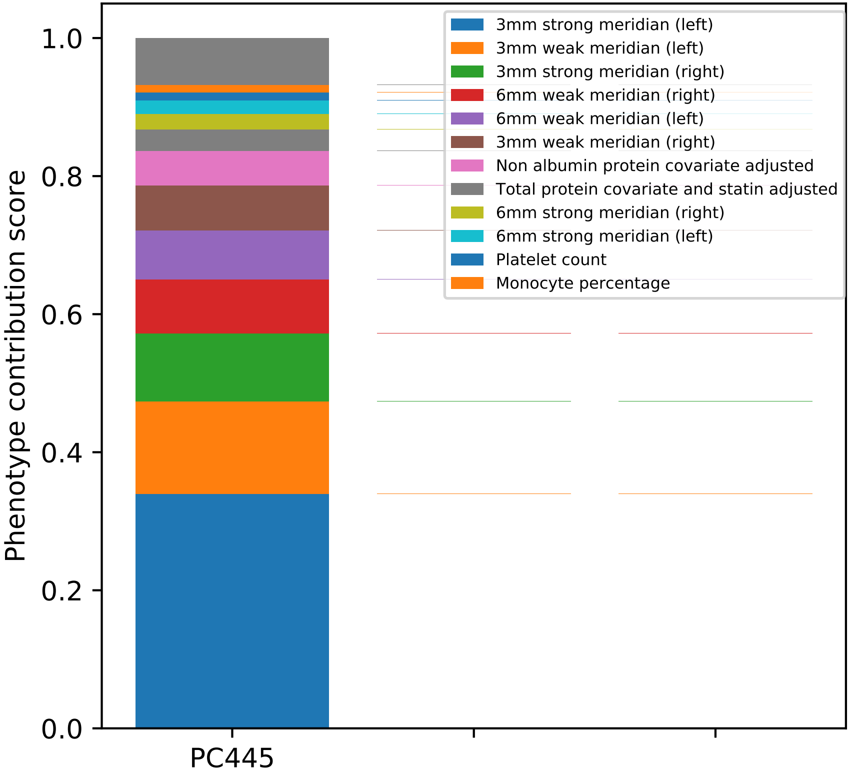

Phenotype contribution score
0.0
0.2
0.4
0.6
0.8
1.0
PC445
3mm strong meridian (left)
3mm weak meridian (left)
3mm strong meridian (right)
6mm weak meridian (right)
6mm weak meridian (left)
3mm weak meridian (right)
Non albumin protein covariate adjusted
Total protein covariate and statin adjusted
6mm strong meridian (right)
6mm strong meridian (left)
Platelet count
Monocyte percentage

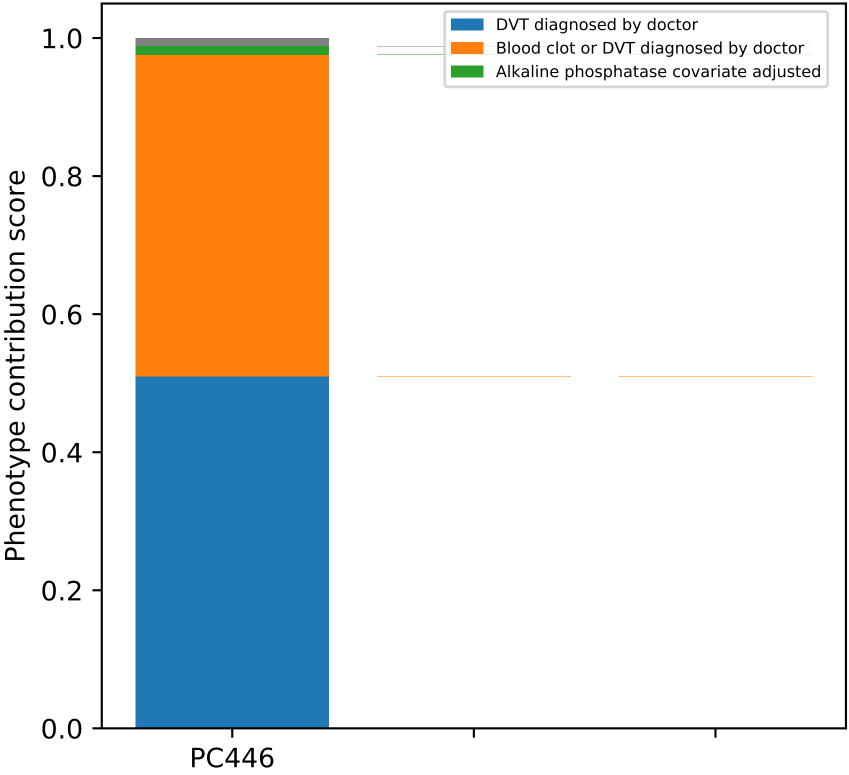

DVT diagnosed by doctor
Blood clot or DVT diagnosed by doctor
Alkaline phosphatase covariate adjusted
Phenotype contribution score
1.0
0.8
0.6
0.4
0.2
0.0
PC446

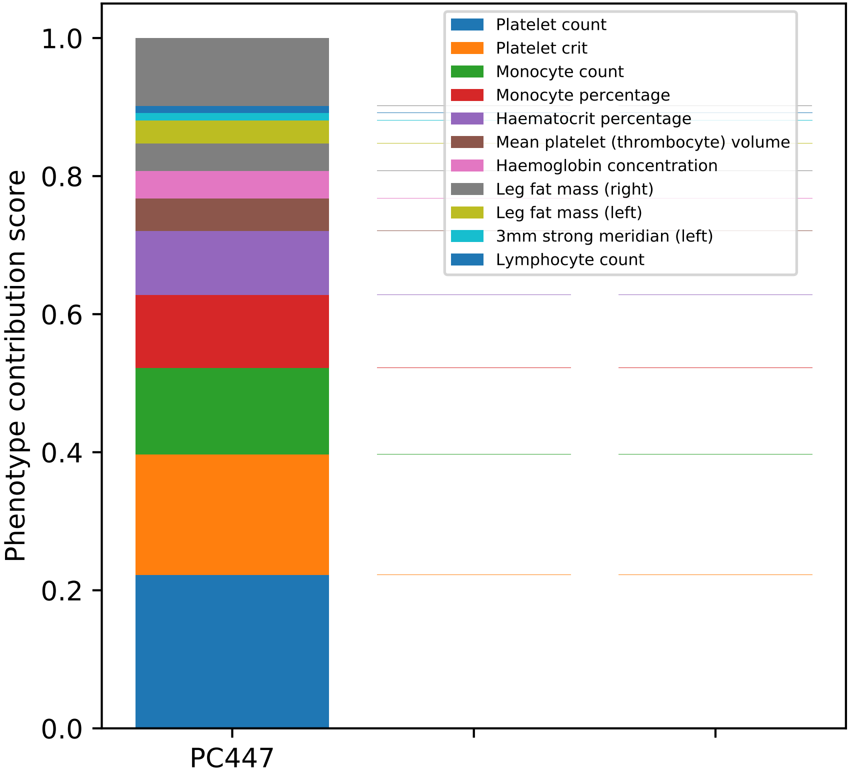
Phenotype contribution score
1.0
0.8
0.6
0.4
0.2
0.0
PC447
Platelet count
Platelet crit
Monocyte count
Monocyte percentage
Haematocrit percentage
Mean platelet (thrombocyte) volume
Haemoglobin concentration
Leg fat mass (right)
Leg fat mass (left)
3mm strong meridian (left)
Lymphocyte count

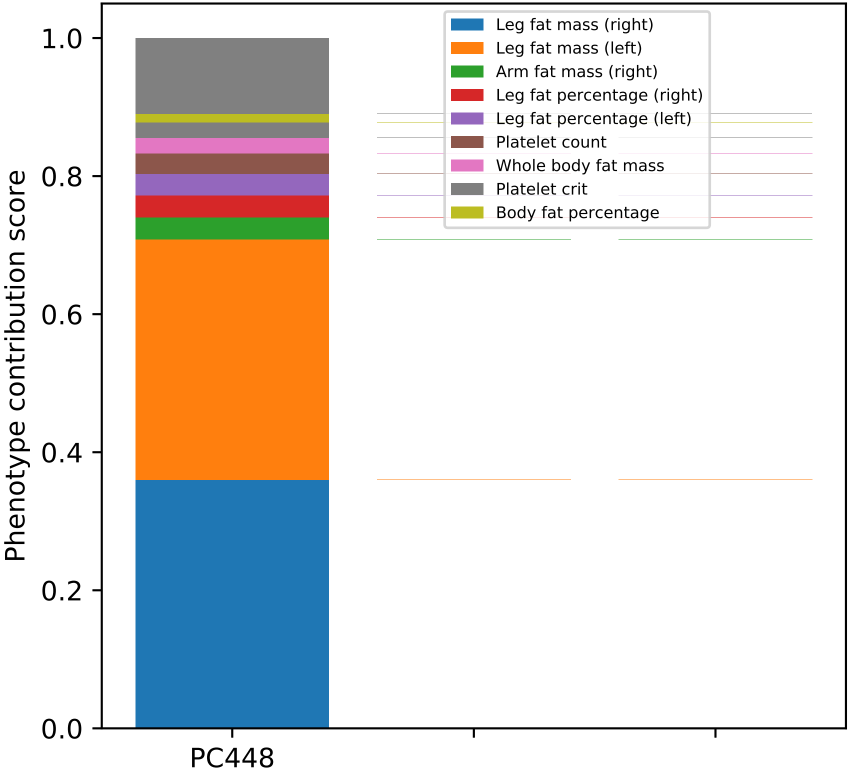
Leg fat mass (right)
Leg fat mass (left)
Arm fat mass (right)
Leg fat percentage (right)
Leg fat percentage (left)
Platelet count
Whole body fat mass
Platelet crit
Body fat percentage
Phenotype contribution score
1.0
0.8
0.6
0.4
0.2
0.0
PC448

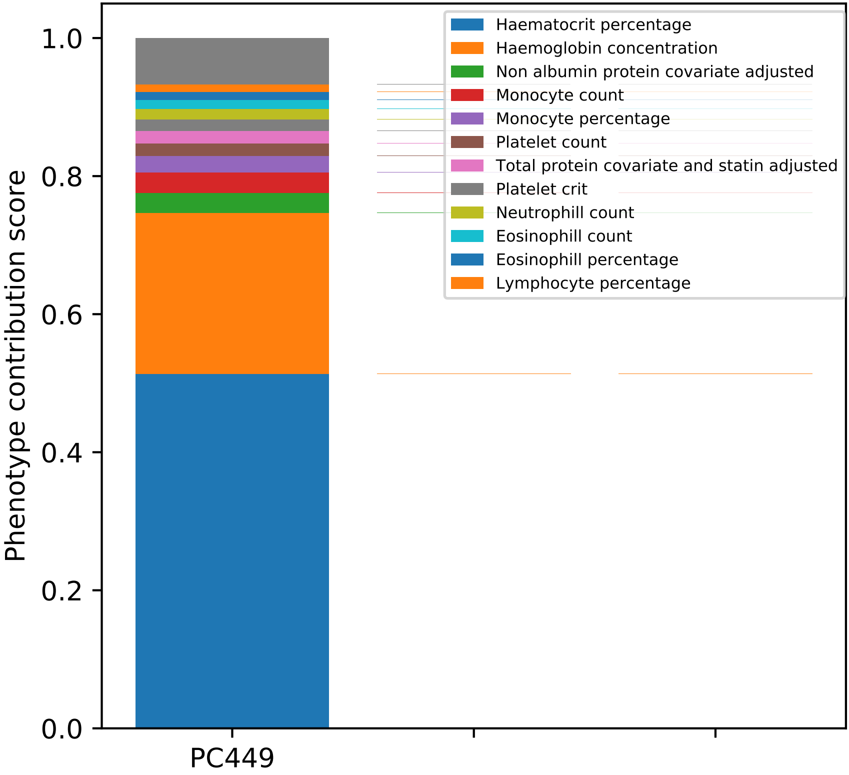

Haematocrit percentage
Haemoglobin concentration
Non albumin protein covariate adjusted
Monocyte count
Monocyte percentage
Platelet count
Total protein covariate and statin adjusted
Platelet crit
Neutrophill count
Eosinophill count
Eosinophill percentage
Lymphocyte percentage
Phenotype contribution score
1.0
0.8
0.6
0.4
0.2
0.0
PC449

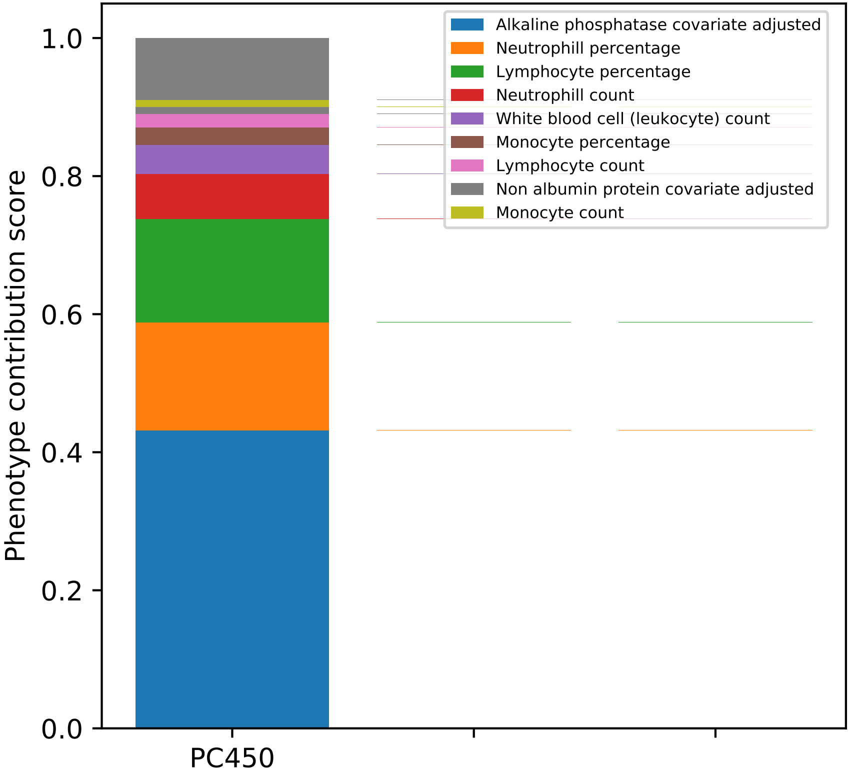

Phenotype contribution score
1.0
0.8
0.6
0.4
0.2
0.0
PC450
Alkaline phosphatase covariate adjusted
Neutrophill percentage
Lymphocyte percentage
Neutrophill count
White blood cell (leukocyte) count
Monocyte percentage
Lymphocyte count
Non albumin protein covariate adjusted
Monocyte count

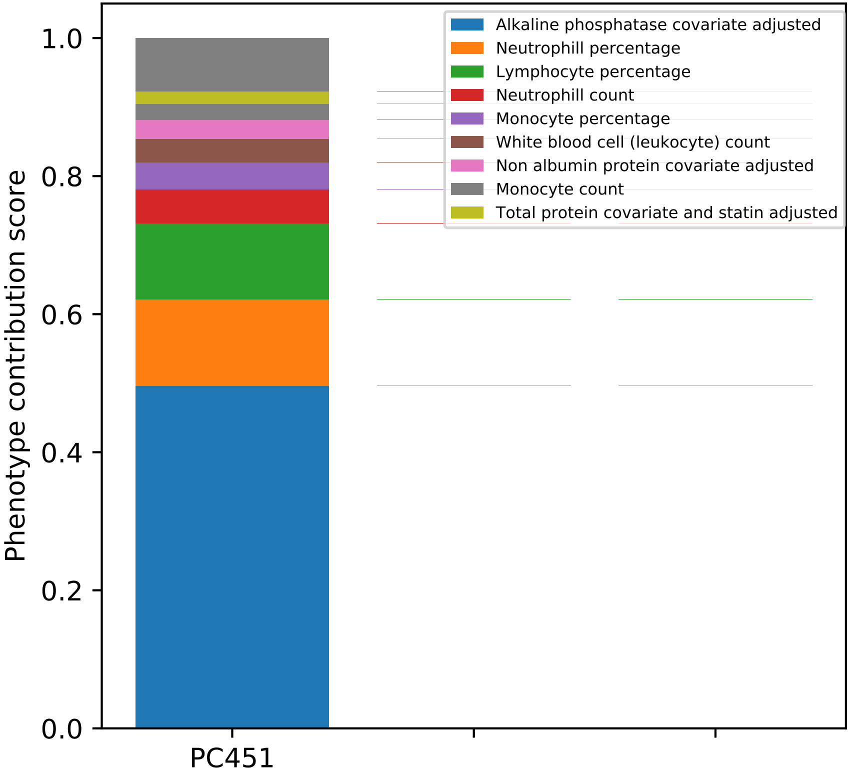
Alkaline phosphatase covariate adjusted
Neutrophill percentage
Lymphocyte percentage
Neutrophill count
Monocyte percentage
White blood cell (leukocyte) count
Non albumin protein covariate adjusted
Monocyte count
Total protein covariate and statin adjusted
Phenotype contribution score
1.0
0.8
0.6
0.4
0.2
0.0
PC451

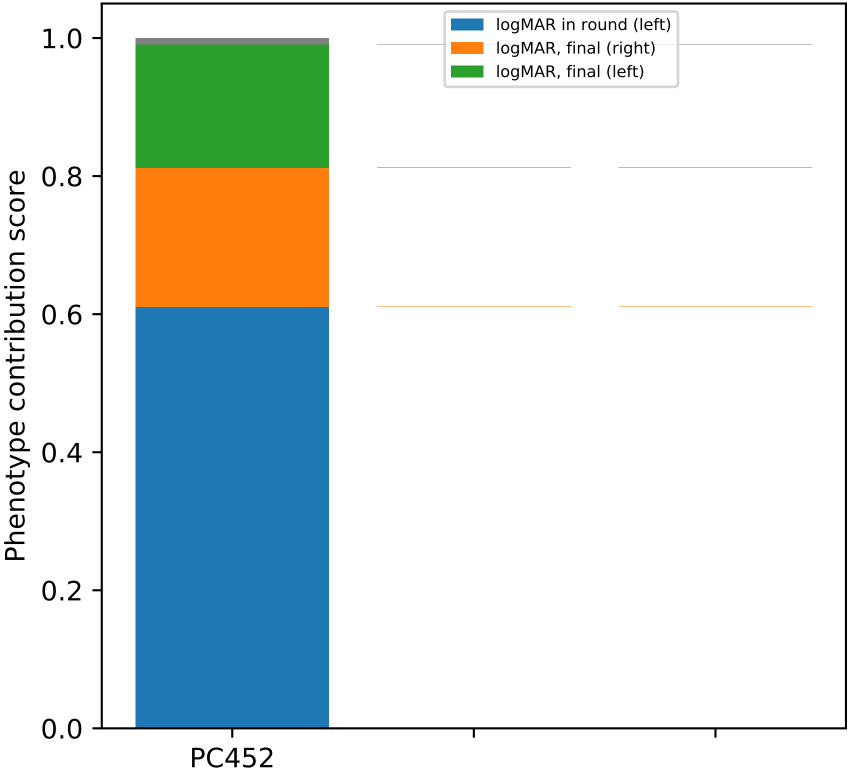
logMAR in round (left)
logMAR, final (right)
logMAR, final (left)
Phenotype contribution score
0.0
0.2
0.4
0.6
0.8
1.0
PC452

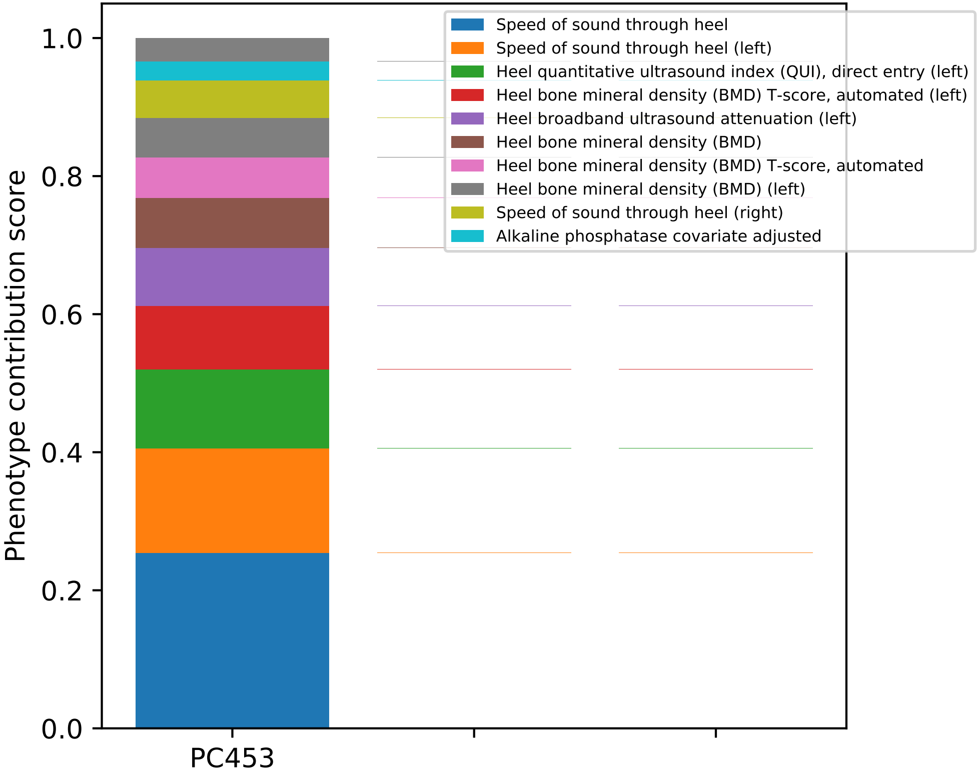

Speed of sound through heel
Speed of sound through heel (left)
Heel quantitative ultrasound index (QUI), direct entry (left)
Heel bone mineral density (BMD) T-score, automated (left)
Heel broadband ultrasound attenuation (left)
Heel bone mineral density (BMD)
Heel bone mineral density (BMD) T-score, automated
Heel bone mineral density (BMD) (left)
Speed of sound through heel (right)
Alkaline phosphatase covariate adjusted
Phenotype contribution score
0.0
0.2
0.4
0.6
0.8
1.0
PC453

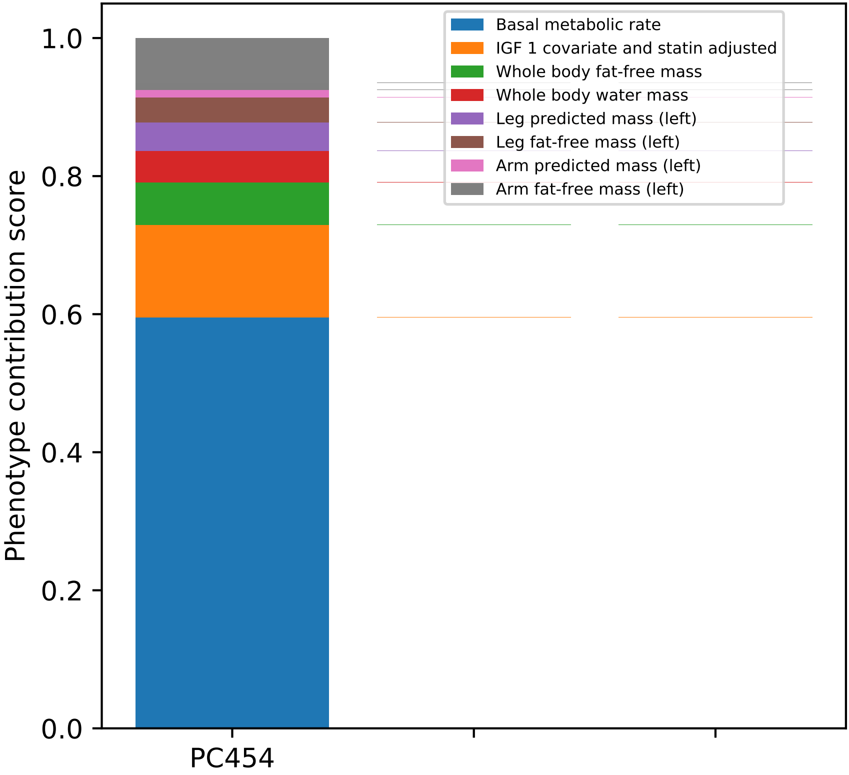

Basal metabolic rate
IGF 1 covariate and statin adjusted
Whole body fat-free mass
Whole body water mass
Leg predicted mass (left)
Leg fat-free mass (left)
Arm predicted mass (left)
Arm fat-free mass (left)
Phenotype contribution score
0.0
0.2
0.4
0.6
0.8
1.0
PC454

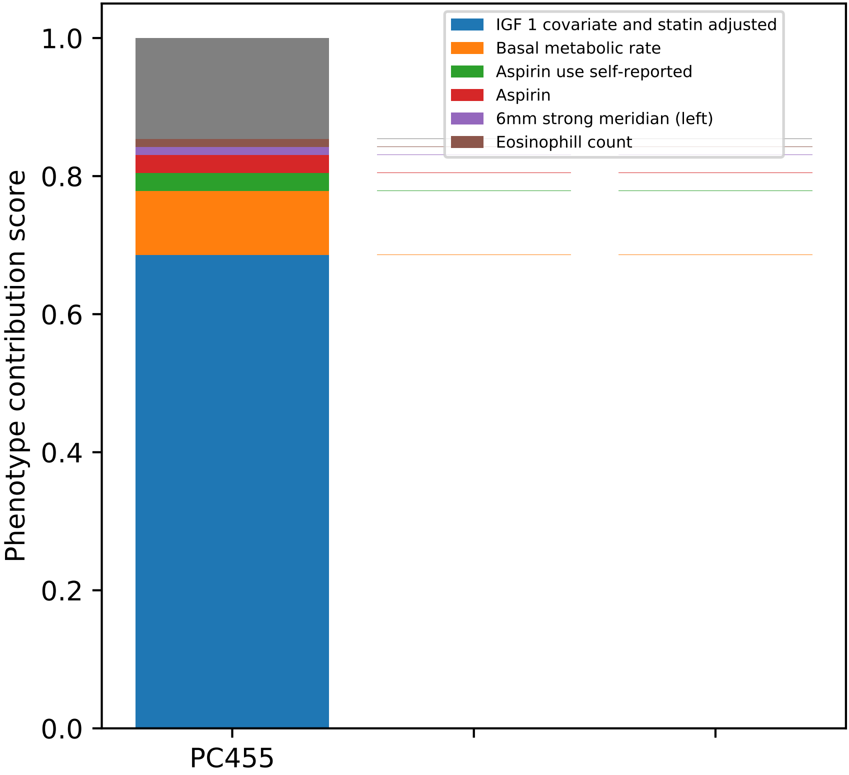

IGF 1 covariate and statin adjusted
Basal metabolic rate
Aspirin use self-reported
Aspirin
6mm strong meridian (left)
Eosinophill count
1.0
0.8
0.6
0.4
0.2
0.0
Phenotype contribution score
PC455

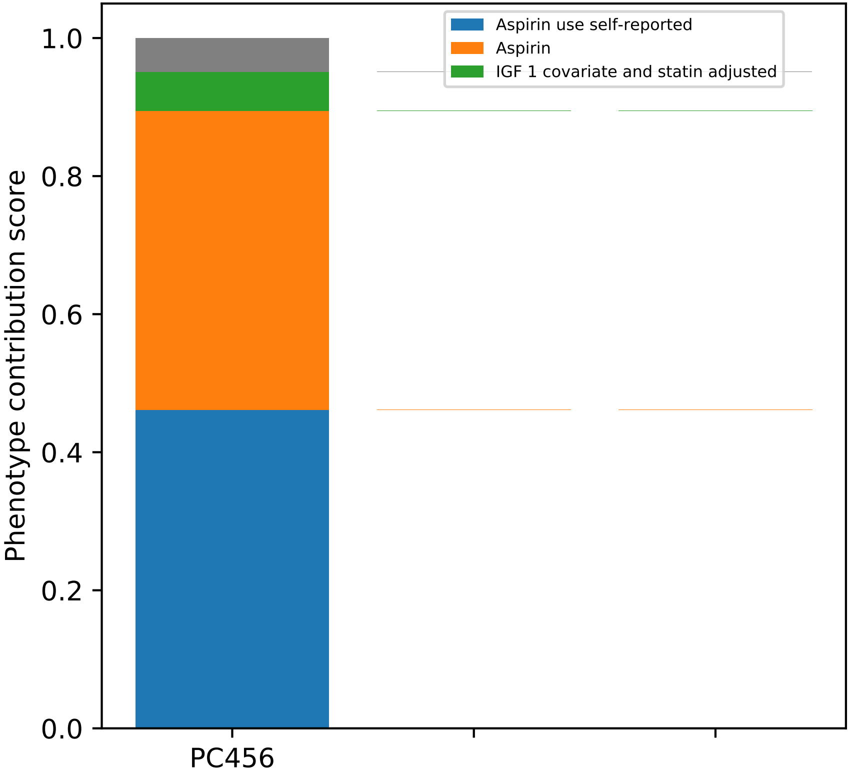

Aspirin use self-reported
Aspirin
IGF 1 covariate and statin adjusted
Phenotype contribution score
1.0
0.8
0.6
0.4
0.2
0.0
PC456

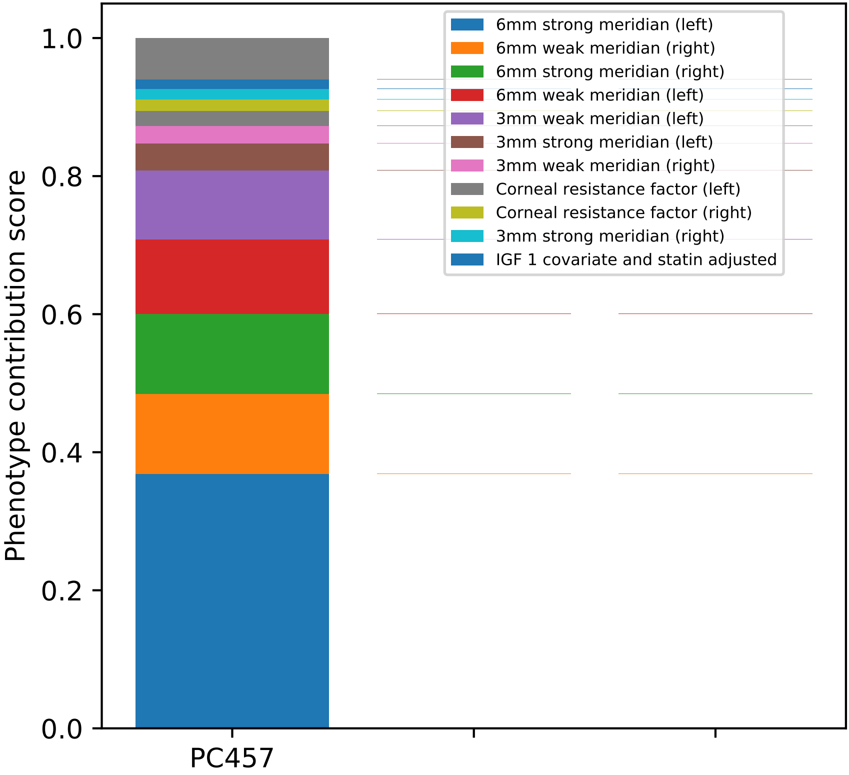

Phenotype contribution score
0.0
0.2
0.4
0.6
0.8
1.0
PC457
6mm strong meridian (left)
6mm weak meridian (right)
6mm strong meridian (right)
6mm weak meridian (left)
3mm weak meridian (left)
3mm strong meridian (left)
3mm weak meridian (right)
Corneal resistance factor (left)
Corneal resistance factor (right)
3mm strong meridian (right)
IGF 1 covariate and statin adjusted

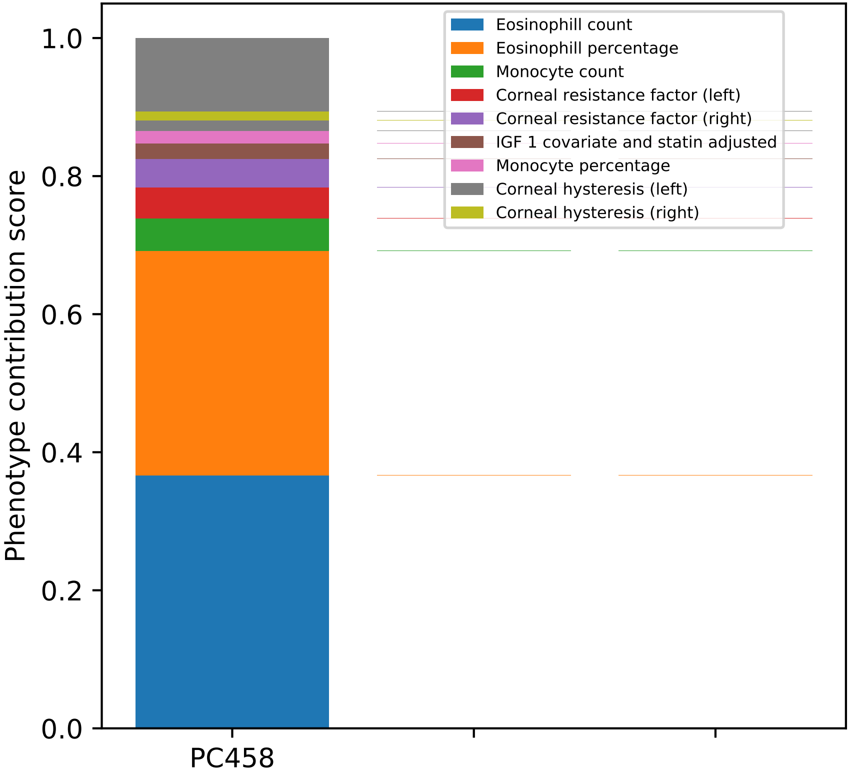

Eosinophill count
Eosinophill percentage
Monocyte count
Corneal resistance factor (left)
Corneal resistance factor (right)
IGF 1 covariate and statin adjusted
Monocyte percentage
Corneal hysteresis (left)
Corneal hysteresis (right)
Phenotype contribution score
1.0
0.8
0.6
0.4
0.2
0.0
PC458

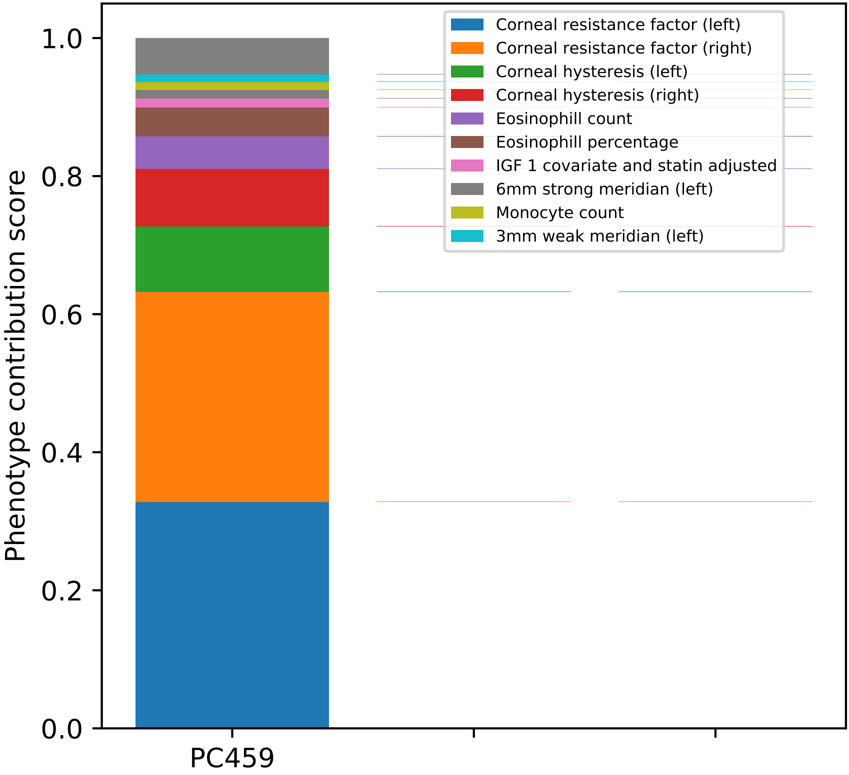

Corneal resistance factor (left)
Corneal resistance factor (right)
Corneal hysteresis (left)
Corneal hysteresis (right)
Eosinophill count
Eosinophill percentage
IGF 1 covariate and statin adjusted
6mm strong meridian (left)
Monocyte count
3mm weak meridian (left)
Phenotype contribution score
0.0
0.2
0.4
0.6
0.8
1.0
PC459

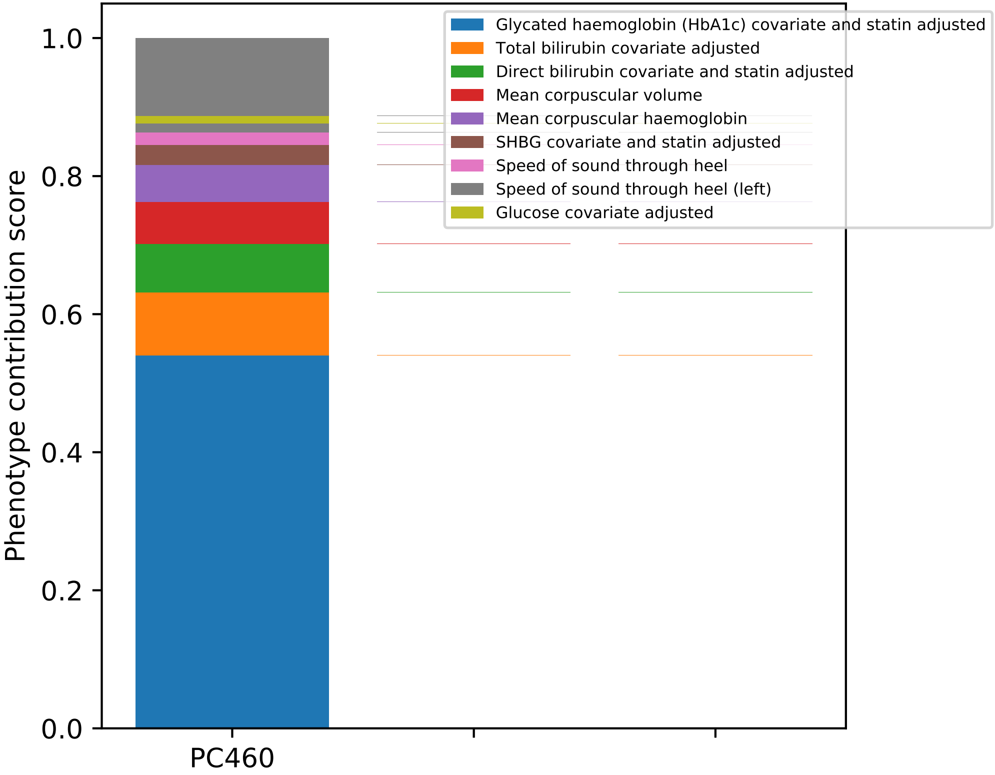

Phenotype contribution score
1.0
0.8
0.6
0.4
0.2
0.0
PC460
Glycated haemoglobin (HbA1c) covariate and statin adjusted
Total bilirubin covariate adjusted
Direct bilirubin covariate and statin adjusted
Mean corpuscular volume
Mean corpuscular haemoglobin
SHBG covariate and statin adjusted
Speed of sound through heel
Speed of sound through heel (left)
Glucose covariate adjusted

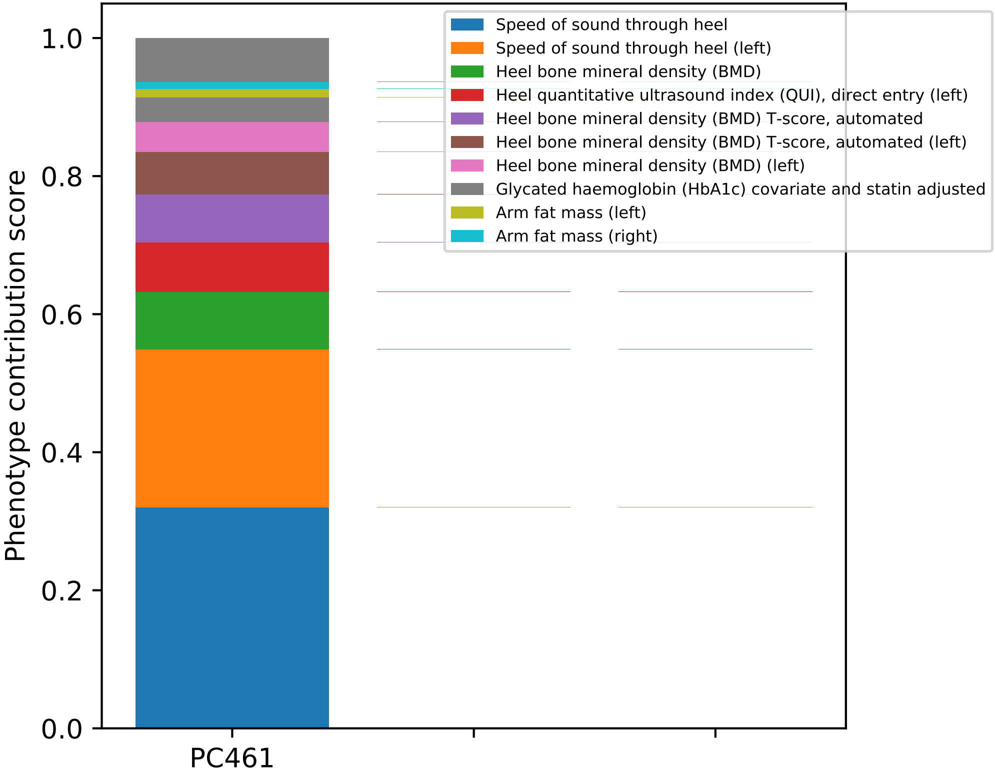

Phenotype contribution score
1.0
0.8
0.6
0.4
0.2
0.0
PC461
Speed of sound through heel
Speed of sound through heel (left)
Heel bone mineral density (BMD)
Heel quantitative ultrasound index (QUI), direct entry (left)
Heel bone mineral density (BMD) T-score, automated
Heel bone mineral density (BMD) T-score, automated (left)
Heel bone mineral density (BMD) (left)
Glycated haemoglobin (HbA1c) covariate and statin adjusted
Arm fat mass (left)
Arm fat mass (right)

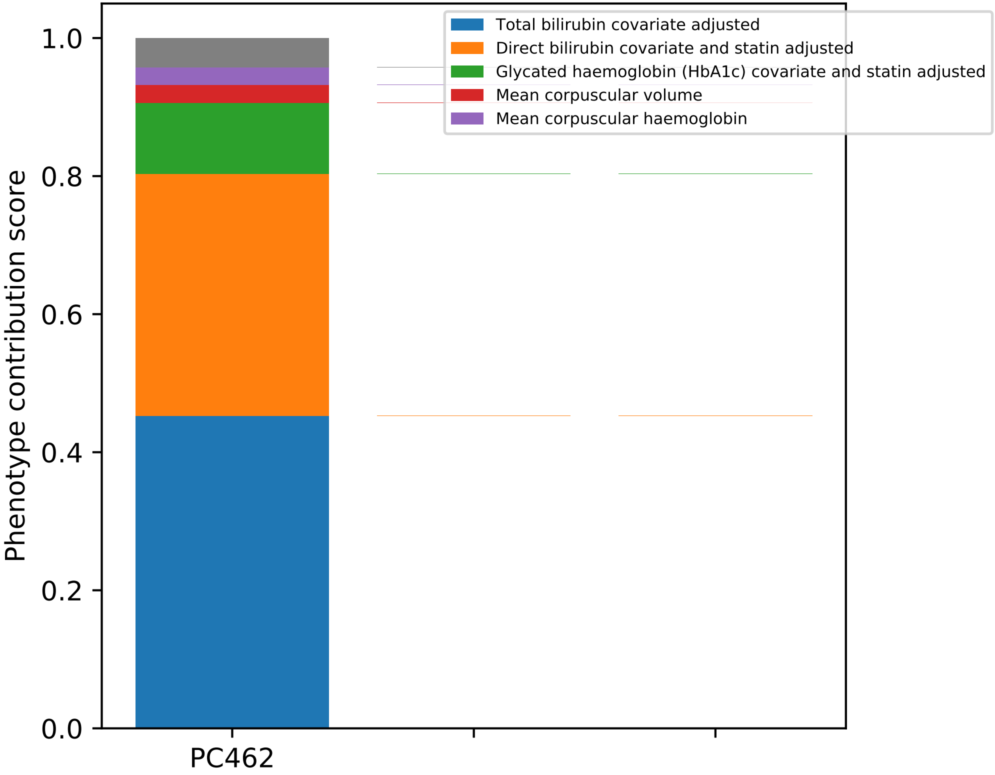

Total bilirubin covariate adjusted
Direct bilirubin covariate and statin adjusted
Glycated haemoglobin (HbA1c) covariate and statin adjusted
Mean corpuscular volume
Mean corpuscular haemoglobin
Phenotype contribution score
0.0
0.2
0.4
0.6
0.8
1.0
PC462

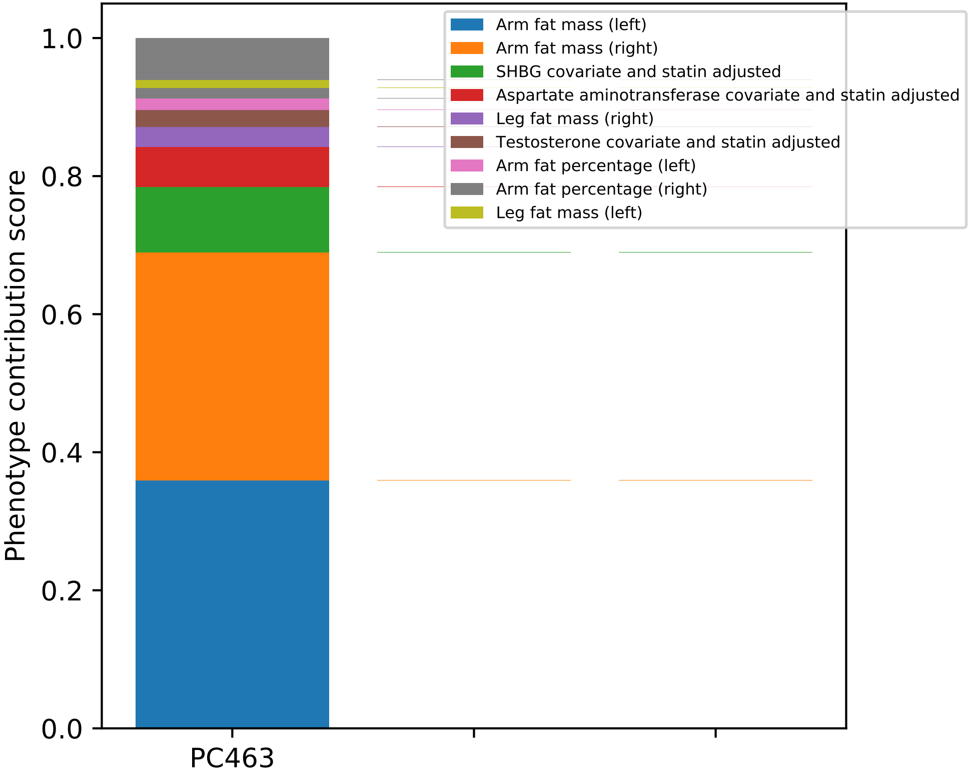
Arm fat mass (left)
Arm fat mass (right)
SHBG covariate and statin adjusted
Aspartate aminotransferase covariate and statin adjusted
Leg fat mass (right)
Testosterone covariate and statin adjusted
Arm fat percentage (left)
Arm fat percentage (right)
Leg fat mass (left)
Phenotype contribution score
1.0
0.8
0.6
0.4
0.2
0.0
PC463

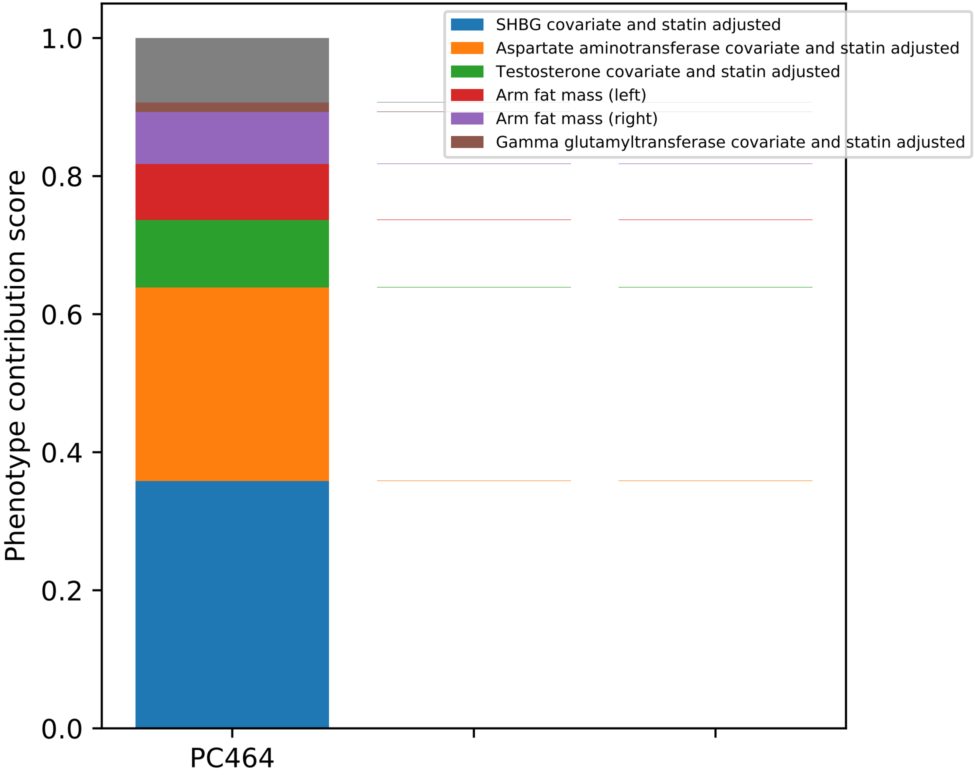

SHBG covariate and statin adjusted
Aspartate aminotransferase covariate and statin adjusted
Testosterone covariate and statin adjusted
Arm fat mass (left)
Arm fat mass (right)
Gamma glutamyltransferase covariate and statin adjusted
Phenotype contribution score
0.0
0.2
0.4
0.6
0.8
1.0
PC464

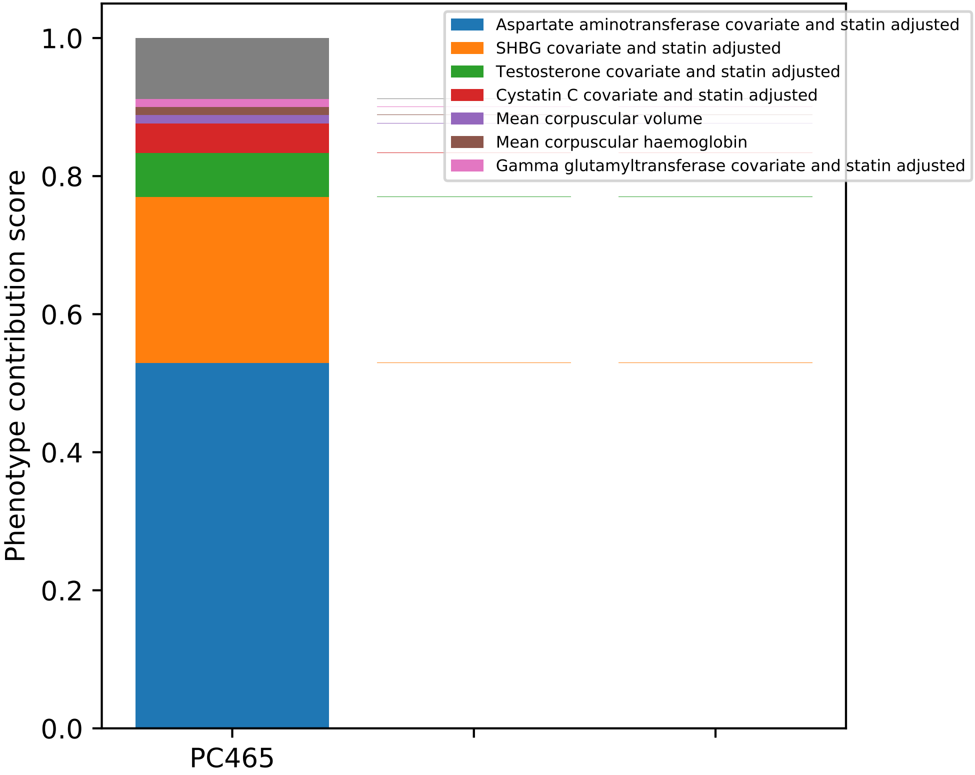

Aspartate aminotransferase covariate and statin adjusted
SHBG covariate and statin adjusted
Testosterone covariate and statin adjusted
Cystatin C covariate and statin adjusted
Mean corpuscular volume
Mean corpuscular haemoglobin
Gamma glutamyltransferase covariate and statin adjusted
Phenotype contribution score
0.0
0.2
0.4
0.6
0.8
1.0
PC465

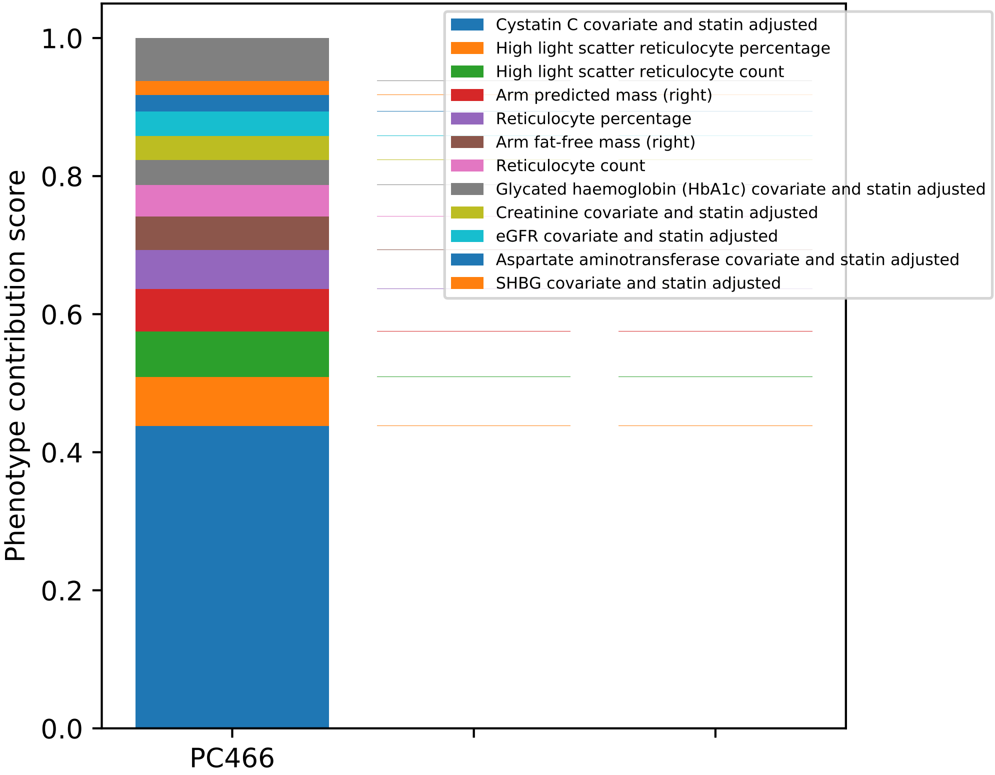

Phenotype contribution score
0.0
0.2
0.4
0.6
0.8
1.0
PC466
Cystatin C covariate and statin adjusted
High light scatter reticulocyte percentage
High light scatter reticulocyte count
Arm predicted mass (right)
Reticulocyte percentage
Arm fat-free mass (right)
Reticulocyte count
Glycated haemoglobin (HbA1c) covariate and statin adjusted
Creatinine covariate and statin adjusted
eGFR covariate and statin adjusted
Aspartate aminotransferase covariate and statin adjusted
SHBG covariate and statin adjusted

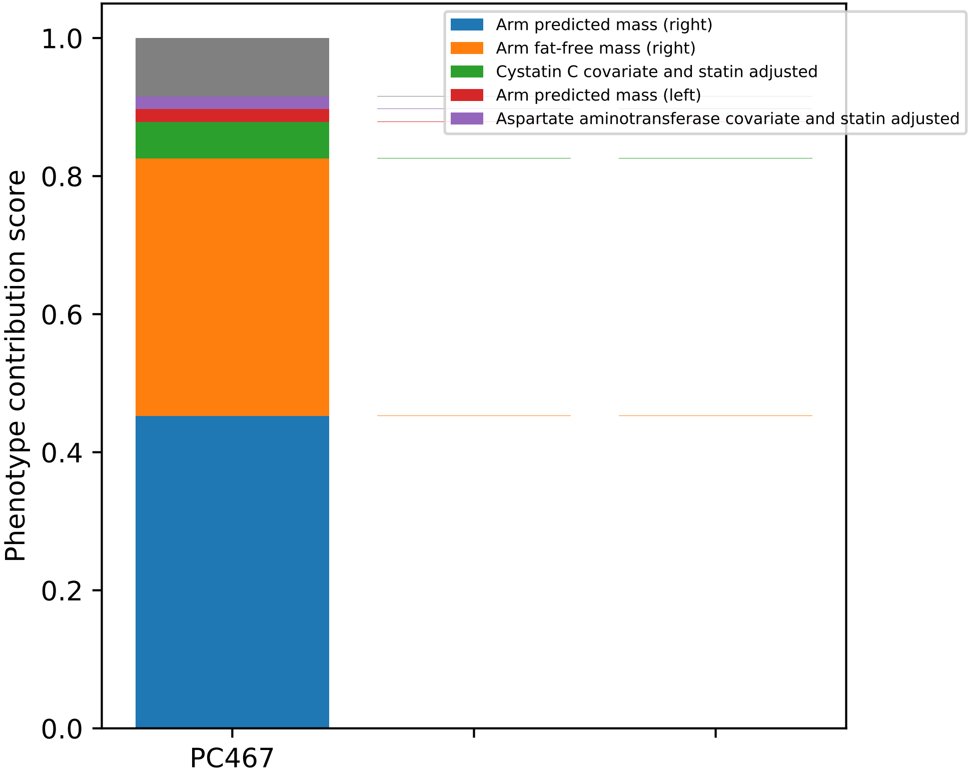
Arm predicted mass (right)
Arm fat-free mass (right)
Cystatin C covariate and statin adjusted
Arm predicted mass (left)
Aspartate aminotransferase covariate and statin adjusted
Phenotype contribution score
0.0
0.2
0.4
0.6
0.8
1.0
PC467

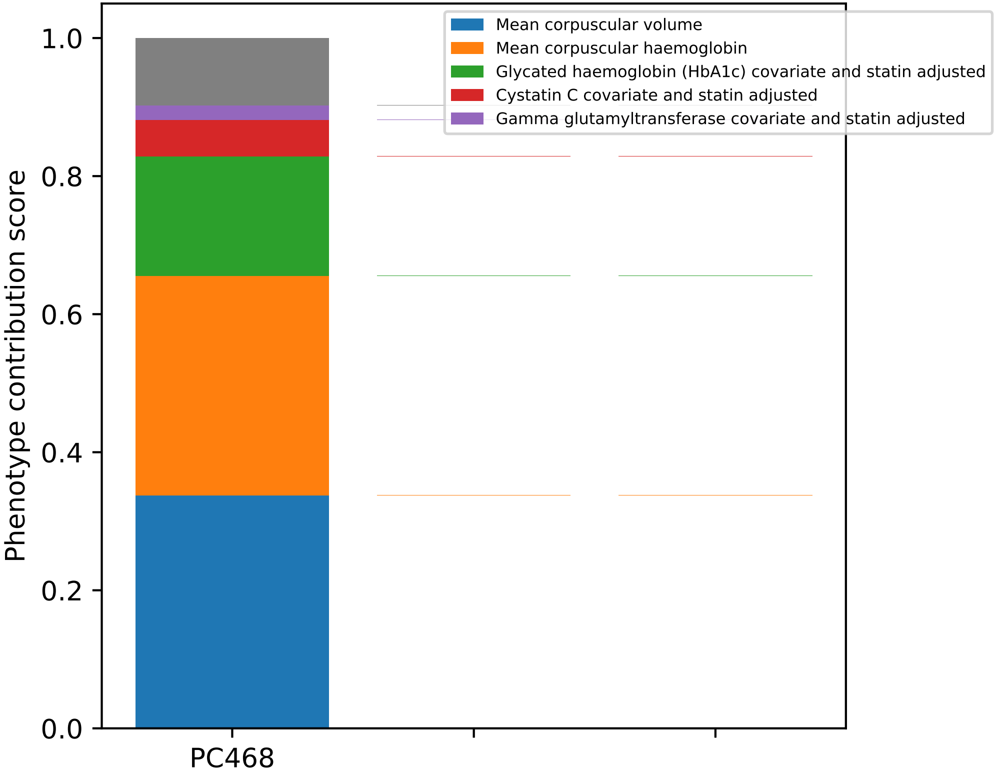
Mean corpuscular volume
Mean corpuscular haemoglobin
Glycated haemoglobin (HbA1c) covariate and statin adjusted
Cystatin C covariate and statin adjusted
Gamma glutamyltransferase covariate and statin adjusted
Phenotype contribution score
1.0
0.8
0.6
0.4
0.2
0.0
PC468

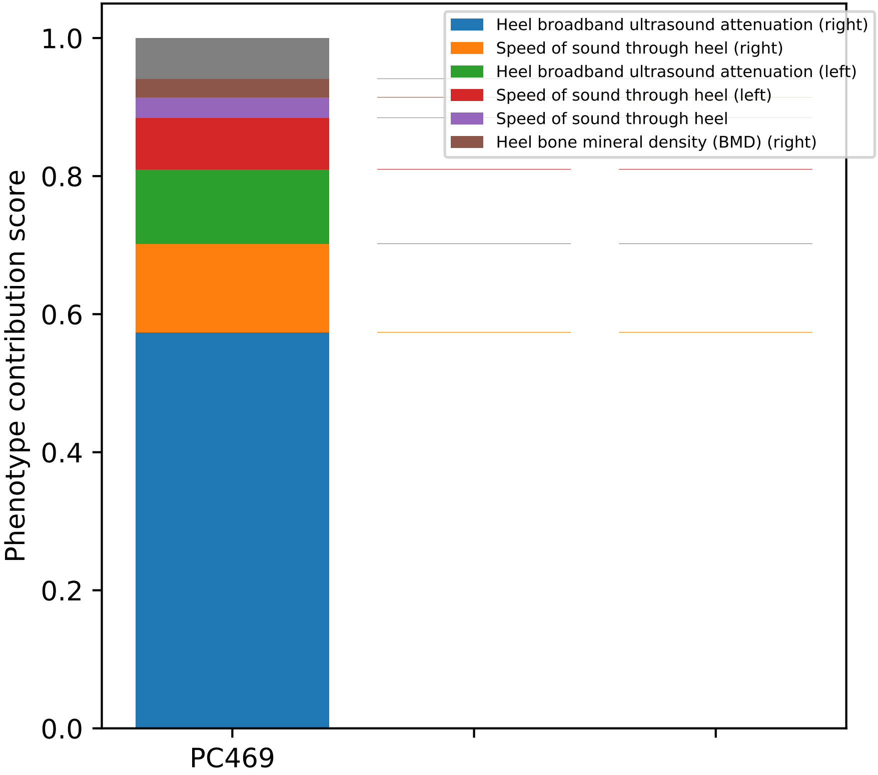

Heel broadband ultrasound attenuation (right)
Speed of sound through heel (right)
Heel broadband ultrasound attenuation (left)
Speed of sound through heel (left)
Speed of sound through heel
Heel bone mineral density (BMD) (right)
Phenotype contribution score
1.0
0.8
0.6
0.4
0.2
0.0
PC469

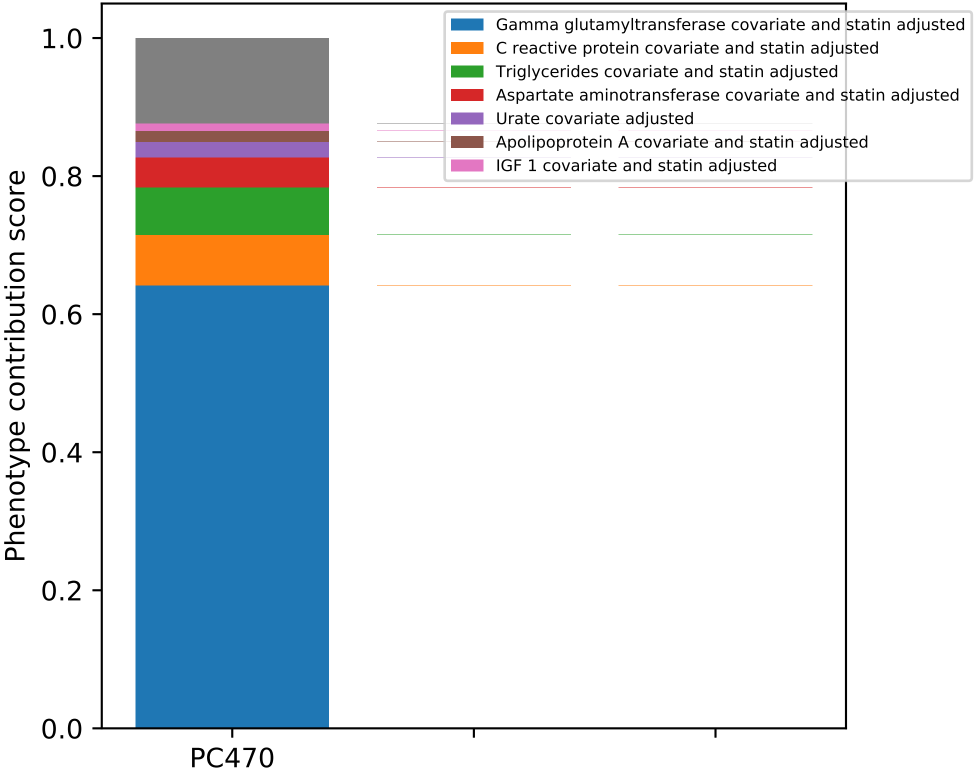

Gamma glutamyltransferase covariate and statin adjusted
C reactive protein covariate and statin adjusted
Triglycerides covariate and statin adjusted
Aspartate aminotransferase covariate and statin adjusted
Urate covariate adjusted
Apolipoprotein A covariate and statin adjusted
IGF 1 covariate and statin adjusted
Phenotype contribution score
1.0
0.8
0.6
0.4
0.2
0.0
PC470

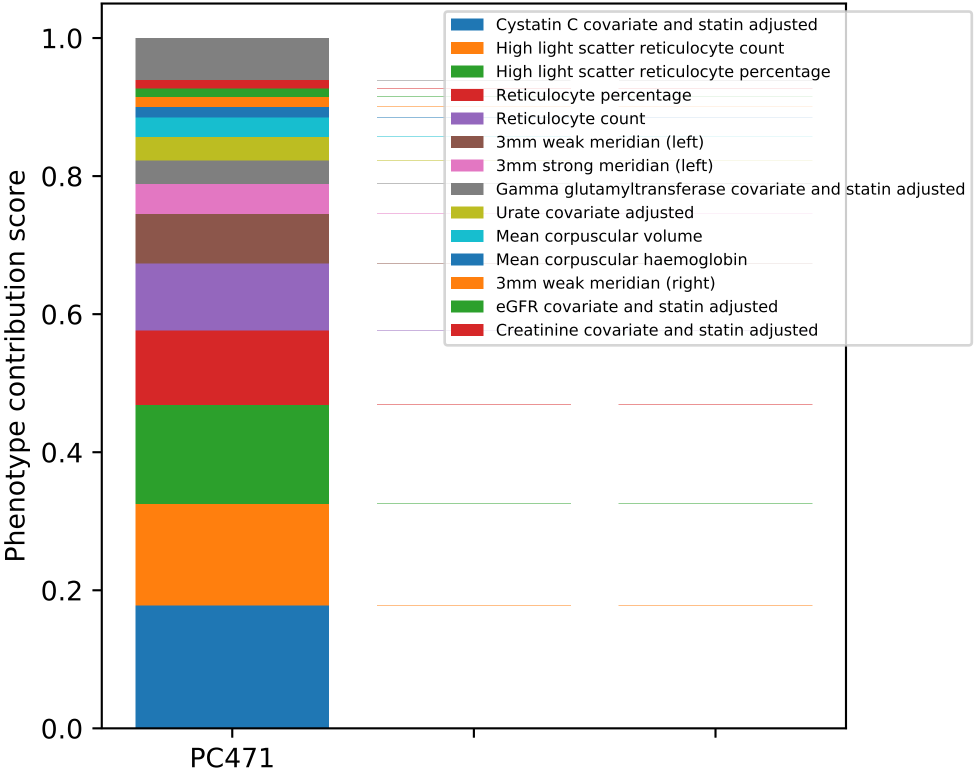
Cystatin C covariate and statin adjusted
High light scatter reticulocyte count
High light scatter reticulocyte percentage
Reticulocyte percentage
Reticulocyte count
3mm weak meridian (left)
3mm strong meridian (left)
Gamma glutamyltransferase covariate and statin adjusted
Urate covariate adjusted
Mean corpuscular volume
Mean corpuscular haemoglobin
3mm weak meridian (right)
eGFR covariate and statin adjusted
Creatinine covariate and statin adjusted
Phenotype contribution score
1.0
0.8
0.6
0.4
0.2
0.0
PC471

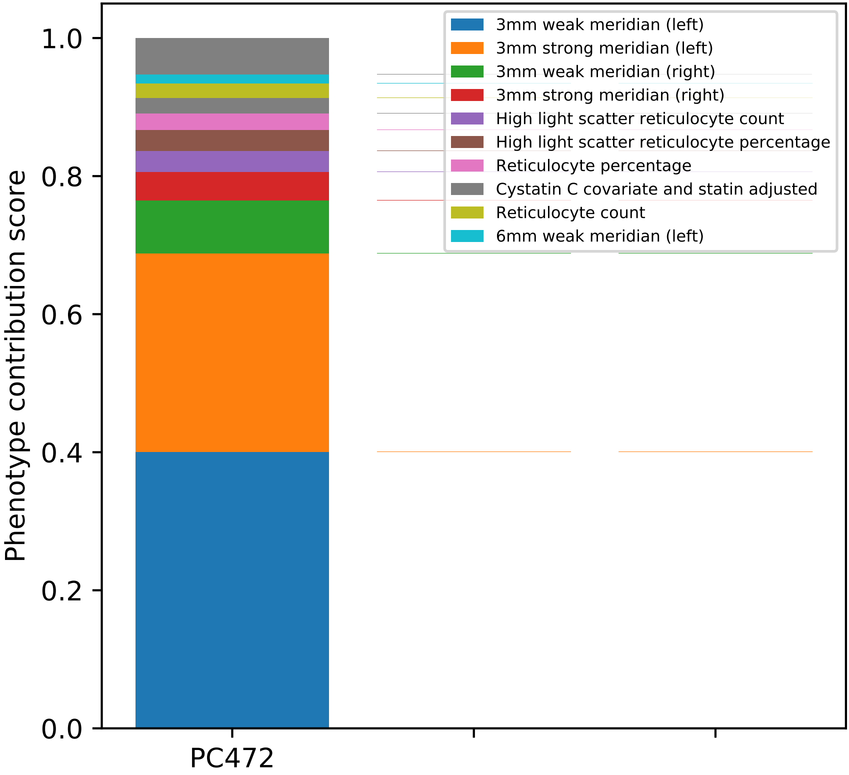

Phenotype contribution score
1.0
0.8
0.6
0.4
0.2
0.0
PC472
3mm weak meridian (left)
3mm strong meridian (left)
3mm weak meridian (right)
3mm strong meridian (right)
High light scatter reticulocyte count
High light scatter reticulocyte percentage
Reticulocyte percentage
Cystatin C covariate and statin adjusted
Reticulocyte count
6mm weak meridian (left)

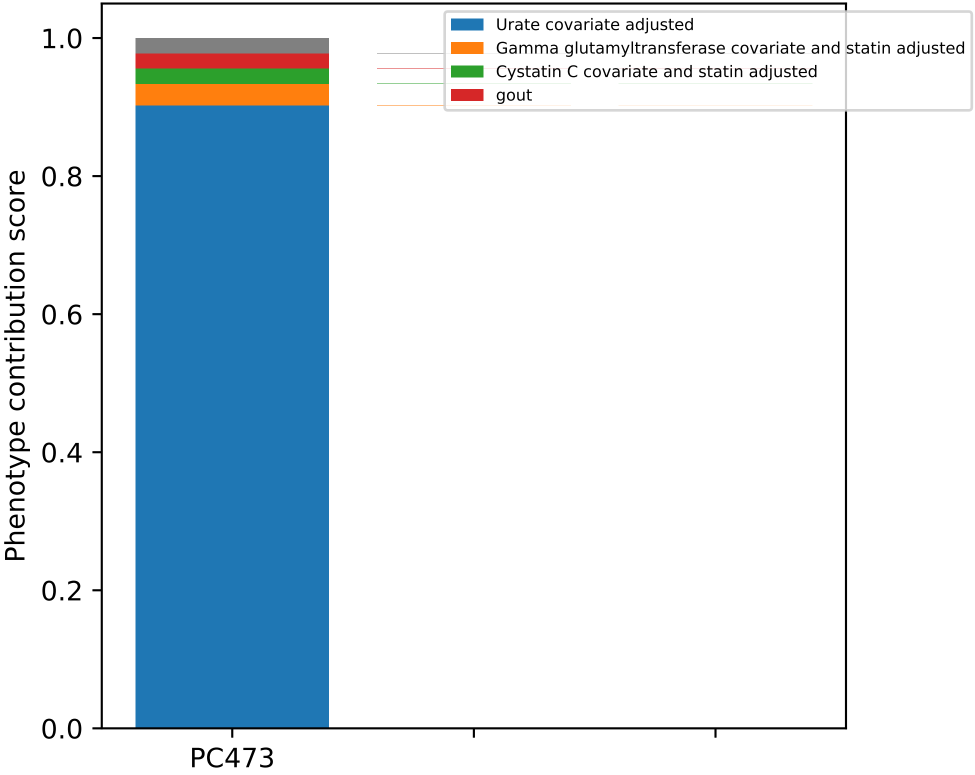
Urate covariate adjusted
Gamma glutamyltransferase covariate and statin adjusted
Cystatin C covariate and statin adjusted
gout
Phenotype contribution score
1.0
0.8
0.6
0.4
0.2
0.0
PC473

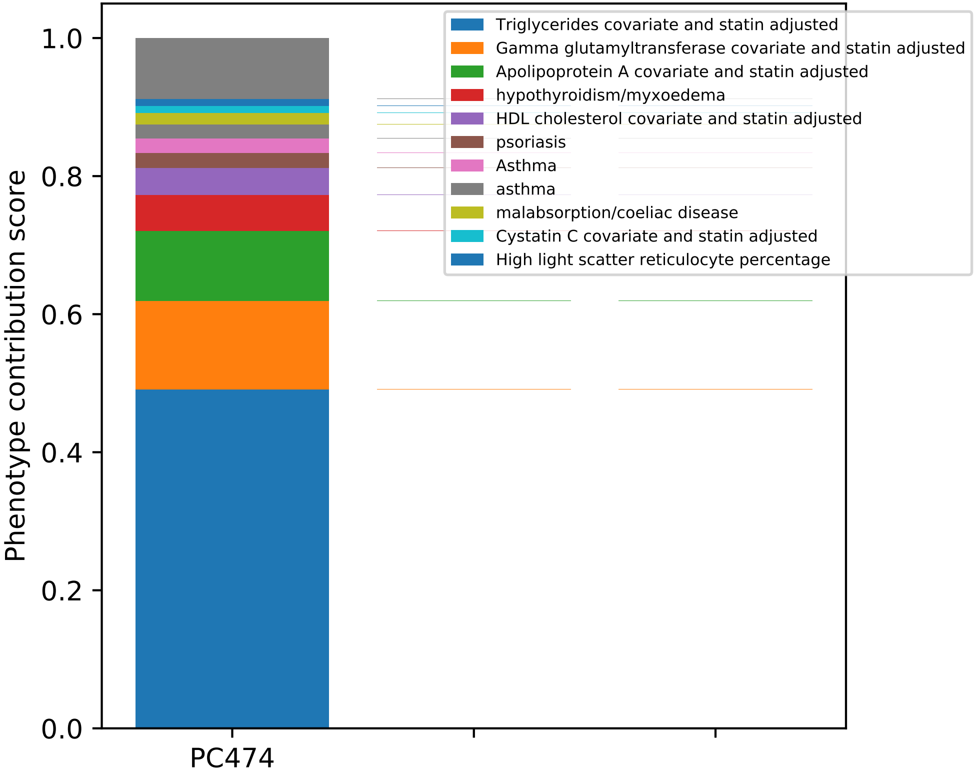
Phenotype contribution score
0.0
0.2
0.4
0.6
0.8
1.0
PC474
Triglycerides covariate and statin adjusted
Gamma glutamyltransferase covariate and statin adjusted
Apolipoprotein A covariate and statin adjusted
hypothyroidism/myxoedema
HDL cholesterol covariate and statin adjusted
psoriasis
Asthma
asthma
malabsorption/coeliac disease
Cystatin C covariate and statin adjusted
High light scatter reticulocyte percentage

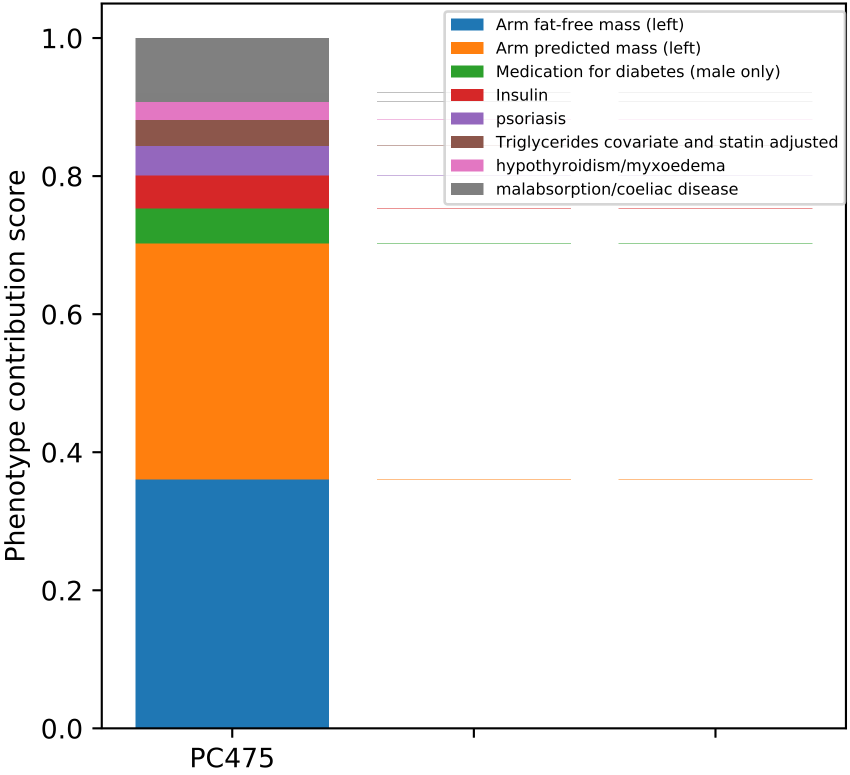
Phenotype contribution score
0.0
0.2
0.4
0.6
0.8
1.0
PC475
Arm fat-free mass (left)
Arm predicted mass (left)
Medication for diabetes (male only)
Insulin
psoriasis
Triglycerides covariate and statin adjusted
hypothyroidism/myxoedema
malabsorption/coeliac disease

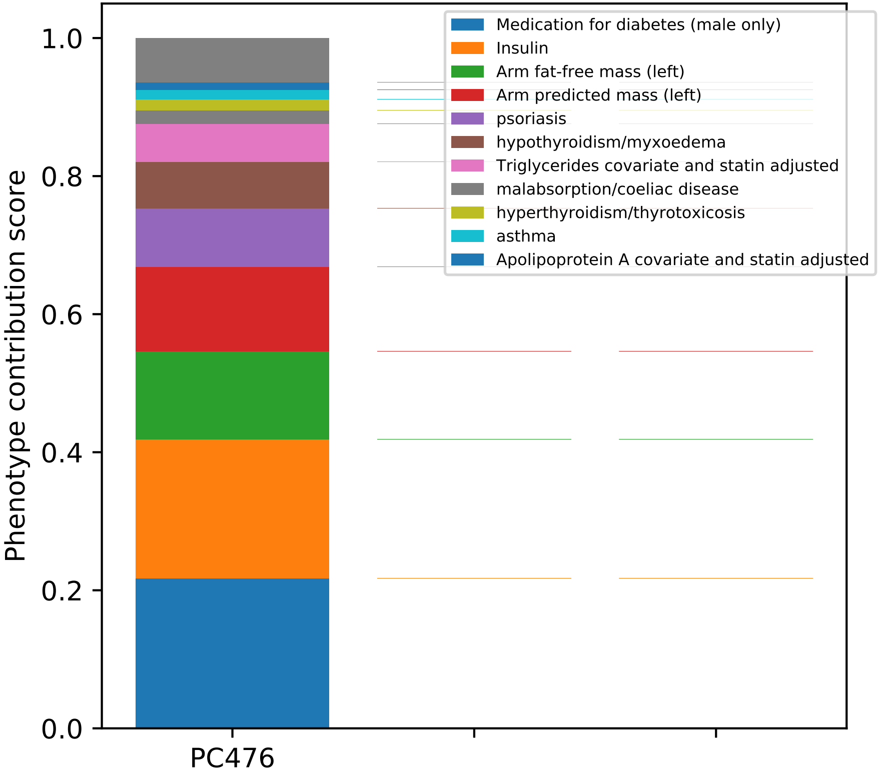
Medication for diabetes (male only)
Insulin
Arm fat-free mass (left)
Arm predicted mass (left)
psoriasis
hypothyroidism/myxoedema
Triglycerides covariate and statin adjusted
malabsorption/coeliac disease
hyperthyroidism/thyrotoxicosis
asthma
Apolipoprotein A covariate and statin adjusted
Phenotype contribution score
1.0
0.8
0.6
0.4
0.2
0.0
PC476

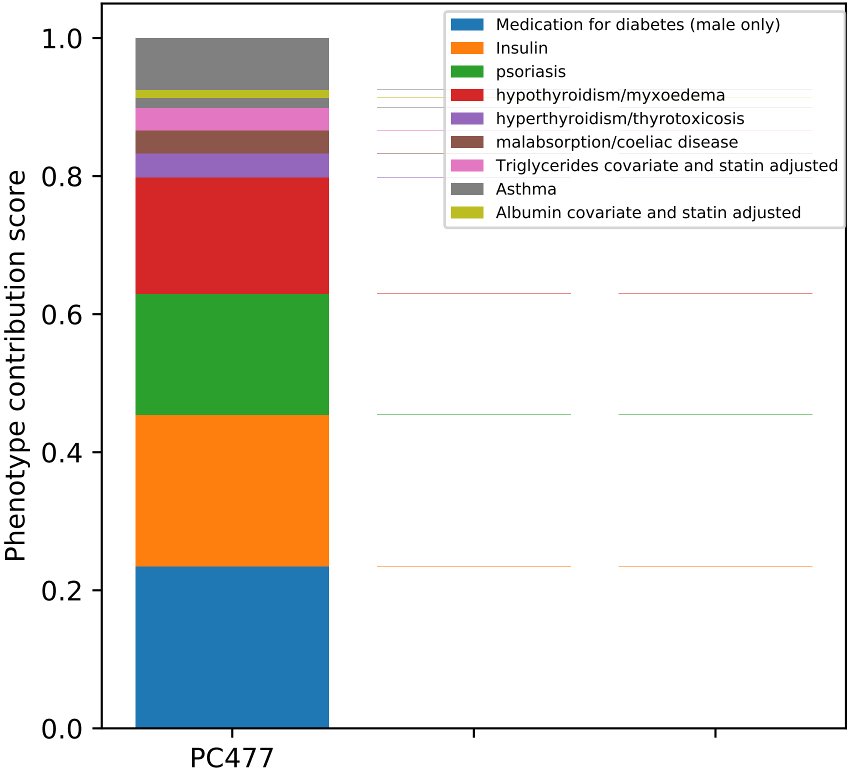

Medication for diabetes (male only)
Insulin
psoriasis
hypothyroidism/myxoedema
hyperthyroidism/thyrotoxicosis
malabsorption/coeliac disease
Triglycerides covariate and statin adjusted
Asthma
Albumin covariate and statin adjusted
Phenotype contribution score
1.0
0.8
0.6
0.4
0.2
0.0
PC477

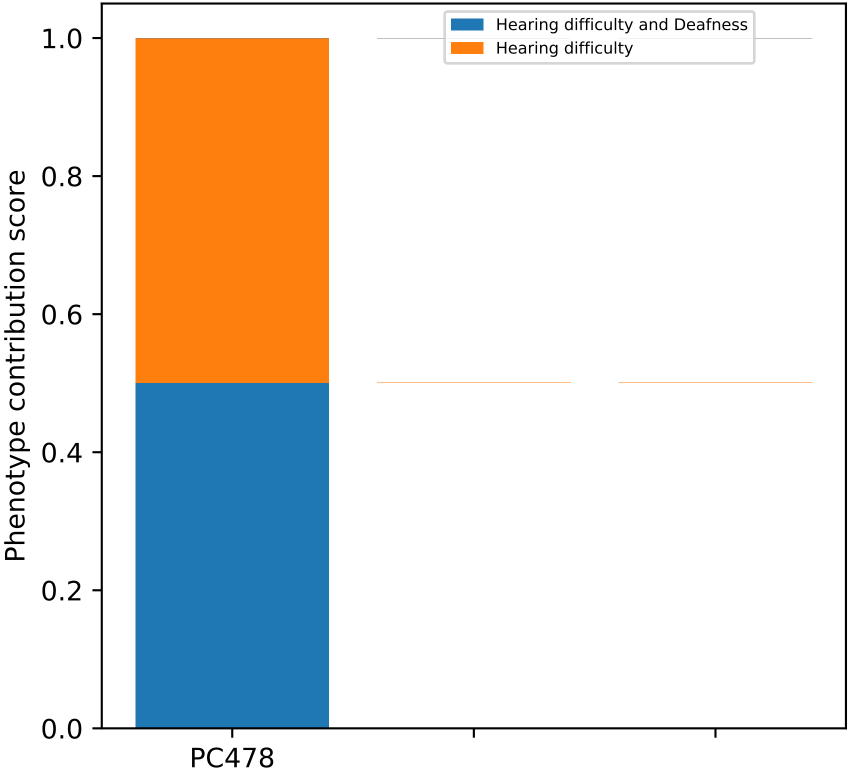
Hearing difficulty and Deafness
Hearing difficulty
1.0
0.8
0.6
0.4
0.2
0.0
Phenotype contribution score
PC478

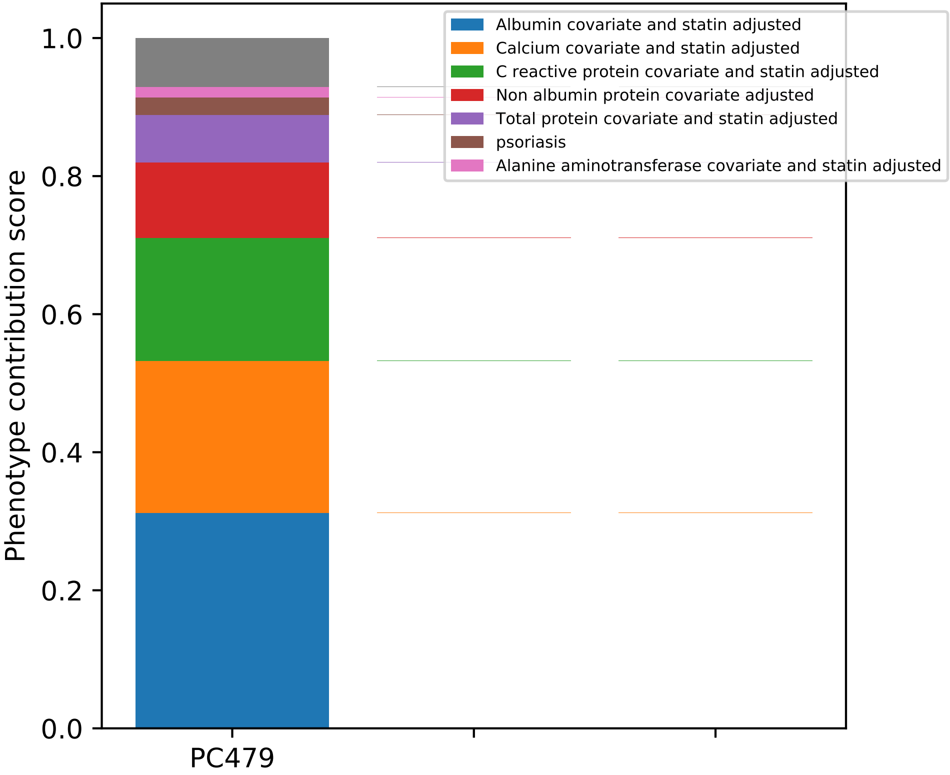
Albumin covariate and statin adjusted
Calcium covariate and statin adjusted
C reactive protein covariate and statin adjusted
Non albumin protein covariate adjusted
Total protein covariate and statin adjusted
psoriasis
Alanine aminotransferase covariate and statin adjusted
Phenotype contribution score
0.0
0.2
0.4
0.6
0.8
1.0
PC479

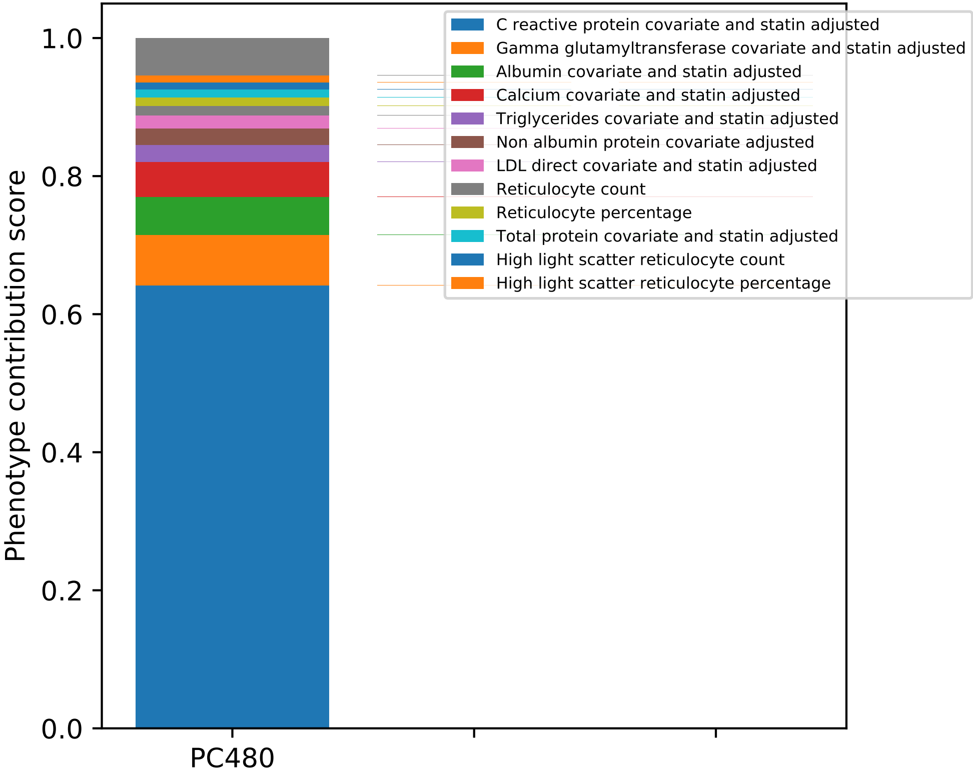

C reactive protein covariate and statin adjusted
Gamma glutamyltransferase covariate and statin adjusted
Albumin covariate and statin adjusted
Calcium covariate and statin adjusted
Triglycerides covariate and statin adjusted
Non albumin protein covariate adjusted
LDL direct covariate and statin adjusted
Reticulocyte count
Reticulocyte percentage
Total protein covariate and statin adjusted
High light scatter reticulocyte count
High light scatter reticulocyte percentage
Phenotype contribution score
1.0
0.8
0.6
0.4
0.2
0.0
PC480

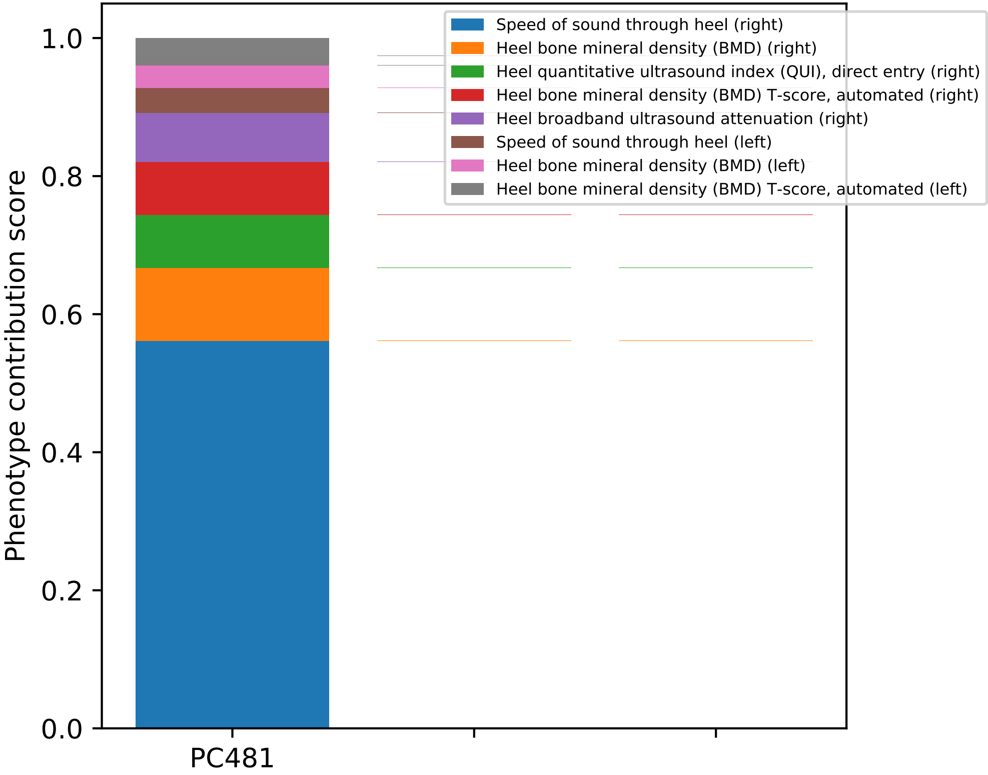

Speed of sound through heel (right)
Heel bone mineral density (BMD) (right)
Heel quantitative ultrasound index (QUI), direct entry (right)
Heel bone mineral density (BMD) T-score, automated (right)
Heel broadband ultrasound attenuation (right)
Speed of sound through heel (left)
Heel bone mineral density (BMD) (left)
Heel bone mineral density (BMD) T-score, automated (left)
Phenotype contribution score
1.0
0.8
0.6
0.4
0.2
0.0
PC481

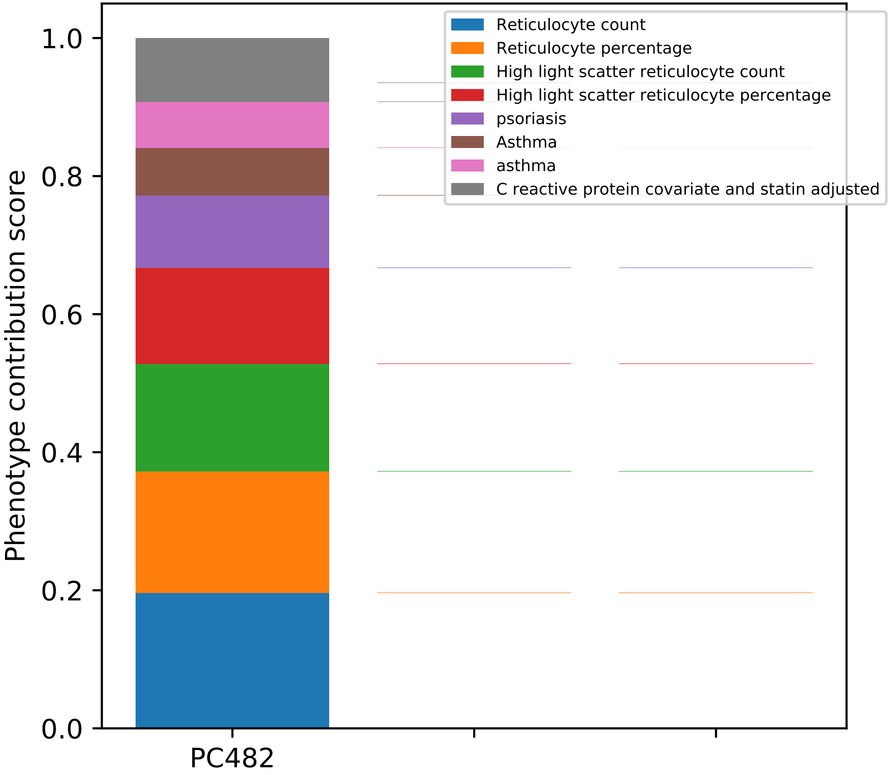
Reticulocyte count
Reticulocyte percentage
High light scatter reticulocyte count
High light scatter reticulocyte percentage
psoriasis
Asthma
asthma
C reactive protein covariate and statin adjusted
Phenotype contribution score
1.0
0.8
0.6
0.4
0.2
0.0
PC482

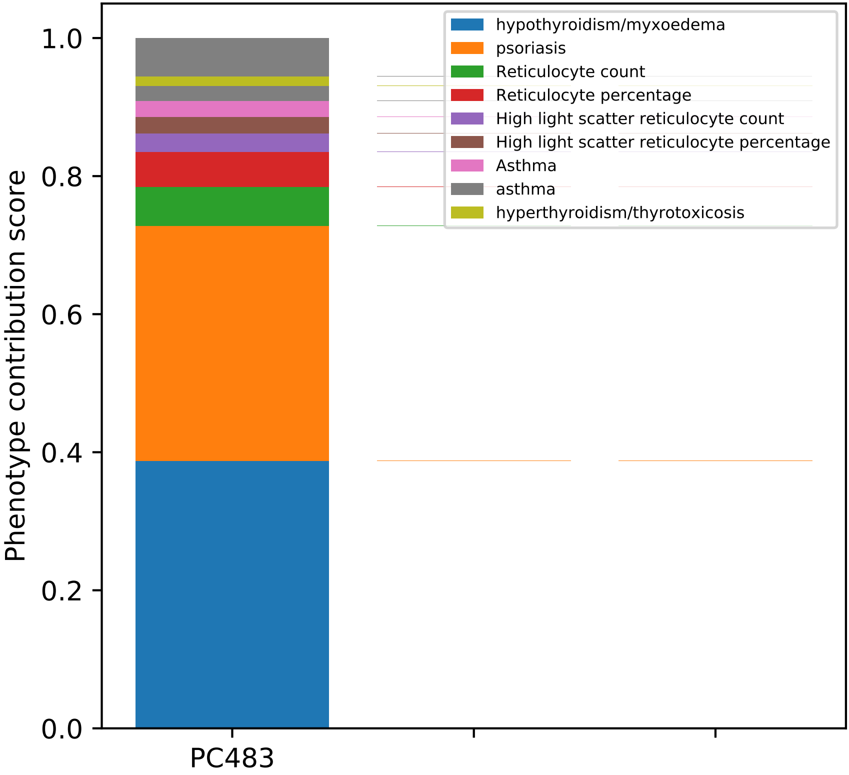

hypothyroidism/myxoedema
psoriasis
Reticulocyte count
Reticulocyte percentage
High light scatter reticulocyte count
High light scatter reticulocyte percentage
Asthma
asthma
hyperthyroidism/thyrotoxicosis
Phenotype contribution score
0.0
0.2
0.4
0.6
0.8
1.0
PC483

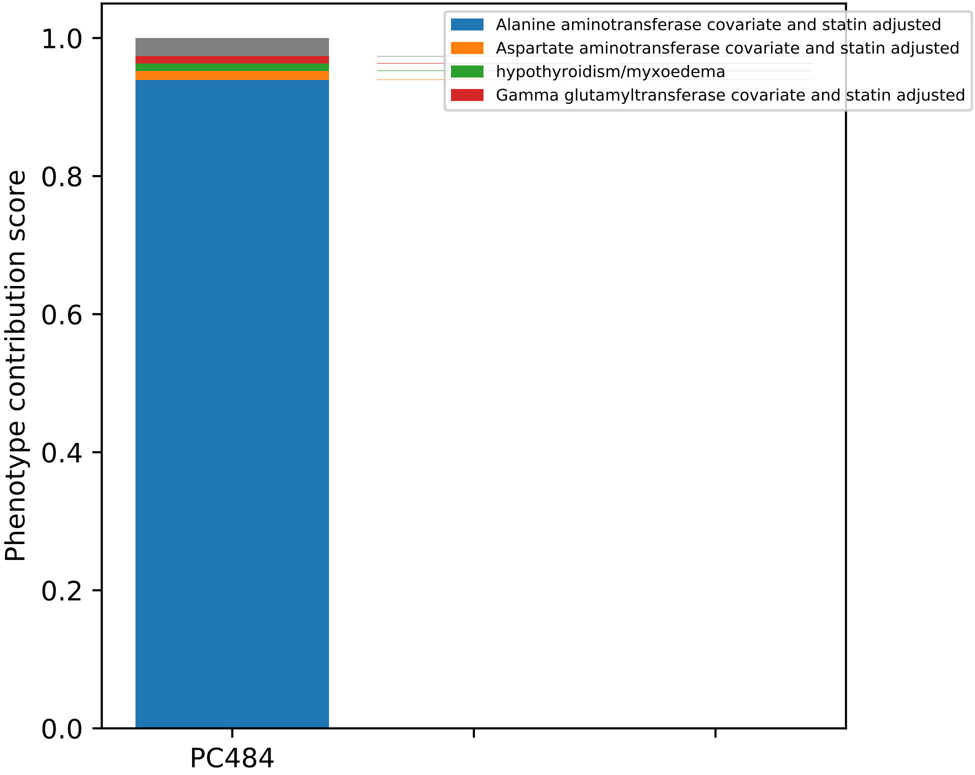
Alanine aminotransferase covariate and statin adjusted
Aspartate aminotransferase covariate and statin adjusted
hypothyroidism/myxoedema
Gamma glutamyltransferase covariate and statin adjusted
Phenotype contribution score
1.0
0.8
0.6
0.4
0.2
0.0
PC484

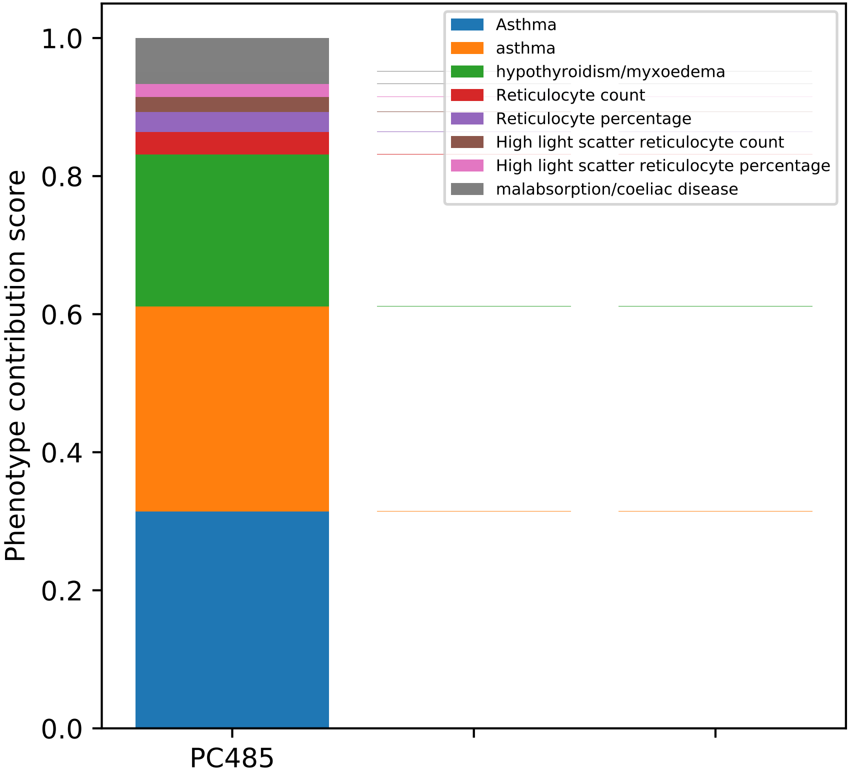
Phenotype contribution score
1.0
0.8
0.6
0.4
0.2
0.0
PC485
Asthma
asthma
hypothyroidism/myxoedema
Reticulocyte count
Reticulocyte percentage
High light scatter reticulocyte count
High light scatter reticulocyte percentage
malabsorption/coeliac disease

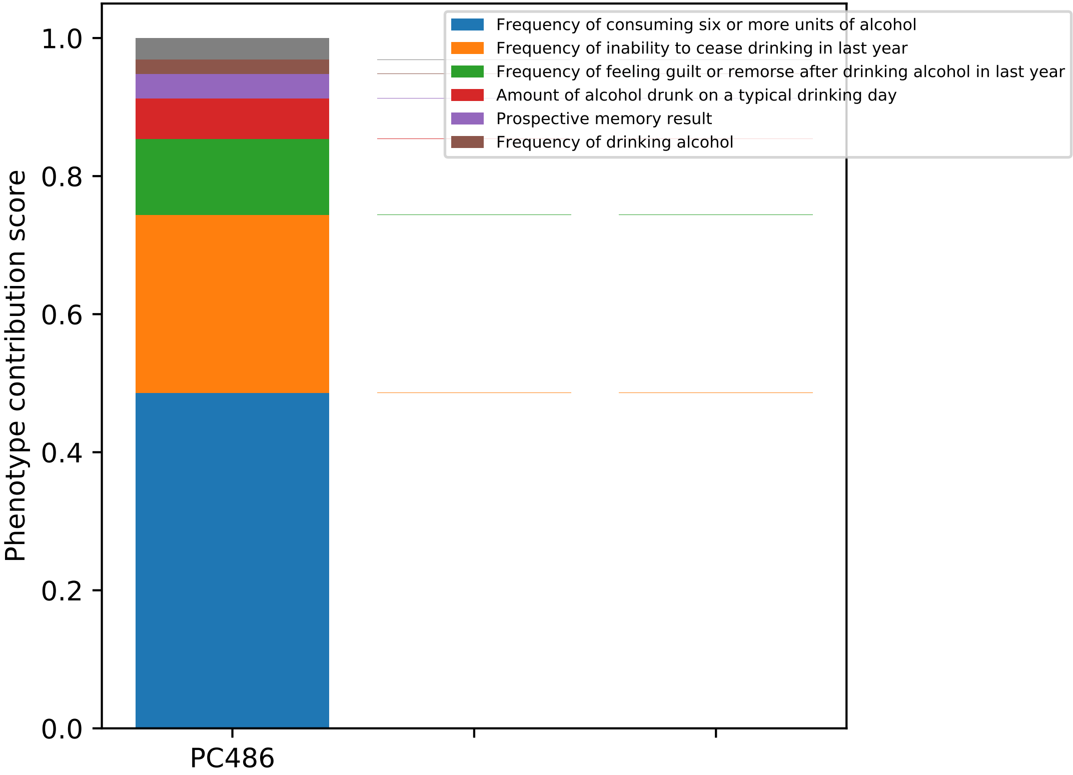
Frequency of consuming six or more units of alcohol
Frequency of inability to cease drinking in last year
Frequency of feeling guilt or remorse after drinking alcohol in last year
Amount of alcohol drunk on a typical drinking day
Prospective memory result
Frequency of drinking alcohol
Phenotype contribution score
1.0
0.8
0.6
0.4
0.2
0.0
PC486

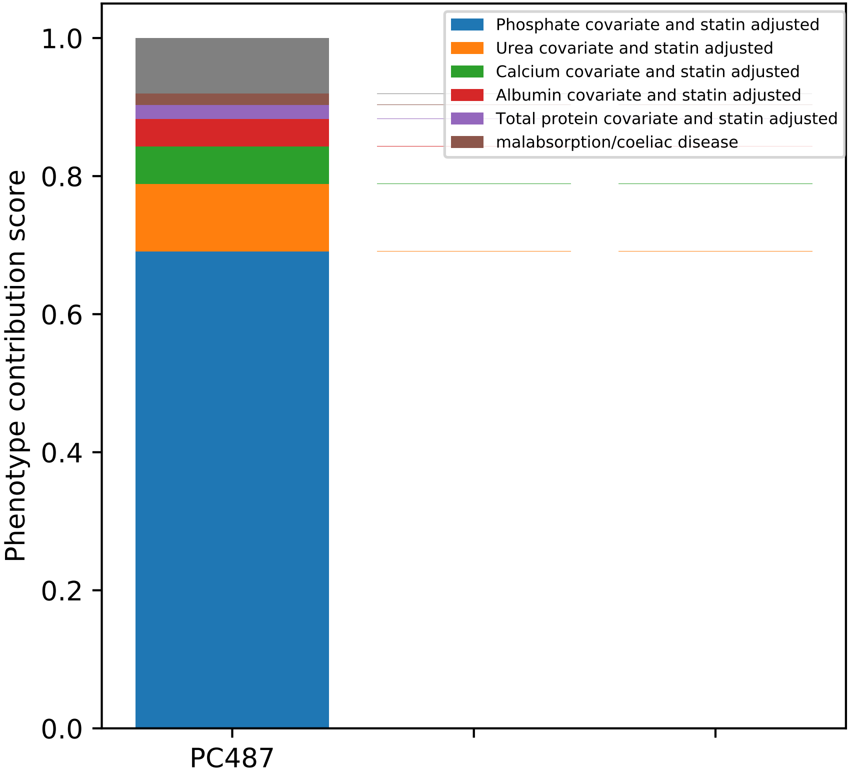

Phosphate covariate and statin adjusted
Urea covariate and statin adjusted
Calcium covariate and statin adjusted
Albumin covariate and statin adjusted
Total protein covariate and statin adjusted
malabsorption/coeliac disease
Phenotype contribution score
0.0
0.2
0.4
0.6
0.8
1.0
PC487

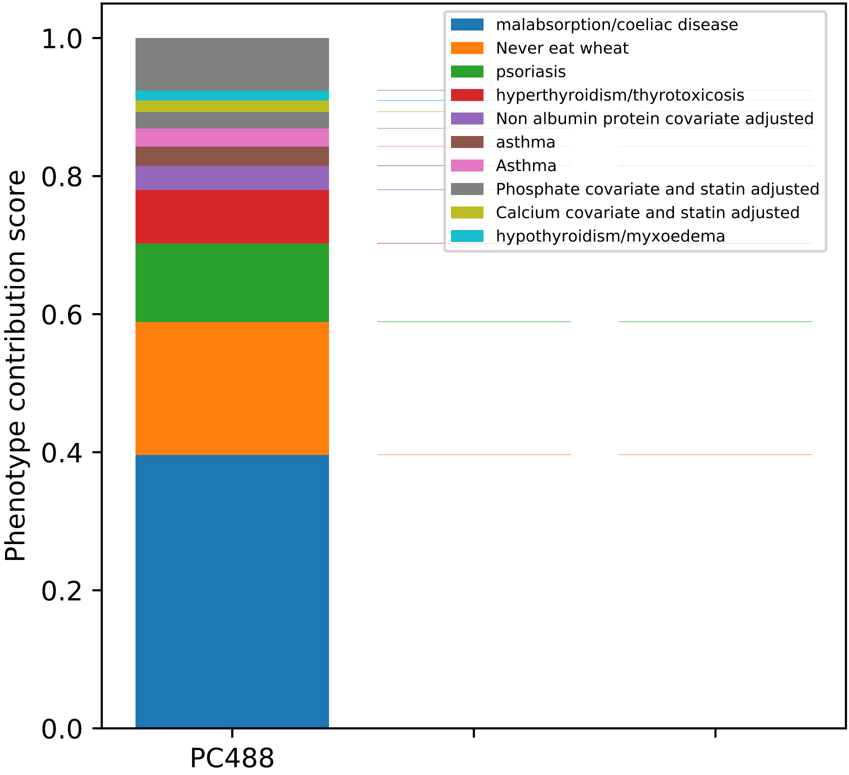

Phenotype contribution score
0.0
0.2
0.4
0.6
0.8
1.0
PC488
malabsorption/coeliac disease
Never eat wheat
psoriasis
hyperthyroidism/thyrotoxicosis
Non albumin protein covariate adjusted
asthma
Asthma
Phosphate covariate and statin adjusted
Calcium covariate and statin adjusted
hypothyroidism/myxoedema

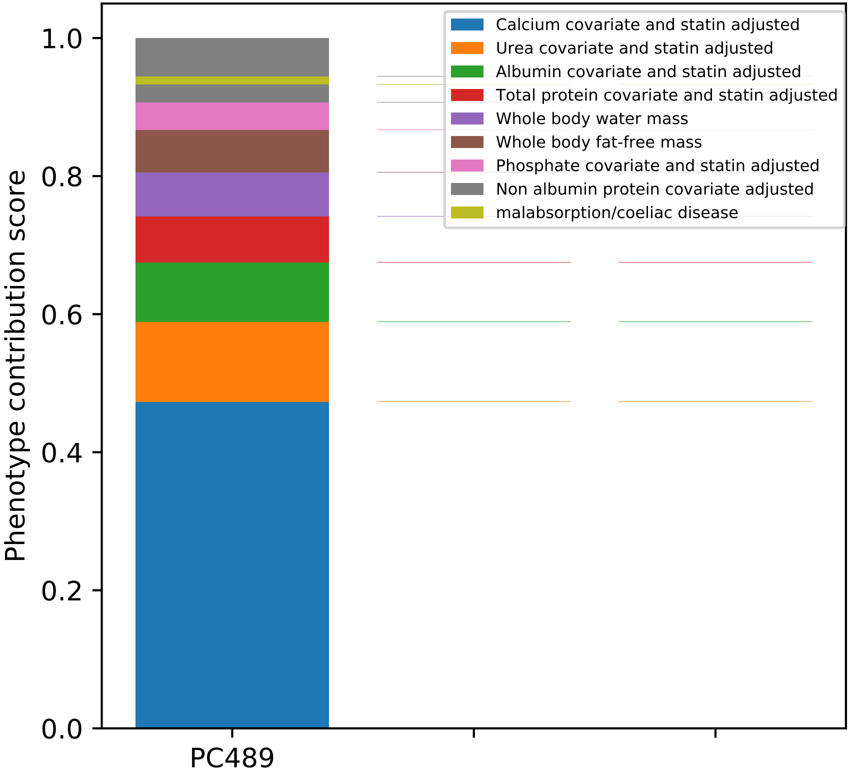

Calcium covariate and statin adjusted
Urea covariate and statin adjusted
Albumin covariate and statin adjusted
Total protein covariate and statin adjusted
Whole body water mass
Whole body fat-free mass
Phosphate covariate and statin adjusted
Non albumin protein covariate adjusted
malabsorption/coeliac disease
Phenotype contribution score
1.0
0.8
0.6
0.4
0.2
0.0
PC489

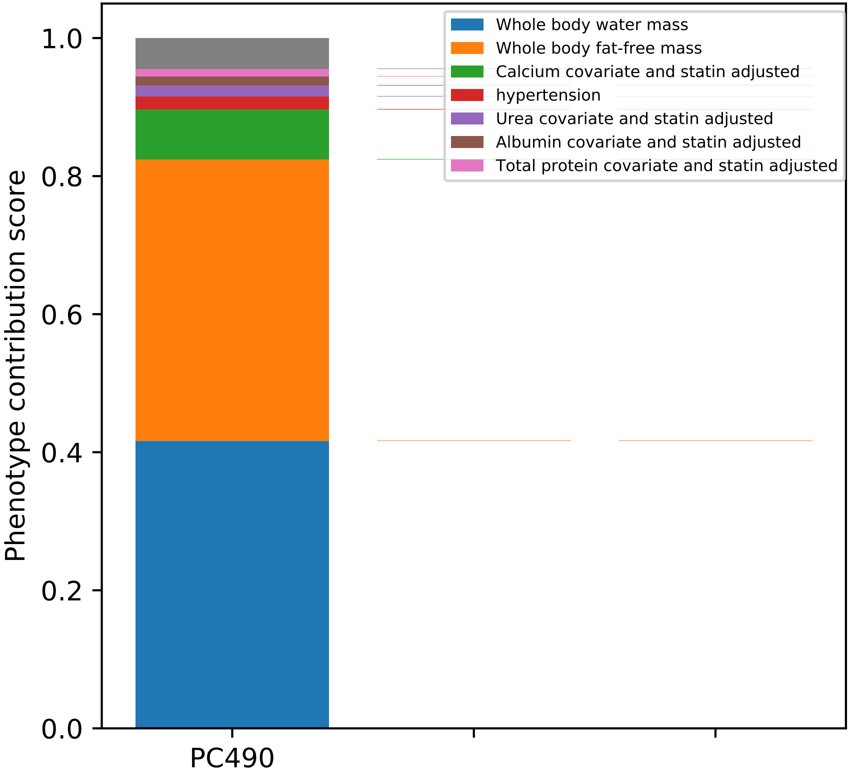

Whole body water mass
Whole body fat-free mass
Calcium covariate and statin adjusted
hypertension
Urea covariate and statin adjusted
Albumin covariate and statin adjusted
Total protein covariate and statin adjusted
Phenotype contribution score
1.0
0.8
0.6
0.4
0.2
0.0
PC490

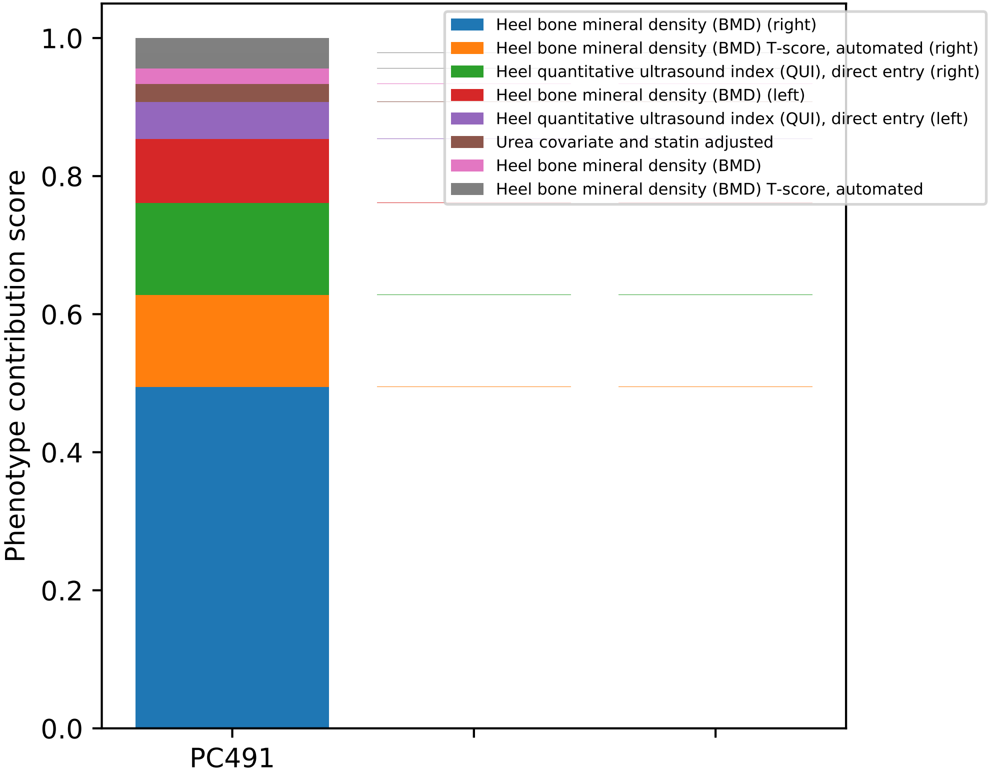
Heel bone mineral density (BMD) (right)
Heel bone mineral density (BMD) T-score, automated (right)
Heel quantitative ultrasound index (QUI), direct entry (right)
Heel bone mineral density (BMD) (left)
Heel quantitative ultrasound index (QUI), direct entry (left)
Urea covariate and statin adjusted
Heel bone mineral density (BMD)
Heel bone mineral density (BMD) T-score, automated
Phenotype contribution score
0.0
0.2
0.4
0.6
0.8
1.0
PC491

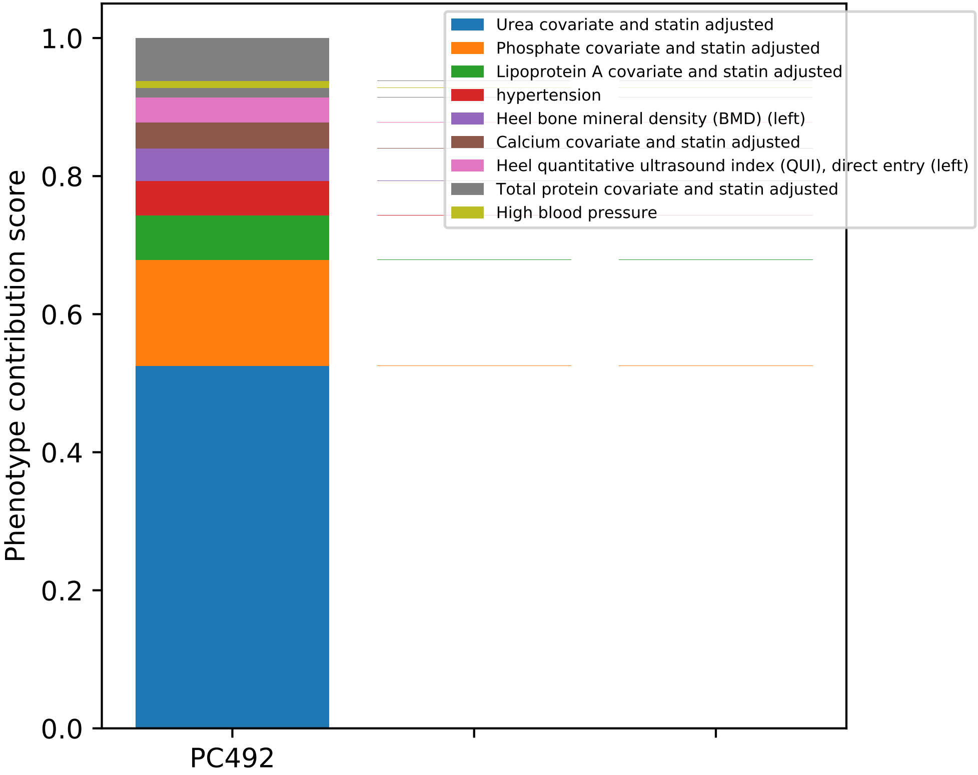

Phenotype contribution score
1.0
0.8
0.6
0.4
0.2
0.0
PC492
Urea covariate and statin adjusted
Phosphate covariate and statin adjusted
Lipoprotein A covariate and statin adjusted
hypertension
Heel bone mineral density (BMD) (left)
Calcium covariate and statin adjusted
Heel quantitative ultrasound index (QUI), direct entry (left)
Total protein covariate and statin adjusted
High blood pressure

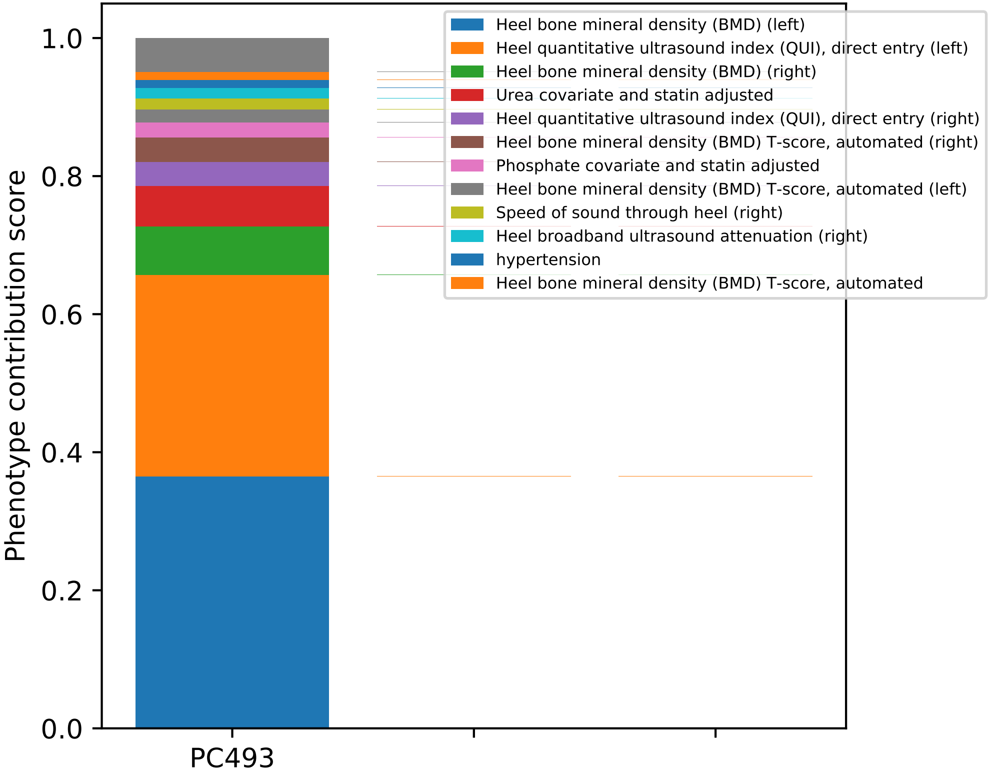
Phenotype contribution score
1.0
0.8
0.6
0.4
0.2
0.0
PC493
Heel bone mineral density (BMD) (left)
Heel quantitative ultrasound index (QUI), direct entry (left)
Heel bone mineral density (BMD) (right)
Urea covariate and statin adjusted
Heel quantitative ultrasound index (QUI), direct entry (right)
Heel bone mineral density (BMD) T-score, automated (right)
Phosphate covariate and statin adjusted
Heel bone mineral density (BMD) T-score, automated (left)
Speed of sound through heel (right)
Heel broadband ultrasound attenuation (right)
hypertension
Heel bone mineral density (BMD) T-score, automated

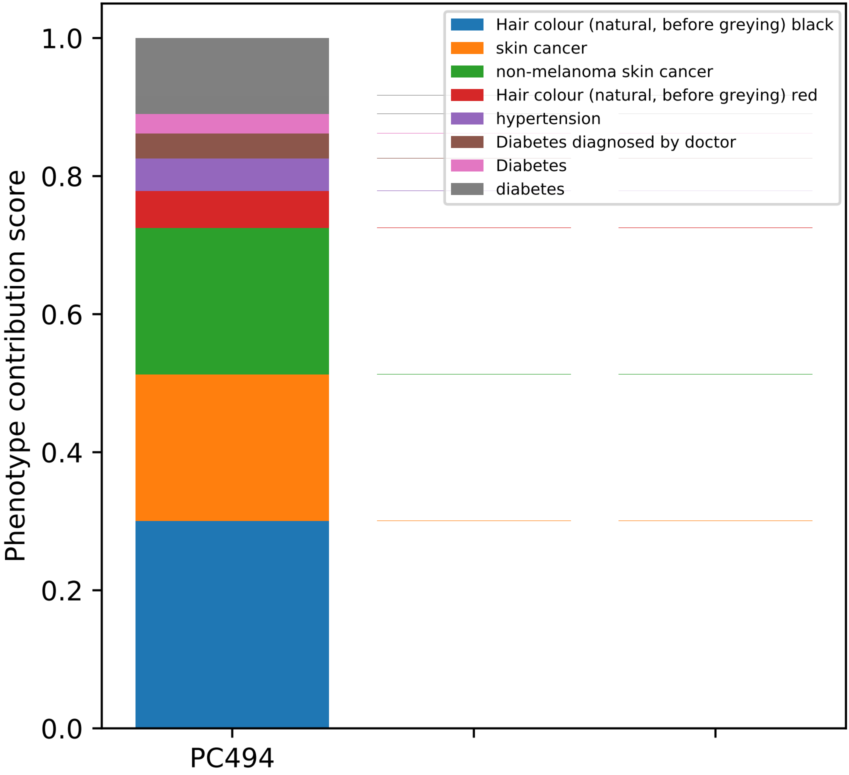
Hair colour (natural, before greying) black
skin cancer
non-melanoma skin cancer
Hair colour (natural, before greying) red
hypertension
Diabetes diagnosed by doctor
Diabetes
diabetes
Phenotype contribution score
1.0
0.8
0.6
0.4
0.2
0.0
PC494

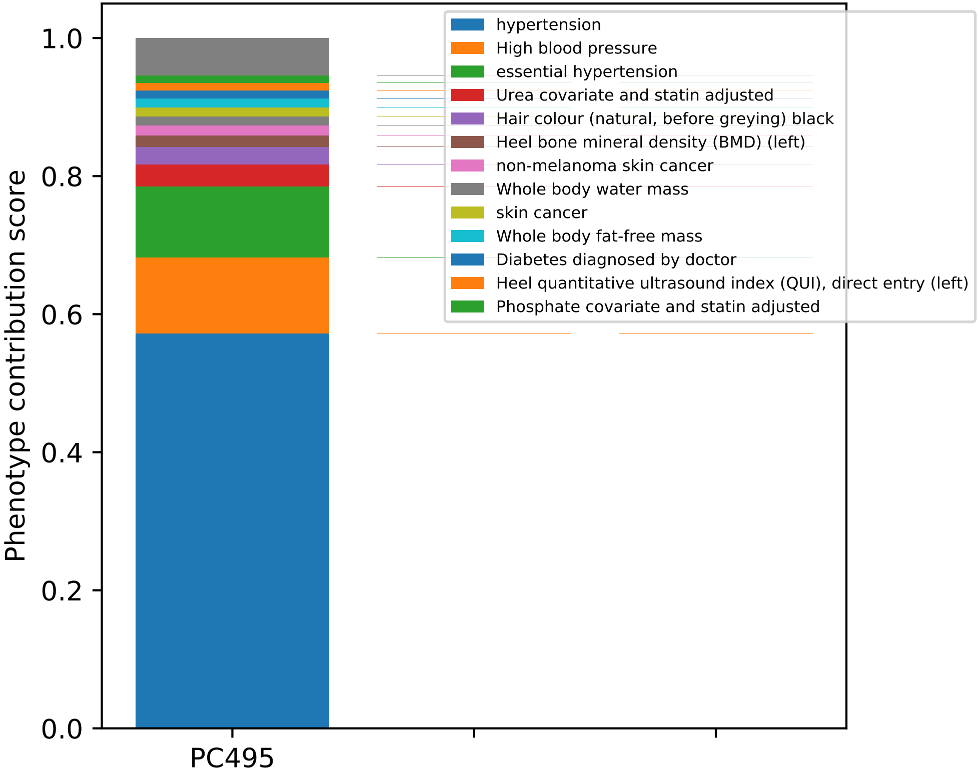

Phenotype contribution score
1.0
0.8
0.6
0.4
0.2
0.0
PC495
hypertension
High blood pressure
essential hypertension
Urea covariate and statin adjusted
Hair colour (natural, before greying) black
Heel bone mineral density (BMD) (left)
non-melanoma skin cancer
Whole body water mass
skin cancer
Whole body fat-free mass
Diabetes diagnosed by doctor
Heel quantitative ultrasound index (QUI), direct entry (left)
Phosphate covariate and statin adjusted

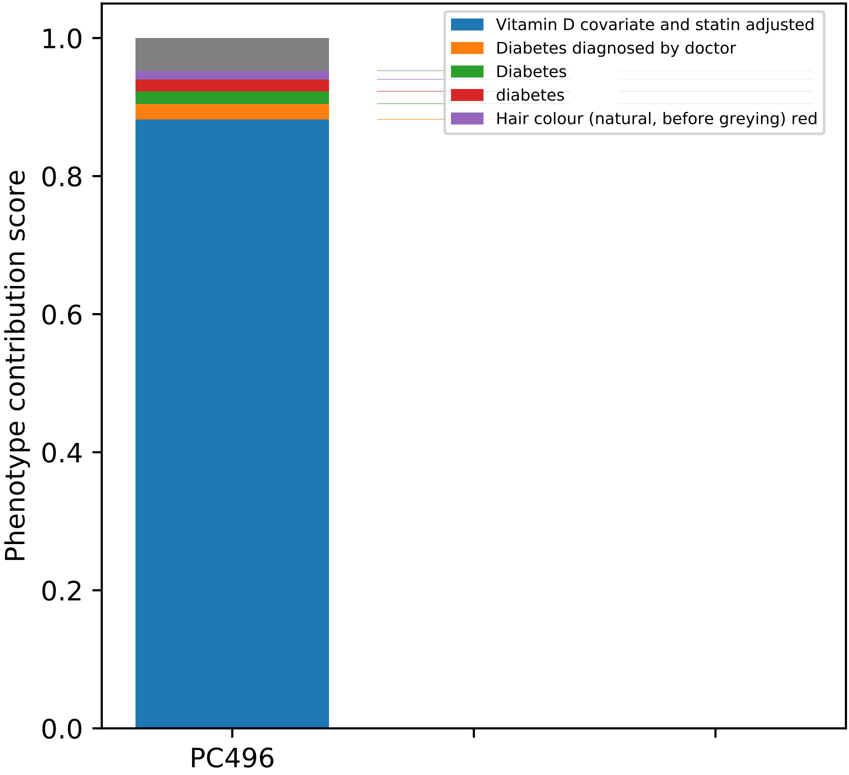
Vitamin D covariate and statin adjusted
Diabetes diagnosed by doctor
Diabetes
diabetes
Hair colour (natural, before greying) red
Phenotype contribution score
0.0
0.2
0.4
0.6
0.8
1.0
PC496

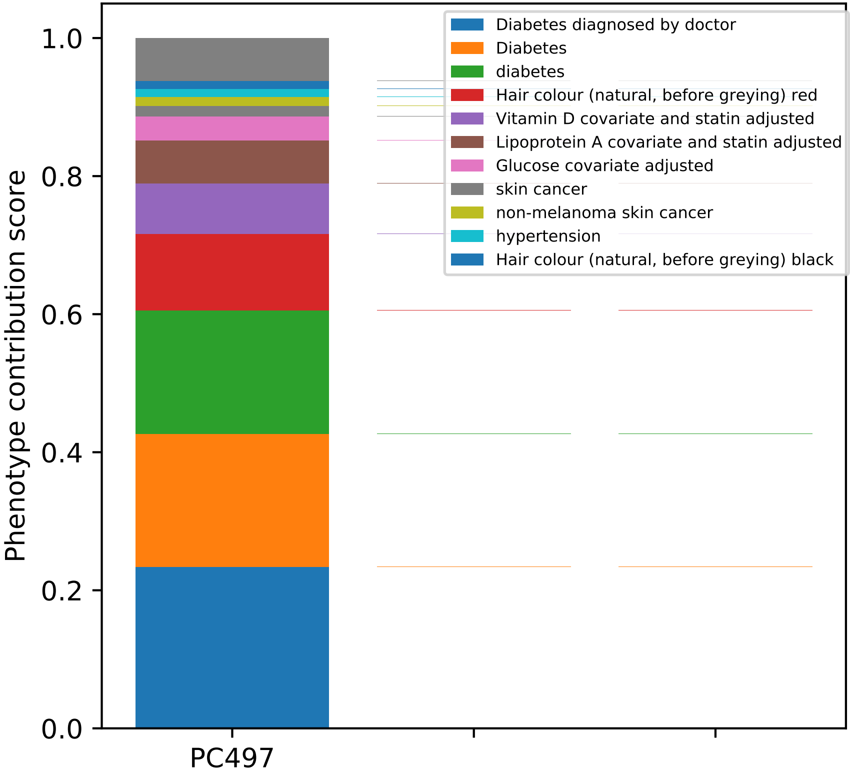

Phenotype contribution score
1.0
0.8
0.6
0.4
0.2
0.0
PC497
Diabetes diagnosed by doctor
Diabetes
diabetes
Hair colour (natural, before greying) red
Vitamin D covariate and statin adjusted
Lipoprotein A covariate and statin adjusted
Glucose covariate adjusted
skin cancer
non-melanoma skin cancer
hypertension
Hair colour (natural, before greying) black

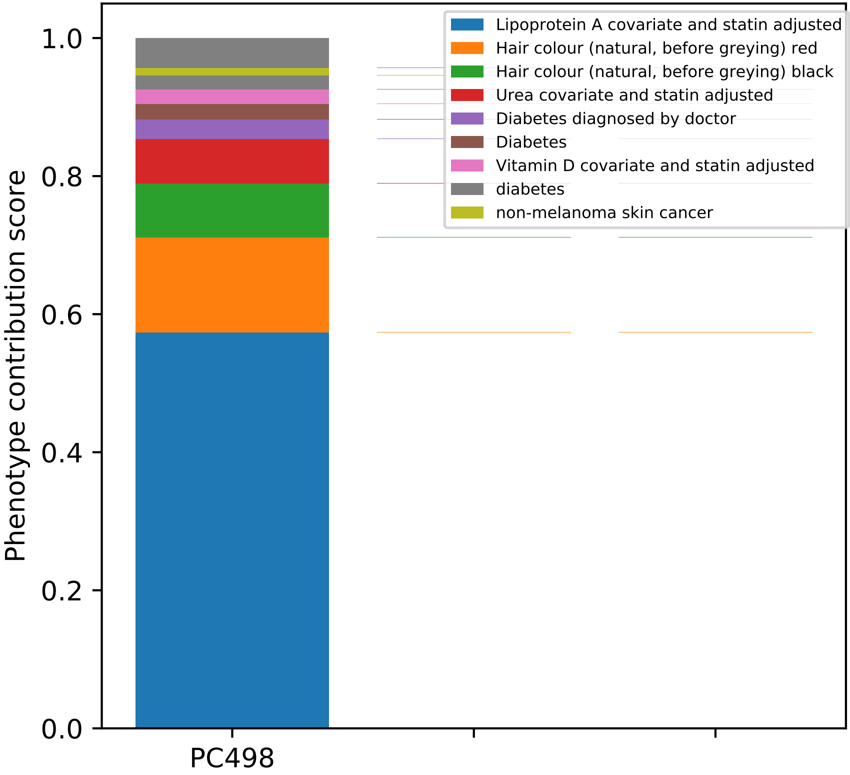

Lipoprotein A covariate and statin adjusted
Hair colour (natural, before greying) red
Hair colour (natural, before greying) black
Urea covariate and statin adjusted
Diabetes diagnosed by doctor
Diabetes
Vitamin D covariate and statin adjusted
diabetes
non-melanoma skin cancer
Phenotype contribution score
1.0
0.8
0.6
0.4
0.2
0.0
PC498

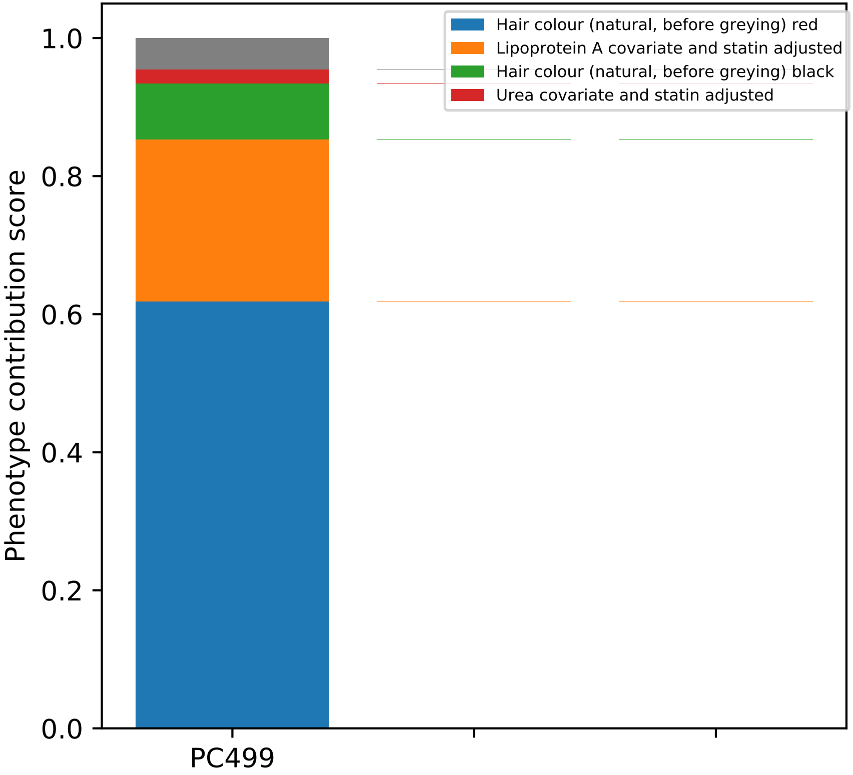
Hair colour (natural, before greying) red
Lipoprotein A covariate and statin adjusted
Hair colour (natural, before greying) black
Urea covariate and statin adjusted
Phenotype contribution score
0.0
0.2
0.4
0.6
0.8
1.0
PC499

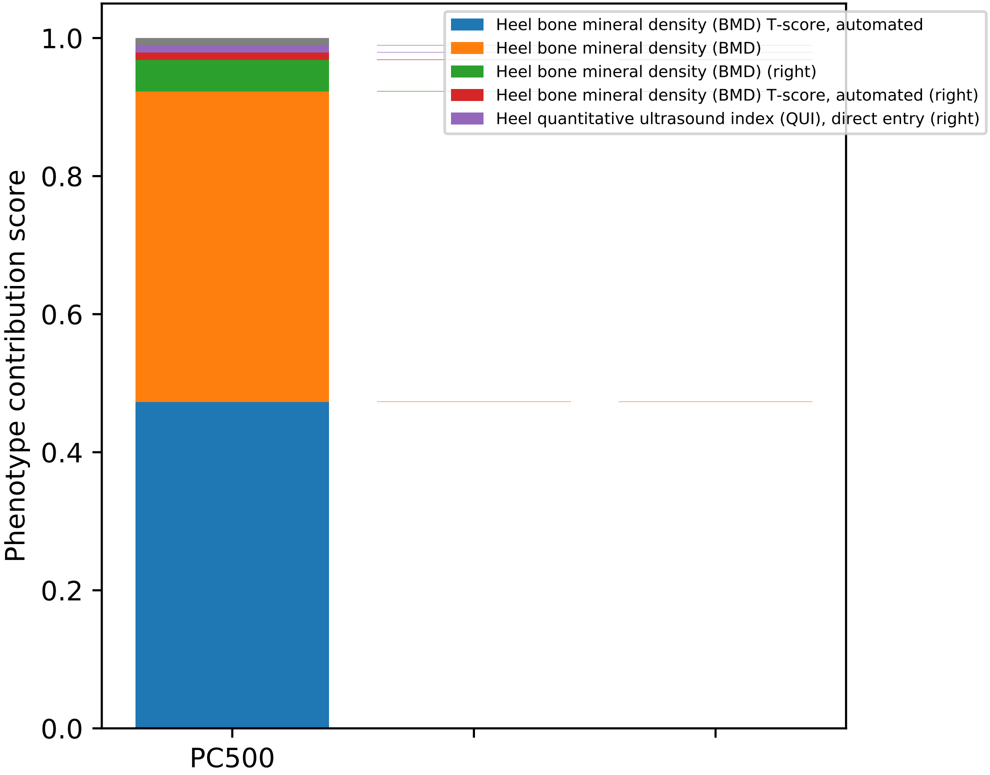

Heel bone mineral density (BMD) T-score, automated
Heel bone mineral density (BMD)
Heel bone mineral density (BMD) (right)
Heel bone mineral density (BMD) T-score, automated (right)
Heel quantitative ultrasound index (QUI), direct entry (right)
Phenotype contribution score
1.0
0.8
0.6
0.4
0.2
0.0
PC500
