## Supplementary material for "Polygenic risk modeling with latent trait-related genetic components": Data S3: Component annotation (genes)

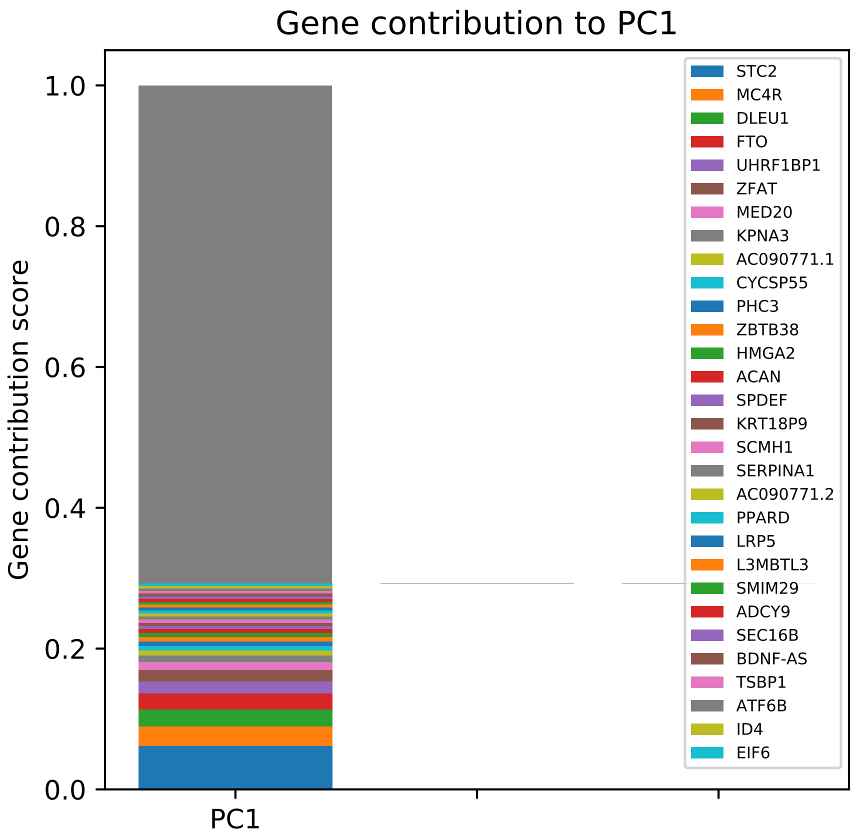
Gene contribution to PC1
Gene contribution score
1.0
0.8
0.6
0.4
0.2
0.0
PC1
STC2
MC4R
DLEU1
FTO
UHRF1BP1
ZFAT
MED20
KPNA3
AC090771.1
CYCSP55
PHC3
ZBTB38
HMGA2
ACAN
SPDEF
KRT18P9
SCMH1
SERPINA1
AC090771.2
PPARD
LRP5
L3MBTL3
SMIM29
ADCY9
SEC16B
BDNF-AS
TSBP1
ATF6B
ID4
EIF6

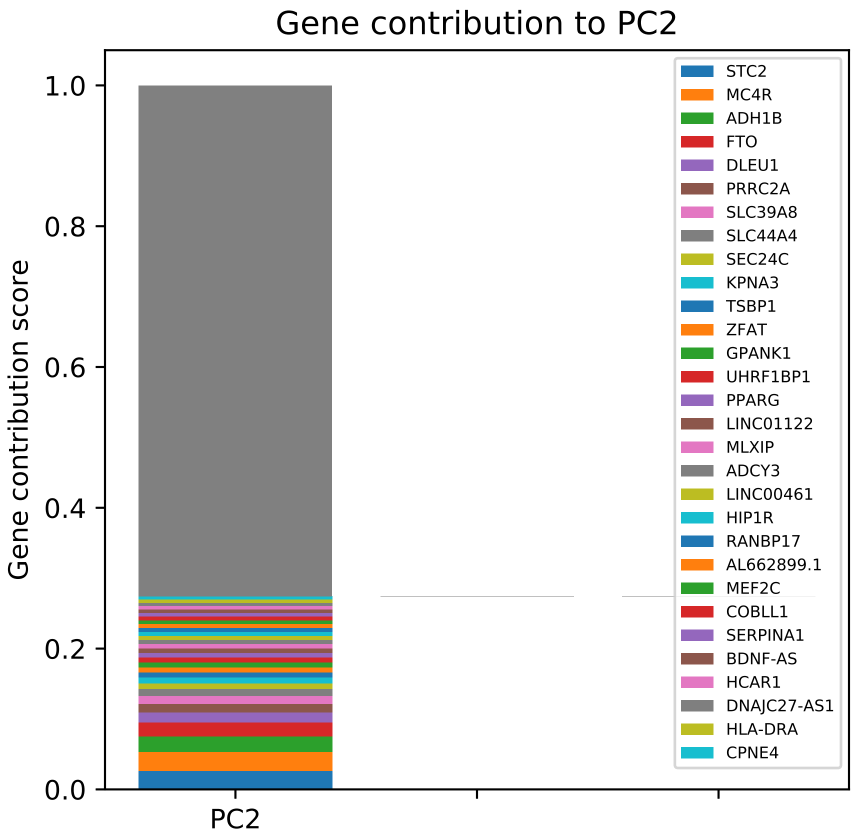
Gene contribution to PC2
Gene contribution score
1.0
0.8
0.6
0.4
0.2
0.0
PC2
STC2
MC4R
ADH1B
FTO
DLEU1
PRRC2A
SLC39A8
SLC44A4
SEC24C
KPNA3
TSBP1
ZFAT
GPANK1
UHRF1BP1
PPARG
LINC01122
MLXIP
ADCY3
LINC00461
HIP1R
RANBP17
AL662899.1
MEF2C
COBLL1
SERPINA1
BDNF-AS
HCAR1
DNAJC27-AS1
HLA-DRA
CPNE4

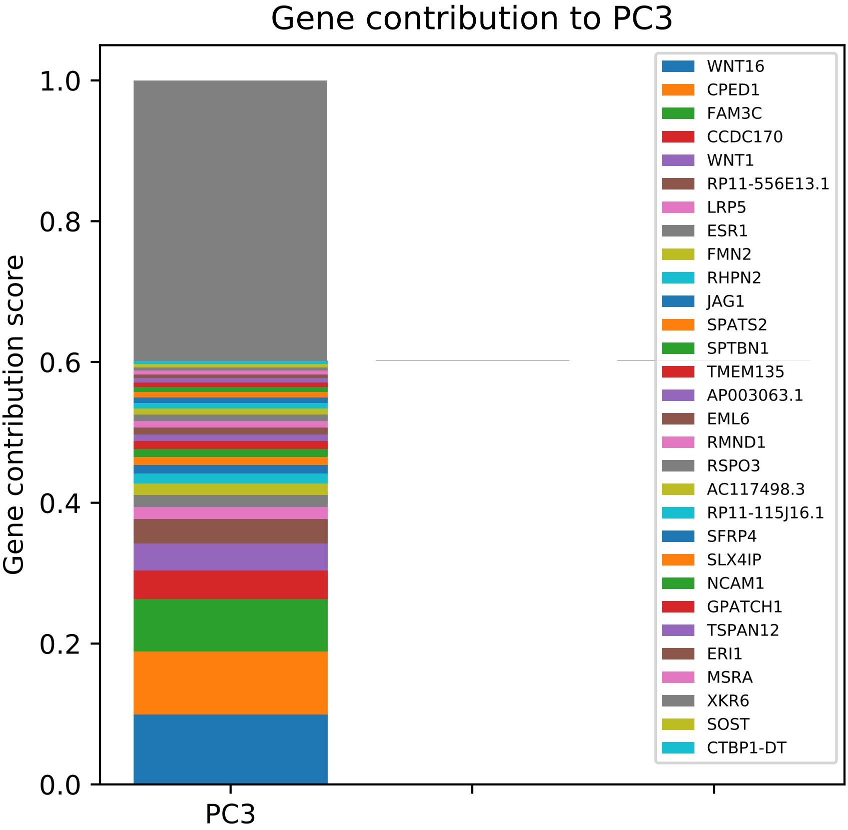

Gene contribution to PC3
Gene contribution score
1.0
0.8
0.6
0.4
0.2
0.0
PC3
WNT16
CPED1
FAM3C
CCDC170
WNT1
RP11-556E13.1
LRP5
ESR1
FMN2
RHPN2
JAG1
SPATS2
SPTBN1
TMEM135
AP003063.1
EML6
RMND1
RSPO3
AC117498.3
RP11-115J16.1
SFRP4
SLX4IP
NCAM1
GPATCH1
TSPAN12
ERI1
MSRA
XKR6
SOST
CTBP1-DT

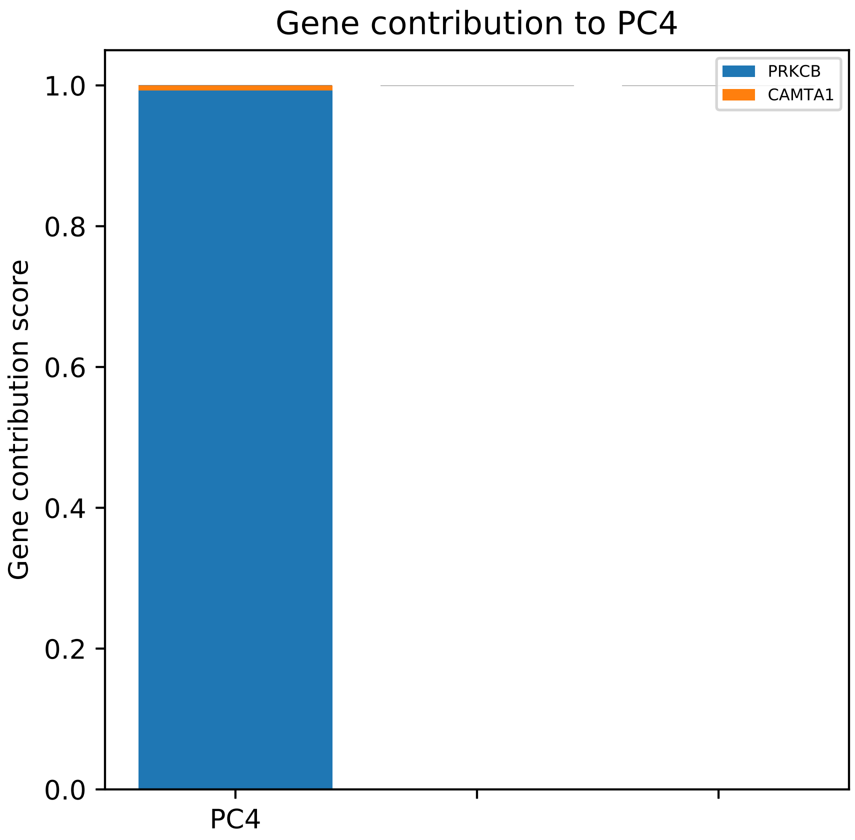
Gene contribution to PC4
PRKCB
CAMTA1
Gene contribution score
1.0
0.8
0.6
0.4
0.2
0.0
PC4

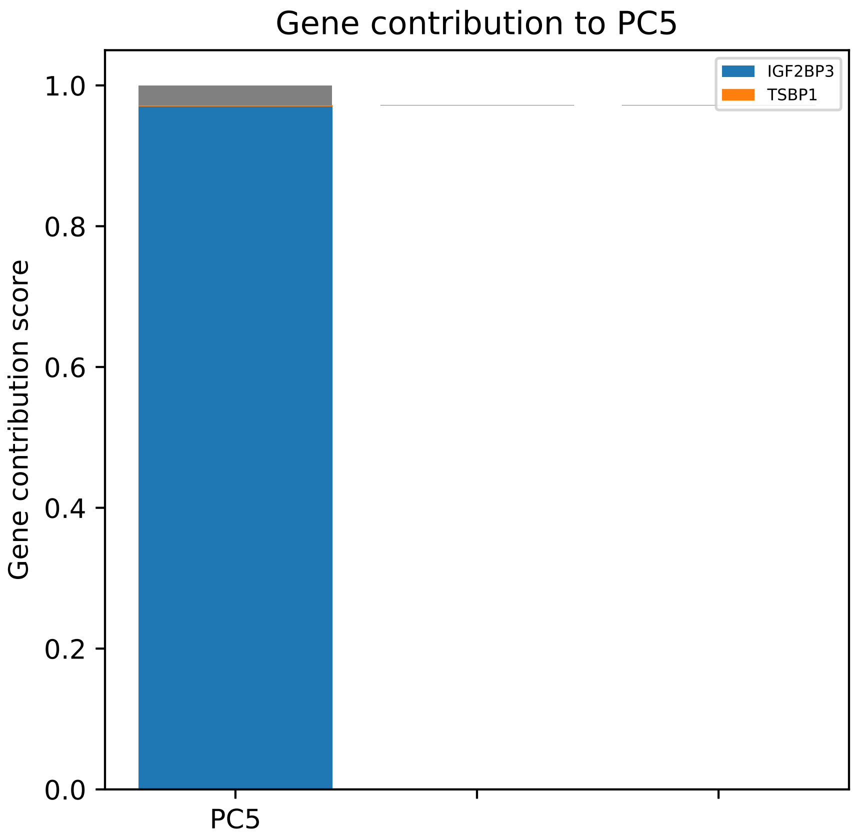

Gene contribution to PC5
IGF2BP3
TSBP1
1.0
0.8
0.6
0.4
0.2
0.0
Gene contribution score
PC5

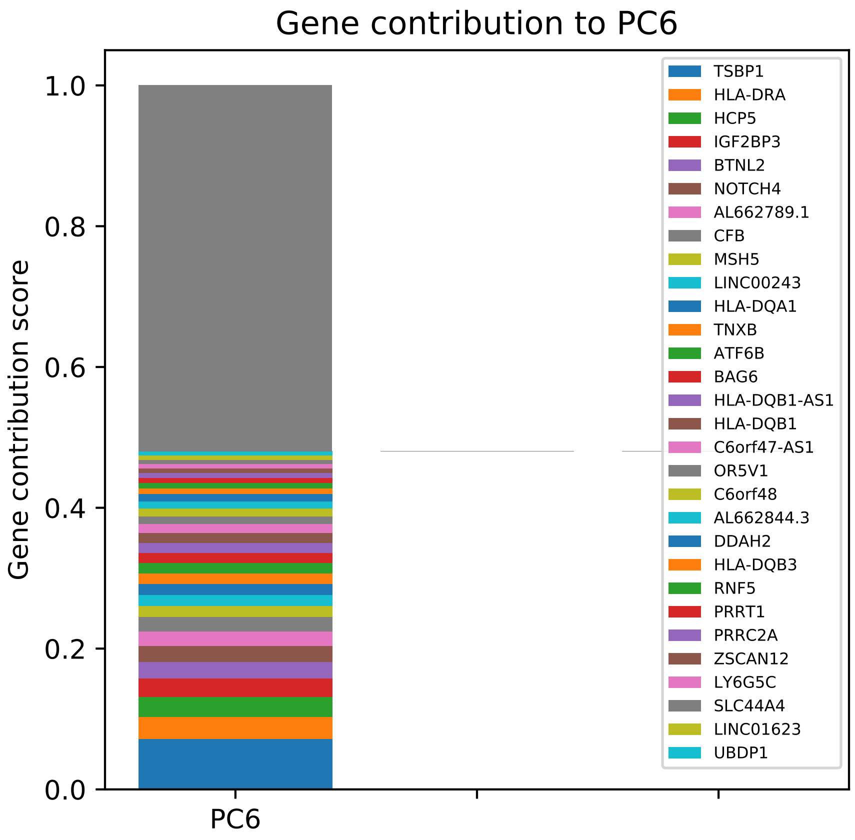

Gene contribution to PC6
Gene contribution score
1.0
0.8
0.6
0.4
0.2
0.0
PC6
TSBP1
HLA-DRA
HCP5
IGF2BP3
BTNL2
NOTCH4
AL662789.1
CFB
MSH5
LINC00243
HLA-DQA1
TNXB
ATF6B
BAG6
HLA-DQB1-AS1
HLA-DQB1
C6orf47-AS1
OR5V1
C6orf48
AL662844.3
DDAH2
HLA-DQB3
RNF5
PRRT1
PRRC2A
ZSCAN12
LY6G5C
SLC44A4
LINC01623
UBDP1

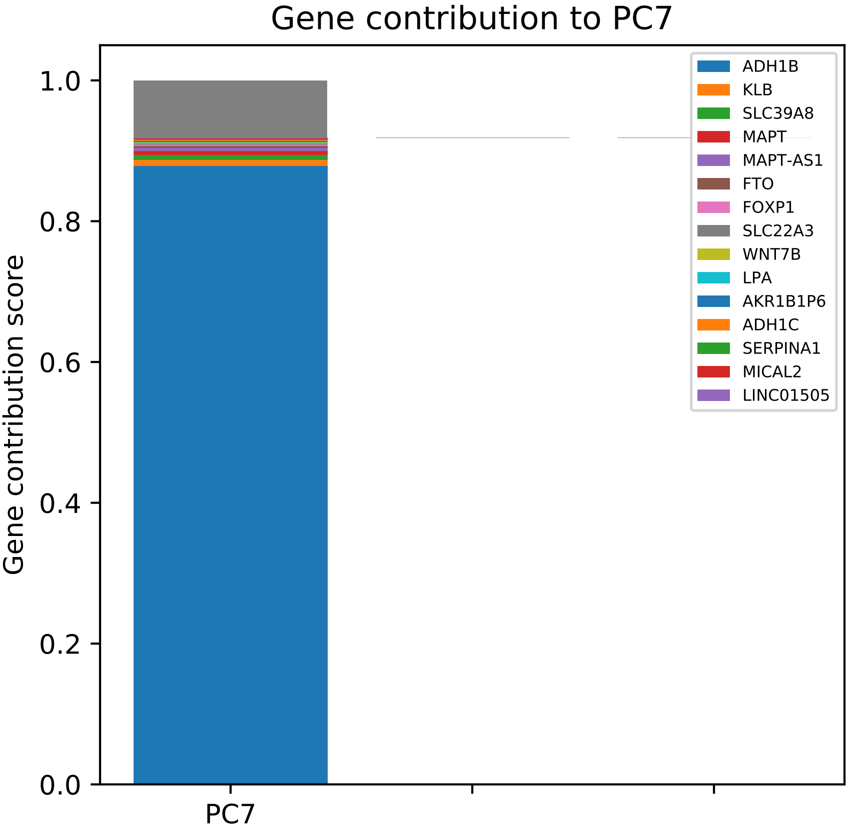

Gene contribution to PC7
Gene contribution score
0.0
0.2
0.4
0.6
0.8
1.0
PC7
ADH1B
KLB
SLC39A8
MAPT
MAPT-AS1
FTO
FOXP1
SLC22A3
WNT7B
LPA
AKR1B1P6
ADH1C
SERPINA1
MICAL2
LINC01505

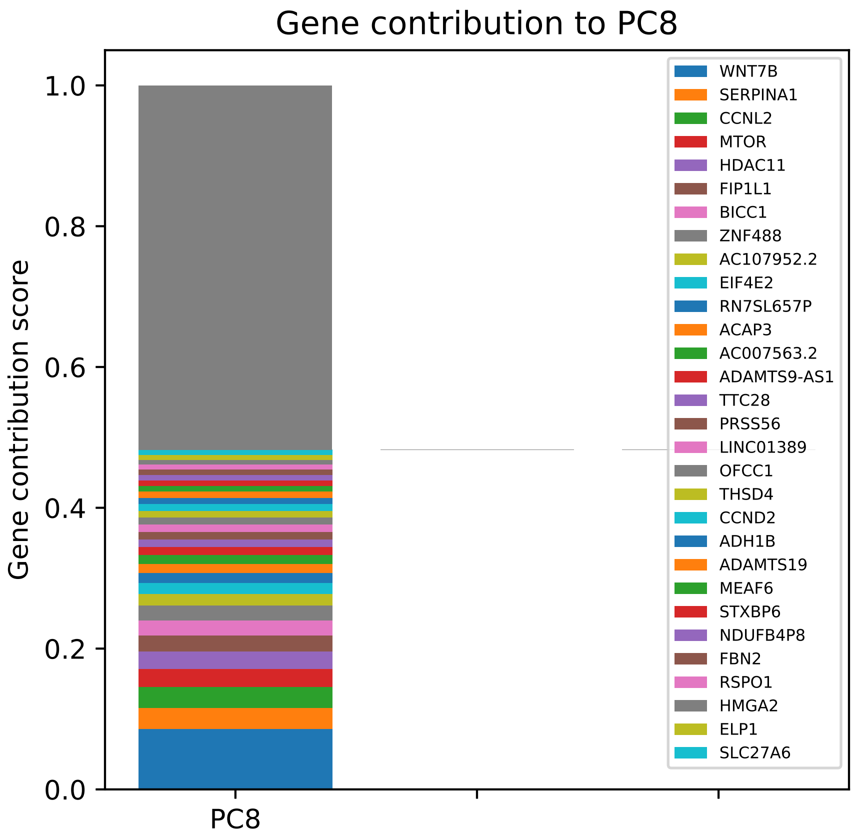

Gene contribution to PC8
Gene contribution score
1.0
0.8
0.6
0.4
0.2
0.0
PC8
WNT7B
SERPINA1
CCNL2
MTOR
HDAC11
FIP1L1
BICC1
ZNF488
AC107952.2
EIF4E2
RN7SL657P
ACAP3
AC007563.2
ADAMTS9-AS1
TTC28
PRSS56
LINC01389
OFCC1
THSD4
CCND2
ADH1B
ADAMTS19
MEAF6
STXBP6
NDUFB4P8
FBN2
RSPO1
HMGA2
ELP1
SLC27A6

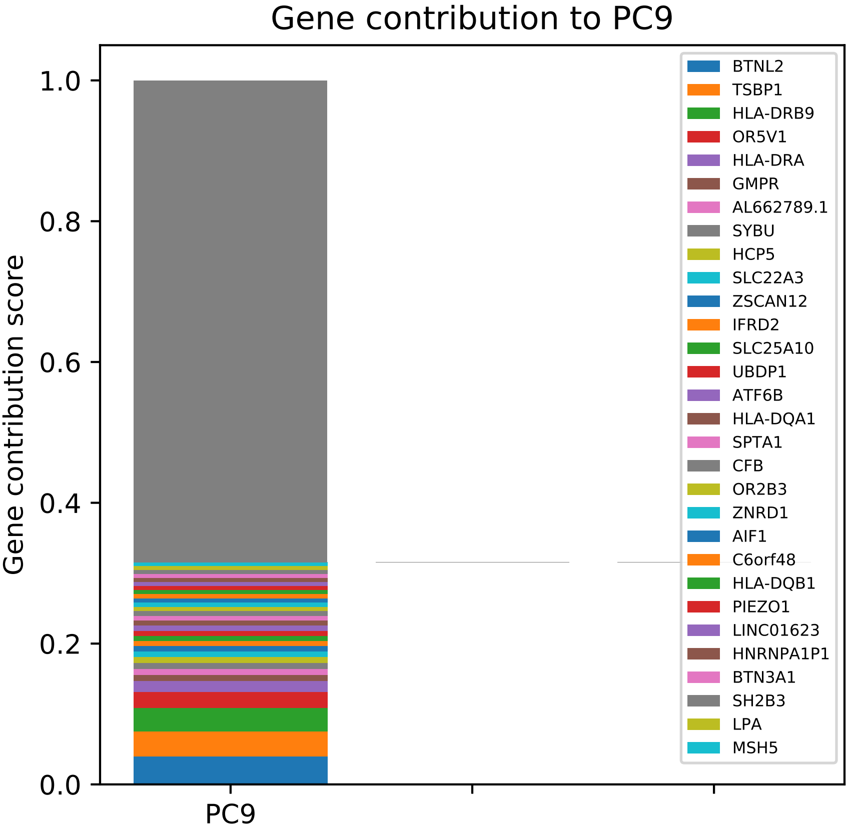

Gene contribution to PC9
Gene contribution score
1.0
0.8
0.6
0.4
0.2
0.0
PC9
BTNL2
TSBP1
HLA-DRB9
OR5V1
HLA-DRA
GMPR
AL662789.1
SYBU
HCP5
SLC22A3
ZSCAN12
IFRD2
SLC25A10
UBDP1
ATF6B
HLA-DQA1
SPTA1
CFB
OR2B3
ZNRD1
AIF1
C6orf48
HLA-DQB1
PIEZO1
LINC01623
HNRNPA1P1
BTN3A1
SH2B3
LPA
MSH5

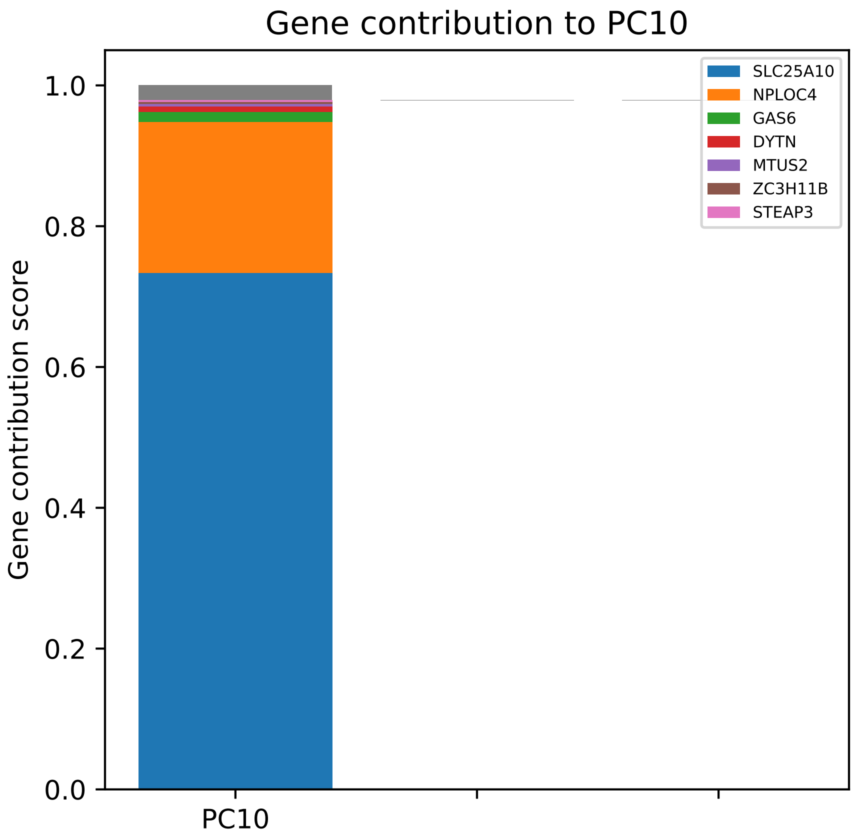
Gene contribution to PC10
Gene contribution score
1.0
0.8
0.6
0.4
0.2
0.0
PC10
SLC25A10
NPLOC4
GAS6
DYTN
MTUS2
ZC3H11B
STEAP3

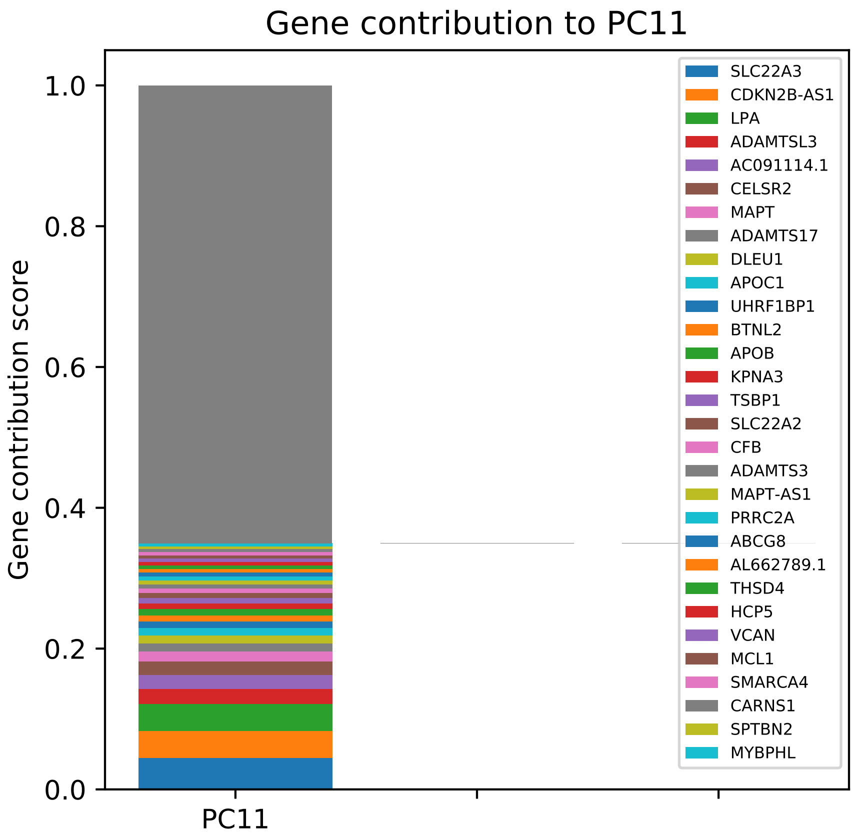

Gene contribution to PC11
Gene contribution score
1.0
0.8
0.6
0.4
0.2
0.0
PC11
SLC22A3
CDKN2B-AS1
LPA
ADAMTSL3
AC091114.1
CELSR2
MAPT
ADAMTS17
DLEU1
APOC1
UHRF1BP1
BTNL2
APOB
KPNA3
TSBP1
SLC22A2
CFB
ADAMTS3
MAPT-AS1
PRRC2A
ABCG8
AL662789.1
THSD4
HCP5
VCAN
MCL1
SMARCA4
CARNS1
SPTBN2
MYBPHL

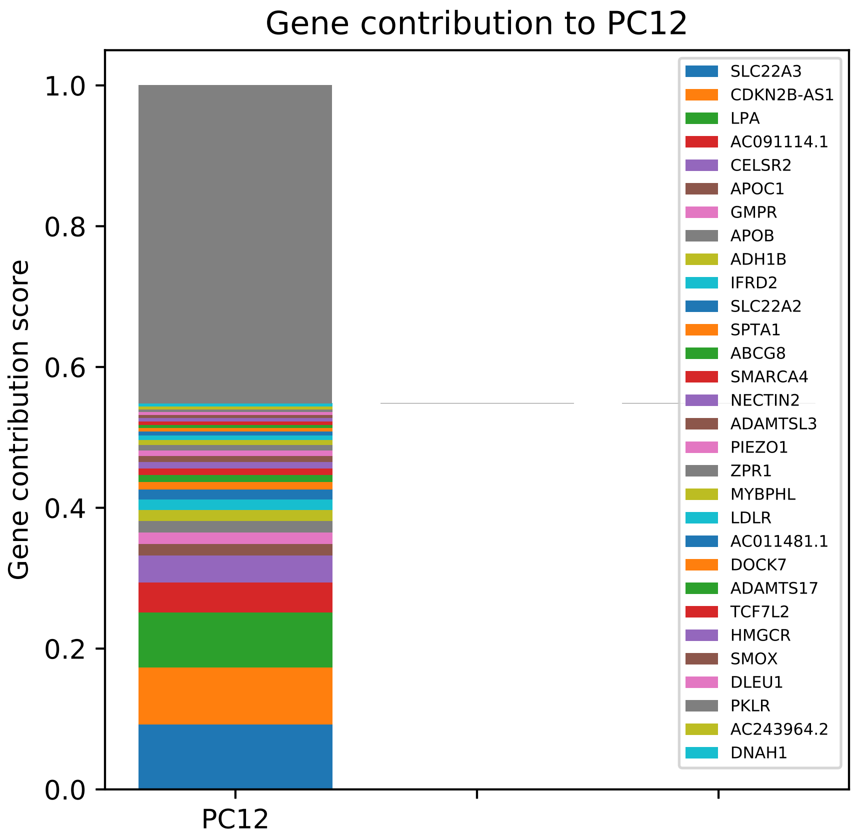
Gene contribution to PC12
Gene contribution score
1.0
0.8
0.6
0.4
0.2
0.0
PC12
SLC22A3
CDKN2B-AS1
LPA
AC091114.1
CELSR2
APOC1
GMPR
APOB
ADH1B
IFRD2
SLC22A2
SPTA1
ABCG8
SMARCA4
NECTIN2
ADAMTSL3
PIEZO1
ZPR1
MYBPHL
LDLR
AC011481.1
DOCK7
ADAMTS17
TCF7L2
HMGCR
SMOX
DLEU1
PKLR
AC243964.2
DNAH1

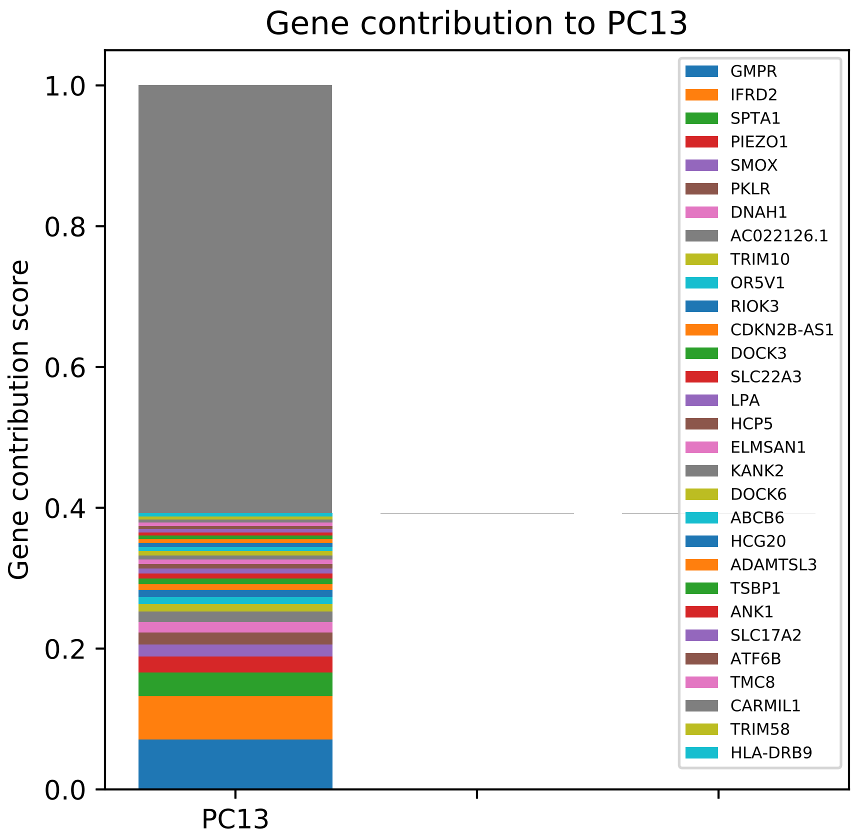
Gene contribution to PC13
Gene contribution score
1.0
0.8
0.6
0.4
0.2
0.0
PC13
GMPR
IFRD2
SPTA1
PIEZO1
SMOX
PKLR
DNAH1
AC022126.1
TRIM10
OR5V1
RIOK3
CDKN2B-AS1
DOCK3
SLC22A3
LPA
HCP5
ELMSAN1
KANK2
DOCK6
ABCB6
HCG20
ADAMTSL3
TSBP1
ANK1
SLC17A2
ATF6B
TMC8
CARMIL1
TRIM58
HLA-DRB9

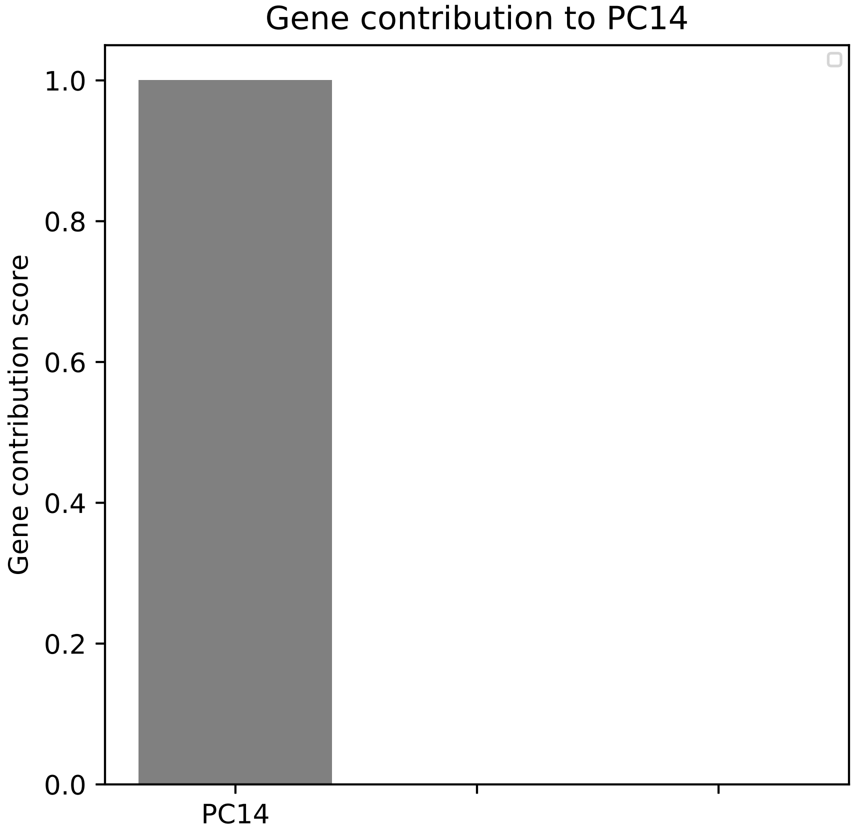
Gene contribution to PC14
Gene contribution score
1.0
0.8
0.6
0.4
0.2
0.0
PC14

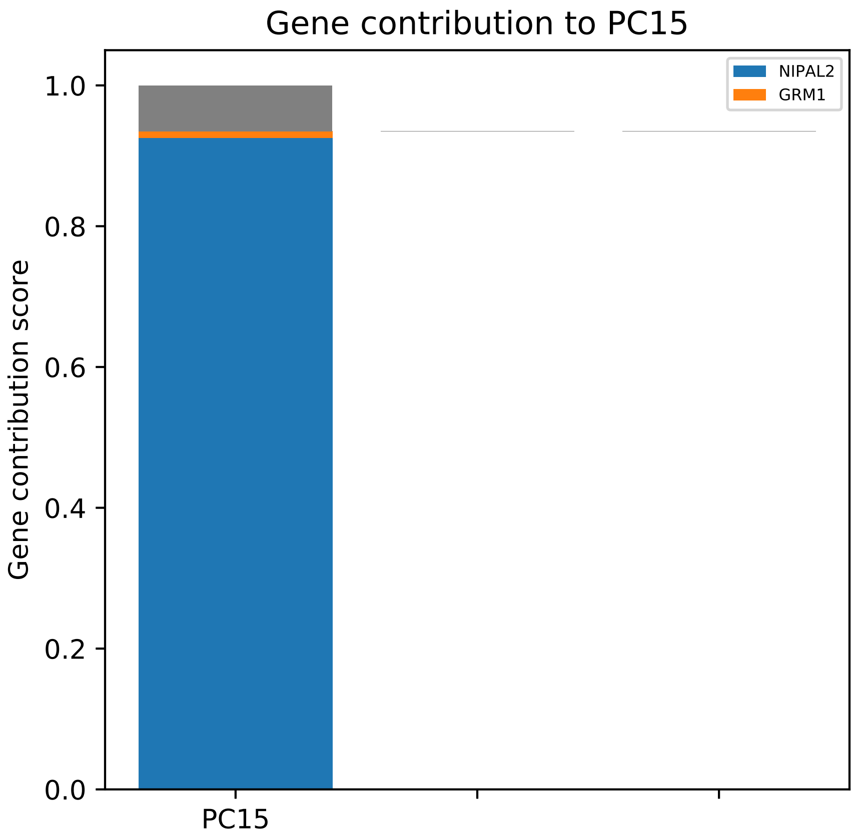
Gene contribution to PC15
NIPAL2
GRM1
1.0
0.8
0.6
0.4
0.2
0.0
Gene contribution score
PC15

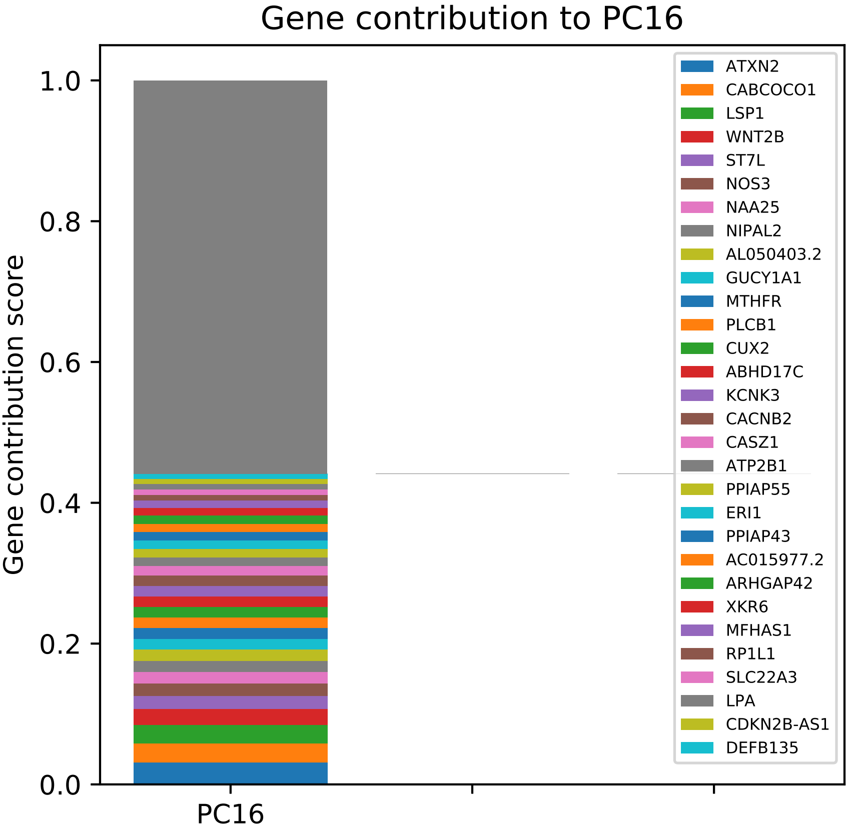
Gene contribution to PC16
Gene contribution score
1.0
0.8
0.6
0.4
0.2
0.0
PC16
ATXN2
CABCOCO1
LSP1
WNT2B
ST7L
NOS3
NAA25
NIPAL2
AL050403.2
GUCY1A1
MTHFR
PLCB1
CUX2
ABHD17C
KCNK3
CACNB2
CASZ1
ATP2B1
PPIAP55
ERI1
PPIAP43
AC015977.2
ARHGAP42
XKR6
MFHAS1
RP1L1
SLC22A3
LPA
CDKN2B-AS1
DEFB135

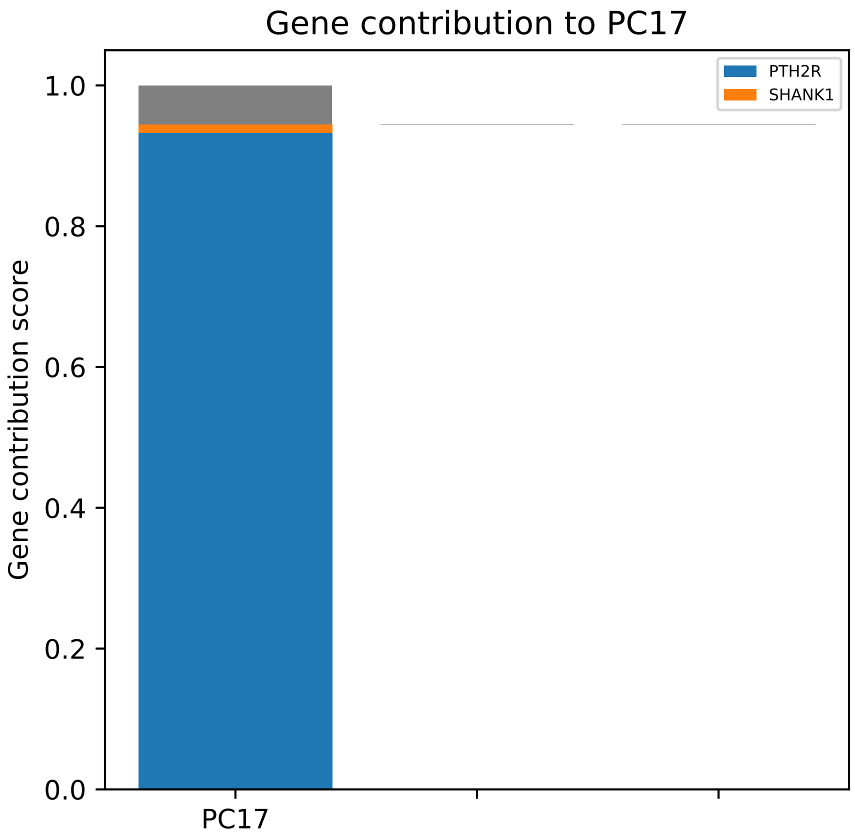

Gene contribution to PC17
Gene contribution score
1.0
0.8
0.6
0.4
0.2
0.0
PC17
PTH2R
SHANK1

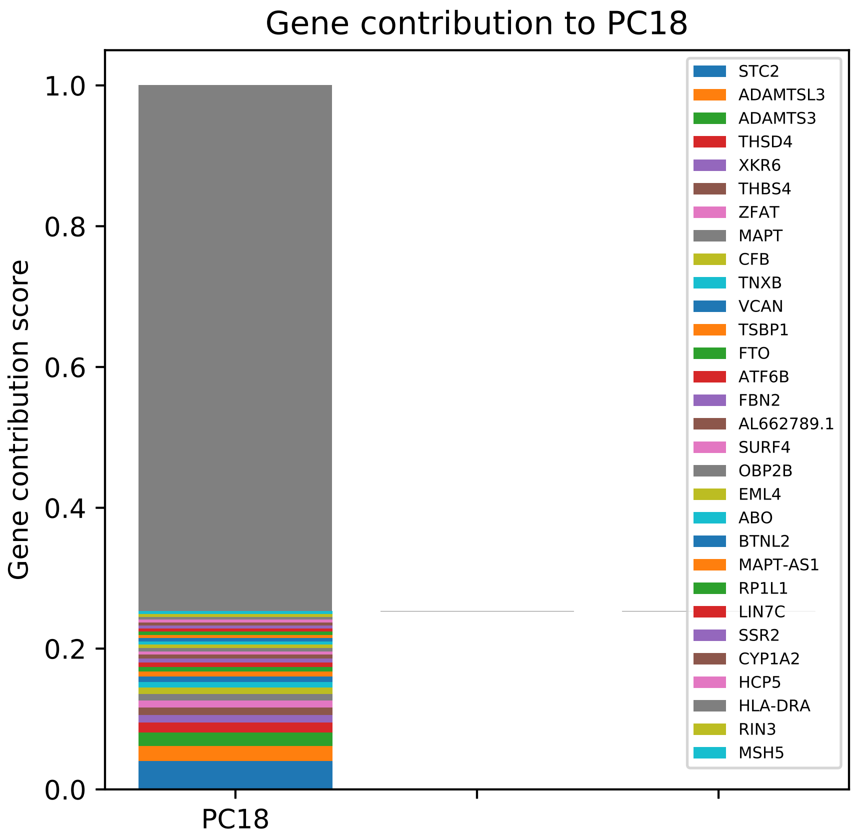
Gene contribution to PC18
Gene contribution score
1.0
0.8
0.6
0.4
0.2
0.0
PC18
STC2
ADAMTSL3
ADAMTS3
THSD4
XKR6
THBS4
ZFAT
MAPT
CFB
TNXB
VCAN
TSBP1
FTO
ATF6B
FBN2
AL662789.1
SURF4
OBP2B
EML4
ABO
BTNL2
MAPT-AS1
RP1L1
LIN7C
SSR2
CYP1A2
HCP5
HLA-DRA
RIN3
MSH5

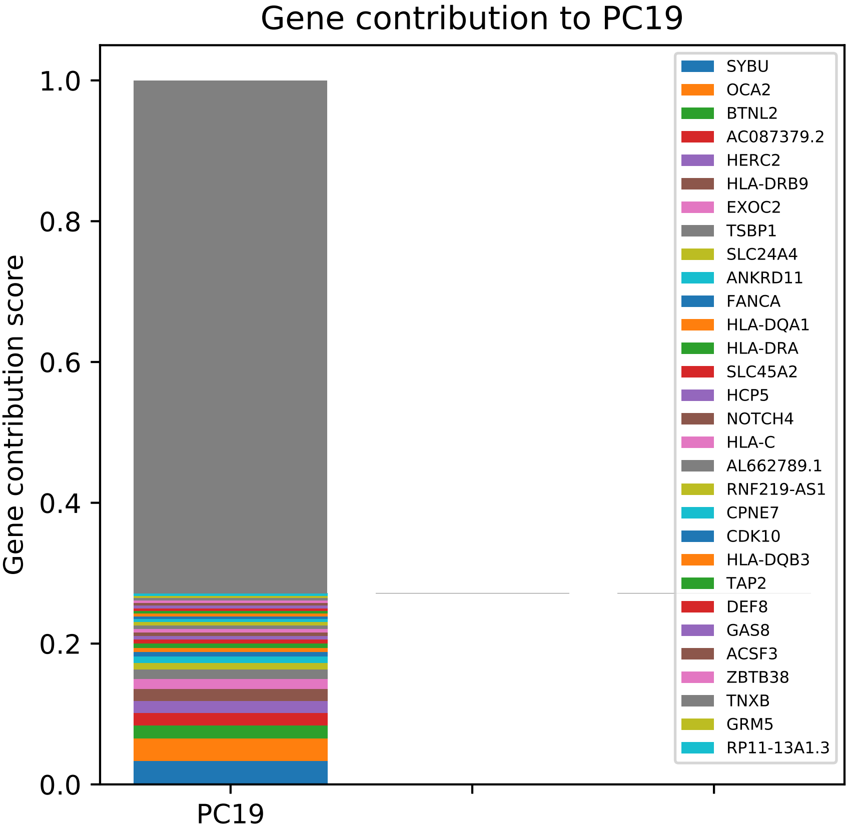

Gene contribution to PC19
Gene contribution score
1.0
0.8
0.6
0.4
0.2
0.0
PC19
SYBU
OCA2
BTNL2
AC087379.2
HERC2
HLA-DRB9
EXOC2
TSBP1
SLC24A4
ANKRD11
FANCA
HLA-DQA1
HLA-DRA
SLC45A2
HCP5
NOTCH4
HLA-C
AL662789.1
RNF219-AS1
CPNE7
CDK10
HLA-DQB3
TAP2
DEF8
GAS8
ACSF3
ZBTB38
TNXB
GRM5
RP11-13A1.3

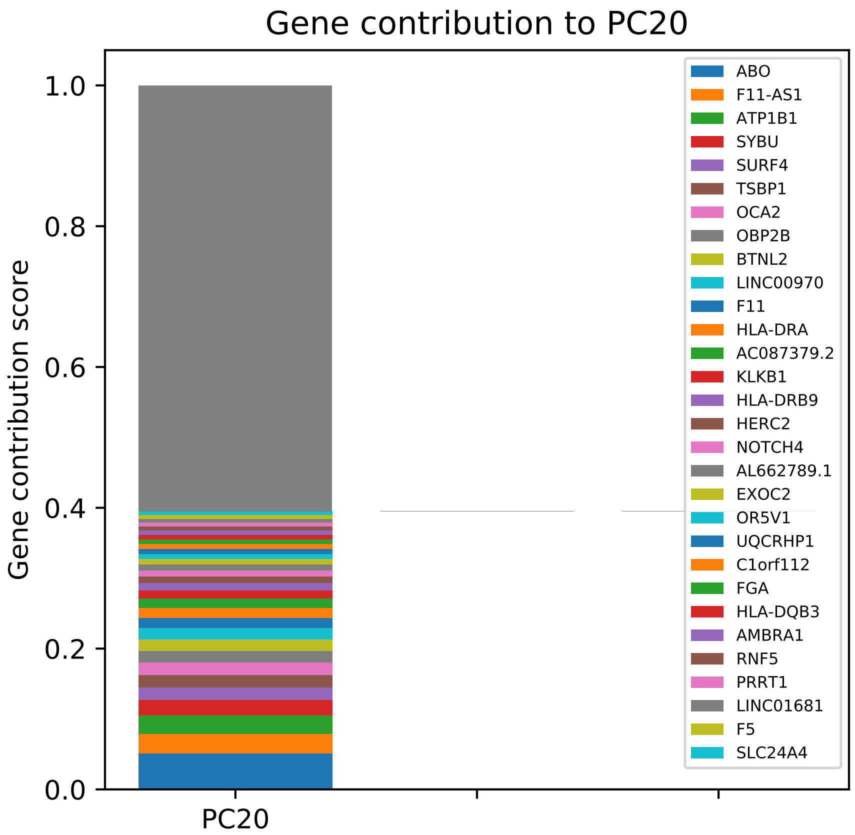

Gene contribution to PC20
Gene contribution score
1.0
0.8
0.6
0.4
0.2
0.0
PC20
ABO
F11-AS1
ATP1B1
SYBU
SURF4
TSBP1
OCA2
OBP2B
BTNL2
LINC00970
F11
HLA-DRA
AC087379.2
KLKB1
HLA-DRB9
HERC2
NOTCH4
AL662789.1
EXOC2
OR5V1
UQCRHP1
C1orf112
FGA
HLA-DQB3
AMBRA1
RNF5
PRRT1
LINC01681
F5
SLC24A4

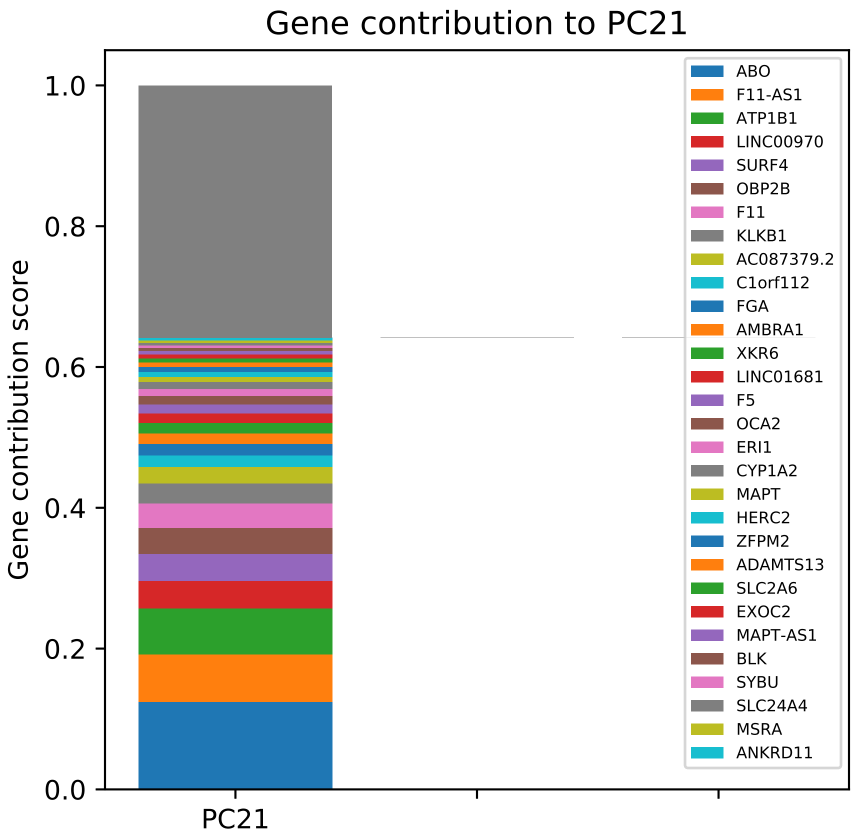

Gene contribution to PC21
Gene contribution score
0.0
0.2
0.4
0.6
0.8
1.0
PC21
ABO
F11-AS1
ATP1B1
LINC00970
SURF4
OBP2B
F11
KLKB1
AC087379.2
C1orf112
FGA
AMBRA1
XKR6
LINC01681
F5
OCA2
ERI1
CYP1A2
MAPT
HERC2
ZFPM2
ADAMTS13
SLC2A6
EXOC2
MAPT-AS1
BLK
SYBU
SLC24A4
MSRA
ANKRD11

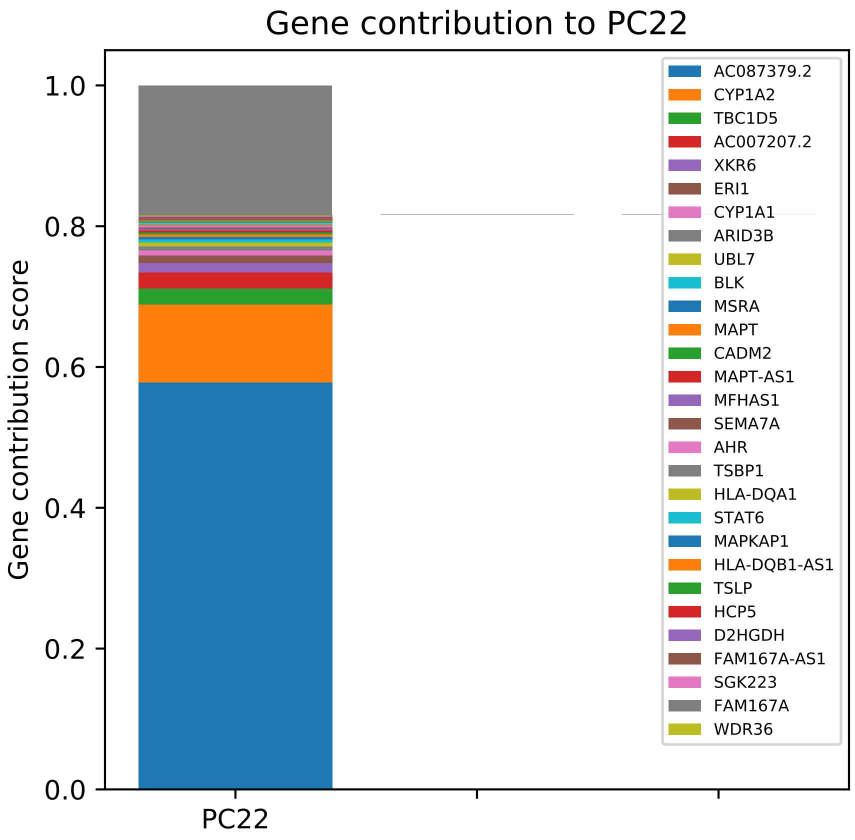

Gene contribution to PC22
Gene contribution score
1.0
0.8
0.6
0.4
0.2
0.0
PC22
AC087379.2
CYP1A2
TBC1D5
AC007207.2
XKR6
ERI1
CYP1A1
ARID3B
UBL7
BLK
MSRA
MAPT
CADM2
MAPT-AS1
MFHAS1
SEMA7A
AHR
TSBP1
HLA-DQA1
STAT6
MAPKAP1
HLA-DQB1-AS1
TSLP
HCP5
D2HGDH
FAM167A-AS1
SGK223
FAM167A
WDR36

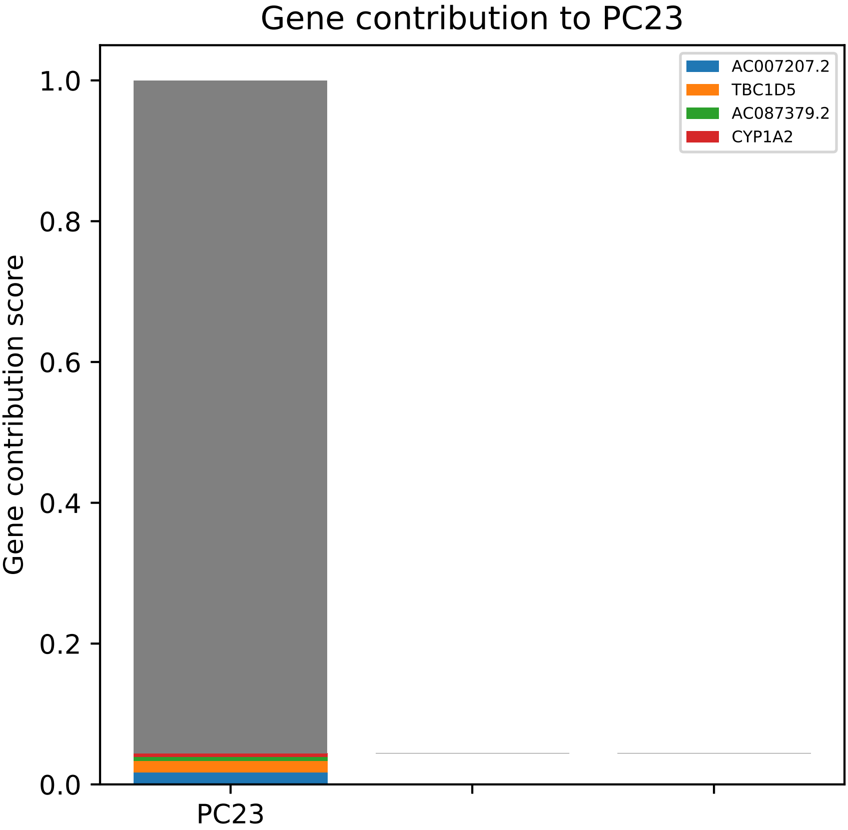
Gene contribution to PC23
Gene contribution score
1.0
0.8
0.6
0.4
0.2
0.0
PC23
AC007207.2
TBC1D5
AC087379.2
CYP1A2

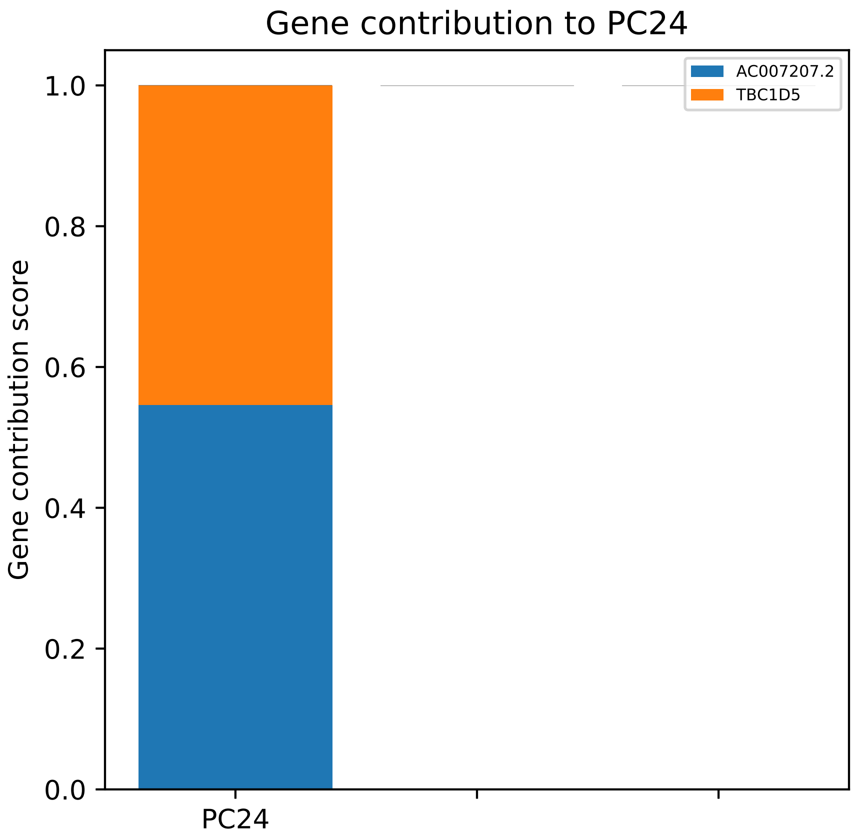

Gene contribution to PC24
Gene contribution score
1.0
0.8
0.6
0.4
0.2
0.0
PC24
AC007207.2
TBC1D5

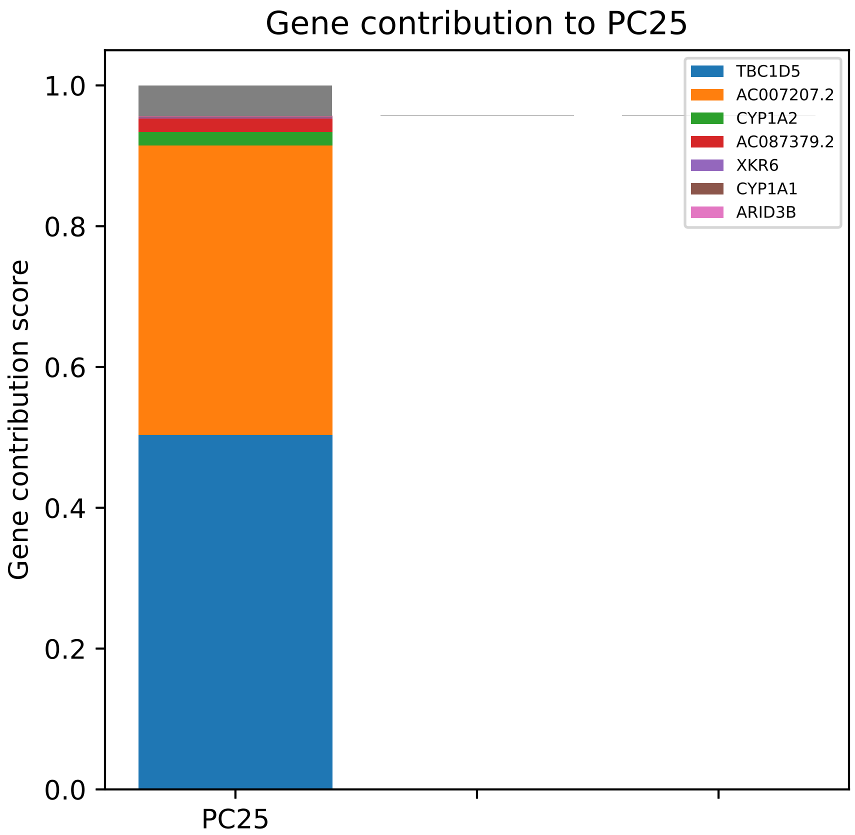

Gene contribution to PC25
Gene contribution score
1.0
0.8
0.6
0.4
0.2
0.0
PC25
TBC1D5
AC007207.2
CYP1A2
AC087379.2
XKR6
CYP1A1
ARID3B

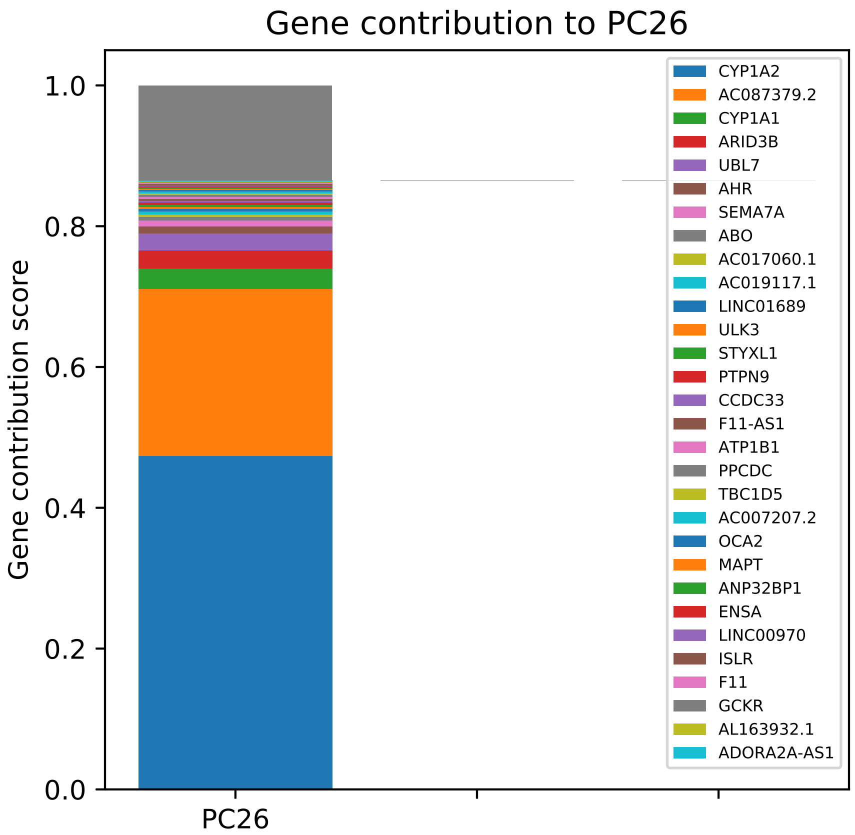

Gene contribution to PC26
Gene contribution score
1.0
0.8
0.6
0.4
0.2
0.0
PC26
CYP1A2
AC087379.2
CYP1A1
ARID3B
UBL7
AHR
SEMA7A
ABO
AC017060.1
AC019117.1
LINC01689
ULK3
STYXL1
PTPN9
CCDC33
F11-AS1
ATP1B1
PPCDC
TBC1D5
AC007207.2
OCA2
MAPT
ANP32BP1
ENSA
LINC00970
ISLR
F11
GCKR
AL163932.1
ADORA2A-AS1

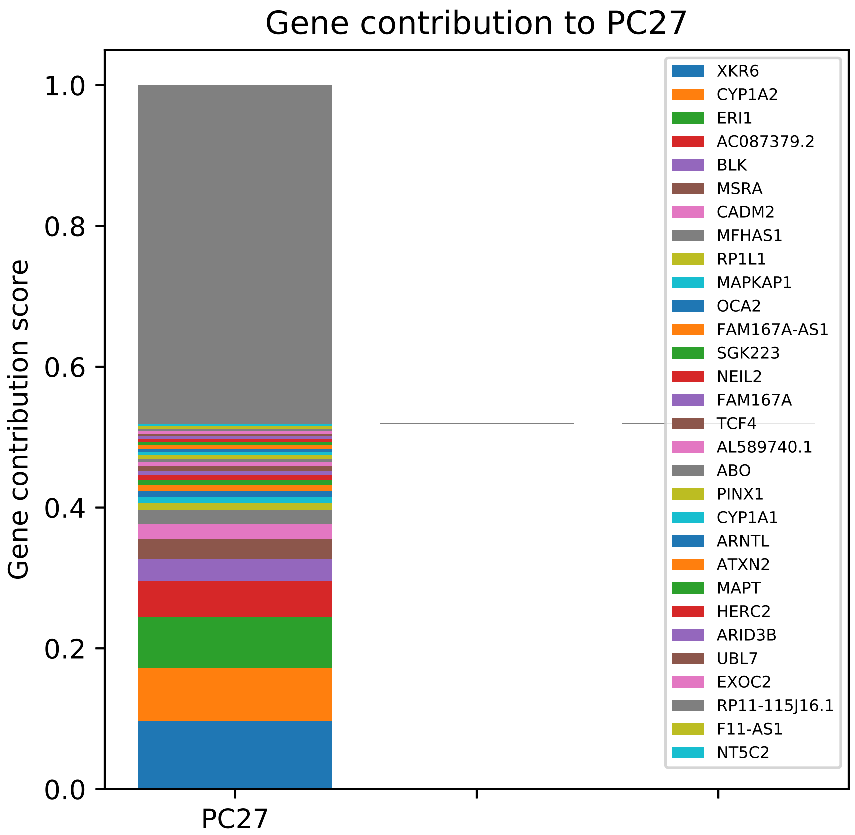

Gene contribution to PC27
Gene contribution score
1.0
0.8
0.6
0.4
0.2
0.0
PC27
XKR6
CYP1A2
ERI1
AC087379.2
BLK
MSRA
CADM2
MFHAS1
RP1L1
MAPKAP1
OCA2
FAM167A-AS1
SGK223
NEIL2
FAM167A
TCF4
AL589740.1
ABO
PINX1
CYP1A1
ARNTL
ATXN2
MAPT
HERC2
ARID3B
UBL7
EXOC2
RP11-115J16.1
F11-AS1
NT5C2

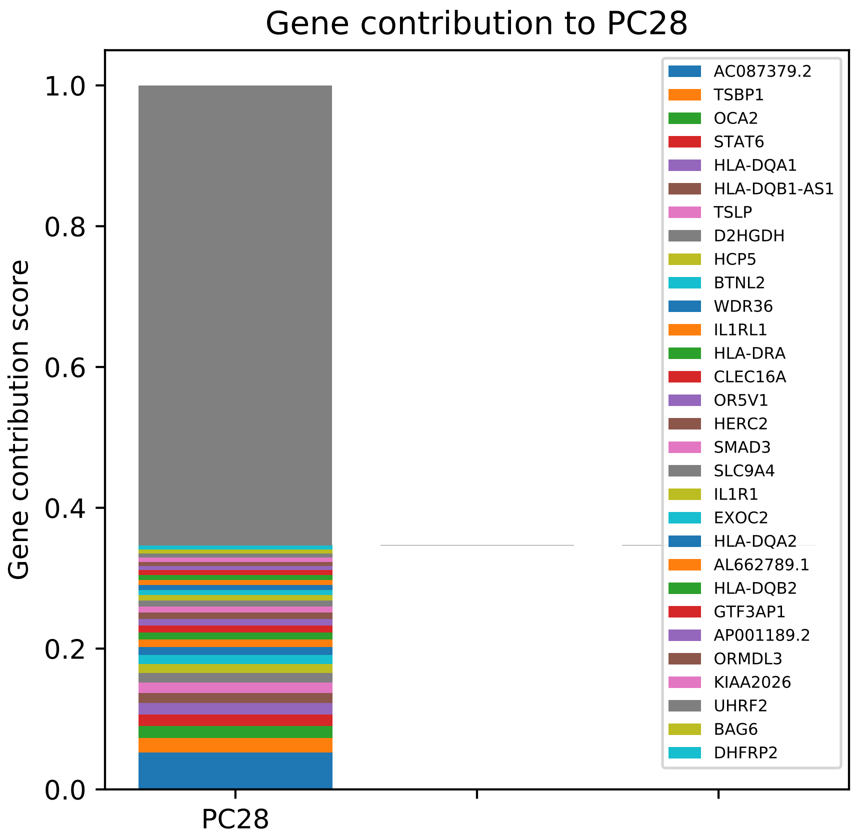
Gene contribution to PC28
Gene contribution score
1.0
0.8
0.6
0.4
0.2
0.0
PC28
AC087379.2
TSBP1
OCA2
STAT6
HLA-DQA1
HLA-DQB1-AS1
TSLP
D2HGDH
HCP5
BTNL2
WDR36
IL1RL1
HLA-DRA
CLEC16A
OR5V1
HERC2
SMAD3
SLC9A4
IL1R1
EXOC2
HLA-DQA2
AL662789.1
HLA-DQB2
GTF3AP1
AP001189.2
ORMDL3
KIAA2026
UHRF2
BAG6
DHFRP2

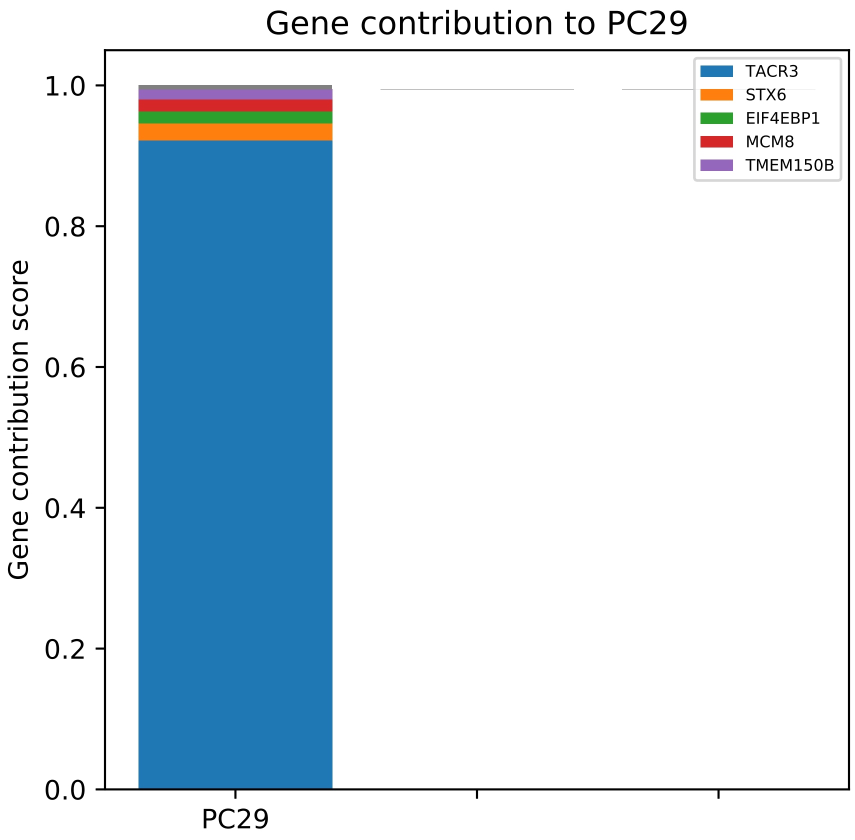

Gene contribution to PC29
Gene contribution score
0.0
0.2
0.4
0.6
0.8
1.0
PC29
TACR3
STX6
EIF4EBP1
MCM8
TMEM150B

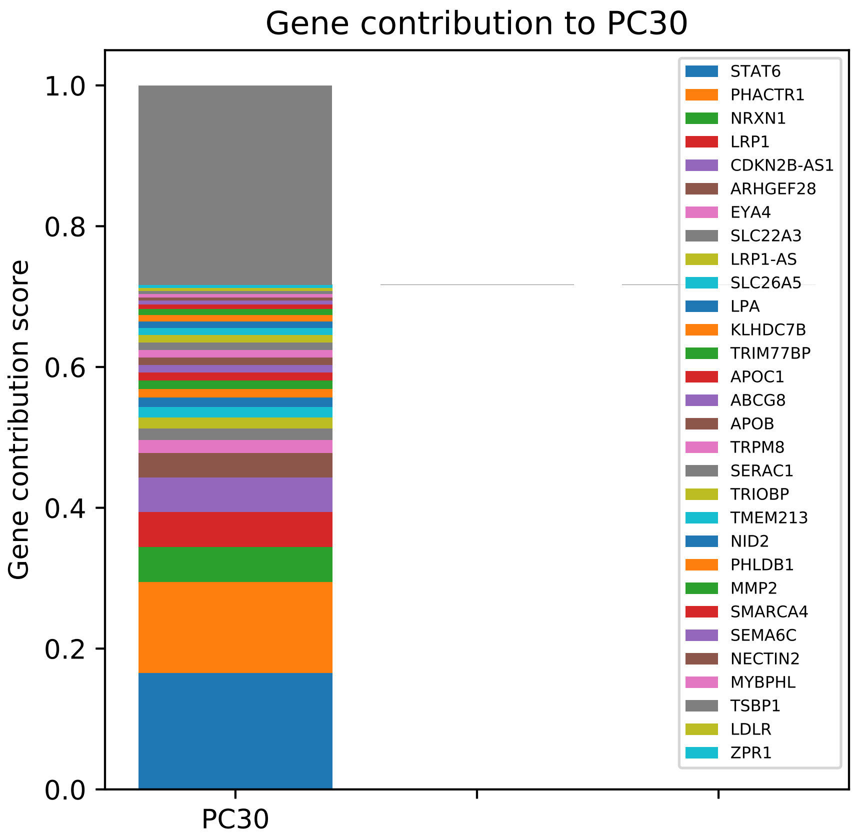
Gene contribution to PC30
Gene contribution score
1.0
0.8
0.6
0.4
0.2
0.0
PC30
STAT6
PHACTR1
NRXN1
LRP1
CDKN2B-AS1
ARHGEF28
EYA4
SLC22A3
LRP1-AS
SLC26A5
LPA
KLHDC7B
TRIM77BP
APOC1
ABCG8
APOB
TRPM8
SERAC1
TRIOBP
TMEM213
NID2
PHLDB1
MMP2
SMARCA4
SEMA6C
NECTIN2
MYBPHL
TSBP1
LDLR
ZPR1

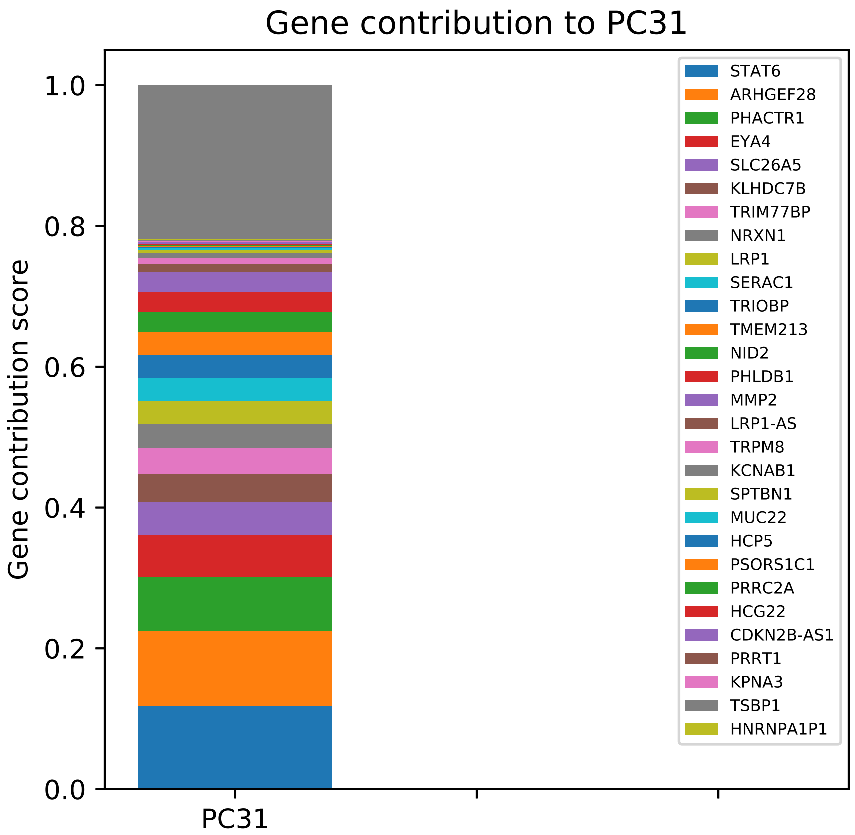

Gene contribution to PC31
Gene contribution score
1.0
0.8
0.6
0.4
0.2
0.0
PC31
STAT6
ARHGEF28
PHACTR1
EYA4
SLC26A5
KLHDC7B
TRIM77BP
NRXN1
LRP1
SERAC1
TRIOBP
TMEM213
NID2
PHLDB1
MMP2
LRP1-AS
TRPM8
KCNAB1
SPTBN1
MUC22
HCP5
PSORS1C1
PRRC2A
HCG22
CDKN2B-AS1
PRRT1
KPNA3
TSBP1
HNRNPA1P1

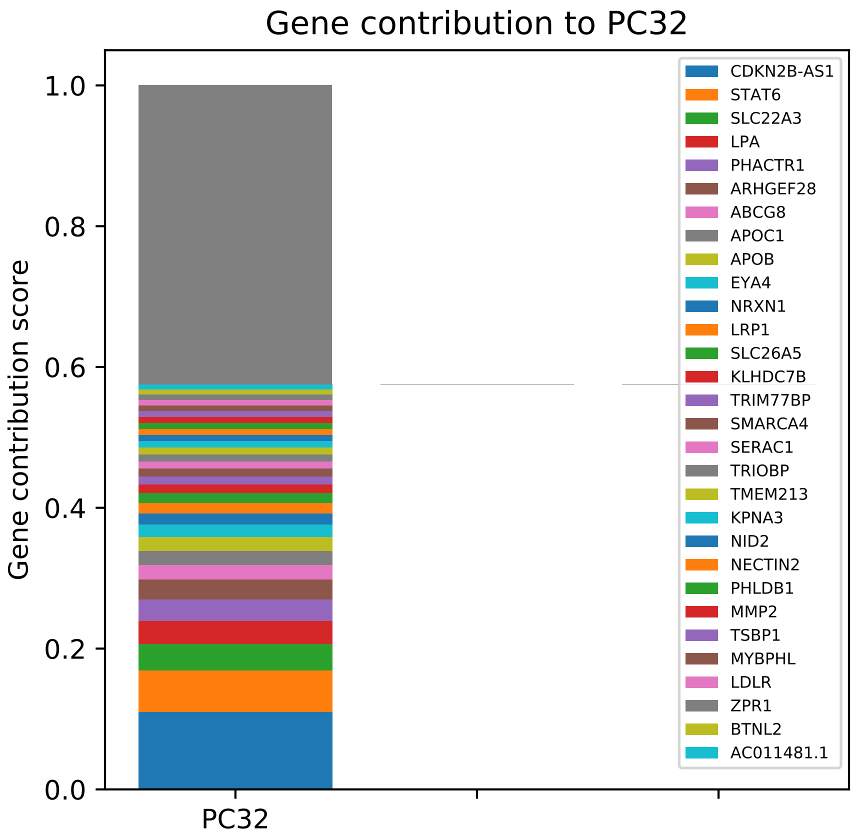

Gene contribution to PC32
Gene contribution score
1.0
0.8
0.6
0.4
0.2
0.0
PC32
CDKN2B-AS1
STAT6
SLC22A3
LPA
PHACTR1
ARHGEF28
ABCG8
APOC1
APOB
EYA4
NRXN1
LRP1
SLC26A5
KLHDC7B
TRIM77BP
SMARCA4
SERAC1
TRIOBP
TMEM213
KPNA3
NID2
NECTIN2
PHLDB1
MMP2
TSBP1
MYBPHL
LDLR
ZPR1
BTNL2
AC011481.1

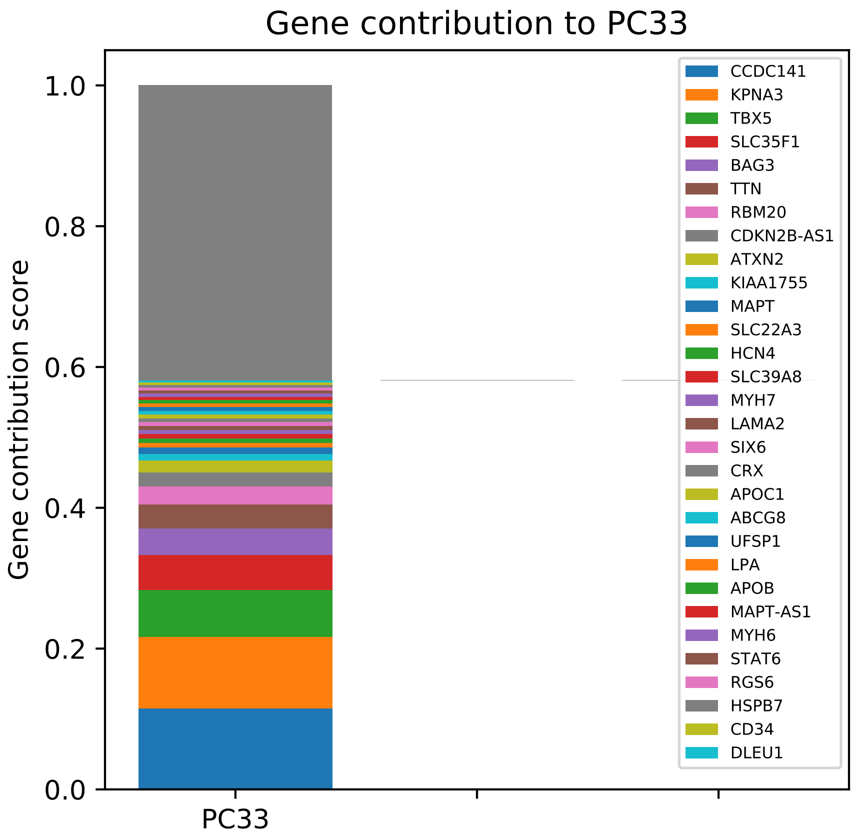
Gene contribution to PC33
Gene contribution score
1.0
0.8
0.6
0.4
0.2
0.0
PC33
CCDC141
KPNA3
TBX5
SLC35F1
BAG3
TTN
RBM20
CDKN2B-AS1
ATXN2
KIAA1755
MAPT
SLC22A3
HCN4
SLC39A8
MYH7
LAMA2
SIX6
CRX
APOC1
ABCG8
UFSP1
LPA
APOB
MAPT-AS1
MYH6
STAT6
RGS6
HSPB7
CD34
DLEU1

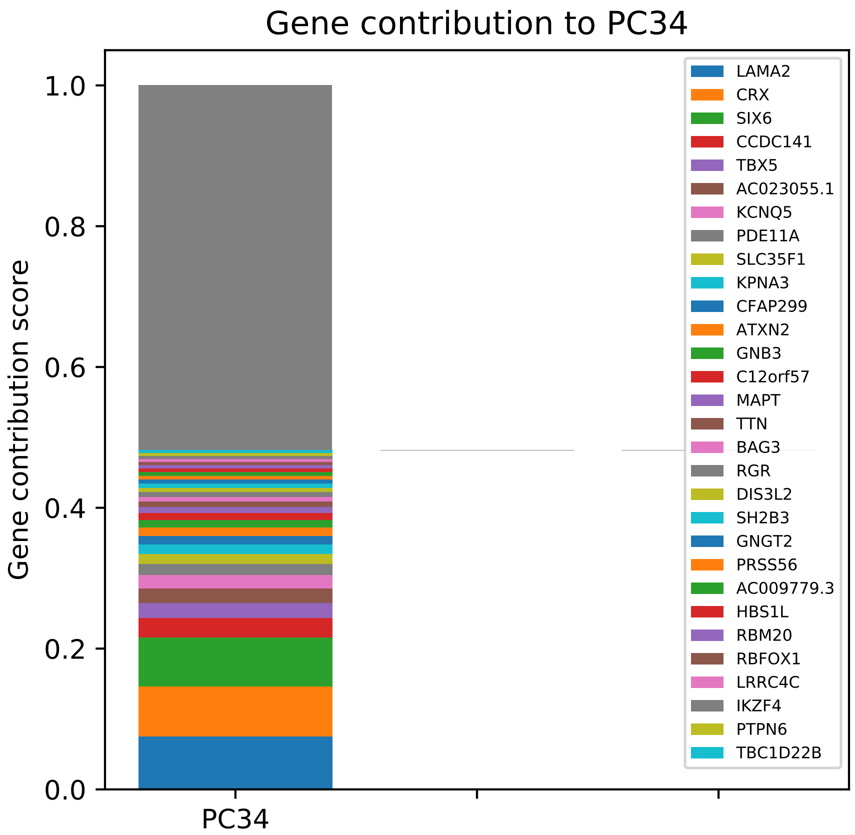
Gene contribution to PC34
Gene contribution score
1.0
0.8
0.6
0.4
0.2
0.0
PC34
LAMA2
CRX
SIX6
CCDC141
TBX5
AC023055.1
KCNQ5
PDE11A
SLC35F1
KPNA3
CFAP299
ATXN2
GNB3
C12orf57
MAPT
TTN
BAG3
RGR
DIS3L2
SH2B3
GNGT2
PRSS56
AC009779.3
HBS1L
RBM20
RBFOX1
LRRC4C
IKZF4
PTPN6
TBC1D22B

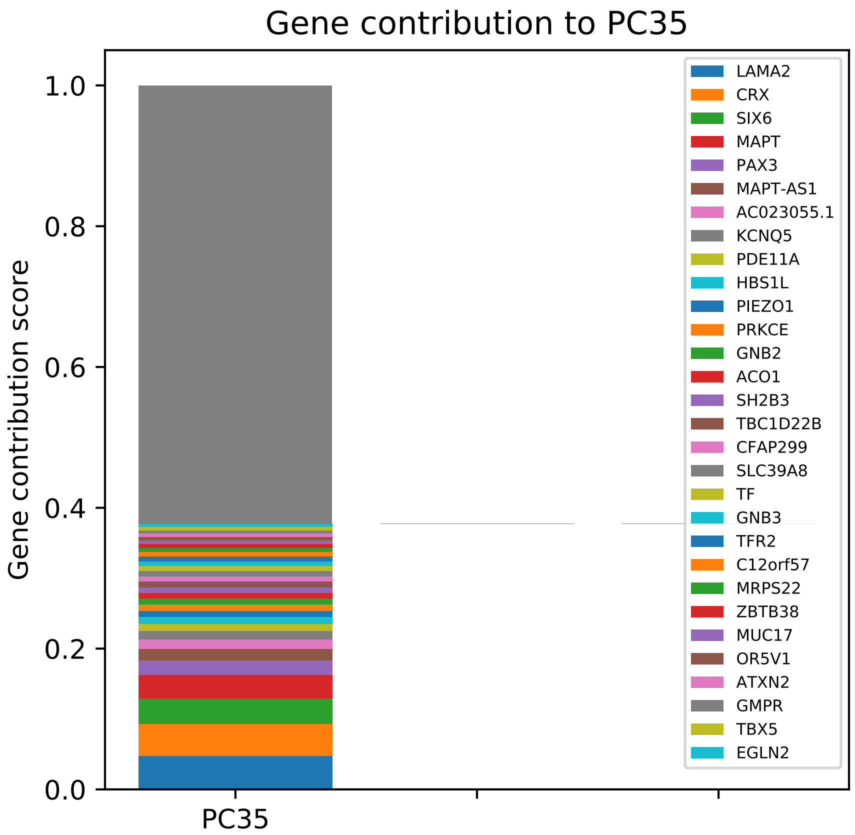

Gene contribution to PC35
Gene contribution score
1.0
0.8
0.6
0.4
0.2
0.0
PC35
LAMA2
CRX
SIX6
MAPT
PAX3
MAPT-AS1
AC023055.1
KCNQ5
PDE11A
HBS1L
PIEZO1
PRKCE
GNB2
ACO1
SH2B3
TBC1D22B
CFAP299
SLC39A8
TF
GNB3
TFR2
C12orf57
MRPS22
ZBTB38
MUC17
OR5V1
ATXN2
GMPR
TBX5
EGLN2

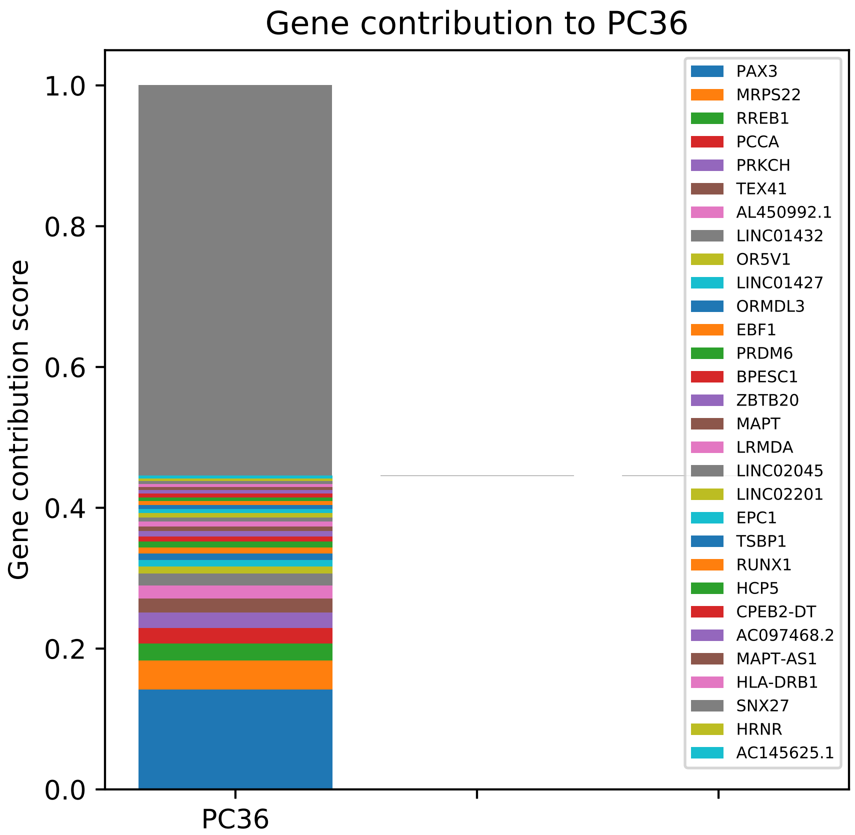

Gene contribution to PC36
Gene contribution score
1.0
0.8
0.6
0.4
0.2
0.0
PC36
PAX3
MRPS22
RREB1
PCCA
PRKCH
TEX41
AL450992.1
LINC01432
OR5V1
LINC01427
ORMDL3
EBF1
PRDM6
BPESC1
ZBTB20
MAPT
LRMDA
LINC02045
LINC02201
EPC1
TSBP1
RUNX1
HCP5
CPEB2-DT
AC097468.2
MAPT-AS1
HLA-DRB1
SNX27
HRNR
AC145625.1

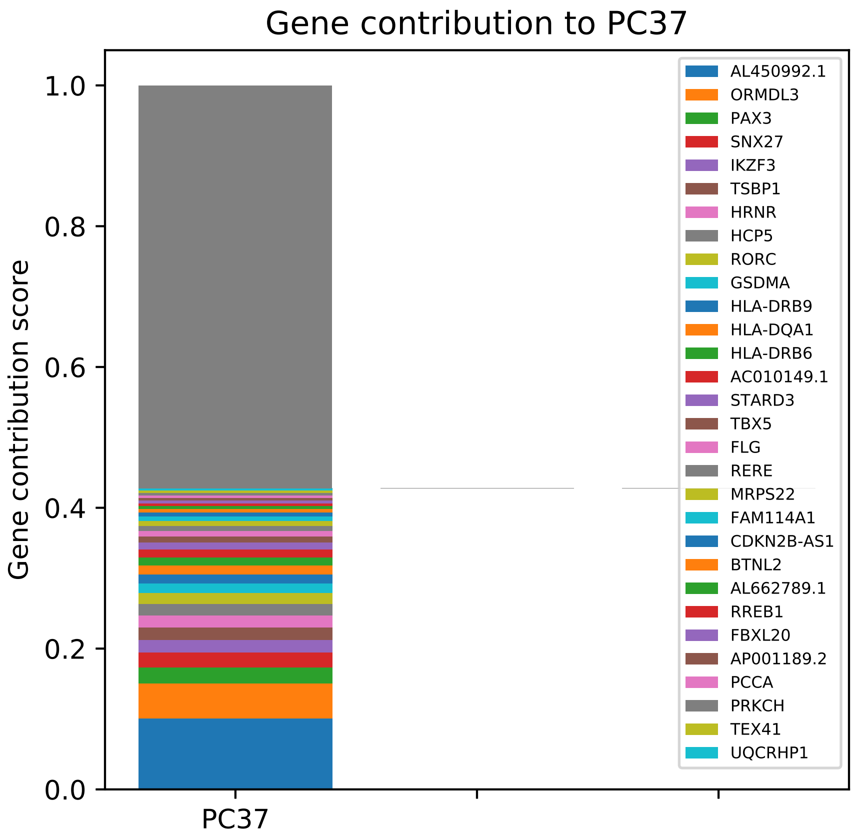
Gene contribution to PC37
Gene contribution score
1.0
0.8
0.6
0.4
0.2
0.0
PC37
AL450992.1
ORMDL3
PAX3
SNX27
IKZF3
TSBP1
HRNR
HCP5
RORC
GSDMA
HLA-DRB9
HLA-DQA1
HLA-DRB6
AC010149.1
STARD3
TBX5
FLG
RERE
MRPS22
FAM114A1
CDKN2B-AS1
BTNL2
AL662789.1
RREB1
FBXL20
AP001189.2
PCCA
PRKCH
TEX41
UQCRHP1

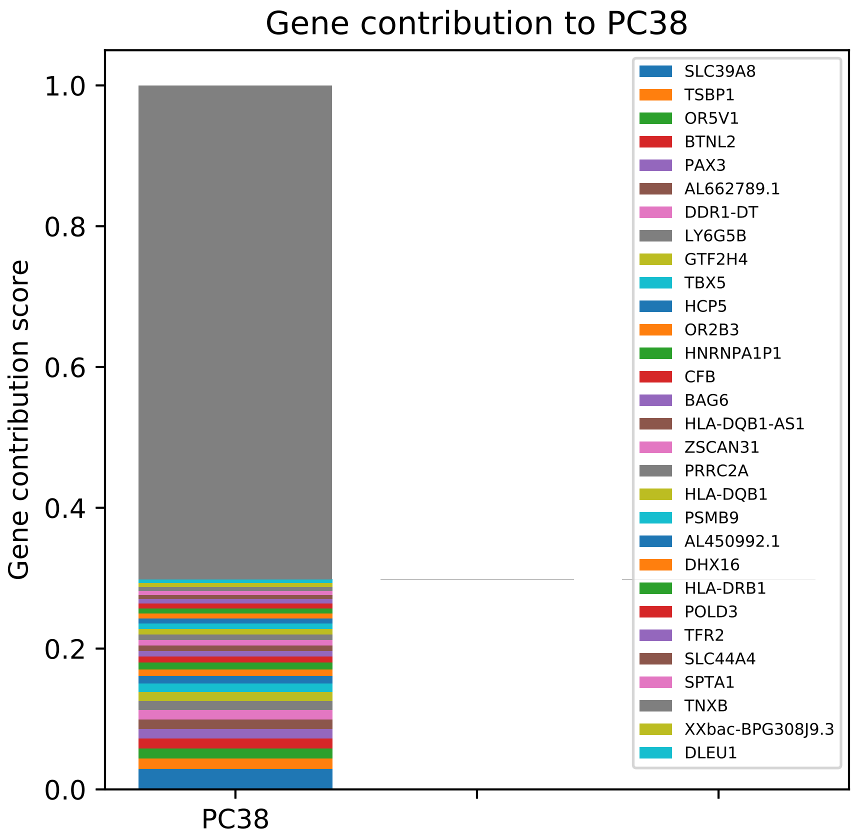

Gene contribution to PC38
Gene contribution score
1.0
0.8
0.6
0.4
0.2
0.0
PC38
SLC39A8
TSBP1
OR5V1
BTNL2
PAX3
AL662789.1
DDR1-DT
LY6G5B
GTF2H4
TBX5
HCP5
OR2B3
HNRNPA1P1
CFB
BAG6
HLA-DQB1-AS1
ZSCAN31
PRRC2A
HLA-DQB1
PSMB9
AL450992.1
DHX16
HLA-DRB1
POLD3
TFR2
SLC44A4
SPTA1
TNXB
XXbac-BPG308J9.3
DLEU1

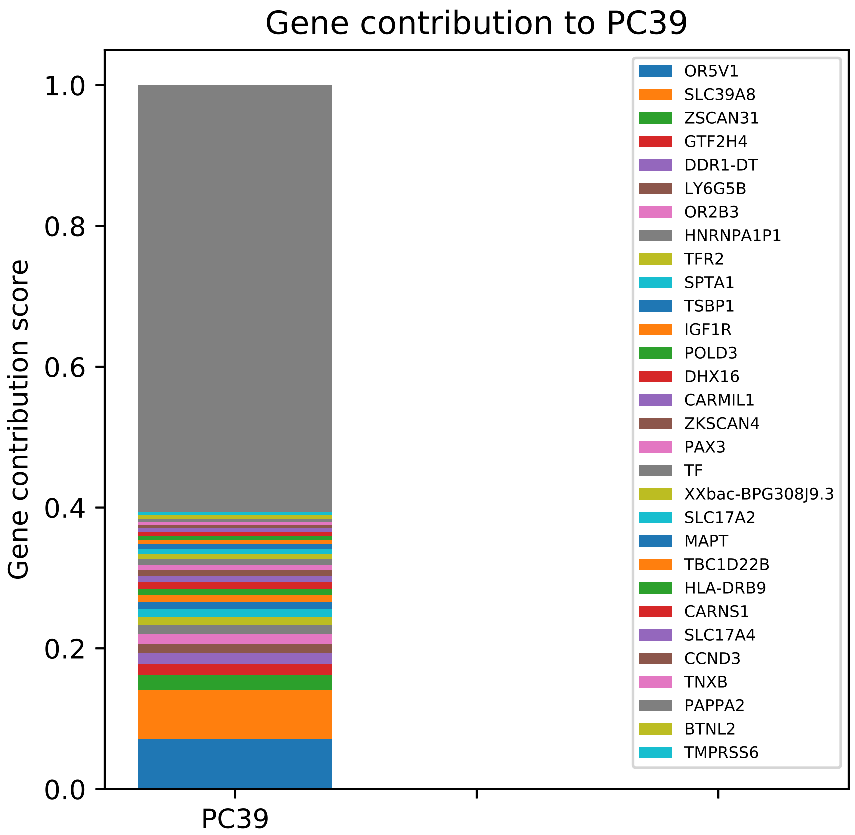

Gene contribution to PC39
Gene contribution score
1.0
0.8
0.6
0.4
0.2
0.0
PC39
OR5V1
SLC39A8
ZSCAN31
GTF2H4
DDR1-DT
LY6G5B
OR2B3
HNRNPA1P1
TFR2
SPTA1
TSBP1
IGF1R
POLD3
DHX16
CARMIL1
ZKSCAN4
PAX3
TF
XXbac-BPG308J9.3
SLC17A2
MAPT
TBC1D22B
HLA-DRB9
CARNS1
SLC17A4
CCND3
TNXB
PAPPA2
BTNL2
TMPRSS6

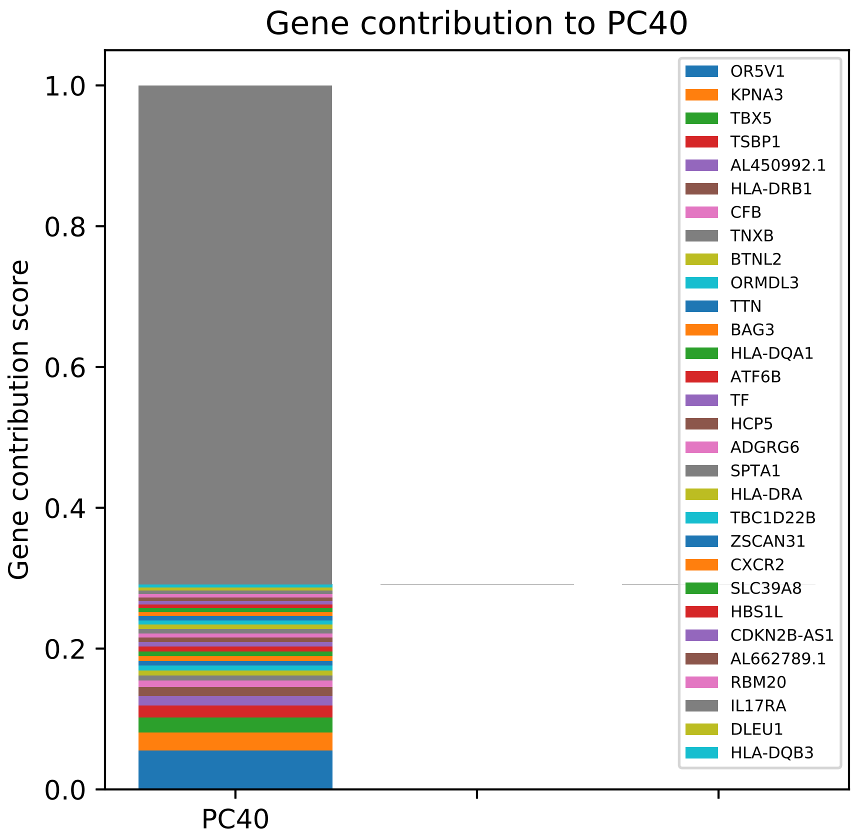
Gene contribution to PC40
Gene contribution score
1.0
0.8
0.6
0.4
0.2
0.0
PC40
OR5V1
KPNA3
TBX5
TSBP1
AL450992.1
HLA-DRB1
CFB
TNXB
BTNL2
ORMDL3
TTN
BAG3
HLA-DQA1
ATF6B
TF
HCP5
ADGRG6
SPTA1
HLA-DRA
TBC1D22B
ZSCAN31
CXCR2
SLC39A8
HBS1L
CDKN2B-AS1
AL662789.1
RBM20
IL17RA
DLEU1
HLA-DQB3

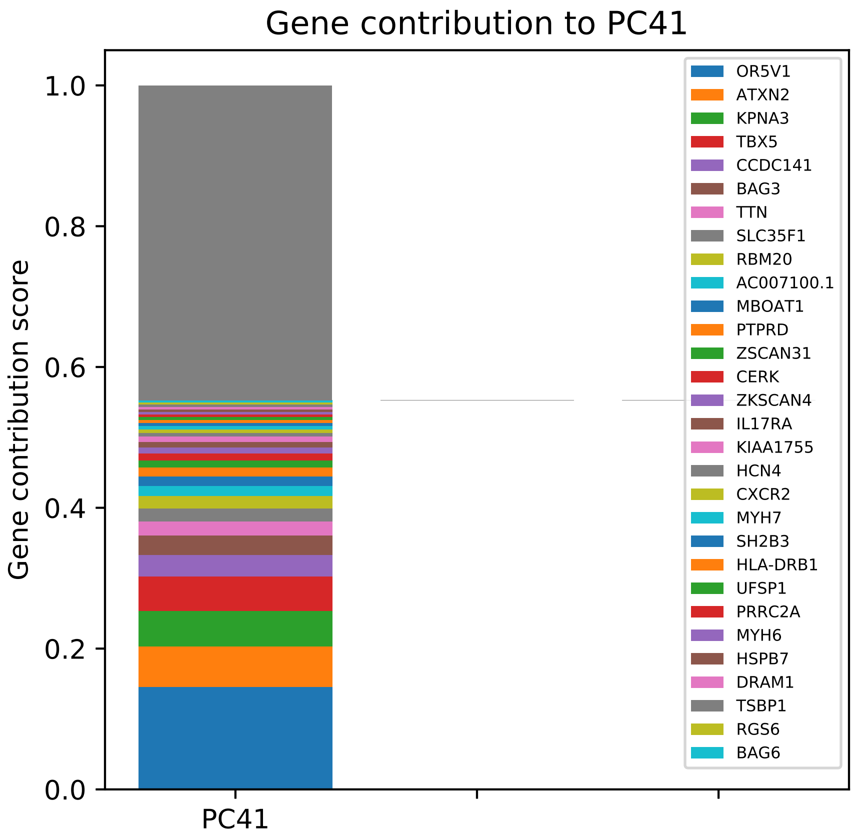
Gene contribution to PC41
Gene contribution score
1.0
0.8
0.6
0.4
0.2
0.0
PC41
OR5V1
ATXN2
KPNA3
TBX5
CCDC141
BAG3
TTN
SLC35F1
RBM20
AC007100.1
MBOAT1
PTPRD
ZSCAN31
CERK
ZKSCAN4
IL17RA
KIAA1755
HCN4
CXCR2
MYH7
SH2B3
HLA-DRB1
UFSP1
PRRC2A
MYH6
HSPB7
DRAM1
TSBP1
RGS6
BAG6

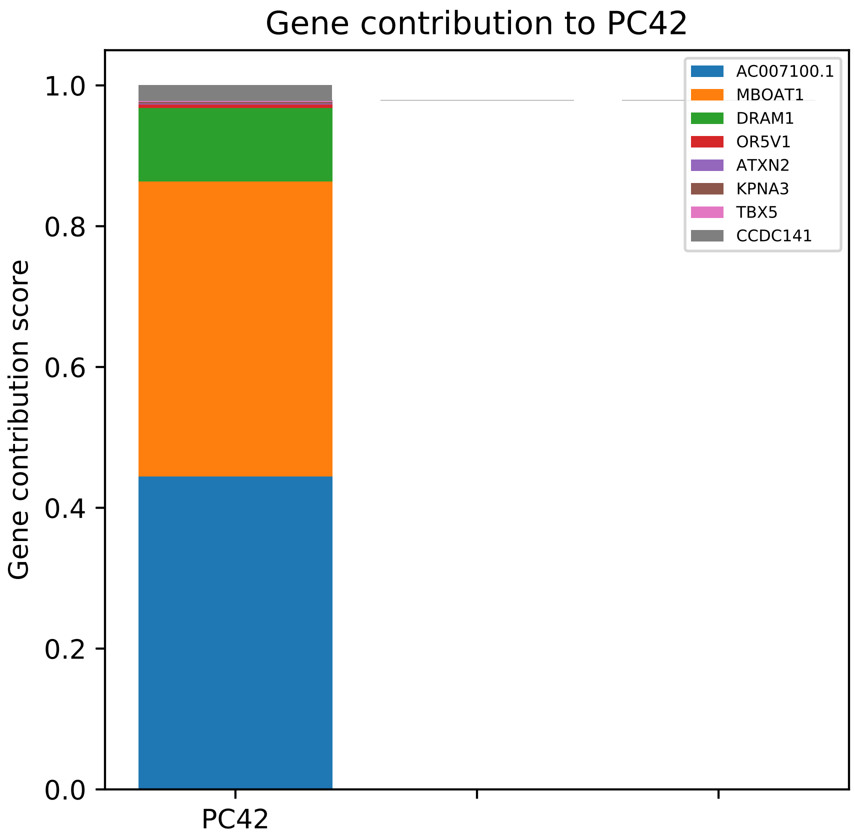

Gene contribution to PC42
Gene contribution score
1.0
0.8
0.6
0.4
0.2
0.0
PC42
AC007100.1
MBOAT1
DRAM1
OR5V1
ATXN2
KPNA3
TBX5
CCDC141

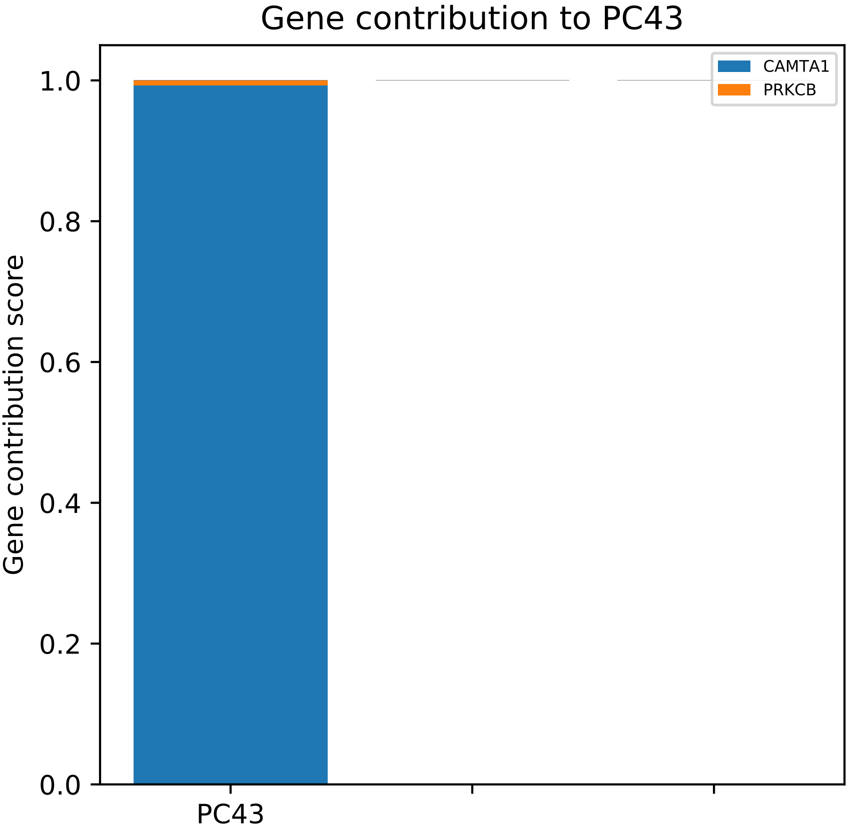
Gene contribution to PC43
Gene contribution score
1.0
0.8
0.6
0.4
0.2
0.0
PC43
CAMTA1
PRKCB

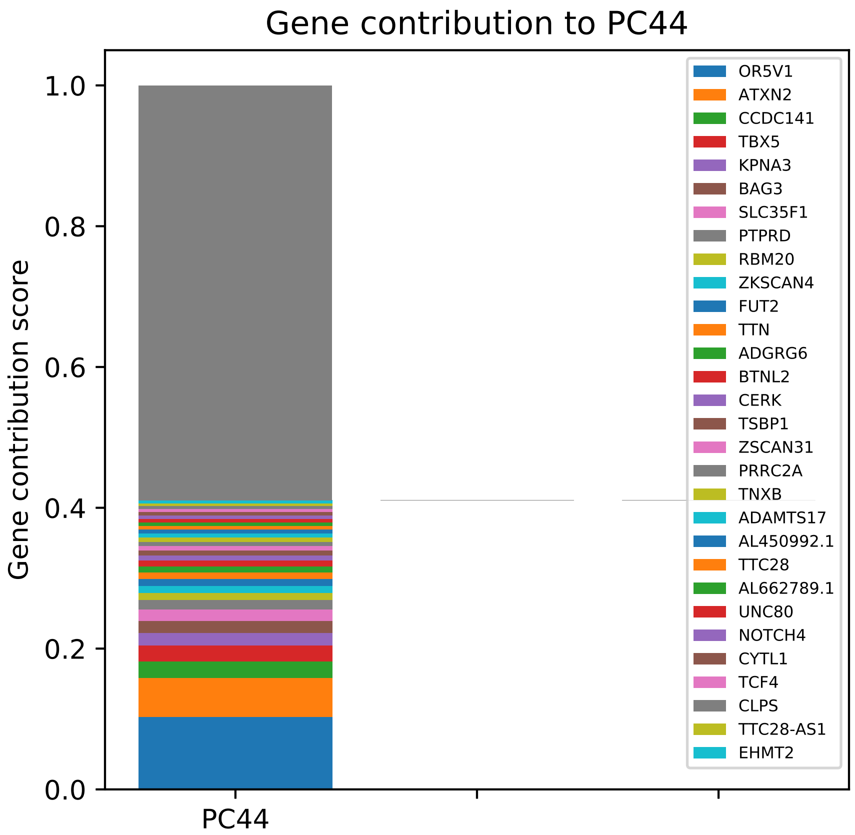

Gene contribution to PC44
Gene contribution score
1.0
0.8
0.6
0.4
0.2
0.0
PC44
OR5V1
ATXN2
CCDC141
TBX5
KPNA3
BAG3
SLC35F1
PTPRD
RBM20
ZKSCAN4
FUT2
TTN
ADGRG6
BTNL2
CERK
TSBP1
ZSCAN31
PRRC2A
TNXB
ADAMTS17
AL450992.1
TTC28
AL662789.1
UNC80
NOTCH4
CYTL1
TCF4
CLPS
TTC28-AS1
EHMT2

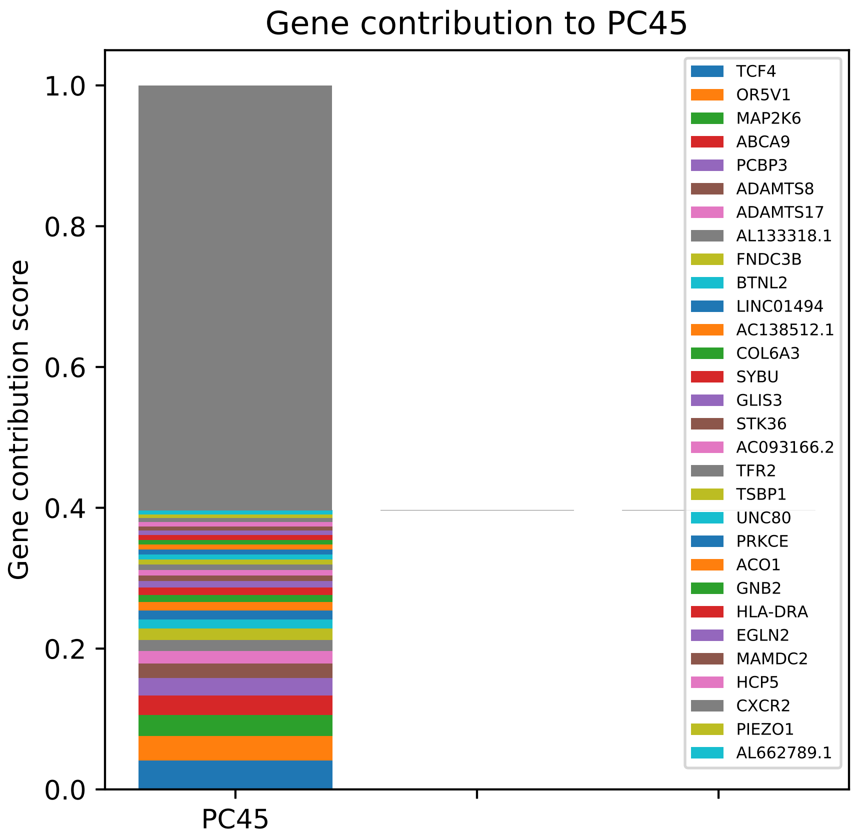

Gene contribution to PC45
Gene contribution score
1.0
0.8
0.6
0.4
0.2
0.0
PC45
TCF4
OR5V1
MAP2K6
ABCA9
PCBP3
ADAMTS8
ADAMTS17
AL133318.1
FNDC3B
BTNL2
LINC01494
AC138512.1
COL6A3
SYBU
GLIS3
STK36
AC093166.2
TFR2
TSBP1
UNC80
PRKCE
ACO1
GNB2
HLA-DRA
EGLN2
MAMDC2
HCP5
CXCR2
PIEZO1
AL662789.1

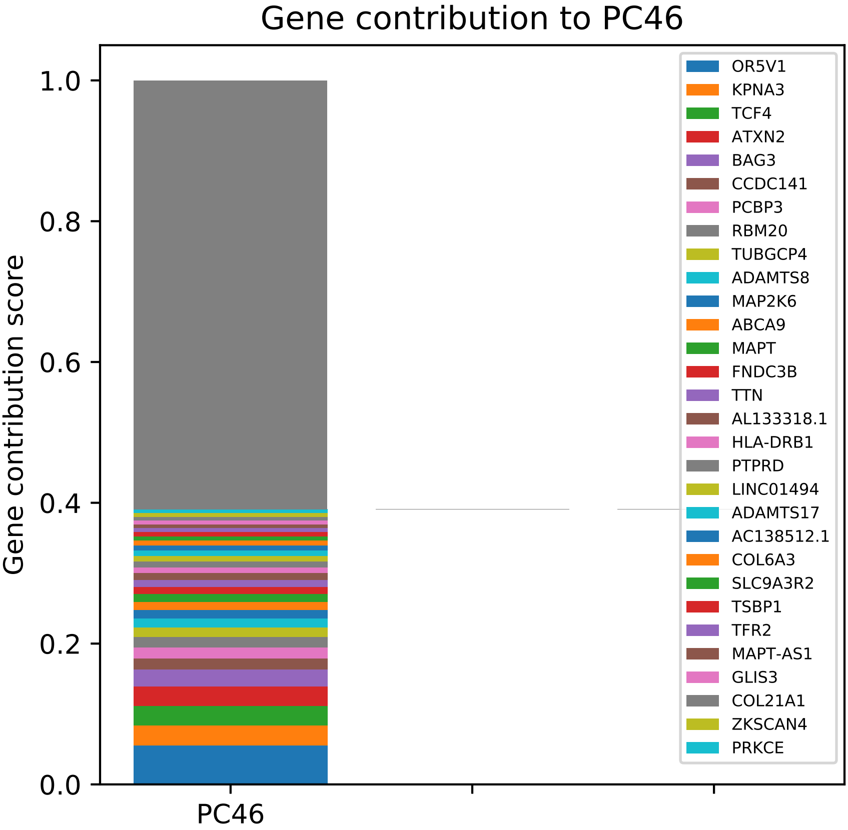

Gene contribution to PC46
Gene contribution score
0.0
0.2
0.4
0.6
0.8
1.0
PC46
OR5V1
KPNA3
TCF4
ATXN2
BAG3
CCDC141
PCBP3
RBM20
TUBGCP4
ADAMTS8
MAP2K6
ABCA9
MAPT
FNDC3B
TTN
AL133318.1
HLA-DRB1
PTPRD
LINC01494
ADAMTS17
AC138512.1
COL6A3
SLC9A3R2
TSBP1
TFR2
MAPT-AS1
GLIS3
COL21A1
ZKSCAN4
PRKCE

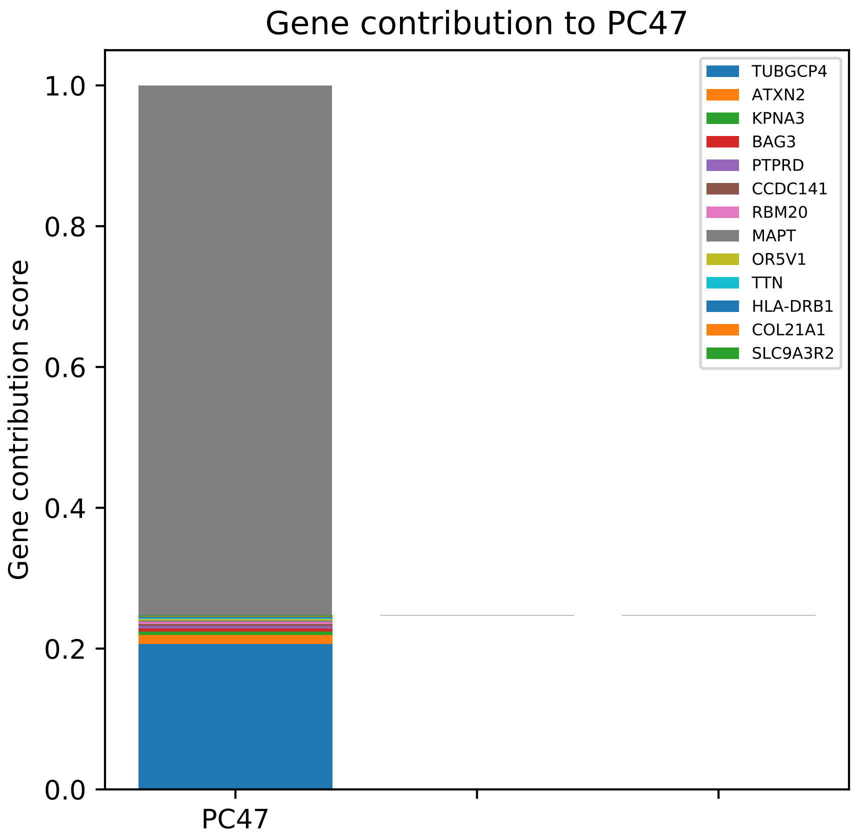
Gene contribution to PC47
Gene contribution score
1.0
0.8
0.6
0.4
0.2
0.0
PC47
TUBGCP4
ATXN2
KPNA3
BAG3
PTPRD
CCDC141
RBM20
MAPT
OR5V1
TTN
HLA-DRB1
COL21A1
SLC9A3R2

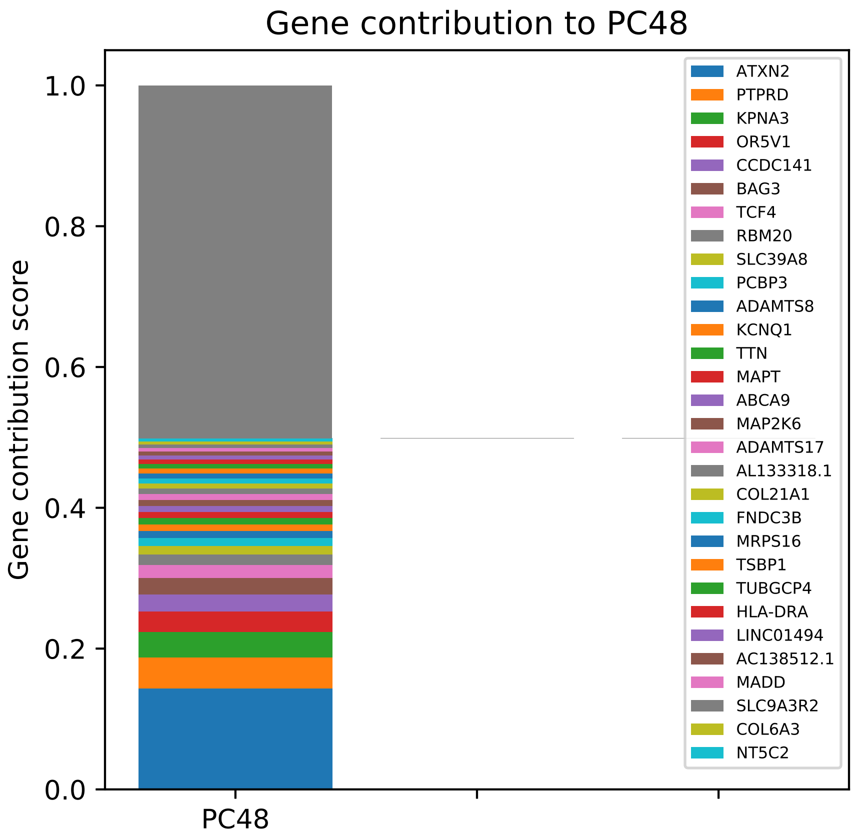

Gene contribution to PC48
Gene contribution score
1.0
0.8
0.6
0.4
0.2
0.0
PC48
ATXN2
PTPRD
KPNA3
OR5V1
CCDC141
BAG3
TCF4
RBM20
SLC39A8
PCBP3
ADAMTS8
KCNQ1
TTN
MAPT
ABCA9
MAP2K6
ADAMTS17
AL133318.1
COL21A1
FNDC3B
MRPS16
TSBP1
TUBGCP4
HLA-DRA
LINC01494
AC138512.1
MADD
SLC9A3R2
COL6A3
NT5C2

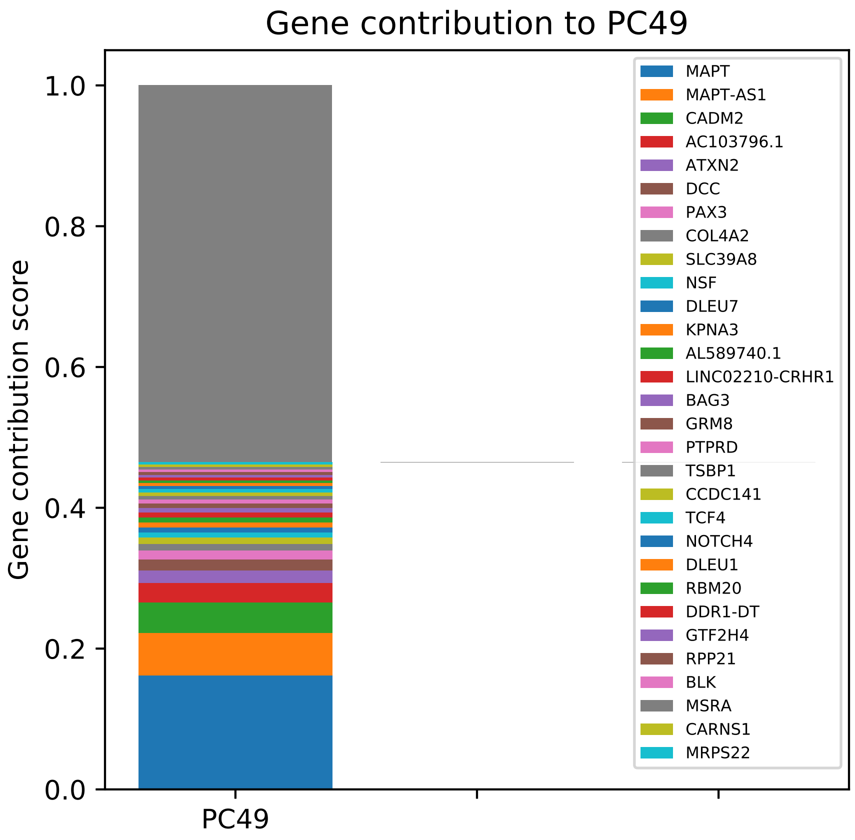
Gene contribution to PC49
Gene contribution score
1.0
0.8
0.6
0.4
0.2
0.0
PC49
MAPT
MAPT-AS1
CADM2
AC103796.1
ATXN2
DCC
PAX3
COL4A2
SLC39A8
NSF
DLEU7
KPNA3
AL589740.1
LINC02210-CRHR1
BAG3
GRM8
PTPRD
TSBP1
CCDC141
TCF4
NOTCH4
DLEU1
RBM20
DDR1-DT
GTF2H4
RPP21
BLK
MSRA
CARNS1
MRPS22

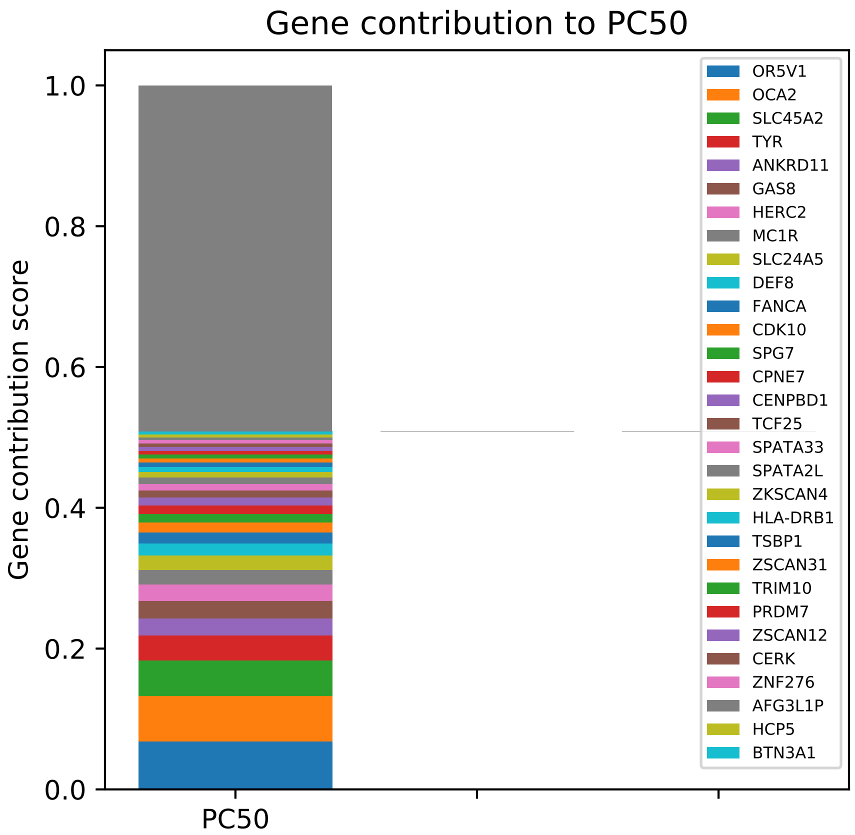
Gene contribution to PC50
Gene contribution score
1.0
0.8
0.6
0.4
0.2
0.0
PC50
OR5V1
OCA2
SLC45A2
TYR
ANKRD11
GAS8
HERC2
MC1R
SLC24A5
DEF8
FANCA
CDK10
SPG7
CPNE7
CENPBD1
TCF25
SPATA33
SPATA2L
ZKSCAN4
HLA-DRB1
TSBP1
ZSCAN31
TRIM10
PRDM7
ZSCAN12
CERK
ZNF276
AFG3L1P
HCP5
BTN3A1

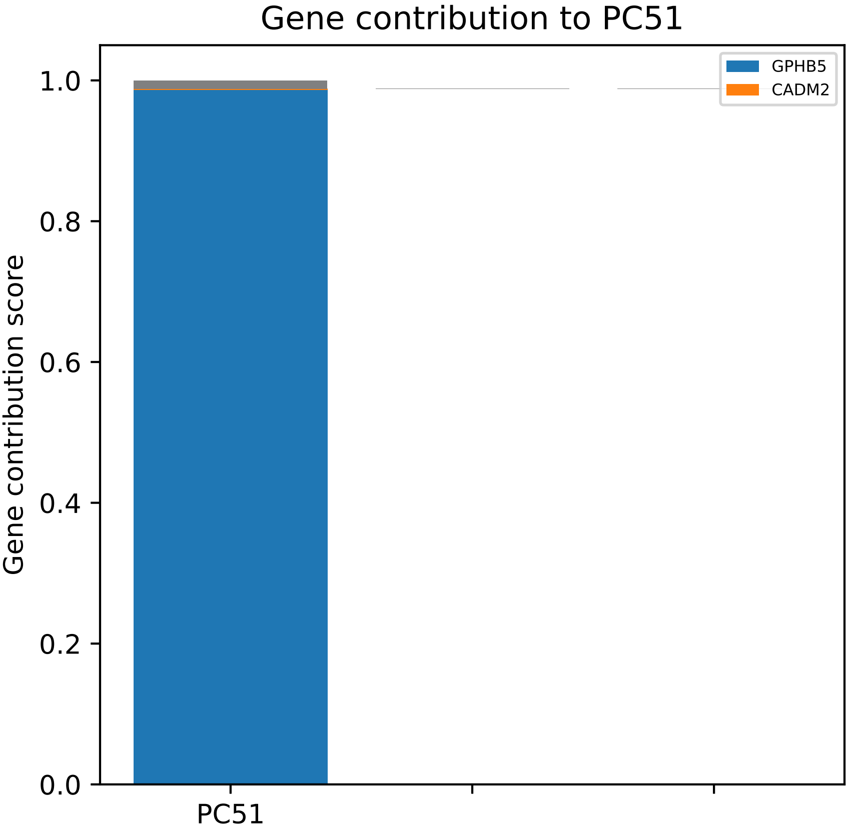
Gene contribution to PC51
Gene contribution score
1.0
0.8
0.6
0.4
0.2
0.0
PC51
GPHB5
CADM2

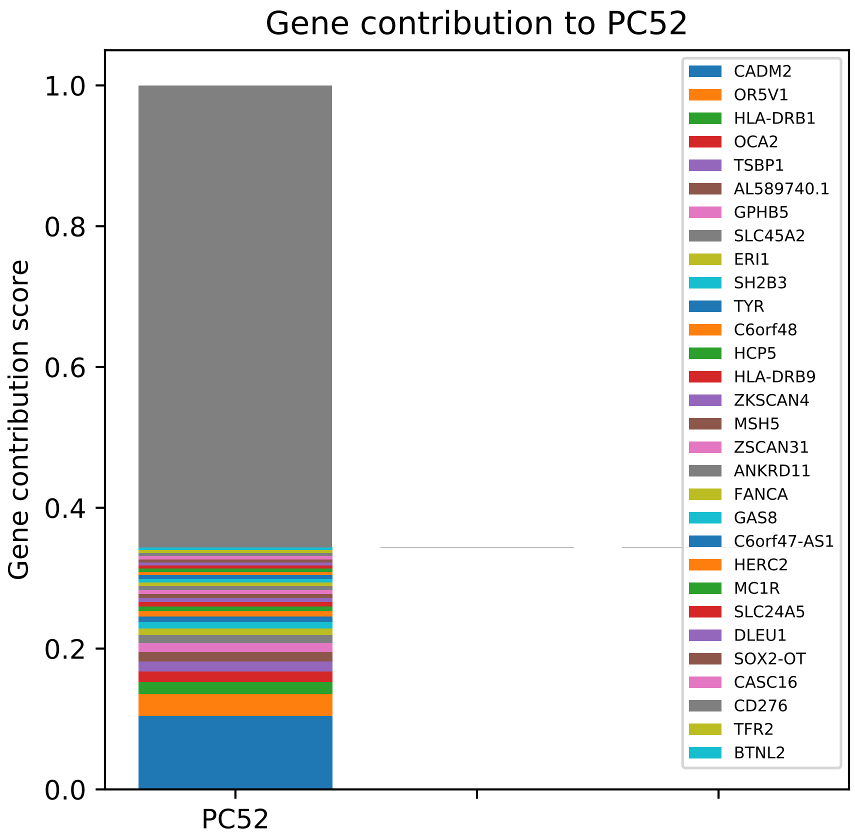

Gene contribution to PC52
Gene contribution score
1.0
0.8
0.6
0.4
0.2
0.0
PC52
CADM2
OR5V1
HLA-DRB1
OCA2
TSBP1
AL589740.1
GPHB5
SLC45A2
ERI1
SH2B3
TYR
C6orf48
HCP5
HLA-DRB9
ZKSCAN4
MSH5
ZSCAN31
ANKRD11
FANCA
GAS8
C6orf47-AS1
HERC2
MC1R
SLC24A5
DLEU1
SOX2-OT
CASC16
CD276
TFR2
BTNL2

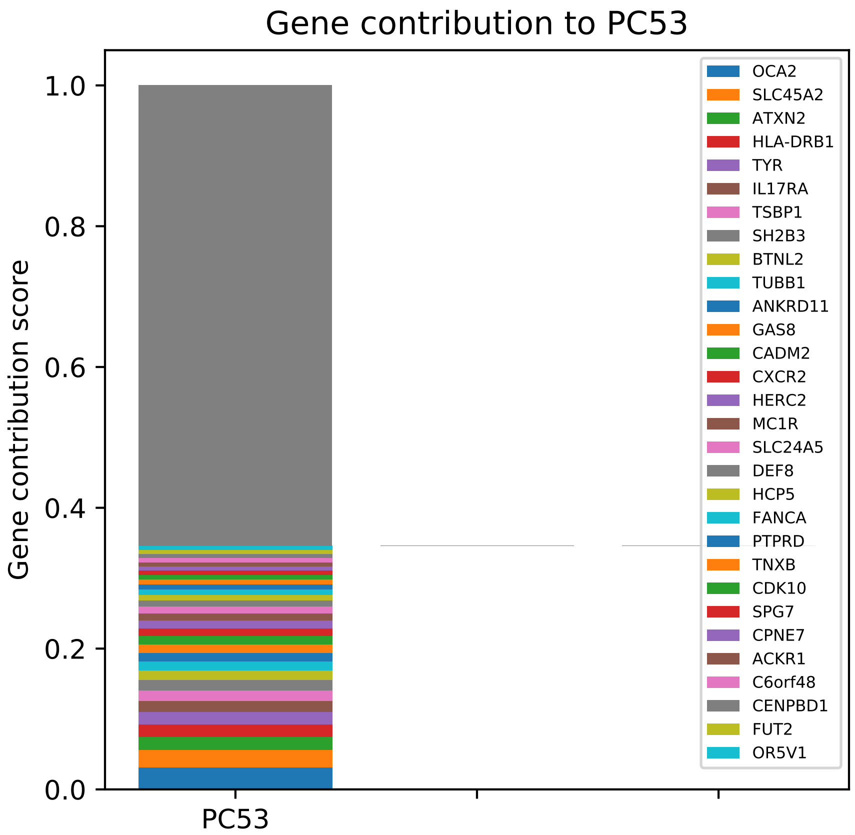

Gene contribution to PC53
Gene contribution score
1.0
0.8
0.6
0.4
0.2
0.0
PC53
OCA2
SLC45A2
ATXN2
HLA-DRB1
TYR
IL17RA
TSBP1
SH2B3
BTNL2
TUBB1
ANKRD11
GAS8
CADM2
CXCR2
HERC2
MC1R
SLC24A5
DEF8
HCP5
FANCA
PTPRD
TNXB
CDK10
SPG7
CPNE7
ACKR1
C6orf48
CENPBD1
FUT2
OR5V1

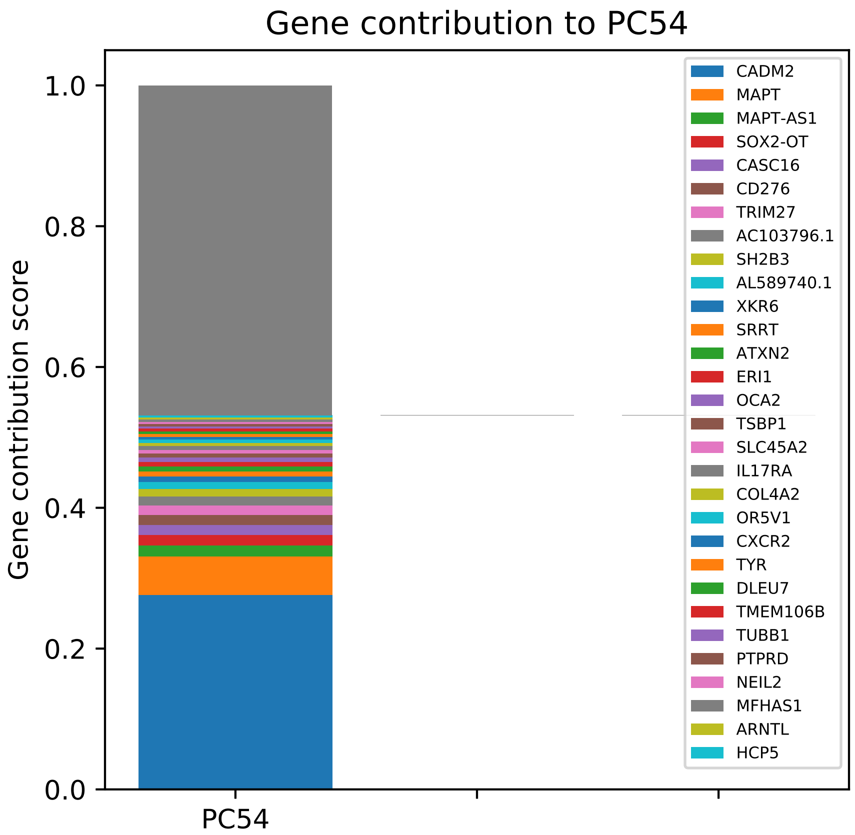

Gene contribution to PC54
Gene contribution score
0.0
0.2
0.4
0.6
0.8
1.0
PC54
CADM2
MAPT
MAPT-AS1
SOX2-OT
CASC16
CD276
TRIM27
AC103796.1
SH2B3
AL589740.1
XKR6
SRRT
ATXN2
ERI1
OCA2
TSBP1
SLC45A2
IL17RA
COL4A2
OR5V1
CXCR2
TYR
DLEU7
TMEM106B
TUBB1
PTPRD
NEIL2
MFHAS1
ARNTL
HCP5

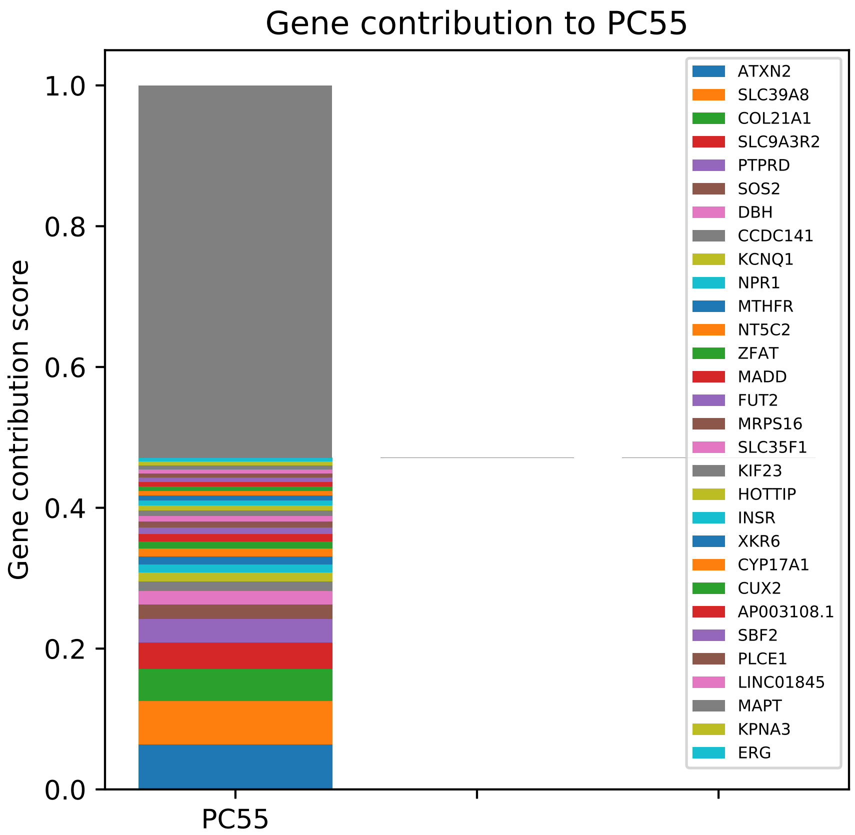

Gene contribution to PC55
Gene contribution score
1.0
0.8
0.6
0.4
0.2
0.0
PC55
ATXN2
SLC39A8
COL21A1
SLC9A3R2
PTPRD
SOS2
DBH
CCDC141
KCNQ1
NPR1
MTHFR
NT5C2
ZFAT
MADD
FUT2
MRPS16
SLC35F1
KIF23
HOTTIP
INSR
XKR6
CYP17A1
CUX2
AP003108.1
SBF2
PLCE1
LINC01845
MAPT
KPNA3
ERG

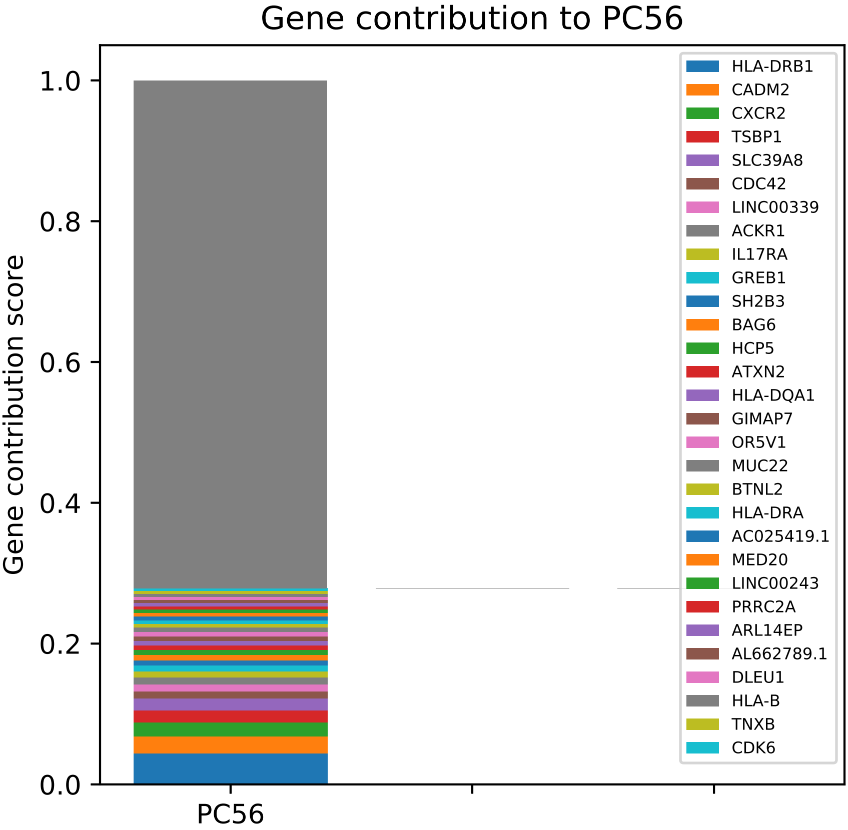
Gene contribution to PC56
Gene contribution score
1.0
0.8
0.6
0.4
0.2
0.0
PC56
HLA-DRB1
CADM2
CXCR2
TSBP1
SLC39A8
CDC42
LINC00339
ACKR1
IL17RA
GREB1
SH2B3
BAG6
HCP5
ATXN2
HLA-DQA1
GIMAP7
OR5V1
MUC22
BTNL2
HLA-DRA
AC025419.1
MED20
LINC00243
PRRC2A
ARL14EP
AL662789.1
DLEU1
HLA-B
TNXB
CDK6

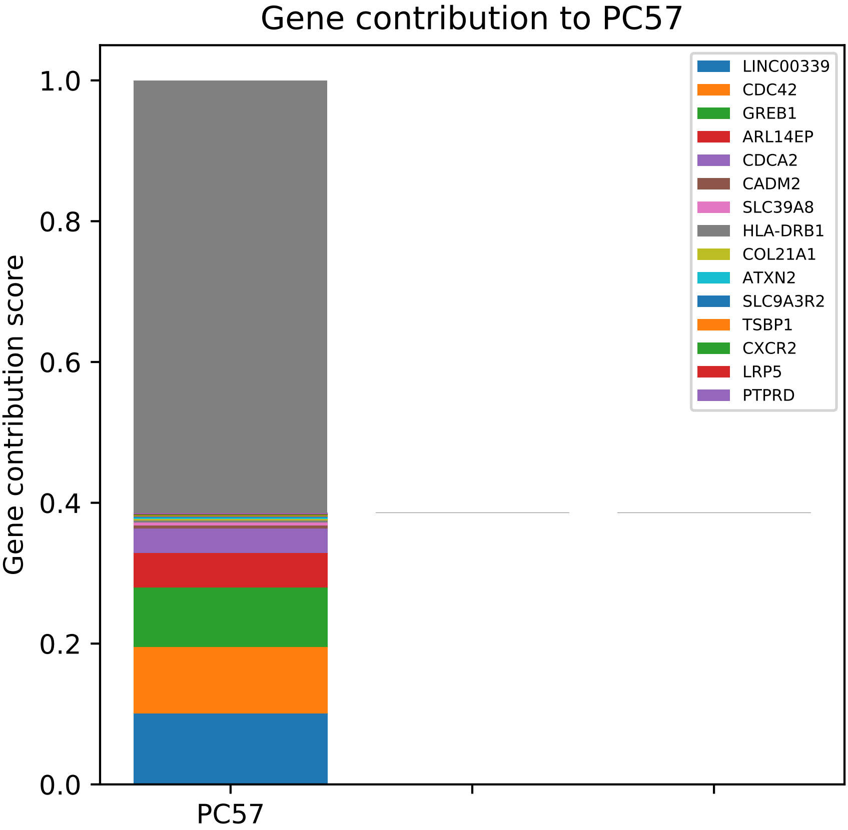

Gene contribution to PC57
Gene contribution score
1.0
0.8
0.6
0.4
0.2
0.0
PC57
LINC00339
CDC42
GREB1
ARL14EP
CDCA2
CADM2
SLC39A8
HLA-DRB1
COL21A1
ATXN2
SLC9A3R2
TSBP1
CXCR2
LRP5
PTPRD

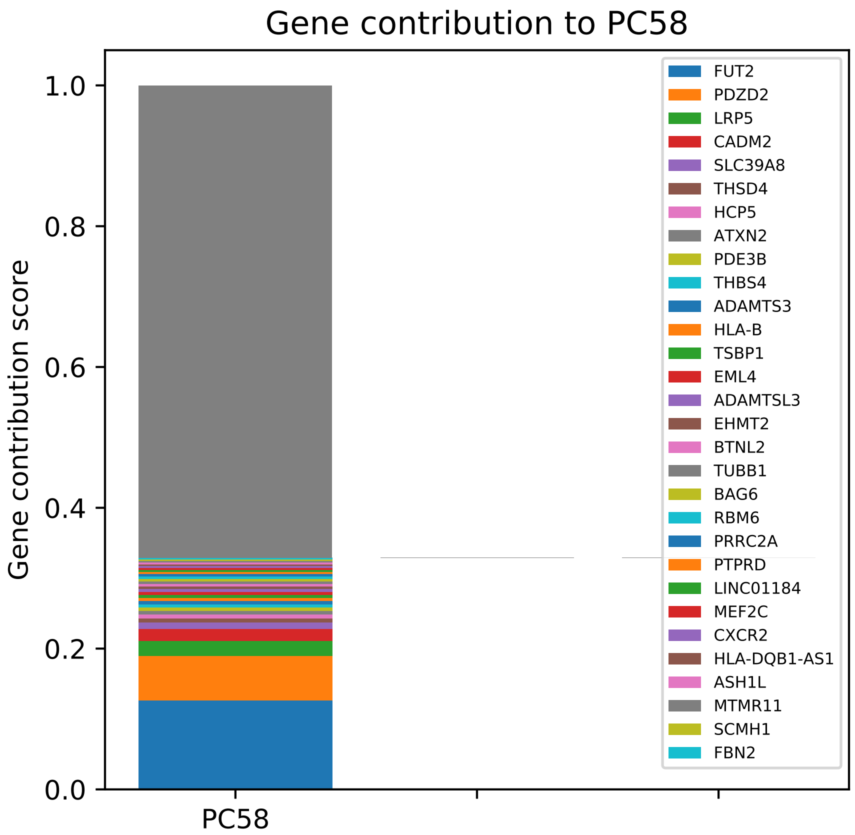

Gene contribution to PC58
Gene contribution score
1.0
0.8
0.6
0.4
0.2
0.0
PC58
FUT2
PDZD2
LRP5
CADM2
SLC39A8
THSD4
HCP5
ATXN2
PDE3B
THBS4
ADAMTS3
HLA-B
TSBP1
EML4
ADAMTSL3
EHMT2
BTNL2
TUBB1
BAG6
RBM6
PRRC2A
PTPRD
LINC01184
MEF2C
CXCR2
HLA-DQB1-AS1
ASH1L
MTMR11
SCMH1
FBN2

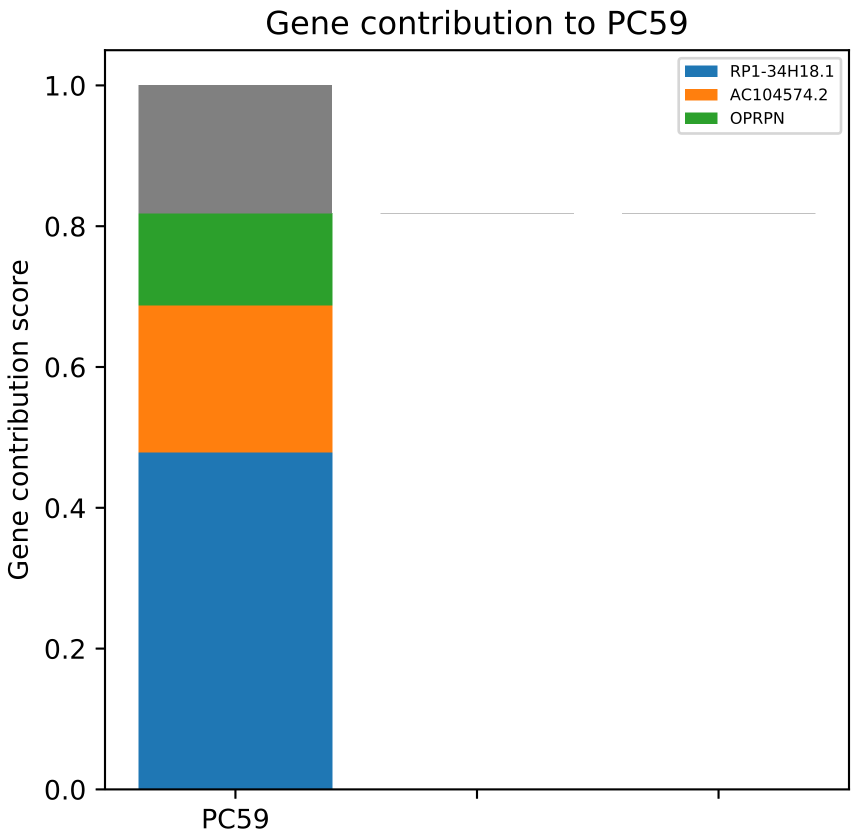

Gene contribution to PC59
RP1-34H18.1
AC104574.2
OPRPN
1.0
0.8
0.6
0.4
0.2
0.0
Gene contribution score
PC59

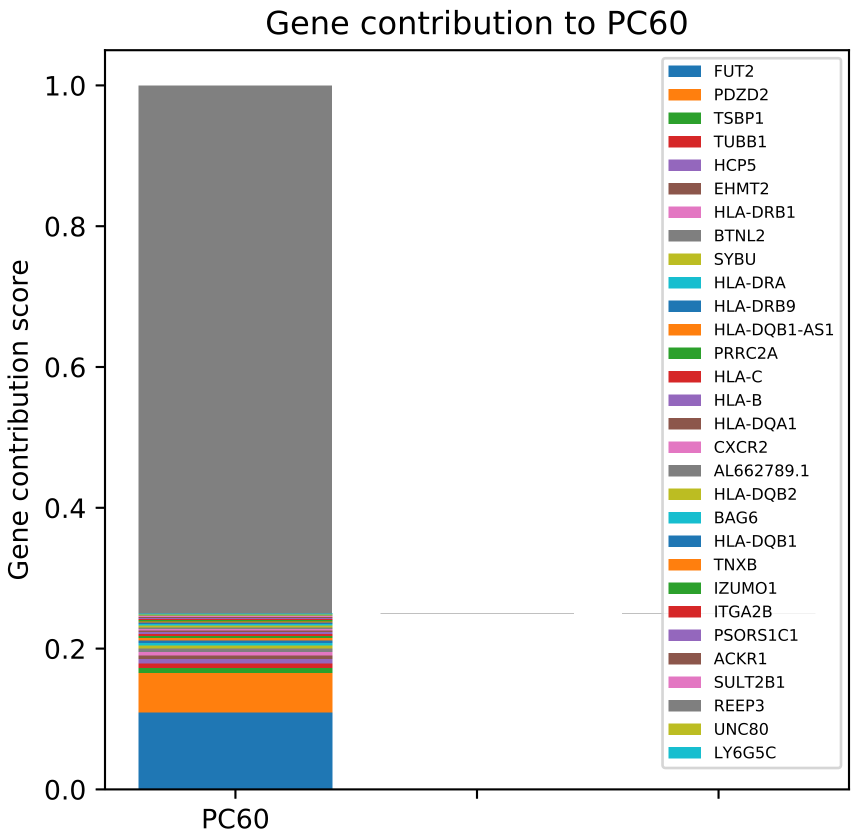
Gene contribution to PC60
Gene contribution score
1.0
0.8
0.6
0.4
0.2
0.0
PC60
FUT2
PDZD2
TSBP1
TUBB1
HCP5
EHMT2
HLA-DRB1
BTNL2
SYBU
HLA-DRA
HLA-DRB9
HLA-DQB1-AS1
PRRC2A
HLA-C
HLA-B
HLA-DQA1
CXCR2
AL662789.1
HLA-DQB2
BAG6
HLA-DQB1
TNXB
IZUMO1
ITGA2B
PSORS1C1
ACKR1
SULT2B1
REEP3
UNC80
LY6G5C

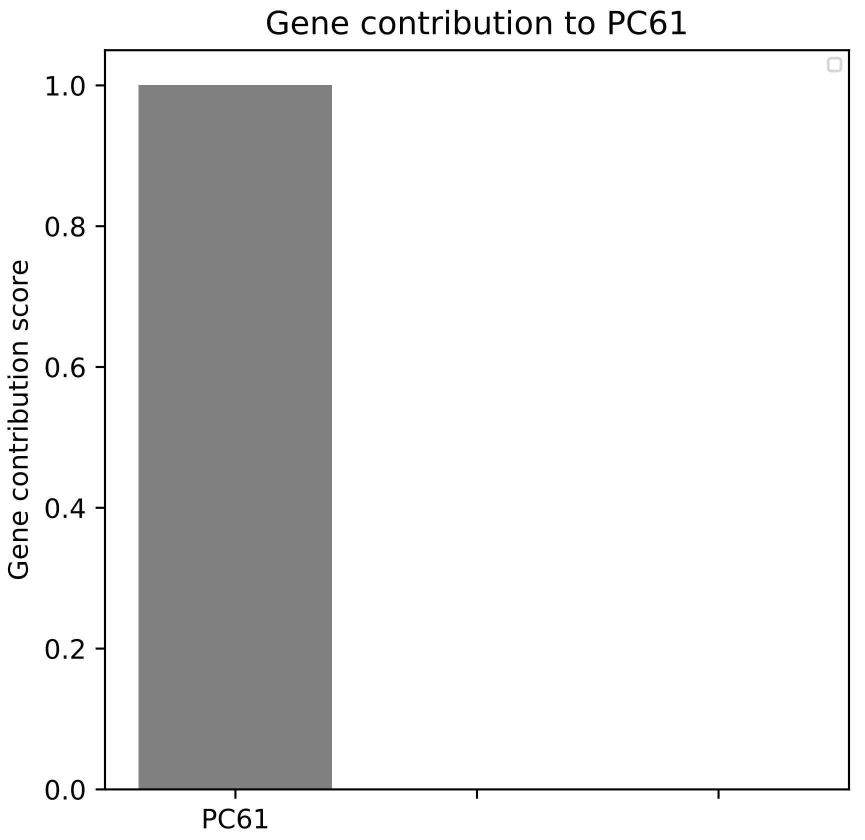

Gene contribution to PC61
Gene contribution score
1.0
0.8
0.6
0.4
0.2
0.0
PC61

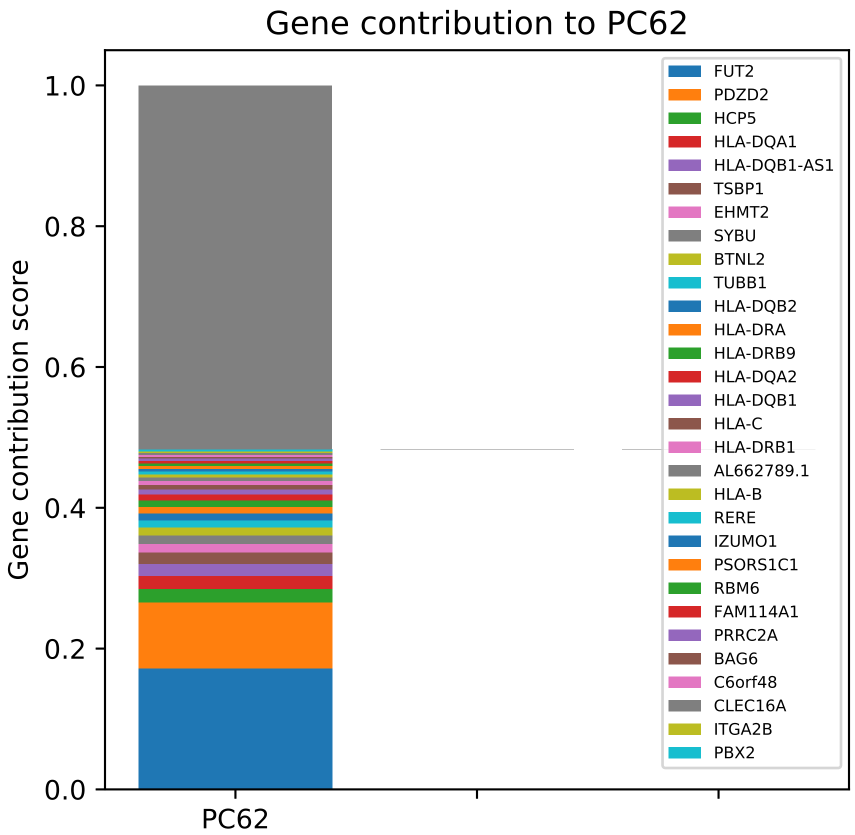

Gene contribution to PC62
Gene contribution score
1.0
0.8
0.6
0.4
0.2
0.0
PC62
FUT2
PDZD2
HCP5
HLA-DQA1
HLA-DQB1-AS1
TSBP1
EHMT2
SYBU
BTNL2
TUBB1
HLA-DQB2
HLA-DRA
HLA-DRB9
HLA-DQA2
HLA-DQB1
HLA-C
HLA-DRB1
AL662789.1
HLA-B
RERE
IZUMO1
PSORS1C1
RBM6
FAM114A1
PRRC2A
BAG6
C6orf48
CLEC16A
ITGA2B
PBX2

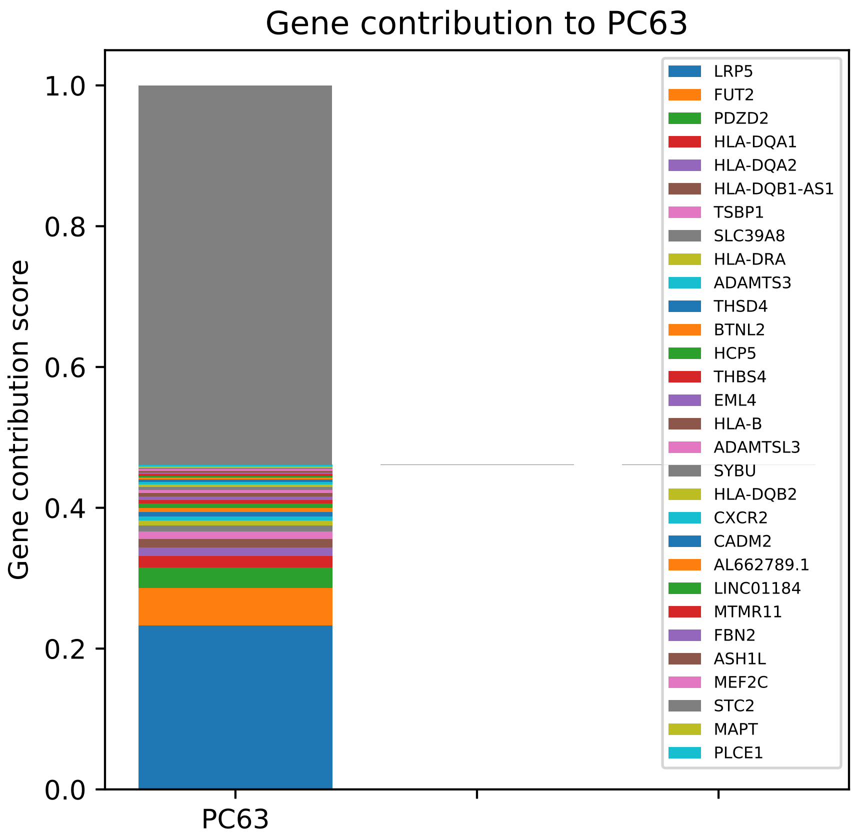
Gene contribution to PC63
Gene contribution score
1.0
0.8
0.6
0.4
0.2
0.0
PC63
LRP5
FUT2
PDZD2
HLA-DQA1
HLA-DQA2
HLA-DQB1-AS1
TSBP1
SLC39A8
HLA-DRA
ADAMTS3
THSD4
BTNL2
HCP5
THBS4
EML4
HLA-B
ADAMTSL3
SYBU
HLA-DQB2
CXCR2
CADM2
AL662789.1
LINC01184
MTMR11
FBN2
ASH1L
MEF2C
STC2
MAPT
PLCE1

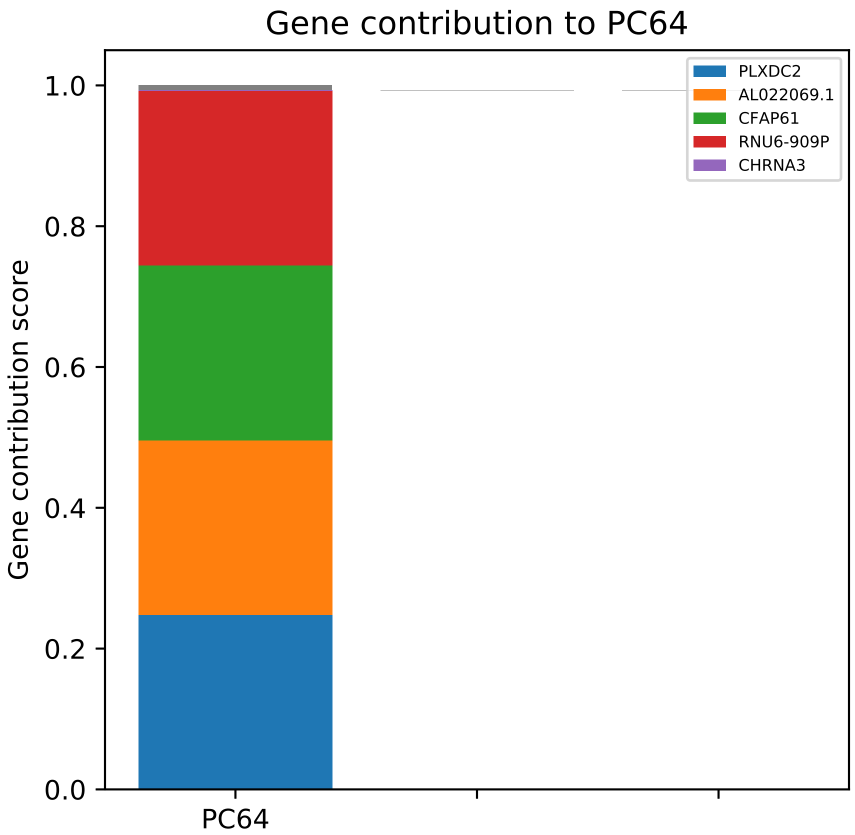

Gene contribution to PC64
Gene contribution score
1.0
0.8
0.6
0.4
0.2
0.0
PC64
PLXDC2
AL022069.1
CFAP61
RNU6-909P
CHRNA3

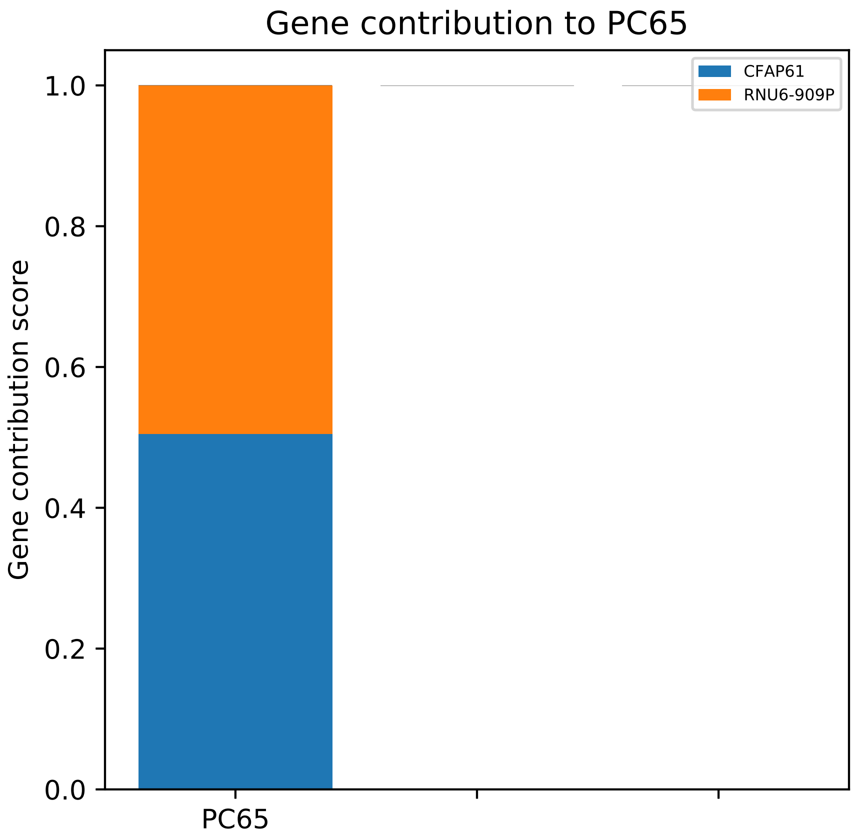
Gene contribution to PC65
Gene contribution score
1.0
0.8
0.6
0.4
0.2
0.0
PC65
CFAP61
RNU6-909P

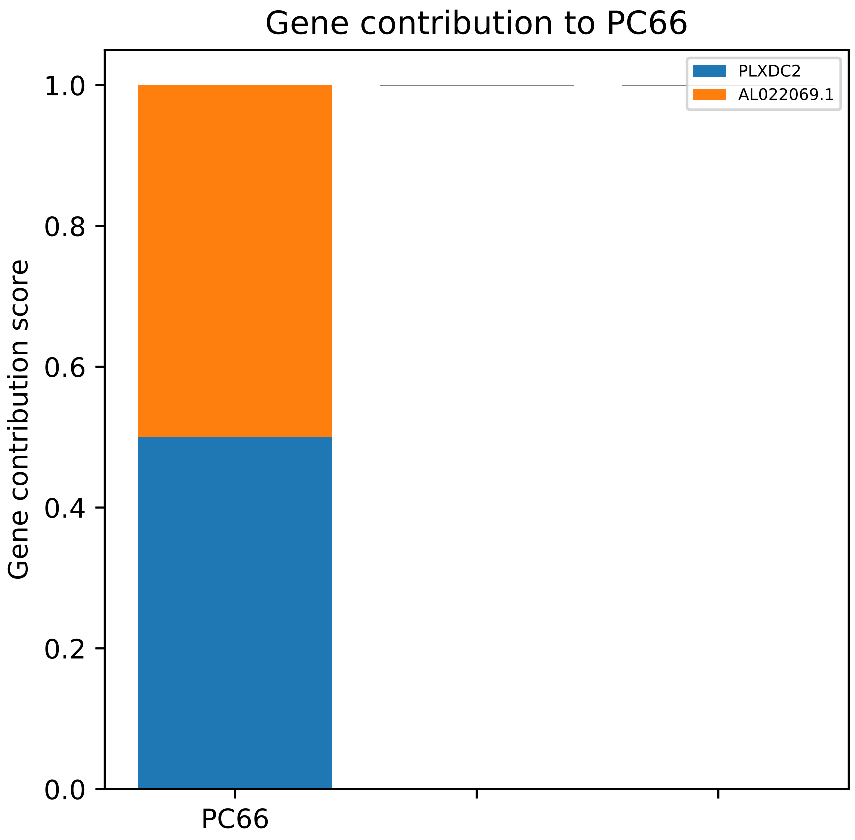

Gene contribution to PC66
PLXDC2
AL022069.1
1.0
0.8
0.6
0.4
0.2
0.0
Gene contribution score
PC66

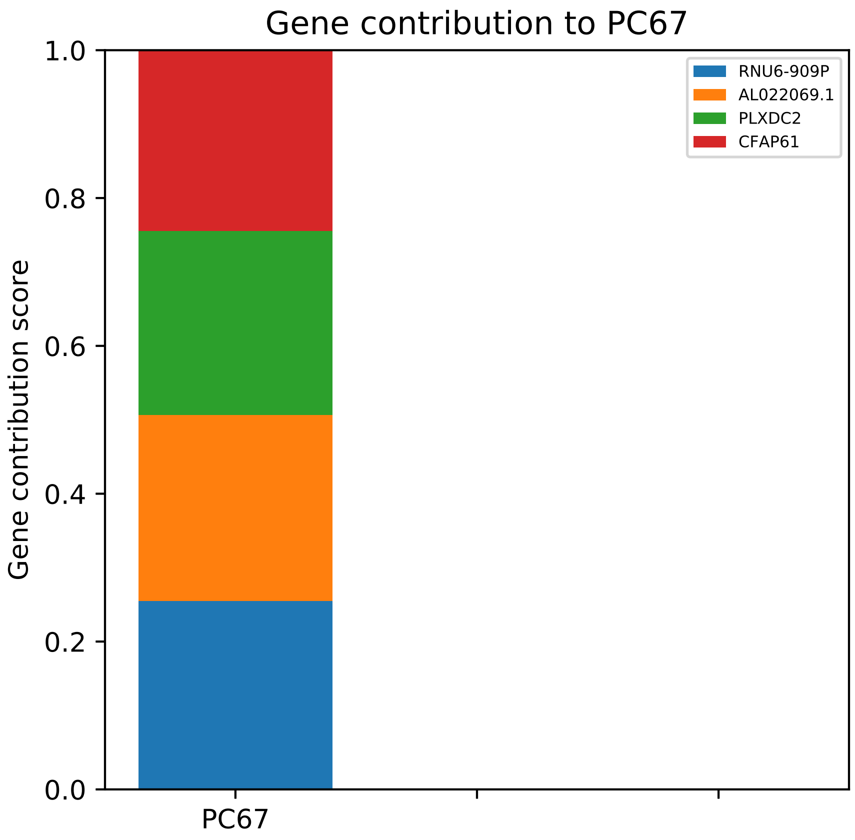
Gene contribution to PC67
1.0
0.8
0.6
0.4
0.2
0.0
Gene contribution score
PC67
RNU6-909P
AL022069.1
PLXDC2
CFAP61

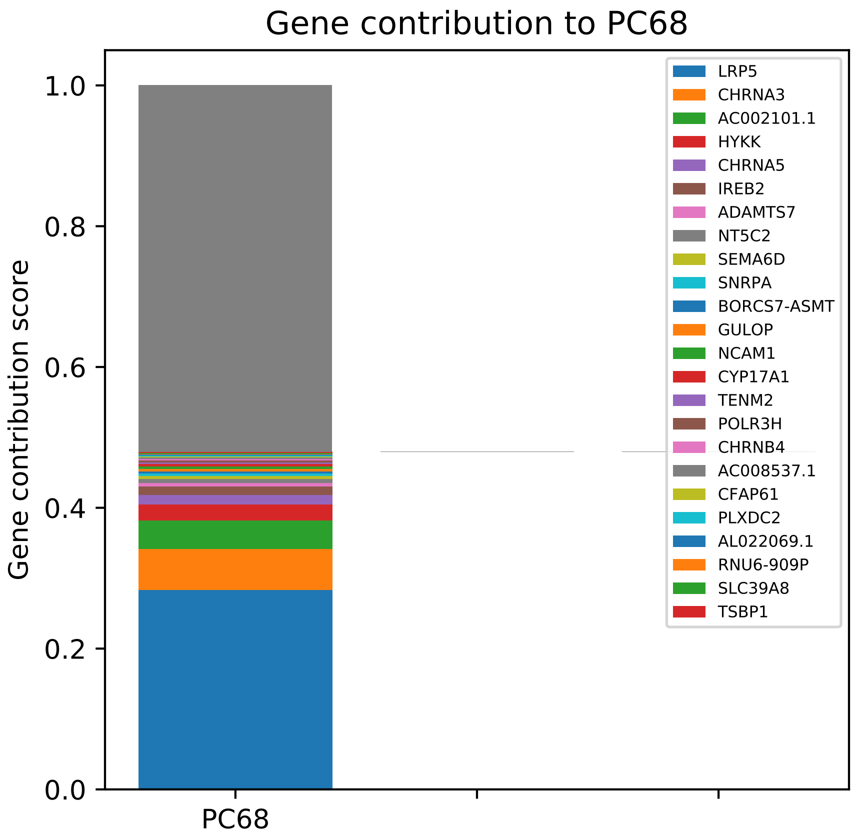

Gene contribution to PC68
Gene contribution score
1.0
0.8
0.6
0.4
0.2
0.0
PC68
LRP5
CHRNA3
AC002101.1
HYKK
CHRNA5
IREB2
ADAMTS7
NT5C2
SEMA6D
SNRPA
BORCS7-ASMT
GULOP
NCAM1
CYP17A1
TENM2
POLR3H
CHRNB4
AC008537.1
CFAP61
PLXDC2
AL022069.1
RNU6-909P
SLC39A8
TSBP1

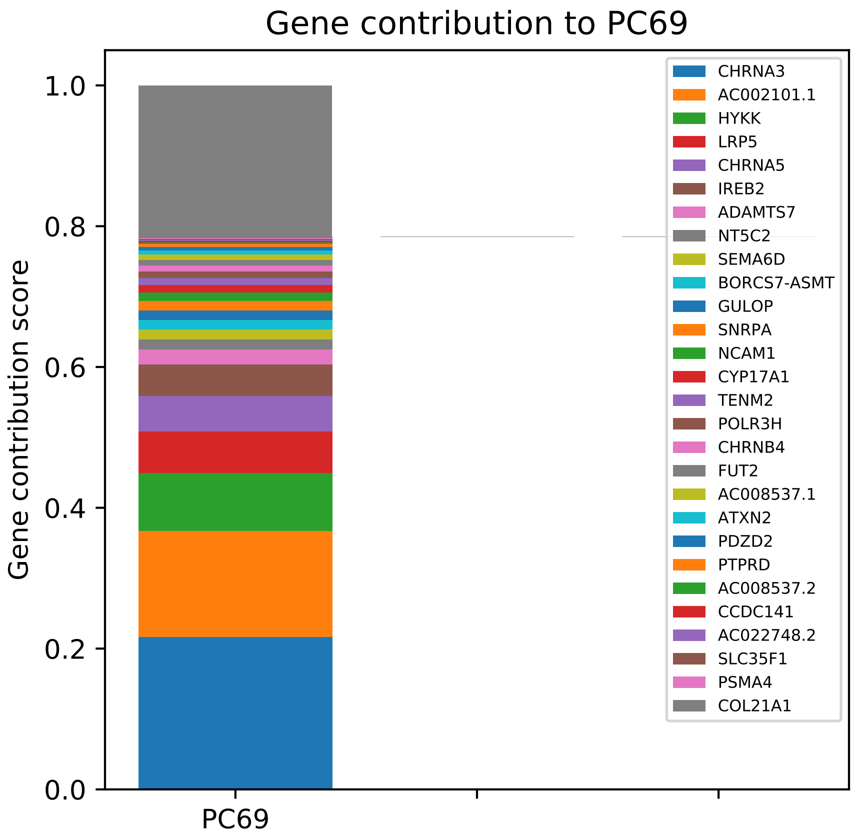

Gene contribution to PC69
Gene contribution score
1.0
0.8
0.6
0.4
0.2
0.0
PC69
CHRNA3
AC002101.1
HYKK
LRP5
CHRNA5
IREB2
ADAMTS7
NT5C2
SEMA6D
BORCS7-ASMT
GULOP
SNRPA
NCAM1
CYP17A1
TENM2
POLR3H
CHRNB4
FUT2
AC008537.1
ATXN2
PDZD2
PTPRD
AC008537.2
CCDC141
AC022748.2
SLC35F1
PSMA4
COL21A1

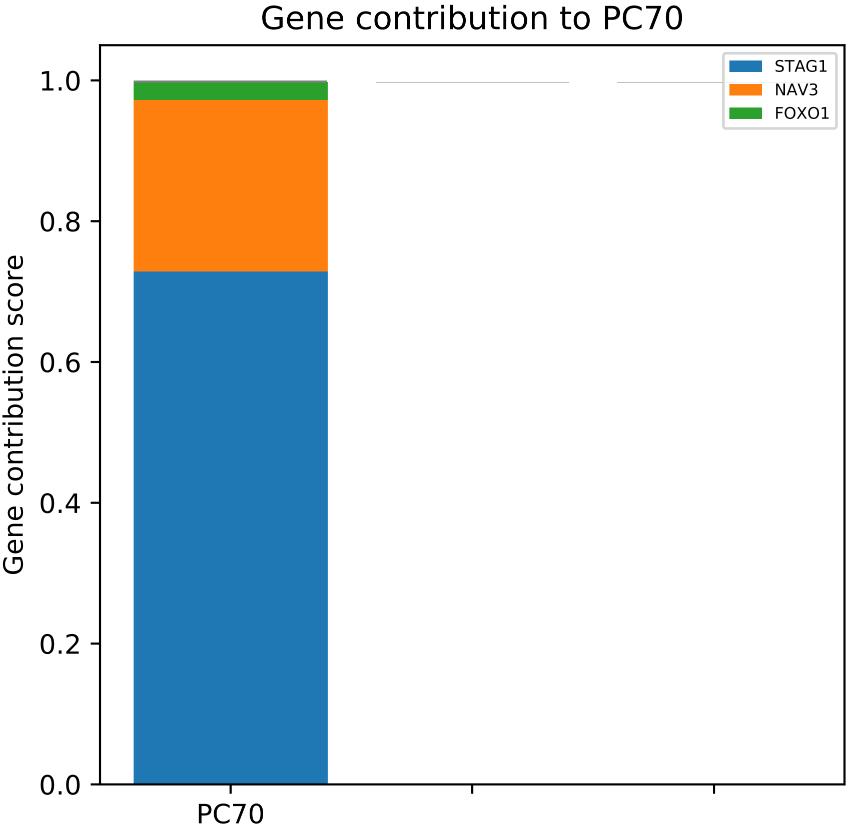

Gene contribution to PC70
Gene contribution score
1.0
0.8
0.6
0.4
0.2
0.0
PC70
STAG1
NAV3
FOXO1

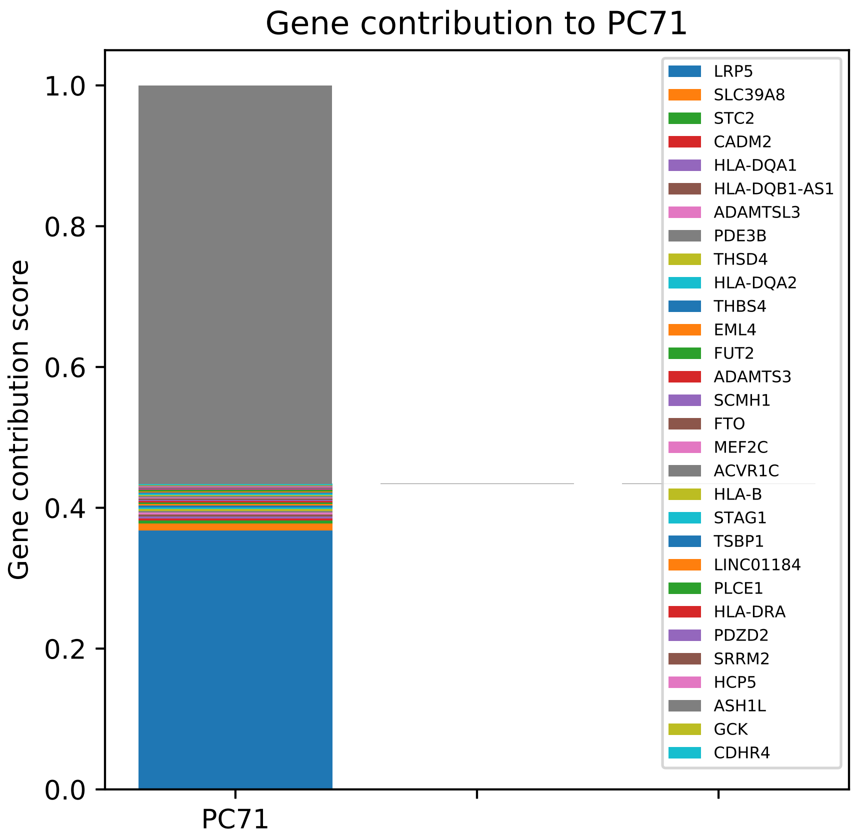
Gene contribution to PC71
Gene contribution score
1.0
0.8
0.6
0.4
0.2
0.0
PC71
LRP5
SLC39A8
STC2
CADM2
HLA-DQA1
HLA-DQB1-AS1
ADAMTSL3
PDE3B
THSD4
HLA-DQA2
THBS4
EML4
FUT2
ADAMTS3
SCMH1
FTO
MEF2C
ACVR1C
HLA-B
STAG1
TSBP1
LINC01184
PLCE1
HLA-DRA
PDZD2
SRRM2
HCP5
ASH1L
GCK
CDHR4

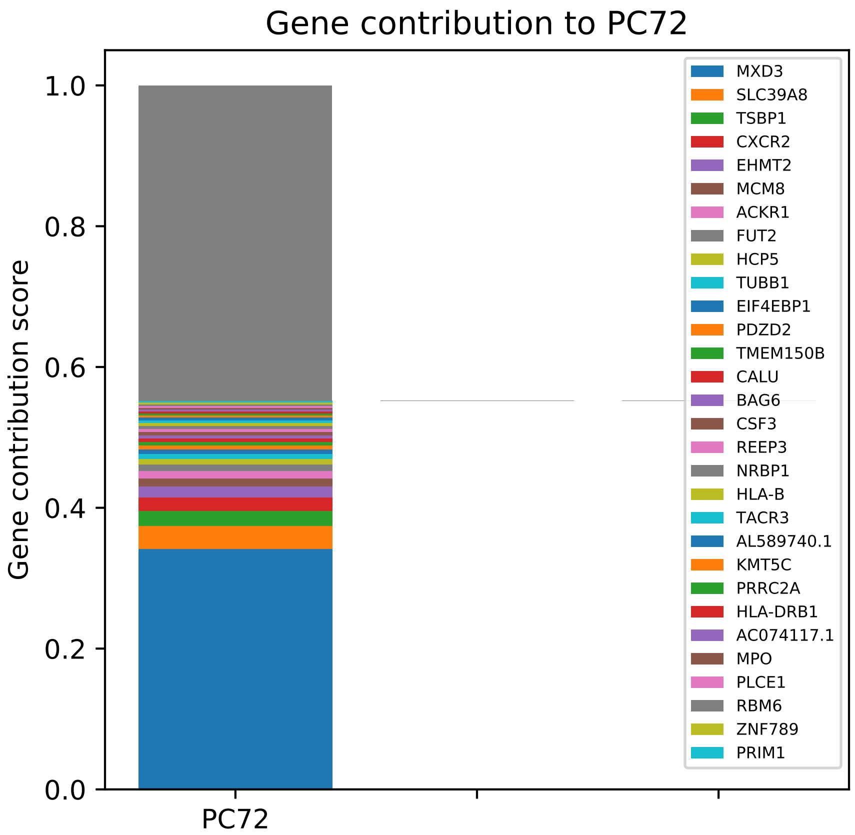

Gene contribution to PC72
Gene contribution score
1.0
0.8
0.6
0.4
0.2
0.0
PC72
MXD3
SLC39A8
TSBP1
CXCR2
EHMT2
MCM8
ACKR1
FUT2
HCP5
TUBB1
EIF4EBP1
PDZD2
TMEM150B
CALU
BAG6
CSF3
REEP3
NRBP1
HLA-B
TACR3
AL589740.1
KMT5C
PRRC2A
HLA-DRB1
AC074117.1
MPO
PLCE1
RBM6
ZNF789
PRIM1

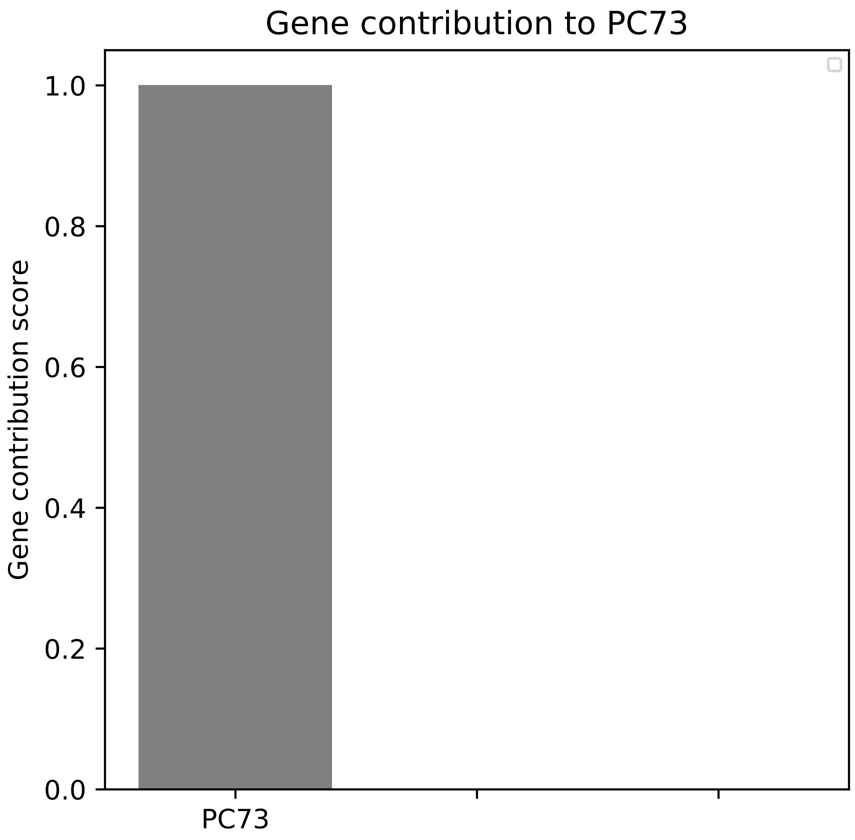

Gene contribution to PC73
Gene contribution score
1.0
0.8
0.6
0.4
0.2
0.0
PC73

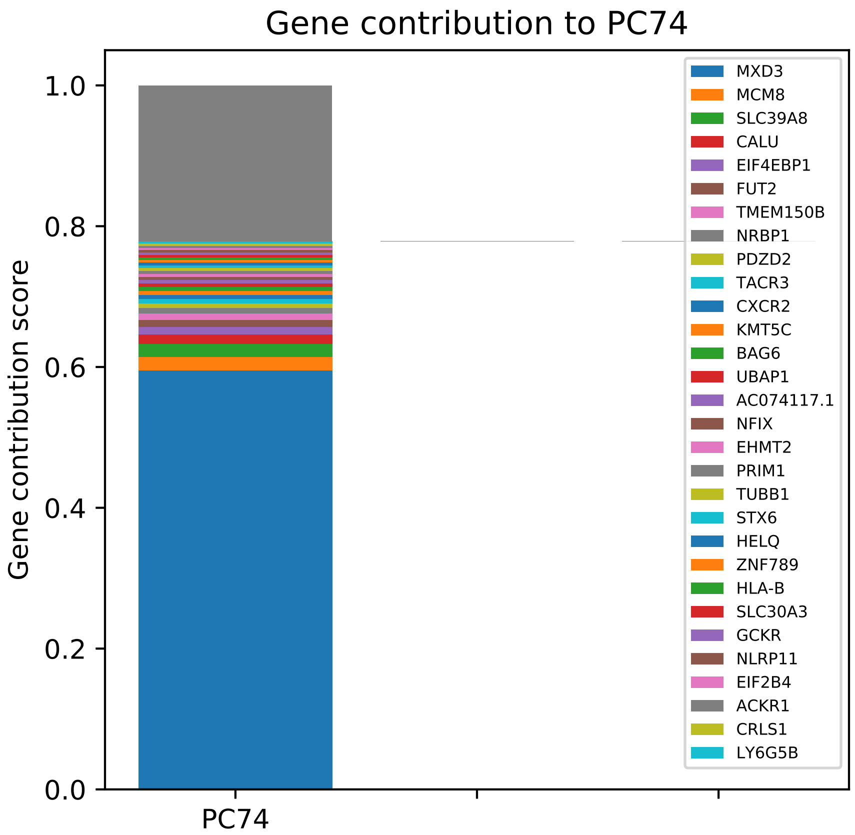

Gene contribution to PC74
Gene contribution score
1.0
0.8
0.6
0.4
0.2
0.0
PC74
MXD3
MCM8
SLC39A8
CALU
EIF4EBP1
FUT2
TMEM150B
NRBP1
PDZD2
TACR3
CXCR2
KMT5C
BAG6
UBAP1
AC074117.1
NFIX
EHMT2
PRIM1
TUBB1
STX6
HELQ
ZNF789
HLA-B
SLC30A3
GCKR
NLRP11
EIF2B4
ACKR1
CRLS1
LY6G5B

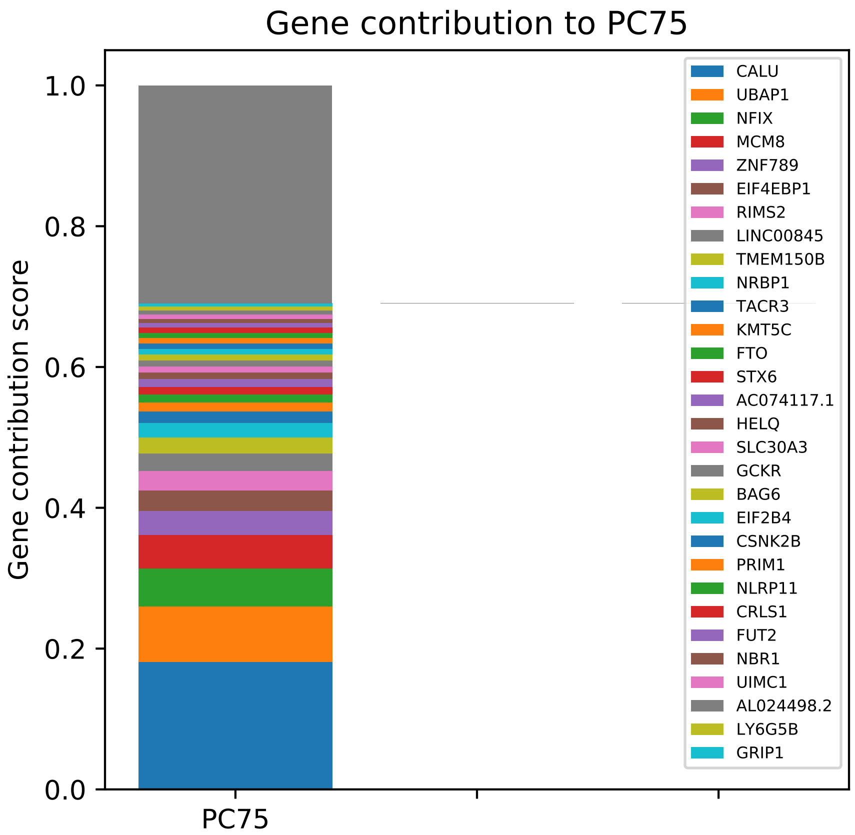

Gene contribution to PC75
Gene contribution score
1.0
0.8
0.6
0.4
0.2
0.0
PC75
CALU
UBAP1
NFIX
MCM8
ZNF789
EIF4EBP1
RIMS2
LINC00845
TMEM150B
NRBP1
TACR3
KMT5C
FTO
STX6
AC074117.1
HELQ
SLC30A3
GCKR
BAG6
EIF2B4
CSNK2B
PRIM1
NLRP11
CRLS1
FUT2
NBR1
UIMC1
AL024498.2
LY6G5B
GRIP1

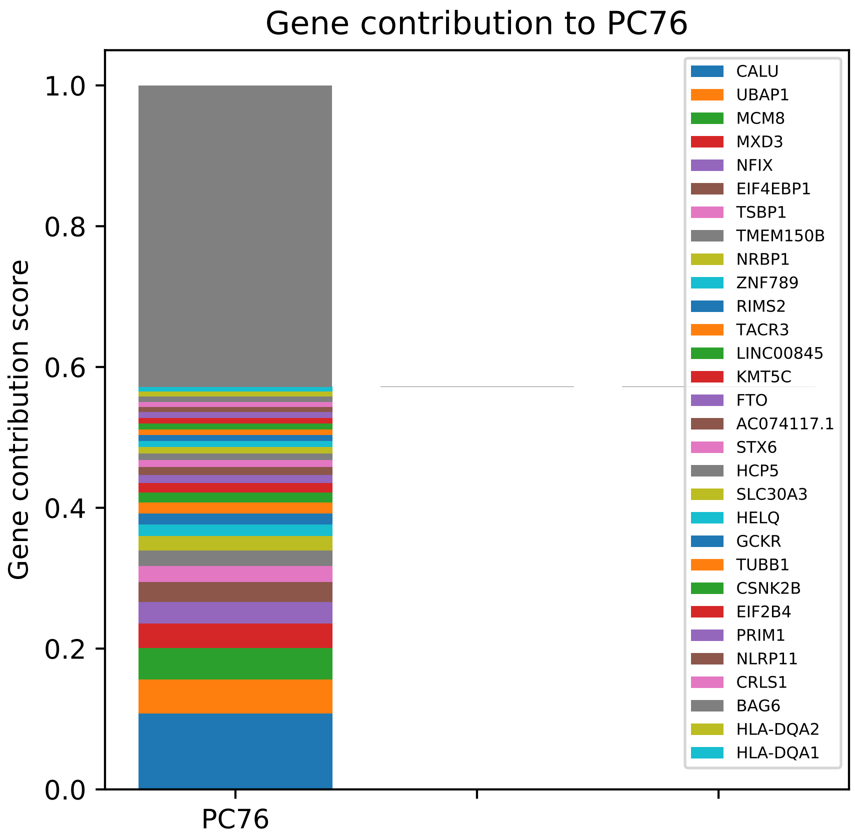
Gene contribution to PC76
Gene contribution score
1.0
0.8
0.6
0.4
0.2
0.0
PC76
CALU
UBAP1
MCM8
MXD3
NFIX
EIF4EBP1
TSBP1
TMEM150B
NRBP1
ZNF789
RIMS2
TACR3
LINC00845
KMT5C
FTO
AC074117.1
STX6
HCP5
SLC30A3
HELQ
GCKR
TUBB1
CSNK2B
EIF2B4
PRIM1
NLRP11
CRLS1
BAG6
HLA-DQA2
HLA-DQA1

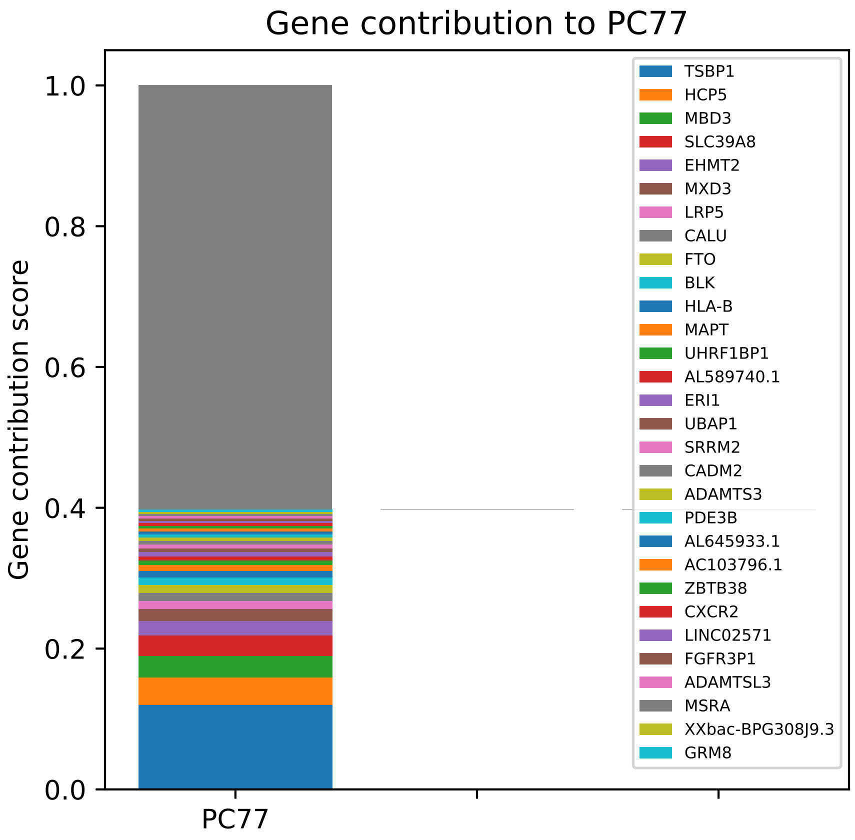

Gene contribution to PC77
Gene contribution score
1.0
0.8
0.6
0.4
0.2
0.0
PC77
TSBP1
HCP5
MBD3
SLC39A8
EHMT2
MXD3
LRP5
CALU
FTO
BLK
HLA-B
MAPT
UHRF1BP1
AL589740.1
ERI1
UBAP1
SRRM2
CADM2
ADAMTS3
PDE3B
AL645933.1
AC103796.1
ZBTB38
CXCR2
LINC02571
FGFR3P1
ADAMTSL3
MSRA
XXbac-BPG308J9.3
GRM8

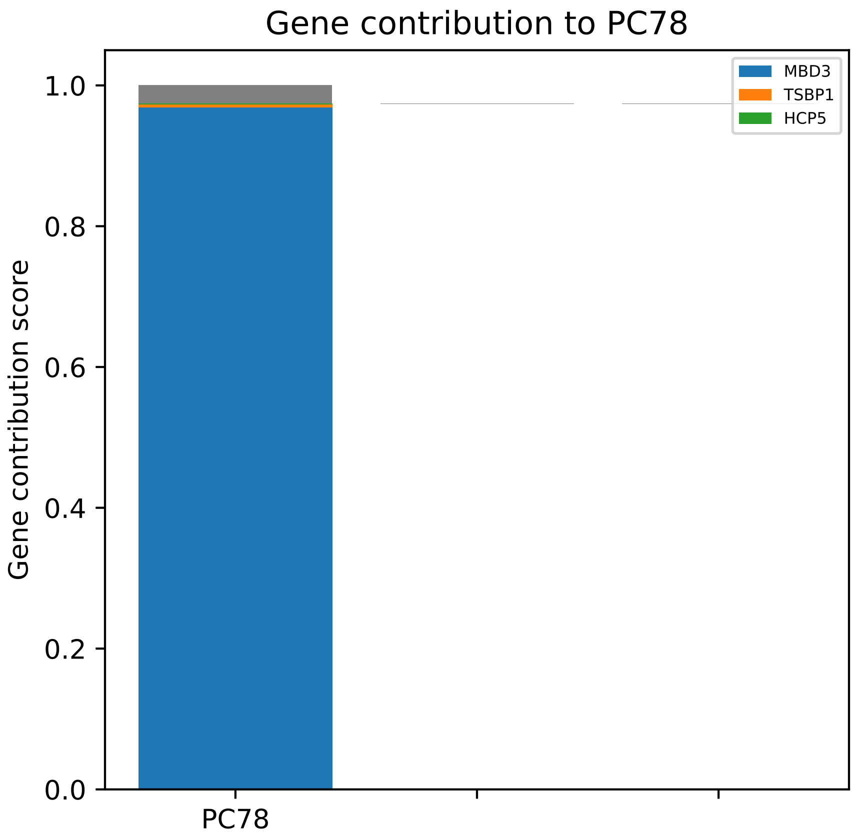

Gene contribution to PC78
Gene contribution score
1.0
0.8
0.6
0.4
0.2
0.0
PC78
MBD3
TSBP1
HCP5

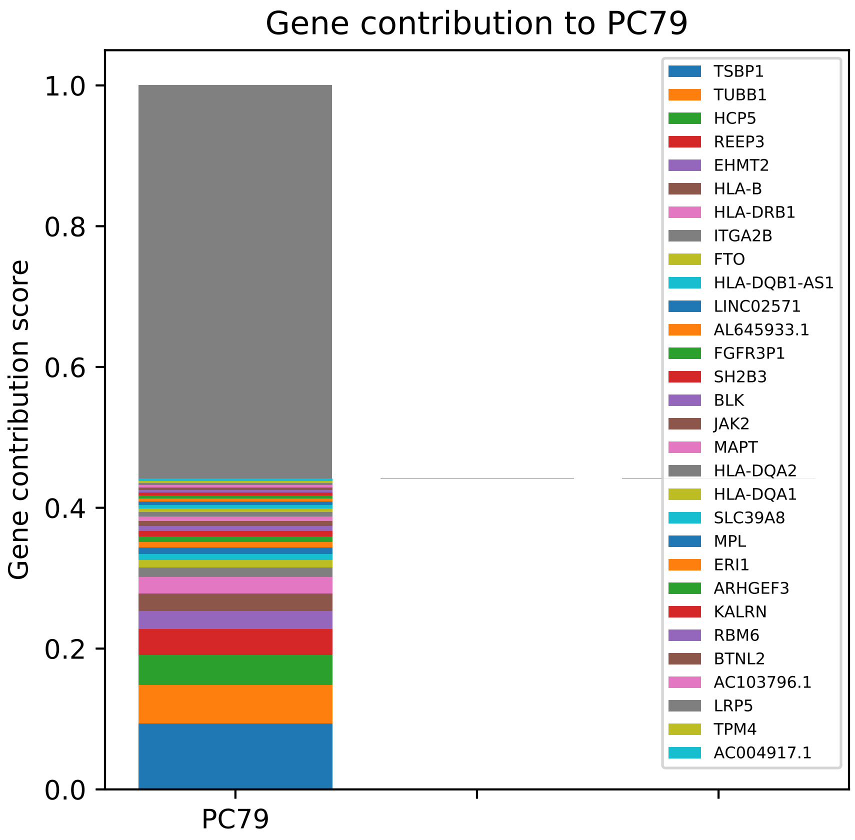

Gene contribution to PC79
Gene contribution score
1.0
0.8
0.6
0.4
0.2
0.0
PC79
TSBP1
TUBB1
HCP5
REEP3
EHMT2
HLA-B
HLA-DRB1
ITGA2B
FTO
HLA-DQB1-AS1
LINC02571
AL645933.1
FGFR3P1
SH2B3
BLK
JAK2
MAPT
HLA-DQA2
HLA-DQA1
SLC39A8
MPL
ERI1
ARHGEF3
KALRN
RBM6
BTNL2
AC103796.1
LRP5
TPM4
AC004917.1

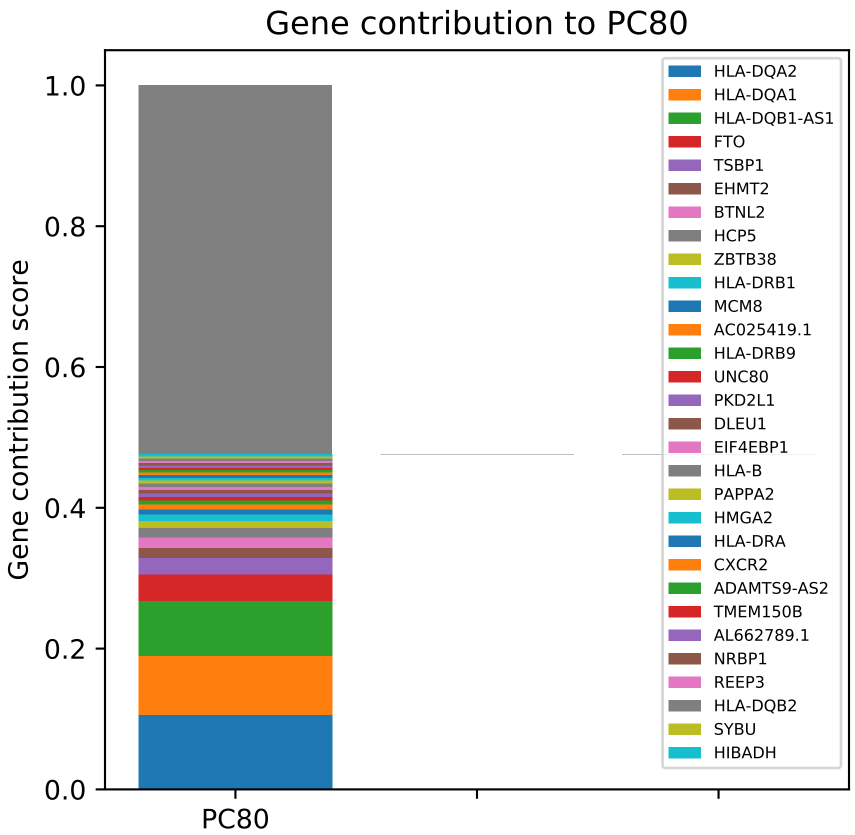

Gene contribution to PC80
Gene contribution score
1.0
0.8
0.6
0.4
0.2
0.0
PC80
HLA-DQA2
HLA-DQA1
HLA-DQB1-AS1
FTO
TSBP1
EHMT2
BTNL2
HCP5
ZBTB38
HLA-DRB1
MCM8
AC025419.1
HLA-DRB9
UNC80
PKD2L1
DLEU1
EIF4EBP1
HLA-B
PAPPA2
HMGA2
HLA-DRA
CXCR2
ADAMTS9-AS2
TMEM150B
AL662789.1
NRBP1
REEP3
HLA-DQB2
SYBU
HIBADH

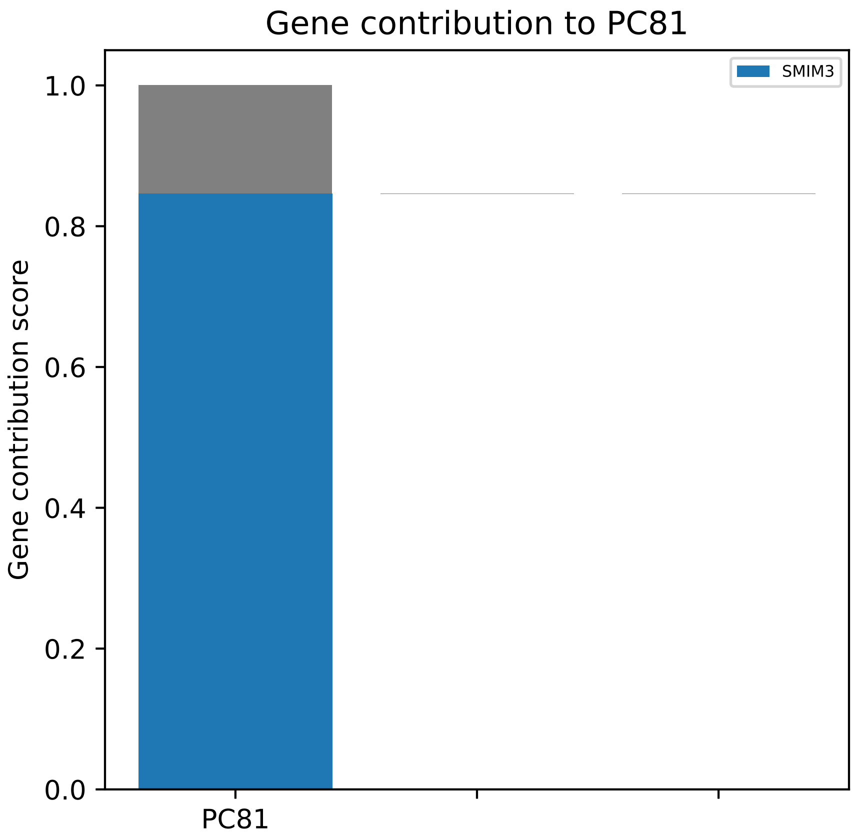

Gene contribution to PC81
Gene contribution score
1.0
0.8
0.6
0.4
0.2
0.0
PC81
SMIM3

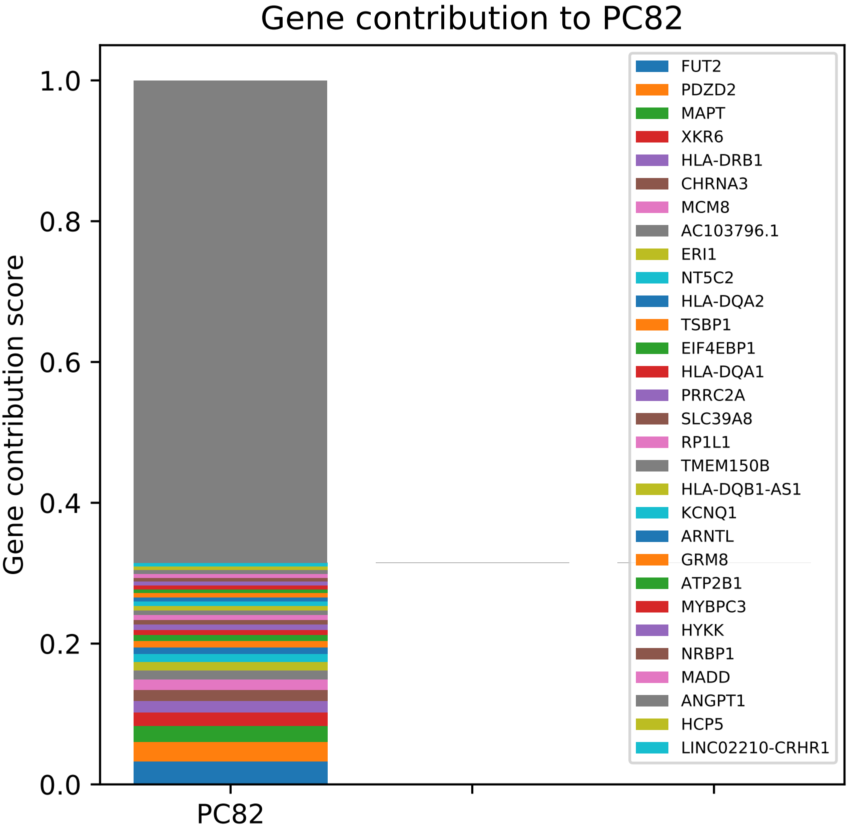

Gene contribution to PC82
Gene contribution score
1.0
0.8
0.6
0.4
0.2
0.0
PC82
FUT2
PDZD2
MAPT
XKR6
HLA-DRB1
CHRNA3
MCM8
AC103796.1
ERI1
NT5C2
HLA-DQA2
TSBP1
EIF4EBP1
HLA-DQA1
PRRC2A
SLC39A8
RP1L1
TMEM150B
HLA-DQB1-AS1
KCNQ1
ARNTL
GRM8
ATP2B1
MYBPC3
HYKK
NRBP1
MADD
ANGPT1
HCP5
LINC02210-CRHR1

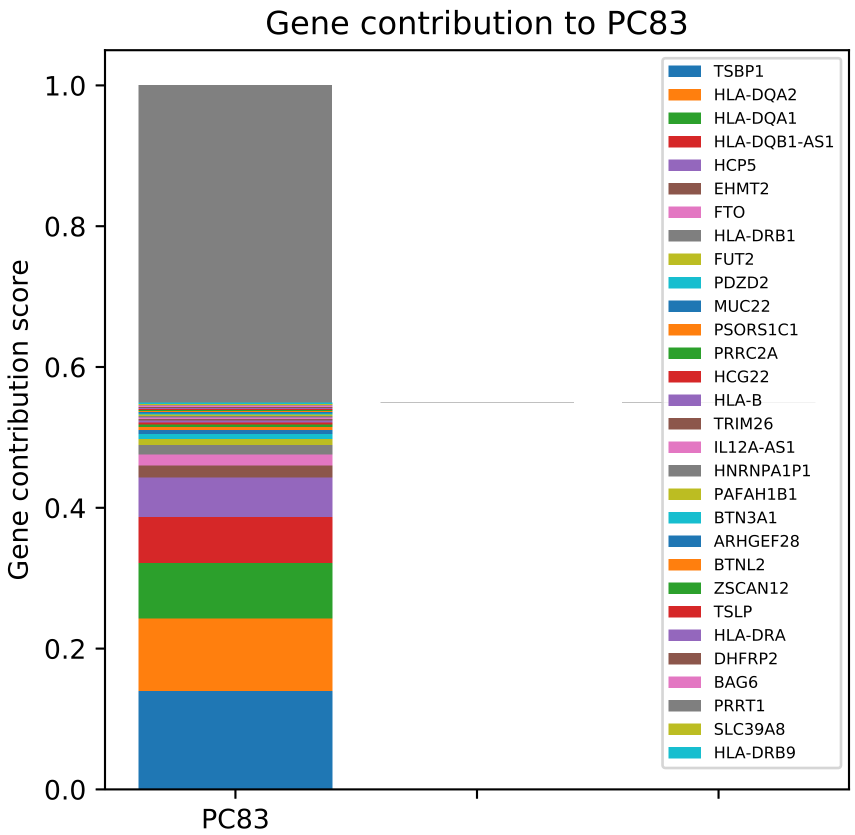

Gene contribution to PC83
Gene contribution score
1.0
0.8
0.6
0.4
0.2
0.0
PC83
TSBP1
HLA-DQA2
HLA-DQA1
HLA-DQB1-AS1
HCP5
EHMT2
FTO
HLA-DRB1
FUT2
PDZD2
MUC22
PSORS1C1
PRRC2A
HCG22
HLA-B
TRIM26
IL12A-AS1
HNRNPA1P1
PAFAH1B1
BTN3A1
ARHGEF28
BTNL2
ZSCAN12
TSLP
HLA-DRA
DHFRP2
BAG6
PRRT1
SLC39A8
HLA-DRB9

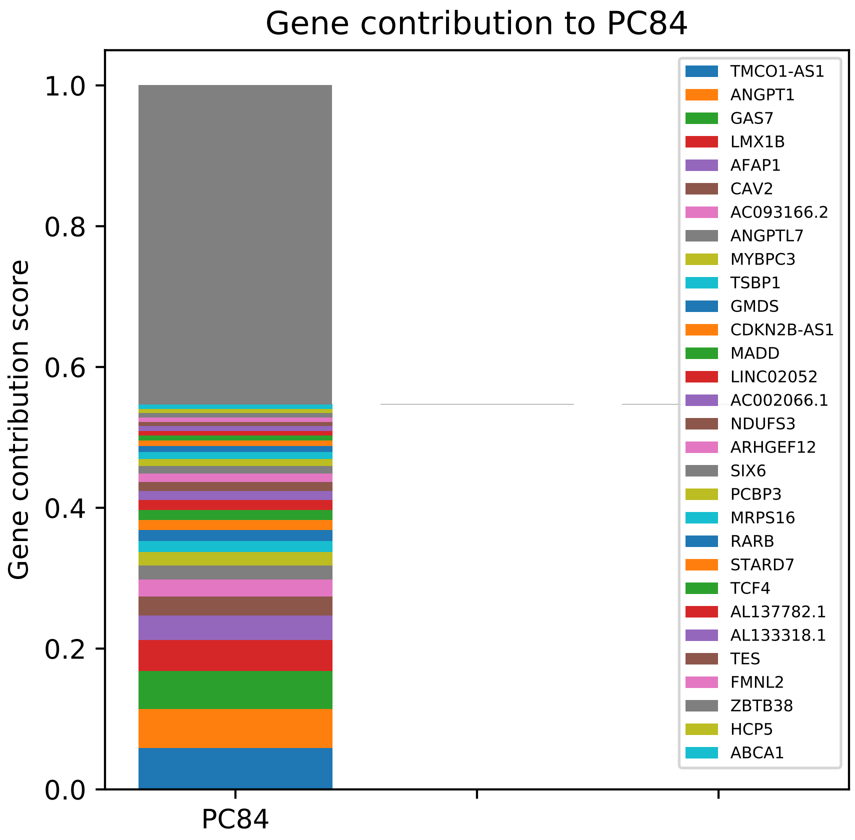

Gene contribution to PC84
Gene contribution score
1.0
0.8
0.6
0.4
0.2
0.0
PC84
TMCO1-AS1
ANGPT1
GAS7
LMX1B
AFAP1
CAV2
AC093166.2
ANGPTL7
MYBPC3
TSBP1
GMDS
CDKN2B-AS1
MADD
LINC02052
AC002066.1
NDUFS3
ARHGEF12
SIX6
PCBP3
MRPS16
RARB
STARD7
TCF4
AL137782.1
AL133318.1
TES
FMNL2
ZBTB38
HCP5
ABCA1

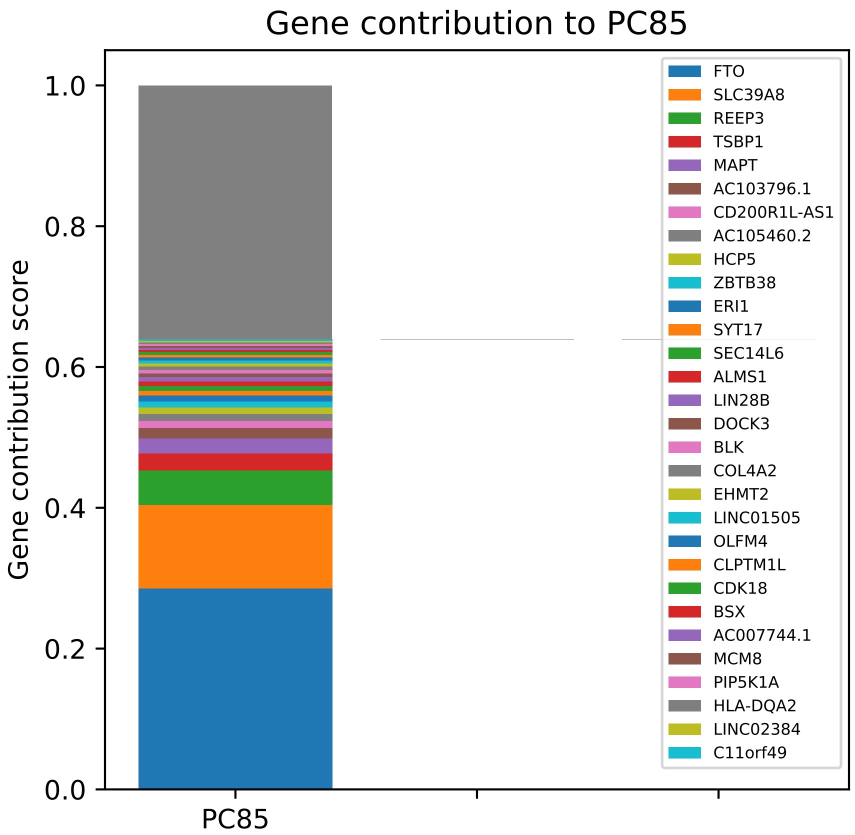

Gene contribution to PC85
Gene contribution score
1.0
0.8
0.6
0.4
0.2
0.0
PC85
FTO
SLC39A8
REEP3
TSBP1
MAPT
AC103796.1
CD200R1L-AS1
AC105460.2
HCP5
ZBTB38
ERI1
SYT17
SEC14L6
ALMS1
LIN28B
DOCK3
BLK
COL4A2
EHMT2
LINC01505
OLFM4
CLPTM1L
CDK18
BSX
AC007744.1
MCM8
PIP5K1A
HLA-DQA2
LINC02384
C11orf49

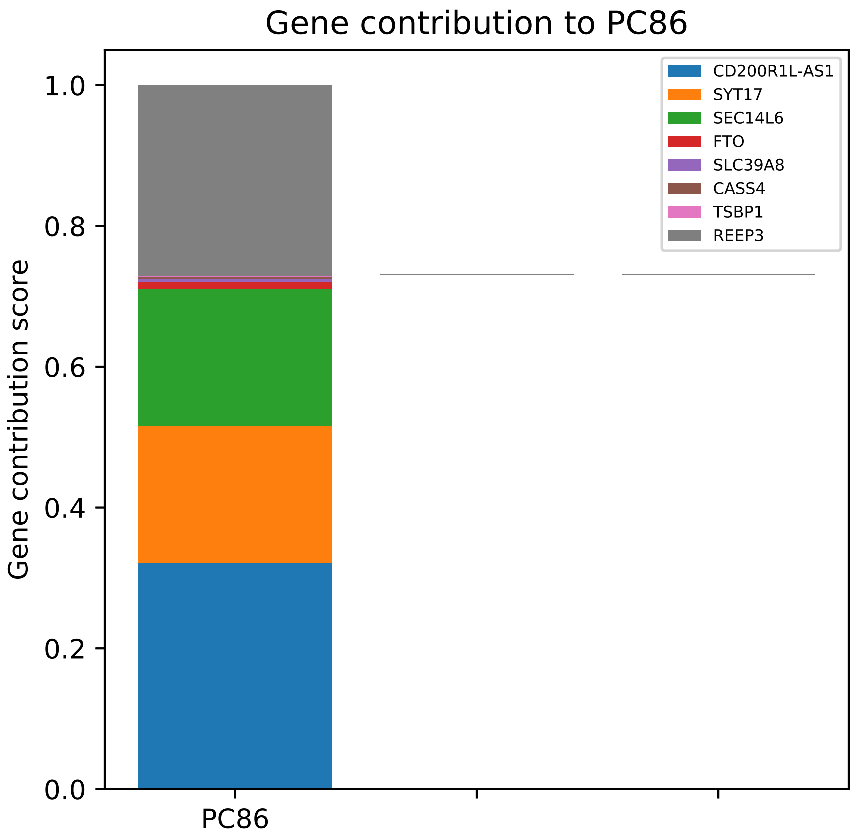

Gene contribution to PC86
Gene contribution score
1.0
0.8
0.6
0.4
0.2
0.0
PC86
CD200R1L-AS1
SYT17
SEC14L6
FTO
SLC39A8
CASS4
TSBP1
REEP3

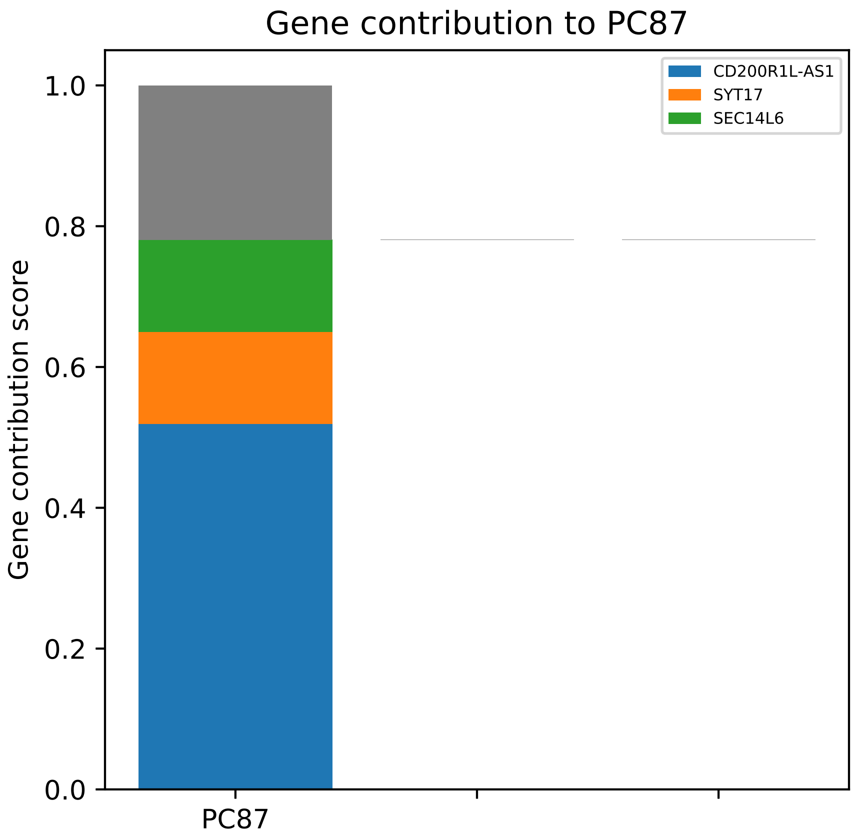

Gene contribution to PC87
CD200R1L-AS1
SYT17
SEC14L6
1.0
0.8
0.6
0.4
0.2
0.0
Gene contribution score
PC87

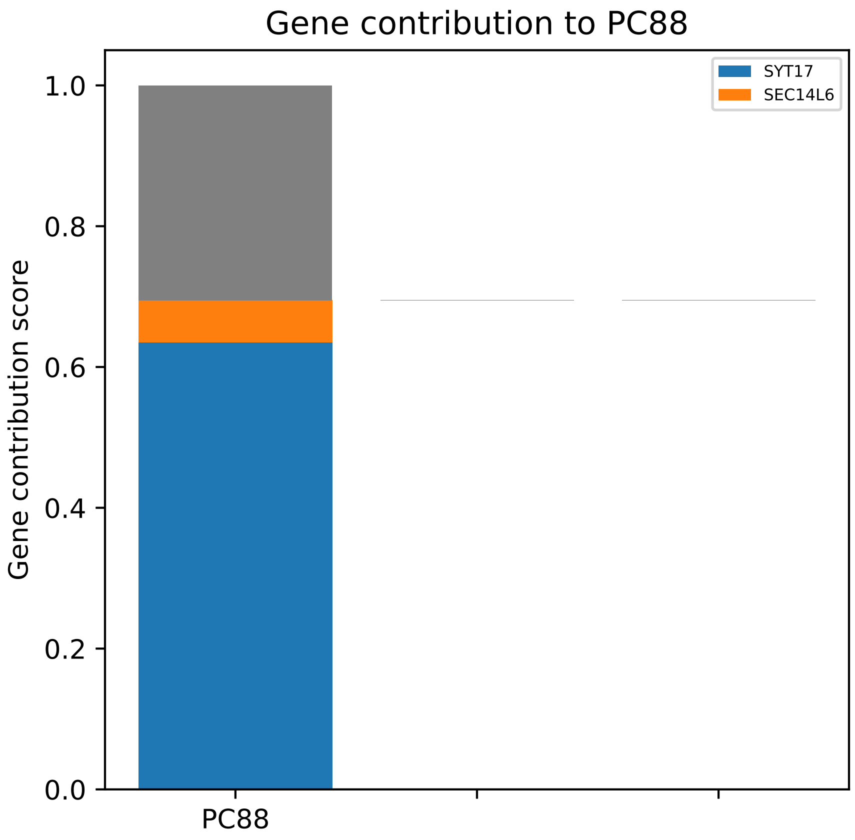
Gene contribution to PC88
SYT17
SEC14L6
Gene contribution score
1.0
0.8
0.6
0.4
0.2
0.0
PC88

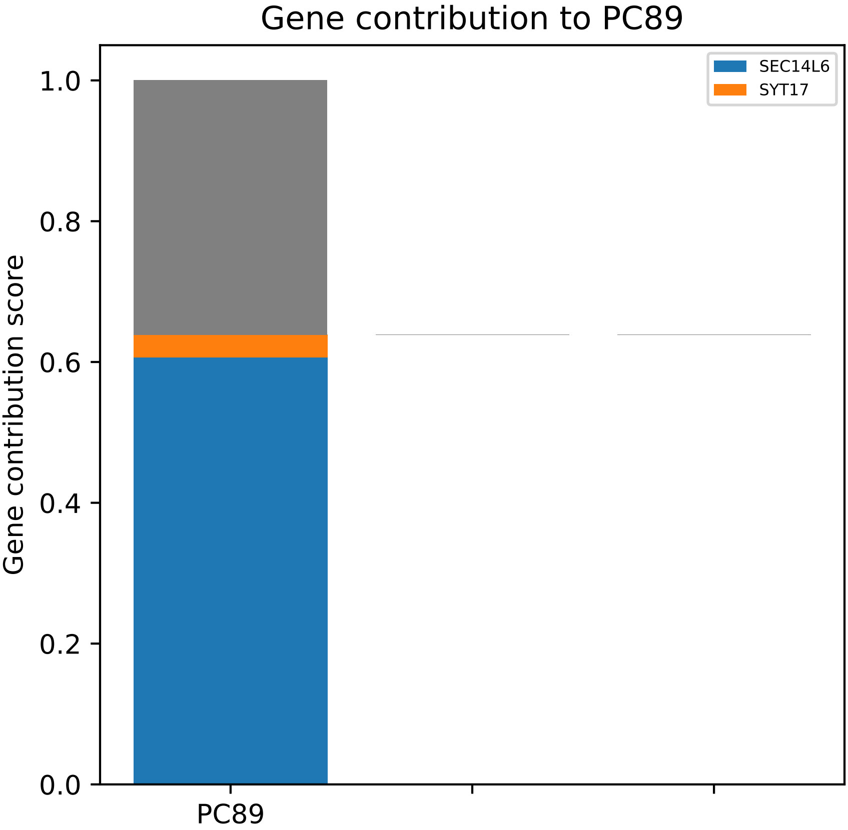
Gene contribution to PC89
SEC14L6
SYT17
Gene contribution score
1.0
0.8
0.6
0.4
0.2
0.0
PC89

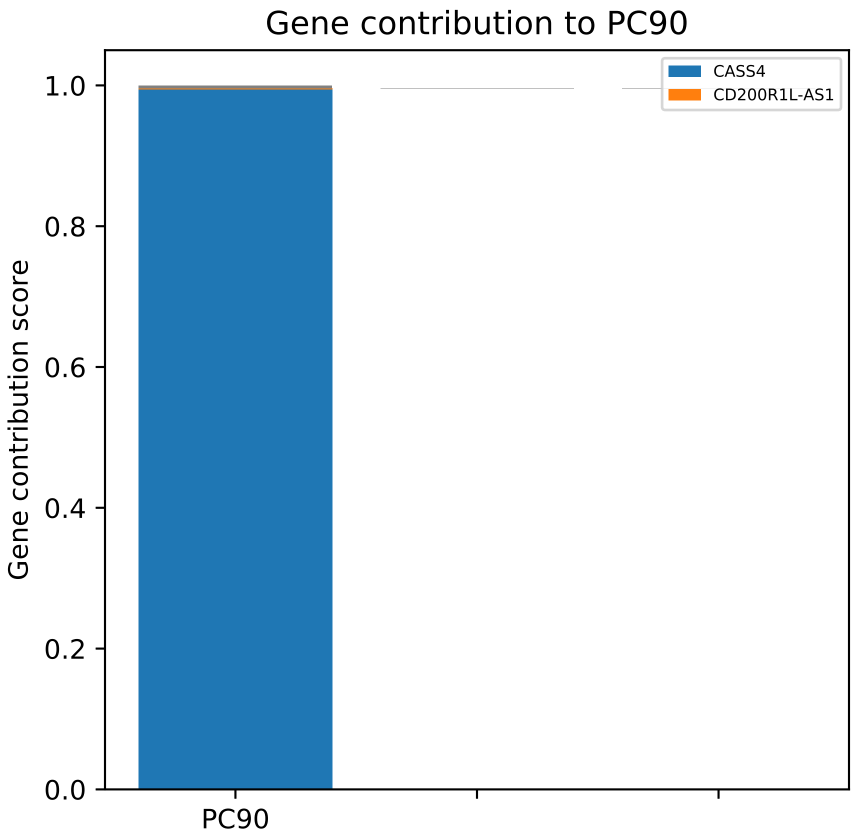

Gene contribution to PC90
Gene contribution score
1.0
0.8
0.6
0.4
0.2
0.0
PC90
CASS4
CD200R1L-AS1

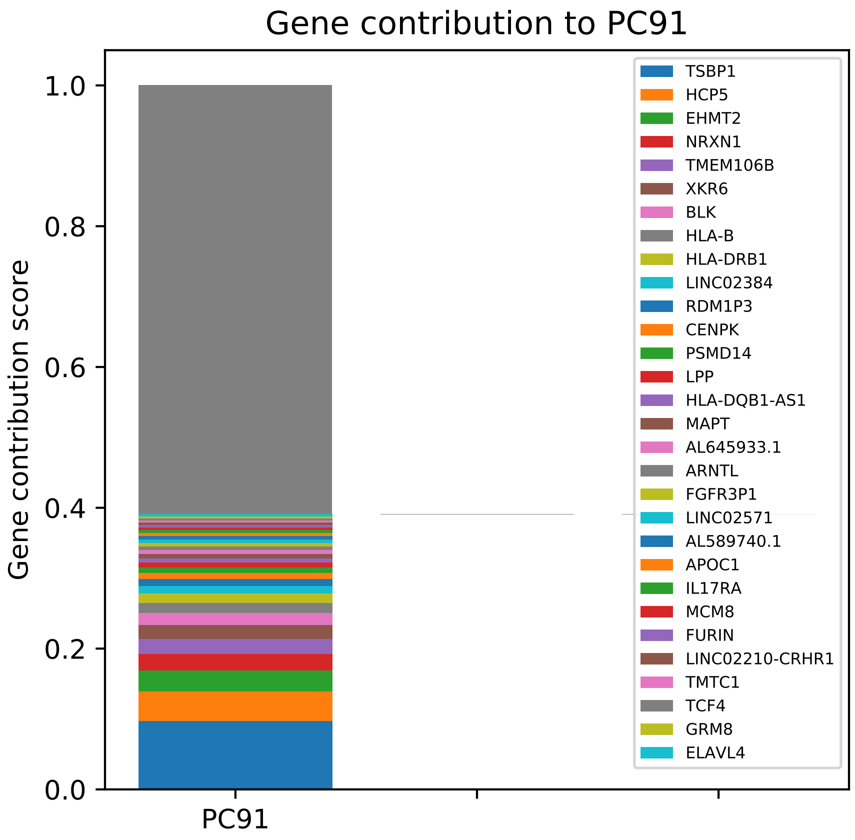

Gene contribution to PC91
Gene contribution score
0.0
0.2
0.4
0.6
0.8
1.0
PC91
TSBP1
HCP5
EHMT2
NRXN1
TMEM106B
XKR6
BLK
HLA-B
HLA-DRB1
LINC02384
RDM1P3
CENPK
PSMD14
LPP
HLA-DQB1-AS1
MAPT
AL645933.1
ARNTL
FGFR3P1
LINC02571
AL589740.1
APOC1
IL17RA
MCM8
FURIN
LINC02210-CRHR1
TMTC1
TCF4
GRM8
ELAVL4

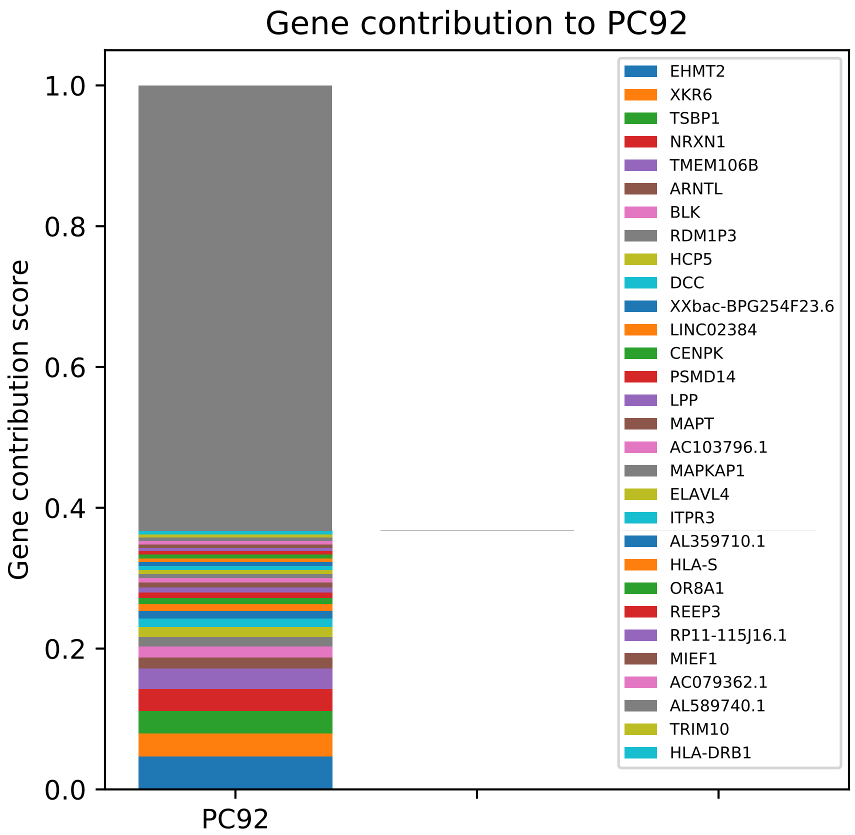

Gene contribution to PC92
Gene contribution score
1.0
0.8
0.6
0.4
0.2
0.0
PC92
EHMT2
XKR6
TSBP1
NRXN1
TMEM106B
ARNTL
BLK
RDM1P3
HCP5
DCC
XXbac-BPG254F23.6
LINC02384
CENPK
PSMD14
LPP
MAPT
AC103796.1
MAPKAP1
ELAVL4
ITPR3
AL359710.1
HLA-S
OR8A1
REEP3
RP11-115J16.1
MIEF1
AC079362.1
AL589740.1
TRIM10
HLA-DRB1

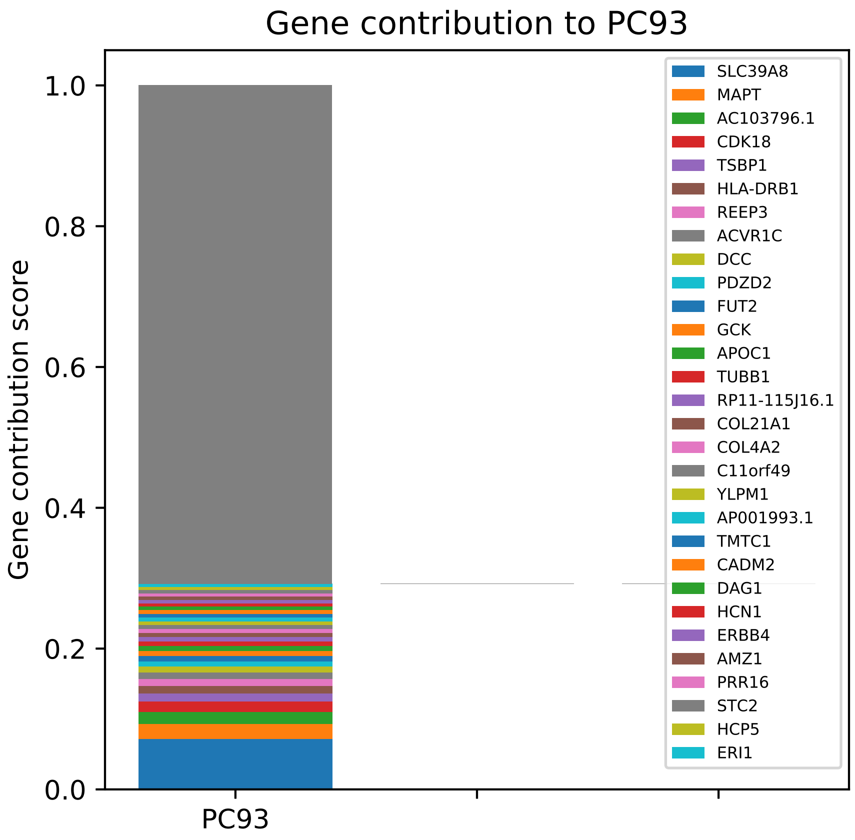
Gene contribution to PC93
Gene contribution score
0.0
0.2
0.4
0.6
0.8
1.0
PC93
SLC39A8
MAPT
AC103796.1
CDK18
TSBP1
HLA-DRB1
REEP3
ACVR1C
DCC
PDZD2
FUT2
GCK
APOC1
TUBB1
RP11-115J16.1
COL21A1
COL4A2
C11orf49
YLPM1
AP001993.1
TMTC1
CADM2
DAG1
HCN1
ERBB4
AMZ1
PRR16
STC2
HCP5
ERI1

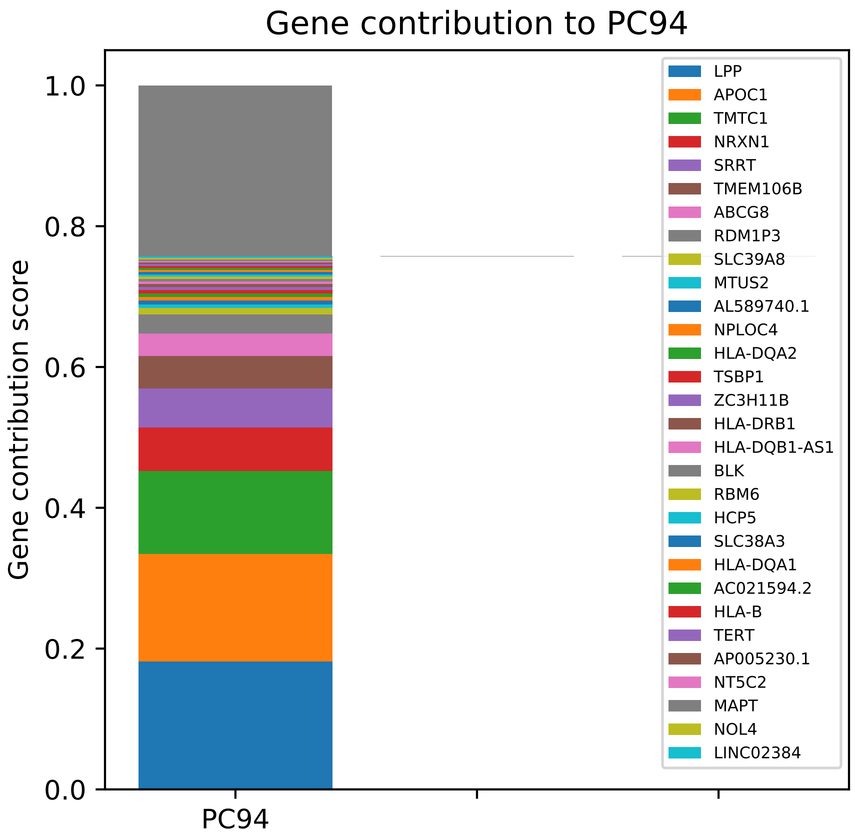

Gene contribution to PC94
Gene contribution score
0.0
0.2
0.4
0.6
0.8
1.0
PC94
LPP
APOC1
TMTC1
NRXN1
SRRT
TMEM106B
ABCG8
RDM1P3
SLC39A8
MTUS2
AL589740.1
NPLOC4
HLA-DQA2
TSBP1
ZC3H11B
HLA-DRB1
HLA-DQB1-AS1
BLK
RBM6
HCP5
SLC38A3
HLA-DQA1
AC021594.2
HLA-B
TERT
AP005230.1
NT5C2
MAPT
NOL4
LINC02384

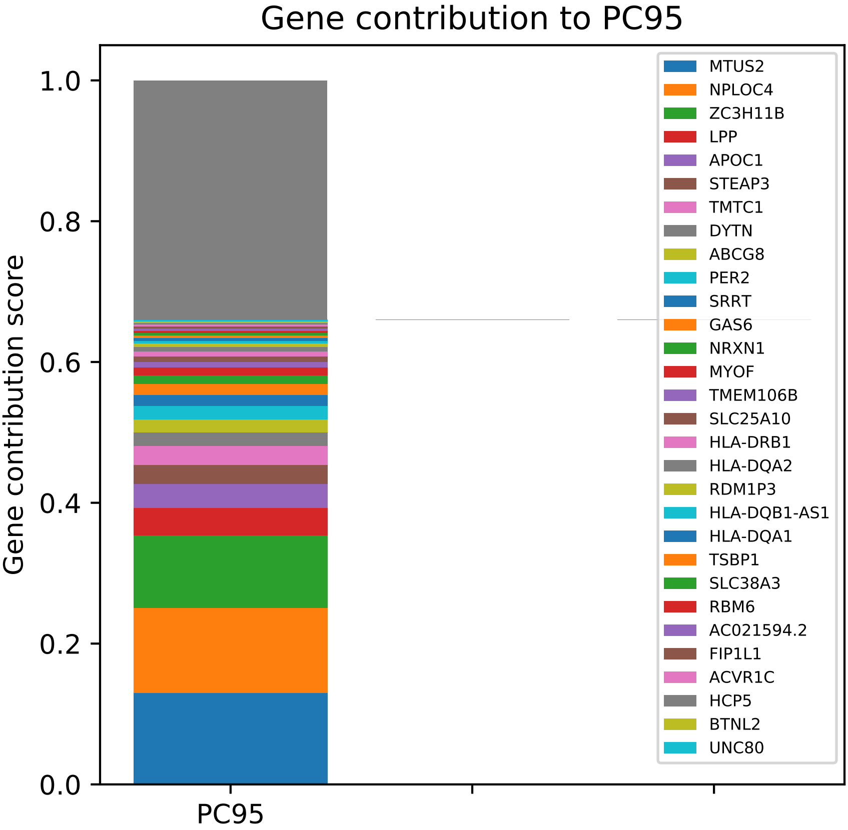
Gene contribution to PC95
Gene contribution score
1.0
0.8
0.6
0.4
0.2
0.0
PC95
MTUS2
NPLOC4
ZC3H11B
LPP
APOC1
STEAP3
TMTC1
DYTN
ABCG8
PER2
SRRT
GAS6
NRXN1
MYOF
TMEM106B
SLC25A10
HLA-DRB1
HLA-DQA2
RDM1P3
HLA-DQB1-AS1
HLA-DQA1
TSBP1
SLC38A3
RBM6
AC021594.2
FIP1L1
ACVR1C
HCP5
BTNL2
UNC80

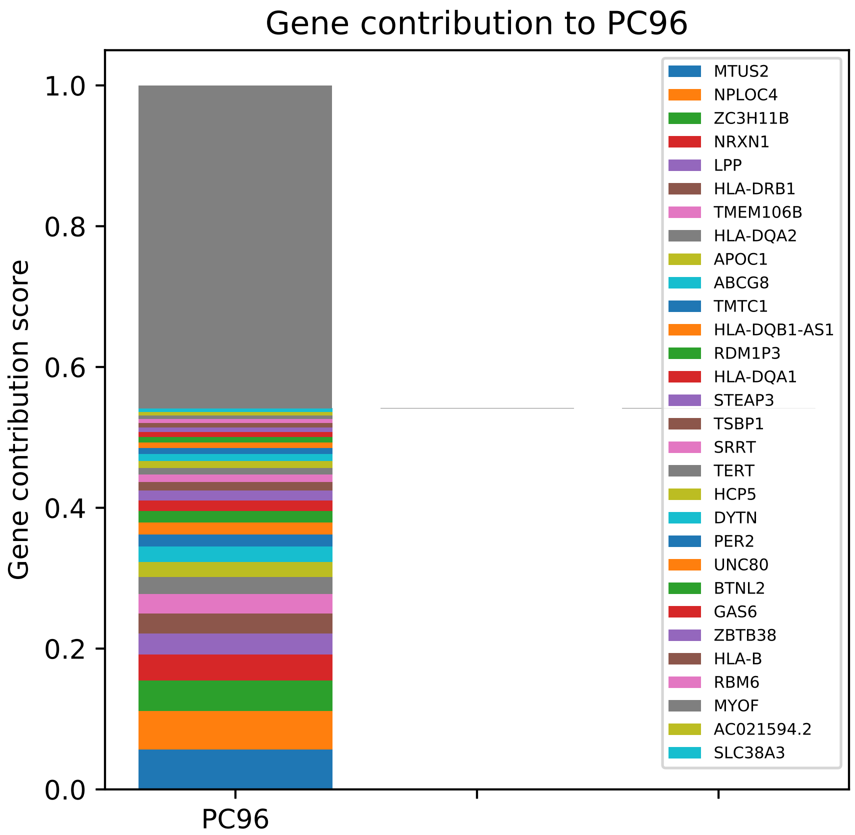

Gene contribution to PC96
Gene contribution score
1.0
0.8
0.6
0.4
0.2
0.0
PC96
MTUS2
NPLOC4
ZC3H11B
NRXN1
LPP
HLA-DRB1
TMEM106B
HLA-DQA2
APOC1
ABCG8
TMTC1
HLA-DQB1-AS1
RDM1P3
HLA-DQA1
STEAP3
TSBP1
SRRT
TERT
HCP5
DYTN
PER2
UNC80
BTNL2
GAS6
ZBTB38
HLA-B
RBM6
MYOF
AC021594.2
SLC38A3

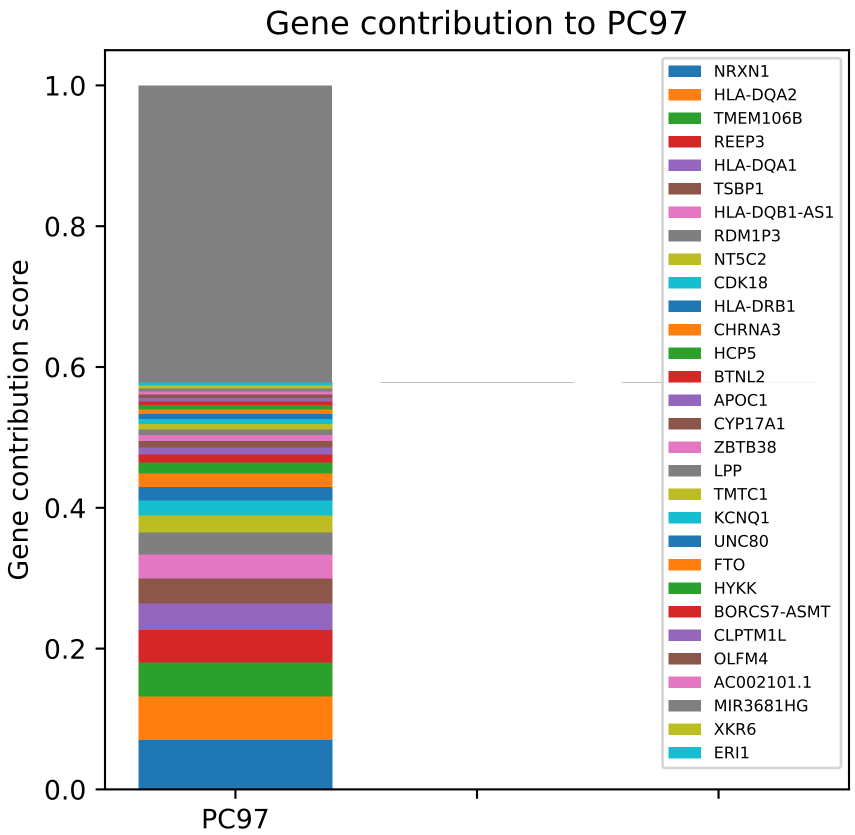
Gene contribution to PC97
Gene contribution score
1.0
0.8
0.6
0.4
0.2
0.0
PC97
NRXN1
HLA-DQA2
TMEM106B
REEP3
HLA-DQA1
TSBP1
HLA-DQB1-AS1
RDM1P3
NT5C2
CDK18
HLA-DRB1
CHRNA3
HCP5
BTNL2
APOC1
CYP17A1
ZBTB38
LPP
TMTC1
KCNQ1
UNC80
FTO
HYKK
BORCS7-ASMT
CLPTM1L
OLFM4
AC002101.1
MIR3681HG
XKR6
ERI1

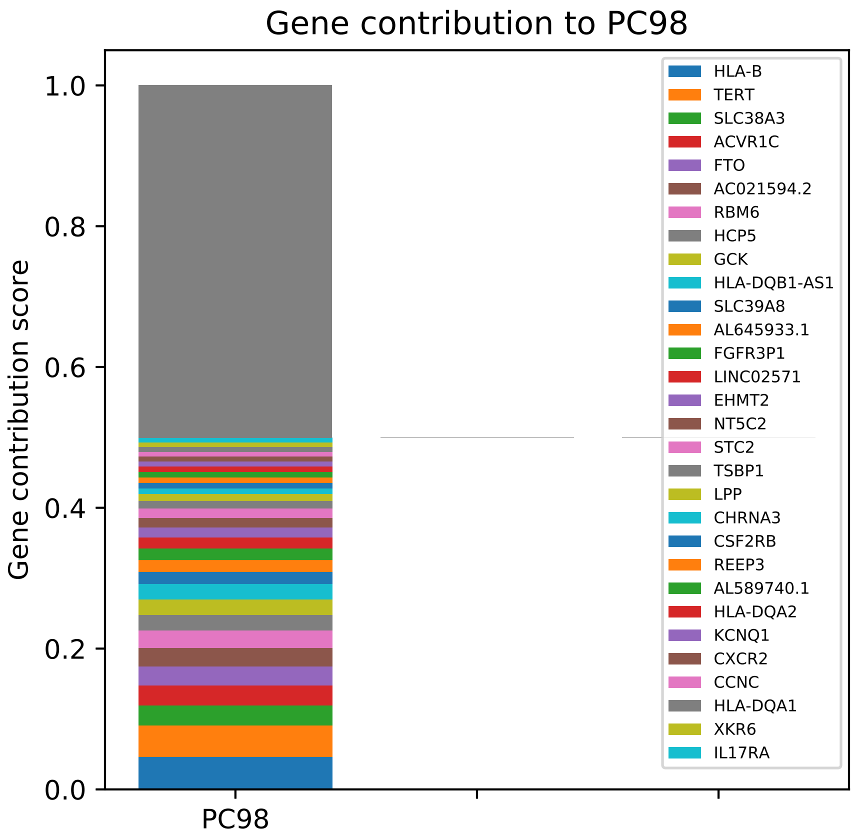

Gene contribution to PC98
Gene contribution score
1.0
0.8
0.6
0.4
0.2
0.0
PC98
HLA-B
TERT
SLC38A3
ACVR1C
FTO
AC021594.2
RBM6
HCP5
GCK
HLA-DQB1-AS1
SLC39A8
AL645933.1
FGFR3P1
LINC02571
EHMT2
NT5C2
STC2
TSBP1
LPP
CHRNA3
CSF2RB
REEP3
AL589740.1
HLA-DQA2
KCNQ1
CXCR2
CCNC
HLA-DQA1
XKR6
IL17RA

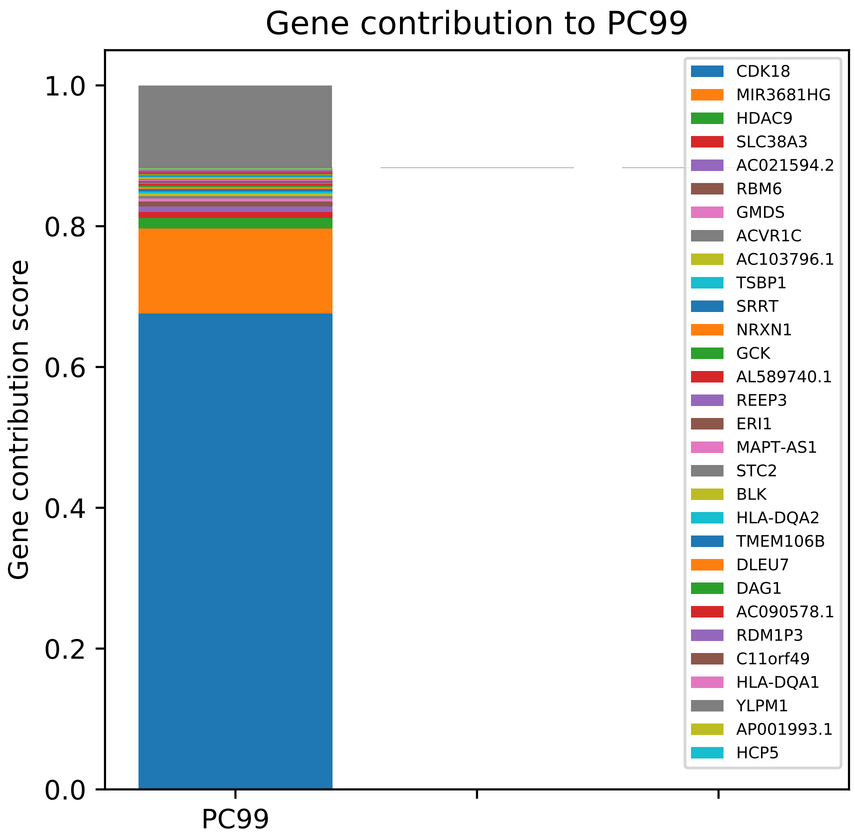

Gene contribution to PC99
Gene contribution score
1.0
0.8
0.6
0.4
0.2
0.0
PC99
CDK18
MIR3681HG
HDAC9
SLC38A3
AC021594.2
RBM6
GMDS
ACVR1C
AC103796.1
TSBP1
SRRT
NRXN1
GCK
AL589740.1
REEP3
ERI1
MAPT-AS1
STC2
BLK
HLA-DQA2
TMEM106B
DLEU7
DAG1
AC090578.1
RDM1P3
C11orf49
HLA-DQA1
YLPM1
AP001993.1
HCP5

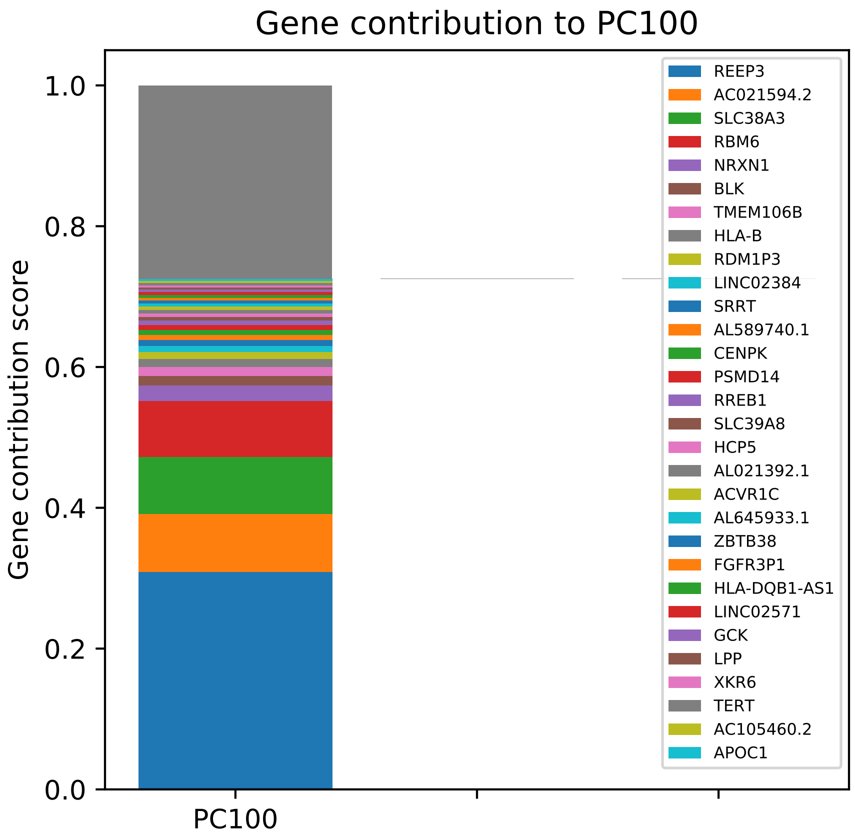

Gene contribution to PC100
Gene contribution score
1.0
0.8
0.6
0.4
0.2
0.0
PC100
REEP3
AC021594.2
SLC38A3
RBM6
NRXN1
BLK
TMEM106B
HLA-B
RDM1P3
LINC02384
SRRT
AL589740.1
CENPK
PSMD14
RREB1
SLC39A8
HCP5
AL021392.1
ACVR1C
AL645933.1
ZBTB38
FGFR3P1
HLA-DQB1-AS1
LINC02571
GCK
LPP
XKR6
TERT
AC105460.2
APOC1

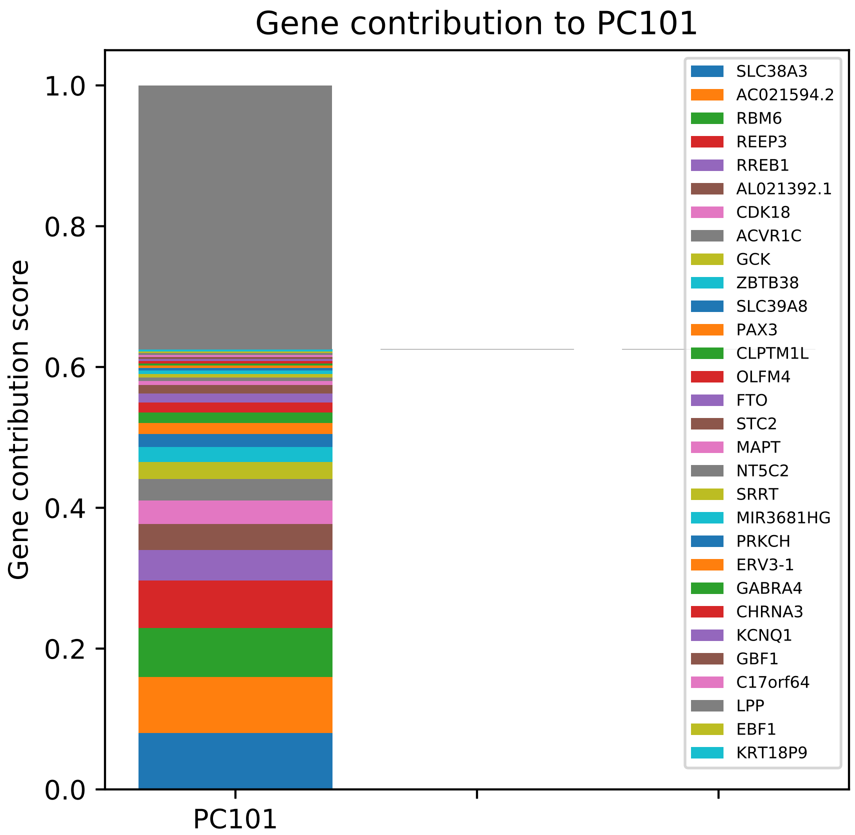

Gene contribution to PC101
Gene contribution score
1.0
0.8
0.6
0.4
0.2
0.0
PC101
SLC38A3
AC021594.2
RBM6
REEP3
RREB1
AL021392.1
CDK18
ACVR1C
GCK
ZBTB38
SLC39A8
PAX3
CLPTM1L
OLFM4
FTO
STC2
MAPT
NT5C2
SRRT
MIR3681HG
PRKCH
ERV3-1
GABRA4
CHRNA3
KCNQ1
GBF1
C17orf64
LPP
EBF1
KRT18P9

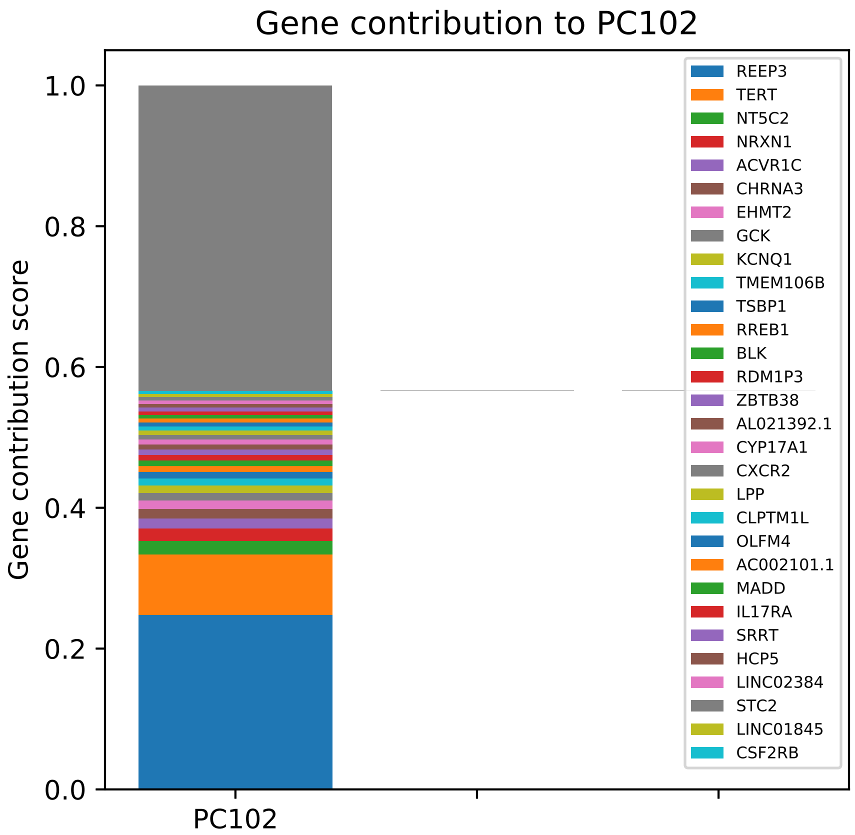

Gene contribution to PC102
Gene contribution score
1.0
0.8
0.6
0.4
0.2
0.0
PC102
REEP3
TERT
NT5C2
NRXN1
ACVR1C
CHRNA3
EHMT2
GCK
KCNQ1
TMEM106B
TSBP1
RREB1
BLK
RDM1P3
ZBTB38
AL021392.1
CYP17A1
CXCR2
LPP
CLPTM1L
OLFM4
AC002101.1
MADD
IL17RA
SRRT
HCP5
LINC02384
STC2
LINC01845
CSF2RB

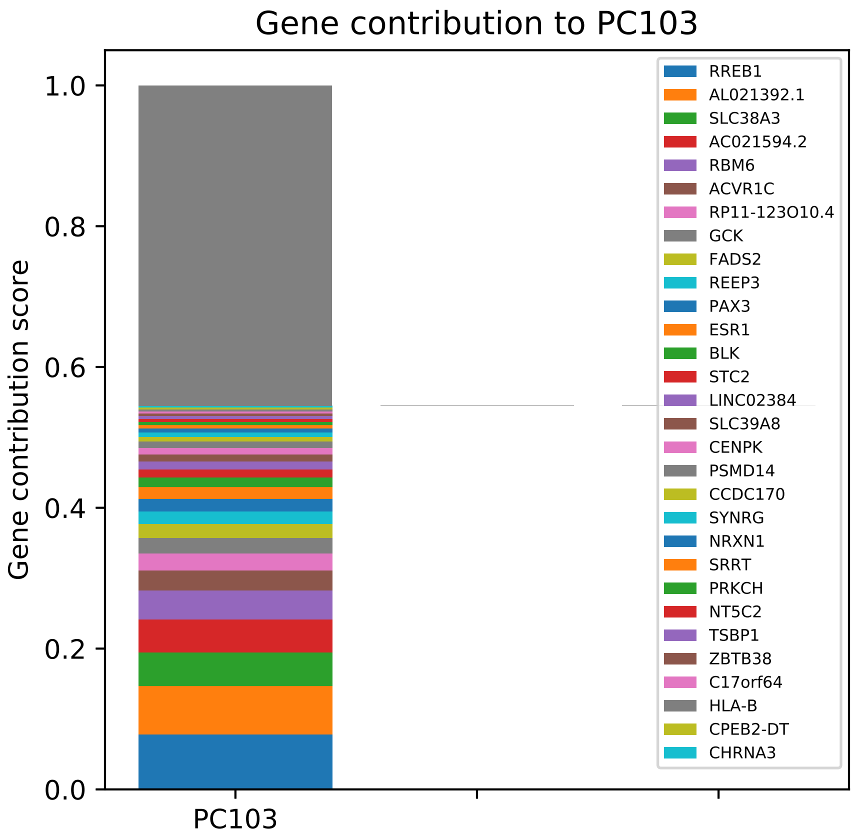

Gene contribution to PC103
Gene contribution score
1.0
0.8
0.6
0.4
0.2
0.0
PC103
RREB1
AL021392.1
SLC38A3
AC021594.2
RBM6
ACVR1C
RP11-123O10.4
GCK
FADS2
REEP3
PAX3
ESR1
BLK
STC2
LINC02384
SLC39A8
CENPK
PSMD14
CCDC170
SYNRG
NRXN1
SRRT
PRKCH
NT5C2
TSBP1
ZBTB38
C17orf64
HLA-B
CPEB2-DT
CHRNA3

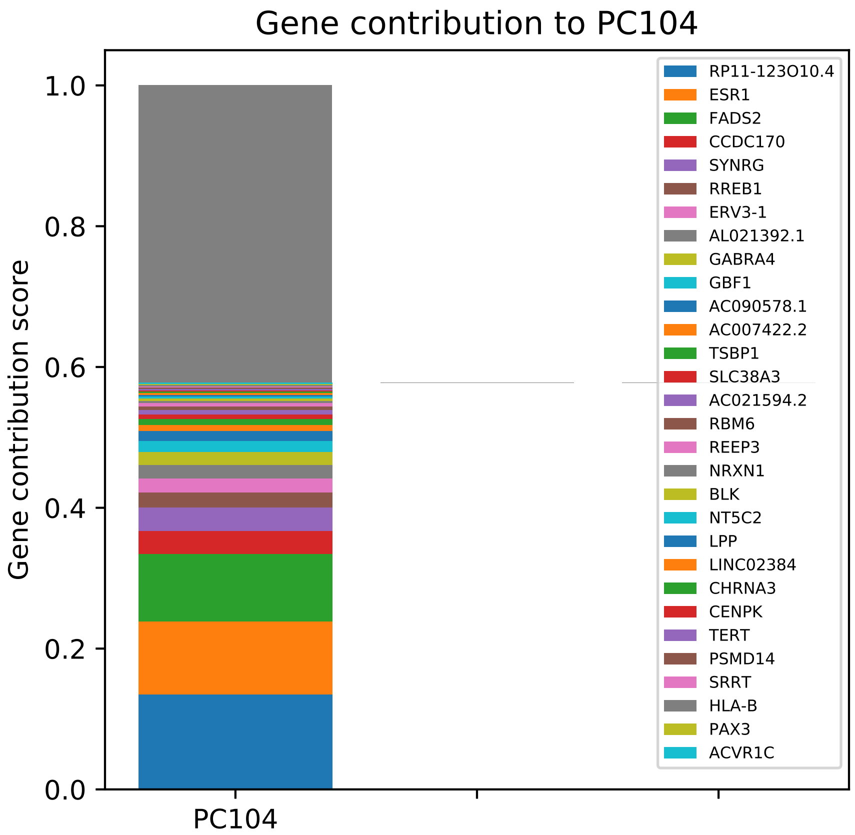

Gene contribution to PC104
Gene contribution score
0.0
0.2
0.4
0.6
0.8
1.0
PC104
RP11-123O10.4
ESR1
FADS2
CCDC170
SYNRG
RREB1
ERV3-1
AL021392.1
GABRA4
GBF1
AC090578.1
AC007422.2
TSBP1
SLC38A3
AC021594.2
RBM6
REEP3
NRXN1
BLK
NT5C2
LPP
LINC02384
CHRNA3
CENPK
TERT
PSMD14
SRRT
HLA-B
PAX3
ACVR1C

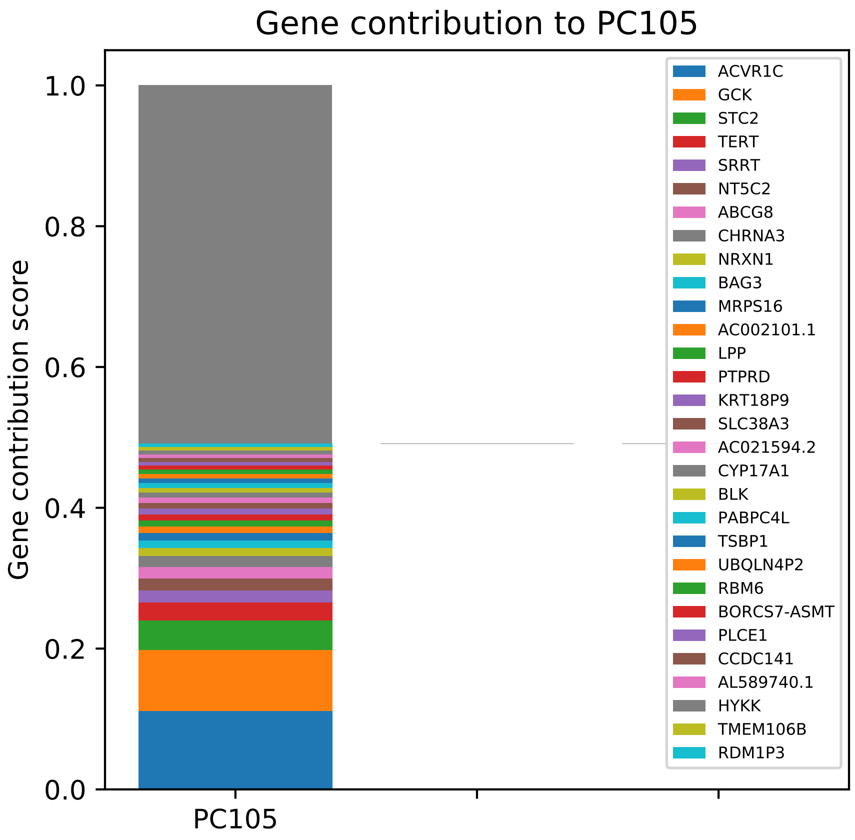
Gene contribution to PC105
Gene contribution score
1.0
0.8
0.6
0.4
0.2
0.0
PC105
ACVR1C
GCK
STC2
TERT
SRRT
NT5C2
ABCG8
CHRNA3
NRXN1
BAG3
MRPS16
AC002101.1
LPP
PTPRD
KRT18P9
SLC38A3
AC021594.2
CYP17A1
BLK
PABPC4L
TSBP1
UBQLN4P2
RBM6
BORCS7-ASMT
PLCE1
CCDC141
AL589740.1
HYKK
TMEM106B
RDM1P3

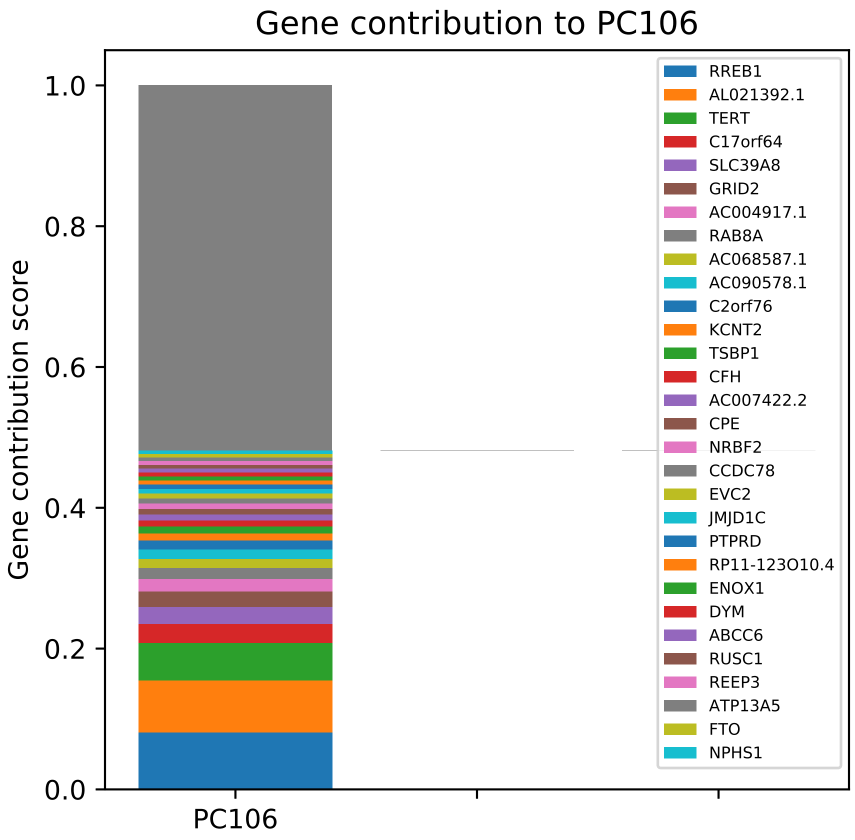
Gene contribution to PC106
Gene contribution score
0.0
0.2
0.4
0.6
0.8
1.0
PC106
RREB1
AL021392.1
TERT
C17orf64
SLC39A8
GRID2
AC004917.1
RAB8A
AC068587.1
AC090578.1
C2orf76
KCNT2
TSBP1
CFH
AC007422.2
CPE
NRBF2
CCDC78
EVC2
JMJD1C
PTPRD
RP11-123O10.4
ENOX1
DYM
ABCC6
RUSC1
REEP3
ATP13A5
FTO
NPHS1

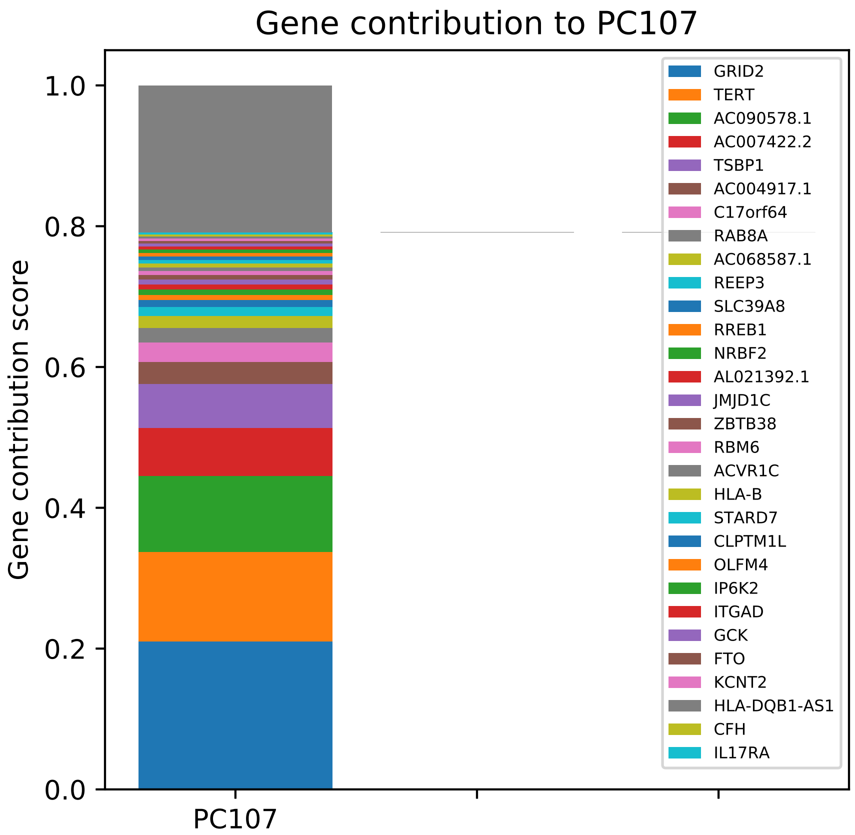

Gene contribution to PC107
Gene contribution score
0.0
0.2
0.4
0.6
0.8
1.0
PC107
GRID2
TERT
AC090578.1
AC007422.2
TSBP1
AC004917.1
C17orf64
RAB8A
AC068587.1
REEP3
SLC39A8
RREB1
NRBF2
AL021392.1
JMJD1C
ZBTB38
RBM6
ACVR1C
HLA-B
STARD7
CLPTM1L
OLFM4
IP6K2
ITGAD
GCK
FTO
KCNT2
HLA-DQB1-AS1
CFH
IL17RA

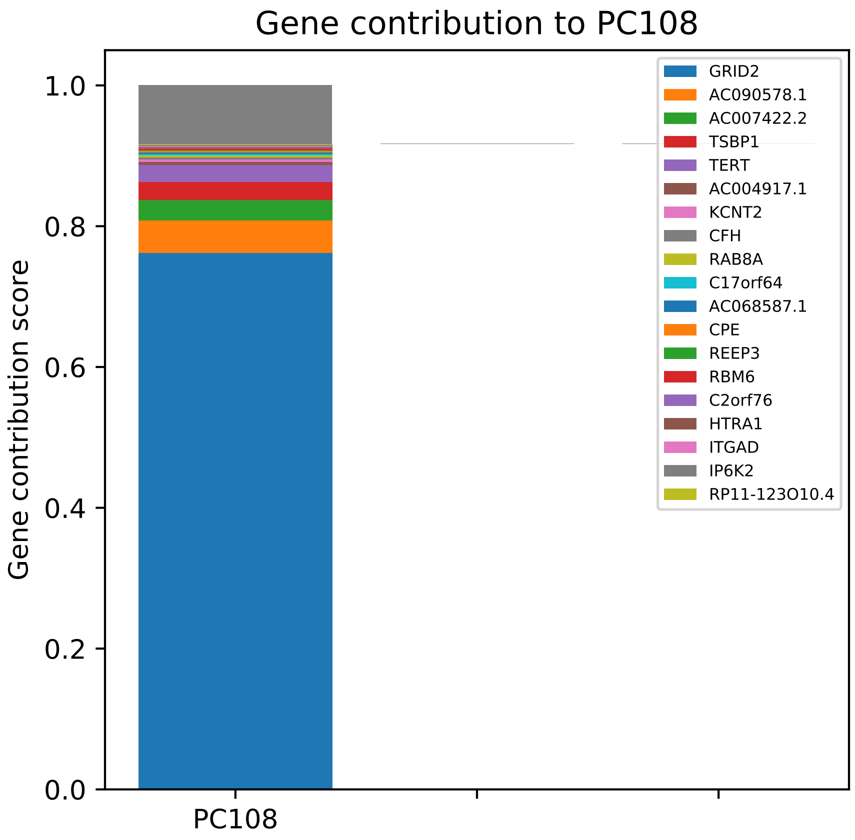

Gene contribution to PC108
Gene contribution score
0.0
0.2
0.4
0.6
0.8
1.0
PC108
GRID2
AC090578.1
AC007422.2
TSBP1
TERT
AC004917.1
KCNT2
CFH
RAB8A
C17orf64
AC068587.1
CPE
REEP3
RBM6
C2orf76
HTRA1
ITGAD
IP6K2
RP11-123O10.4

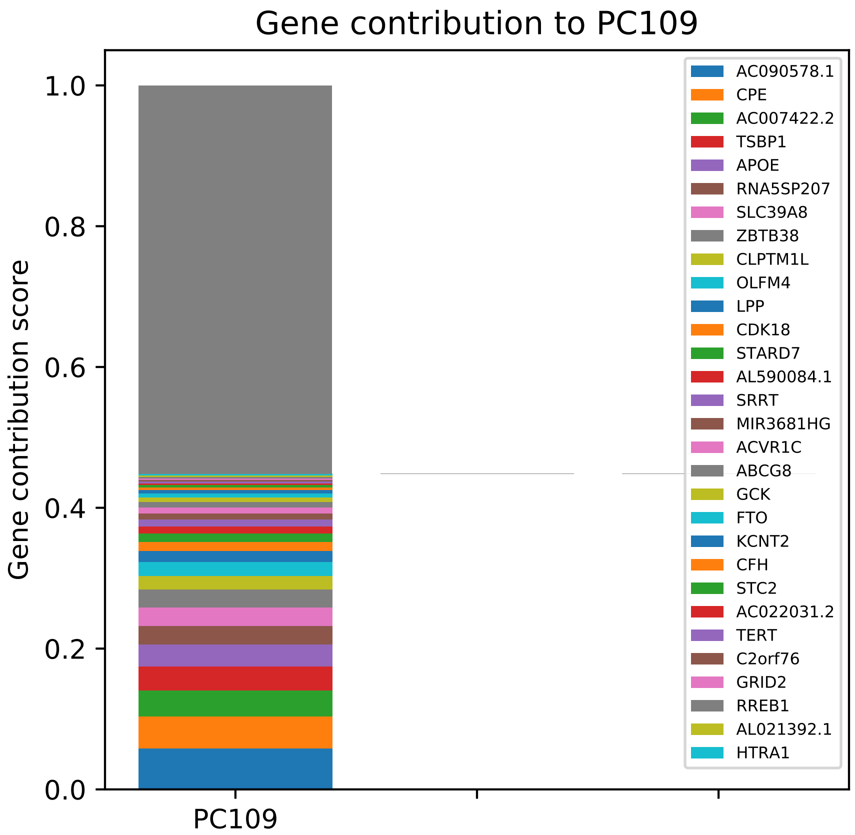
Gene contribution to PC109
Gene contribution score
1.0
0.8
0.6
0.4
0.2
0.0
PC109
AC090578.1
CPE
AC007422.2
TSBP1
APOE
RNA5SP207
SLC39A8
ZBTB38
CLPTM1L
OLFM4
LPP
CDK18
STARD7
AL590084.1
SRRT
MIR3681HG
ACVR1C
ABCG8
GCK
FTO
KCNT2
CFH
STC2
AC022031.2
TERT
C2orf76
GRID2
RREB1
AL021392.1
HTRA1

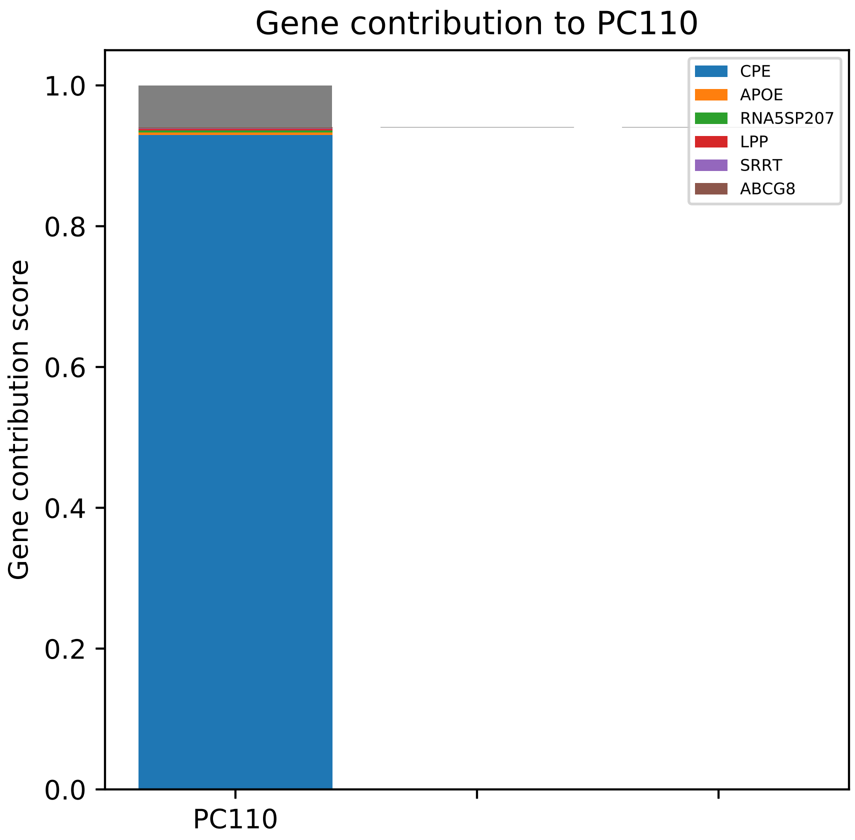
Gene contribution to PC110
CPE
APOE
RNA5SP207
LPP
SRRT
ABCG8
1.0
0.8
0.6
0.4
0.2
0.0
Gene contribution score
PC110

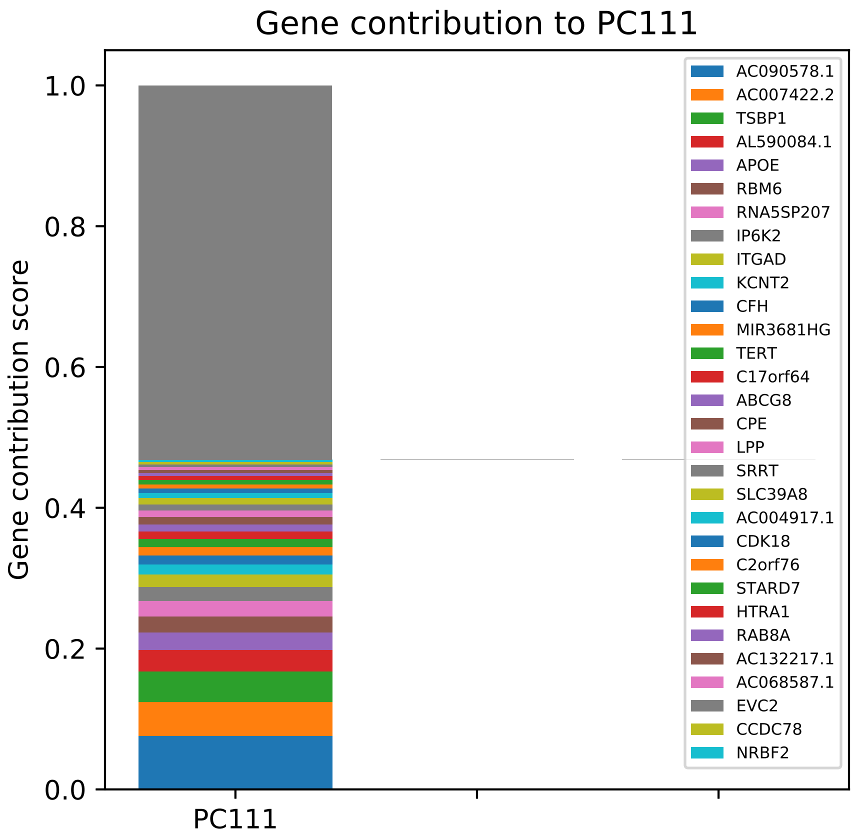

Gene contribution to PC111
Gene contribution score
1.0
0.8
0.6
0.4
0.2
0.0
PC111
AC090578.1
AC007422.2
TSBP1
AL590084.1
APOE
RBM6
RNA5SP207
IP6K2
ITGAD
KCNT2
CFH
MIR3681HG
TERT
C17orf64
ABCG8
CPE
LPP
SRRT
SLC39A8
AC004917.1
CDK18
C2orf76
STARD7
HTRA1
RAB8A
AC132217.1
AC068587.1
EVC2
CCDC78
NRBF2

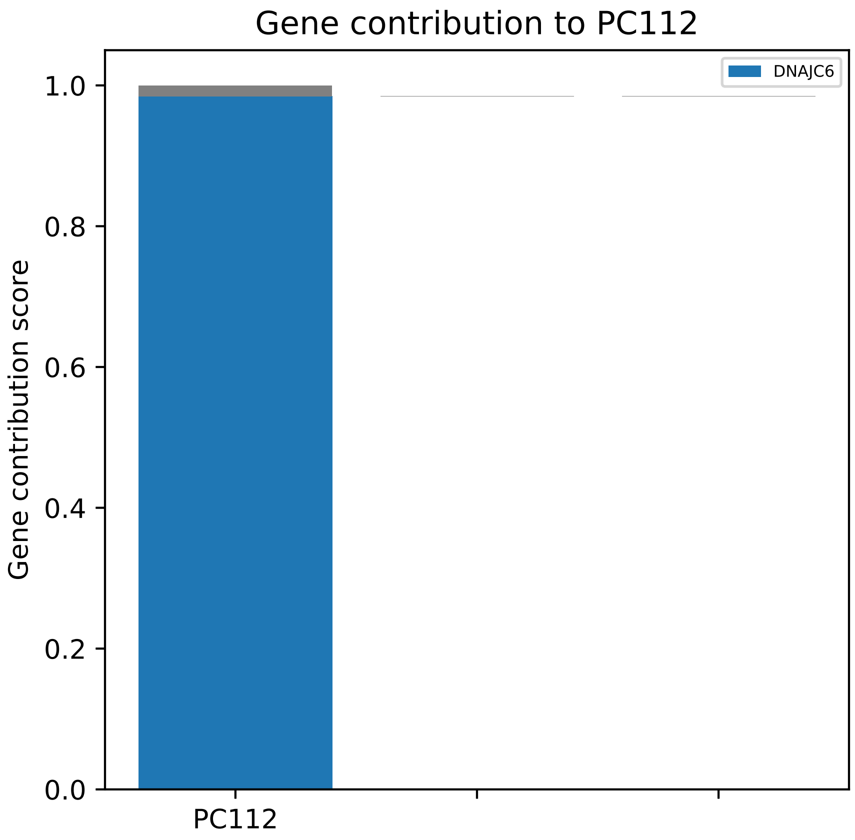

Gene contribution to PC112
DNAJC6
1.0
0.8
0.6
0.4
0.2
0.0
Gene contribution score
PC112

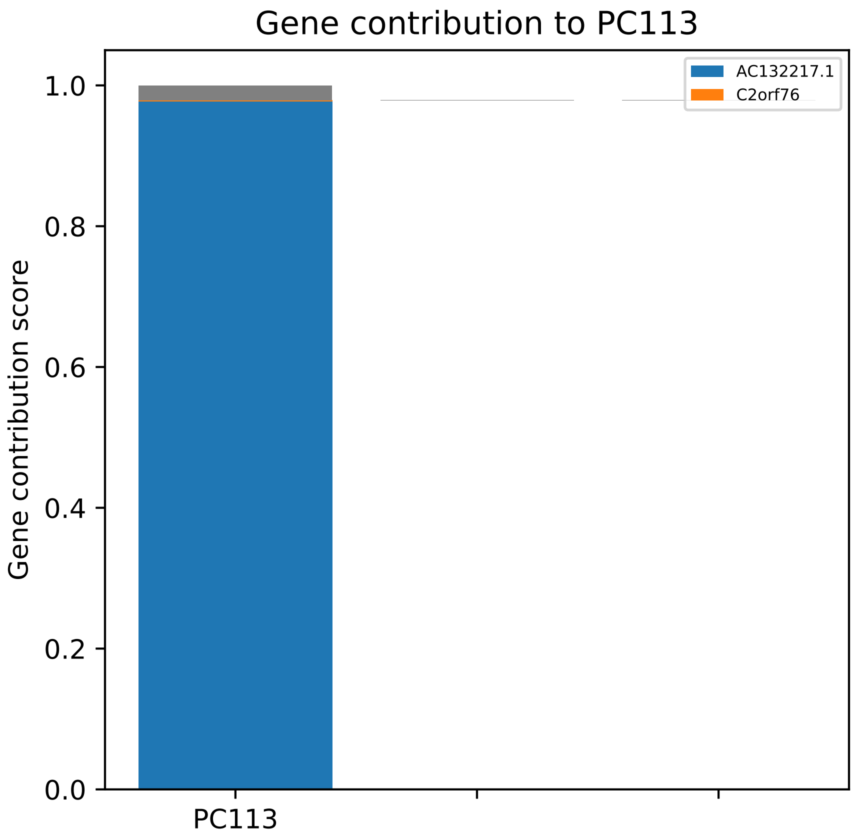

Gene contribution to PC113
Gene contribution score
1.0
0.8
0.6
0.4
0.2
0.0
PC113
AC132217.1
C2orf76

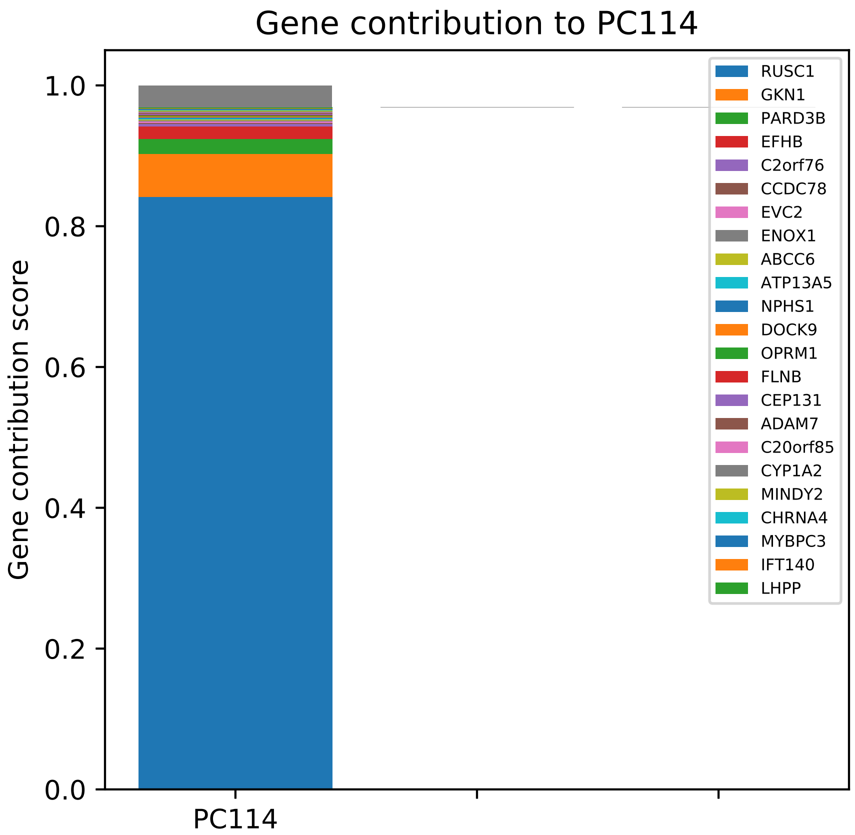
Gene contribution to PC114
Gene contribution score
1.0
0.8
0.6
0.4
0.2
0.0
PC114
RUSC1
GKN1
PARD3B
EFHB
C2orf76
CCDC78
EVC2
ENOX1
ABCC6
ATP13A5
NPHS1
DOCK9
OPRM1
FLNB
CEP131
ADAM7
C20orf85
CYP1A2
MINDY2
CHRNA4
MYBPC3
IFT140
LHPP

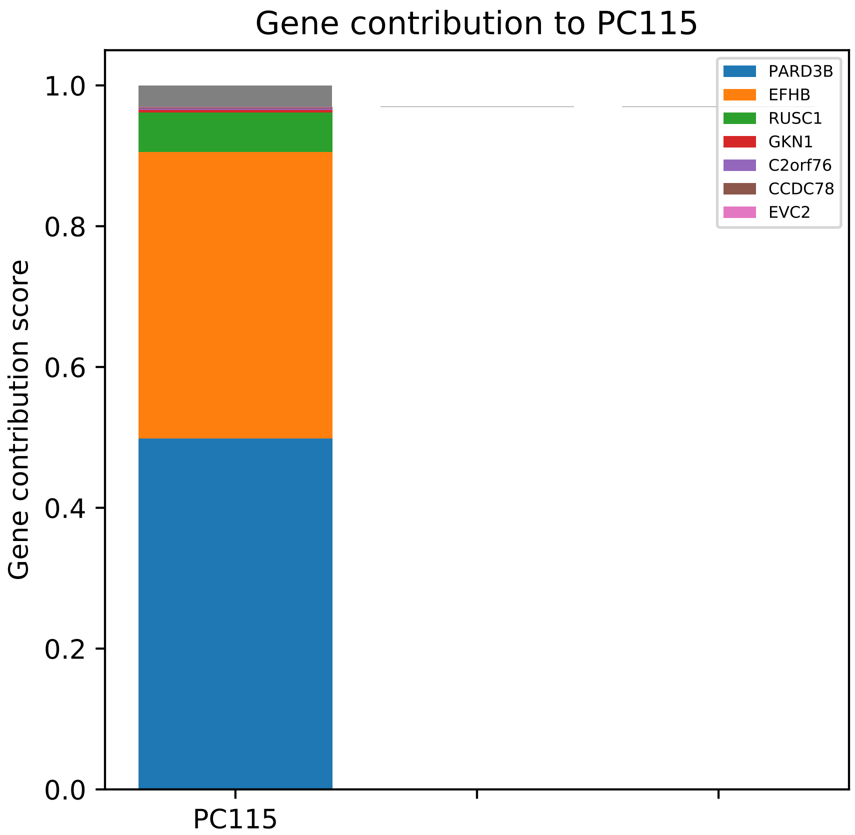
Gene contribution to PC115
Gene contribution score
1.0
0.8
0.6
0.4
0.2
0.0
PC115
PARD3B
EFHB
RUSC1
GKN1
C2orf76
CCDC78
EVC2

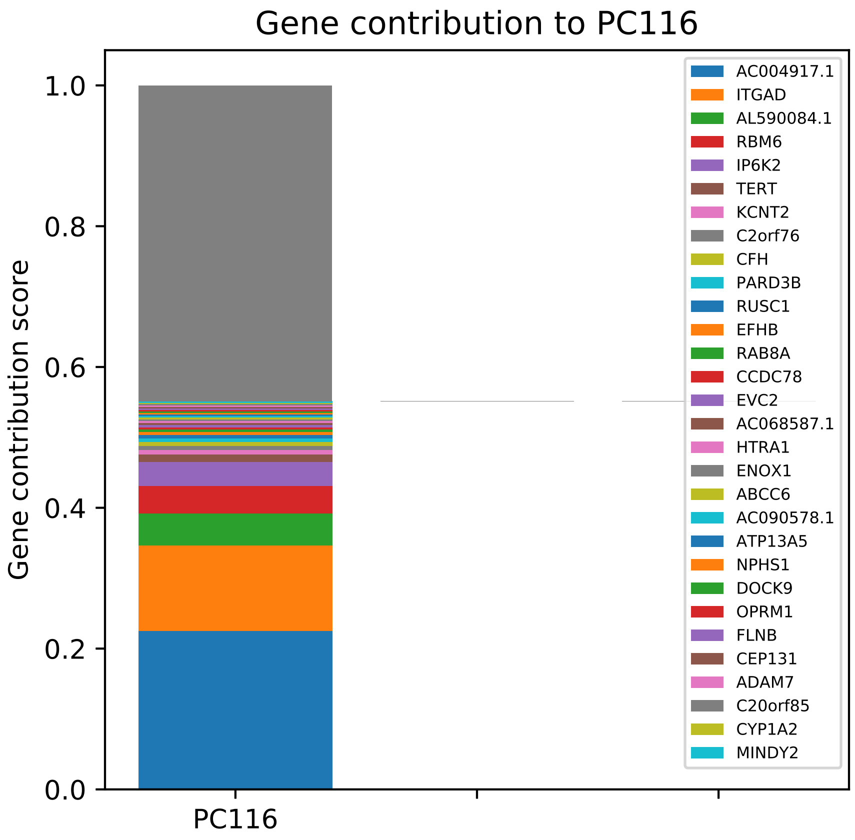

Gene contribution to PC116
Gene contribution score
1.0
0.8
0.6
0.4
0.2
0.0
PC116
AC004917.1
ITGAD
AL590084.1
RBM6
IP6K2
TERT
KCNT2
C2orf76
CFH
PARD3B
RUSC1
EFHB
RAB8A
CCDC78
EVC2
AC068587.1
HTRA1
ENOX1
ABCC6
AC090578.1
ATP13A5
NPHS1
DOCK9
OPRM1
FLNB
CEP131
ADAM7
C20orf85
CYP1A2
MINDY2

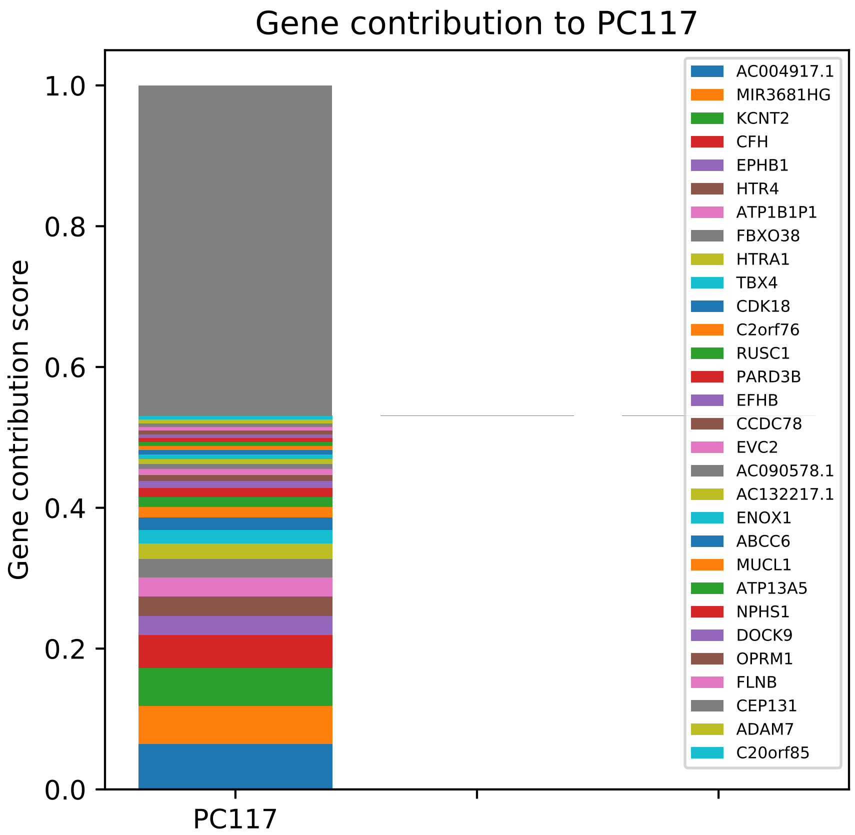

Gene contribution to PC117
Gene contribution score
0.0
0.2
0.4
0.6
0.8
1.0
PC117
AC004917.1
MIR3681HG
KCNT2
CFH
EPHB1
HTR4
ATP1B1P1
FBXO38
HTRA1
TBX4
CDK18
C2orf76
RUSC1
PARD3B
EFHB
CCDC78
EVC2
AC090578.1
AC132217.1
ENOX1
ABCC6
MUCL1
ATP13A5
NPHS1
DOCK9
OPRM1
FLNB
CEP131
ADAM7
C20orf85

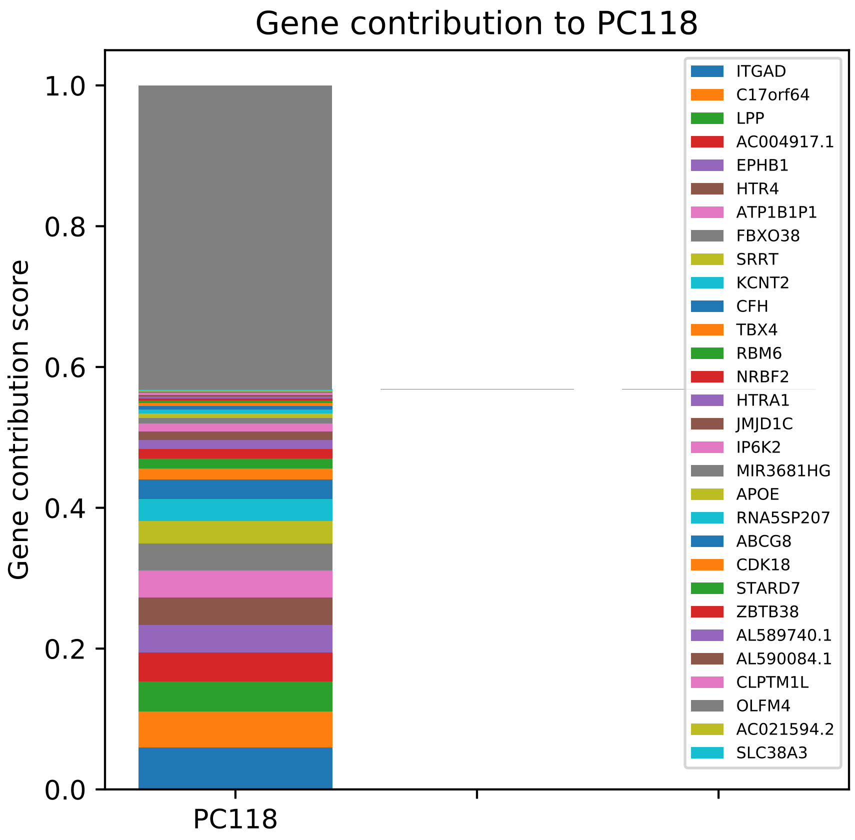

Gene contribution to PC118
Gene contribution score
1.0
0.8
0.6
0.4
0.2
0.0
PC118
ITGAD
C17orf64
LPP
AC004917.1
EPHB1
HTR4
ATP1B1P1
FBXO38
SRRT
KCNT2
CFH
TBX4
RBM6
NRBF2
HTRA1
JMJD1C
IP6K2
MIR3681HG
APOE
RNA5SP207
ABCG8
CDK18
STARD7
ZBTB38
AL589740.1
AL590084.1
CLPTM1L
OLFM4
AC021594.2
SLC38A3

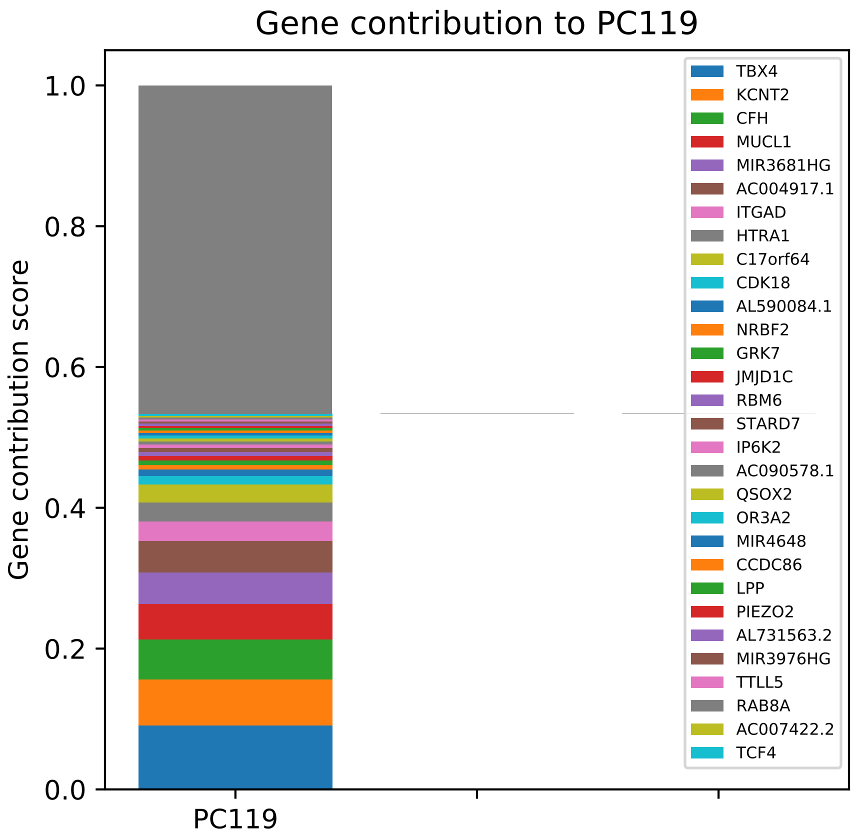

Gene contribution to PC119
Gene contribution score
1.0
0.8
0.6
0.4
0.2
0.0
PC119
TBX4
KCNT2
CFH
MUCL1
MIR3681HG
AC004917.1
ITGAD
HTRA1
C17orf64
CDK18
AL590084.1
NRBF2
GRK7
JMJD1C
RBM6
STARD7
IP6K2
AC090578.1
QSOX2
OR3A2
MIR4648
CCDC86
LPP
PIEZO2
AL731563.2
MIR3976HG
TTLL5
RAB8A
AC007422.2
TCF4

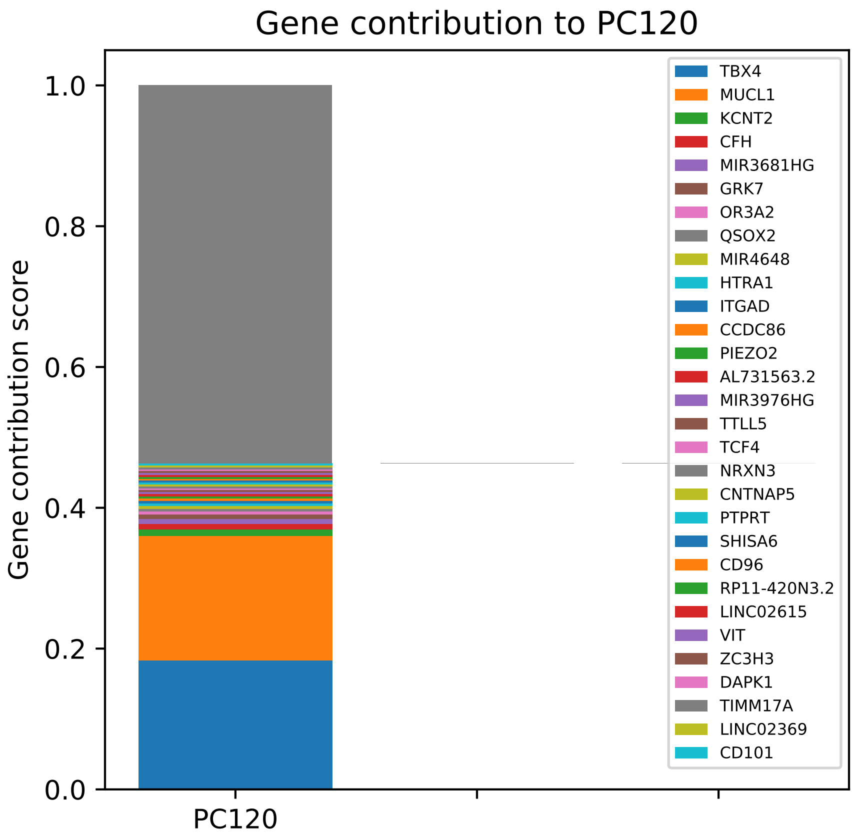
Gene contribution to PC120
Gene contribution score
1.0
0.8
0.6
0.4
0.2
0.0
PC120
TBX4
MUCL1
KCNT2
CFH
MIR3681HG
GRK7
OR3A2
QSOX2
MIR4648
HTRA1
ITGAD
CCDC86
PIEZO2
AL731563.2
MIR3976HG
TTLL5
TCF4
NRXN3
CNTNAP5
PTPRT
SHISA6
CD96
RP11-420N3.2
LINC02615
VIT
ZC3H3
DAPK1
TIMM17A
LINC02369
CD101

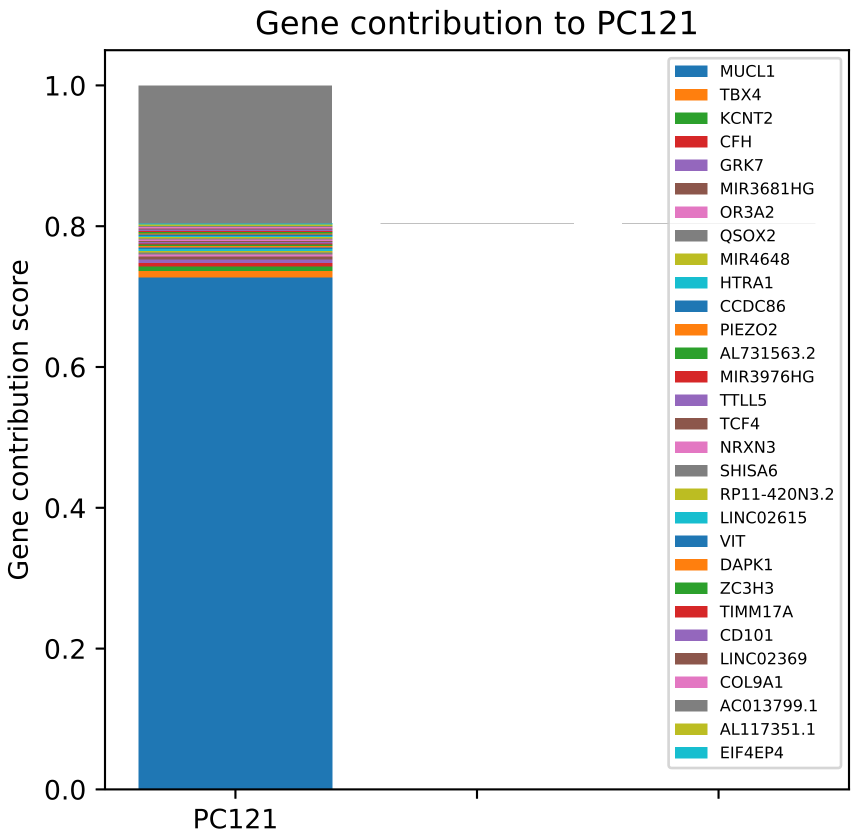

Gene contribution to PC121
Gene contribution score
1.0
0.8
0.6
0.4
0.2
0.0
PC121
MUCL1
TBX4
KCNT2
CFH
GRK7
MIR3681HG
OR3A2
QSOX2
MIR4648
HTRA1
CCDC86
PIEZO2
AL731563.2
MIR3976HG
TTLL5
TCF4
NRXN3
SHISA6
RP11-420N3.2
LINC02615
VIT
DAPK1
ZC3H3
TIMM17A
CD101
LINC02369
COL9A1
AC013799.1
AL117351.1
EIF4EP4

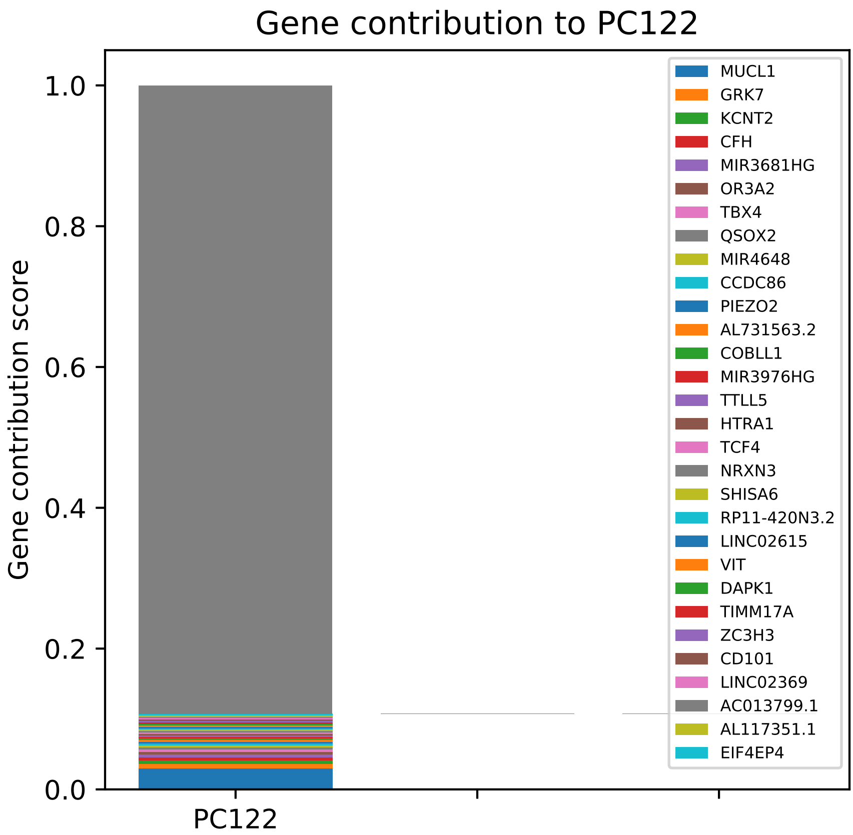

Gene contribution to PC122
Gene contribution score
1.0
0.8
0.6
0.4
0.2
0.0
PC122
MUCL1
GRK7
KCNT2
CFH
MIR3681HG
OR3A2
TBX4
QSOX2
MIR4648
CCDC86
PIEZO2
AL731563.2
COBLL1
MIR3976HG
TTLL5
HTRA1
TCF4
NRXN3
SHISA6
RP11-420N3.2
LINC02615
VIT
DAPK1
TIMM17A
ZC3H3
CD101
LINC02369
AC013799.1
AL117351.1
EIF4EP4

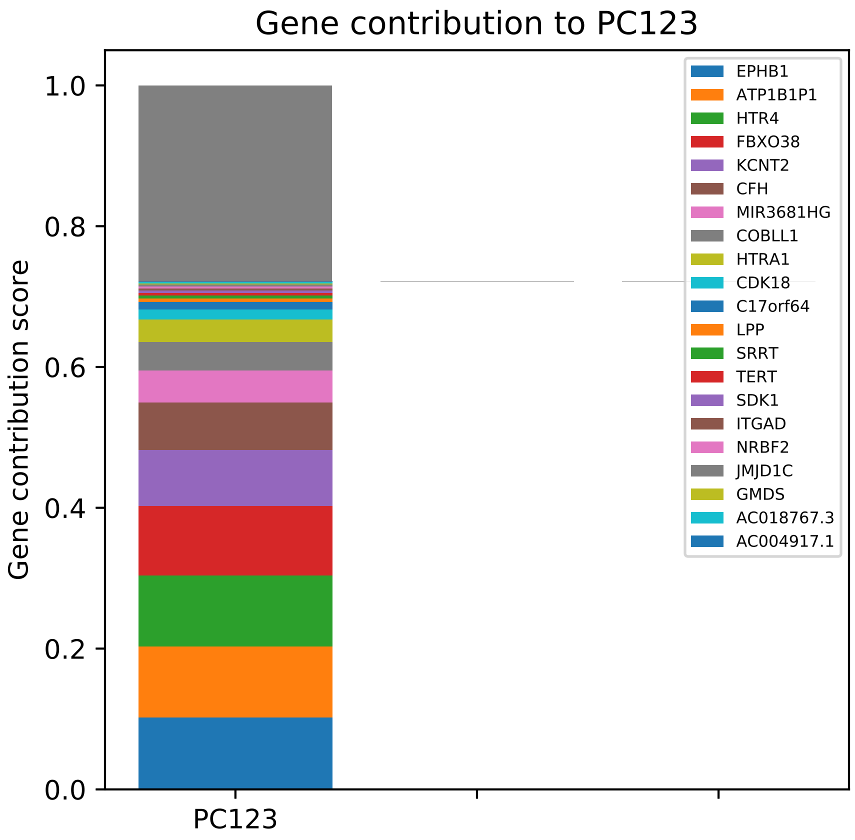

Gene contribution to PC123
Gene contribution score
1.0
0.8
0.6
0.4
0.2
0.0
PC123
EPHB1
ATP1B1P1
HTR4
FBXO38
KCNT2
CFH
MIR3681HG
COBLL1
HTRA1
CDK18
C17orf64
LPP
SRRT
TERT
SDK1
ITGAD
NRBF2
JMJD1C
GMDS
AC018767.3
AC004917.1

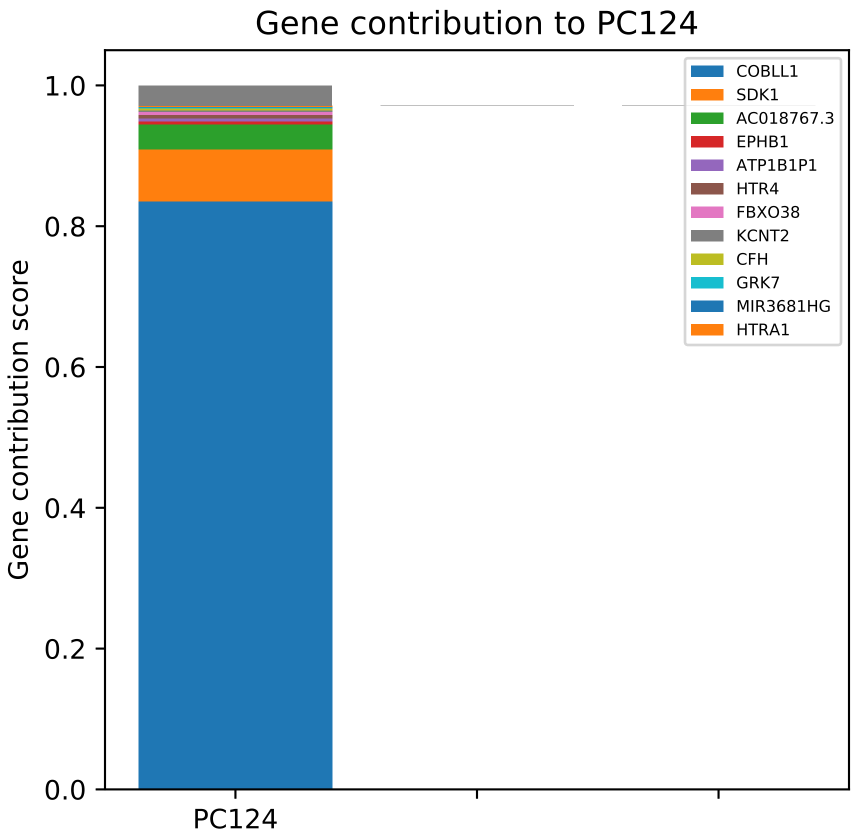

Gene contribution to PC124
Gene contribution score
1.0
0.8
0.6
0.4
0.2
0.0
PC124
COBLL1
SDK1
AC018767.3
EPHB1
ATP1B1P1
HTR4
FBXO38
KCNT2
CFH
GRK7
MIR3681HG
HTRA1

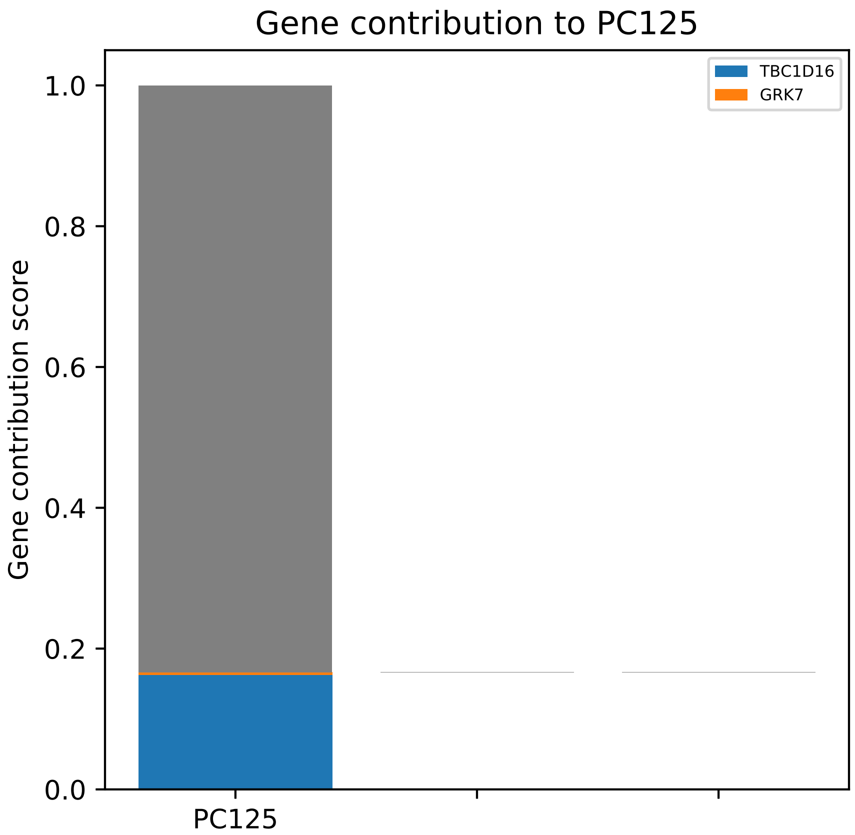
Gene contribution to PC125
Gene contribution score
1.0
0.8
0.6
0.4
0.2
0.0
PC125
TBC1D16
GRK7

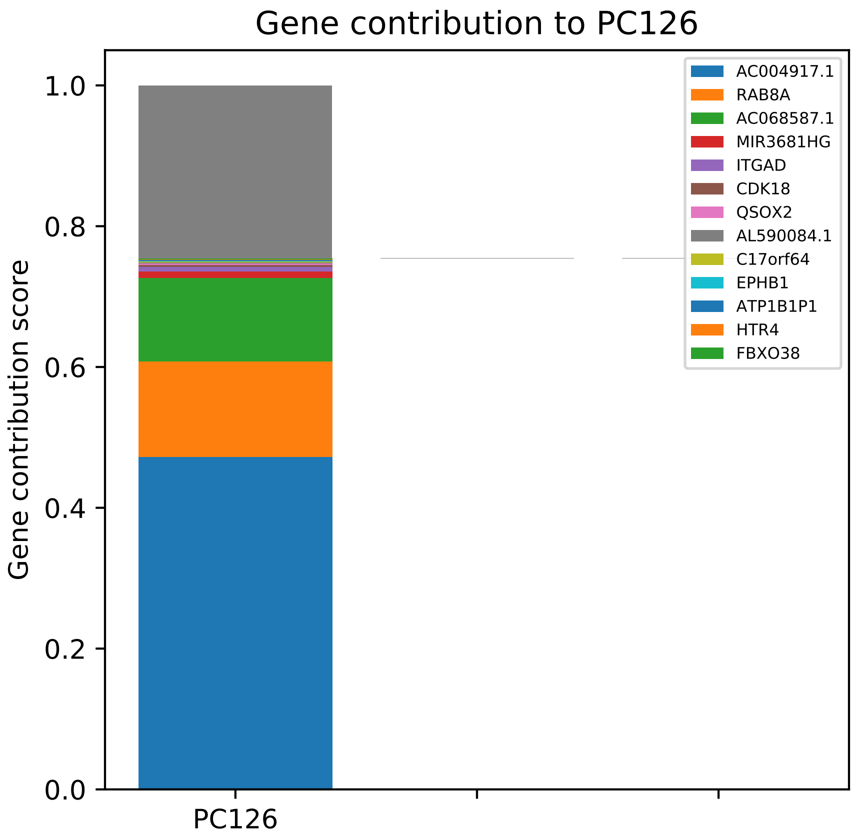
Gene contribution to PC126
Gene contribution score
0.0
0.2
0.4
0.6
0.8
1.0
PC126
AC004917.1
RAB8A
AC068587.1
MIR3681HG
ITGAD
CDK18
QSOX2
AL590084.1
C17orf64
EPHB1
ATP1B1P1
HTR4
FBXO38

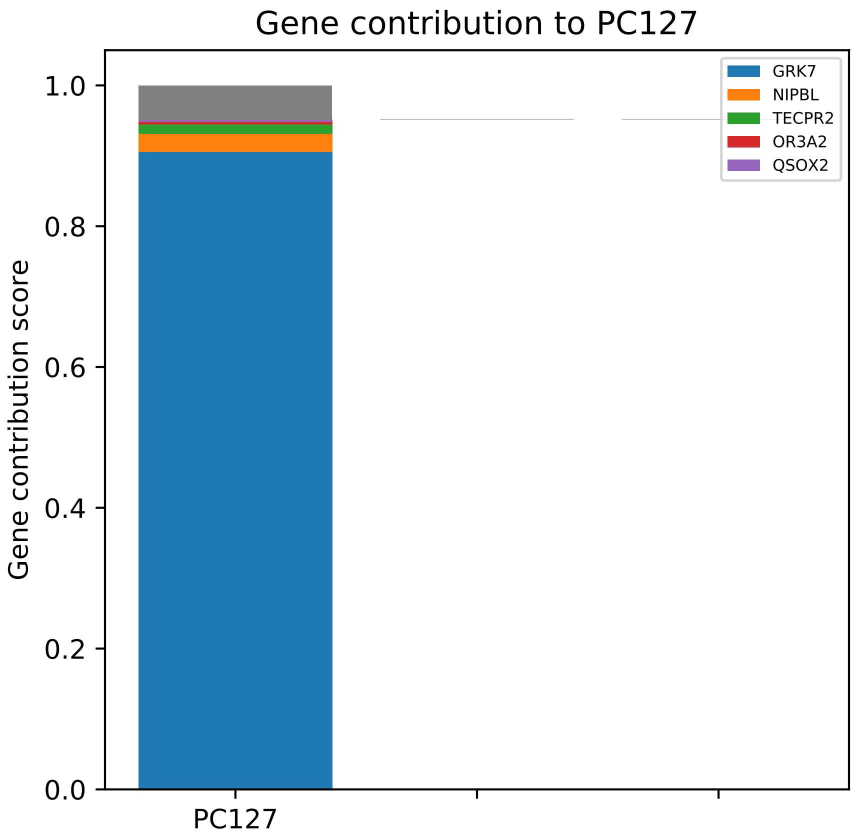

Gene contribution to PC127
Gene contribution score
0.0
0.2
0.4
0.6
0.8
1.0
PC127
GRK7
NIPBL
TECPR2
OR3A2
QSOX2

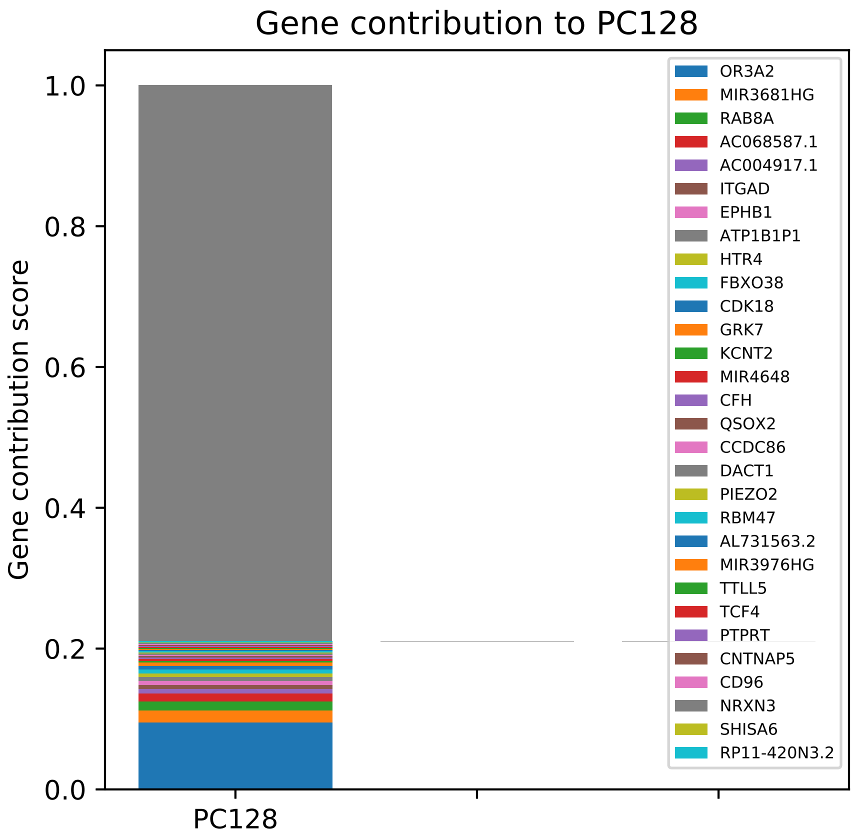

Gene contribution to PC128
Gene contribution score
1.0
0.8
0.6
0.4
0.2
0.0
PC128
OR3A2
MIR3681HG
RAB8A
AC068587.1
AC004917.1
ITGAD
EPHB1
ATP1B1P1
HTR4
FBXO38
CDK18
GRK7
KCNT2
MIR4648
CFH
QSOX2
CCDC86
DACT1
PIEZO2
RBM47
AL731563.2
MIR3976HG
TTLL5
TCF4
PTPRT
CNTNAP5
CD96
NRXN3
SHISA6
RP11-420N3.2

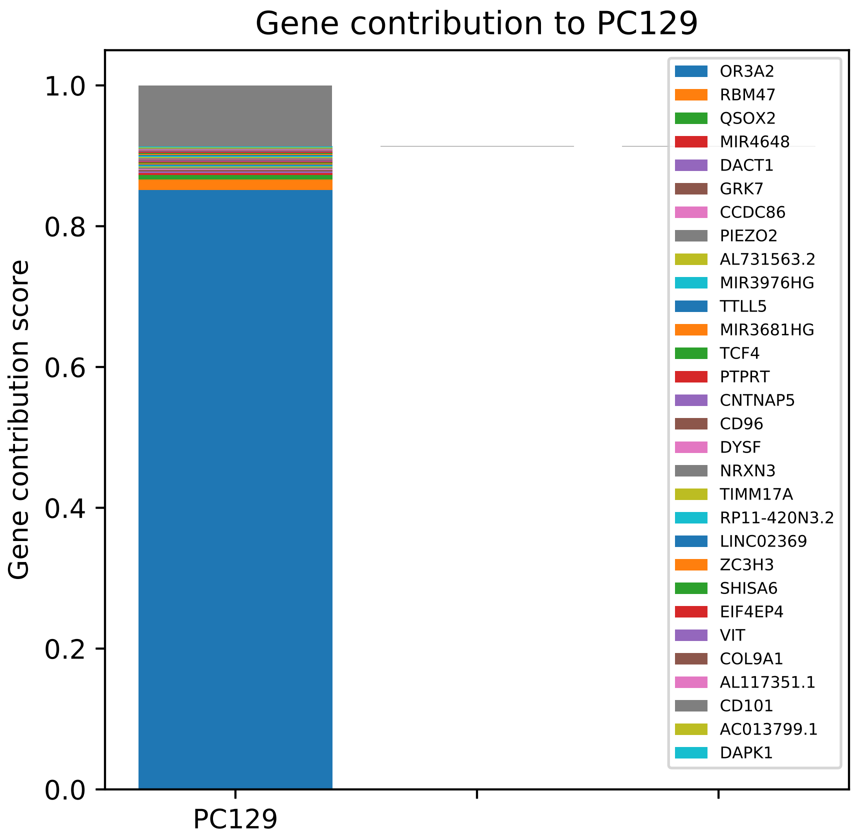
Gene contribution to PC129
Gene contribution score
1.0
0.8
0.6
0.4
0.2
0.0
PC129
OR3A2
RBM47
QSOX2
MIR4648
DACT1
GRK7
CCDC86
PIEZO2
AL731563.2
MIR3976HG
TTLL5
MIR3681HG
TCF4
PTPRT
CNTNAP5
CD96
DYSF
NRXN3
TIMM17A
RP11-420N3.2
LINC02369
ZC3H3
SHISA6
EIF4EP4
VIT
COL9A1
AL117351.1
CD101
AC013799.1
DAPK1

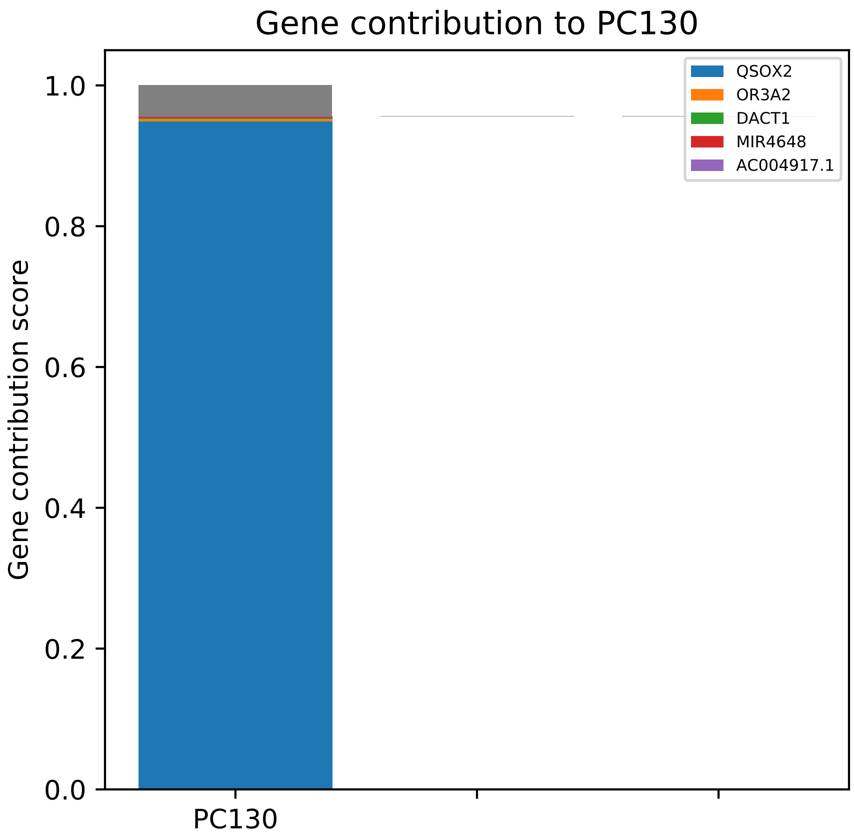
Gene contribution to PC130
Gene contribution score
0.0
0.2
0.4
0.6
0.8
1.0
PC130
QSOX2
OR3A2
DACT1
MIR4648
AC004917.1

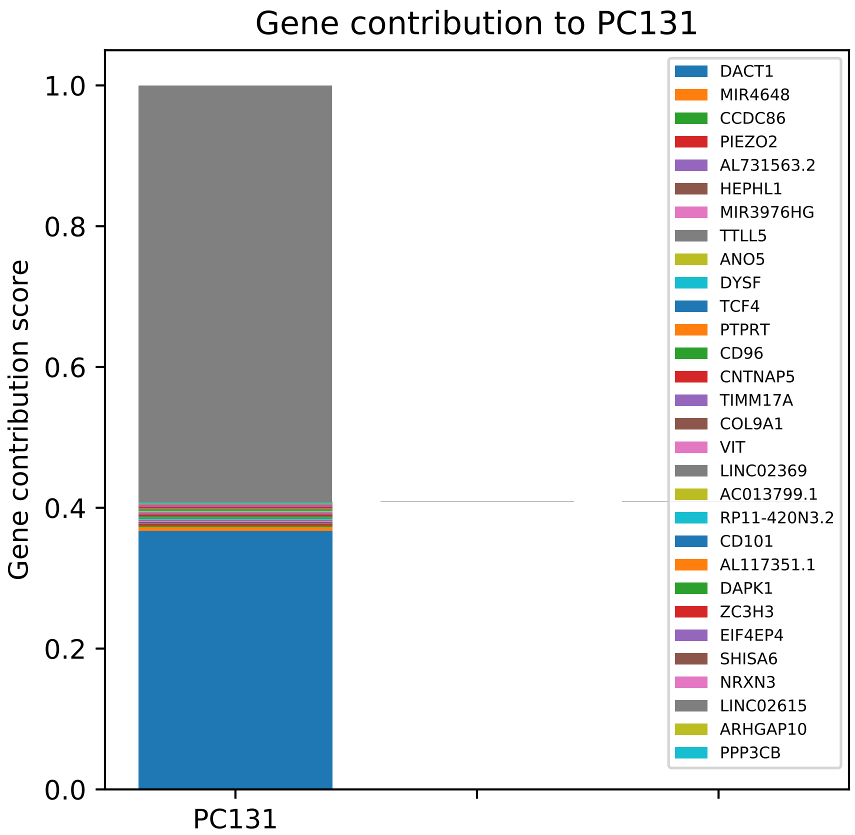
Gene contribution to PC131
Gene contribution score
1.0
0.8
0.6
0.4
0.2
0.0
PC131
DACT1
MIR4648
CCDC86
PIEZO2
AL731563.2
HEPHL1
MIR3976HG
TTLL5
ANO5
DYSF
TCF4
PTPRT
CD96
CNTNAP5
TIMM17A
COL9A1
VIT
LINC02369
AC013799.1
RP11-420N3.2
CD101
AL117351.1
DAPK1
ZC3H3
EIF4EP4
SHISA6
NRXN3
LINC02615
ARHGAP10
PPP3CB

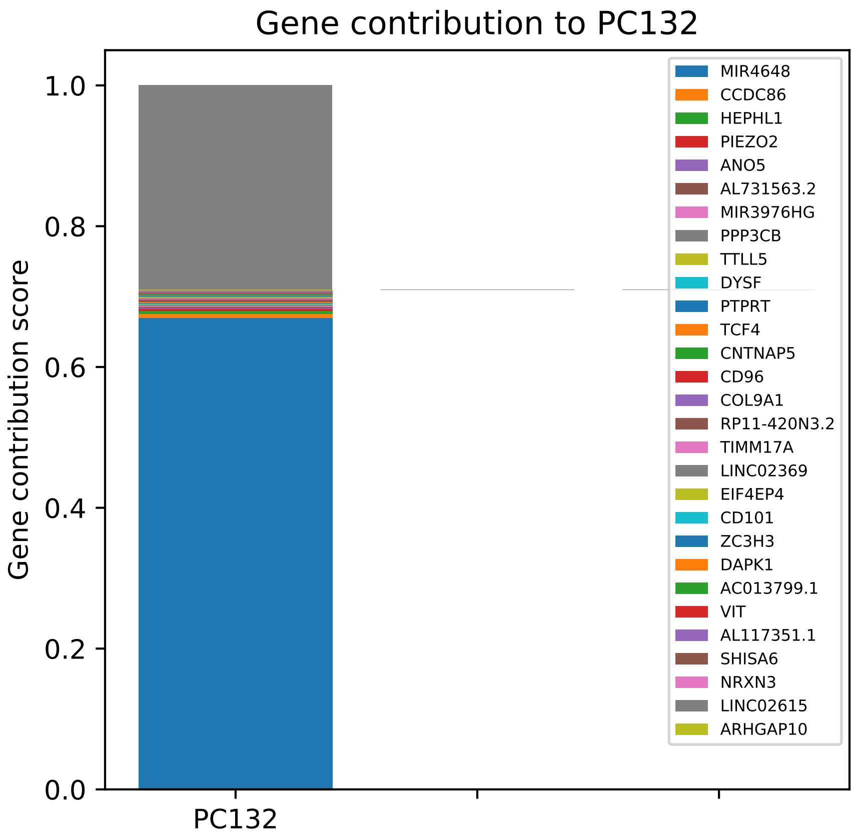

Gene contribution to PC132
Gene contribution score
1.0
0.8
0.6
0.4
0.2
0.0
PC132
MIR4648
CCDC86
HEPHL1
PIEZO2
ANO5
AL731563.2
MIR3976HG
PPP3CB
TTLL5
DYSF
PTPRT
TCF4
CNTNAP5
CD96
COL9A1
RP11-420N3.2
TIMM17A
LINC02369
EIF4EP4
CD101
ZC3H3
DAPK1
AC013799.1
VIT
AL117351.1
SHISA6
NRXN3
LINC02615
ARHGAP10

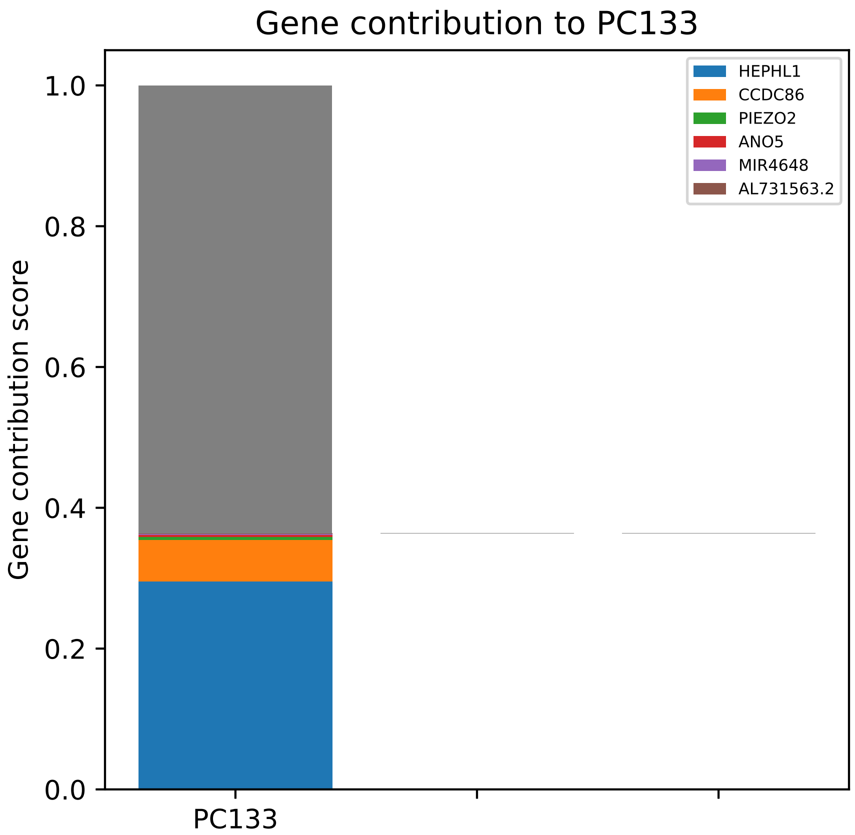
Gene contribution to PC133
Gene contribution score
1.0
0.8
0.6
0.4
0.2
0.0
PC133
HEPHL1
CCDC86
PIEZO2
ANO5
MIR4648
AL731563.2

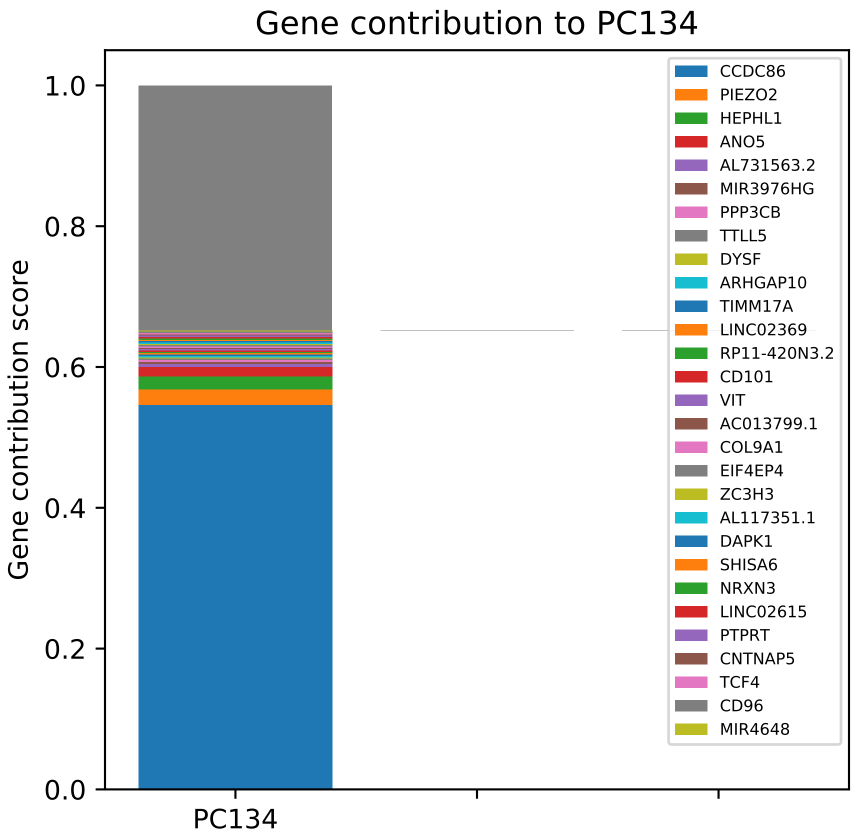

Gene contribution to PC134
Gene contribution score
0.0
0.2
0.4
0.6
0.8
1.0
PC134
CCDC86
PIEZO2
HEPHL1
ANO5
AL731563.2
MIR3976HG
PPP3CB
TTLL5
DYSF
ARHGAP10
TIMM17A
LINC02369
RP11-420N3.2
CD101
VIT
AC013799.1
COL9A1
EIF4EP4
ZC3H3
AL117351.1
DAPK1
SHISA6
NRXN3
LINC02615
PTPRT
CNTNAP5
TCF4
CD96
MIR4648

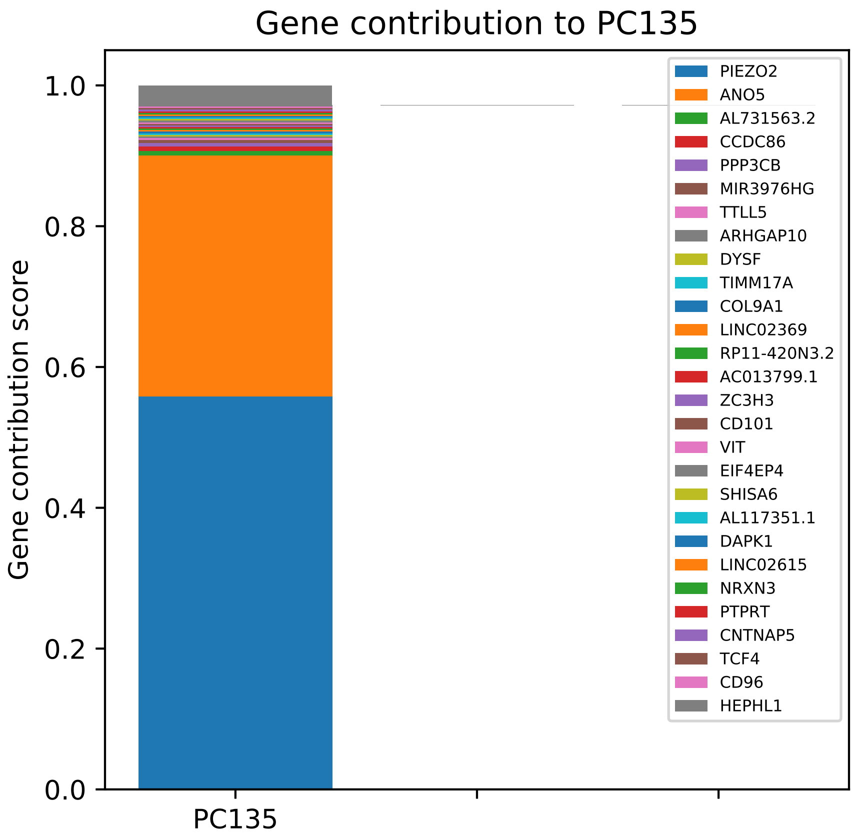

Gene contribution to PC135
Gene contribution score
1.0
0.8
0.6
0.4
0.2
0.0
PC135
PIEZO2
ANO5
AL731563.2
CCDC86
PPP3CB
MIR3976HG
TTLL5
ARHGAP10
DYSF
TIMM17A
COL9A1
LINC02369
RP11-420N3.2
AC013799.1
ZC3H3
CD101
VIT
EIF4EP4
SHISA6
AL117351.1
DAPK1
LINC02615
NRXN3
PTPRT
CNTNAP5
TCF4
CD96
HEPHL1

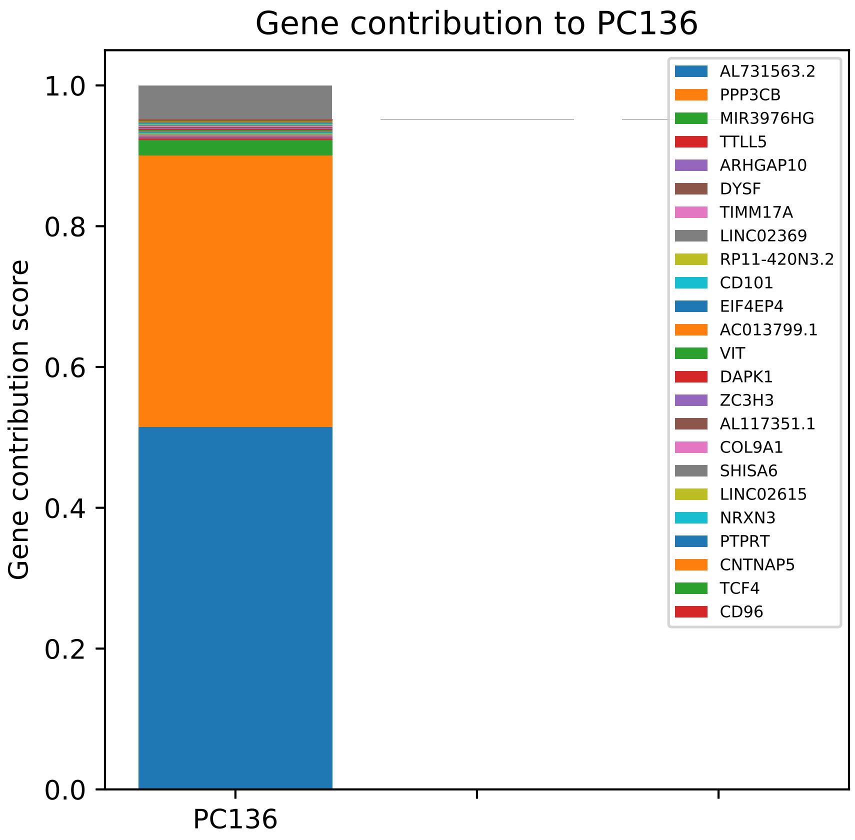
Gene contribution to PC136
Gene contribution score
1.0
0.8
0.6
0.4
0.2
0.0
PC136
AL731563.2
PPP3CB
MIR3976HG
TTLL5
ARHGAP10
DYSF
TIMM17A
LINC02369
RP11-420N3.2
CD101
EIF4EP4
AC013799.1
VIT
DAPK1
ZC3H3
AL117351.1
COL9A1
SHISA6
LINC02615
NRXN3
PTPRT
CNTNAP5
TCF4
CD96

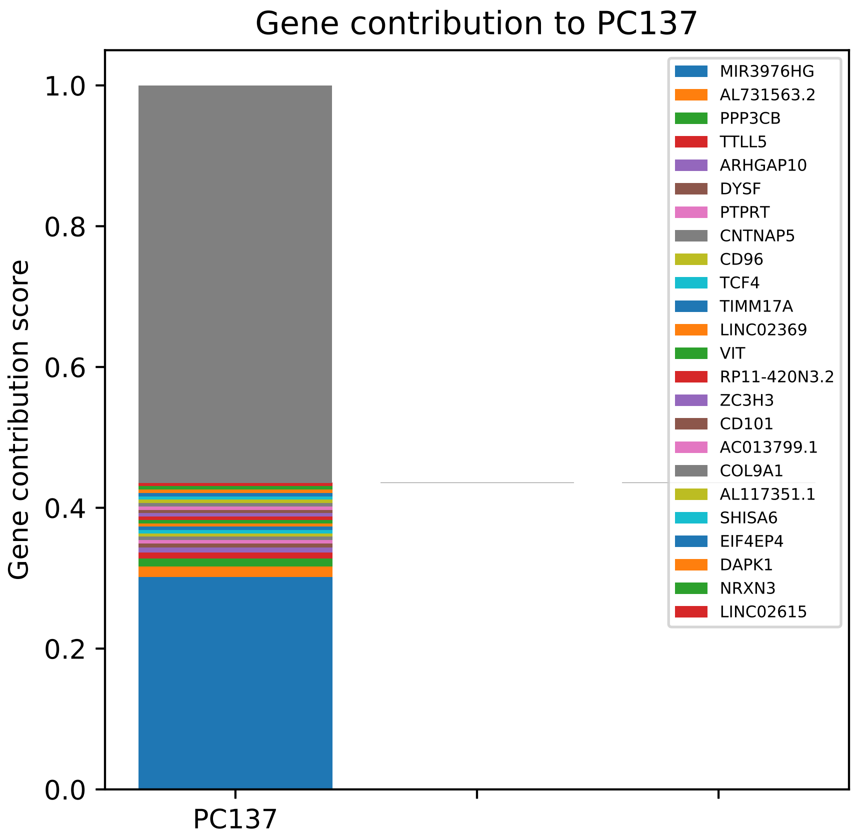

Gene contribution to PC137
Gene contribution score
1.0
0.8
0.6
0.4
0.2
0.0
PC137
MIR3976HG
AL731563.2
PPP3CB
TTLL5
ARHGAP10
DYSF
PTPRT
CNTNAP5
CD96
TCF4
TIMM17A
LINC02369
VIT
RP11-420N3.2
ZC3H3
CD101
AC013799.1
COL9A1
AL117351.1
SHISA6
EIF4EP4
DAPK1
NRXN3
LINC02615

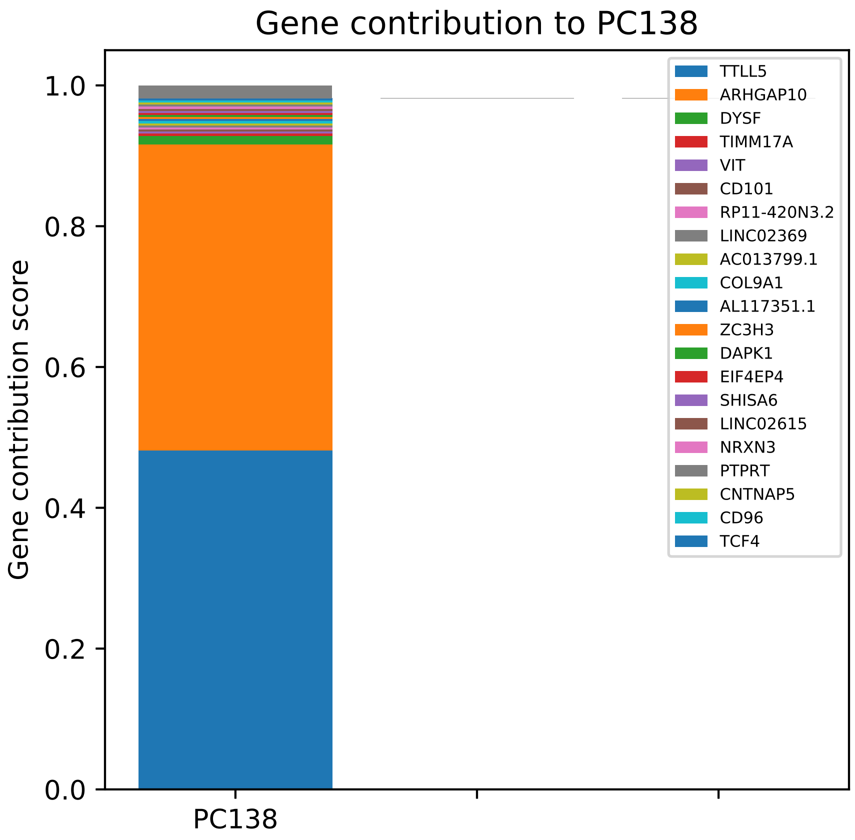
Gene contribution to PC138
Gene contribution score
0.0
0.2
0.4
0.6
0.8
1.0
PC138
TTLL5
ARHGAP10
DYSF
TIMM17A
VIT
CD101
RP11-420N3.2
LINC02369
AC013799.1
COL9A1
AL117351.1
ZC3H3
DAPK1
EIF4EP4
SHISA6
LINC02615
NRXN3
PTPRT
CNTNAP5
CD96
TCF4

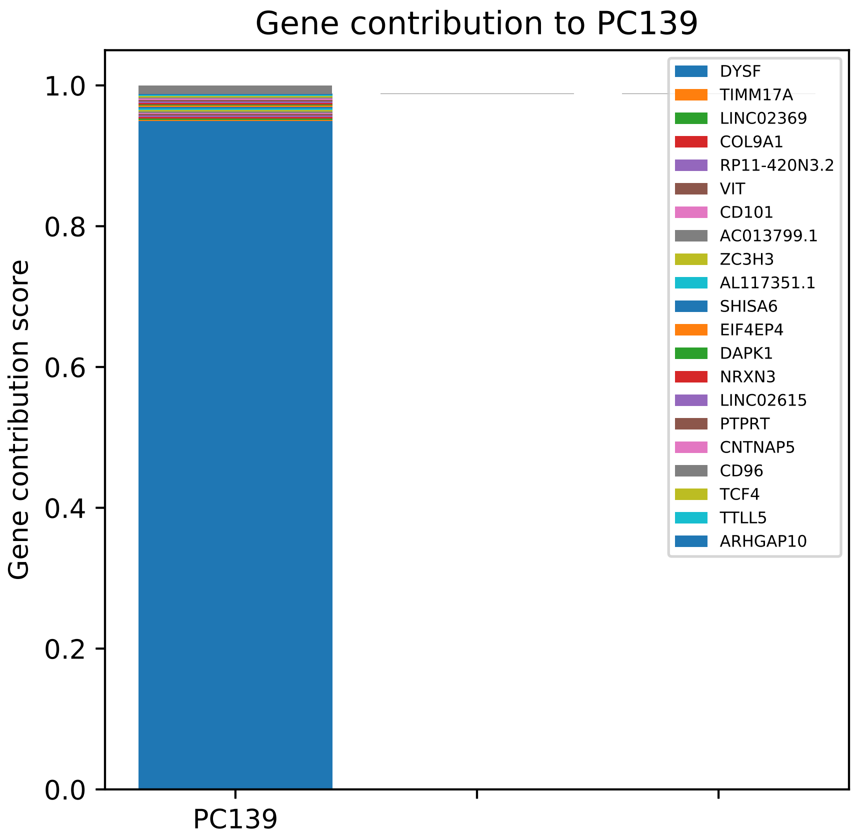
Gene contribution to PC139
Gene contribution score
0.0
0.2
0.4
0.6
0.8
1.0
PC139
DYSF
TIMM17A
LINC02369
COL9A1
RP11-420N3.2
VIT
CD101
AC013799.1
ZC3H3
AL117351.1
SHISA6
EIF4EP4
DAPK1
NRXN3
LINC02615
PTPRT
CNTNAP5
CD96
TCF4
TTLL5
ARHGAP10

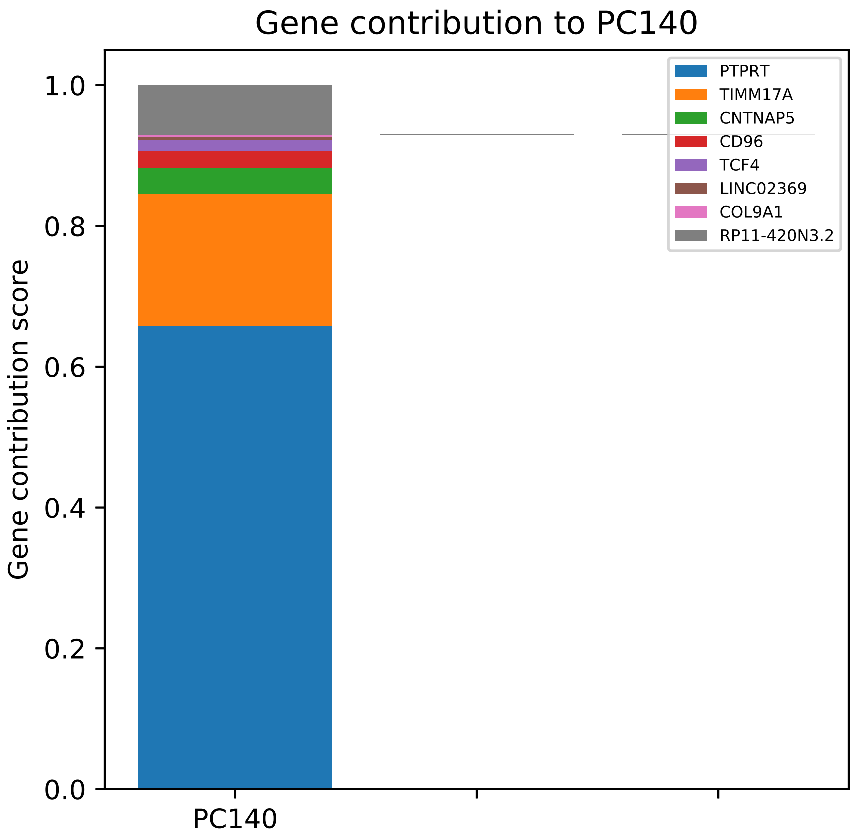

Gene contribution to PC140
Gene contribution score
1.0
0.8
0.6
0.4
0.2
0.0
PC140
PTPRT
TIMM17A
CNTNAP5
CD96
TCF4
LINC02369
COL9A1
RP11-420N3.2

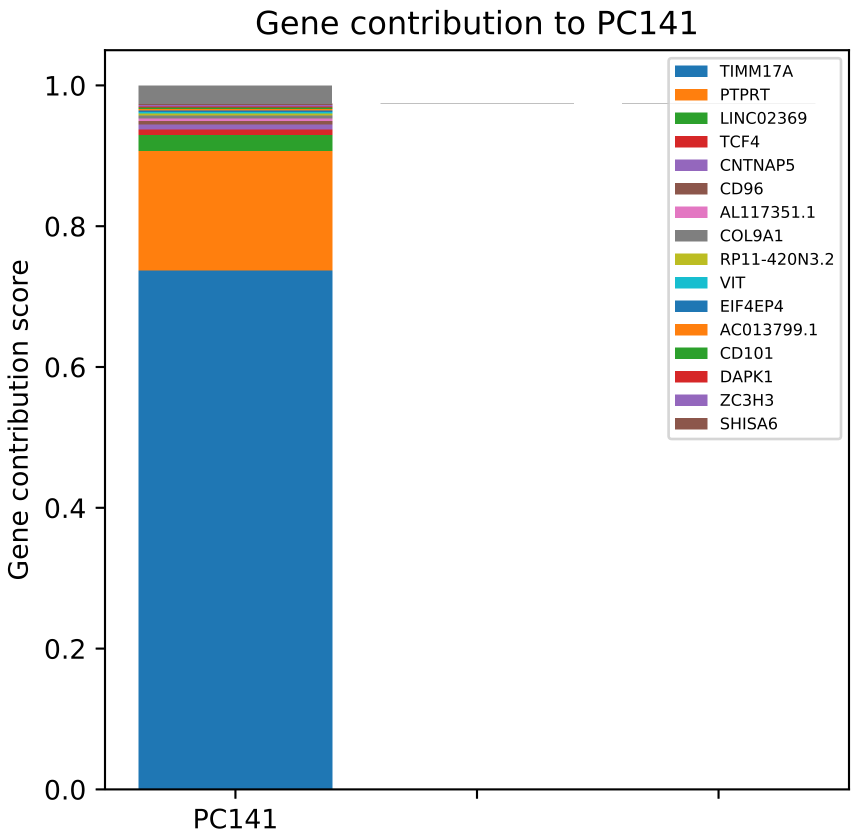

Gene contribution to PC141
Gene contribution score
1.0
0.8
0.6
0.4
0.2
0.0
PC141
TIMM17A
PTPRT
LINC02369
TCF4
CNTNAP5
CD96
AL117351.1
COL9A1
RP11-420N3.2
VIT
EIF4EP4
AC013799.1
CD101
DAPK1
ZC3H3
SHISA6

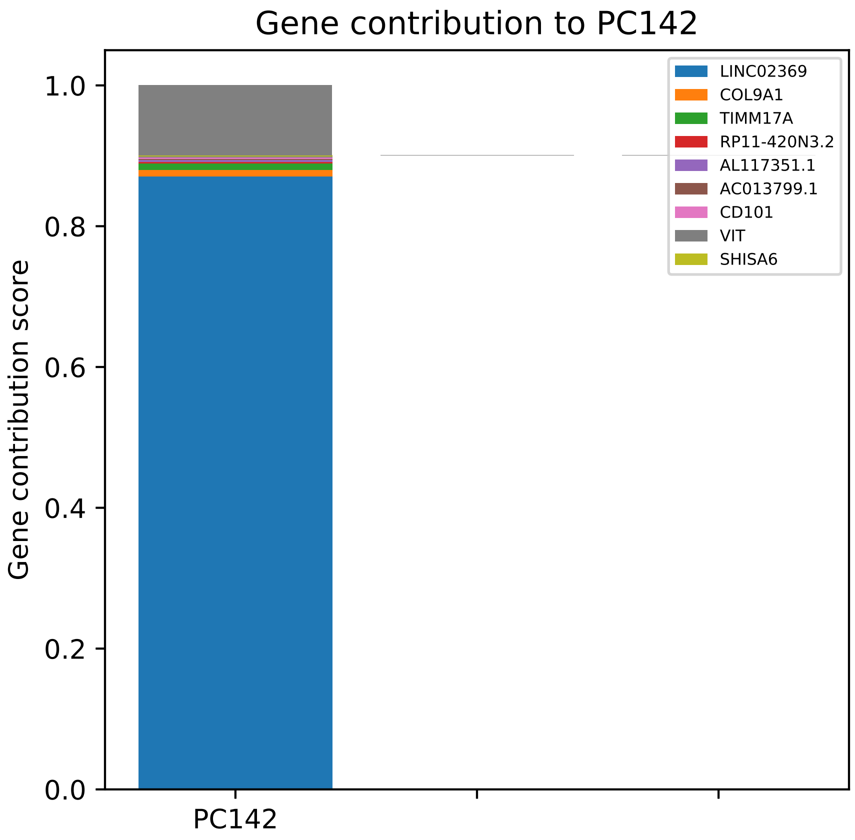
Gene contribution to PC142
Gene contribution score
1.0
0.8
0.6
0.4
0.2
0.0
PC142
LINC02369
COL9A1
TIMM17A
RP11-420N3.2
AL117351.1
AC013799.1
CD101
VIT
SHISA6

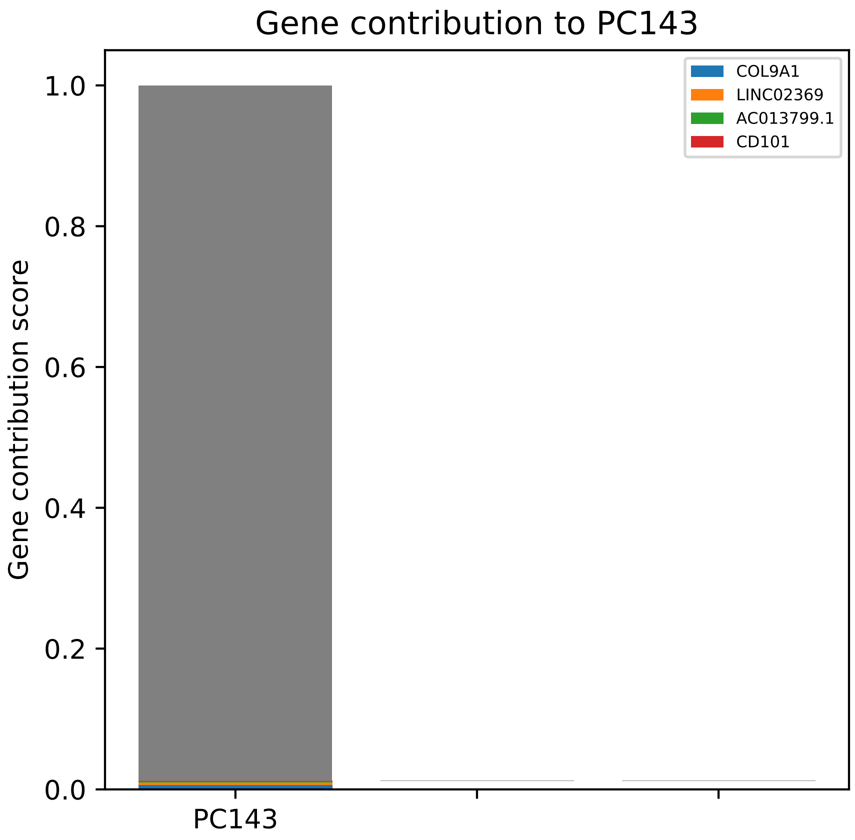

Gene contribution to PC143
Gene contribution score
1.0
0.8
0.6
0.4
0.2
0.0
PC143
COL9A1
LINC02369
AC013799.1
CD101

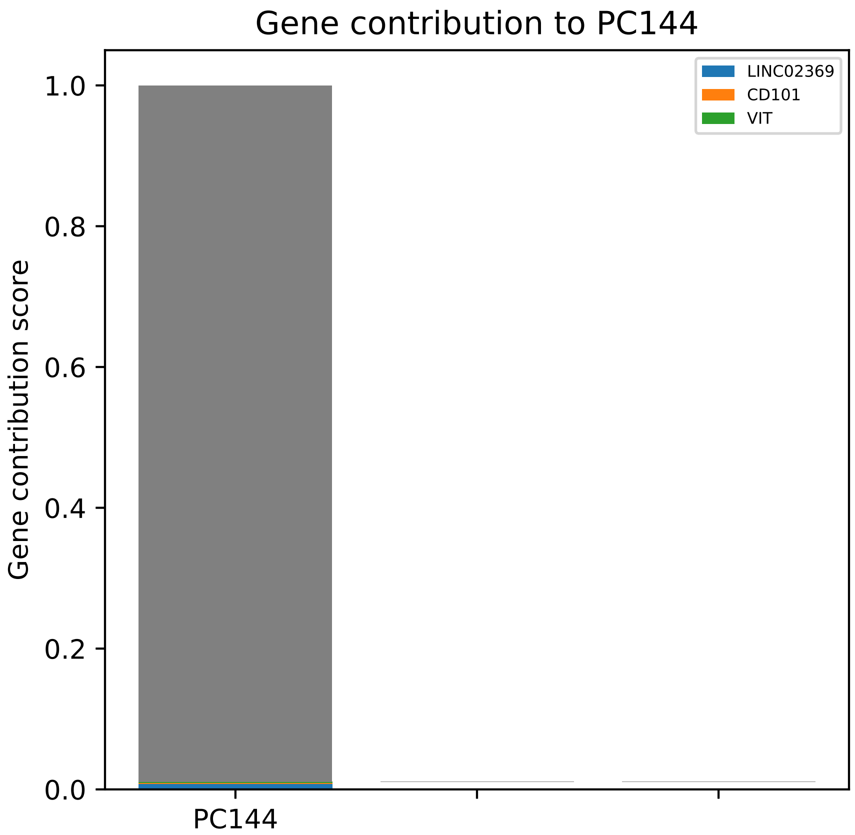

Gene contribution to PC144
LINC02369
CD101
VIT
Gene contribution score
1.0
0.8
0.6
0.4
0.2
0.0
PC144

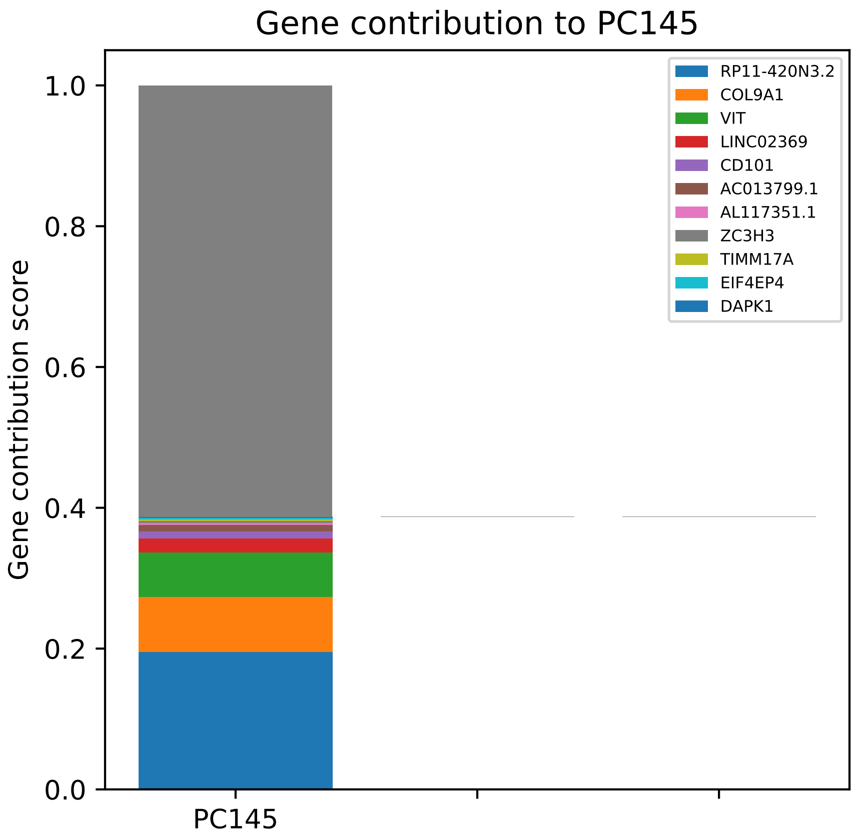
Gene contribution to PC145
Gene contribution score
0.0
0.2
0.4
0.6
0.8
1.0
PC145
RP11-420N3.2
COL9A1
VIT
LINC02369
CD101
AC013799.1
AL117351.1
ZC3H3
TIMM17A
EIF4EP4
DAPK1

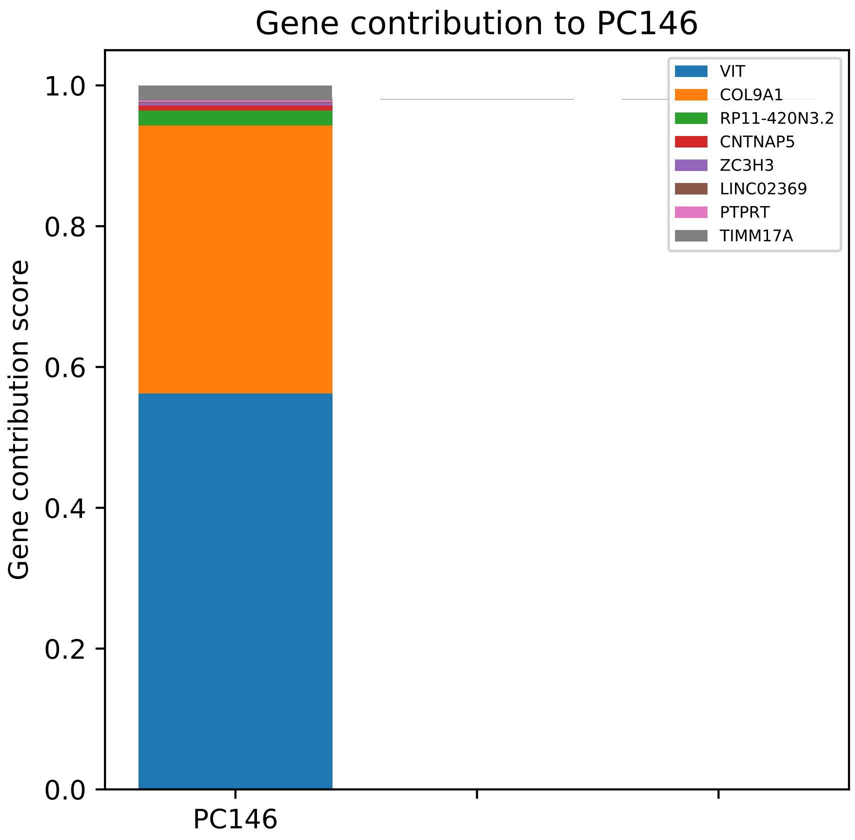

Gene contribution to PC146
Gene contribution score
1.0
0.8
0.6
0.4
0.2
0.0
PC146
VIT
COL9A1
RP11-420N3.2
CNTNAP5
ZC3H3
LINC02369
PTPRT
TIMM17A

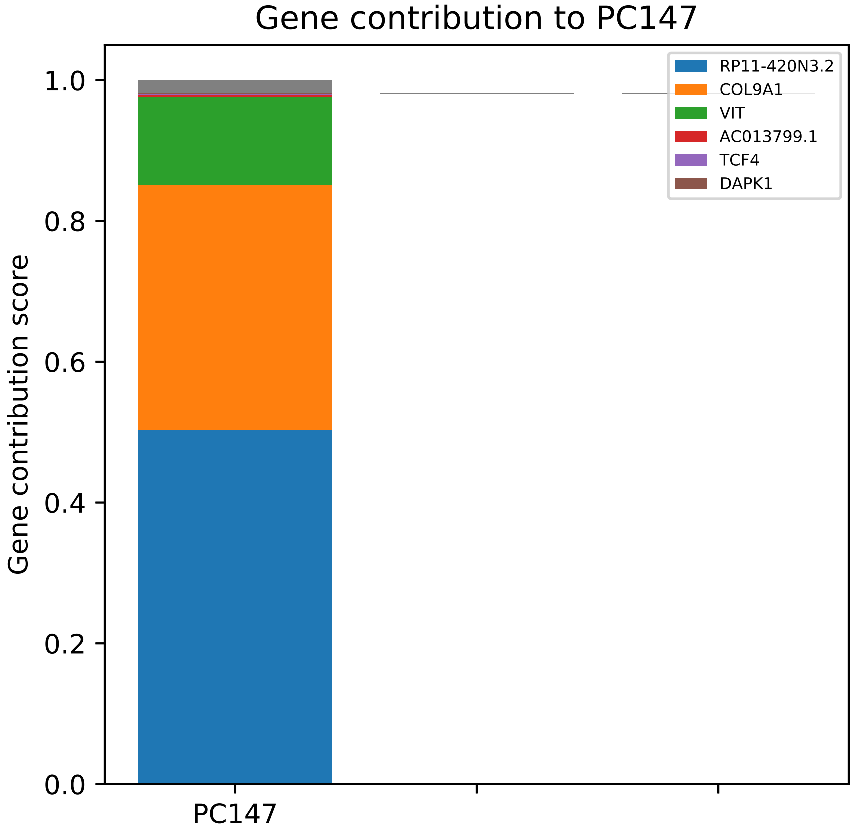

Gene contribution to PC147
Gene contribution score
1.0
0.8
0.6
0.4
0.2
0.0
PC147
RP11-420N3.2
COL9A1
VIT
AC013799.1
TCF4
DAPK1

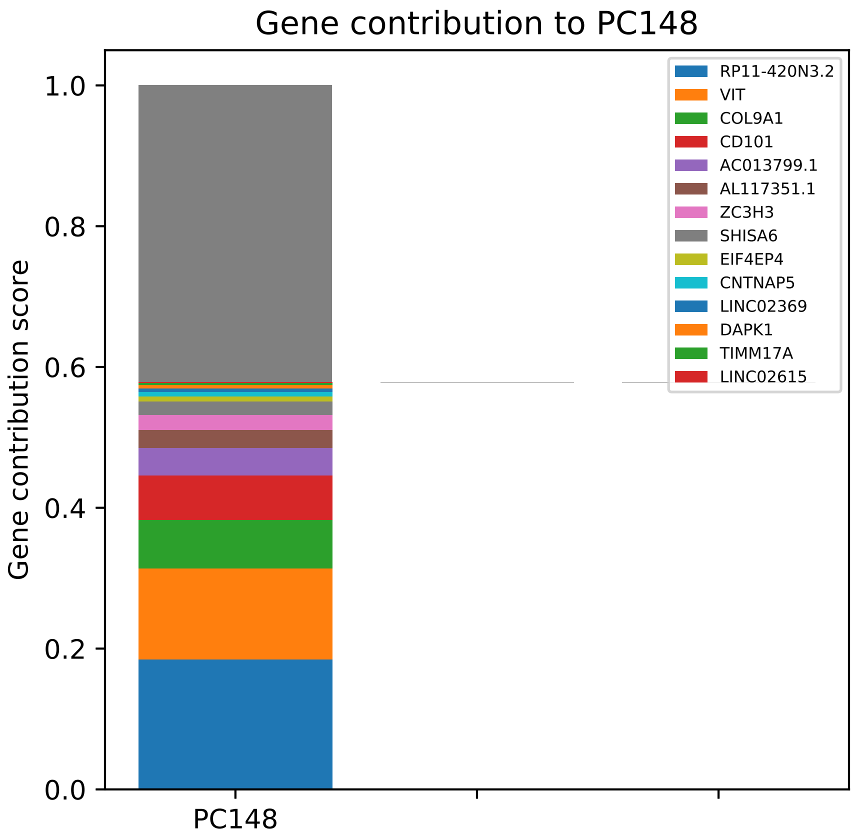

Gene contribution to PC148
Gene contribution score
1.0
0.8
0.6
0.4
0.2
0.0
PC148
RP11-420N3.2
VIT
COL9A1
CD101
AC013799.1
AL117351.1
ZC3H3
SHISA6
EIF4EP4
CNTNAP5
LINC02369
DAPK1
TIMM17A
LINC02615

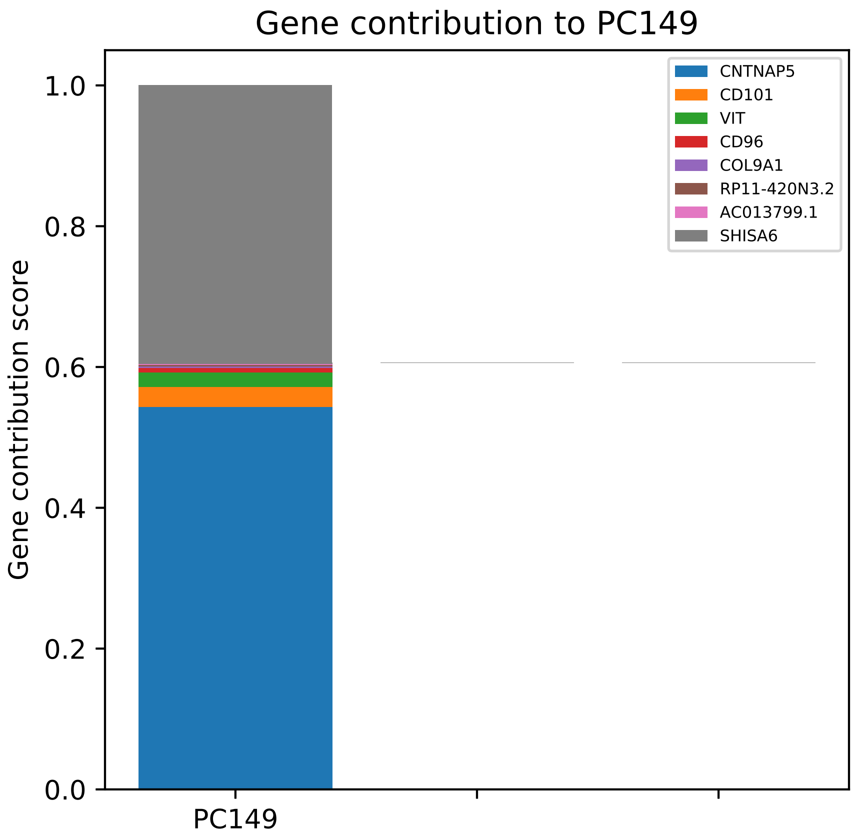

Gene contribution to PC149
Gene contribution score
1.0
0.8
0.6
0.4
0.2
0.0
PC149
CNTNAP5
CD101
VIT
CD96
COL9A1
RP11-420N3.2
AC013799.1
SHISA6

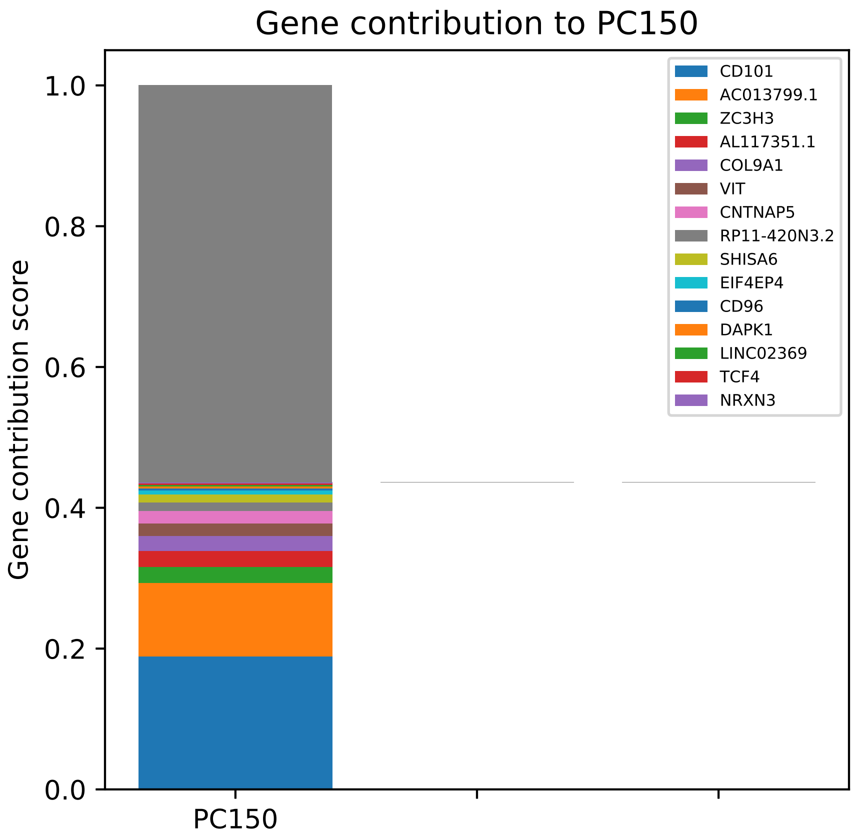

Gene contribution to PC150
Gene contribution score
1.0
0.8
0.6
0.4
0.2
0.0
PC150
CD101
AC013799.1
ZC3H3
AL117351.1
COL9A1
VIT
CNTNAP5
RP11-420N3.2
SHISA6
EIF4EP4
CD96
DAPK1
LINC02369
TCF4
NRXN3

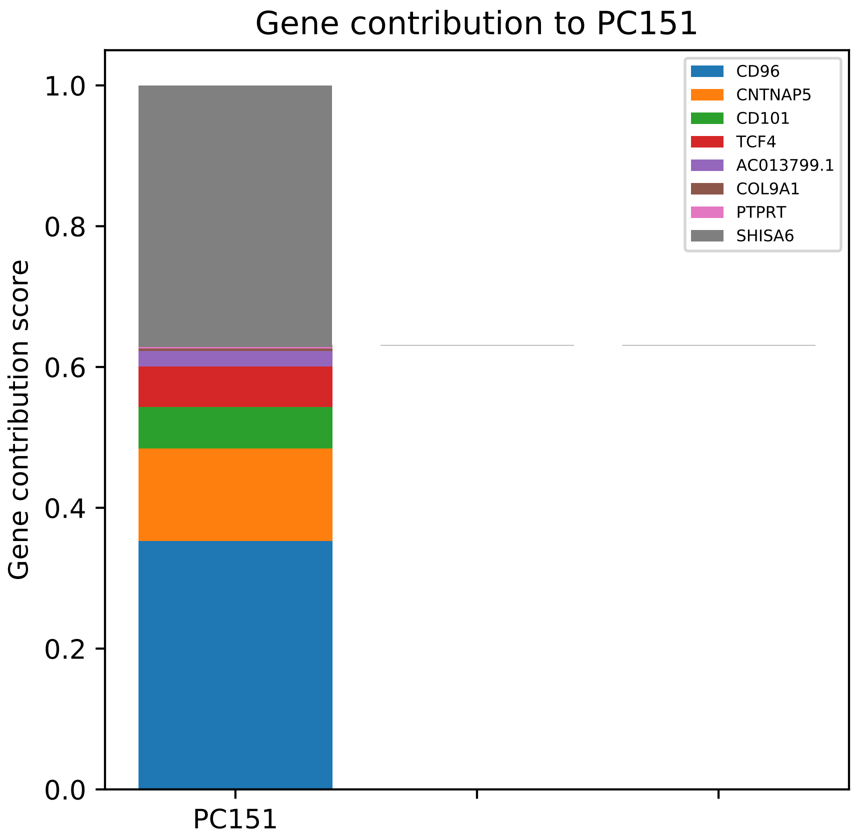

Gene contribution to PC151
Gene contribution score
1.0
0.8
0.6
0.4
0.2
0.0
PC151
CD96
CNTNAP5
CD101
TCF4
AC013799.1
COL9A1
PTPRT
SHISA6

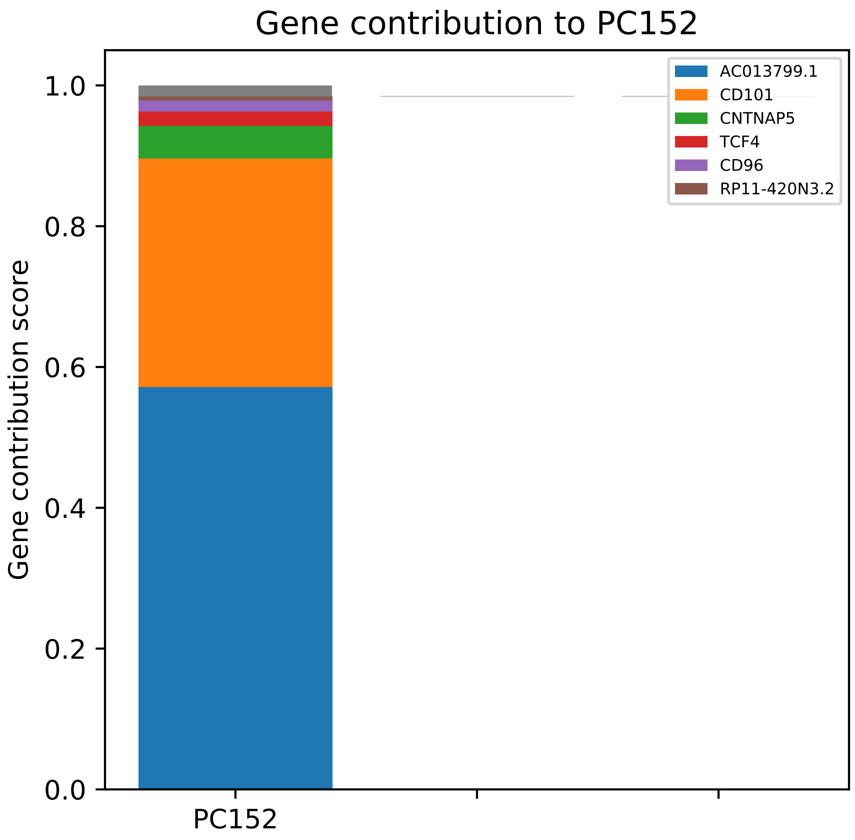

Gene contribution to PC152
Gene contribution score
1.0
0.8
0.6
0.4
0.2
0.0
PC152
AC013799.1
CD101
CNTNAP5
TCF4
CD96
RP11-420N3.2

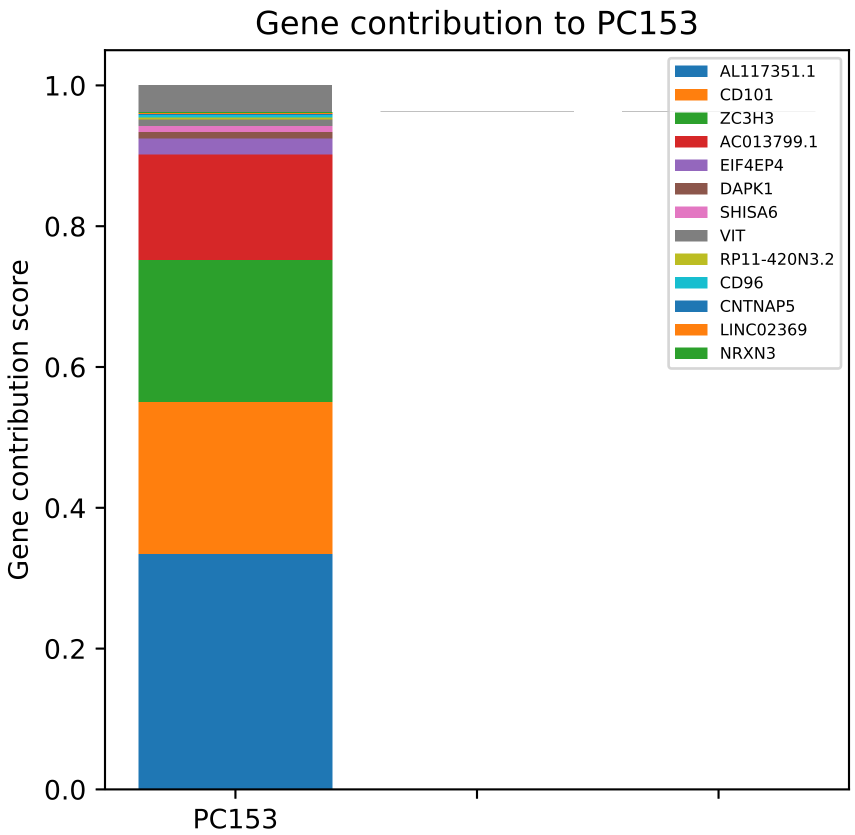

Gene contribution to PC153
Gene contribution score
1.0
0.8
0.6
0.4
0.2
0.0
PC153
AL117351.1
CD101
ZC3H3
AC013799.1
EIF4EP4
DAPK1
SHISA6
VIT
RP11-420N3.2
CD96
CNTNAP5
LINC02369
NRXN3

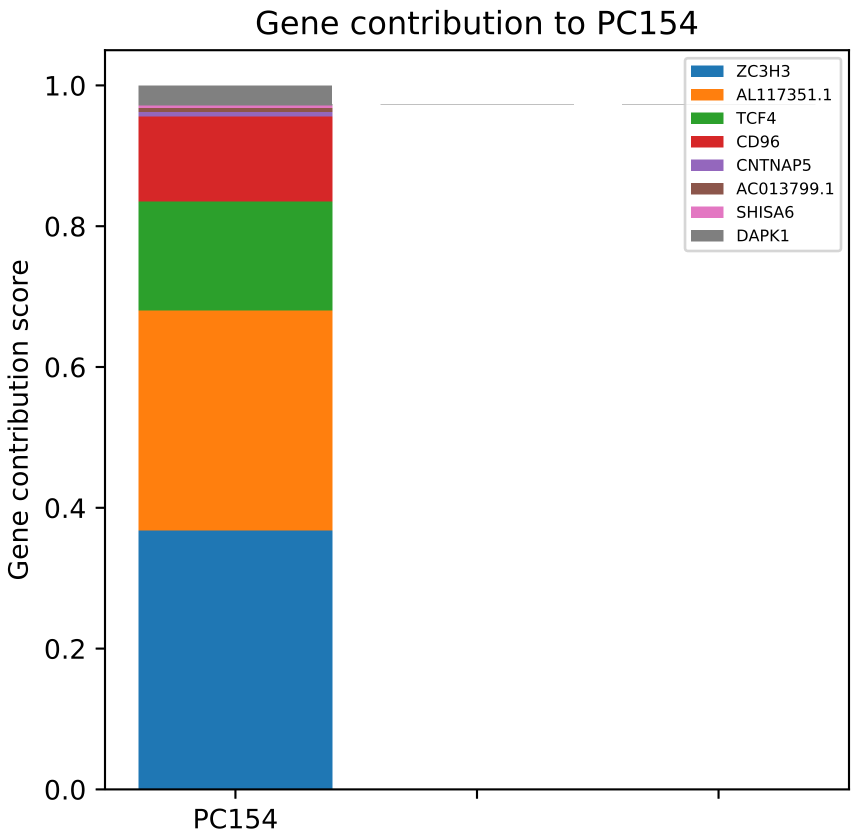

Gene contribution to PC154
Gene contribution score
1.0
0.8
0.6
0.4
0.2
0.0
PC154
ZC3H3
AL117351.1
TCF4
CD96
CNTNAP5
AC013799.1
SHISA6
DAPK1

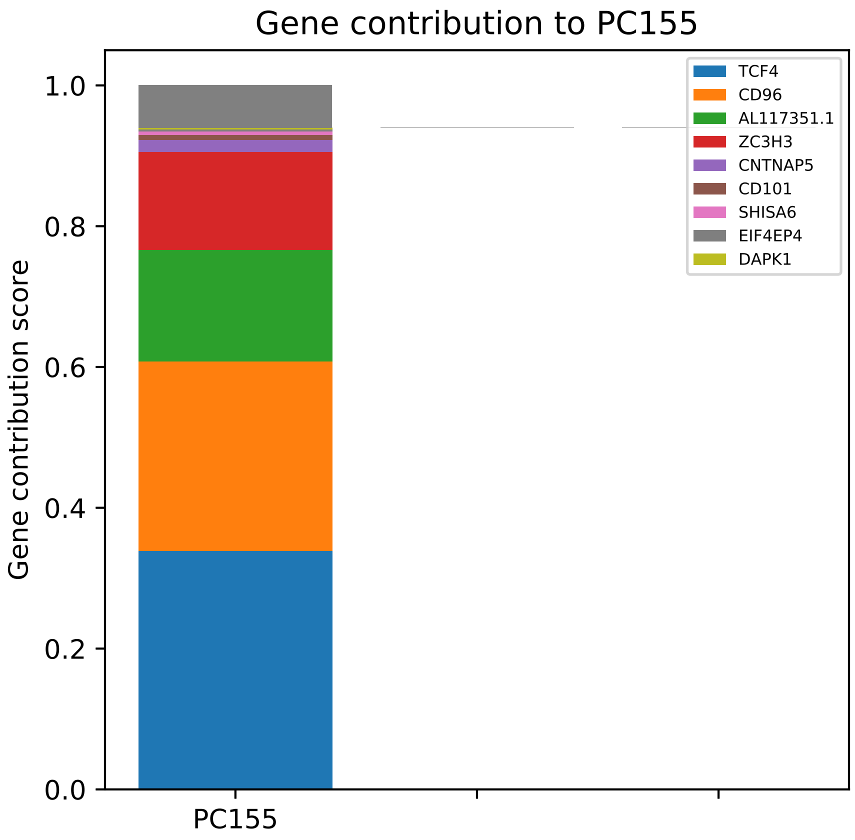
Gene contribution to PC155
Gene contribution score
1.0
0.8
0.6
0.4
0.2
0.0
PC155
TCF4
CD96
AL117351.1
ZC3H3
CNTNAP5
CD101
SHISA6
EIF4EP4
DAPK1

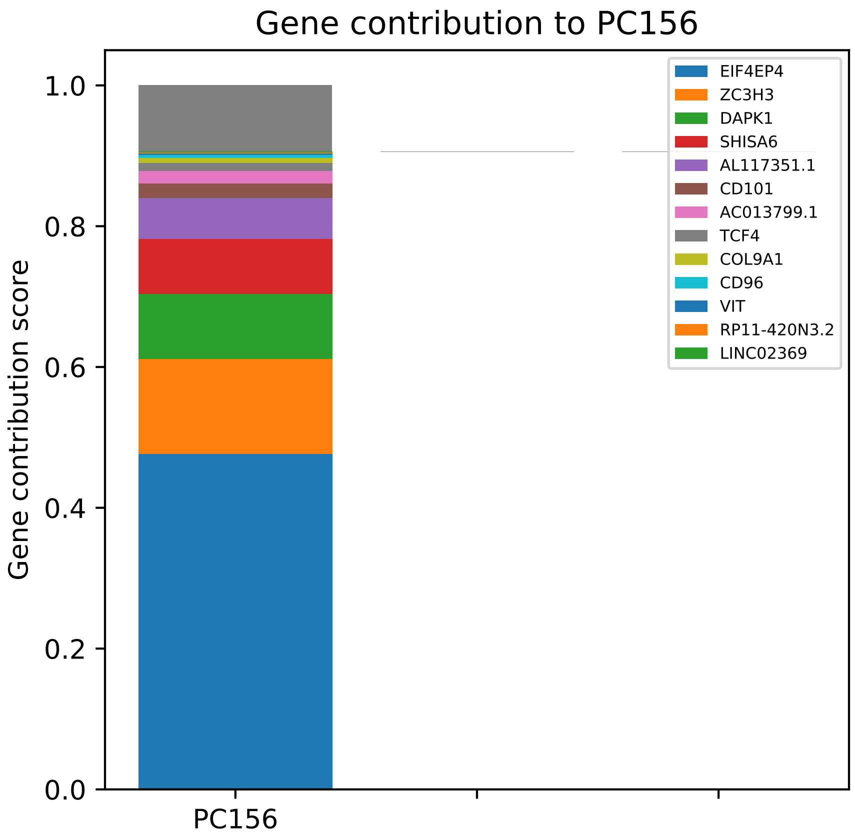

Gene contribution to PC156
Gene contribution score
1.0
0.8
0.6
0.4
0.2
0.0
PC156
EIF4EP4
ZC3H3
DAPK1
SHISA6
AL117351.1
CD101
AC013799.1
TCF4
COL9A1
CD96
VIT
RP11-420N3.2
LINC02369

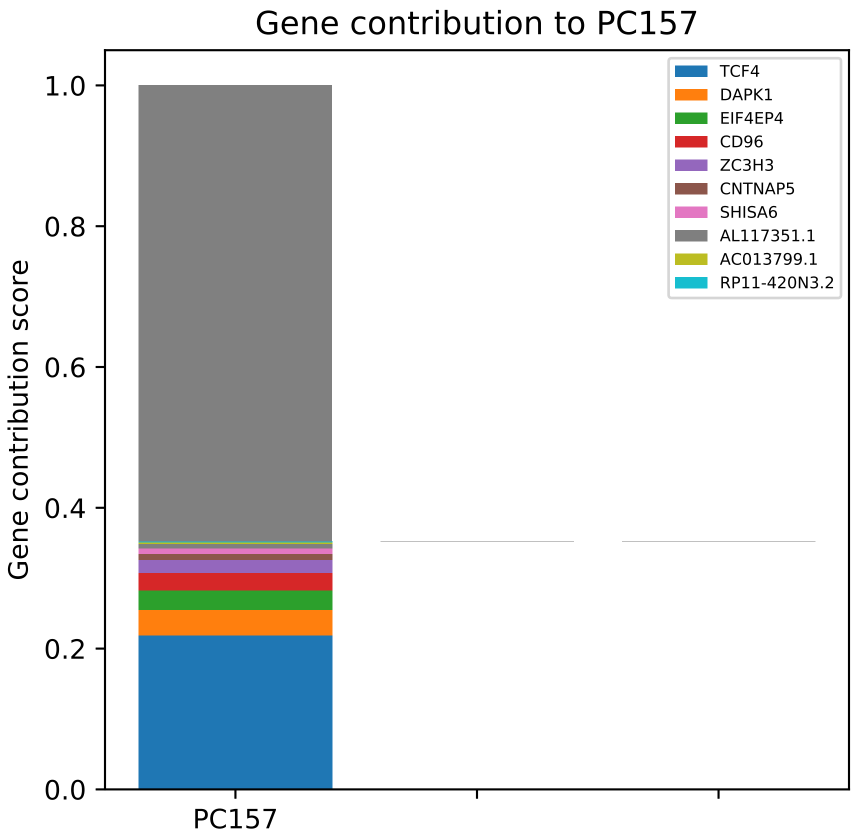

Gene contribution to PC157
Gene contribution score
1.0
0.8
0.6
0.4
0.2
0.0
PC157
TCF4
DAPK1
EIF4EP4
CD96
ZC3H3
CNTNAP5
SHISA6
AL117351.1
AC013799.1
RP11-420N3.2

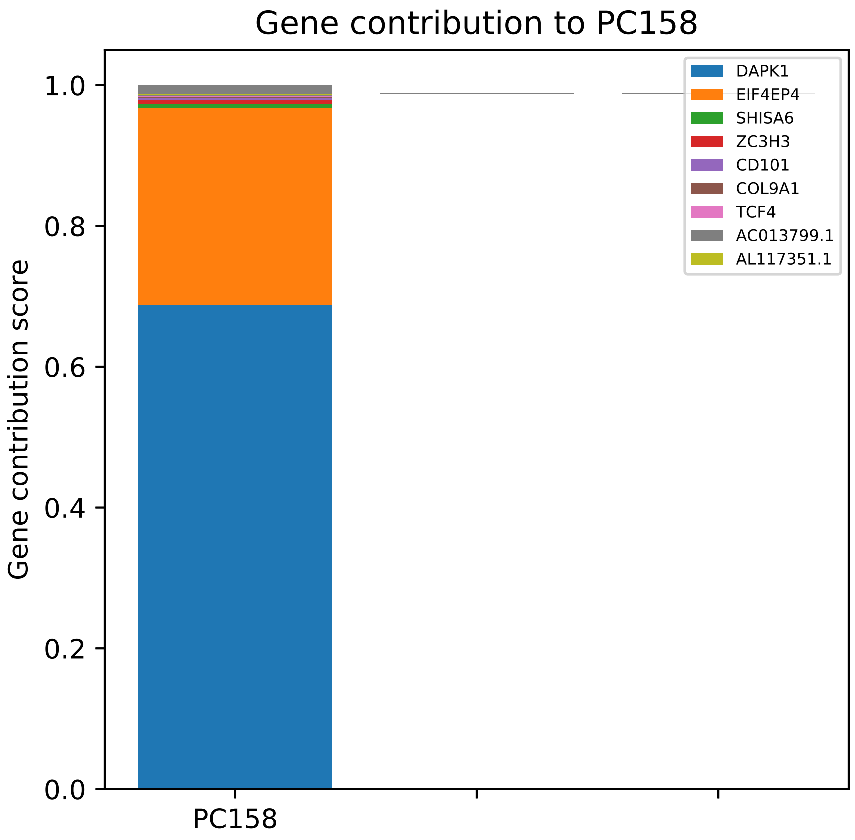

Gene contribution to PC158
Gene contribution score
0.0
0.2
0.4
0.6
0.8
1.0
PC158
DAPK1
EIF4EP4
SHISA6
ZC3H3
CD101
COL9A1
TCF4
AC013799.1
AL117351.1

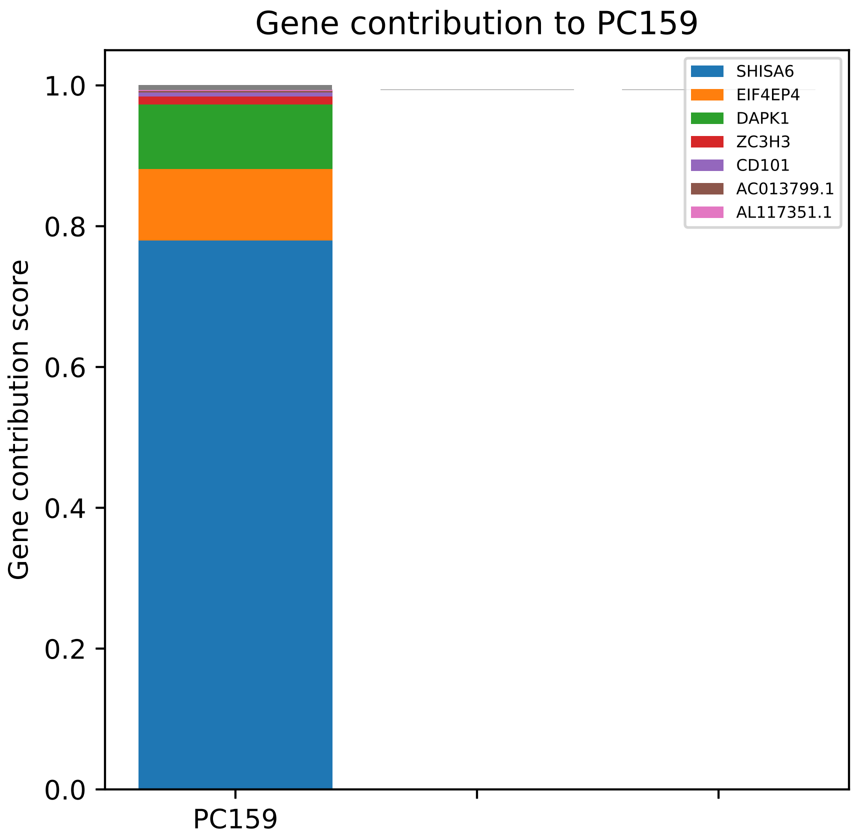

Gene contribution to PC159
Gene contribution score
1.0
0.8
0.6
0.4
0.2
0.0
PC159
SHISA6
EIF4EP4
DAPK1
ZC3H3
CD101
AC013799.1
AL117351.1

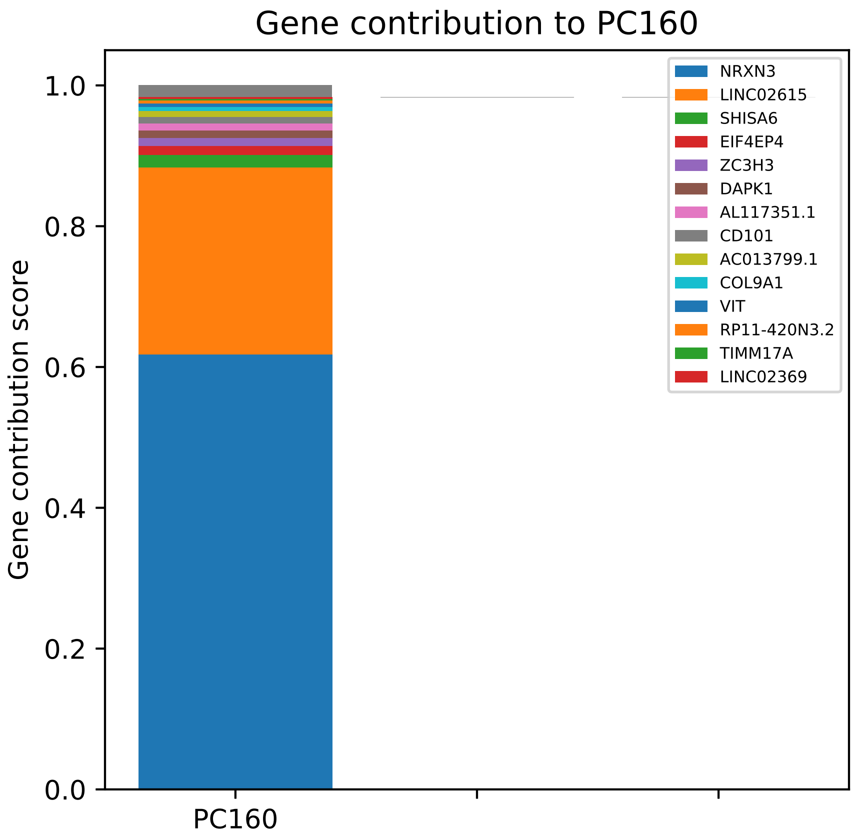
Gene contribution to PC160
Gene contribution score
1.0
0.8
0.6
0.4
0.2
0.0
PC160
NRXN3
LINC02615
SHISA6
EIF4EP4
ZC3H3
DAPK1
AL117351.1
CD101
AC013799.1
COL9A1
VIT
RP11-420N3.2
TIMM17A
LINC02369

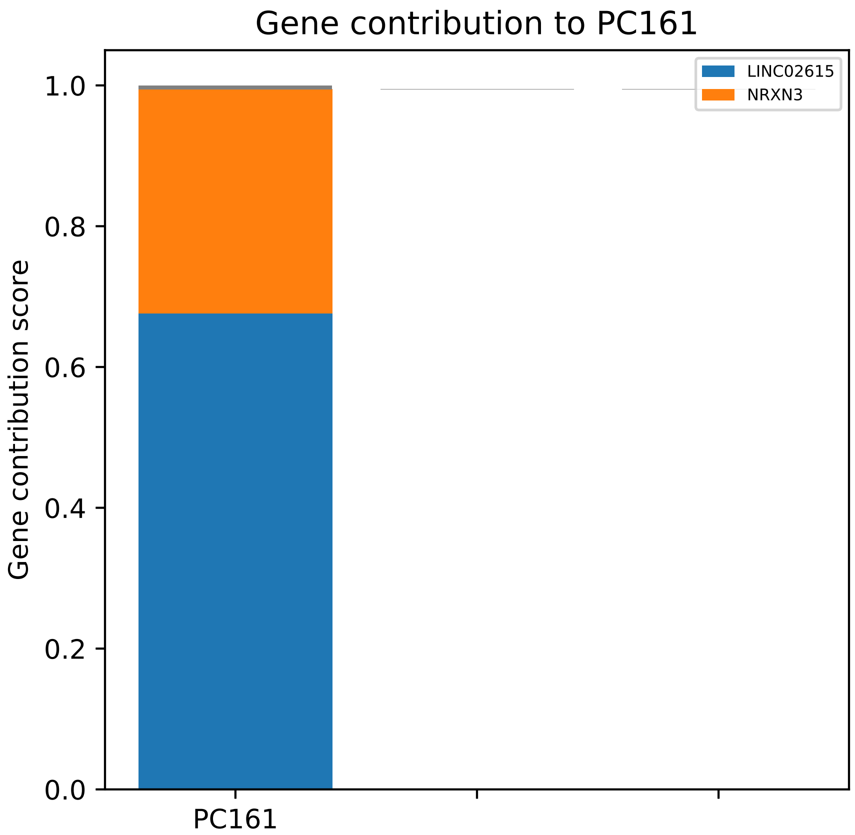
Gene contribution to PC161
LINC02615
NRXN3
Gene contribution score
1.0
0.8
0.6
0.4
0.2
0.0
PC161

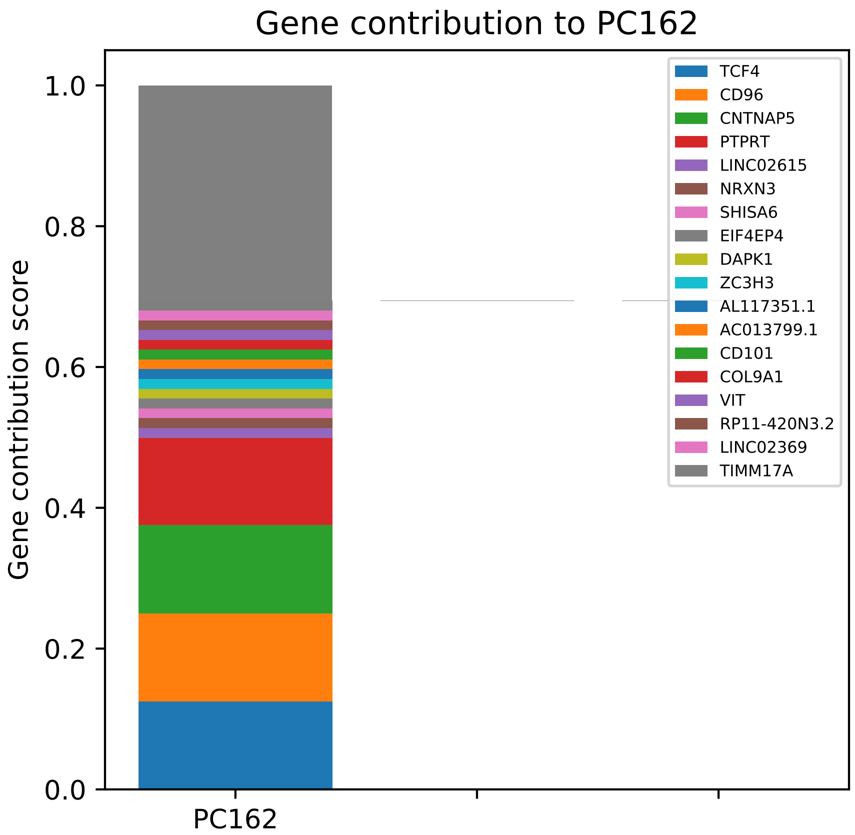

Gene contribution to PC162
Gene contribution score
1.0
0.8
0.6
0.4
0.2
0.0
PC162
TCF4
CD96
CNTNAP5
PTPRT
LINC02615
NRXN3
SHISA6
EIF4EP4
DAPK1
ZC3H3
AL117351.1
AC013799.1
CD101
COL9A1
VIT
RP11-420N3.2
LINC02369
TIMM17A

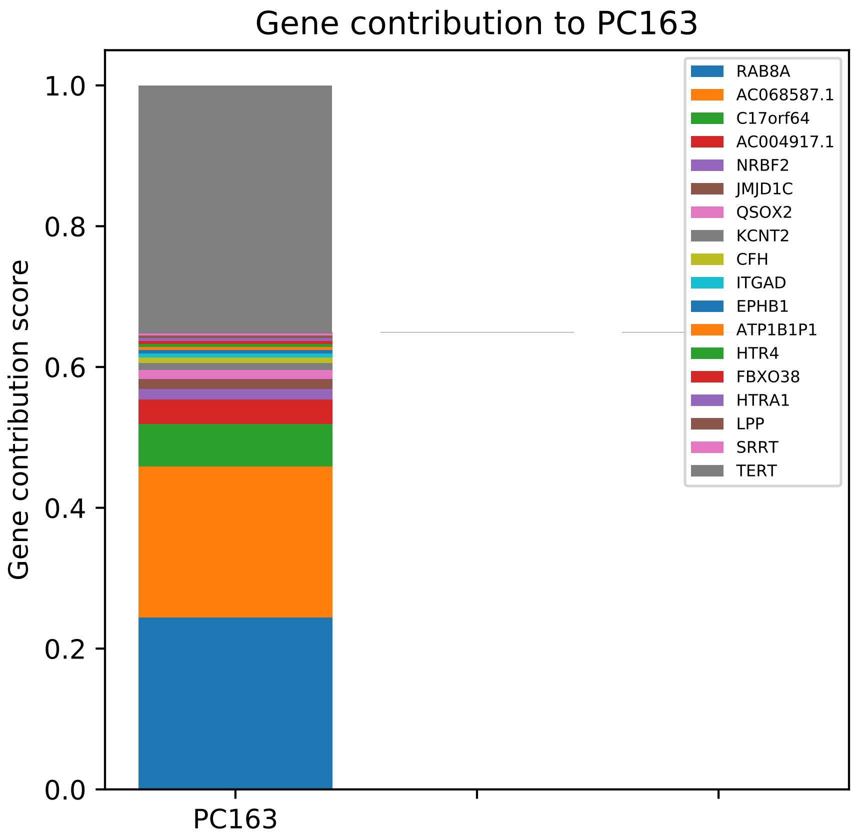

Gene contribution to PC163
Gene contribution score
0.0
0.2
0.4
0.6
0.8
1.0
PC163
RAB8A
AC068587.1
C17orf64
AC004917.1
NRBF2
JMJD1C
QSOX2
KCNT2
CFH
ITGAD
EPHB1
ATP1B1P1
HTR4
FBXO38
HTRA1
LPP
SRRT
TERT

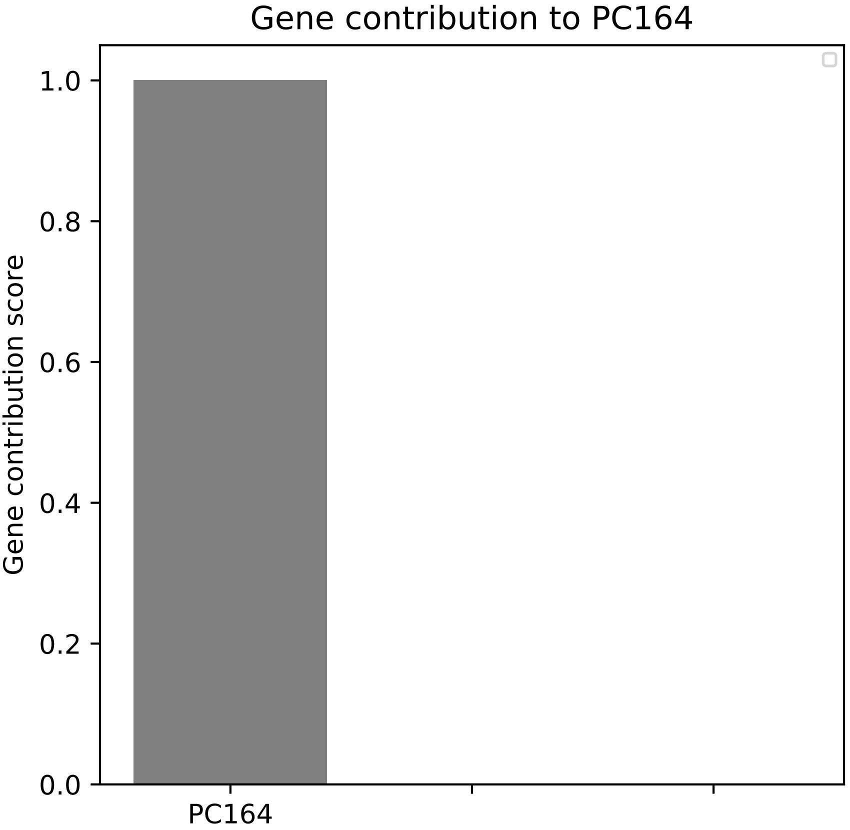
Gene contribution to PC164
Gene contribution score
1.0
0.8
0.6
0.4
0.2
0.0
PC164

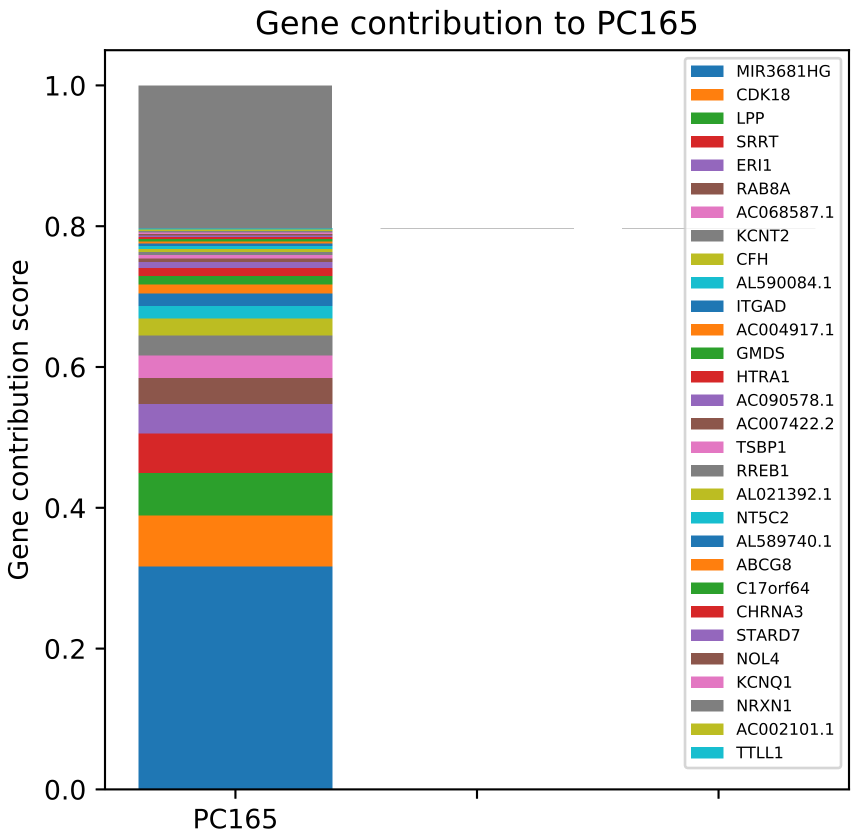
Gene contribution to PC165
Gene contribution score
1.0
0.8
0.6
0.4
0.2
0.0
PC165
MIR3681HG
CDK18
LPP
SRRT
ERI1
RAB8A
AC068587.1
KCNT2
CFH
AL590084.1
ITGAD
AC004917.1
GMDS
HTRA1
AC090578.1
AC007422.2
TSBP1
RREB1
AL021392.1
NT5C2
AL589740.1
ABCG8
C17orf64
CHRNA3
STARD7
NOL4
KCNQ1
NRXN1
AC002101.1
TTLL1

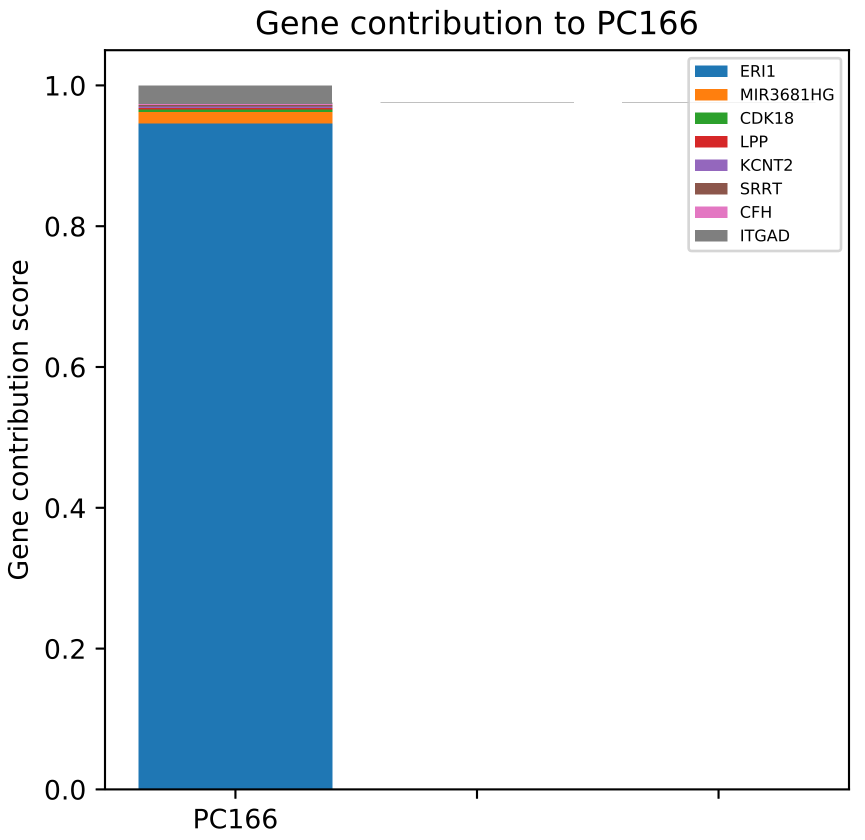
Gene contribution to PC166
Gene contribution score
0.0
0.2
0.4
0.6
0.8
1.0
PC166
ERI1
MIR3681HG
CDK18
LPP
KCNT2
SRRT
CFH
ITGAD

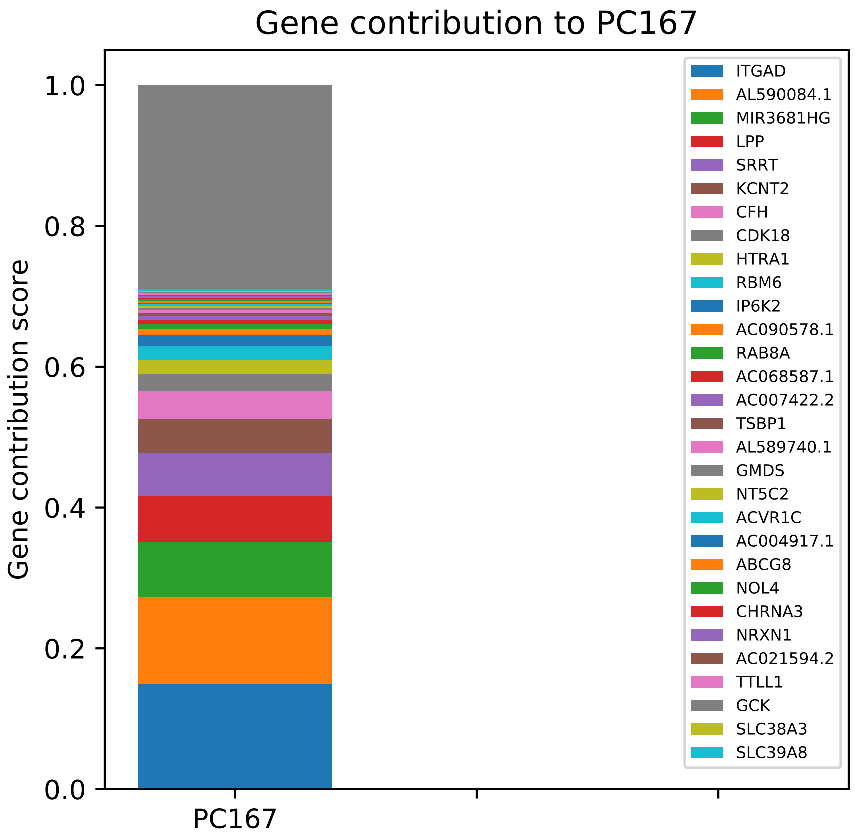

Gene contribution to PC167
Gene contribution score
1.0
0.8
0.6
0.4
0.2
0.0
PC167
ITGAD
AL590084.1
MIR3681HG
LPP
SRRT
KCNT2
CFH
CDK18
HTRA1
RBM6
IP6K2
AC090578.1
RAB8A
AC068587.1
AC007422.2
TSBP1
AL589740.1
GMDS
NT5C2
ACVR1C
AC004917.1
ABCG8
NOL4
CHRNA3
NRXN1
AC021594.2
TTLL1
GCK
SLC38A3
SLC39A8

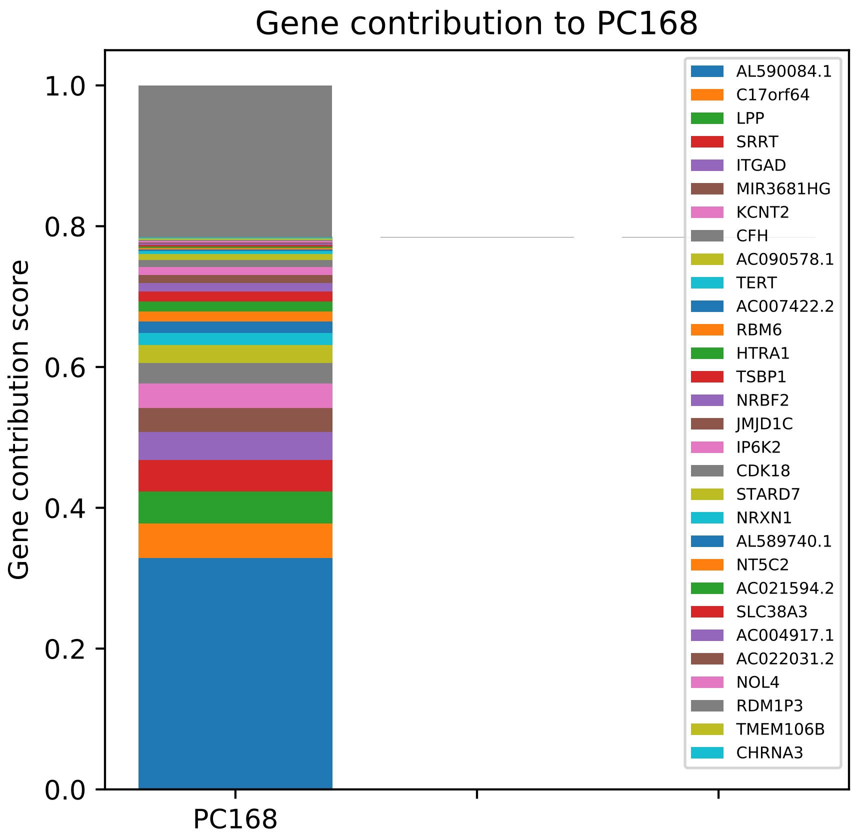

Gene contribution to PC168
Gene contribution score
1.0
0.8
0.6
0.4
0.2
0.0
PC168
AL590084.1
C17orf64
LPP
SRRT
ITGAD
MIR3681HG
KCNT2
CFH
AC090578.1
TERT
AC007422.2
RBM6
HTRA1
TSBP1
NRBF2
JMJD1C
IP6K2
CDK18
STARD7
NRXN1
AL589740.1
NT5C2
AC021594.2
SLC38A3
AC004917.1
AC022031.2
NOL4
RDM1P3
TMEM106B
CHRNA3

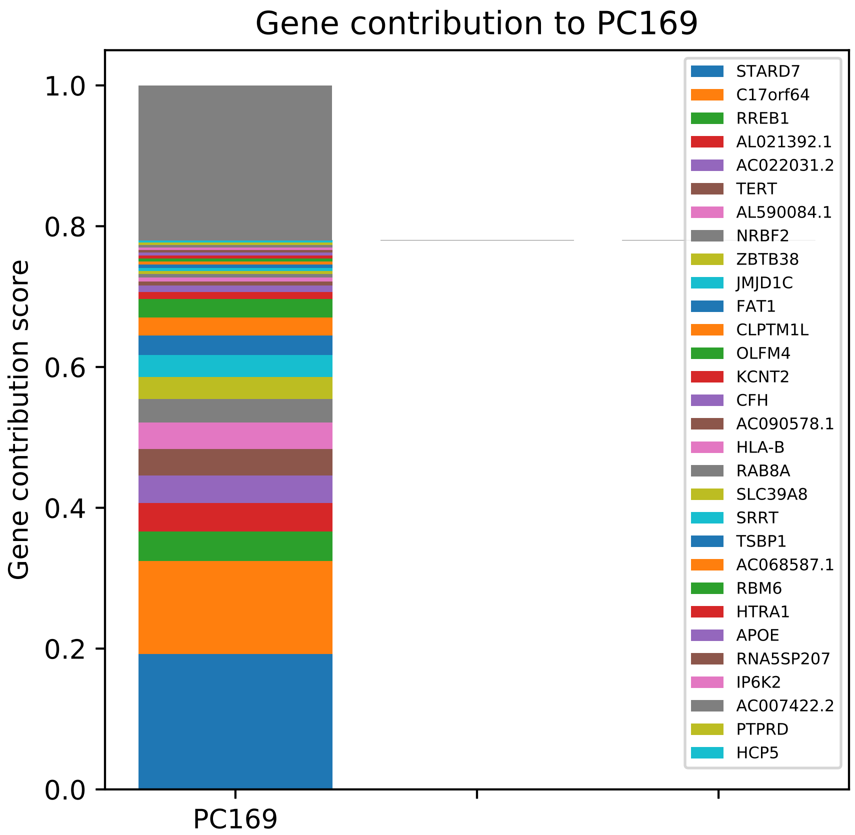
Gene contribution to PC169
Gene contribution score
0.0
0.2
0.4
0.6
0.8
1.0
PC169
STARD7
C17orf64
RREB1
AL021392.1
AC022031.2
TERT
AL590084.1
NRBF2
ZBTB38
JMJD1C
FAT1
CLPTM1L
OLFM4
KCNT2
CFH
AC090578.1
HLA-B
RAB8A
SLC39A8
SRRT
TSBP1
AC068587.1
RBM6
HTRA1
APOE
RNA5SP207
IP6K2
AC007422.2
PTPRD
HCP5

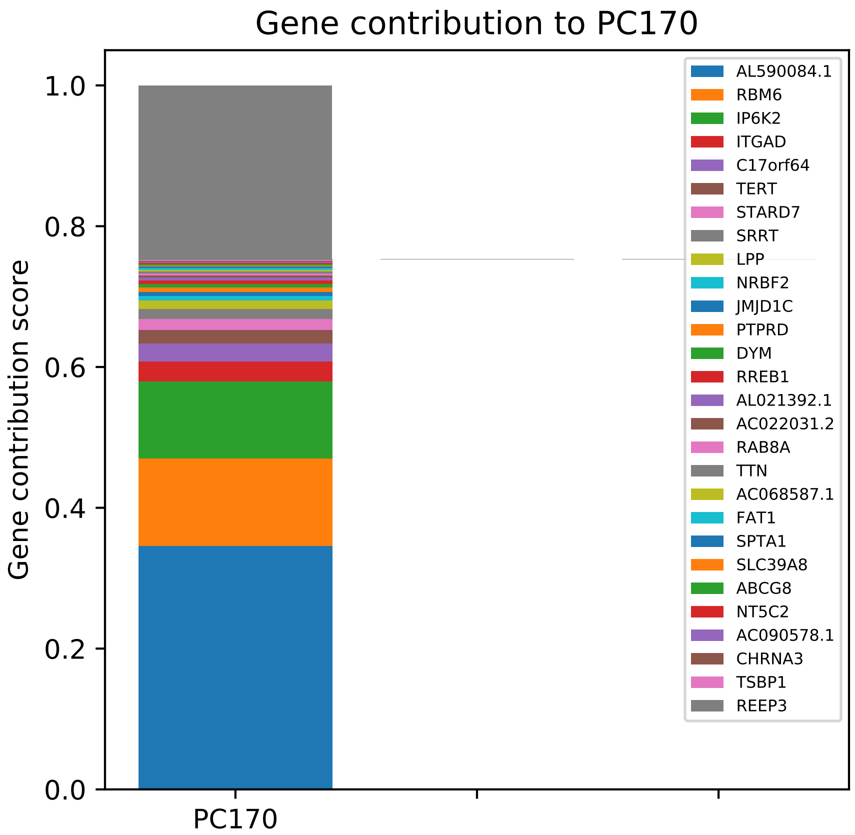
Gene contribution to PC170
Gene contribution score
1.0
0.8
0.6
0.4
0.2
0.0
PC170
AL590084.1
RBM6
IP6K2
ITGAD
C17orf64
TERT
STARD7
SRRT
LPP
NRBF2
JMJD1C
PTPRD
DYM
RREB1
AL021392.1
AC022031.2
RAB8A
TTN
AC068587.1
FAT1
SPTA1
SLC39A8
ABCG8
NT5C2
AC090578.1
CHRNA3
TSBP1
REEP3

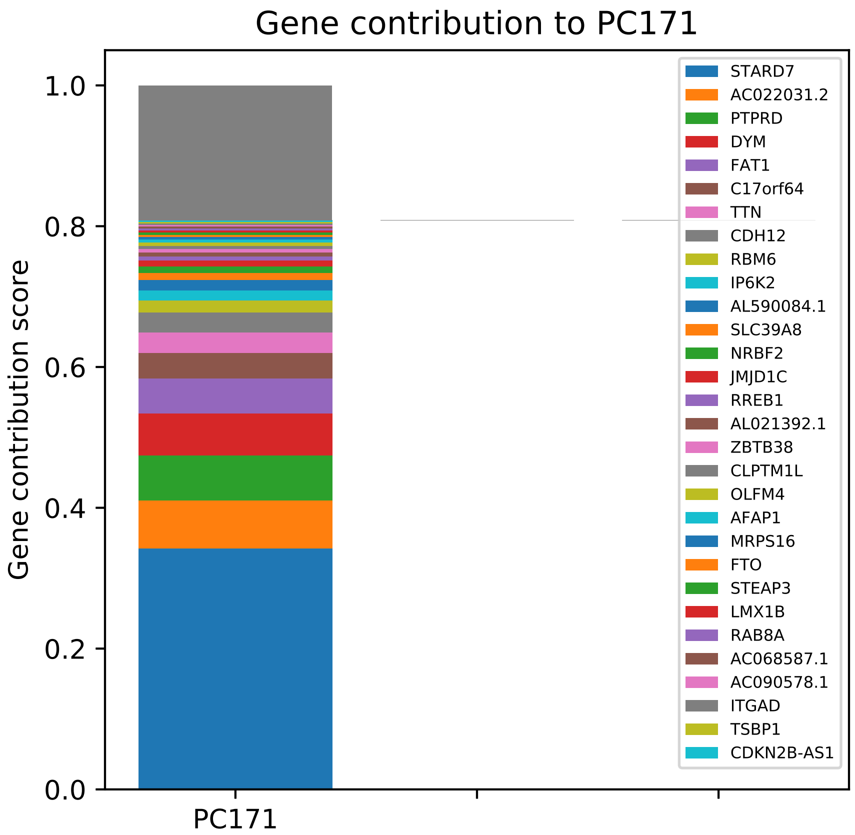

Gene contribution to PC171
Gene contribution score
0.0
0.2
0.4
0.6
0.8
1.0
PC171
STARD7
AC022031.2
PTPRD
DYM
FAT1
C17orf64
TTN
CDH12
RBM6
IP6K2
AL590084.1
SLC39A8
NRBF2
JMJD1C
RREB1
AL021392.1
ZBTB38
CLPTM1L
OLFM4
AFAP1
MRPS16
FTO
STEAP3
LMX1B
RAB8A
AC068587.1
AC090578.1
ITGAD
TSBP1
CDKN2B-AS1

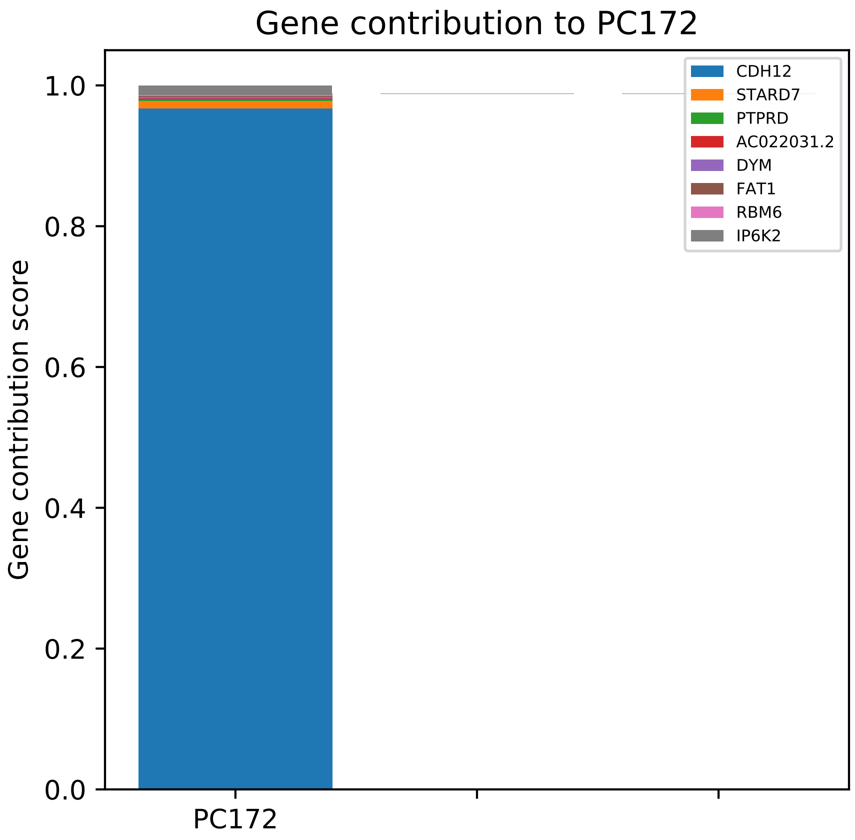

Gene contribution to PC172
Gene contribution score
1.0
0.8
0.6
0.4
0.2
0.0
PC172
CDH12
STARD7
PTPRD
AC022031.2
DYM
FAT1
RBM6
IP6K2

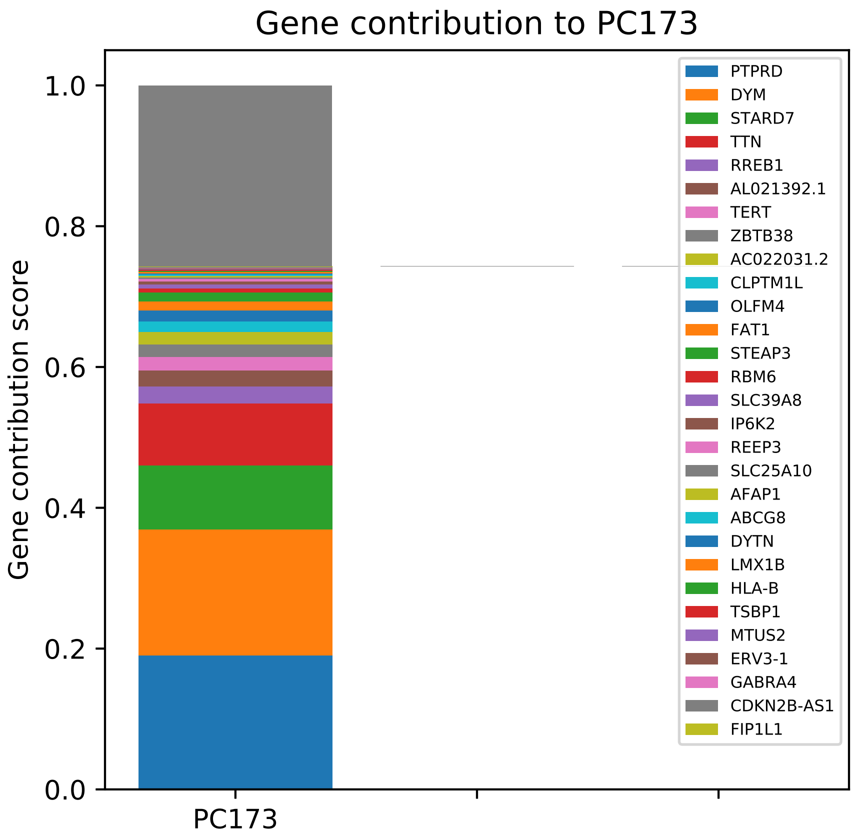
Gene contribution to PC173
Gene contribution score
1.0
0.8
0.6
0.4
0.2
0.0
PC173
PTPRD
DYM
STARD7
TTN
RREB1
AL021392.1
TERT
ZBTB38
AC022031.2
CLPTM1L
OLFM4
FAT1
STEAP3
RBM6
SLC39A8
IP6K2
REEP3
SLC25A10
AFAP1
ABCG8
DYTN
LMX1B
HLA-B
TSBP1
MTUS2
ERV3-1
GABRA4
CDKN2B-AS1
FIP1L1

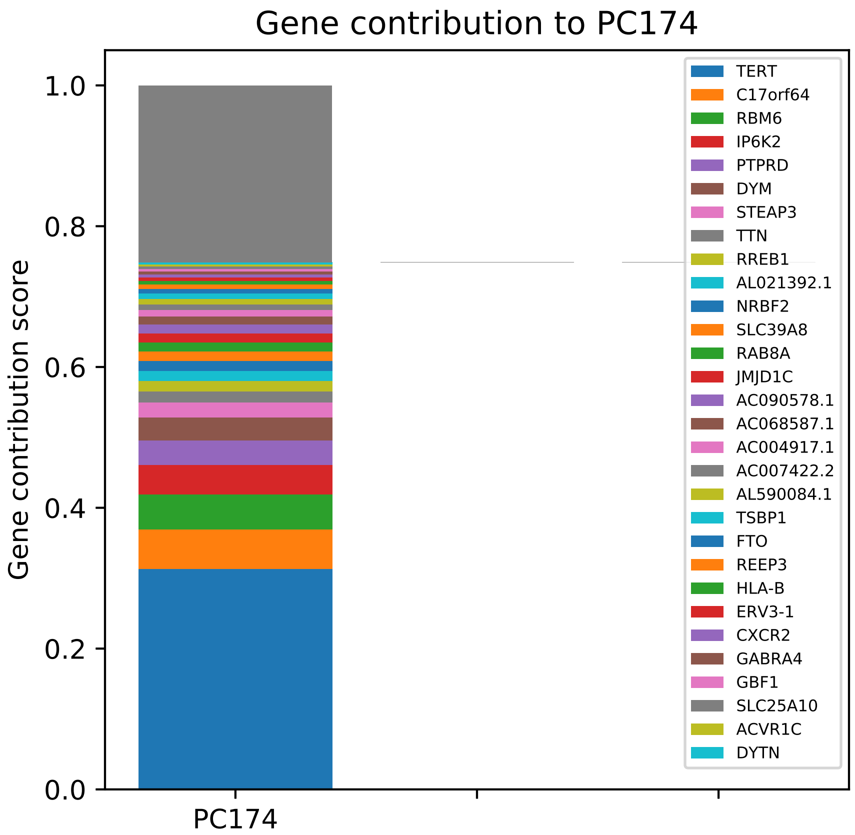
Gene contribution to PC174
Gene contribution score
1.0
0.8
0.6
0.4
0.2
0.0
PC174
TERT
C17orf64
RBM6
IP6K2
PTPRD
DYM
STEAP3
TTN
RREB1
AL021392.1
NRBF2
SLC39A8
RAB8A
JMJD1C
AC090578.1
AC068587.1
AC004917.1
AC007422.2
AL590084.1
TSBP1
FTO
REEP3
HLA-B
ERV3-1
CXCR2
GABRA4
GBF1
SLC25A10
ACVR1C
DYTN

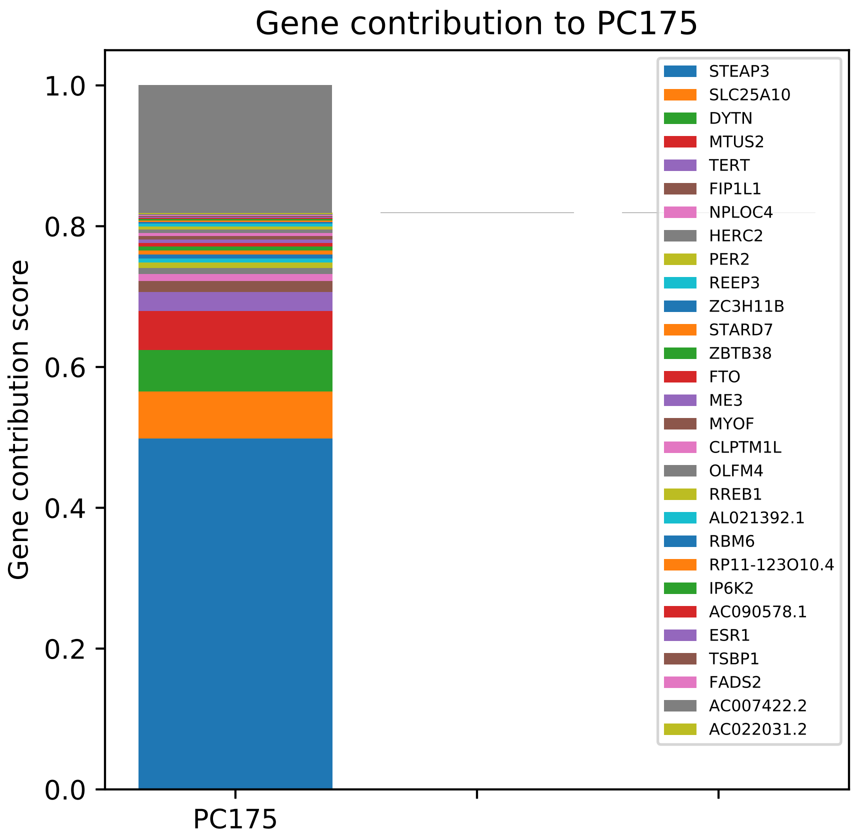
Gene contribution to PC175
Gene contribution score
1.0
0.8
0.6
0.4
0.2
0.0
PC175
STEAP3
SLC25A10
DYTN
MTUS2
TERT
FIP1L1
NPLOC4
HERC2
PER2
REEP3
ZC3H11B
STARD7
ZBTB38
FTO
ME3
MYOF
CLPTM1L
OLFM4
RREB1
AL021392.1
RBM6
RP11-123O10.4
IP6K2
AC090578.1
ESR1
TSBP1
FADS2
AC007422.2
AC022031.2

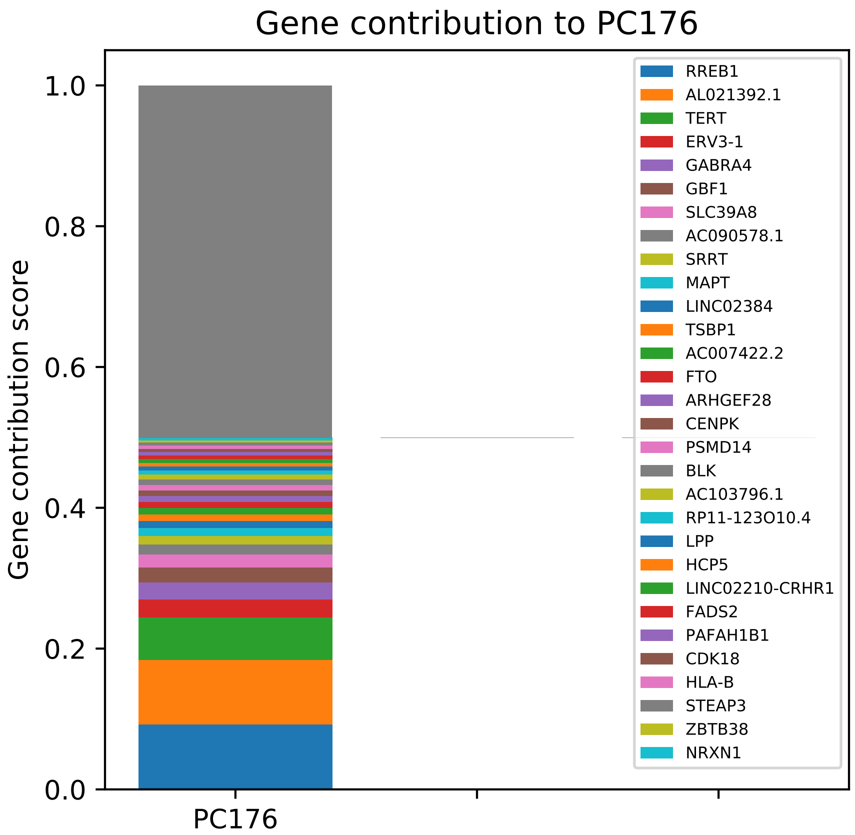

Gene contribution to PC176
Gene contribution score
1.0
0.8
0.6
0.4
0.2
0.0
PC176
RREB1
AL021392.1
TERT
ERV3-1
GABRA4
GBF1
SLC39A8
AC090578.1
SRRT
MAPT
LINC02384
TSBP1
AC007422.2
FTO
ARHGEF28
CENPK
PSMD14
BLK
AC103796.1
RP11-123O10.4
LPP
HCP5
LINC02210-CRHR1
FADS2
PAFAH1B1
CDK18
HLA-B
STEAP3
ZBTB38
NRXN1

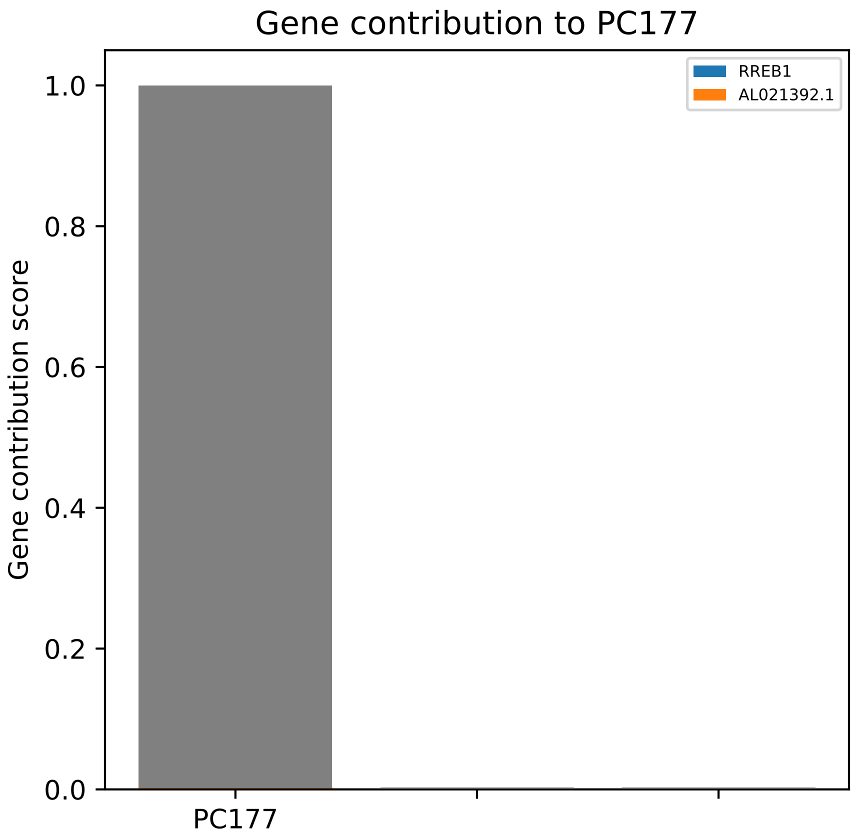

Gene contribution to PC177
RREB1
AL021392.1
1.0
0.8
0.6
0.4
0.2
0.0
Gene contribution score
PC177

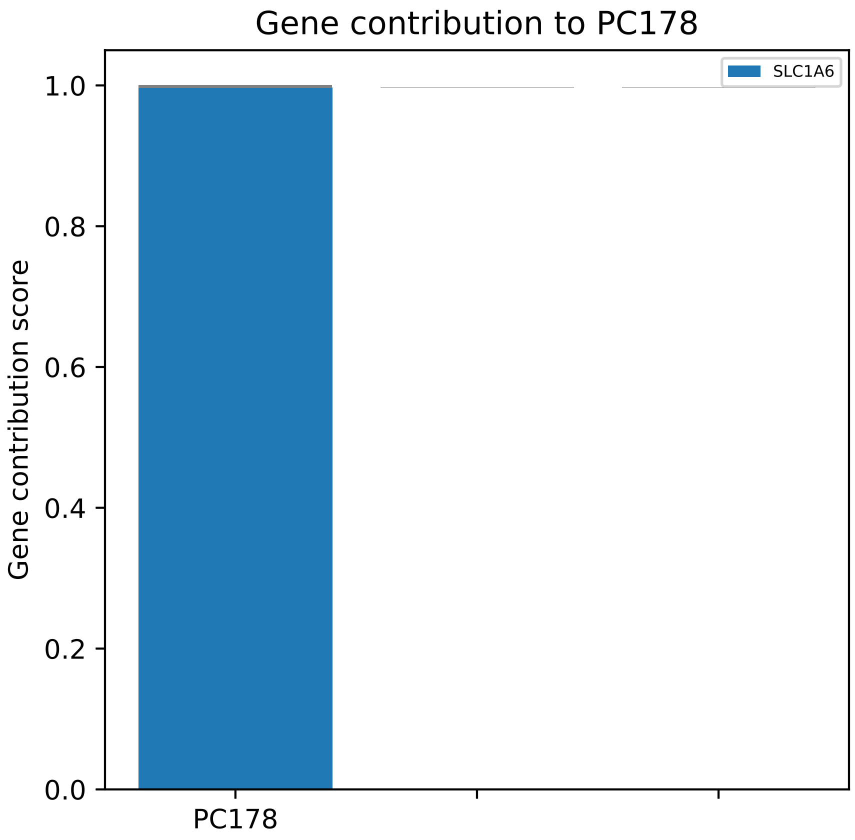

Gene contribution to PC178
Gene contribution score
1.0
0.8
0.6
0.4
0.2
0.0
PC178
SLC1A6

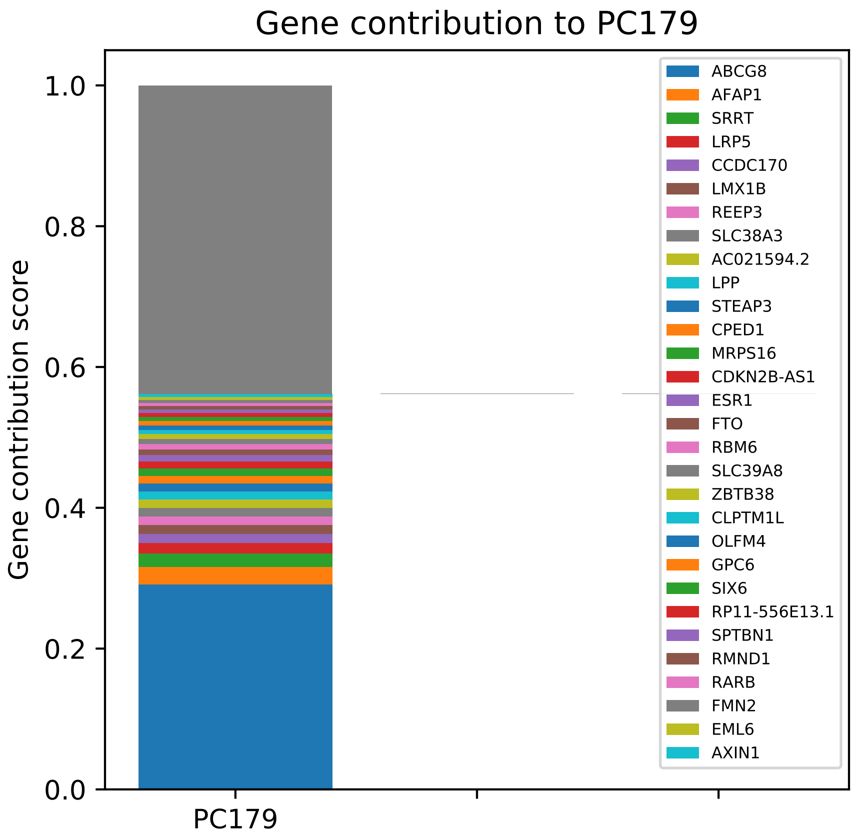

Gene contribution to PC179
Gene contribution score
1.0
0.8
0.6
0.4
0.2
0.0
PC179
ABCG8
AFAP1
SRRT
LRP5
CCDC170
LMX1B
REEP3
SLC38A3
AC021594.2
LPP
STEAP3
CPED1
MRPS16
CDKN2B-AS1
ESR1
FTO
RBM6
SLC39A8
ZBTB38
CLPTM1L
OLFM4
GPC6
SIX6
RP11-556E13.1
SPTBN1
RMND1
RARB
FMN2
EML6
AXIN1

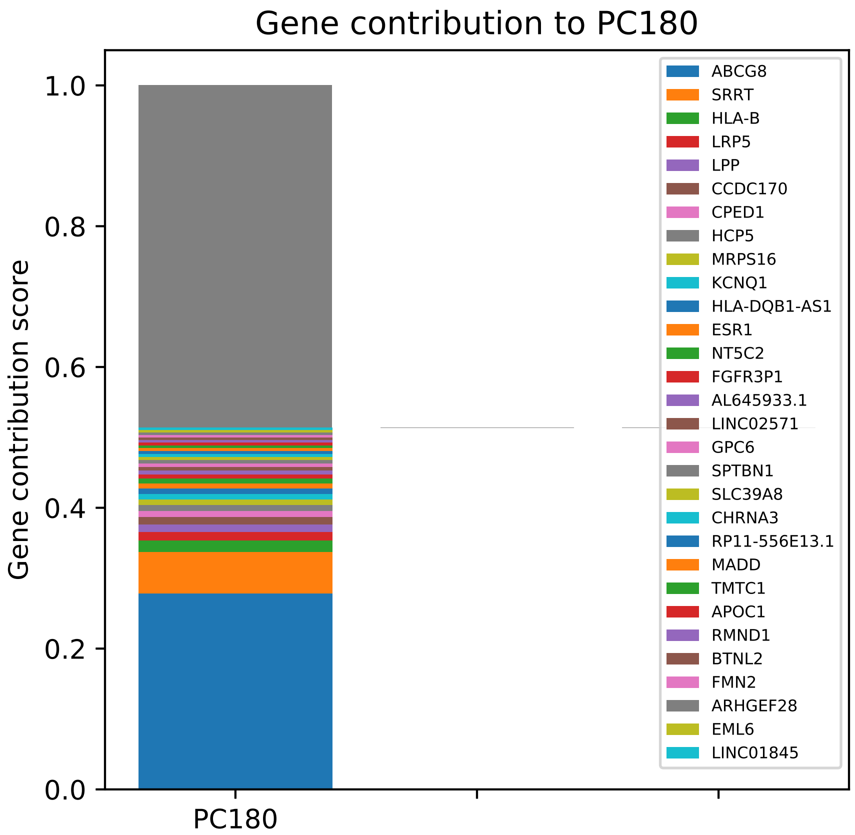

Gene contribution to PC180
Gene contribution score
1.0
0.8
0.6
0.4
0.2
0.0
PC180
ABCG8
SRRT
HLA-B
LRP5
LPP
CCDC170
CPED1
HCP5
MRPS16
KCNQ1
HLA-DQB1-AS1
ESR1
NT5C2
FGFR3P1
AL645933.1
LINC02571
GPC6
SPTBN1
SLC39A8
CHRNA3
RP11-556E13.1
MADD
TMTC1
APOC1
RMND1
BTNL2
FMN2
ARHGEF28
EML6
LINC01845

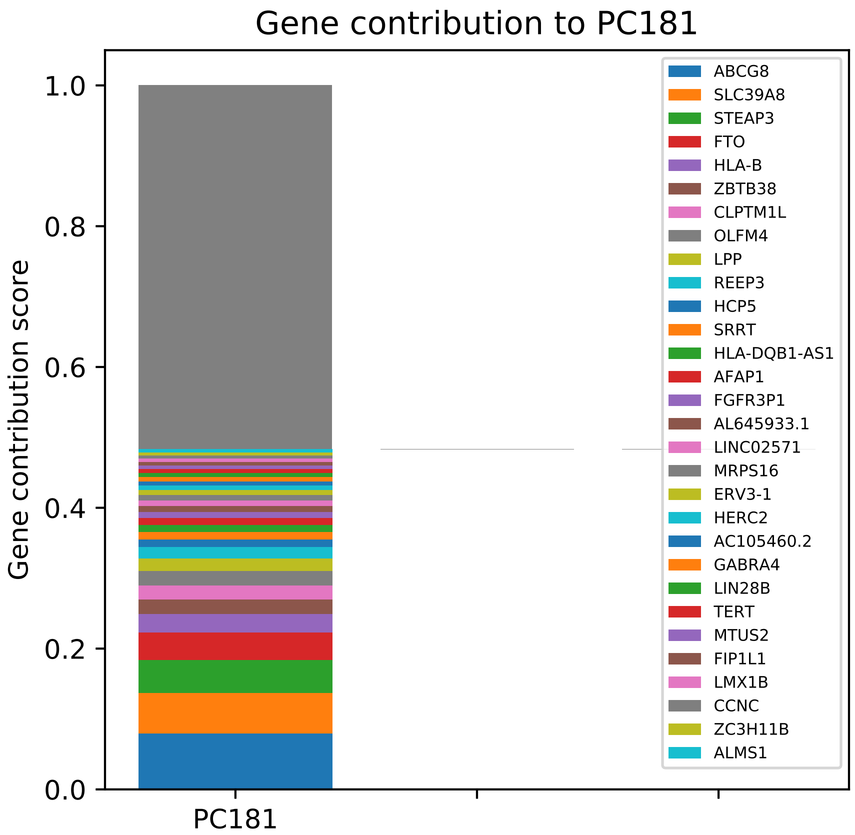

Gene contribution to PC181
Gene contribution score
1.0
0.8
0.6
0.4
0.2
0.0
PC181
ABCG8
SLC39A8
STEAP3
FTO
HLA-B
ZBTB38
CLPTM1L
OLFM4
LPP
REEP3
HCP5
SRRT
HLA-DQB1-AS1
AFAP1
FGFR3P1
AL645933.1
LINC02571
MRPS16
ERV3-1
HERC2
AC105460.2
GABRA4
LIN28B
TERT
MTUS2
FIP1L1
LMX1B
CCNC
ZC3H11B
ALMS1

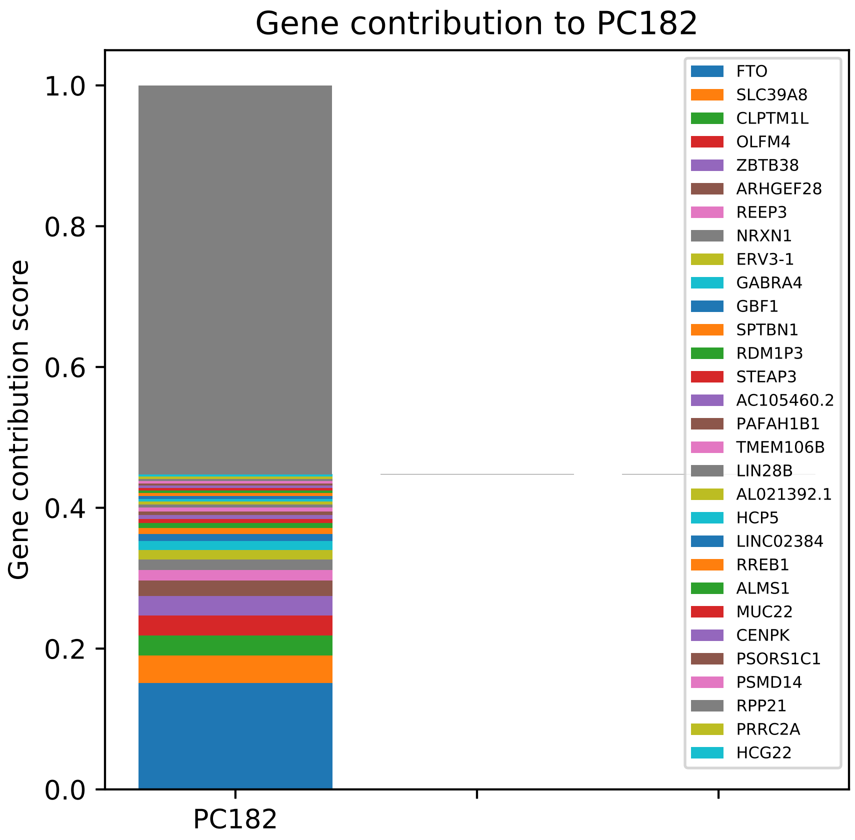
Gene contribution to PC182
Gene contribution score
1.0
0.8
0.6
0.4
0.2
0.0
PC182
FTO
SLC39A8
CLPTM1L
OLFM4
ZBTB38
ARHGEF28
REEP3
NRXN1
ERV3-1
GABRA4
GBF1
SPTBN1
RDM1P3
STEAP3
AC105460.2
PAFAH1B1
TMEM106B
LIN28B
AL021392.1
HCP5
LINC02384
RREB1
ALMS1
MUC22
CENPK
PSORS1C1
PSMD14
RPP21
PRRC2A
HCG22

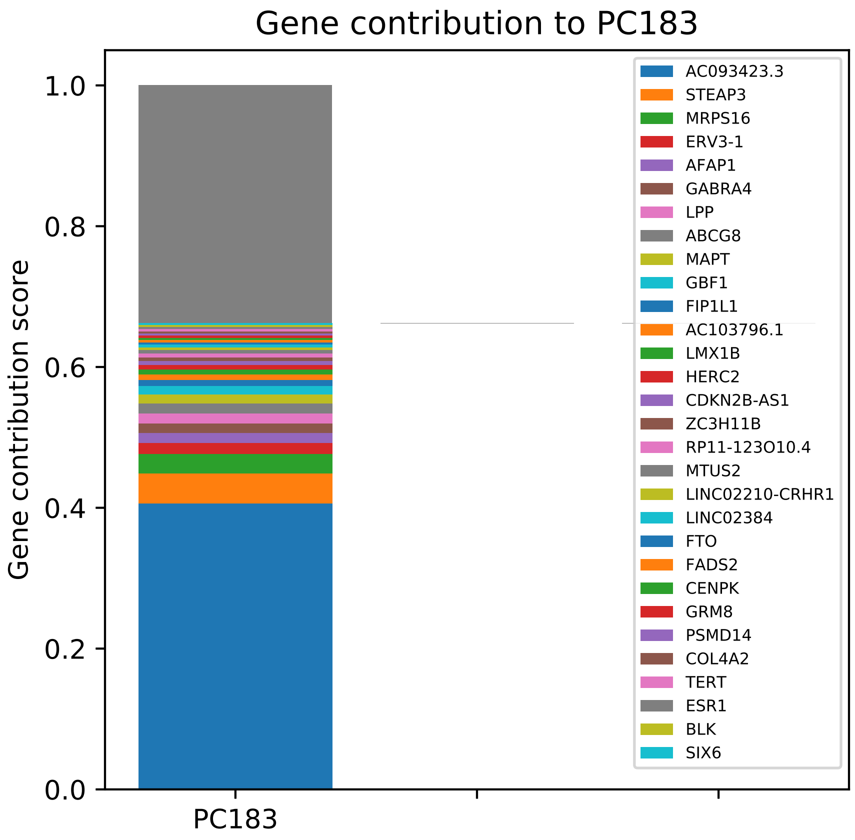

Gene contribution to PC183
Gene contribution score
1.0
0.8
0.6
0.4
0.2
0.0
PC183
AC093423.3
STEAP3
MRPS16
ERV3-1
AFAP1
GABRA4
LPP
ABCG8
MAPT
GBF1
FIP1L1
AC103796.1
LMX1B
HERC2
CDKN2B-AS1
ZC3H11B
RP11-123O10.4
MTUS2
LINC02210-CRHR1
LINC02384
FTO
FADS2
CENPK
GRM8
PSMD14
COL4A2
TERT
ESR1
BLK
SIX6

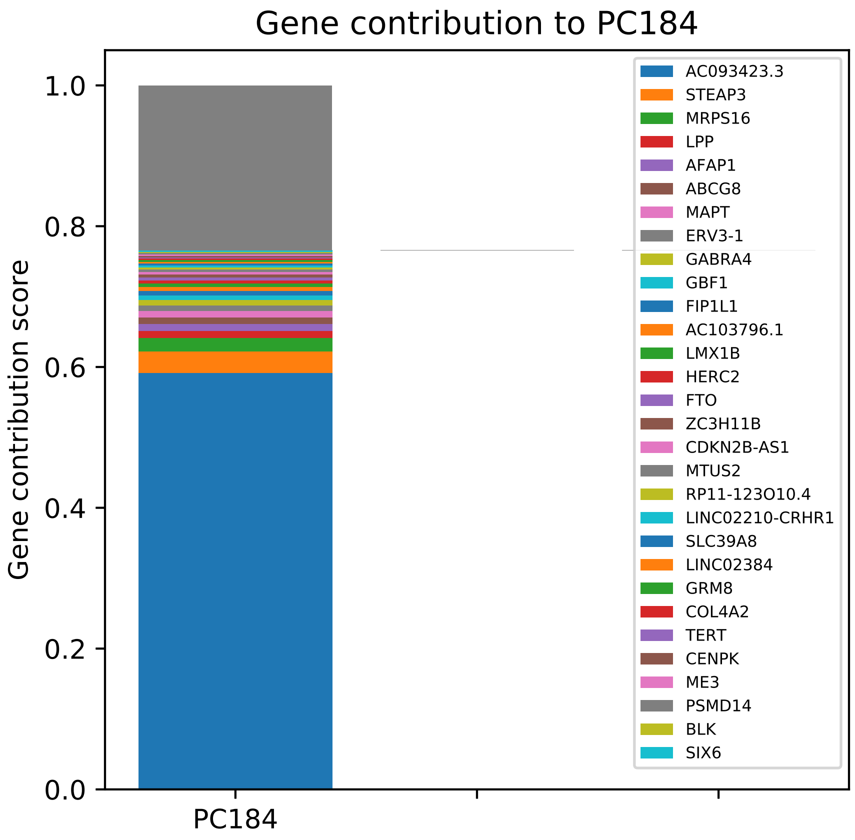

Gene contribution to PC184
Gene contribution score
0.0
0.2
0.4
0.6
0.8
1.0
PC184
AC093423.3
STEAP3
MRPS16
LPP
AFAP1
ABCG8
MAPT
ERV3-1
GABRA4
GBF1
FIP1L1
AC103796.1
LMX1B
HERC2
FTO
ZC3H11B
CDKN2B-AS1
MTUS2
RP11-123O10.4
LINC02210-CRHR1
SLC39A8
LINC02384
GRM8
COL4A2
TERT
CENPK
ME3
PSMD14
BLK
SIX6

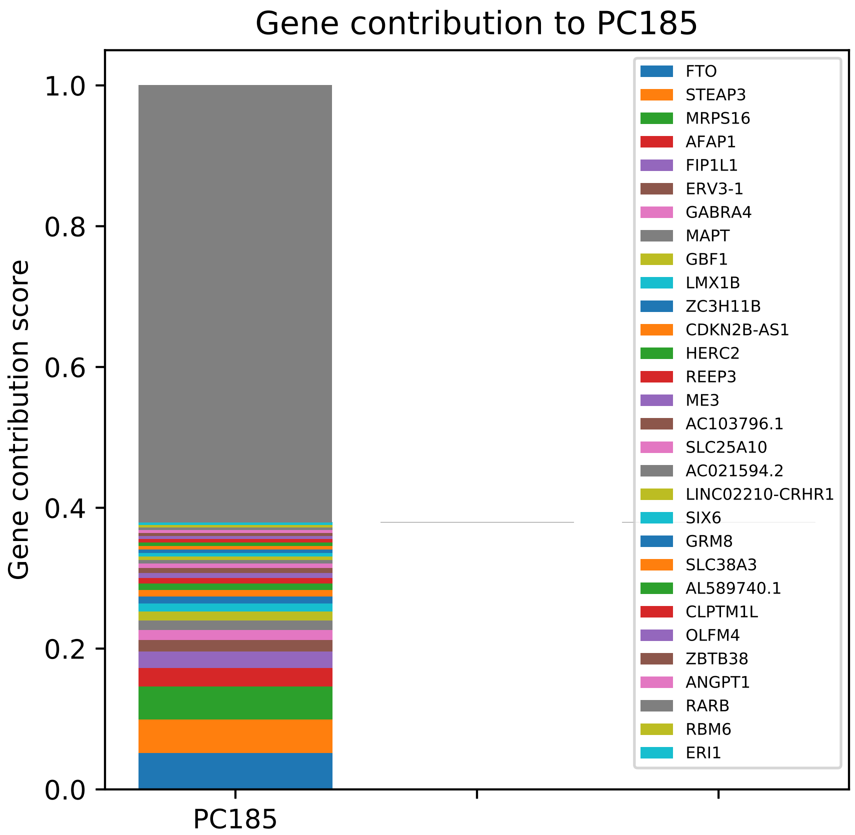
Gene contribution to PC185
Gene contribution score
1.0
0.8
0.6
0.4
0.2
0.0
PC185
FTO
STEAP3
MRPS16
AFAP1
FIP1L1
ERV3-1
GABRA4
MAPT
GBF1
LMX1B
ZC3H11B
CDKN2B-AS1
HERC2
REEP3
ME3
AC103796.1
SLC25A10
AC021594.2
LINC02210-CRHR1
SIX6
GRM8
SLC38A3
AL589740.1
CLPTM1L
OLFM4
ZBTB38
ANGPT1
RARB
RBM6
ERI1

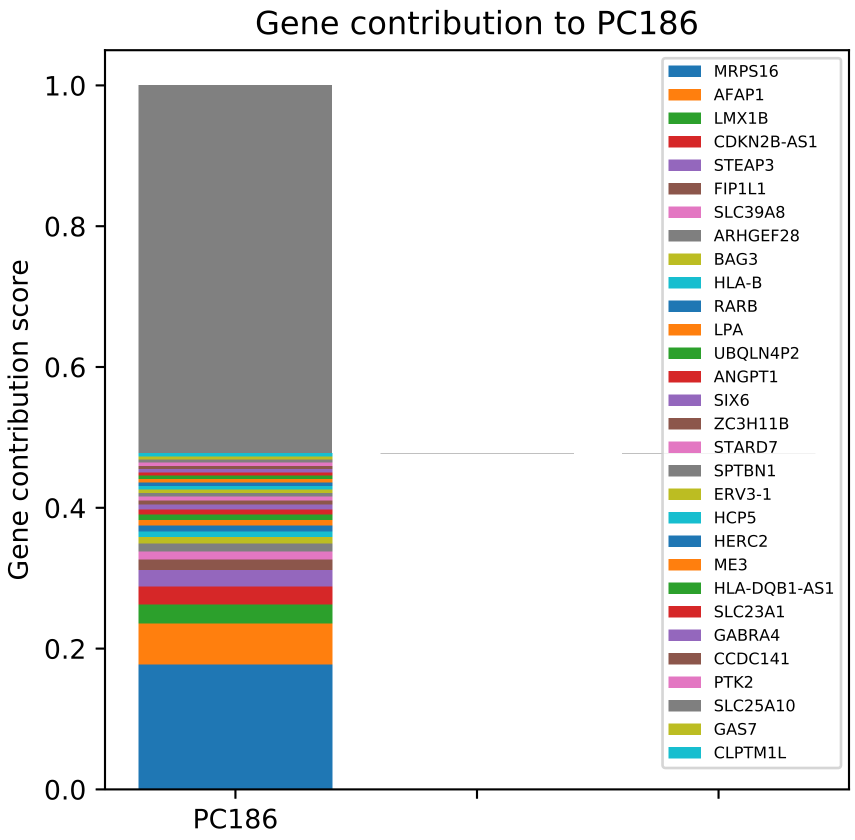

Gene contribution to PC186
Gene contribution score
1.0
0.8
0.6
0.4
0.2
0.0
PC186
MRPS16
AFAP1
LMX1B
CDKN2B-AS1
STEAP3
FIP1L1
SLC39A8
ARHGEF28
BAG3
HLA-B
RARB
LPA
UBQLN4P2
ANGPT1
SIX6
ZC3H11B
STARD7
SPTBN1
ERV3-1
HCP5
HERC2
ME3
HLA-DQB1-AS1
SLC23A1
GABRA4
CCDC141
PTK2
SLC25A10
GAS7
CLPTM1L

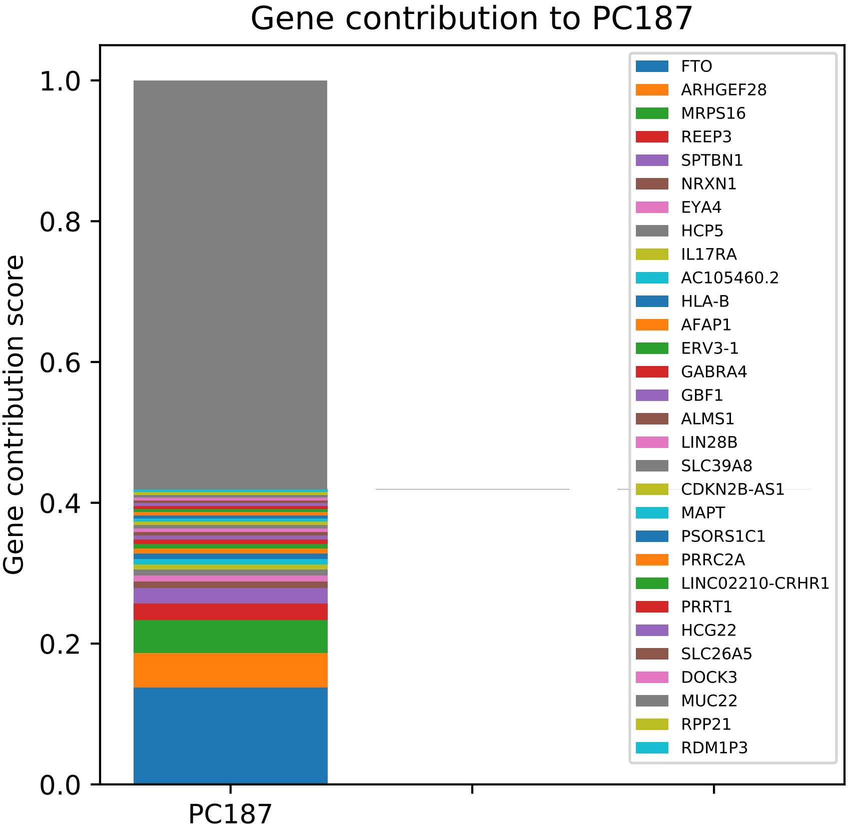
Gene contribution to PC187
Gene contribution score
1.0
0.8
0.6
0.4
0.2
0.0
PC187
FTO
ARHGEF28
MRPS16
REEP3
SPTBN1
NRXN1
EYA4
HCP5
IL17RA
AC105460.2
HLA-B
AFAP1
ERV3-1
GABRA4
GBF1
ALMS1
LIN28B
SLC39A8
CDKN2B-AS1
MAPT
PSORS1C1
PRRC2A
LINC02210-CRHR1
PRRT1
HCG22
SLC26A5
DOCK3
MUC22
RPP21
RDM1P3

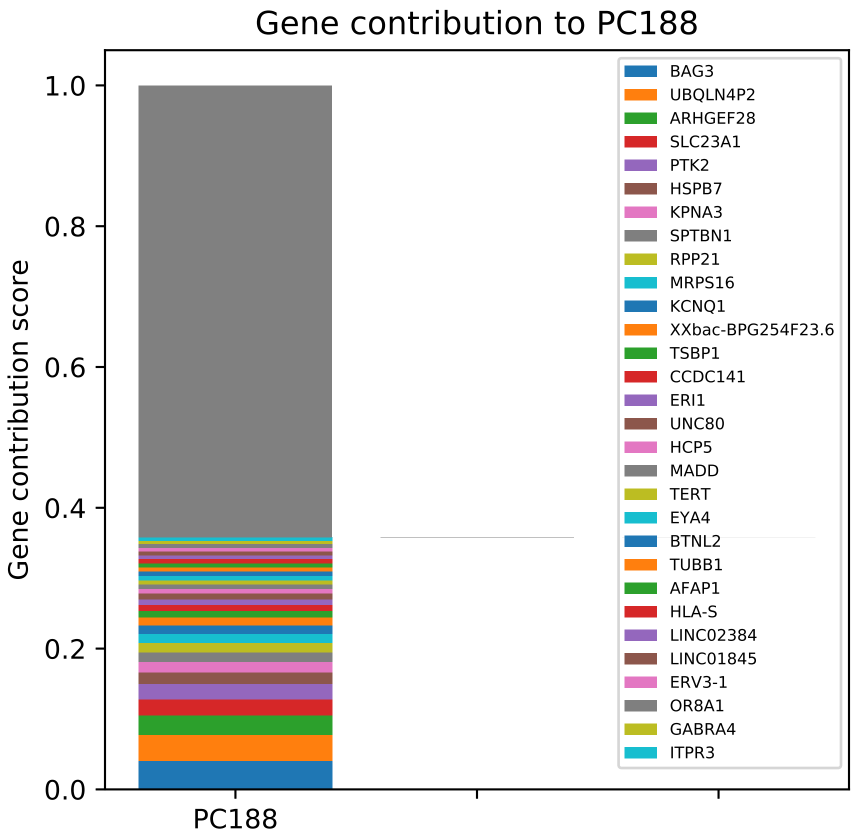

Gene contribution to PC188
Gene contribution score
1.0
0.8
0.6
0.4
0.2
0.0
PC188
BAG3
UBQLN4P2
ARHGEF28
SLC23A1
PTK2
HSPB7
KPNA3
SPTBN1
RPP21
MRPS16
KCNQ1
XXbac-BPG254F23.6
TSBP1
CCDC141
ERI1
UNC80
HCP5
MADD
TERT
EYA4
BTNL2
TUBB1
AFAP1
HLA-S
LINC02384
LINC01845
ERV3-1
OR8A1
GABRA4
ITPR3

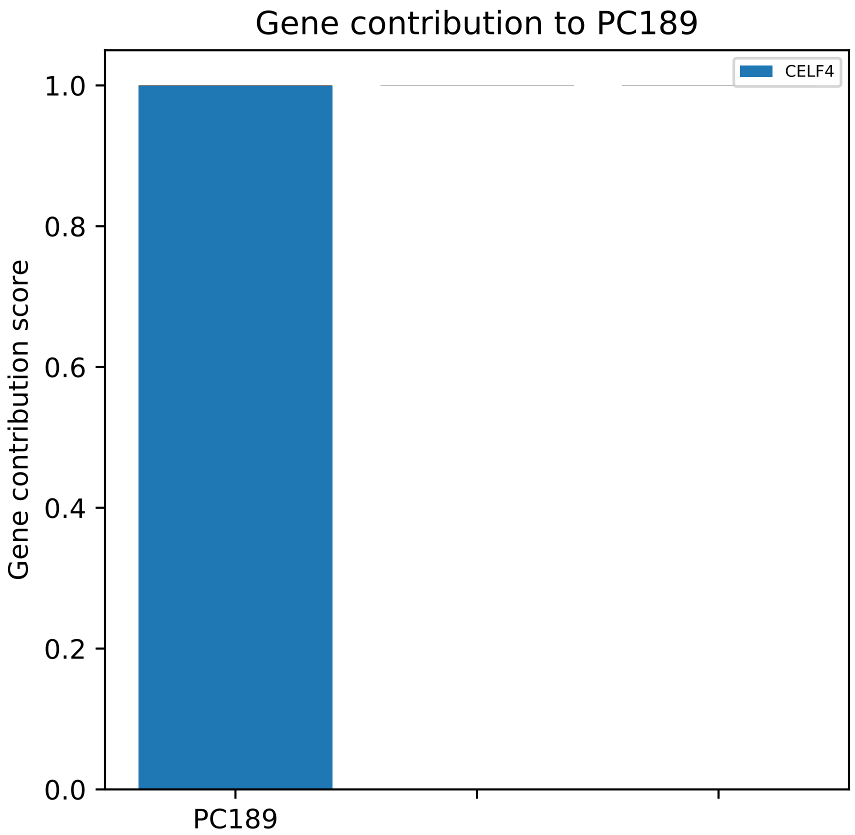
Gene contribution to PC189
Gene contribution score
CELF4
1.0
0.8
0.6
0.4
0.2
0.0
PC189

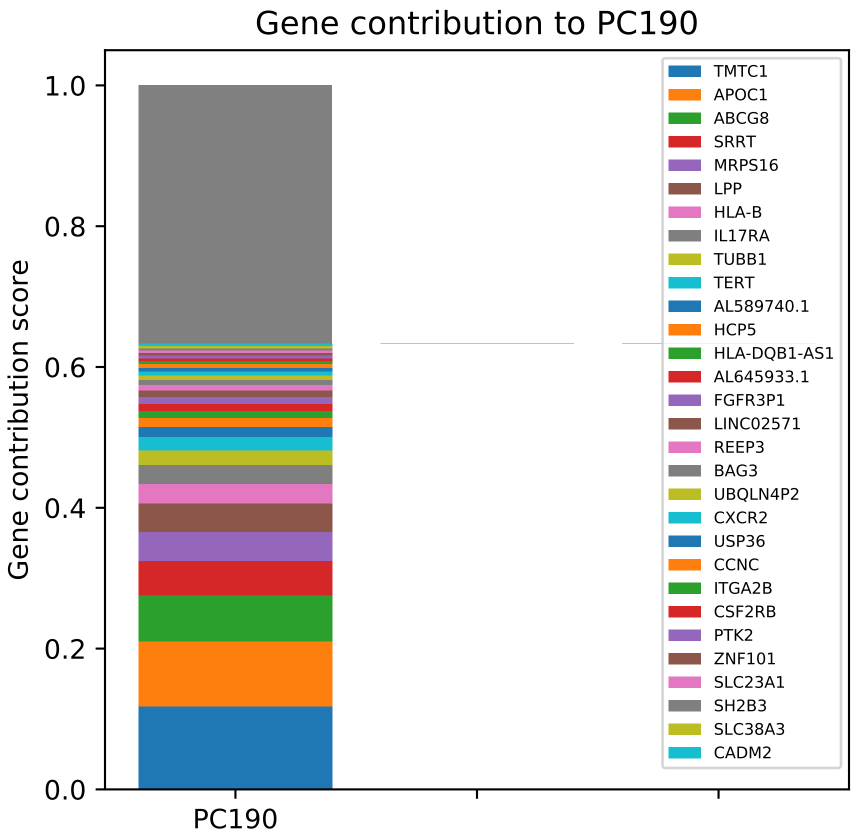

Gene contribution to PC190
Gene contribution score
1.0
0.8
0.6
0.4
0.2
0.0
PC190
TMTC1
APOC1
ABCG8
SRRT
MRPS16
LPP
HLA-B
IL17RA
TUBB1
TERT
AL589740.1
HCP5
HLA-DQB1-AS1
AL645933.1
FGFR3P1
LINC02571
REEP3
BAG3
UBQLN4P2
CXCR2
USP36
CCNC
ITGA2B
CSF2RB
PTK2
ZNF101
SLC23A1
SH2B3
SLC38A3
CADM2

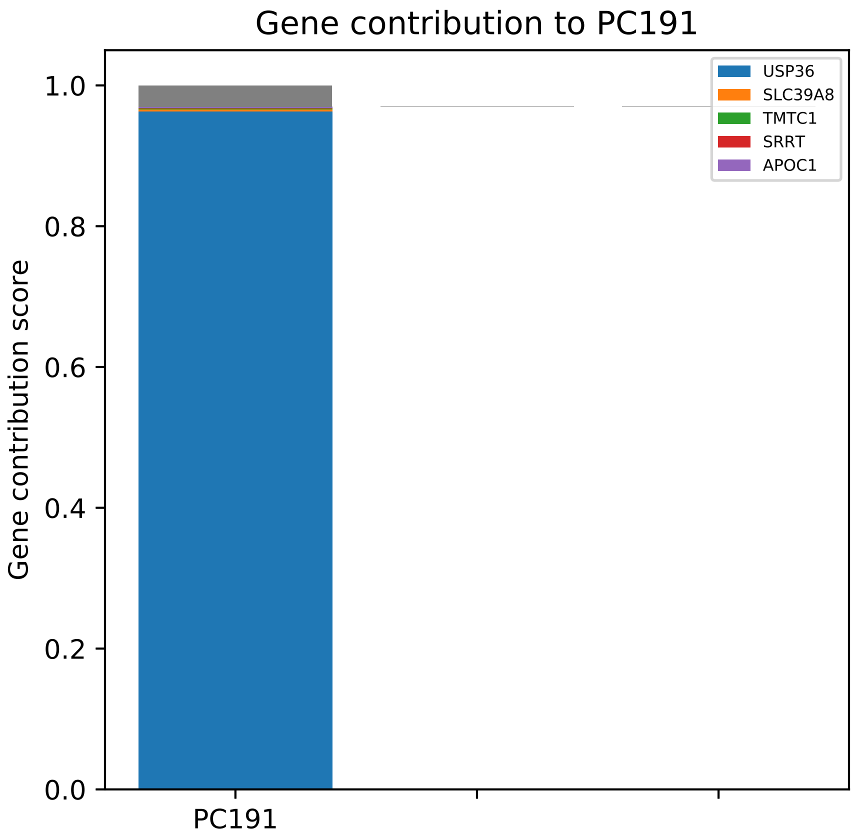
Gene contribution to PC191
Gene contribution score
1.0
0.8
0.6
0.4
0.2
0.0
PC191
USP36
SLC39A8
TMTC1
SRRT
APOC1

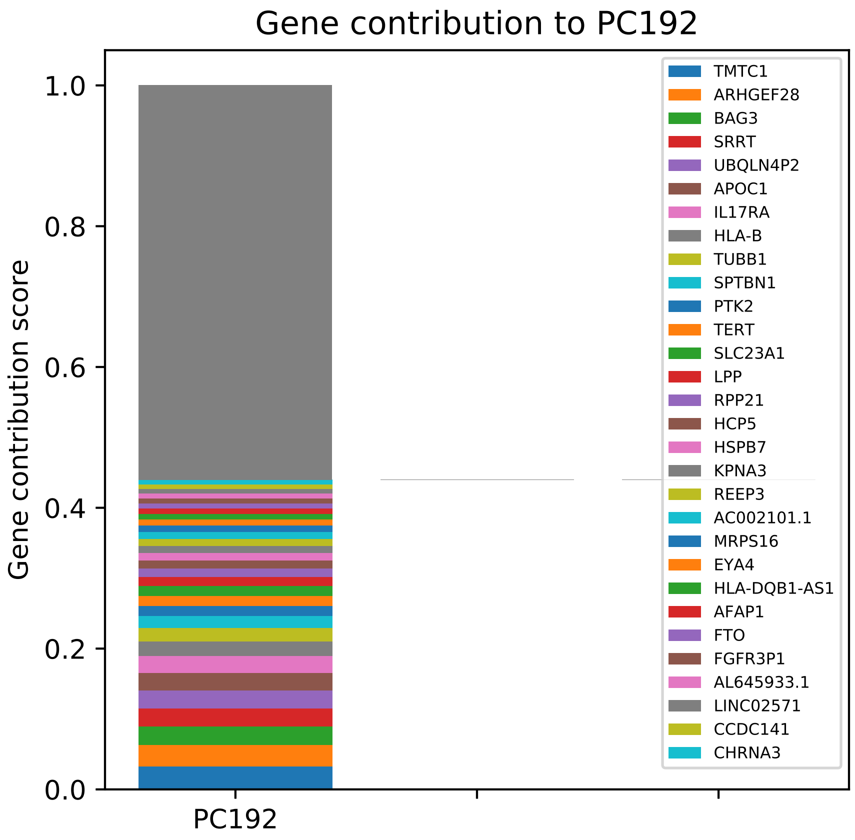

Gene contribution to PC192
Gene contribution score
1.0
0.8
0.6
0.4
0.2
0.0
PC192
TMTC1
ARHGEF28
BAG3
SRRT
UBQLN4P2
APOC1
IL17RA
HLA-B
TUBB1
SPTBN1
PTK2
TERT
SLC23A1
LPP
RPP21
HCP5
HSPB7
KPNA3
REEP3
AC002101.1
MRPS16
EYA4
HLA-DQB1-AS1
AFAP1
FTO
FGFR3P1
AL645933.1
LINC02571
CCDC141
CHRNA3

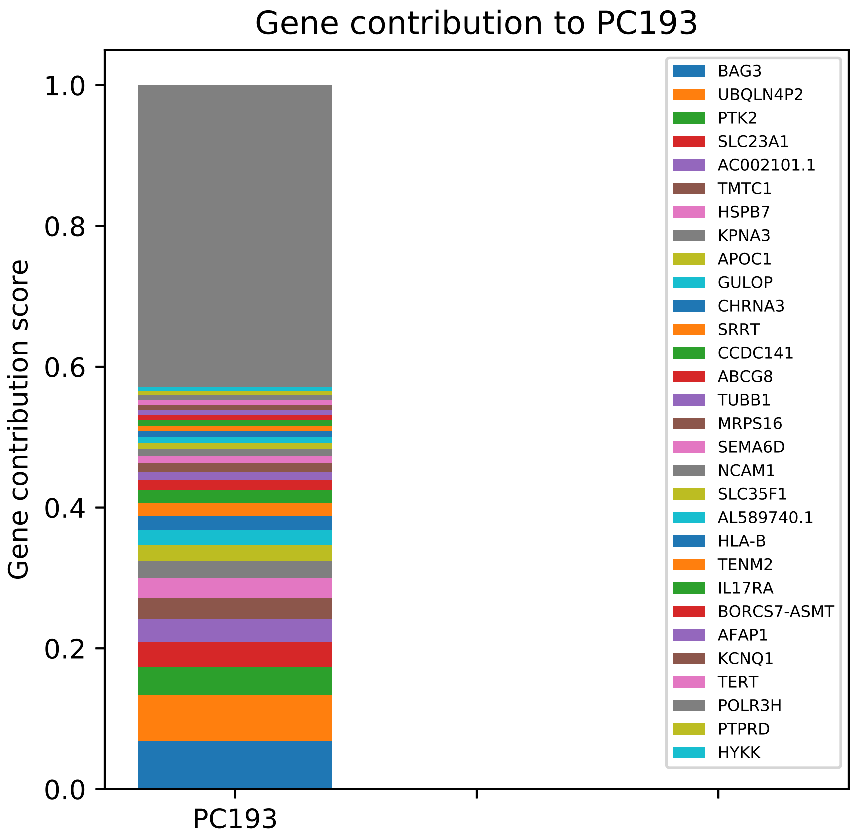

Gene contribution to PC193
Gene contribution score
0.0
0.2
0.4
0.6
0.8
1.0
PC193
BAG3
UBQLN4P2
PTK2
SLC23A1
AC002101.1
TMTC1
HSPB7
KPNA3
APOC1
GULOP
CHRNA3
SRRT
CCDC141
ABCG8
TUBB1
MRPS16
SEMA6D
NCAM1
SLC35F1
AL589740.1
HLA-B
TENM2
IL17RA
BORCS7-ASMT
AFAP1
KCNQ1
TERT
POLR3H
PTPRD
HYKK

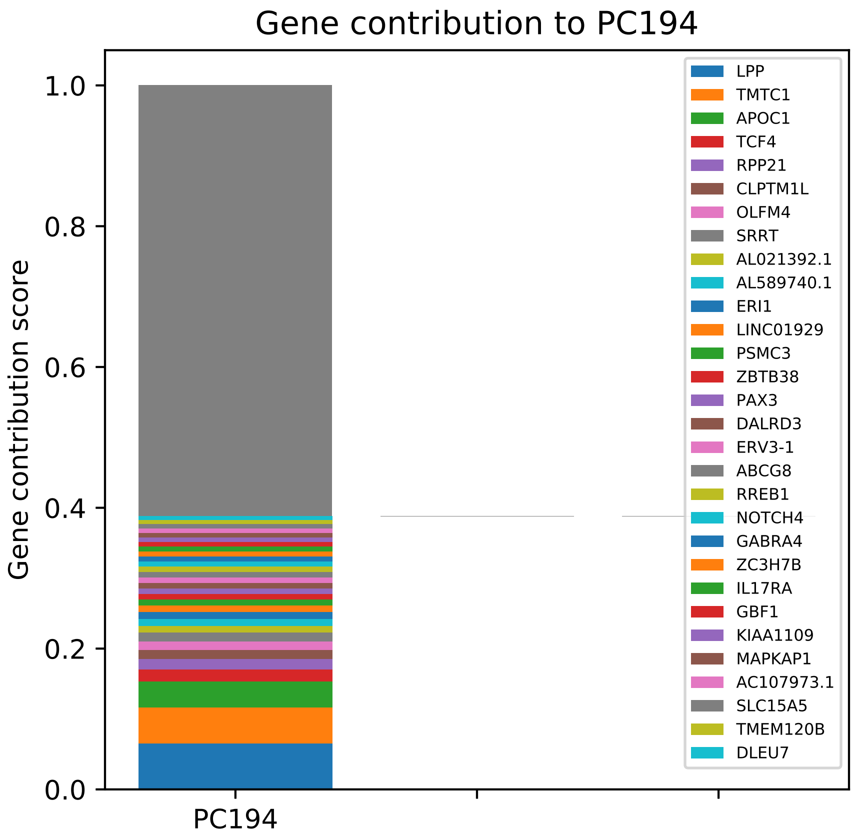
Gene contribution to PC194
Gene contribution score
1.0
0.8
0.6
0.4
0.2
0.0
PC194
LPP
TMTC1
APOC1
TCF4
RPP21
CLPTM1L
OLFM4
SRRT
AL021392.1
AL589740.1
ERI1
LINC01929
PSMC3
ZBTB38
PAX3
DALRD3
ERV3-1
ABCG8
RREB1
NOTCH4
GABRA4
ZC3H7B
IL17RA
GBF1
KIAA1109
MAPKAP1
AC107973.1
SLC15A5
TMEM120B
DLEU7

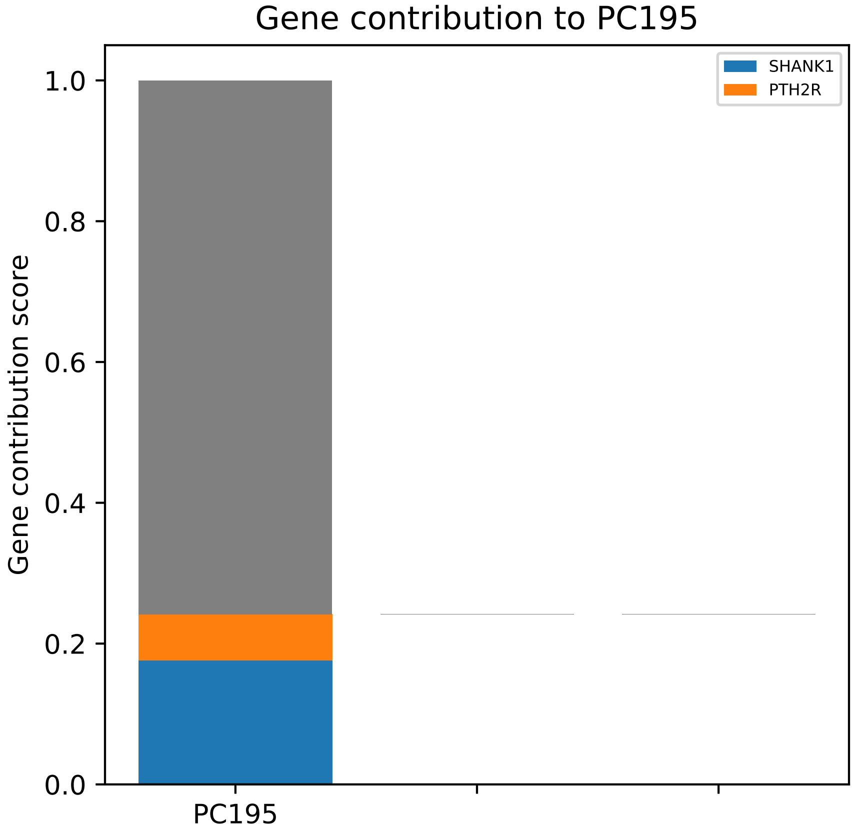
Gene contribution to PC195
SHANK1
PTH2R
Gene contribution score
1.0
0.8
0.6
0.4
0.2
0.0
PC195

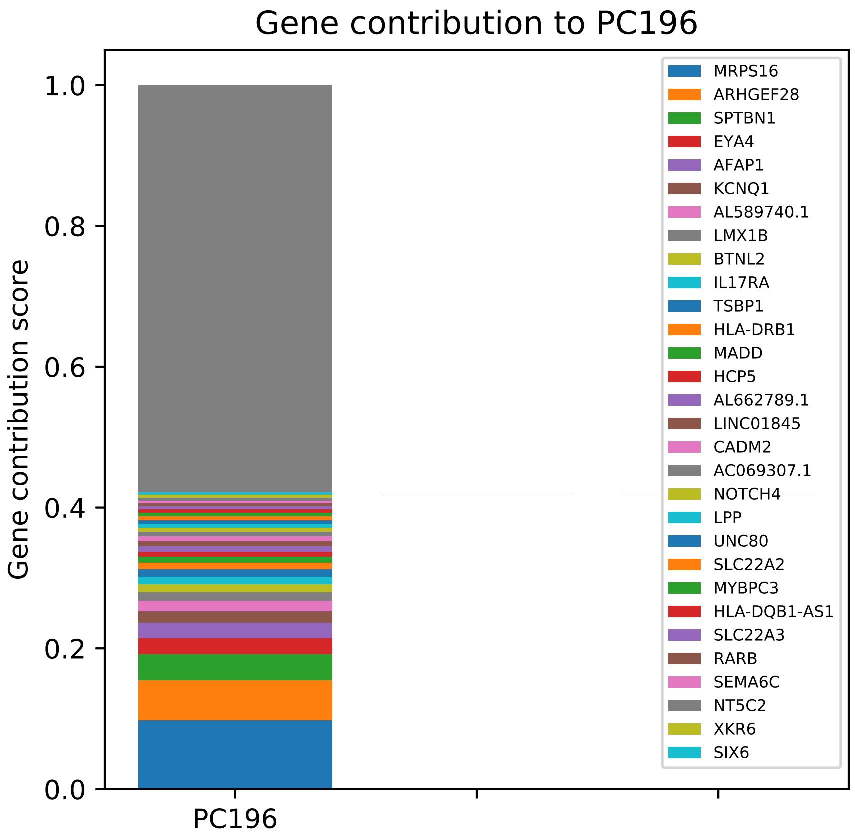
Gene contribution to PC196
Gene contribution score
1.0
0.8
0.6
0.4
0.2
0.0
PC196
MRPS16
ARHGEF28
SPTBN1
EYA4
AFAP1
KCNQ1
AL589740.1
LMX1B
BTNL2
IL17RA
TSBP1
HLA-DRB1
MADD
HCP5
AL662789.1
LINC01845
CADM2
AC069307.1
NOTCH4
LPP
UNC80
SLC22A2
MYBPC3
HLA-DQB1-AS1
SLC22A3
RARB
SEMA6C
NT5C2
XKR6
SIX6

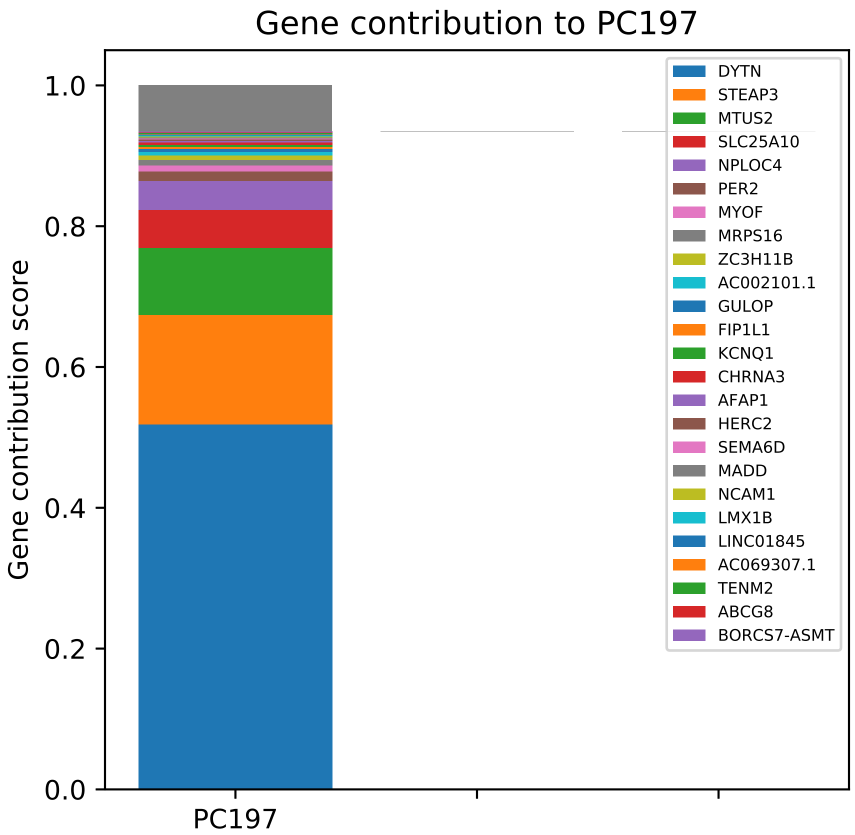

Gene contribution to PC197
Gene contribution score
1.0
0.8
0.6
0.4
0.2
0.0
PC197
DYTN
STEAP3
MTUS2
SLC25A10
NPLOC4
PER2
MYOF
MRPS16
ZC3H11B
AC002101.1
GULOP
FIP1L1
KCNQ1
CHRNA3
AFAP1
HERC2
SEMA6D
MADD
NCAM1
LMX1B
LINC01845
AC069307.1
TENM2
ABCG8
BORCS7-ASMT

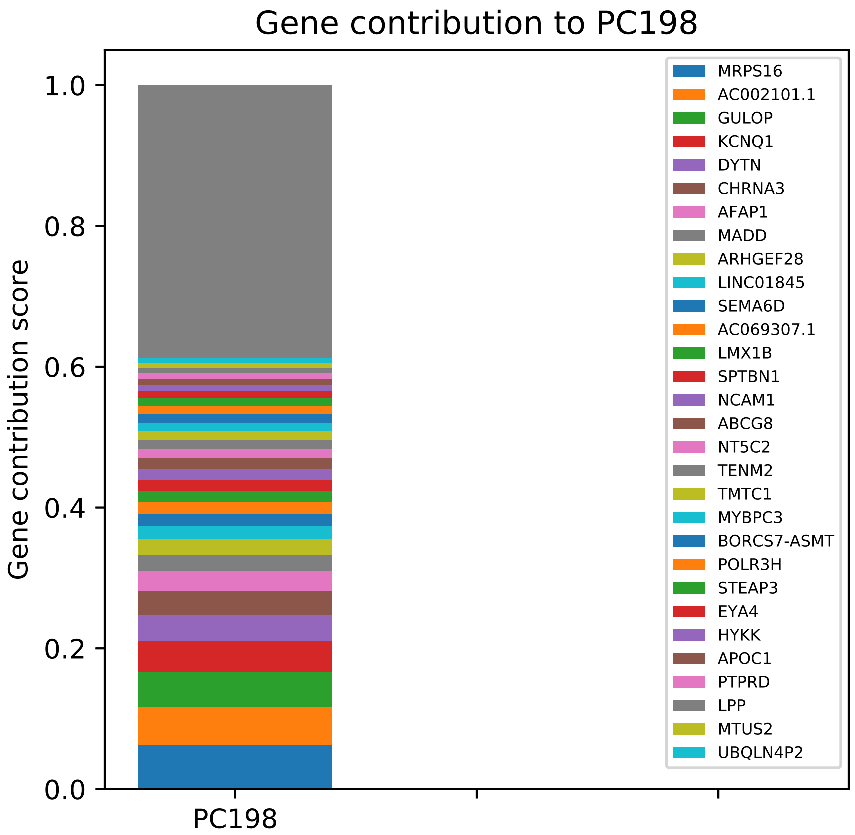

Gene contribution to PC198
Gene contribution score
1.0
0.8
0.6
0.4
0.2
0.0
PC198
MRPS16
AC002101.1
GULOP
KCNQ1
DYTN
CHRNA3
AFAP1
MADD
ARHGEF28
LINC01845
SEMA6D
AC069307.1
LMX1B
SPTBN1
NCAM1
ABCG8
NT5C2
TENM2
TMTC1
MYBPC3
BORCS7-ASMT
POLR3H
STEAP3
EYA4
HYKK
APOC1
PTPRD
LPP
MTUS2
UBQLN4P2

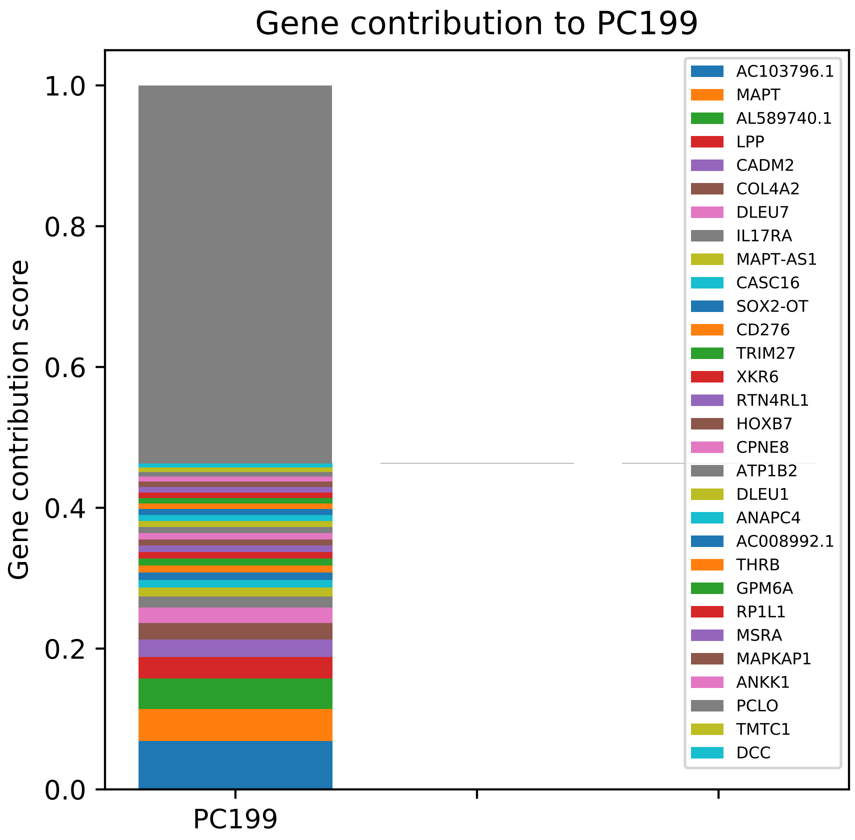

Gene contribution to PC199
Gene contribution score
1.0
0.8
0.6
0.4
0.2
0.0
PC199
AC103796.1
MAPT
AL589740.1
LPP
CADM2
COL4A2
DLEU7
IL17RA
MAPT-AS1
CASC16
SOX2-OT
CD276
TRIM27
XKR6
RTN4RL1
HOXB7
CPNE8
ATP1B2
DLEU1
ANAPC4
AC008992.1
THRB
GPM6A
RP1L1
MSRA
MAPKAP1
ANKK1
PCLO
TMTC1
DCC

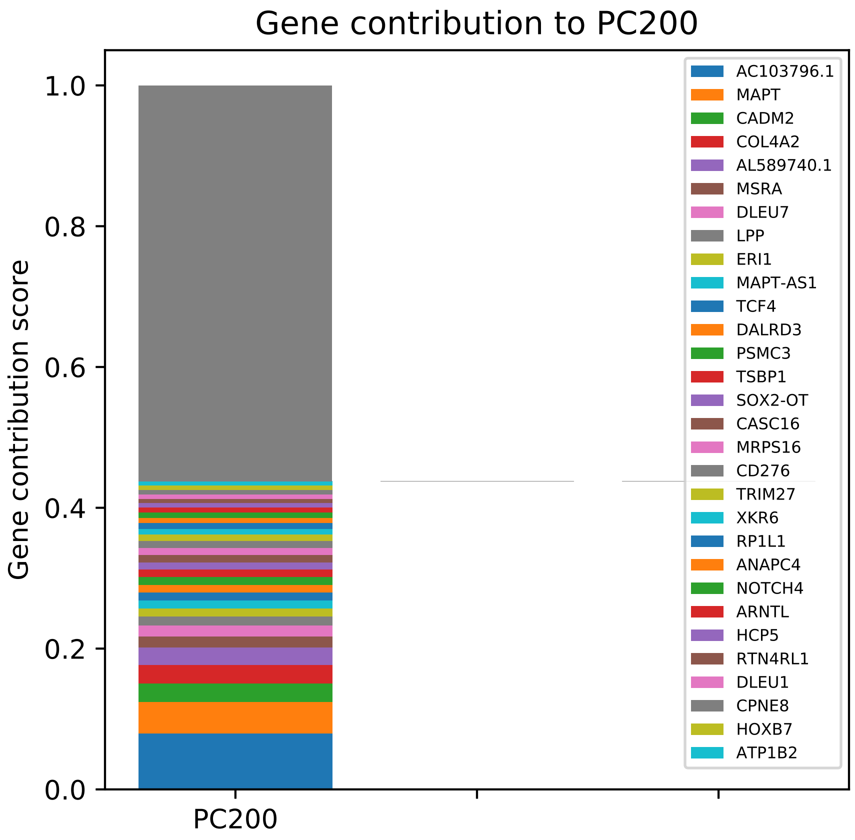

Gene contribution to PC200
Gene contribution score
1.0
0.8
0.6
0.4
0.2
0.0
PC200
AC103796.1
MAPT
CADM2
COL4A2
AL589740.1
MSRA
DLEU7
LPP
ERI1
MAPT-AS1
TCF4
DALRD3
PSMC3
TSBP1
SOX2-OT
CASC16
MRPS16
CD276
TRIM27
XKR6
RP1L1
ANAPC4
NOTCH4
ARNTL
HCP5
RTN4RL1
DLEU1
CPNE8
HOXB7
ATP1B2

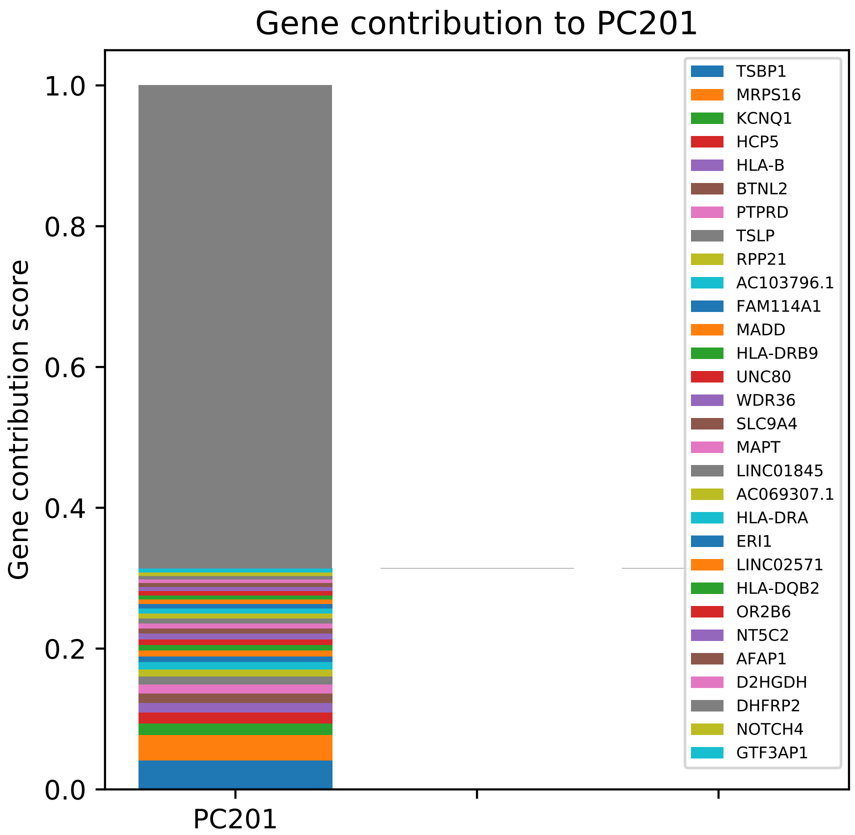

Gene contribution to PC201
Gene contribution score
1.0
0.8
0.6
0.4
0.2
0.0
PC201
TSBP1
MRPS16
KCNQ1
HCP5
HLA-B
BTNL2
PTPRD
TSLP
RPP21
AC103796.1
FAM114A1
MADD
HLA-DRB9
UNC80
WDR36
SLC9A4
MAPT
LINC01845
AC069307.1
HLA-DRA
ERI1
LINC02571
HLA-DQB2
OR2B6
NT5C2
AFAP1
D2HGDH
DHFRP2
NOTCH4
GTF3AP1

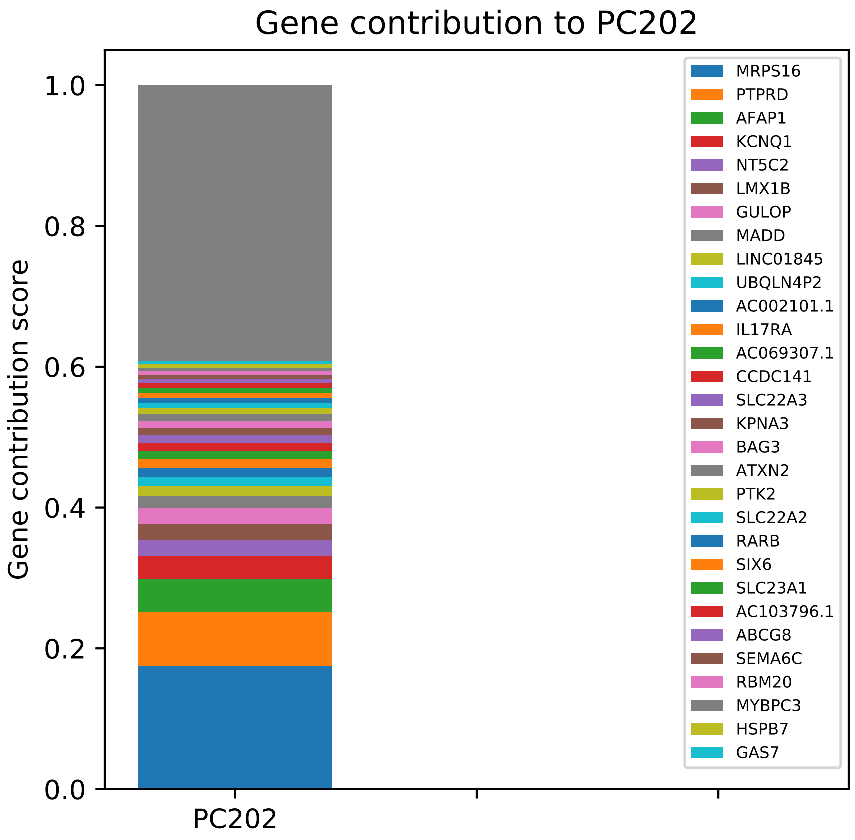

Gene contribution to PC202
Gene contribution score
1.0
0.8
0.6
0.4
0.2
0.0
PC202
MRPS16
PTPRD
AFAP1
KCNQ1
NT5C2
LMX1B
GULOP
MADD
LINC01845
UBQLN4P2
AC002101.1
IL17RA
AC069307.1
CCDC141
SLC22A3
KPNA3
BAG3
ATXN2
PTK2
SLC22A2
RARB
SIX6
SLC23A1
AC103796.1
ABCG8
SEMA6C
RBM20
MYBPC3
HSPB7
GAS7

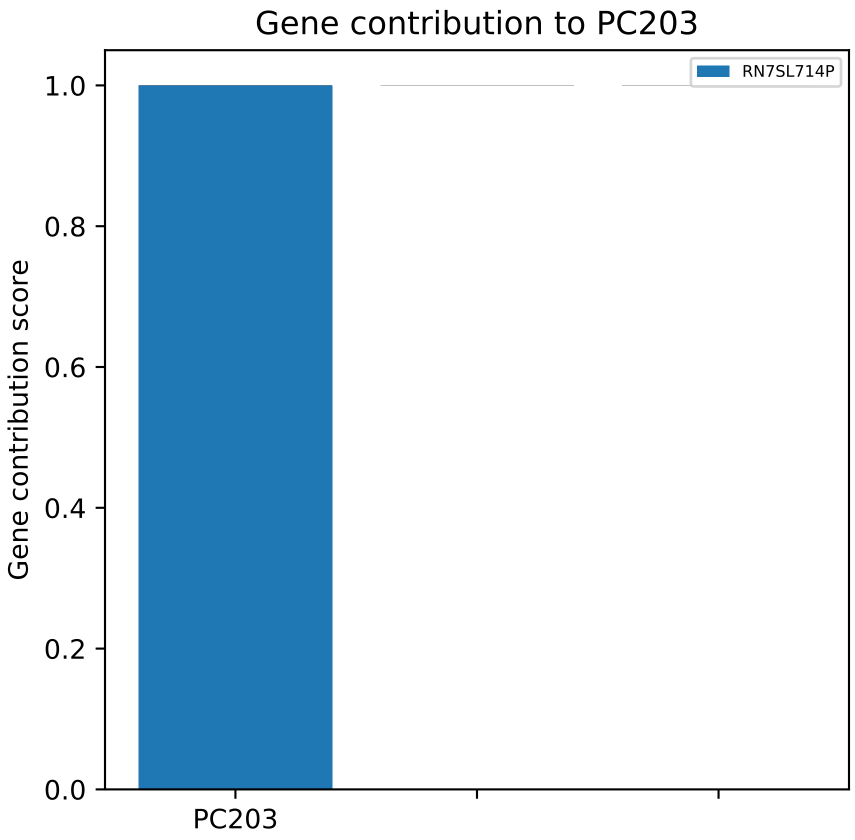
Gene contribution to PC203
RN7SL714P
Gene contribution score
1.0
0.8
0.6
0.4
0.2
0.0
PC203

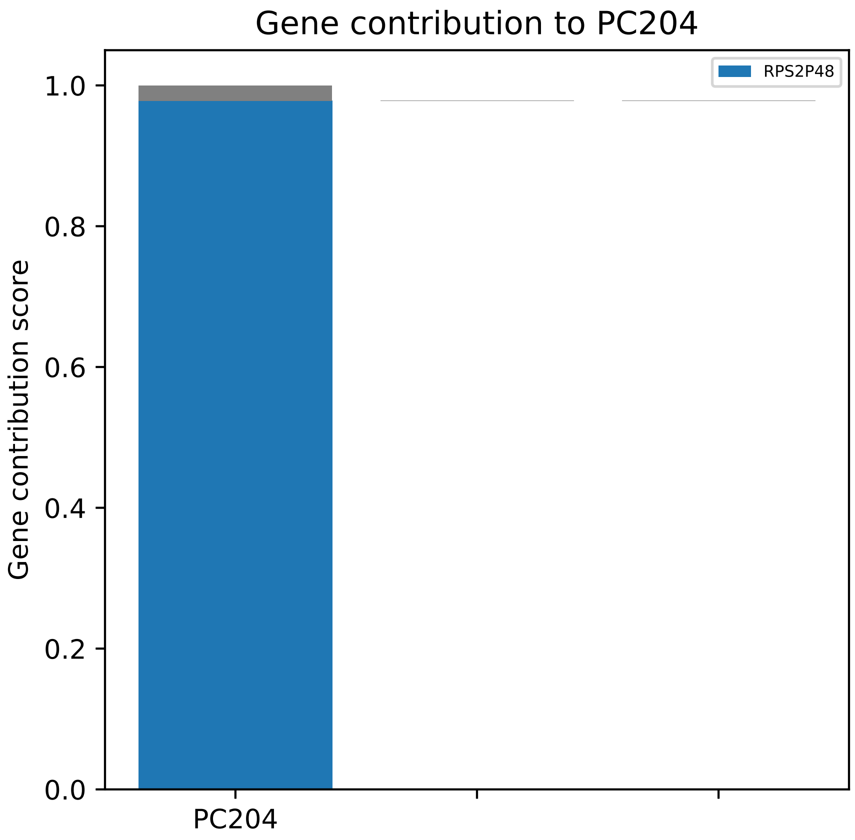

Gene contribution to PC204
RPS2P48
Gene contribution score
1.0
0.8
0.6
0.4
0.2
0.0
PC204

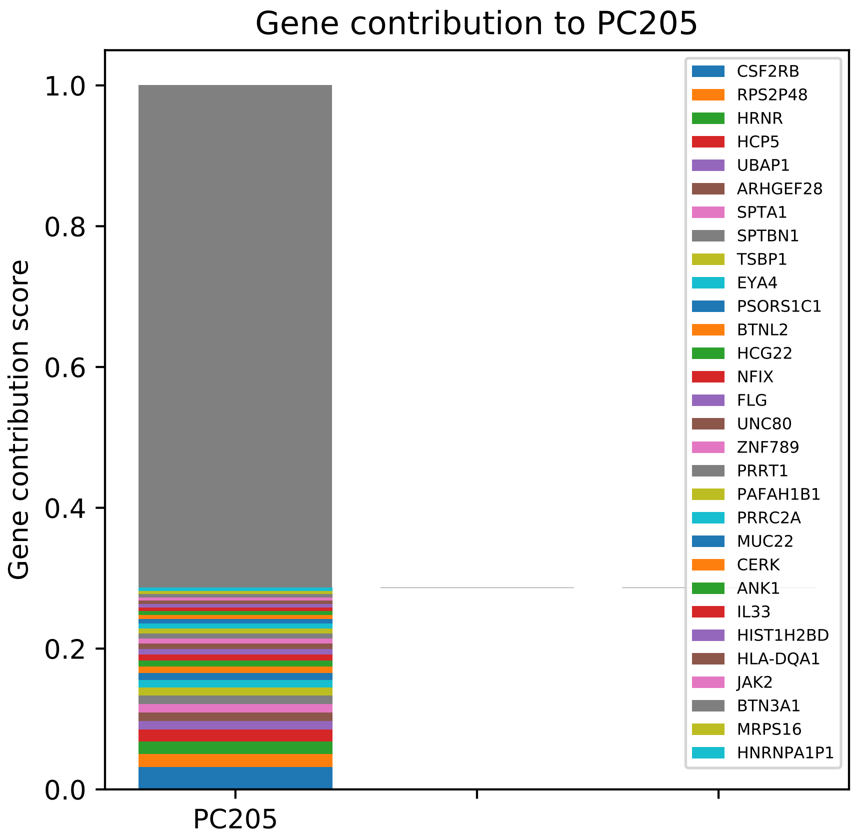

Gene contribution to PC205
Gene contribution score
1.0
0.8
0.6
0.4
0.2
0.0
PC205
CSF2RB
RPS2P48
HRNR
HCP5
UBAP1
ARHGEF28
SPTA1
SPTBN1
TSBP1
EYA4
PSORS1C1
BTNL2
HCG22
NFIX
FLG
UNC80
ZNF789
PRRT1
PAFAH1B1
PRRC2A
MUC22
CERK
ANK1
IL33
HIST1H2BD
HLA-DQA1
JAK2
BTN3A1
MRPS16
HNRNPA1P1

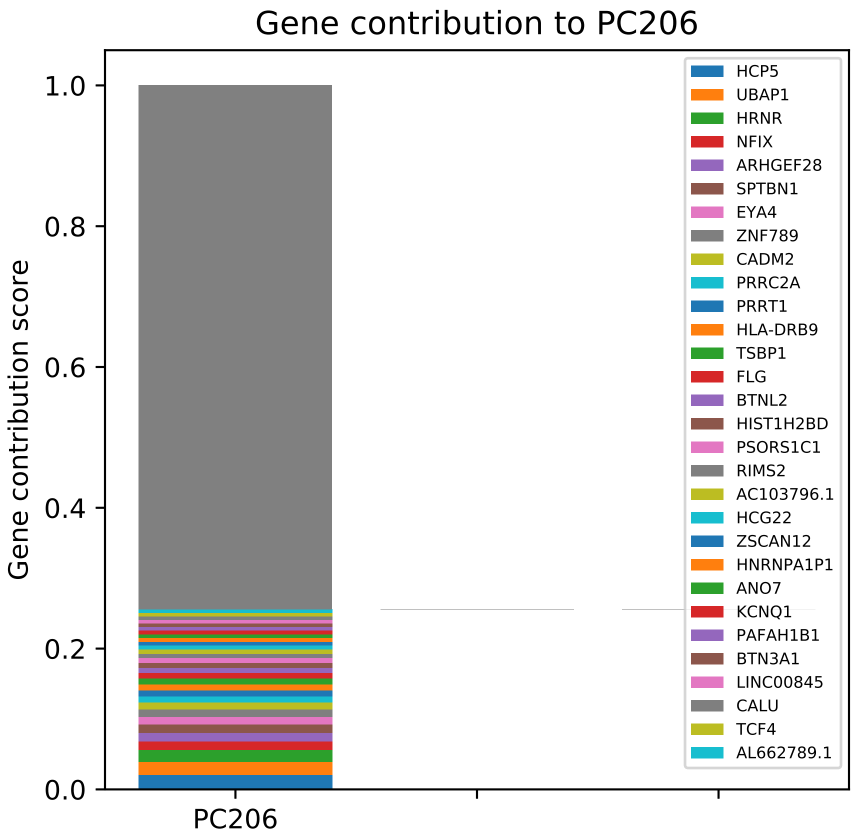
Gene contribution to PC206
Gene contribution score
1.0
0.8
0.6
0.4
0.2
0.0
PC206
HCP5
UBAP1
HRNR
NFIX
ARHGEF28
SPTBN1
EYA4
ZNF789
CADM2
PRRC2A
PRRT1
HLA-DRB9
TSBP1
FLG
BTNL2
HIST1H2BD
PSORS1C1
RIMS2
AC103796.1
HCG22
ZSCAN12
HNRNPA1P1
ANO7
KCNQ1
PAFAH1B1
BTN3A1
LINC00845
CALU
TCF4
AL662789.1

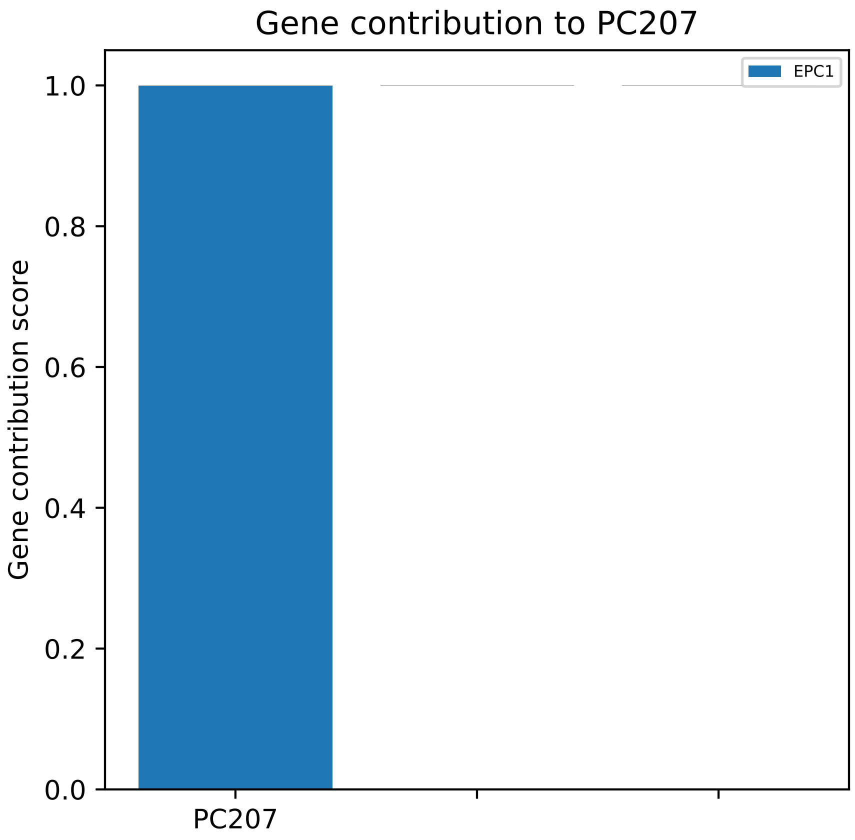
Gene contribution to PC207
EPC1
Gene contribution score
1.0
0.8
0.6
0.4
0.2
0.0
PC207

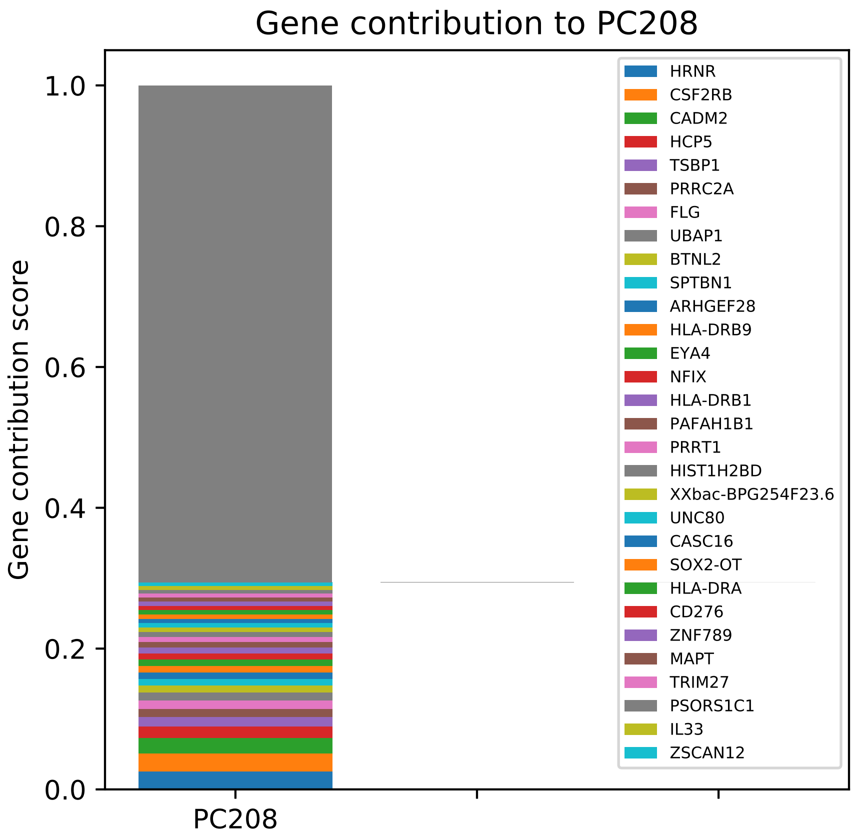

Gene contribution to PC208
Gene contribution score
1.0
0.8
0.6
0.4
0.2
0.0
PC208
HRNR
CSF2RB
CADM2
HCP5
TSBP1
PRRC2A
FLG
UBAP1
BTNL2
SPTBN1
ARHGEF28
HLA-DRB9
EYA4
NFIX
HLA-DRB1
PAFAH1B1
PRRT1
HIST1H2BD
XXbac-BPG254F23.6
UNC80
CASC16
SOX2-OT
HLA-DRA
CD276
ZNF789
MAPT
TRIM27
PSORS1C1
IL33
ZSCAN12

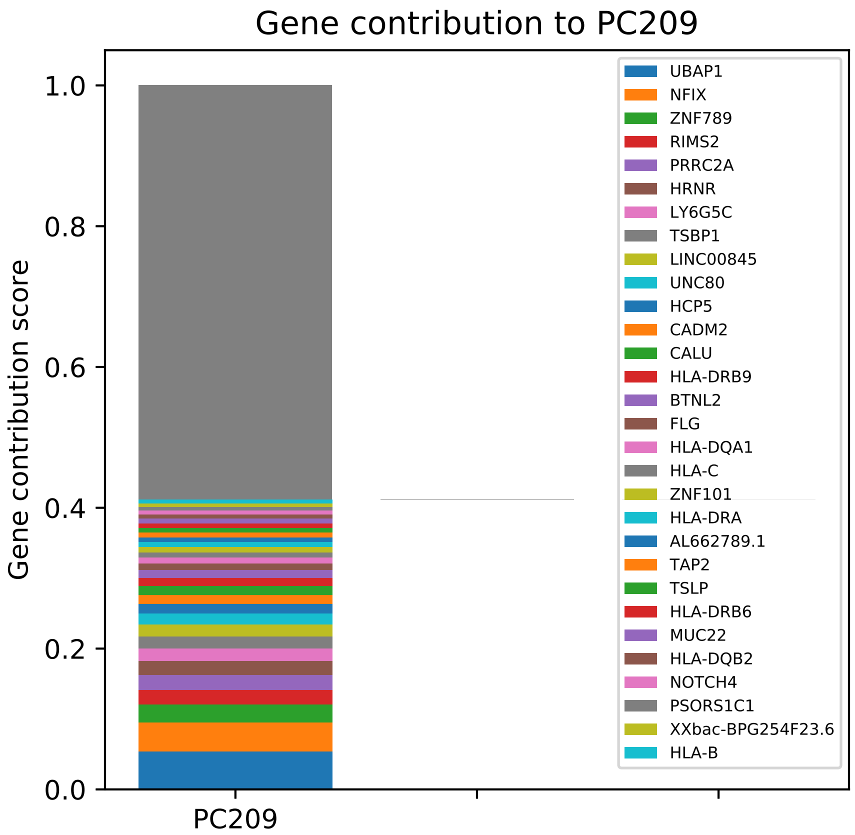

Gene contribution to PC209
Gene contribution score
1.0
0.8
0.6
0.4
0.2
0.0
PC209
UBAP1
NFIX
ZNF789
RIMS2
PRRC2A
HRNR
LY6G5C
TSBP1
LINC00845
UNC80
HCP5
CADM2
CALU
HLA-DRB9
BTNL2
FLG
HLA-DQA1
HLA-C
ZNF101
HLA-DRA
AL662789.1
TAP2
TSLP
HLA-DRB6
MUC22
HLA-DQB2
NOTCH4
PSORS1C1
XXbac-BPG254F23.6
HLA-B

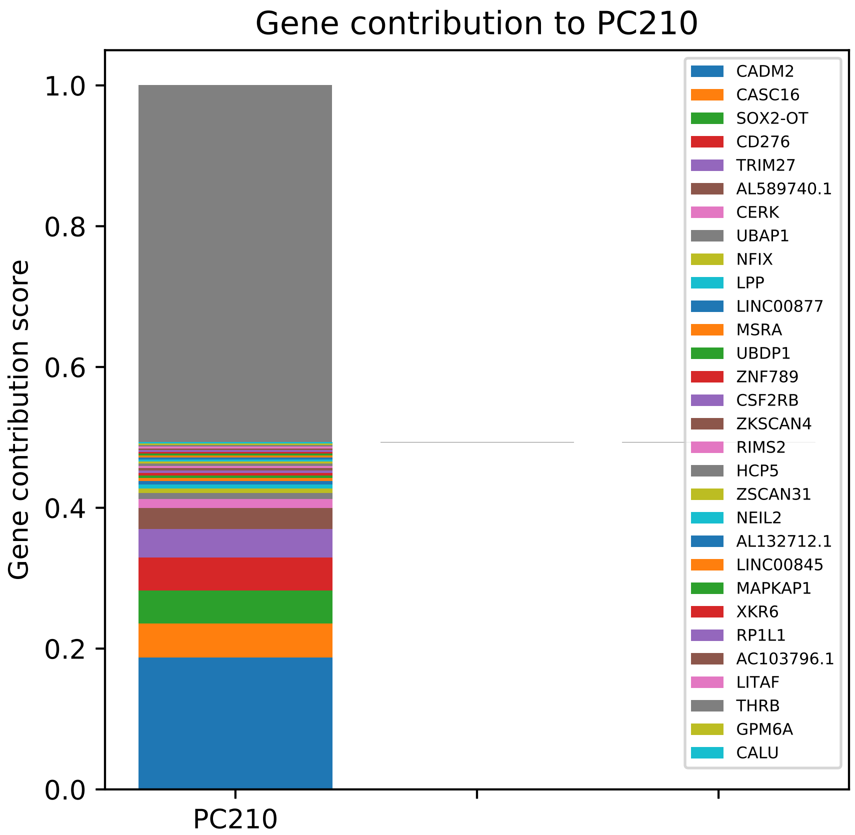
Gene contribution to PC210
Gene contribution score
1.0
0.8
0.6
0.4
0.2
0.0
PC210
CADM2
CASC16
SOX2-OT
CD276
TRIM27
AL589740.1
CERK
UBAP1
NFIX
LPP
LINC00877
MSRA
UBDP1
ZNF789
CSF2RB
ZKSCAN4
RIMS2
HCP5
ZSCAN31
NEIL2
AL132712.1
LINC00845
MAPKAP1
XKR6
RP1L1
AC103796.1
LITAF
THRB
GPM6A
CALU

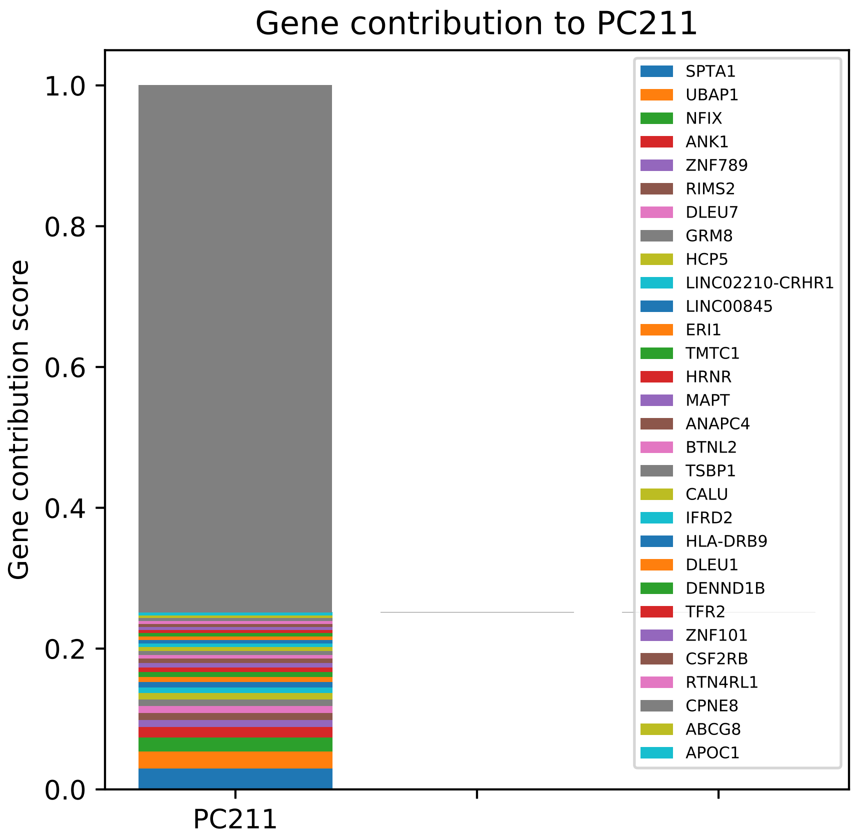
Gene contribution to PC211
Gene contribution score
PC211
0.0
0.2
0.4
0.6
0.8
1.0
SPTA1
UBAP1
NFIX
ANK1
ZNF789
RIMS2
DLEU7
GRM8
HCP5
LINC02210-CRHR1
LINC00845
ERI1
TMTC1
HRNR
MAPT
ANAPC4
BTNL2
TSBP1
CALU
IFRD2
HLA-DRB9
DLEU1
DENND1B
TFR2
ZNF101
CSF2RB
RTN4RL1
CPNE8
ABCG8
APOC1

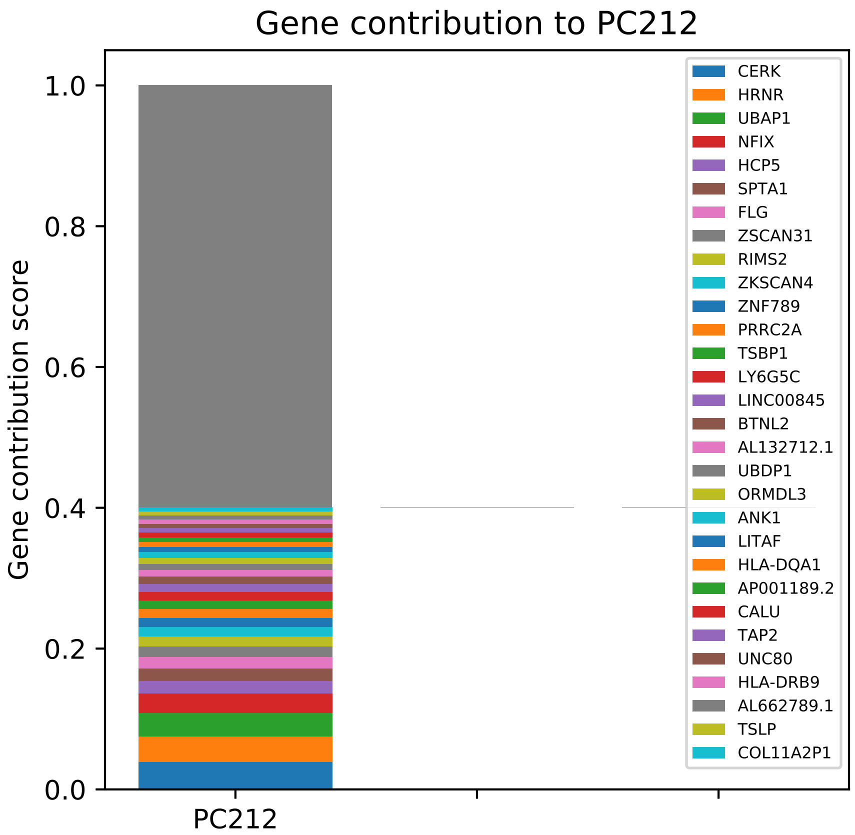

Gene contribution to PC212
Gene contribution score
1.0
0.8
0.6
0.4
0.2
0.0
PC212
CERK
HRNR
UBAP1
NFIX
HCP5
SPTA1
FLG
ZSCAN31
RIMS2
ZKSCAN4
ZNF789
PRRC2A
TSBP1
LY6G5C
LINC00845
BTNL2
AL132712.1
UBDP1
ORMDL3
ANK1
LITAF
HLA-DQA1
AP001189.2
CALU
TAP2
UNC80
HLA-DRB9
AL662789.1
TSLP
COL11A2P1

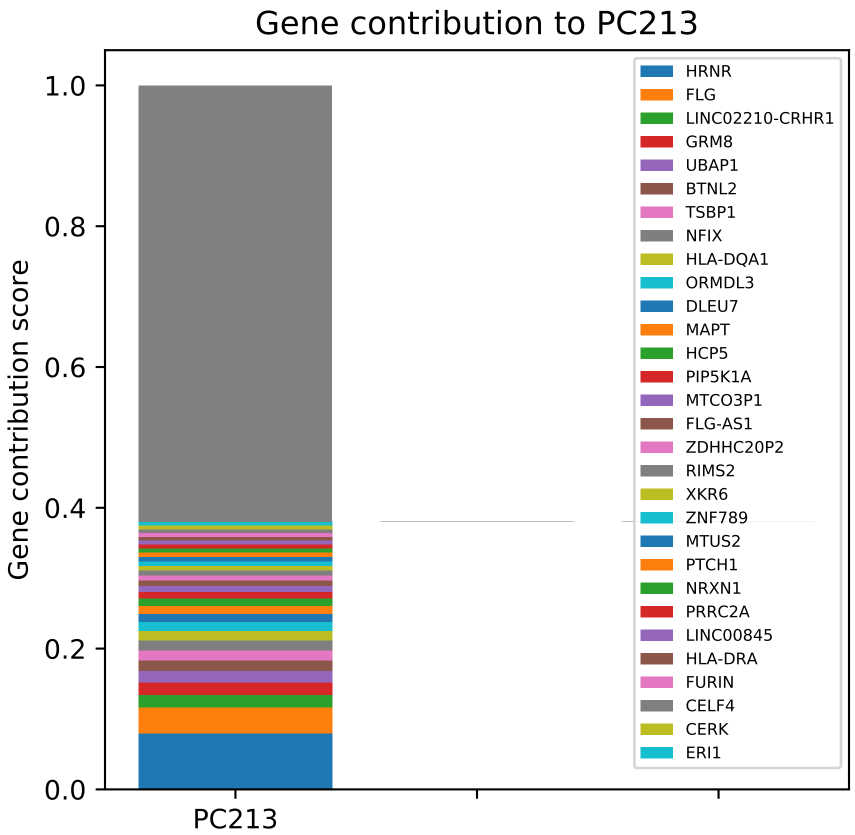

Gene contribution to PC213
Gene contribution score
1.0
0.8
0.6
0.4
0.2
0.0
PC213
HRNR
FLG
LINC02210-CRHR1
GRM8
UBAP1
BTNL2
TSBP1
NFIX
HLA-DQA1
ORMDL3
DLEU7
MAPT
HCP5
PIP5K1A
MTCO3P1
FLG-AS1
ZDHHC20P2
RIMS2
XKR6
ZNF789
MTUS2
PTCH1
NRXN1
PRRC2A
LINC00845
HLA-DRA
FURIN
CELF4
CERK
ERI1

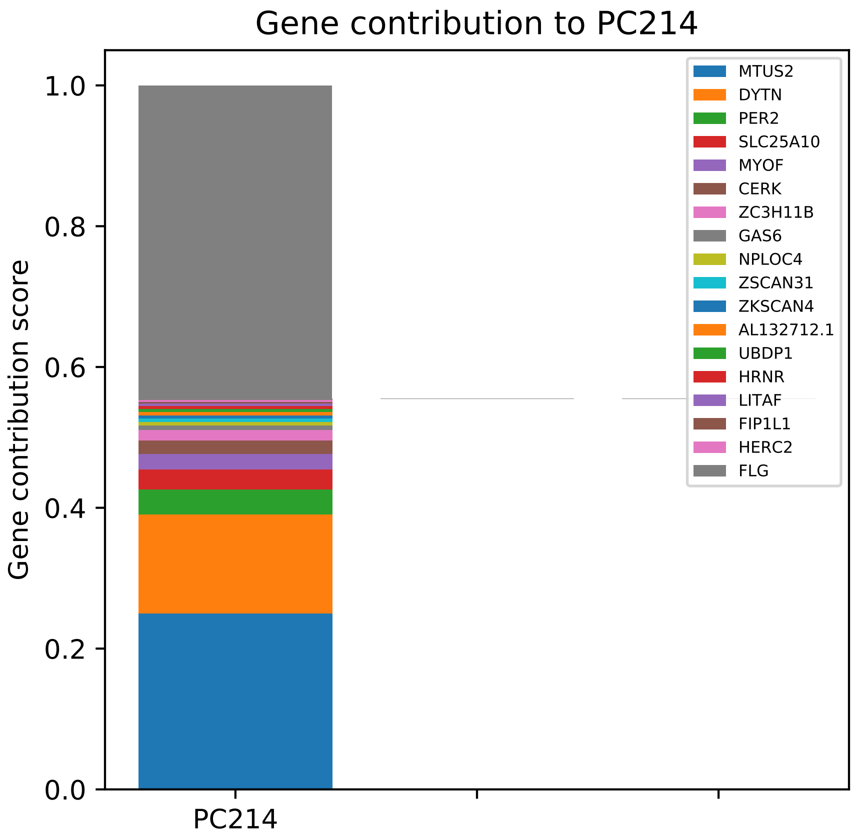

Gene contribution to PC214
Gene contribution score
1.0
0.8
0.6
0.4
0.2
0.0
PC214
MTUS2
DYTN
PER2
SLC25A10
MYOF
CERK
ZC3H11B
GAS6
NPLOC4
ZSCAN31
ZKSCAN4
AL132712.1
UBDP1
HRNR
LITAF
FIP1L1
HERC2
FLG

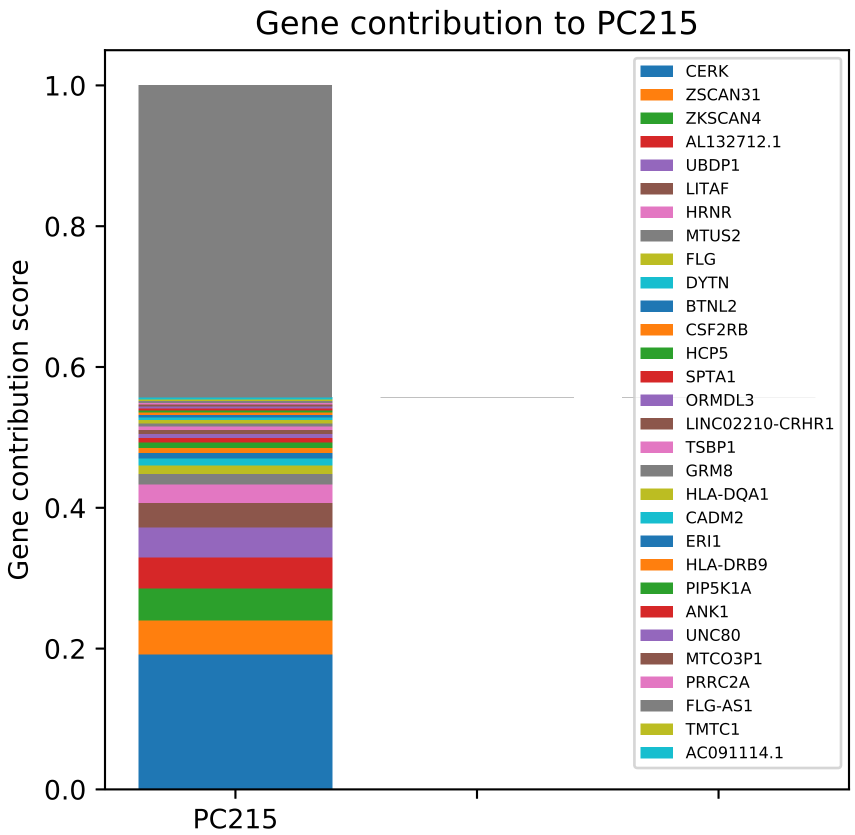
Gene contribution to PC215
Gene contribution score
1.0
0.8
0.6
0.4
0.2
0.0
PC215
CERK
ZSCAN31
ZKSCAN4
AL132712.1
UBDP1
LITAF
HRNR
MTUS2
FLG
DYTN
BTNL2
CSF2RB
HCP5
SPTA1
ORMDL3
LINC02210-CRHR1
TSBP1
GRM8
HLA-DQA1
CADM2
ERI1
HLA-DRB9
PIP5K1A
ANK1
UNC80
MTCO3P1
PRRC2A
FLG-AS1
TMTC1
AC091114.1

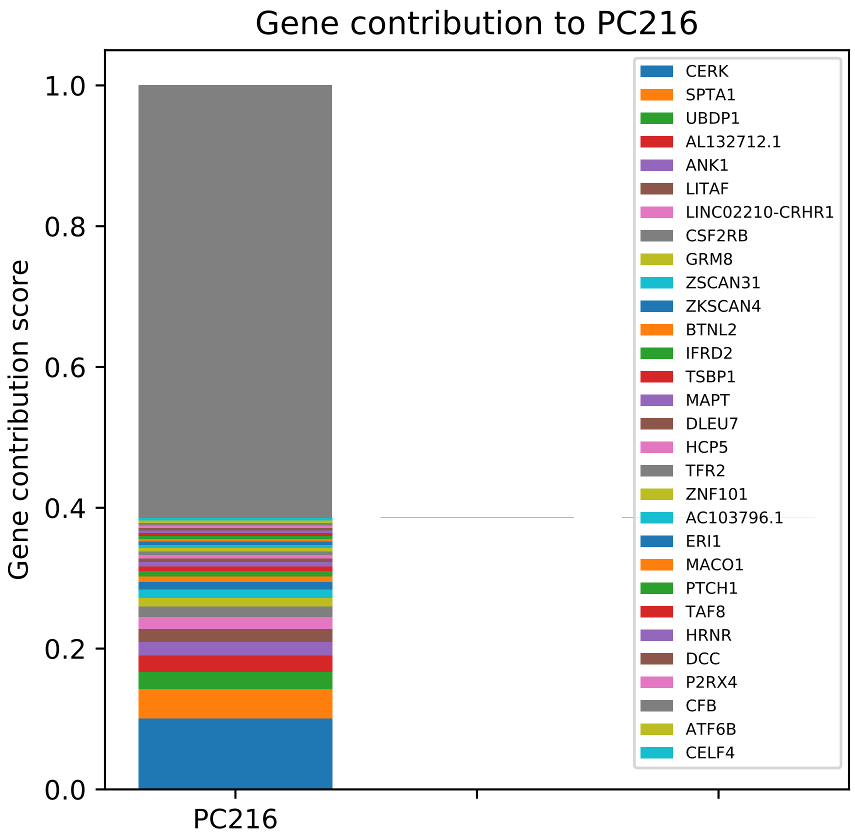

Gene contribution to PC216
Gene contribution score
1.0
0.8
0.6
0.4
0.2
0.0
PC216
CERK
SPTA1
UBDP1
AL132712.1
ANK1
LITAF
LINC02210-CRHR1
CSF2RB
GRM8
ZSCAN31
ZKSCAN4
BTNL2
IFRD2
TSBP1
MAPT
DLEU7
HCP5
TFR2
ZNF101
AC103796.1
ERI1
MACO1
PTCH1
TAF8
HRNR
DCC
P2RX4
CFB
ATF6B
CELF4

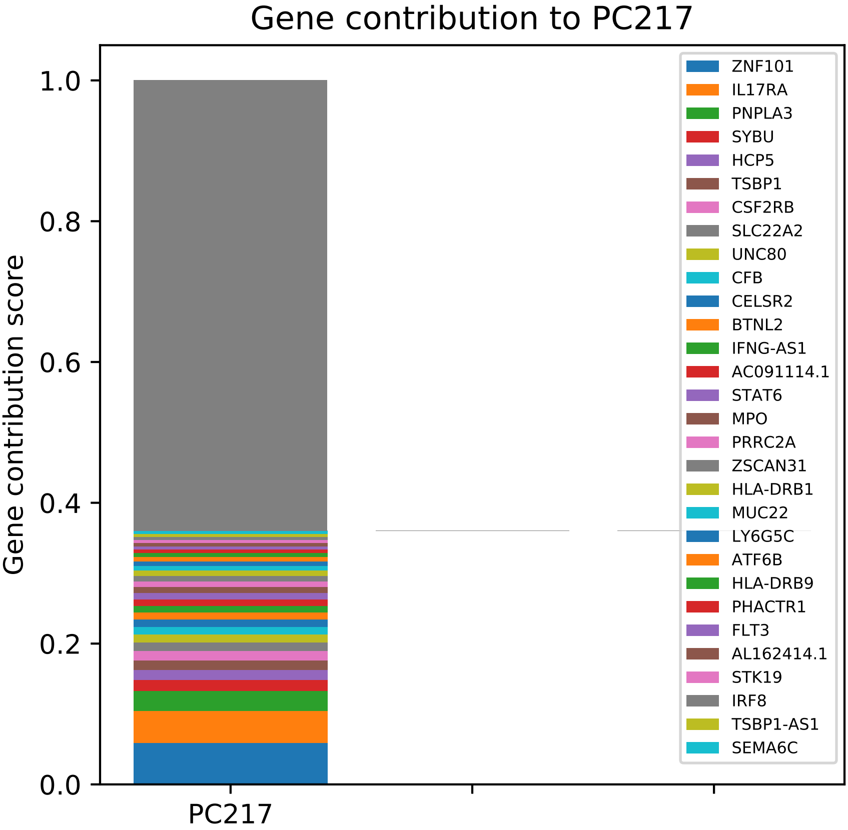

Gene contribution to PC217
Gene contribution score
1.0
0.8
0.6
0.4
0.2
0.0
PC217
ZNF101
IL17RA
PNPLA3
SYBU
HCP5
TSBP1
CSF2RB
SLC22A2
UNC80
CFB
CELSR2
BTNL2
IFNG-AS1
AC091114.1
STAT6
MPO
PRRC2A
ZSCAN31
HLA-DRB1
MUC22
LY6G5C
ATF6B
HLA-DRB9
PHACTR1
FLT3
AL162414.1
STK19
IRF8
TSBP1-AS1
SEMA6C

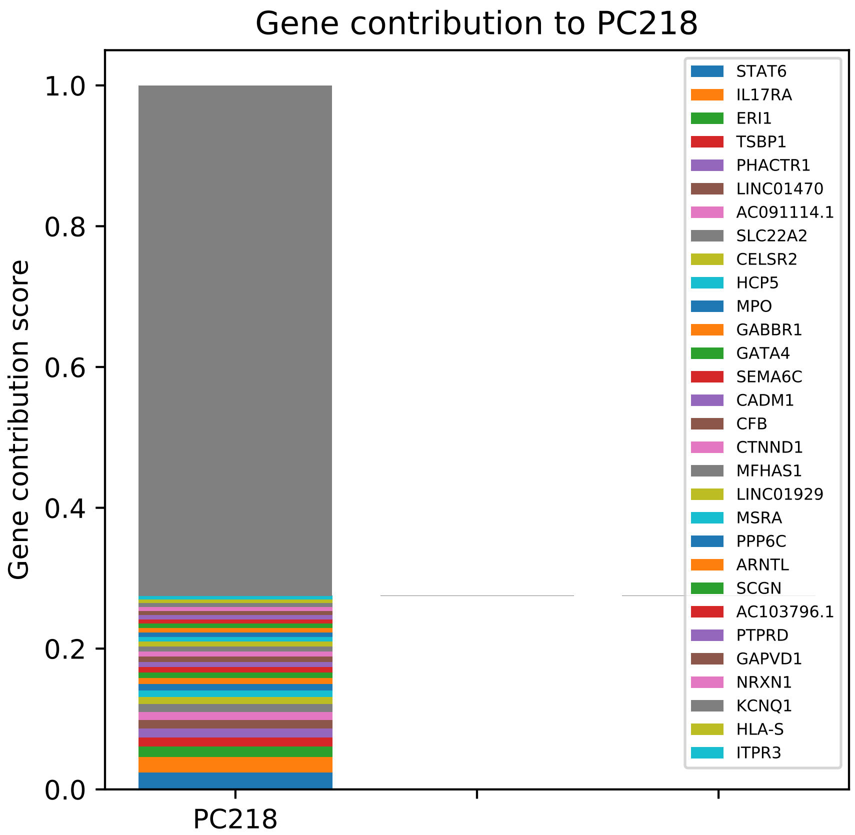

Gene contribution to PC218
Gene contribution score
1.0
0.8
0.6
0.4
0.2
0.0
PC218
STAT6
IL17RA
ERI1
TSBP1
PHACTR1
LINC01470
AC091114.1
SLC22A2
CELSR2
HCP5
MPO
GABBR1
GATA4
SEMA6C
CADM1
CFB
CTNND1
MFHAS1
LINC01929
MSRA
PPP6C
ARNTL
SCGN
AC103796.1
PTPRD
GAPVD1
NRXN1
KCNQ1
HLA-S
ITPR3

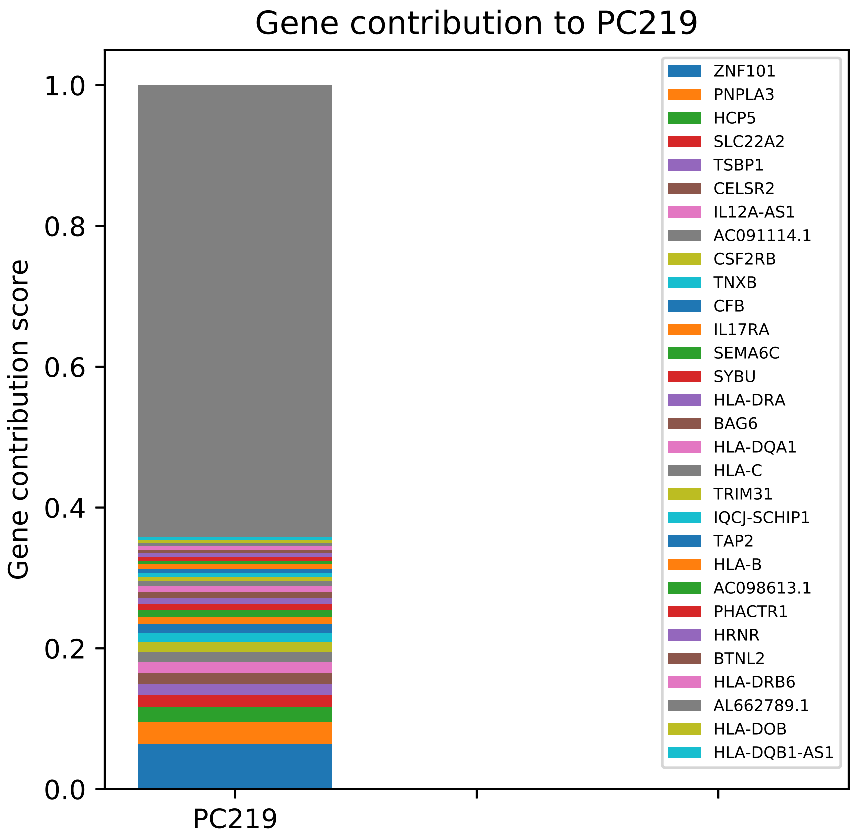

Gene contribution to PC219
Gene contribution score
1.0
0.8
0.6
0.4
0.2
0.0
PC219
ZNF101
PNPLA3
HCP5
SLC22A2
TSBP1
CELSR2
IL12A-AS1
AC091114.1
CSF2RB
TNXB
CFB
IL17RA
SEMA6C
SYBU
HLA-DRA
BAG6
HLA-DQA1
HLA-C
TRIM31
IQCJ-SCHIP1
TAP2
HLA-B
AC098613.1
PHACTR1
HRNR
BTNL2
HLA-DRB6
AL662789.1
HLA-DOB
HLA-DQB1-AS1

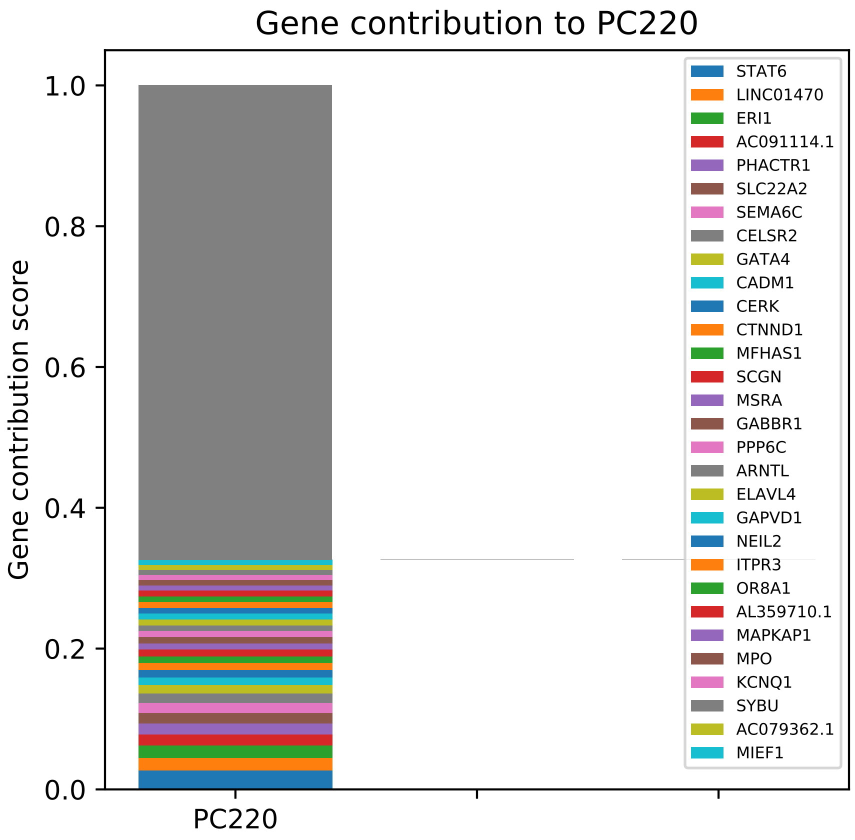
Gene contribution to PC220
Gene contribution score
1.0
0.8
0.6
0.4
0.2
0.0
PC220
STAT6
LINC01470
ERI1
AC091114.1
PHACTR1
SLC22A2
SEMA6C
CELSR2
GATA4
CADM1
CERK
CTNND1
MFHAS1
SCGN
MSRA
GABBR1
PPP6C
ARNTL
ELAVL4
GAPVD1
NEIL2
ITPR3
OR8A1
AL359710.1
MAPKAP1
MPO
KCNQ1
SYBU
AC079362.1
MIEF1

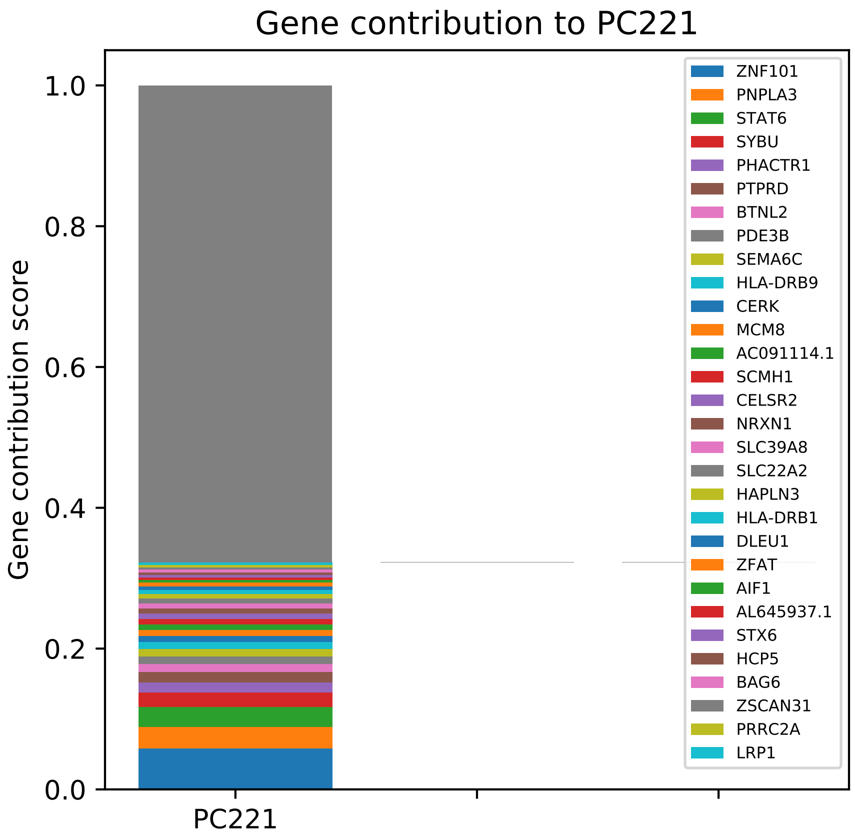

Gene contribution to PC221
Gene contribution score
1.0
0.8
0.6
0.4
0.2
0.0
PC221
ZNF101
PNPLA3
STAT6
SYBU
PHACTR1
PTPRD
BTNL2
PDE3B
SEMA6C
HLA-DRB9
CERK
MCM8
AC091114.1
SCMH1
CELSR2
NRXN1
SLC39A8
SLC22A2
HAPLN3
HLA-DRB1
DLEU1
ZFAT
AIF1
AL645937.1
STX6
HCP5
BAG6
ZSCAN31
PRRC2A
LRP1

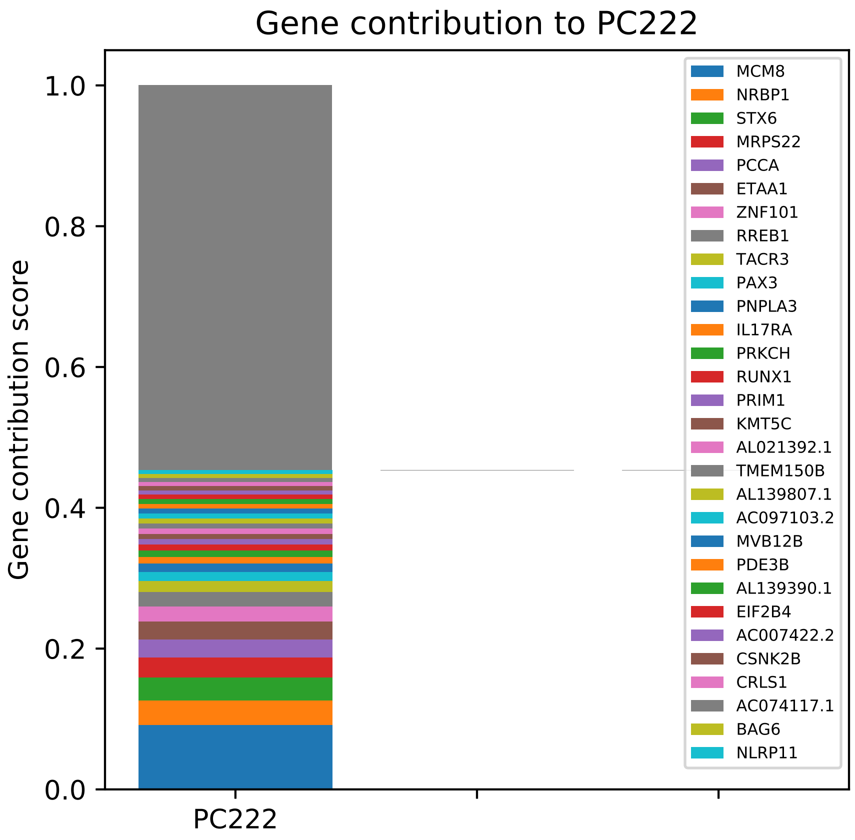
Gene contribution to PC222
Gene contribution score
0.0
0.2
0.4
0.6
0.8
1.0
PC222
MCM8
NRBP1
STX6
MRPS22
PCCA
ETAA1
ZNF101
RREB1
TACR3
PAX3
PNPLA3
IL17RA
PRKCH
RUNX1
PRIM1
KMT5C
AL021392.1
TMEM150B
AL139807.1
AC097103.2
MVB12B
PDE3B
AL139390.1
EIF2B4
AC007422.2
CSNK2B
CRLS1
AC074117.1
BAG6
NLRP11

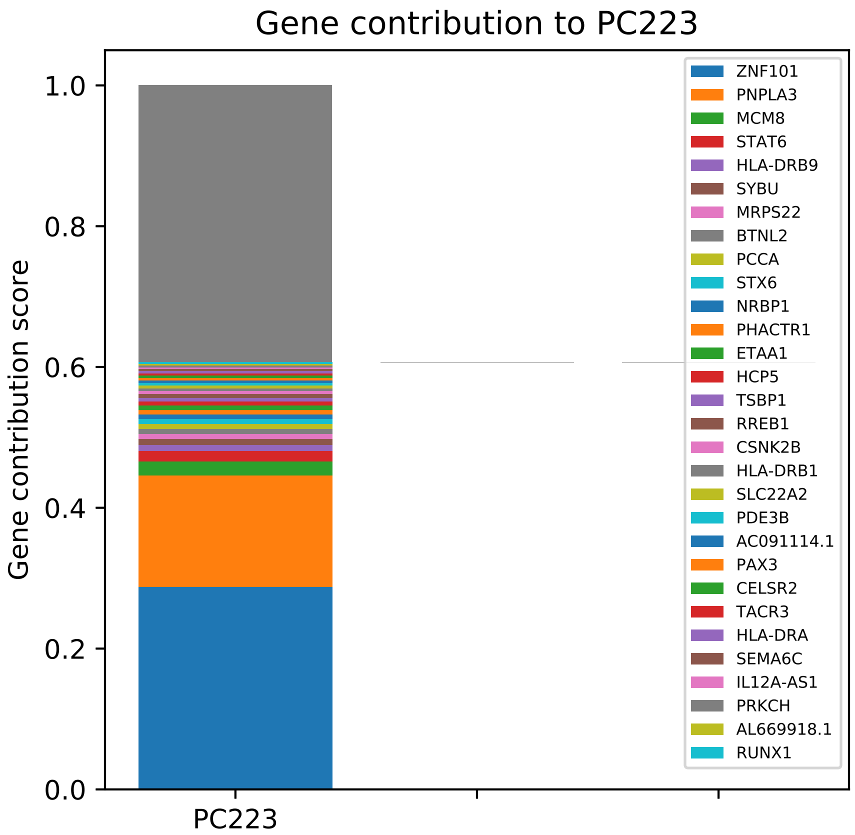

Gene contribution to PC223
Gene contribution score
1.0
0.8
0.6
0.4
0.2
0.0
PC223
ZNF101
PNPLA3
MCM8
STAT6
HLA-DRB9
SYBU
MRPS22
BTNL2
PCCA
STX6
NRBP1
PHACTR1
ETAA1
HCP5
TSBP1
RREB1
CSNK2B
HLA-DRB1
SLC22A2
PDE3B
AC091114.1
PAX3
CELSR2
TACR3
HLA-DRA
SEMA6C
IL12A-AS1
PRKCH
AL669918.1
RUNX1

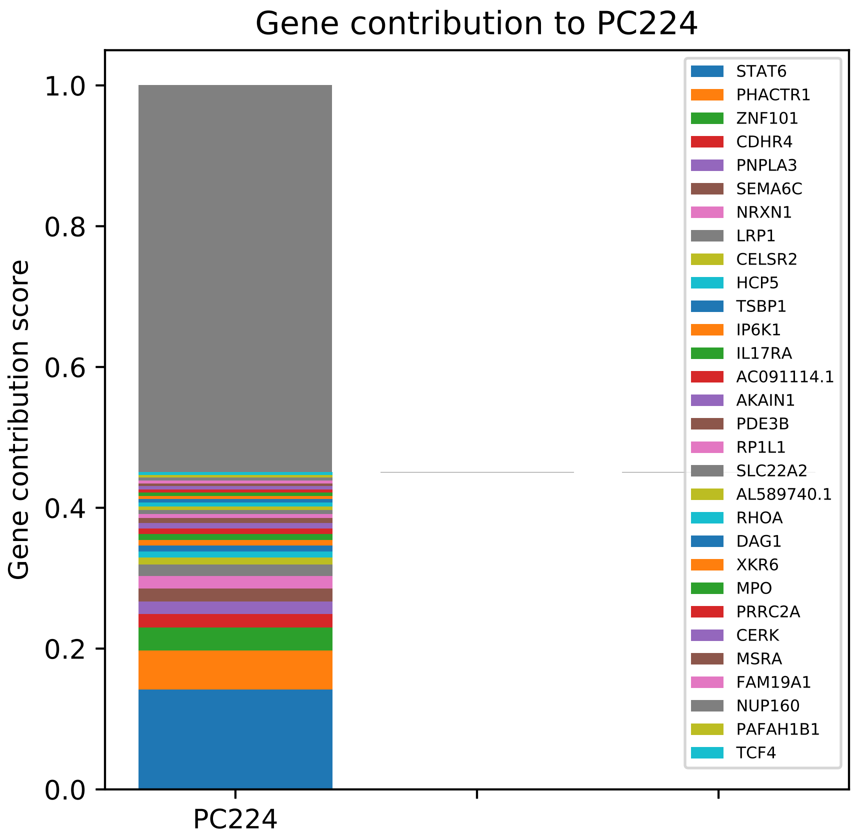

Gene contribution to PC224
Gene contribution score
1.0
0.8
0.6
0.4
0.2
0.0
PC224
STAT6
PHACTR1
ZNF101
CDHR4
PNPLA3
SEMA6C
NRXN1
LRP1
CELSR2
HCP5
TSBP1
IP6K1
IL17RA
AC091114.1
AKAIN1
PDE3B
RP1L1
SLC22A2
AL589740.1
RHOA
DAG1
XKR6
MPO
PRRC2A
CERK
MSRA
FAM19A1
NUP160
PAFAH1B1
TCF4

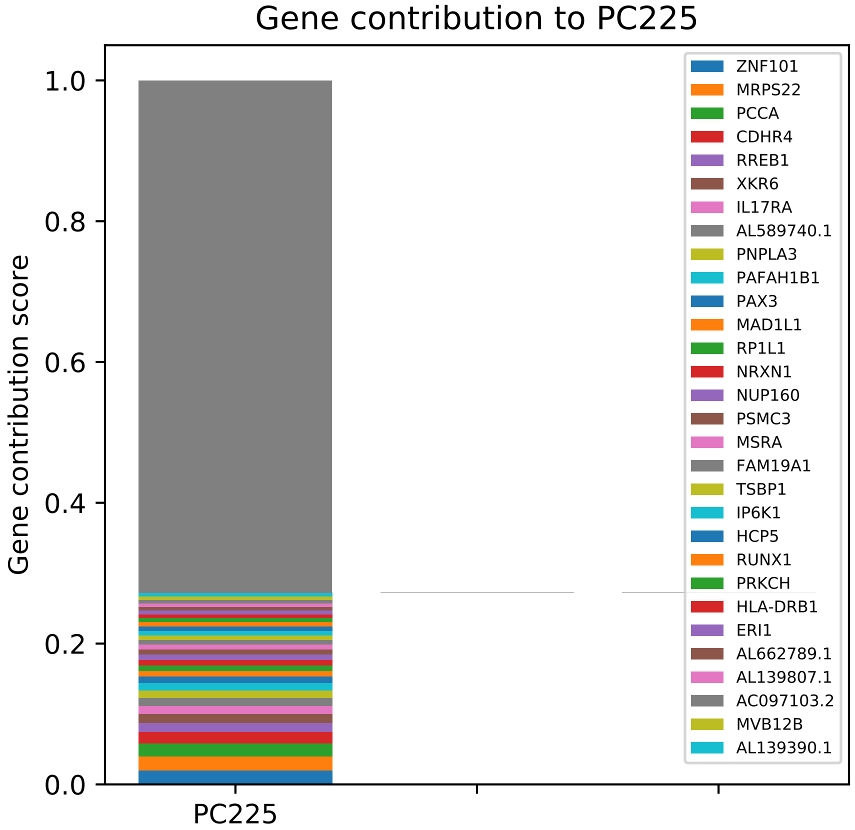

Gene contribution to PC225
Gene contribution score
1.0
0.8
0.6
0.4
0.2
0.0
PC225
ZNF101
MRPS22
PCCA
CDHR4
RREB1
XKR6
IL17RA
AL589740.1
PNPLA3
PAFAH1B1
PAX3
MAD1L1
RP1L1
NRXN1
NUP160
PSMC3
MSRA
FAM19A1
TSBP1
IP6K1
HCP5
RUNX1
PRKCH
HLA-DRB1
ERI1
AL662789.1
AL139807.1
AC097103.2
MVB12B
AL139390.1

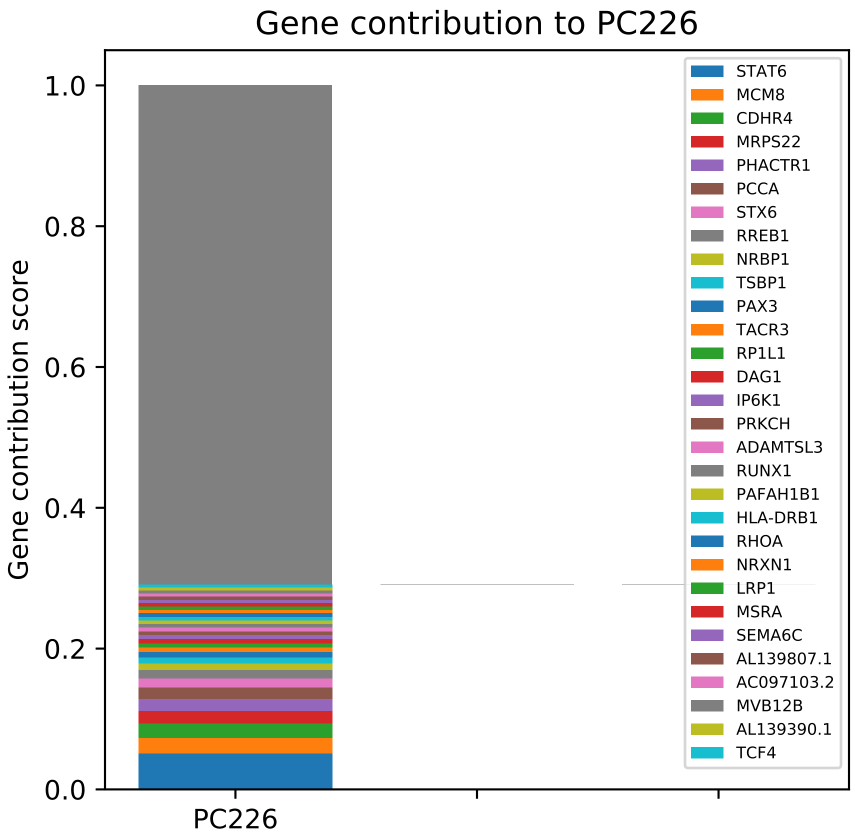

Gene contribution to PC226
Gene contribution score
1.0
0.8
0.6
0.4
0.2
0.0
PC226
STAT6
MCM8
CDHR4
MRPS22
PHACTR1
PCCA
STX6
RREB1
NRBP1
TSBP1
PAX3
TACR3
RP1L1
DAG1
IP6K1
PRKCH
ADAMTSL3
RUNX1
PAFAH1B1
HLA-DRB1
RHOA
NRXN1
LRP1
MSRA
SEMA6C
AL139807.1
AC097103.2
MVB12B
AL139390.1
TCF4

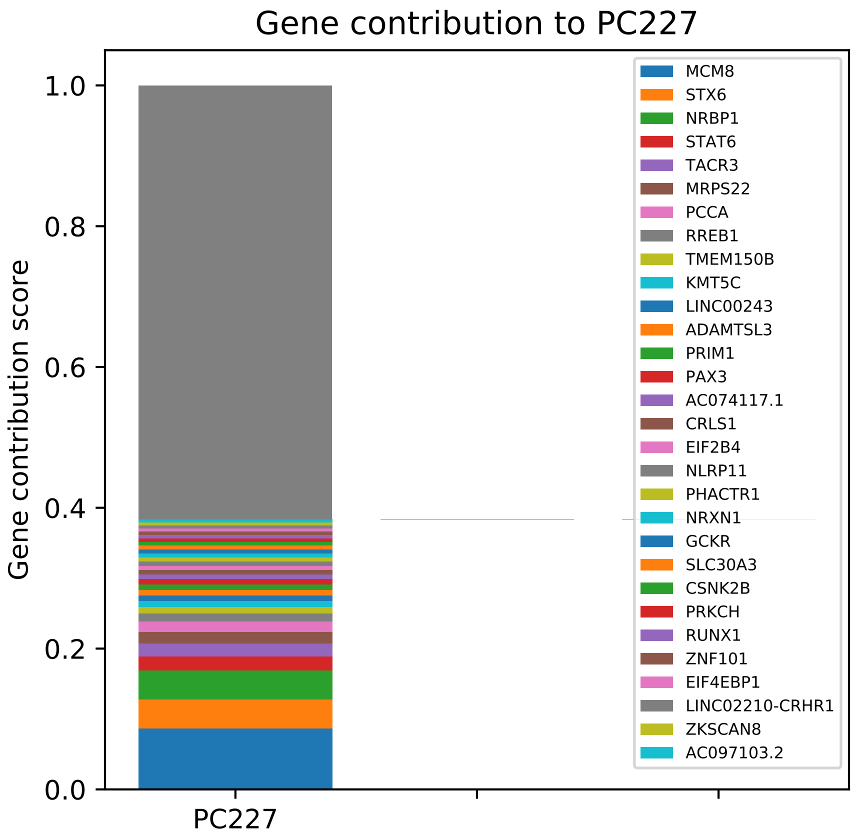
Gene contribution to PC227
Gene contribution score
1.0
0.8
0.6
0.4
0.2
0.0
PC227
MCM8
STX6
NRBP1
STAT6
TACR3
MRPS22
PCCA
RREB1
TMEM150B
KMT5C
LINC00243
ADAMTSL3
PRIM1
PAX3
AC074117.1
CRLS1
EIF2B4
NLRP11
PHACTR1
NRXN1
GCKR
SLC30A3
CSNK2B
PRKCH
RUNX1
ZNF101
EIF4EBP1
LINC02210-CRHR1
ZKSCAN8
AC097103.2

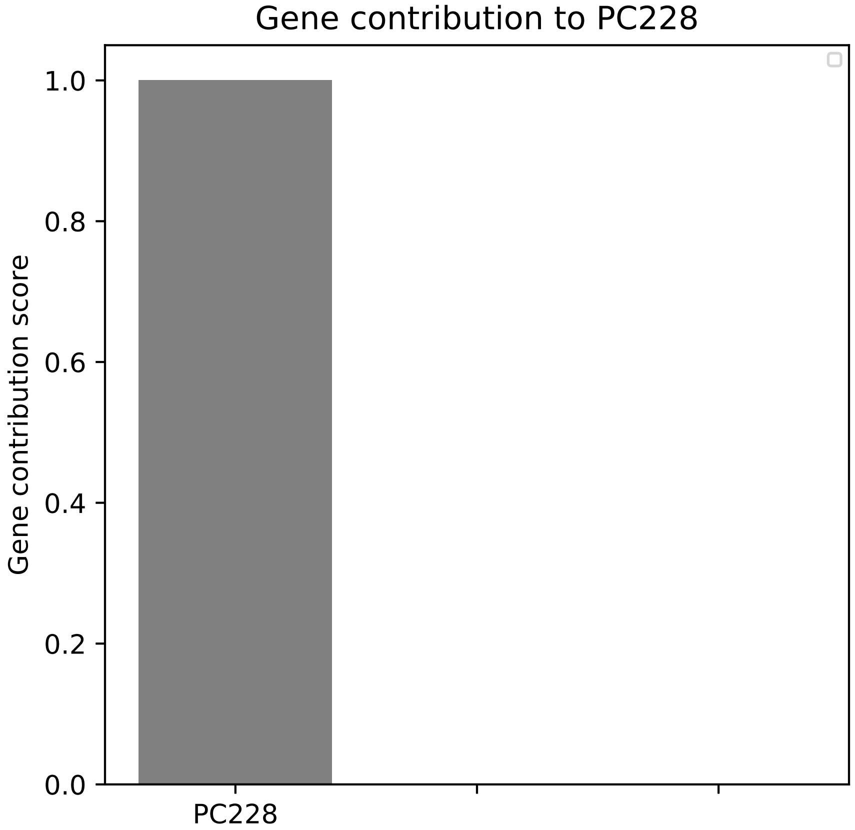
Gene contribution to PC228
Gene contribution score
1.0
0.8
0.6
0.4
0.2
0.0
PC228

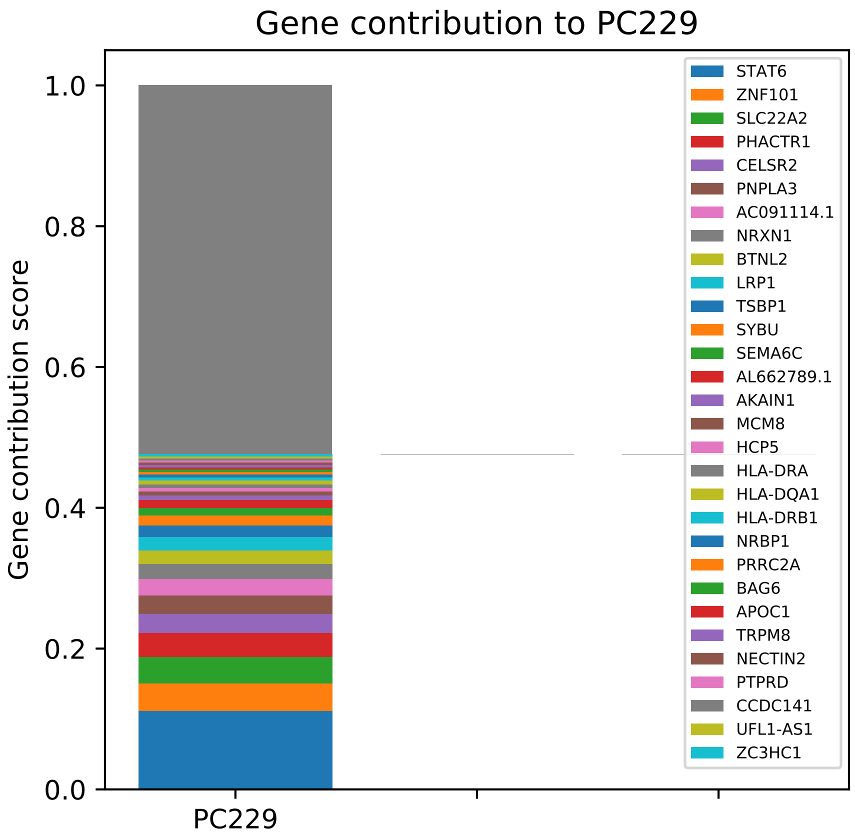

Gene contribution to PC229
Gene contribution score
1.0
0.8
0.6
0.4
0.2
0.0
PC229
STAT6
ZNF101
SLC22A2
PHACTR1
CELSR2
PNPLA3
AC091114.1
NRXN1
BTNL2
LRP1
TSBP1
SYBU
SEMA6C
AL662789.1
AKAIN1
MCM8
HCP5
HLA-DRA
HLA-DQA1
HLA-DRB1
NRBP1
PRRC2A
BAG6
APOC1
TRPM8
NECTIN2
PTPRD
CCDC141
UFL1-AS1
ZC3HC1

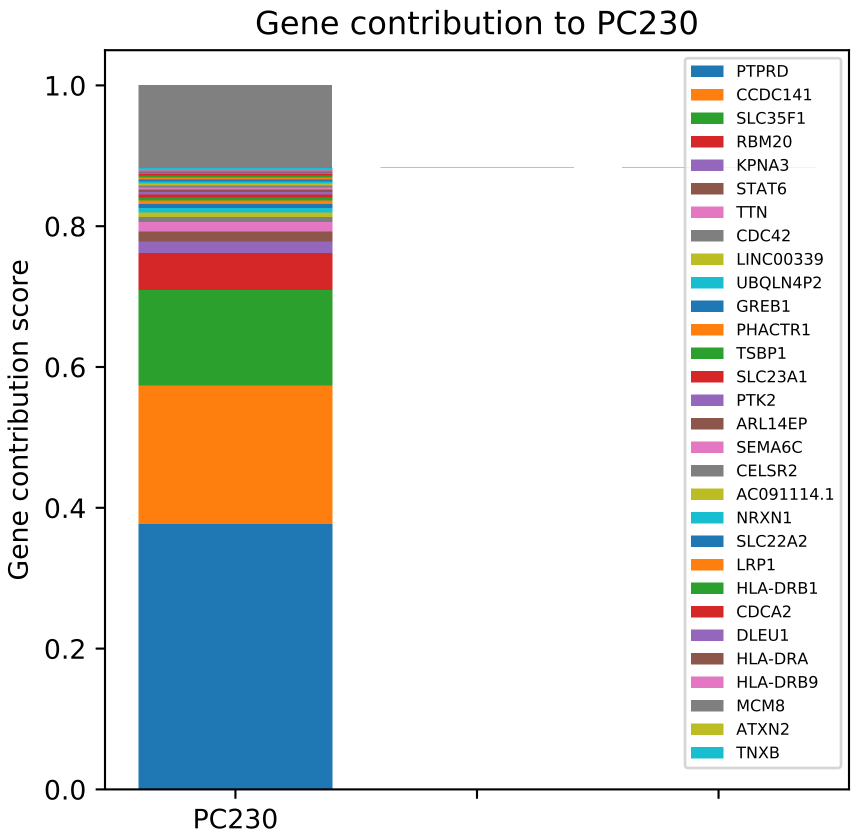
Gene contribution to PC230
Gene contribution score
0.0
0.2
0.4
0.6
0.8
1.0
PC230
PTPRD
CCDC141
SLC35F1
RBM20
KPNA3
STAT6
TTN
CDC42
LINC00339
UBQLN4P2
GREB1
PHACTR1
TSBP1
SLC23A1
PTK2
ARL14EP
SEMA6C
CELSR2
AC091114.1
NRXN1
SLC22A2
LRP1
HLA-DRB1
CDCA2
DLEU1
HLA-DRA
HLA-DRB9
MCM8
ATXN2
TNXB

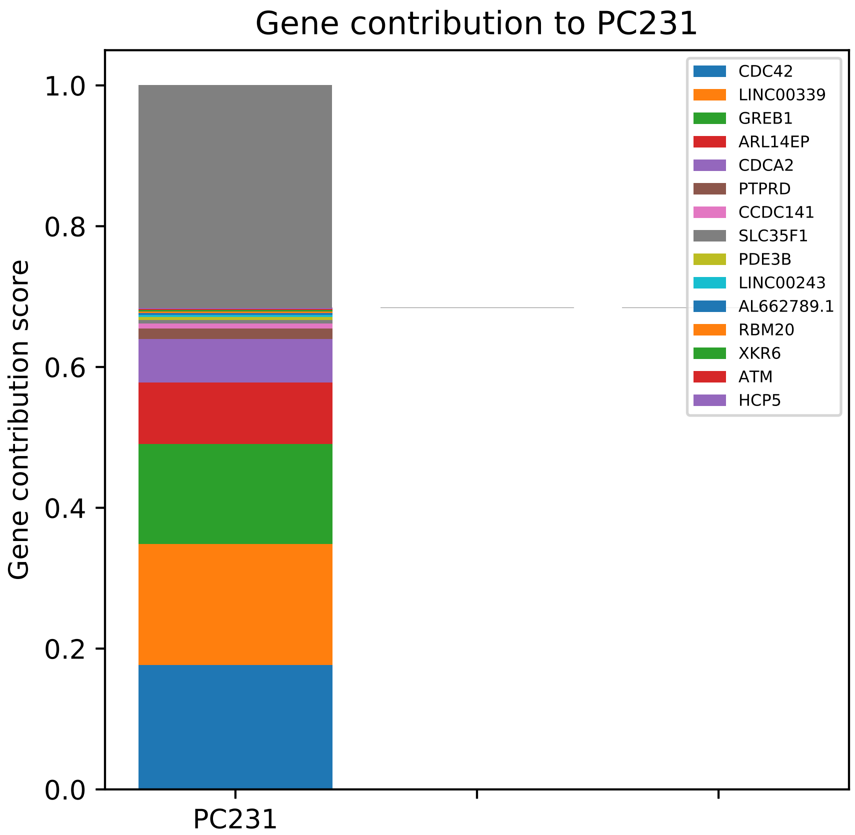

Gene contribution to PC231
Gene contribution score
1.0
0.8
0.6
0.4
0.2
0.0
PC231
CDC42
LINC00339
GREB1
ARL14EP
CDCA2
PTPRD
CCDC141
SLC35F1
PDE3B
LINC00243
AL662789.1
RBM20
XKR6
ATM
HCP5

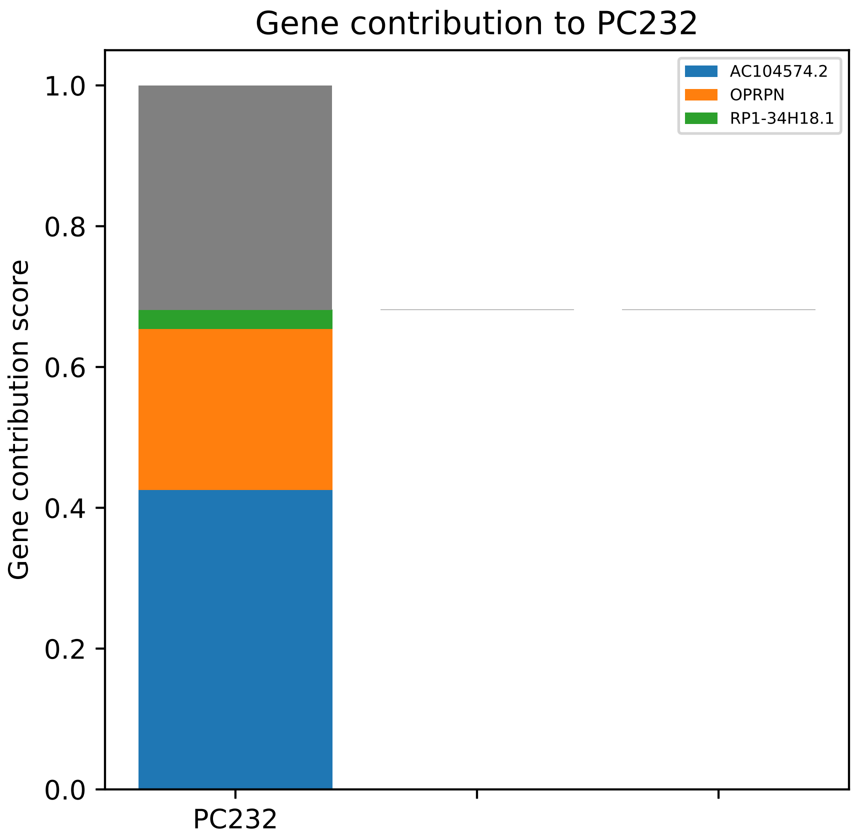

Gene contribution to PC232
AC104574.2
OPRPN
RP1-34H18.1
1.0
0.8
0.6
0.4
0.2
0.0
Gene contribution score
PC232

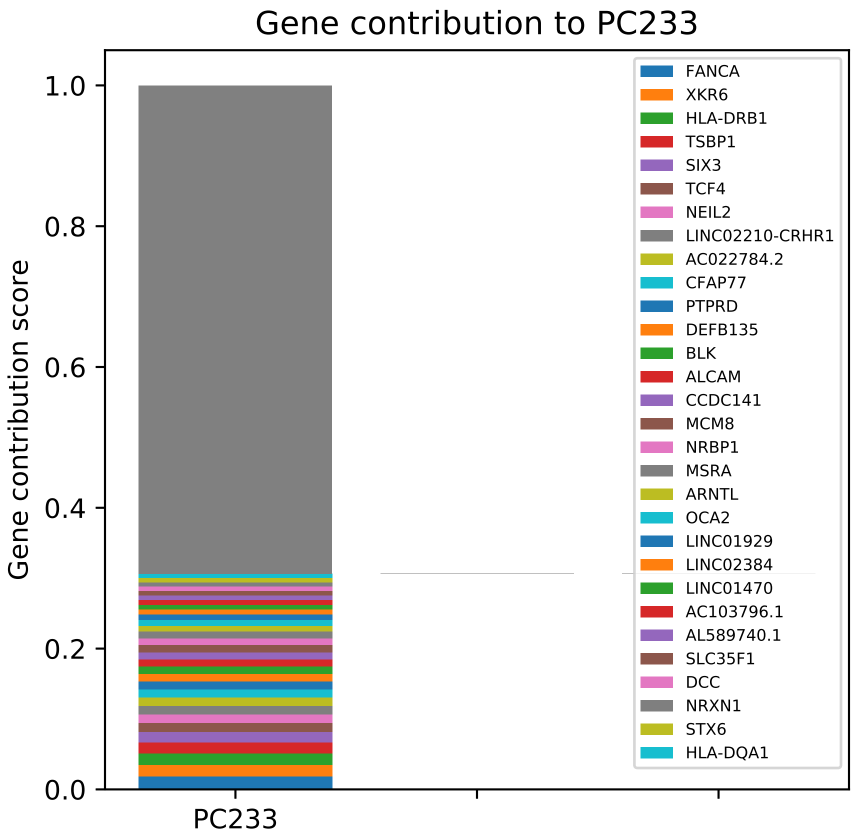

Gene contribution to PC233
Gene contribution score
1.0
0.8
0.6
0.4
0.2
0.0
PC233
FANCA
XKR6
HLA-DRB1
TSBP1
SIX3
TCF4
NEIL2
LINC02210-CRHR1
AC022784.2
CFAP77
PTPRD
DEFB135
BLK
ALCAM
CCDC141
MCM8
NRBP1
MSRA
ARNTL
OCA2
LINC01929
LINC02384
LINC01470
AC103796.1
AL589740.1
SLC35F1
DCC
NRXN1
STX6
HLA-DQA1

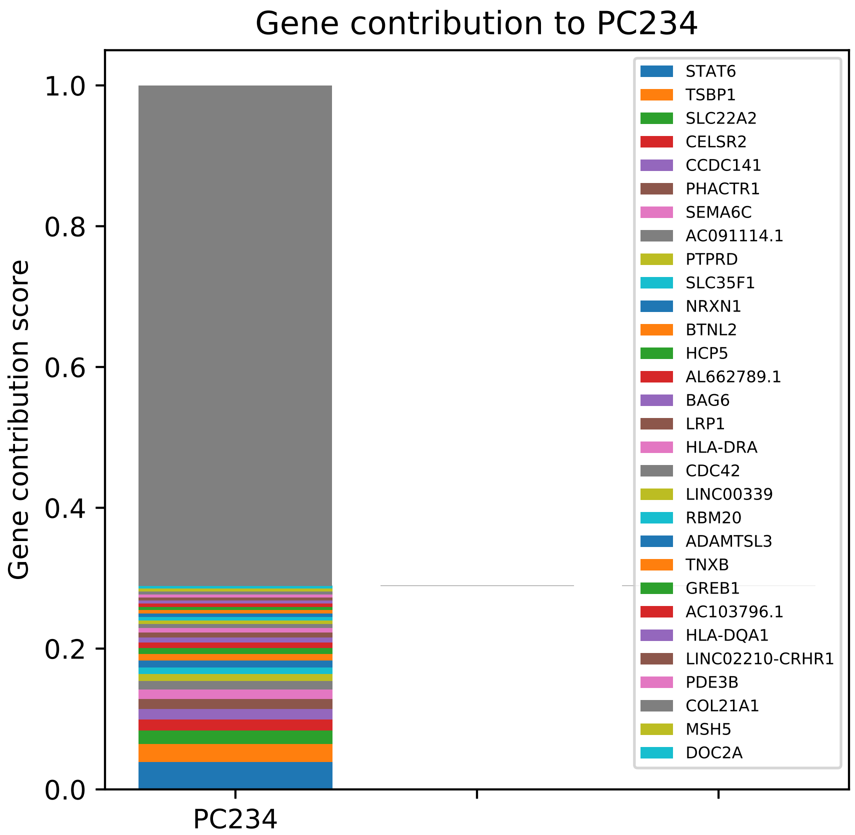

Gene contribution to PC234
Gene contribution score
1.0
0.8
0.6
0.4
0.2
0.0
PC234
STAT6
TSBP1
SLC22A2
CELSR2
CCDC141
PHACTR1
SEMA6C
AC091114.1
PTPRD
SLC35F1
NRXN1
BTNL2
HCP5
AL662789.1
BAG6
LRP1
HLA-DRA
CDC42
LINC00339
RBM20
ADAMTSL3
TNXB
GREB1
AC103796.1
HLA-DQA1
LINC02210-CRHR1
PDE3B
COL21A1
MSH5
DOC2A

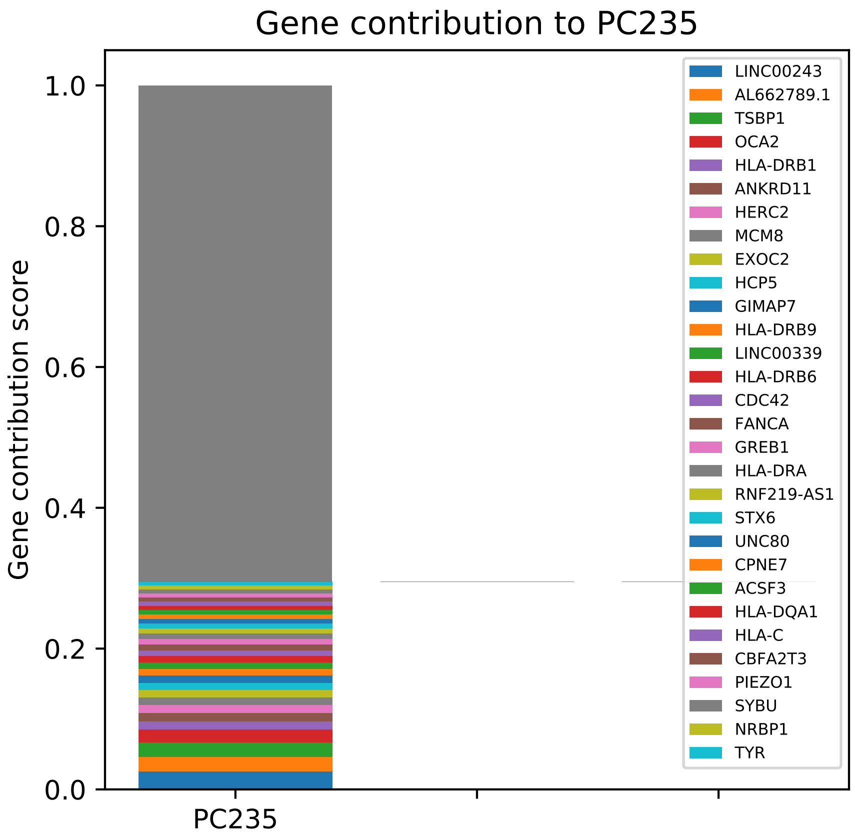

Gene contribution to PC235
Gene contribution score
1.0
0.8
0.6
0.4
0.2
0.0
PC235
LINC00243
AL662789.1
TSBP1
OCA2
HLA-DRB1
ANKRD11
HERC2
MCM8
EXOC2
HCP5
GIMAP7
HLA-DRB9
LINC00339
HLA-DRB6
CDC42
FANCA
GREB1
HLA-DRA
RNF219-AS1
STX6
UNC80
CPNE7
ACSF3
HLA-DQA1
HLA-C
CBFA2T3
PIEZO1
SYBU
NRBP1
TYR

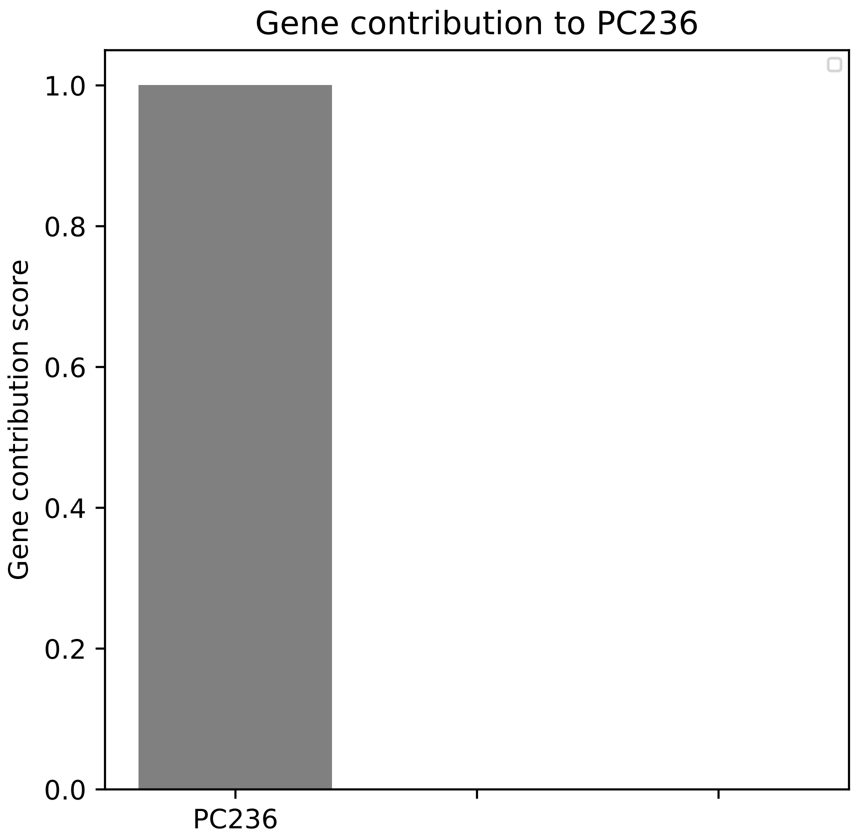
Gene contribution to PC236
Gene contribution score
1.0
0.8
0.6
0.4
0.2
0.0
PC236

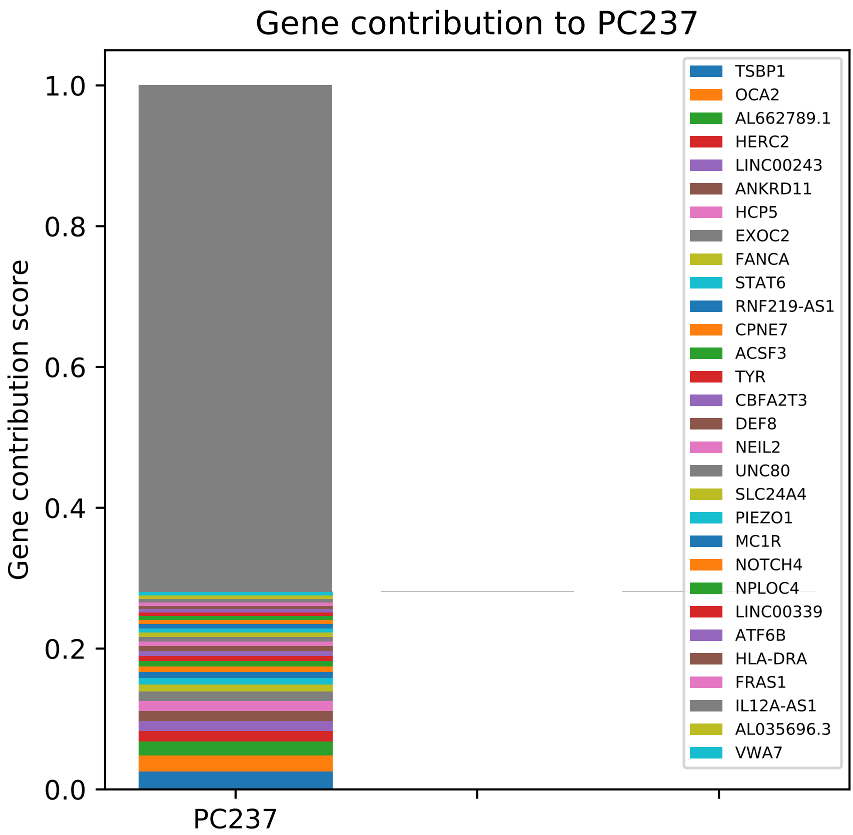

Gene contribution to PC237
Gene contribution score
1.0
0.8
0.6
0.4
0.2
0.0
PC237
TSBP1
OCA2
AL662789.1
HERC2
LINC00243
ANKRD11
HCP5
EXOC2
FANCA
STAT6
RNF219-AS1
CPNE7
ACSF3
TYR
CBFA2T3
DEF8
NEIL2
UNC80
SLC24A4
PIEZO1
MC1R
NOTCH4
NPLOC4
LINC00339
ATF6B
HLA-DRA
FRAS1
IL12A-AS1
AL035696.3
VWA7

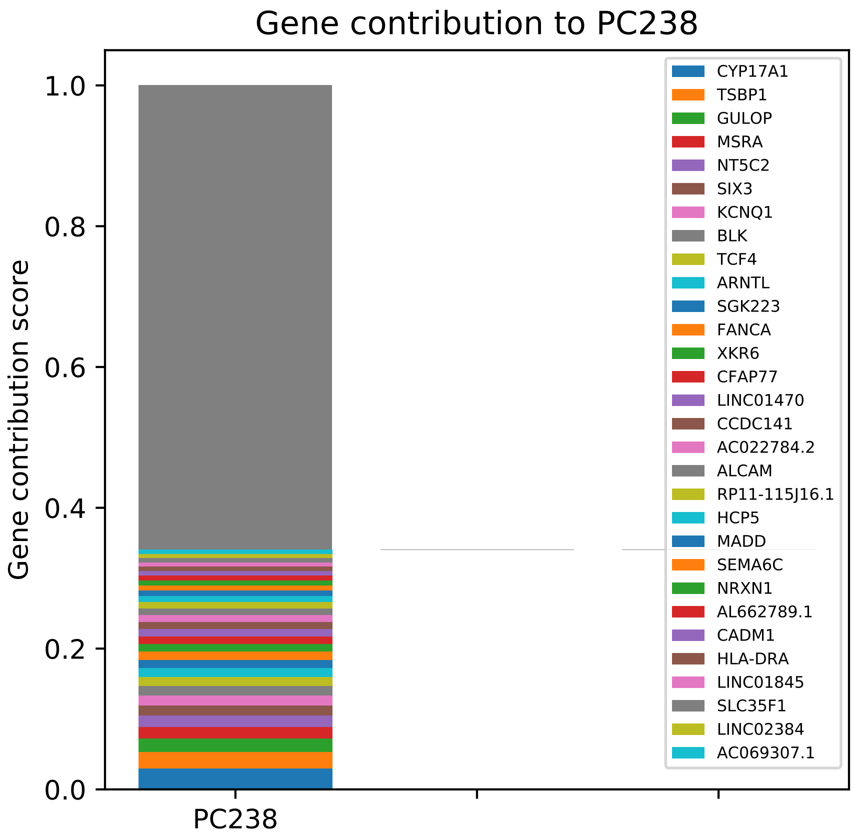

Gene contribution to PC238
Gene contribution score
1.0
0.8
0.6
0.4
0.2
0.0
PC238
CYP17A1
TSBP1
GULOP
MSRA
NT5C2
SIX3
KCNQ1
BLK
TCF4
ARNTL
SGK223
FANCA
XKR6
CFAP77
LINC01470
CCDC141
AC022784.2
ALCAM
RP11-115J16.1
HCP5
MADD
SEMA6C
NRXN1
AL662789.1
CADM1
HLA-DRA
LINC01845
SLC35F1
LINC02384
AC069307.1

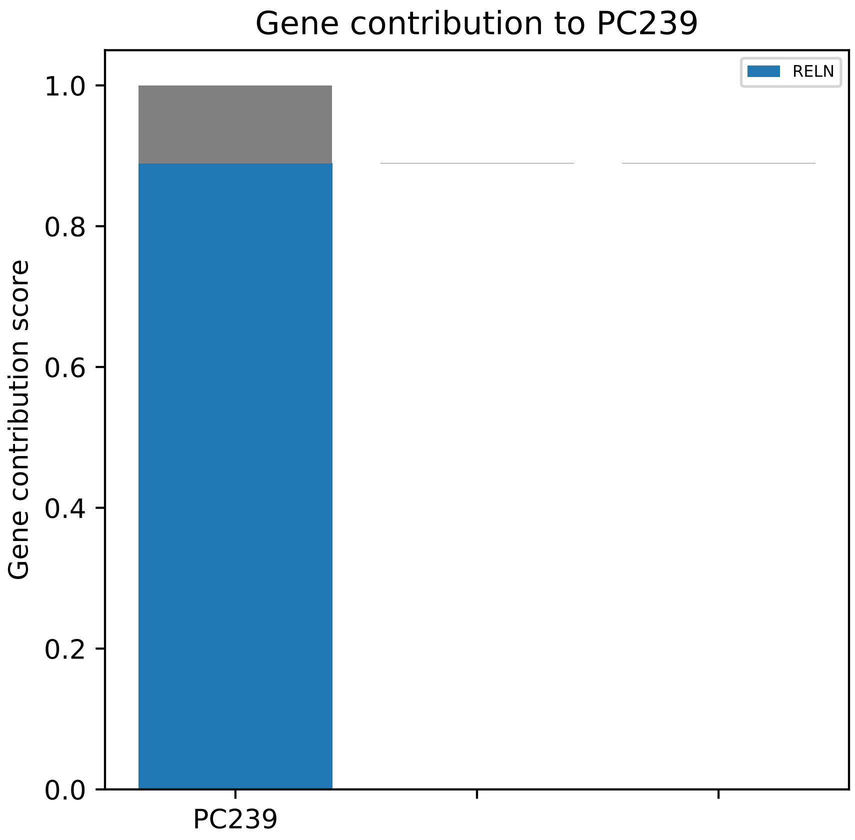

Gene contribution to PC239
RELN
Gene contribution score
1.0
0.8
0.6
0.4
0.2
0.0
PC239

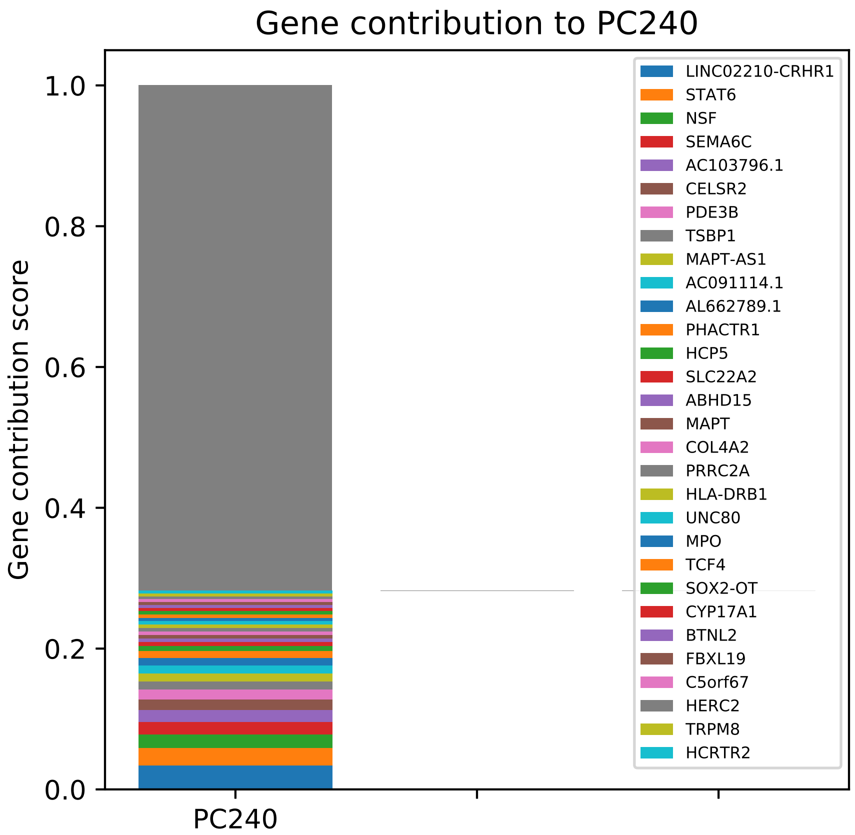

Gene contribution to PC240
Gene contribution score
1.0
0.8
0.6
0.4
0.2
0.0
PC240
LINC02210-CRHR1
STAT6
NSF
SEMA6C
AC103796.1
CELSR2
PDE3B
TSBP1
MAPT-AS1
AC091114.1
AL662789.1
PHACTR1
HCP5
SLC22A2
ABHD15
MAPT
COL4A2
PRRC2A
HLA-DRB1
UNC80
MPO
TCF4
SOX2-OT
CYP17A1
BTNL2
FBXL19
C5orf67
HERC2
TRPM8
HCRTR2

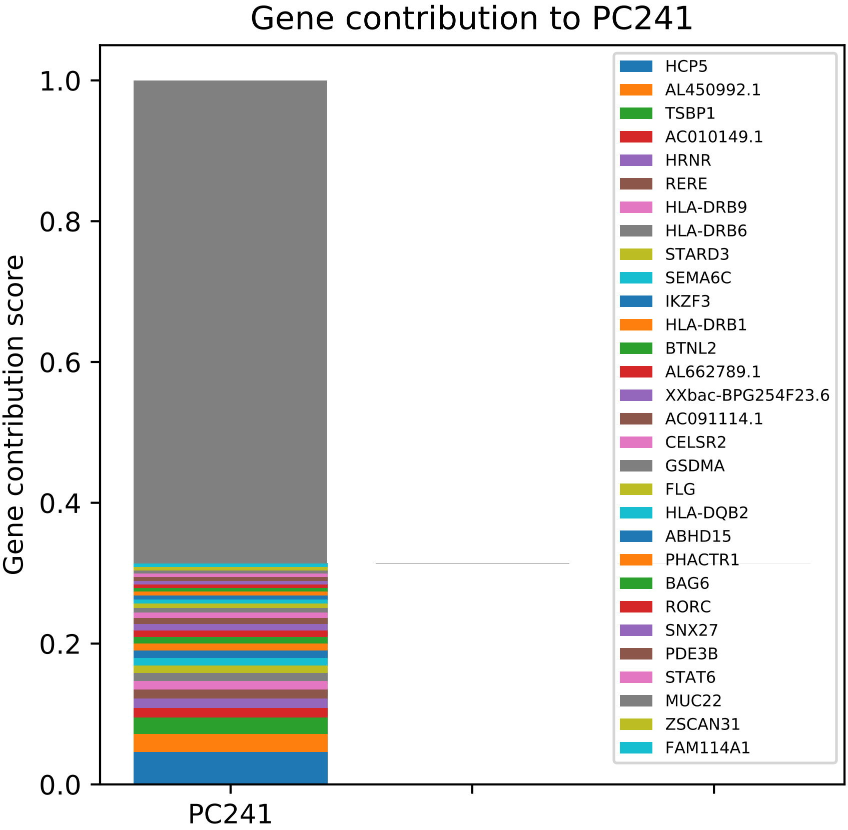
Gene contribution to PC241
Gene contribution score
0.0
0.2
0.4
0.6
0.8
1.0
PC241
HCP5
AL450992.1
TSBP1
AC010149.1
HRNR
RERE
HLA-DRB9
HLA-DRB6
STARD3
SEMA6C
IKZF3
HLA-DRB1
BTNL2
AL662789.1
XXbac-BPG254F23.6
AC091114.1
CELSR2
GSDMA
FLG
HLA-DQB2
ABHD15
PHACTR1
BAG6
RORC
SNX27
PDE3B
STAT6
MUC22
ZSCAN31
FAM114A1

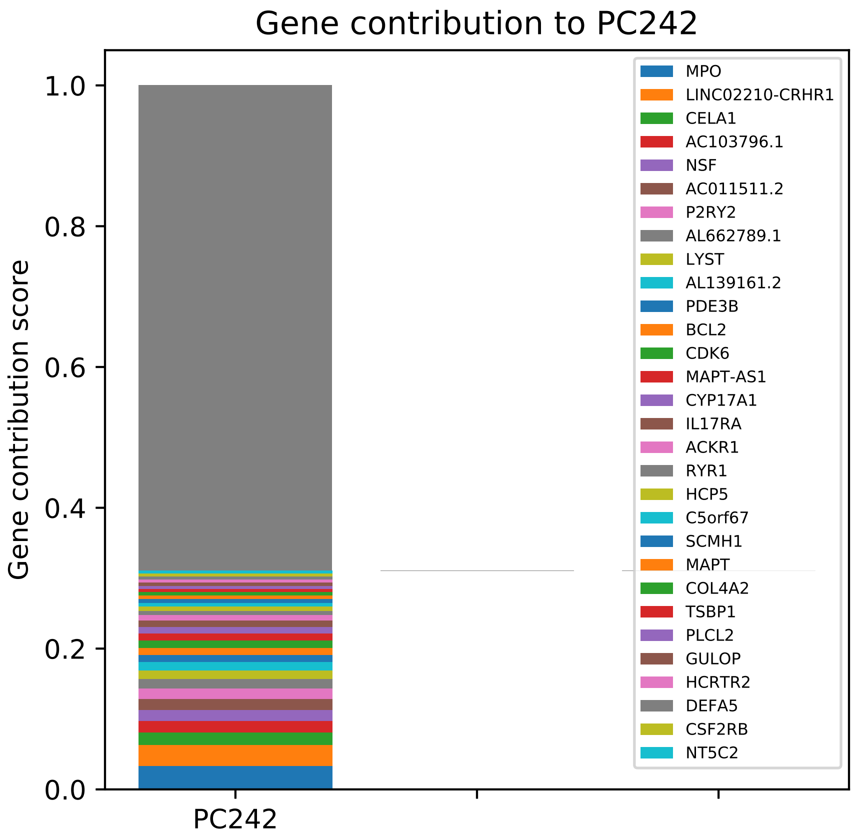
Gene contribution to PC242
Gene contribution score
1.0
0.8
0.6
0.4
0.2
0.0
PC242
MPO
LINC02210-CRHR1
CELA1
AC103796.1
NSF
AC011511.2
P2RY2
AL662789.1
LYST
AL139161.2
PDE3B
BCL2
CDK6
MAPT-AS1
CYP17A1
IL17RA
ACKR1
RYR1
HCP5
C5orf67
SCMH1
MAPT
COL4A2
TSBP1
PLCL2
GULOP
HCRTR2
DEFA5
CSF2RB
NT5C2

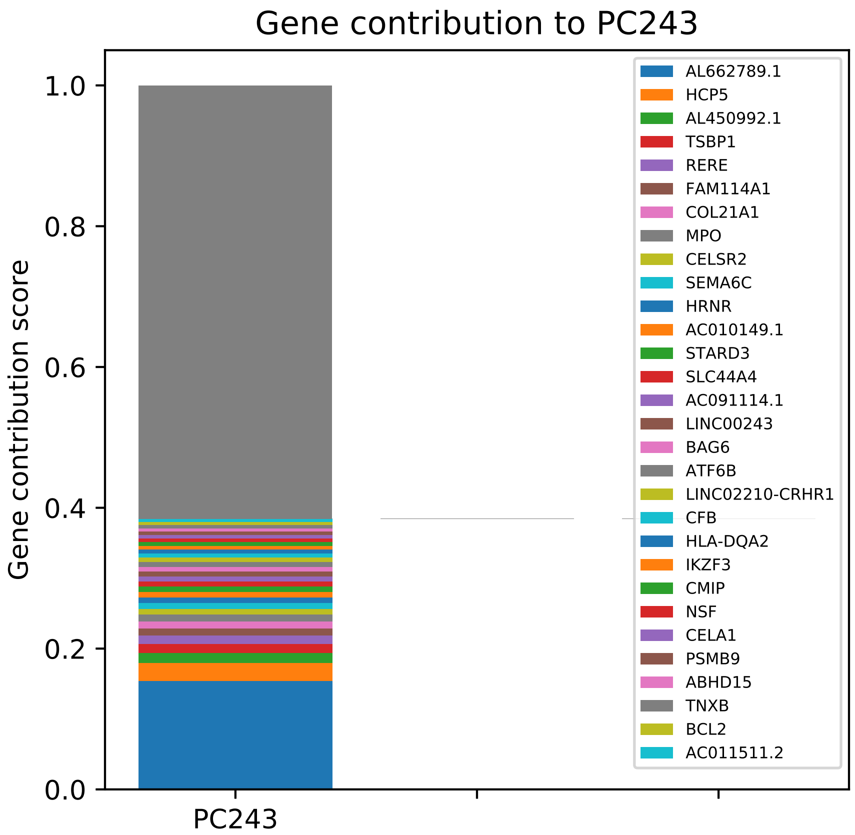

Gene contribution to PC243
Gene contribution score
1.0
0.8
0.6
0.4
0.2
0.0
PC243
AL662789.1
HCP5
AL450992.1
TSBP1
RERE
FAM114A1
COL21A1
MPO
CELSR2
SEMA6C
HRNR
AC010149.1
STARD3
SLC44A4
AC091114.1
LINC00243
BAG6
ATF6B
LINC02210-CRHR1
CFB
HLA-DQA2
IKZF3
CMIP
NSF
CELA1
PSMB9
ABHD15
TNXB
BCL2
AC011511.2

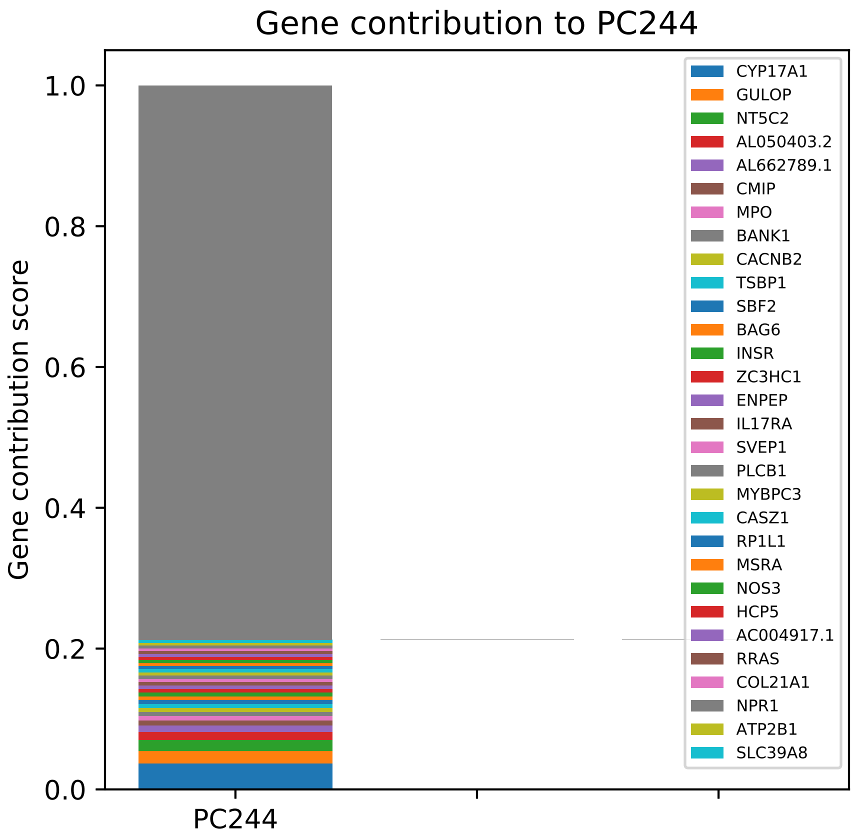

Gene contribution to PC244
Gene contribution score
1.0
0.8
0.6
0.4
0.2
0.0
PC244
CYP17A1
GULOP
NT5C2
AL050403.2
AL662789.1
CMIP
MPO
BANK1
CACNB2
TSBP1
SBF2
BAG6
INSR
ZC3HC1
ENPEP
IL17RA
SVEP1
PLCB1
MYBPC3
CASZ1
RP1L1
MSRA
NOS3
HCP5
AC004917.1
RRAS
COL21A1
NPR1
ATP2B1
SLC39A8

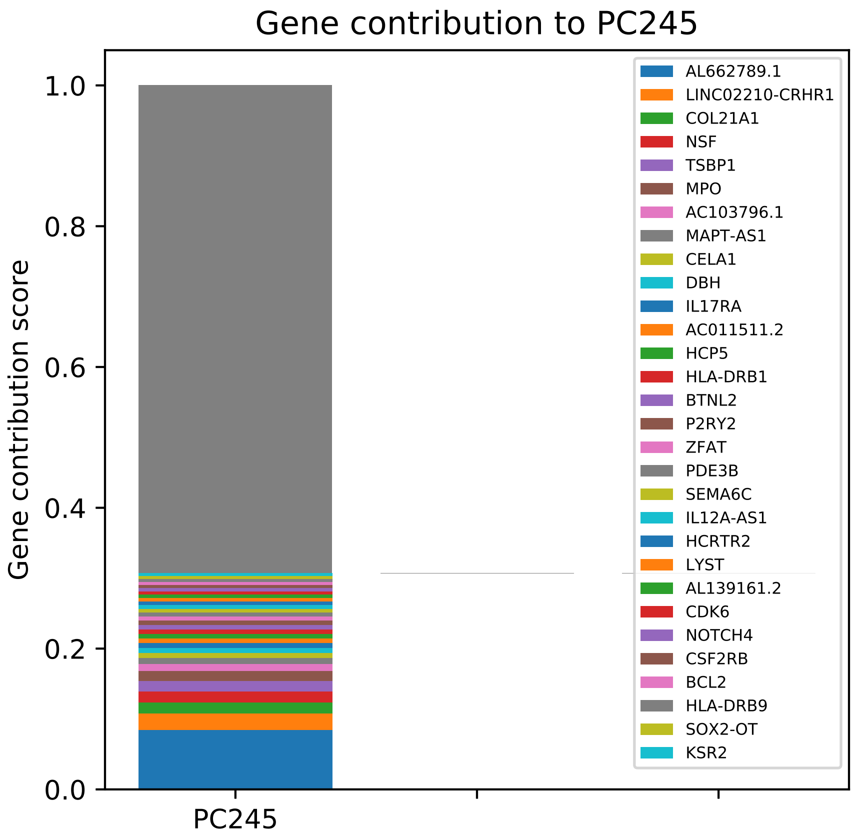

Gene contribution to PC245
Gene contribution score
1.0
0.8
0.6
0.4
0.2
0.0
PC245
AL662789.1
LINC02210-CRHR1
COL21A1
NSF
TSBP1
MPO
AC103796.1
MAPT-AS1
CELA1
DBH
IL17RA
AC011511.2
HCP5
HLA-DRB1
BTNL2
P2RY2
ZFAT
PDE3B
SEMA6C
IL12A-AS1
HCRTR2
LYST
AL139161.2
CDK6
NOTCH4
CSF2RB
BCL2
HLA-DRB9
SOX2-OT
KSR2

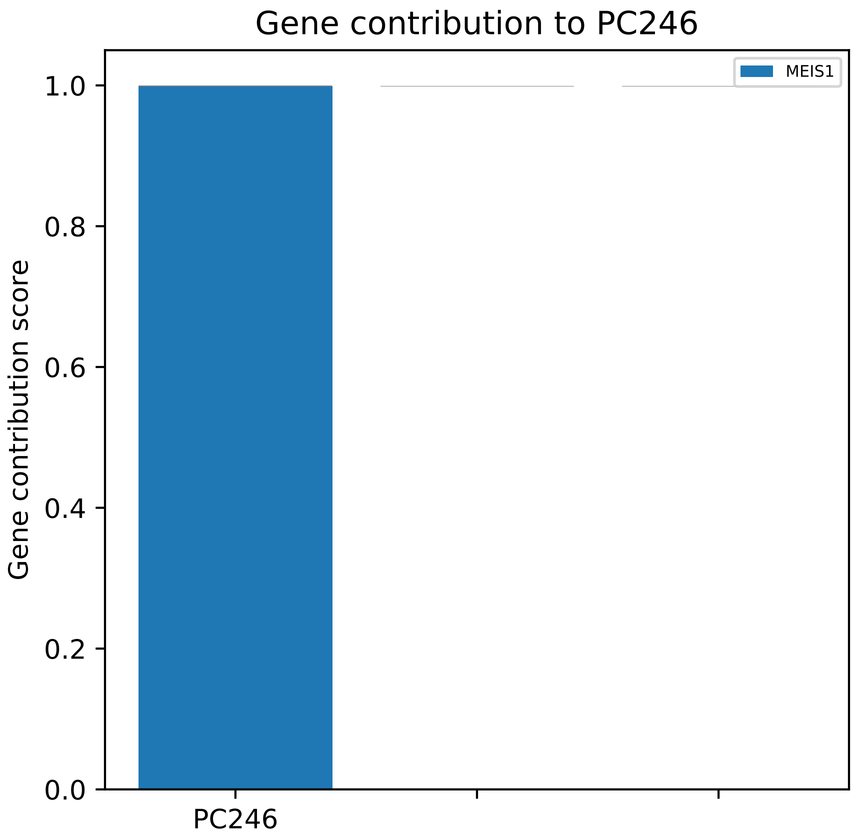

Gene contribution to PC246
MEIS1
Gene contribution score
1.0
0.8
0.6
0.4
0.2
0.0
PC246

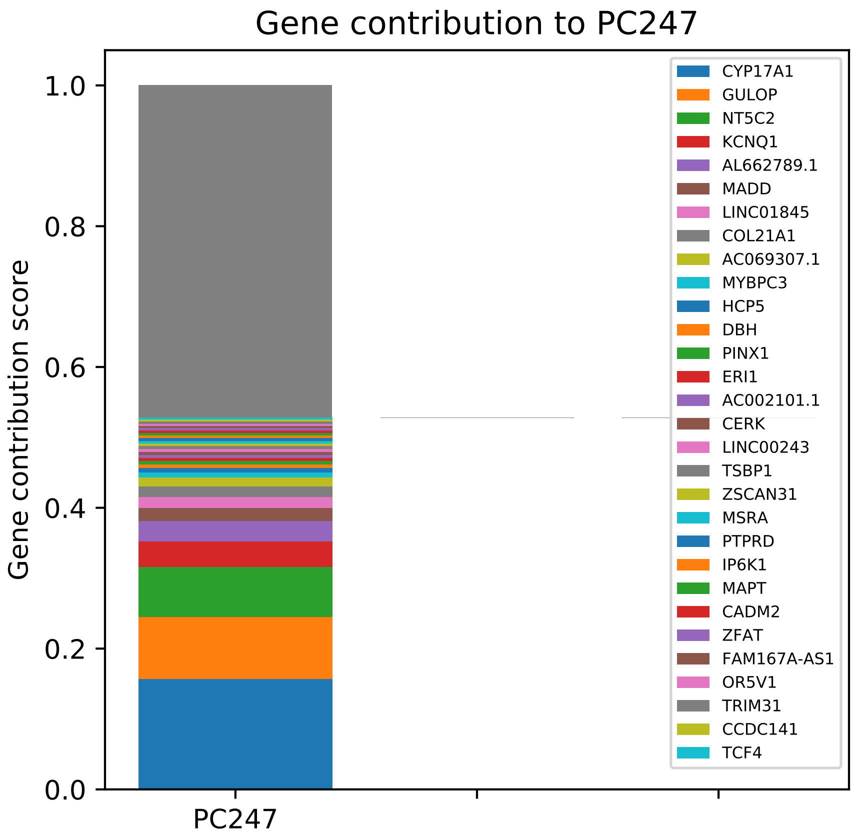

Gene contribution to PC247
1.0
0.8
0.6
0.4
0.2
0.0
Gene contribution score
PC247
CYP17A1
GULOP
NT5C2
KCNQ1
AL662789.1
MADD
LINC01845
COL21A1
AC069307.1
MYBPC3
HCP5
DBH
PINX1
ERI1
AC002101.1
CERK
LINC00243
TSBP1
ZSCAN31
MSRA
PTPRD
IP6K1
MAPT
CADM2
ZFAT
FAM167A-AS1
OR5V1
TRIM31
CCDC141
TCF4

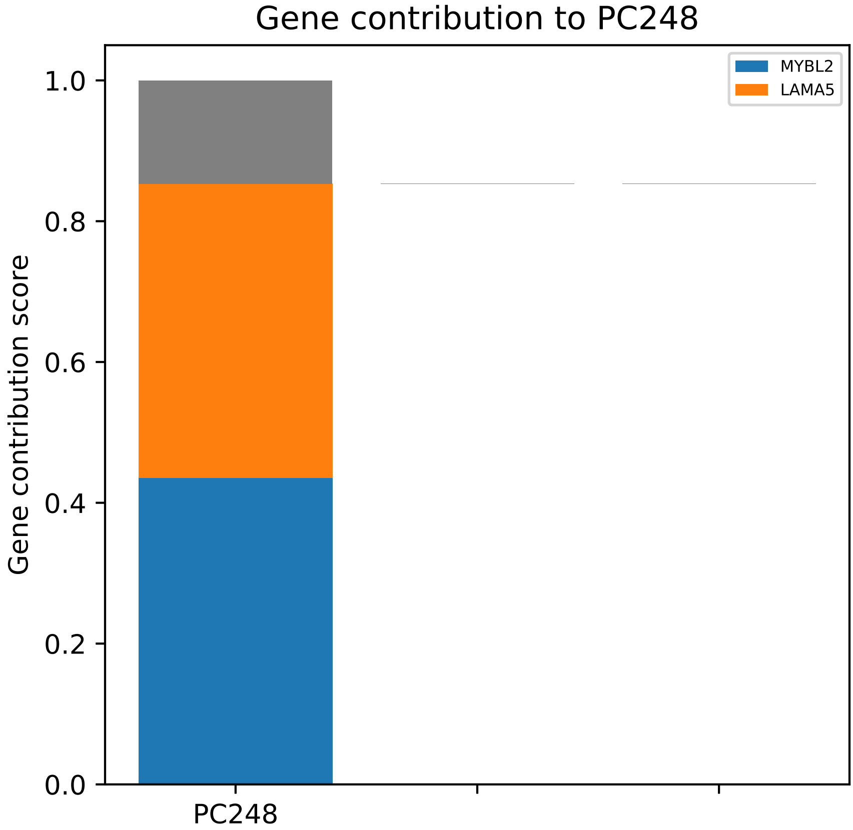

Gene contribution to PC248
Gene contribution score
1.0
0.8
0.6
0.4
0.2
0.0
PC248
MYBL2
LAMA5

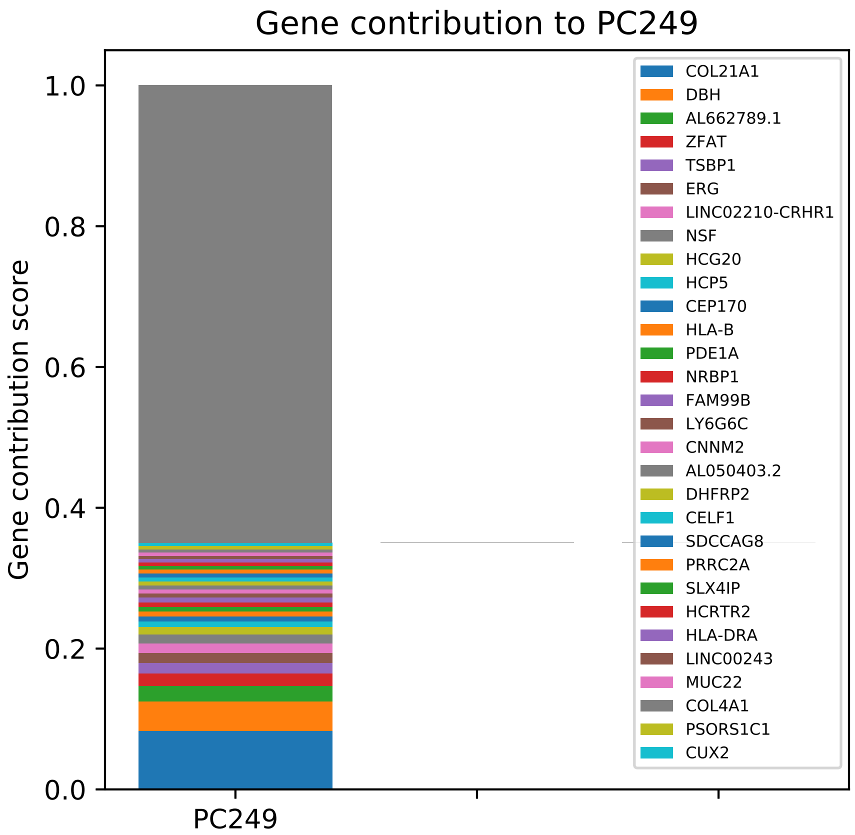
Gene contribution to PC249
Gene contribution score
1.0
0.8
0.6
0.4
0.2
0.0
PC249
COL21A1
DBH
AL662789.1
ZFAT
TSBP1
ERG
LINC02210-CRHR1
NSF
HCG20
HCP5
CEP170
HLA-B
PDE1A
NRBP1
FAM99B
LY6G6C
CNNM2
AL050403.2
DHFRP2
CELF1
SDCCAG8
PRRC2A
SLX4IP
HCRTR2
HLA-DRA
LINC00243
MUC22
COL4A1
PSORS1C1
CUX2

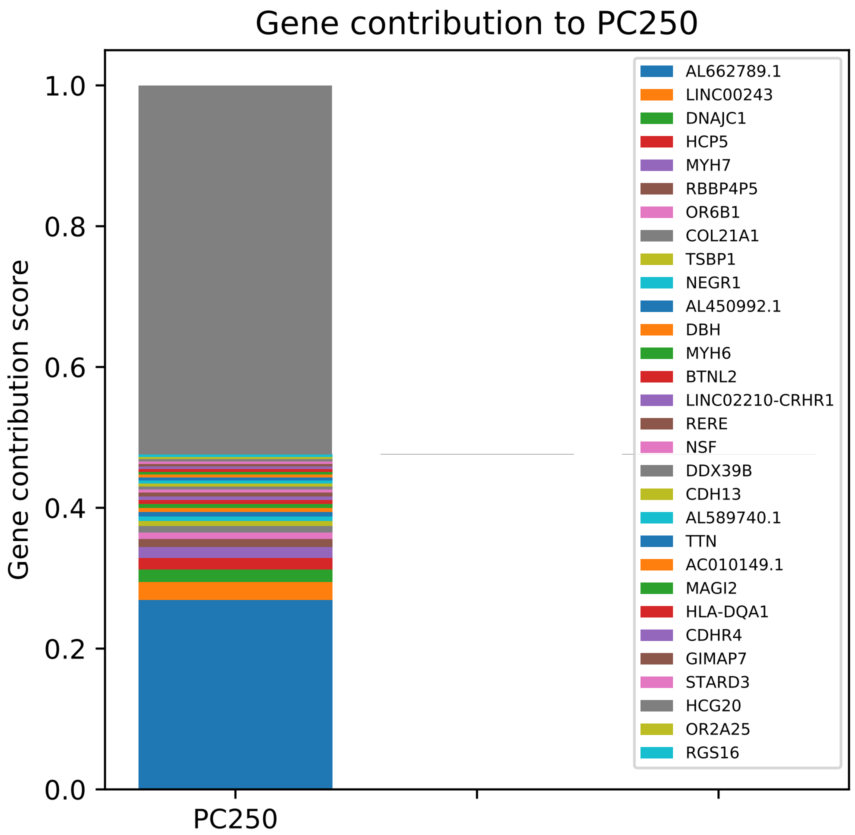

Gene contribution to PC250
Gene contribution score
1.0
0.8
0.6
0.4
0.2
0.0
PC250
AL662789.1
LINC00243
DNAJC1
HCP5
MYH7
RBBP4P5
OR6B1
COL21A1
TSBP1
NEGR1
AL450992.1
DBH
MYH6
BTNL2
LINC02210-CRHR1
RERE
NSF
DDX39B
CDH13
AL589740.1
TTN
AC010149.1
MAGI2
HLA-DQA1
CDHR4
GIMAP7
STARD3
HCG20
OR2A25
RGS16

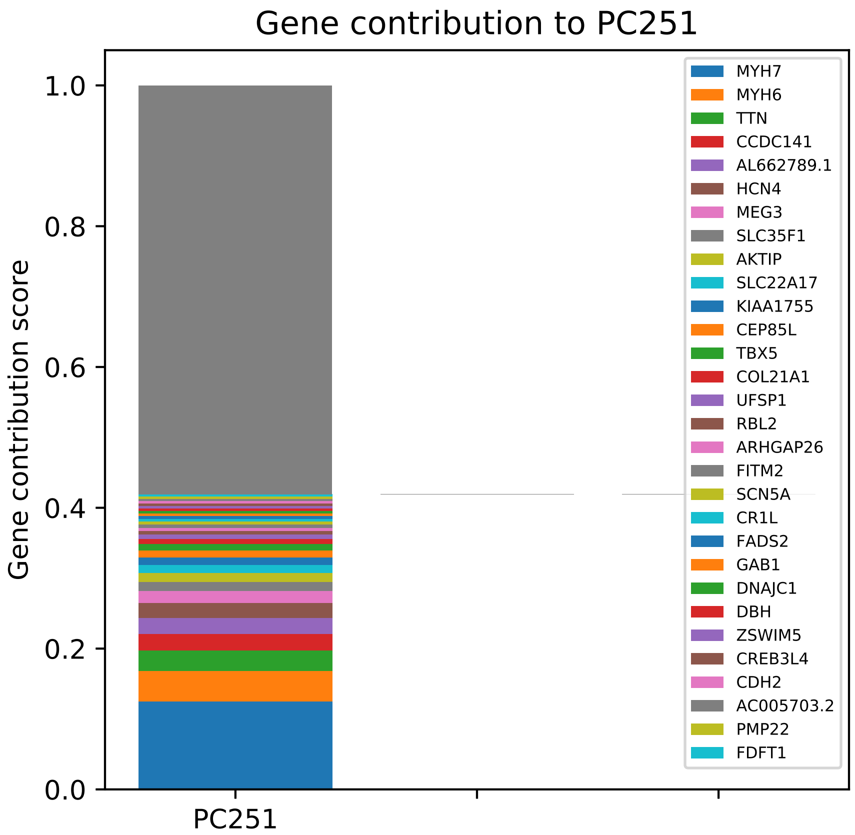

Gene contribution to PC251
Gene contribution score
1.0
0.8
0.6
0.4
0.2
0.0
PC251
MYH7
MYH6
TTN
CCDC141
AL662789.1
HCN4
MEG3
SLC35F1
AKTIP
SLC22A17
KIAA1755
CEP85L
TBX5
COL21A1
UFSP1
RBL2
ARHGAP26
FITM2
SCN5A
CR1L
FADS2
GAB1
DNAJC1
DBH
ZSWIM5
CREB3L4
CDH2
AC005703.2
PMP22
FDFT1

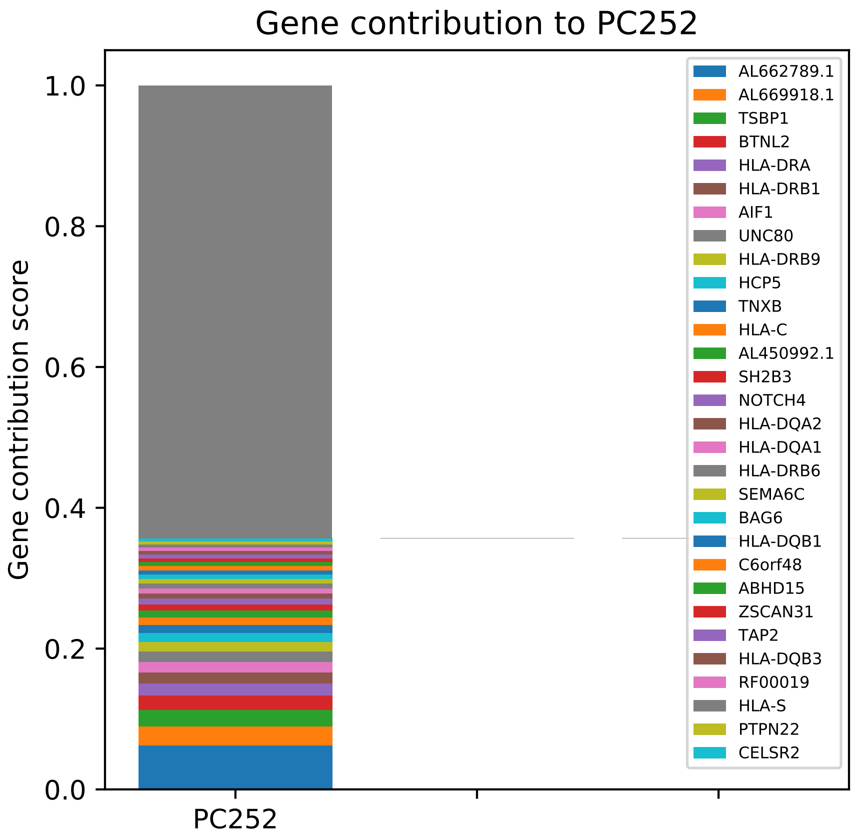
Gene contribution to PC252
Gene contribution score
1.0
0.8
0.6
0.4
0.2
0.0
PC252
AL662789.1
AL669918.1
TSBP1
BTNL2
HLA-DRA
HLA-DRB1
AIF1
UNC80
HLA-DRB9
HCP5
TNXB
HLA-C
AL450992.1
SH2B3
NOTCH4
HLA-DQA2
HLA-DQA1
HLA-DRB6
SEMA6C
BAG6
HLA-DQB1
C6orf48
ABHD15
ZSCAN31
TAP2
HLA-DQB3
RF00019
HLA-S
PTPN22
CELSR2

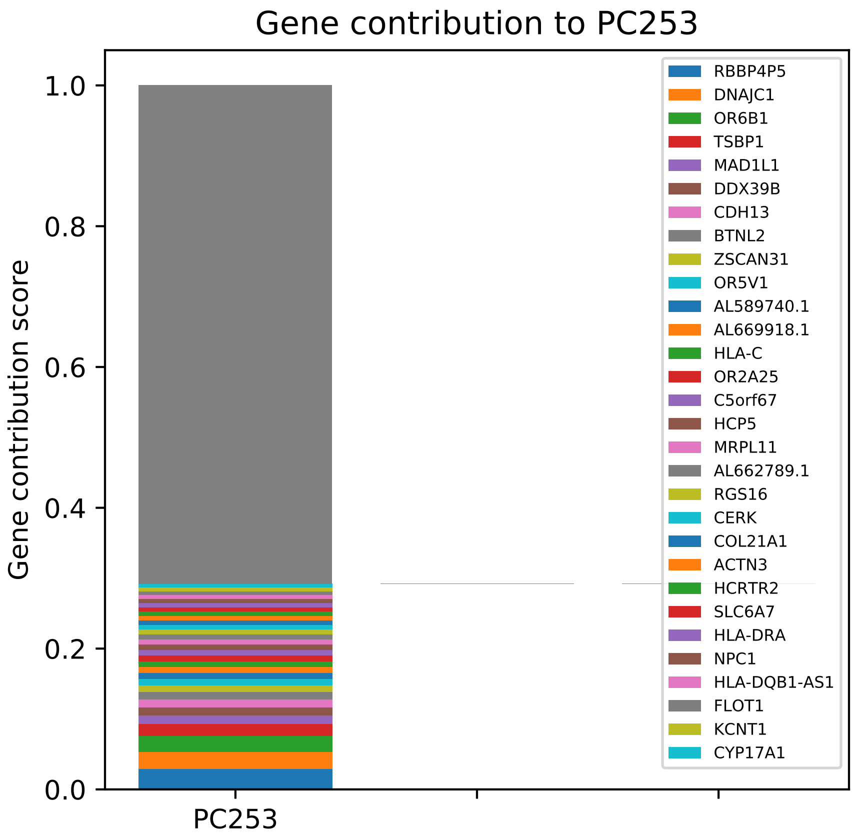
Gene contribution to PC253
Gene contribution score
1.0
0.8
0.6
0.4
0.2
0.0
PC253
RBBP4P5
DNAJC1
OR6B1
TSBP1
MAD1L1
DDX39B
CDH13
BTNL2
ZSCAN31
OR5V1
AL589740.1
AL669918.1
HLA-C
OR2A25
C5orf67
HCP5
MRPL11
AL662789.1
RGS16
CERK
COL21A1
ACTN3
HCRTR2
SLC6A7
HLA-DRA
NPC1
HLA-DQB1-AS1
FLOT1
KCNT1
CYP17A1

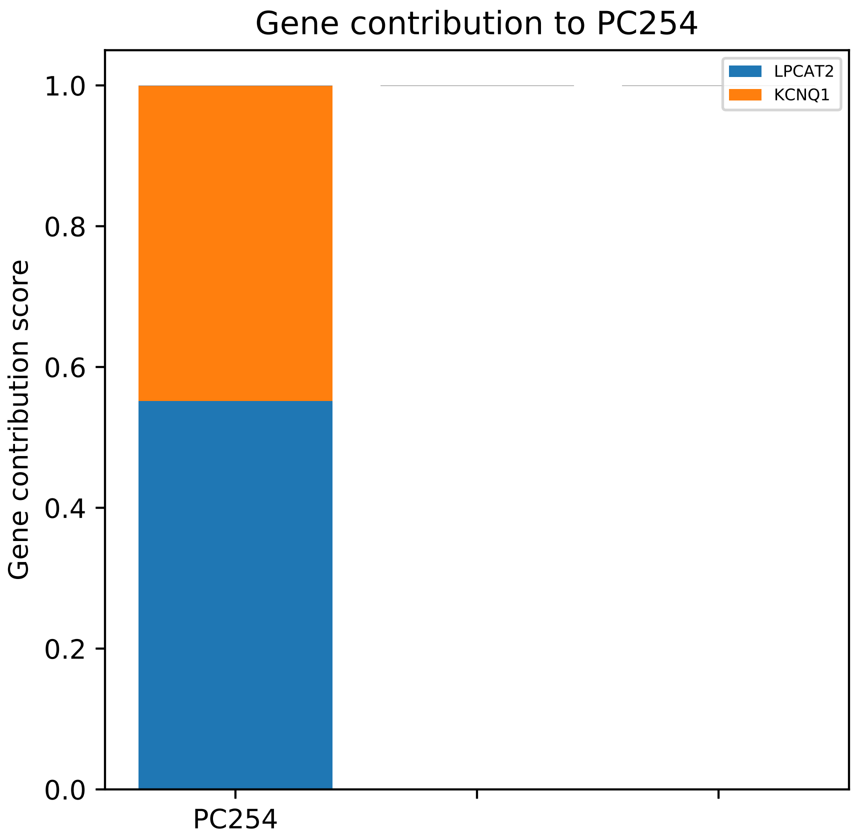

Gene contribution to PC254
Gene contribution score
1.0
0.8
0.6
0.4
0.2
0.0
PC254
LPCAT2
KCNQ1

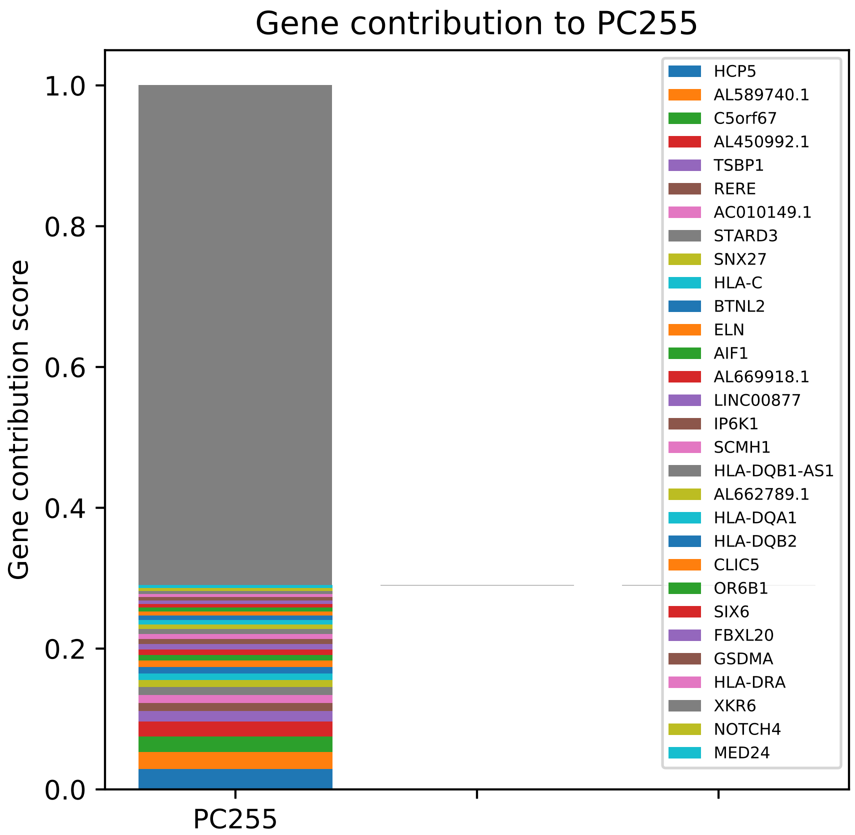

Gene contribution to PC255
Gene contribution score
1.0
0.8
0.6
0.4
0.2
0.0
PC255
HCP5
AL589740.1
C5orf67
AL450992.1
TSBP1
RERE
AC010149.1
STARD3
SNX27
HLA-C
BTNL2
ELN
AIF1
AL669918.1
LINC00877
IP6K1
SCMH1
HLA-DQB1-AS1
AL662789.1
HLA-DQA1
HLA-DQB2
CLIC5
OR6B1
SIX6
FBXL20
GSDMA
HLA-DRA
XKR6
NOTCH4
MED24

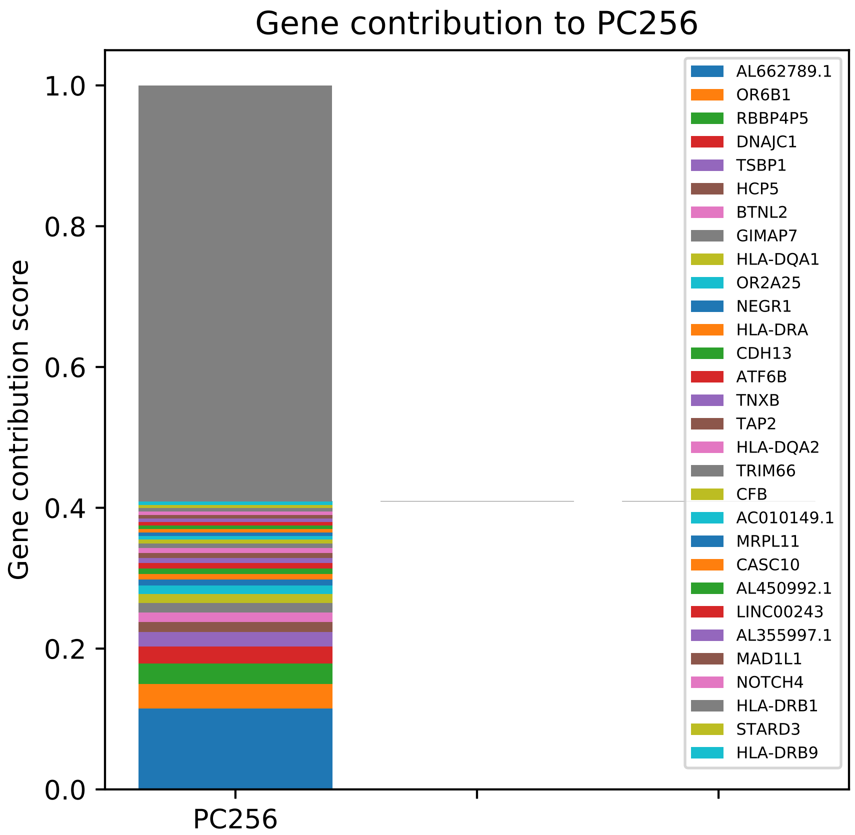
Gene contribution to PC256
Gene contribution score
1.0
0.8
0.6
0.4
0.2
0.0
PC256
AL662789.1
OR6B1
RBBP4P5
DNAJC1
TSBP1
HCP5
BTNL2
GIMAP7
HLA-DQA1
OR2A25
NEGR1
HLA-DRA
CDH13
ATF6B
TNXB
TAP2
HLA-DQA2
TRIM66
CFB
AC010149.1
MRPL11
CASC10
AL450992.1
LINC00243
AL355997.1
MAD1L1
NOTCH4
HLA-DRB1
STARD3
HLA-DRB9

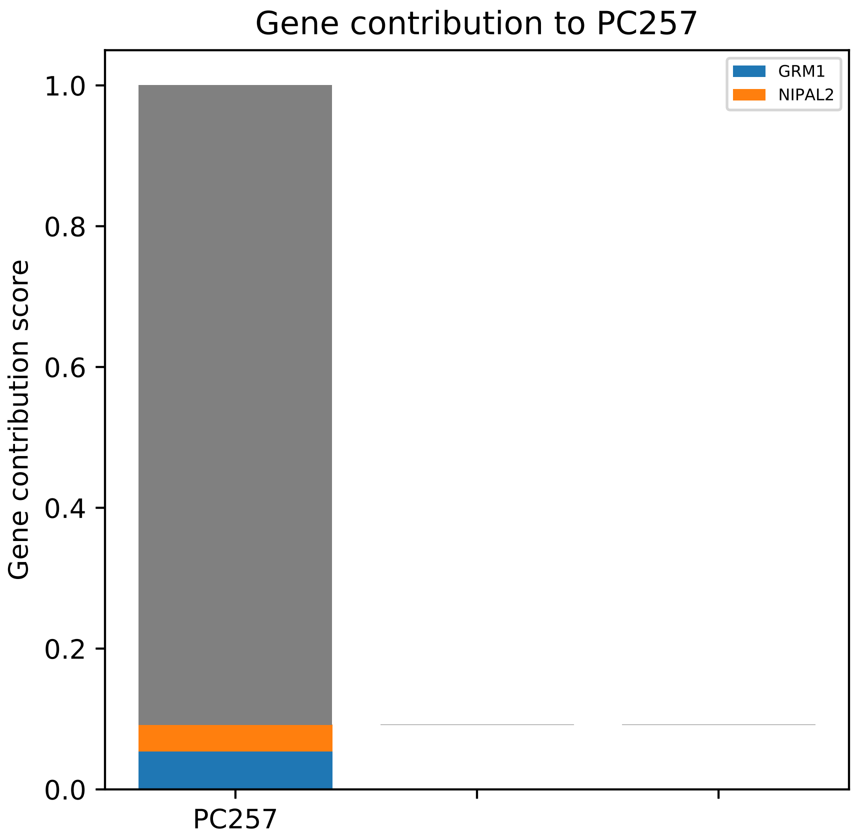
Gene contribution to PC257
GRM1
NIPAL2
Gene contribution score
1.0
0.8
0.6
0.4
0.2
0.0
PC257

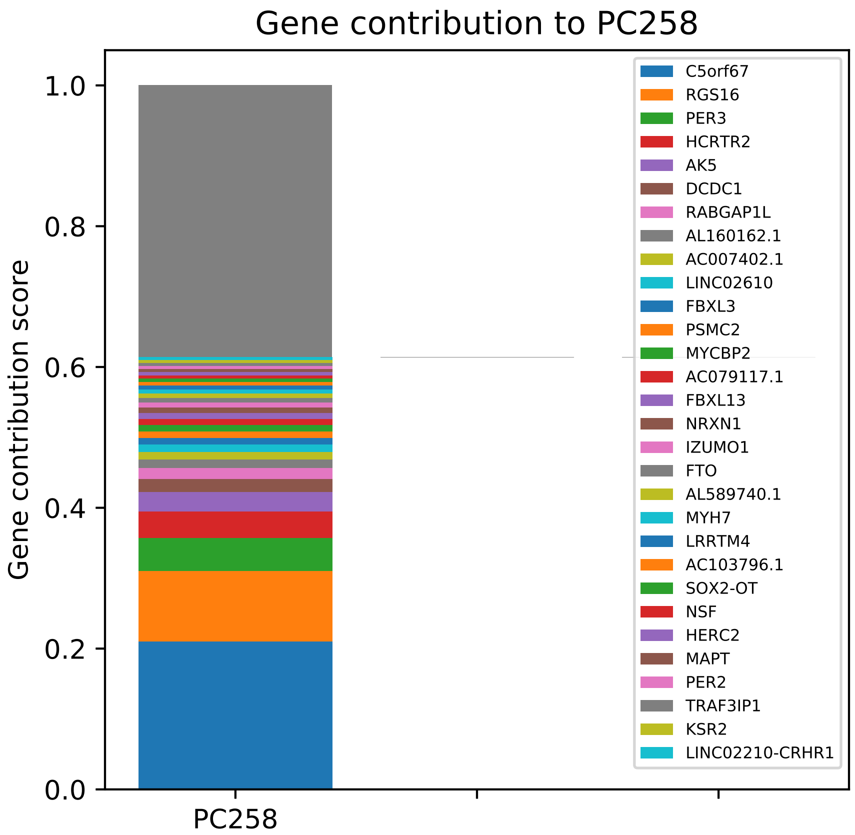
Gene contribution to PC258
Gene contribution score
1.0
0.8
0.6
0.4
0.2
0.0
PC258
C5orf67
RGS16
PER3
HCRTR2
AK5
DCDC1
RABGAP1L
AL160162.1
AC007402.1
LINC02610
FBXL3
PSMC2
MYCBP2
AC079117.1
FBXL13
NRXN1
IZUMO1
FTO
AL589740.1
MYH7
LRRTM4
AC103796.1
SOX2-OT
NSF
HERC2
MAPT
PER2
TRAF3IP1
KSR2
LINC02210-CRHR1

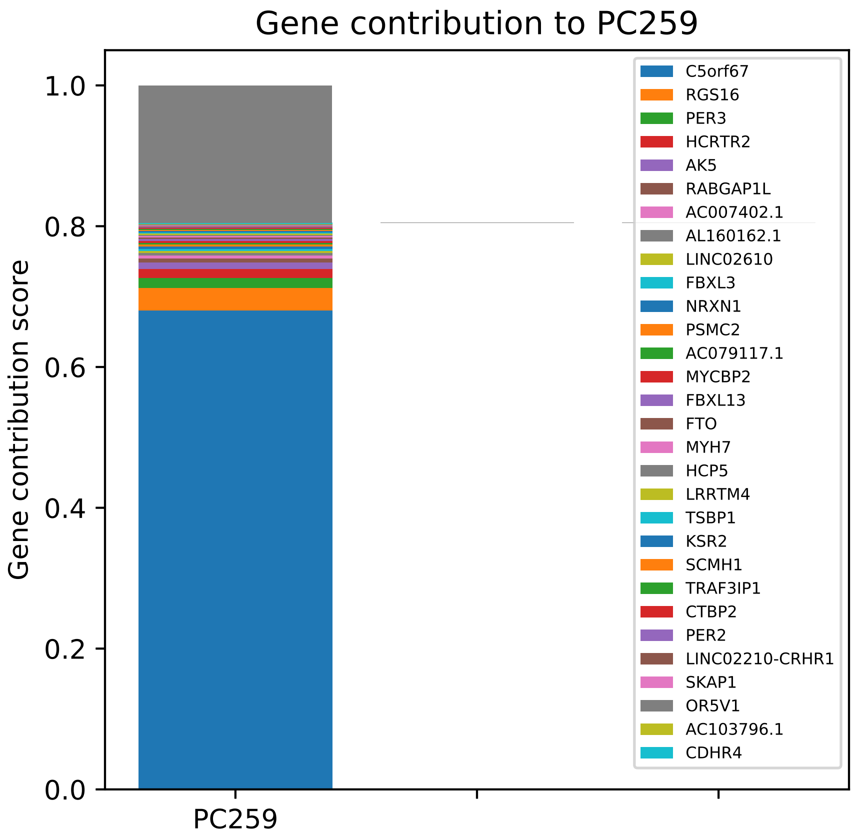

Gene contribution to PC259
Gene contribution score
1.0
0.8
0.6
0.4
0.2
0.0
PC259
C5orf67
RGS16
PER3
HCRTR2
AK5
RABGAP1L
AC007402.1
AL160162.1
LINC02610
FBXL3
NRXN1
PSMC2
AC079117.1
MYCBP2
FBXL13
FTO
MYH7
HCP5
LRRTM4
TSBP1
KSR2
SCMH1
TRAF3IP1
CTBP2
PER2
LINC02210-CRHR1
SKAP1
OR5V1
AC103796.1
CDHR4

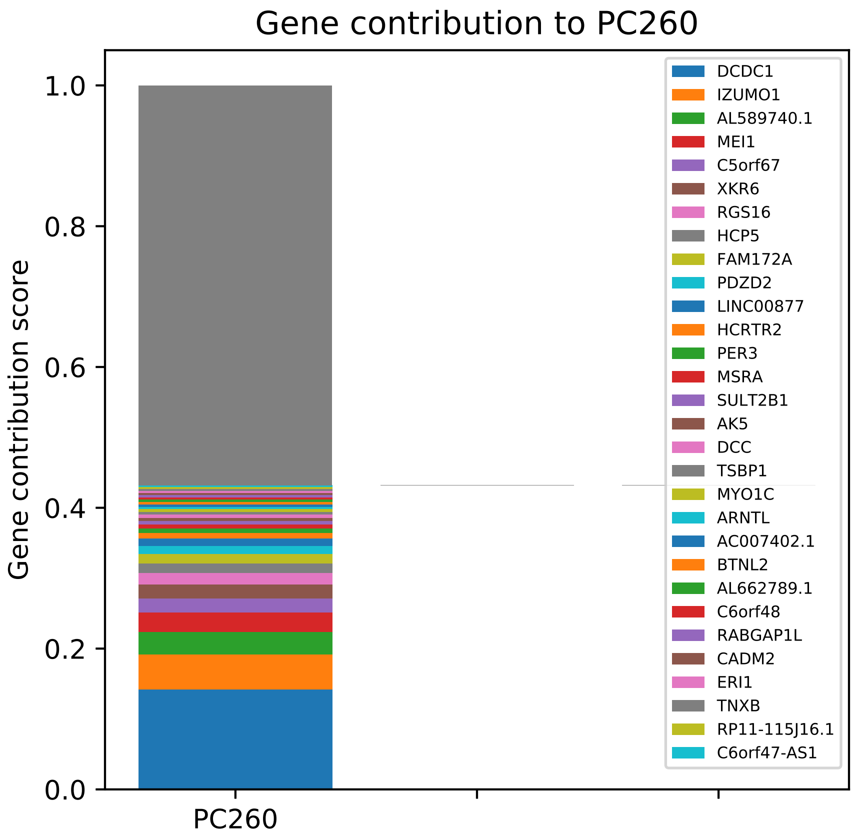

Gene contribution to PC260
Gene contribution score
1.0
0.8
0.6
0.4
0.2
0.0
PC260
DCDC1
IZUMO1
AL589740.1
MEI1
C5orf67
XKR6
RGS16
HCP5
FAM172A
PDZD2
LINC00877
HCRTR2
PER3
MSRA
SULT2B1
AK5
DCC
TSBP1
MYO1C
ARNTL
AC007402.1
BTNL2
AL662789.1
C6orf48
RABGAP1L
CADM2
ERI1
TNXB
RP11-115J16.1
C6orf47-AS1

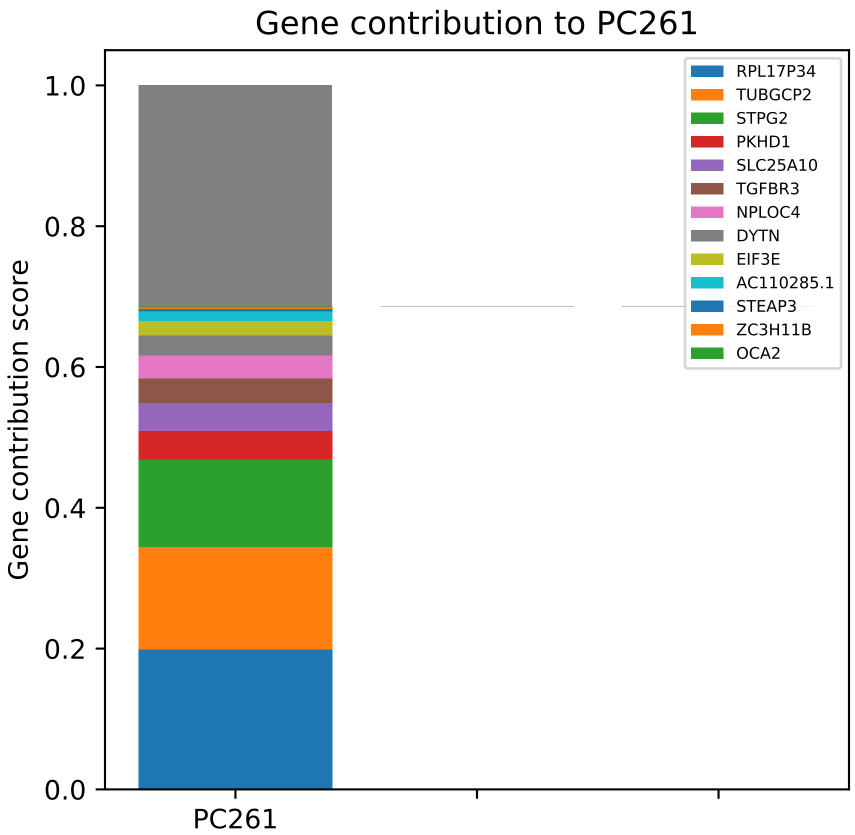
Gene contribution to PC261
Gene contribution score
1.0
0.8
0.6
0.4
0.2
0.0
PC261
RPL17P34
TUBGCP2
STPG2
PKHD1
SLC25A10
TGFBR3
NPLOC4
DYTN
EIF3E
AC110285.1
STEAP3
ZC3H11B
OCA2

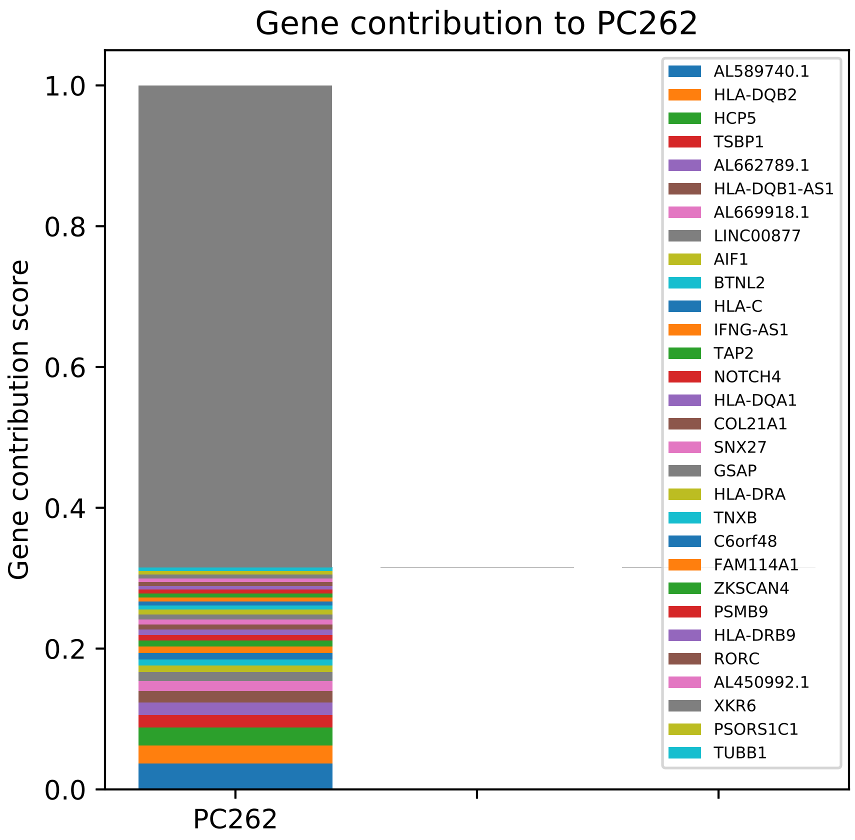

Gene contribution to PC262
Gene contribution score
1.0
0.8
0.6
0.4
0.2
0.0
PC262
AL589740.1
HLA-DQB2
HCP5
TSBP1
AL662789.1
HLA-DQB1-AS1
AL669918.1
LINC00877
AIF1
BTNL2
HLA-C
IFNG-AS1
TAP2
NOTCH4
HLA-DQA1
COL21A1
SNX27
GSAP
HLA-DRA
TNXB
C6orf48
FAM114A1
ZKSCAN4
PSMB9
HLA-DRB9
RORC
AL450992.1
XKR6
PSORS1C1
TUBB1

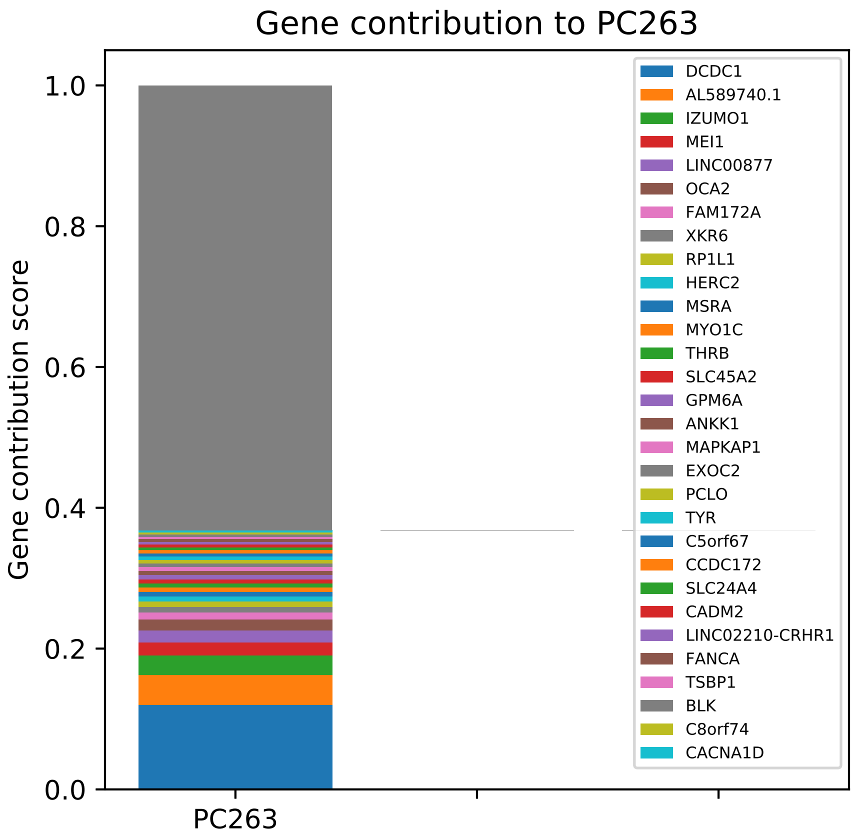
Gene contribution to PC263
Gene contribution score
1.0
0.8
0.6
0.4
0.2
0.0
PC263
DCDC1
AL589740.1
IZUMO1
MEI1
LINC00877
OCA2
FAM172A
XKR6
RP1L1
HERC2
MSRA
MYO1C
THRB
SLC45A2
GPM6A
ANKK1
MAPKAP1
EXOC2
PCLO
TYR
C5orf67
CCDC172
SLC24A4
CADM2
LINC02210-CRHR1
FANCA
TSBP1
BLK
C8orf74
CACNA1D

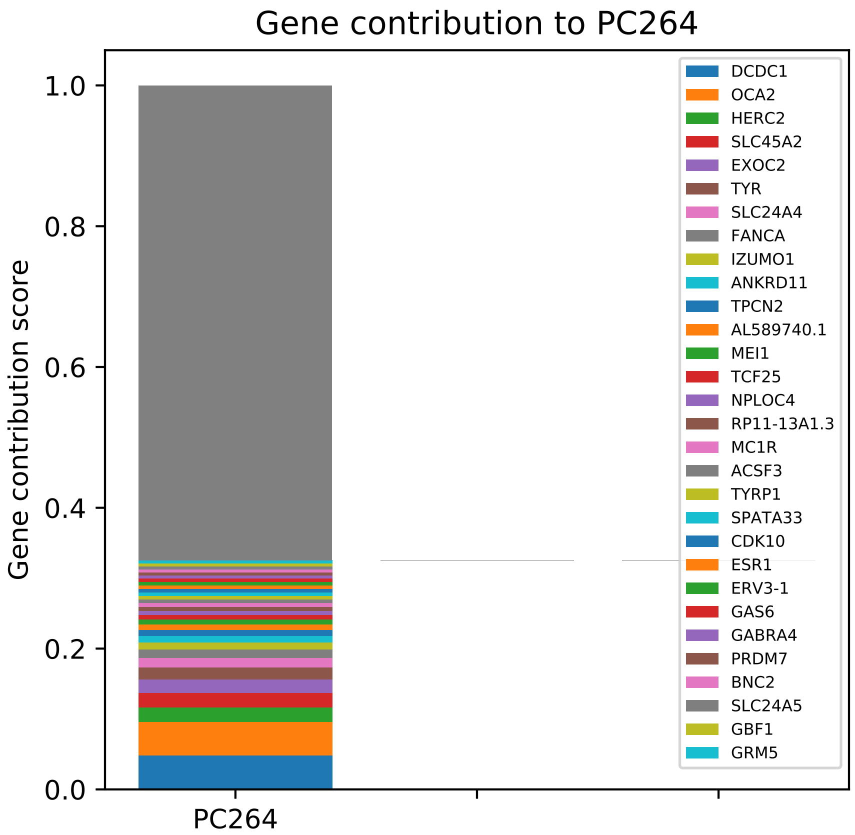

Gene contribution to PC264
Gene contribution score
1.0
0.8
0.6
0.4
0.2
0.0
PC264
DCDC1
OCA2
HERC2
SLC45A2
EXOC2
TYR
SLC24A4
FANCA
IZUMO1
ANKRD11
TPCN2
AL589740.1
MEI1
TCF25
NPLOC4
RP11-13A1.3
MC1R
ACSF3
TYRP1
SPATA33
CDK10
ESR1
ERV3-1
GAS6
GABRA4
PRDM7
BNC2
SLC24A5
GBF1
GRM5

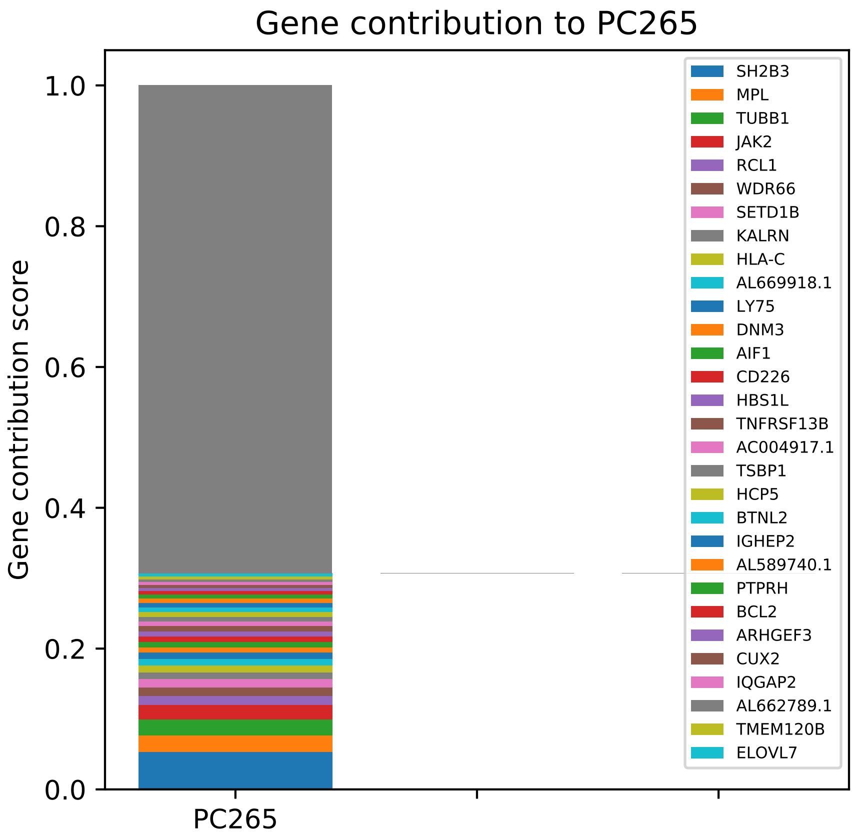
Gene contribution to PC265
Gene contribution score
1.0
0.8
0.6
0.4
0.2
0.0
PC265
SH2B3
MPL
TUBB1
JAK2
RCL1
WDR66
SETD1B
KALRN
HLA-C
AL669918.1
LY75
DNM3
AIF1
CD226
HBS1L
TNFRSF13B
AC004917.1
TSBP1
HCP5
BTNL2
IGHEP2
AL589740.1
PTPRH
BCL2
ARHGEF3
CUX2
IQGAP2
AL662789.1
TMEM120B
ELOVL7

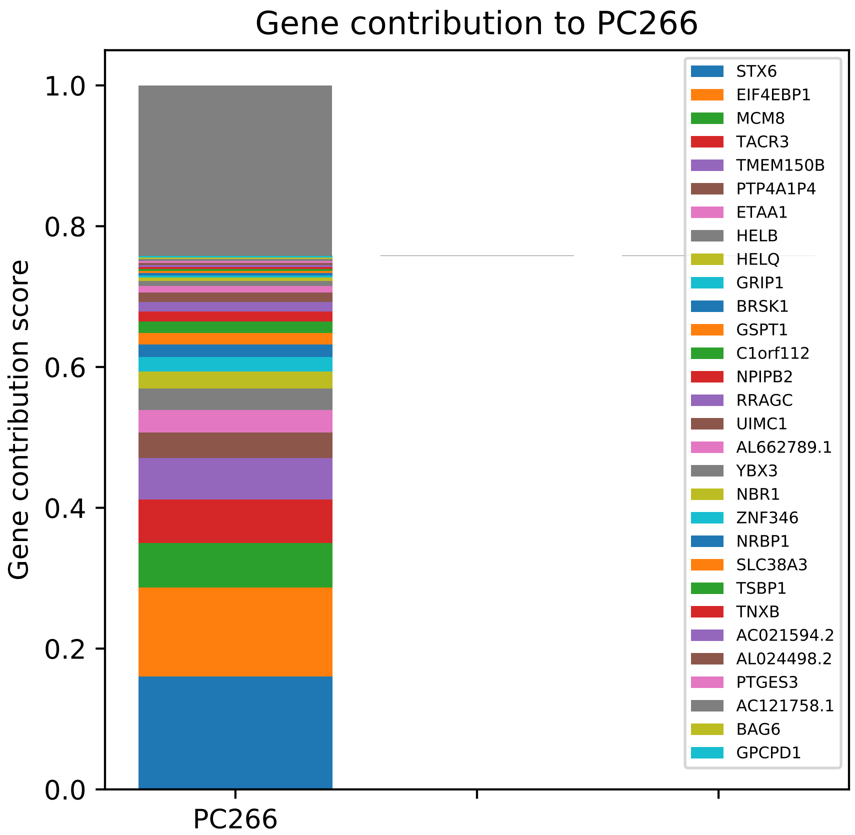
Gene contribution to PC266
Gene contribution score
1.0
0.8
0.6
0.4
0.2
0.0
PC266
STX6
EIF4EBP1
MCM8
TACR3
TMEM150B
PTP4A1P4
ETAA1
HELB
HELQ
GRIP1
BRSK1
GSPT1
C1orf112
NPIPB2
RRAGC
UIMC1
AL662789.1
YBX3
NBR1
ZNF346
NRBP1
SLC38A3
TSBP1
TNXB
AC021594.2
AL024498.2
PTGES3
AC121758.1
BAG6
GPCPD1

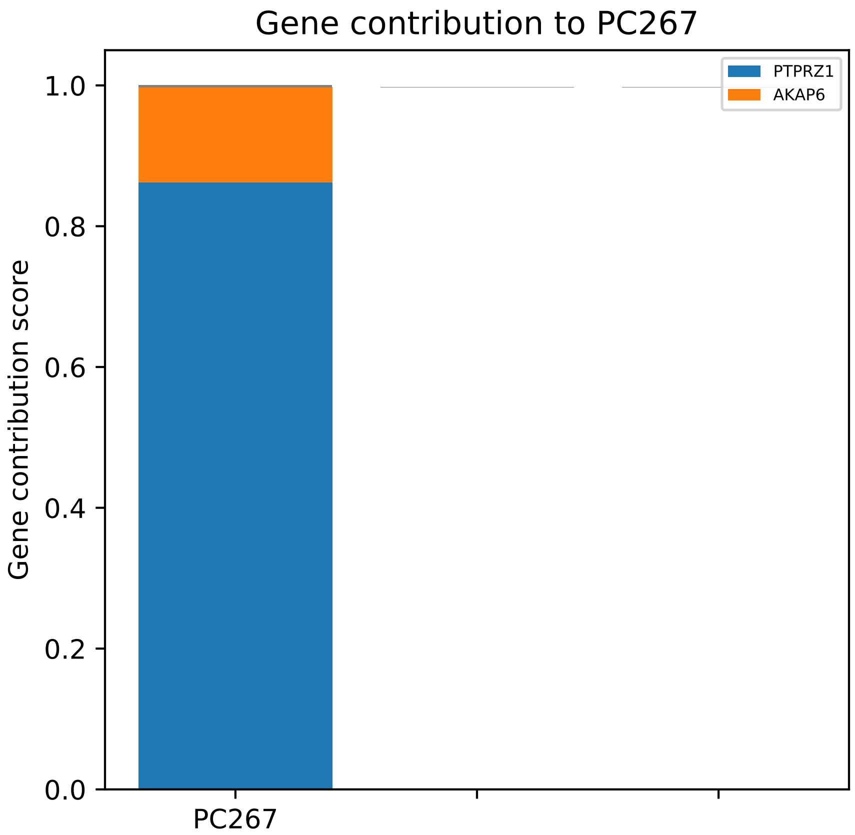
Gene contribution to PC267
Gene contribution score
1.0
0.8
0.6
0.4
0.2
0.0
PC267
PTPRZ1
AKAP6

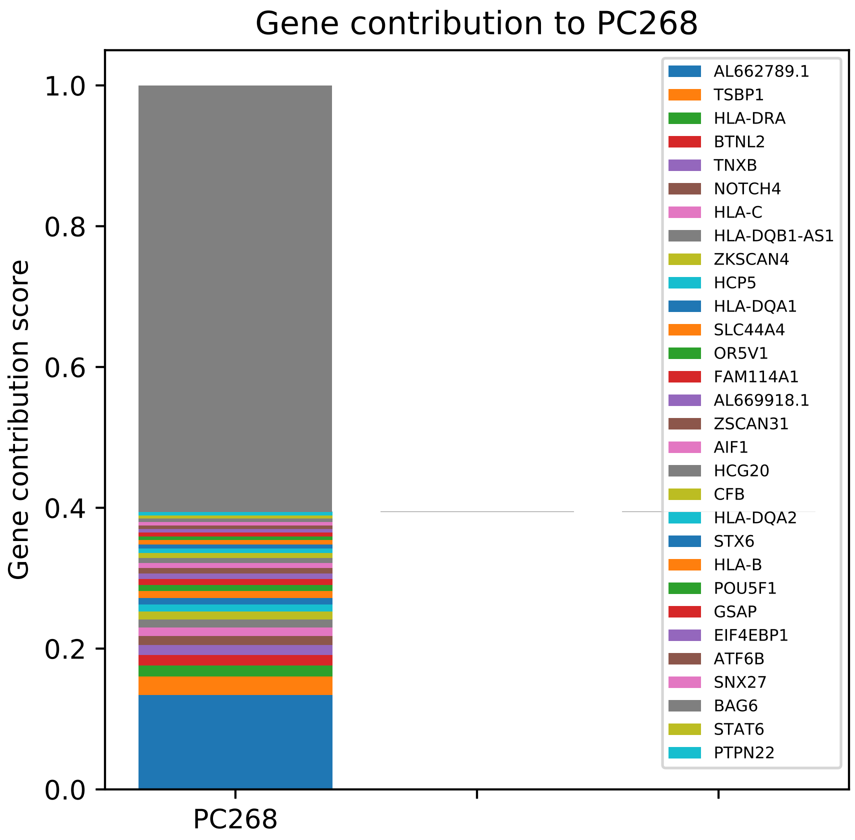
Gene contribution to PC268
Gene contribution score
1.0
0.8
0.6
0.4
0.2
0.0
PC268
AL662789.1
TSBP1
HLA-DRA
BTNL2
TNXB
NOTCH4
HLA-C
HLA-DQB1-AS1
ZKSCAN4
HCP5
HLA-DQA1
SLC44A4
OR5V1
FAM114A1
AL669918.1
ZSCAN31
AIF1
HCG20
CFB
HLA-DQA2
STX6
HLA-B
POU5F1
GSAP
EIF4EBP1
ATF6B
SNX27
BAG6
STAT6
PTPN22

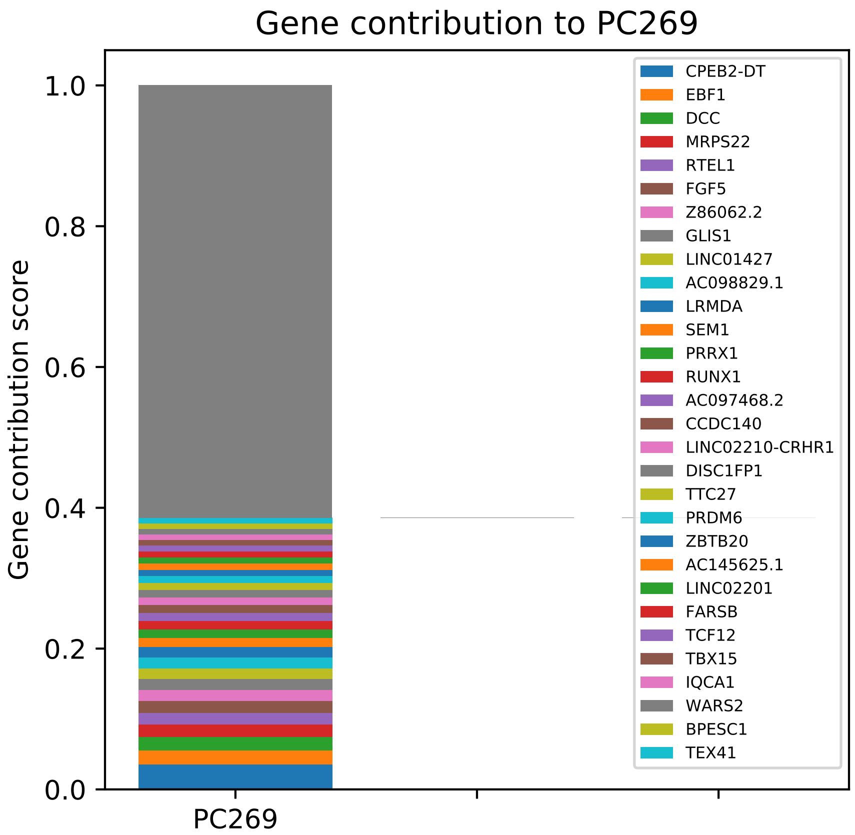

Gene contribution to PC269
Gene contribution score
1.0
0.8
0.6
0.4
0.2
0.0
PC269
CPEB2-DT
EBF1
DCC
MRPS22
RTEL1
FGF5
Z86062.2
GLIS1
LINC01427
AC098829.1
LRMDA
SEM1
PRRX1
RUNX1
AC097468.2
CCDC140
LINC02210-CRHR1
DISC1FP1
TTC27
PRDM6
ZBTB20
AC145625.1
LINC02201
FARSB
TCF12
TBX15
IQCA1
WARS2
BPESC1
TEX41

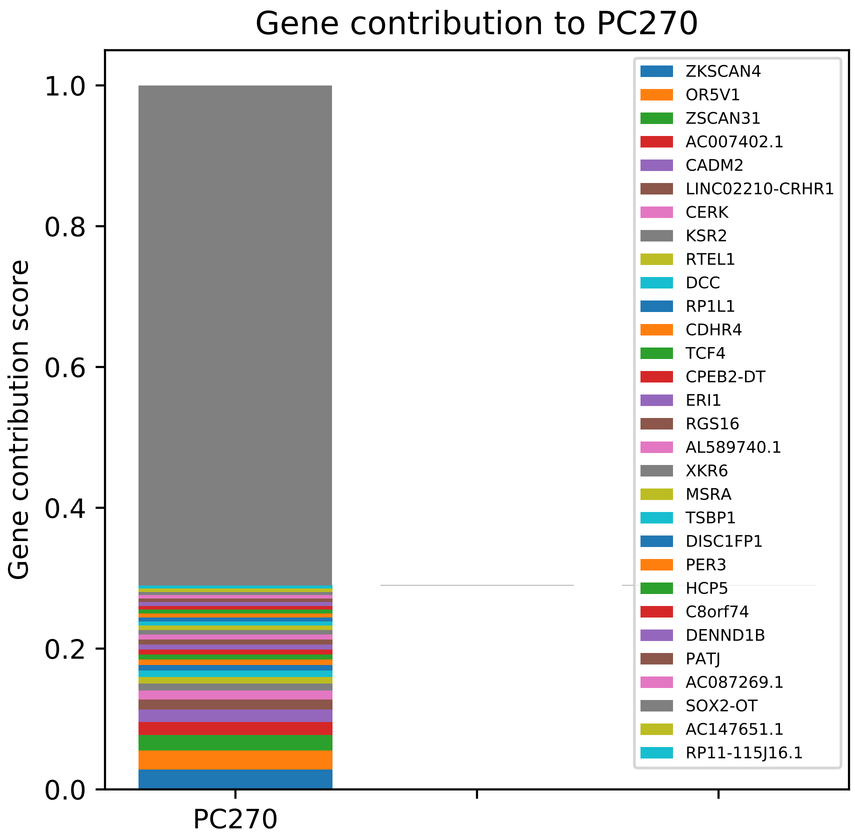
Gene contribution to PC270
Gene contribution score
1.0
0.8
0.6
0.4
0.2
0.0
PC270
ZKSCAN4
OR5V1
ZSCAN31
AC007402.1
CADM2
LINC02210-CRHR1
CERK
KSR2
RTEL1
DCC
RP1L1
CDHR4
TCF4
CPEB2-DT
ERI1
RGS16
AL589740.1
XKR6
MSRA
TSBP1
DISC1FP1
PER3
HCP5
C8orf74
DENND1B
PATJ
AC087269.1
SOX2-OT
AC147651.1
RP11-115J16.1

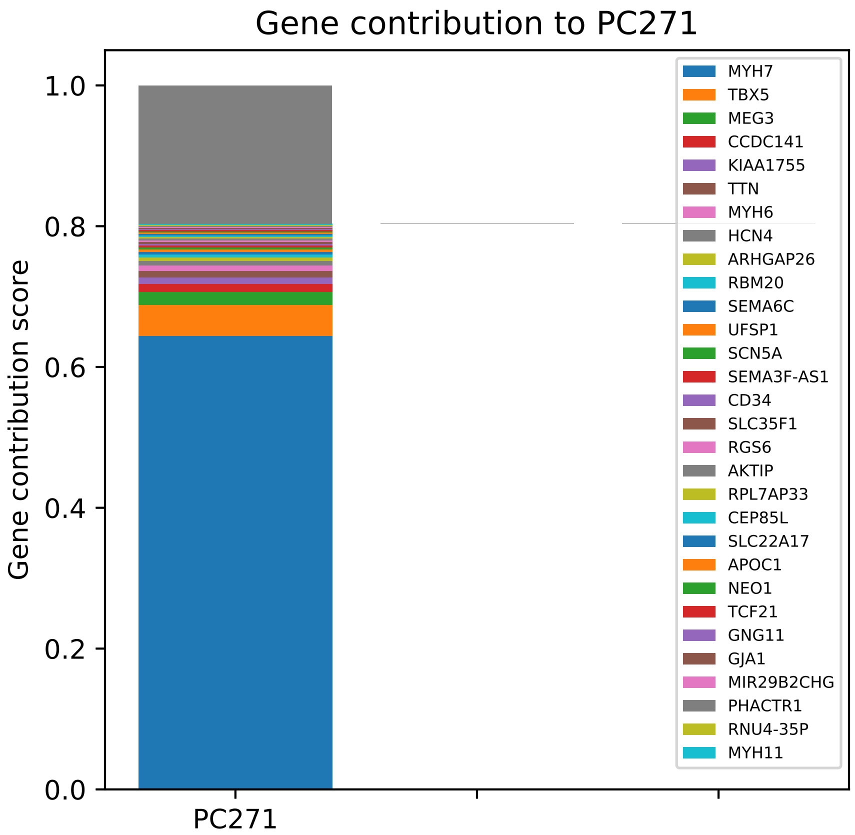
Gene contribution to PC271
Gene contribution score
1.0
0.8
0.6
0.4
0.2
0.0
PC271
MYH7
TBX5
MEG3
CCDC141
KIAA1755
TTN
MYH6
HCN4
ARHGAP26
RBM20
SEMA6C
UFSP1
SCN5A
SEMA3F-AS1
CD34
SLC35F1
RGS6
AKTIP
RPL7AP33
CEP85L
SLC22A17
APOC1
NEO1
TCF21
GNG11
GJA1
MIR29B2CHG
PHACTR1
RNU4-35P
MYH11

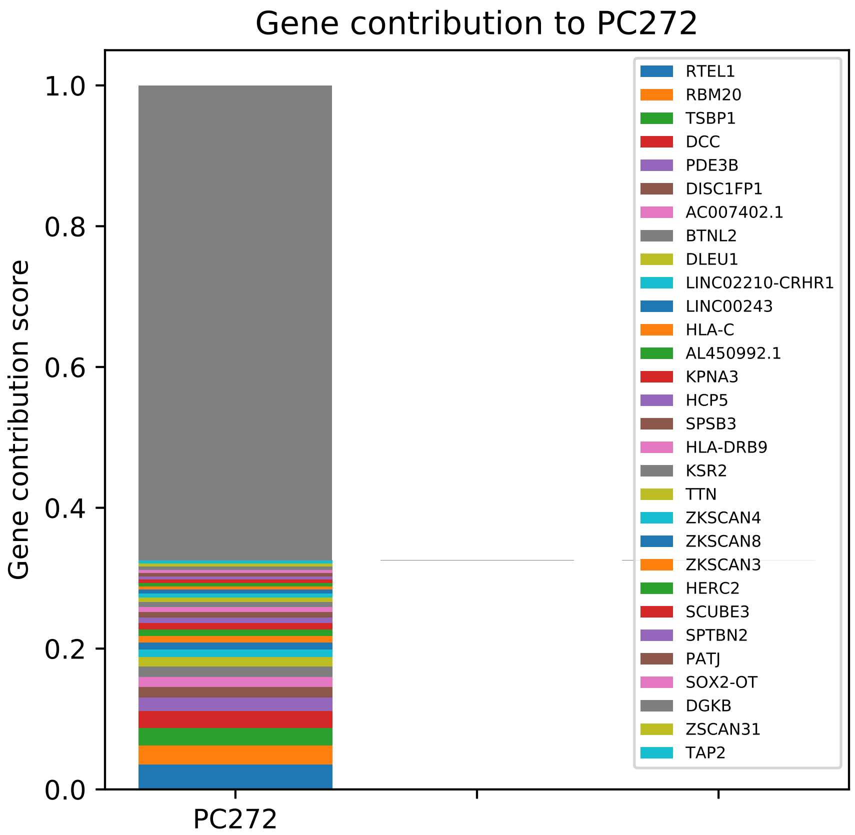

Gene contribution to PC272
Gene contribution score
1.0
0.8
0.6
0.4
0.2
0.0
PC272
RTEL1
RBM20
TSBP1
DCC
PDE3B
DISC1FP1
AC007402.1
BTNL2
DLEU1
LINC02210-CRHR1
LINC00243
HLA-C
AL450992.1
KPNA3
HCP5
SPSB3
HLA-DRB9
KSR2
TTN
ZKSCAN4
ZKSCAN8
ZKSCAN3
HERC2
SCUBE3
SPTBN2
PATJ
SOX2-OT
DGKB
ZSCAN31
TAP2

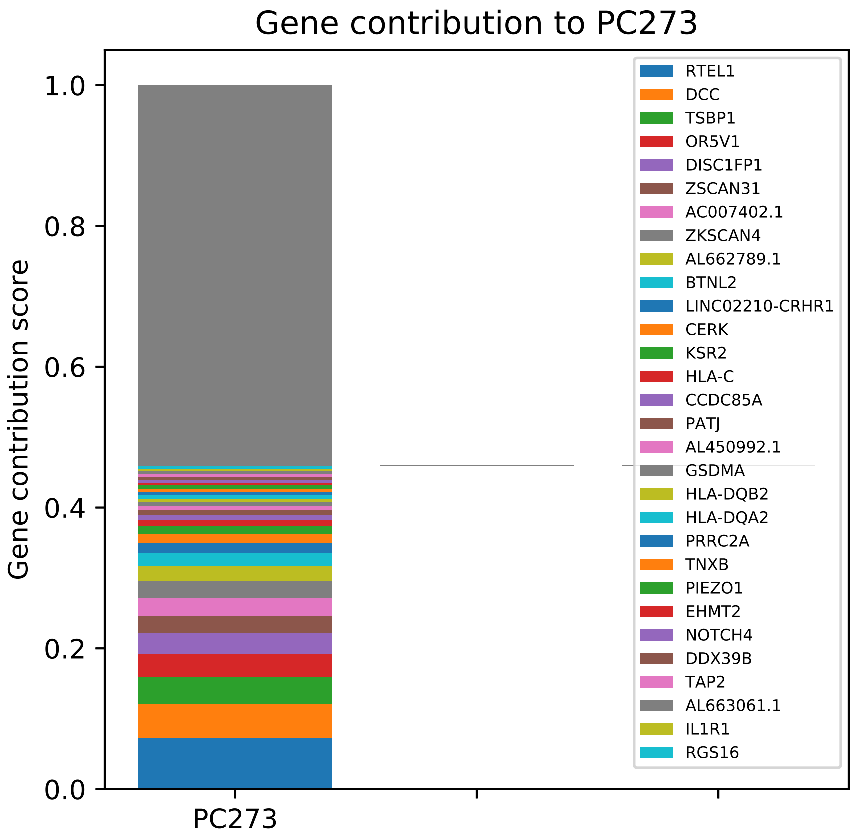

Gene contribution to PC273
Gene contribution score
0.0
0.2
0.4
0.6
0.8
1.0
PC273
RTEL1
DCC
TSBP1
OR5V1
DISC1FP1
ZSCAN31
AC007402.1
ZKSCAN4
AL662789.1
BTNL2
LINC02210-CRHR1
CERK
KSR2
HLA-C
CCDC85A
PATJ
AL450992.1
GSDMA
HLA-DQB2
HLA-DQA2
PRRC2A
TNXB
PIEZO1
EHMT2
NOTCH4
DDX39B
TAP2
AL663061.1
IL1R1
RGS16

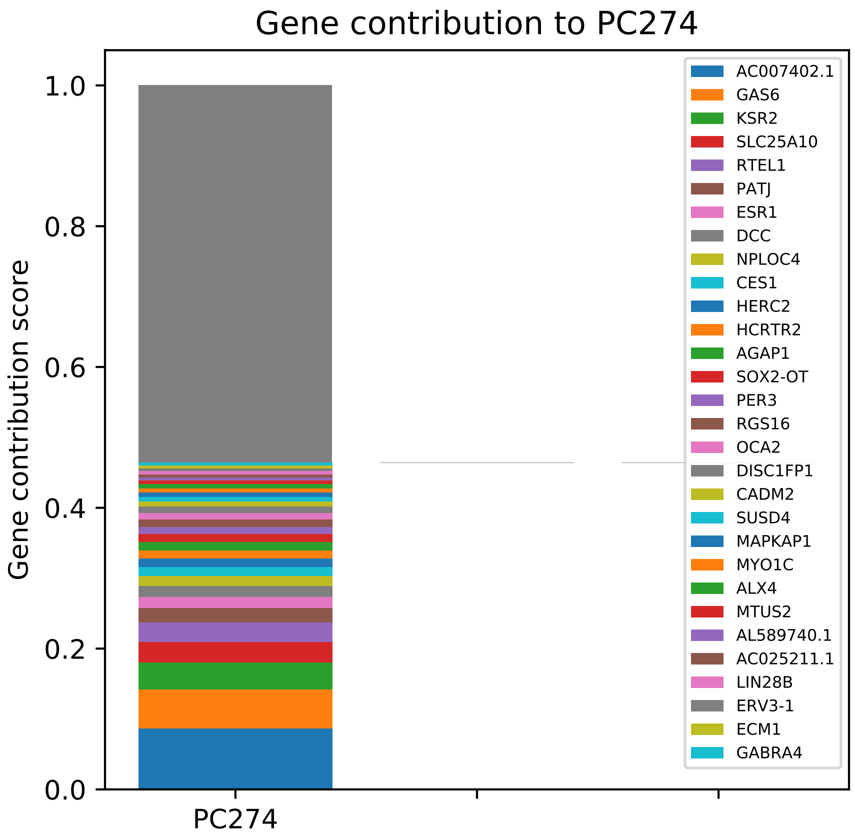
Gene contribution to PC274
Gene contribution score
1.0
0.8
0.6
0.4
0.2
0.0
PC274
AC007402.1
GAS6
KSR2
SLC25A10
RTEL1
PATJ
ESR1
DCC
NPLOC4
CES1
HERC2
HCRTR2
AGAP1
SOX2-OT
PER3
RGS16
OCA2
DISC1FP1
CADM2
SUSD4
MAPKAP1
MYO1C
ALX4
MTUS2
AL589740.1
AC025211.1
LIN28B
ERV3-1
ECM1
GABRA4

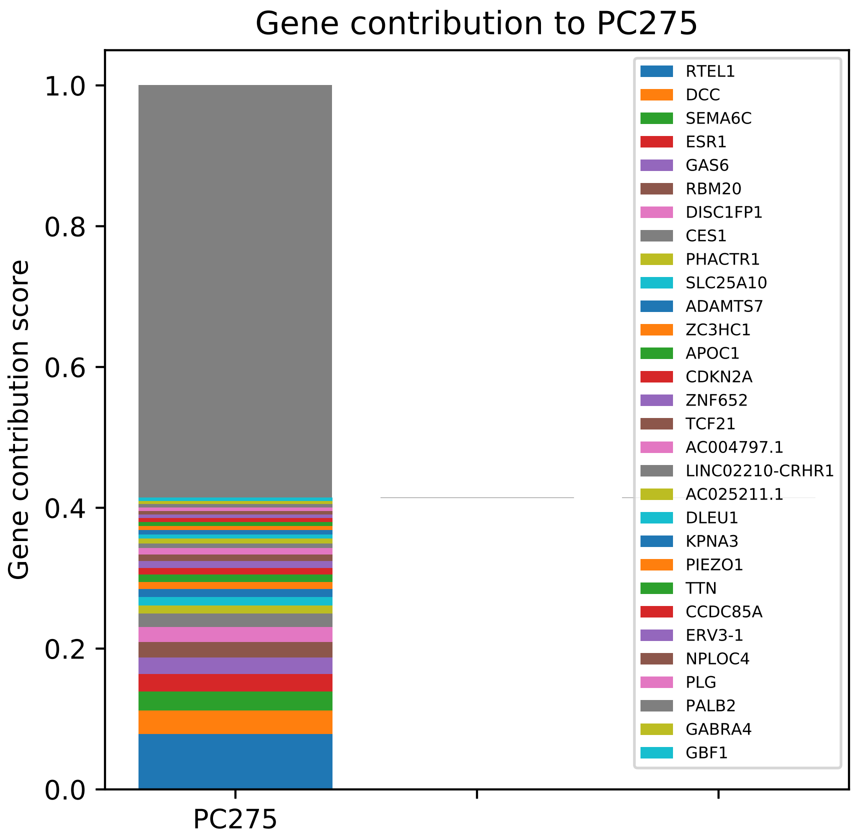
Gene contribution to PC275
Gene contribution score
1.0
0.8
0.6
0.4
0.2
0.0
PC275
RTEL1
DCC
SEMA6C
ESR1
GAS6
RBM20
DISC1FP1
CES1
PHACTR1
SLC25A10
ADAMTS7
ZC3HC1
APOC1
CDKN2A
ZNF652
TCF21
AC004797.1
LINC02210-CRHR1
AC025211.1
DLEU1
KPNA3
PIEZO1
TTN
CCDC85A
ERV3-1
NPLOC4
PLG
PALB2
GABRA4
GBF1

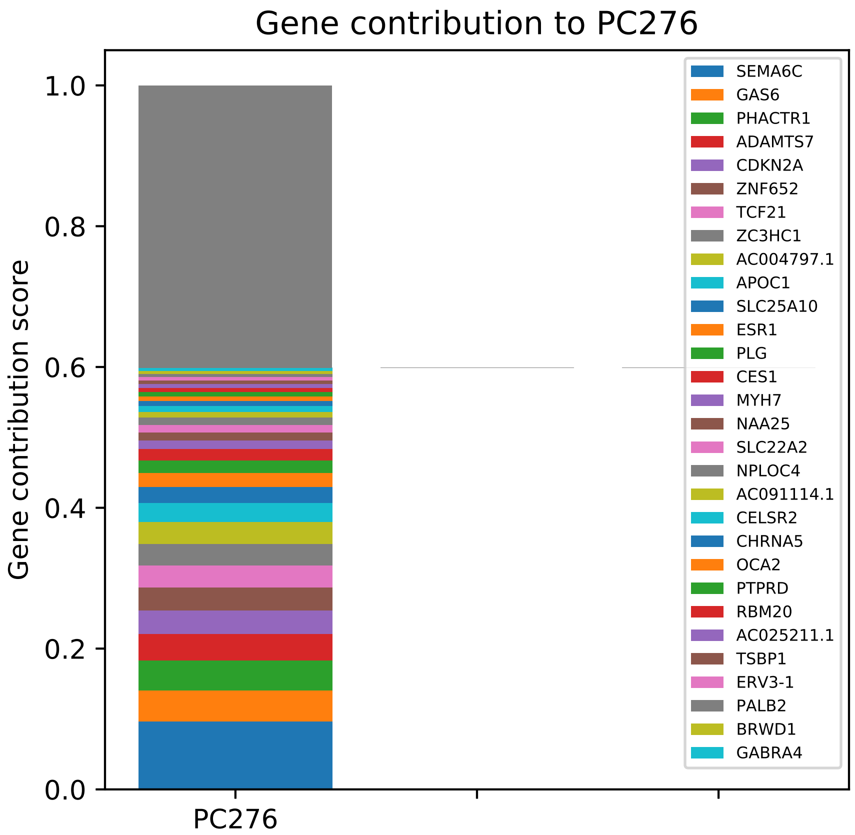

Gene contribution to PC276
Gene contribution score
1.0
0.8
0.6
0.4
0.2
0.0
PC276
SEMA6C
GAS6
PHACTR1
ADAMTS7
CDKN2A
ZNF652
TCF21
ZC3HC1
AC004797.1
APOC1
SLC25A10
ESR1
PLG
CES1
MYH7
NAA25
SLC22A2
NPLOC4
AC091114.1
CELSR2
CHRNA5
OCA2
PTPRD
RBM20
AC025211.1
TSBP1
ERV3-1
PALB2
BRWD1
GABRA4

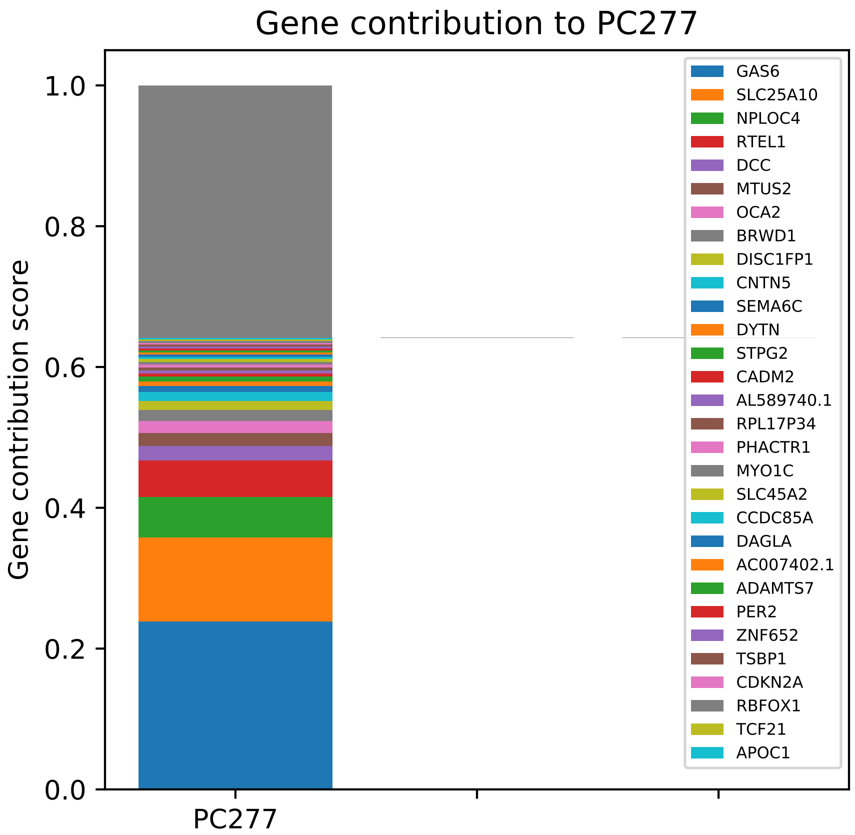

Gene contribution to PC277
Gene contribution score
1.0
0.8
0.6
0.4
0.2
0.0
PC277
GAS6
SLC25A10
NPLOC4
RTEL1
DCC
MTUS2
OCA2
BRWD1
DISC1FP1
CNTN5
SEMA6C
DYTN
STPG2
CADM2
AL589740.1
RPL17P34
PHACTR1
MYO1C
SLC45A2
CCDC85A
DAGLA
AC007402.1
ADAMTS7
PER2
ZNF652
TSBP1
CDKN2A
RBFOX1
TCF21
APOC1

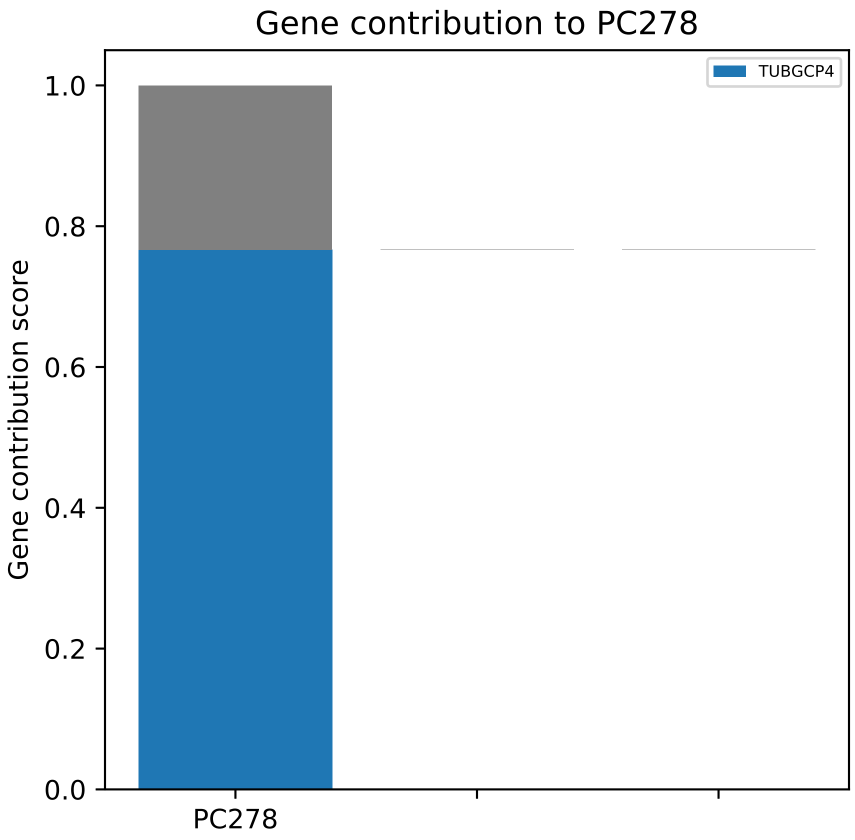

Gene contribution to PC278
TUBGCP4
1.0
0.8
0.6
0.4
0.2
0.0
Gene contribution score
PC278

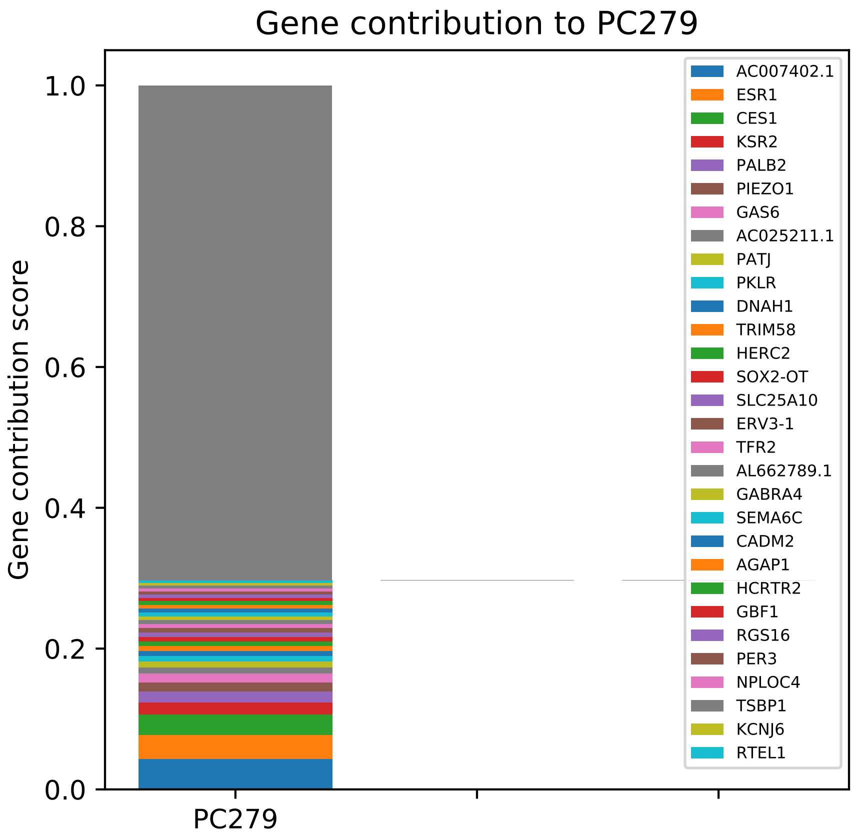
Gene contribution to PC279
Gene contribution score
1.0
0.8
0.6
0.4
0.2
0.0
PC279
AC007402.1
ESR1
CES1
KSR2
PALB2
PIEZO1
GAS6
AC025211.1
PATJ
PKLR
DNAH1
TRIM58
HERC2
SOX2-OT
SLC25A10
ERV3-1
TFR2
AL662789.1
GABRA4
SEMA6C
CADM2
AGAP1
HCRTR2
GBF1
RGS16
PER3
NPLOC4
TSBP1
KCNJ6
RTEL1

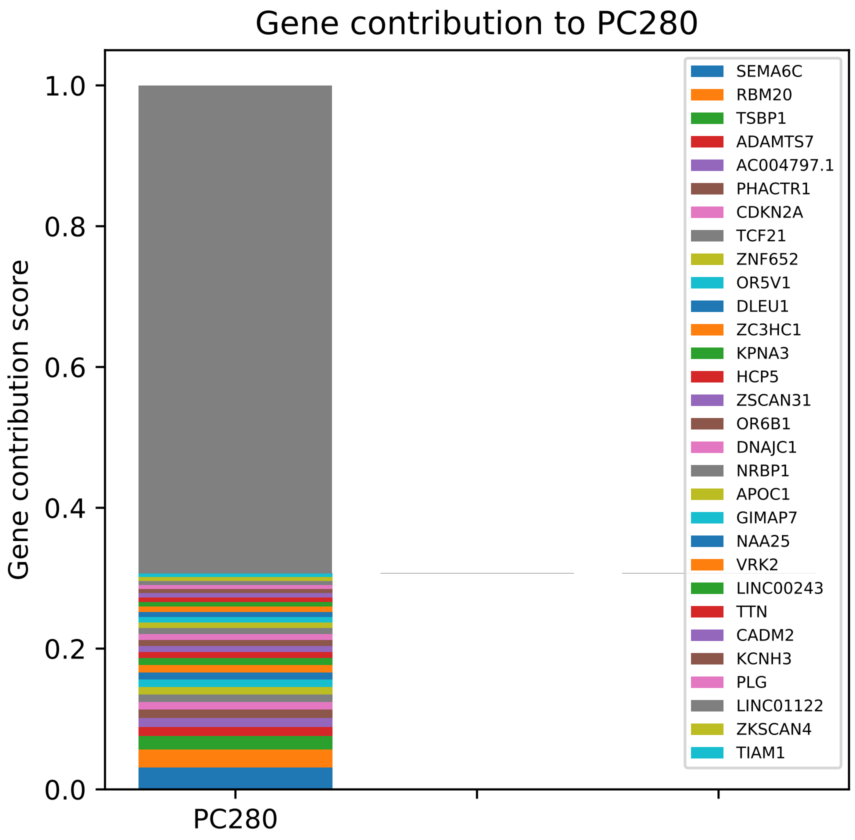

Gene contribution to PC280
Gene contribution score
1.0
0.8
0.6
0.4
0.2
0.0
PC280
SEMA6C
RBM20
TSBP1
ADAMTS7
AC004797.1
PHACTR1
CDKN2A
TCF21
ZNF652
OR5V1
DLEU1
ZC3HC1
KPNA3
HCP5
ZSCAN31
OR6B1
DNAJC1
NRBP1
APOC1
GIMAP7
NAA25
VRK2
LINC00243
TTN
CADM2
KCNH3
PLG
LINC01122
ZKSCAN4
TIAM1

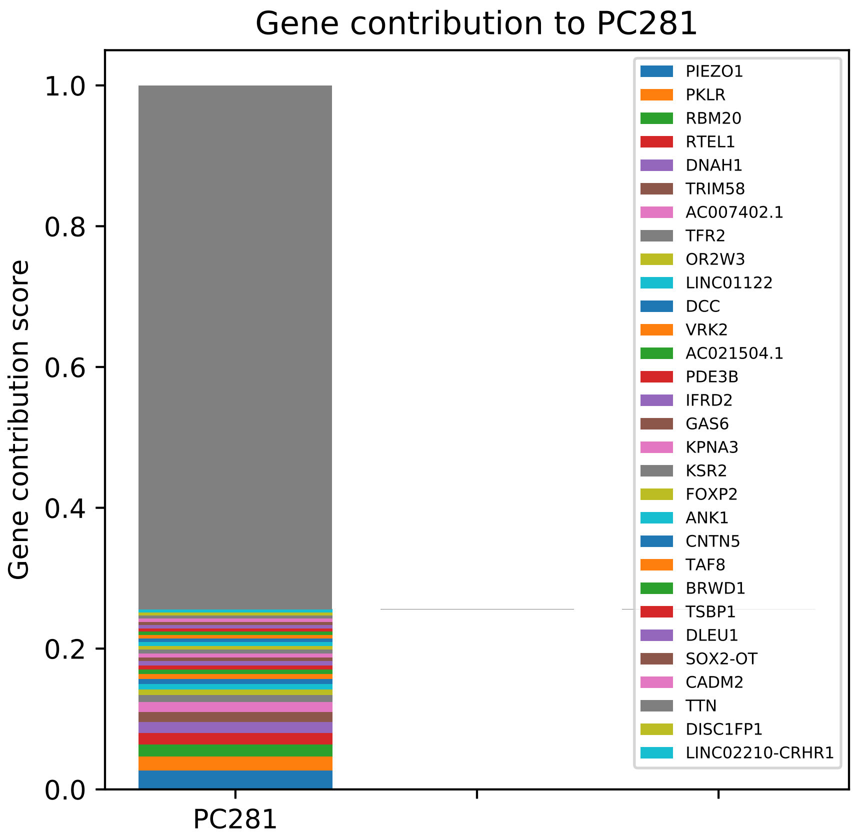

Gene contribution to PC281
Gene contribution score
1.0
0.8
0.6
0.4
0.2
0.0
PC281
PIEZO1
PKLR
RBM20
RTEL1
DNAH1
TRIM58
AC007402.1
TFR2
OR2W3
LINC01122
DCC
VRK2
AC021504.1
PDE3B
IFRD2
GAS6
KPNA3
KSR2
FOXP2
ANK1
CNTN5
TAF8
BRWD1
TSBP1
DLEU1
SOX2-OT
CADM2
TTN
DISC1FP1
LINC02210-CRHR1

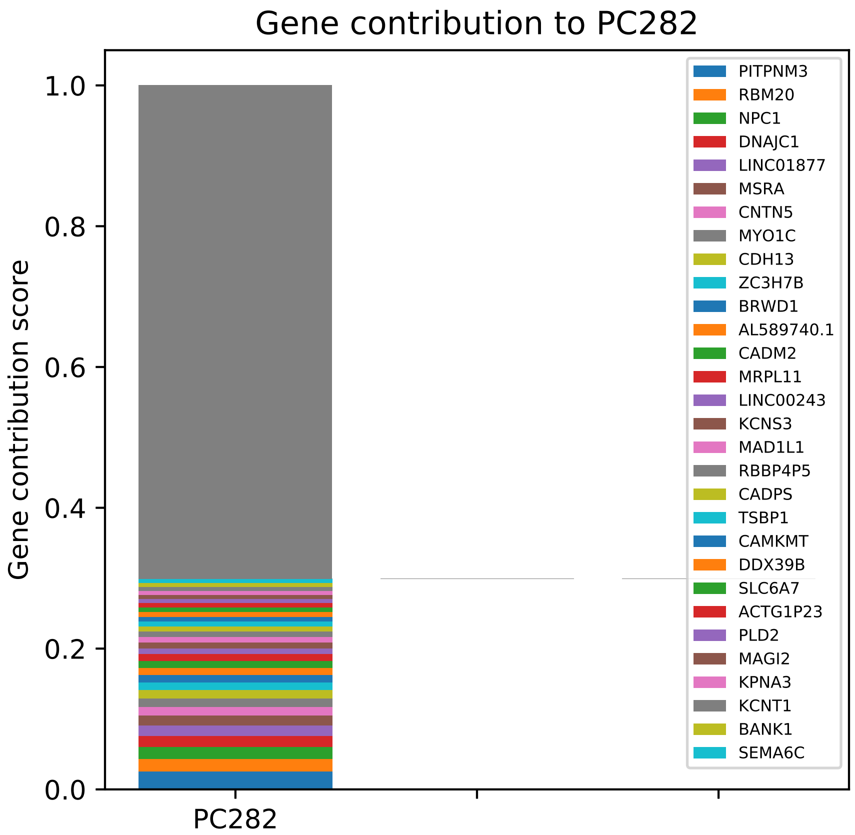

Gene contribution to PC282
Gene contribution score
1.0
0.8
0.6
0.4
0.2
0.0
PC282
PITPNM3
RBM20
NPC1
DNAJC1
LINC01877
MSRA
CNTN5
MYO1C
CDH13
ZC3H7B
BRWD1
AL589740.1
CADM2
MRPL11
LINC00243
KCNS3
MAD1L1
RBBP4P5
CADPS
TSBP1
CAMKMT
DDX39B
SLC6A7
ACTG1P23
PLD2
MAGI2
KPNA3
KCNT1
BANK1
SEMA6C

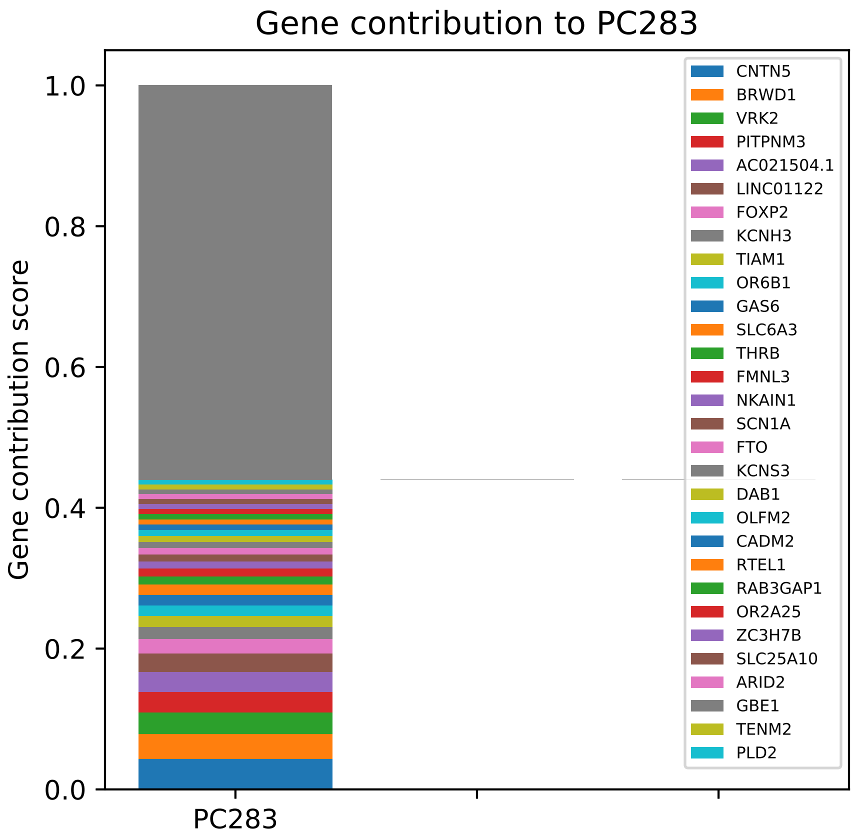
Gene contribution to PC283
Gene contribution score
1.0
0.8
0.6
0.4
0.2
0.0
PC283
CNTN5
BRWD1
VRK2
PITPNM3
AC021504.1
LINC01122
FOXP2
KCNH3
TIAM1
OR6B1
GAS6
SLC6A3
THRB
FMNL3
NKAIN1
SCN1A
FTO
KCNS3
DAB1
OLFM2
CADM2
RTEL1
RAB3GAP1
OR2A25
ZC3H7B
SLC25A10
ARID2
GBE1
TENM2
PLD2

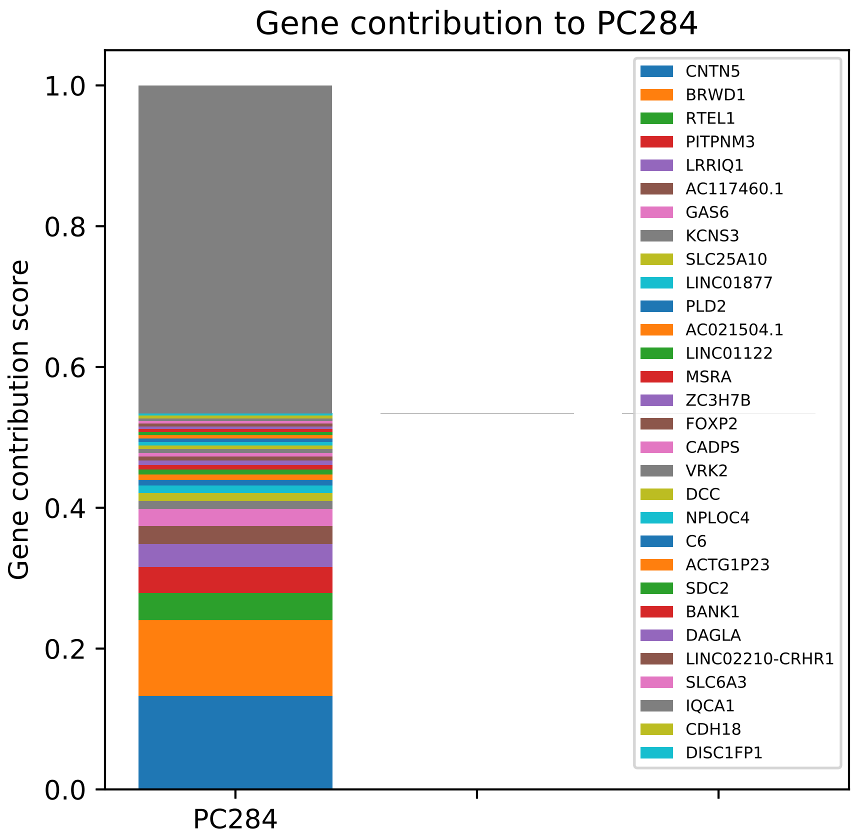
Gene contribution to PC284
Gene contribution score
0.0
0.2
0.4
0.6
0.8
1.0
PC284
CNTN5
BRWD1
RTEL1
PITPNM3
LRRIQ1
AC117460.1
GAS6
KCNS3
SLC25A10
LINC01877
PLD2
AC021504.1
LINC01122
MSRA
ZC3H7B
FOXP2
CADPS
VRK2
DCC
NPLOC4
C6
ACTG1P23
SDC2
BANK1
DAGLA
LINC02210-CRHR1
SLC6A3
IQCA1
CDH18
DISC1FP1

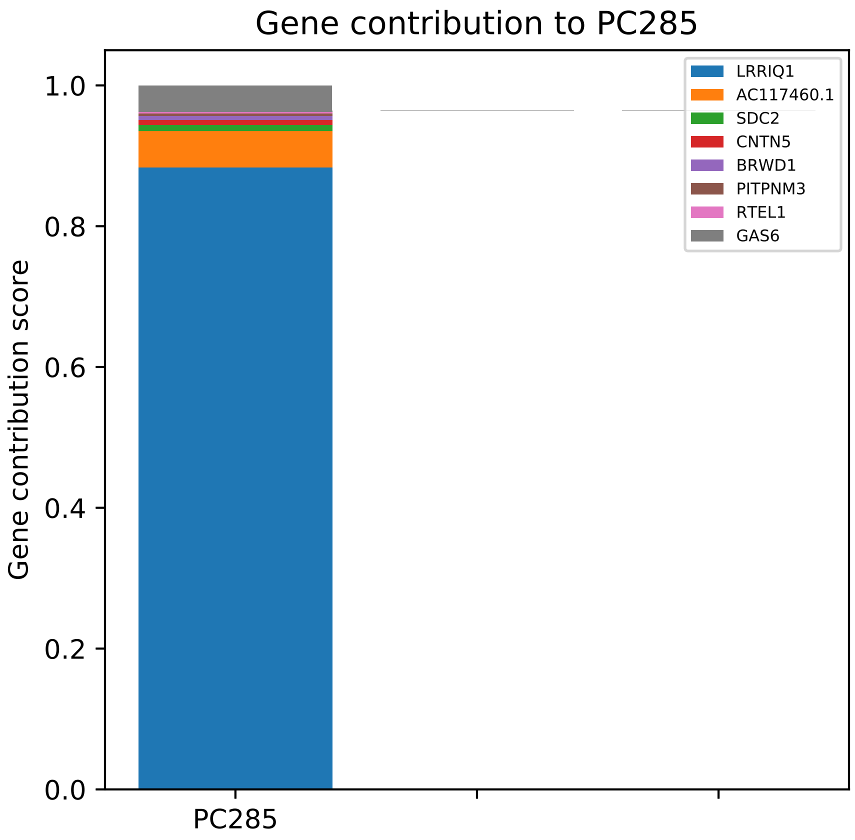

Gene contribution to PC285
Gene contribution score
1.0
0.8
0.6
0.4
0.2
0.0
PC285
LRRIQ1
AC117460.1
SDC2
CNTN5
BRWD1
PITPNM3
RTEL1
GAS6

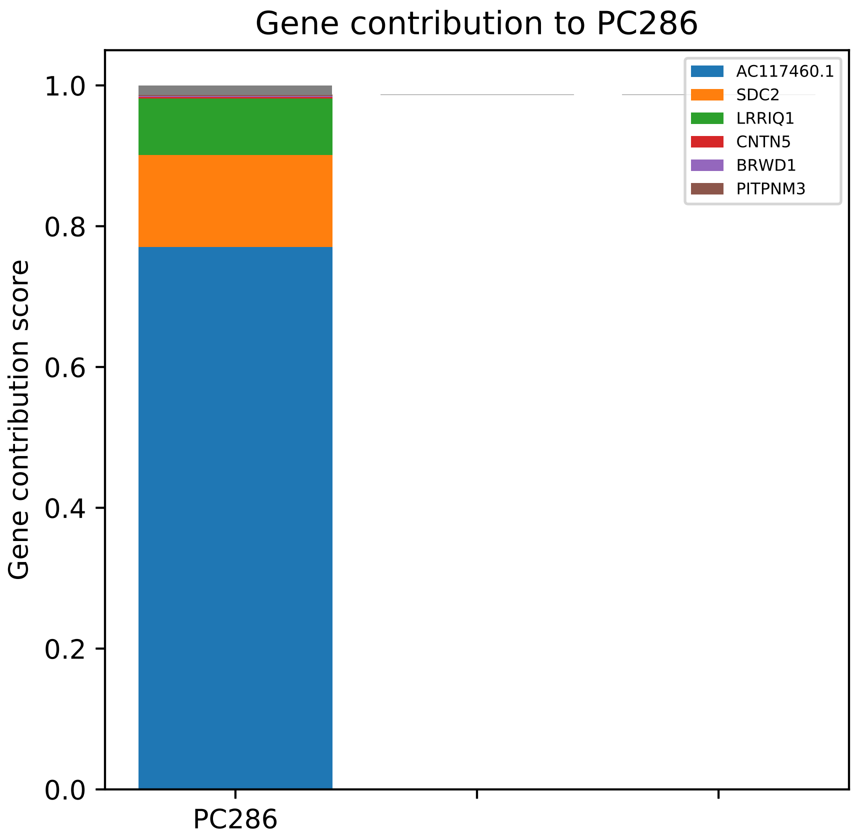

Gene contribution to PC286
Gene contribution score
1.0
0.8
0.6
0.4
0.2
0.0
PC286
AC117460.1
SDC2
LRRIQ1
CNTN5
BRWD1
PITPNM3

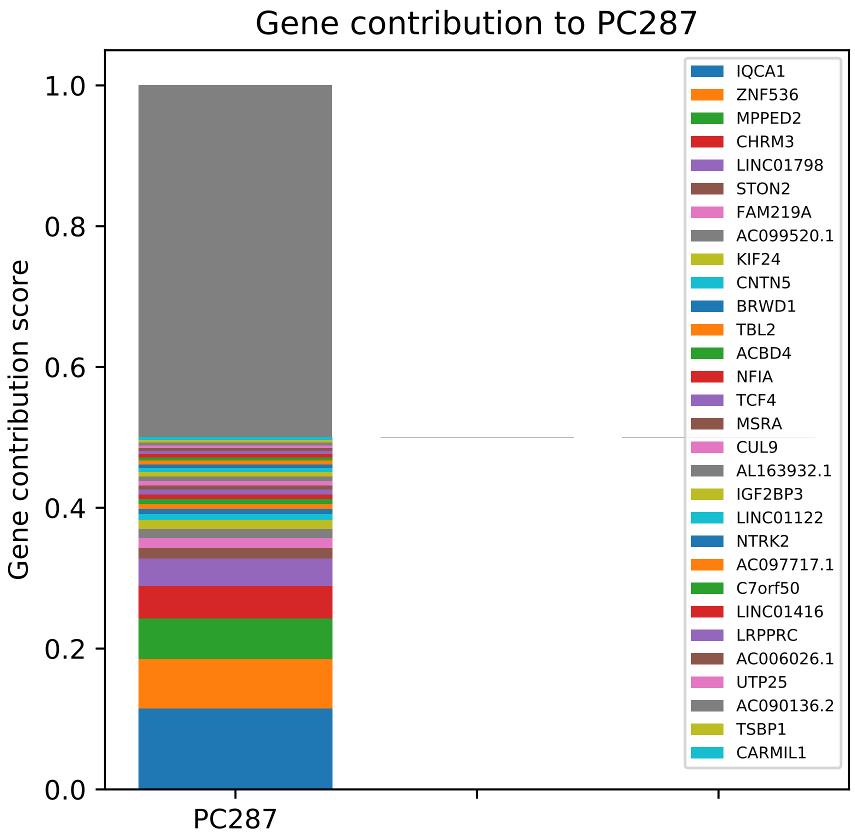
Gene contribution to PC287
Gene contribution score
1.0
0.8
0.6
0.4
0.2
0.0
PC287
IQCA1
ZNF536
MPPED2
CHRM3
LINC01798
STON2
FAM219A
AC099520.1
KIF24
CNTN5
BRWD1
TBL2
ACBD4
NFIA
TCF4
MSRA
CUL9
AL163932.1
IGF2BP3
LINC01122
NTRK2
AC097717.1
C7orf50
LINC01416
LRPPRC
AC006026.1
UTP25
AC090136.2
TSBP1
CARMIL1

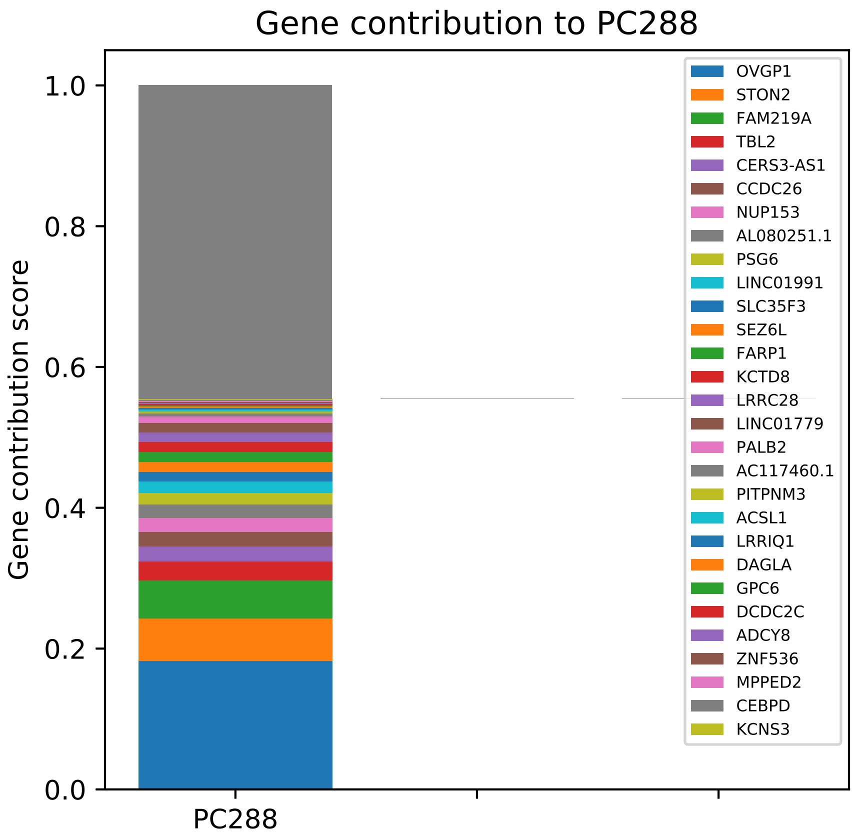

Gene contribution to PC288
Gene contribution score
0.0
0.2
0.4
0.6
0.8
1.0
PC288
OVGP1
STON2
FAM219A
TBL2
CERS3-AS1
CCDC26
NUP153
AL080251.1
PSG6
LINC01991
SLC35F3
SEZ6L
FARP1
KCTD8
LRRC28
LINC01779
PALB2
AC117460.1
PITPNM3
ACSL1
LRRIQ1
DAGLA
GPC6
DCDC2C
ADCY8
ZNF536
MPPED2
CEBPD
KCNS3

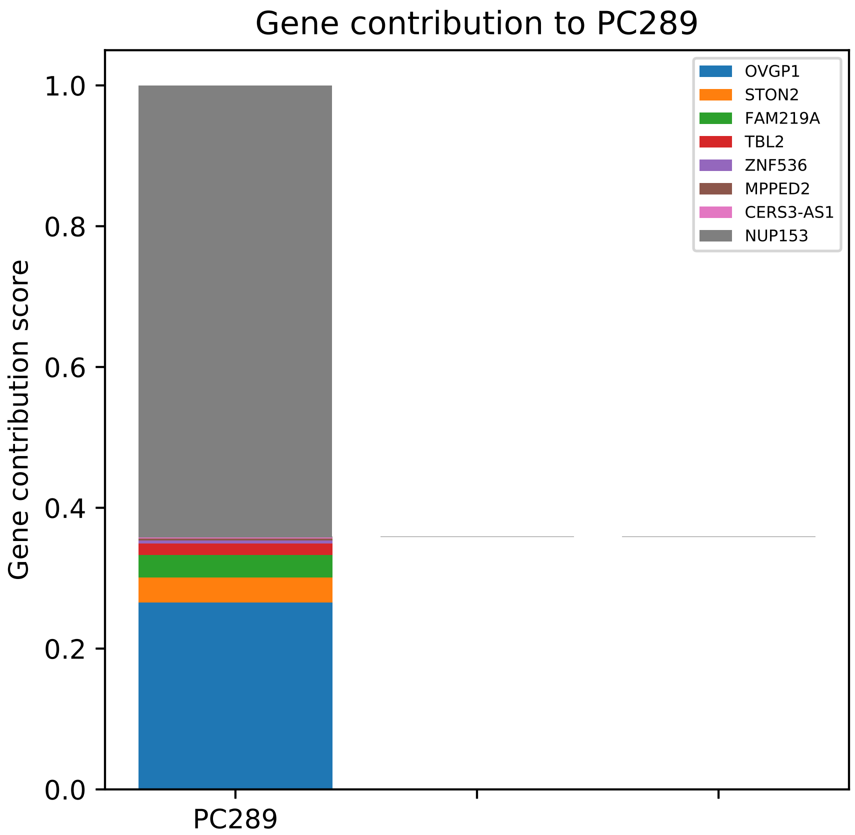

Gene contribution to PC289
Gene contribution score
1.0
0.8
0.6
0.4
0.2
0.0
PC289
OVGP1
STON2
FAM219A
TBL2
ZNF536
MPPED2
CERS3-AS1
NUP153

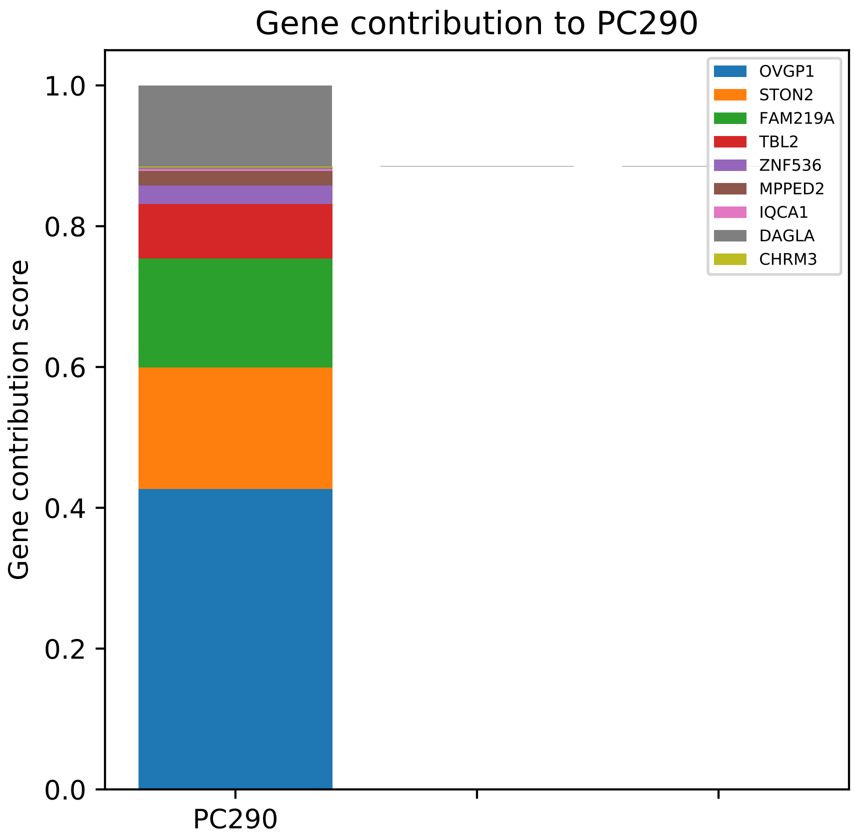
Gene contribution to PC290
Gene contribution score
1.0
0.8
0.6
0.4
0.2
0.0
PC290
OVGP1
STON2
FAM219A
TBL2
ZNF536
MPPED2
IQCA1
DAGLA
CHRM3

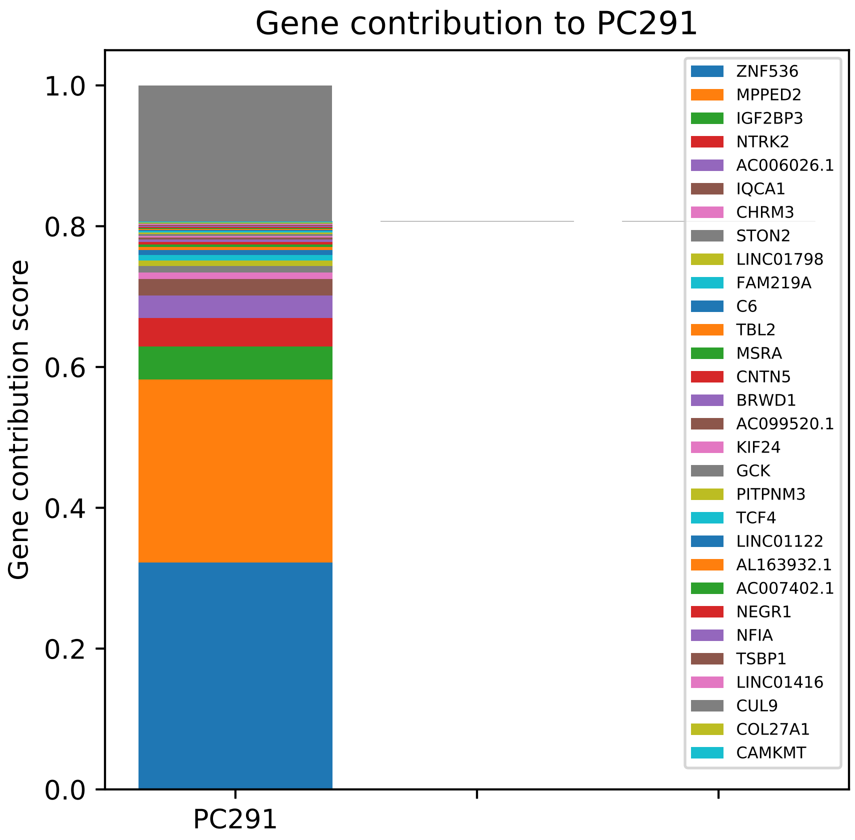

Gene contribution to PC291
Gene contribution score
1.0
0.8
0.6
0.4
0.2
0.0
PC291
ZNF536
MPPED2
IGF2BP3
NTRK2
AC006026.1
IQCA1
CHRM3
STON2
LINC01798
FAM219A
C6
TBL2
MSRA
CNTN5
BRWD1
AC099520.1
KIF24
GCK
PITPNM3
TCF4
LINC01122
AL163932.1
AC007402.1
NEGR1
NFIA
TSBP1
LINC01416
CUL9
COL27A1
CAMKMT

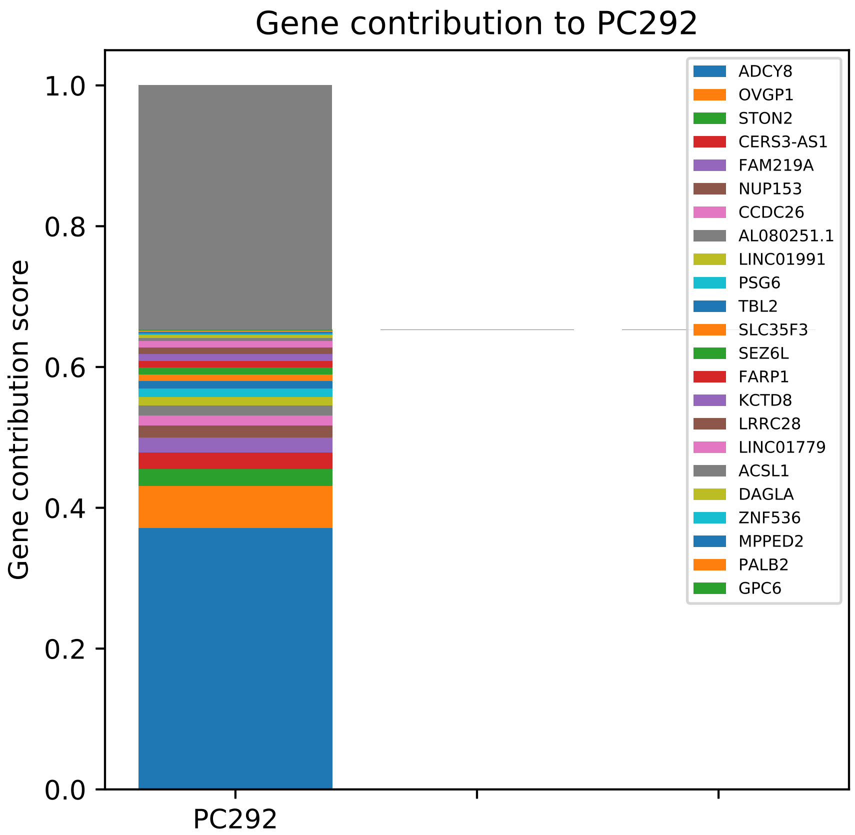

Gene contribution to PC292
Gene contribution score
1.0
0.8
0.6
0.4
0.2
0.0
PC292
ADCY8
OVGP1
STON2
CERS3-AS1
FAM219A
NUP153
CCDC26
AL080251.1
LINC01991
PSG6
TBL2
SLC35F3
SEZ6L
FARP1
KCTD8
LRRC28
LINC01779
ACSL1
DAGLA
ZNF536
MPPED2
PALB2
GPC6

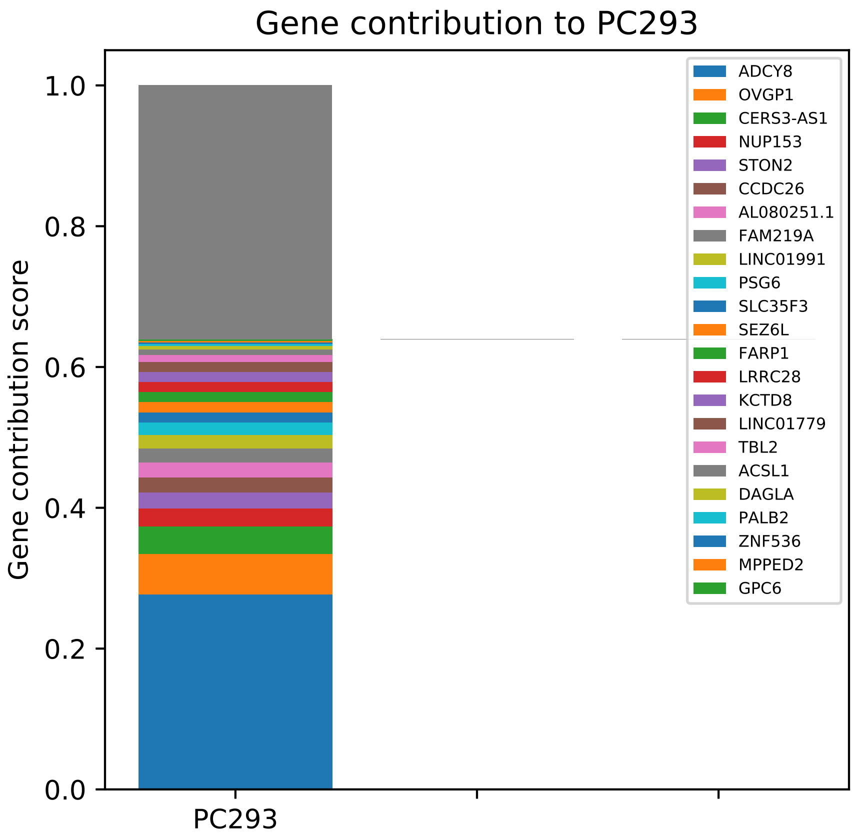

Gene contribution to PC293
Gene contribution score
1.0
0.8
0.6
0.4
0.2
0.0
PC293
ADCY8
OVGP1
CERS3-AS1
NUP153
STON2
CCDC26
AL080251.1
FAM219A
LINC01991
PSG6
SLC35F3
SEZ6L
FARP1
LRRC28
KCTD8
LINC01779
TBL2
ACSL1
DAGLA
PALB2
ZNF536
MPPED2
GPC6

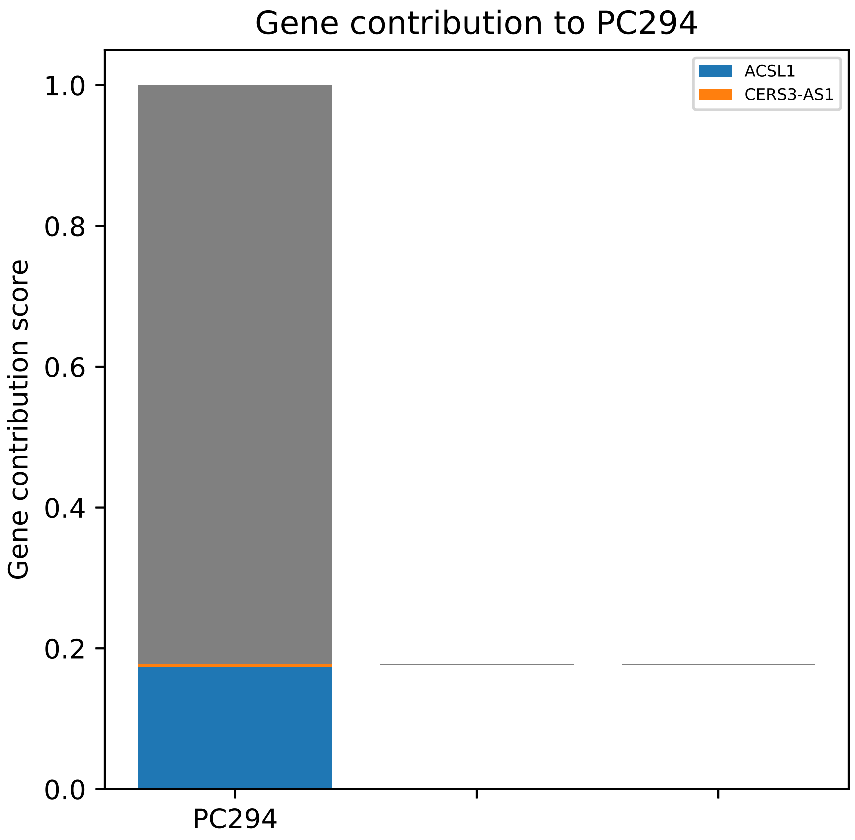

Gene contribution to PC294
Gene contribution score
1.0
0.8
0.6
0.4
0.2
0.0
PC294
ACSL1
CERS3-AS1

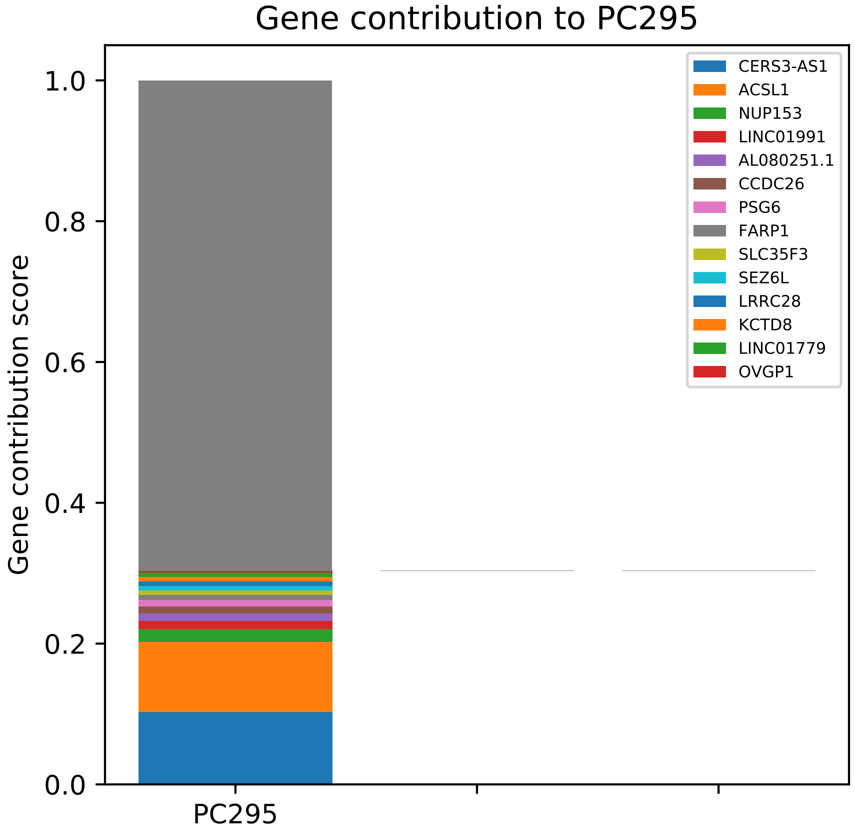

Gene contribution to PC295
Gene contribution score
1.0
0.8
0.6
0.4
0.2
0.0
PC295
CERS3-AS1
ACSL1
NUP153
LINC01991
AL080251.1
CCDC26
PSG6
FARP1
SLC35F3
SEZ6L
LRRC28
KCTD8
LINC01779
OVGP1

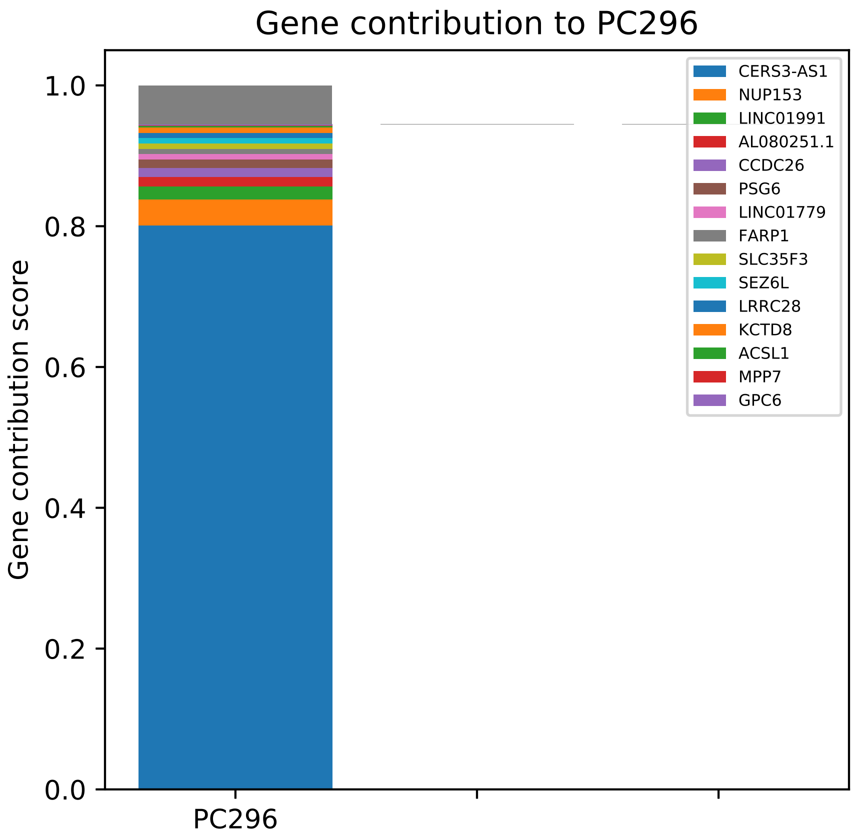

Gene contribution to PC296
Gene contribution score
0.0
0.2
0.4
0.6
0.8
1.0
PC296
CERS3-AS1
NUP153
LINC01991
AL080251.1
CCDC26
PSG6
LINC01779
FARP1
SLC35F3
SEZ6L
LRRC28
KCTD8
ACSL1
MPP7
GPC6

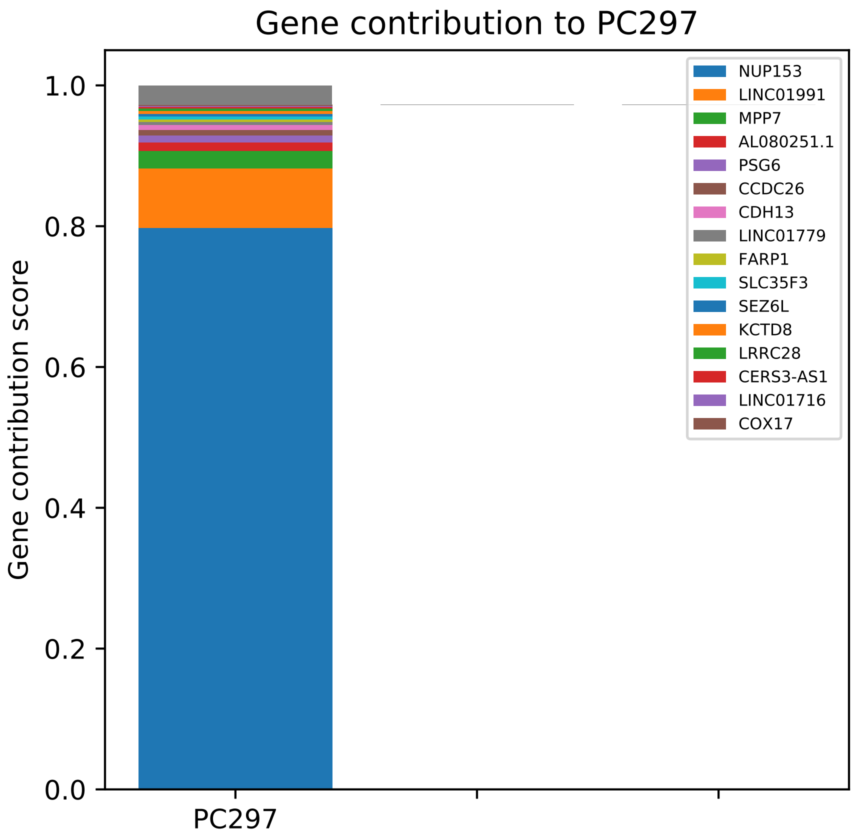

Gene contribution to PC297
Gene contribution score
1.0
0.8
0.6
0.4
0.2
0.0
PC297
NUP153
LINC01991
MPP7
AL080251.1
PSG6
CCDC26
CDH13
LINC01779
FARP1
SLC35F3
SEZ6L
KCTD8
LRRC28
CERS3-AS1
LINC01716
COX17

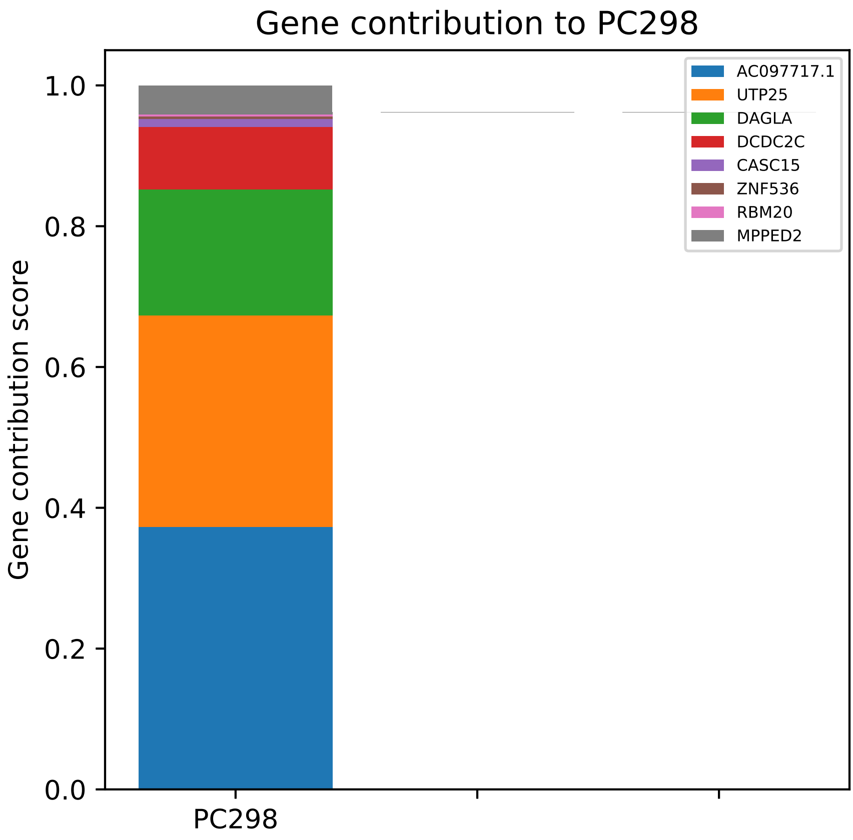

Gene contribution to PC298
Gene contribution score
0.0
0.2
0.4
0.6
0.8
1.0
PC298
AC097717.1
UTP25
DAGLA
DCDC2C
CASC15
ZNF536
RBM20
MPPED2

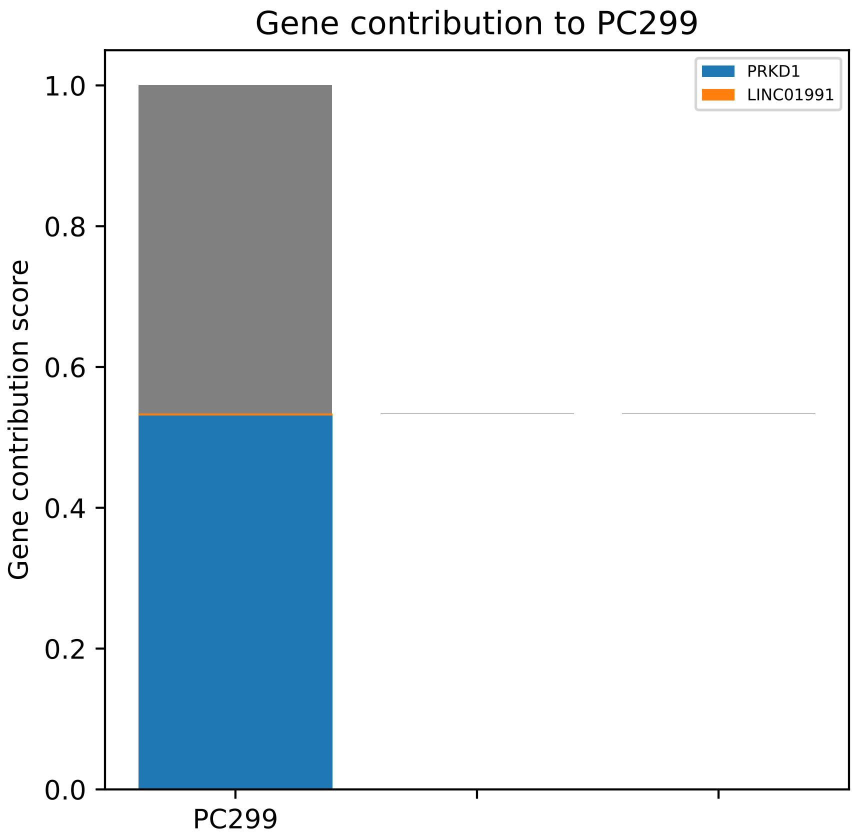
Gene contribution to PC299
PRKD1
LINC01991
Gene contribution score
1.0
0.8
0.6
0.4
0.2
0.0
PC299

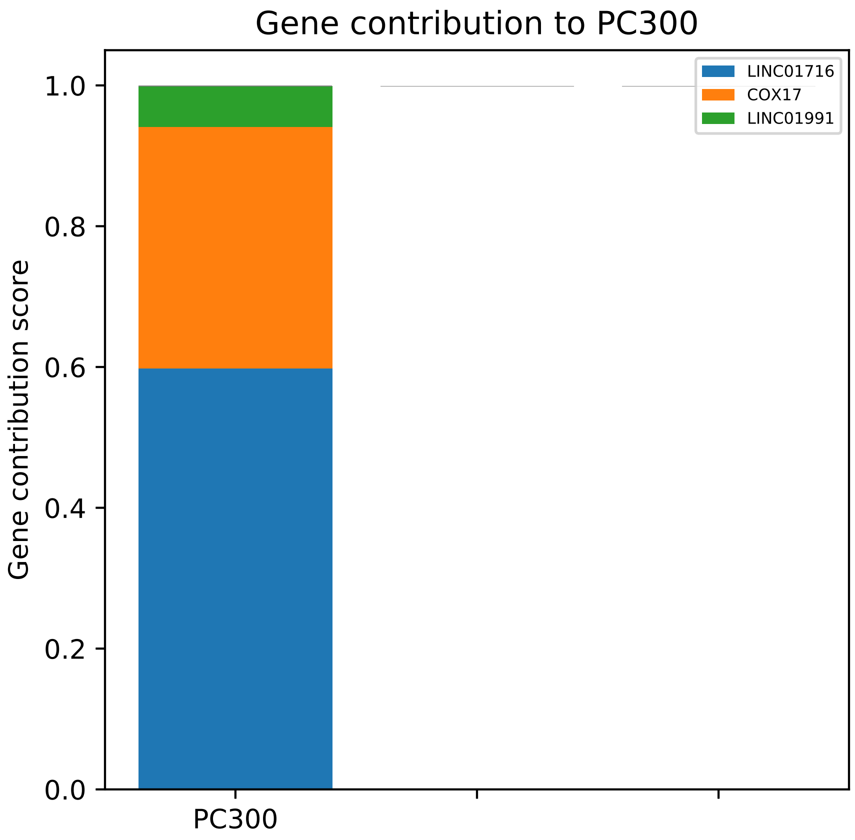

Gene contribution to PC300
Gene contribution score
1.0
0.8
0.6
0.4
0.2
0.0
PC300
LINC01716
COX17
LINC01991

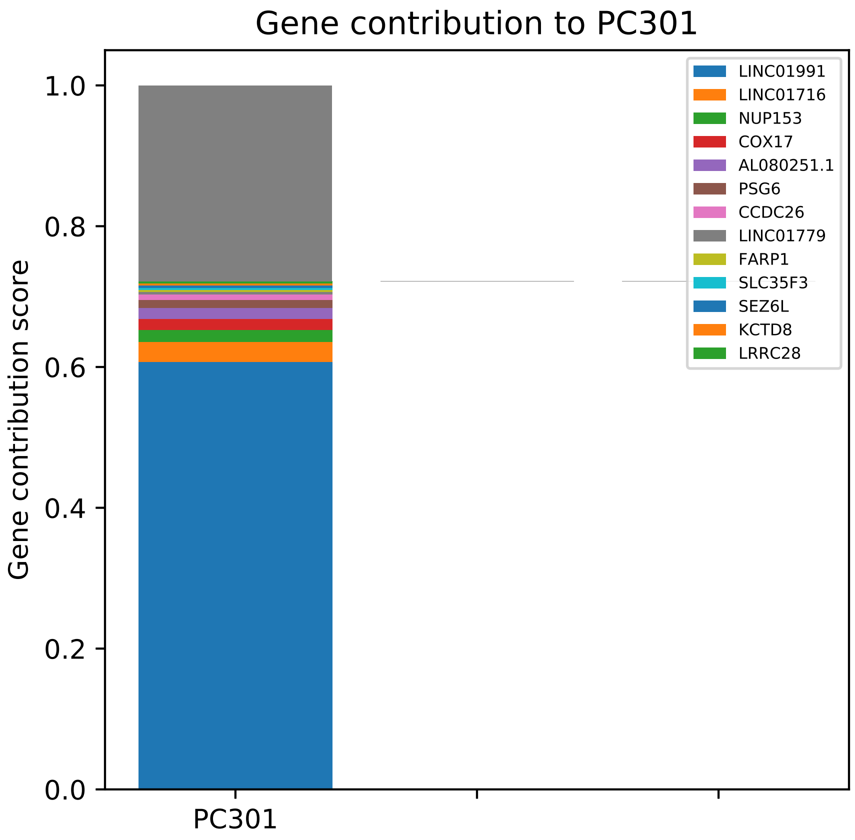
Gene contribution to PC301
Gene contribution score
1.0
0.8
0.6
0.4
0.2
0.0
PC301
LINC01991
LINC01716
NUP153
COX17
AL080251.1
PSG6
CCDC26
LINC01779
FARP1
SLC35F3
SEZ6L
KCTD8
LRRC28

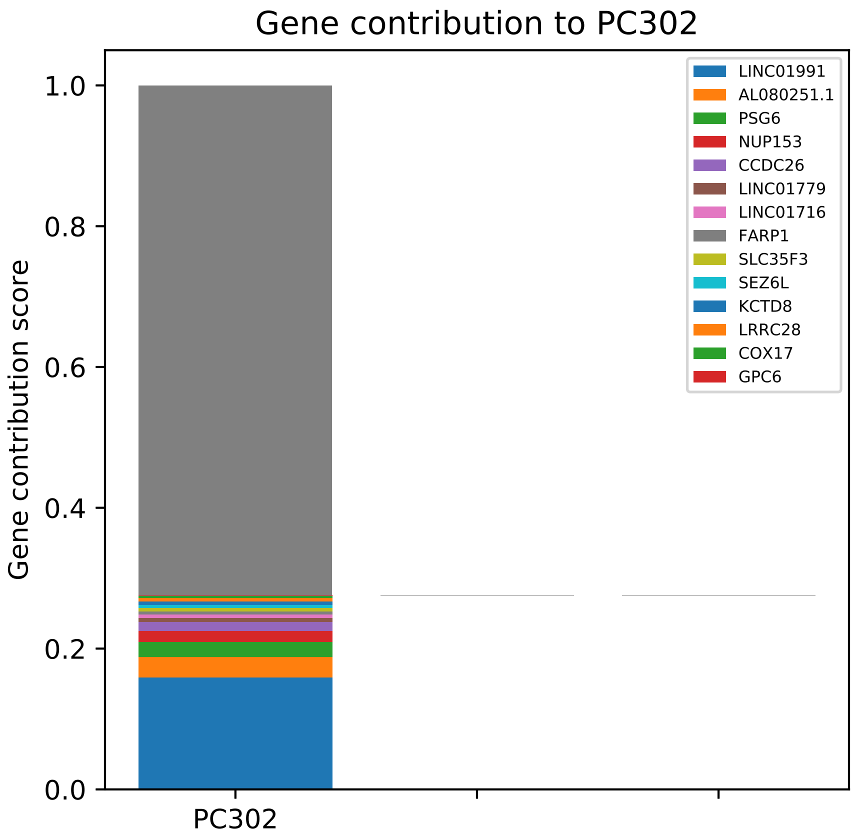
Gene contribution to PC302
Gene contribution score
1.0
0.8
0.6
0.4
0.2
0.0
PC302
LINC01991
AL080251.1
PSG6
NUP153
CCDC26
LINC01779
LINC01716
FARP1
SLC35F3
SEZ6L
KCTD8
LRRC28
COX17
GPC6

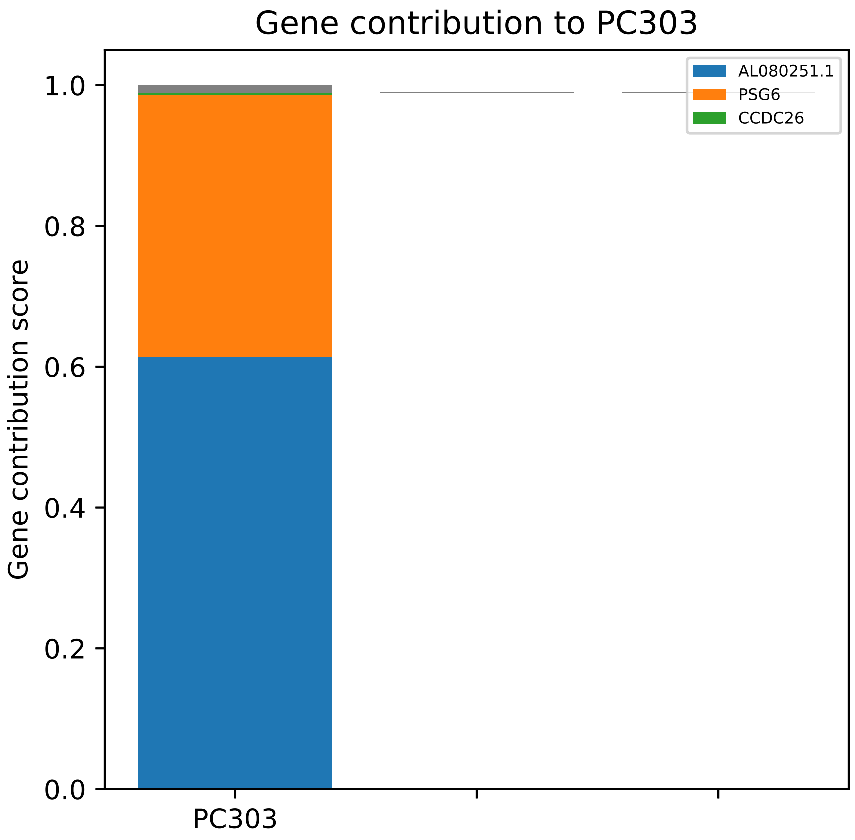

Gene contribution to PC303
Gene contribution score
1.0
0.8
0.6
0.4
0.2
0.0
PC303
AL080251.1
PSG6
CCDC26

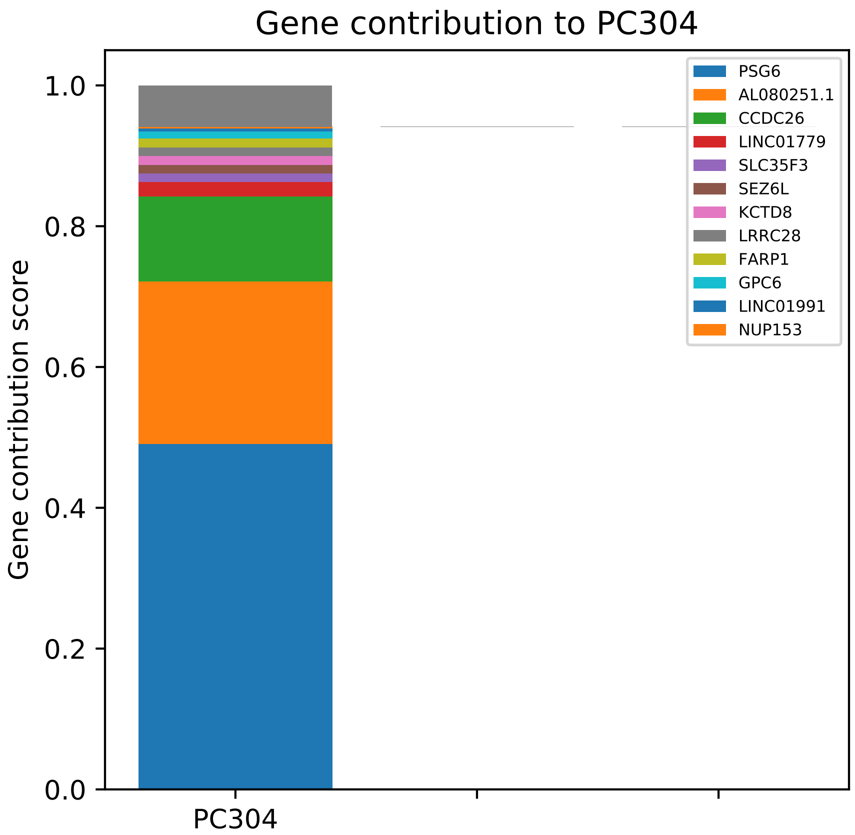

Gene contribution to PC304
Gene contribution score
0.0
0.2
0.4
0.6
0.8
1.0
PC304
PSG6
AL080251.1
CCDC26
LINC01779
SLC35F3
SEZ6L
KCTD8
LRRC28
FARP1
GPC6
LINC01991
NUP153

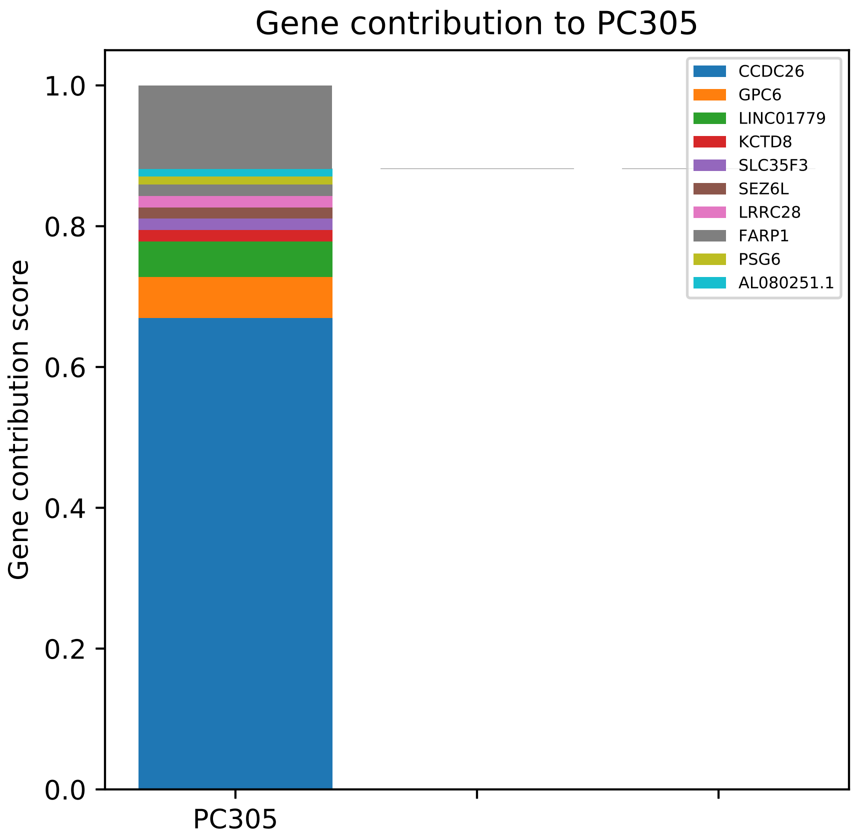

Gene contribution to PC305
Gene contribution score
1.0
0.8
0.6
0.4
0.2
0.0
PC305
CCDC26
GPC6
LINC01779
KCTD8
SLC35F3
SEZ6L
LRRC28
FARP1
PSG6
AL080251.1

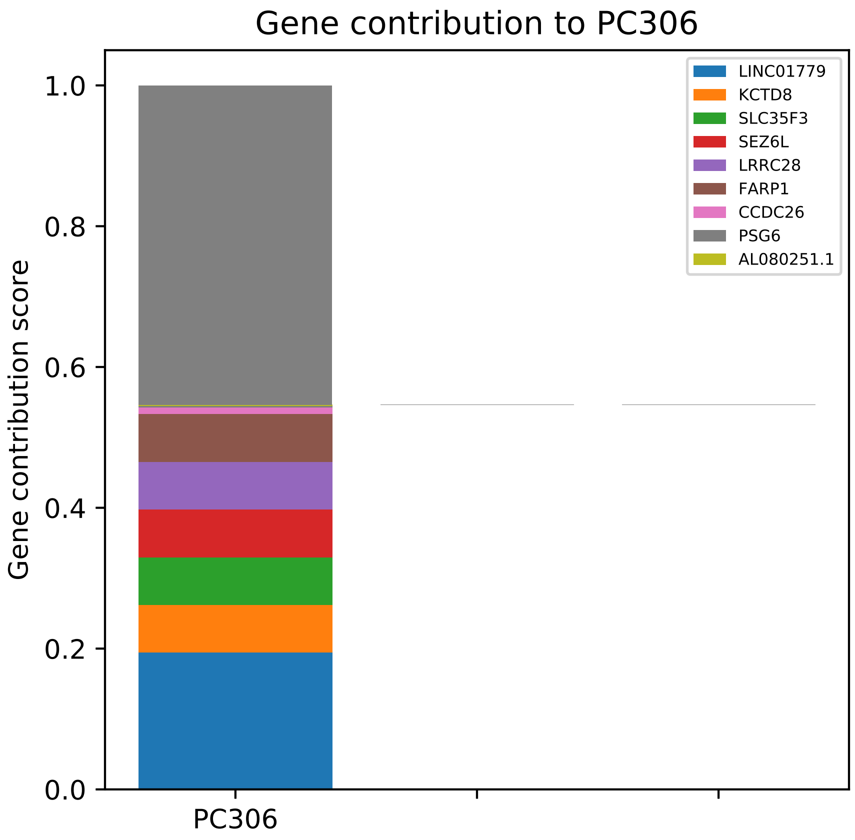

Gene contribution to PC306
Gene contribution score
1.0
0.8
0.6
0.4
0.2
0.0
PC306
LINC01779
KCTD8
SLC35F3
SEZ6L
LRRC28
FARP1
CCDC26
PSG6
AL080251.1

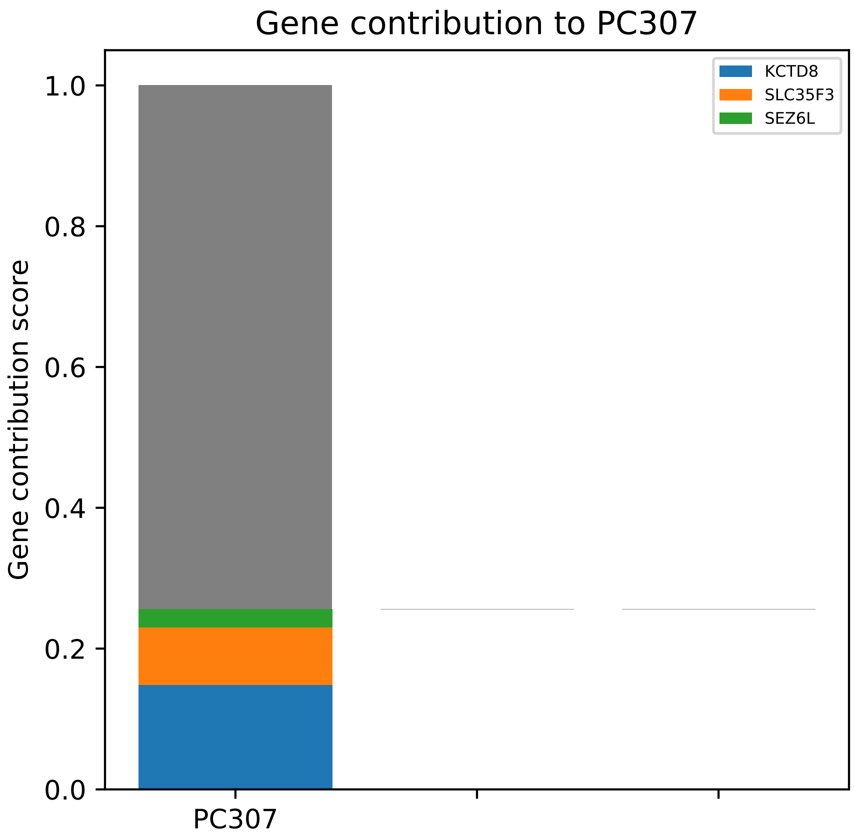

Gene contribution to PC307
Gene contribution score
1.0
0.8
0.6
0.4
0.2
0.0
PC307
KCTD8
SLC35F3
SEZ6L

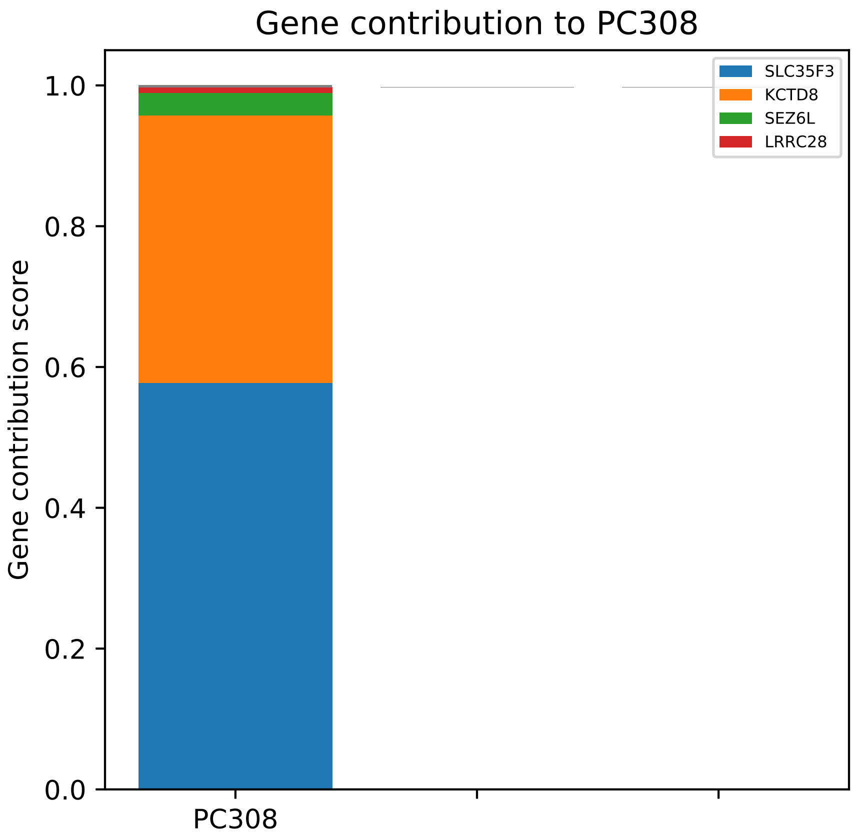

Gene contribution to PC308
Gene contribution score
1.0
0.8
0.6
0.4
0.2
0.0
PC308
SLC35F3
KCTD8
SEZ6L
LRRC28

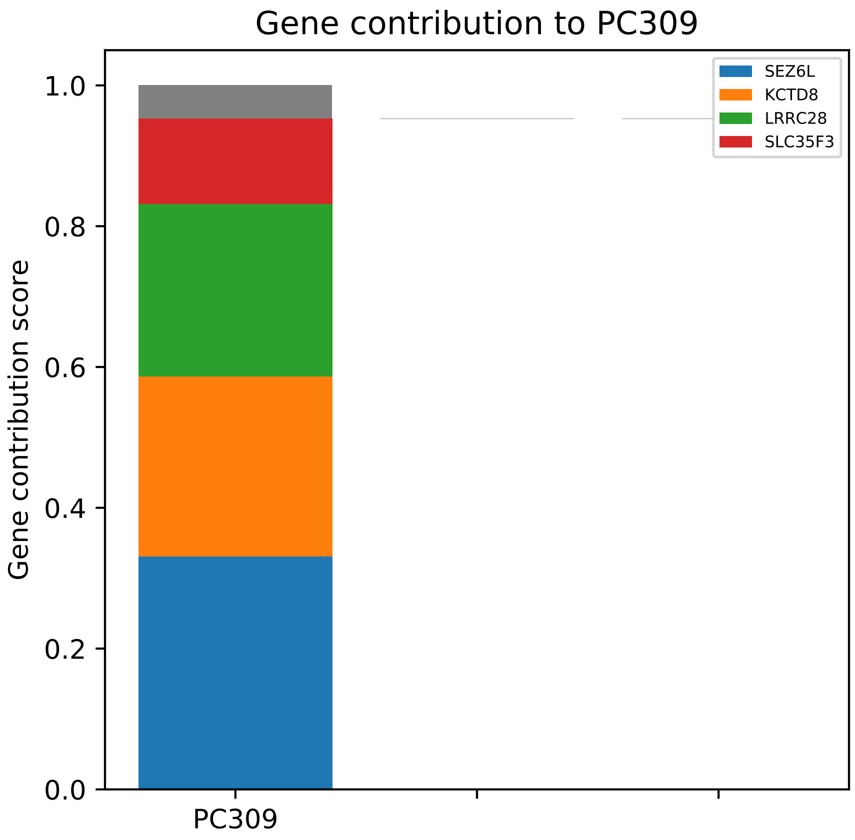

Gene contribution to PC309
Gene contribution score
1.0
0.8
0.6
0.4
0.2
0.0
PC309
SEZ6L
KCTD8
LRRC28
SLC35F3

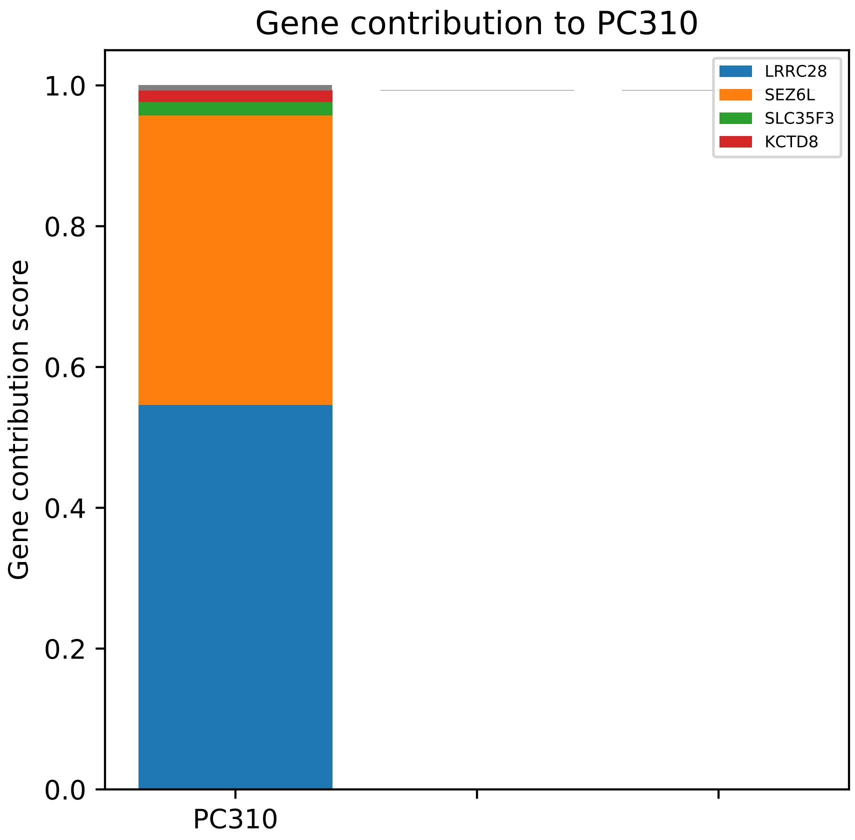
Gene contribution to PC310
Gene contribution score
1.0
0.8
0.6
0.4
0.2
0.0
PC310
LRRC28
SEZ6L
SLC35F3
KCTD8

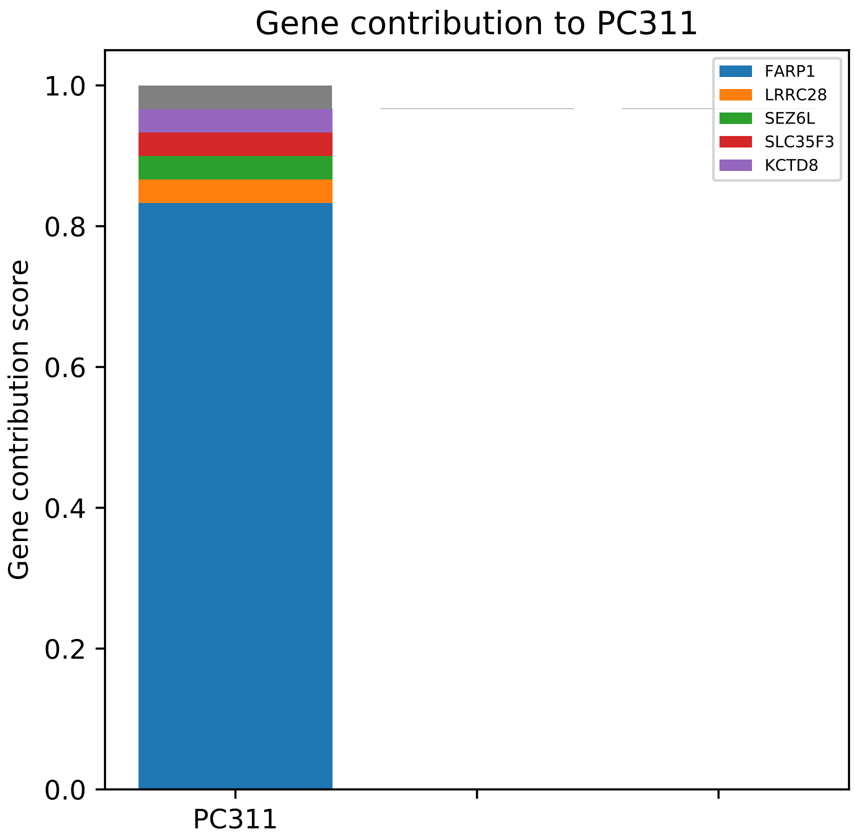
Gene contribution to PC311
Gene contribution score
1.0
0.8
0.6
0.4
0.2
0.0
PC311
FARP1
LRRC28
SEZ6L
SLC35F3
KCTD8

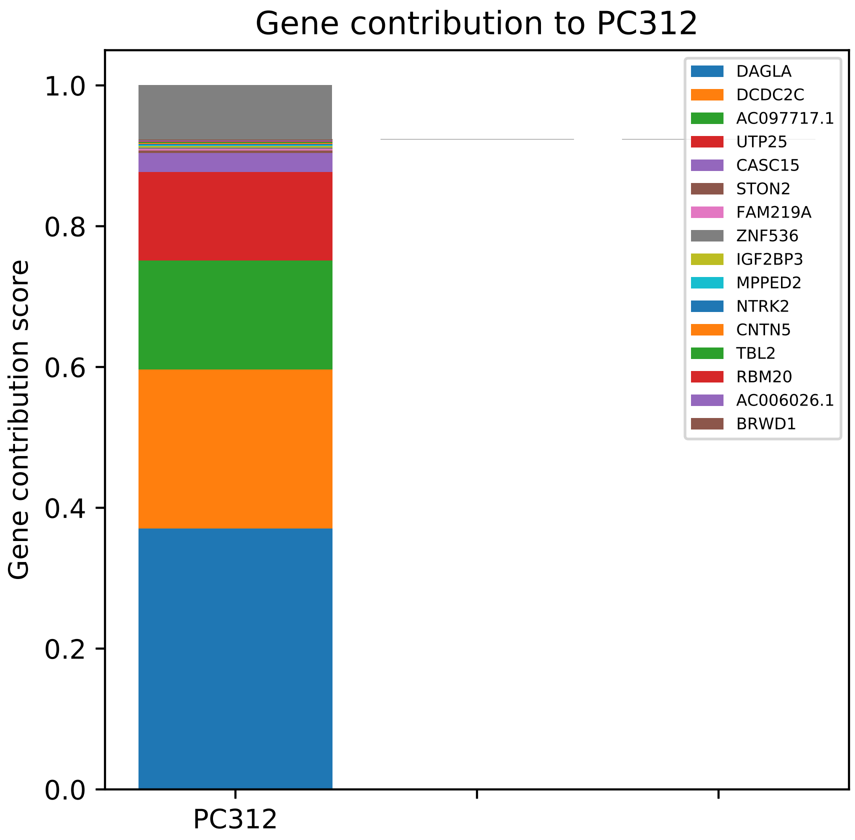

Gene contribution to PC312
Gene contribution score
1.0
0.8
0.6
0.4
0.2
0.0
PC312
DAGLA
DCDC2C
AC097717.1
UTP25
CASC15
STON2
FAM219A
ZNF536
IGF2BP3
MPPED2
NTRK2
CNTN5
TBL2
RBM20
AC006026.1
BRWD1

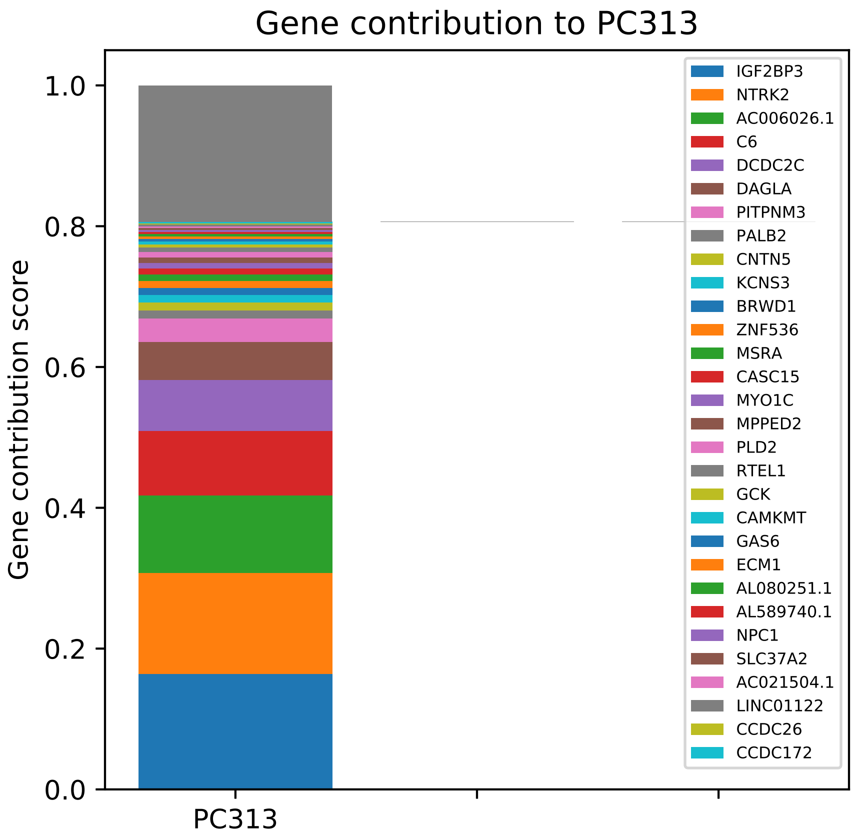

Gene contribution to PC313
Gene contribution score
1.0
0.8
0.6
0.4
0.2
0.0
PC313
IGF2BP3
NTRK2
AC006026.1
C6
DCDC2C
DAGLA
PITPNM3
PALB2
CNTN5
KCNS3
BRWD1
ZNF536
MSRA
CASC15
MYO1C
MPPED2
PLD2
RTEL1
GCK
CAMKMT
GAS6
ECM1
AL080251.1
AL589740.1
NPC1
SLC37A2
AC021504.1
LINC01122
CCDC26
CCDC172

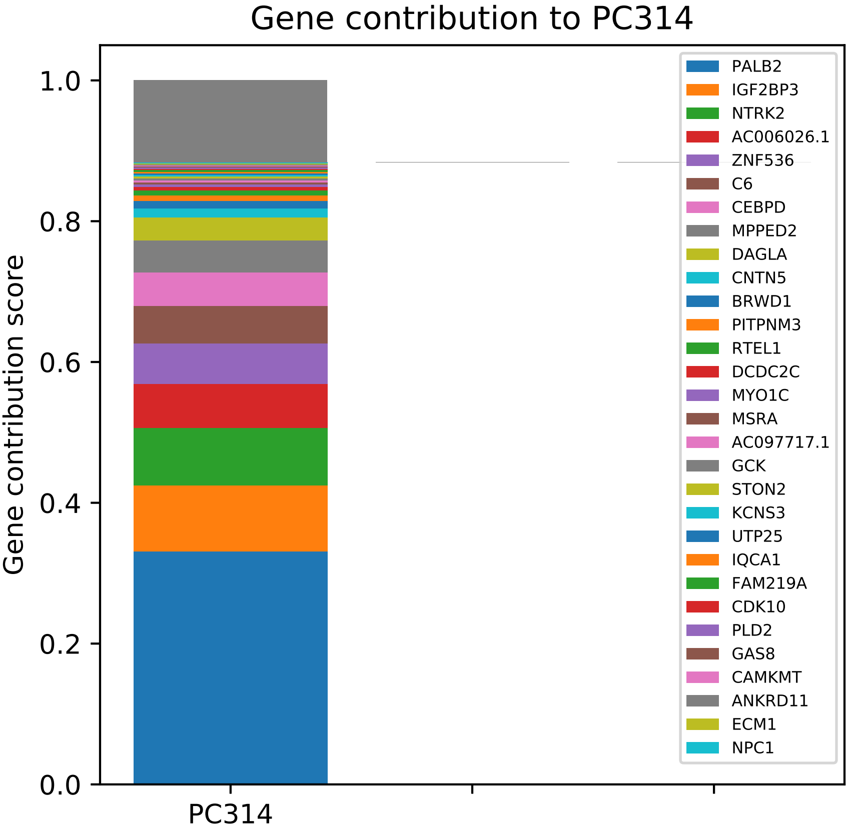
Gene contribution to PC314
Gene contribution score
1.0
0.8
0.6
0.4
0.2
0.0
PC314
PALB2
IGF2BP3
NTRK2
AC006026.1
ZNF536
C6
CEBPD
MPPED2
DAGLA
CNTN5
BRWD1
PITPNM3
RTEL1
DCDC2C
MYO1C
MSRA
AC097717.1
GCK
STON2
KCNS3
UTP25
IQCA1
FAM219A
CDK10
PLD2
GAS8
CAMKMT
ANKRD11
ECM1
NPC1

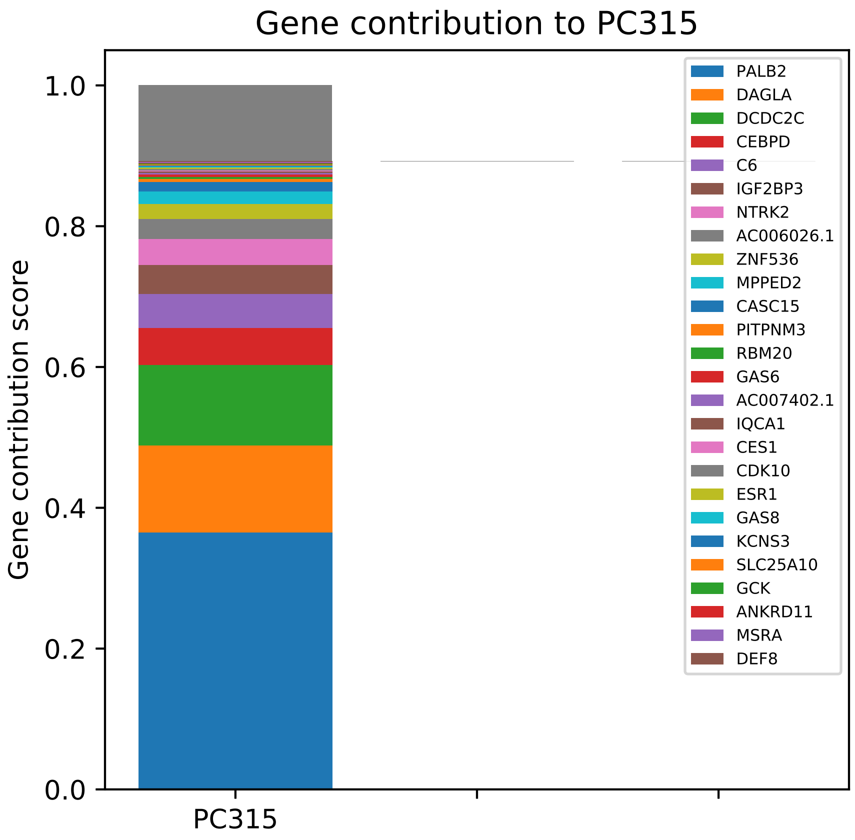
Gene contribution to PC315
Gene contribution score
1.0
0.8
0.6
0.4
0.2
0.0
PC315
PALB2
DAGLA
DCDC2C
CEBPD
C6
IGF2BP3
NTRK2
AC006026.1
ZNF536
MPPED2
CASC15
PITPNM3
RBM20
GAS6
AC007402.1
IQCA1
CES1
CDK10
ESR1
GAS8
KCNS3
SLC25A10
GCK
ANKRD11
MSRA
DEF8

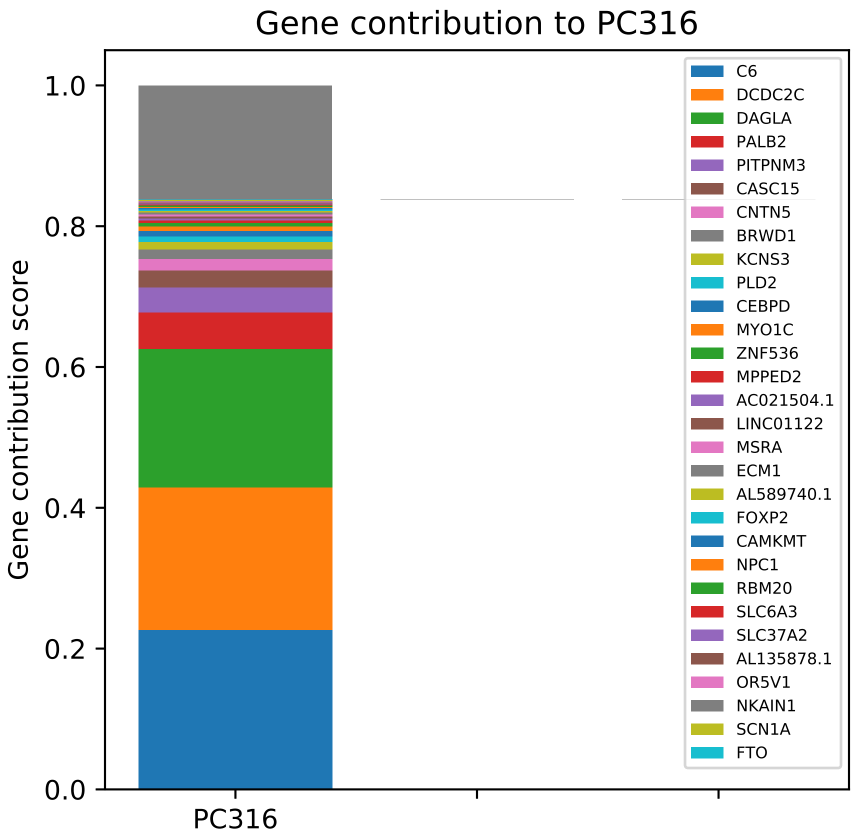
Gene contribution to PC316
Gene contribution score
0.0
0.2
0.4
0.6
0.8
1.0
PC316
C6
DCDC2C
DAGLA
PALB2
PITPNM3
CASC15
CNTN5
BRWD1
KCNS3
PLD2
CEBPD
MYO1C
ZNF536
MPPED2
AC021504.1
LINC01122
MSRA
ECM1
AL589740.1
FOXP2
CAMKMT
NPC1
RBM20
SLC6A3
SLC37A2
AL135878.1
OR5V1
NKAIN1
SCN1A
FTO

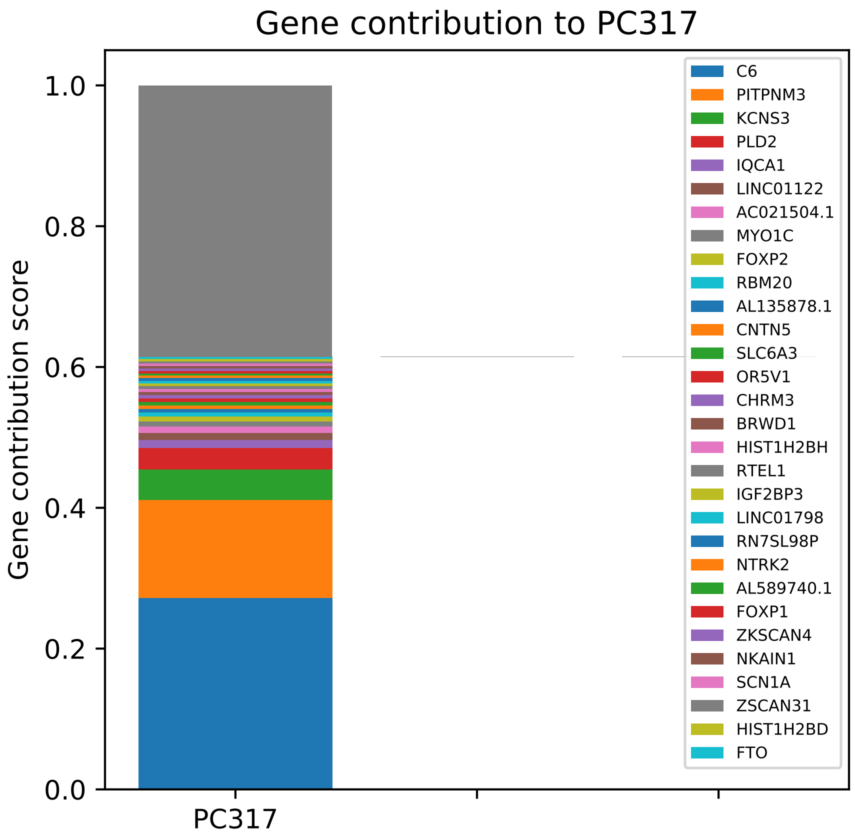

Gene contribution to PC317
Gene contribution score
1.0
0.8
0.6
0.4
0.2
0.0
PC317
C6
PITPNM3
KCNS3
PLD2
IQCA1
LINC01122
AC021504.1
MYO1C
FOXP2
RBM20
AL135878.1
CNTN5
SLC6A3
OR5V1
CHRM3
BRWD1
HIST1H2BH
RTEL1
IGF2BP3
LINC01798
RN7SL98P
NTRK2
AL589740.1
FOXP1
ZKSCAN4
NKAIN1
SCN1A
ZSCAN31
HIST1H2BD
FTO

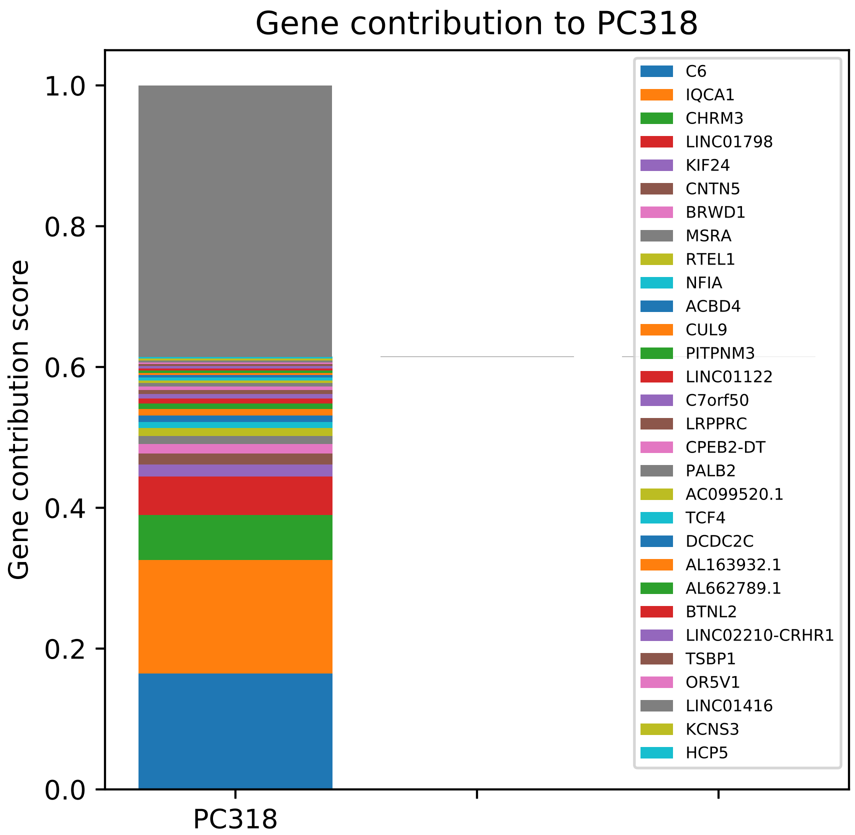

Gene contribution to PC318
Gene contribution score
1.0
0.8
0.6
0.4
0.2
0.0
PC318
C6
IQCA1
CHRM3
LINC01798
KIF24
CNTN5
BRWD1
MSRA
RTEL1
NFIA
ACBD4
CUL9
PITPNM3
LINC01122
C7orf50
LRPPRC
CPEB2-DT
PALB2
AC099520.1
TCF4
DCDC2C
AL163932.1
AL662789.1
BTNL2
LINC02210-CRHR1
TSBP1
OR5V1
LINC01416
KCNS3
HCP5

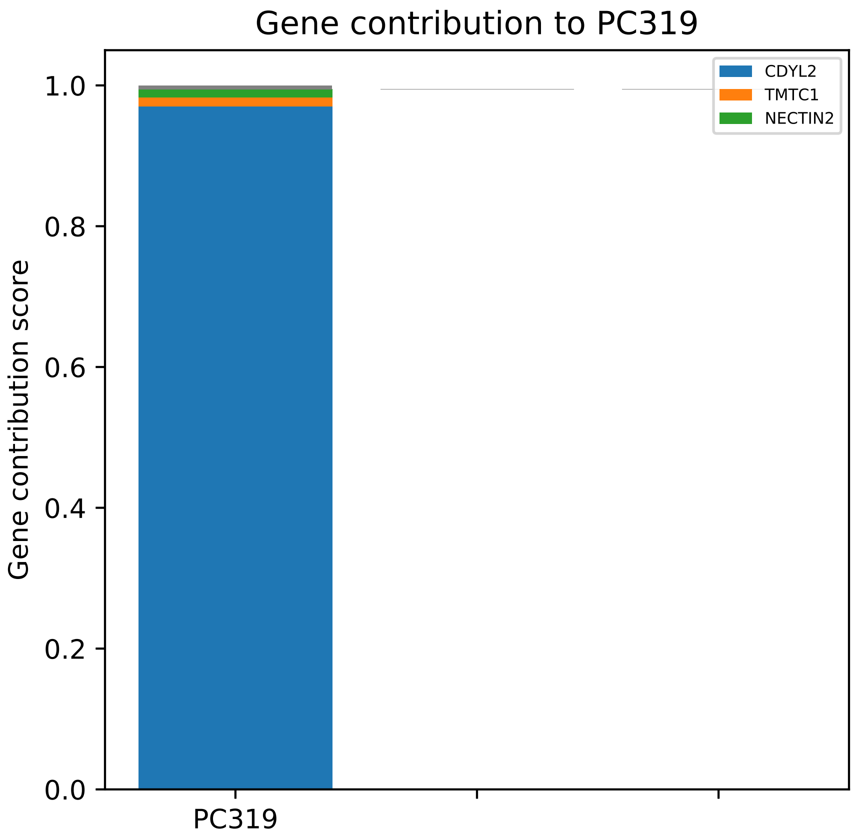
Gene contribution to PC319
Gene contribution score
1.0
0.8
0.6
0.4
0.2
0.0
PC319
CDYL2
TMTC1
NECTIN2

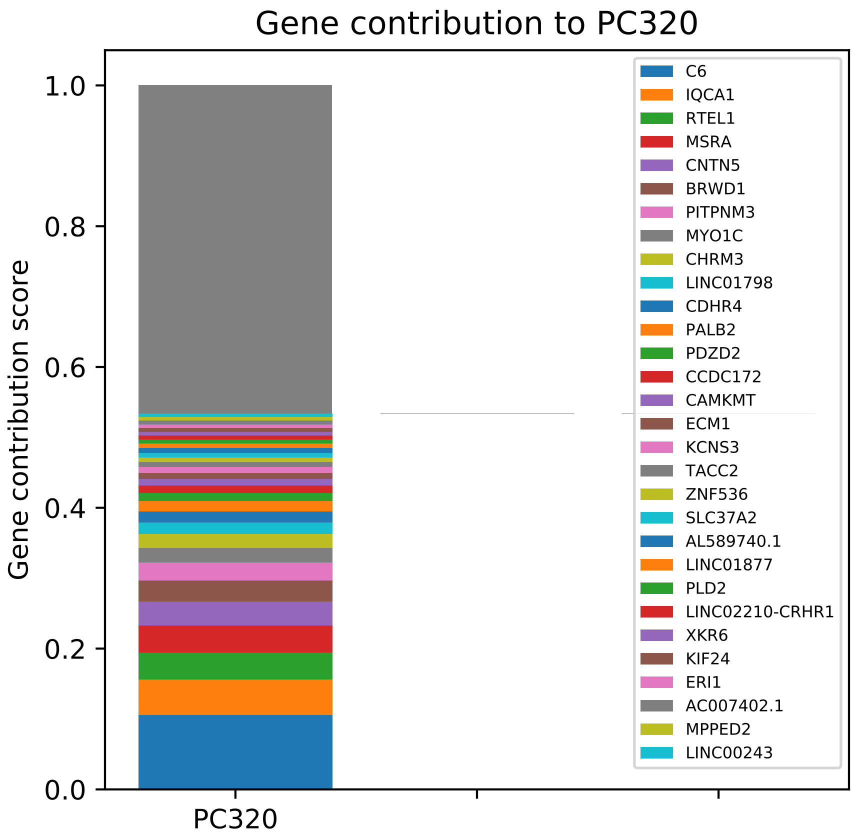
Gene contribution to PC320
Gene contribution score
1.0
0.8
0.6
0.4
0.2
0.0
PC320
C6
IQCA1
RTEL1
MSRA
CNTN5
BRWD1
PITPNM3
MYO1C
CHRM3
LINC01798
CDHR4
PALB2
PDZD2
CCDC172
CAMKMT
ECM1
KCNS3
TACC2
ZNF536
SLC37A2
AL589740.1
LINC01877
PLD2
LINC02210-CRHR1
XKR6
KIF24
ERI1
AC007402.1
MPPED2
LINC00243

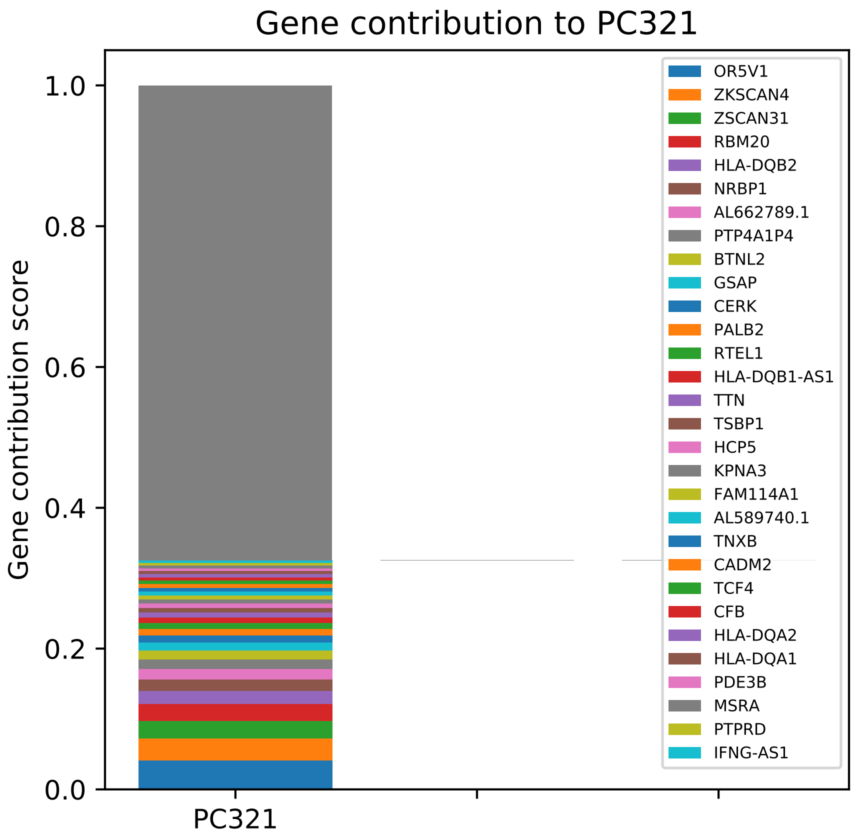
Gene contribution to PC321
Gene contribution score
0.0
0.2
0.4
0.6
0.8
1.0
PC321
OR5V1
ZKSCAN4
ZSCAN31
RBM20
HLA-DQB2
NRBP1
AL662789.1
PTP4A1P4
BTNL2
GSAP
CERK
PALB2
RTEL1
HLA-DQB1-AS1
TTN
TSBP1
HCP5
KPNA3
FAM114A1
AL589740.1
TNXB
CADM2
TCF4
CFB
HLA-DQA2
HLA-DQA1
PDE3B
MSRA
PTPRD
IFNG-AS1

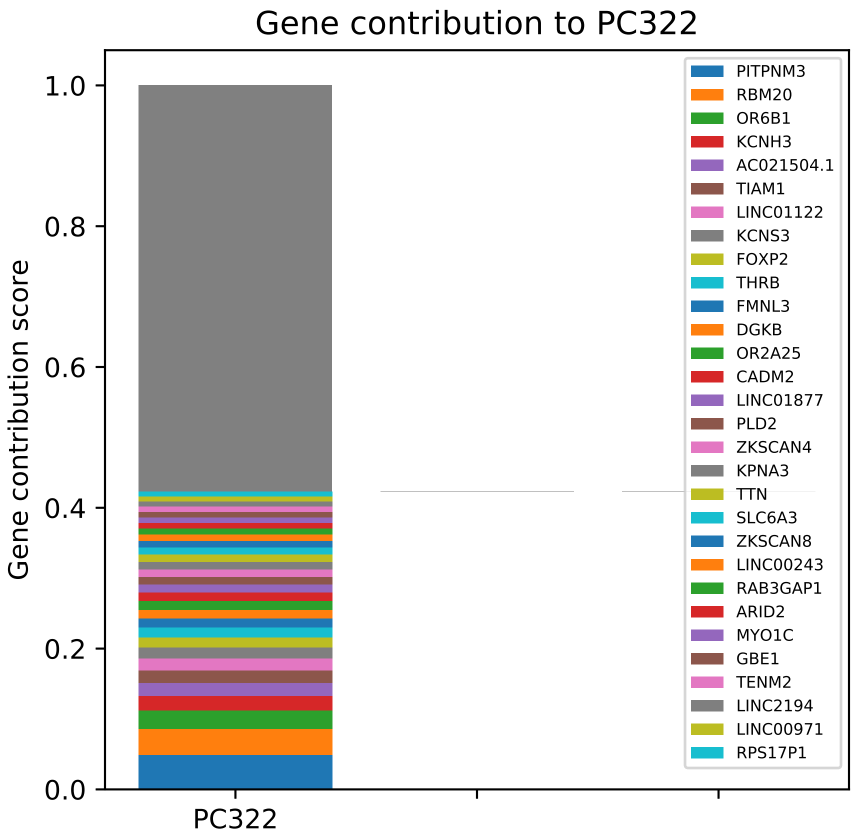

Gene contribution to PC322
Gene contribution score
1.0
0.8
0.6
0.4
0.2
0.0
PC322
PITPNM3
RBM20
OR6B1
KCNH3
AC021504.1
TIAM1
LINC01122
KCNS3
FOXP2
THRB
FMNL3
DGKB
OR2A25
CADM2
LINC01877
PLD2
ZKSCAN4
KPNA3
TTN
SLC6A3
ZKSCAN8
LINC00243
RAB3GAP1
ARID2
MYO1C
GBE1
TENM2
LINC2194
LINC00971
RPS17P1

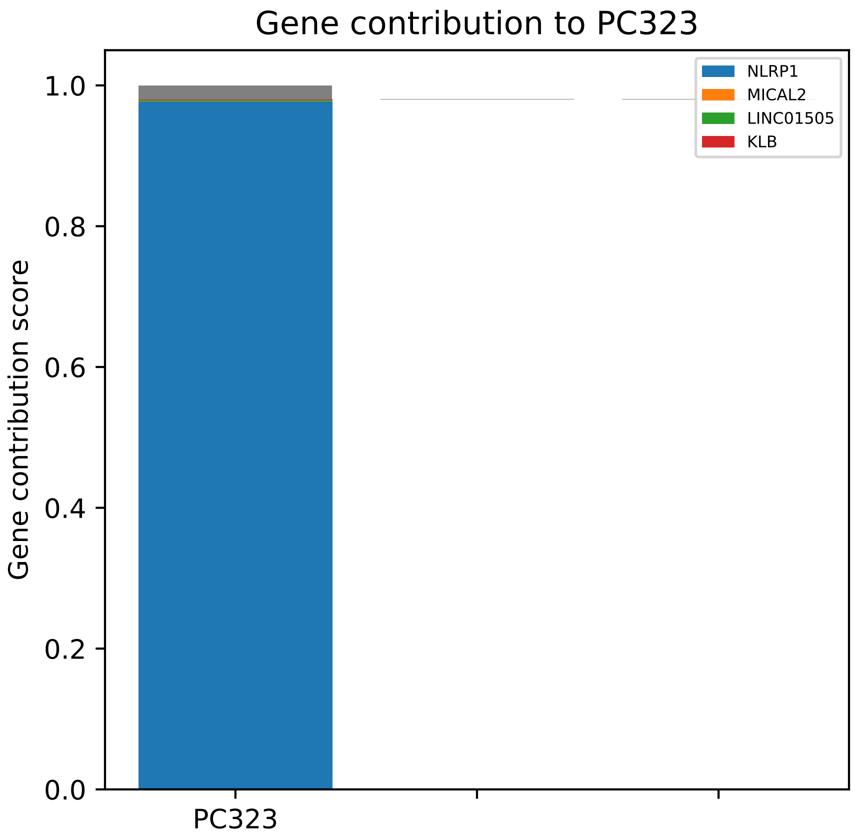

Gene contribution to PC323
Gene contribution score
1.0
0.8
0.6
0.4
0.2
0.0
PC323
NLRP1
MICAL2
LINC01505
KLB

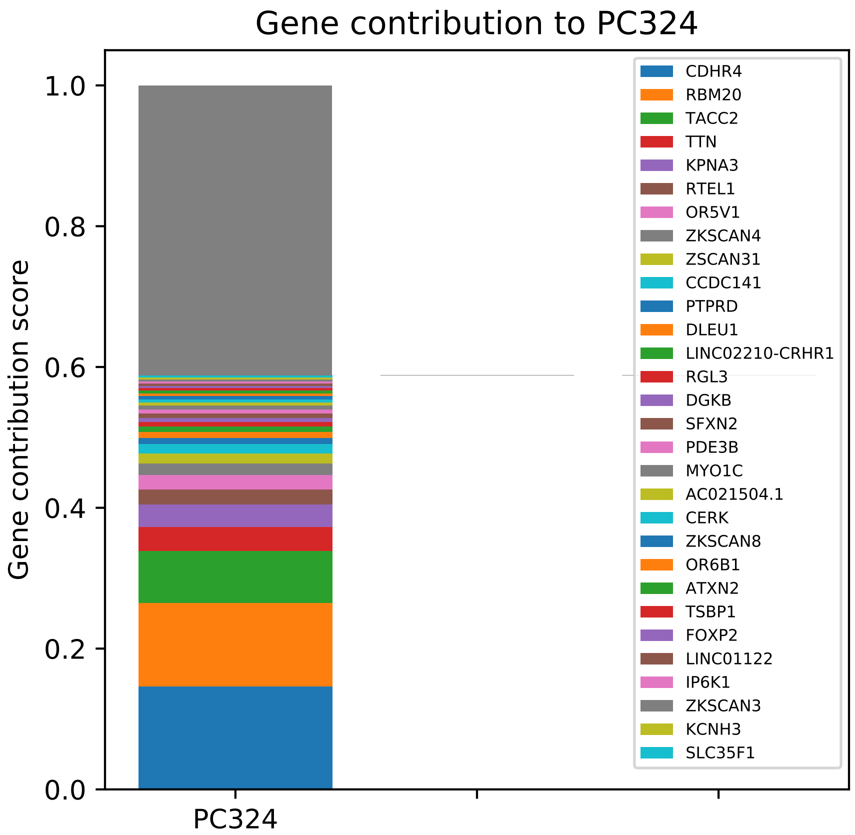

Gene contribution to PC324
Gene contribution score
1.0
0.8
0.6
0.4
0.2
0.0
PC324
CDHR4
RBM20
TACC2
TTN
KPNA3
RTEL1
OR5V1
ZKSCAN4
ZSCAN31
CCDC141
PTPRD
DLEU1
LINC02210-CRHR1
RGL3
DGKB
SFXN2
PDE3B
MYO1C
AC021504.1
CERK
ZKSCAN8
OR6B1
ATXN2
TSBP1
FOXP2
LINC01122
IP6K1
ZKSCAN3
KCNH3
SLC35F1

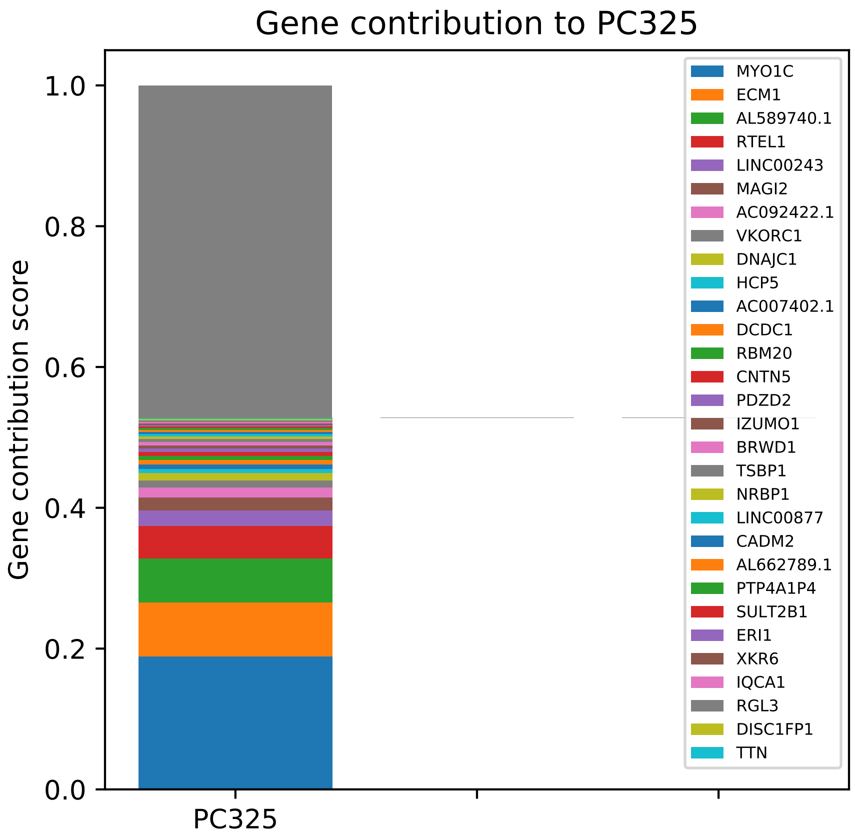

Gene contribution to PC325
Gene contribution score
0.0
0.2
0.4
0.6
0.8
1.0
PC325
MYO1C
ECM1
AL589740.1
RTEL1
LINC00243
MAGI2
AC092422.1
VKORC1
DNAJC1
HCP5
AC007402.1
DCDC1
RBM20
CNTN5
PDZD2
IZUMO1
BRWD1
TSBP1
NRBP1
LINC00877
CADM2
AL662789.1
PTP4A1P4
SULT2B1
ERI1
XKR6
IQCA1
RGL3
DISC1FP1
TTN

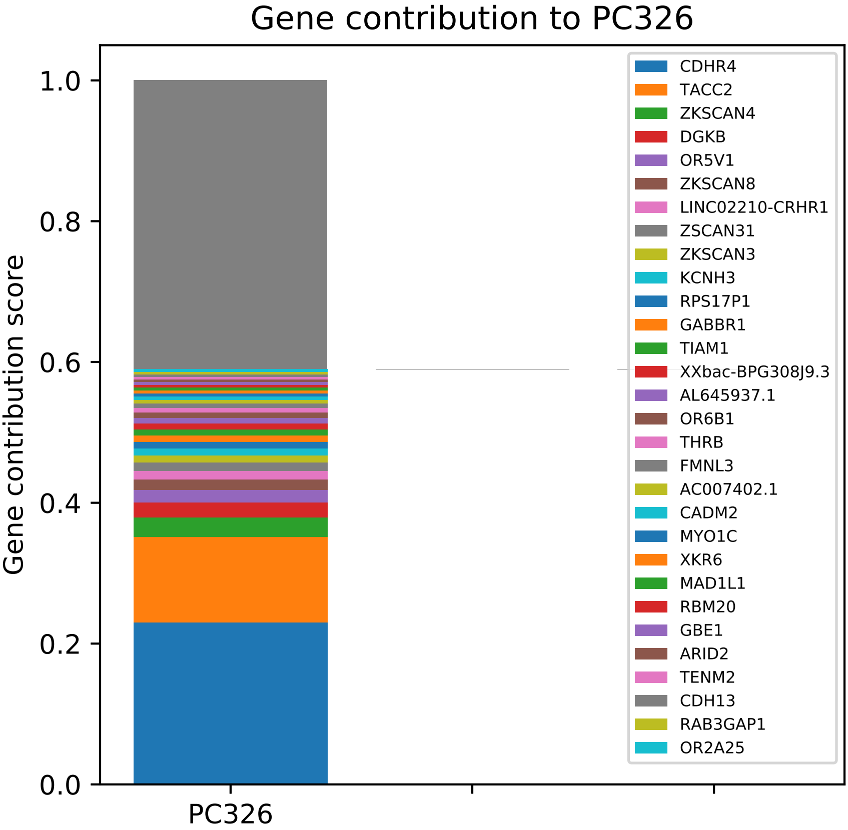
Gene contribution to PC326
Gene contribution score
1.0
0.8
0.6
0.4
0.2
0.0
PC326
CDHR4
TACC2
ZKSCAN4
DGKB
OR5V1
ZKSCAN8
LINC02210-CRHR1
ZSCAN31
ZKSCAN3
KCNH3
RPS17P1
GABBR1
TIAM1
XXbac-BPG308J9.3
AL645937.1
OR6B1
THRB
FMNL3
AC007402.1
CADM2
MYO1C
XKR6
MAD1L1
RBM20
GBE1
ARID2
TENM2
CDH13
RAB3GAP1
OR2A25

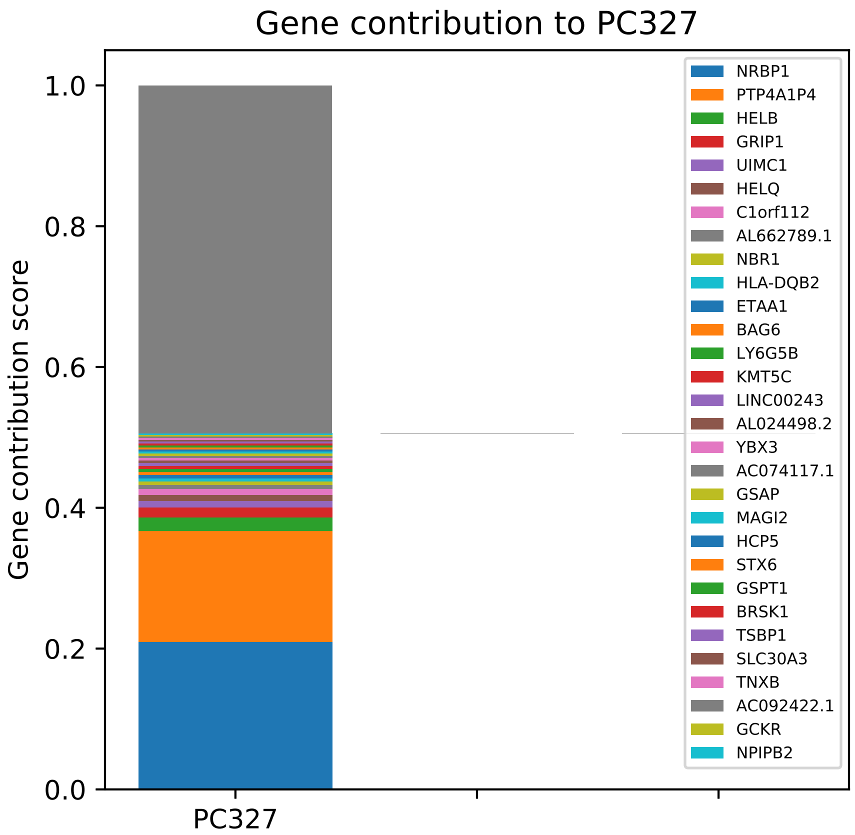

Gene contribution to PC327
Gene contribution score
1.0
0.8
0.6
0.4
0.2
0.0
PC327
NRBP1
PTP4A1P4
HELB
GRIP1
UIMC1
HELQ
C1orf112
AL662789.1
NBR1
HLA-DQB2
ETAA1
BAG6
LY6G5B
KMT5C
LINC00243
AL024498.2
YBX3
AC074117.1
GSAP
MAGI2
HCP5
STX6
GSPT1
BRSK1
TSBP1
SLC30A3
TNXB
AC092422.1
GCKR
NPIPB2

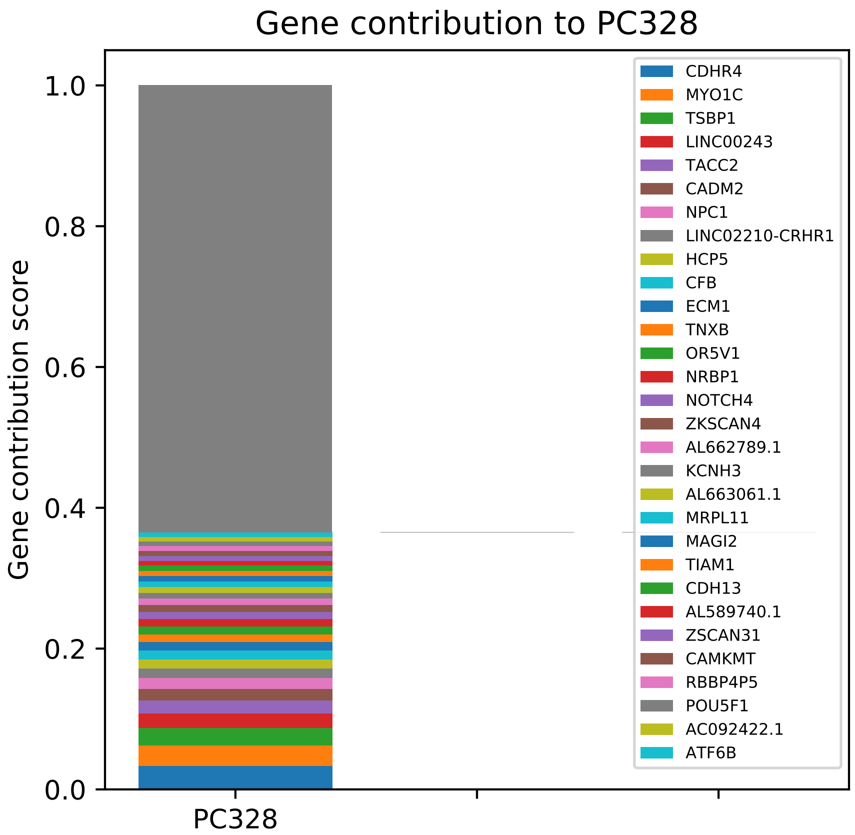
Gene contribution to PC328
Gene contribution score
1.0
0.8
0.6
0.4
0.2
0.0
PC328
CDHR4
MYO1C
TSBP1
LINC00243
TACC2
CADM2
NPC1
LINC02210-CRHR1
HCP5
CFB
ECM1
TNXB
OR5V1
NRBP1
NOTCH4
ZKSCAN4
AL662789.1
KCNH3
AL663061.1
MRPL11
MAGI2
TIAM1
CDH13
AL589740.1
ZSCAN31
CAMKMT
RBBP4P5
POU5F1
AC092422.1
ATF6B

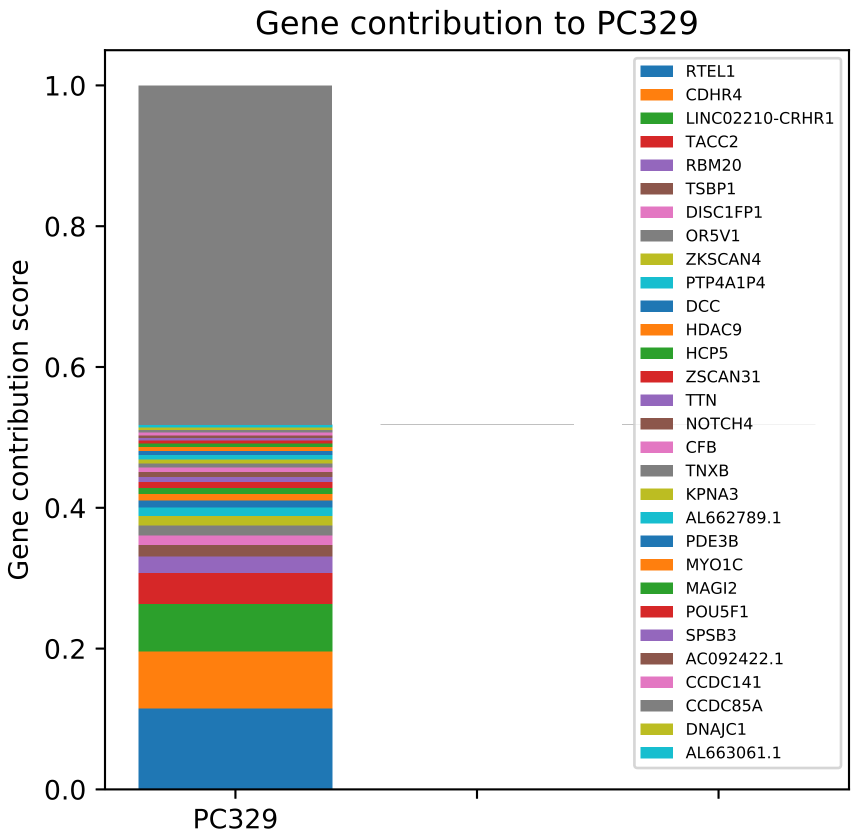
Gene contribution to PC329
Gene contribution score
0.0
0.2
0.4
0.6
0.8
1.0
PC329
RTEL1
CDHR4
LINC02210-CRHR1
TACC2
RBM20
TSBP1
DISC1FP1
OR5V1
ZKSCAN4
PTP4A1P4
DCC
HDAC9
HCP5
ZSCAN31
TTN
NOTCH4
CFB
TNXB
KPNA3
AL662789.1
PDE3B
MYO1C
MAGI2
POU5F1
SPSB3
AC092422.1
CCDC141
CCDC85A
DNAJC1
AL663061.1

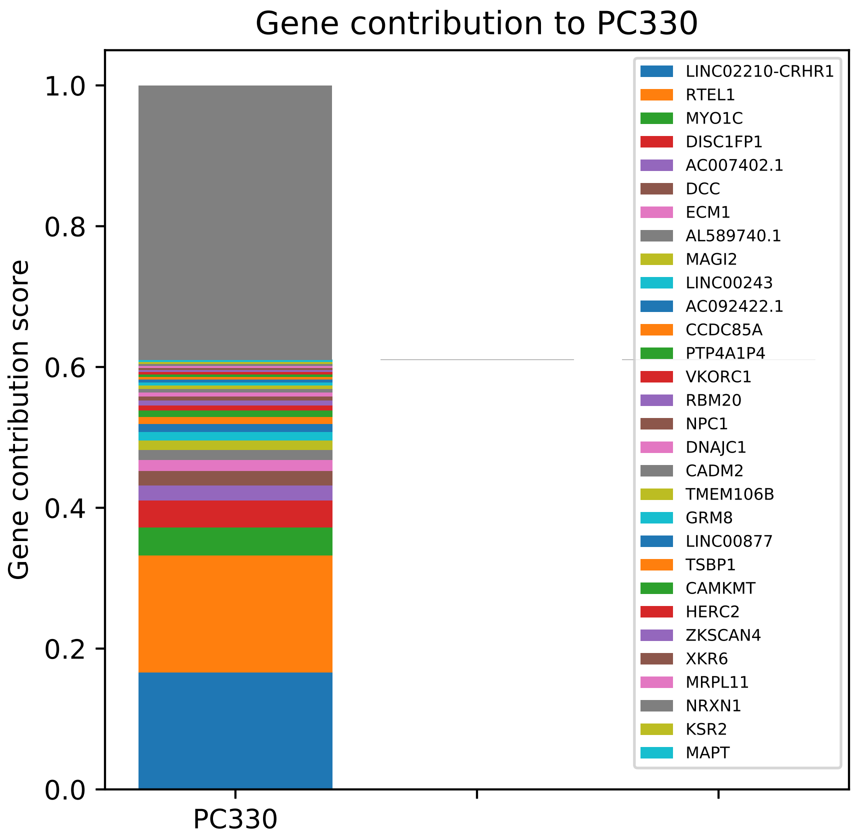
Gene contribution to PC330
Gene contribution score
1.0
0.8
0.6
0.4
0.2
0.0
PC330
LINC02210-CRHR1
RTEL1
MYO1C
DISC1FP1
AC007402.1
DCC
ECM1
AL589740.1
MAGI2
LINC00243
AC092422.1
CCDC85A
PTP4A1P4
VKORC1
RBM20
NPC1
DNAJC1
CADM2
TMEM106B
GRM8
LINC00877
TSBP1
CAMKMT
HERC2
ZKSCAN4
XKR6
MRPL11
NRXN1
KSR2
MAPT

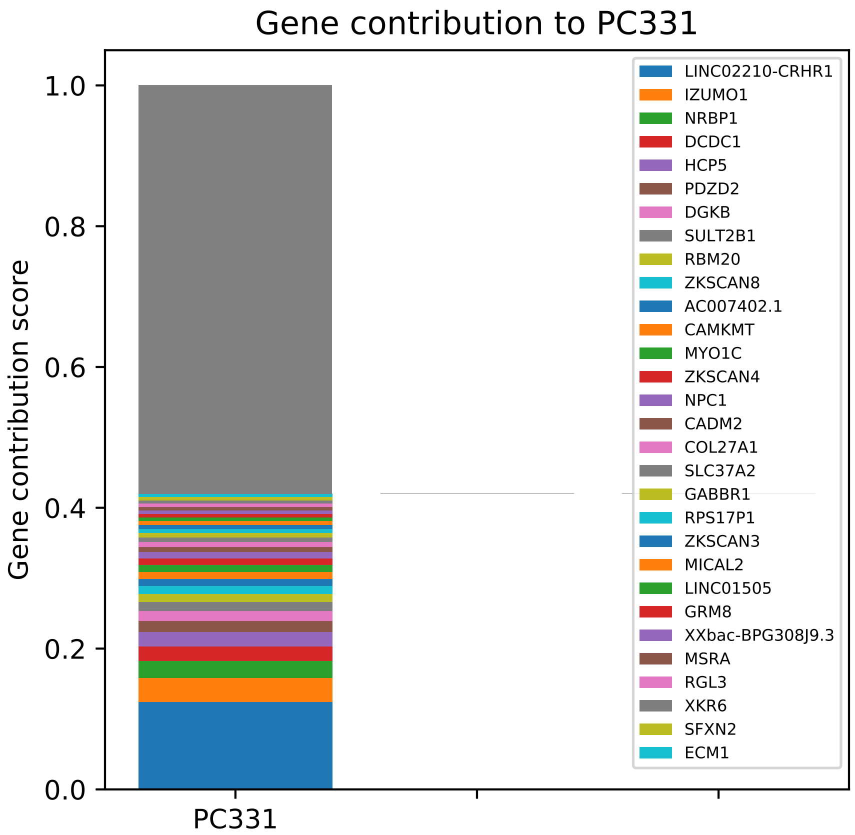

Gene contribution to PC331
Gene contribution score
1.0
0.8
0.6
0.4
0.2
0.0
PC331
LINC02210-CRHR1
IZUMO1
NRBP1
DCDC1
HCP5
PDZD2
DGKB
SULT2B1
RBM20
ZKSCAN8
AC007402.1
CAMKMT
MYO1C
ZKSCAN4
NPC1
CADM2
COL27A1
SLC37A2
GABBR1
RPS17P1
ZKSCAN3
MICAL2
LINC01505
GRM8
XXbac-BPG308J9.3
MSRA
RGL3
XKR6
SFXN2
ECM1

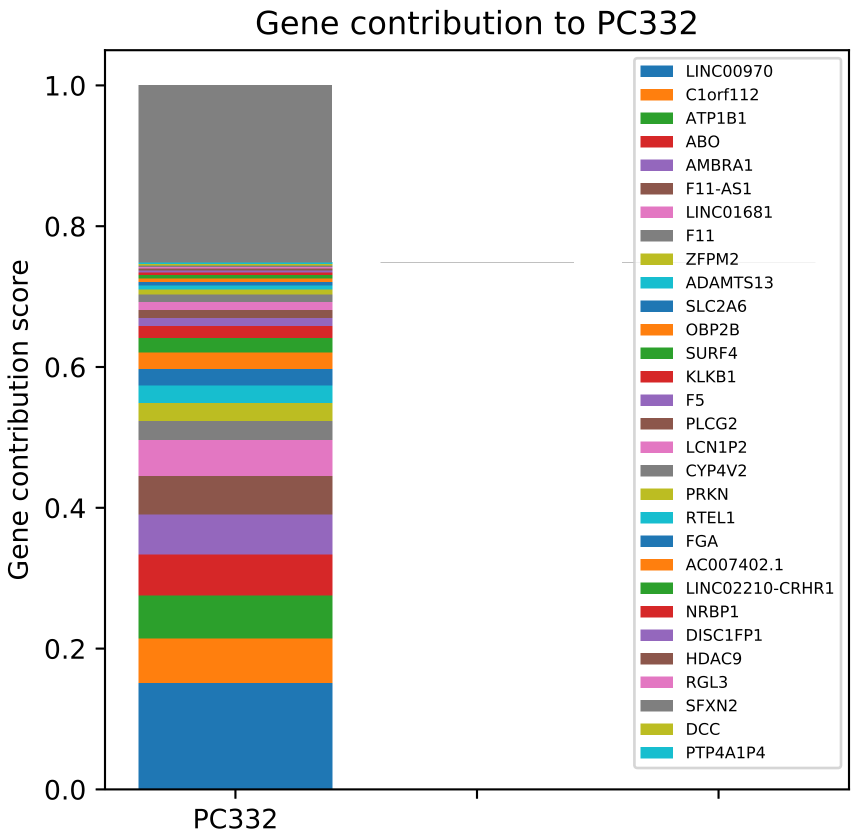
Gene contribution to PC332
Gene contribution score
1.0
0.8
0.6
0.4
0.2
0.0
PC332
LINC00970
C1orf112
ATP1B1
ABO
AMBRA1
F11-AS1
LINC01681
F11
ZFPM2
ADAMTS13
SLC2A6
OBP2B
SURF4
KLKB1
F5
PLCG2
LCN1P2
CYP4V2
PRKN
RTEL1
FGA
AC007402.1
LINC02210-CRHR1
NRBP1
DISC1FP1
HDAC9
RGL3
SFXN2
DCC
PTP4A1P4

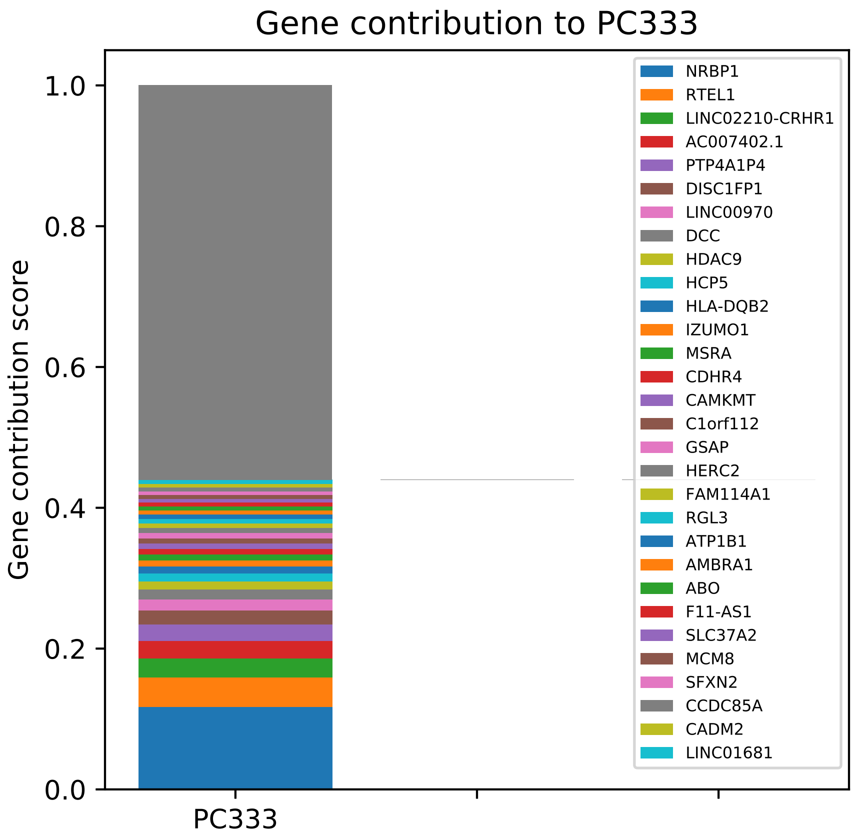
Gene contribution to PC333
Gene contribution score
1.0
0.8
0.6
0.4
0.2
0.0
PC333
NRBP1
RTEL1
LINC02210-CRHR1
AC007402.1
PTP4A1P4
DISC1FP1
LINC00970
DCC
HDAC9
HCP5
HLA-DQB2
IZUMO1
MSRA
CDHR4
CAMKMT
C1orf112
GSAP
HERC2
FAM114A1
RGL3
ATP1B1
AMBRA1
ABO
F11-AS1
SLC37A2
MCM8
SFXN2
CCDC85A
CADM2
LINC01681

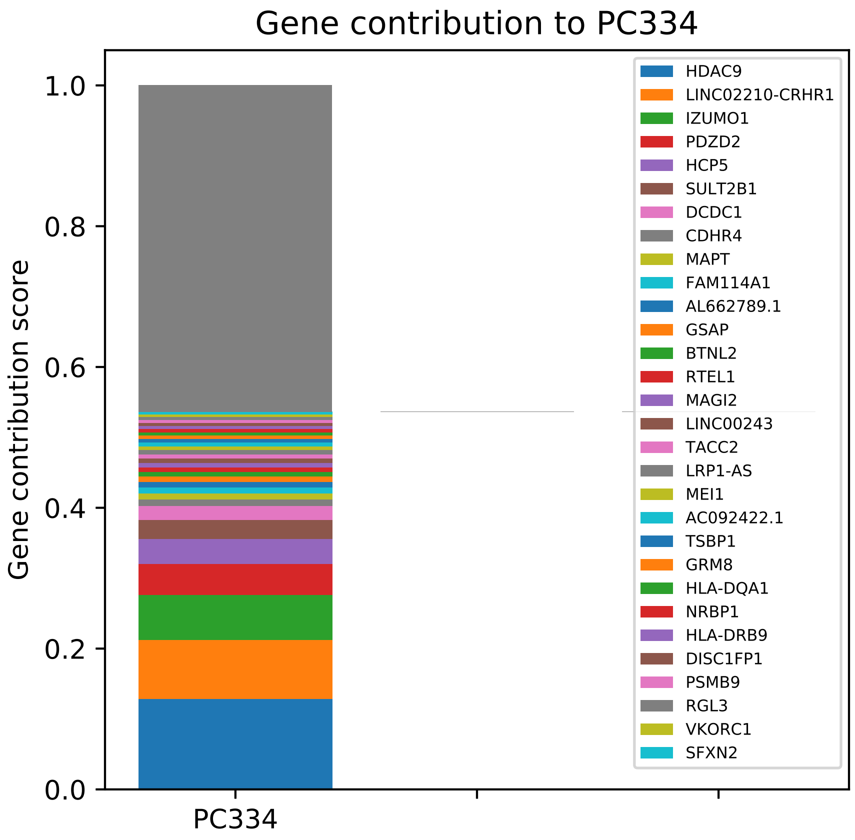

Gene contribution to PC334
Gene contribution score
1.0
0.8
0.6
0.4
0.2
0.0
PC334
HDAC9
LINC02210-CRHR1
IZUMO1
PDZD2
HCP5
SULT2B1
DCDC1
CDHR4
MAPT
FAM114A1
AL662789.1
GSAP
BTNL2
RTEL1
MAGI2
LINC00243
TACC2
LRP1-AS
MEI1
AC092422.1
TSBP1
GRM8
HLA-DQA1
NRBP1
HLA-DRB9
DISC1FP1
PSMB9
RGL3
VKORC1
SFXN2

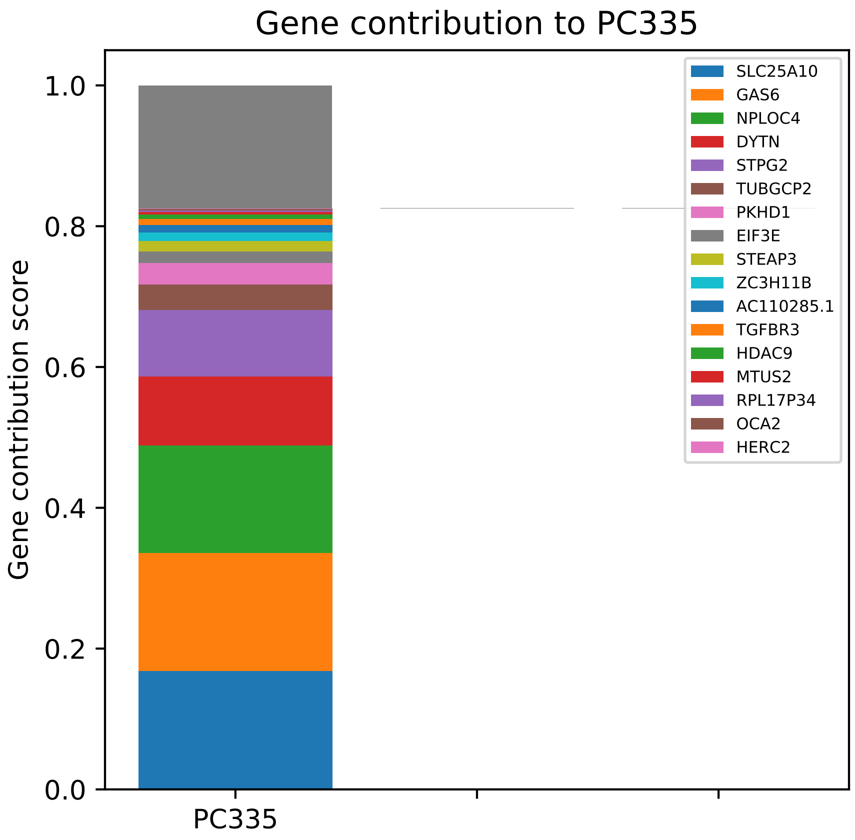

Gene contribution to PC335
Gene contribution score
1.0
0.8
0.6
0.4
0.2
0.0
PC335
SLC25A10
GAS6
NPLOC4
DYTN
STPG2
TUBGCP2
PKHD1
EIF3E
STEAP3
ZC3H11B
AC110285.1
TGFBR3
HDAC9
MTUS2
RPL17P34
OCA2
HERC2

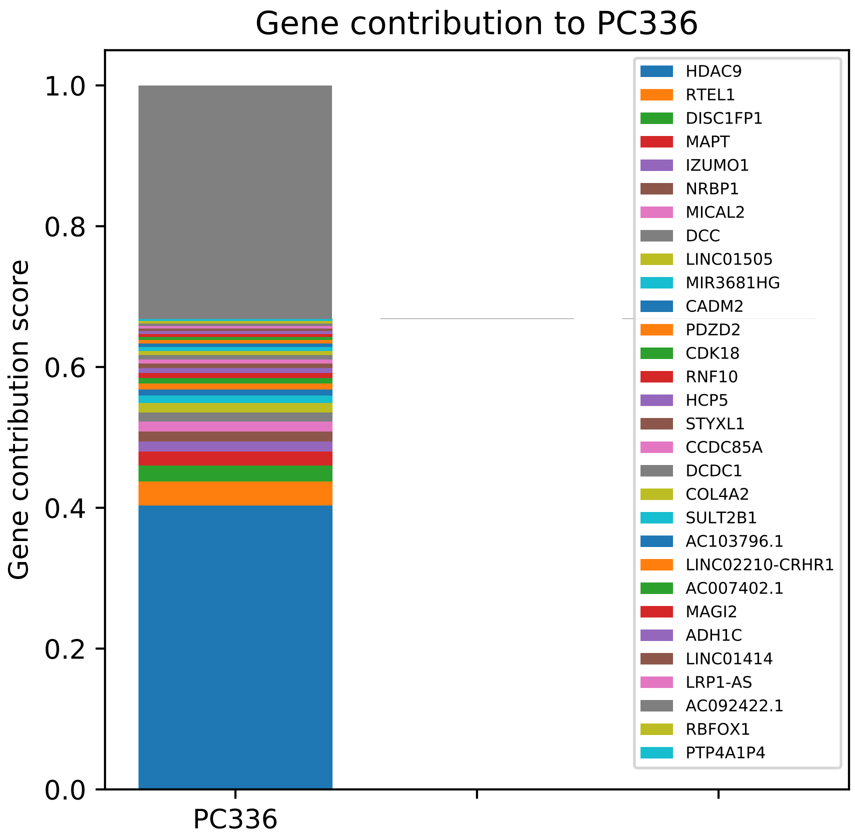

Gene contribution to PC336
Gene contribution score
1.0
0.8
0.6
0.4
0.2
0.0
PC336
HDAC9
RTEL1
DISC1FP1
MAPT
IZUMO1
NRBP1
MICAL2
DCC
LINC01505
MIR3681HG
CADM2
PDZD2
CDK18
RNF10
HCP5
STYXL1
CCDC85A
DCDC1
COL4A2
SULT2B1
AC103796.1
LINC02210-CRHR1
AC007402.1
MAGI2
ADH1C
LINC01414
LRP1-AS
AC092422.1
RBFOX1
PTP4A1P4

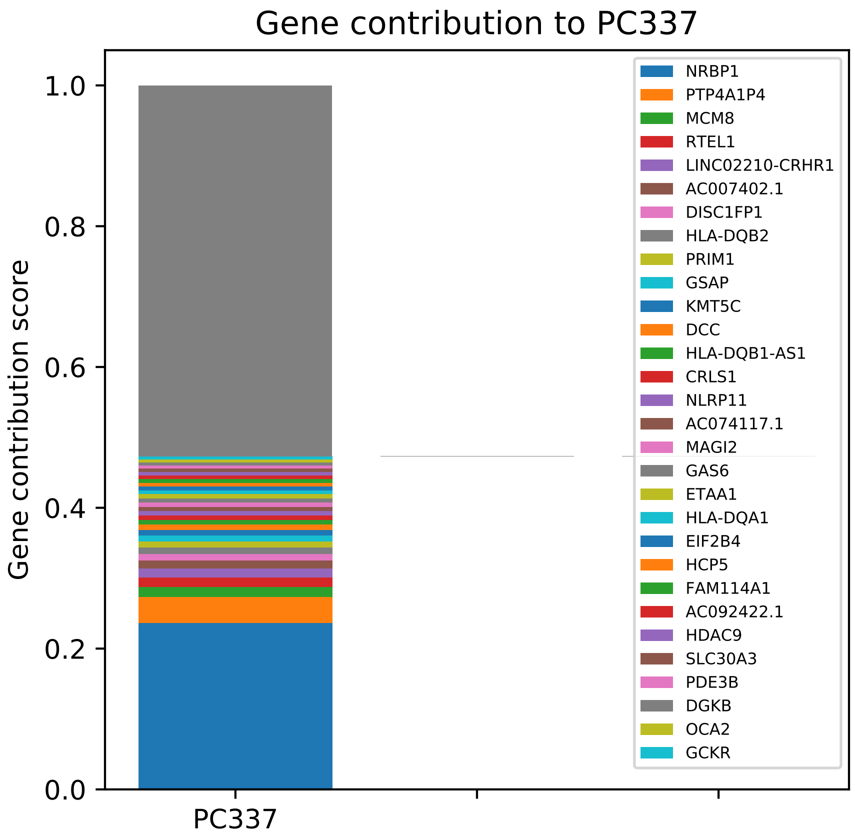

Gene contribution to PC337
Gene contribution score
1.0
0.8
0.6
0.4
0.2
0.0
PC337
NRBP1
PTP4A1P4
MCM8
RTEL1
LINC02210-CRHR1
AC007402.1
DISC1FP1
HLA-DQB2
PRIM1
GSAP
KMT5C
DCC
HLA-DQB1-AS1
CRLS1
NLRP11
AC074117.1
MAGI2
GAS6
ETAA1
HLA-DQA1
EIF2B4
HCP5
FAM114A1
AC092422.1
HDAC9
SLC30A3
PDE3B
DGKB
OCA2
GCKR

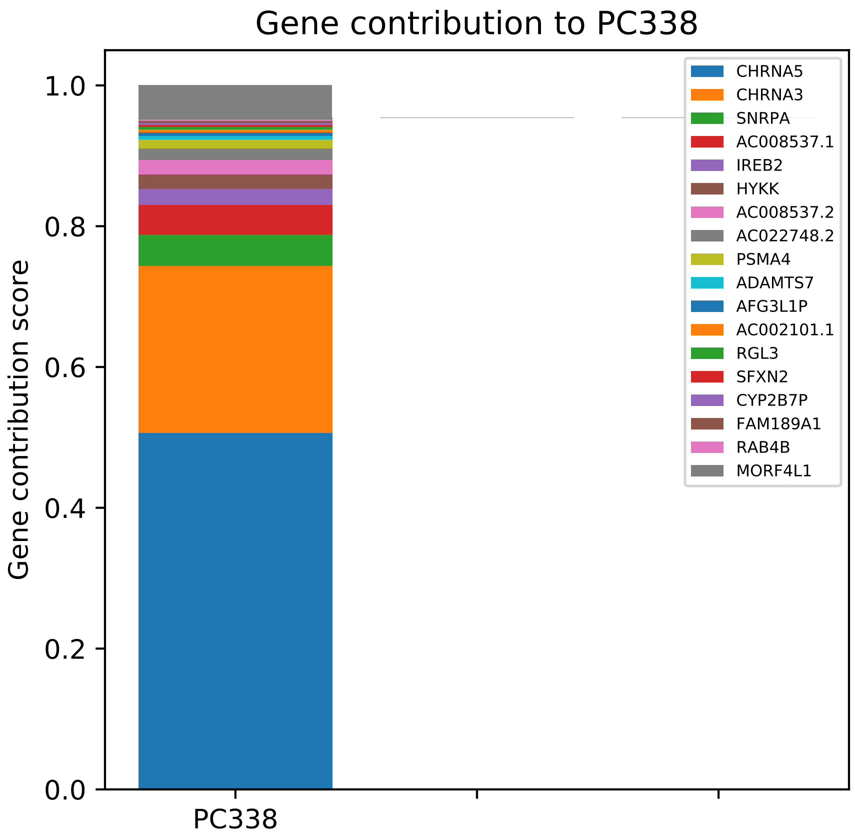

Gene contribution to PC338
Gene contribution score
1.0
0.8
0.6
0.4
0.2
0.0
PC338
CHRNA5
CHRNA3
SNRPA
AC008537.1
IREB2
HYKK
AC008537.2
AC022748.2
PSMA4
ADAMTS7
AFG3L1P
AC002101.1
RGL3
SFXN2
CYP2B7P
FAM189A1
RAB4B
MORF4L1

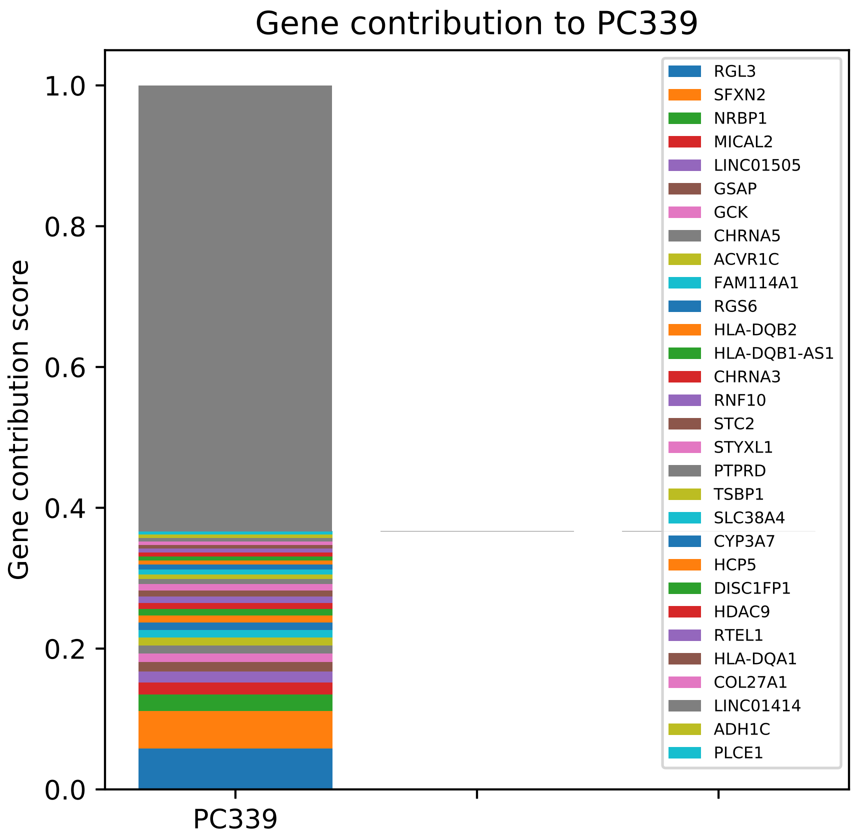

Gene contribution to PC339
Gene contribution score
1.0
0.8
0.6
0.4
0.2
0.0
PC339
RGL3
SFXN2
NRBP1
MICAL2
LINC01505
GSAP
GCK
CHRNA5
ACVR1C
FAM114A1
RGS6
HLA-DQB2
HLA-DQB1-AS1
CHRNA3
RNF10
STC2
STYXL1
PTPRD
TSBP1
SLC38A4
CYP3A7
HCP5
DISC1FP1
HDAC9
RTEL1
HLA-DQA1
COL27A1
LINC01414
ADH1C
PLCE1

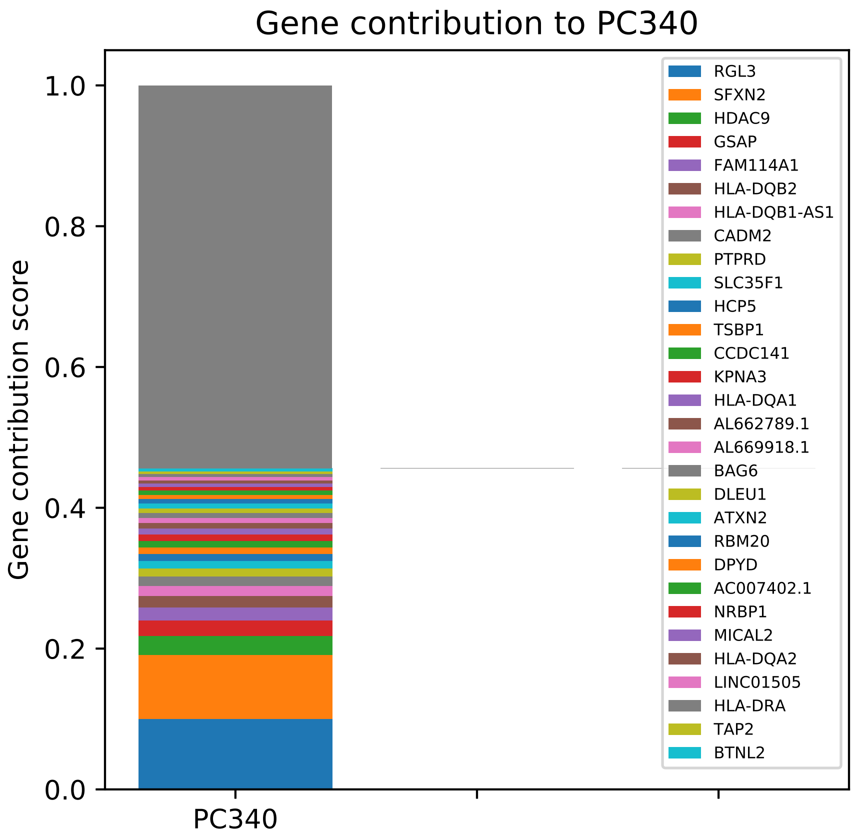

Gene contribution to PC340
Gene contribution score
1.0
0.8
0.6
0.4
0.2
0.0
PC340
RGL3
SFXN2
HDAC9
GSAP
FAM114A1
HLA-DQB2
HLA-DQB1-AS1
CADM2
PTPRD
SLC35F1
HCP5
TSBP1
CCDC141
KPNA3
HLA-DQA1
AL662789.1
AL669918.1
BAG6
DLEU1
ATXN2
RBM20
DPYD
AC007402.1
NRBP1
MICAL2
HLA-DQA2
LINC01505
HLA-DRA
TAP2
BTNL2

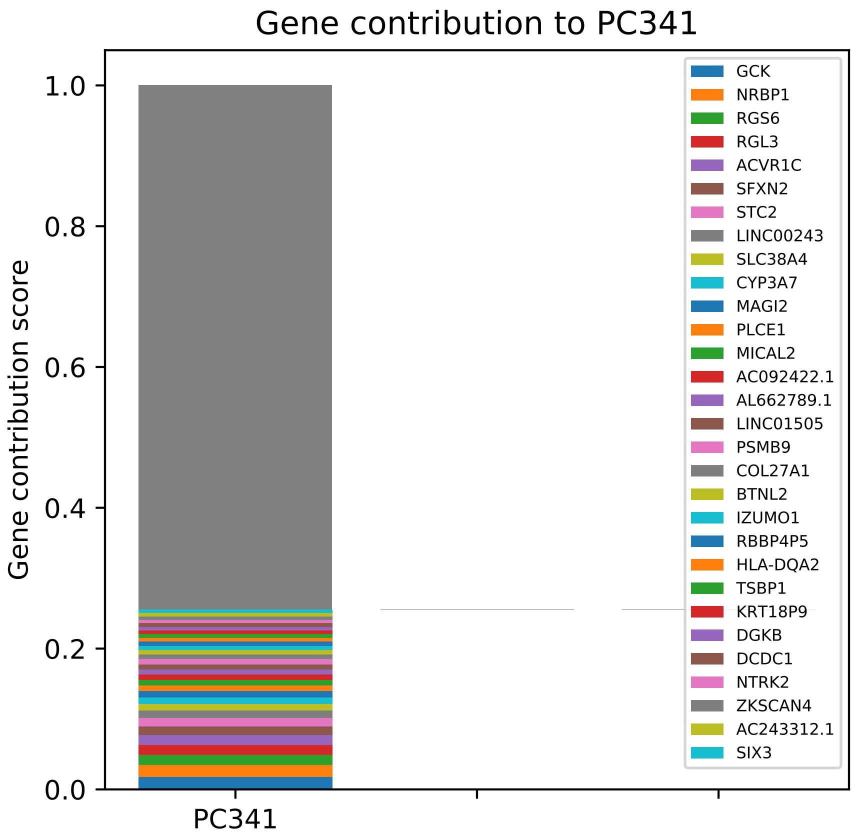

Gene contribution to PC341
Gene contribution score
1.0
0.8
0.6
0.4
0.2
0.0
PC341
GCK
NRBP1
RGS6
RGL3
ACVR1C
SFXN2
STC2
LINC00243
SLC38A4
CYP3A7
MAGI2
PLCE1
MICAL2
AC092422.1
AL662789.1
LINC01505
PSMB9
COL27A1
BTNL2
IZUMO1
RBBP4P5
HLA-DQA2
TSBP1
KRT18P9
DGKB
DCDC1
NTRK2
ZKSCAN4
AC243312.1
SIX3

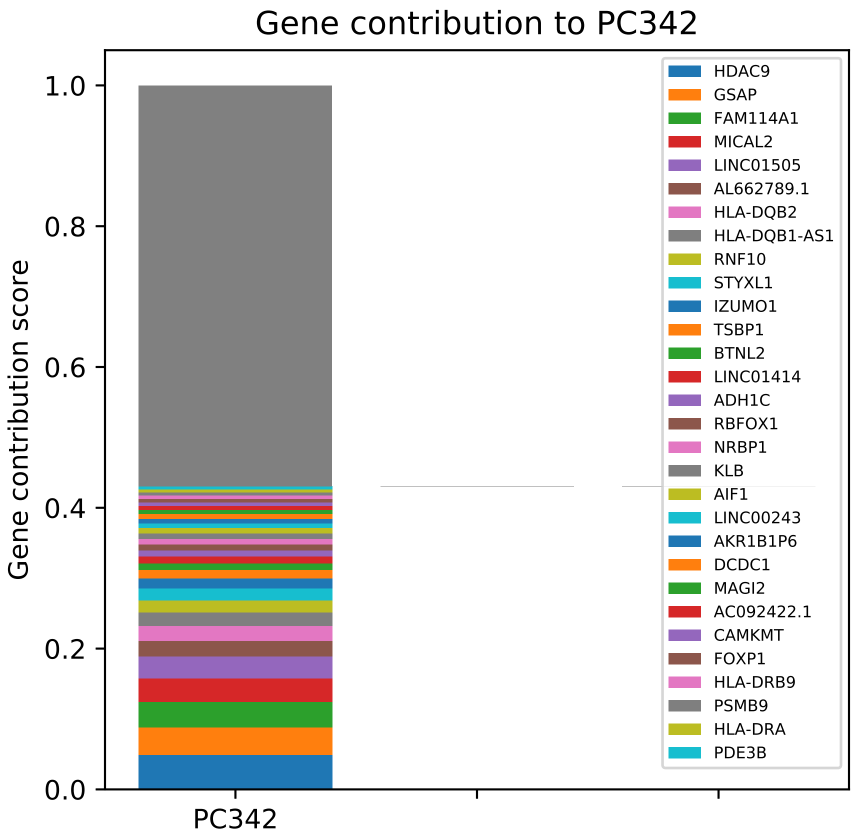
Gene contribution to PC342
Gene contribution score
1.0
0.8
0.6
0.4
0.2
0.0
PC342
HDAC9
GSAP
FAM114A1
MICAL2
LINC01505
AL662789.1
HLA-DQB2
HLA-DQB1-AS1
RNF10
STYXL1
IZUMO1
TSBP1
BTNL2
LINC01414
ADH1C
RBFOX1
NRBP1
KLB
AIF1
LINC00243
AKR1B1P6
DCDC1
MAGI2
AC092422.1
CAMKMT
FOXP1
HLA-DRB9
PSMB9
HLA-DRA
PDE3B

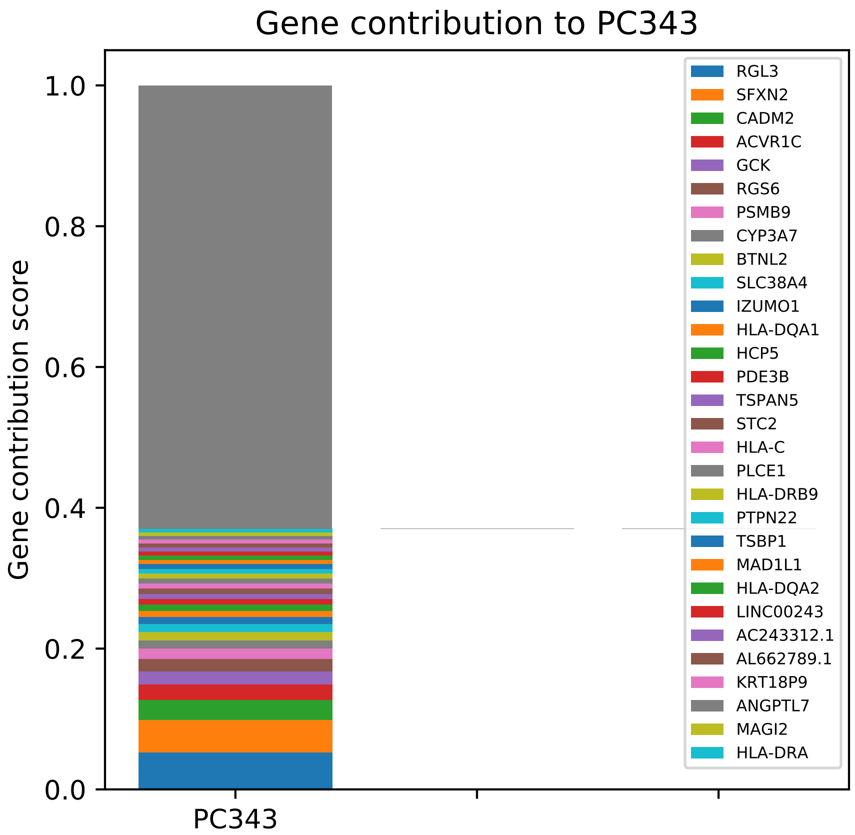

Gene contribution to PC343
Gene contribution score
0.0
0.2
0.4
0.6
0.8
1.0
PC343
RGL3
SFXN2
CADM2
ACVR1C
GCK
RGS6
PSMB9
CYP3A7
BTNL2
SLC38A4
IZUMO1
HLA-DQA1
HCP5
PDE3B
TSPAN5
STC2
HLA-C
PLCE1
HLA-DRB9
PTPN22
TSBP1
MAD1L1
HLA-DQA2
LINC00243
AC243312.1
AL662789.1
KRT18P9
ANGPTL7
MAGI2
HLA-DRA

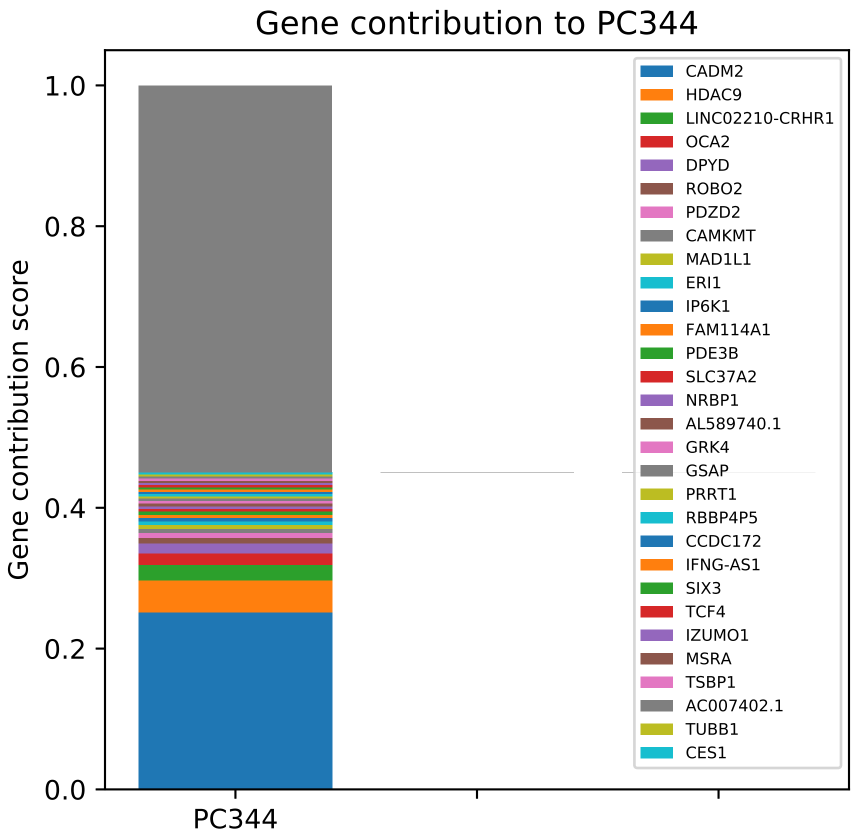

Gene contribution to PC344
Gene contribution score
1.0
0.8
0.6
0.4
0.2
0.0
PC344
CADM2
HDAC9
LINC02210-CRHR1
OCA2
DPYD
ROBO2
PDZD2
CAMKMT
MAD1L1
ERI1
IP6K1
FAM114A1
PDE3B
SLC37A2
NRBP1
AL589740.1
GRK4
GSAP
PRRT1
RBBP4P5
CCDC172
IFNG-AS1
SIX3
TCF4
IZUMO1
MSRA
TSBP1
AC007402.1
TUBB1
CES1

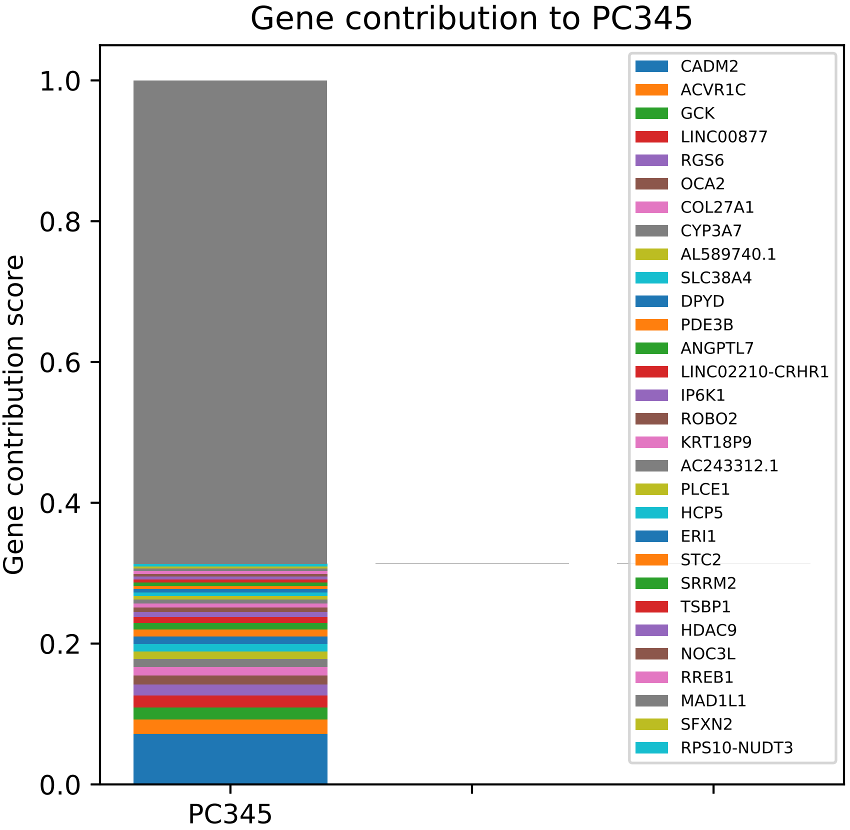

Gene contribution to PC345
Gene contribution score
1.0
0.8
0.6
0.4
0.2
0.0
PC345
CADM2
ACVR1C
GCK
LINC00877
RGS6
OCA2
COL27A1
CYP3A7
AL589740.1
SLC38A4
DPYD
PDE3B
ANGPTL7
LINC02210-CRHR1
IP6K1
ROBO2
KRT18P9
AC243312.1
PLCE1
HCP5
ERI1
STC2
SRRM2
TSBP1
HDAC9
NOC3L
RREB1
MAD1L1
SFXN2
RPS10-NUDT3

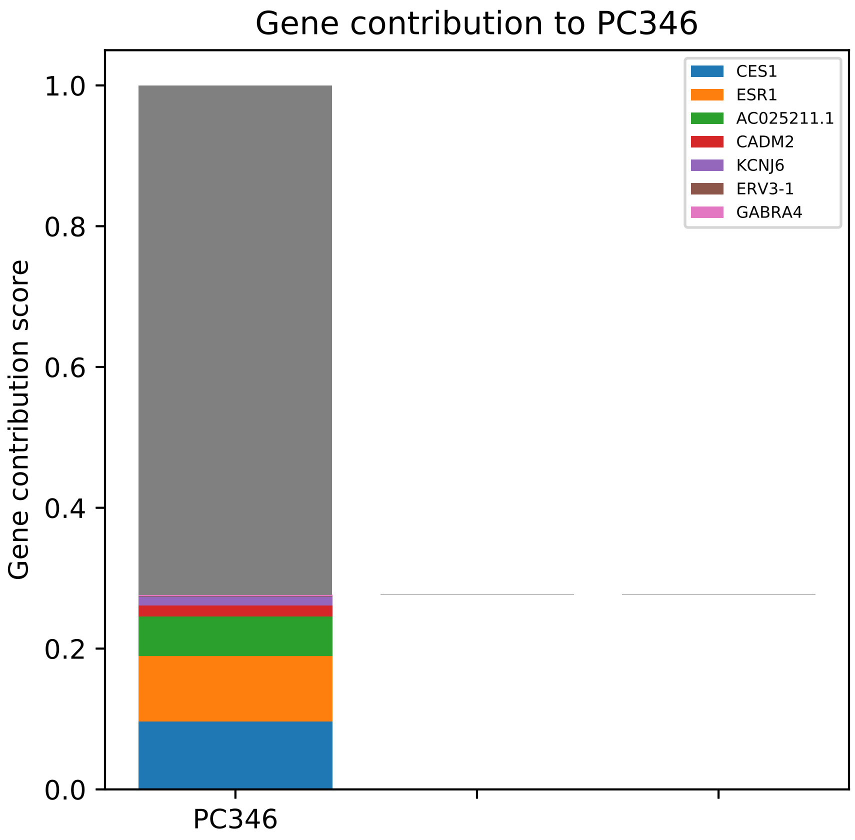

Gene contribution to PC346
Gene contribution score
1.0
0.8
0.6
0.4
0.2
0.0
PC346
CES1
ESR1
AC025211.1
CADM2
KCNJ6
ERV3-1
GABRA4

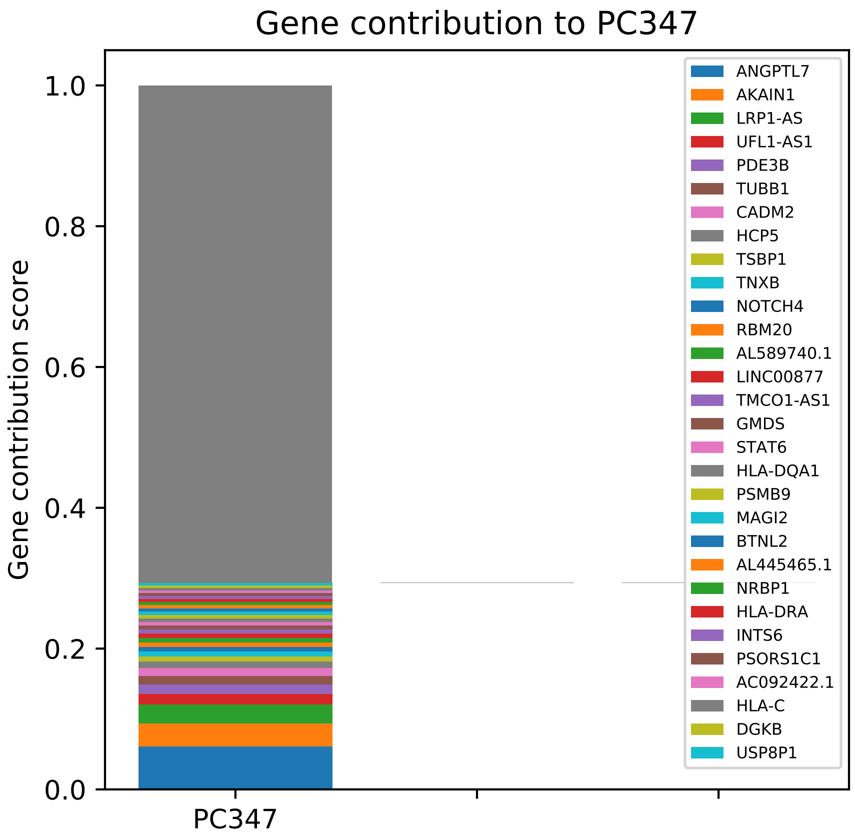

Gene contribution to PC347
Gene contribution score
1.0
0.8
0.6
0.4
0.2
0.0
PC347
ANGPTL7
AKAIN1
LRP1-AS
UFL1-AS1
PDE3B
TUBB1
CADM2
HCP5
TSBP1
TNXB
NOTCH4
RBM20
AL589740.1
LINC00877
TMCO1-AS1
GMDS
STAT6
HLA-DQA1
PSMB9
MAGI2
BTNL2
AL445465.1
NRBP1
HLA-DRA
INTS6
PSORS1C1
AC092422.1
HLA-C
DGKB
USP8P1

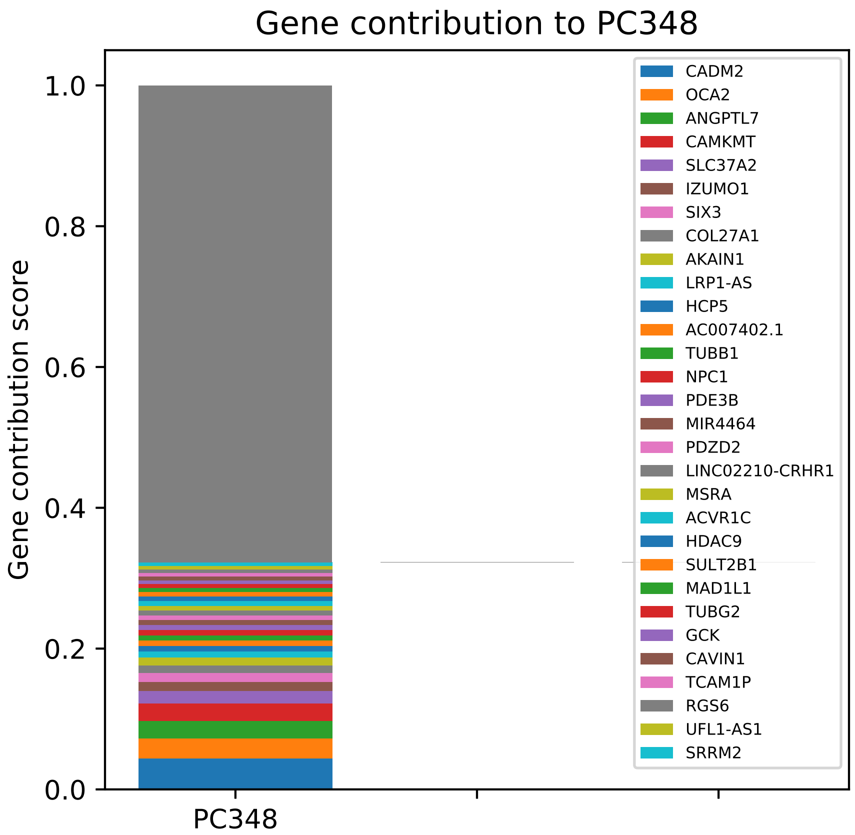

Gene contribution to PC348
Gene contribution score
1.0
0.8
0.6
0.4
0.2
0.0
PC348
CADM2
OCA2
ANGPTL7
CAMKMT
SLC37A2
IZUMO1
SIX3
COL27A1
AKAIN1
LRP1-AS
HCP5
AC007402.1
TUBB1
NPC1
PDE3B
MIR4464
PDZD2
LINC02210-CRHR1
MSRA
ACVR1C
HDAC9
SULT2B1
MAD1L1
TUBG2
GCK
CAVIN1
TCAM1P
RGS6
UFL1-AS1
SRRM2

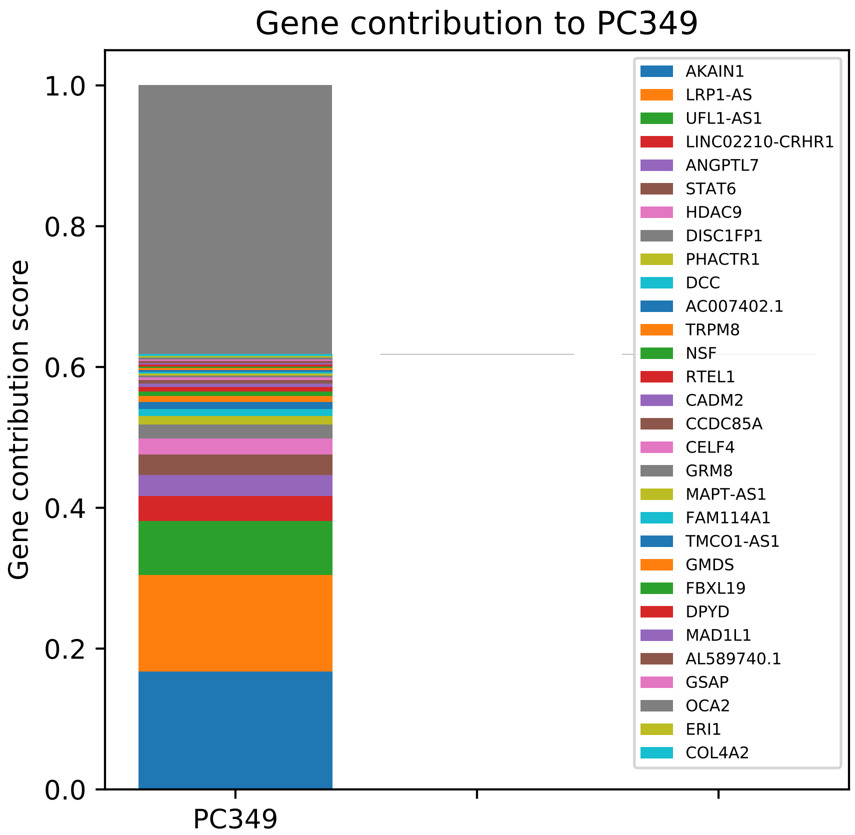

Gene contribution to PC349
Gene contribution score
1.0
0.8
0.6
0.4
0.2
0.0
PC349
AKAIN1
LRP1-AS
UFL1-AS1
LINC02210-CRHR1
ANGPTL7
STAT6
HDAC9
DISC1FP1
PHACTR1
DCC
AC007402.1
TRPM8
NSF
RTEL1
CADM2
CCDC85A
CELF4
GRM8
MAPT-AS1
FAM114A1
TMCO1-AS1
GMDS
FBXL19
DPYD
MAD1L1
AL589740.1
GSAP
OCA2
ERI1
COL4A2

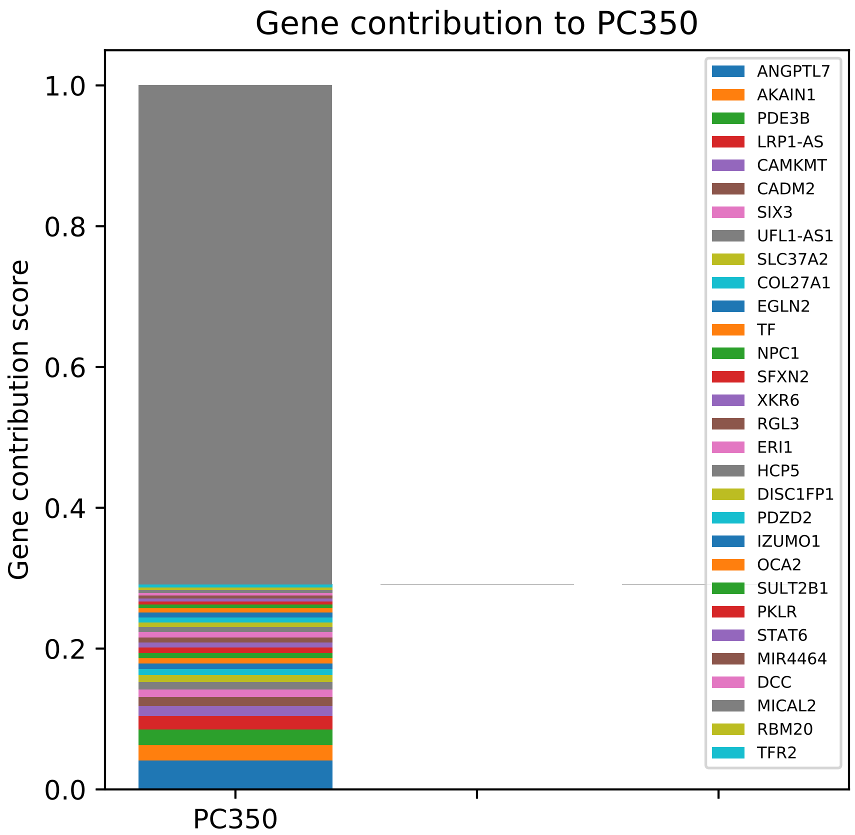
Gene contribution to PC350
Gene contribution score
0.0
0.2
0.4
0.6
0.8
1.0
PC350
ANGPTL7
AKAIN1
PDE3B
LRP1-AS
CAMKMT
CADM2
SIX3
UFL1-AS1
SLC37A2
COL27A1
EGLN2
TF
NPC1
SFXN2
XKR6
RGL3
ERI1
HCP5
DISC1FP1
PDZD2
IZUMO1
OCA2
SULT2B1
PKLR
STAT6
MIR4464
DCC
MICAL2
RBM20
TFR2

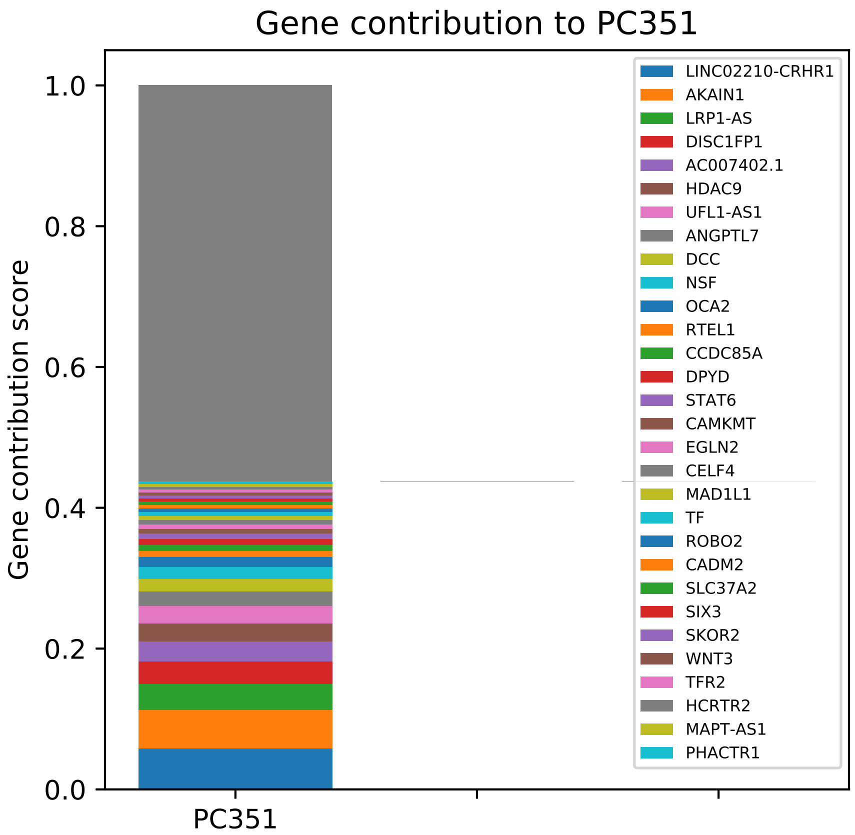
Gene contribution to PC351
Gene contribution score
1.0
0.8
0.6
0.4
0.2
0.0
PC351
LINC02210-CRHR1
AKAIN1
LRP1-AS
DISC1FP1
AC007402.1
HDAC9
UFL1-AS1
ANGPTL7
DCC
NSF
OCA2
RTEL1
CCDC85A
DPYD
STAT6
CAMKMT
EGLN2
CELF4
MAD1L1
TF
ROBO2
CADM2
SLC37A2
SIX3
SKOR2
WNT3
TFR2
HCRTR2
MAPT-AS1
PHACTR1

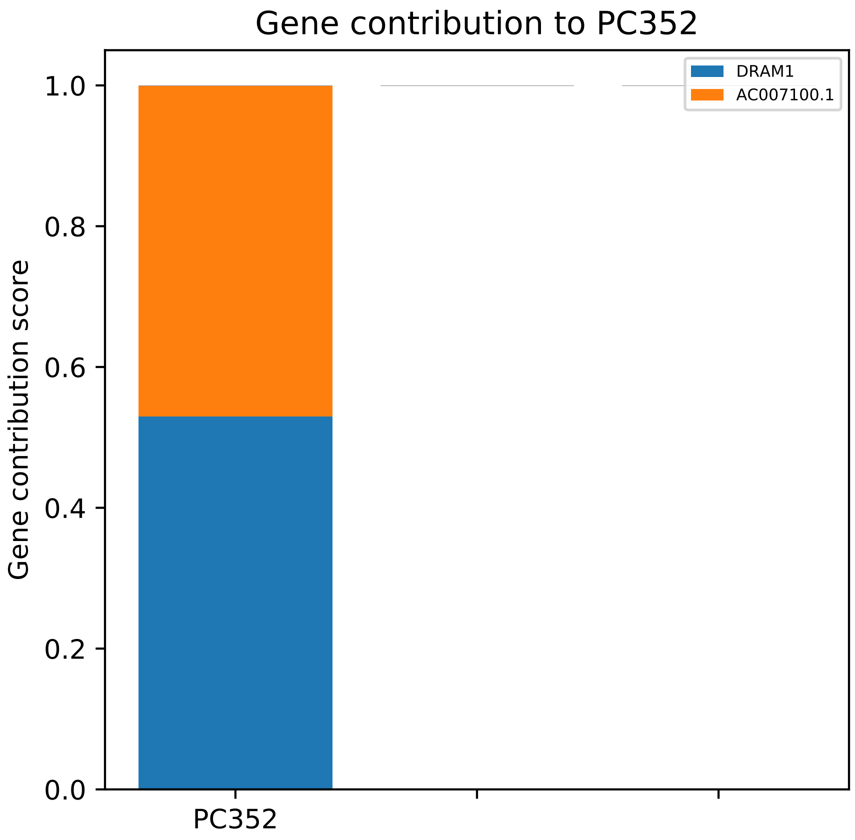
Gene contribution to PC352
DRAM1
AC007100.1
Gene contribution score
1.0
0.8
0.6
0.4
0.2
0.0
PC352

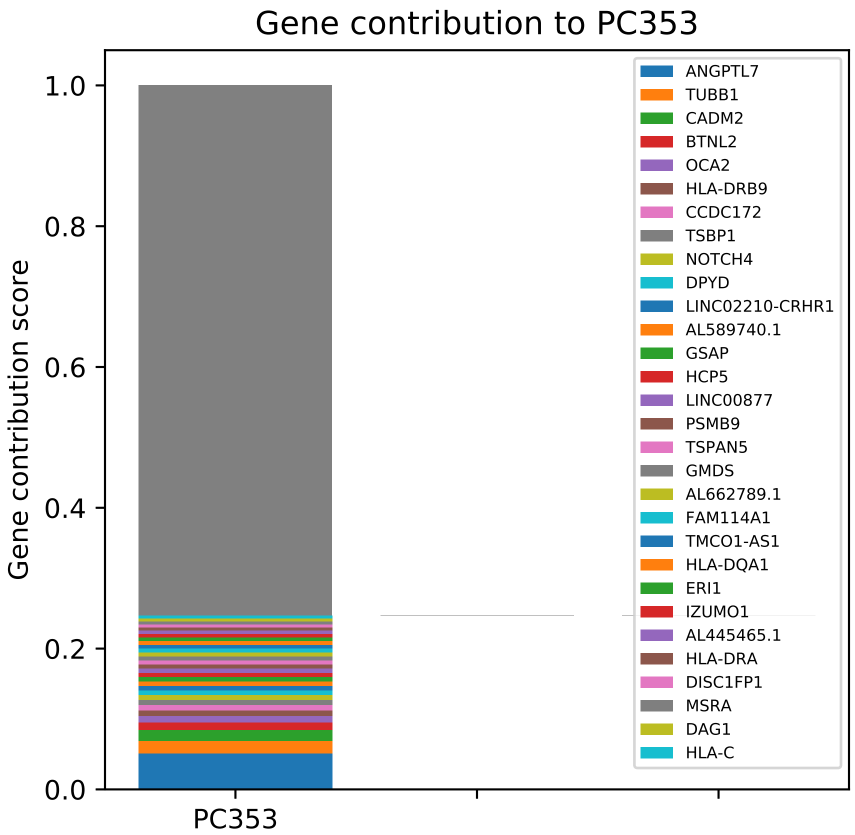
Gene contribution to PC353
Gene contribution score
1.0
0.8
0.6
0.4
0.2
0.0
PC353
ANGPTL7
TUBB1
CADM2
BTNL2
OCA2
HLA-DRB9
CCDC172
TSBP1
NOTCH4
DPYD
LINC02210-CRHR1
AL589740.1
GSAP
HCP5
LINC00877
PSMB9
TSPAN5
GMDS
AL662789.1
FAM114A1
TMCO1-AS1
HLA-DQA1
ERI1
IZUMO1
AL445465.1
HLA-DRA
DISC1FP1
MSRA
DAG1
HLA-C

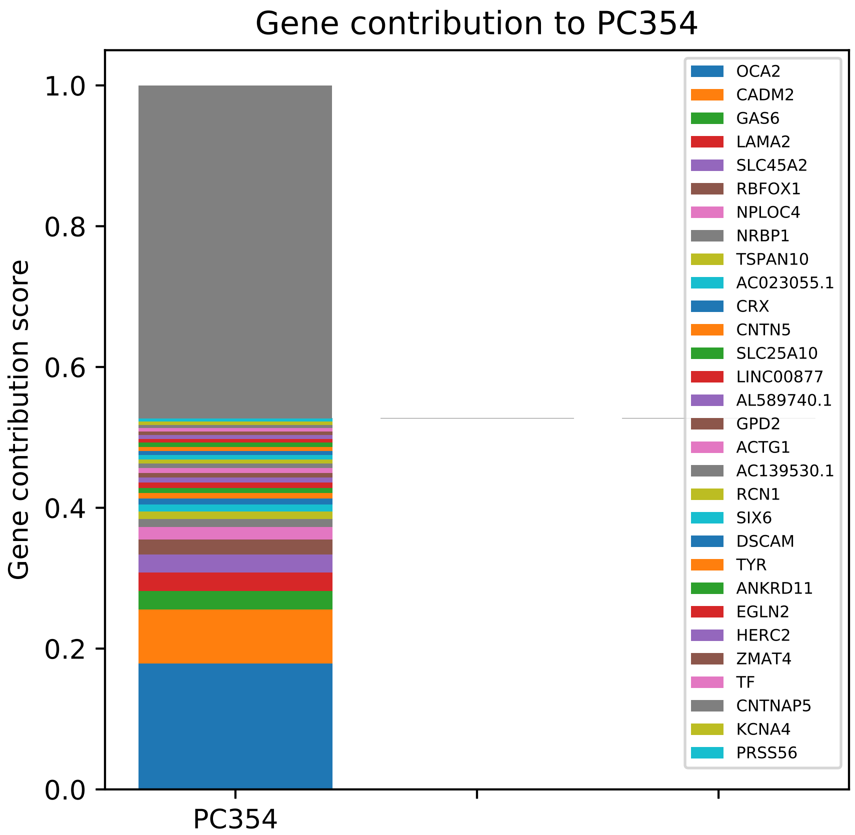
Gene contribution to PC354
Gene contribution score
1.0
0.8
0.6
0.4
0.2
0.0
PC354
OCA2
CADM2
GAS6
LAMA2
SLC45A2
RBFOX1
NPLOC4
NRBP1
TSPAN10
AC023055.1
CRX
CNTN5
SLC25A10
LINC00877
AL589740.1
GPD2
ACTG1
AC139530.1
RCN1
SIX6
DSCAM
TYR
ANKRD11
EGLN2
HERC2
ZMAT4
TF
CNTNAP5
KCNA4
PRSS56

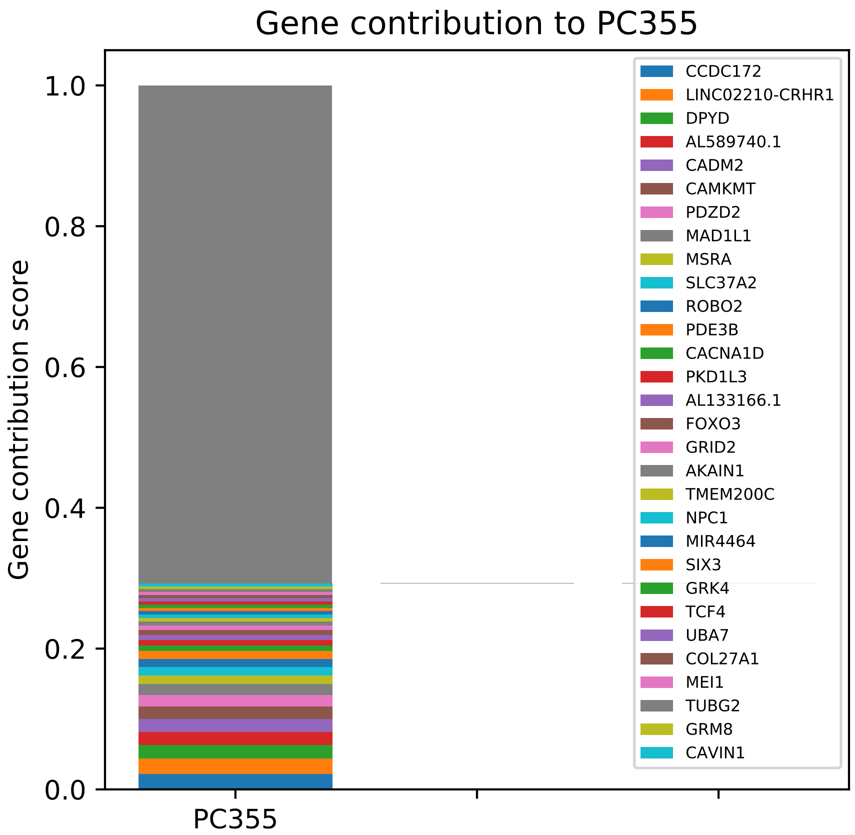

Gene contribution to PC355
Gene contribution score
1.0
0.8
0.6
0.4
0.2
0.0
PC355
CCDC172
LINC02210-CRHR1
DPYD
AL589740.1
CADM2
CAMKMT
PDZD2
MAD1L1
MSRA
SLC37A2
ROBO2
PDE3B
CACNA1D
PKD1L3
AL133166.1
FOXO3
GRID2
AKAIN1
TMEM200C
NPC1
MIR4464
SIX3
GRK4
TCF4
UBA7
COL27A1
MEI1
TUBG2
GRM8
CAVIN1

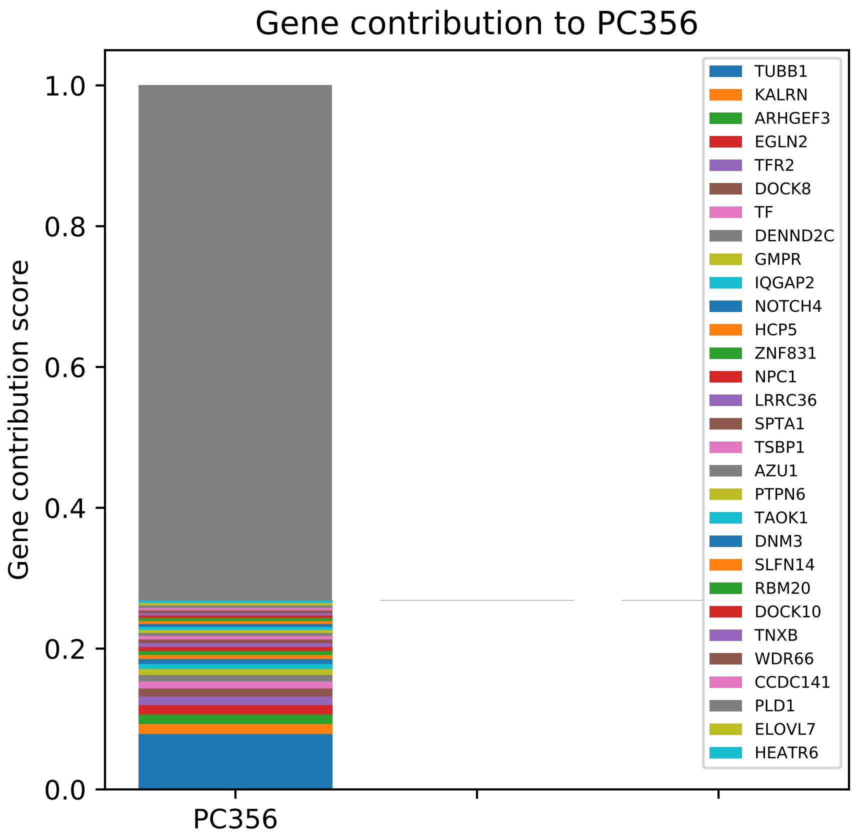
Gene contribution to PC356
Gene contribution score
1.0
0.8
0.6
0.4
0.2
0.0
PC356
TUBB1
KALRN
ARHGEF3
EGLN2
TFR2
DOCK8
TF
DENND2C
GMPR
IQGAP2
NOTCH4
HCP5
ZNF831
NPC1
LRRC36
SPTA1
TSBP1
AZU1
PTPN6
TAOK1
DNM3
SLFN14
RBM20
DOCK10
TNXB
WDR66
CCDC141
PLD1
ELOVL7
HEATR6

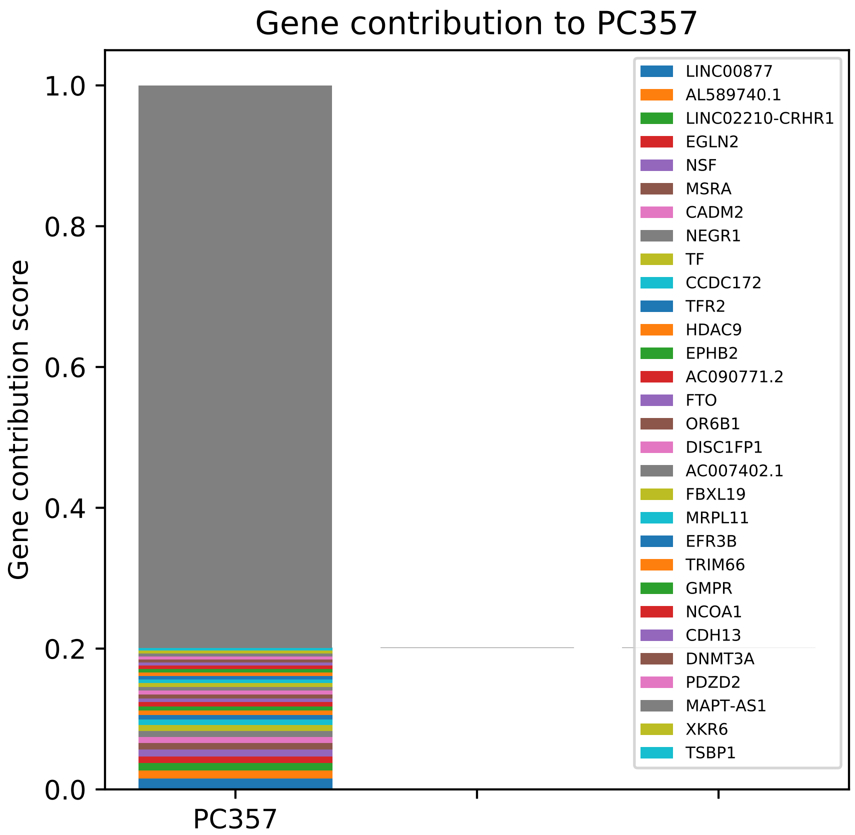

Gene contribution to PC357
Gene contribution score
1.0
0.8
0.6
0.4
0.2
0.0
PC357
LINC00877
AL589740.1
LINC02210-CRHR1
EGLN2
NSF
MSRA
CADM2
NEGR1
TF
CCDC172
TFR2
HDAC9
EPHB2
AC090771.2
FTO
OR6B1
DISC1FP1
AC007402.1
FBXL19
MRPL11
EFR3B
TRIM66
GMPR
NCOA1
CDH13
DNMT3A
PDZD2
MAPT-AS1
XKR6
TSBP1

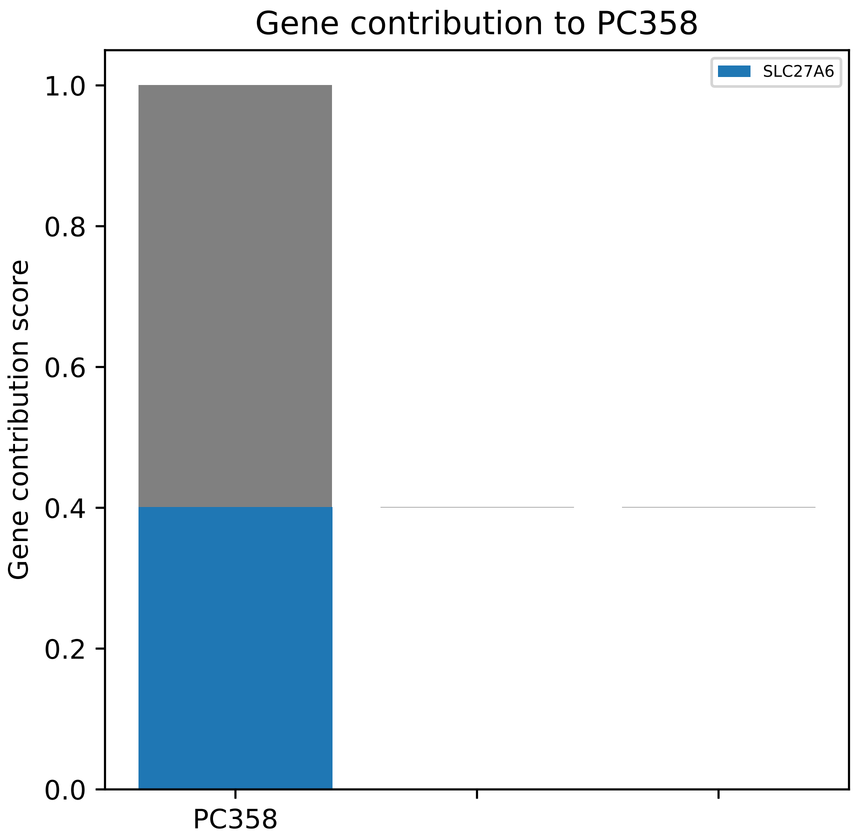
Gene contribution to PC358
SLC27A6
Gene contribution score
1.0
0.8
0.6
0.4
0.2
0.0
PC358

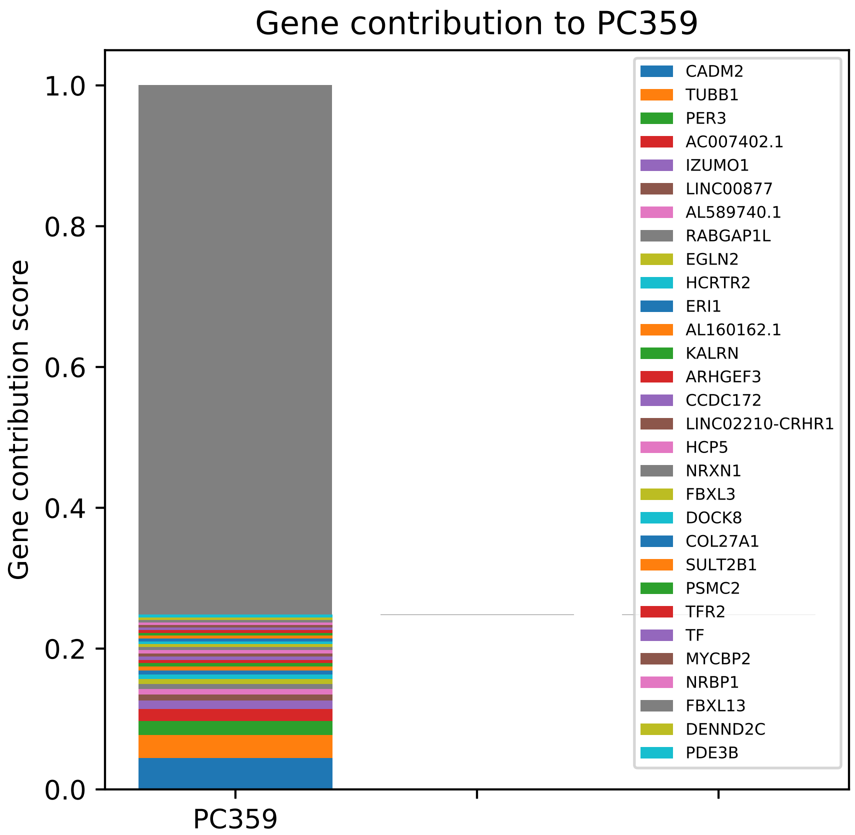

Gene contribution to PC359
Gene contribution score
1.0
0.8
0.6
0.4
0.2
0.0
PC359
CADM2
TUBB1
PER3
AC007402.1
IZUMO1
LINC00877
AL589740.1
RABGAP1L
EGLN2
HCRTR2
ERI1
AL160162.1
KALRN
ARHGEF3
CCDC172
LINC02210-CRHR1
HCP5
NRXN1
FBXL3
DOCK8
COL27A1
SULT2B1
PSMC2
TFR2
TF
MYCBP2
NRBP1
FBXL13
DENND2C
PDE3B

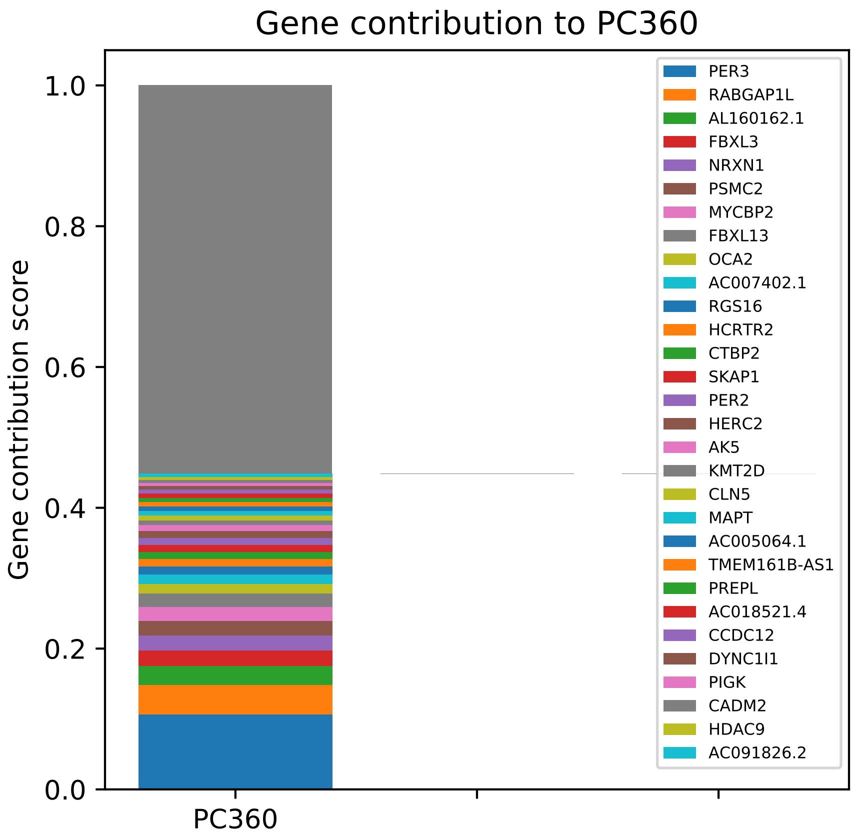
Gene contribution to PC360
Gene contribution score
1.0
0.8
0.6
0.4
0.2
0.0
PC360
PER3
RABGAP1L
AL160162.1
FBXL3
NRXN1
PSMC2
MYCBP2
FBXL13
OCA2
AC007402.1
RGS16
HCRTR2
CTBP2
SKAP1
PER2
HERC2
AK5
KMT2D
CLN5
MAPT
AC005064.1
TMEM161B-AS1
PREPL
AC018521.4
CCDC12
DYNC1I1
PIGK
CADM2
HDAC9
AC091826.2

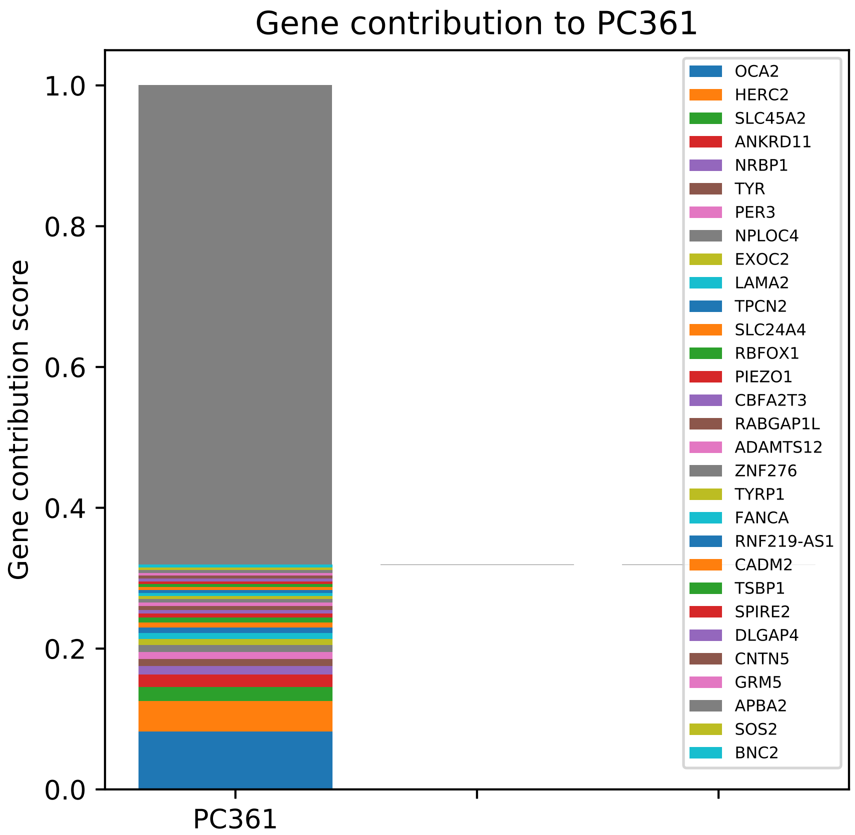
Gene contribution to PC361
Gene contribution score
1.0
0.8
0.6
0.4
0.2
0.0
PC361
OCA2
HERC2
SLC45A2
ANKRD11
NRBP1
TYR
PER3
NPLOC4
EXOC2
LAMA2
TPCN2
SLC24A4
RBFOX1
PIEZO1
CBFA2T3
RABGAP1L
ADAMTS12
ZNF276
TYRP1
FANCA
RNF219-AS1
CADM2
TSBP1
SPIRE2
DLGAP4
CNTN5
GRM5
APBA2
SOS2
BNC2

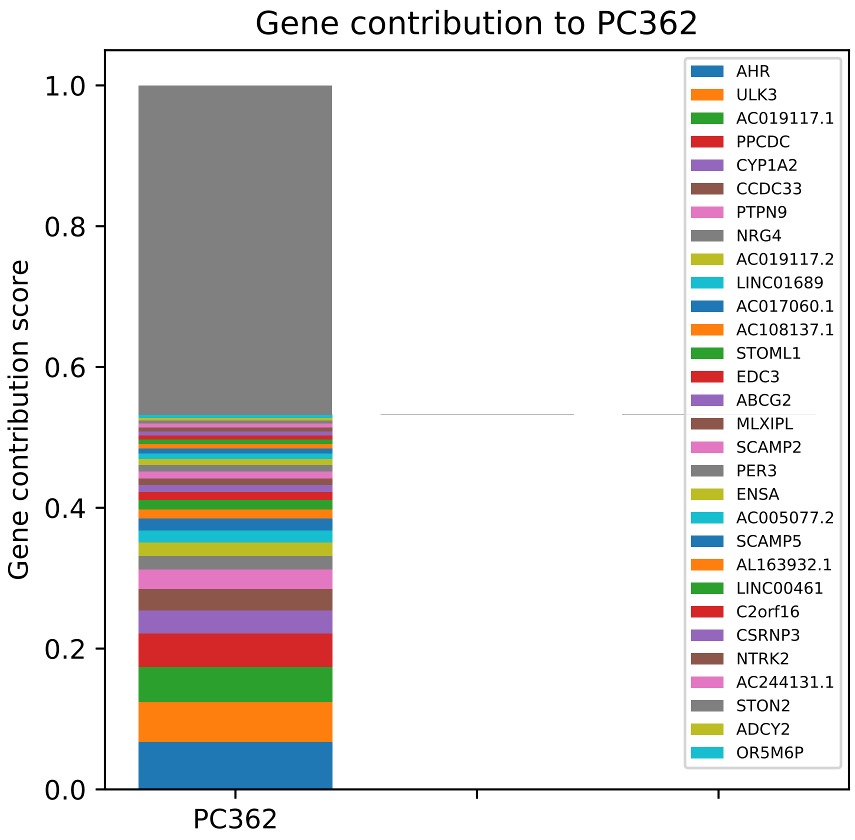

Gene contribution to PC362
Gene contribution score
1.0
0.8
0.6
0.4
0.2
0.0
PC362
AHR
ULK3
AC019117.1
PPCDC
CYP1A2
CCDC33
PTPN9
NRG4
AC019117.2
LINC01689
AC017060.1
AC108137.1
STOML1
EDC3
ABCG2
MLXIPL
SCAMP2
PER3
ENSA
AC005077.2
SCAMP5
AL163932.1
LINC00461
C2orf16
CSRNP3
NTRK2
AC244131.1
STON2
ADCY2
OR5M6P

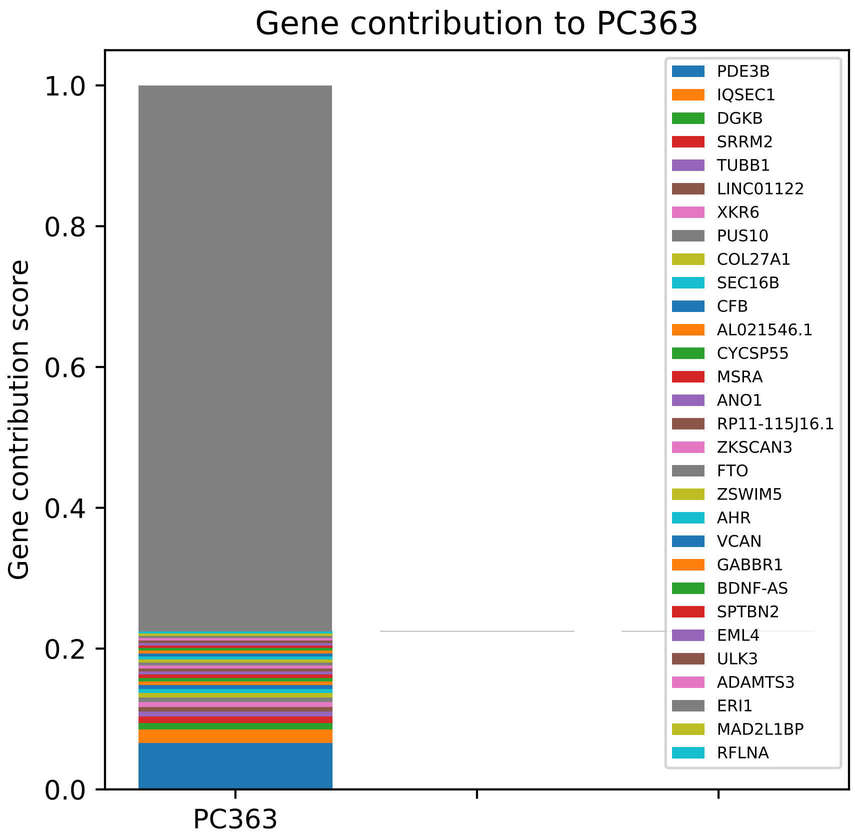

Gene contribution to PC363
Gene contribution score
1.0
0.8
0.6
0.4
0.2
0.0
PC363
PDE3B
IQSEC1
DGKB
SRRM2
TUBB1
LINC01122
XKR6
PUS10
COL27A1
SEC16B
CFB
AL021546.1
CYCSP55
MSRA
ANO1
RP11-115J16.1
ZKSCAN3
FTO
ZSWIM5
AHR
VCAN
GABBR1
BDNF-AS
SPTBN2
EML4
ULK3
ADAMTS3
ERI1
MAD2L1BP
RFLNA

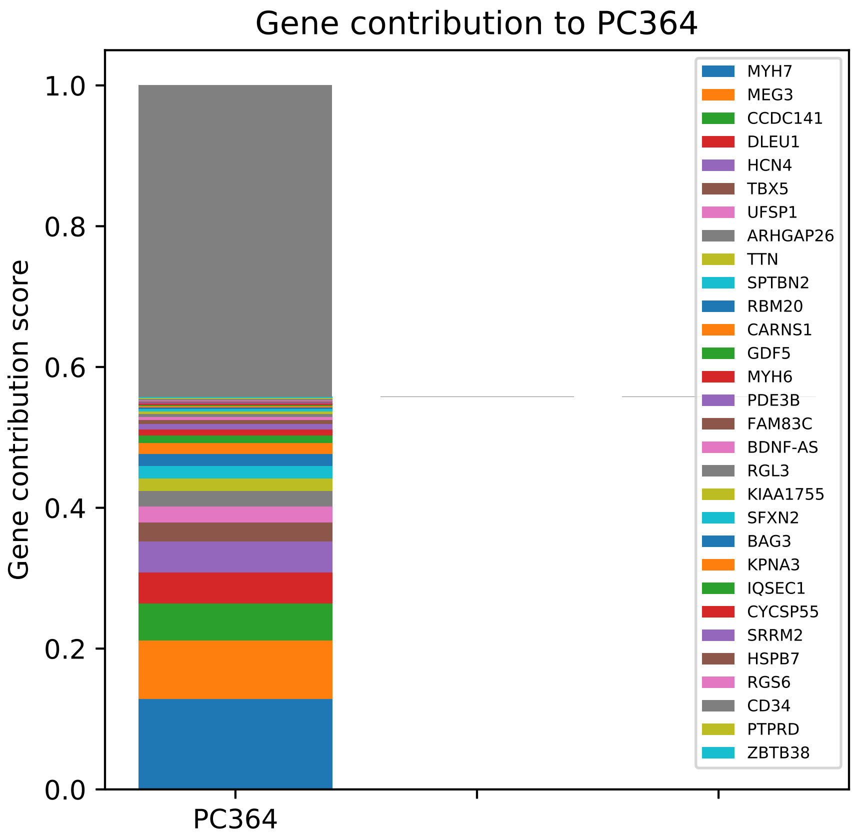

Gene contribution to PC364
Gene contribution score
1.0
0.8
0.6
0.4
0.2
0.0
PC364
MYH7
MEG3
CCDC141
DLEU1
HCN4
TBX5
UFSP1
ARHGAP26
TTN
SPTBN2
RBM20
CARNS1
GDF5
MYH6
PDE3B
FAM83C
BDNF-AS
RGL3
KIAA1755
SFXN2
BAG3
KPNA3
IQSEC1
CYCSP55
SRRM2
HSPB7
RGS6
CD34
PTPRD
ZBTB38

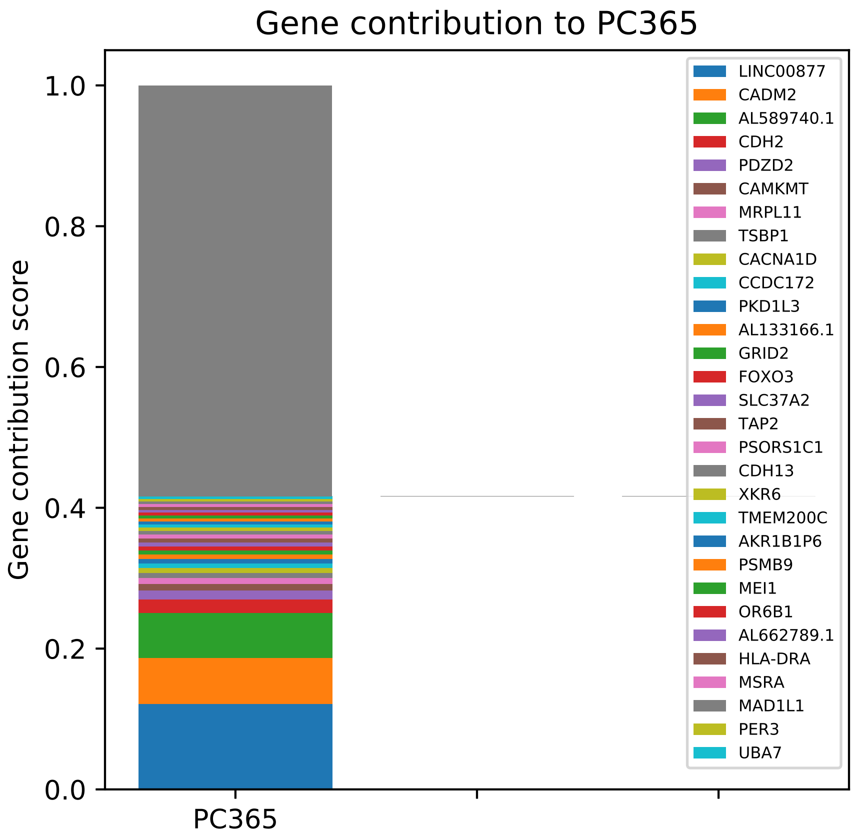

Gene contribution to PC365
Gene contribution score
1.0
0.8
0.6
0.4
0.2
0.0
PC365
LINC00877
CADM2
AL589740.1
CDH2
PDZD2
CAMKMT
MRPL11
TSBP1
CACNA1D
CCDC172
PKD1L3
AL133166.1
GRID2
FOXO3
SLC37A2
TAP2
PSORS1C1
CDH13
XKR6
TMEM200C
AKR1B1P6
PSMB9
MEI1
OR6B1
AL662789.1
HLA-DRA
MSRA
MAD1L1
PER3
UBA7

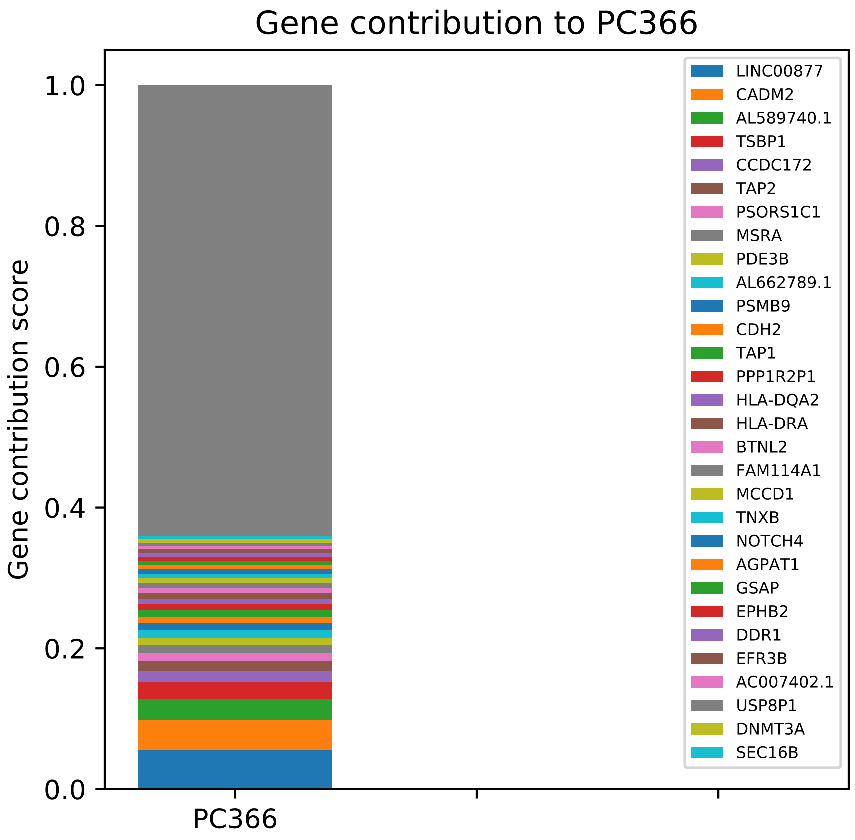

Gene contribution to PC366
Gene contribution score
1.0
0.8
0.6
0.4
0.2
0.0
PC366
LINC00877
CADM2
AL589740.1
TSBP1
CCDC172
TAP2
PSORS1C1
MSRA
PDE3B
AL662789.1
PSMB9
CDH2
TAP1
PPP1R2P1
HLA-DQA2
HLA-DRA
BTNL2
FAM114A1
MCCD1
TNXB
NOTCH4
AGPAT1
GSAP
EPHB2
DDR1
EFR3B
AC007402.1
USP8P1
DNMT3A
SEC16B

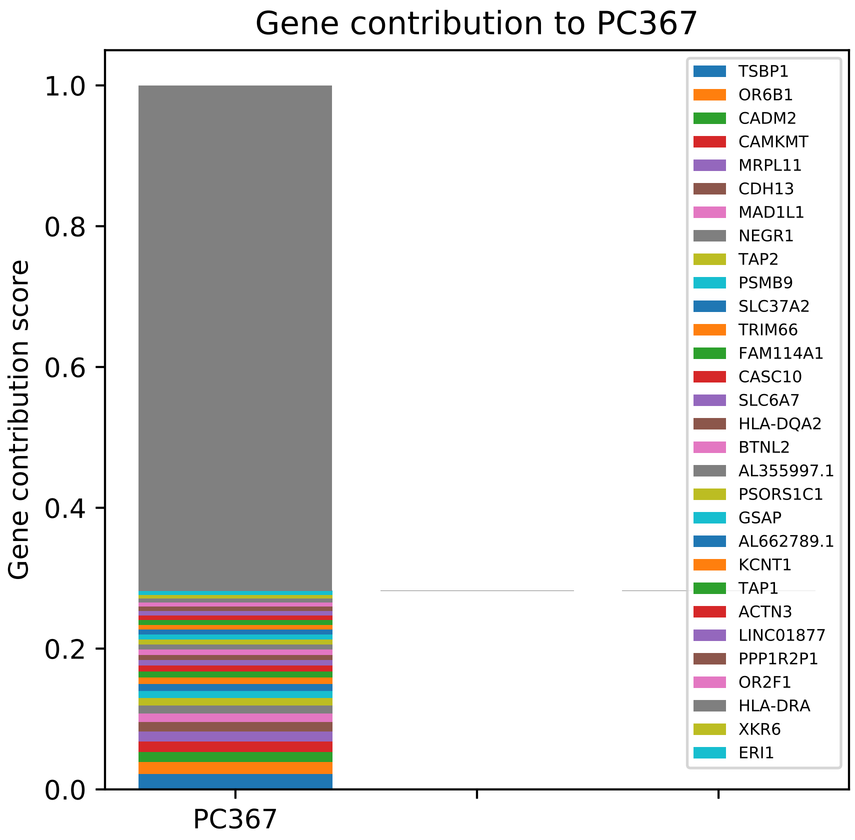
Gene contribution to PC367
Gene contribution score
0.0
0.2
0.4
0.6
0.8
1.0
PC367
TSBP1
OR6B1
CADM2
CAMKMT
MRPL11
CDH13
MAD1L1
NEGR1
TAP2
PSMB9
SLC37A2
TRIM66
FAM114A1
CASC10
SLC6A7
HLA-DQA2
BTNL2
AL355997.1
PSORS1C1
GSAP
AL662789.1
KCNT1
TAP1
ACTN3
LINC01877
PPP1R2P1
OR2F1
HLA-DRA
XKR6
ERI1

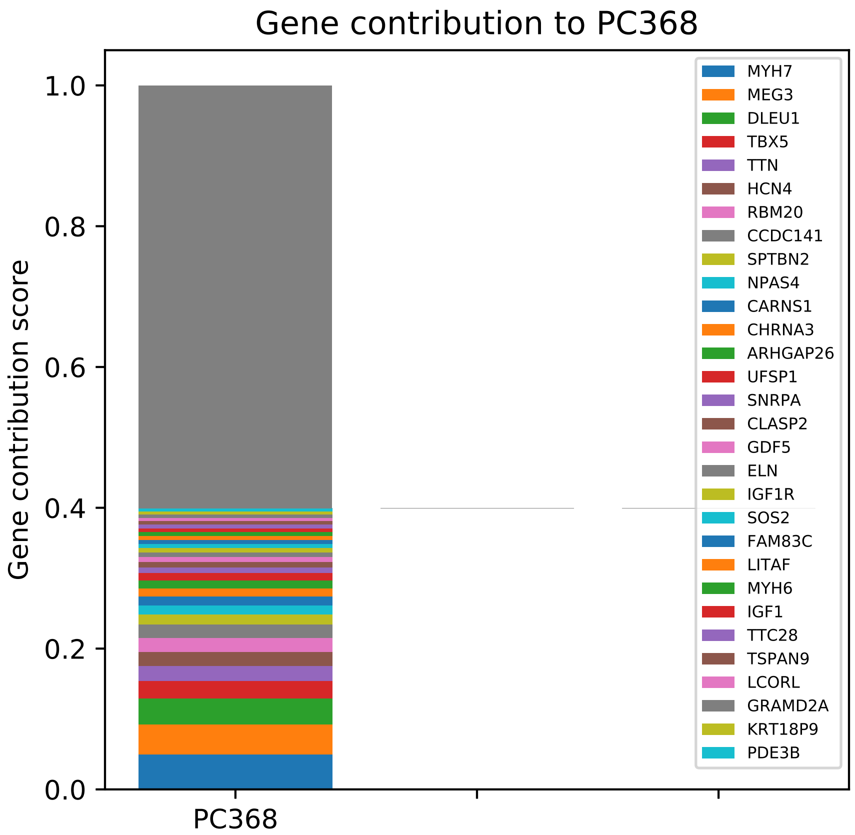
Gene contribution to PC368
Gene contribution score
1.0
0.8
0.6
0.4
0.2
0.0
PC368
MYH7
MEG3
DLEU1
TBX5
TTN
HCN4
RBM20
CCDC141
SPTBN2
NPAS4
CARNS1
CHRNA3
ARHGAP26
UFSP1
SNRPA
CLASP2
GDF5
ELN
IGF1R
SOS2
FAM83C
LITAF
MYH6
IGF1
TTC28
TSPAN9
LCORL
GRAMD2A
KRT18P9
PDE3B

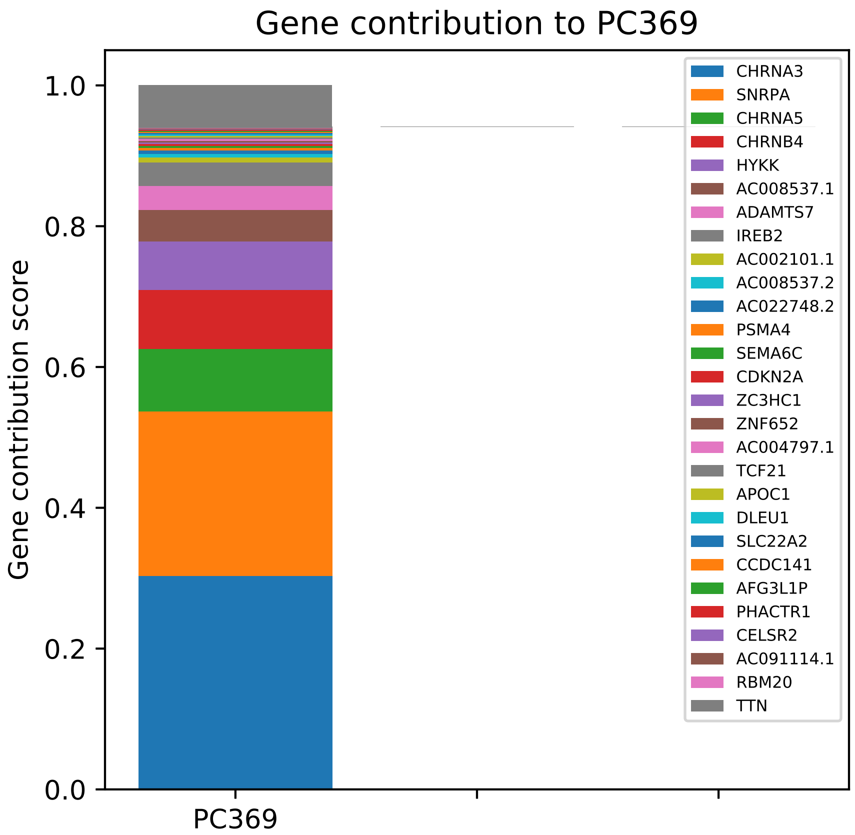
Gene contribution to PC369
Gene contribution score
1.0
0.8
0.6
0.4
0.2
0.0
PC369
CHRNA3
SNRPA
CHRNA5
CHRNB4
HYKK
AC008537.1
ADAMTS7
IREB2
AC002101.1
AC008537.2
AC022748.2
PSMA4
SEMA6C
CDKN2A
ZC3HC1
ZNF652
AC004797.1
TCF21
APOC1
DLEU1
SLC22A2
CCDC141
AFG3L1P
PHACTR1
CELSR2
AC091114.1
RBM20
TTN

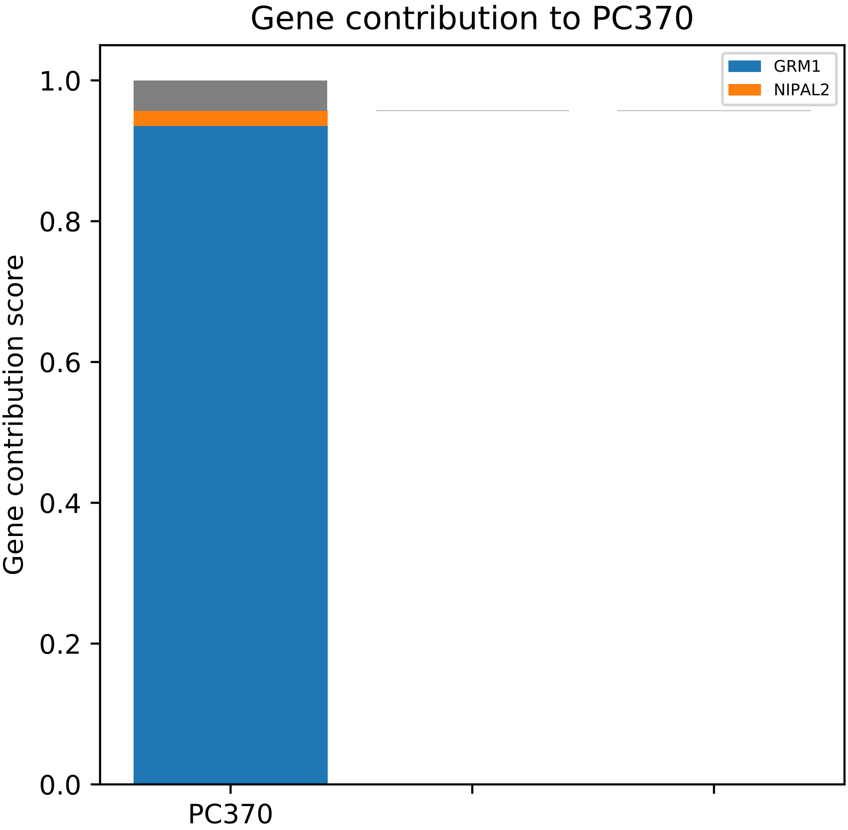
Gene contribution to PC370
GRM1
NIPAL2
Gene contribution score
1.0
0.8
0.6
0.4
0.2
0.0
PC370

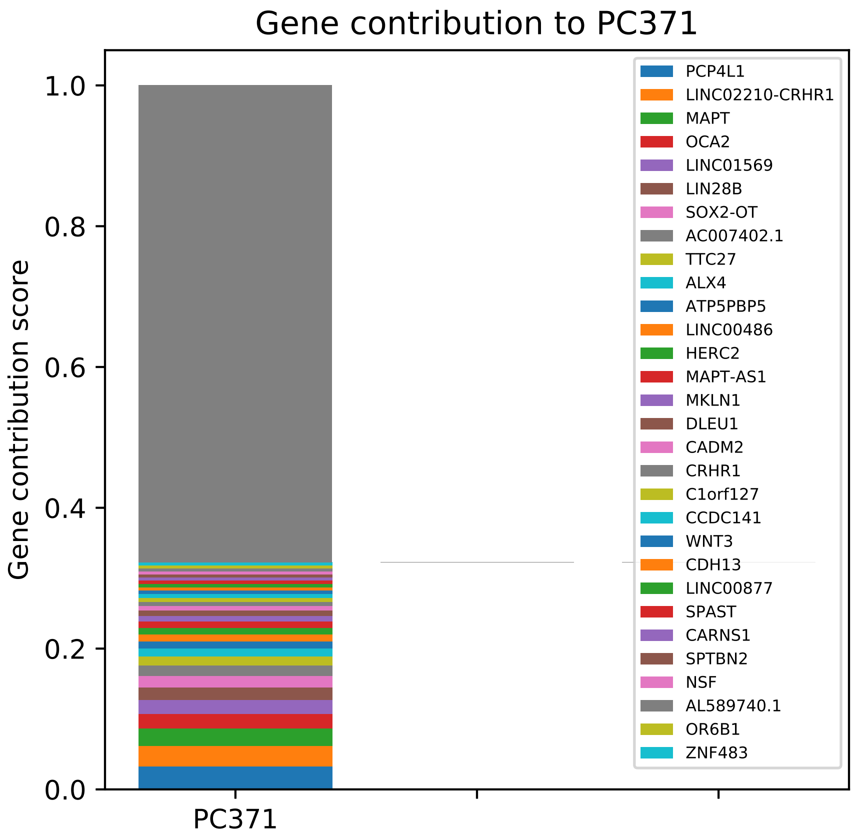

Gene contribution to PC371
Gene contribution score
1.0
0.8
0.6
0.4
0.2
0.0
PC371
PCP4L1
LINC02210-CRHR1
MAPT
OCA2
LINC01569
LIN28B
SOX2-OT
AC007402.1
TTC27
ALX4
ATP5PBP5
LINC00486
HERC2
MAPT-AS1
MKLN1
DLEU1
CADM2
CRHR1
C1orf127
CCDC141
WNT3
CDH13
LINC00877
SPAST
CARNS1
SPTBN2
NSF
AL589740.1
OR6B1
ZNF483

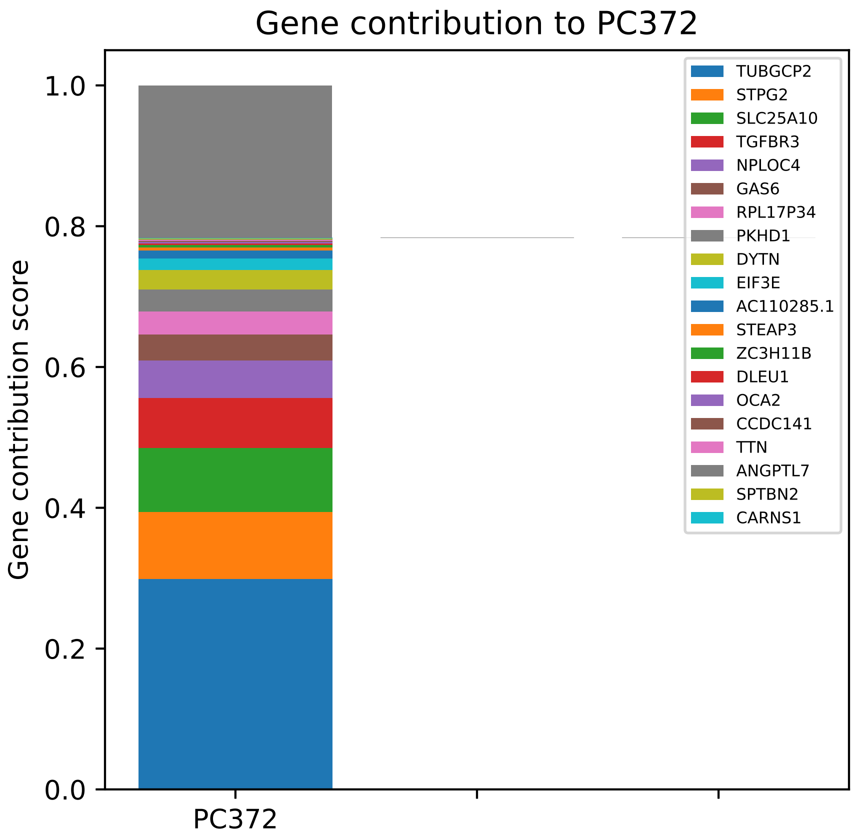
Gene contribution to PC372
Gene contribution score
1.0
0.8
0.6
0.4
0.2
0.0
PC372
TUBGCP2
STPG2
SLC25A10
TGFBR3
NPLOC4
GAS6
RPL17P34
PKHD1
DYTN
EIF3E
AC110285.1
STEAP3
ZC3H11B
DLEU1
OCA2
CCDC141
TTN
ANGPTL7
SPTBN2
CARNS1

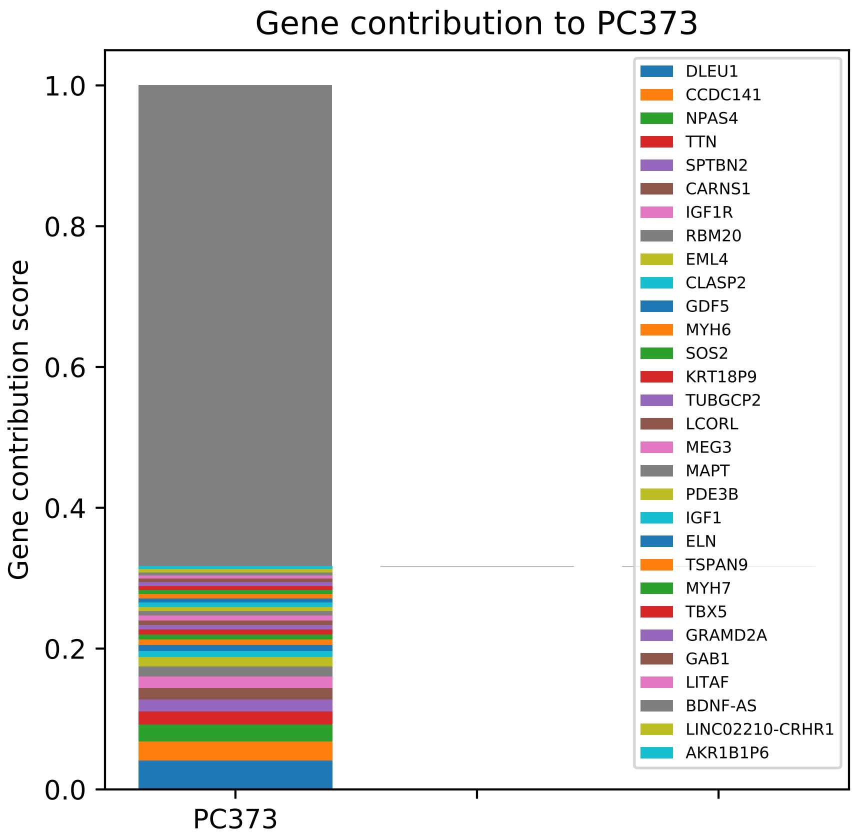

Gene contribution to PC373
Gene contribution score
1.0
0.8
0.6
0.4
0.2
0.0
PC373
DLEU1
CCDC141
NPAS4
TTN
SPTBN2
CARNS1
IGF1R
RBM20
EML4
CLASP2
GDF5
MYH6
SOS2
KRT18P9
TUBGCP2
LCORL
MEG3
MAPT
PDE3B
IGF1
ELN
TSPAN9
MYH7
TBX5
GRAMD2A
GAB1
LITAF
BDNF-AS
LINC02210-CRHR1
AKR1B1P6

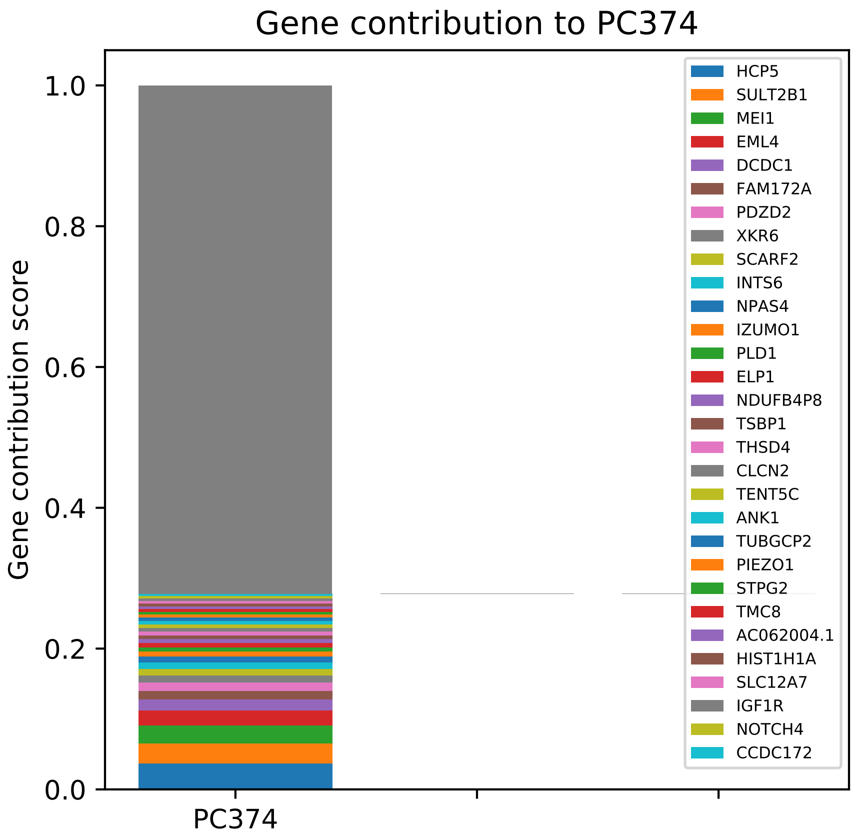
Gene contribution to PC374
Gene contribution score
1.0
0.8
0.6
0.4
0.2
0.0
PC374
HCP5
SULT2B1
MEI1
EML4
DCDC1
FAM172A
PDZD2
XKR6
SCARF2
INTS6
NPAS4
IZUMO1
PLD1
ELP1
NDUFB4P8
TSBP1
THSD4
CLCN2
TENT5C
ANK1
TUBGCP2
PIEZO1
STPG2
TMC8
AC062004.1
HIST1H1A
SLC12A7
IGF1R
NOTCH4
CCDC172

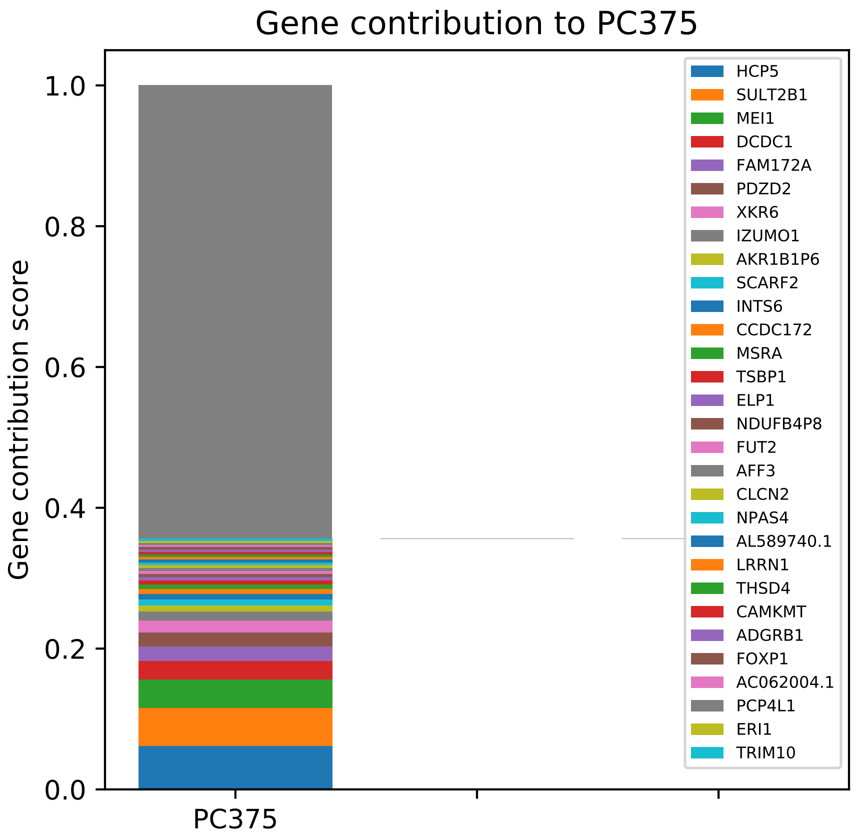
Gene contribution to PC375
Gene contribution score
1.0
0.8
0.6
0.4
0.2
0.0
PC375
HCP5
SULT2B1
MEI1
DCDC1
FAM172A
PDZD2
XKR6
IZUMO1
AKR1B1P6
SCARF2
INTS6
CCDC172
MSRA
TSBP1
ELP1
NDUFB4P8
FUT2
AFF3
CLCN2
NPAS4
AL589740.1
LRRN1
THSD4
CAMKMT
ADGRB1
FOXP1
AC062004.1
PCP4L1
ERI1
TRIM10

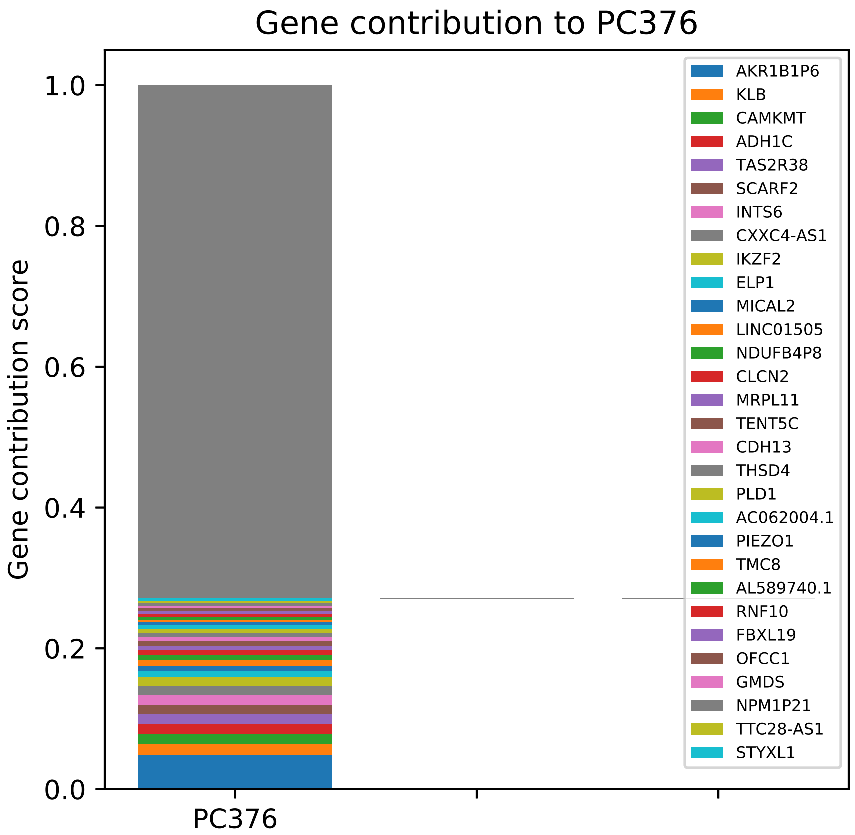

Gene contribution to PC376
Gene contribution score
1.0
0.8
0.6
0.4
0.2
0.0
PC376
AKR1B1P6
KLB
CAMKMT
ADH1C
TAS2R38
SCARF2
INTS6
CXXC4-AS1
IKZF2
ELP1
MICAL2
LINC01505
NDUFB4P8
CLCN2
MRPL11
TENT5C
CDH13
THSD4
PLD1
AC062004.1
PIEZO1
TMC8
AL589740.1
RNF10
FBXL19
OFCC1
GMDS
NPM1P21
TTC28-AS1
STYXL1

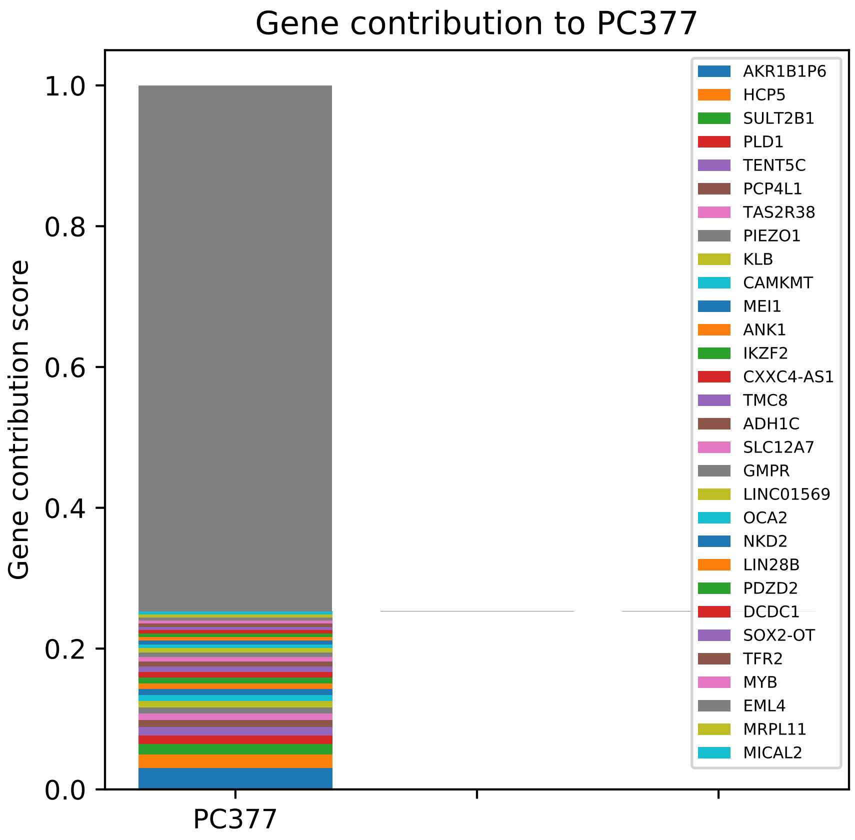

Gene contribution to PC377
Gene contribution score
1.0
0.8
0.6
0.4
0.2
0.0
PC377
AKR1B1P6
HCP5
SULT2B1
PLD1
TENT5C
PCP4L1
TAS2R38
PIEZO1
KLB
CAMKMT
MEI1
ANK1
IKZF2
CXXC4-AS1
TMC8
ADH1C
SLC12A7
GMPR
LINC01569
OCA2
NKD2
LIN28B
PDZD2
DCDC1
SOX2-OT
TFR2
MYB
EML4
MRPL11
MICAL2

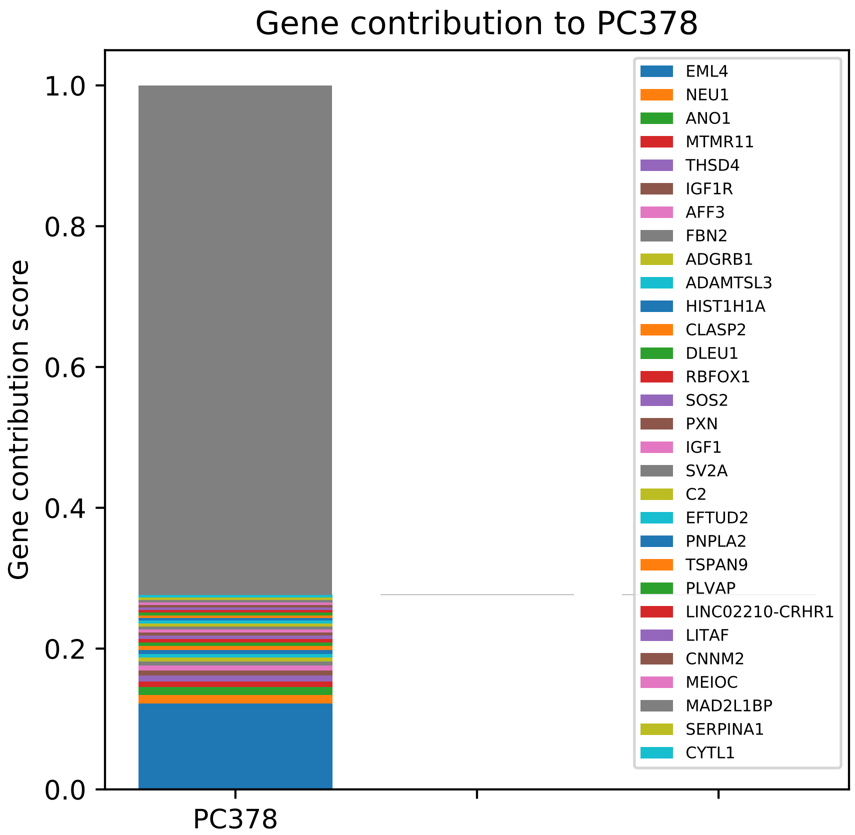

Gene contribution to PC378
Gene contribution score
1.0
0.8
0.6
0.4
0.2
0.0
PC378
EML4
NEU1
ANO1
MTMR11
THSD4
IGF1R
AFF3
FBN2
ADGRB1
ADAMTSL3
HIST1H1A
CLASP2
DLEU1
RBFOX1
SOS2
PXN
IGF1
SV2A
C2
EFTUD2
PNPLA2
TSPAN9
PLVAP
LINC02210-CRHR1
LITAF
CNNM2
MEIOC
MAD2L1BP
SERPINA1
CYTL1

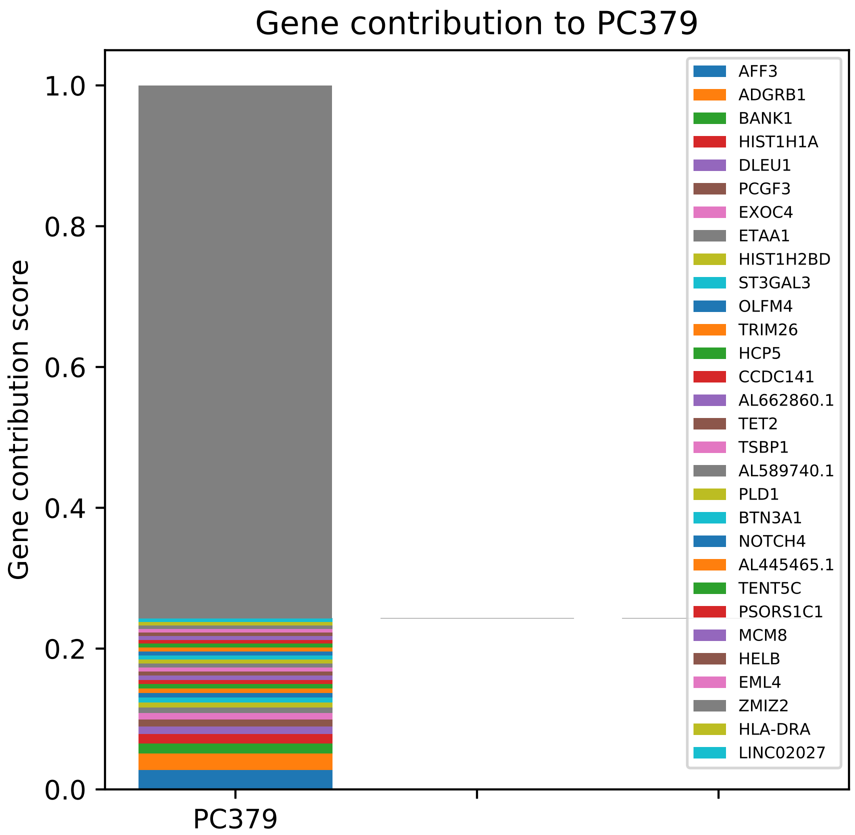
Gene contribution to PC379
Gene contribution score
1.0
0.8
0.6
0.4
0.2
0.0
PC379
AFF3
ADGRB1
BANK1
HIST1H1A
DLEU1
PCGF3
EXOC4
ETAA1
HIST1H2BD
ST3GAL3
OLFM4
TRIM26
HCP5
CCDC141
AL662860.1
TET2
TSBP1
AL589740.1
PLD1
BTN3A1
NOTCH4
AL445465.1
TENT5C
PSORS1C1
MCM8
HELB
EML4
ZMIZ2
HLA-DRA
LINC02027

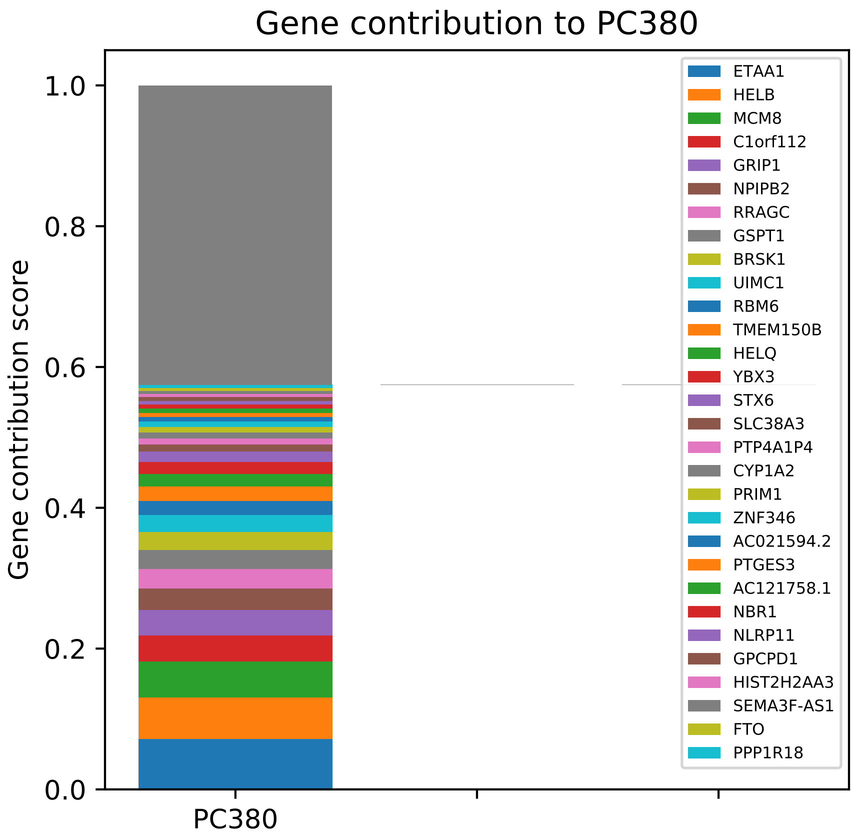

Gene contribution to PC380
Gene contribution score
1.0
0.8
0.6
0.4
0.2
0.0
PC380
ETAA1
HELB
MCM8
C1orf112
GRIP1
NPIPB2
RRAGC
GSPT1
BRSK1
UIMC1
RBM6
TMEM150B
HELQ
YBX3
STX6
SLC38A3
PTP4A1P4
CYP1A2
PRIM1
ZNF346
AC021594.2
PTGES3
AC121758.1
NBR1
NLRP11
GPCPD1
HIST2H2AA3
SEMA3F-AS1
FTO
PPP1R18

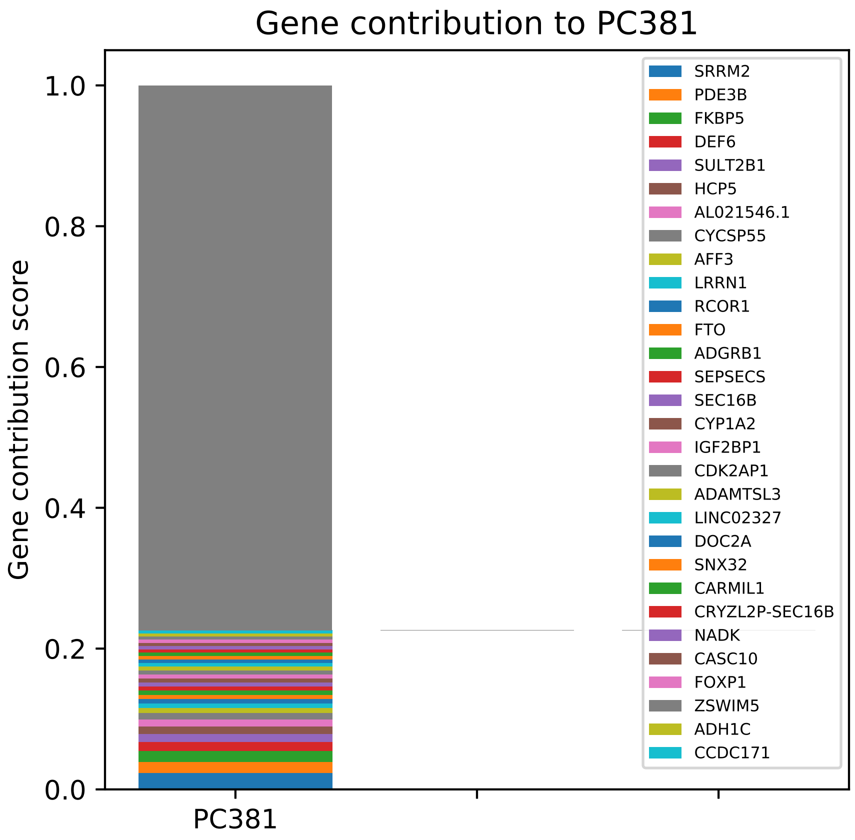

Gene contribution to PC381
Gene contribution score
0.0
0.2
0.4
0.6
0.8
1.0
PC381
SRRM2
PDE3B
FKBP5
DEF6
SULT2B1
HCP5
AL021546.1
CYCSP55
AFF3
LRRN1
RCOR1
FTO
ADGRB1
SEPSECS
SEC16B
CYP1A2
IGF2BP1
CDK2AP1
ADAMTSL3
LINC02327
DOC2A
SNX32
CARMIL1
CRYZL2P-SEC16B
NADK
CASC10
FOXP1
ZSWIM5
ADH1C
CCDC171

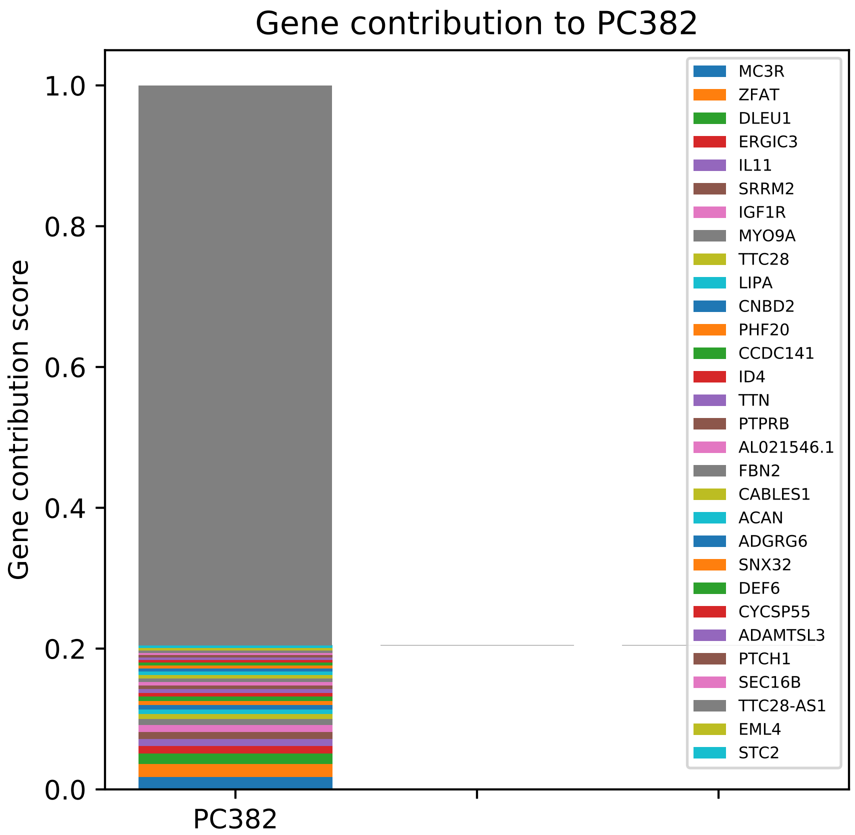

Gene contribution to PC382
Gene contribution score
1.0
0.8
0.6
0.4
0.2
0.0
PC382
MC3R
ZFAT
DLEU1
ERGIC3
IL11
SRRM2
IGF1R
MYO9A
TTC28
LIPA
CNBD2
PHF20
CCDC141
ID4
TTN
PTPRB
AL021546.1
FBN2
CABLES1
ACAN
ADGRG6
SNX32
DEF6
CYCSP55
ADAMTSL3
PTCH1
SEC16B
TTC28-AS1
EML4
STC2

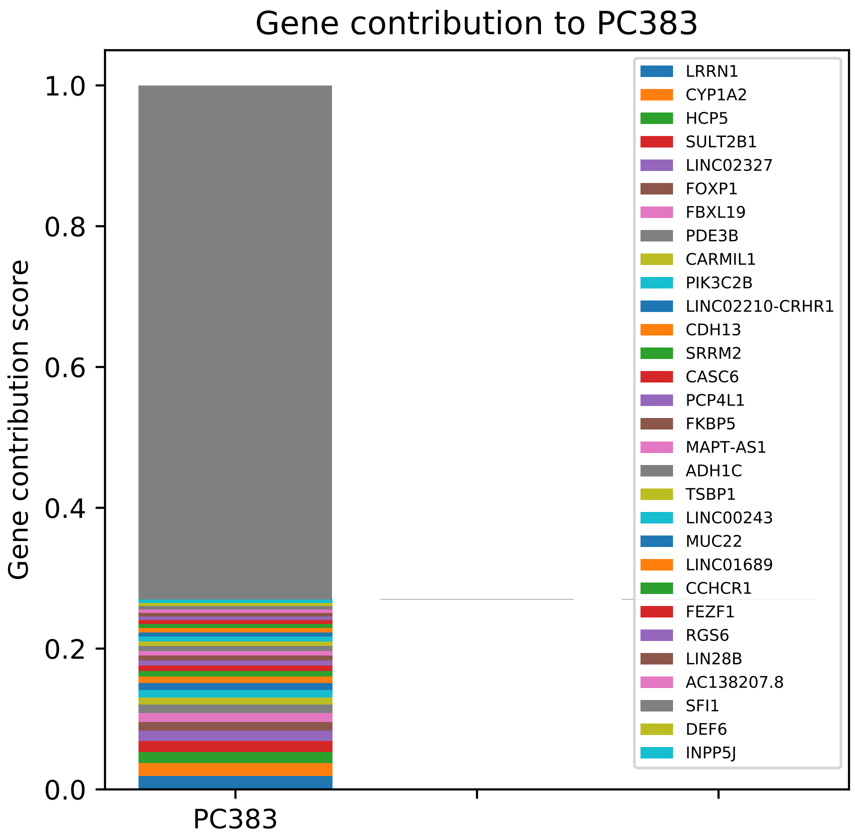

Gene contribution to PC383
Gene contribution score
1.0
0.8
0.6
0.4
0.2
0.0
PC383
LRRN1
CYP1A2
HCP5
SULT2B1
LINC02327
FOXP1
FBXL19
PDE3B
CARMIL1
PIK3C2B
LINC02210-CRHR1
CDH13
SRRM2
CASC6
PCP4L1
FKBP5
MAPT-AS1
ADH1C
TSBP1
LINC00243
MUC22
LINC01689
CCHCR1
FEZF1
RGS6
LIN28B
AC138207.8
SFI1
DEF6
INPP5J

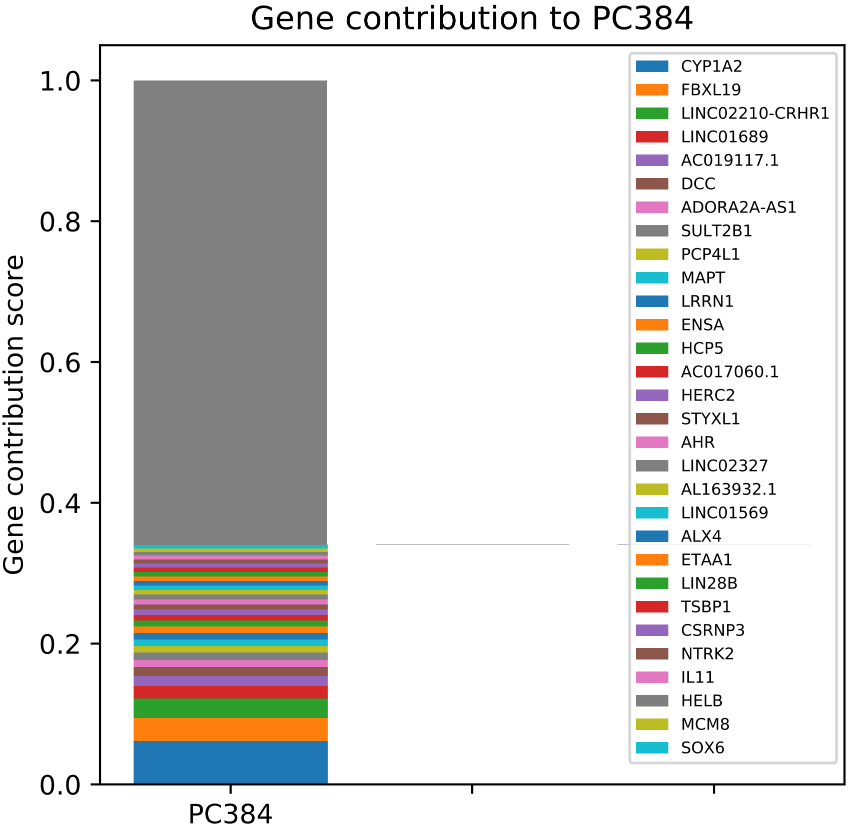

Gene contribution to PC384
Gene contribution score
1.0
0.8
0.6
0.4
0.2
0.0
PC384
CYP1A2
FBXL19
LINC02210-CRHR1
LINC01689
AC019117.1
DCC
ADORA2A-AS1
SULT2B1
PCP4L1
MAPT
LRRN1
ENSA
HCP5
AC017060.1
HERC2
STYXL1
AHR
LINC02327
AL163932.1
LINC01569
ALX4
ETAA1
LIN28B
TSBP1
CSRNP3
NTRK2
IL11
HELB
MCM8
SOX6

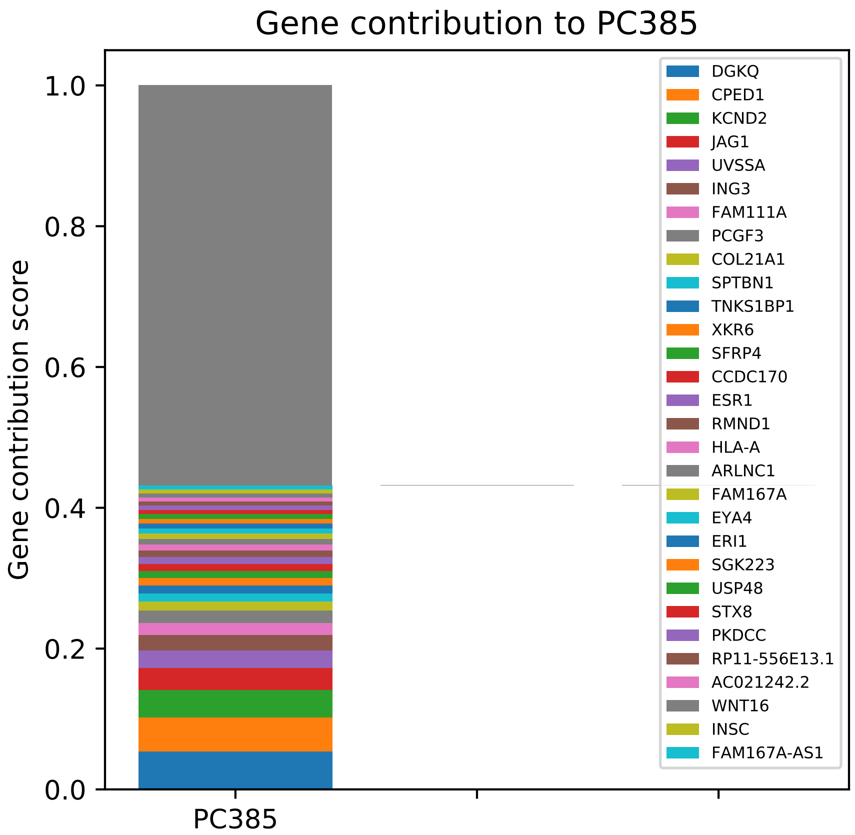
Gene contribution to PC385
Gene contribution score
1.0
0.8
0.6
0.4
0.2
0.0
PC385
DGKQ
CPED1
KCND2
JAG1
UVSSA
ING3
FAM111A
PCGF3
COL21A1
SPTBN1
TNKS1BP1
XKR6
SFRP4
CCDC170
ESR1
RMND1
HLA-A
ARLNC1
FAM167A
EYA4
ERI1
SGK223
USP48
STX8
PKDCC
RP11-556E13.1
AC021242.2
WNT16
INSC
FAM167A-AS1

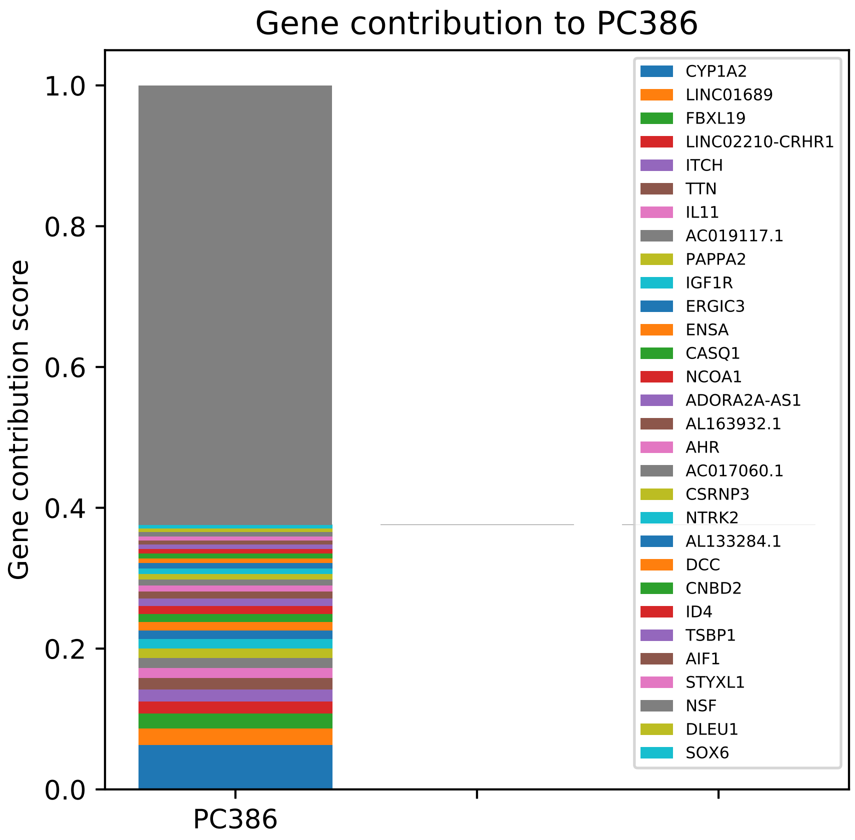
Gene contribution to PC386
Gene contribution score
1.0
0.8
0.6
0.4
0.2
0.0
PC386
CYP1A2
LINC01689
FBXL19
LINC02210-CRHR1
ITCH
TTN
IL11
AC019117.1
PAPPA2
IGF1R
ERGIC3
ENSA
CASQ1
NCOA1
ADORA2A-AS1
AL163932.1
AHR
AC017060.1
CSRNP3
NTRK2
AL133284.1
DCC
CNBD2
ID4
TSBP1
AIF1
STYXL1
NSF
DLEU1
SOX6

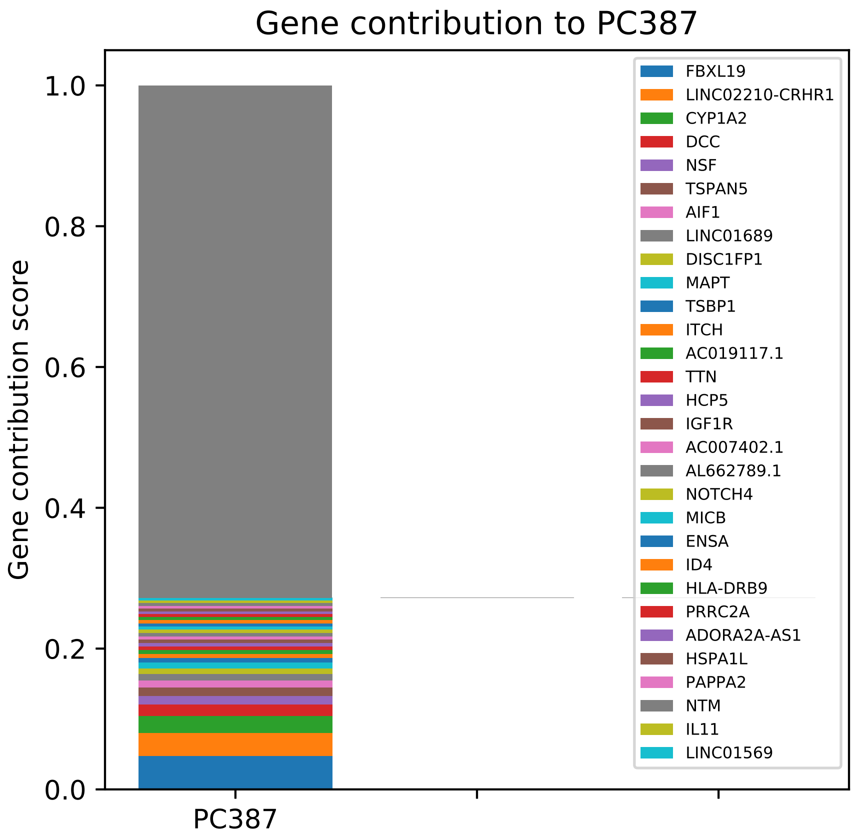

Gene contribution to PC387
Gene contribution score
1.0
0.8
0.6
0.4
0.2
0.0
PC387
FBXL19
LINC02210-CRHR1
CYP1A2
DCC
NSF
TSPAN5
AIF1
LINC01689
DISC1FP1
MAPT
TSBP1
ITCH
AC019117.1
TTN
HCP5
IGF1R
AC007402.1
AL662789.1
NOTCH4
MICB
ENSA
ID4
HLA-DRB9
PRRC2A
ADORA2A-AS1
HSPA1L
PAPPA2
NTM
IL11
LINC01569

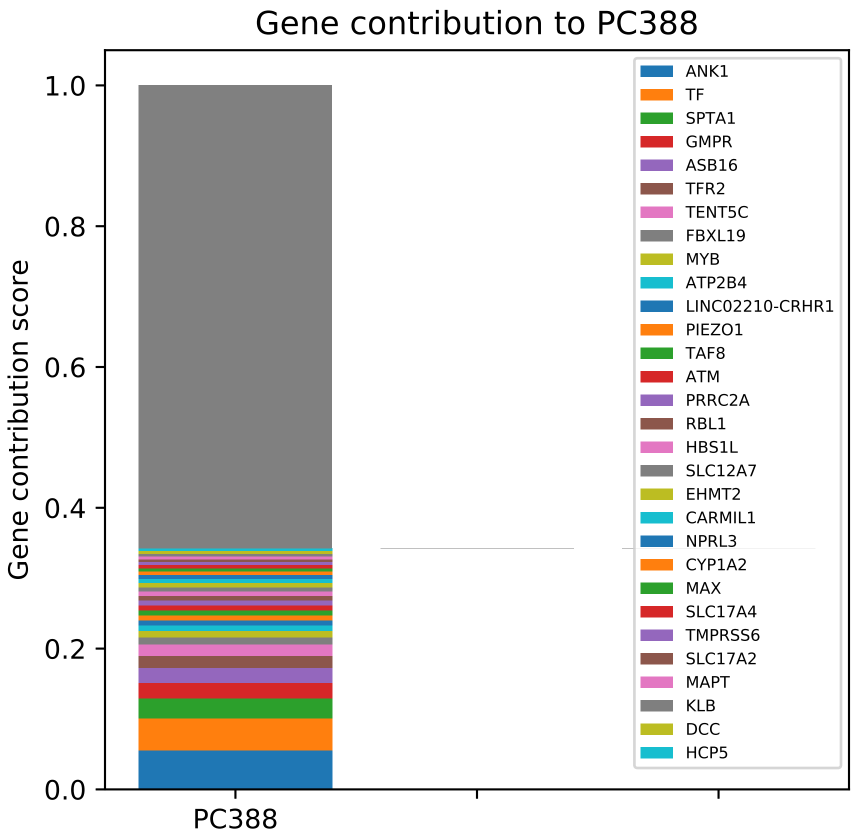

Gene contribution to PC388
Gene contribution score
1.0
0.8
0.6
0.4
0.2
0.0
PC388
ANK1
TF
SPTA1
GMPR
ASB16
TFR2
TENT5C
FBXL19
MYB
ATP2B4
LINC02210-CRHR1
PIEZO1
TAF8
ATM
PRRC2A
RBL1
HBS1L
SLC12A7
EHMT2
CARMIL1
NPRL3
CYP1A2
MAX
SLC17A4
TMPRSS6
SLC17A2
MAPT
KLB
DCC
HCP5

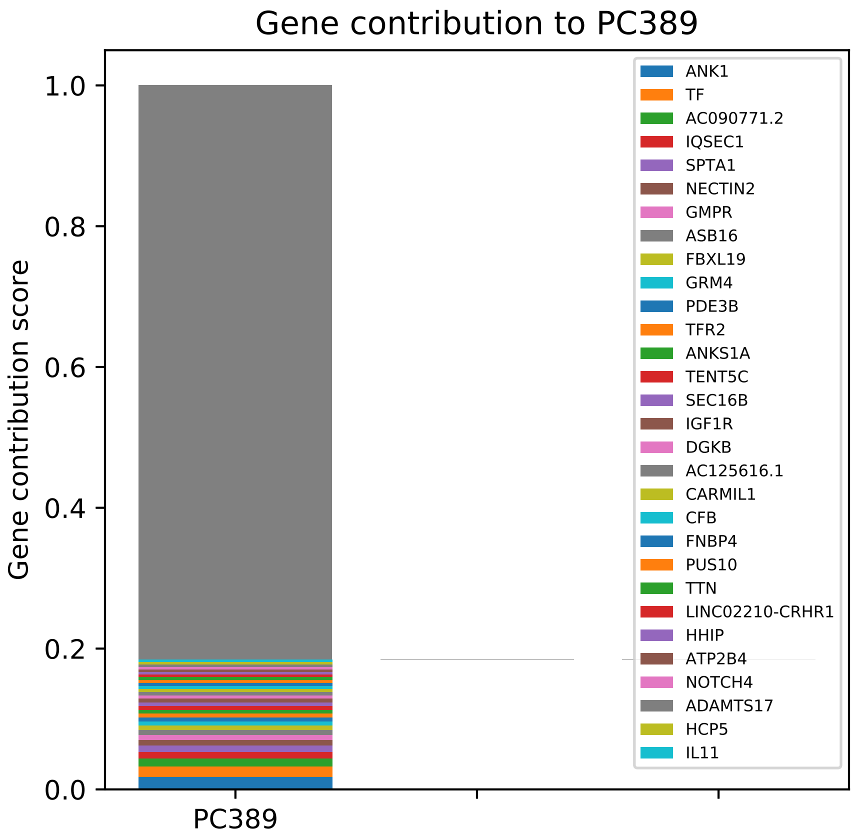
Gene contribution to PC389
Gene contribution score
1.0
0.8
0.6
0.4
0.2
0.0
PC389
ANK1
TF
AC090771.2
IQSEC1
SPTA1
NECTIN2
GMPR
ASB16
FBXL19
GRM4
PDE3B
TFR2
ANKS1A
TENT5C
SEC16B
IGF1R
DGKB
AC125616.1
CARMIL1
CFB
FNBP4
PUS10
TTN
LINC02210-CRHR1
HHIP
ATP2B4
NOTCH4
ADAMTS17
HCP5
IL11

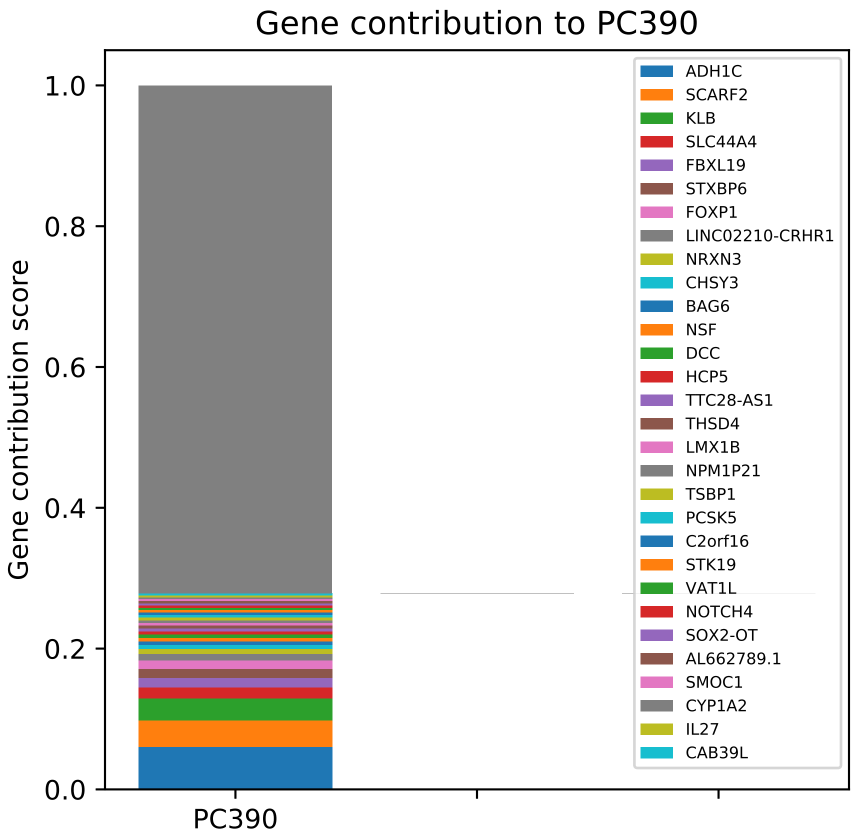
Gene contribution to PC390
Gene contribution score
1.0
0.8
0.6
0.4
0.2
0.0
PC390
ADH1C
SCARF2
KLB
SLC44A4
FBXL19
STXBP6
FOXP1
LINC02210-CRHR1
NRXN3
CHSY3
BAG6
NSF
DCC
HCP5
TTC28-AS1
THSD4
LMX1B
NPM1P21
TSBP1
PCSK5
C2orf16
STK19
VAT1L
NOTCH4
SOX2-OT
AL662789.1
SMOC1
CYP1A2
IL27
CAB39L

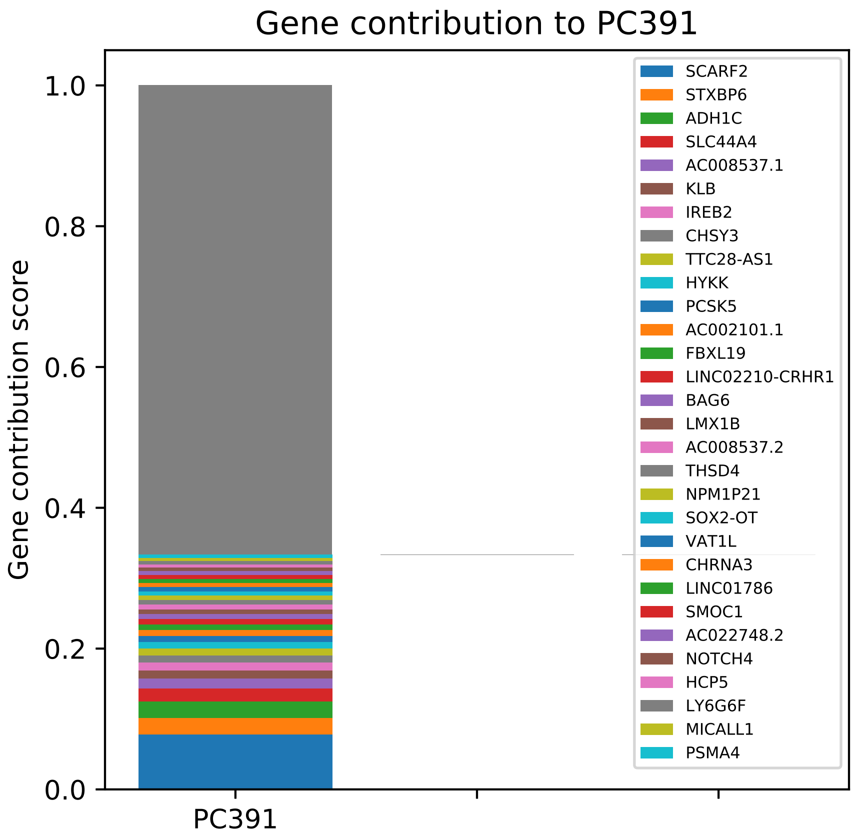

Gene contribution to PC391
Gene contribution score
1.0
0.8
0.6
0.4
0.2
0.0
PC391
SCARF2
STXBP6
ADH1C
SLC44A4
AC008537.1
KLB
IREB2
CHSY3
TTC28-AS1
HYKK
PCSK5
AC002101.1
FBXL19
LINC02210-CRHR1
BAG6
LMX1B
AC008537.2
THSD4
NPM1P21
SOX2-OT
VAT1L
CHRNA3
LINC01786
SMOC1
AC022748.2
NOTCH4
HCP5
LY6G6F
MICALL1
PSMA4

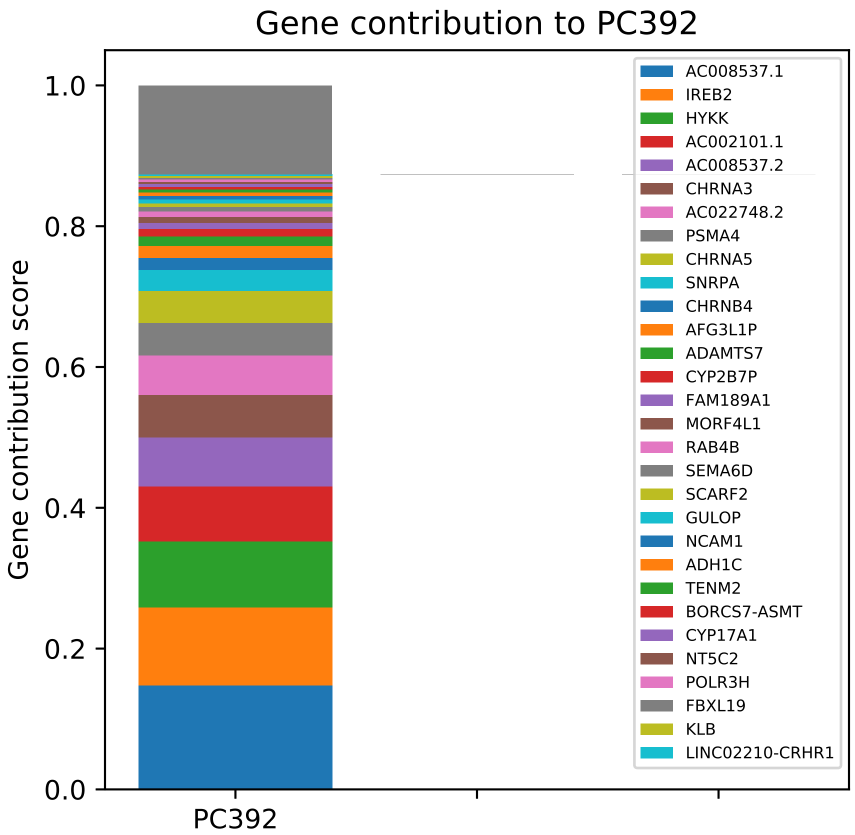

Gene contribution to PC392
Gene contribution score
0.0
0.2
0.4
0.6
0.8
1.0
PC392
AC008537.1
IREB2
HYKK
AC002101.1
AC008537.2
CHRNA3
AC022748.2
PSMA4
CHRNA5
SNRPA
CHRNB4
AFG3L1P
ADAMTS7
CYP2B7P
FAM189A1
MORF4L1
RAB4B
SEMA6D
SCARF2
GULOP
NCAM1
ADH1C
TENM2
BORCS7-ASMT
CYP17A1
NT5C2
POLR3H
FBXL19
KLB
LINC02210-CRHR1

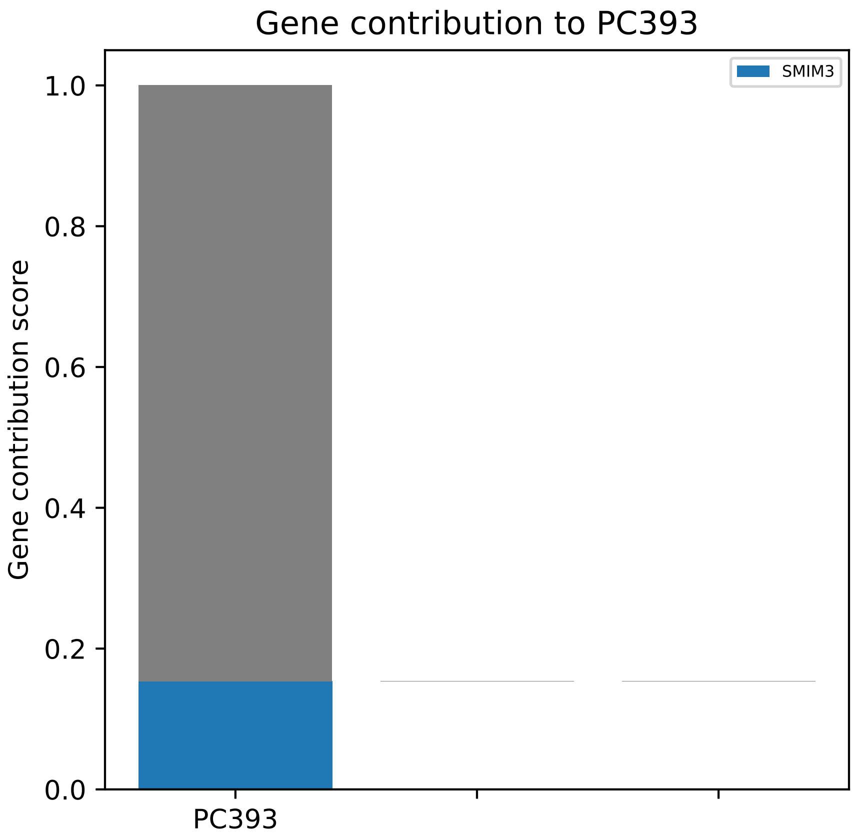

Gene contribution to PC393
Gene contribution score
1.0
0.8
0.6
0.4
0.2
0.0
PC393
SMIM3

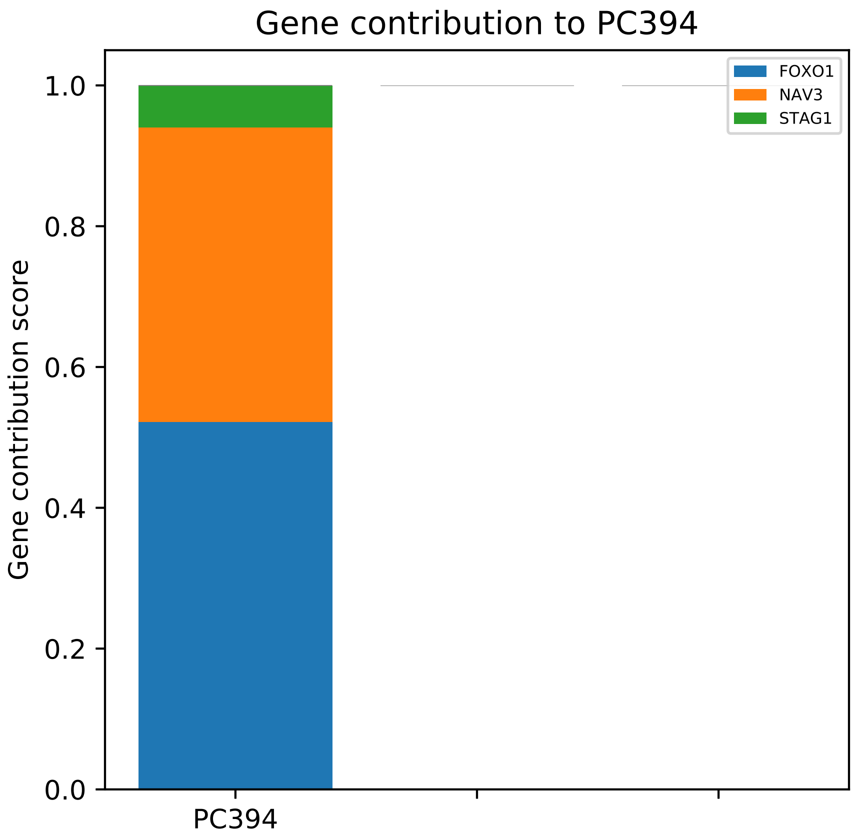

Gene contribution to PC394
1.0
0.8
0.6
0.4
0.2
0.0
Gene contribution score
PC394
FOXO1
NAV3
STAG1

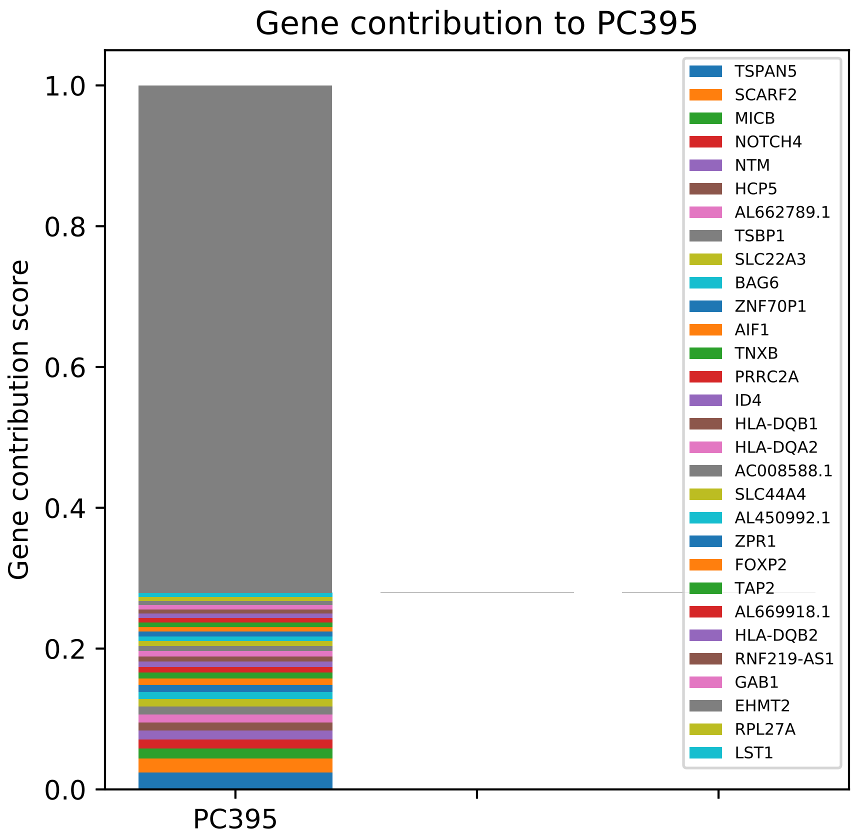

Gene contribution to PC395
Gene contribution score
1.0
0.8
0.6
0.4
0.2
0.0
PC395
TSPAN5
SCARF2
MICB
NOTCH4
NTM
HCP5
AL662789.1
TSBP1
SLC22A3
BAG6
ZNF70P1
AIF1
TNXB
PRRC2A
ID4
HLA-DQB1
HLA-DQA2
AC008588.1
SLC44A4
AL450992.1
ZPR1
FOXP2
TAP2
AL669918.1
HLA-DQB2
RNF219-AS1
GAB1
EHMT2
RPL27A
LST1

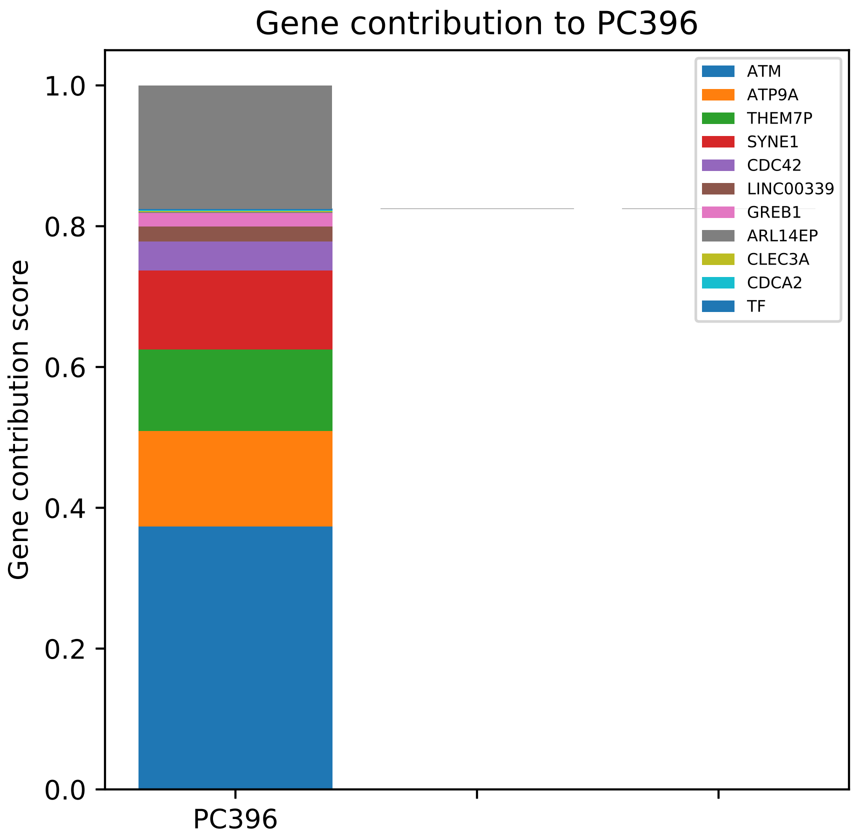

Gene contribution to PC396
Gene contribution score
1.0
0.8
0.6
0.4
0.2
0.0
PC396
ATM
ATP9A
THEM7P
SYNE1
CDC42
LINC00339
GREB1
ARL14EP
CLEC3A
CDCA2
TF

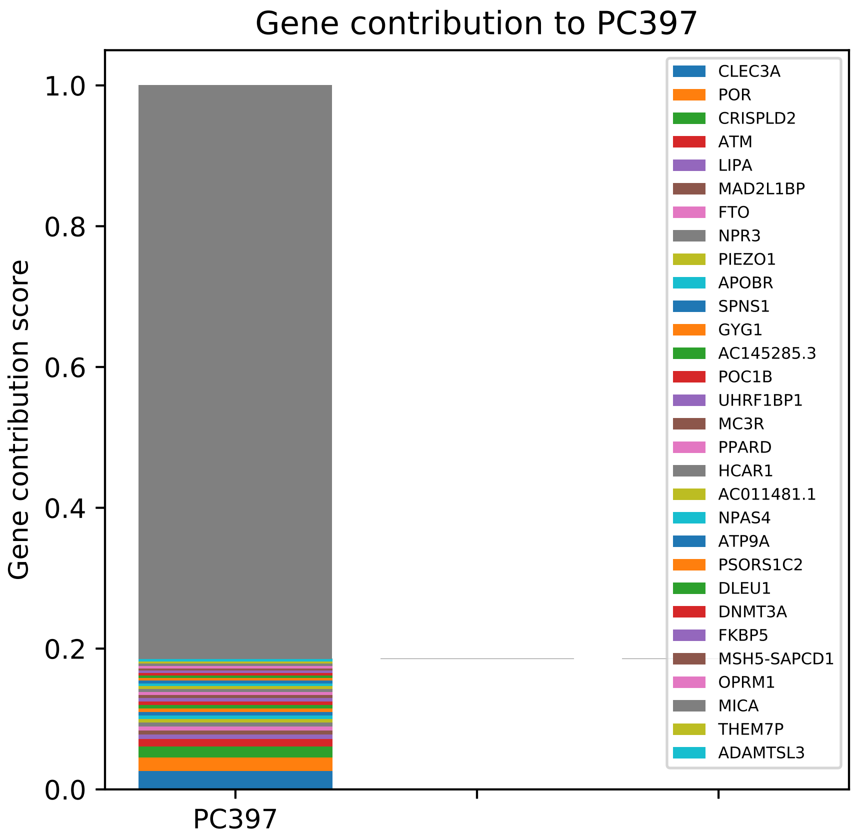
Gene contribution to PC397
Gene contribution score
1.0
0.8
0.6
0.4
0.2
0.0
PC397
CLEC3A
POR
CRISPLD2
ATM
LIPA
MAD2L1BP
FTO
NPR3
PIEZO1
APOBR
SPNS1
GYG1
AC145285.3
POC1B
UHRF1BP1
MC3R
PPARD
HCAR1
AC011481.1
NPAS4
ATP9A
PSORS1C2
DLEU1
DNMT3A
FKBP5
MSH5-SAPCD1
OPRM1
MICA
THEM7P
ADAMTSL3

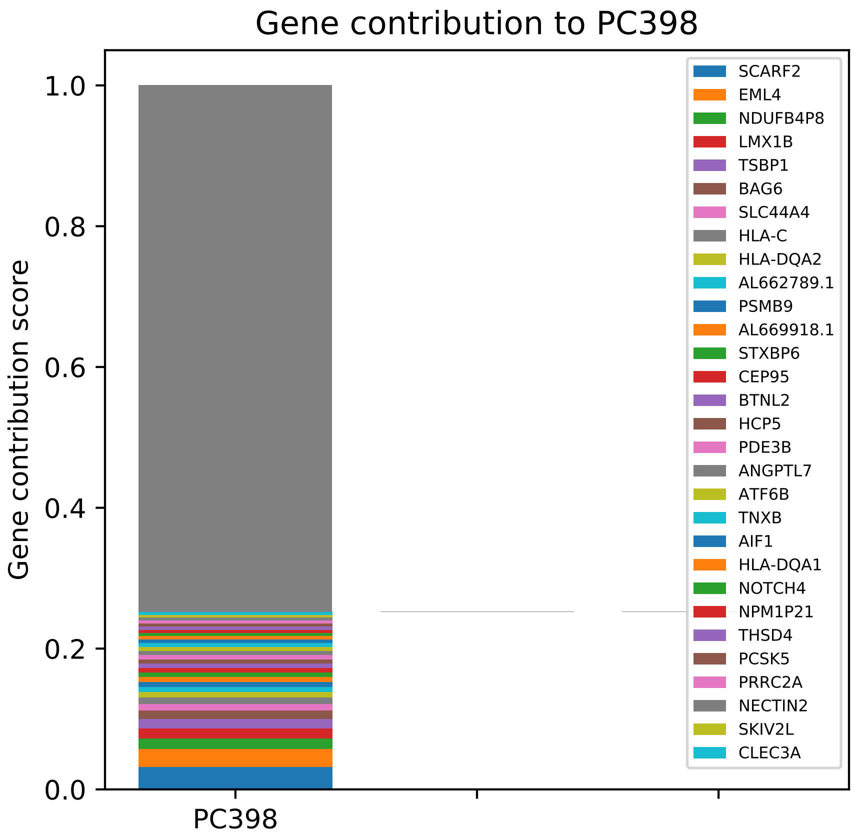
Gene contribution to PC398
Gene contribution score
1.0
0.8
0.6
0.4
0.2
0.0
PC398
SCARF2
EML4
NDUFB4P8
LMX1B
TSBP1
BAG6
SLC44A4
HLA-C
HLA-DQA2
AL662789.1
PSMB9
AL669918.1
STXBP6
CEP95
BTNL2
HCP5
PDE3B
ANGPTL7
ATF6B
TNXB
AIF1
HLA-DQA1
NOTCH4
NPM1P21
THSD4
PCSK5
PRRC2A
NECTIN2
SKIV2L
CLEC3A

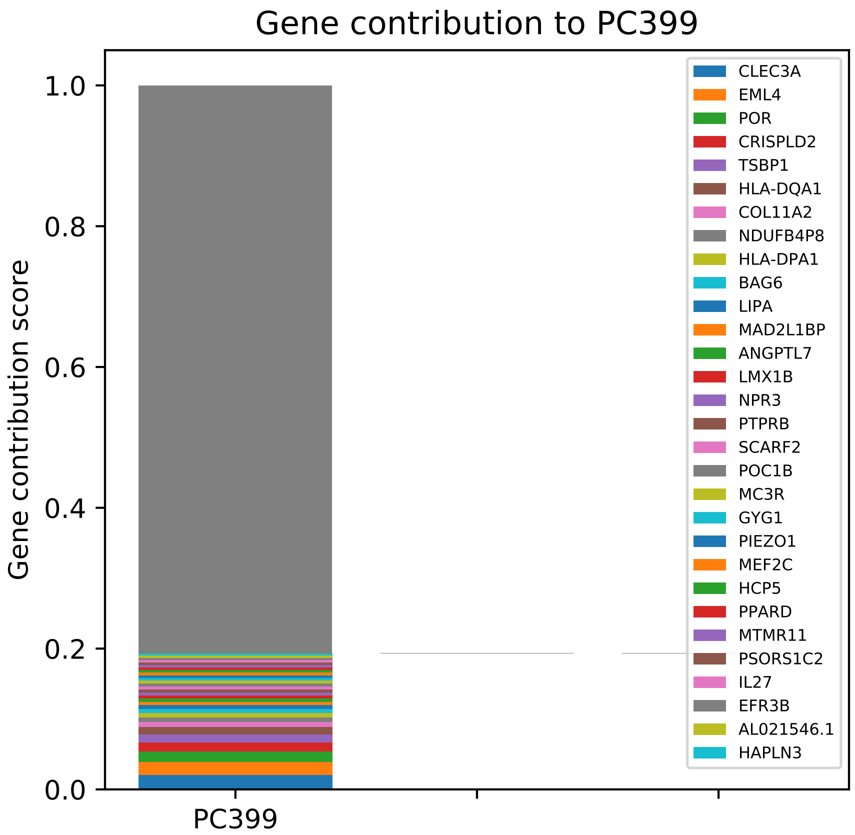
Gene contribution to PC399
Gene contribution score
1.0
0.8
0.6
0.4
0.2
0.0
PC399
CLEC3A
EML4
POR
CRISPLD2
TSBP1
HLA-DQA1
COL11A2
NDUFB4P8
HLA-DPA1
BAG6
LIPA
MAD2L1BP
ANGPTL7
LMX1B
NPR3
PTPRB
SCARF2
POC1B
MC3R
GYG1
PIEZO1
MEF2C
HCP5
PPARD
MTMR11
PSORS1C2
IL27
EFR3B
AL021546.1
HAPLN3

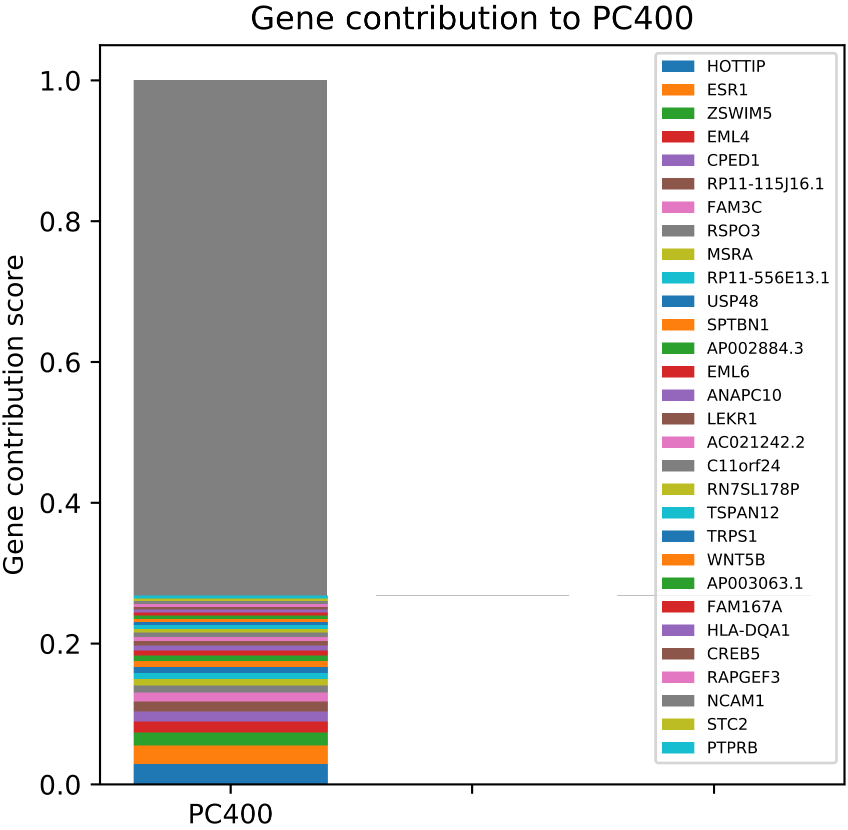
Gene contribution to PC400
Gene contribution score
1.0
0.8
0.6
0.4
0.2
0.0
PC400
HOTTIP
ESR1
ZSWIM5
EML4
CPED1
RP11-115J16.1
FAM3C
RSPO3
MSRA
RP11-556E13.1
USP48
SPTBN1
AP002884.3
EML6
ANAPC10
LEKR1
AC021242.2
C11orf24
RN7SL178P
TSPAN12
TRPS1
WNT5B
AP003063.1
FAM167A
HLA-DQA1
CREB5
RAPGEF3
NCAM1
STC2
PTPRB

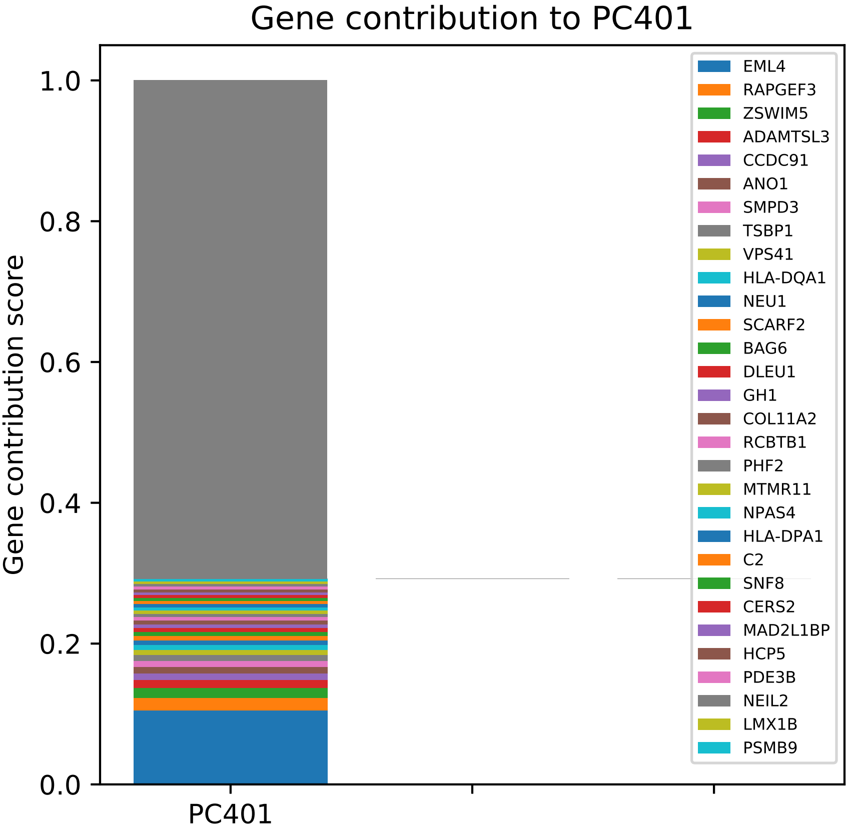

Gene contribution to PC401
Gene contribution score
1.0
0.8
0.6
0.4
0.2
0.0
PC401
EML4
RAPGEF3
ZSWIM5
ADAMTSL3
CCDC91
ANO1
SMPD3
TSBP1
VPS41
HLA-DQA1
NEU1
SCARF2
BAG6
DLEU1
GH1
COL11A2
RCBTB1
PHF2
MTMR11
NPAS4
HLA-DPA1
C2
SNF8
CERS2
MAD2L1BP
HCP5
PDE3B
NEIL2
LMX1B
PSMB9

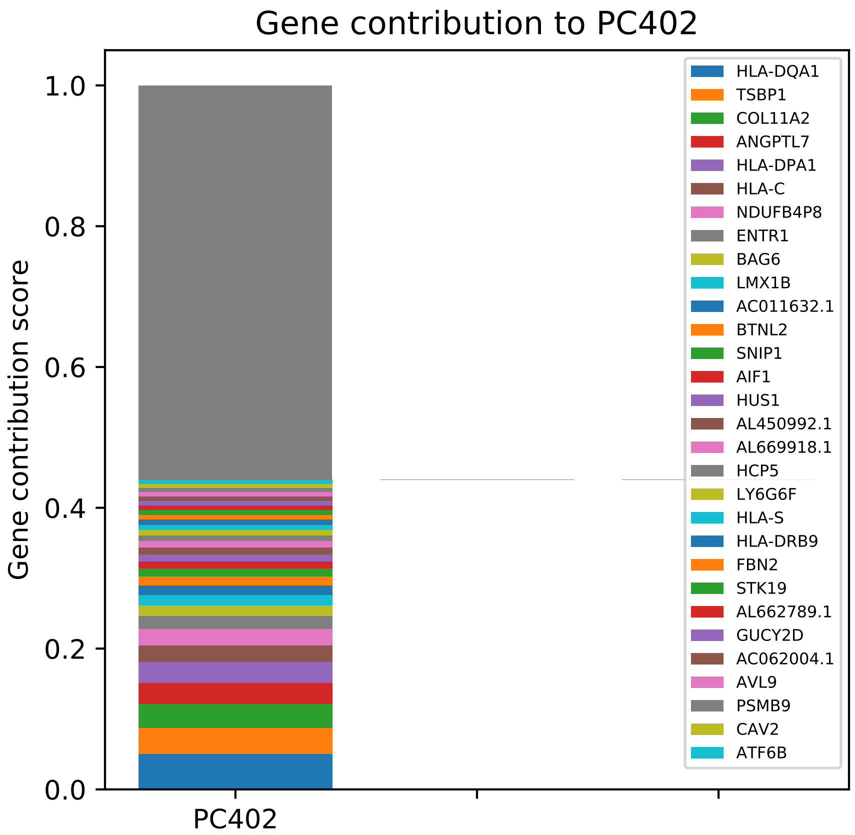
Gene contribution to PC402
Gene contribution score
1.0
0.8
0.6
0.4
0.2
0.0
PC402
HLA-DQA1
TSBP1
COL11A2
ANGPTL7
HLA-DPA1
HLA-C
NDUFB4P8
ENTR1
BAG6
LMX1B
AC011632.1
BTNL2
SNIP1
AIF1
HUS1
AL450992.1
AL669918.1
HCP5
LY6G6F
HLA-S
HLA-DRB9
FBN2
STK19
AL662789.1
GUCY2D
AC062004.1
AVL9
PSMB9
CAV2
ATF6B

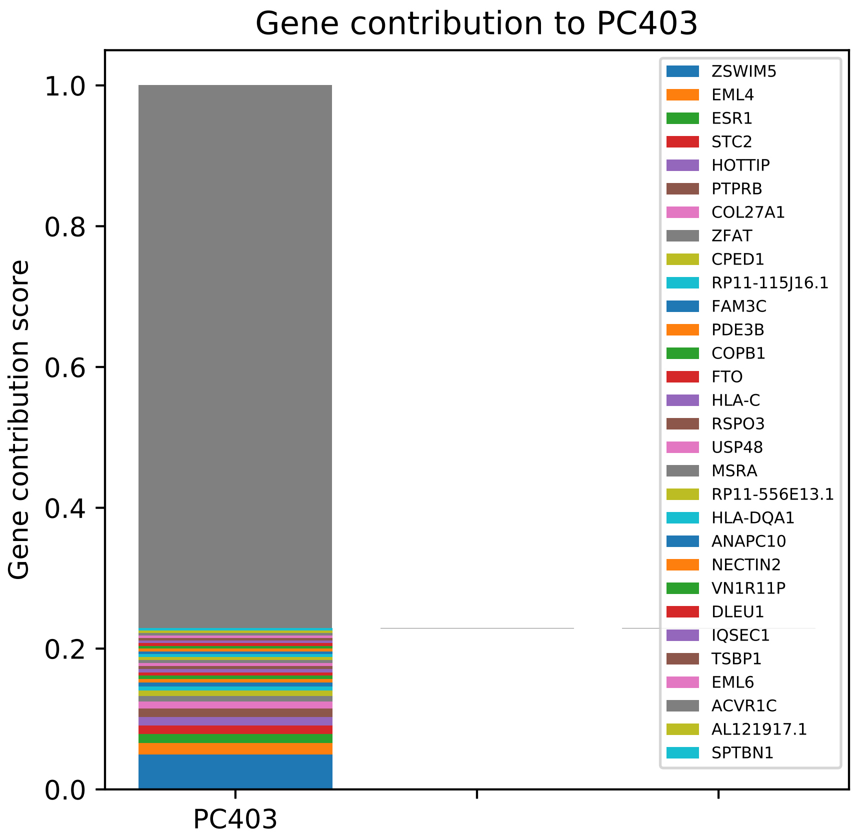

Gene contribution to PC403
Gene contribution score
1.0
0.8
0.6
0.4
0.2
0.0
PC403
ZSWIM5
EML4
ESR1
STC2
HOTTIP
PTPRB
COL27A1
ZFAT
CPED1
RP11-115J16.1
FAM3C
PDE3B
COPB1
FTO
HLA-C
RSPO3
USP48
MSRA
RP11-556E13.1
HLA-DQA1
ANAPC10
NECTIN2
VN1R11P
DLEU1
IQSEC1
TSBP1
EML6
ACVR1C
AL121917.1
SPTBN1

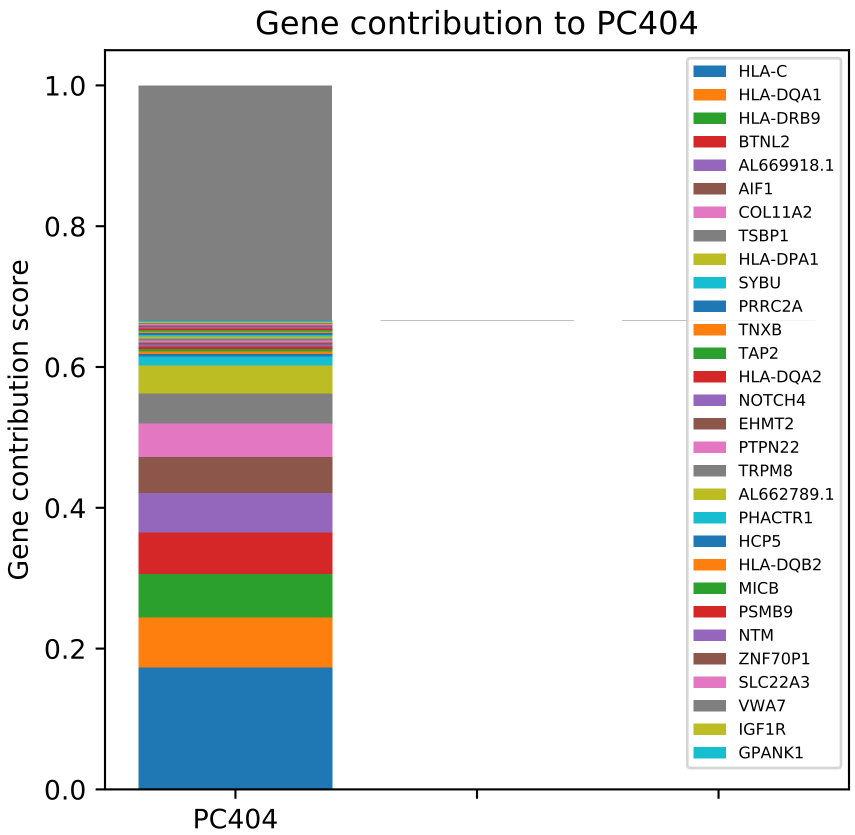

Gene contribution to PC404
Gene contribution score
0.0
0.2
0.4
0.6
0.8
1.0
PC404
HLA-C
HLA-DQA1
HLA-DRB9
BTNL2
AL669918.1
AIF1
COL11A2
TSBP1
HLA-DPA1
SYBU
PRRC2A
TNXB
TAP2
HLA-DQA2
NOTCH4
EHMT2
PTPN22
TRPM8
AL662789.1
PHACTR1
HCP5
HLA-DQB2
MICB
PSMB9
NTM
ZNF70P1
SLC22A3
VWA7
IGF1R
GPANK1

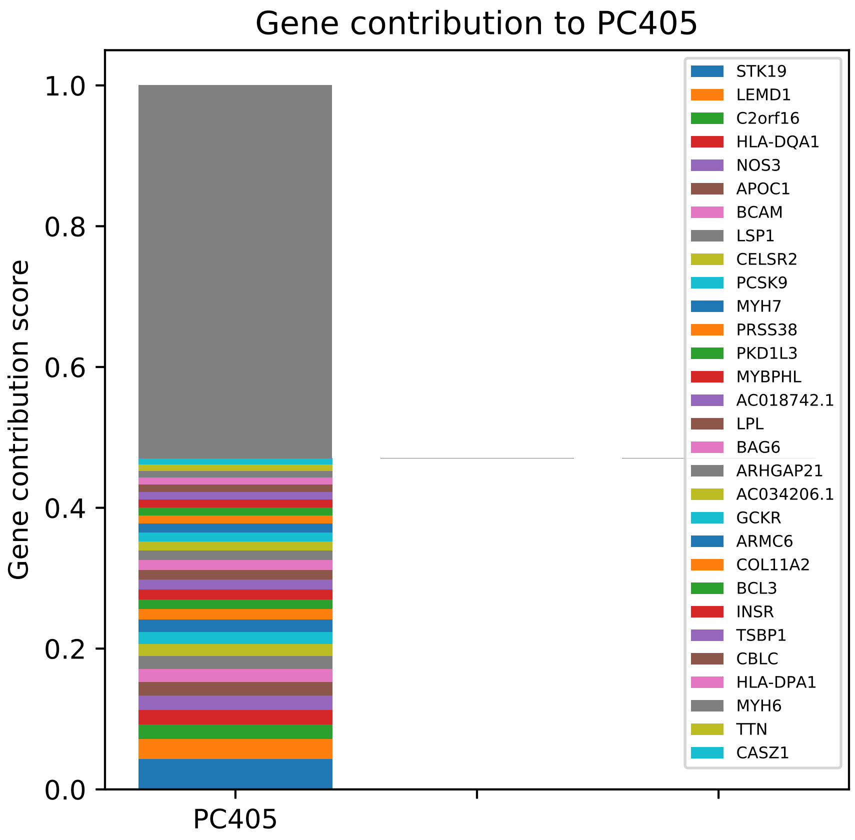

Gene contribution to PC405
Gene contribution score
1.0
0.8
0.6
0.4
0.2
0.0
PC405
STK19
LEMD1
C2orf16
HLA-DQA1
NOS3
APOC1
BCAM
LSP1
CELSR2
PCSK9
MYH7
PRSS38
PKD1L3
MYBPHL
AC018742.1
LPL
BAG6
ARHGAP21
AC034206.1
GCKR
ARMC6
COL11A2
BCL3
INSR
TSBP1
CBLC
HLA-DPA1
MYH6
TTN
CASZ1

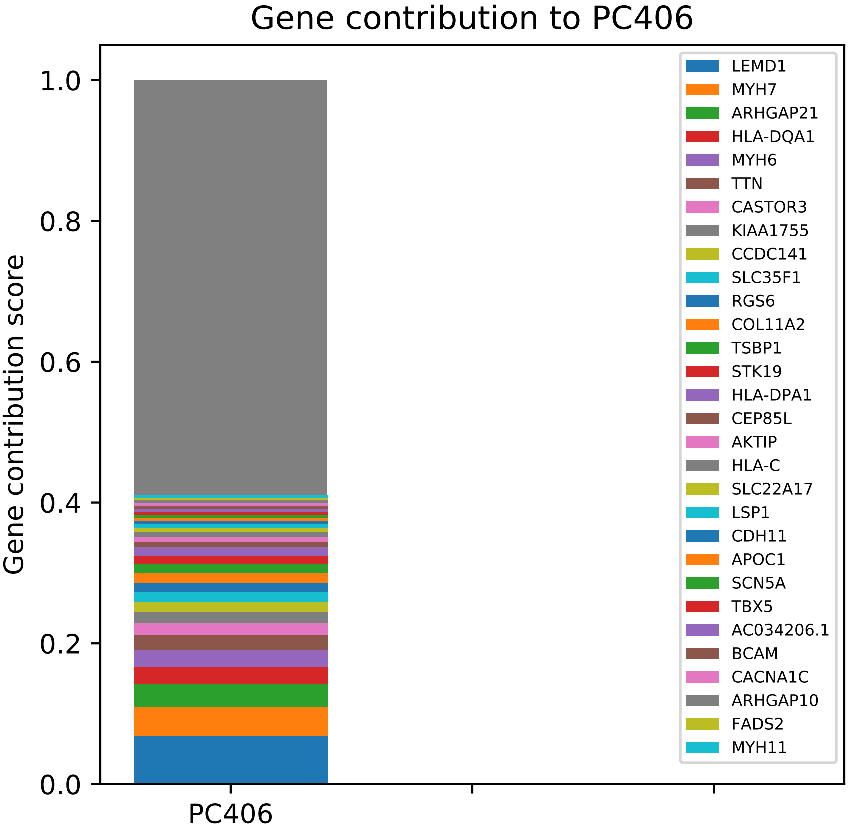
Gene contribution to PC406
Gene contribution score
1.0
0.8
0.6
0.4
0.2
0.0
PC406
LEMD1
MYH7
ARHGAP21
HLA-DQA1
MYH6
TTN
CASTOR3
KIAA1755
CCDC141
SLC35F1
RGS6
COL11A2
TSBP1
STK19
HLA-DPA1
CEP85L
AKTIP
HLA-C
SLC22A17
LSP1
CDH11
APOC1
SCN5A
TBX5
AC034206.1
BCAM
CACNA1C
ARHGAP10
FADS2
MYH11

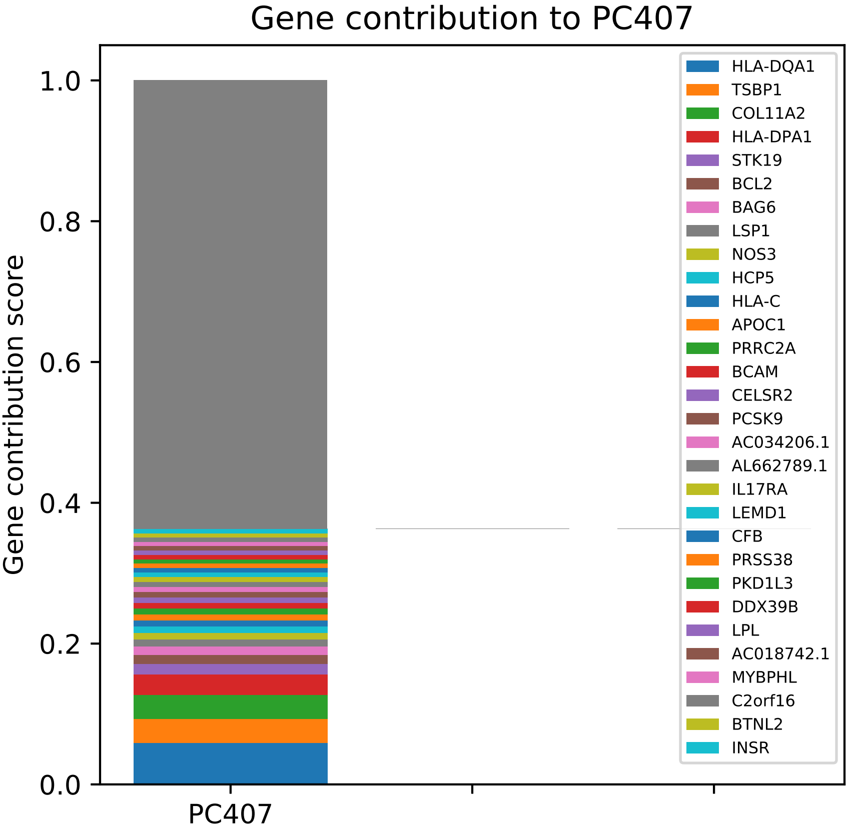
Gene contribution to PC407
Gene contribution score
1.0
0.8
0.6
0.4
0.2
0.0
PC407
HLA-DQA1
TSBP1
COL11A2
HLA-DPA1
STK19
BCL2
BAG6
LSP1
NOS3
HCP5
HLA-C
APOC1
PRRC2A
BCAM
CELSR2
PCSK9
AC034206.1
AL662789.1
IL17RA
LEMD1
CFB
PRSS38
PKD1L3
DDX39B
LPL
AC018742.1
MYBPHL
C2orf16
BTNL2
INSR

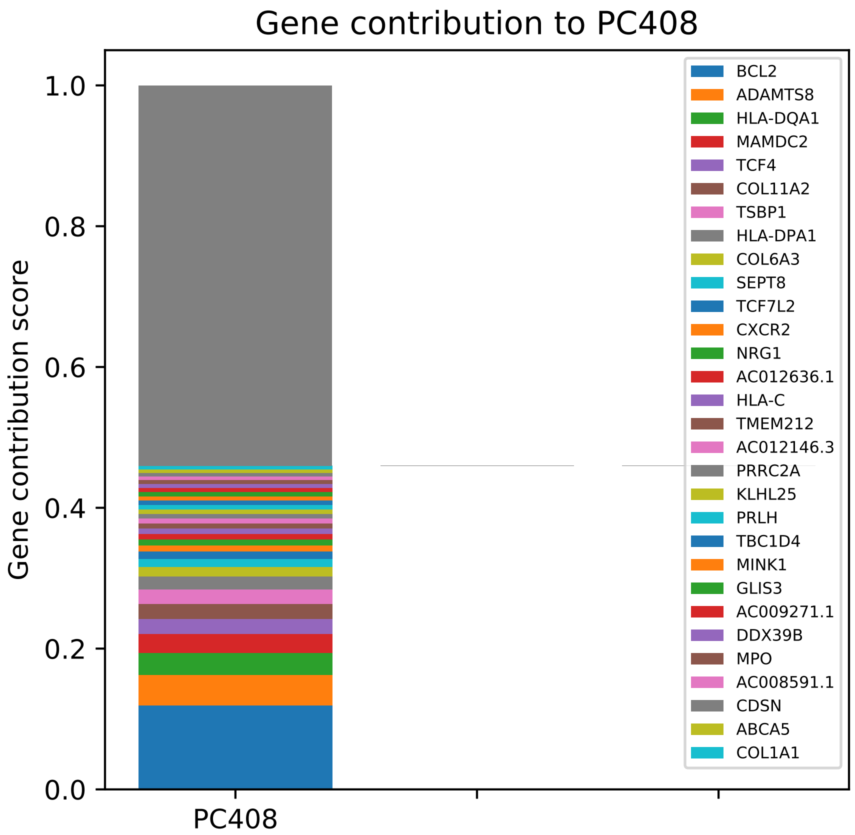

Gene contribution to PC408
Gene contribution score
1.0
0.8
0.6
0.4
0.2
0.0
PC408
BCL2
ADAMTS8
HLA-DQA1
MAMDC2
TCF4
COL11A2
TSBP1
HLA-DPA1
COL6A3
SEPT8
TCF7L2
CXCR2
NRG1
AC012636.1
HLA-C
TMEM212
AC012146.3
PRRC2A
KLHL25
PRLH
TBC1D4
MINK1
GLIS3
AC009271.1
DDX39B
MPO
AC008591.1
CDSN
ABCA5
COL1A1

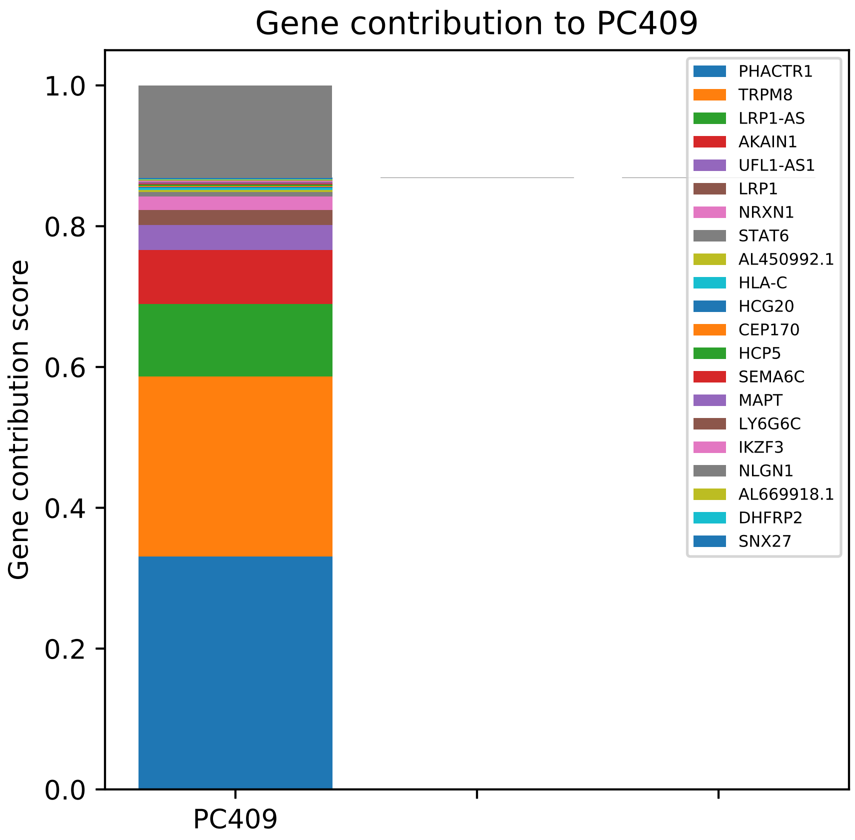

Gene contribution to PC409
Gene contribution score
1.0
0.8
0.6
0.4
0.2
0.0
PC409
PHACTR1
TRPM8
LRP1-AS
AKAIN1
UFL1-AS1
LRP1
NRXN1
STAT6
AL450992.1
HLA-C
HCG20
CEP170
HCP5
SEMA6C
MAPT
LY6G6C
IKZF3
NLGN1
AL669918.1
DHFRP2
SNX27

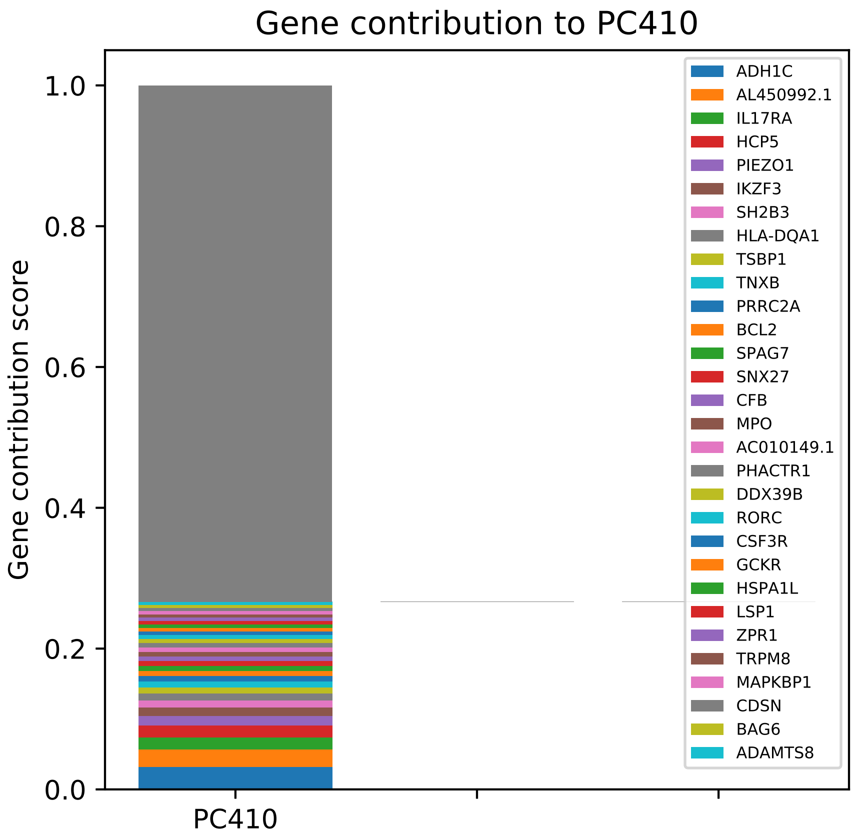
Gene contribution to PC410
Gene contribution score
1.0
0.8
0.6
0.4
0.2
0.0
PC410
ADH1C
AL450992.1
IL17RA
HCP5
PIEZO1
IKZF3
SH2B3
HLA-DQA1
TSBP1
TNXB
PRRC2A
BCL2
SPAG7
SNX27
CFB
MPO
AC010149.1
PHACTR1
DDX39B
RORC
CSF3R
GCKR
HSPA1L
LSP1
ZPR1
TRPM8
MAPKBP1
CDSN
BAG6
ADAMTS8

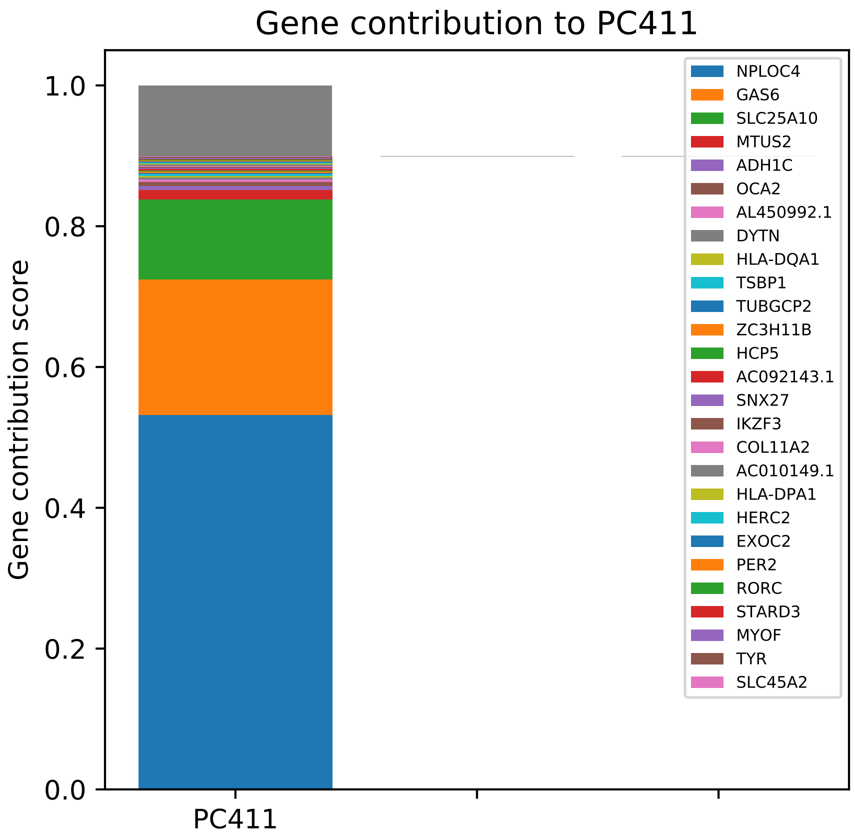
Gene contribution to PC411
Gene contribution score
1.0
0.8
0.6
0.4
0.2
0.0
PC411
NPLOC4
GAS6
SLC25A10
MTUS2
ADH1C
OCA2
AL450992.1
DYTN
HLA-DQA1
TSBP1
TUBGCP2
ZC3H11B
HCP5
AC092143.1
SNX27
IKZF3
COL11A2
AC010149.1
HLA-DPA1
HERC2
EXOC2
PER2
RORC
STARD3
MYOF
TYR
SLC45A2

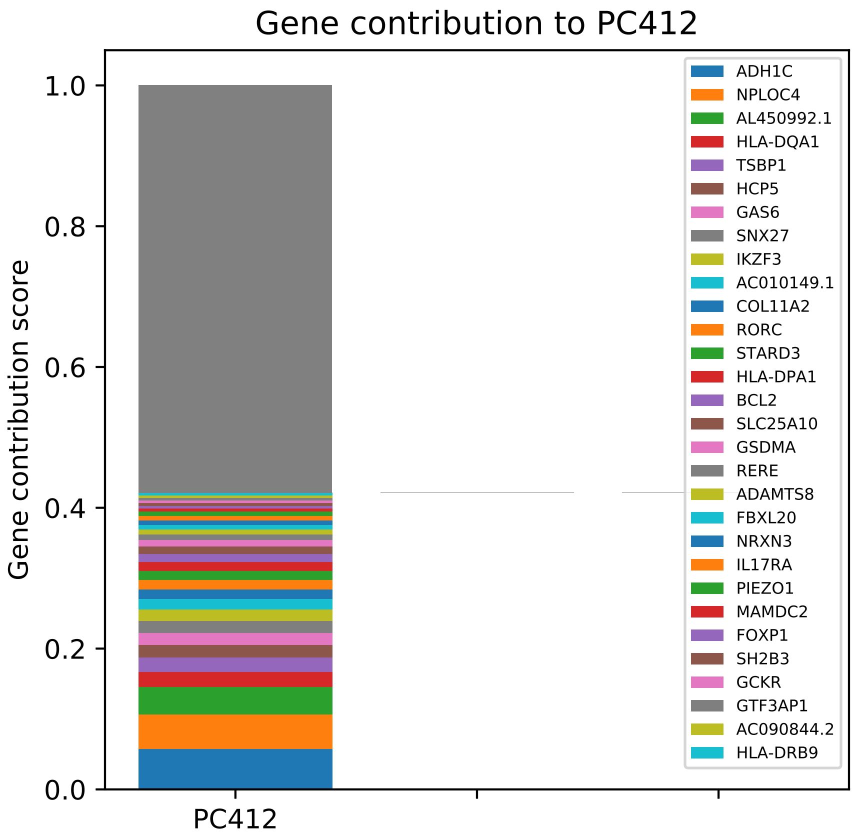
Gene contribution to PC412
Gene contribution score
1.0
0.8
0.6
0.4
0.2
0.0
PC412
ADH1C
NPLOC4
AL450992.1
HLA-DQA1
TSBP1
HCP5
GAS6
SNX27
IKZF3
AC010149.1
COL11A2
RORC
STARD3
HLA-DPA1
BCL2
SLC25A10
GSDMA
RERE
ADAMTS8
FBXL20
NRXN3
IL17RA
PIEZO1
MAMDC2
FOXP1
SH2B3
GCKR
GTF3AP1
AC090844.2
HLA-DRB9

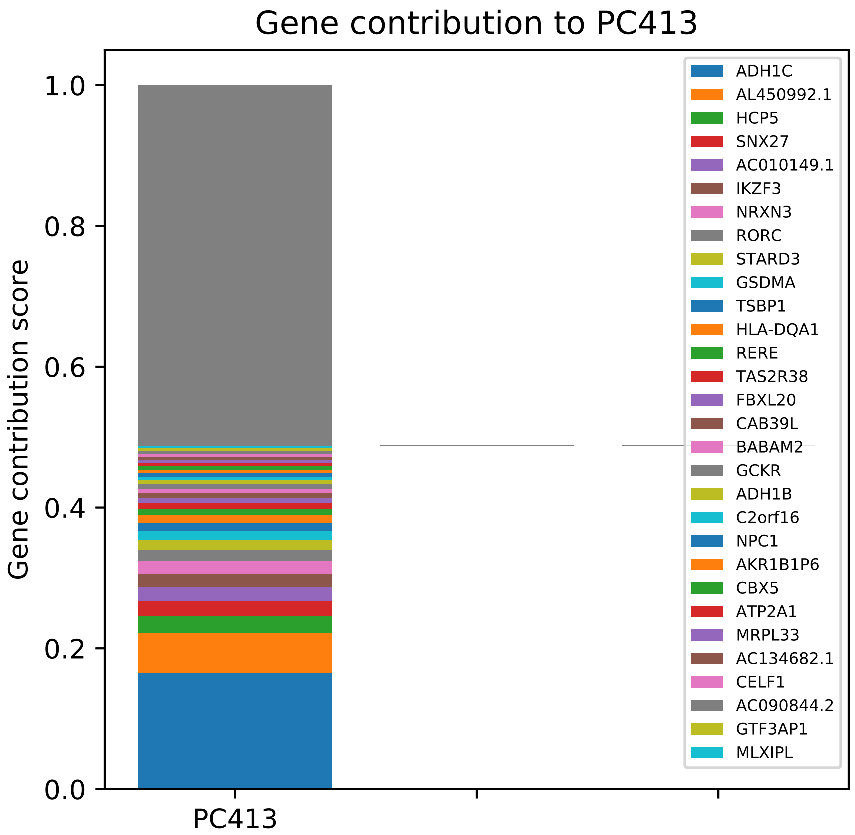

Gene contribution to PC413
Gene contribution score
1.0
0.8
0.6
0.4
0.2
0.0
PC413
ADH1C
AL450992.1
HCP5
SNX27
AC010149.1
IKZF3
NRXN3
RORC
STARD3
GSDMA
TSBP1
HLA-DQA1
RERE
TAS2R38
FBXL20
CAB39L
BABAM2
GCKR
ADH1B
C2orf16
NPC1
AKR1B1P6
CBX5
ATP2A1
MRPL33
AC134682.1
CELF1
AC090844.2
GTF3AP1
MLXIPL

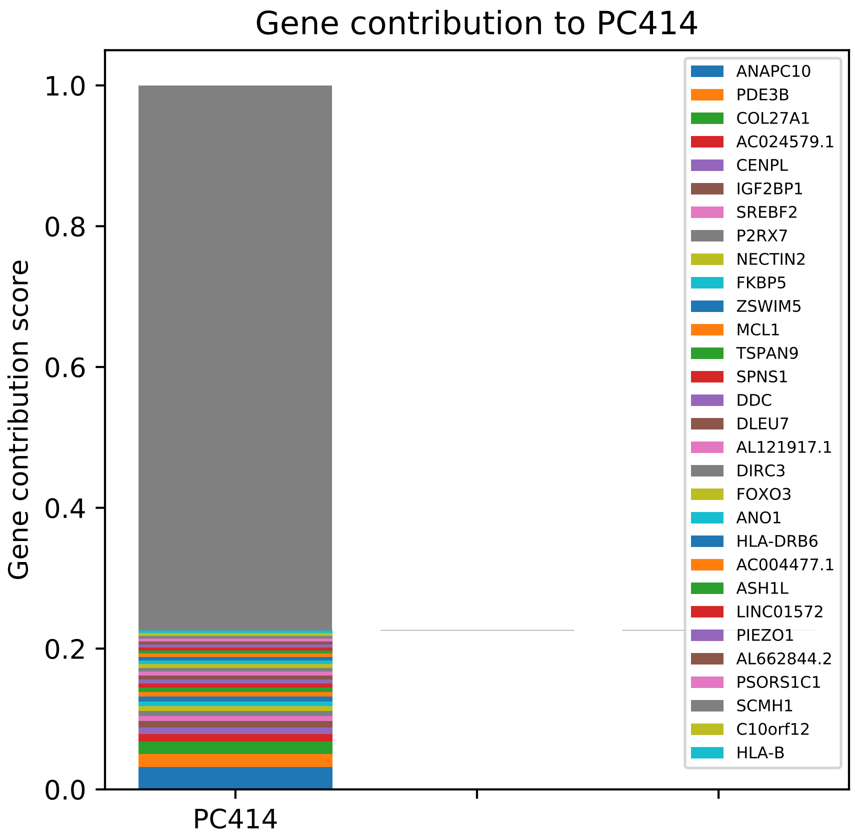

Gene contribution to PC414
Gene contribution score
1.0
0.8
0.6
0.4
0.2
0.0
PC414
ANAPC10
PDE3B
COL27A1
AC024579.1
CENPL
IGF2BP1
SREBF2
P2RX7
NECTIN2
FKBP5
ZSWIM5
MCL1
TSPAN9
SPNS1
DDC
DLEU7
AL121917.1
DIRC3
FOXO3
ANO1
HLA-DRB6
AC004477.1
ASH1L
LINC01572
PIEZO1
AL662844.2
PSORS1C1
SCMH1
C10orf12
HLA-B

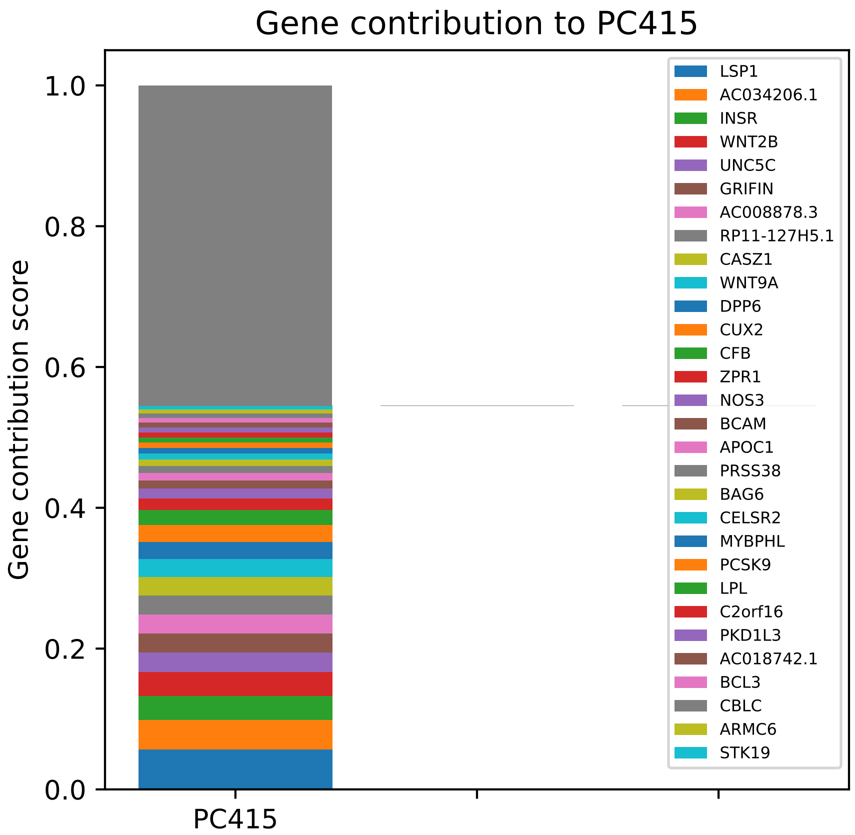

Gene contribution to PC415
Gene contribution score
1.0
0.8
0.6
0.4
0.2
0.0
PC415
LSP1
AC034206.1
INSR
WNT2B
UNC5C
GRIFIN
AC008878.3
RP11-127H5.1
CASZ1
WNT9A
DPP6
CUX2
CFB
ZPR1
NOS3
BCAM
APOC1
PRSS38
BAG6
CELSR2
MYBPHL
PCSK9
LPL
C2orf16
PKD1L3
AC018742.1
BCL3
CBLC
ARMC6
STK19

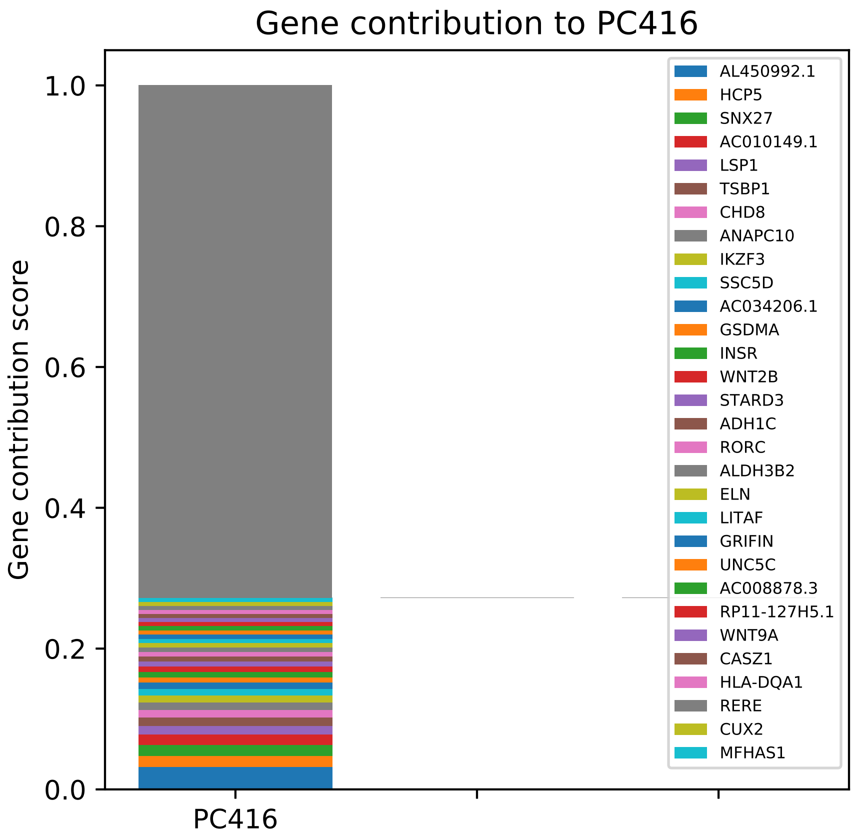

Gene contribution to PC416
Gene contribution score
1.0
0.8
0.6
0.4
0.2
0.0
PC416
AL450992.1
HCP5
SNX27
AC010149.1
LSP1
TSBP1
CHD8
ANAPC10
IKZF3
SSC5D
AC034206.1
GSDMA
INSR
WNT2B
STARD3
ADH1C
RORC
ALDH3B2
ELN
LITAF
GRIFIN
UNC5C
AC008878.3
RP11-127H5.1
WNT9A
CASZ1
HLA-DQA1
RERE
CUX2
MFHAS1

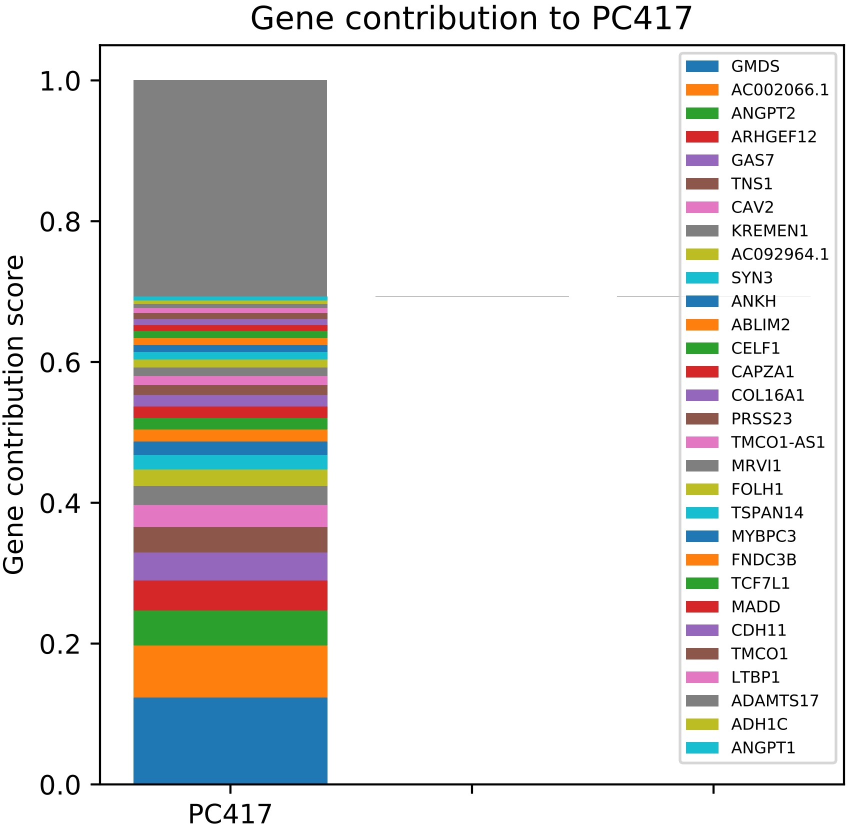
Gene contribution to PC417
Gene contribution score
1.0
0.8
0.6
0.4
0.2
0.0
PC417
GMDS
AC002066.1
ANGPT2
ARHGEF12
GAS7
TNS1
CAV2
KREMEN1
AC092964.1
SYN3
ANKH
ABLIM2
CELF1
CAPZA1
COL16A1
PRSS23
TMCO1-AS1
MRVI1
FOLH1
TSPAN14
MYBPC3
FNDC3B
TCF7L1
MADD
CDH11
TMCO1
LTBP1
ADAMTS17
ADH1C
ANGPT1

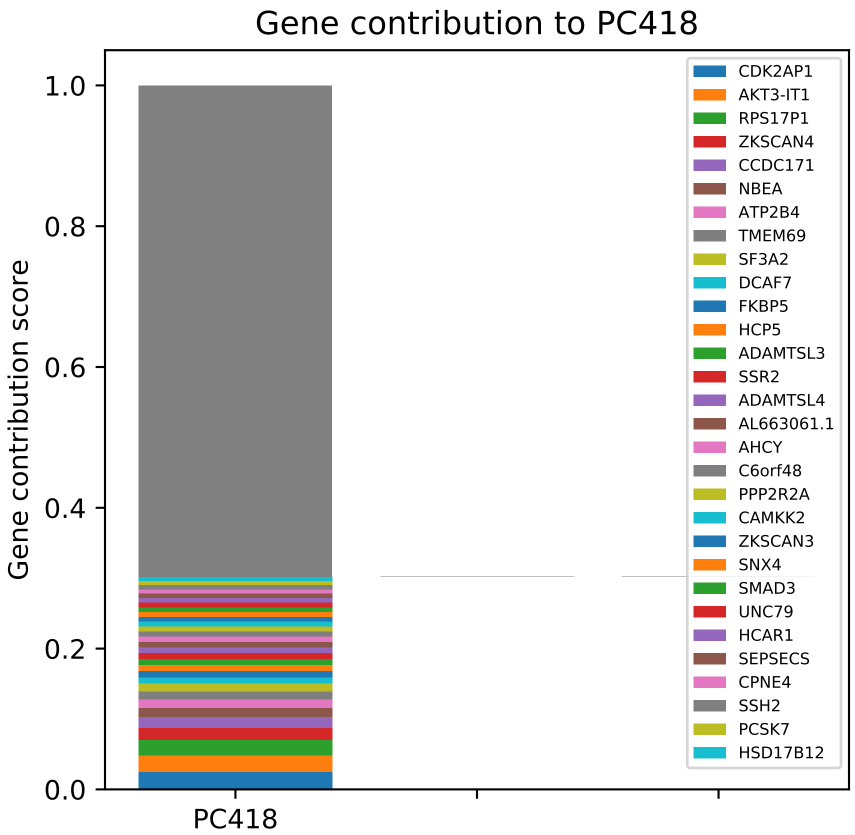
Gene contribution to PC418
Gene contribution score
1.0
0.8
0.6
0.4
0.2
0.0
PC418
CDK2AP1
AKT3-IT1
RPS17P1
ZKSCAN4
CCDC171
NBEA
ATP2B4
TMEM69
SF3A2
DCAF7
FKBP5
HCP5
ADAMTSL3
SSR2
ADAMTSL4
AL663061.1
AHCY
C6orf48
PPP2R2A
CAMKK2
ZKSCAN3
SNX4
SMAD3
UNC79
HCAR1
SEPSECS
CPNE4
SSH2
PCSK7
HSD17B12

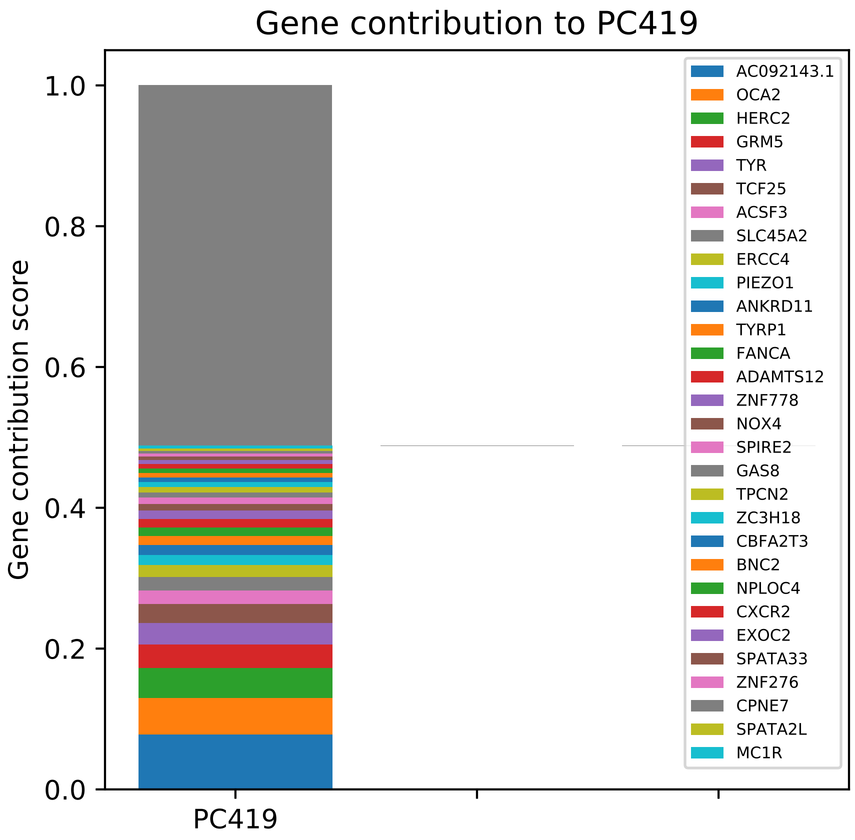
Gene contribution to PC419
Gene contribution score
1.0
0.8
0.6
0.4
0.2
0.0
PC419
AC092143.1
OCA2
HERC2
GRM5
TYR
TCF25
ACSF3
SLC45A2
ERCC4
PIEZO1
ANKRD11
TYRP1
FANCA
ADAMTS12
ZNF778
NOX4
SPIRE2
GAS8
TPCN2
ZC3H18
CBFA2T3
BNC2
NPLOC4
CXCR2
EXOC2
SPATA33
ZNF276
CPNE7
SPATA2L
MC1R

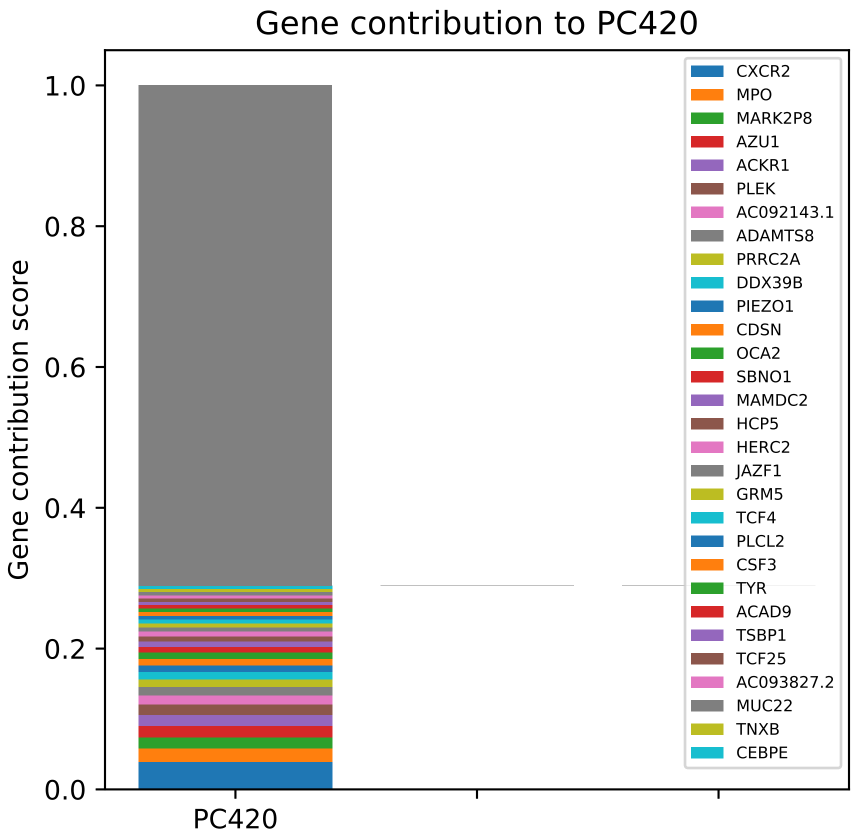
Gene contribution to PC420
Gene contribution score
1.0
0.8
0.6
0.4
0.2
0.0
PC420
CXCR2
MPO
MARK2P8
AZU1
ACKR1
PLEK
AC092143.1
ADAMTS8
PRRC2A
DDX39B
PIEZO1
CDSN
OCA2
SBNO1
MAMDC2
HCP5
HERC2
JAZF1
GRM5
TCF4
PLCL2
CSF3
TYR
ACAD9
TSBP1
TCF25
AC093827.2
MUC22
TNXB
CEBPE

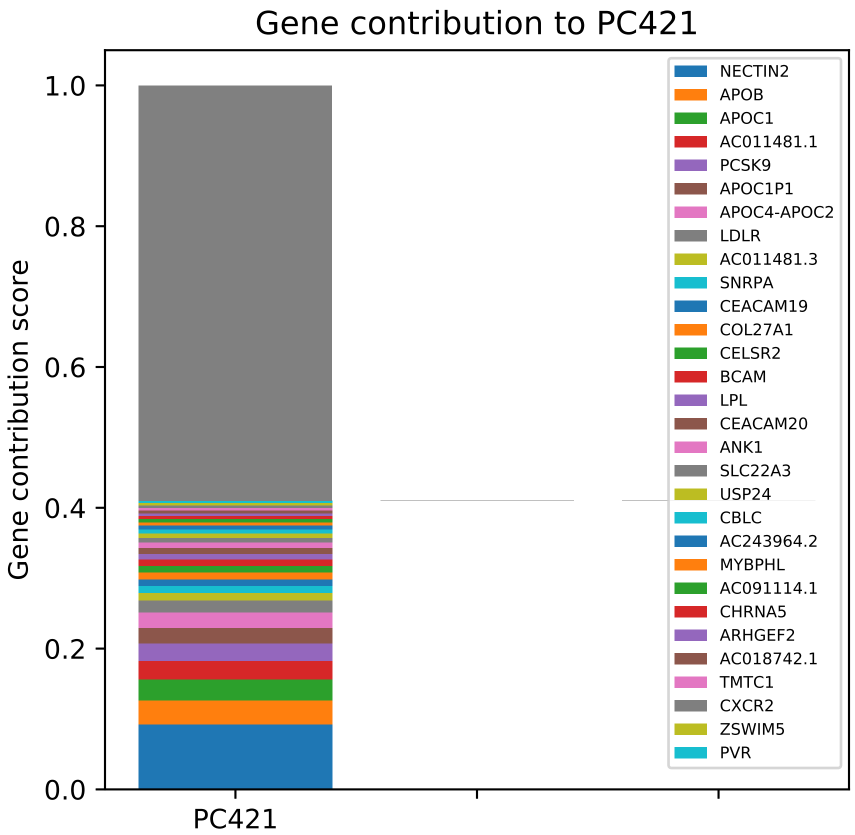
Gene contribution to PC421
Gene contribution score
1.0
0.8
0.6
0.4
0.2
0.0
PC421
NECTIN2
APOB
APOC1
AC011481.1
PCSK9
APOC1P1
APOC4-APOC2
LDLR
AC011481.3
SNRPA
CEACAM19
COL27A1
CELSR2
BCAM
LPL
CEACAM20
ANK1
SLC22A3
USP24
CBLC
AC243964.2
MYBPHL
AC091114.1
CHRNA5
ARHGEF2
AC018742.1
TMTC1
CXCR2
ZSWIM5
PVR

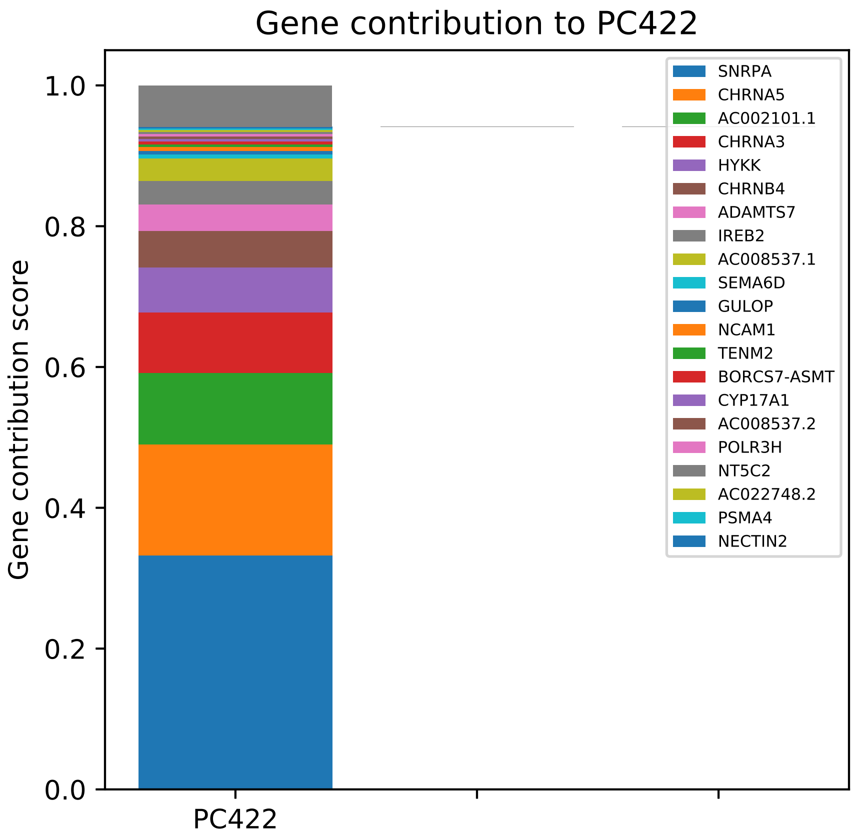

Gene contribution to PC422
Gene contribution score
1.0
0.8
0.6
0.4
0.2
0.0
PC422
SNRPA
CHRNA5
AC002101.1
CHRNA3
HYKK
CHRNB4
ADAMTS7
IREB2
AC008537.1
SEMA6D
GULOP
NCAM1
TENM2
BORCS7-ASMT
CYP17A1
AC008537.2
POLR3H
NT5C2
AC022748.2
PSMA4
NECTIN2

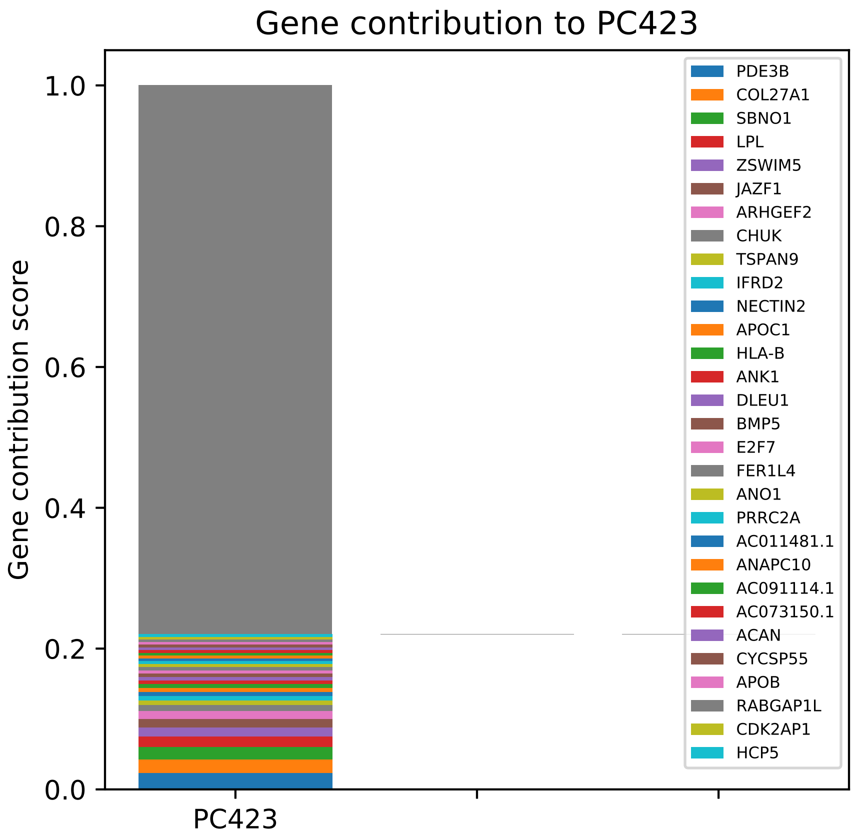

Gene contribution to PC423
1.0
0.8
0.6
0.4
0.2
0.0
Gene contribution score
PC423
PDE3B
COL27A1
SBNO1
LPL
ZSWIM5
JAZF1
ARHGEF2
CHUK
TSPAN9
IFRD2
NECTIN2
APOC1
HLA-B
ANK1
DLEU1
BMP5
E2F7
FER1L4
ANO1
PRRC2A
AC011481.1
ANAPC10
AC091114.1
AC073150.1
ACAN
CYCSP55
APOB
RABGAP1L
CDK2AP1
HCP5

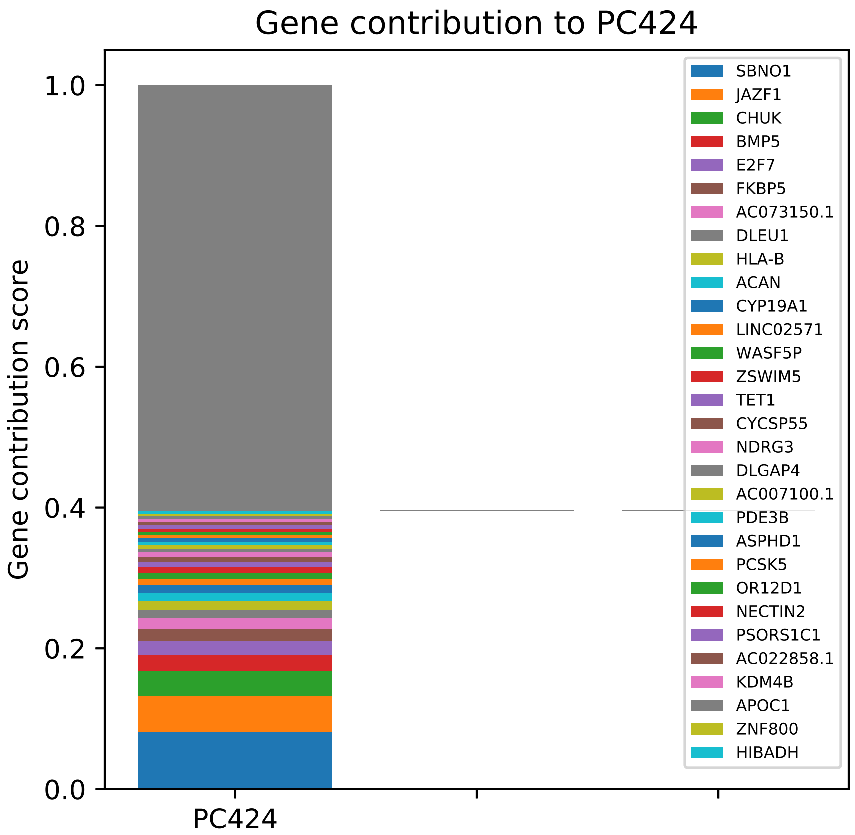
Gene contribution to PC424
Gene contribution score
1.0
0.8
0.6
0.4
0.2
0.0
PC424
SBNO1
JAZF1
CHUK
BMP5
E2F7
FKBP5
AC073150.1
DLEU1
HLA-B
ACAN
CYP19A1
LINC02571
WASF5P
ZSWIM5
TET1
CYCSP55
NDRG3
DLGAP4
AC007100.1
PDE3B
ASPHD1
PCSK5
OR12D1
NECTIN2
PSORS1C1
AC022858.1
KDM4B
APOC1
ZNF800
HIBADH

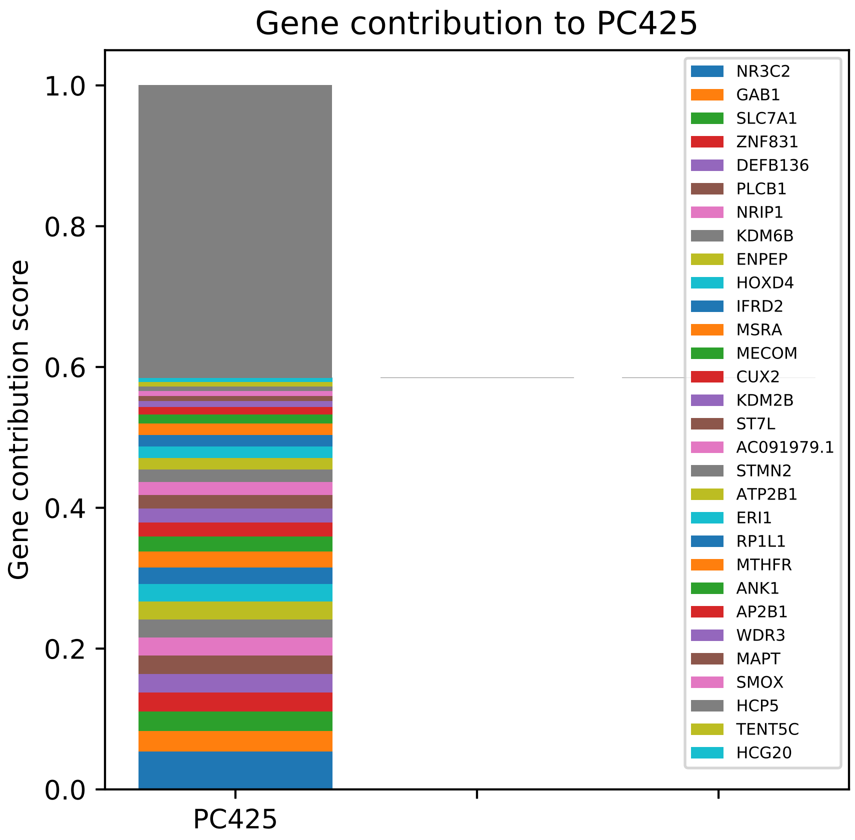

Gene contribution to PC425
Gene contribution score
1.0
0.8
0.6
0.4
0.2
0.0
PC425
NR3C2
GAB1
SLC7A1
ZNF831
DEFB136
PLCB1
NRIP1
KDM6B
ENPEP
HOXD4
IFRD2
MSRA
MECOM
CUX2
KDM2B
ST7L
AC091979.1
STMN2
ATP2B1
ERI1
RP1L1
MTHFR
ANK1
AP2B1
WDR3
MAPT
SMOX
HCP5
TENT5C
HCG20

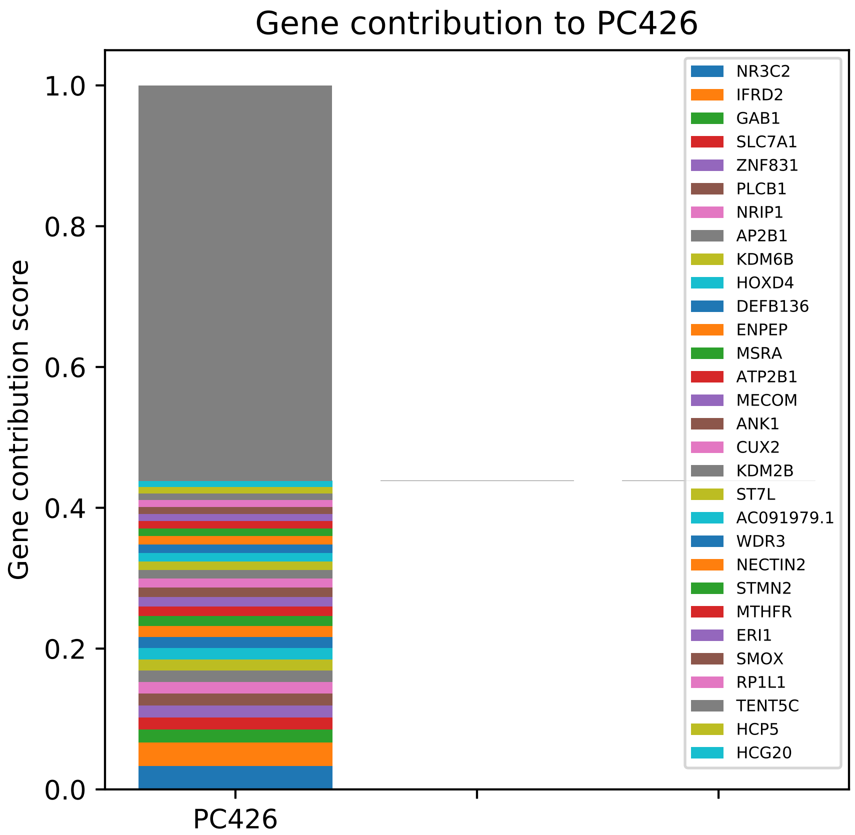

Gene contribution to PC426
Gene contribution score
1.0
0.8
0.6
0.4
0.2
0.0
PC426
NR3C2
IFRD2
GAB1
SLC7A1
ZNF831
PLCB1
NRIP1
AP2B1
KDM6B
HOXD4
DEFB136
ENPEP
MSRA
ATP2B1
MECOM
ANK1
CUX2
KDM2B
ST7L
AC091979.1
WDR3
NECTIN2
STMN2
MTHFR
ERI1
SMOX
RP1L1
TENT5C
HCP5
HCG20

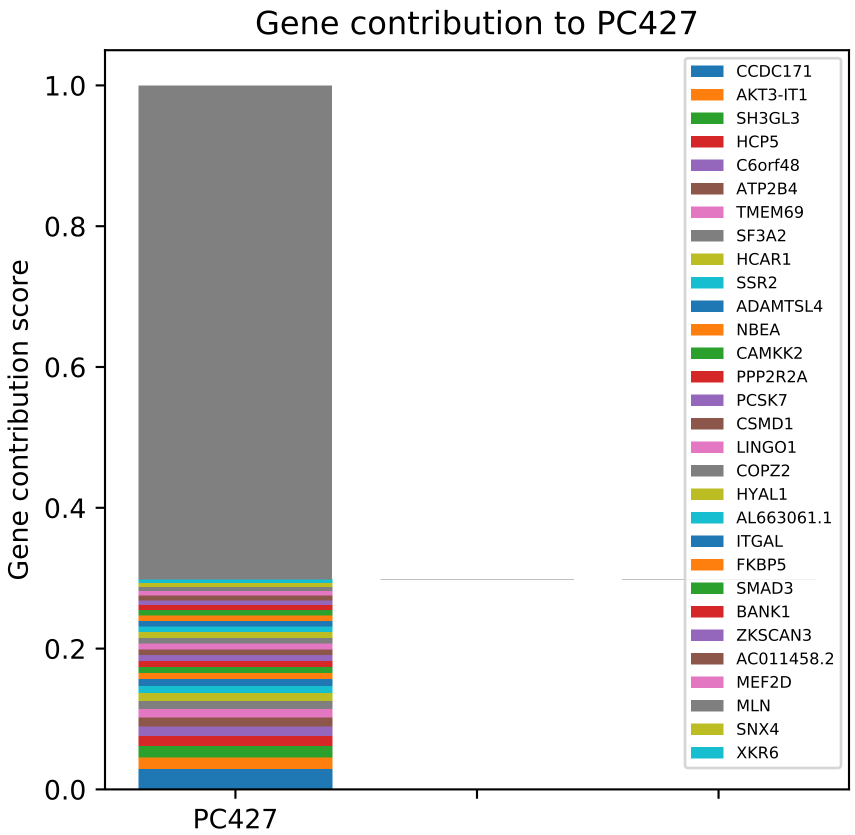

Gene contribution to PC427
Gene contribution score
1.0
0.8
0.6
0.4
0.2
0.0
PC427
CCDC171
AKT3-IT1
SH3GL3
HCP5
C6orf48
ATP2B4
TMEM69
SF3A2
HCAR1
SSR2
ADAMTSL4
NBEA
CAMKK2
PPP2R2A
PCSK7
CSMD1
LINGO1
COPZ2
HYAL1
AL663061.1
ITGAL
FKBP5
SMAD3
BANK1
ZKSCAN3
AC011458.2
MEF2D
MLN
SNX4
XKR6

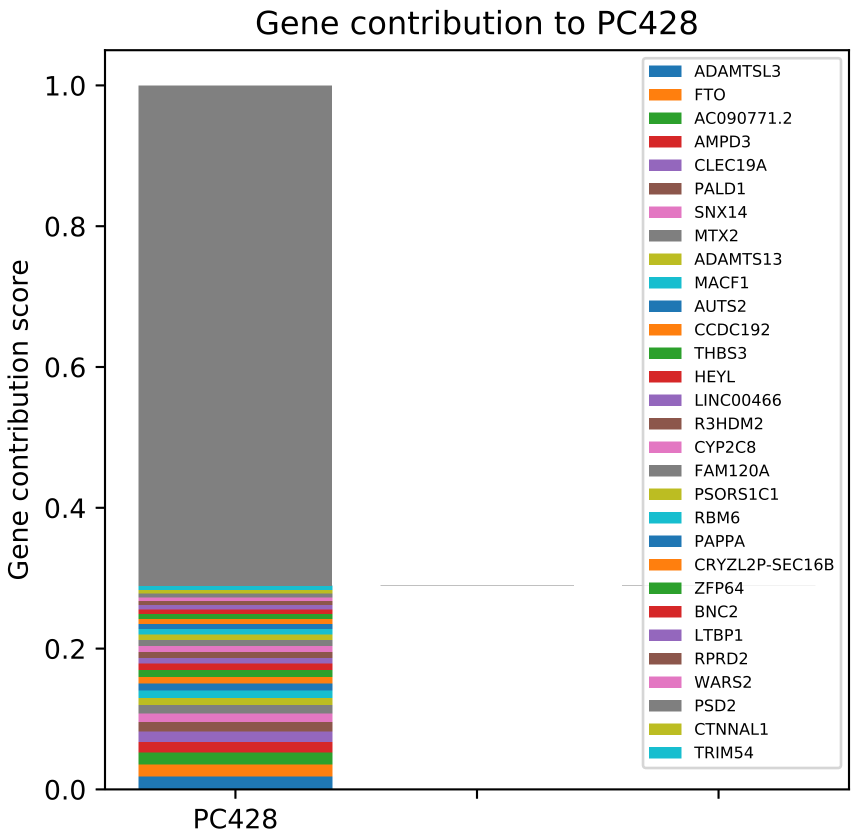
Gene contribution to PC428
Gene contribution score
1.0
0.8
0.6
0.4
0.2
0.0
PC428
ADAMTSL3
FTO
AC090771.2
AMPD3
CLEC19A
PALD1
SNX14
MTX2
ADAMTS13
MACF1
AUTS2
CCDC192
THBS3
HEYL
LINC00466
R3HDM2
CYP2C8
FAM120A
PSORS1C1
RBM6
PAPPA
CRYZL2P-SEC16B
ZFP64
BNC2
LTBP1
RPRD2
WARS2
PSD2
CTNNAL1
TRIM54

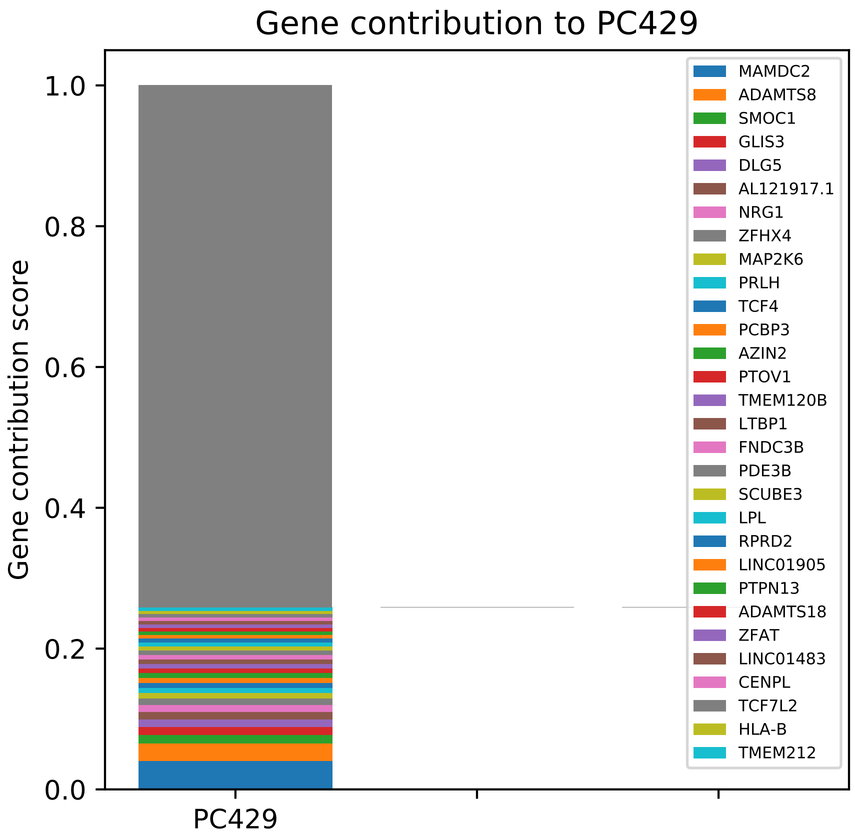

Gene contribution to PC429
Gene contribution score
1.0
0.8
0.6
0.4
0.2
0.0
PC429
MAMDC2
ADAMTS8
SMOC1
GLIS3
DLG5
AL121917.1
NRG1
ZFHX4
MAP2K6
PRLH
TCF4
PCBP3
AZIN2
PTOV1
TMEM120B
LTBP1
FNDC3B
PDE3B
SCUBE3
LPL
RPRD2
LINC01905
PTPN13
ADAMTS18
ZFAT
LINC01483
CENPL
TCF7L2
HLA-B
TMEM212

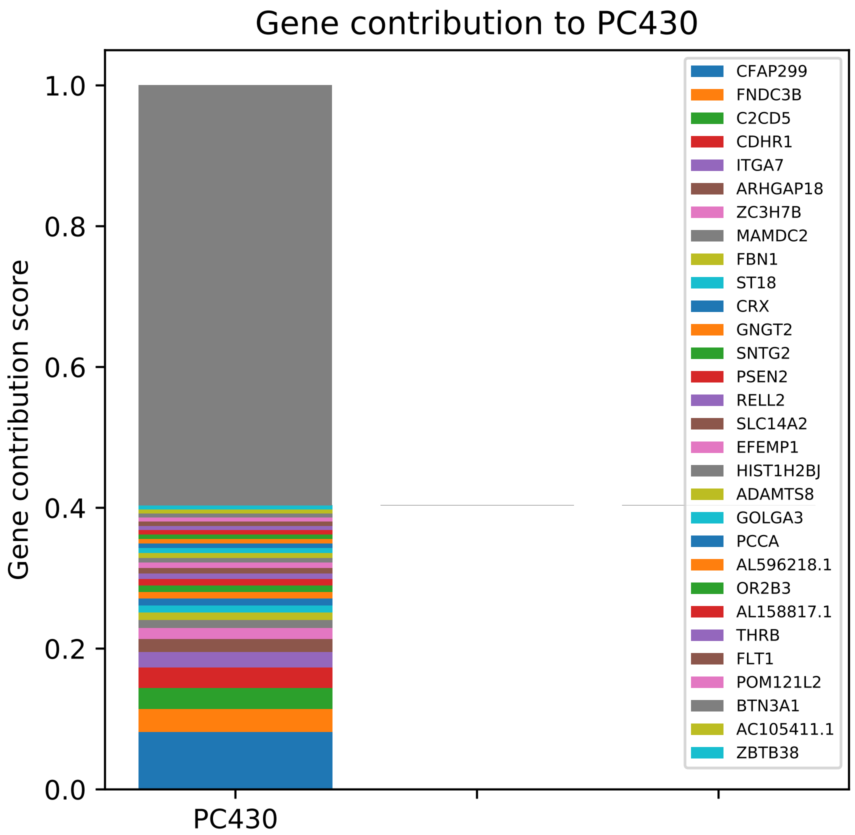

Gene contribution to PC430
Gene contribution score
1.0
0.8
0.6
0.4
0.2
0.0
PC430
CFAP299
FNDC3B
C2CD5
CDHR1
ITGA7
ARHGAP18
ZC3H7B
MAMDC2
FBN1
ST18
CRX
GNGT2
SNTG2
PSEN2
RELL2
SLC14A2
EFEMP1
HIST1H2BJ
ADAMTS8
GOLGA3
PCCA
AL596218.1
OR2B3
AL158817.1
THRB
FLT1
POM121L2
BTN3A1
AC105411.1
ZBTB38

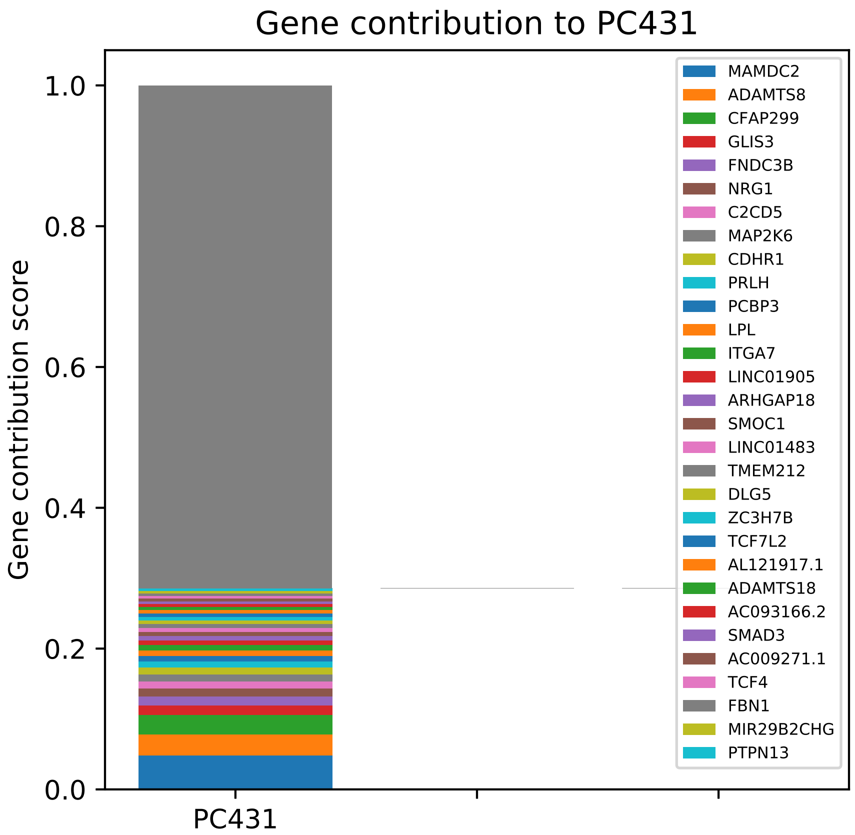

Gene contribution to PC431
Gene contribution score
1.0
0.8
0.6
0.4
0.2
0.0
PC431
MAMDC2
ADAMTS8
CFAP299
GLIS3
FNDC3B
NRG1
C2CD5
MAP2K6
CDHR1
PRLH
PCBP3
LPL
ITGA7
LINC01905
ARHGAP18
SMOC1
LINC01483
TMEM212
DLG5
ZC3H7B
TCF7L2
AL121917.1
ADAMTS18
AC093166.2
SMAD3
AC009271.1
TCF4
FBN1
MIR29B2CHG
PTPN13

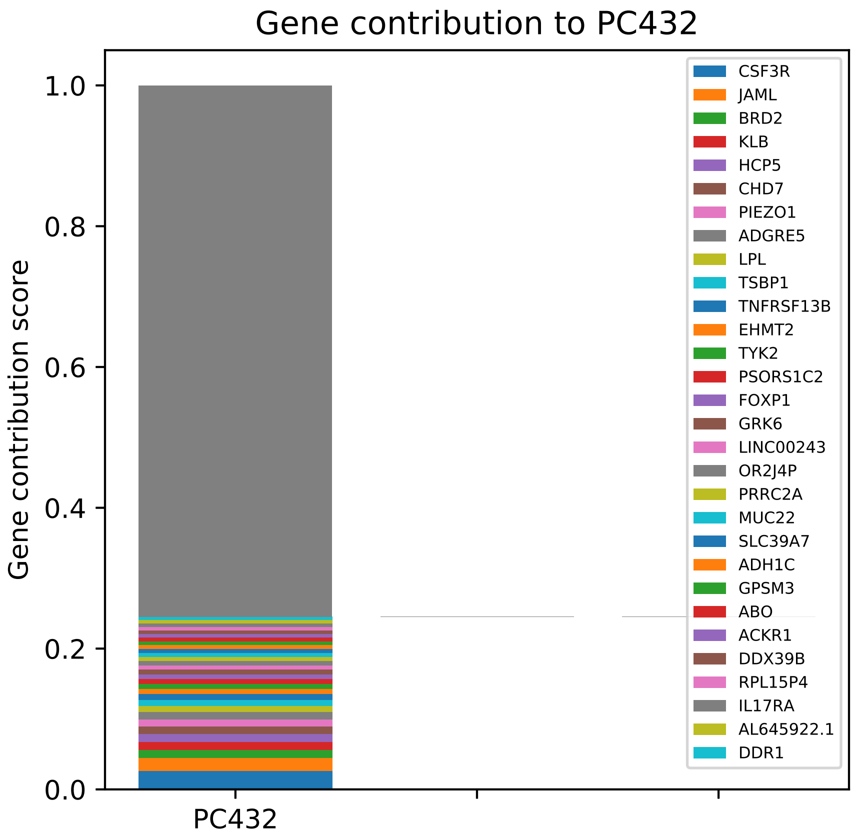

Gene contribution to PC432
Gene contribution score
1.0
0.8
0.6
0.4
0.2
0.0
PC432
CSF3R
JAML
BRD2
KLB
HCP5
CHD7
PIEZO1
ADGRE5
LPL
TSBP1
TNFRSF13B
EHMT2
TYK2
PSORS1C2
FOXP1
GRK6
LINC00243
OR2J4P
PRRC2A
MUC22
SLC39A7
ADH1C
GPSM3
ABO
ACKR1
DDX39B
RPL15P4
IL17RA
AL645922.1
DDR1

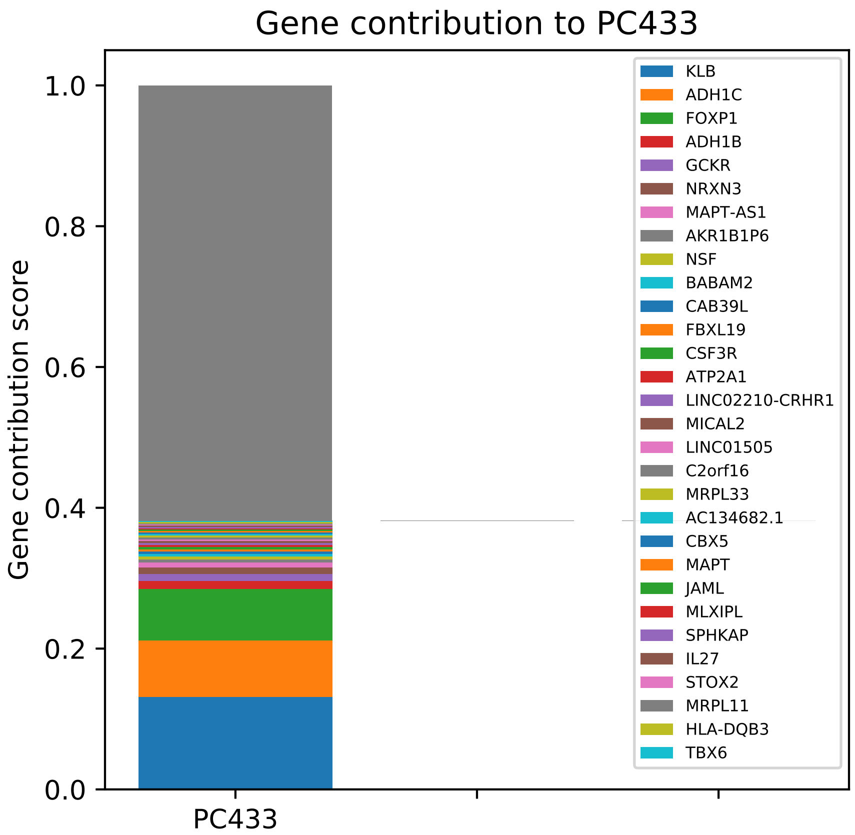
Gene contribution to PC433
Gene contribution score
1.0
0.8
0.6
0.4
0.2
0.0
PC433
KLB
ADH1C
FOXP1
ADH1B
GCKR
NRXN3
MAPT-AS1
AKR1B1P6
NSF
BABAM2
CAB39L
FBXL19
CSF3R
ATP2A1
LINC02210-CRHR1
MICAL2
LINC01505
C2orf16
MRPL33
AC134682.1
CBX5
MAPT
JAML
MLXIPL
SPHKAP
IL27
STOX2
MRPL11
HLA-DQB3
TBX6

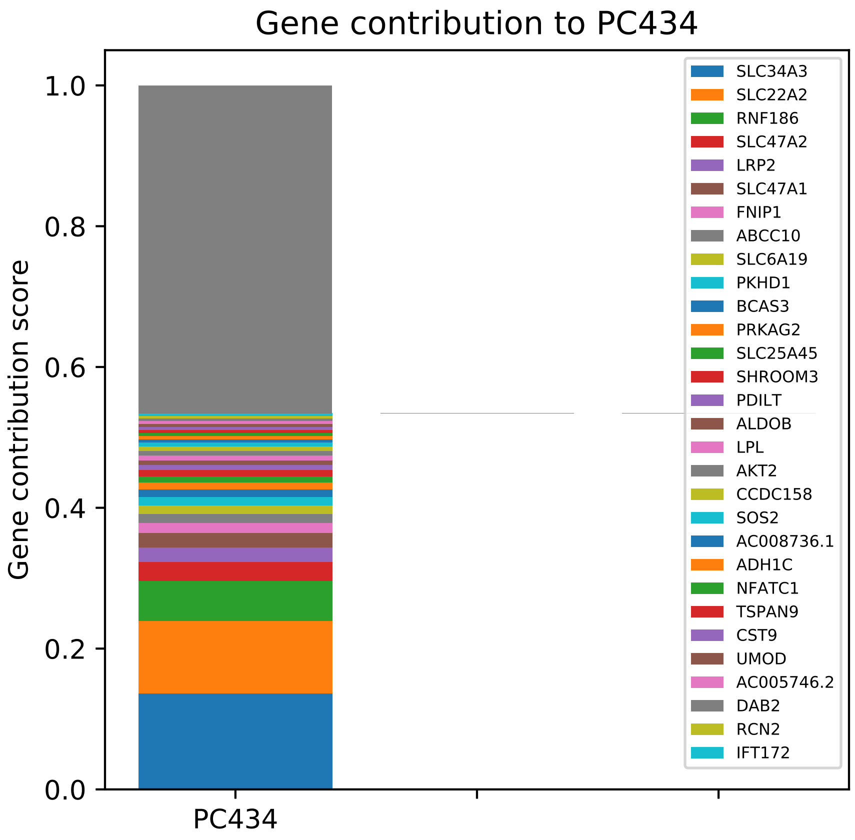

Gene contribution to PC434
Gene contribution score
1.0
0.8
0.6
0.4
0.2
0.0
PC434
SLC34A3
SLC22A2
RNF186
SLC47A2
LRP2
SLC47A1
FNIP1
ABCC10
SLC6A19
PKHD1
BCAS3
PRKAG2
SLC25A45
SHROOM3
PDILT
ALDOB
LPL
AKT2
CCDC158
SOS2
AC008736.1
ADH1C
NFATC1
TSPAN9
CST9
UMOD
AC005746.2
DAB2
RCN2
IFT172

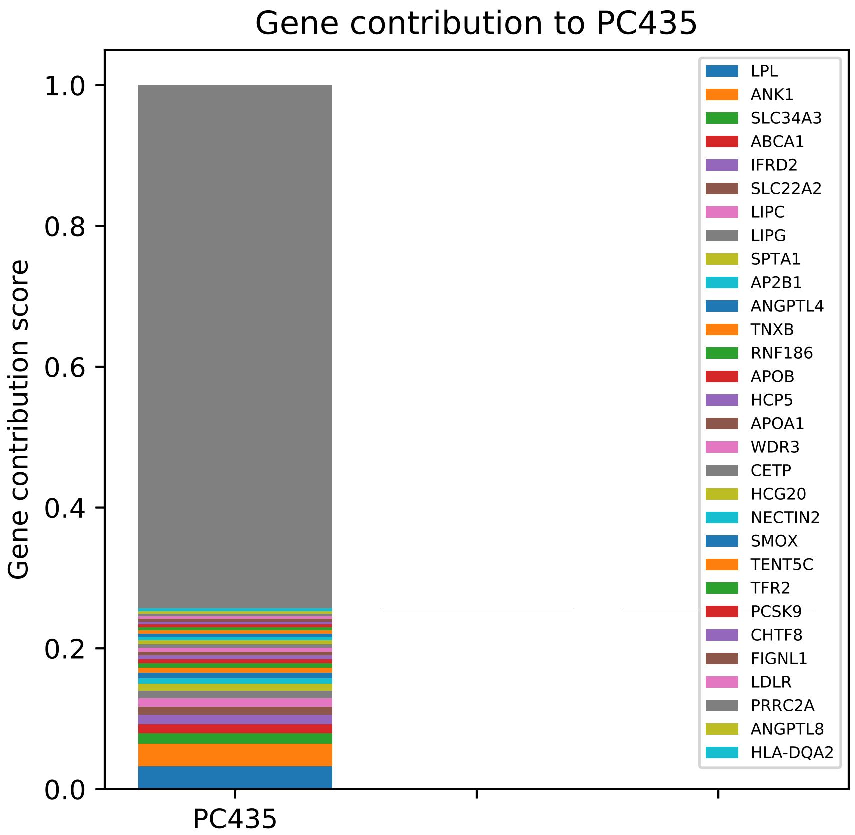

Gene contribution to PC435
Gene contribution score
1.0
0.8
0.6
0.4
0.2
0.0
PC435
LPL
ANK1
SLC34A3
ABCA1
IFRD2
SLC22A2
LIPC
LIPG
SPTA1
AP2B1
ANGPTL4
TNXB
RNF186
APOB
HCP5
APOA1
WDR3
CETP
HCG20
NECTIN2
SMOX
TENT5C
TFR2
PCSK9
CHTF8
FIGNL1
LDLR
PRRC2A
ANGPTL8
HLA-DQA2

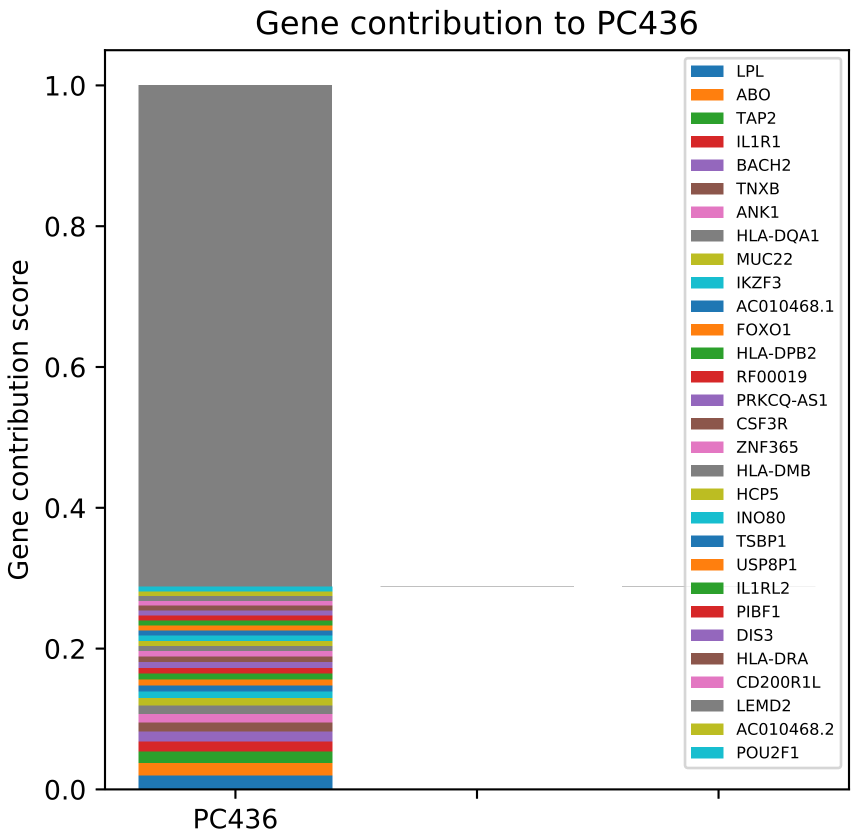
Gene contribution to PC436
Gene contribution score
1.0
0.8
0.6
0.4
0.2
0.0
PC436
LPL
ABO
TAP2
IL1R1
BACH2
TNXB
ANK1
HLA-DQA1
MUC22
IKZF3
AC010468.1
FOXO1
HLA-DPB2
RF00019
PRKCQ-AS1
CSF3R
ZNF365
HLA-DMB
HCP5
INO80
TSBP1
USP8P1
IL1RL2
PIBF1
DIS3
HLA-DRA
CD200R1L
LEMD2
AC010468.2
POU2F1

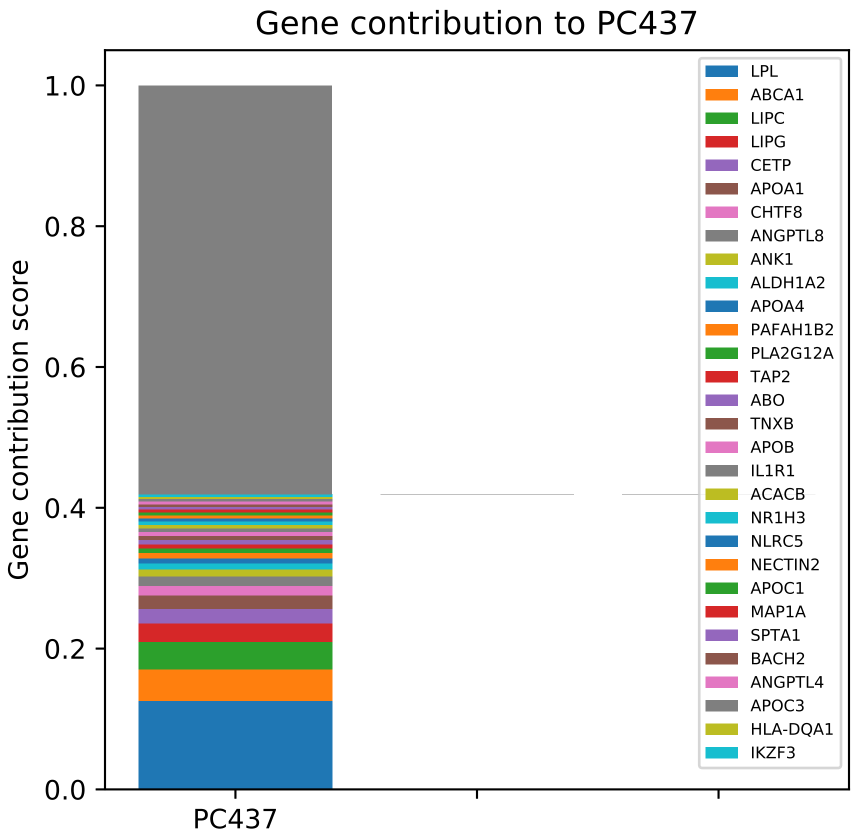

Gene contribution to PC437
Gene contribution score
1.0
0.8
0.6
0.4
0.2
0.0
PC437
LPL
ABCA1
LIPC
LIPG
CETP
APOA1
CHTF8
ANGPTL8
ANK1
ALDH1A2
APOA4
PAFAH1B2
PLA2G12A
TAP2
ABO
TNXB
APOB
IL1R1
ACACB
NR1H3
NLRC5
NECTIN2
APOC1
MAP1A
SPTA1
BACH2
ANGPTL4
APOC3
HLA-DQA1
IKZF3

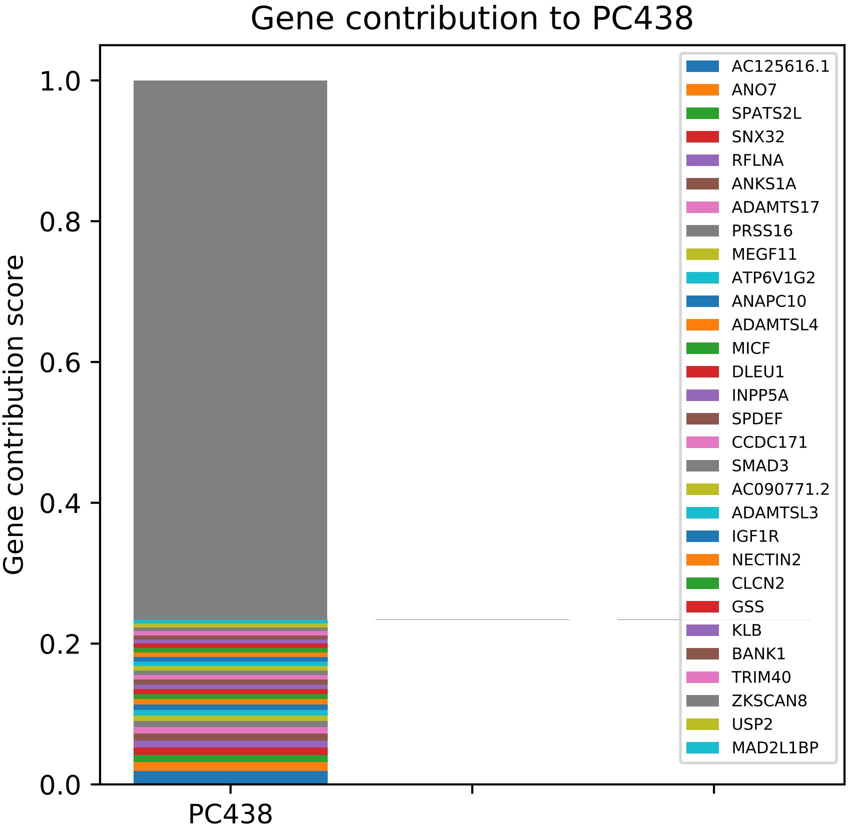

Gene contribution to PC438
Gene contribution score
1.0
0.8
0.6
0.4
0.2
0.0
PC438
AC125616.1
ANO7
SPATS2L
SNX32
RFLNA
ANKS1A
ADAMTS17
PRSS16
MEGF11
ATP6V1G2
ANAPC10
ADAMTSL4
MICF
DLEU1
INPP5A
SPDEF
CCDC171
SMAD3
AC090771.2
ADAMTSL3
IGF1R
NECTIN2
CLCN2
GSS
KLB
BANK1
TRIM40
ZKSCAN8
USP2
MAD2L1BP

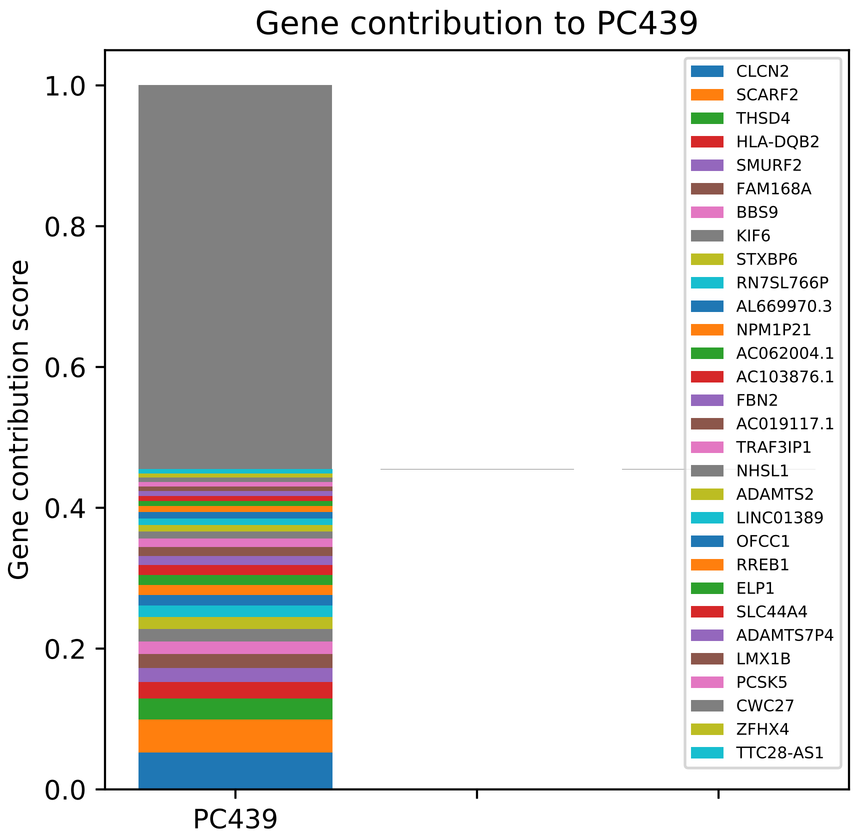

Gene contribution to PC439
Gene contribution score
1.0
0.8
0.6
0.4
0.2
0.0
PC439
CLCN2
SCARF2
THSD4
HLA-DQB2
SMURF2
FAM168A
BBS9
KIF6
STXBP6
RN7SL766P
AL669970.3
NPM1P21
AC062004.1
AC103876.1
FBN2
AC019117.1
TRAF3IP1
NHSL1
ADAMTS2
LINC01389
OFCC1
RREB1
ELP1
SLC44A4
ADAMTS7P4
LMX1B
PCSK5
CWC27
ZFHX4
TTC28-AS1

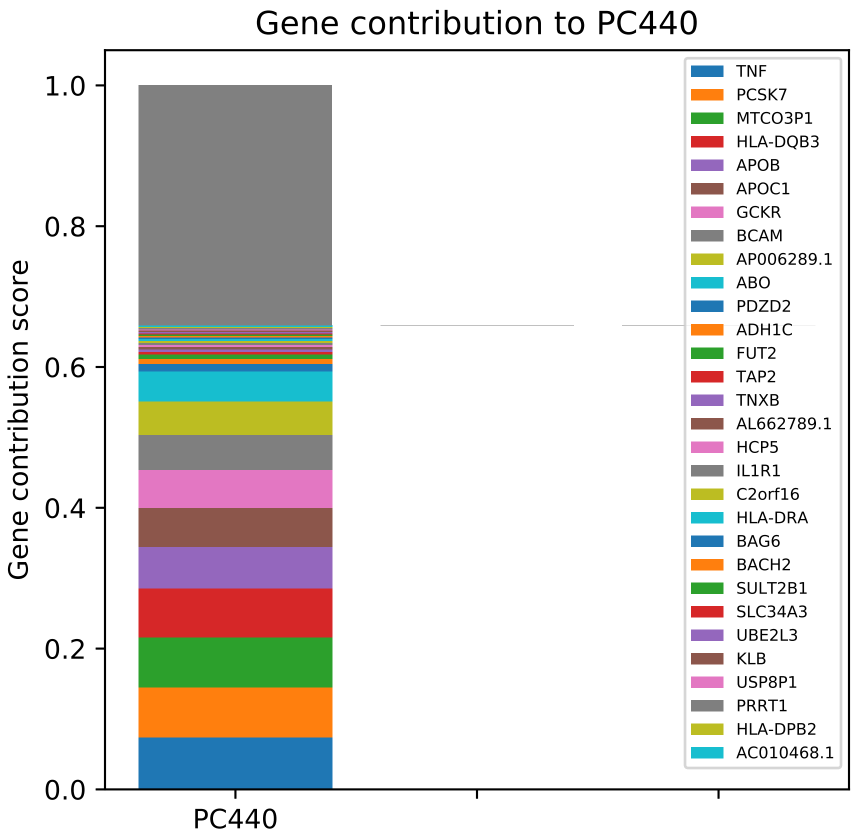
Gene contribution to PC440
Gene contribution score
PC440
0.0
0.2
0.4
0.6
0.8
1.0
TNF
PCSK7
MTCO3P1
HLA-DQB3
APOB
APOC1
GCKR
BCAM
AP006289.1
ABO
PDZD2
ADH1C
FUT2
TAP2
TNXB
AL662789.1
HCP5
IL1R1
C2orf16
HLA-DRA
BAG6
BACH2
SULT2B1
SLC34A3
UBE2L3
KLB
USP8P1
PRRT1
HLA-DPB2
AC010468.1

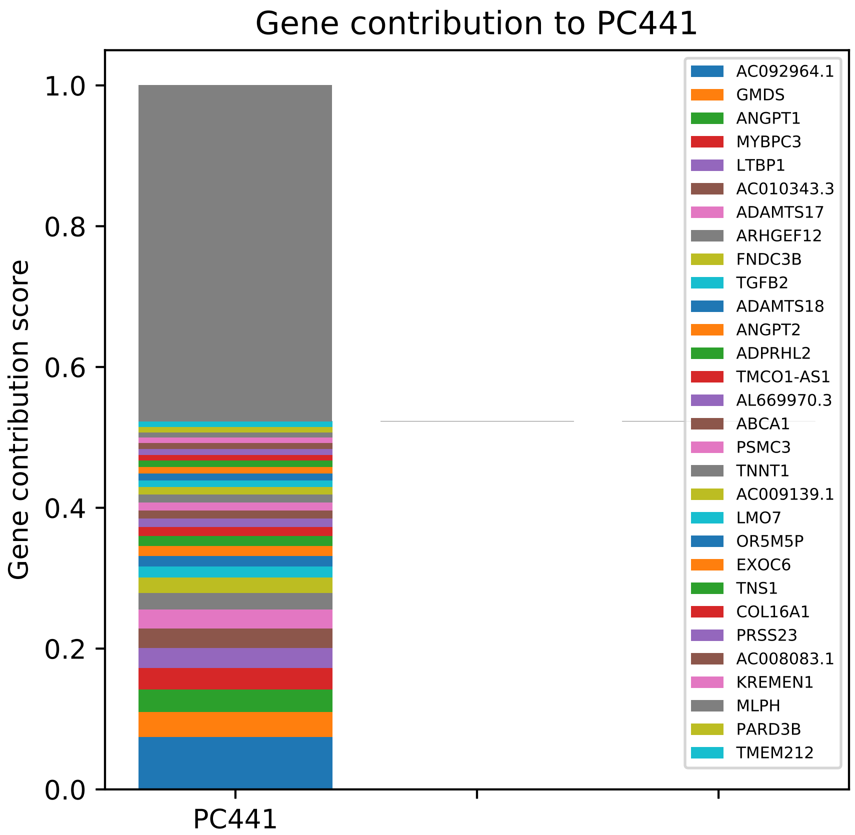

Gene contribution to PC441
Gene contribution score
1.0
0.8
0.6
0.4
0.2
0.0
PC441
AC092964.1
GMDS
ANGPT1
MYBPC3
LTBP1
AC010343.3
ADAMTS17
ARHGEF12
FNDC3B
TGFB2
ADAMTS18
ANGPT2
ADPRHL2
TMCO1-AS1
AL669970.3
ABCA1
PSMC3
TNNT1
AC009139.1
LMO7
OR5M5P
EXOC6
TNS1
COL16A1
PRSS23
AC008083.1
KREMEN1
MLPH
PARD3B
TMEM212

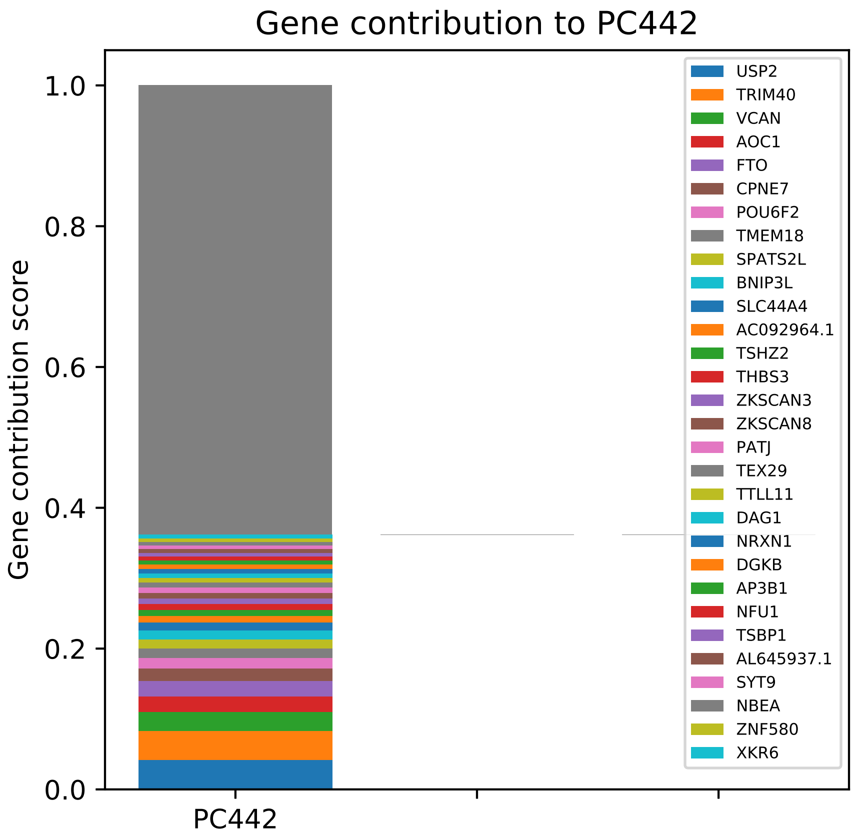
Gene contribution to PC442
Gene contribution score
1.0
0.8
0.6
0.4
0.2
0.0
PC442
USP2
TRIM40
VCAN
AOC1
FTO
CPNE7
POU6F2
TMEM18
SPATS2L
BNIP3L
SLC44A4
AC092964.1
TSHZ2
THBS3
ZKSCAN3
ZKSCAN8
PATJ
TEX29
TTLL11
DAG1
NRXN1
DGKB
AP3B1
NFU1
TSBP1
AL645937.1
SYT9
NBEA
ZNF580
XKR6

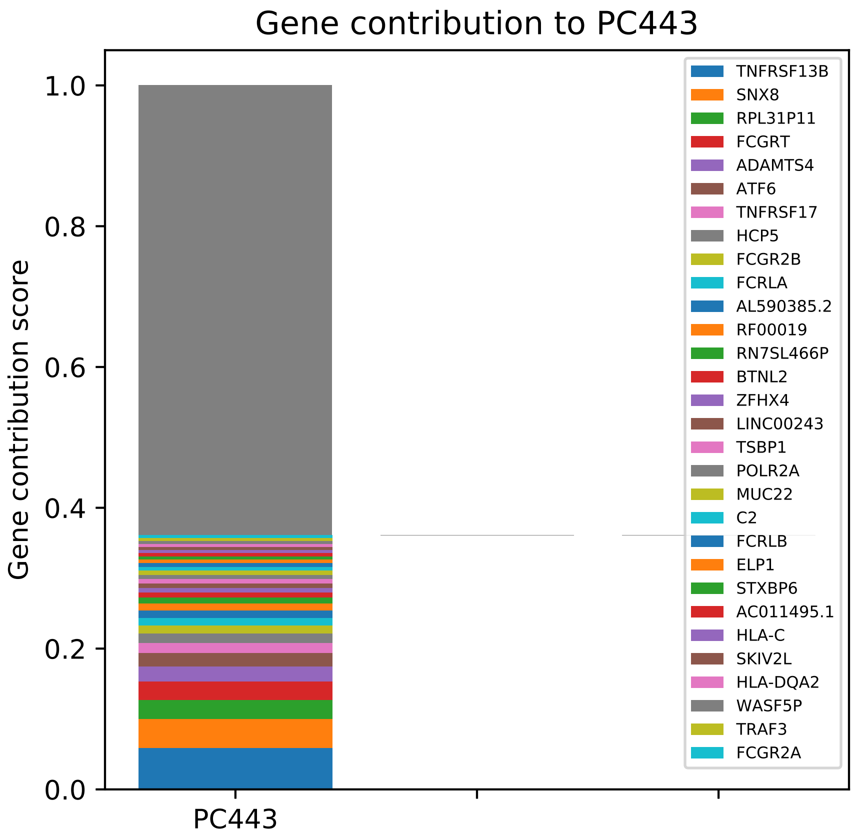

Gene contribution to PC443
Gene contribution score
1.0
0.8
0.6
0.4
0.2
0.0
PC443
TNFRSF13B
SNX8
RPL31P11
FCGRT
ADAMTS4
ATF6
TNFRSF17
HCP5
FCGR2B
FCRLA
AL590385.2
RF00019
RN7SL466P
BTNL2
ZFHX4
LINC00243
TSBP1
POLR2A
MUC22
C2
FCRLB
ELP1
STXBP6
AC011495.1
HLA-C
SKIV2L
HLA-DQA2
WASF5P
TRAF3
FCGR2A

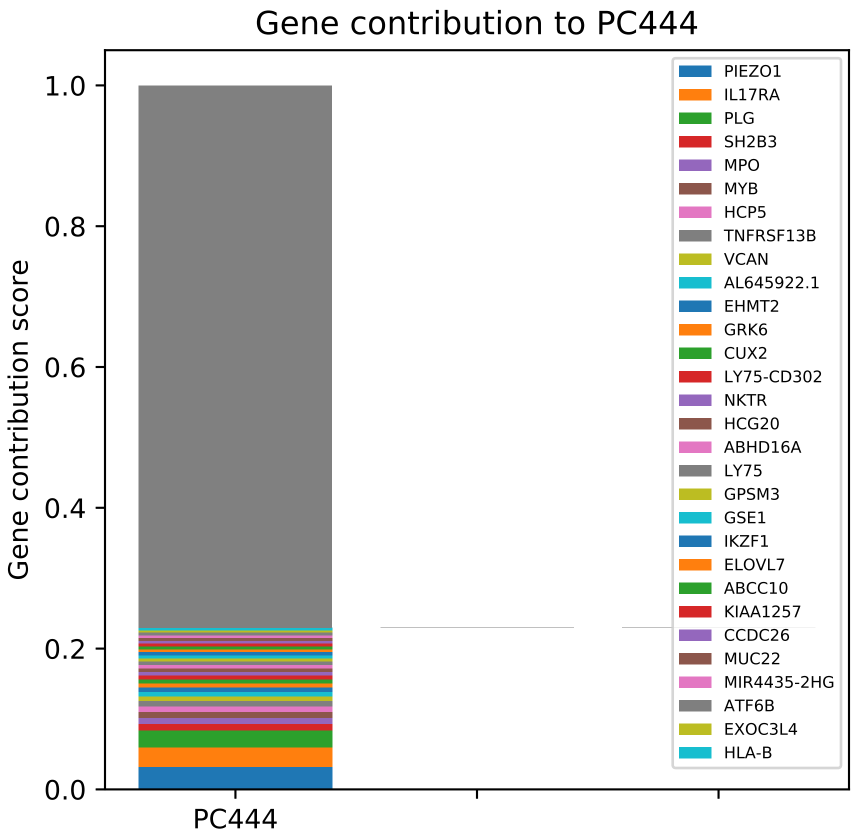

Gene contribution to PC444
Gene contribution score
1.0
0.8
0.6
0.4
0.2
0.0
PC444
PIEZO1
IL17RA
PLG
SH2B3
MPO
MYB
HCP5
TNFRSF13B
VCAN
AL645922.1
EHMT2
GRK6
CUX2
LY75-CD302
NKTR
HCG20
ABHD16A
LY75
GPSM3
GSE1
IKZF1
ELOVL7
ABCC10
KIAA1257
CCDC26
MUC22
MIR4435-2HG
ATF6B
EXOC3L4
HLA-B

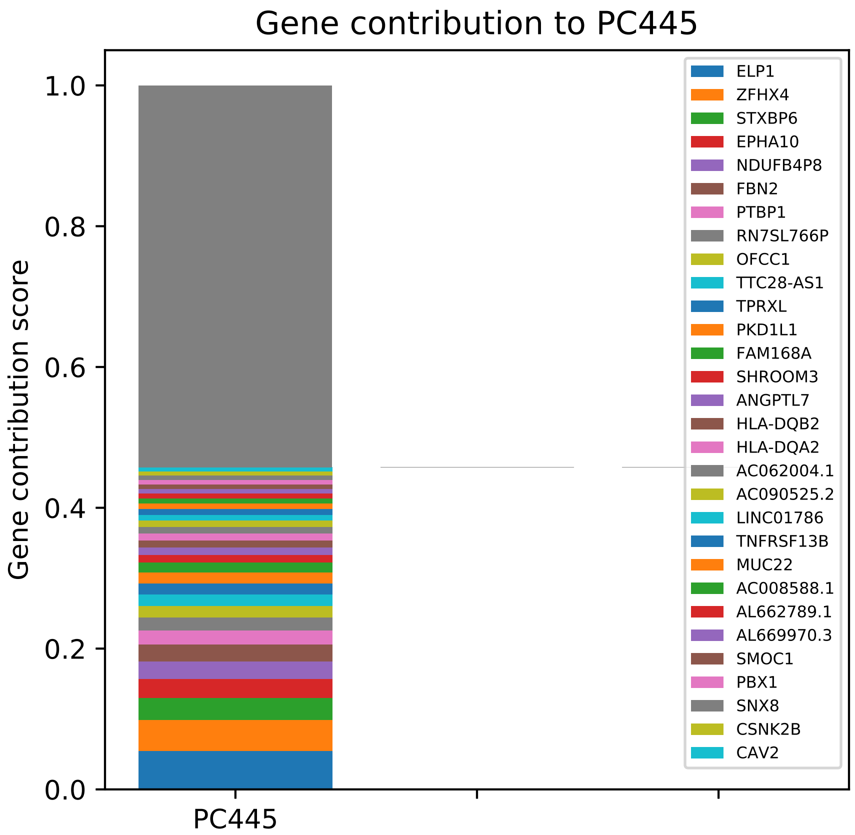

Gene contribution to PC445
Gene contribution score
1.0
0.8
0.6
0.4
0.2
0.0
PC445
ELP1
ZFHX4
STXBP6
EPHA10
NDUFB4P8
FBN2
PTBP1
RN7SL766P
OFCC1
TTC28-AS1
TPRXL
PKD1L1
FAM168A
SHROOM3
ANGPTL7
HLA-DQB2
HLA-DQA2
AC062004.1
AC090525.2
LINC01786
TNFRSF13B
MUC22
AC008588.1
AL662789.1
AL669970.3
SMOC1
PBX1
SNX8
CSNK2B
CAV2

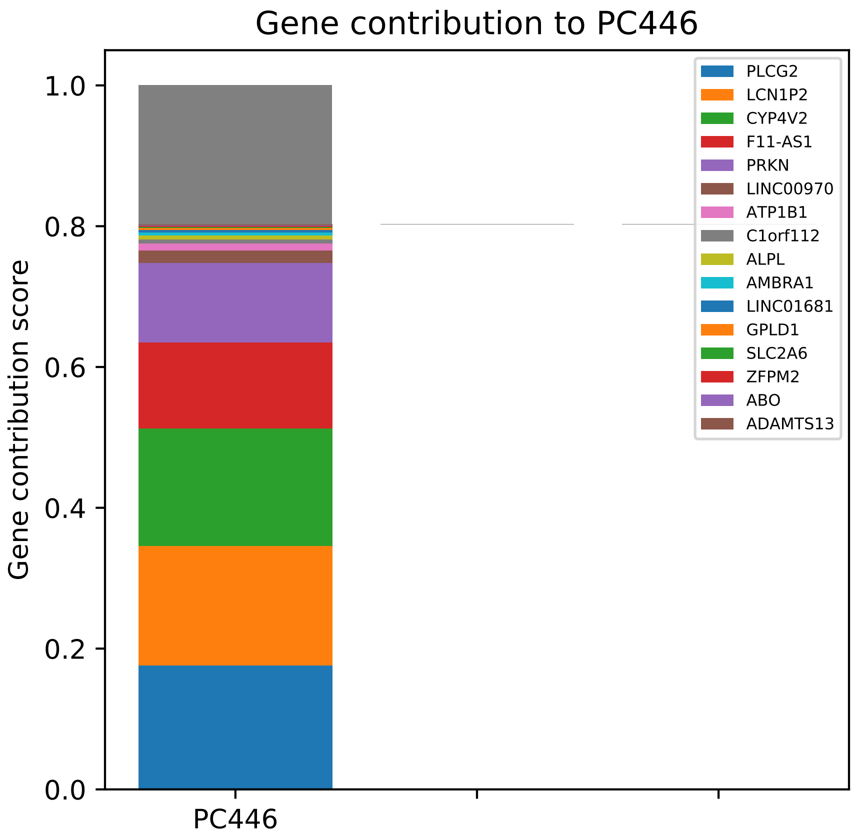
Gene contribution to PC446
Gene contribution score
0.0
0.2
0.4
0.6
0.8
1.0
PC446
PLCG2
LCN1P2
CYP4V2
F11-AS1
PRKN
LINC00970
ATP1B1
C1orf112
ALPL
AMBRA1
LINC01681
GPLD1
SLC2A6
ZFPM2
ABO
ADAMTS13

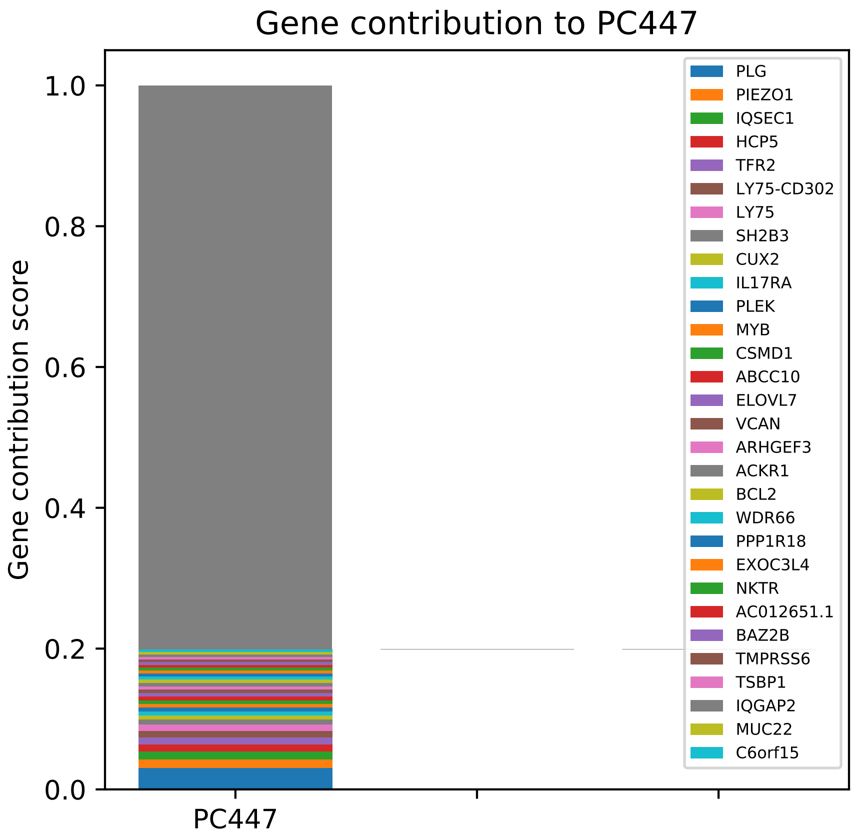

Gene contribution to PC447
Gene contribution score
1.0
0.8
0.6
0.4
0.2
0.0
PC447
PLG
PIEZO1
IQSEC1
HCP5
TFR2
LY75-CD302
LY75
SH2B3
CUX2
IL17RA
PLEK
MYB
CSMD1
ABCC10
ELOVL7
VCAN
ARHGEF3
ACKR1
BCL2
WDR66
PPP1R18
EXOC3L4
NKTR
AC012651.1
BAZ2B
TMPRSS6
TSBP1
IQGAP2
MUC22
C6orf15

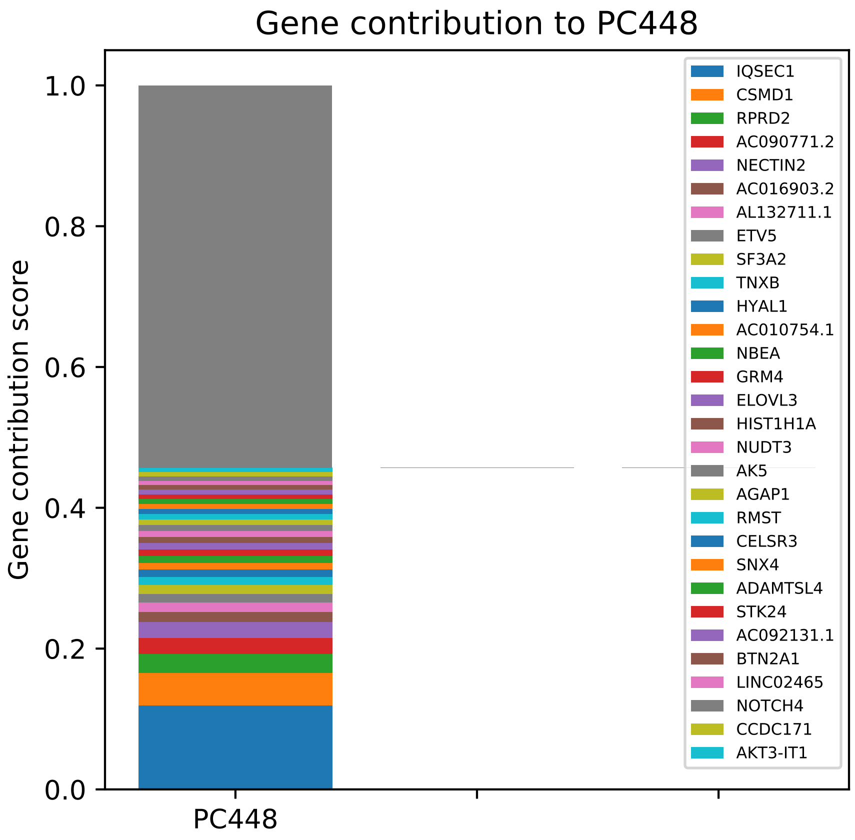

Gene contribution to PC448
Gene contribution score
1.0
0.8
0.6
0.4
0.2
0.0
PC448
IQSEC1
CSMD1
RPRD2
AC090771.2
NECTIN2
AC016903.2
AL132711.1
ETV5
SF3A2
TNXB
HYAL1
AC010754.1
NBEA
GRM4
ELOVL3
HIST1H1A
NUDT3
AK5
AGAP1
RMST
CELSR3
SNX4
ADAMTSL4
STK24
AC092131.1
BTN2A1
LINC02465
NOTCH4
CCDC171
AKT3-IT1

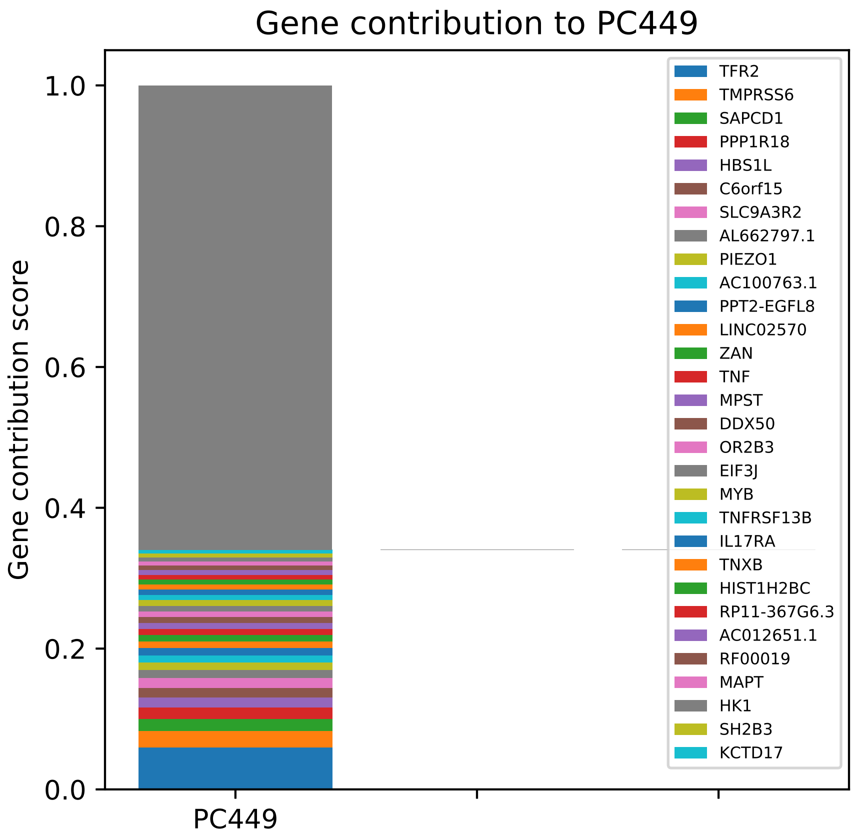

Gene contribution to PC449
Gene contribution score
1.0
0.8
0.6
0.4
0.2
0.0
PC449
TFR2
TMPRSS6
SAPCD1
PPP1R18
HBS1L
C6orf15
SLC9A3R2
AL662797.1
PIEZO1
AC100763.1
PPT2-EGFL8
LINC02570
ZAN
TNF
MPST
DDX50
OR2B3
EIF3J
MYB
TNFRSF13B
IL17RA
TNXB
HIST1H2BC
RP11-367G6.3
AC012651.1
RF00019
MAPT
HK1
SH2B3
KCTD17

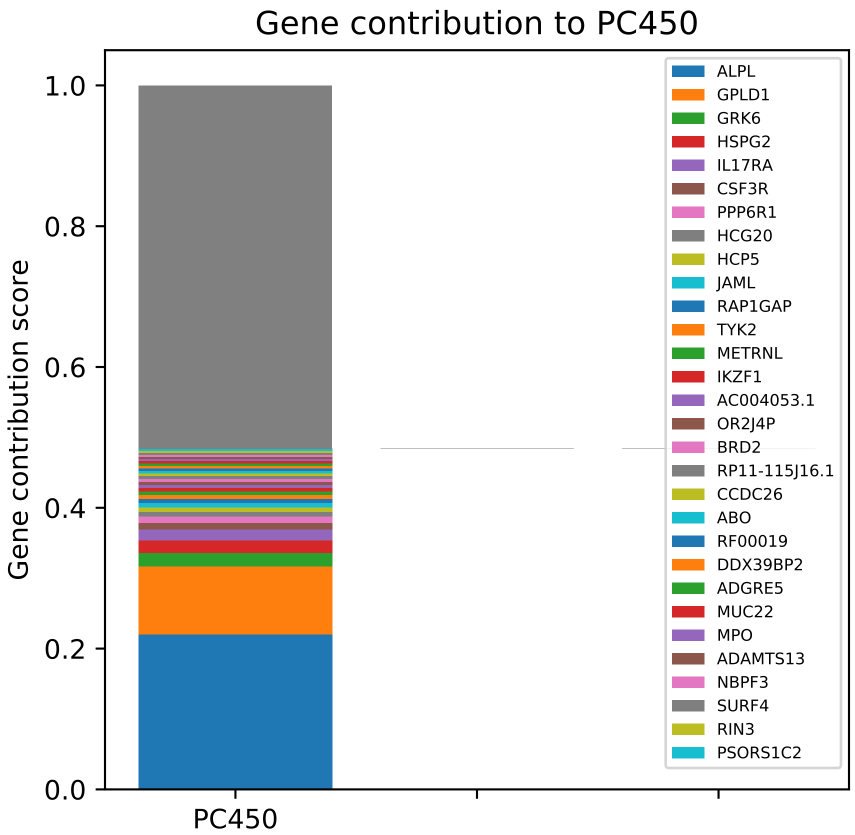
Gene contribution to PC450
Gene contribution score
1.0
0.8
0.6
0.4
0.2
0.0
PC450
ALPL
GPLD1
GRK6
HSPG2
IL17RA
CSF3R
PPP6R1
HCG20
HCP5
JAML
RAP1GAP
TYK2
METRNL
IKZF1
AC004053.1
OR2J4P
BRD2
RP11-115J16.1
CCDC26
ABO
RF00019
DDX39BP2
ADGRE5
MUC22
MPO
ADAMTS13
NBPF3
SURF4
RIN3
PSORS1C2

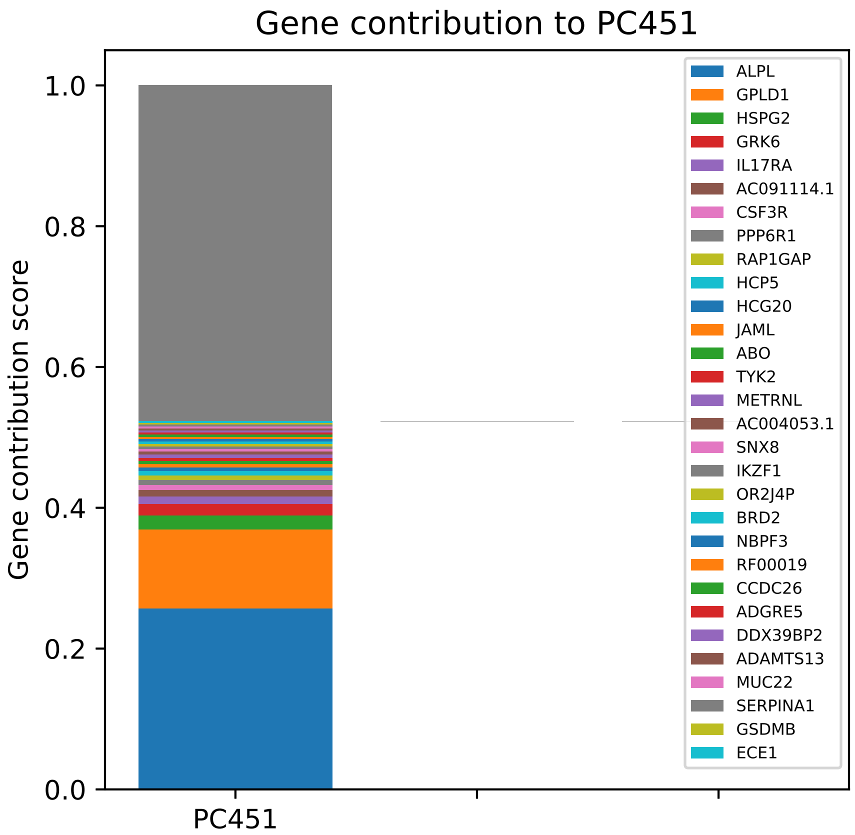

Gene contribution to PC451
Gene contribution score
1.0
0.8
0.6
0.4
0.2
0.0
PC451
ALPL
GPLD1
HSPG2
GRK6
IL17RA
AC091114.1
CSF3R
PPP6R1
RAP1GAP
HCP5
HCG20
JAML
ABO
TYK2
METRNL
AC004053.1
SNX8
IKZF1
OR2J4P
BRD2
NBPF3
RF00019
CCDC26
ADGRE5
DDX39BP2
ADAMTS13
MUC22
SERPINA1
GSDMB
ECE1

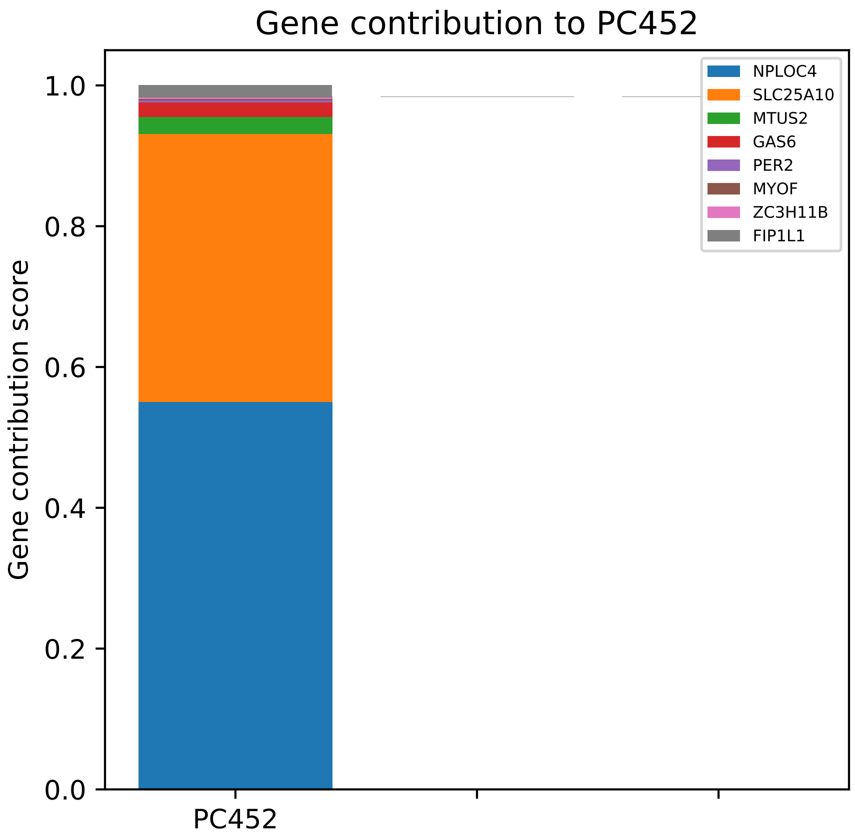

Gene contribution to PC452
Gene contribution score
0.0
0.2
0.4
0.6
0.8
1.0
PC452
NPLOC4
SLC25A10
MTUS2
GAS6
PER2
MYOF
ZC3H11B
FIP1L1

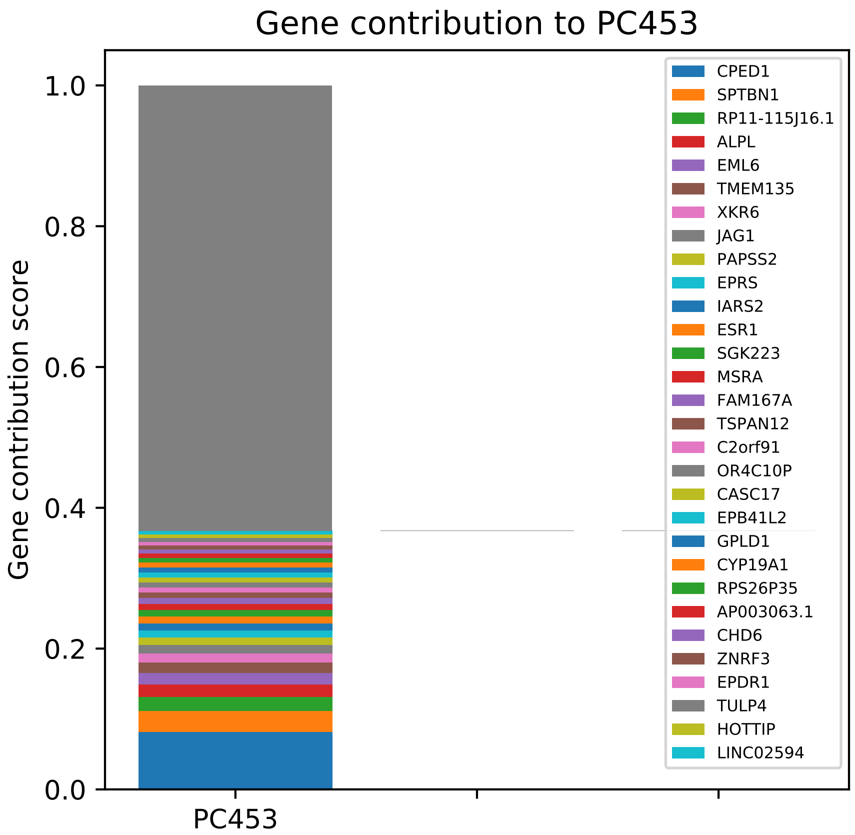

Gene contribution to PC453
Gene contribution score
1.0
0.8
0.6
0.4
0.2
0.0
PC453
CPED1
SPTBN1
RP11-115J16.1
ALPL
EML6
TMEM135
XKR6
JAG1
PAPSS2
EPRS
IARS2
ESR1
SGK223
MSRA
FAM167A
TSPAN12
C2orf91
OR4C10P
CASC17
EPB41L2
GPLD1
CYP19A1
RPS26P35
AP003063.1
CHD6
ZNRF3
EPDR1
TULP4
HOTTIP
LINC02594

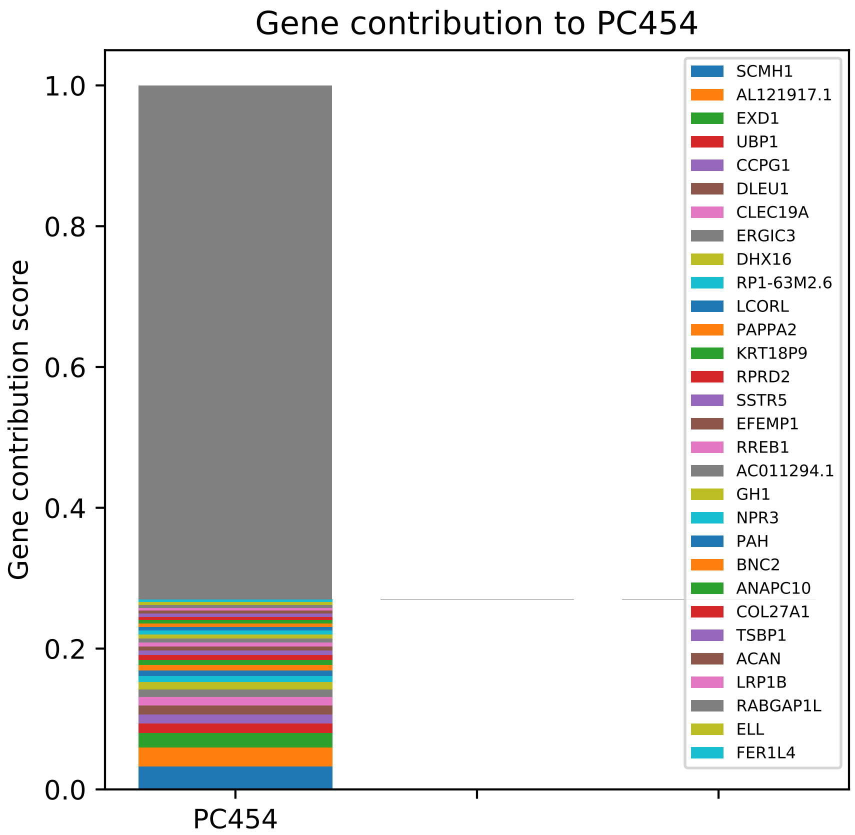

Gene contribution to PC454
Gene contribution score
1.0
0.8
0.6
0.4
0.2
0.0
PC454
SCMH1
AL121917.1
EXD1
UBP1
CCPG1
DLEU1
CLEC19A
ERGIC3
DHX16
RP1-63M2.6
LCORL
PAPPA2
KRT18P9
RPRD2
SSTR5
EFEMP1
RREB1
AC011294.1
GH1
NPR3
PAH
BNC2
ANAPC10
COL27A1
TSBP1
ACAN
LRP1B
RABGAP1L
ELL
FER1L4

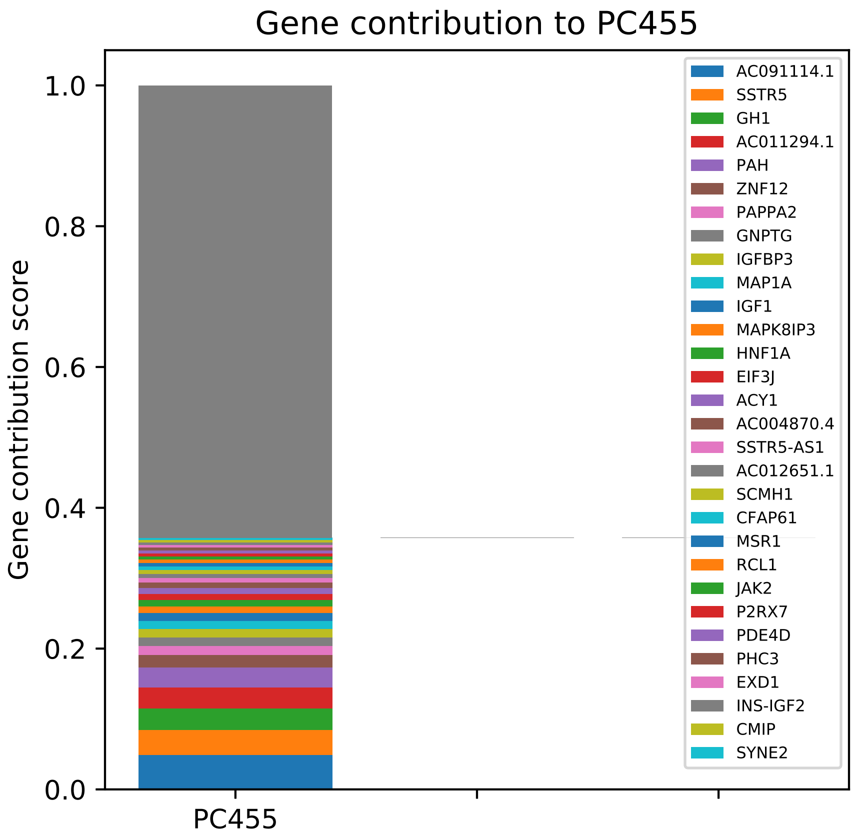

Gene contribution to PC455
Gene contribution score
1.0
0.8
0.6
0.4
0.2
0.0
PC455
AC091114.1
SSTR5
GH1
AC011294.1
PAH
ZNF12
PAPPA2
GNPTG
IGFBP3
MAP1A
IGF1
MAPK8IP3
HNF1A
EIF3J
ACY1
AC004870.4
SSTR5-AS1
AC012651.1
SCMH1
CFAP61
MSR1
RCL1
JAK2
P2RX7
PDE4D
PHC3
EXD1
INS-IGF2
CMIP
SYNE2

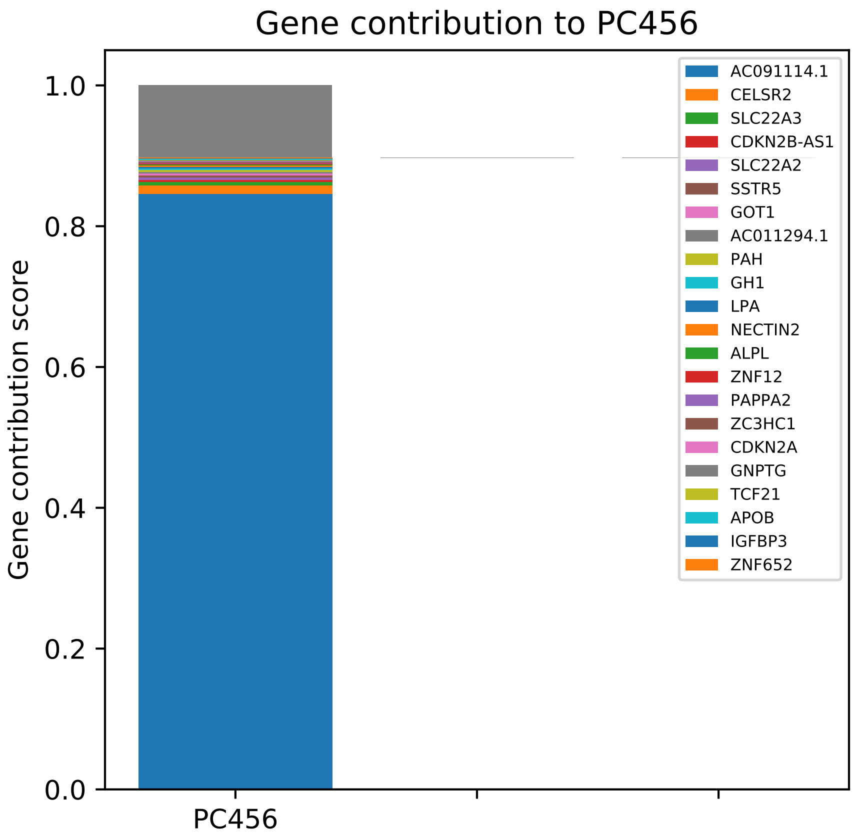

Gene contribution to PC456
Gene contribution score
1.0
0.8
0.6
0.4
0.2
0.0
PC456
AC091114.1
CELSR2
SLC22A3
CDKN2B-AS1
SLC22A2
SSTR5
GOT1
AC011294.1
PAH
GH1
LPA
NECTIN2
ALPL
ZNF12
PAPPA2
ZC3HC1
CDKN2A
GNPTG
TCF21
APOB
IGFBP3
ZNF652

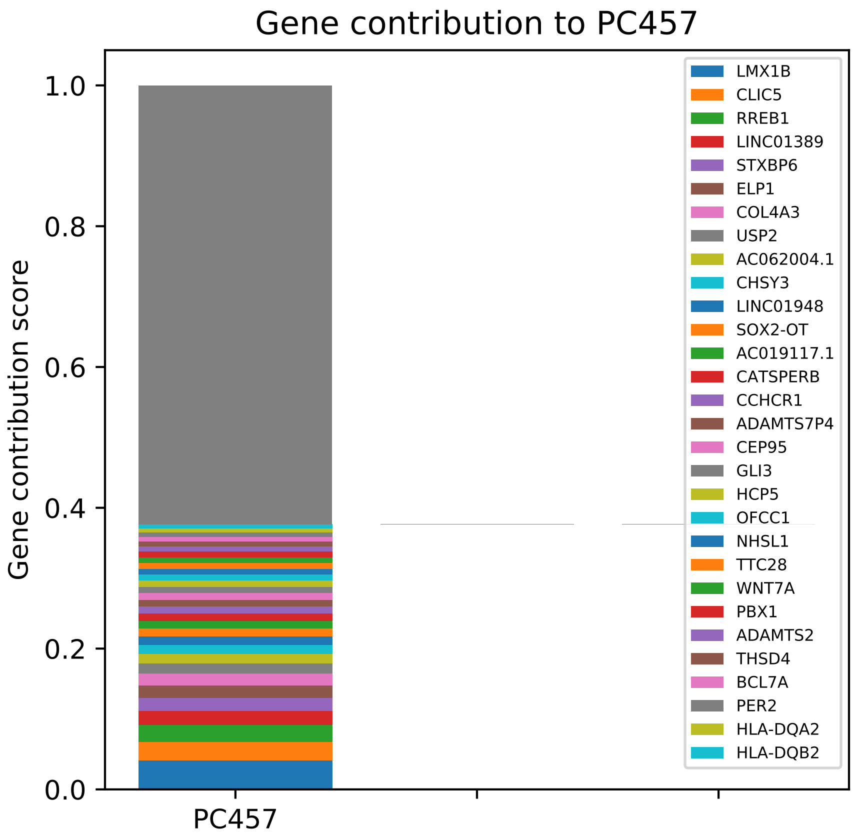
Gene contribution to PC457
Gene contribution score
1.0
0.8
0.6
0.4
0.2
0.0
PC457
LMX1B
CLIC5
RREB1
LINC01389
STXBP6
ELP1
COL4A3
USP2
AC062004.1
CHSY3
LINC01948
SOX2-OT
AC019117.1
CATSPERB
CCHCR1
ADAMTS7P4
CEP95
GLI3
HCP5
OFCC1
NHSL1
TTC28
WNT7A
PBX1
ADAMTS2
THSD4
BCL7A
PER2
HLA-DQA2
HLA-DQB2

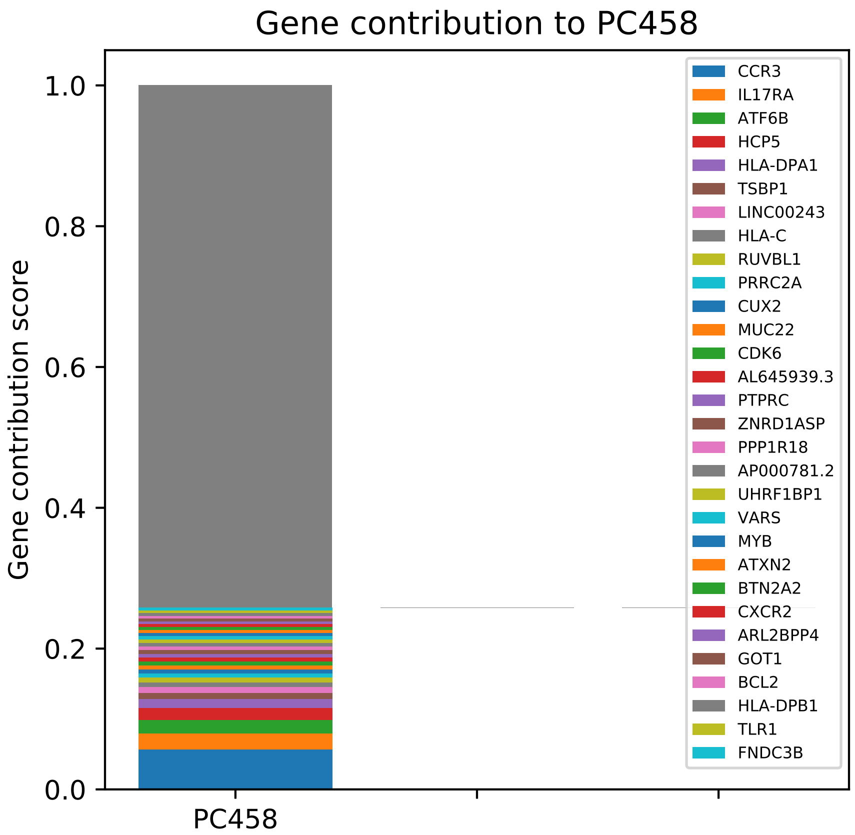
Gene contribution to PC458
Gene contribution score
0.0
0.2
0.4
0.6
0.8
1.0
PC458
CCR3
IL17RA
ATF6B
HCP5
HLA-DPA1
TSBP1
LINC00243
HLA-C
RUVBL1
PRRC2A
CUX2
MUC22
CDK6
AL645939.3
PTPRC
ZNRD1ASP
PPP1R18
AP000781.2
UHRF1BP1
VARS
MYB
ATXN2
BTN2A2
CXCR2
ARL2BPP4
GOT1
BCL2
HLA-DPB1
TLR1
FNDC3B

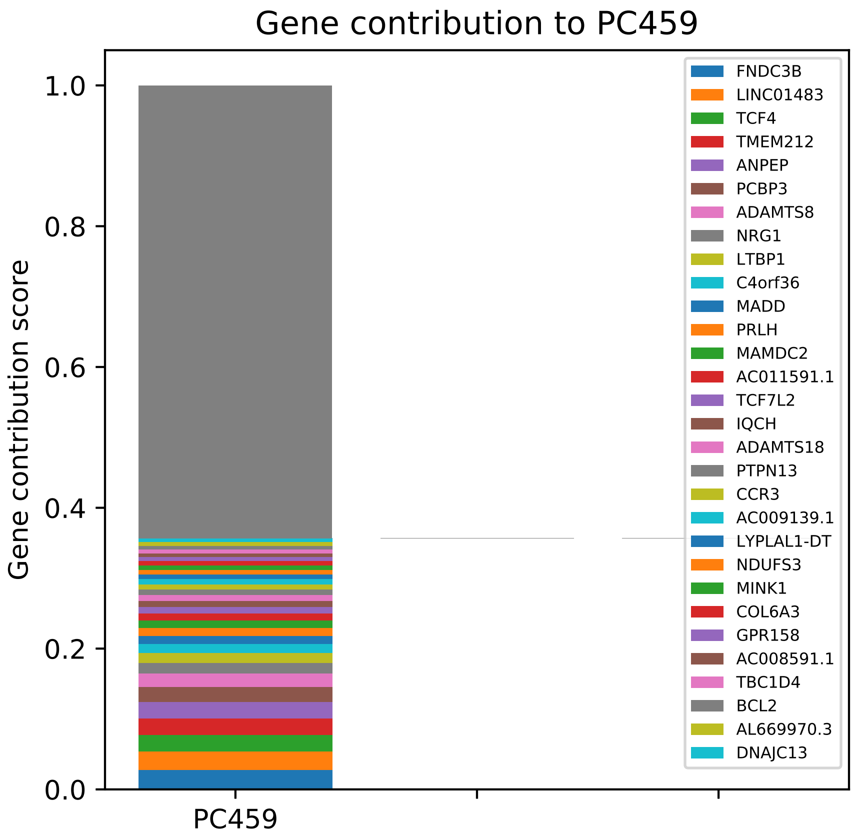
Gene contribution to PC459
Gene contribution score
1.0
0.8
0.6
0.4
0.2
0.0
PC459
FNDC3B
LINC01483
TCF4
TMEM212
ANPEP
PCBP3
ADAMTS8
NRG1
LTBP1
C4orf36
MADD
PRLH
MAMDC2
AC011591.1
TCF7L2
IQCH
ADAMTS18
PTPN13
CCR3
AC009139.1
LYPLAL1-DT
NDUFS3
MINK1
COL6A3
GPR158
AC008591.1
TBC1D4
BCL2
AL669970.3
DNAJC13

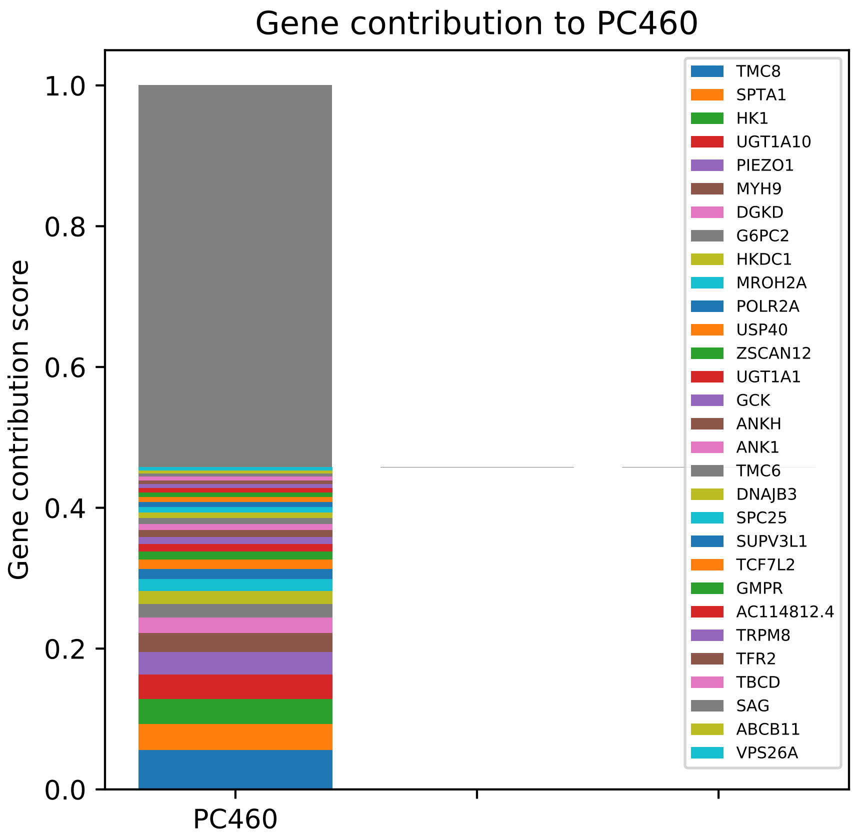

Gene contribution to PC460
Gene contribution score
1.0
0.8
0.6
0.4
0.2
0.0
PC460
TMC8
SPTA1
HK1
UGT1A10
PIEZO1
MYH9
DGKD
G6PC2
HKDC1
MROH2A
POLR2A
USP40
ZSCAN12
UGT1A1
GCK
ANKH
ANK1
TMC6
DNAJB3
SPC25
SUPV3L1
TCF7L2
GMPR
AC114812.4
TRPM8
TFR2
TBCD
SAG
ABCB11
VPS26A

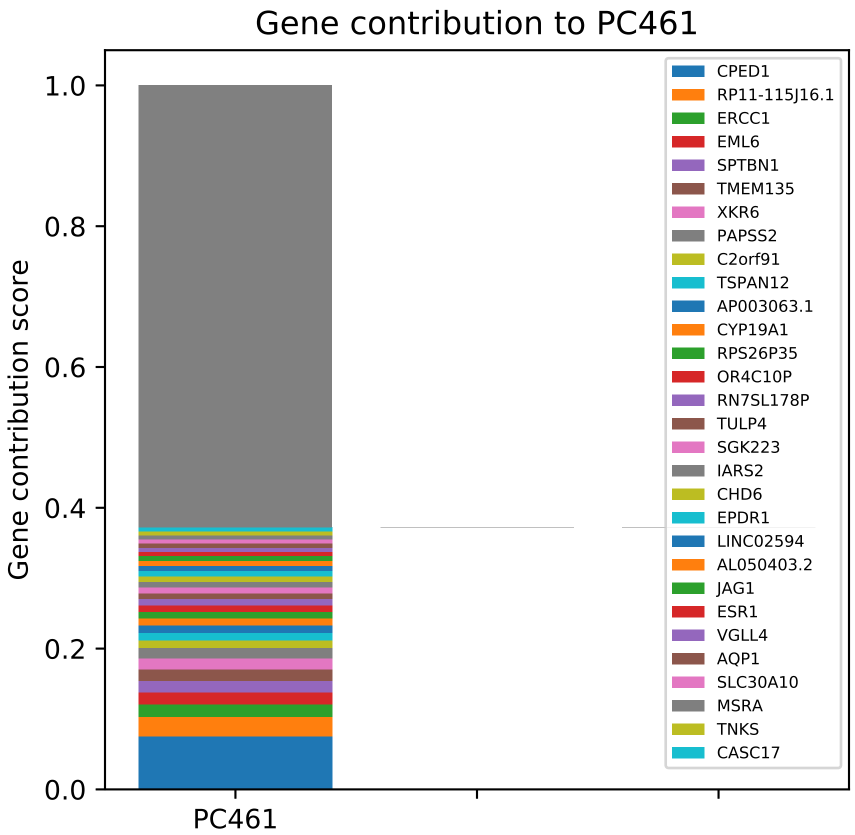

Gene contribution to PC461
Gene contribution score
1.0
0.8
0.6
0.4
0.2
0.0
PC461
CPED1
RP11-115J16.1
ERCC1
EML6
SPTBN1
TMEM135
XKR6
PAPSS2
C2orf91
TSPAN12
AP003063.1
CYP19A1
RPS26P35
OR4C10P
RN7SL178P
TULP4
SGK223
IARS2
CHD6
EPDR1
LINC02594
AL050403.2
JAG1
ESR1
VGLL4
AQP1
SLC30A10
MSRA
TNKS
CASC17

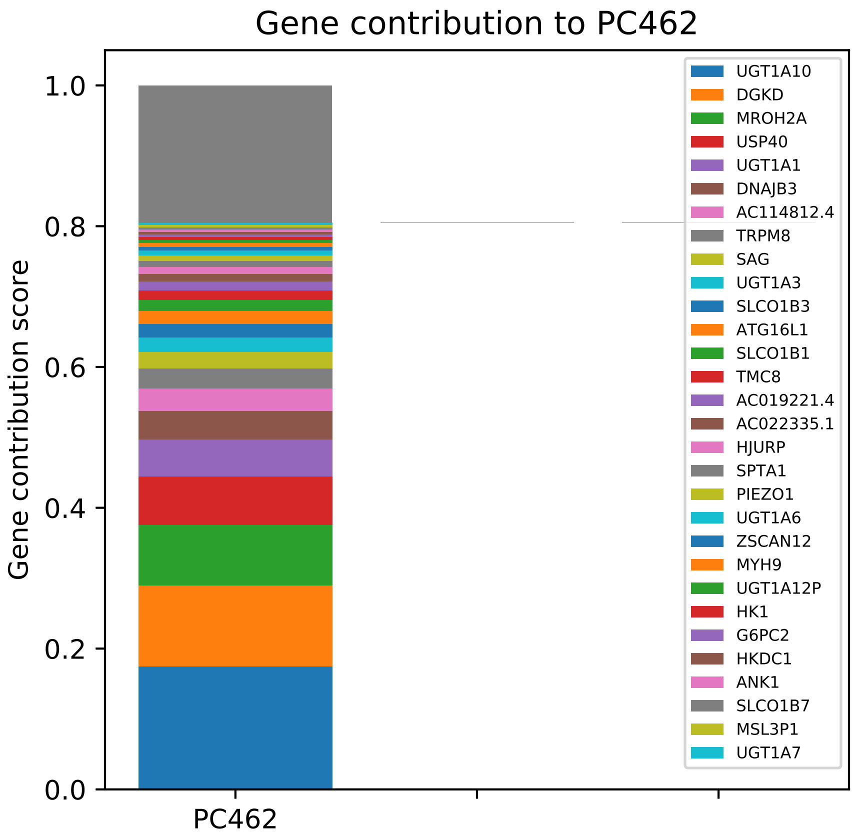

Gene contribution to PC462
Gene contribution score
1.0
0.8
0.6
0.4
0.2
0.0
PC462
UGT1A10
DGKD
MROH2A
USP40
UGT1A1
DNAJB3
AC114812.4
TRPM8
SAG
UGT1A3
SLCO1B3
ATG16L1
SLCO1B1
TMC8
AC019221.4
AC022335.1
HJURP
SPTA1
PIEZO1
UGT1A6
ZSCAN12
MYH9
UGT1A12P
HK1
G6PC2
HKDC1
ANK1
SLCO1B7
MSL3P1
UGT1A7

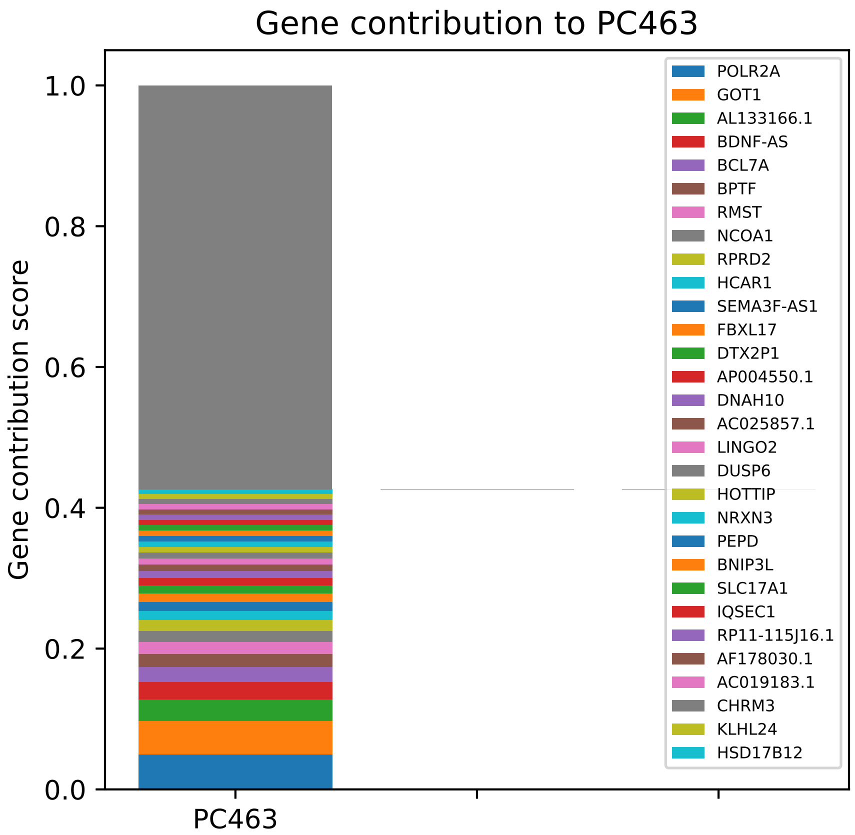

Gene contribution to PC463
Gene contribution score
1.0
0.8
0.6
0.4
0.2
0.0
PC463
POLR2A
GOT1
AL133166.1
BDNF-AS
BCL7A
BPTF
RMST
NCOA1
RPRD2
HCAR1
SEMA3F-AS1
FBXL17
DTX2P1
AP004550.1
DNAH10
AC025857.1
LINGO2
DUSP6
HOTTIP
NRXN3
PEPD
BNIP3L
SLC17A1
IQSEC1
RP11-115J16.1
AF178030.1
AC019183.1
CHRM3
KLHL24
HSD17B12

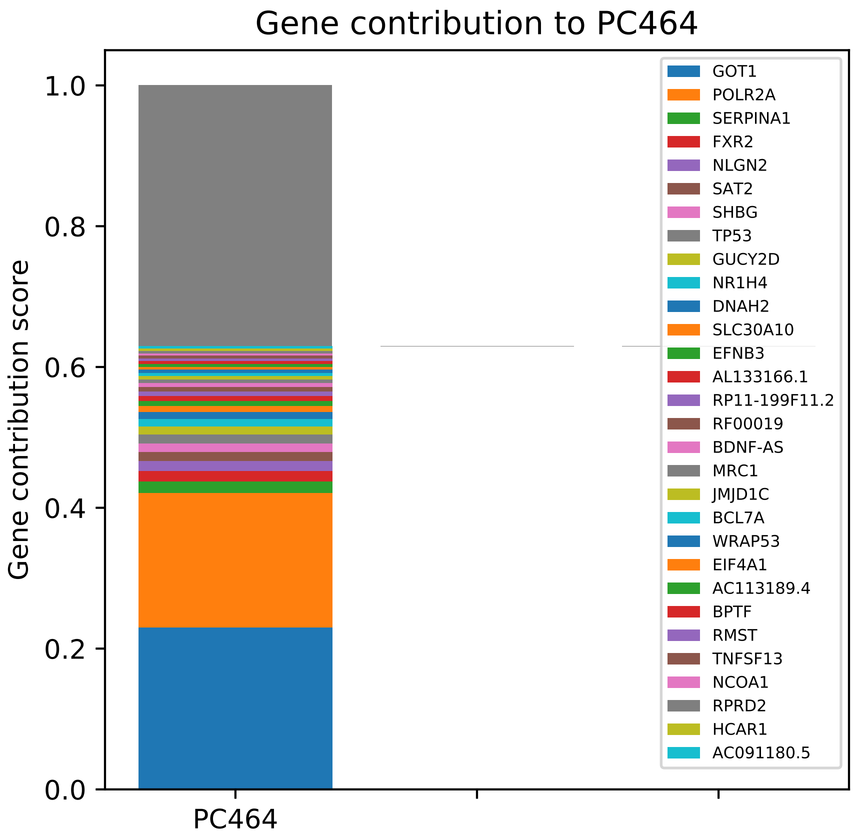

Gene contribution to PC464
Gene contribution score
0.0
0.2
0.4
0.6
0.8
1.0
PC464
GOT1
POLR2A
SERPINA1
FXR2
NLGN2
SAT2
SHBG
TP53
GUCY2D
NR1H4
DNAH2
SLC30A10
EFNB3
AL133166.1
RP11-199F11.2
RF00019
BDNF-AS
MRC1
JMJD1C
BCL7A
WRAP53
EIF4A1
AC113189.4
BPTF
RMST
TNFSF13
NCOA1
RPRD2
HCAR1
AC091180.5

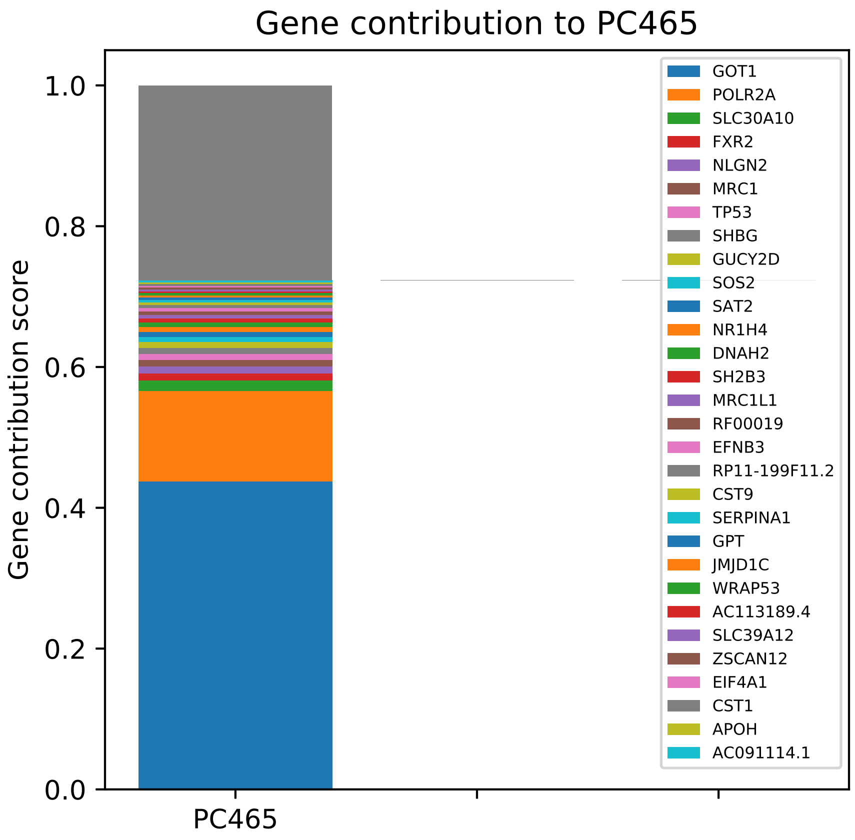

Gene contribution to PC465
Gene contribution score
1.0
0.8
0.6
0.4
0.2
0.0
PC465
GOT1
POLR2A
SLC30A10
FXR2
NLGN2
MRC1
TP53
SHBG
GUCY2D
SOS2
SAT2
NR1H4
DNAH2
SH2B3
MRC1L1
RF00019
EFNB3
RP11-199F11.2
CST9
SERPINA1
GPT
JMJD1C
WRAP53
AC113189.4
SLC39A12
ZSCAN12
EIF4A1
CST1
APOH
AC091114.1

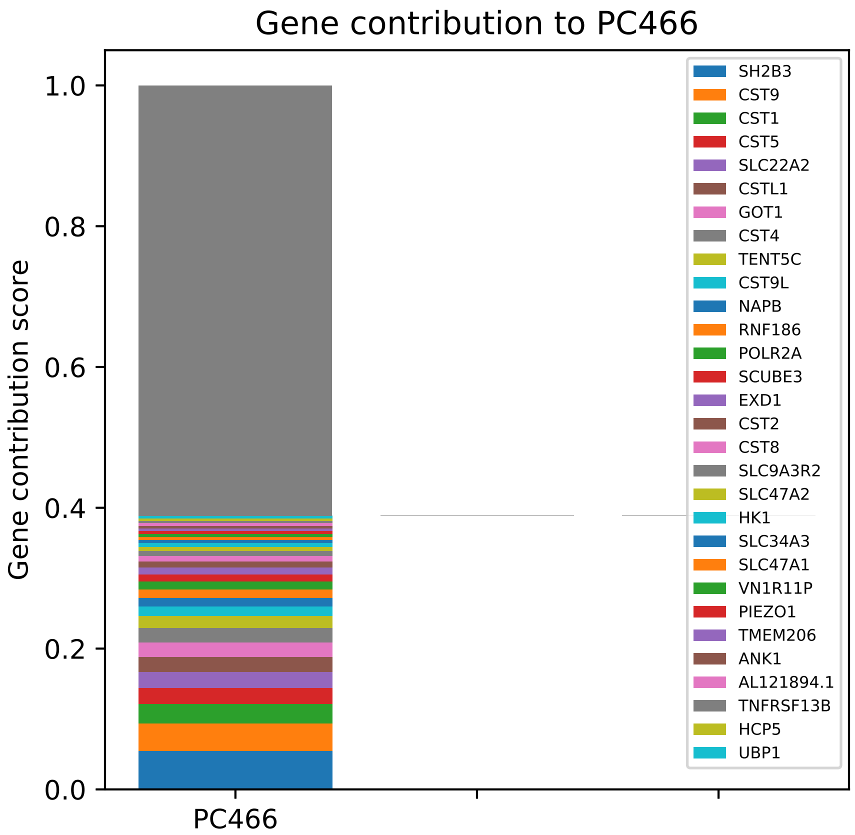
Gene contribution to PC466
Gene contribution score
1.0
0.8
0.6
0.4
0.2
0.0
PC466
SH2B3
CST9
CST1
CST5
SLC22A2
CSTL1
GOT1
CST4
TENT5C
CST9L
NAPB
RNF186
POLR2A
SCUBE3
EXD1
CST2
CST8
SLC9A3R2
SLC47A2
HK1
SLC34A3
SLC47A1
VN1R11P
PIEZO1
TMEM206
ANK1
AL121894.1
TNFRSF13B
HCP5
UBP1

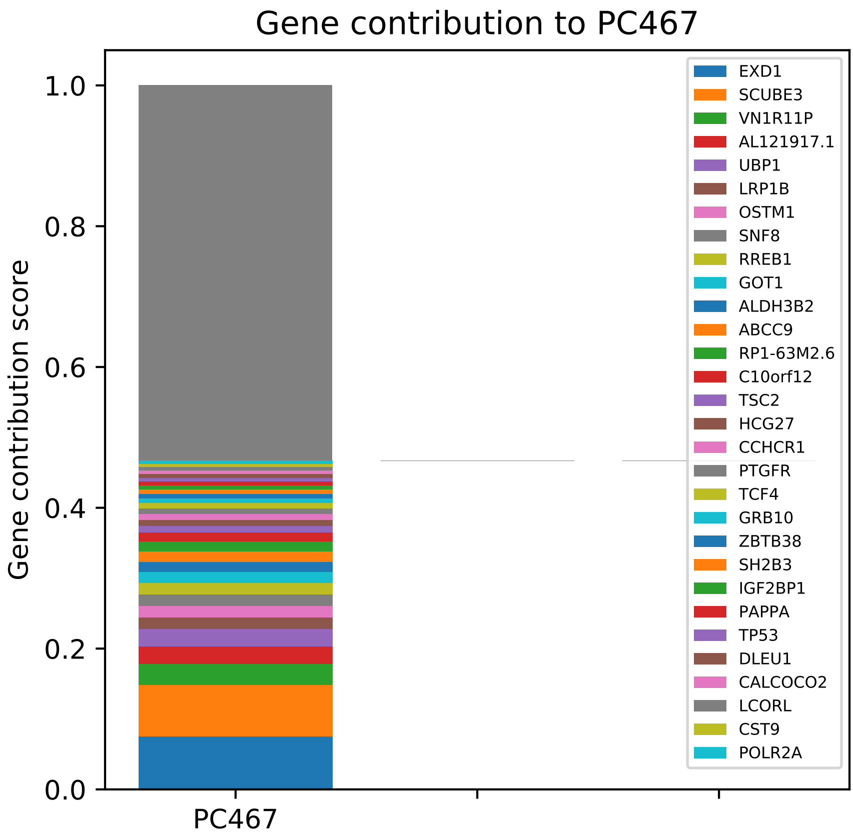

Gene contribution to PC467
Gene contribution score
0.0
0.2
0.4
0.6
0.8
1.0
PC467
EXD1
SCUBE3
VN1R11P
AL121917.1
UBP1
LRP1B
OSTM1
SNF8
RREB1
GOT1
ALDH3B2
ABCC9
RP1-63M2.6
C10orf12
TSC2
HCG27
CCHCR1
PTGFR
TCF4
GRB10
ZBTB38
SH2B3
IGF2BP1
PAPPA
TP53
DLEU1
CALCOCO2
LCORL
CST9
POLR2A

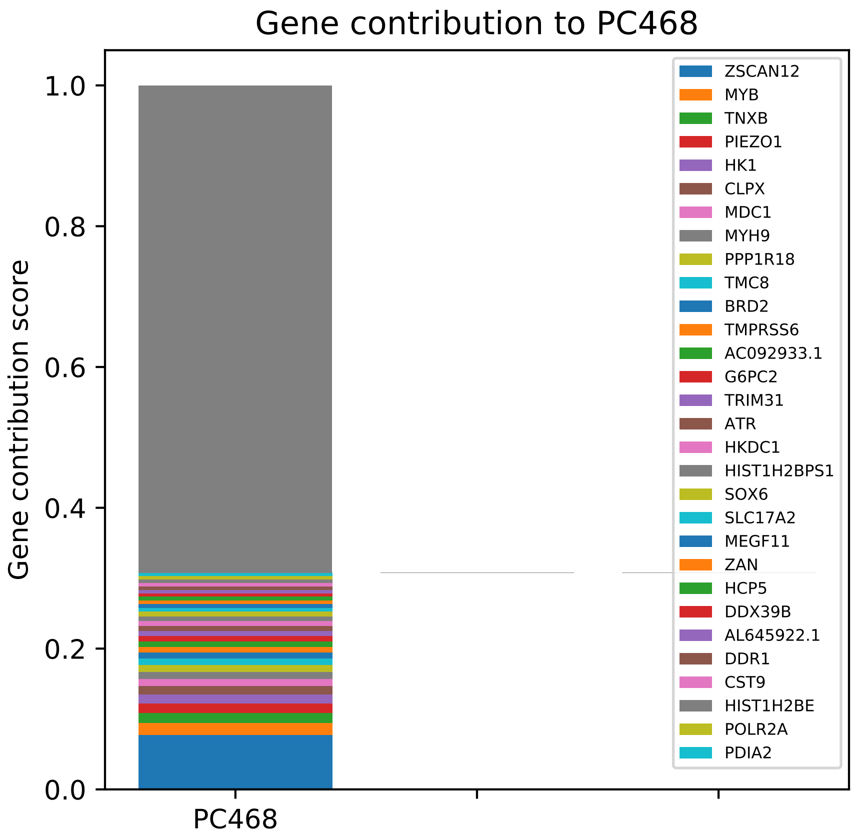

Gene contribution to PC468
Gene contribution score
1.0
0.8
0.6
0.4
0.2
0.0
PC468
ZSCAN12
MYB
TNXB
PIEZO1
HK1
CLPX
MDC1
MYH9
PPP1R18
TMC8
BRD2
TMPRSS6
AC092933.1
G6PC2
TRIM31
ATR
HKDC1
HIST1H2BPS1
SOX6
SLC17A2
MEGF11
ZAN
HCP5
DDX39B
AL645922.1
DDR1
CST9
HIST1H2BE
POLR2A
PDIA2

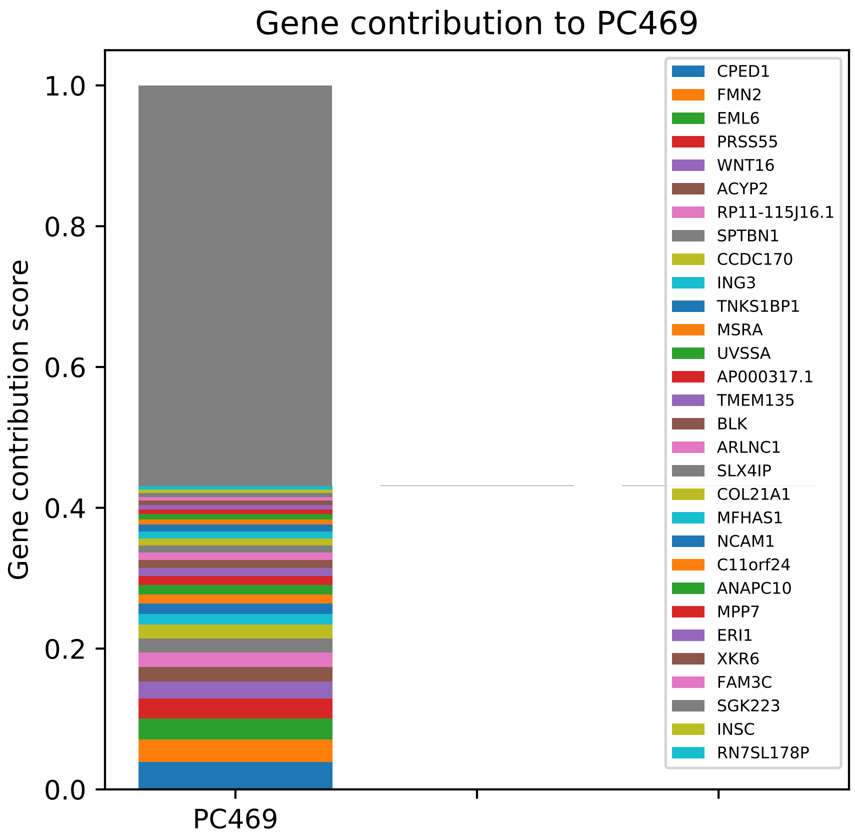

Gene contribution to PC469
Gene contribution score
1.0
0.8
0.6
0.4
0.2
0.0
PC469
CPED1
FMN2
EML6
PRSS55
WNT16
ACYP2
RP11-115J16.1
SPTBN1
CCDC170
ING3
TNKS1BP1
MSRA
UVSSA
AP000317.1
TMEM135
BLK
ARLNC1
SLX4IP
COL21A1
MFHAS1
NCAM1
C11orf24
ANAPC10
MPP7
ERI1
XKR6
FAM3C
SGK223
INSC
RN7SL178P

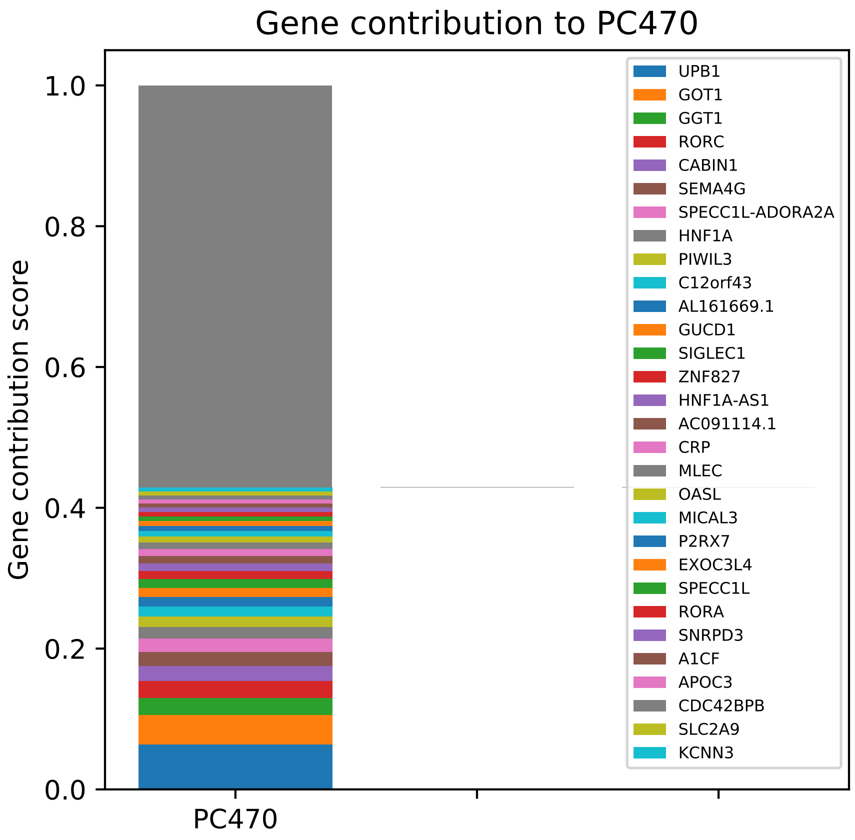

Gene contribution to PC470
Gene contribution score
1.0
0.8
0.6
0.4
0.2
0.0
PC470
UPB1
GOT1
GGT1
RORC
CABIN1
SEMA4G
SPECC1L-ADORA2A
HNF1A
PIWIL3
C12orf43
AL161669.1
GUCD1
SIGLEC1
ZNF827
HNF1A-AS1
AC091114.1
CRP
MLEC
OASL
MICAL3
P2RX7
EXOC3L4
SPECC1L
RORA
SNRPD3
A1CF
APOC3
CDC42BPB
SLC2A9
KCNN3

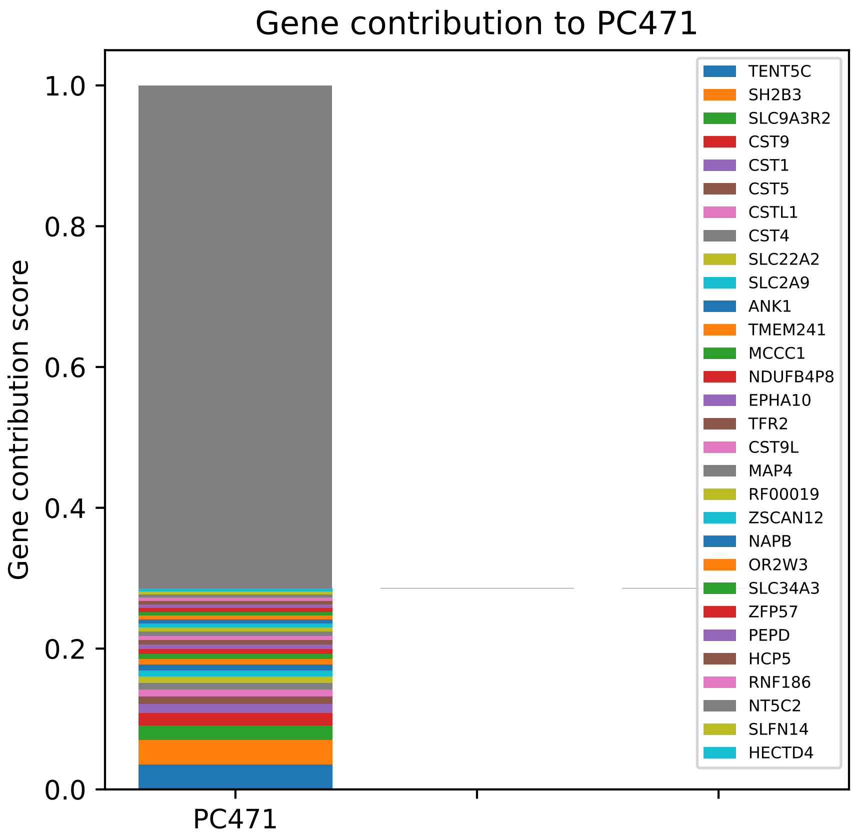

Gene contribution to PC471
Gene contribution score
1.0
0.8
0.6
0.4
0.2
0.0
PC471
TENT5C
SH2B3
SLC9A3R2
CST9
CST1
CST5
CSTL1
CST4
SLC22A2
SLC2A9
ANK1
TMEM241
MCCC1
NDUFB4P8
EPHA10
TFR2
CST9L
MAP4
RF00019
ZSCAN12
NAPB
OR2W3
SLC34A3
ZFP57
PEPD
HCP5
RNF186
NT5C2
SLFN14
HECTD4

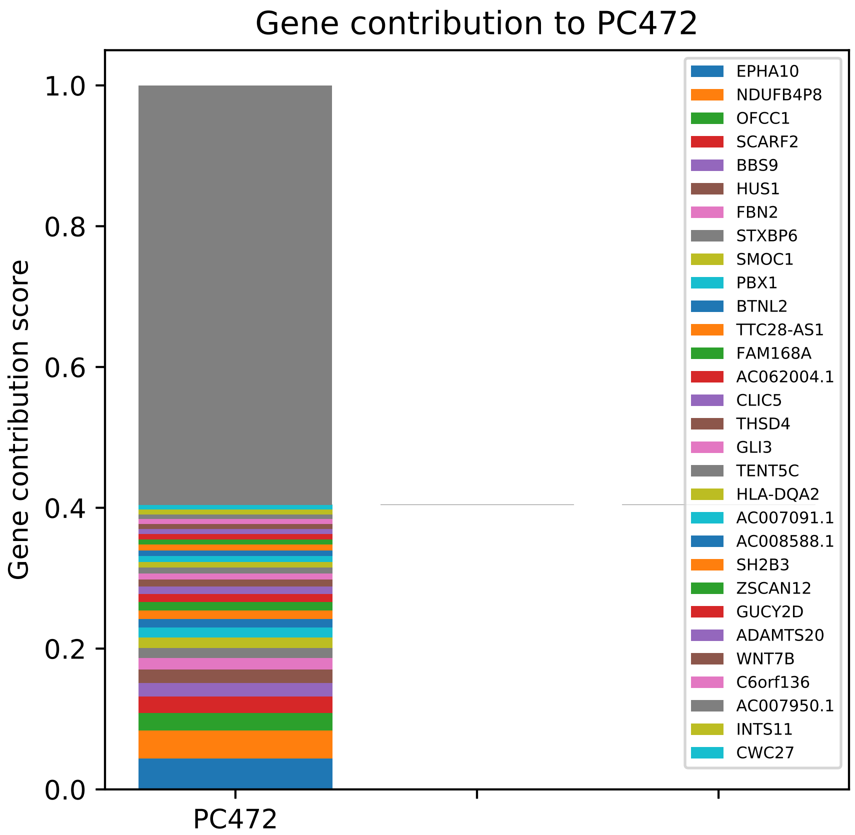

Gene contribution to PC472
Gene contribution score
1.0
0.8
0.6
0.4
0.2
0.0
PC472
EPHA10
NDUFB4P8
OFCC1
SCARF2
BBS9
HUS1
FBN2
STXBP6
SMOC1
PBX1
BTNL2
TTC28-AS1
FAM168A
AC062004.1
CLIC5
THSD4
GLI3
TENT5C
HLA-DQA2
AC007091.1
AC008588.1
SH2B3
ZSCAN12
GUCY2D
ADAMTS20
WNT7B
C6orf136
AC007950.1
INTS11
CWC27

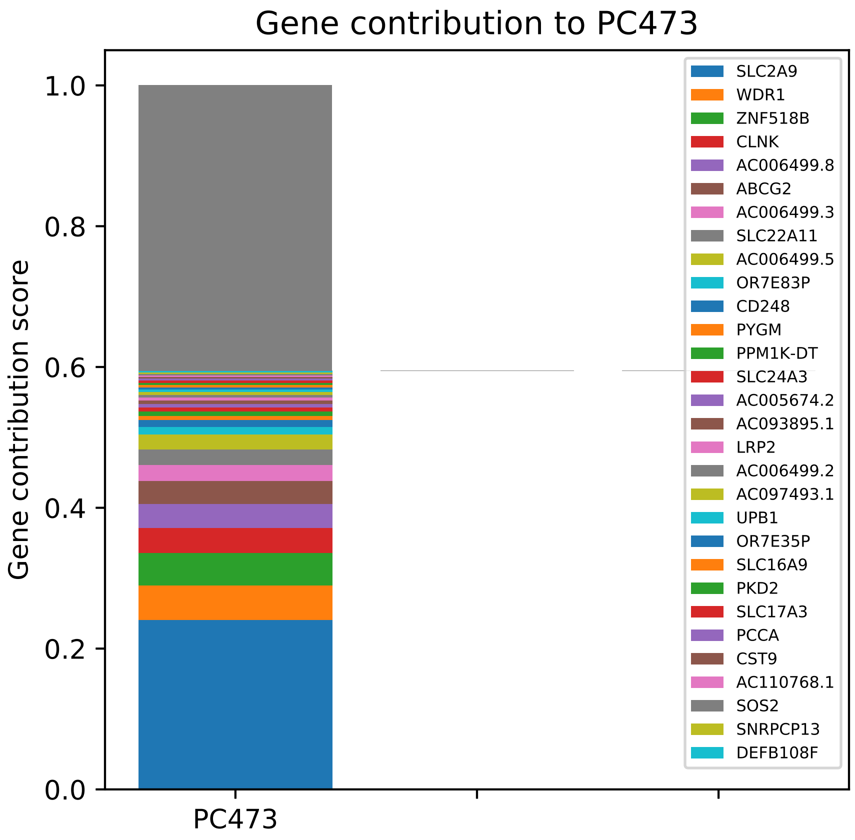

Gene contribution to PC473
Gene contribution score
1.0
0.8
0.6
0.4
0.2
0.0
PC473
SLC2A9
WDR1
ZNF518B
CLNK
AC006499.8
ABCG2
AC006499.3
SLC22A11
AC006499.5
OR7E83P
CD248
PYGM
PPM1K-DT
SLC24A3
AC005674.2
AC093895.1
LRP2
AC006499.2
AC097493.1
UPB1
OR7E35P
SLC16A9
PKD2
SLC17A3
PCCA
CST9
AC110768.1
SOS2
SNRPCP13
DEFB108F

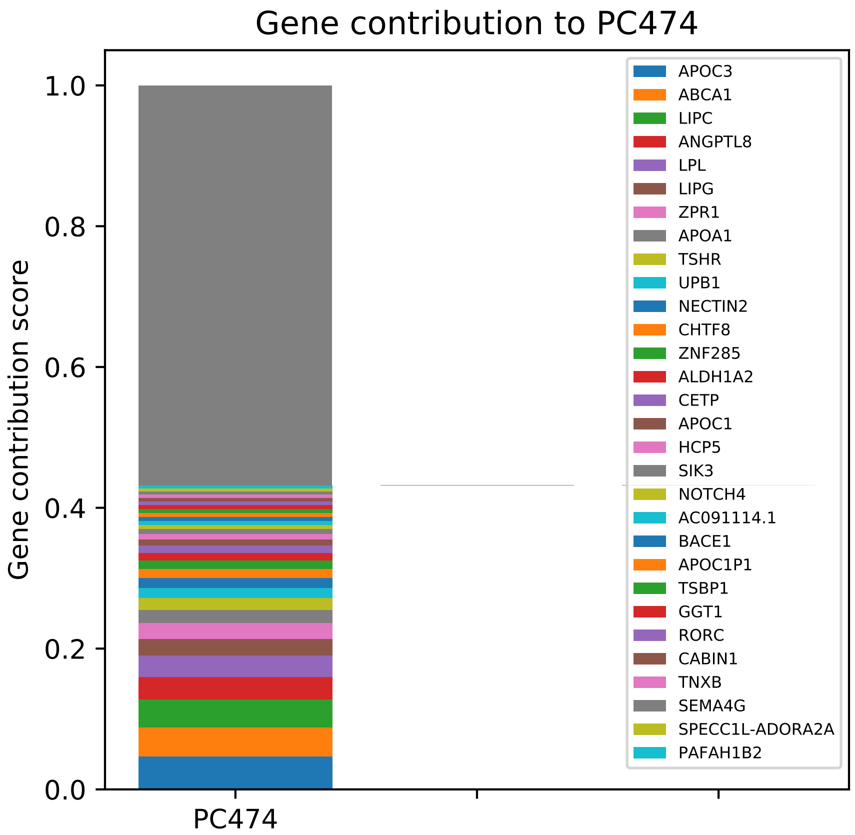

Gene contribution to PC474
Gene contribution score
1.0
0.8
0.6
0.4
0.2
0.0
PC474
APOC3
ABCA1
LIPC
ANGPTL8
LPL
LIPG
ZPR1
APOA1
TSHR
UPB1
NECTIN2
CHTF8
ZNF285
ALDH1A2
CETP
APOC1
HCP5
SIK3
NOTCH4
AC091114.1
BACE1
APOC1P1
TSBP1
GGT1
RORC
CABIN1
TNXB
SEMA4G
SPECC1L-ADORA2A
PAFAH1B2

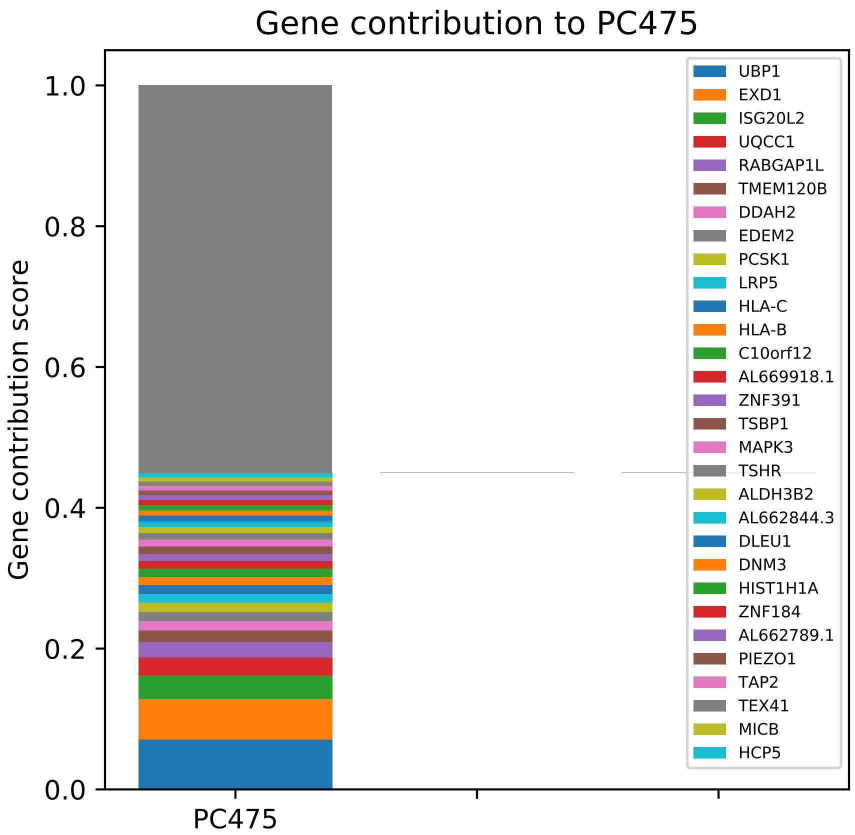
Gene contribution to PC475
Gene contribution score
0.0
0.2
0.4
0.6
0.8
1.0
PC475
UBP1
EXD1
ISG20L2
UQCC1
RABGAP1L
TMEM120B
DDAH2
EDEM2
PCSK1
LRP5
HLA-C
HLA-B
C10orf12
AL669918.1
ZNF391
TSBP1
MAPK3
TSHR
ALDH3B2
AL662844.3
DLEU1
DNM3
HIST1H1A
ZNF184
AL662789.1
PIEZO1
TAP2
TEX41
MICB
HCP5

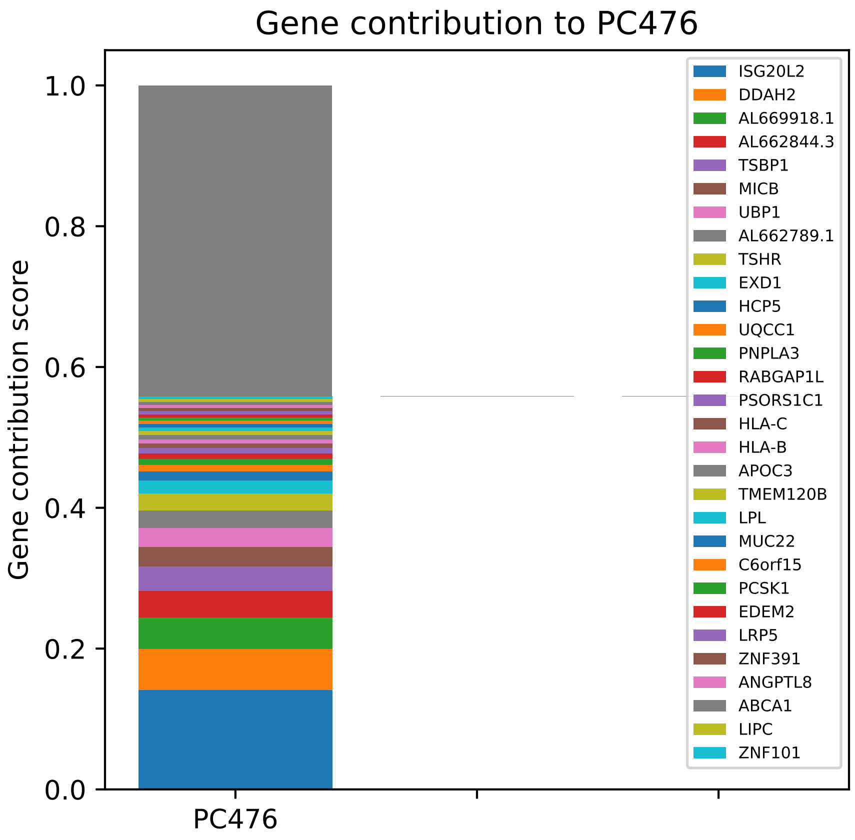
Gene contribution to PC476
Gene contribution score
1.0
0.8
0.6
0.4
0.2
0.0
PC476
ISG20L2
DDAH2
AL669918.1
AL662844.3
TSBP1
MICB
UBP1
AL662789.1
TSHR
EXD1
HCP5
UQCC1
PNPLA3
RABGAP1L
PSORS1C1
HLA-C
HLA-B
APOC3
TMEM120B
LPL
MUC22
C6orf15
PCSK1
EDEM2
LRP5
ZNF391
ANGPTL8
ABCA1
LIPC
ZNF101

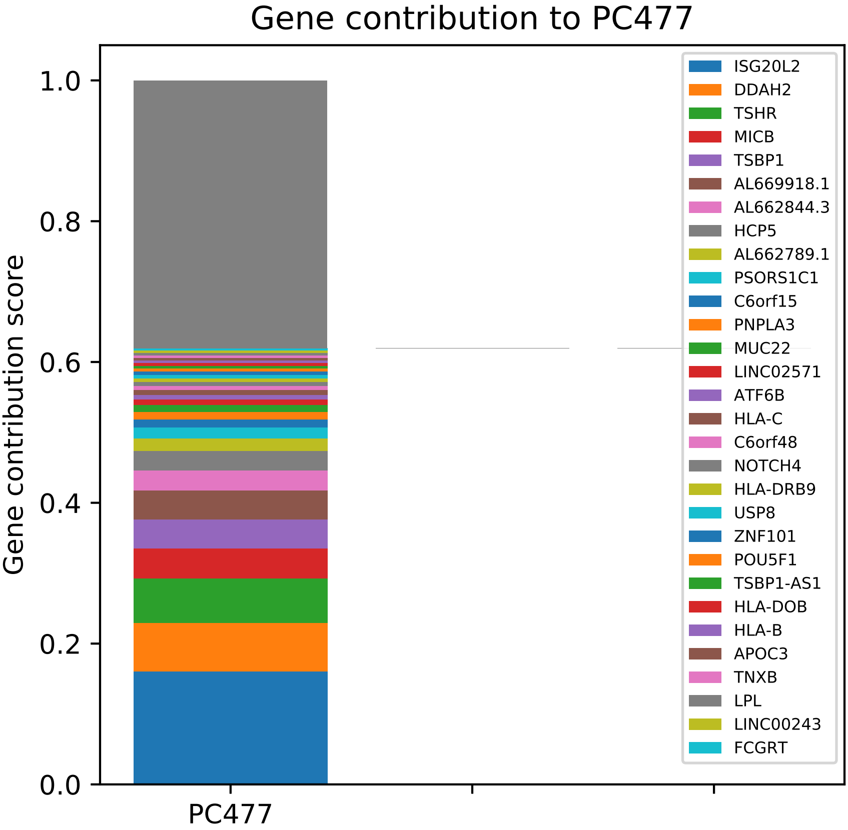
Gene contribution to PC477
Gene contribution score
1.0
0.8
0.6
0.4
0.2
0.0
PC477
ISG20L2
DDAH2
TSHR
MICB
TSBP1
AL669918.1
AL662844.3
HCP5
AL662789.1
PSORS1C1
C6orf15
PNPLA3
MUC22
LINC02571
ATF6B
HLA-C
C6orf48
NOTCH4
HLA-DRB9
USP8
ZNF101
POU5F1
TSBP1-AS1
HLA-DOB
HLA-B
APOC3
TNXB
LPL
LINC00243
FCGRT

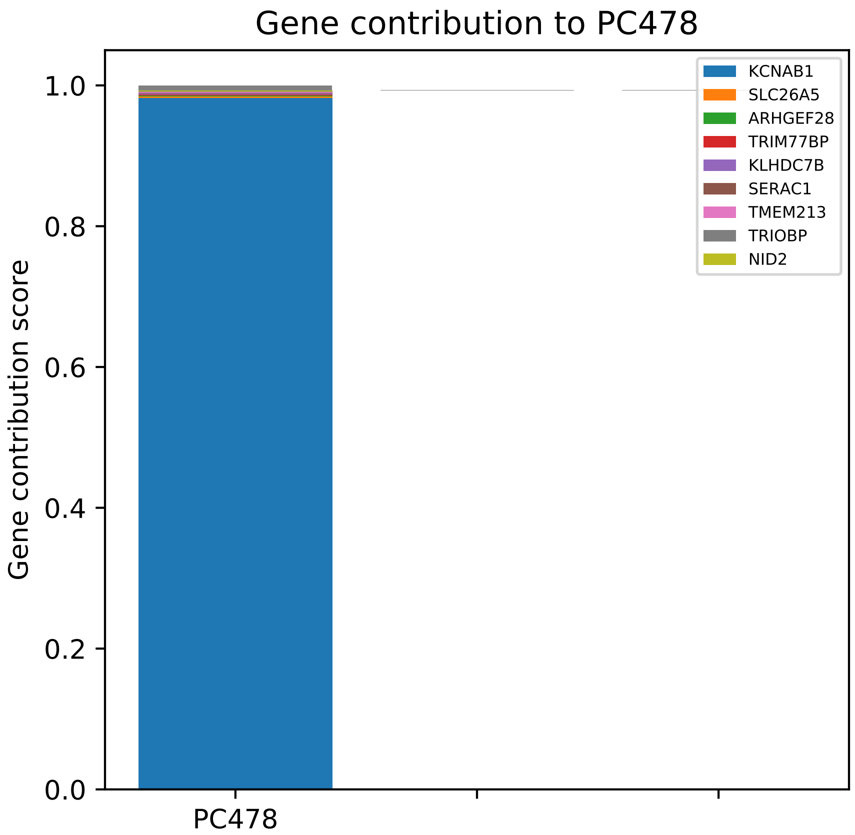

Gene contribution to PC478
Gene contribution score
1.0
0.8
0.6
0.4
0.2
0.0
PC478
KCNAB1
SLC26A5
ARHGEF28
TRIM77BP
KLHDC7B
SERAC1
TMEM213
TRIOBP
NID2

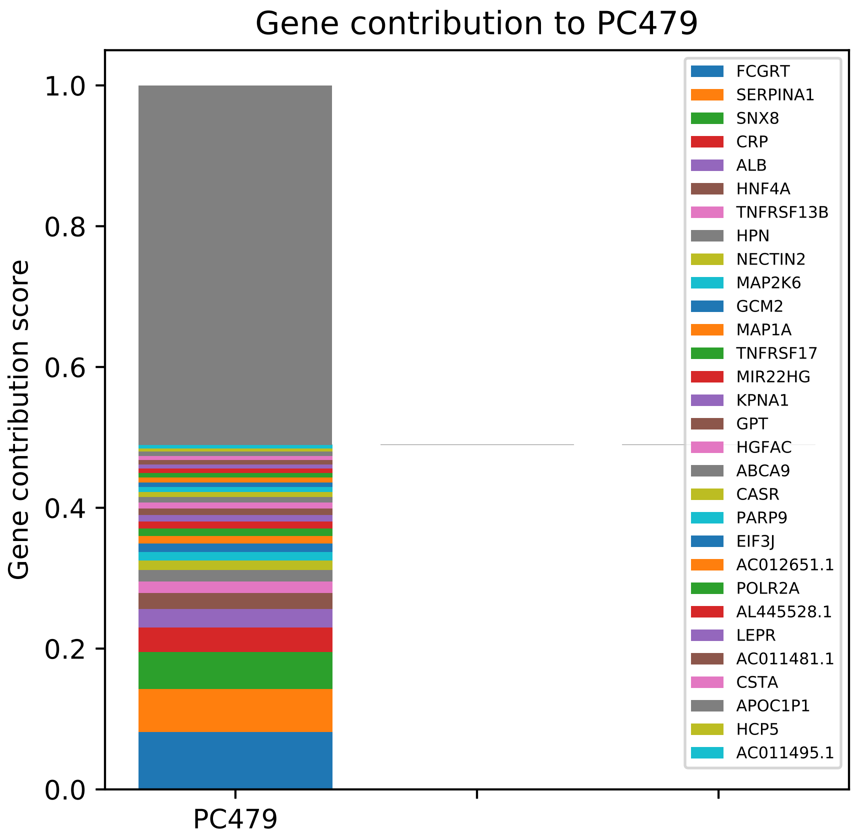

Gene contribution to PC479
Gene contribution score
1.0
0.8
0.6
0.4
0.2
0.0
PC479
FCGRT
SERPINA1
SNX8
CRP
ALB
HNF4A
TNFRSF13B
HPN
NECTIN2
MAP2K6
GCM2
MAP1A
TNFRSF17
MIR22HG
KPNA1
GPT
HGFAC
ABCA9
CASR
PARP9
EIF3J
AC012651.1
POLR2A
AL445528.1
LEPR
AC011481.1
CSTA
APOC1P1
HCP5
AC011495.1

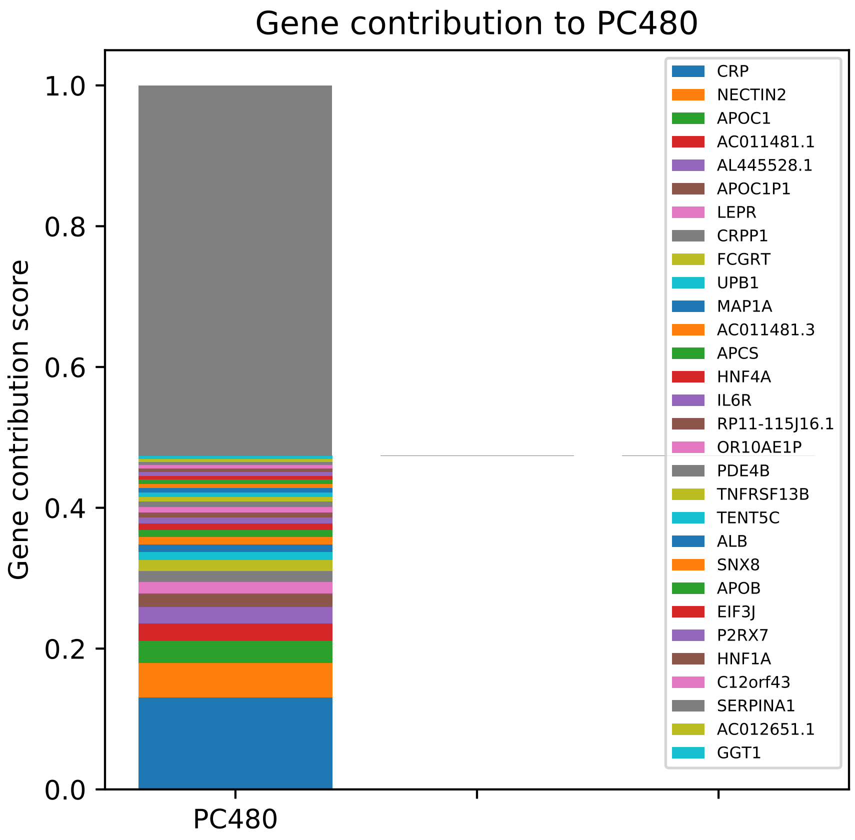
Gene contribution to PC480
Gene contribution score
1.0
0.8
0.6
0.4
0.2
0.0
PC480
CRP
NECTIN2
APOC1
AC011481.1
AL445528.1
APOC1P1
LEPR
CRPP1
FCGRT
UPB1
MAP1A
AC011481.3
APCS
HNF4A
IL6R
RP11-115J16.1
OR10AE1P
PDE4B
TNFRSF13B
TENT5C
ALB
SNX8
APOB
EIF3J
P2RX7
HNF1A
C12orf43
SERPINA1
AC012651.1
GGT1

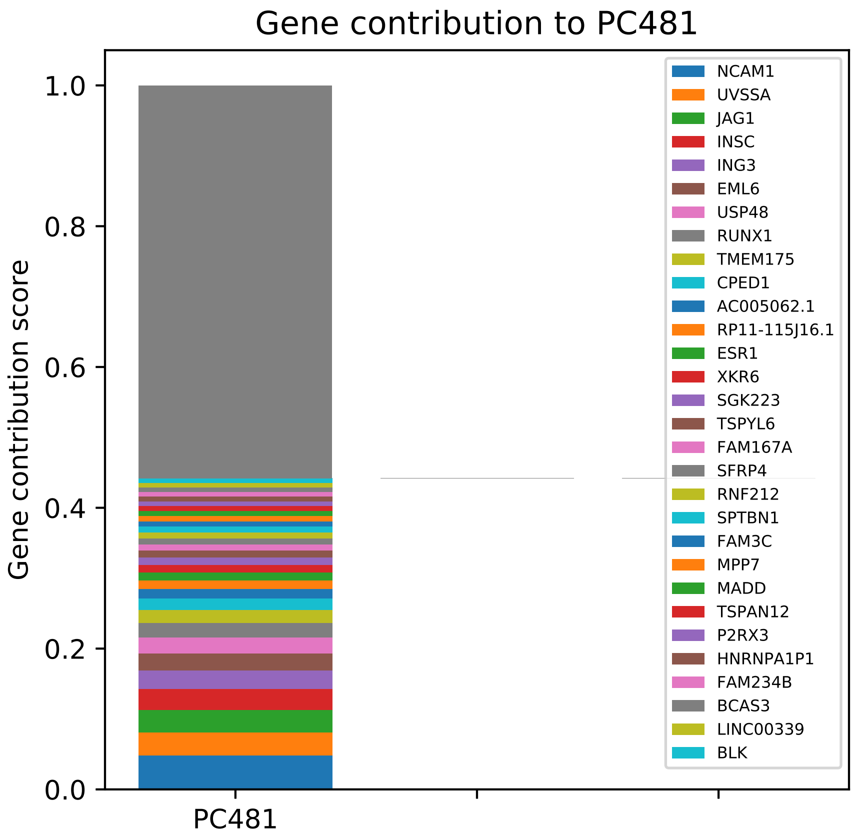

Gene contribution to PC481
Gene contribution score
1.0
0.8
0.6
0.4
0.2
0.0
PC481
NCAM1
UVSSA
JAG1
INSC
ING3
EML6
USP48
RUNX1
TMEM175
CPED1
AC005062.1
RP11-115J16.1
ESR1
XKR6
SGK223
TSPYL6
FAM167A
SFRP4
RNF212
SPTBN1
FAM3C
MPP7
MADD
TSPAN12
P2RX3
HNRNPA1P1
FAM234B
BCAS3
LINC00339
BLK

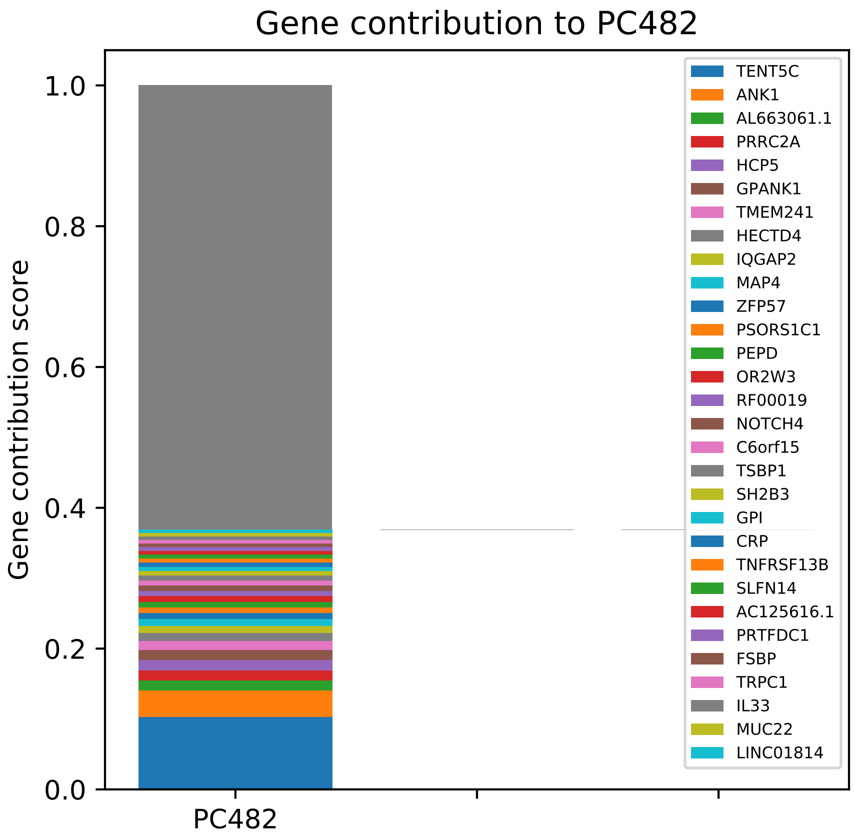
Gene contribution to PC482
Gene contribution score
1.0
0.8
0.6
0.4
0.2
0.0
PC482
TENT5C
ANK1
AL663061.1
PRRC2A
HCP5
GPANK1
TMEM241
HECTD4
IQGAP2
MAP4
ZFP57
PSORS1C1
PEPD
OR2W3
RF00019
NOTCH4
C6orf15
TSBP1
SH2B3
GPI
CRP
TNFRSF13B
SLFN14
AC125616.1
PRTFDC1
FSBP
TRPC1
IL33
MUC22
LINC01814

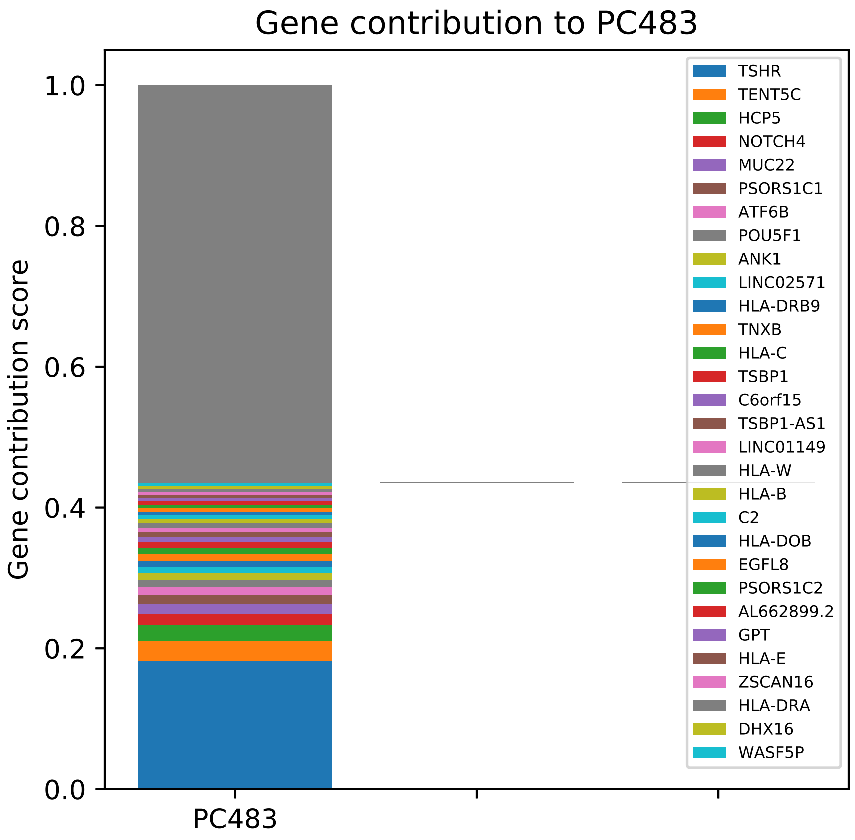
Gene contribution to PC483
Gene contribution score
0.0
0.2
0.4
0.6
0.8
1.0
PC483
TSHR
TENT5C
HCP5
NOTCH4
MUC22
PSORS1C1
ATF6B
POU5F1
ANK1
LINC02571
HLA-DRB9
TNXB
HLA-C
TSBP1
C6orf15
TSBP1-AS1
LINC01149
HLA-W
HLA-B
C2
HLA-DOB
EGFL8
PSORS1C2
AL662899.2
GPT
HLA-E
ZSCAN16
HLA-DRA
DHX16
WASF5P

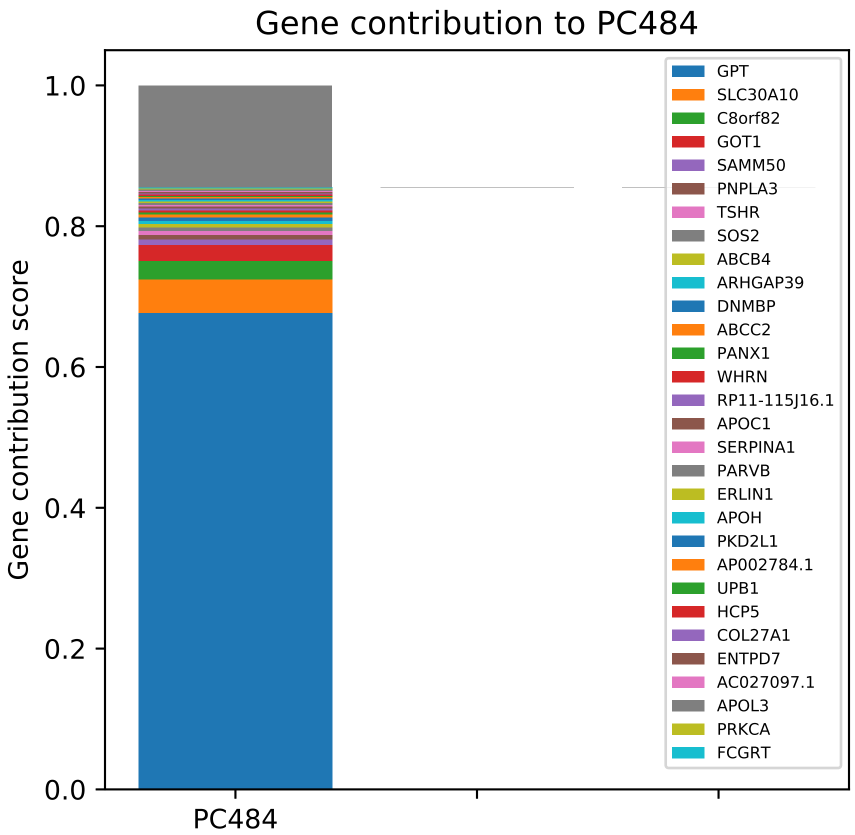

Gene contribution to PC484
Gene contribution score
0.0
0.2
0.4
0.6
0.8
1.0
PC484
GPT
SLC30A10
C8orf82
GOT1
SAMM50
PNPLA3
TSHR
SOS2
ABCB4
ARHGAP39
DNMBP
ABCC2
PANX1
WHRN
RP11-115J16.1
APOC1
SERPINA1
PARVB
ERLIN1
APOH
PKD2L1
AP002784.1
UPB1
HCP5
COL27A1
ENTPD7
AC027097.1
APOL3
PRKCA
FCGRT

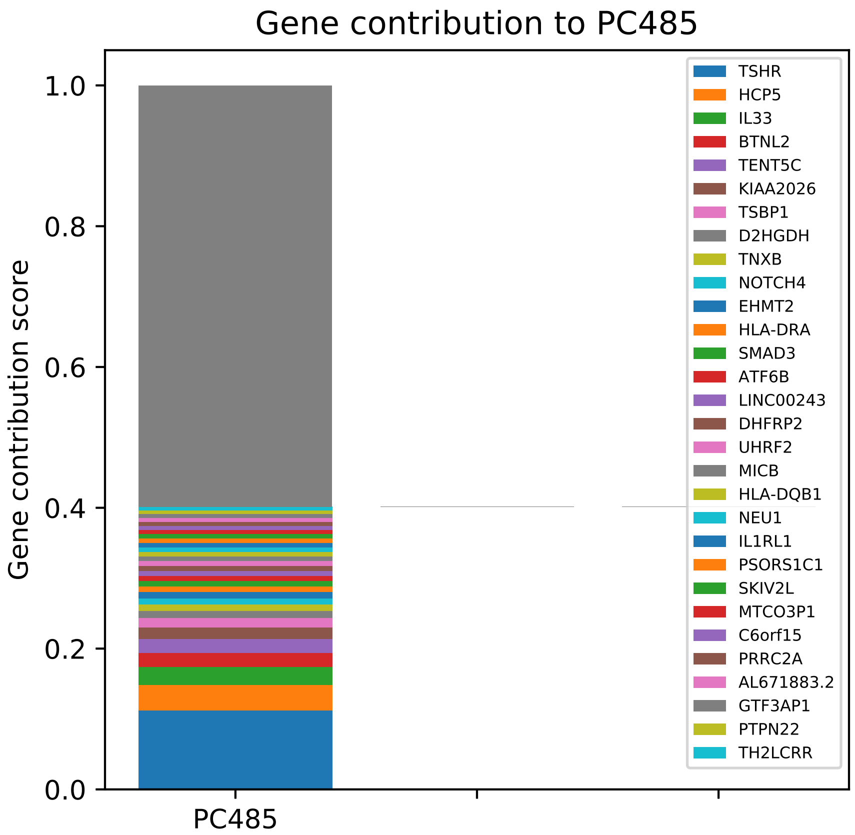
Gene contribution to PC485
Gene contribution score
1.0
0.8
0.6
0.4
0.2
0.0
PC485
TSHR
HCP5
IL33
BTNL2
TENT5C
KIAA2026
TSBP1
D2HGDH
TNXB
NOTCH4
EHMT2
HLA-DRA
SMAD3
ATF6B
LINC00243
DHFRP2
UHRF2
MICB
HLA-DQB1
NEU1
IL1RL1
PSORS1C1
SKIV2L
MTCO3P1
C6orf15
PRRC2A
AL671883.2
GTF3AP1
PTPN22
TH2LCRR

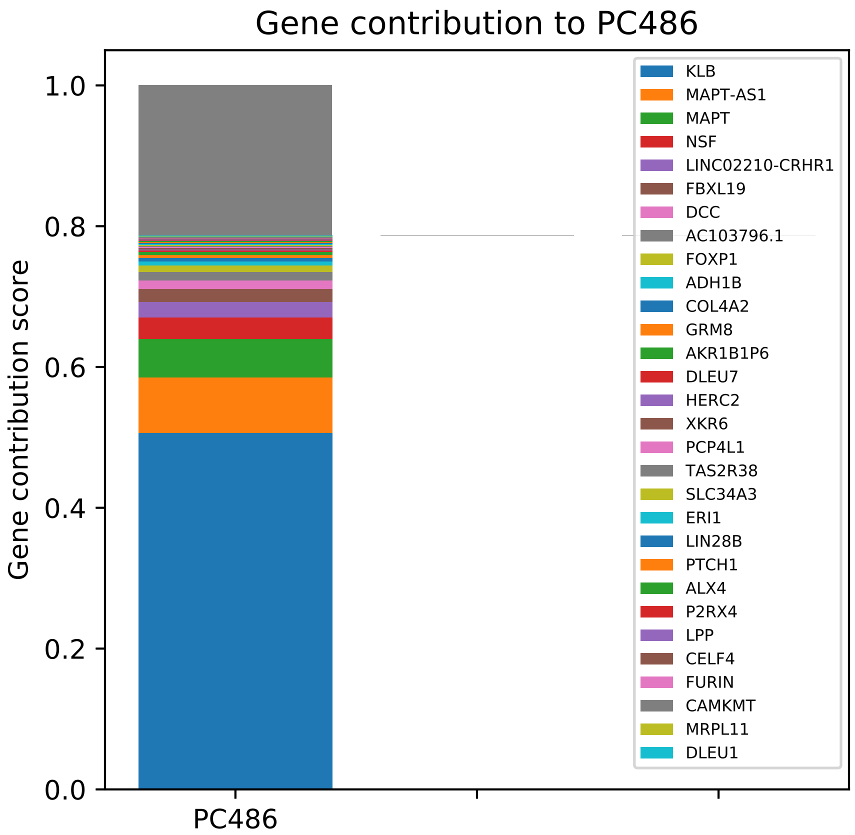

Gene contribution to PC486
Gene contribution score
1.0
0.8
0.6
0.4
0.2
0.0
PC486
KLB
MAPT-AS1
MAPT
NSF
LINC02210-CRHR1
FBXL19
DCC
AC103796.1
FOXP1
ADH1B
COL4A2
GRM8
AKR1B1P6
DLEU7
HERC2
XKR6
PCP4L1
TAS2R38
SLC34A3
ERI1
LIN28B
PTCH1
ALX4
P2RX4
LPP
CELF4
FURIN
CAMKMT
MRPL11
DLEU1

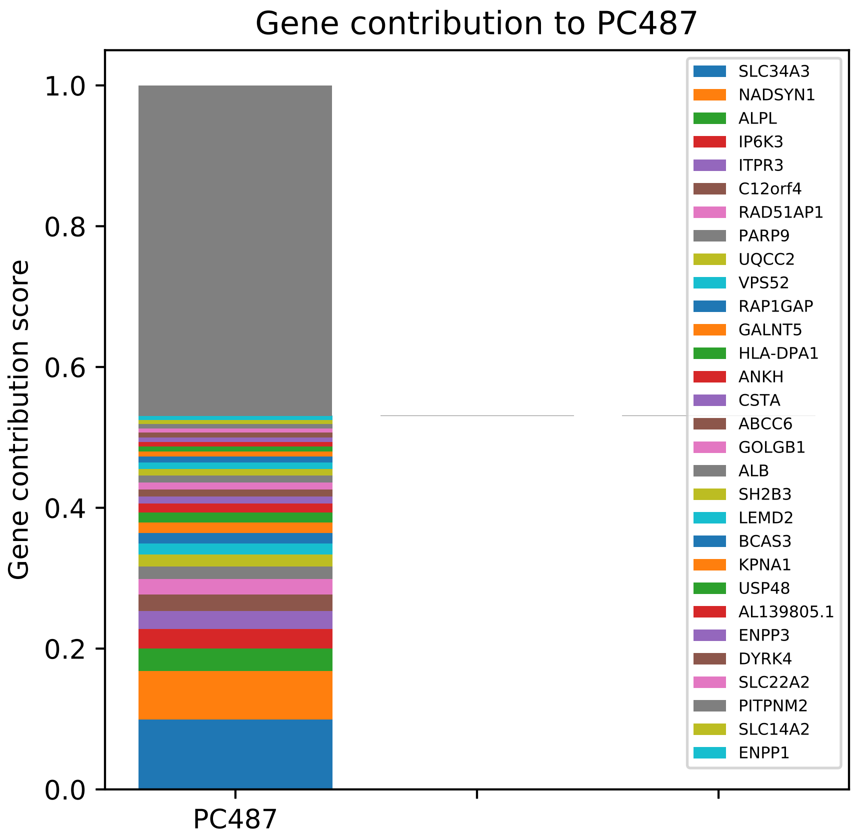

Gene contribution to PC487
Gene contribution score
1.0
0.8
0.6
0.4
0.2
0.0
PC487
SLC34A3
NADSYN1
ALPL
IP6K3
ITPR3
C12orf4
RAD51AP1
PARP9
UQCC2
VPS52
RAP1GAP
GALNT5
HLA-DPA1
ANKH
CSTA
ABCC6
GOLGB1
ALB
SH2B3
LEMD2
BCAS3
KPNA1
USP48
AL139805.1
ENPP3
DYRK4
SLC22A2
PITPNM2
SLC14A2
ENPP1

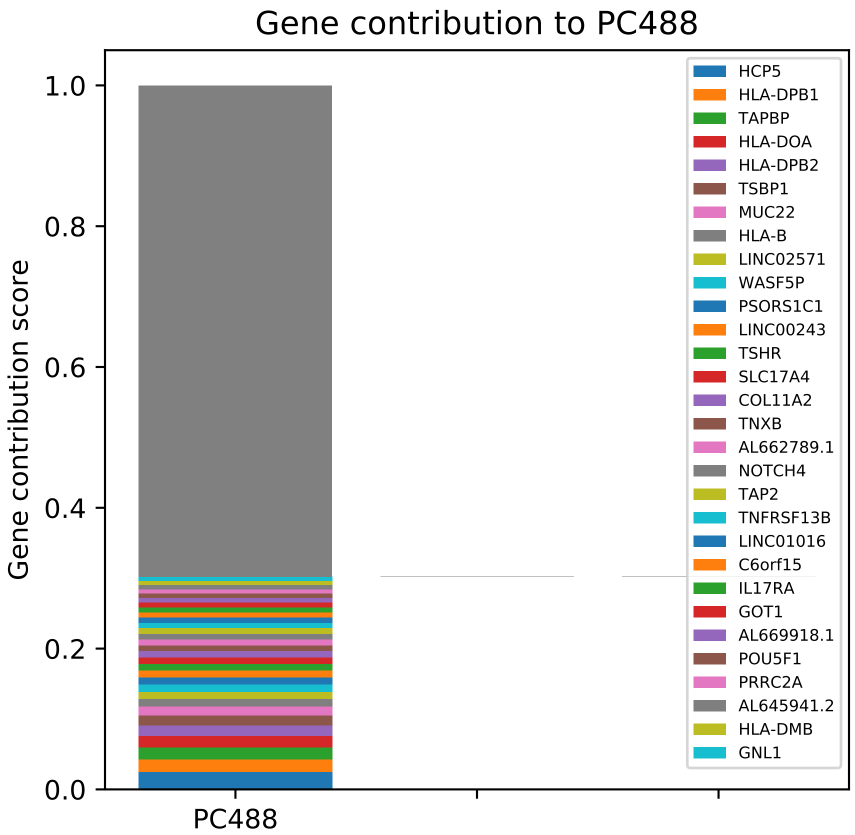
Gene contribution to PC488
Gene contribution score
1.0
0.8
0.6
0.4
0.2
0.0
PC488
HCP5
HLA-DPB1
TAPBP
HLA-DOA
HLA-DPB2
TSBP1
MUC22
HLA-B
LINC02571
WASF5P
PSORS1C1
LINC00243
TSHR
SLC17A4
COL11A2
TNXB
AL662789.1
NOTCH4
TAP2
TNFRSF13B
LINC01016
C6orf15
IL17RA
GOT1
AL669918.1
POU5F1
PRRC2A
AL645941.2
HLA-DMB
GNL1

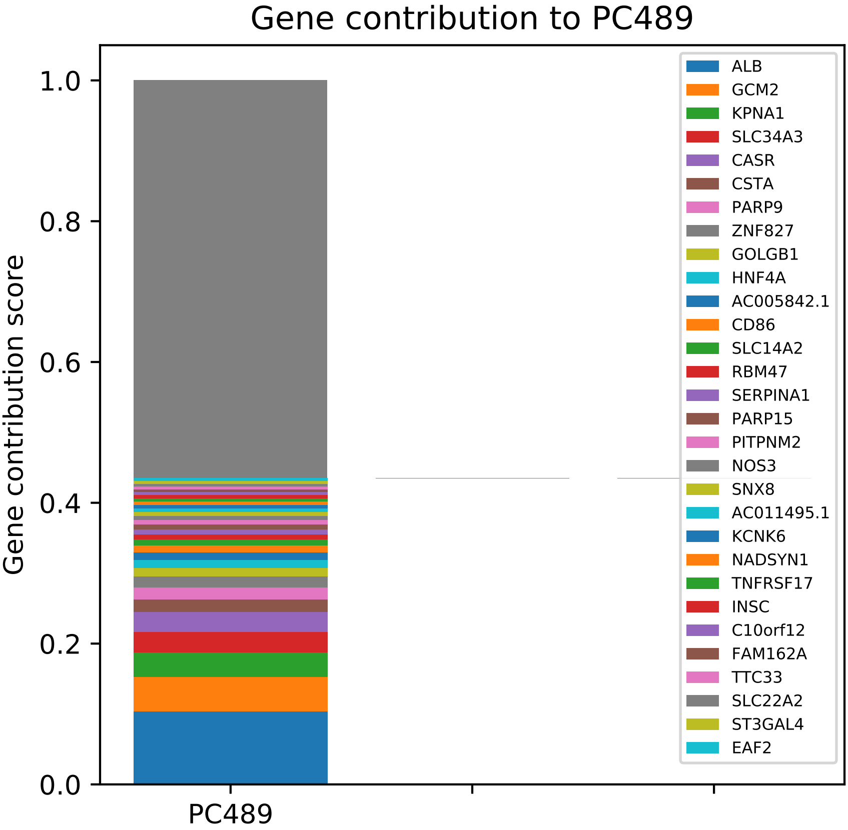

Gene contribution to PC489
Gene contribution score
1.0
0.8
0.6
0.4
0.2
0.0
PC489
ALB
GCM2
KPNA1
SLC34A3
CASR
CSTA
PARP9
ZNF827
GOLGB1
HNF4A
AC005842.1
CD86
SLC14A2
RBM47
SERPINA1
PARP15
PITPNM2
NOS3
SNX8
AC011495.1
KCNK6
NADSYN1
TNFRSF17
INSC
C10orf12
FAM162A
TTC33
SLC22A2
ST3GAL4
EAF2

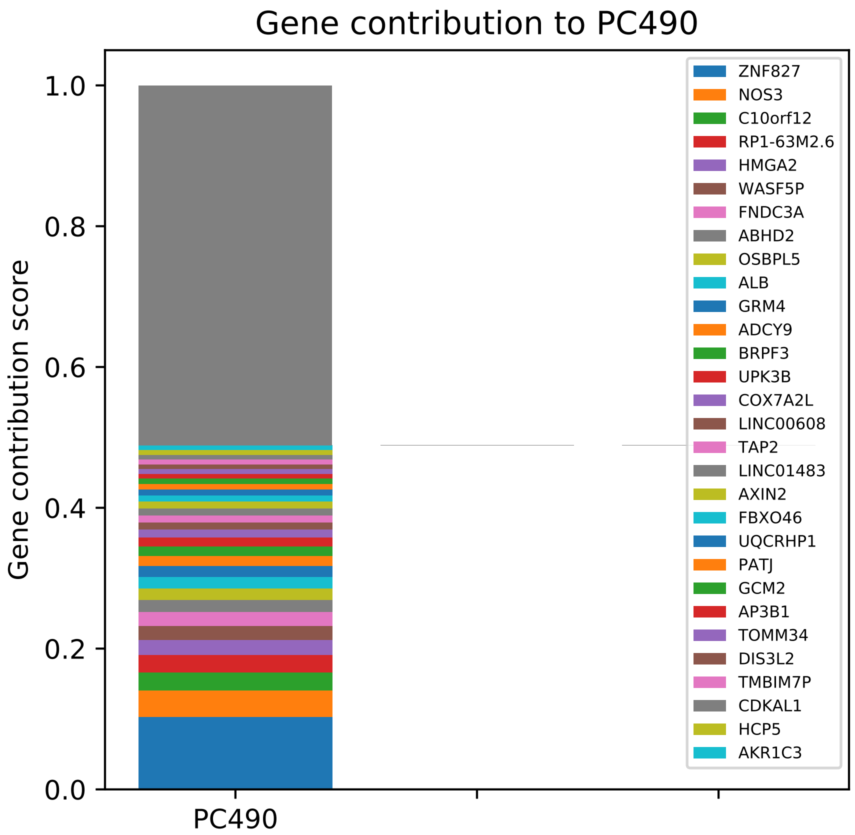

Gene contribution to PC490
Gene contribution score
1.0
0.8
0.6
0.4
0.2
0.0
PC490
ZNF827
NOS3
C10orf12
RP1-63M2.6
HMGA2
WASF5P
FNDC3A
ABHD2
OSBPL5
ALB
GRM4
ADCY9
BRPF3
UPK3B
COX7A2L
LINC00608
TAP2
LINC01483
AXIN2
FBXO46
UQCRHP1
PATJ
GCM2
AP3B1
TOMM34
DIS3L2
TMBIM7P
CDKAL1
HCP5
AKR1C3

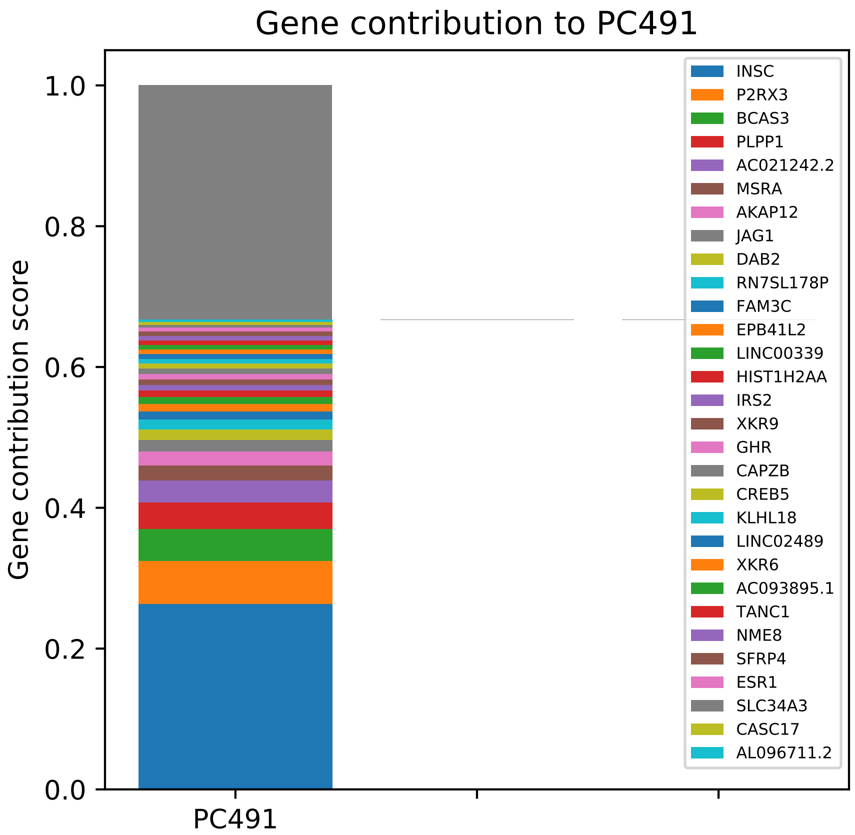

Gene contribution to PC491
Gene contribution score
0.0
0.2
0.4
0.6
0.8
1.0
PC491
INSC
P2RX3
BCAS3
PLPP1
AC021242.2
MSRA
AKAP12
JAG1
DAB2
RN7SL178P
FAM3C
EPB41L2
LINC00339
HIST1H2AA
IRS2
XKR9
GHR
CAPZB
CREB5
KLHL18
LINC02489
XKR6
AC093895.1
TANC1
NME8
SFRP4
ESR1
SLC34A3
CASC17
AL096711.2

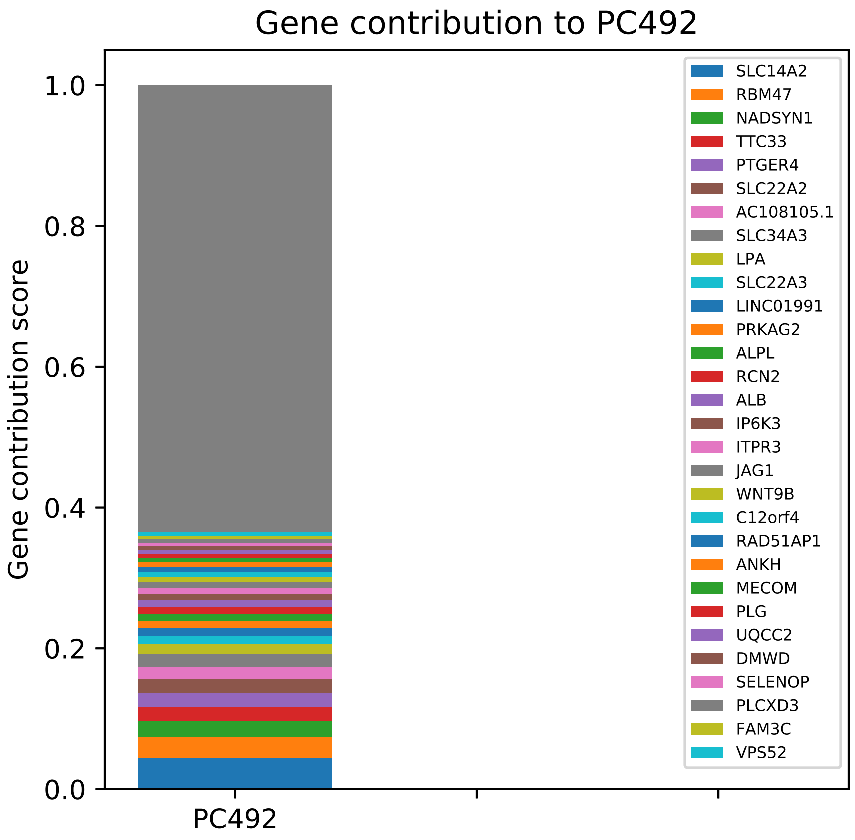

Gene contribution to PC492
Gene contribution score
1.0
0.8
0.6
0.4
0.2
0.0
PC492
SLC14A2
RBM47
NADSYN1
TTC33
PTGER4
SLC22A2
AC108105.1
SLC34A3
LPA
SLC22A3
LINC01991
PRKAG2
ALPL
RCN2
ALB
IP6K3
ITPR3
JAG1
WNT9B
C12orf4
RAD51AP1
ANKH
MECOM
PLG
UQCC2
DMWD
SELENOP
PLCXD3
FAM3C
VPS52

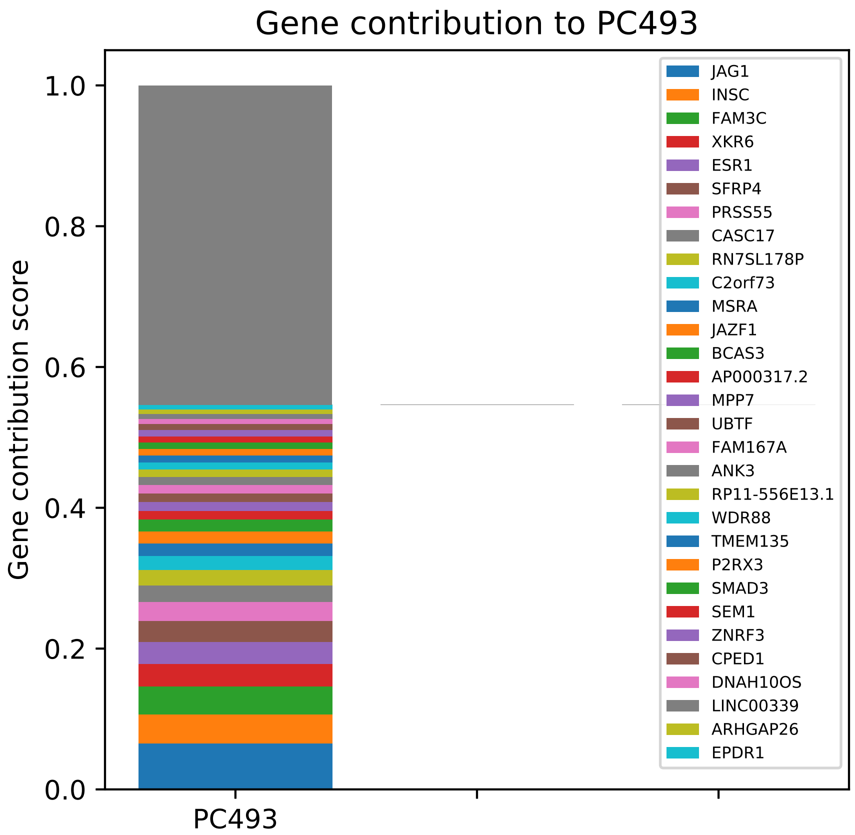
Gene contribution to PC493
Gene contribution score
1.0
0.8
0.6
0.4
0.2
0.0
PC493
JAG1
INSC
FAM3C
XKR6
ESR1
SFRP4
PRSS55
CASC17
RN7SL178P
C2orf73
MSRA
JAZF1
BCAS3
AP000317.2
MPP7
UBTF
FAM167A
ANK3
RP11-556E13.1
WDR88
TMEM135
P2RX3
SMAD3
SEM1
ZNRF3
CPED1
DNAH10OS
LINC00339
ARHGAP26
EPDR1

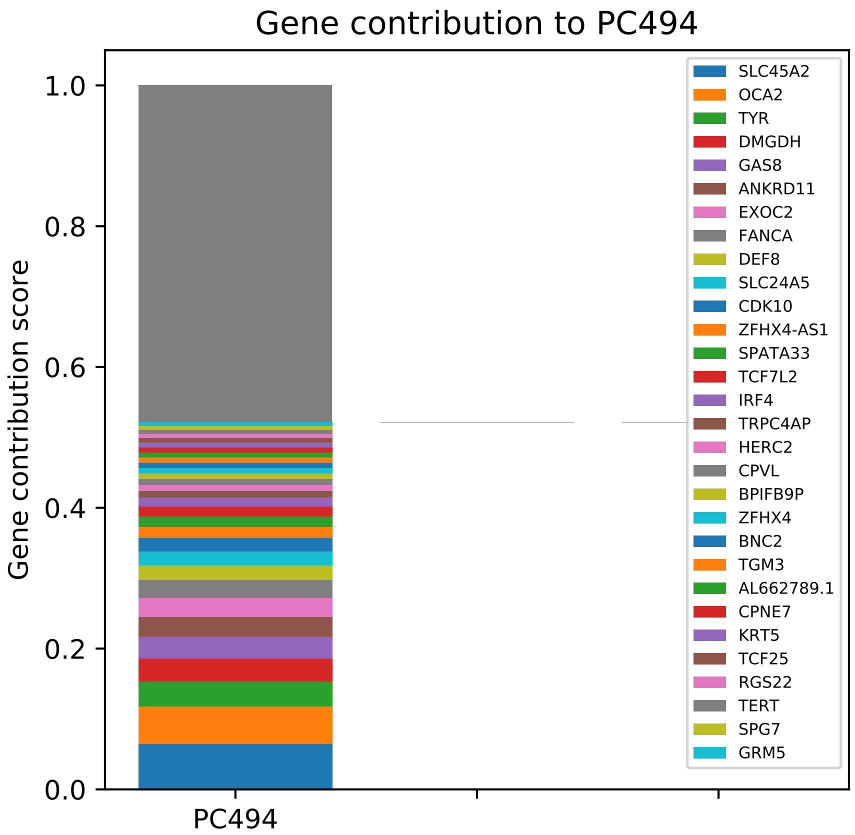

Gene contribution to PC494
Gene contribution score
0.0
0.2
0.4
0.6
0.8
1.0
PC494
SLC45A2
OCA2
TYR
DMGDH
GAS8
ANKRD11
EXOC2
FANCA
DEF8
SLC24A5
CDK10
ZFHX4-AS1
SPATA33
TCF7L2
IRF4
TRPC4AP
HERC2
CPVL
BPIFB9P
ZFHX4
BNC2
TGM3
AL662789.1
CPNE7
KRT5
TCF25
RGS22
TERT
SPG7
GRM5

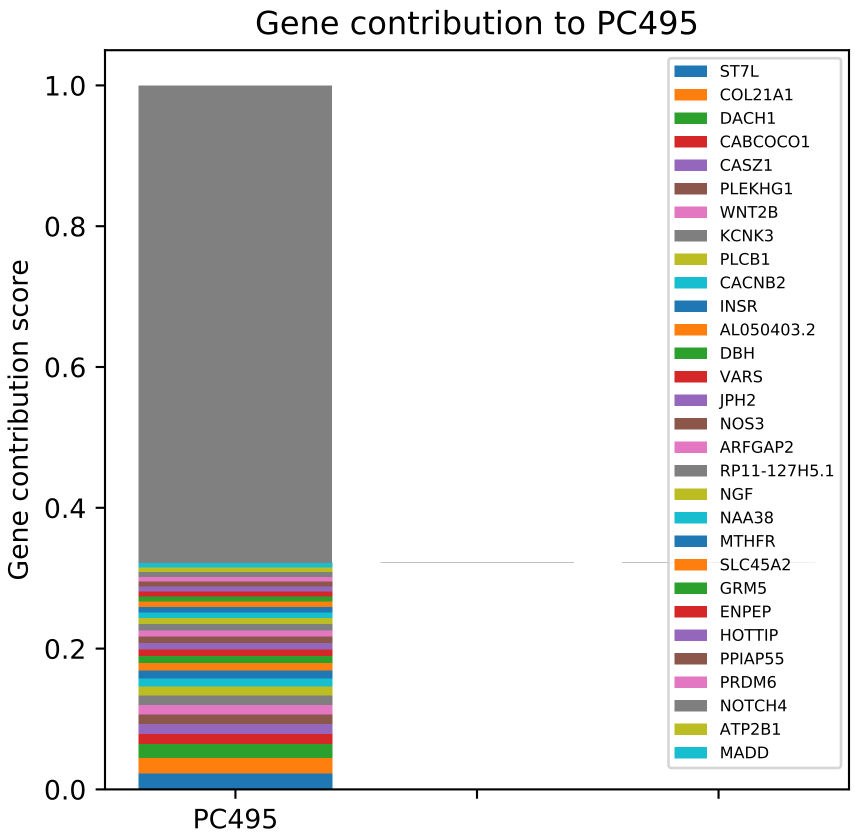

Gene contribution to PC495
Gene contribution score
1.0
0.8
0.6
0.4
0.2
0.0
PC495
ST7L
COL21A1
DACH1
CABCOCO1
CASZ1
PLEKHG1
WNT2B
KCNK3
PLCB1
CACNB2
INSR
AL050403.2
DBH
VARS
JPH2
NOS3
ARFGAP2
RP11-127H5.1
NGF
NAA38
MTHFR
SLC45A2
GRM5
ENPEP
HOTTIP
PPIAP55
PRDM6
NOTCH4
ATP2B1
MADD

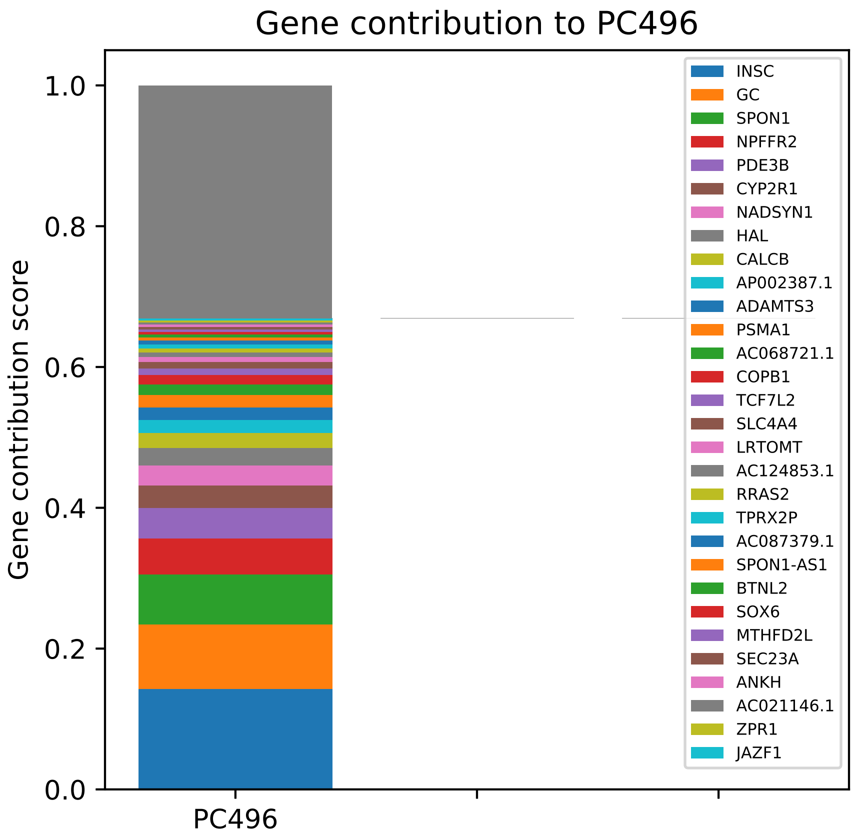

Gene contribution to PC496
Gene contribution score
0.0
0.2
0.4
0.6
0.8
1.0
PC496
INSC
GC
SPON1
NPFFR2
PDE3B
CYP2R1
NADSYN1
HAL
CALCB
AP002387.1
ADAMTS3
PSMA1
AC068721.1
COPB1
TCF7L2
SLC4A4
LRTOMT
AC124853.1
RRAS2
TPRX2P
AC087379.1
SPON1-AS1
BTNL2
SOX6
MTHFD2L
SEC23A
ANKH
AC021146.1
ZPR1
JAZF1

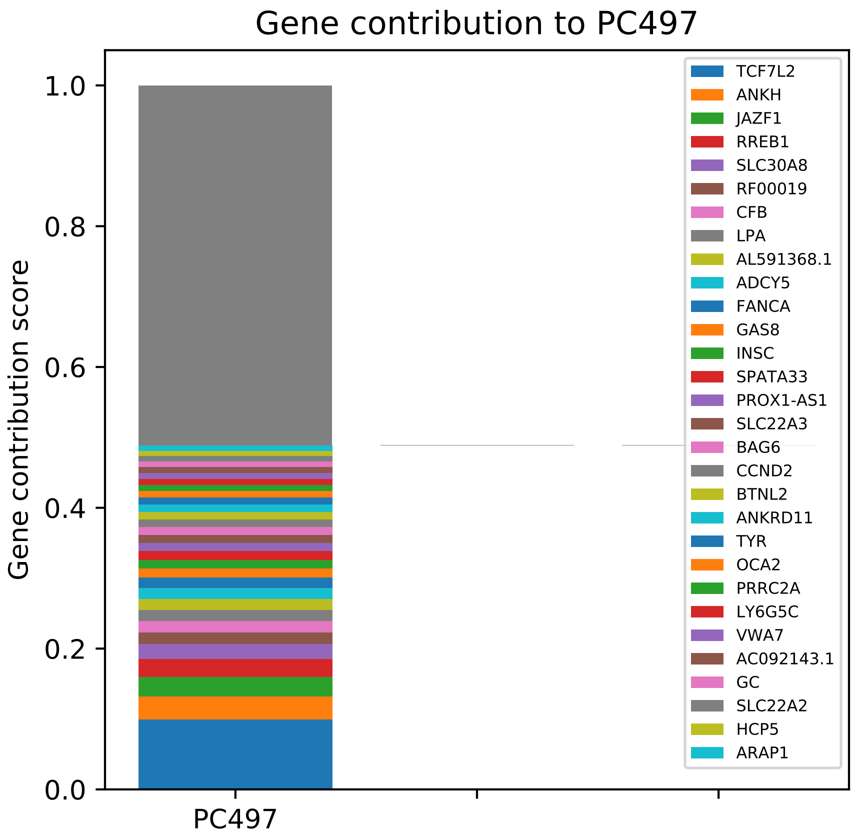

Gene contribution to PC497
Gene contribution score
0.0
0.2
0.4
0.6
0.8
1.0
PC497
TCF7L2
ANKH
JAZF1
RREB1
SLC30A8
RF00019
CFB
LPA
AL591368.1
ADCY5
FANCA
GAS8
INSC
SPATA33
PROX1-AS1
SLC22A3
BAG6
CCND2
BTNL2
ANKRD11
TYR
OCA2
PRRC2A
LY6G5C
VWA7
AC092143.1
GC
SLC22A2
HCP5
ARAP1

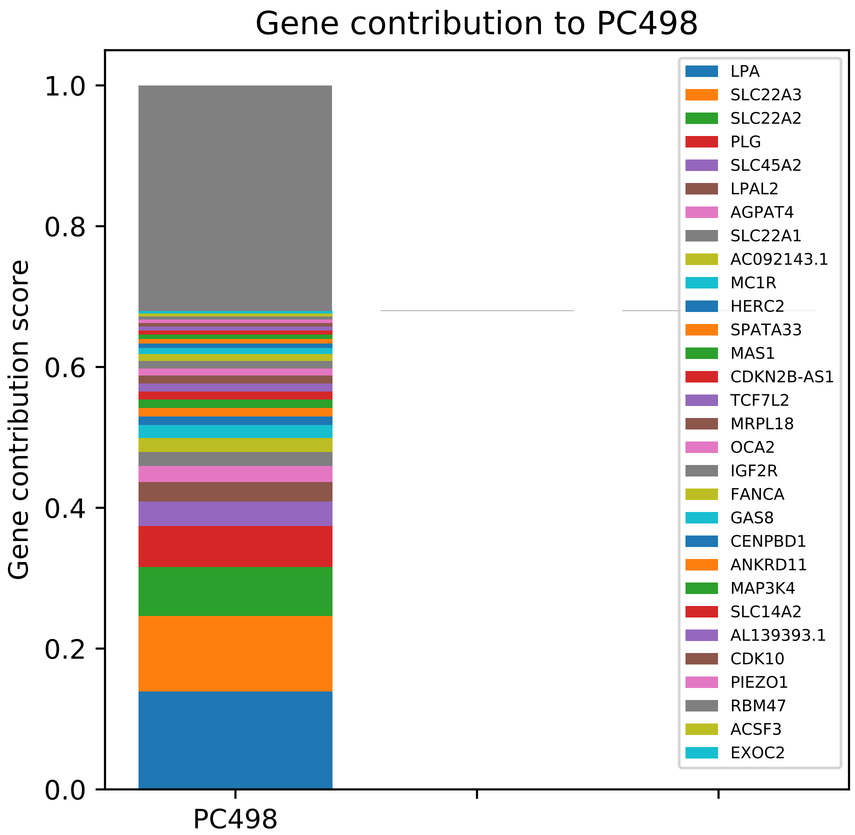

Gene contribution to PC498
Gene contribution score
0.0
0.2
0.4
0.6
0.8
1.0
PC498
LPA
SLC22A3
SLC22A2
PLG
SLC45A2
LPAL2
AGPAT4
SLC22A1
AC092143.1
MC1R
HERC2
SPATA33
MAS1
CDKN2B-AS1
TCF7L2
MRPL18
OCA2
IGF2R
FANCA
GAS8
CENPBD1
ANKRD11
MAP3K4
SLC14A2
AL139393.1
CDK10
PIEZO1
RBM47
ACSF3
EXOC2

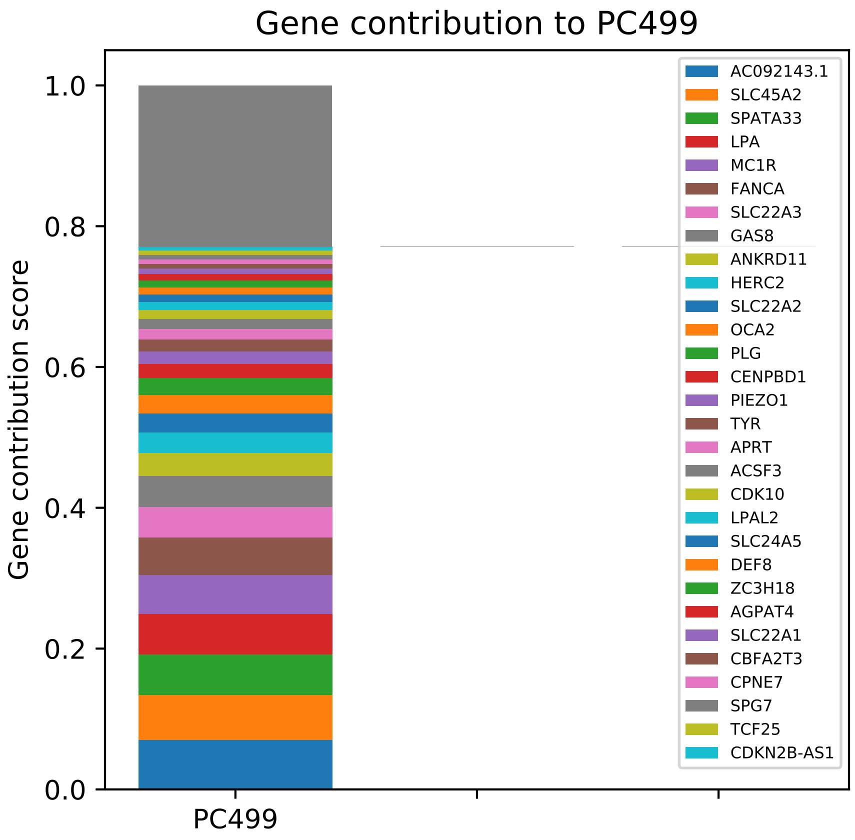
Gene contribution to PC499
Gene contribution score
0.0
0.2
0.4
0.6
0.8
1.0
PC499
AC092143.1
SLC45A2
SPATA33
LPA
MC1R
FANCA
SLC22A3
GAS8
ANKRD11
HERC2
SLC22A2
OCA2
PLG
CENPBD1
PIEZO1
TYR
APRT
ACSF3
CDK10
LPAL2
SLC24A5
DEF8
ZC3H18
AGPAT4
SLC22A1
CBFA2T3
CPNE7
SPG7
TCF25
CDKN2B-AS1

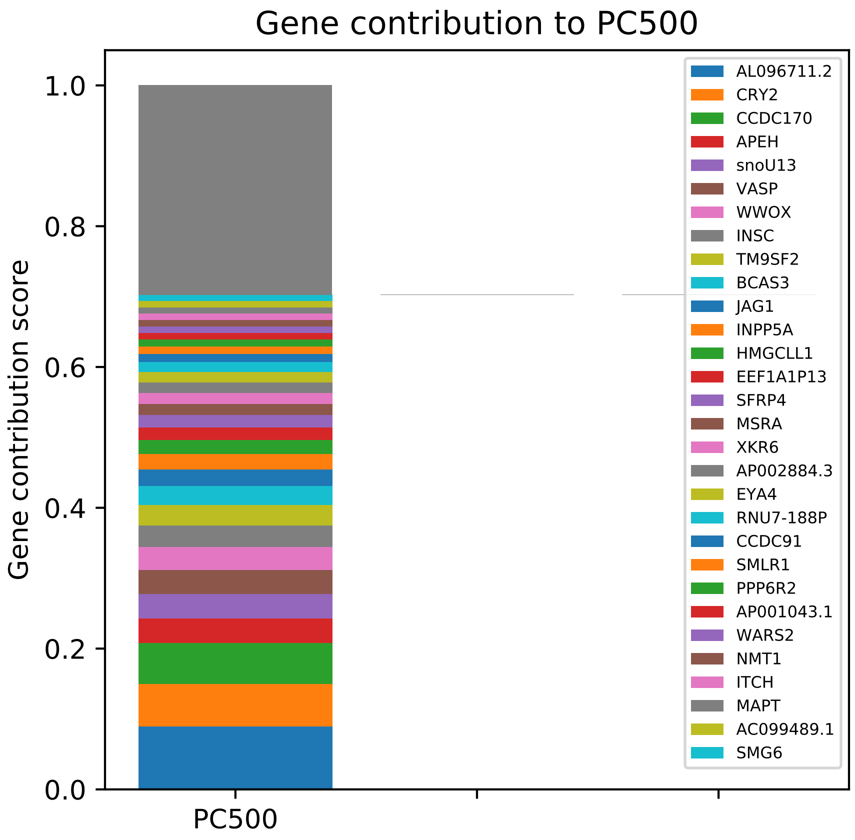
Gene contribution to PC500
Gene contribution score
0.0
0.2
0.4
0.6
0.8
1.0
PC500
AL096711.2
CRY2
CCDC170
APEH
snoU13
VASP
WWOX
INSC
TM9SF2
BCAS3
JAG1
INPP5A
HMGCLL1
EEF1A1P13
SFRP4
MSRA
XKR6
AP002884.3
EYA4
RNU7-188P
CCDC91
SMLR1
PPP6R2
AP001043.1
WARS2
NMT1
ITCH
MAPT
AC099489.1
SMG6
